# Charting the Insect Biodiversity of Crete: Insights from a Pilot Metabarcoding Survey

**DOI:** 10.64898/2026.06.05.730060

**Authors:** Georgios D. Koutsovoulos, Martin Sorg, Thomas Hörren, Dominik Buchner, Sarah J. Bourlat, Kathrin Langen, Apostolos Trichas, Florian Leese, Alexandros Stamatakis

**Affiliations:** Biodiversity Computing Group, Institute of Computer Science, Foundation for Research and Technology Hellas, 100 Nikolaou Plastira, 70013 Heraklion, Crete, Greece; Entomological Society Krefeld, Magdeburger Straße 38-40, 47800 Krefeld, Germany; University of Duisburg-Essen, Faculty of Biology & Centre for Water and Environmental Research, Universitätsstraße 5, 45141 Essen, Germany; Leibniz Institute for the Analysis of Biodiversity Change, Museum Koenig Bonn, Adenauerallee 160, 53113 Bonn, Germany; Natural History Museum of Crete, University of Crete, Heraklion, Crete, Greece; Computational Molecular Evolution Group, Heidelberg Institute for Theoretical Studies, Schloss-Wolfsbrunnenweg 35, 69118 Heidelberg, Baden-Württemberg, Germany; Institute for Theoretical Informatics, Karlsruhe Institute of Technology, Kaiserstraße 12, 76131 Karlsruhe, Baden-Württemberg, Germany

**Keywords:** biodiversity monitoring, DNA metabarcoding, insect diversity, Malaise trap, citizen science

## Abstract

Among eukaryotes, insects are by far the most diverse organisms on Earth, yet their global decline threatens ecosystem stability. Understanding local and regional biodiversity patterns is critical for conservation planning, ecosystem management, and predicting responses to environmental change, but traditional surveys for assessing insect diversity (e.g., manual collection, morphological identification, and counting) are highly labor-intensive, time-consuming, and often require rare or simply unavailable dedicated taxonomic expertise. DNA metabarcoding offers an efficient, high-resolution alternative to assess insect communities. Here, we report on the first insect metabarcoding survey on Crete that spans two years of sample collection between 2021 and 2023 from a small area in Southern Central Crete in the context of a citizen science project. A total of 29 samples yielded 10,865 Exact Sequence Variants (ESVs), 10,516 of which were assigned to insects, covering 988 species, 900 genera, and 227 families across 14 orders. A comparison with the existing observation records reveals 406 potential newly-observed species and an estimated 690 unclassified species, indicating substantial cryptic diversity. Our results demonstrate that even small-scale sampling can unravel substantial insect diversity and highlight critical gaps in barcode reference databases. Our study demonstrates how DNA metabarcoding can accelerate biodiversity discovery and monitoring in understudied regions.

## Introduction

Insects are an extremely important and by far the most species-rich component of global biodiversity. They account for around three-quarters of the estimated 8 million animal and plant species on Earth (IPBES, 2019). They contribute to a plethora of essential environmental activities including pollination, decomposition, pest control, soil health, and food chains. However, the current scientific consensus is that insect populations have been declining dramatically over the last years with potential severe ecological consequences (Hallmann et al., 2017; Wagner et al., 2021; Raven and Wagner, 2021). While this decline can predominantly be attributed to human activity (e.g., habitat loss, pesticides, climate change), other factors, such as ecosystem dynamics, may also impact insect populations. Furthermore, the number of known insect species is constantly increasing. These new members confirm our limited estimates of distinct insect species numbers (estimated at 5.5 million, with only *∼* 20% having been described so far (Stork, 2017)).

The most comprehensive approach to assess insect biodiversity is by analysing samples via classical taxonomic analyses. However, this requires taxonomic expertise and numerous man-hours. Alternatively, to quickly and reliably assess and monitor a region’s insect biodiversity, one can apply DNA metabarcoding methods to samples that have been obtained via insect traps (e.g., Malaise traps) (Buchner et al., 2025; Thomas et al., 2025). The availability of extensive barcode reference databases such as BOLD (Ratnasingham et al., 2024) facilitates high resolution taxonomic assignments that often accurately reflect species richness. Metabarcoding has proven particularly effective for detecting cryptic species and characterizing complex communities (Semmouri et al., 2021; Noguerales et al., 2023), thus, efficiently complementing or even replacing classical taxonomy in biodiversity research and monitoring (Keck et al., 2022).

Notwithstanding its effectiveness, DNA metabarcoding also exhibits limitations. Deviations in sample processing and sequencing between laboratories may induce inconsistent taxon identifications (McGee et al., 2019). Furthermore, because barcode sequences are relatively short, they often contain limited informative sequence variance, which can render distinguishing between closely related species challenging. In addition, reference database inaccuracies or errors are likely to compromise taxon identification accuracy and consequently induce erroneous conclusions. Finally, computational analyses introduce inherent biases at the workflow/pipeline stage (e.g., primer trimming parameters, choice of clustering algorithm (Fasolo et al., 2024)). However, current computational research strives to quantify and reduce some of the potential intrinsic errors induced by DNA metabarcoding analyses (Wahl et al., 2025).

Crete is the fifth largest island in the Mediterranean and the largest Greek island. Because of its varied landscape, Crete’s climate gradually changes bidirectionally from west to east and from north to south, resulting in a plethora of distinct microclimates across the island. Over 30% of the land is used for agriculture, while the rest is dominated by mountain ranges and drylands, with but a few forests. Crete’s unique geological history, location, climate, and geography led to an increased level of endemism both in flora and fauna. Within Greece, it is one of *the* hotspots in terms of endemic invertebrate biodiversity (Sfenthourakis and Legakis, 2001). Thus, Crete constitutes a priority location for biodiversity research (Schüle, 1993; Myers et al., 2000).

Insect biodiversity surveys of Crete have been conducted over the years (de Jong et al., 2014; Sfenthourakis et al., 2018; Bolanakis et al., 2024). While comprehensive barcoding studies have only recently been published, they exclusively focus on butterflies (order Lepidoptera) (Huemer et al., 2025). Here, we present the first metabarcoding study of insects on Crete conducted over a two-year period in a small region around the village of Listaros. This pilot project was chosen partly due to its practicality, as one of the authors resides in the village, and because it also embraces a citizen science approach to engage the local community in biodiversity monitoring. Listaros is located in southern central Crete and is surrounded by olive groves. It is known by locals in the Messara plain for its exceptional climate.

## Results

### Metabarcoding pipeline results

We sequenced 32 metabarcoding samples spanning 2 years from 3 Malaise traps located in or near Listaros village on Crete. The samples were split into two datasets denoted by A and B to reflect the distinct sample preparation and DNA extraction methods deployed (see Materials & Methods). The samples from dataset A and dataset B were processed independently by two different labs without prior communication which explains the differences in sample processing (e.g. sieving in 2 size fractions), the different protocols used for library preparation, the distinct primers pairs used, and the widely varying sequencing depths. Three samples from dataset A failed to produce a sufficient amount of barcodes, and we did therefore not include them in subsequent downstream analyses. For dataset A, the primers used were *fwhF2* and *Fol-degen-rev* to amplify a 313 bp fragment of the mitochondrial cytochrome c oxidase I (COX1). For dataset B, the primers *fwhF2* and *fwhR2n* were used to amplify a 205 bp fragment of COX1. Furthermore, dataset B, where each sample was sequenced in two technical replicates, exhibited a substantially increased sequencing depth (Table 1).

**Table 1.** Sequencing statistics.

|  | Dataset A | Dataset B |
| --- | --- | --- |
| Period | 04/21–05/22 | 05/22–08/23 |
| Samples | 11 | 18 |
| Total Reads | 6,851,865 | 294,458,406 |
| After QC | 6,549,210 | 293,701,763 |
| Percentage | 95.58 | 99.74 |

After dereplication, denoising, and chimera removal, a total of 10,865 Exact Sequence Variants (ESVs) remained. We classified these ESVs with our recently released raxtax (Wahl et al., 2025) tool and applied a confidence cutoff of 70 at each taxonomic level. This assigned 10,566 ESVs to insects. We subsequently applied additional filtering to these assignments, reducing the total number of Insecta-classified ESVs to 10,516 (see Materials & Methods for filtering details). In total, the taxonomic classification assigned the ESVs to 988 species, 900 genera, and 227 families belonging to 14 insect orders (Table 2). Despite the increased sequencing depth in dataset B, dataset A yields a more diverse insect representation. One explanation for this surprising discrepancy despite the sequence depth difference can be attributed to size sorting prior to sequencing. Dataset B was not sorted and therefore yielded more reads from species with a high biomass. In contrast, in dataset A, the two size fractions “small” and “large” were processed separately until sequencing, resulting in the recovery of more small species with a low biomass. However, it is challenging to determine whether the different methods or distinct collection windows, or a combination thereof best explain this difference.

**Table 2.** ESVs statistics.

|  | Total | Dataset A | Dataset B | Common |
| --- | --- | --- | --- | --- |
| ESVs | 10,865 | 3,689 | 7,176 |  |
| Insects |  |  |  |  |
| ESVs | 10,516 | 3,531 | 6,985 |  |
| Orders | 14 | 14 | 11 | 11 |
| Families | 227 | 181 | 183 | 137 |
| Genera | 900 | 624 | 584 | 309 |
| Species | 988 | 688 | 580 | 280 |

### ESV distribution

The most common orders found are Diptera, Orthoptera, Hymenoptera, Lepidoptera, Hemiptera, and Coleoptera (more than 200 ESVs assigned to each) (Fig. 1), while 60 or less ESVs are assigned to the orders Neuroptera, Mantodea, Psocodea, Blattodea, Thysanoptera (SFig. SF1). Finally, dataset A yielded only one ESV each for Dermaptera, Mecoptera, and Trichoptera. Additionally, 318 ESVs from dataset A and 282 ESVs from dataset B are broadly assigned to Insecta, but with low order or rank confidences. This indicates that they might be unknown taxa or that reference data are missing, even from taxonomically related species. Only 42.7% Dipteran ESVs are assigned to named species, while 70.5% Lepidopteran ESVs are identified at species-level. We observe the lowest percentage for Orthoptera ESVs (34.8%) which highlights the need to sample additional reference sequences for this group. ESVs that are not assigned to species names, but to higher taxonomic ranks, likely represent, either undescribed species, or incomplete reference data. In some cases, the ESV assignment is restricted to the genus level due to barcoding-related limitations that hinder differentiation between species from the same genus. For instance, B_ESV_115 with a read abundance of *∼* 300k was an exact match to moths *Bryophila rectilinea* and *Bryophila tephrocharis*, both of which have been observed on Crete (Huemer et al., 2025).

**Figure 1.**
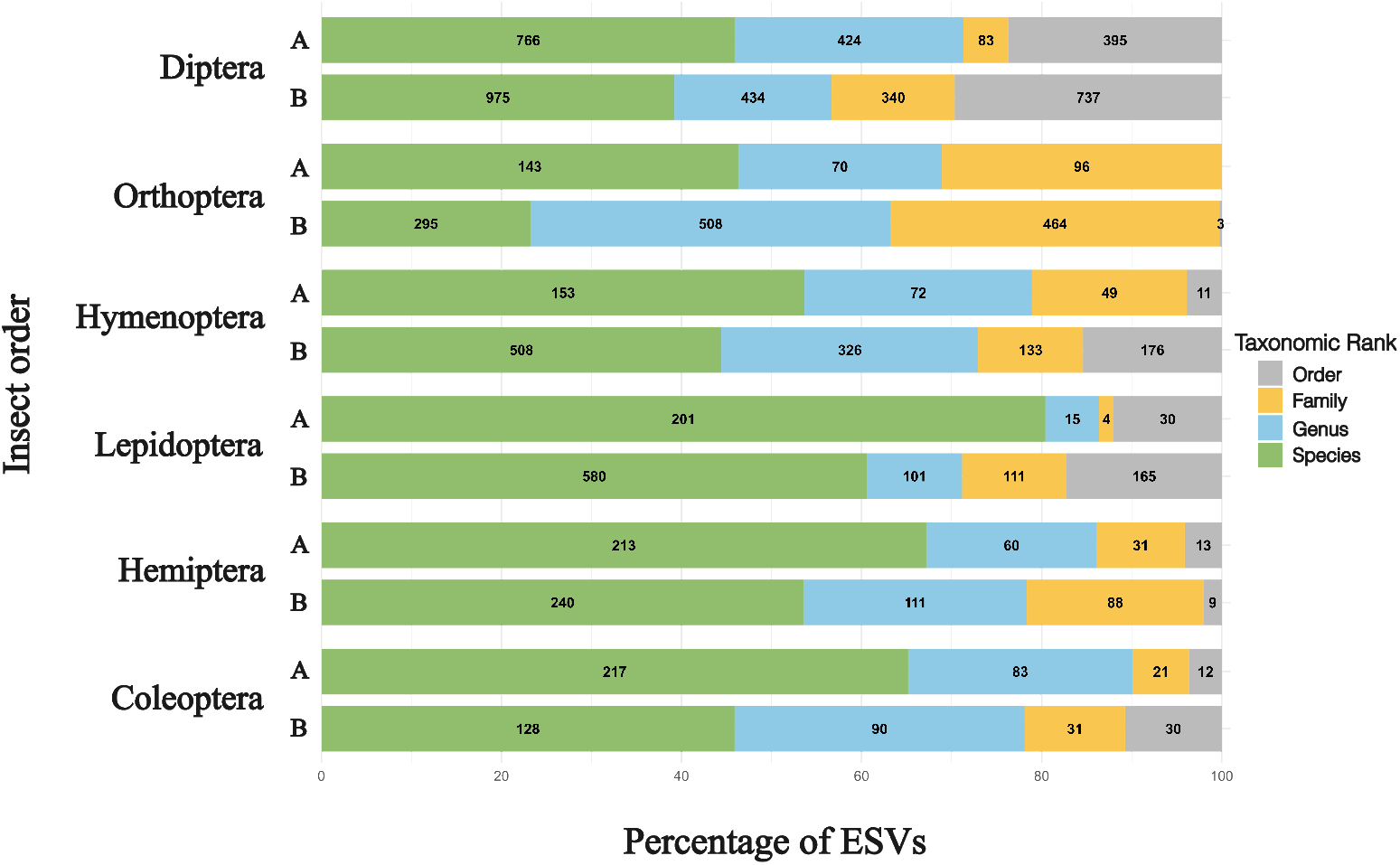
ESV assignment. Percentage of ESVs assigned up to which taxonomic rank within the most common insect orders for each dataset. The actual amount of the ESVs is shown within each box.

### Time-series results

Each sample contains a varying number of ESVs (SFig SF2). Since we collected the samples over a period of two years and used two different sample preparation and sequencing strategies, we observed differences in the proportion of ESVs assigned to a specific order. Figure 2 shows the percentage of ESVs that belong to the six most common orders per sample. Dataset B exhibits a distinctly different order composition compared to dataset A. Notably, the overall proportion of ESVs assigned to Diptera or Coleoptera is substantially lower than in dataset A, whereas the proportion of ESVs assigned to Lepidoptera is higher. Given the differing sample preparation strategies between dataset A and dataset B, a direct comparison of ESV counts across datasets is not feasible. Differences in observed order composition can therefore only be interpreted separately on a per-dataset basis.

**Figure 2.**
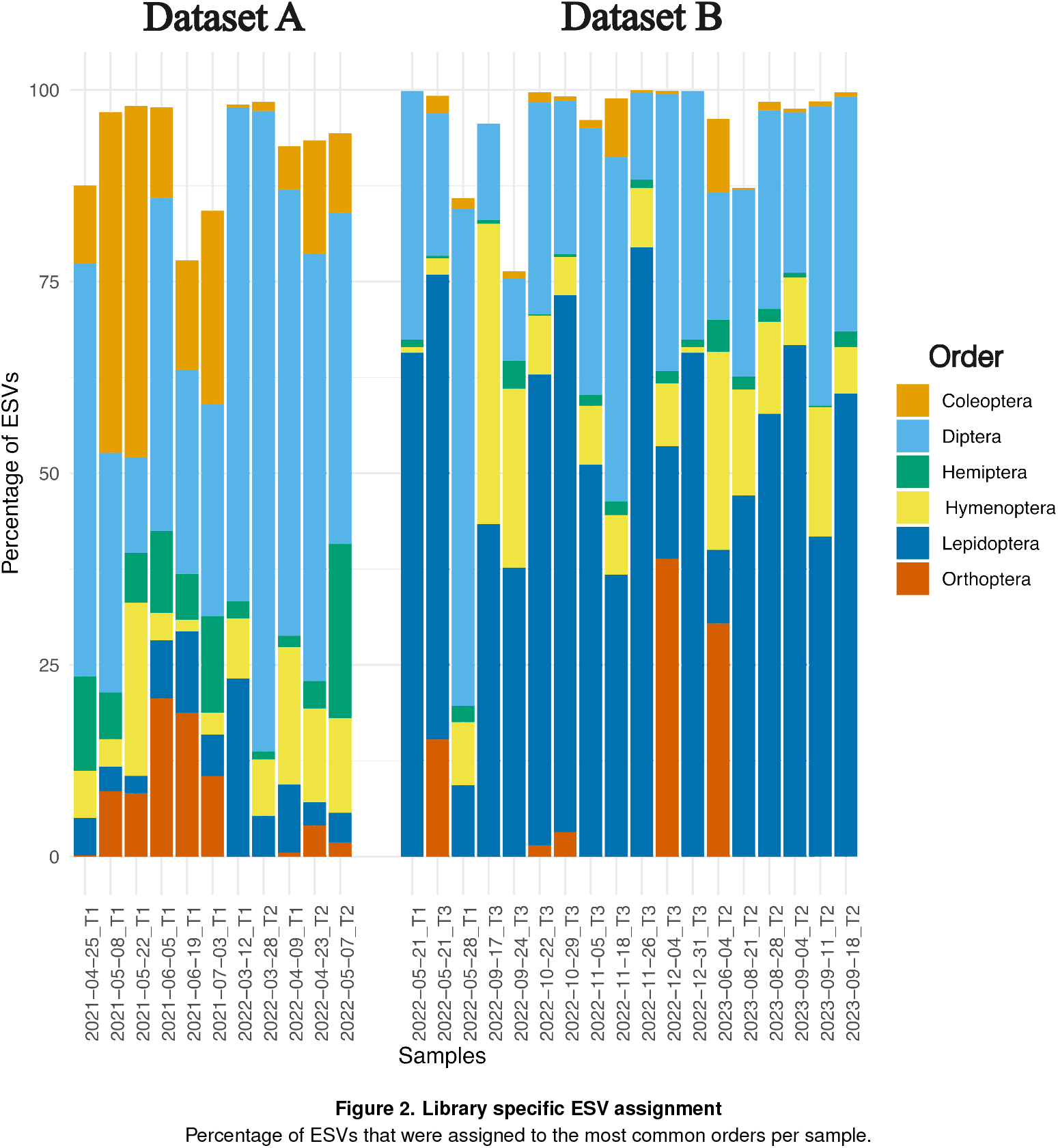
Library specific ESV assignment. Percentage of ESVs that were assigned to the most common orders per sample.

### Taxonomic assignment

We compiled an Insect Species Checklist of Crete (ISCC) for the 14 insect orders by integrating data from an earlier checklist curated by the Natural History Museum of Crete and from more recent checklists published in the scientific literature (Müller et al., 2023; Vujić et al., 2019; Ivković et al., 2017; Angelova et al., 2026; Salata et al., 2020; Huemer et al., 2025; Kotitsa et al., 2025; Willemse et al., 2018; Sziráki, 2013; Chatzimanolis et al., 2026). The complete ISCC is available in the supplementary materials. We use this checklist to cross-reference the species detected via taxonomic ESV assignments and further extend the ISCC by an additional 108 species using GBIF (GBIF.org) occurrence records.

We identify 63% of the described families and 24% of the observed genera in our samples. We also discover 33 new families and 286 new genera that are not listed in the ISCC. We provide a breakdown of these per-order findings in Supplementary Table S1. Since the area where our traps are installed is small, the low percentage of observed genera compared to the ISCC can be attributed to spatial insect distribution variation. Furthermore, the amplification of the DNA region may fail for a small percentage of species due to primer incompatibilities (Elbrecht et al., 2019). However, the observed prevalence of new families and genera in the samples highlights our elliptic knowledge of Cretan insect biodiversity.

A total of 4,685 insect ESVs were assigned with high confidence (>70%) to 988 species. Surprisingly, only 682 (59%) of the corresponding species names appear in the ISCC. The order Lepidoptera exhibits the largest overlap between our results and the ISCC, with 183 shared species, whereas for Diptera, we observe the largest number of species (193 species) that are absent from the ISCC. In addition, we could not assign a total of 6,127 ESVs to a species rank. Using the mean number of ESVs for species that were specifically classified within each order (Supplementary Table S2) and subsequently calculating the average across both datasets, this corresponds to 690 unclassified species. The inability to correctly identify these 690 species can be attributed to three main factors, i) missing reference sequences, ii) insufficient information/signal in the barcode sequence to differentiate between closely relatedtaxa, and iii) dark taxa.

For the 14 orders, 30% of the species listed in the ISCC lack a corresponding BOLD database barcode. This indicates that at least part of the unclassified species can be explained by the absence of reference sequences. The results are summarised in Figure 3 for the most common orders and in Supp. Figure SF3 for the uncommon ones. Overall, our results suggest a total of 1,678 species, of which 682 are included in the ISCC, 406 are absent from it, and 690 remain unclassified.

**Figure 3.**
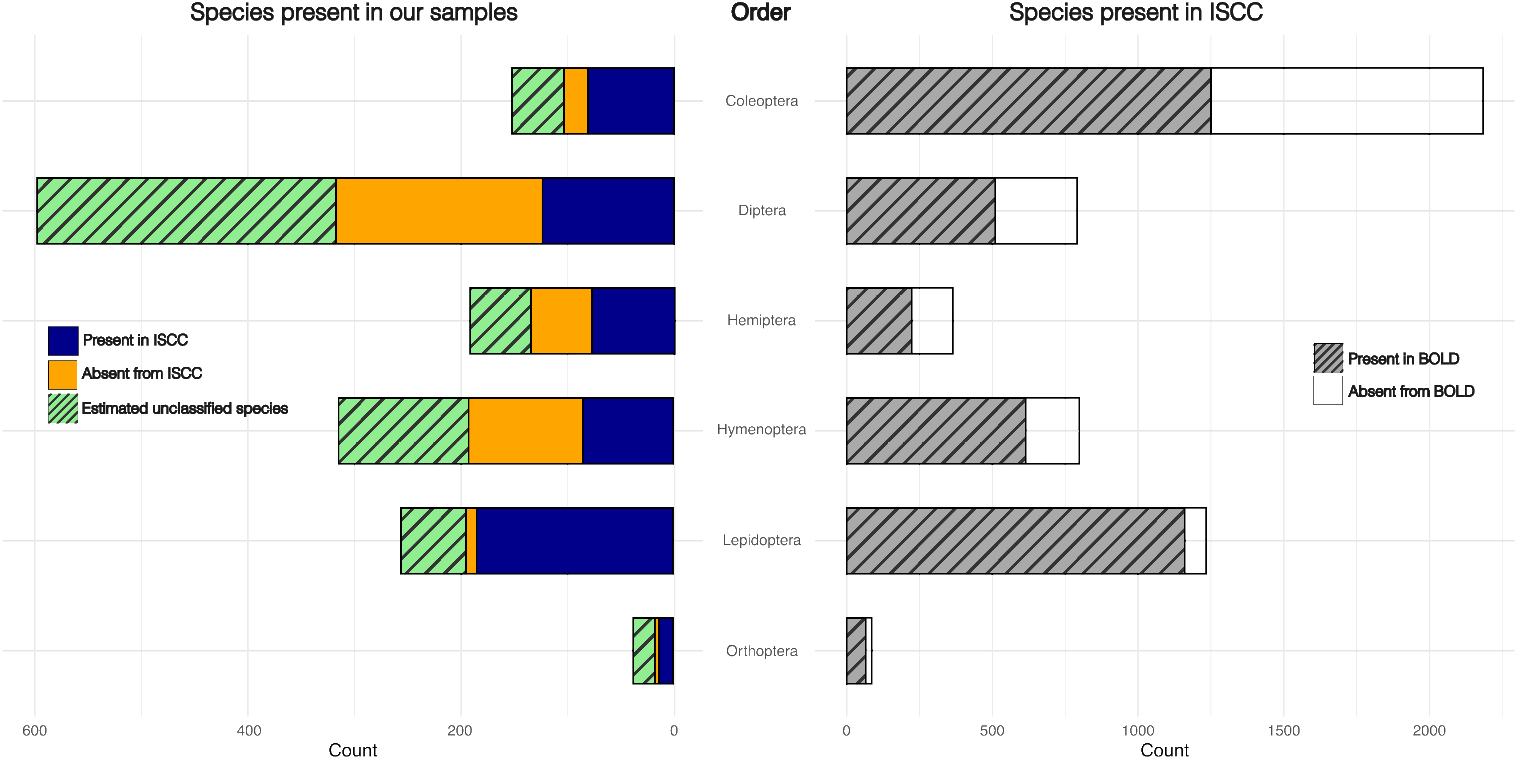
Species assignment stats. Comparison of species assignment based on the ISCC and our assignment results for the six most common insect orders.

### Rank-specific information

Taxon-specific comprehensive species checklists for Crete exist for the order Coleoptera (Chatzimanolis et al., 2026), for the family Syrphidae (order Diptera) (Vujić et al., 2019), for the suborder Auchenorrhyncha (order Hemiptera) (Angelova et al., 2026), for the family Formicidae (order Hymenoptera) (Salata et al., 2020), for the order Lepidoptera (Huemer et al., 2025), and for the order Orthoptera (Kotitsa et al., 2025; Willemse et al., 2018).

#### Coleoptera

A total of 612 ESVs were assigned to the order Coleoptera resulting in 103 species-level assignments and an estimated 49 unclassified species. A total of 81 species are in the checklist, and we report 22 species as potential new records for Crete and thereby further broaden the extensive knowledge about this order on the island. Given that almost half of the species in the checklist of more than 2,000 entries lack corresponding barcode sequences in BOLD, the estimated number of 49 unclassified species can predominantly be attributed to missing reference barcodes.

#### Diptera: Syrphidae

The taxonomic analysis resulted in 20 species-level classifications, including *Callicera fagesii* and *Paragus haemorrhous* as potential new records for Crete. Given their previously known geographic distribution in Greece, their metabarcoding-based detection on Crete is highly likely to constitute the first documented record of these species on the island.

#### Hemiptera: Auchenorrhyncha

For the following families of this suborder, our results show:

i. Cicadellidae – 21 known species, 17 potential new records
ii. Cicadidae – 1 potential new record
iii. Cixiidae – 1 known species, 1 potential new record
iv. Delphacidae – 2 known species
v. Heterogastridae – 1 known species
vi. Issidae – 5 known species, 1 potential new record

#### Hymenoptera: Formicidae

All 13 species found in our samples are in the checklist.

#### Lepidoptera

Our metabarcoding analysis recovered 1,207 ESVs assigned to the order Lepidoptera. These ESVs yielded 194 species-level identifications with an estimated number of 61 taxa exhibiting uncertain assignments. The latter group of uncertain identifications is particularly noteworthy, as it may either represent species that have not yet been described or documented on Crete, or species that have already been recorded, but lack corresponding reference barcode sequences.

Among the 194 identified species, we report 10 as potential new records for Crete, thereby expanding the known Lepidoptera diversity. This finding underscores the value of DNA metabarcoding in complementing traditional survey approaches. While classical taxonomy provides the necessary baseline, molecular methods allow for detecting cryptic diversity, highlighting gaps in reference libraries, and accelerating the discovery of novel, or previously unrecorded, taxa. The substantial proportion of uncertain taxa (61 cases) further emphasizes the need for continued taxonomic work and the importance of enriching barcode databases to better capture the regional Lepidopteran fauna.

#### Orthoptera

The analysis identified 13 species that are also present in the checklist and 4 potential new records. One of these is the species *Calliptamus abbreviatus* which has a primarily East Palearctic geographic distribution. However, the phylogenetic tree for this genus based on BOLD barcodes shows the associated ESVs to be nested within a monophyletic *C. abbreviatus* clade. Whether this constitutes a true observation or a metabarcoding artefact remains to be determined.

## Discussion

Here, we have presented the first metabarcoding analysis of Cretan insects and obtained an estimated 1,678 distinct taxa. Nearly 43% of the ESVs are assigned to 988 species with high confidence. However, it is evident that 41% of the species are assigned to previously unobserved taxa. Furthermore, our results suggest the presence of an estimated 690 unclassified species. Whether this high number merely reflects missing reference sequences in public databases or authentic dark taxa, will require additional species description and sampling efforts. Even when prudently taking into account assignment errors, our findings highlight that Cretan insect biodiversity is substantially under-sampled and under-studied.

The majority of ESVs are assigned to the six dominant orders Diptera, Hymenoptera, Lepidoptera, Orthoptera, Hemiptera, and Coleoptera, whereas several other orders are represented by but a few ESVs. Assignment success varied across groups, reflecting targeted order-specific sampling efforts. Overall, Diptera exhibits the most expansive species diversity in our samples, while Lepidoptera yields the most precise taxonomic assignments. Cross-referencing our results with the ISCC unravels a substantial knowledge gap about Cretan insects.

Across libraries, the number and composition of ESVs varies, reflecting both temporal and taxonomic differences among samples. While Diptera consistently dominates, in terms of ESV representation, other groups such as Orthoptera, Hymenoptera, and Lepidoptera exhibit fluctuations across libraries.

The sampled area, though relatively small, harbors a diverse insect community. Crete is a large island with a plethora of highly distinct microclimates that are likely to host numerous unsampled species. We thus plan to expand our research by deploying insect traps across more geographically as well as climatically diverse locations on the island to obtain a broader assessment of its insect biodiversity. Given the results from this pilot project, we have only captured a small fraction of the total insect species that are present on Crete.

## Materials & Methods

### Data collection

To measure the diversity values of insects and other invertebrates, standardized Townes-type Malaise traps from the Entomological Society of Krefeld were used (Hallmann et al., 2017; Ssymank et al., 2018). This is intended to ensure that the results obtained are comparable with other locations across Europe that have been examined using identical methodology. We installed and operated three traps, in, or near Listaros village, Crete (Supplementary Table S3). Samples were taken weekly and stored at refrigeration temperature with ethanol until their transport to Germany. The wet biomass was measured and documented according to the protocol by Hallmann et al. (2017). The samples were then delivered to the two different laboratories for metabarcoding. Dataset A containing samples obtained between April 2021 and May 2022 was prepared and sequenced with Method 1, while dataset B containing samples from May 2022 until August 2023 was prepared and sequenced via Method 2 (Supplementary Table S3).

### Sample preparation

#### Dataset A

After biomass measurements, the preservative ethanol was removed from the Malaise trap samples using a tea sieve. Wet specimens were sieved through a 4 x 4 mm mesh (wire diameter 0.5 mm, untreated stainless steel) into two size fractions: S (small,≤ 4 mm) and L (large, *>* 4 mm). Size fractions were transferred to either disposable grinding chambers (IKA, 40 or 100 mL according to sample volume) or 30 mL Nalgene tubes with metal beads (5 mm in diameter) and dried in an incubator at 50 °C for up to 7 days until complete ethanol evaporation. Dried insect tissue was homogenised for 3 minutes either with a batch mill (Tube Mill 100 Control, IKA) at 25,000 rpm or a mixer mill (MM400, Retsch) at 30 Hz for 5 minutes.

#### Dataset B

The Malaise trap samples were homogenized for 30 seconds in the Ethanol used for storage as described in Buchner et al. (2021a). One mL of the homogenate was used for subsequent DNA extraction, two additional mL were kept for long-term storage. In addition to the samples, two negative controls were generated at the homogenization step by filling the mixer with 100 mL of 96 percent ethanol. Negative controls samples were treated as regular samples for the subsequent laboratory and bioinformatic workflow.

### DNA extraction

#### Dataset A

Approximately 20 mg of finely homogenised tissue was transferred to a 1.5 mL Eppendorf tube. For tissue lysis 180 µL ATL buffer and 20 µL Proteinase K (Qiagen, Hilden, Germany) were added per sample. Samples were lysed overnight at 56 °C on a shaking incubator at 110 rpm (INCU Line ILS 6, VWR, Radnor, PA, USA). Twelve negative controls (200 µL ATL lysis buffer) were added per 84 samples to enable parallel processing of 96 samples in plate format. DNA was extracted for each size fraction (S and L) separately with the DNeasy 96 Blood and Tissue Kit (Quiagen, Hilden, Germany) following the manufacturer’s instructions. Extraction success and DNA quality were checked on a 1% agarose gel.

Ground tissue samples, DNA extracts as well as sample metadata were submitted to the LIB biobank and are stored under accession numbers: ZFMK-TIS-63870 - ZFMK-TIS-63881 (ZFMK-Tissue-Bank_1451) and ZFMK-TIS-68987 - ZFMK-TIS-69002 (ZFMK-Tissue-Bank_1499) for the tissue samples, and boxes ZFMK-DNA-Bank_1922_SA00936062 and ZFMK-DNA-Bank_1912_SA00936037 for the DNA extracts.

#### Dataset B

The homogenate was lysed as described in Buchner (2022a) for 20 minutes at 56 °C. 300 µL of the resulting lysate was used for DNA extraction using a silica spin-column based protocol as described in Buchner (2025a) with an elution volume of 100 µL. Extraction success was validated via 1 percent agarose gel. To control for possible cross-contamination during laboratory processing, each sample was processed in duplicate as proposed in Buchner et al. (2021b).

### Sequencing

#### Dataset A

A two-step PCR protocol was used for amplicon library preparation (Bourlat et al., 2016) using standard Illumina Nextera primers for dual indexing of samples. The first PCR was performed with the PCR Multiplex Plus Kit (Qiagen, Hilden, Germany) using 12.5 µL master mix, 1 µL of DNA template, 0.2 µM of the fwhF2 forward (GGDACWGGWTGAACWGTWTAY-CCHCC, (Vamos et al., 2017)) and Fol_degen_rev reverse (TANACYTCNGGRTGNCCRAARAAYCA, (Yu et al., 2012)) primers respectively, and 10.5 µL ddH_2_O to make up a 25 µL final reaction volume. The primer pair targets a 313 bp long stretch of the COX1 DNA barcode region and ensures a sufficient fragment overlap during paired-end merging after 2 x 250 bp sequencing. The following PCR program was applied on a 2720 Thermal Cycler (Applied Biosystems): initial denaturation at 95 °C for 5 min; 30 cycles of: 30s at 95 °C, 30s at 50 °C and 50s at 72 °C; final extension of 5 min at 72 °C. In the first step (PCR 1), two positive controls were included in sample processing (P1: 435 species of morphologically determined arthropod species distributed in Germany; P2: 26 species of morphologically determined species distributed on the Canary Islands with no known occurrences in Germany).

In the second PCR, combinatorial dual indexing using a set of forward and reverse primers with unique identifiers was applied to guarantee the assignment of sequences to the sample of origin. The reaction included 1 µL PCR 1 product template, 0.2 µM of each tagging primer (Nextera, Illumina, San Diego, USA), 12.5 µL master mix (PCR Multiplex Plus Kit, Qiagen, Hilden, Germany), and 9.5 µL ddH2O. The PCR 2, was run with the same program as PCR 1, but with 15 instead of 25 cycles. The two size fractions were kept separate for sequencing.

PCR success was evaluated on a 1 % agarose gel, before PCR products were normalised via a Sequal-Prep normalisation plate (Thermo Fisher Scientific, MA, USA) with sequential elution resulting in an end concentration of 25 ng per sample (100 µL). For each sample, 10 µL product was pooled, and left-sided size selection was applied on the sample pool with magnetic beads to remove residual primers (ratio 0.70x, SPRIselect Beckman Coulter). Library concentration was measured with a Quantus fluorometer (Promega, Madison, USA) and on a FragmentAnalyzer (Agilent Technologies, Santa Clara, CA, USA). The pool was sequenced via two Illumina Hiseq 2500 runs (2 x 250 bp) by Macrogen Europe B.V. in the Netherlands.

#### Dataset B

A two-step protocol was used for amplicon library preparation using a dual-twin indexing approach. Samples were tagged with a unique combination of tags in the first PCR as well as in the second PCR, yielding a unique tag pair per sample. DNA was amplified in the first PCR according to Buchner (2025b) with the fwh2 primer pair (Vamos et al., 2017) with the AccuStart II PCR supermix (QuantaBio, Beverly, USA). PCR products were cleaned up as described in Buchner (2022b) with a ratio of 0.8x. The second PCR was performed according to Buchner (2025c) with the same polymerase as the first PCR. PCR success was validated via 1 percent agarose gel. PCR products were normalized with a ratio of 0.7x as described in Buchner (2022c). To remove over-amplified PCR products a reconditioning PCR was performed according to Buchner (2025d). The final library pool was sequenced on a NovaSeq 6000 (105 Gb data package, 300 cycles) at Genewiz (Leipzig, Germany).

### Bioinformatics workflow

Each dataset was processed independently using its corresponding primer pair. The demultiplexed raw reads were analysed through a custom Python pipeline (available at https://github.com/GDKO/R2V2) that is described in detail below. Paired-end reads were merged using PEAR (Zhang et al., 2013), and primers were trimmed with cutadapt (Martin, 2011). For dataset A, we retained reads ranging from 282 bp to 344 bp, while for dataset B we kept reads between 185 bp and 225 bp. Global dereplication was performed separately for each dataset. For dataset B, only sequences detected in both technical replicates were retained, and reads present in negative controls were removed. We then filtered sequences by keeping those with a minimum abundance of 4 and a mean maximum expected error below 1. We only kept sequences containing a valid open reading frame (ORF). We validated the correctness of ORFs via HMMER (Eddy, 2011) using the HMM profile from (Porter and Hajibabaei, 2018). Finally, we used VSEARCH (Rognes et al., 2016) to denoise the reads into exact sequence variants (ESVs) and to detect and remove chimeras.

### Taxonomic assignment

We downloaded the BOLD Public Data Package (BOLD Systems, 2026) and further selected all the COX1 sequences that contained taxonomic information at the species rank. For each dataset, BOLD was processed separately by trimming the sequences with the corresponding primer pairs. We selected entries with a sequence length between 185 bp and 344 bp to be used for dataset A, and between 185 bp and 225 bp to be used for dataset B. We used our raxtax tool (Wahl et al., 2025) to query our ESVs against these reduced BOLD databases, and only considered taxonomic information with a confidence score ≥ 70. We selected 70 as a robust near-optimal cutoff because it lies on the plateau of the True Positive %-False Positive % curve across two cross-validation runs, where performance is nearly unchanged despite small shifts in the exact optimum. For species that were not present in ISCC, we extracted sequences belonging to the same genus from the reduced BOLD database and created multiple sequence alignments (MSAs) along with the associated ESVs using muscle (Edgar, 2022). We constructed phylogenetic trees using raxml-ng v2.0.0 (Kozlov et al., 2019) with automatic model selection and used these to filter unreliable species assignments. This resulted in 244 species-level assignments being downgraded to genus rank (25 of these due to barcode limitations), 44 to family rank, and 8 to be entirely removed as fungal or bacterial contamination. We then used these filtered results for all subsequent analyses.

### Visualisation

To enhance accessibility and facilitate result exploration, we provide both, a Krona plot (Ondov et al., 2011), and a Lifemap HTML file (de Vienne, 2016) as supplementary materials.

## Data availability

The sequencing data generated in this study have been deposited in the European Nucleotide Archive (ENA) under BioProject accession number PRJEB114132.

## Author Contributions

G.D.K. and A.S. conceptualized the study; M.S. and T.H. provided resources for the traps; A.S. collected most of the samples; D.B., S.J.B., K.L., and F.L. performed the laboratory investigation; G.D.K., A.T., and A.S. performed formal analysis and interpretation of the results; all authors contributed to writing and reviewing the manuscript.

## Acknowledgments

We thank the people of Listaros, especially Vaggelis Chourdakis, Manolis Fragkioudakis, and Jannis Kampourakis. We are also grateful to The Cultural Committee of Listaros Village (Klimatogi) for their support.

## Funding

This work was funded by the European Union’s Horizon Europe ERA Chair program under grant agreement No. 101087081 (Comp-Biodiv-GR) and by the Klaus Tschira Foundation via the Heidelberg Institute of Theoretical Studies.

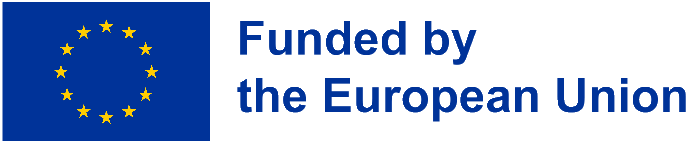

## Supplementary Figures

**Figure SF1.**
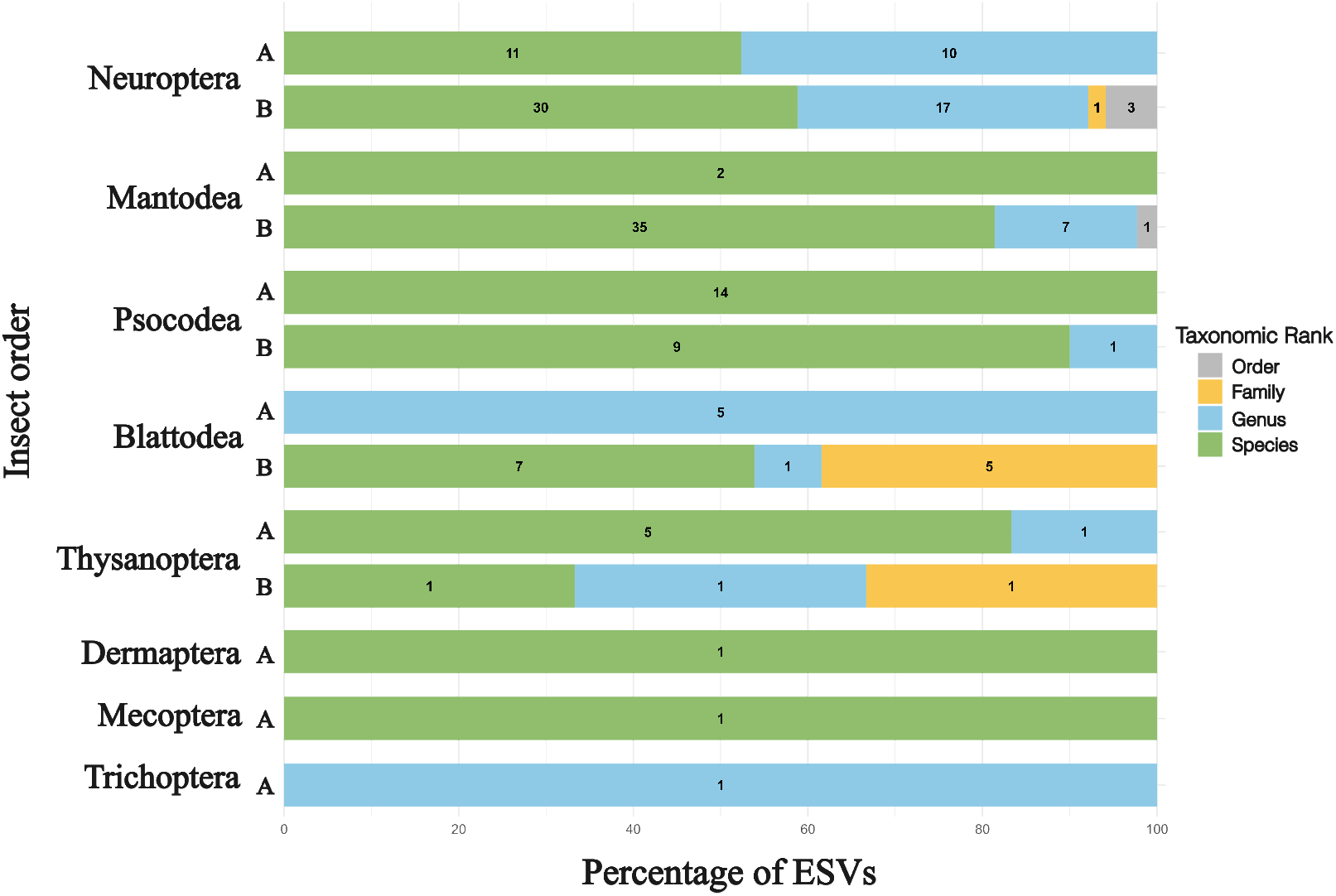
Percentage of ESVs assigned up to which taxonomic rank within the most uncommon insect orders for each dataset. The actual amount of the ESVs is shown within each box.

**Figure SF2.**
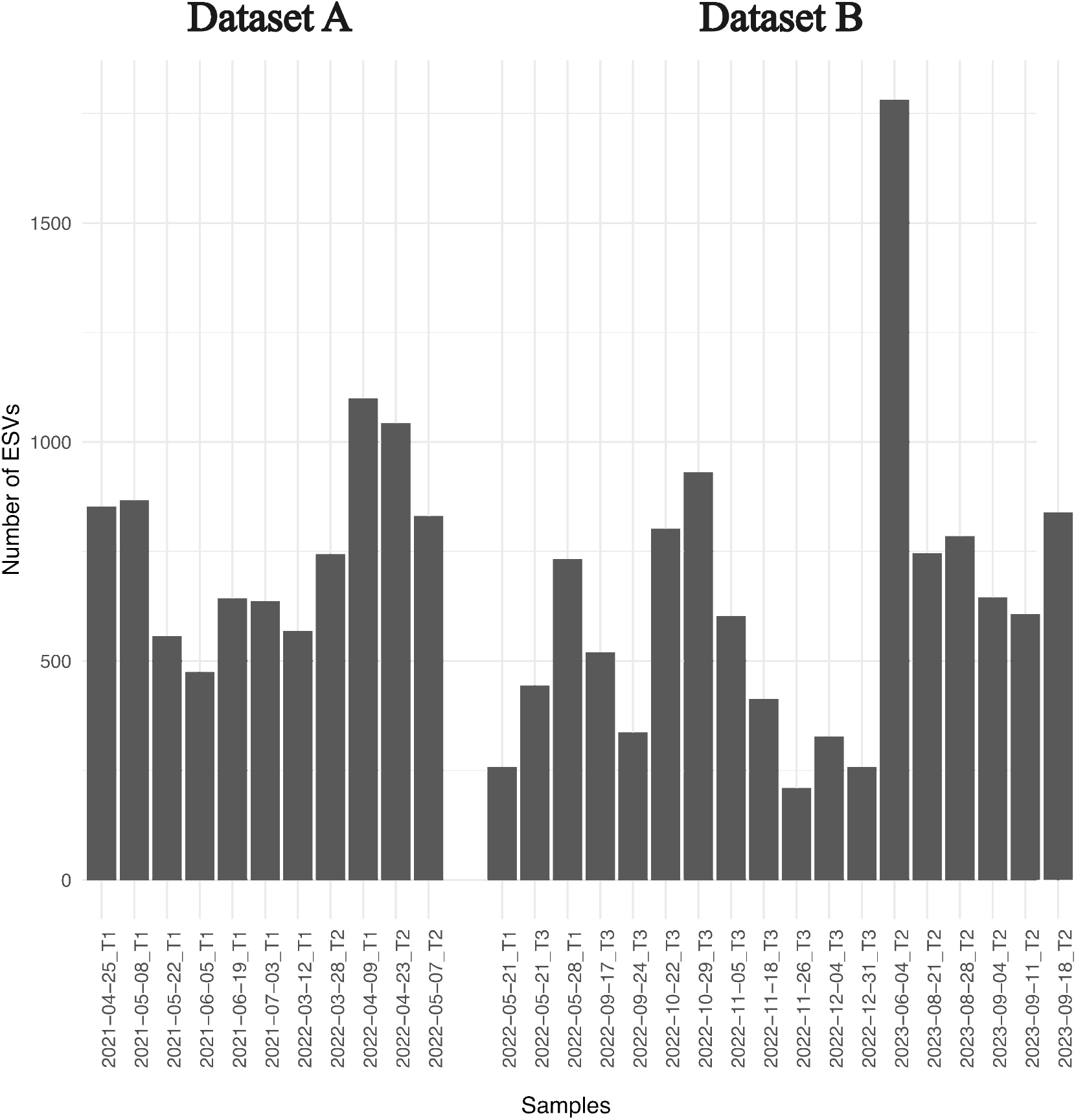
Number of ESVs per library.

**Figure SF3.**
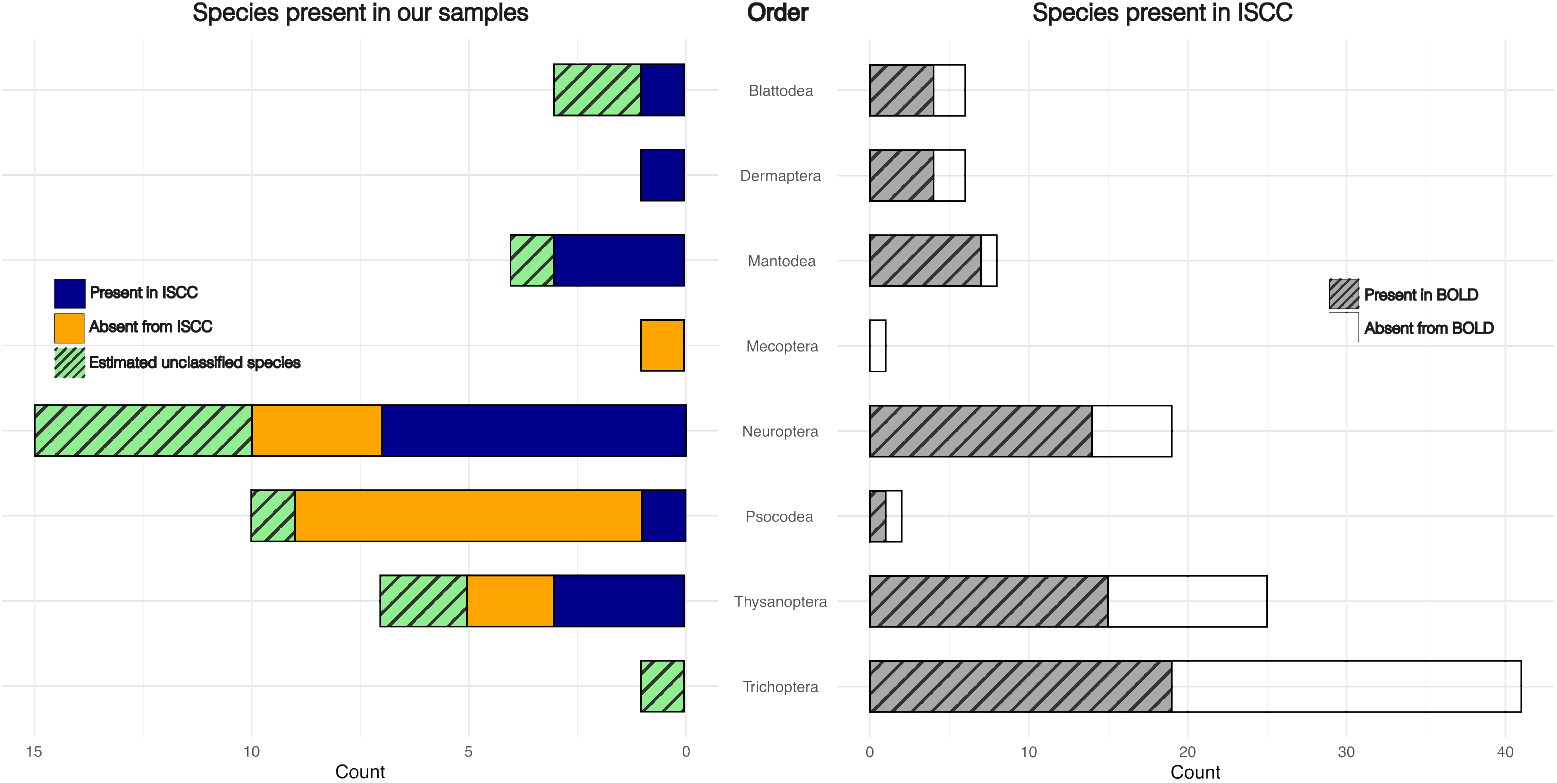
Species assignment stats. Comparison of species assignment based on the ISCC and our assignment results for the uncommon insect orders.

## Supplementary Tables

**Table S1.** Families and Genera statistics for each order.

| Order | Rank | Reference | Common | New |
| --- | --- | --- | --- | --- |
| Blattodea | Families | 4 | 2 | 1 |
|  | Genera | 5 | 1 | 2 |
| Coleoptera | Families | 63 | 30 | 0 |
|  | Genera | 895 | 101 | 11 |
| Dermaptera | Families | 3 | 1 | 0 |
|  | Genera | 4 | 1 | 0 |
| Diptera | Families | 58 | 45 | 9 |
|  | Genera | 380 | 138 | 108 |
| Hemiptera | Families | 38 | 27 | 6 |
|  | Genera | 246 | 89 | 38 |
| Hymenoptera | Families | 35 | 27 | 16 |
|  | Genera | 262 | 88 | 110 |
| Lepidoptera | Families | 62 | 41 | 0 |
|  | Genera | 604 | 162 | 5 |
| Mantodea | Families | 5 | 2 | 0 |
|  | Genera | 6 | 2 | 0 |
| Mecoptera | Families | 1 | 1 | 0 |
|  | Genera | 1 | 1 | 0 |
| Neuroptera | Families | 6 | 4 | 0 |
|  | Genera | 14 | 6 | 1 |
| Orthoptera | Families | 11 | 3 | 1 |
|  | Genera | 51 | 14 | 8 |
| Psocodea | Families | 2 | 2 | 0 |
|  | Genera | 2 | 2 | 0 |
| Thysanoptera | Families | 4 | 4 | 0 |
|  | Genera | 12 | 3 | 3 |
| Trichoptera | Families | 11 | 1 | 0 |
|  | Genera | 21 | 1 | 0 |

**Table S2.**
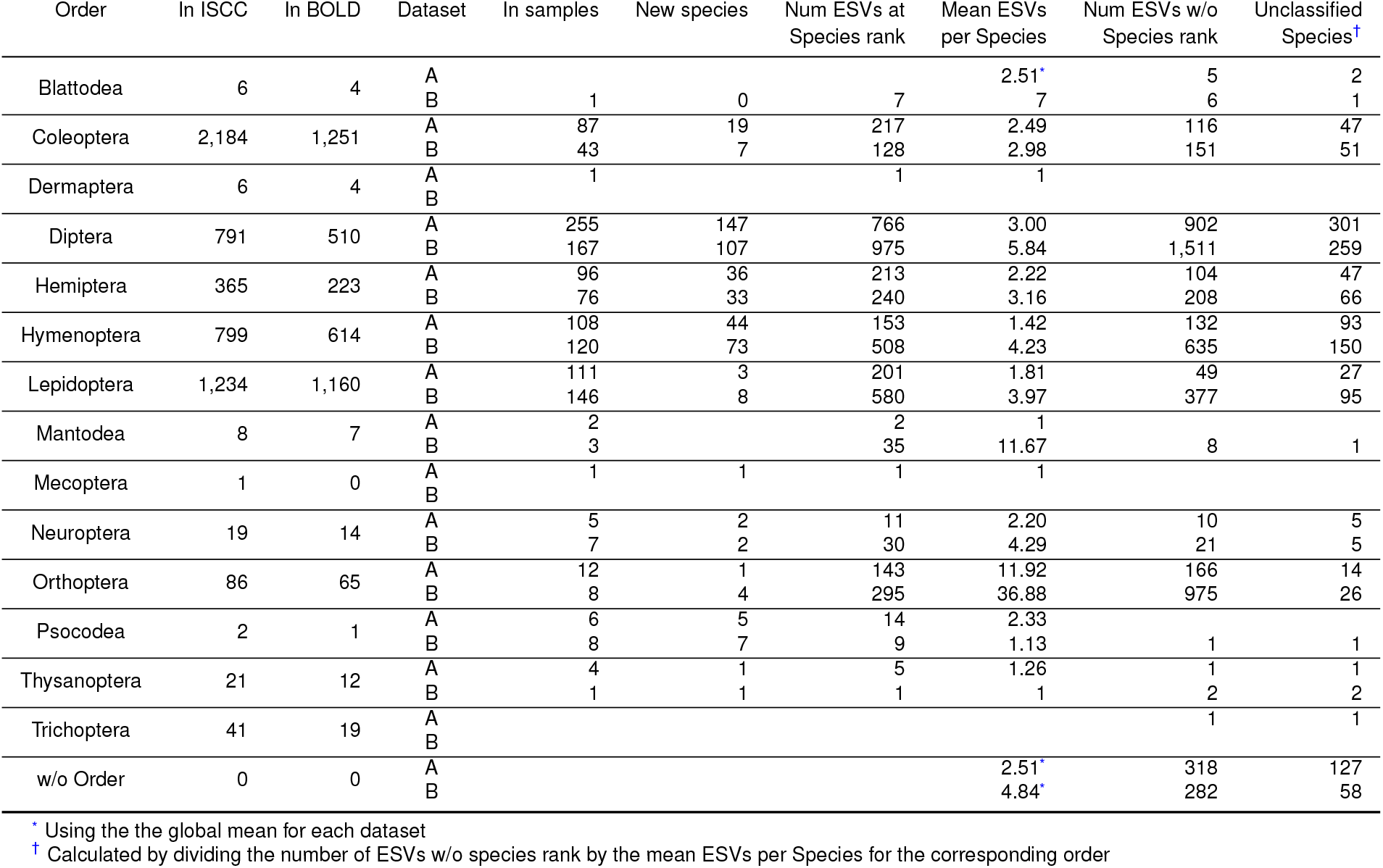
Species statistics for each order.

**Table S3.** Sampling per trap.

| Location | Trap 1 (T1)<br>34.999861N<br>24.817528E | Trap 2 (T2)<br>35.001472N<br>24.826306E | Trap 3 (T3)<br>35.000028N<br>24.817944E |
| --- | --- | --- | --- |
| Dataset A |  |  |  |
| 2021-04-25_T1 | X |  |  |
| 2021-05-08_T1 | X |  |  |
| 2021-05-22_T1 | X |  |  |
| 2021-06-05_T1 | X |  |  |
| 2021-06-19_T1 | X |  |  |
| 2021-07-03_T1 | X |  |  |
| 2022-03-12_T1 | X |  |  |
| 2022-03-28_T2 |  | X |  |
| 2022-04-09_T1 | X |  |  |
| 2022-04-23_T2 |  | X |  |
| 2022-05-07_T2 |  | X |  |
| Dataset B |  |  |  |
| 2022-05-21_T1 | X |  |  |
| 2022-05-21_T3 |  |  | X |
| 2022-05-28_T1 | X |  |  |
| 2022-09-17_T3 |  |  | X |
| 2022-09-24_T3 |  |  | X |
| 2022-10-22_T3 |  |  | X |
| 2022-10-29_T3 |  |  | X |
| 2022-11-05_T3 |  |  | X |
| 2022-11-18_T3 |  |  | X |
| 2022-11-26_T3 |  |  | X |
| 2022-12-04_T3 |  |  | X |
| 2022-12-31_T3 |  |  | X |
| 2023-06-04_T2 |  | X |  |
| 2023-08-21_T2 |  | X |  |
| 2023-08-28_T2 |  | X |  |
| 2023-09-04_T2 |  | X |  |
| 2023-09-11_T2 |  | X |  |
| 2023-09-18_T2 |  | X |  |

