## Supplementary material for "Charting the Insect Biodiversity of Crete: Insights from a Pilot Metabarcoding Survey": Krona_abundances.html

Javascript must be enabled to view this page.

xml version="1.0" ?


abundance
conf

All
2021-04-25\_T1
2021-05-08\_T1
2021-05-22\_T1
2021-06-05\_T1
2021-06-19\_T1
2021-07-03\_T1
2022-03-12\_T1
2022-03-28\_T2
2022-04-09\_T1
2022-04-23\_T2
2022-05-07\_T2
2022-05-21\_T1
2022-05-21\_T3
2022-05-28\_T1
2022-09-17\_T3
2022-09-24\_T3
2022-10-22\_T3
2022-10-29\_T3
2022-11-05\_T3
2022-11-18\_T3
2022-11-26\_T3
2022-12-04\_T3
2022-12-31\_T3
2023-06-04\_T2
2023-08-21\_T2
2023-08-28\_T2
2023-09-04\_T2
2023-09-11\_T2
2023-09-18\_T2


125025813.0
157121.0
162859.0
149056.0
192438.0
239742.0
213604.0
508680.0
410124.0
444302.0
389236.0
386830.0
4782175.0
8676947.0
6441233.0
9079494.0
6434214.0
9810061.0
8345763.0
7348485.0
4472399.0
7373623.0
6029481.0
4782175.0
6099283.0
8663641.0
6646384.0
5777704.0
6622645.0
4386114.0


125022749.0
156942.0
162758.0
148985.0
192334.0
239479.0
213140.0
508468.0
409602.0
443806.0
389009.0
386709.0
4782164.0
8676947.0
6441233.0
9079494.0
6434211.0
9810032.0
8345739.0
7348464.0
4472390.0
7373619.0
6029467.0
4782164.0
6099212.0
8663617.0
6646343.0
5777699.0
6622636.0
4386086.0

1.0
1.0
1.0
1.0
1.0
1.0
1.0
1.0
1.0
1.0
1.0
1.0
1.0
1.0
1.0
1.0
1.0
1.0
1.0
1.0
1.0
1.0
1.0
1.0
1.0
1.0
1.0
1.0
1.0
1.0


124614653.0
156887.0
160017.0
148909.0
189336.0
238254.0
197178.0
505123.0
409548.0
443764.0
388236.0
361397.0
4782164.0
8674844.0
6441226.0
9079494.0
6434211.0
9809975.0
8345739.0
7348444.0
4273382.0
7373251.0
6029389.0
4782164.0
5945754.0
8663278.0
6646300.0
5777671.0
6622632.0
4386086.0

1.0
1.0
1.0
1.0
1.0
1.0
1.0
1.0
1.0
1.0
1.0
1.0
1.0
1.0
1.0
1.0
1.0
1.0
1.0
1.0
1.0
1.0
1.0
1.0
1.0
1.0
1.0
1.0
1.0
1.0


124457537.0
156375.0
159756.0
148725.0
189129.0
237942.0
197090.0
504276.0
409343.0
443550.0
387096.0
360198.0
4773958.0
8665736.0
6424827.0
9049889.0
6428025.0
9806141.0
8344983.0
7347654.0
4270715.0
7373190.0
6027587.0
4773958.0
5907292.0
8658731.0
6636365.0
5771738.0
6618952.0
4384316.0

1.0
1.0
1.0
1.0
1.0
1.0
1.0
1.0
1.0
1.0
1.0
1.0
1.0
1.0
1.0
1.0
1.0
1.0
1.0
1.0
1.0
1.0
1.0
1.0
1.0
1.0
1.0
1.0
1.0
1.0


34147368.0
84196.0
49957.0
18531.0
82153.0
63434.0
54536.0
325130.0
341819.0
258191.0
215912.0
155700.0
1544449.0
1608138.0
4164906.0
1140146.0
691648.0
2705000.0
1670496.0
2557448.0
1916508.0
831672.0
2173827.0
1544449.0
981169.0
2113789.0
1719502.0
1208379.0
2585441.0
1340842.0

1.0
1.0
1.0
1.0
1.0
1.0
1.0
1.0
1.0
1.0
1.0
1.0
1.0
1.0
1.0
1.0
1.0
1.0
1.0
1.0
1.0
1.0
1.0
1.0
1.0
1.0
1.0
1.0
1.0
1.0


336196.0
246.0
47.0
77.0
83.0
47.0
110.0
70731.0
81831.0
17186.0
3460.0
519.0
832.0
1515.0
3915.0
3.0
8.0
1784.0
18130.0
60379.0
37635.0
708.0
33628.0
832.0
119.0
115.0
630.0
99.0
1393.0
134.0

1.0
1.0
1.0
1.0
1.0
1.0
1.0
1.0
1.0
1.0
1.0
1.0
1.0
1.0
1.0
1.0
1.0
1.0
1.0
1.0
1.0
1.0
1.0
1.0
1.0
1.0
1.0
1.0
1.0
1.0


236725.0
148.0
15.0
3.0
3.0
4.0
0.0
69243.0
81522.0
15559.0
2747.0
130.0
794.0
0.0
61.0
3.0
0.0
512.0
7901.0
40339.0
16496.0
358.0
0.0
794.0
0.0
0.0
0.0
93.0
0.0
0.0

1.0
1.0
1.0
1.0
1.0
1.0
1.0
1.0
1.0
1.0
1.0
1.0
1.0
1.0
1.0
1.0
1.0
1.0
1.0
1.0
1.0
1.0
1.0
1.0
1.0
1.0
1.0
1.0
1.0
1.0


228329.0
120.0
7.0
0.0
0.0
2.0
0.0
69079.0
81205.0
15332.0
2693.0
116.0
794.0
0.0
0.0
0.0
0.0
449.0
3170.0
37941.0
16496.0
131.0
0.0
794.0
0.0
0.0
0.0
0.0
0.0
0.0

1.0
1.0
1.0
1.0
1.0
1.0
1.0
1.0
1.0
1.0
1.0
1.0
1.0
1.0
1.0
1.0
1.0
1.0
1.0
1.0
1.0
1.0
1.0
1.0
1.0
1.0
1.0
1.0
1.0
1.0


286.0
3.0
0.0
0.0
0.0
0.0
0.0
71.0
145.0
51.0
13.0
3.0
0.0
0.0
0.0
0.0
0.0
0.0
0.0
0.0
0.0
0.0
0.0
0.0
0.0
0.0
0.0
0.0
0.0
0.0

1.0
1.0
1.0
1.0
1.0
1.0
1.0
1.0
1.0
1.0
1.0
1.0
1.0
1.0
1.0
1.0
1.0
1.0
1.0
1.0
1.0
1.0
1.0
1.0
1.0
1.0
1.0
1.0
1.0
1.0


218.0
0.0
2.0
0.0
0.0
0.0
0.0
0.0
0.0
109.0
4.0
10.0
0.0
0.0
0.0
0.0
0.0
0.0
0.0
0.0
0.0
0.0
0.0
0.0
0.0
0.0
0.0
93.0
0.0
0.0

1.0
1.0
1.0
1.0
1.0
1.0
1.0
1.0
1.0
1.0
1.0
1.0
1.0
1.0
1.0
1.0
1.0
1.0
1.0
1.0
1.0
1.0
1.0
1.0
1.0
1.0
1.0
1.0
1.0
1.0


88.0
17.0
0.0
0.0
0.0
0.0
0.0
58.0
0.0
0.0
13.0
0.0
0.0
0.0
0.0
0.0
0.0
0.0
0.0
0.0
0.0
0.0
0.0
0.0
0.0
0.0
0.0
0.0
0.0
0.0

0.97
0.97
0.97
0.97
0.97
0.97
0.97
0.97
0.97
0.97
0.97
0.97
0.97
0.97
0.97
0.97
0.97
0.97
0.97
0.97
0.97
0.97
0.97
0.97
0.97
0.97
0.97
0.97
0.97
0.97


119.0
1.0
5.0
0.0
0.0
0.0
0.0
26.0
21.0
5.0
0.0
0.0
0.0
0.0
61.0
0.0
0.0
0.0
0.0
0.0
0.0
0.0
0.0
0.0
0.0
0.0
0.0
0.0
0.0
0.0

1.0
1.0
1.0
1.0
1.0
1.0
1.0
1.0
1.0
1.0
1.0
1.0
1.0
1.0
1.0
1.0
1.0
1.0
1.0
1.0
1.0
1.0
1.0
1.0
1.0
1.0
1.0
1.0
1.0
1.0


3543.0
38.0
2.0
8.0
0.0
0.0
0.0
593.0
5.0
1023.0
350.0
184.0
0.0
0.0
1340.0
0.0
0.0
0.0
0.0
0.0
0.0
0.0
0.0
0.0
0.0
0.0
0.0
0.0
0.0
0.0


3543.0
38.0
2.0
8.0
0.0
0.0
0.0
593.0
5.0
1023.0
350.0
184.0
0.0
0.0
1340.0
0.0
0.0
0.0
0.0
0.0
0.0
0.0
0.0
0.0
0.0
0.0
0.0
0.0
0.0
0.0

1.0
1.0
1.0
1.0
1.0
1.0
1.0
1.0
1.0
1.0
1.0
1.0
1.0
1.0
1.0
1.0
1.0
1.0
1.0
1.0
1.0
1.0
1.0
1.0
1.0
1.0
1.0
1.0
1.0
1.0


1020.0
31.0
20.0
60.0
14.0
18.0
90.0
0.0
0.0
0.0
0.0
0.0
0.0
0.0
0.0
0.0
8.0
391.0
0.0
36.0
3.0
349.0
0.0
0.0
0.0
0.0
0.0
0.0
0.0
0.0

1.0
1.0
1.0
1.0
1.0
1.0
1.0
1.0
1.0
1.0
1.0
1.0
1.0
1.0
1.0
1.0
1.0
1.0
1.0
1.0
1.0
1.0
1.0
1.0
1.0
1.0
1.0
1.0
1.0
1.0


609.0
0.0
1.0
0.0
0.0
3.0
2.0
190.0
123.0
162.0
54.0
1.0
0.0
0.0
0.0
0.0
0.0
0.0
0.0
63.0
10.0
0.0
0.0
0.0
0.0
0.0
0.0
0.0
0.0
0.0

0.99
0.99
0.99
0.99
0.99
0.99
0.99
0.99
0.99
0.99
0.99
0.99
0.99
0.99
0.99
0.99
0.99
0.99
0.99
0.99
0.99
0.99
0.99
0.99
0.99
0.99
0.99
0.99
0.99
0.99


136.0
0.0
0.0
0.0
0.0
3.0
2.0
48.0
0.0
82.0
0.0
1.0
0.0
0.0
0.0
0.0
0.0
0.0
0.0
0.0
0.0
0.0
0.0
0.0
0.0
0.0
0.0
0.0
0.0
0.0

1.0
1.0
1.0
1.0
1.0
1.0
1.0
1.0
1.0
1.0
1.0
1.0
1.0
1.0
1.0
1.0
1.0
1.0
1.0
1.0
1.0
1.0
1.0
1.0
1.0
1.0
1.0
1.0
1.0
1.0


25638.0
17.0
0.0
0.0
19.0
0.0
0.0
184.0
42.0
82.0
62.0
6.0
0.0
0.0
2514.0
0.0
0.0
22.0
80.0
15.0
20651.0
0.0
1944.0
0.0
0.0
0.0
0.0
0.0
0.0
0.0

0.93
0.93
0.93
0.93
0.93
0.93
0.93
0.93
0.93
0.93
0.93
0.93
0.93
0.93
0.93
0.93
0.93
0.93
0.93
0.93
0.93
0.93
0.93
0.93
0.93
0.93
0.93
0.93
0.93
0.93


229.0
0.0
0.0
0.0
0.0
0.0
0.0
184.0
29.0
9.0
7.0
0.0
0.0
0.0
0.0
0.0
0.0
0.0
0.0
0.0
0.0
0.0
0.0
0.0
0.0
0.0
0.0
0.0
0.0
0.0

1.0
1.0
1.0
1.0
1.0
1.0
1.0
1.0
1.0
1.0
1.0
1.0
1.0
1.0
1.0
1.0
1.0
1.0
1.0
1.0
1.0
1.0
1.0
1.0
1.0
1.0
1.0
1.0
1.0
1.0


2756.0
14.0
0.0
0.0
14.0
0.0
0.0
0.0
0.0
65.0
47.0
0.0
0.0
0.0
2514.0
0.0
0.0
22.0
80.0
0.0
0.0
0.0
0.0
0.0
0.0
0.0
0.0
0.0
0.0
0.0

1.0
1.0
1.0
1.0
1.0
1.0
1.0
1.0
1.0
1.0
1.0
1.0
1.0
1.0
1.0
1.0
1.0
1.0
1.0
1.0
1.0
1.0
1.0
1.0
1.0
1.0
1.0
1.0
1.0
1.0


58.0
3.0
0.0
0.0
5.0
0.0
0.0
0.0
13.0
8.0
8.0
6.0
0.0
0.0
0.0
0.0
0.0
0.0
0.0
15.0
0.0
0.0
0.0
0.0
0.0
0.0
0.0
0.0
0.0
0.0

1.0
1.0
1.0
1.0
1.0
1.0
1.0
1.0
1.0
1.0
1.0
1.0
1.0
1.0
1.0
1.0
1.0
1.0
1.0
1.0
1.0
1.0
1.0
1.0
1.0
1.0
1.0
1.0
1.0
1.0


155.0
12.0
5.0
0.0
0.0
0.0
0.0
0.0
0.0
0.0
32.0
80.0
0.0
0.0
0.0
0.0
0.0
0.0
8.0
2.0
16.0
0.0
0.0
0.0
0.0
0.0
0.0
0.0
0.0
0.0

1.0
1.0
1.0
1.0
1.0
1.0
1.0
1.0
1.0
1.0
1.0
1.0
1.0
1.0
1.0
1.0
1.0
1.0
1.0
1.0
1.0
1.0
1.0
1.0
1.0
1.0
1.0
1.0
1.0
1.0


517953.0
39969.0
4826.0
1.0
3.0
0.0
18101.0
1.0
23011.0
11587.0
33871.0
46784.0
0.0
0.0
0.0
0.0
0.0
0.0
0.0
0.0
0.0
0.0
0.0
0.0
24813.0
0.0
117892.0
197094.0
0.0
0.0


160001.0
39969.0
4774.0
1.0
3.0
0.0
0.0
1.0
23011.0
11587.0
33871.0
46784.0
0.0
0.0
0.0
0.0
0.0
0.0
0.0
0.0
0.0
0.0
0.0
0.0
0.0
0.0
0.0
0.0
0.0
0.0

0.75
0.75
0.75
0.75
0.75
0.75
0.75
0.75
0.75
0.75
0.75
0.75
0.75
0.75
0.75
0.75
0.75
0.75
0.75
0.75
0.75
0.75
0.75
0.75
0.75
0.75
0.75
0.75
0.75
0.75


125401.0
39969.0
4774.0
1.0
3.0
0.0
0.0
0.0
0.0
1.0
33870.0
46783.0
0.0
0.0
0.0
0.0
0.0
0.0
0.0
0.0
0.0
0.0
0.0
0.0
0.0
0.0
0.0
0.0
0.0
0.0

1.0
1.0
1.0
1.0
1.0
1.0
1.0
1.0
1.0
1.0
1.0
1.0
1.0
1.0
1.0
1.0
1.0
1.0
1.0
1.0
1.0
1.0
1.0
1.0
1.0
1.0
1.0
1.0
1.0
1.0


25311.0
0.0
0.0
0.0
0.0
0.0
0.0
0.0
16534.0
8775.0
1.0
1.0
0.0
0.0
0.0
0.0
0.0
0.0
0.0
0.0
0.0
0.0
0.0
0.0
0.0
0.0
0.0
0.0
0.0
0.0

0.99
0.99
0.99
0.99
0.99
0.99
0.99
0.99
0.99
0.99
0.99
0.99
0.99
0.99
0.99
0.99
0.99
0.99
0.99
0.99
0.99
0.99
0.99
0.99
0.99
0.99
0.99
0.99
0.99
0.99


333087.0
0.0
0.0
0.0
0.0
0.0
18101.0
0.0
0.0
0.0
0.0
0.0
0.0
0.0
0.0
0.0
0.0
0.0
0.0
0.0
0.0
0.0
0.0
0.0
0.0
0.0
117892.0
197094.0
0.0
0.0


333087.0
0.0
0.0
0.0
0.0
0.0
18101.0
0.0
0.0
0.0
0.0
0.0
0.0
0.0
0.0
0.0
0.0
0.0
0.0
0.0
0.0
0.0
0.0
0.0
0.0
0.0
117892.0
197094.0
0.0
0.0

1.0
1.0
1.0
1.0
1.0
1.0
1.0
1.0
1.0
1.0
1.0
1.0
1.0
1.0
1.0
1.0
1.0
1.0
1.0
1.0
1.0
1.0
1.0
1.0
1.0
1.0
1.0
1.0
1.0
1.0


24865.0
0.0
52.0
0.0
0.0
0.0
0.0
0.0
0.0
0.0
0.0
0.0
0.0
0.0
0.0
0.0
0.0
0.0
0.0
0.0
0.0
0.0
0.0
0.0
24813.0
0.0
0.0
0.0
0.0
0.0


52.0
0.0
52.0
0.0
0.0
0.0
0.0
0.0
0.0
0.0
0.0
0.0
0.0
0.0
0.0
0.0
0.0
0.0
0.0
0.0
0.0
0.0
0.0
0.0
0.0
0.0
0.0
0.0
0.0
0.0

0.99
0.99
0.99
0.99
0.99
0.99
0.99
0.99
0.99
0.99
0.99
0.99
0.99
0.99
0.99
0.99
0.99
0.99
0.99
0.99
0.99
0.99
0.99
0.99
0.99
0.99
0.99
0.99
0.99
0.99


24813.0
0.0
0.0
0.0
0.0
0.0
0.0
0.0
0.0
0.0
0.0
0.0
0.0
0.0
0.0
0.0
0.0
0.0
0.0
0.0
0.0
0.0
0.0
0.0
24813.0
0.0
0.0
0.0
0.0
0.0

0.84
0.84
0.84
0.84
0.84
0.84
0.84
0.84
0.84
0.84
0.84
0.84
0.84
0.84
0.84
0.84
0.84
0.84
0.84
0.84
0.84
0.84
0.84
0.84
0.84
0.84
0.84
0.84
0.84
0.84


5900557.0
3514.0
1540.0
9.0
2.0
3.0
1.0
82757.0
55861.0
32546.0
32970.0
3326.0
492008.0
0.0
22.0
337432.0
383610.0
153049.0
32689.0
253517.0
399266.0
215987.0
125650.0
492008.0
0.0
295577.0
781390.0
534430.0
810125.0
381268.0

0.93
0.93
0.93
0.93
0.93
0.93
0.93
0.93
0.93
0.93
0.93
0.93
0.93
0.93
0.93
0.93
0.93
0.93
0.93
0.93
0.93
0.93
0.93
0.93
0.93
0.93
0.93
0.93
0.93
0.93


4488506.0
894.0
831.0
3.0
0.0
0.0
0.0
50033.0
33815.0
10410.0
8874.0
239.0
34519.0
0.0
1.0
299658.0
366179.0
117513.0
26302.0
253517.0
225950.0
204385.0
124159.0
34519.0
0.0
295544.0
781177.0
502891.0
797036.0
320057.0

1.0
1.0
1.0
1.0
1.0
1.0
1.0
1.0
1.0
1.0
1.0
1.0
1.0
1.0
1.0
1.0
1.0
1.0
1.0
1.0
1.0
1.0
1.0
1.0
1.0
1.0
1.0
1.0
1.0
1.0


1964037.0
885.0
814.0
0.0
0.0
0.0
0.0
49901.0
33619.0
10116.0
8266.0
239.0
34519.0
0.0
1.0
70246.0
111920.0
41564.0
26300.0
124594.0
193902.0
178980.0
124158.0
34519.0
0.0
218980.0
328828.0
167703.0
68275.0
135708.0

1.0
1.0
1.0
1.0
1.0
1.0
1.0
1.0
1.0
1.0
1.0
1.0
1.0
1.0
1.0
1.0
1.0
1.0
1.0
1.0
1.0
1.0
1.0
1.0
1.0
1.0
1.0
1.0
1.0
1.0


625874.0
9.0
2.0
3.0
0.0
0.0
0.0
132.0
196.0
294.0
270.0
0.0
0.0
0.0
0.0
66692.0
127283.0
20292.0
0.0
0.0
0.0
25405.0
0.0
0.0
0.0
0.0
140261.0
27706.0
163596.0
53733.0

1.0
1.0
1.0
1.0
1.0
1.0
1.0
1.0
1.0
1.0
1.0
1.0
1.0
1.0
1.0
1.0
1.0
1.0
1.0
1.0
1.0
1.0
1.0
1.0
1.0
1.0
1.0
1.0
1.0
1.0


1895448.0
0.0
15.0
0.0
0.0
0.0
0.0
0.0
0.0
0.0
338.0
0.0
0.0
0.0
0.0
162521.0
126786.0
55539.0
2.0
128809.0
32015.0
0.0
1.0
0.0
0.0
76520.0
310524.0
307156.0
564866.0
130356.0

1.0
1.0
1.0
1.0
1.0
1.0
1.0
1.0
1.0
1.0
1.0
1.0
1.0
1.0
1.0
1.0
1.0
1.0
1.0
1.0
1.0
1.0
1.0
1.0
1.0
1.0
1.0
1.0
1.0
1.0


282769.0
978.0
467.0
3.0
2.0
3.0
1.0
24891.0
20445.0
13719.0
20150.0
2822.0
3244.0
0.0
0.0
0.0
0.0
0.0
6387.0
0.0
173316.0
11602.0
1491.0
3244.0
0.0
0.0
0.0
0.0
0.0
4.0

1.0
1.0
1.0
1.0
1.0
1.0
1.0
1.0
1.0
1.0
1.0
1.0
1.0
1.0
1.0
1.0
1.0
1.0
1.0
1.0
1.0
1.0
1.0
1.0
1.0
1.0
1.0
1.0
1.0
1.0


80694.0
978.0
467.0
3.0
2.0
3.0
1.0
20089.0
16559.0
11809.0
19817.0
2502.0
3244.0
0.0
0.0
0.0
0.0
0.0
481.0
0.0
0.0
0.0
1491.0
3244.0
0.0
0.0
0.0
0.0
0.0
4.0

1.0
1.0
1.0
1.0
1.0
1.0
1.0
1.0
1.0
1.0
1.0
1.0
1.0
1.0
1.0
1.0
1.0
1.0
1.0
1.0
1.0
1.0
1.0
1.0
1.0
1.0
1.0
1.0
1.0
1.0


928096.0
1545.0
211.0
0.0
0.0
0.0
0.0
0.0
0.0
6995.0
3497.0
228.0
359616.0
0.0
0.0
37763.0
17431.0
35525.0
0.0
0.0
0.0
0.0
0.0
359616.0
0.0
0.0
0.0
31387.0
13089.0
61193.0


928096.0
1545.0
211.0
0.0
0.0
0.0
0.0
0.0
0.0
6995.0
3497.0
228.0
359616.0
0.0
0.0
37763.0
17431.0
35525.0
0.0
0.0
0.0
0.0
0.0
359616.0
0.0
0.0
0.0
31387.0
13089.0
61193.0

1.0
1.0
1.0
1.0
1.0
1.0
1.0
1.0
1.0
1.0
1.0
1.0
1.0
1.0
1.0
1.0
1.0
1.0
1.0
1.0
1.0
1.0
1.0
1.0
1.0
1.0
1.0
1.0
1.0
1.0


10521.0
0.0
0.0
1.0
0.0
0.0
0.0
7394.0
1601.0
1126.0
399.0
0.0
0.0
0.0
0.0
0.0
0.0
0.0
0.0
0.0
0.0
0.0
0.0
0.0
0.0
0.0
0.0
0.0
0.0
0.0


10521.0
0.0
0.0
1.0
0.0
0.0
0.0
7394.0
1601.0
1126.0
399.0
0.0
0.0
0.0
0.0
0.0
0.0
0.0
0.0
0.0
0.0
0.0
0.0
0.0
0.0
0.0
0.0
0.0
0.0
0.0

1.0
1.0
1.0
1.0
1.0
1.0
1.0
1.0
1.0
1.0
1.0
1.0
1.0
1.0
1.0
1.0
1.0
1.0
1.0
1.0
1.0
1.0
1.0
1.0
1.0
1.0
1.0
1.0
1.0
1.0


481.0
42.0
0.0
0.0
0.0
0.0
0.0
439.0
0.0
0.0
0.0
0.0
0.0
0.0
0.0
0.0
0.0
0.0
0.0
0.0
0.0
0.0
0.0
0.0
0.0
0.0
0.0
0.0
0.0
0.0

0.93
0.93
0.93
0.93
0.93
0.93
0.93
0.93
0.93
0.93
0.93
0.93
0.93
0.93
0.93
0.93
0.93
0.93
0.93
0.93
0.93
0.93
0.93
0.93
0.93
0.93
0.93
0.93
0.93
0.93


42.0
42.0
0.0
0.0
0.0
0.0
0.0
0.0
0.0
0.0
0.0
0.0
0.0
0.0
0.0
0.0
0.0
0.0
0.0
0.0
0.0
0.0
0.0
0.0
0.0
0.0
0.0
0.0
0.0
0.0

0.91
0.91
0.91
0.91
0.91
0.91
0.91
0.91
0.91
0.91
0.91
0.91
0.91
0.91
0.91
0.91
0.91
0.91
0.91
0.91
0.91
0.91
0.91
0.91
0.91
0.91
0.91
0.91
0.91
0.91


195.0
0.0
0.0
0.0
0.0
0.0
0.0
0.0
0.0
195.0
0.0
0.0
0.0
0.0
0.0
0.0
0.0
0.0
0.0
0.0
0.0
0.0
0.0
0.0
0.0
0.0
0.0
0.0
0.0
0.0


195.0
0.0
0.0
0.0
0.0
0.0
0.0
0.0
0.0
195.0
0.0
0.0
0.0
0.0
0.0
0.0
0.0
0.0
0.0
0.0
0.0
0.0
0.0
0.0
0.0
0.0
0.0
0.0
0.0
0.0

1.0
1.0
1.0
1.0
1.0
1.0
1.0
1.0
1.0
1.0
1.0
1.0
1.0
1.0
1.0
1.0
1.0
1.0
1.0
1.0
1.0
1.0
1.0
1.0
1.0
1.0
1.0
1.0
1.0
1.0


189244.0
0.0
0.0
0.0
0.0
0.0
0.0
0.0
0.0
0.0
0.0
0.0
94622.0
0.0
0.0
0.0
0.0
0.0
0.0
0.0
0.0
0.0
0.0
94622.0
0.0
0.0
0.0
0.0
0.0
0.0


189244.0
0.0
0.0
0.0
0.0
0.0
0.0
0.0
0.0
0.0
0.0
0.0
94622.0
0.0
0.0
0.0
0.0
0.0
0.0
0.0
0.0
0.0
0.0
94622.0
0.0
0.0
0.0
0.0
0.0
0.0

0.88
0.88
0.88
0.88
0.88
0.88
0.88
0.88
0.88
0.88
0.88
0.88
0.88
0.88
0.88
0.88
0.88
0.88
0.88
0.88
0.88
0.88
0.88
0.88
0.88
0.88
0.88
0.88
0.88
0.88


6451473.0
3958.0
327.0
53.0
59.0
120.0
78.0
72182.0
72849.0
16982.0
17474.0
6374.0
659587.0
931.0
169874.0
55461.0
2939.0
211582.0
266596.0
77074.0
102847.0
0.0
691699.0
659587.0
26777.0
1152250.0
304688.0
208134.0
1270901.0
400090.0

0.98
0.98
0.98
0.98
0.98
0.98
0.98
0.98
0.98
0.98
0.98
0.98
0.98
0.98
0.98
0.98
0.98
0.98
0.98
0.98
0.98
0.98
0.98
0.98
0.98
0.98
0.98
0.98
0.98
0.98


130245.0
648.0
0.0
0.0
0.0
0.0
0.0
60169.0
60227.0
3892.0
2667.0
2642.0
0.0
0.0
0.0
0.0
0.0
0.0
0.0
0.0
0.0
0.0
0.0
0.0
0.0
0.0
0.0
0.0
0.0
0.0


121337.0
0.0
0.0
0.0
0.0
0.0
0.0
55103.0
57033.0
3892.0
2667.0
2642.0
0.0
0.0
0.0
0.0
0.0
0.0
0.0
0.0
0.0
0.0
0.0
0.0
0.0
0.0
0.0
0.0
0.0
0.0

1.0
1.0
1.0
1.0
1.0
1.0
1.0
1.0
1.0
1.0
1.0
1.0
1.0
1.0
1.0
1.0
1.0
1.0
1.0
1.0
1.0
1.0
1.0
1.0
1.0
1.0
1.0
1.0
1.0
1.0


8602.0
342.0
0.0
0.0
0.0
0.0
0.0
5066.0
3194.0
0.0
0.0
0.0
0.0
0.0
0.0
0.0
0.0
0.0
0.0
0.0
0.0
0.0
0.0
0.0
0.0
0.0
0.0
0.0
0.0
0.0

1.0
1.0
1.0
1.0
1.0
1.0
1.0
1.0
1.0
1.0
1.0
1.0
1.0
1.0
1.0
1.0
1.0
1.0
1.0
1.0
1.0
1.0
1.0
1.0
1.0
1.0
1.0
1.0
1.0
1.0


306.0
306.0
0.0
0.0
0.0
0.0
0.0
0.0
0.0
0.0
0.0
0.0
0.0
0.0
0.0
0.0
0.0
0.0
0.0
0.0
0.0
0.0
0.0
0.0
0.0
0.0
0.0
0.0
0.0
0.0

1.0
1.0
1.0
1.0
1.0
1.0
1.0
1.0
1.0
1.0
1.0
1.0
1.0
1.0
1.0
1.0
1.0
1.0
1.0
1.0
1.0
1.0
1.0
1.0
1.0
1.0
1.0
1.0
1.0
1.0


1375563.0
1720.0
0.0
0.0
0.0
0.0
0.0
9760.0
9758.0
2768.0
0.0
1304.0
114731.0
0.0
0.0
1.0
0.0
114920.0
0.0
0.0
0.0
0.0
691699.0
114731.0
0.0
298263.0
15908.0
0.0
0.0
0.0

0.99
0.99
0.99
0.99
0.99
0.99
0.99
0.99
0.99
0.99
0.99
0.99
0.99
0.99
0.99
0.99
0.99
0.99
0.99
0.99
0.99
0.99
0.99
0.99
0.99
0.99
0.99
0.99
0.99
0.99


1276241.0
156.0
0.0
0.0
0.0
0.0
0.0
9760.0
9758.0
2768.0
0.0
1304.0
92577.0
0.0
0.0
1.0
0.0
114920.0
0.0
0.0
0.0
0.0
638249.0
92577.0
0.0
298263.0
15908.0
0.0
0.0
0.0

1.0
1.0
1.0
1.0
1.0
1.0
1.0
1.0
1.0
1.0
1.0
1.0
1.0
1.0
1.0
1.0
1.0
1.0
1.0
1.0
1.0
1.0
1.0
1.0
1.0
1.0
1.0
1.0
1.0
1.0


93150.0
1564.0
0.0
0.0
0.0
0.0
0.0
0.0
0.0
0.0
0.0
0.0
20817.0
0.0
0.0
0.0
0.0
0.0
0.0
0.0
0.0
0.0
49952.0
20817.0
0.0
0.0
0.0
0.0
0.0
0.0

1.0
1.0
1.0
1.0
1.0
1.0
1.0
1.0
1.0
1.0
1.0
1.0
1.0
1.0
1.0
1.0
1.0
1.0
1.0
1.0
1.0
1.0
1.0
1.0
1.0
1.0
1.0
1.0
1.0
1.0


53661.0
872.0
95.0
0.0
0.0
0.0
0.0
0.0
1577.0
5816.0
10763.0
810.0
15943.0
0.0
1832.0
0.0
0.0
0.0
0.0
0.0
0.0
0.0
0.0
15943.0
0.0
10.0
0.0
0.0
0.0
0.0

0.98
0.98
0.98
0.98
0.98
0.98
0.98
0.98
0.98
0.98
0.98
0.98
0.98
0.98
0.98
0.98
0.98
0.98
0.98
0.98
0.98
0.98
0.98
0.98
0.98
0.98
0.98
0.98
0.98
0.98


202813.0
8.0
0.0
53.0
0.0
116.0
28.0
0.0
157.0
3288.0
1597.0
819.0
0.0
931.0
160978.0
0.0
0.0
0.0
0.0
0.0
0.0
0.0
0.0
0.0
26777.0
0.0
0.0
0.0
8061.0
0.0

0.7
0.7
0.7
0.7
0.7
0.7
0.7
0.7
0.7
0.7
0.7
0.7
0.7
0.7
0.7
0.7
0.7
0.7
0.7
0.7
0.7
0.7
0.7
0.7
0.7
0.7
0.7
0.7
0.7
0.7


17674.0
0.0
0.0
0.0
0.0
0.0
28.0
0.0
0.0
2221.0
254.0
124.0
0.0
0.0
4879.0
0.0
0.0
0.0
0.0
0.0
0.0
0.0
0.0
0.0
10168.0
0.0
0.0
0.0
0.0
0.0

1.0
1.0
1.0
1.0
1.0
1.0
1.0
1.0
1.0
1.0
1.0
1.0
1.0
1.0
1.0
1.0
1.0
1.0
1.0
1.0
1.0
1.0
1.0
1.0
1.0
1.0
1.0
1.0
1.0
1.0


164930.0
8.0
0.0
0.0
0.0
69.0
0.0
0.0
77.0
878.0
1084.0
660.0
0.0
745.0
144800.0
0.0
0.0
0.0
0.0
0.0
0.0
0.0
0.0
0.0
16609.0
0.0
0.0
0.0
0.0
0.0

1.0
1.0
1.0
1.0
1.0
1.0
1.0
1.0
1.0
1.0
1.0
1.0
1.0
1.0
1.0
1.0
1.0
1.0
1.0
1.0
1.0
1.0
1.0
1.0
1.0
1.0
1.0
1.0
1.0
1.0


475.0
0.0
0.0
0.0
0.0
0.0
0.0
0.0
80.0
136.0
259.0
0.0
0.0
0.0
0.0
0.0
0.0
0.0
0.0
0.0
0.0
0.0
0.0
0.0
0.0
0.0
0.0
0.0
0.0
0.0

1.0
1.0
1.0
1.0
1.0
1.0
1.0
1.0
1.0
1.0
1.0
1.0
1.0
1.0
1.0
1.0
1.0
1.0
1.0
1.0
1.0
1.0
1.0
1.0
1.0
1.0
1.0
1.0
1.0
1.0


19718.0
0.0
0.0
53.0
0.0
47.0
0.0
0.0
0.0
53.0
0.0
35.0
0.0
186.0
11283.0
0.0
0.0
0.0
0.0
0.0
0.0
0.0
0.0
0.0
0.0
0.0
0.0
0.0
8061.0
0.0

1.0
1.0
1.0
1.0
1.0
1.0
1.0
1.0
1.0
1.0
1.0
1.0
1.0
1.0
1.0
1.0
1.0
1.0
1.0
1.0
1.0
1.0
1.0
1.0
1.0
1.0
1.0
1.0
1.0
1.0


103971.0
230.0
73.0
0.0
0.0
0.0
0.0
1426.0
231.0
345.0
155.0
0.0
0.0
0.0
0.0
0.0
0.0
2.0
101501.0
8.0
0.0
0.0
0.0
0.0
0.0
0.0
0.0
0.0
0.0
0.0

0.94
0.94
0.94
0.94
0.94
0.94
0.94
0.94
0.94
0.94
0.94
0.94
0.94
0.94
0.94
0.94
0.94
0.94
0.94
0.94
0.94
0.94
0.94
0.94
0.94
0.94
0.94
0.94
0.94
0.94


103170.0
230.0
73.0
0.0
0.0
0.0
0.0
1426.0
231.0
345.0
155.0
0.0
0.0
0.0
0.0
0.0
0.0
2.0
100708.0
0.0
0.0
0.0
0.0
0.0
0.0
0.0
0.0
0.0
0.0
0.0

1.0
1.0
1.0
1.0
1.0
1.0
1.0
1.0
1.0
1.0
1.0
1.0
1.0
1.0
1.0
1.0
1.0
1.0
1.0
1.0
1.0
1.0
1.0
1.0
1.0
1.0
1.0
1.0
1.0
1.0


293752.0
97.0
29.0
0.0
0.0
0.0
0.0
821.0
0.0
335.0
164.0
0.0
94730.0
0.0
0.0
0.0
0.0
0.0
0.0
0.0
102846.0
0.0
0.0
94730.0
0.0
0.0
0.0
0.0
0.0
0.0


293752.0
97.0
29.0
0.0
0.0
0.0
0.0
821.0
0.0
335.0
164.0
0.0
94730.0
0.0
0.0
0.0
0.0
0.0
0.0
0.0
102846.0
0.0
0.0
94730.0
0.0
0.0
0.0
0.0
0.0
0.0

1.0
1.0
1.0
1.0
1.0
1.0
1.0
1.0
1.0
1.0
1.0
1.0
1.0
1.0
1.0
1.0
1.0
1.0
1.0
1.0
1.0
1.0
1.0
1.0
1.0
1.0
1.0
1.0
1.0
1.0


3487011.0
0.0
0.0
0.0
0.0
0.0
0.0
0.0
0.0
0.0
1161.0
0.0
434155.0
0.0
0.0
2.0
2.0
0.0
3.0
2.0
1.0
0.0
0.0
434155.0
0.0
710640.0
252169.0
137364.0
1205338.0
312019.0


3486618.0
0.0
0.0
0.0
0.0
0.0
0.0
0.0
0.0
0.0
768.0
0.0
434155.0
0.0
0.0
2.0
2.0
0.0
3.0
2.0
1.0
0.0
0.0
434155.0
0.0
710640.0
252169.0
137364.0
1205338.0
312019.0

1.0
1.0
1.0
1.0
1.0
1.0
1.0
1.0
1.0
1.0
1.0
1.0
1.0
1.0
1.0
1.0
1.0
1.0
1.0
1.0
1.0
1.0
1.0
1.0
1.0
1.0
1.0
1.0
1.0
1.0


393.0
0.0
0.0
0.0
0.0
0.0
0.0
0.0
0.0
0.0
393.0
0.0
0.0
0.0
0.0
0.0
0.0
0.0
0.0
0.0
0.0
0.0
0.0
0.0
0.0
0.0
0.0
0.0
0.0
0.0

1.0
1.0
1.0
1.0
1.0
1.0
1.0
1.0
1.0
1.0
1.0
1.0
1.0
1.0
1.0
1.0
1.0
1.0
1.0
1.0
1.0
1.0
1.0
1.0
1.0
1.0
1.0
1.0
1.0
1.0


63388.0
33.0
0.0
0.0
0.0
0.0
0.0
0.0
655.0
0.0
254.0
716.0
0.0
0.0
0.0
35684.0
0.0
15580.0
0.0
0.0
0.0
0.0
0.0
0.0
0.0
6190.0
0.0
0.0
4276.0
0.0

1.0
1.0
1.0
1.0
1.0
1.0
1.0
1.0
1.0
1.0
1.0
1.0
1.0
1.0
1.0
1.0
1.0
1.0
1.0
1.0
1.0
1.0
1.0
1.0
1.0
1.0
1.0
1.0
1.0
1.0


11154.0
33.0
0.0
0.0
0.0
0.0
0.0
0.0
655.0
0.0
0.0
0.0
0.0
0.0
0.0
0.0
0.0
0.0
0.0
0.0
0.0
0.0
0.0
0.0
0.0
6190.0
0.0
0.0
4276.0
0.0

1.0
1.0
1.0
1.0
1.0
1.0
1.0
1.0
1.0
1.0
1.0
1.0
1.0
1.0
1.0
1.0
1.0
1.0
1.0
1.0
1.0
1.0
1.0
1.0
1.0
1.0
1.0
1.0
1.0
1.0


901.0
0.0
0.0
0.0
0.0
0.0
0.0
0.0
0.0
0.0
219.0
682.0
0.0
0.0
0.0
0.0
0.0
0.0
0.0
0.0
0.0
0.0
0.0
0.0
0.0
0.0
0.0
0.0
0.0
0.0

1.0
1.0
1.0
1.0
1.0
1.0
1.0
1.0
1.0
1.0
1.0
1.0
1.0
1.0
1.0
1.0
1.0
1.0
1.0
1.0
1.0
1.0
1.0
1.0
1.0
1.0
1.0
1.0
1.0
1.0


51264.0
0.0
0.0
0.0
0.0
0.0
0.0
0.0
0.0
0.0
0.0
0.0
0.0
0.0
0.0
35684.0
0.0
15580.0
0.0
0.0
0.0
0.0
0.0
0.0
0.0
0.0
0.0
0.0
0.0
0.0

0.94
0.94
0.94
0.94
0.94
0.94
0.94
0.94
0.94
0.94
0.94
0.94
0.94
0.94
0.94
0.94
0.94
0.94
0.94
0.94
0.94
0.94
0.94
0.94
0.94
0.94
0.94
0.94
0.94
0.94


200215.0
51.0
82.0
0.0
0.0
0.0
1.0
0.0
238.0
250.0
233.0
23.0
0.0
0.0
0.0
0.0
0.0
47719.0
119019.0
25965.0
0.0
0.0
0.0
0.0
0.0
0.0
0.0
0.0
0.0
6634.0

1.0
1.0
1.0
1.0
1.0
1.0
1.0
1.0
1.0
1.0
1.0
1.0
1.0
1.0
1.0
1.0
1.0
1.0
1.0
1.0
1.0
1.0
1.0
1.0
1.0
1.0
1.0
1.0
1.0
1.0


62812.0
51.0
54.0
0.0
0.0
0.0
0.0
0.0
87.0
250.0
141.0
0.0
0.0
0.0
0.0
0.0
0.0
40711.0
0.0
14884.0
0.0
0.0
0.0
0.0
0.0
0.0
0.0
0.0
0.0
6634.0

1.0
1.0
1.0
1.0
1.0
1.0
1.0
1.0
1.0
1.0
1.0
1.0
1.0
1.0
1.0
1.0
1.0
1.0
1.0
1.0
1.0
1.0
1.0
1.0
1.0
1.0
1.0
1.0
1.0
1.0


119230.0
0.0
1.0
0.0
0.0
0.0
1.0
0.0
151.0
0.0
92.0
0.0
0.0
0.0
0.0
0.0
0.0
0.0
118985.0
0.0
0.0
0.0
0.0
0.0
0.0
0.0
0.0
0.0
0.0
0.0

0.97
0.97
0.97
0.97
0.97
0.97
0.97
0.97
0.97
0.97
0.97
0.97
0.97
0.97
0.97
0.97
0.97
0.97
0.97
0.97
0.97
0.97
0.97
0.97
0.97
0.97
0.97
0.97
0.97
0.97


199708.0
0.0
0.0
0.0
0.0
0.0
0.0
0.0
0.0
73.0
82.0
60.0
0.0
0.0
0.0
0.0
0.0
0.0
0.0
0.0
0.0
0.0
0.0
0.0
0.0
19455.0
33687.0
70678.0
33729.0
41944.0

0.99
0.99
0.99
0.99
0.99
0.99
0.99
0.99
0.99
0.99
0.99
0.99
0.99
0.99
0.99
0.99
0.99
0.99
0.99
0.99
0.99
0.99
0.99
0.99
0.99
0.99
0.99
0.99
0.99
0.99


33096.0
0.0
0.0
0.0
0.0
0.0
0.0
0.0
0.0
73.0
82.0
60.0
0.0
0.0
0.0
0.0
0.0
0.0
0.0
0.0
0.0
0.0
0.0
0.0
0.0
0.0
32881.0
0.0
0.0
0.0

1.0
1.0
1.0
1.0
1.0
1.0
1.0
1.0
1.0
1.0
1.0
1.0
1.0
1.0
1.0
1.0
1.0
1.0
1.0
1.0
1.0
1.0
1.0
1.0
1.0
1.0
1.0
1.0
1.0
1.0


123229.0
0.0
0.0
0.0
0.0
0.0
0.0
0.0
0.0
0.0
0.0
0.0
0.0
0.0
0.0
0.0
0.0
0.0
0.0
0.0
0.0
0.0
0.0
0.0
0.0
23.0
0.0
47533.0
33729.0
41944.0

1.0
1.0
1.0
1.0
1.0
1.0
1.0
1.0
1.0
1.0
1.0
1.0
1.0
1.0
1.0
1.0
1.0
1.0
1.0
1.0
1.0
1.0
1.0
1.0
1.0
1.0
1.0
1.0
1.0
1.0


23145.0
0.0
0.0
0.0
0.0
0.0
0.0
0.0
0.0
0.0
0.0
0.0
0.0
0.0
0.0
0.0
0.0
0.0
0.0
0.0
0.0
0.0
0.0
0.0
0.0
0.0
0.0
23145.0
0.0
0.0

0.97
0.97
0.97
0.97
0.97
0.97
0.97
0.97
0.97
0.97
0.97
0.97
0.97
0.97
0.97
0.97
0.97
0.97
0.97
0.97
0.97
0.97
0.97
0.97
0.97
0.97
0.97
0.97
0.97
0.97


140589.0
21.0
48.0
0.0
59.0
0.0
0.0
0.0
0.0
0.0
0.0
0.0
0.0
0.0
0.0
0.0
0.0
0.0
46073.0
0.0
0.0
0.0
0.0
0.0
0.0
58878.0
0.0
14.0
3237.0
32259.0

1.0
1.0
1.0
1.0
1.0
1.0
1.0
1.0
1.0
1.0
1.0
1.0
1.0
1.0
1.0
1.0
1.0
1.0
1.0
1.0
1.0
1.0
1.0
1.0
1.0
1.0
1.0
1.0
1.0
1.0


140429.0
0.0
0.0
0.0
0.0
0.0
0.0
0.0
0.0
0.0
0.0
0.0
0.0
0.0
0.0
0.0
0.0
0.0
46073.0
0.0
0.0
0.0
0.0
0.0
0.0
58878.0
0.0
0.0
3237.0
32241.0

0.86
0.86
0.86
0.86
0.86
0.86
0.86
0.86
0.86
0.86
0.86
0.86
0.86
0.86
0.86
0.86
0.86
0.86
0.86
0.86
0.86
0.86
0.86
0.86
0.86
0.86
0.86
0.86
0.86
0.86


126.0
0.0
0.0
0.0
0.0
0.0
0.0
0.0
0.0
0.0
126.0
0.0
0.0
0.0
0.0
0.0
0.0
0.0
0.0
0.0
0.0
0.0
0.0
0.0
0.0
0.0
0.0
0.0
0.0
0.0


126.0
0.0
0.0
0.0
0.0
0.0
0.0
0.0
0.0
0.0
126.0
0.0
0.0
0.0
0.0
0.0
0.0
0.0
0.0
0.0
0.0
0.0
0.0
0.0
0.0
0.0
0.0
0.0
0.0
0.0

1.0
1.0
1.0
1.0
1.0
1.0
1.0
1.0
1.0
1.0
1.0
1.0
1.0
1.0
1.0
1.0
1.0
1.0
1.0
1.0
1.0
1.0
1.0
1.0
1.0
1.0
1.0
1.0
1.0
1.0


40057.0
0.0
0.0
0.0
0.0
0.0
0.0
0.0
0.0
0.0
0.0
0.0
0.0
0.0
6960.0
0.0
0.0
33097.0
0.0
0.0
0.0
0.0
0.0
0.0
0.0
0.0
0.0
0.0
0.0
0.0

0.8
0.8
0.8
0.8
0.8
0.8
0.8
0.8
0.8
0.8
0.8
0.8
0.8
0.8
0.8
0.8
0.8
0.8
0.8
0.8
0.8
0.8
0.8
0.8
0.8
0.8
0.8
0.8
0.8
0.8


49.0
0.0
0.0
0.0
0.0
0.0
0.0
0.0
0.0
0.0
0.0
0.0
0.0
0.0
0.0
0.0
5.0
0.0
0.0
0.0
0.0
0.0
0.0
0.0
0.0
0.0
7.0
21.0
16.0
0.0

0.89
0.89
0.89
0.89
0.89
0.89
0.89
0.89
0.89
0.89
0.89
0.89
0.89
0.89
0.89
0.89
0.89
0.89
0.89
0.89
0.89
0.89
0.89
0.89
0.89
0.89
0.89
0.89
0.89
0.89


14.0
0.0
0.0
0.0
0.0
0.0
0.0
0.0
0.0
0.0
0.0
0.0
7.0
0.0
0.0
0.0
0.0
0.0
0.0
0.0
0.0
0.0
0.0
7.0
0.0
0.0
0.0
0.0
0.0
0.0

0.8
0.8
0.8
0.8
0.8
0.8
0.8
0.8
0.8
0.8
0.8
0.8
0.8
0.8
0.8
0.8
0.8
0.8
0.8
0.8
0.8
0.8
0.8
0.8
0.8
0.8
0.8
0.8
0.8
0.8


17.0
0.0
0.0
0.0
0.0
0.0
0.0
0.0
0.0
0.0
0.0
0.0
0.0
0.0
0.0
0.0
0.0
0.0
0.0
0.0
0.0
0.0
0.0
0.0
0.0
0.0
17.0
0.0
0.0
0.0

1.0
1.0
1.0
1.0
1.0
1.0
1.0
1.0
1.0
1.0
1.0
1.0
1.0
1.0
1.0
1.0
1.0
1.0
1.0
1.0
1.0
1.0
1.0
1.0
1.0
1.0
1.0
1.0
1.0
1.0


3587858.0
3248.0
3153.0
384.0
1757.0
3578.0
370.0
649.0
23742.0
22504.0
12744.0
19768.0
33748.0
108145.0
94965.0
12098.0
134119.0
1475567.0
639657.0
483142.0
161099.0
75448.0
0.0
33748.0
34120.0
26412.0
70913.0
26142.0
71113.0
15525.0

1.0
1.0
1.0
1.0
1.0
1.0
1.0
1.0
1.0
1.0
1.0
1.0
1.0
1.0
1.0
1.0
1.0
1.0
1.0
1.0
1.0
1.0
1.0
1.0
1.0
1.0
1.0
1.0
1.0
1.0


349973.0
1190.0
593.0
0.0
0.0
0.0
0.0
0.0
18767.0
14881.0
9100.0
17741.0
0.0
0.0
0.0
0.0
0.0
164808.0
0.0
0.0
122893.0
0.0
0.0
0.0
0.0
0.0
0.0
0.0
0.0
0.0

1.0
1.0
1.0
1.0
1.0
1.0
1.0
1.0
1.0
1.0
1.0
1.0
1.0
1.0
1.0
1.0
1.0
1.0
1.0
1.0
1.0
1.0
1.0
1.0
1.0
1.0
1.0
1.0
1.0
1.0


2912.0
0.0
0.0
0.0
0.0
0.0
0.0
0.0
0.0
955.0
0.0
1957.0
0.0
0.0
0.0
0.0
0.0
0.0
0.0
0.0
0.0
0.0
0.0
0.0
0.0
0.0
0.0
0.0
0.0
0.0

1.0
1.0
1.0
1.0
1.0
1.0
1.0
1.0
1.0
1.0
1.0
1.0
1.0
1.0
1.0
1.0
1.0
1.0
1.0
1.0
1.0
1.0
1.0
1.0
1.0
1.0
1.0
1.0
1.0
1.0


34603.0
322.0
773.0
0.0
0.0
0.0
0.0
649.0
2127.0
1631.0
1107.0
941.0
0.0
0.0
0.0
0.0
0.0
0.0
4906.0
2054.0
0.0
20093.0
0.0
0.0
0.0
0.0
0.0
0.0
0.0
0.0

1.0
1.0
1.0
1.0
1.0
1.0
1.0
1.0
1.0
1.0
1.0
1.0
1.0
1.0
1.0
1.0
1.0
1.0
1.0
1.0
1.0
1.0
1.0
1.0
1.0
1.0
1.0
1.0
1.0
1.0


4939.0
0.0
0.0
0.0
0.0
0.0
0.0
0.0
1487.0
3452.0
0.0
0.0
0.0
0.0
0.0
0.0
0.0
0.0
0.0
0.0
0.0
0.0
0.0
0.0
0.0
0.0
0.0
0.0
0.0
0.0


4939.0
0.0
0.0
0.0
0.0
0.0
0.0
0.0
1487.0
3452.0
0.0
0.0
0.0
0.0
0.0
0.0
0.0
0.0
0.0
0.0
0.0
0.0
0.0
0.0
0.0
0.0
0.0
0.0
0.0
0.0

1.0
1.0
1.0
1.0
1.0
1.0
1.0
1.0
1.0
1.0
1.0
1.0
1.0
1.0
1.0
1.0
1.0
1.0
1.0
1.0
1.0
1.0
1.0
1.0
1.0
1.0
1.0
1.0
1.0
1.0


254302.0
0.0
664.0
39.0
0.0
3222.0
0.0
0.0
0.0
0.0
0.0
38.0
0.0
108145.0
0.0
0.0
126806.0
0.0
0.0
0.0
0.0
0.0
0.0
0.0
0.0
0.0
0.0
0.0
0.0
15388.0


111990.0
0.0
584.0
39.0
0.0
3222.0
0.0
0.0
0.0
0.0
0.0
0.0
0.0
108145.0
0.0
0.0
0.0
0.0
0.0
0.0
0.0
0.0
0.0
0.0
0.0
0.0
0.0
0.0
0.0
0.0

0.98
0.98
0.98
0.98
0.98
0.98
0.98
0.98
0.98
0.98
0.98
0.98
0.98
0.98
0.98
0.98
0.98
0.98
0.98
0.98
0.98
0.98
0.98
0.98
0.98
0.98
0.98
0.98
0.98
0.98


118.0
0.0
80.0
0.0
0.0
0.0
0.0
0.0
0.0
0.0
0.0
38.0
0.0
0.0
0.0
0.0
0.0
0.0
0.0
0.0
0.0
0.0
0.0
0.0
0.0
0.0
0.0
0.0
0.0
0.0

1.0
1.0
1.0
1.0
1.0
1.0
1.0
1.0
1.0
1.0
1.0
1.0
1.0
1.0
1.0
1.0
1.0
1.0
1.0
1.0
1.0
1.0
1.0
1.0
1.0
1.0
1.0
1.0
1.0
1.0


142194.0
0.0
0.0
0.0
0.0
0.0
0.0
0.0
0.0
0.0
0.0
0.0
0.0
0.0
0.0
0.0
126806.0
0.0
0.0
0.0
0.0
0.0
0.0
0.0
0.0
0.0
0.0
0.0
0.0
15388.0

1.0
1.0
1.0
1.0
1.0
1.0
1.0
1.0
1.0
1.0
1.0
1.0
1.0
1.0
1.0
1.0
1.0
1.0
1.0
1.0
1.0
1.0
1.0
1.0
1.0
1.0
1.0
1.0
1.0
1.0


2392.0
125.0
0.0
0.0
0.0
0.0
0.0
0.0
1143.0
480.0
644.0
0.0
0.0
0.0
0.0
0.0
0.0
0.0
0.0
0.0
0.0
0.0
0.0
0.0
0.0
0.0
0.0
0.0
0.0
0.0


2392.0
125.0
0.0
0.0
0.0
0.0
0.0
0.0
1143.0
480.0
644.0
0.0
0.0
0.0
0.0
0.0
0.0
0.0
0.0
0.0
0.0
0.0
0.0
0.0
0.0
0.0
0.0
0.0
0.0
0.0

1.0
1.0
1.0
1.0
1.0
1.0
1.0
1.0
1.0
1.0
1.0
1.0
1.0
1.0
1.0
1.0
1.0
1.0
1.0
1.0
1.0
1.0
1.0
1.0
1.0
1.0
1.0
1.0
1.0
1.0


1136.0
0.0
67.0
44.0
1025.0
0.0
0.0
0.0
0.0
0.0
0.0
0.0
0.0
0.0
0.0
0.0
0.0
0.0
0.0
0.0
0.0
0.0
0.0
0.0
0.0
0.0
0.0
0.0
0.0
0.0


1136.0
0.0
67.0
44.0
1025.0
0.0
0.0
0.0
0.0
0.0
0.0
0.0
0.0
0.0
0.0
0.0
0.0
0.0
0.0
0.0
0.0
0.0
0.0
0.0
0.0
0.0
0.0
0.0
0.0
0.0

1.0
1.0
1.0
1.0
1.0
1.0
1.0
1.0
1.0
1.0
1.0
1.0
1.0
1.0
1.0
1.0
1.0
1.0
1.0
1.0
1.0
1.0
1.0
1.0
1.0
1.0
1.0
1.0
1.0
1.0


752.0
0.0
151.0
0.0
0.0
0.0
0.0
0.0
213.0
388.0
0.0
0.0
0.0
0.0
0.0
0.0
0.0
0.0
0.0
0.0
0.0
0.0
0.0
0.0
0.0
0.0
0.0
0.0
0.0
0.0

1.0
1.0
1.0
1.0
1.0
1.0
1.0
1.0
1.0
1.0
1.0
1.0
1.0
1.0
1.0
1.0
1.0
1.0
1.0
1.0
1.0
1.0
1.0
1.0
1.0
1.0
1.0
1.0
1.0
1.0


1234.0
321.0
112.0
0.0
0.0
0.0
0.0
0.0
0.0
50.0
751.0
0.0
0.0
0.0
0.0
0.0
0.0
0.0
0.0
0.0
0.0
0.0
0.0
0.0
0.0
0.0
0.0
0.0
0.0
0.0

1.0
1.0
1.0
1.0
1.0
1.0
1.0
1.0
1.0
1.0
1.0
1.0
1.0
1.0
1.0
1.0
1.0
1.0
1.0
1.0
1.0
1.0
1.0
1.0
1.0
1.0
1.0
1.0
1.0
1.0


1363.0
211.0
57.0
0.0
0.0
0.0
0.0
0.0
0.0
831.0
264.0
0.0
0.0
0.0
0.0
0.0
0.0
0.0
0.0
0.0
0.0
0.0
0.0
0.0
0.0
0.0
0.0
0.0
0.0
0.0


1363.0
211.0
57.0
0.0
0.0
0.0
0.0
0.0
0.0
831.0
264.0
0.0
0.0
0.0
0.0
0.0
0.0
0.0
0.0
0.0
0.0
0.0
0.0
0.0
0.0
0.0
0.0
0.0
0.0
0.0

1.0
1.0
1.0
1.0
1.0
1.0
1.0
1.0
1.0
1.0
1.0
1.0
1.0
1.0
1.0
1.0
1.0
1.0
1.0
1.0
1.0
1.0
1.0
1.0
1.0
1.0
1.0
1.0
1.0
1.0


21897.0
0.0
0.0
0.0
153.0
299.0
119.0
0.0
0.0
0.0
565.0
64.0
0.0
0.0
0.0
0.0
0.0
0.0
0.0
0.0
0.0
0.0
0.0
0.0
0.0
8621.0
12076.0
0.0
0.0
0.0

1.0
1.0
1.0
1.0
1.0
1.0
1.0
1.0
1.0
1.0
1.0
1.0
1.0
1.0
1.0
1.0
1.0
1.0
1.0
1.0
1.0
1.0
1.0
1.0
1.0
1.0
1.0
1.0
1.0
1.0


744.0
0.0
65.0
0.0
0.0
0.0
0.0
0.0
0.0
271.0
0.0
267.0
0.0
0.0
0.0
0.0
0.0
0.0
0.0
0.0
0.0
0.0
0.0
0.0
141.0
0.0
0.0
0.0
0.0
0.0


744.0
0.0
65.0
0.0
0.0
0.0
0.0
0.0
0.0
271.0
0.0
267.0
0.0
0.0
0.0
0.0
0.0
0.0
0.0
0.0
0.0
0.0
0.0
0.0
141.0
0.0
0.0
0.0
0.0
0.0

1.0
1.0
1.0
1.0
1.0
1.0
1.0
1.0
1.0
1.0
1.0
1.0
1.0
1.0
1.0
1.0
1.0
1.0
1.0
1.0
1.0
1.0
1.0
1.0
1.0
1.0
1.0
1.0
1.0
1.0


920.0
244.0
0.0
0.0
0.0
0.0
0.0
0.0
0.0
382.0
0.0
294.0
0.0
0.0
0.0
0.0
0.0
0.0
0.0
0.0
0.0
0.0
0.0
0.0
0.0
0.0
0.0
0.0
0.0
0.0


920.0
244.0
0.0
0.0
0.0
0.0
0.0
0.0
0.0
382.0
0.0
294.0
0.0
0.0
0.0
0.0
0.0
0.0
0.0
0.0
0.0
0.0
0.0
0.0
0.0
0.0
0.0
0.0
0.0
0.0

1.0
1.0
1.0
1.0
1.0
1.0
1.0
1.0
1.0
1.0
1.0
1.0
1.0
1.0
1.0
1.0
1.0
1.0
1.0
1.0
1.0
1.0
1.0
1.0
1.0
1.0
1.0
1.0
1.0
1.0


82188.0
159.0
285.0
234.0
66.0
57.0
72.0
0.0
0.0
0.0
0.0
247.0
33748.0
0.0
6485.0
7087.0
0.0
0.0
0.0
0.0
0.0
0.0
0.0
33748.0
0.0
0.0
0.0
0.0
0.0
0.0

1.0
1.0
1.0
1.0
1.0
1.0
1.0
1.0
1.0
1.0
1.0
1.0
1.0
1.0
1.0
1.0
1.0
1.0
1.0
1.0
1.0
1.0
1.0
1.0
1.0
1.0
1.0
1.0
1.0
1.0


569.0
0.0
182.0
140.0
0.0
0.0
0.0
0.0
0.0
0.0
0.0
247.0
0.0
0.0
0.0
0.0
0.0
0.0
0.0
0.0
0.0
0.0
0.0
0.0
0.0
0.0
0.0
0.0
0.0
0.0

0.93
0.93
0.93
0.93
0.93
0.93
0.93
0.93
0.93
0.93
0.93
0.93
0.93
0.93
0.93
0.93
0.93
0.93
0.93
0.93
0.93
0.93
0.93
0.93
0.93
0.93
0.93
0.93
0.93
0.93


340.0
340.0
0.0
0.0
0.0
0.0
0.0
0.0
0.0
0.0
0.0
0.0
0.0
0.0
0.0
0.0
0.0
0.0
0.0
0.0
0.0
0.0
0.0
0.0
0.0
0.0
0.0
0.0
0.0
0.0


340.0
340.0
0.0
0.0
0.0
0.0
0.0
0.0
0.0
0.0
0.0
0.0
0.0
0.0
0.0
0.0
0.0
0.0
0.0
0.0
0.0
0.0
0.0
0.0
0.0
0.0
0.0
0.0
0.0
0.0

1.0
1.0
1.0
1.0
1.0
1.0
1.0
1.0
1.0
1.0
1.0
1.0
1.0
1.0
1.0
1.0
1.0
1.0
1.0
1.0
1.0
1.0
1.0
1.0
1.0
1.0
1.0
1.0
1.0
1.0


160580.0
0.0
15.0
0.0
189.0
0.0
179.0
0.0
0.0
0.0
0.0
0.0
0.0
0.0
0.0
5006.0
0.0
148482.0
0.0
0.0
0.0
0.0
0.0
0.0
0.0
6709.0
0.0
0.0
0.0
0.0

1.0
1.0
1.0
1.0
1.0
1.0
1.0
1.0
1.0
1.0
1.0
1.0
1.0
1.0
1.0
1.0
1.0
1.0
1.0
1.0
1.0
1.0
1.0
1.0
1.0
1.0
1.0
1.0
1.0
1.0


148497.0
0.0
15.0
0.0
0.0
0.0
0.0
0.0
0.0
0.0
0.0
0.0
0.0
0.0
0.0
0.0
0.0
148482.0
0.0
0.0
0.0
0.0
0.0
0.0
0.0
0.0
0.0
0.0
0.0
0.0

1.0
1.0
1.0
1.0
1.0
1.0
1.0
1.0
1.0
1.0
1.0
1.0
1.0
1.0
1.0
1.0
1.0
1.0
1.0
1.0
1.0
1.0
1.0
1.0
1.0
1.0
1.0
1.0
1.0
1.0


145864.0
103.0
25.0
0.0
47.0
0.0
0.0
0.0
0.0
33.0
100.0
0.0
0.0
0.0
0.0
0.0
0.0
42736.0
0.0
0.0
0.0
0.0
0.0
0.0
31586.0
0.0
0.0
0.0
71098.0
136.0

1.0
1.0
1.0
1.0
1.0
1.0
1.0
1.0
1.0
1.0
1.0
1.0
1.0
1.0
1.0
1.0
1.0
1.0
1.0
1.0
1.0
1.0
1.0
1.0
1.0
1.0
1.0
1.0
1.0
1.0


261.0
103.0
25.0
0.0
0.0
0.0
0.0
0.0
0.0
33.0
100.0
0.0
0.0
0.0
0.0
0.0
0.0
0.0
0.0
0.0
0.0
0.0
0.0
0.0
0.0
0.0
0.0
0.0
0.0
0.0

0.99
0.99
0.99
0.99
0.99
0.99
0.99
0.99
0.99
0.99
0.99
0.99
0.99
0.99
0.99
0.99
0.99
0.99
0.99
0.99
0.99
0.99
0.99
0.99
0.99
0.99
0.99
0.99
0.99
0.99


144906.0
0.0
0.0
0.0
47.0
0.0
0.0
0.0
0.0
0.0
0.0
0.0
0.0
0.0
0.0
0.0
0.0
42736.0
0.0
0.0
0.0
0.0
0.0
0.0
31335.0
0.0
0.0
0.0
70653.0
135.0

0.99
0.99
0.99
0.99
0.99
0.99
0.99
0.99
0.99
0.99
0.99
0.99
0.99
0.99
0.99
0.99
0.99
0.99
0.99
0.99
0.99
0.99
0.99
0.99
0.99
0.99
0.99
0.99
0.99
0.99


148204.0
0.0
22.0
0.0
277.0
0.0
0.0
0.0
0.0
0.0
0.0
0.0
0.0
0.0
0.0
0.0
7313.0
24160.0
127.0
116305.0
0.0
0.0
0.0
0.0
0.0
0.0
0.0
0.0
0.0
0.0


148204.0
0.0
22.0
0.0
277.0
0.0
0.0
0.0
0.0
0.0
0.0
0.0
0.0
0.0
0.0
0.0
7313.0
24160.0
127.0
116305.0
0.0
0.0
0.0
0.0
0.0
0.0
0.0
0.0
0.0
0.0

1.0
1.0
1.0
1.0
1.0
1.0
1.0
1.0
1.0
1.0
1.0
1.0
1.0
1.0
1.0
1.0
1.0
1.0
1.0
1.0
1.0
1.0
1.0
1.0
1.0
1.0
1.0
1.0
1.0
1.0


2608.0
42.0
91.0
8.0
0.0
0.0
0.0
0.0
0.0
0.0
0.0
82.0
0.0
0.0
0.0
0.0
0.0
0.0
0.0
0.0
0.0
0.0
0.0
0.0
2385.0
0.0
0.0
0.0
0.0
0.0


2608.0
42.0
91.0
8.0
0.0
0.0
0.0
0.0
0.0
0.0
0.0
82.0
0.0
0.0
0.0
0.0
0.0
0.0
0.0
0.0
0.0
0.0
0.0
0.0
2385.0
0.0
0.0
0.0
0.0
0.0

1.0
1.0
1.0
1.0
1.0
1.0
1.0
1.0
1.0
1.0
1.0
1.0
1.0
1.0
1.0
1.0
1.0
1.0
1.0
1.0
1.0
1.0
1.0
1.0
1.0
1.0
1.0
1.0
1.0
1.0


164147.0
0.0
6.0
0.0
0.0
0.0
0.0
0.0
0.0
0.0
119.0
0.0
0.0
0.0
0.0
0.0
0.0
0.0
25296.0
46697.0
36674.0
55355.0
0.0
0.0
0.0
0.0
0.0
0.0
0.0
0.0

1.0
1.0
1.0
1.0
1.0
1.0
1.0
1.0
1.0
1.0
1.0
1.0
1.0
1.0
1.0
1.0
1.0
1.0
1.0
1.0
1.0
1.0
1.0
1.0
1.0
1.0
1.0
1.0
1.0
1.0


161137.0
0.0
6.0
0.0
0.0
0.0
0.0
0.0
0.0
0.0
119.0
0.0
0.0
0.0
0.0
0.0
0.0
0.0
24722.0
45900.0
35857.0
54533.0
0.0
0.0
0.0
0.0
0.0
0.0
0.0
0.0

1.0
1.0
1.0
1.0
1.0
1.0
1.0
1.0
1.0
1.0
1.0
1.0
1.0
1.0
1.0
1.0
1.0
1.0
1.0
1.0
1.0
1.0
1.0
1.0
1.0
1.0
1.0
1.0
1.0
1.0


105.0
0.0
0.0
0.0
0.0
0.0
0.0
0.0
0.0
105.0
0.0
0.0
0.0
0.0
0.0
0.0
0.0
0.0
0.0
0.0
0.0
0.0
0.0
0.0
0.0
0.0
0.0
0.0
0.0
0.0


105.0
0.0
0.0
0.0
0.0
0.0
0.0
0.0
0.0
105.0
0.0
0.0
0.0
0.0
0.0
0.0
0.0
0.0
0.0
0.0
0.0
0.0
0.0
0.0
0.0
0.0
0.0
0.0
0.0
0.0

0.99
0.99
0.99
0.99
0.99
0.99
0.99
0.99
0.99
0.99
0.99
0.99
0.99
0.99
0.99
0.99
0.99
0.99
0.99
0.99
0.99
0.99
0.99
0.99
0.99
0.99
0.99
0.99
0.99
0.99


94.0
0.0
0.0
0.0
0.0
0.0
0.0
0.0
0.0
0.0
94.0
0.0
0.0
0.0
0.0
0.0
0.0
0.0
0.0
0.0
0.0
0.0
0.0
0.0
0.0
0.0
0.0
0.0
0.0
0.0


94.0
0.0
0.0
0.0
0.0
0.0
0.0
0.0
0.0
0.0
94.0
0.0
0.0
0.0
0.0
0.0
0.0
0.0
0.0
0.0
0.0
0.0
0.0
0.0
0.0
0.0
0.0
0.0
0.0
0.0

1.0
1.0
1.0
1.0
1.0
1.0
1.0
1.0
1.0
1.0
1.0
1.0
1.0
1.0
1.0
1.0
1.0
1.0
1.0
1.0
1.0
1.0
1.0
1.0
1.0
1.0
1.0
1.0
1.0
1.0


136.0
78.0
58.0
0.0
0.0
0.0
0.0
0.0
0.0
0.0
0.0
0.0
0.0
0.0
0.0
0.0
0.0
0.0
0.0
0.0
0.0
0.0
0.0
0.0
0.0
0.0
0.0
0.0
0.0
0.0

0.95
0.95
0.95
0.95
0.95
0.95
0.95
0.95
0.95
0.95
0.95
0.95
0.95
0.95
0.95
0.95
0.95
0.95
0.95
0.95
0.95
0.95
0.95
0.95
0.95
0.95
0.95
0.95
0.95
0.95


58.0
0.0
58.0
0.0
0.0
0.0
0.0
0.0
0.0
0.0
0.0
0.0
0.0
0.0
0.0
0.0
0.0
0.0
0.0
0.0
0.0
0.0
0.0
0.0
0.0
0.0
0.0
0.0
0.0
0.0

0.89
0.89
0.89
0.89
0.89
0.89
0.89
0.89
0.89
0.89
0.89
0.89
0.89
0.89
0.89
0.89
0.89
0.89
0.89
0.89
0.89
0.89
0.89
0.89
0.89
0.89
0.89
0.89
0.89
0.89


92.0
32.0
60.0
0.0
0.0
0.0
0.0
0.0
0.0
0.0
0.0
0.0
0.0
0.0
0.0
0.0
0.0
0.0
0.0
0.0
0.0
0.0
0.0
0.0
0.0
0.0
0.0
0.0
0.0
0.0


73.0
32.0
41.0
0.0
0.0
0.0
0.0
0.0
0.0
0.0
0.0
0.0
0.0
0.0
0.0
0.0
0.0
0.0
0.0
0.0
0.0
0.0
0.0
0.0
0.0
0.0
0.0
0.0
0.0
0.0

1.0
1.0
1.0
1.0
1.0
1.0
1.0
1.0
1.0
1.0
1.0
1.0
1.0
1.0
1.0
1.0
1.0
1.0
1.0
1.0
1.0
1.0
1.0
1.0
1.0
1.0
1.0
1.0
1.0
1.0


19.0
0.0
19.0
0.0
0.0
0.0
0.0
0.0
0.0
0.0
0.0
0.0
0.0
0.0
0.0
0.0
0.0
0.0
0.0
0.0
0.0
0.0
0.0
0.0
0.0
0.0
0.0
0.0
0.0
0.0

1.0
1.0
1.0
1.0
1.0
1.0
1.0
1.0
1.0
1.0
1.0
1.0
1.0
1.0
1.0
1.0
1.0
1.0
1.0
1.0
1.0
1.0
1.0
1.0
1.0
1.0
1.0
1.0
1.0
1.0


61.0
0.0
61.0
0.0
0.0
0.0
0.0
0.0
0.0
0.0
0.0
0.0
0.0
0.0
0.0
0.0
0.0
0.0
0.0
0.0
0.0
0.0
0.0
0.0
0.0
0.0
0.0
0.0
0.0
0.0

1.0
1.0
1.0
1.0
1.0
1.0
1.0
1.0
1.0
1.0
1.0
1.0
1.0
1.0
1.0
1.0
1.0
1.0
1.0
1.0
1.0
1.0
1.0
1.0
1.0
1.0
1.0
1.0
1.0
1.0


88503.0
0.0
13.0
27.0
0.0
0.0
0.0
0.0
5.0
0.0
0.0
0.0
0.0
0.0
88458.0
0.0
0.0
0.0
0.0
0.0
0.0
0.0
0.0
0.0
0.0
0.0
0.0
0.0
0.0
0.0


88503.0
0.0
13.0
27.0
0.0
0.0
0.0
0.0
5.0
0.0
0.0
0.0
0.0
0.0
88458.0
0.0
0.0
0.0
0.0
0.0
0.0
0.0
0.0
0.0
0.0
0.0
0.0
0.0
0.0
0.0

1.0
1.0
1.0
1.0
1.0
1.0
1.0
1.0
1.0
1.0
1.0
1.0
1.0
1.0
1.0
1.0
1.0
1.0
1.0
1.0
1.0
1.0
1.0
1.0
1.0
1.0
1.0
1.0
1.0
1.0


1704075.0
0.0
0.0
0.0
0.0
0.0
0.0
0.0
0.0
0.0
0.0
0.0
0.0
0.0
0.0
1.0
0.0
780863.0
606336.0
316871.0
0.0
0.0
0.0
0.0
0.0
0.0
1.0
0.0
2.0
1.0

0.75
0.75
0.75
0.75
0.75
0.75
0.75
0.75
0.75
0.75
0.75
0.75
0.75
0.75
0.75
0.75
0.75
0.75
0.75
0.75
0.75
0.75
0.75
0.75
0.75
0.75
0.75
0.75
0.75
0.75


69631.0
0.0
0.0
0.0
0.0
0.0
0.0
0.0
0.0
0.0
0.0
0.0
0.0
0.0
0.0
0.0
0.0
69631.0
0.0
0.0
0.0
0.0
0.0
0.0
0.0
0.0
0.0
0.0
0.0
0.0

0.96
0.96
0.96
0.96
0.96
0.96
0.96
0.96
0.96
0.96
0.96
0.96
0.96
0.96
0.96
0.96
0.96
0.96
0.96
0.96
0.96
0.96
0.96
0.96
0.96
0.96
0.96
0.96
0.96
0.96


69627.0
0.0
0.0
0.0
0.0
0.0
0.0
0.0
0.0
0.0
0.0
0.0
0.0
0.0
0.0
0.0
0.0
69627.0
0.0
0.0
0.0
0.0
0.0
0.0
0.0
0.0
0.0
0.0
0.0
0.0

1.0
1.0
1.0
1.0
1.0
1.0
1.0
1.0
1.0
1.0
1.0
1.0
1.0
1.0
1.0
1.0
1.0
1.0
1.0
1.0
1.0
1.0
1.0
1.0
1.0
1.0
1.0
1.0
1.0
1.0


26142.0
0.0
0.0
0.0
0.0
0.0
0.0
0.0
0.0
0.0
0.0
0.0
0.0
0.0
0.0
0.0
0.0
0.0
0.0
0.0
0.0
0.0
0.0
0.0
0.0
0.0
0.0
26142.0
0.0
0.0


26142.0
0.0
0.0
0.0
0.0
0.0
0.0
0.0
0.0
0.0
0.0
0.0
0.0
0.0
0.0
0.0
0.0
0.0
0.0
0.0
0.0
0.0
0.0
0.0
0.0
0.0
0.0
26142.0
0.0
0.0

1.0
1.0
1.0
1.0
1.0
1.0
1.0
1.0
1.0
1.0
1.0
1.0
1.0
1.0
1.0
1.0
1.0
1.0
1.0
1.0
1.0
1.0
1.0
1.0
1.0
1.0
1.0
1.0
1.0
1.0


2082.0
0.0
0.0
0.0
0.0
0.0
0.0
0.0
0.0
0.0
0.0
0.0
0.0
0.0
0.0
0.0
0.0
556.0
0.0
0.0
1526.0
0.0
0.0
0.0
0.0
0.0
0.0
0.0
0.0
0.0


2082.0
0.0
0.0
0.0
0.0
0.0
0.0
0.0
0.0
0.0
0.0
0.0
0.0
0.0
0.0
0.0
0.0
556.0
0.0
0.0
1526.0
0.0
0.0
0.0
0.0
0.0
0.0
0.0
0.0
0.0

1.0
1.0
1.0
1.0
1.0
1.0
1.0
1.0
1.0
1.0
1.0
1.0
1.0
1.0
1.0
1.0
1.0
1.0
1.0
1.0
1.0
1.0
1.0
1.0
1.0
1.0
1.0
1.0
1.0
1.0


22.0
0.0
0.0
0.0
0.0
0.0
0.0
0.0
0.0
0.0
0.0
0.0
0.0
0.0
22.0
0.0
0.0
0.0
0.0
0.0
0.0
0.0
0.0
0.0
0.0
0.0
0.0
0.0
0.0
0.0

0.7
0.7
0.7
0.7
0.7
0.7
0.7
0.7
0.7
0.7
0.7
0.7
0.7
0.7
0.7
0.7
0.7
0.7
0.7
0.7
0.7
0.7
0.7
0.7
0.7
0.7
0.7
0.7
0.7
0.7


2040598.0
2073.0
13182.0
9206.0
50038.0
37732.0
19594.0
0.0
931.0
3511.0
957.0
172.0
0.0
0.0
1743.0
419031.0
109493.0
111647.0
0.0
221274.0
124097.0
178623.0
222756.0
0.0
64917.0
103943.0
108864.0
15251.0
218967.0
2596.0

0.92
0.92
0.92
0.92
0.92
0.92
0.92
0.92
0.92
0.92
0.92
0.92
0.92
0.92
0.92
0.92
0.92
0.92
0.92
0.92
0.92
0.92
0.92
0.92
0.92
0.92
0.92
0.92
0.92
0.92


1712518.0
1897.0
13153.0
9206.0
50038.0
37233.0
18635.0
0.0
931.0
2466.0
141.0
172.0
0.0
0.0
1.0
363565.0
109246.0
111647.0
0.0
221274.0
124097.0
178623.0
222756.0
0.0
64917.0
53782.0
92046.0
14124.0
22568.0
0.0

1.0
1.0
1.0
1.0
1.0
1.0
1.0
1.0
1.0
1.0
1.0
1.0
1.0
1.0
1.0
1.0
1.0
1.0
1.0
1.0
1.0
1.0
1.0
1.0
1.0
1.0
1.0
1.0
1.0
1.0


454347.0
0.0
5774.0
4288.0
40041.0
31123.0
11675.0
0.0
0.0
1228.0
0.0
172.0
0.0
0.0
0.0
235979.0
61865.0
0.0
0.0
0.0
0.0
0.0
0.0
0.0
6842.0
48.0
55299.0
13.0
0.0
0.0

1.0
1.0
1.0
1.0
1.0
1.0
1.0
1.0
1.0
1.0
1.0
1.0
1.0
1.0
1.0
1.0
1.0
1.0
1.0
1.0
1.0
1.0
1.0
1.0
1.0
1.0
1.0
1.0
1.0
1.0


305307.0
1125.0
6397.0
3686.0
8385.0
4951.0
4878.0
0.0
0.0
0.0
0.0
0.0
0.0
0.0
0.0
44602.0
42409.0
0.0
0.0
0.0
124090.0
0.0
0.0
0.0
57446.0
1880.0
0.0
5458.0
0.0
0.0

1.0
1.0
1.0
1.0
1.0
1.0
1.0
1.0
1.0
1.0
1.0
1.0
1.0
1.0
1.0
1.0
1.0
1.0
1.0
1.0
1.0
1.0
1.0
1.0
1.0
1.0
1.0
1.0
1.0
1.0


4183.0
0.0
625.0
607.0
634.0
886.0
1431.0
0.0
0.0
0.0
0.0
0.0
0.0
0.0
0.0
0.0
0.0
0.0
0.0
0.0
0.0
0.0
0.0
0.0
0.0
0.0
0.0
0.0
0.0
0.0

0.94
0.94
0.94
0.94
0.94
0.94
0.94
0.94
0.94
0.94
0.94
0.94
0.94
0.94
0.94
0.94
0.94
0.94
0.94
0.94
0.94
0.94
0.94
0.94
0.94
0.94
0.94
0.94
0.94
0.94


2369.0
535.0
0.0
404.0
358.0
0.0
0.0
0.0
931.0
0.0
141.0
0.0
0.0
0.0
0.0
0.0
0.0
0.0
0.0
0.0
0.0
0.0
0.0
0.0
0.0
0.0
0.0
0.0
0.0
0.0

1.0
1.0
1.0
1.0
1.0
1.0
1.0
1.0
1.0
1.0
1.0
1.0
1.0
1.0
1.0
1.0
1.0
1.0
1.0
1.0
1.0
1.0
1.0
1.0
1.0
1.0
1.0
1.0
1.0
1.0


1426.0
0.0
35.0
174.0
420.0
101.0
130.0
0.0
0.0
566.0
0.0
0.0
0.0
0.0
0.0
0.0
0.0
0.0
0.0
0.0
0.0
0.0
0.0
0.0
0.0
0.0
0.0
0.0
0.0
0.0

1.0
1.0
1.0
1.0
1.0
1.0
1.0
1.0
1.0
1.0
1.0
1.0
1.0
1.0
1.0
1.0
1.0
1.0
1.0
1.0
1.0
1.0
1.0
1.0
1.0
1.0
1.0
1.0
1.0
1.0


333.0
30.0
0.0
0.0
0.0
0.0
0.0
0.0
0.0
303.0
0.0
0.0
0.0
0.0
0.0
0.0
0.0
0.0
0.0
0.0
0.0
0.0
0.0
0.0
0.0
0.0
0.0
0.0
0.0
0.0

1.0
1.0
1.0
1.0
1.0
1.0
1.0
1.0
1.0
1.0
1.0
1.0
1.0
1.0
1.0
1.0
1.0
1.0
1.0
1.0
1.0
1.0
1.0
1.0
1.0
1.0
1.0
1.0
1.0
1.0


92270.0
0.0
0.0
0.0
0.0
0.0
0.0
0.0
0.0
0.0
0.0
0.0
0.0
0.0
0.0
0.0
0.0
0.0
0.0
0.0
0.0
0.0
0.0
0.0
0.0
51497.0
27923.0
8203.0
4647.0
0.0

1.0
1.0
1.0
1.0
1.0
1.0
1.0
1.0
1.0
1.0
1.0
1.0
1.0
1.0
1.0
1.0
1.0
1.0
1.0
1.0
1.0
1.0
1.0
1.0
1.0
1.0
1.0
1.0
1.0
1.0


1328.0
0.0
0.0
0.0
0.0
0.0
0.0
0.0
0.0
0.0
0.0
0.0
0.0
0.0
0.0
0.0
0.0
0.0
0.0
1328.0
0.0
0.0
0.0
0.0
0.0
0.0
0.0
0.0
0.0
0.0

0.89
0.89
0.89
0.89
0.89
0.89
0.89
0.89
0.89
0.89
0.89
0.89
0.89
0.89
0.89
0.89
0.89
0.89
0.89
0.89
0.89
0.89
0.89
0.89
0.89
0.89
0.89
0.89
0.89
0.89


1137.0
0.0
0.0
0.0
0.0
0.0
0.0
0.0
0.0
0.0
0.0
0.0
0.0
0.0
0.0
0.0
0.0
284.0
0.0
0.0
0.0
270.0
583.0
0.0
0.0
0.0
0.0
0.0
0.0
0.0

0.97
0.97
0.97
0.97
0.97
0.97
0.97
0.97
0.97
0.97
0.97
0.97
0.97
0.97
0.97
0.97
0.97
0.97
0.97
0.97
0.97
0.97
0.97
0.97
0.97
0.97
0.97
0.97
0.97
0.97


314374.0
160.0
0.0
0.0
0.0
0.0
800.0
0.0
0.0
1045.0
816.0
0.0
0.0
0.0
0.0
55439.0
209.0
0.0
0.0
0.0
0.0
0.0
0.0
0.0
0.0
50161.0
9345.0
0.0
196399.0
0.0

1.0
1.0
1.0
1.0
1.0
1.0
1.0
1.0
1.0
1.0
1.0
1.0
1.0
1.0
1.0
1.0
1.0
1.0
1.0
1.0
1.0
1.0
1.0
1.0
1.0
1.0
1.0
1.0
1.0
1.0


1613.0
0.0
0.0
0.0
0.0
499.0
0.0
0.0
0.0
0.0
0.0
0.0
0.0
0.0
0.0
0.0
0.0
0.0
0.0
0.0
0.0
0.0
0.0
0.0
0.0
0.0
0.0
1114.0
0.0
0.0


1613.0
0.0
0.0
0.0
0.0
499.0
0.0
0.0
0.0
0.0
0.0
0.0
0.0
0.0
0.0
0.0
0.0
0.0
0.0
0.0
0.0
0.0
0.0
0.0
0.0
0.0
0.0
1114.0
0.0
0.0

1.0
1.0
1.0
1.0
1.0
1.0
1.0
1.0
1.0
1.0
1.0
1.0
1.0
1.0
1.0
1.0
1.0
1.0
1.0
1.0
1.0
1.0
1.0
1.0
1.0
1.0
1.0
1.0
1.0
1.0


4532.0
16.0
29.0
0.0
0.0
0.0
149.0
0.0
0.0
0.0
0.0
0.0
0.0
0.0
1742.0
0.0
0.0
0.0
0.0
0.0
0.0
0.0
0.0
0.0
0.0
0.0
0.0
0.0
0.0
2596.0


4532.0
16.0
29.0
0.0
0.0
0.0
149.0
0.0
0.0
0.0
0.0
0.0
0.0
0.0
1742.0
0.0
0.0
0.0
0.0
0.0
0.0
0.0
0.0
0.0
0.0
0.0
0.0
0.0
0.0
2596.0

1.0
1.0
1.0
1.0
1.0
1.0
1.0
1.0
1.0
1.0
1.0
1.0
1.0
1.0
1.0
1.0
1.0
1.0
1.0
1.0
1.0
1.0
1.0
1.0
1.0
1.0
1.0
1.0
1.0
1.0


23.0
0.0
0.0
0.0
0.0
0.0
10.0
0.0
0.0
0.0
0.0
0.0
0.0
0.0
0.0
0.0
0.0
0.0
0.0
0.0
0.0
0.0
0.0
0.0
0.0
0.0
0.0
13.0
0.0
0.0

0.82
0.82
0.82
0.82
0.82
0.82
0.82
0.82
0.82
0.82
0.82
0.82
0.82
0.82
0.82
0.82
0.82
0.82
0.82
0.82
0.82
0.82
0.82
0.82
0.82
0.82
0.82
0.82
0.82
0.82


10.0
0.0
0.0
0.0
0.0
0.0
10.0
0.0
0.0
0.0
0.0
0.0
0.0
0.0
0.0
0.0
0.0
0.0
0.0
0.0
0.0
0.0
0.0
0.0
0.0
0.0
0.0
0.0
0.0
0.0

1.0
1.0
1.0
1.0
1.0
1.0
1.0
1.0
1.0
1.0
1.0
1.0
1.0
1.0
1.0
1.0
1.0
1.0
1.0
1.0
1.0
1.0
1.0
1.0
1.0
1.0
1.0
1.0
1.0
1.0


7473.0
0.0
0.0
0.0
0.0
0.0
0.0
0.0
0.0
0.0
0.0
0.0
0.0
0.0
0.0
0.0
0.0
0.0
0.0
0.0
0.0
0.0
0.0
0.0
0.0
0.0
7473.0
0.0
0.0
0.0

1.0
1.0
1.0
1.0
1.0
1.0
1.0
1.0
1.0
1.0
1.0
1.0
1.0
1.0
1.0
1.0
1.0
1.0
1.0
1.0
1.0
1.0
1.0
1.0
1.0
1.0
1.0
1.0
1.0
1.0


6029.0
0.0
0.0
0.0
0.0
0.0
0.0
0.0
0.0
0.0
0.0
0.0
0.0
0.0
0.0
0.0
0.0
0.0
0.0
0.0
0.0
0.0
0.0
0.0
0.0
0.0
6029.0
0.0
0.0
0.0

1.0
1.0
1.0
1.0
1.0
1.0
1.0
1.0
1.0
1.0
1.0
1.0
1.0
1.0
1.0
1.0
1.0
1.0
1.0
1.0
1.0
1.0
1.0
1.0
1.0
1.0
1.0
1.0
1.0
1.0


212781.0
2054.0
1497.0
0.0
0.0
2054.0
90.0
12014.0
2996.0
18885.0
7164.0
1065.0
11476.0
0.0
702.0
0.0
0.0
0.0
6816.0
1274.0
0.0
0.0
0.0
11476.0
0.0
64850.0
40221.0
0.0
1064.0
27083.0

1.0
1.0
1.0
1.0
1.0
1.0
1.0
1.0
1.0
1.0
1.0
1.0
1.0
1.0
1.0
1.0
1.0
1.0
1.0
1.0
1.0
1.0
1.0
1.0
1.0
1.0
1.0
1.0
1.0
1.0


199567.0
1798.0
1287.0
0.0
0.0
1993.0
90.0
12014.0
2908.0
18710.0
7115.0
1030.0
11476.0
0.0
0.0
0.0
0.0
0.0
0.0
0.0
0.0
0.0
0.0
11476.0
0.0
64850.0
40221.0
0.0
1064.0
23535.0

1.0
1.0
1.0
1.0
1.0
1.0
1.0
1.0
1.0
1.0
1.0
1.0
1.0
1.0
1.0
1.0
1.0
1.0
1.0
1.0
1.0
1.0
1.0
1.0
1.0
1.0
1.0
1.0
1.0
1.0


196299.0
1788.0
1274.0
0.0
0.0
1983.0
90.0
11967.0
2896.0
18642.0
7078.0
1026.0
11166.0
0.0
0.0
0.0
0.0
0.0
0.0
0.0
0.0
0.0
0.0
11166.0
0.0
63966.0
39056.0
0.0
1019.0
23182.0

1.0
1.0
1.0
1.0
1.0
1.0
1.0
1.0
1.0
1.0
1.0
1.0
1.0
1.0
1.0
1.0
1.0
1.0
1.0
1.0
1.0
1.0
1.0
1.0
1.0
1.0
1.0
1.0
1.0
1.0


87.0
3.0
8.0
0.0
0.0
5.0
0.0
13.0
10.0
28.0
19.0
1.0
0.0
0.0
0.0
0.0
0.0
0.0
0.0
0.0
0.0
0.0
0.0
0.0
0.0
0.0
0.0
0.0
0.0
0.0

0.71
0.71
0.71
0.71
0.71
0.71
0.71
0.71
0.71
0.71
0.71
0.71
0.71
0.71
0.71
0.71
0.71
0.71
0.71
0.71
0.71
0.71
0.71
0.71
0.71
0.71
0.71
0.71
0.71
0.71


244.0
25.0
95.0
0.0
0.0
0.0
0.0
0.0
0.0
73.0
0.0
0.0
0.0
0.0
51.0
0.0
0.0
0.0
0.0
0.0
0.0
0.0
0.0
0.0
0.0
0.0
0.0
0.0
0.0
0.0


244.0
25.0
95.0
0.0
0.0
0.0
0.0
0.0
0.0
73.0
0.0
0.0
0.0
0.0
51.0
0.0
0.0
0.0
0.0
0.0
0.0
0.0
0.0
0.0
0.0
0.0
0.0
0.0
0.0
0.0

0.99
0.99
0.99
0.99
0.99
0.99
0.99
0.99
0.99
0.99
0.99
0.99
0.99
0.99
0.99
0.99
0.99
0.99
0.99
0.99
0.99
0.99
0.99
0.99
0.99
0.99
0.99
0.99
0.99
0.99


6999.0
16.0
0.0
0.0
0.0
0.0
0.0
0.0
88.0
30.0
49.0
0.0
0.0
0.0
0.0
0.0
0.0
0.0
6816.0
0.0
0.0
0.0
0.0
0.0
0.0
0.0
0.0
0.0
0.0
0.0

0.8
0.8
0.8
0.8
0.8
0.8
0.8
0.8
0.8
0.8
0.8
0.8
0.8
0.8
0.8
0.8
0.8
0.8
0.8
0.8
0.8
0.8
0.8
0.8
0.8
0.8
0.8
0.8
0.8
0.8


16.0
16.0
0.0
0.0
0.0
0.0
0.0
0.0
0.0
0.0
0.0
0.0
0.0
0.0
0.0
0.0
0.0
0.0
0.0
0.0
0.0
0.0
0.0
0.0
0.0
0.0
0.0
0.0
0.0
0.0

1.0
1.0
1.0
1.0
1.0
1.0
1.0
1.0
1.0
1.0
1.0
1.0
1.0
1.0
1.0
1.0
1.0
1.0
1.0
1.0
1.0
1.0
1.0
1.0
1.0
1.0
1.0
1.0
1.0
1.0


8.0
0.0
0.0
0.0
0.0
0.0
0.0
0.0
0.0
8.0
0.0
0.0
0.0
0.0
0.0
0.0
0.0
0.0
0.0
0.0
0.0
0.0
0.0
0.0
0.0
0.0
0.0
0.0
0.0
0.0

1.0
1.0
1.0
1.0
1.0
1.0
1.0
1.0
1.0
1.0
1.0
1.0
1.0
1.0
1.0
1.0
1.0
1.0
1.0
1.0
1.0
1.0
1.0
1.0
1.0
1.0
1.0
1.0
1.0
1.0


113.0
110.0
3.0
0.0
0.0
0.0
0.0
0.0
0.0
0.0
0.0
0.0
0.0
0.0
0.0
0.0
0.0
0.0
0.0
0.0
0.0
0.0
0.0
0.0
0.0
0.0
0.0
0.0
0.0
0.0


113.0
110.0
3.0
0.0
0.0
0.0
0.0
0.0
0.0
0.0
0.0
0.0
0.0
0.0
0.0
0.0
0.0
0.0
0.0
0.0
0.0
0.0
0.0
0.0
0.0
0.0
0.0
0.0
0.0
0.0

0.89
0.89
0.89
0.89
0.89
0.89
0.89
0.89
0.89
0.89
0.89
0.89
0.89
0.89
0.89
0.89
0.89
0.89
0.89
0.89
0.89
0.89
0.89
0.89
0.89
0.89
0.89
0.89
0.89
0.89


3655.0
27.0
19.0
0.0
0.0
61.0
0.0
0.0
0.0
0.0
0.0
0.0
0.0
0.0
0.0
0.0
0.0
0.0
0.0
0.0
0.0
0.0
0.0
0.0
0.0
0.0
0.0
0.0
0.0
3548.0

1.0
1.0
1.0
1.0
1.0
1.0
1.0
1.0
1.0
1.0
1.0
1.0
1.0
1.0
1.0
1.0
1.0
1.0
1.0
1.0
1.0
1.0
1.0
1.0
1.0
1.0
1.0
1.0
1.0
1.0


3407.0
27.0
19.0
0.0
0.0
61.0
0.0
0.0
0.0
0.0
0.0
0.0
0.0
0.0
0.0
0.0
0.0
0.0
0.0
0.0
0.0
0.0
0.0
0.0
0.0
0.0
0.0
0.0
0.0
3300.0

1.0
1.0
1.0
1.0
1.0
1.0
1.0
1.0
1.0
1.0
1.0
1.0
1.0
1.0
1.0
1.0
1.0
1.0
1.0
1.0
1.0
1.0
1.0
1.0
1.0
1.0
1.0
1.0
1.0
1.0


105.0
0.0
0.0
0.0
0.0
0.0
0.0
0.0
0.0
0.0
0.0
0.0
0.0
0.0
0.0
0.0
0.0
0.0
0.0
0.0
0.0
0.0
0.0
0.0
0.0
0.0
0.0
0.0
0.0
105.0

0.98
0.98
0.98
0.98
0.98
0.98
0.98
0.98
0.98
0.98
0.98
0.98
0.98
0.98
0.98
0.98
0.98
0.98
0.98
0.98
0.98
0.98
0.98
0.98
0.98
0.98
0.98
0.98
0.98
0.98


107.0
0.0
0.0
0.0
0.0
0.0
0.0
0.0
0.0
72.0
0.0
35.0
0.0
0.0
0.0
0.0
0.0
0.0
0.0
0.0
0.0
0.0
0.0
0.0
0.0
0.0
0.0
0.0
0.0
0.0


107.0
0.0
0.0
0.0
0.0
0.0
0.0
0.0
0.0
72.0
0.0
35.0
0.0
0.0
0.0
0.0
0.0
0.0
0.0
0.0
0.0
0.0
0.0
0.0
0.0
0.0
0.0
0.0
0.0
0.0

0.92
0.92
0.92
0.92
0.92
0.92
0.92
0.92
0.92
0.92
0.92
0.92
0.92
0.92
0.92
0.92
0.92
0.92
0.92
0.92
0.92
0.92
0.92
0.92
0.92
0.92
0.92
0.92
0.92
0.92


78.0
78.0
0.0
0.0
0.0
0.0
0.0
0.0
0.0
0.0
0.0
0.0
0.0
0.0
0.0
0.0
0.0
0.0
0.0
0.0
0.0
0.0
0.0
0.0
0.0
0.0
0.0
0.0
0.0
0.0


78.0
78.0
0.0
0.0
0.0
0.0
0.0
0.0
0.0
0.0
0.0
0.0
0.0
0.0
0.0
0.0
0.0
0.0
0.0
0.0
0.0
0.0
0.0
0.0
0.0
0.0
0.0
0.0
0.0
0.0

0.96
0.96
0.96
0.96
0.96
0.96
0.96
0.96
0.96
0.96
0.96
0.96
0.96
0.96
0.96
0.96
0.96
0.96
0.96
0.96
0.96
0.96
0.96
0.96
0.96
0.96
0.96
0.96
0.96
0.96


68.0
0.0
68.0
0.0
0.0
0.0
0.0
0.0
0.0
0.0
0.0
0.0
0.0
0.0
0.0
0.0
0.0
0.0
0.0
0.0
0.0
0.0
0.0
0.0
0.0
0.0
0.0
0.0
0.0
0.0

1.0
1.0
1.0
1.0
1.0
1.0
1.0
1.0
1.0
1.0
1.0
1.0
1.0
1.0
1.0
1.0
1.0
1.0
1.0
1.0
1.0
1.0
1.0
1.0
1.0
1.0
1.0
1.0
1.0
1.0


1299.0
0.0
25.0
0.0
0.0
0.0
0.0
0.0
0.0
0.0
0.0
0.0
0.0
0.0
0.0
0.0
0.0
0.0
0.0
1274.0
0.0
0.0
0.0
0.0
0.0
0.0
0.0
0.0
0.0
0.0


1299.0
0.0
25.0
0.0
0.0
0.0
0.0
0.0
0.0
0.0
0.0
0.0
0.0
0.0
0.0
0.0
0.0
0.0
0.0
1274.0
0.0
0.0
0.0
0.0
0.0
0.0
0.0
0.0
0.0
0.0

1.0
1.0
1.0
1.0
1.0
1.0
1.0
1.0
1.0
1.0
1.0
1.0
1.0
1.0
1.0
1.0
1.0
1.0
1.0
1.0
1.0
1.0
1.0
1.0
1.0
1.0
1.0
1.0
1.0
1.0


239.0
0.0
0.0
0.0
0.0
0.0
0.0
0.0
0.0
0.0
0.0
0.0
0.0
0.0
239.0
0.0
0.0
0.0
0.0
0.0
0.0
0.0
0.0
0.0
0.0
0.0
0.0
0.0
0.0
0.0


239.0
0.0
0.0
0.0
0.0
0.0
0.0
0.0
0.0
0.0
0.0
0.0
0.0
0.0
239.0
0.0
0.0
0.0
0.0
0.0
0.0
0.0
0.0
0.0
0.0
0.0
0.0
0.0
0.0
0.0

1.0
1.0
1.0
1.0
1.0
1.0
1.0
1.0
1.0
1.0
1.0
1.0
1.0
1.0
1.0
1.0
1.0
1.0
1.0
1.0
1.0
1.0
1.0
1.0
1.0
1.0
1.0
1.0
1.0
1.0


32903.0
323.0
1568.0
1.0
0.0
0.0
0.0
0.0
0.0
0.0
6258.0
24753.0
0.0
0.0
0.0
0.0
0.0
0.0
0.0
0.0
0.0
0.0
0.0
0.0
0.0
0.0
0.0
0.0
0.0
0.0

0.78
0.78
0.78
0.78
0.78
0.78
0.78
0.78
0.78
0.78
0.78
0.78
0.78
0.78
0.78
0.78
0.78
0.78
0.78
0.78
0.78
0.78
0.78
0.78
0.78
0.78
0.78
0.78
0.78
0.78


142641.0
109.0
21.0
0.0
0.0
4.0
12.0
10241.0
7555.0
7856.0
2141.0
271.0
4787.0
0.0
22639.0
0.0
870.0
4710.0
11477.0
1928.0
0.0
0.0
0.0
4787.0
6.0
32587.0
20529.0
498.0
23.0
9590.0

0.91
0.91
0.91
0.91
0.91
0.91
0.91
0.91
0.91
0.91
0.91
0.91
0.91
0.91
0.91
0.91
0.91
0.91
0.91
0.91
0.91
0.91
0.91
0.91
0.91
0.91
0.91
0.91
0.91
0.91


138976.0
108.0
21.0
0.0
0.0
4.0
12.0
9475.0
6691.0
7333.0
1714.0
271.0
4787.0
0.0
22592.0
0.0
870.0
4710.0
11477.0
1928.0
0.0
0.0
0.0
4787.0
6.0
32066.0
20260.0
388.0
0.0
9476.0

1.0
1.0
1.0
1.0
1.0
1.0
1.0
1.0
1.0
1.0
1.0
1.0
1.0
1.0
1.0
1.0
1.0
1.0
1.0
1.0
1.0
1.0
1.0
1.0
1.0
1.0
1.0
1.0
1.0
1.0


25582.0
108.0
0.0
0.0
0.0
4.0
12.0
9475.0
6665.0
7333.0
1714.0
271.0
0.0
0.0
0.0
0.0
0.0
0.0
0.0
0.0
0.0
0.0
0.0
0.0
0.0
0.0
0.0
0.0
0.0
0.0

0.99
0.99
0.99
0.99
0.99
0.99
0.99
0.99
0.99
0.99
0.99
0.99
0.99
0.99
0.99
0.99
0.99
0.99
0.99
0.99
0.99
0.99
0.99
0.99
0.99
0.99
0.99
0.99
0.99
0.99


2992.0
0.0
21.0
0.0
0.0
0.0
0.0
0.0
0.0
0.0
0.0
0.0
0.0
0.0
2971.0
0.0
0.0
0.0
0.0
0.0
0.0
0.0
0.0
0.0
0.0
0.0
0.0
0.0
0.0
0.0

1.0
1.0
1.0
1.0
1.0
1.0
1.0
1.0
1.0
1.0
1.0
1.0
1.0
1.0
1.0
1.0
1.0
1.0
1.0
1.0
1.0
1.0
1.0
1.0
1.0
1.0
1.0
1.0
1.0
1.0


251.0
0.0
0.0
0.0
0.0
0.0
0.0
0.0
17.0
0.0
0.0
0.0
0.0
0.0
0.0
0.0
11.0
0.0
223.0
0.0
0.0
0.0
0.0
0.0
0.0
0.0
0.0
0.0
0.0
0.0

1.0
1.0
1.0
1.0
1.0
1.0
1.0
1.0
1.0
1.0
1.0
1.0
1.0
1.0
1.0
1.0
1.0
1.0
1.0
1.0
1.0
1.0
1.0
1.0
1.0
1.0
1.0
1.0
1.0
1.0


9.0
0.0
0.0
0.0
0.0
0.0
0.0
0.0
9.0
0.0
0.0
0.0
0.0
0.0
0.0
0.0
0.0
0.0
0.0
0.0
0.0
0.0
0.0
0.0
0.0
0.0
0.0
0.0
0.0
0.0

1.0
1.0
1.0
1.0
1.0
1.0
1.0
1.0
1.0
1.0
1.0
1.0
1.0
1.0
1.0
1.0
1.0
1.0
1.0
1.0
1.0
1.0
1.0
1.0
1.0
1.0
1.0
1.0
1.0
1.0


11.0
0.0
0.0
0.0
0.0
0.0
0.0
0.0
0.0
0.0
0.0
0.0
0.0
0.0
0.0
0.0
0.0
0.0
0.0
0.0
0.0
0.0
0.0
0.0
6.0
0.0
0.0
0.0
0.0
5.0

1.0
1.0
1.0
1.0
1.0
1.0
1.0
1.0
1.0
1.0
1.0
1.0
1.0
1.0
1.0
1.0
1.0
1.0
1.0
1.0
1.0
1.0
1.0
1.0
1.0
1.0
1.0
1.0
1.0
1.0


1534.0
0.0
0.0
0.0
0.0
0.0
0.0
93.0
320.0
90.0
275.0
0.0
0.0
0.0
47.0
0.0
0.0
0.0
0.0
0.0
0.0
0.0
0.0
0.0
0.0
243.0
242.0
110.0
0.0
114.0

1.0
1.0
1.0
1.0
1.0
1.0
1.0
1.0
1.0
1.0
1.0
1.0
1.0
1.0
1.0
1.0
1.0
1.0
1.0
1.0
1.0
1.0
1.0
1.0
1.0
1.0
1.0
1.0
1.0
1.0


386.0
0.0
0.0
0.0
0.0
0.0
0.0
0.0
17.0
15.0
32.0
0.0
0.0
0.0
0.0
0.0
0.0
0.0
0.0
0.0
0.0
0.0
0.0
0.0
0.0
64.0
166.0
0.0
0.0
92.0

1.0
1.0
1.0
1.0
1.0
1.0
1.0
1.0
1.0
1.0
1.0
1.0
1.0
1.0
1.0
1.0
1.0
1.0
1.0
1.0
1.0
1.0
1.0
1.0
1.0
1.0
1.0
1.0
1.0
1.0


387.0
0.0
0.0
0.0
0.0
0.0
0.0
0.0
0.0
0.0
0.0
0.0
0.0
0.0
0.0
0.0
0.0
0.0
0.0
0.0
0.0
0.0
0.0
0.0
0.0
179.0
76.0
110.0
0.0
22.0

1.0
1.0
1.0
1.0
1.0
1.0
1.0
1.0
1.0
1.0
1.0
1.0
1.0
1.0
1.0
1.0
1.0
1.0
1.0
1.0
1.0
1.0
1.0
1.0
1.0
1.0
1.0
1.0
1.0
1.0


47.0
0.0
0.0
0.0
0.0
0.0
0.0
0.0
0.0
0.0
0.0
0.0
0.0
0.0
47.0
0.0
0.0
0.0
0.0
0.0
0.0
0.0
0.0
0.0
0.0
0.0
0.0
0.0
0.0
0.0

0.96
0.96
0.96
0.96
0.96
0.96
0.96
0.96
0.96
0.96
0.96
0.96
0.96
0.96
0.96
0.96
0.96
0.96
0.96
0.96
0.96
0.96
0.96
0.96
0.96
0.96
0.96
0.96
0.96
0.96


195.0
0.0
0.0
0.0
0.0
0.0
0.0
56.0
139.0
0.0
0.0
0.0
0.0
0.0
0.0
0.0
0.0
0.0
0.0
0.0
0.0
0.0
0.0
0.0
0.0
0.0
0.0
0.0
0.0
0.0

1.0
1.0
1.0
1.0
1.0
1.0
1.0
1.0
1.0
1.0
1.0
1.0
1.0
1.0
1.0
1.0
1.0
1.0
1.0
1.0
1.0
1.0
1.0
1.0
1.0
1.0
1.0
1.0
1.0
1.0


132.0
0.0
0.0
0.0
0.0
0.0
0.0
0.0
132.0
0.0
0.0
0.0
0.0
0.0
0.0
0.0
0.0
0.0
0.0
0.0
0.0
0.0
0.0
0.0
0.0
0.0
0.0
0.0
0.0
0.0

1.0
1.0
1.0
1.0
1.0
1.0
1.0
1.0
1.0
1.0
1.0
1.0
1.0
1.0
1.0
1.0
1.0
1.0
1.0
1.0
1.0
1.0
1.0
1.0
1.0
1.0
1.0
1.0
1.0
1.0


386.0
0.0
0.0
0.0
0.0
0.0
0.0
0.0
0.0
49.0
9.0
0.0
0.0
0.0
0.0
0.0
0.0
0.0
0.0
0.0
0.0
0.0
0.0
0.0
0.0
278.0
27.0
0.0
23.0
0.0

1.0
1.0
1.0
1.0
1.0
1.0
1.0
1.0
1.0
1.0
1.0
1.0
1.0
1.0
1.0
1.0
1.0
1.0
1.0
1.0
1.0
1.0
1.0
1.0
1.0
1.0
1.0
1.0
1.0
1.0


58.0
0.0
0.0
0.0
0.0
0.0
0.0
0.0
0.0
49.0
9.0
0.0
0.0
0.0
0.0
0.0
0.0
0.0
0.0
0.0
0.0
0.0
0.0
0.0
0.0
0.0
0.0
0.0
0.0
0.0

0.98
0.98
0.98
0.98
0.98
0.98
0.98
0.98
0.98
0.98
0.98
0.98
0.98
0.98
0.98
0.98
0.98
0.98
0.98
0.98
0.98
0.98
0.98
0.98
0.98
0.98
0.98
0.98
0.98
0.98


121.0
0.0
0.0
0.0
0.0
0.0
0.0
0.0
0.0
0.0
0.0
0.0
0.0
0.0
0.0
0.0
0.0
0.0
0.0
0.0
0.0
0.0
0.0
0.0
0.0
94.0
27.0
0.0
0.0
0.0

0.96
0.96
0.96
0.96
0.96
0.96
0.96
0.96
0.96
0.96
0.96
0.96
0.96
0.96
0.96
0.96
0.96
0.96
0.96
0.96
0.96
0.96
0.96
0.96
0.96
0.96
0.96
0.96
0.96
0.96


23.0
0.0
0.0
0.0
0.0
0.0
0.0
0.0
0.0
0.0
0.0
0.0
0.0
0.0
0.0
0.0
0.0
0.0
0.0
0.0
0.0
0.0
0.0
0.0
0.0
0.0
0.0
0.0
23.0
0.0

0.9
0.9
0.9
0.9
0.9
0.9
0.9
0.9
0.9
0.9
0.9
0.9
0.9
0.9
0.9
0.9
0.9
0.9
0.9
0.9
0.9
0.9
0.9
0.9
0.9
0.9
0.9
0.9
0.9
0.9


28070.0
145.0
19.0
0.0
0.0
0.0
0.0
1113.0
2767.0
10798.0
7847.0
475.0
2453.0
0.0
0.0
0.0
0.0
0.0
0.0
0.0
0.0
0.0
0.0
2453.0
0.0
0.0
0.0
0.0
0.0
0.0


23042.0
133.0
12.0
0.0
0.0
0.0
0.0
1113.0
2719.0
10772.0
7818.0
475.0
0.0
0.0
0.0
0.0
0.0
0.0
0.0
0.0
0.0
0.0
0.0
0.0
0.0
0.0
0.0
0.0
0.0
0.0

1.0
1.0
1.0
1.0
1.0
1.0
1.0
1.0
1.0
1.0
1.0
1.0
1.0
1.0
1.0
1.0
1.0
1.0
1.0
1.0
1.0
1.0
1.0
1.0
1.0
1.0
1.0
1.0
1.0
1.0


305.0
0.0
0.0
0.0
0.0
0.0
0.0
0.0
0.0
230.0
75.0
0.0
0.0
0.0
0.0
0.0
0.0
0.0
0.0
0.0
0.0
0.0
0.0
0.0
0.0
0.0
0.0
0.0
0.0
0.0

1.0
1.0
1.0
1.0
1.0
1.0
1.0
1.0
1.0
1.0
1.0
1.0
1.0
1.0
1.0
1.0
1.0
1.0
1.0
1.0
1.0
1.0
1.0
1.0
1.0
1.0
1.0
1.0
1.0
1.0


65.0
0.0
0.0
0.0
0.0
0.0
0.0
0.0
0.0
0.0
27.0
38.0
0.0
0.0
0.0
0.0
0.0
0.0
0.0
0.0
0.0
0.0
0.0
0.0
0.0
0.0
0.0
0.0
0.0
0.0

1.0
1.0
1.0
1.0
1.0
1.0
1.0
1.0
1.0
1.0
1.0
1.0
1.0
1.0
1.0
1.0
1.0
1.0
1.0
1.0
1.0
1.0
1.0
1.0
1.0
1.0
1.0
1.0
1.0
1.0


18.0
11.0
0.0
0.0
0.0
0.0
0.0
0.0
0.0
7.0
0.0
0.0
0.0
0.0
0.0
0.0
0.0
0.0
0.0
0.0
0.0
0.0
0.0
0.0
0.0
0.0
0.0
0.0
0.0
0.0

1.0
1.0
1.0
1.0
1.0
1.0
1.0
1.0
1.0
1.0
1.0
1.0
1.0
1.0
1.0
1.0
1.0
1.0
1.0
1.0
1.0
1.0
1.0
1.0
1.0
1.0
1.0
1.0
1.0
1.0


103.0
0.0
0.0
0.0
0.0
0.0
0.0
0.0
48.0
26.0
29.0
0.0
0.0
0.0
0.0
0.0
0.0
0.0
0.0
0.0
0.0
0.0
0.0
0.0
0.0
0.0
0.0
0.0
0.0
0.0

1.0
1.0
1.0
1.0
1.0
1.0
1.0
1.0
1.0
1.0
1.0
1.0
1.0
1.0
1.0
1.0
1.0
1.0
1.0
1.0
1.0
1.0
1.0
1.0
1.0
1.0
1.0
1.0
1.0
1.0


9.0
0.0
0.0
0.0
0.0
0.0
0.0
0.0
0.0
9.0
0.0
0.0
0.0
0.0
0.0
0.0
0.0
0.0
0.0
0.0
0.0
0.0
0.0
0.0
0.0
0.0
0.0
0.0
0.0
0.0

0.82
0.82
0.82
0.82
0.82
0.82
0.82
0.82
0.82
0.82
0.82
0.82
0.82
0.82
0.82
0.82
0.82
0.82
0.82
0.82
0.82
0.82
0.82
0.82
0.82
0.82
0.82
0.82
0.82
0.82


4925.0
12.0
7.0
0.0
0.0
0.0
0.0
0.0
0.0
0.0
0.0
0.0
2453.0
0.0
0.0
0.0
0.0
0.0
0.0
0.0
0.0
0.0
0.0
2453.0
0.0
0.0
0.0
0.0
0.0
0.0


4925.0
12.0
7.0
0.0
0.0
0.0
0.0
0.0
0.0
0.0
0.0
0.0
2453.0
0.0
0.0
0.0
0.0
0.0
0.0
0.0
0.0
0.0
0.0
2453.0
0.0
0.0
0.0
0.0
0.0
0.0

1.0
1.0
1.0
1.0
1.0
1.0
1.0
1.0
1.0
1.0
1.0
1.0
1.0
1.0
1.0
1.0
1.0
1.0
1.0
1.0
1.0
1.0
1.0
1.0
1.0
1.0
1.0
1.0
1.0
1.0


47106.0
2746.0
0.0
0.0
0.0
0.0
0.0
5627.0
0.0
19199.0
17255.0
2279.0
0.0
0.0
0.0
0.0
0.0
0.0
0.0
0.0
0.0
0.0
0.0
0.0
0.0
0.0
0.0
0.0
0.0
0.0


47106.0
2746.0
0.0
0.0
0.0
0.0
0.0
5627.0
0.0
19199.0
17255.0
2279.0
0.0
0.0
0.0
0.0
0.0
0.0
0.0
0.0
0.0
0.0
0.0
0.0
0.0
0.0
0.0
0.0
0.0
0.0

1.0
1.0
1.0
1.0
1.0
1.0
1.0
1.0
1.0
1.0
1.0
1.0
1.0
1.0
1.0
1.0
1.0
1.0
1.0
1.0
1.0
1.0
1.0
1.0
1.0
1.0
1.0
1.0
1.0
1.0


1365221.0
5121.0
2624.0
2758.0
1423.0
2384.0
1473.0
12375.0
13688.0
12498.0
9866.0
8183.0
65094.0
18859.0
49868.0
24163.0
8834.0
340380.0
253394.0
166422.0
67236.0
89748.0
46951.0
65094.0
17100.0
32503.0
8821.0
8960.0
19521.0
9880.0

1.0
1.0
1.0
1.0
1.0
1.0
1.0
1.0
1.0
1.0
1.0
1.0
1.0
1.0
1.0
1.0
1.0
1.0
1.0
1.0
1.0
1.0
1.0
1.0
1.0
1.0
1.0
1.0
1.0
1.0


1009502.0
861.0
436.0
359.0
459.0
724.0
760.0
10612.0
12826.0
8442.0
5526.0
1470.0
65094.0
2115.0
24376.0
4276.0
1820.0
287240.0
175944.0
155212.0
61063.0
28628.0
46951.0
65094.0
7771.0
16166.0
7220.0
4811.0
9478.0
3768.0

1.0
1.0
1.0
1.0
1.0
1.0
1.0
1.0
1.0
1.0
1.0
1.0
1.0
1.0
1.0
1.0
1.0
1.0
1.0
1.0
1.0
1.0
1.0
1.0
1.0
1.0
1.0
1.0
1.0
1.0


29222.0
39.0
28.0
7.0
0.0
0.0
0.0
1951.0
5768.0
3505.0
2616.0
121.0
5479.0
0.0
0.0
3.0
0.0
109.0
3964.0
0.0
0.0
0.0
82.0
5479.0
41.0
0.0
0.0
0.0
30.0
0.0

1.0
1.0
1.0
1.0
1.0
1.0
1.0
1.0
1.0
1.0
1.0
1.0
1.0
1.0
1.0
1.0
1.0
1.0
1.0
1.0
1.0
1.0
1.0
1.0
1.0
1.0
1.0
1.0
1.0
1.0


23700.0
192.0
13.0
3.0
20.0
0.0
0.0
2731.0
1327.0
720.0
562.0
86.0
0.0
0.0
15254.0
0.0
0.0
593.0
2199.0
0.0
0.0
0.0
0.0
0.0
0.0
0.0
0.0
0.0
0.0
0.0

1.0
1.0
1.0
1.0
1.0
1.0
1.0
1.0
1.0
1.0
1.0
1.0
1.0
1.0
1.0
1.0
1.0
1.0
1.0
1.0
1.0
1.0
1.0
1.0
1.0
1.0
1.0
1.0
1.0
1.0


115452.0
25.0
0.0
0.0
0.0
7.0
0.0
99.0
392.0
1197.0
416.0
40.0
81.0
0.0
0.0
0.0
0.0
108010.0
633.0
372.0
605.0
2154.0
1340.0
81.0
0.0
0.0
0.0
0.0
0.0
0.0

1.0
1.0
1.0
1.0
1.0
1.0
1.0
1.0
1.0
1.0
1.0
1.0
1.0
1.0
1.0
1.0
1.0
1.0
1.0
1.0
1.0
1.0
1.0
1.0
1.0
1.0
1.0
1.0
1.0
1.0


6404.0
31.0
0.0
0.0
0.0
0.0
0.0
153.0
132.0
77.0
8.0
0.0
1663.0
0.0
0.0
0.0
0.0
0.0
0.0
0.0
0.0
0.0
2677.0
1663.0
0.0
0.0
0.0
0.0
0.0
0.0

1.0
1.0
1.0
1.0
1.0
1.0
1.0
1.0
1.0
1.0
1.0
1.0
1.0
1.0
1.0
1.0
1.0
1.0
1.0
1.0
1.0
1.0
1.0
1.0
1.0
1.0
1.0
1.0
1.0
1.0


950.0
4.0
0.0
0.0
0.0
0.0
0.0
88.0
114.0
211.0
91.0
2.0
0.0
0.0
0.0
0.0
0.0
330.0
110.0
0.0
0.0
0.0
0.0
0.0
0.0
0.0
0.0
0.0
0.0
0.0

1.0
1.0
1.0
1.0
1.0
1.0
1.0
1.0
1.0
1.0
1.0
1.0
1.0
1.0
1.0
1.0
1.0
1.0
1.0
1.0
1.0
1.0
1.0
1.0
1.0
1.0
1.0
1.0
1.0
1.0


116271.0
12.0
6.0
0.0
0.0
0.0
0.0
0.0
159.0
393.0
228.0
119.0
536.0
0.0
0.0
0.0
0.0
52520.0
18553.0
35041.0
65.0
4070.0
4033.0
536.0
0.0
0.0
0.0
0.0
0.0
0.0

1.0
1.0
1.0
1.0
1.0
1.0
1.0
1.0
1.0
1.0
1.0
1.0
1.0
1.0
1.0
1.0
1.0
1.0
1.0
1.0
1.0
1.0
1.0
1.0
1.0
1.0
1.0
1.0
1.0
1.0


1569.0
26.0
39.0
0.0
0.0
0.0
0.0
86.0
0.0
7.0
128.0
6.0
0.0
0.0
0.0
0.0
0.0
111.0
0.0
0.0
1101.0
0.0
0.0
0.0
65.0
0.0
0.0
0.0
0.0
0.0

1.0
1.0
1.0
1.0
1.0
1.0
1.0
1.0
1.0
1.0
1.0
1.0
1.0
1.0
1.0
1.0
1.0
1.0
1.0
1.0
1.0
1.0
1.0
1.0
1.0
1.0
1.0
1.0
1.0
1.0


2848.0
1.0
0.0
0.0
0.0
0.0
0.0
90.0
91.0
58.0
0.0
0.0
0.0
68.0
0.0
0.0
0.0
0.0
0.0
2351.0
14.0
0.0
0.0
0.0
0.0
0.0
0.0
0.0
0.0
175.0

1.0
1.0
1.0
1.0
1.0
1.0
1.0
1.0
1.0
1.0
1.0
1.0
1.0
1.0
1.0
1.0
1.0
1.0
1.0
1.0
1.0
1.0
1.0
1.0
1.0
1.0
1.0
1.0
1.0
1.0


29645.0
4.0
2.0
0.0
0.0
0.0
0.0
167.0
19.0
13.0
27.0
1.0
0.0
0.0
0.0
0.0
0.0
10117.0
5852.0
0.0
42.0
4008.0
9393.0
0.0
0.0
0.0
0.0
0.0
0.0
0.0

1.0
1.0
1.0
1.0
1.0
1.0
1.0
1.0
1.0
1.0
1.0
1.0
1.0
1.0
1.0
1.0
1.0
1.0
1.0
1.0
1.0
1.0
1.0
1.0
1.0
1.0
1.0
1.0
1.0
1.0


85291.0
56.0
6.0
0.0
0.0
0.0
0.0
0.0
63.0
51.0
45.0
377.0
41994.0
705.0
0.0
0.0
0.0
0.0
0.0
0.0
0.0
0.0
0.0
41994.0
0.0
0.0
0.0
0.0
0.0
0.0

1.0
1.0
1.0
1.0
1.0
1.0
1.0
1.0
1.0
1.0
1.0
1.0
1.0
1.0
1.0
1.0
1.0
1.0
1.0
1.0
1.0
1.0
1.0
1.0
1.0
1.0
1.0
1.0
1.0
1.0


186.0
1.0
6.0
0.0
17.0
0.0
0.0
0.0
10.0
31.0
107.0
14.0
0.0
0.0
0.0
0.0
0.0
0.0
0.0
0.0
0.0
0.0
0.0
0.0
0.0
0.0
0.0
0.0
0.0
0.0

0.93
0.93
0.93
0.93
0.93
0.93
0.93
0.93
0.93
0.93
0.93
0.93
0.93
0.93
0.93
0.93
0.93
0.93
0.93
0.93
0.93
0.93
0.93
0.93
0.93
0.93
0.93
0.93
0.93
0.93


3274.0
0.0
1.0
1.0
47.0
44.0
2.0
0.0
0.0
0.0
0.0
0.0
0.0
117.0
195.0
0.0
174.0
321.0
69.0
0.0
253.0
0.0
0.0
0.0
1845.0
31.0
0.0
0.0
153.0
21.0

1.0
1.0
1.0
1.0
1.0
1.0
1.0
1.0
1.0
1.0
1.0
1.0
1.0
1.0
1.0
1.0
1.0
1.0
1.0
1.0
1.0
1.0
1.0
1.0
1.0
1.0
1.0
1.0
1.0
1.0


202.0
40.0
6.0
0.0
0.0
0.0
0.0
0.0
34.0
3.0
0.0
6.0
0.0
0.0
0.0
0.0
0.0
0.0
5.0
0.0
0.0
0.0
0.0
0.0
108.0
0.0
0.0
0.0
0.0
0.0

1.0
1.0
1.0
1.0
1.0
1.0
1.0
1.0
1.0
1.0
1.0
1.0
1.0
1.0
1.0
1.0
1.0
1.0
1.0
1.0
1.0
1.0
1.0
1.0
1.0
1.0
1.0
1.0
1.0
1.0


8548.0
0.0
2.0
0.0
14.0
29.0
16.0
0.0
0.0
1.0
0.0
0.0
0.0
0.0
6.0
681.0
1000.0
1898.0
529.0
0.0
0.0
0.0
0.0
0.0
439.0
2421.0
658.0
0.0
798.0
56.0

1.0
1.0
1.0
1.0
1.0
1.0
1.0
1.0
1.0
1.0
1.0
1.0
1.0
1.0
1.0
1.0
1.0
1.0
1.0
1.0
1.0
1.0
1.0
1.0
1.0
1.0
1.0
1.0
1.0
1.0


33.0
4.0
9.0
1.0
0.0
0.0
0.0
0.0
3.0
5.0
11.0
0.0
0.0
0.0
0.0
0.0
0.0
0.0
0.0
0.0
0.0
0.0
0.0
0.0
0.0
0.0
0.0
0.0
0.0
0.0

0.71
0.71
0.71
0.71
0.71
0.71
0.71
0.71
0.71
0.71
0.71
0.71
0.71
0.71
0.71
0.71
0.71
0.71
0.71
0.71
0.71
0.71
0.71
0.71
0.71
0.71
0.71
0.71
0.71
0.71


8314.0
2.0
0.0
0.0
0.0
0.0
0.0
0.0
0.0
4.0
24.0
0.0
0.0
0.0
0.0
0.0
0.0
1630.0
6654.0
0.0
0.0
0.0
0.0
0.0
0.0
0.0
0.0
0.0
0.0
0.0

1.0
1.0
1.0
1.0
1.0
1.0
1.0
1.0
1.0
1.0
1.0
1.0
1.0
1.0
1.0
1.0
1.0
1.0
1.0
1.0
1.0
1.0
1.0
1.0
1.0
1.0
1.0
1.0
1.0
1.0


23.0
0.0
0.0
0.0
0.0
0.0
0.0
23.0
0.0
0.0
0.0
0.0
0.0
0.0
0.0
0.0
0.0
0.0
0.0
0.0
0.0
0.0
0.0
0.0
0.0
0.0
0.0
0.0
0.0
0.0

1.0
1.0
1.0
1.0
1.0
1.0
1.0
1.0
1.0
1.0
1.0
1.0
1.0
1.0
1.0
1.0
1.0
1.0
1.0
1.0
1.0
1.0
1.0
1.0
1.0
1.0
1.0
1.0
1.0
1.0


12.0
5.0
7.0
0.0
0.0
0.0
0.0
0.0
0.0
0.0
0.0
0.0
0.0
0.0
0.0
0.0
0.0
0.0
0.0
0.0
0.0
0.0
0.0
0.0
0.0
0.0
0.0
0.0
0.0
0.0

1.0
1.0
1.0
1.0
1.0
1.0
1.0
1.0
1.0
1.0
1.0
1.0
1.0
1.0
1.0
1.0
1.0
1.0
1.0
1.0
1.0
1.0
1.0
1.0
1.0
1.0
1.0
1.0
1.0
1.0


162036.0
0.0
0.0
0.0
0.0
0.0
0.0
0.0
0.0
0.0
0.0
0.0
1892.0
0.0
0.0
0.0
0.0
9808.0
20366.0
83908.0
42990.0
1180.0
0.0
1892.0
0.0
0.0
0.0
0.0
0.0
0.0

1.0
1.0
1.0
1.0
1.0
1.0
1.0
1.0
1.0
1.0
1.0
1.0
1.0
1.0
1.0
1.0
1.0
1.0
1.0
1.0
1.0
1.0
1.0
1.0
1.0
1.0
1.0
1.0
1.0
1.0


23572.0
0.0
0.0
0.0
0.0
0.0
0.0
0.0
0.0
0.0
0.0
0.0
0.0
0.0
0.0
0.0
0.0
0.0
23572.0
0.0
0.0
0.0
0.0
0.0
0.0
0.0
0.0
0.0
0.0
0.0

1.0
1.0
1.0
1.0
1.0
1.0
1.0
1.0
1.0
1.0
1.0
1.0
1.0
1.0
1.0
1.0
1.0
1.0
1.0
1.0
1.0
1.0
1.0
1.0
1.0
1.0
1.0
1.0
1.0
1.0


5454.0
0.0
0.0
0.0
0.0
0.0
0.0
0.0
0.0
0.0
0.0
0.0
0.0
0.0
0.0
0.0
0.0
0.0
280.0
3575.0
777.0
822.0
0.0
0.0
0.0
0.0
0.0
0.0
0.0
0.0

1.0
1.0
1.0
1.0
1.0
1.0
1.0
1.0
1.0
1.0
1.0
1.0
1.0
1.0
1.0
1.0
1.0
1.0
1.0
1.0
1.0
1.0
1.0
1.0
1.0
1.0
1.0
1.0
1.0
1.0


563.0
0.0
0.0
0.0
0.0
0.0
0.0
0.0
0.0
0.0
0.0
0.0
0.0
0.0
0.0
0.0
0.0
0.0
563.0
0.0
0.0
0.0
0.0
0.0
0.0
0.0
0.0
0.0
0.0
0.0

1.0
1.0
1.0
1.0
1.0
1.0
1.0
1.0
1.0
1.0
1.0
1.0
1.0
1.0
1.0
1.0
1.0
1.0
1.0
1.0
1.0
1.0
1.0
1.0
1.0
1.0
1.0
1.0
1.0
1.0


9.0
0.0
0.0
0.0
0.0
0.0
0.0
0.0
0.0
0.0
0.0
0.0
0.0
0.0
0.0
0.0
0.0
0.0
9.0
0.0
0.0
0.0
0.0
0.0
0.0
0.0
0.0
0.0
0.0
0.0

0.71
0.71
0.71
0.71
0.71
0.71
0.71
0.71
0.71
0.71
0.71
0.71
0.71
0.71
0.71
0.71
0.71
0.71
0.71
0.71
0.71
0.71
0.71
0.71
0.71
0.71
0.71
0.71
0.71
0.71


216460.0
3237.0
1483.0
1698.0
379.0
857.0
282.0
0.0
535.0
1457.0
3324.0
4113.0
0.0
8566.0
16685.0
9679.0
4035.0
28208.0
63061.0
2107.0
0.0
31850.0
0.0
0.0
9231.0
16337.0
544.0
984.0
5791.0
2017.0

1.0
1.0
1.0
1.0
1.0
1.0
1.0
1.0
1.0
1.0
1.0
1.0
1.0
1.0
1.0
1.0
1.0
1.0
1.0
1.0
1.0
1.0
1.0
1.0
1.0
1.0
1.0
1.0
1.0
1.0


3338.0
418.0
174.0
30.0
0.0
0.0
0.0
0.0
535.0
910.0
590.0
681.0
0.0
0.0
0.0
0.0
0.0
0.0
0.0
0.0
0.0
0.0
0.0
0.0
0.0
0.0
0.0
0.0
0.0
0.0

1.0
1.0
1.0
1.0
1.0
1.0
1.0
1.0
1.0
1.0
1.0
1.0
1.0
1.0
1.0
1.0
1.0
1.0
1.0
1.0
1.0
1.0
1.0
1.0
1.0
1.0
1.0
1.0
1.0
1.0


5664.0
36.0
49.0
6.0
25.0
61.0
0.0
1199.0
273.0
348.0
0.0
124.0
0.0
3543.0
0.0
0.0
0.0
0.0
0.0
0.0
0.0
0.0
0.0
0.0
0.0
0.0
0.0
0.0
0.0
0.0

0.7
0.7
0.7
0.7
0.7
0.7
0.7
0.7
0.7
0.7
0.7
0.7
0.7
0.7
0.7
0.7
0.7
0.7
0.7
0.7
0.7
0.7
0.7
0.7
0.7
0.7
0.7
0.7
0.7
0.7


4442.0
36.0
49.0
6.0
25.0
61.0
0.0
0.0
273.0
325.0
0.0
124.0
0.0
3543.0
0.0
0.0
0.0
0.0
0.0
0.0
0.0
0.0
0.0
0.0
0.0
0.0
0.0
0.0
0.0
0.0

1.0
1.0
1.0
1.0
1.0
1.0
1.0
1.0
1.0
1.0
1.0
1.0
1.0
1.0
1.0
1.0
1.0
1.0
1.0
1.0
1.0
1.0
1.0
1.0
1.0
1.0
1.0
1.0
1.0
1.0


10982.0
53.0
0.0
0.0
0.0
0.0
0.0
417.0
0.0
1352.0
543.0
0.0
0.0
0.0
0.0
0.0
0.0
0.0
0.0
0.0
0.0
8617.0
0.0
0.0
0.0
0.0
0.0
0.0
0.0
0.0

1.0
1.0
1.0
1.0
1.0
1.0
1.0
1.0
1.0
1.0
1.0
1.0
1.0
1.0
1.0
1.0
1.0
1.0
1.0
1.0
1.0
1.0
1.0
1.0
1.0
1.0
1.0
1.0
1.0
1.0


77675.0
61.0
63.0
109.0
97.0
407.0
17.0
0.0
0.0
15.0
140.0
149.0
0.0
4412.0
8503.0
10208.0
2979.0
12791.0
14389.0
9103.0
1647.0
259.0
0.0
0.0
98.0
0.0
1057.0
3165.0
3928.0
4078.0


77675.0
61.0
63.0
109.0
97.0
407.0
17.0
0.0
0.0
15.0
140.0
149.0
0.0
4412.0
8503.0
10208.0
2979.0
12791.0
14389.0
9103.0
1647.0
259.0
0.0
0.0
98.0
0.0
1057.0
3165.0
3928.0
4078.0

1.0
1.0
1.0
1.0
1.0
1.0
1.0
1.0
1.0
1.0
1.0
1.0
1.0
1.0
1.0
1.0
1.0
1.0
1.0
1.0
1.0
1.0
1.0
1.0
1.0
1.0
1.0
1.0
1.0
1.0


13226.0
18.0
48.0
9.0
1.0
166.0
12.0
0.0
23.0
25.0
0.0
42.0
0.0
223.0
304.0
0.0
0.0
12141.0
0.0
0.0
0.0
0.0
0.0
0.0
0.0
0.0
0.0
0.0
197.0
17.0


13226.0
18.0
48.0
9.0
1.0
166.0
12.0
0.0
23.0
25.0
0.0
42.0
0.0
223.0
304.0
0.0
0.0
12141.0
0.0
0.0
0.0
0.0
0.0
0.0
0.0
0.0
0.0
0.0
197.0
17.0

1.0
1.0
1.0
1.0
1.0
1.0
1.0
1.0
1.0
1.0
1.0
1.0
1.0
1.0
1.0
1.0
1.0
1.0
1.0
1.0
1.0
1.0
1.0
1.0
1.0
1.0
1.0
1.0
1.0
1.0


107.0
0.0
0.0
0.0
0.0
0.0
0.0
0.0
0.0
0.0
0.0
107.0
0.0
0.0
0.0
0.0
0.0
0.0
0.0
0.0
0.0
0.0
0.0
0.0
0.0
0.0
0.0
0.0
0.0
0.0

1.0
1.0
1.0
1.0
1.0
1.0
1.0
1.0
1.0
1.0
1.0
1.0
1.0
1.0
1.0
1.0
1.0
1.0
1.0
1.0
1.0
1.0
1.0
1.0
1.0
1.0
1.0
1.0
1.0
1.0


1268214.0
3209.0
335.0
0.0
0.0
0.0
0.0
7424.0
3533.0
3600.0
6235.0
3672.0
109414.0
0.0
0.0
0.0
0.0
0.0
0.0
346576.0
228192.0
154963.0
256802.0
109414.0
14818.0
0.0
0.0
0.0
0.0
20027.0


1268214.0
3209.0
335.0
0.0
0.0
0.0
0.0
7424.0
3533.0
3600.0
6235.0
3672.0
109414.0
0.0
0.0
0.0
0.0
0.0
0.0
346576.0
228192.0
154963.0
256802.0
109414.0
14818.0
0.0
0.0
0.0
0.0
20027.0

1.0
1.0
1.0
1.0
1.0
1.0
1.0
1.0
1.0
1.0
1.0
1.0
1.0
1.0
1.0
1.0
1.0
1.0
1.0
1.0
1.0
1.0
1.0
1.0
1.0
1.0
1.0
1.0
1.0
1.0


511729.0
1187.0
0.0
0.0
0.0
0.0
0.0
7424.0
3533.0
2154.0
1833.0
0.0
109412.0
0.0
0.0
0.0
0.0
0.0
0.0
0.0
0.0
0.0
256747.0
109412.0
0.0
0.0
0.0
0.0
0.0
20027.0

1.0
1.0
1.0
1.0
1.0
1.0
1.0
1.0
1.0
1.0
1.0
1.0
1.0
1.0
1.0
1.0
1.0
1.0
1.0
1.0
1.0
1.0
1.0
1.0
1.0
1.0
1.0
1.0
1.0
1.0


21315.0
0.0
0.0
0.0
0.0
0.0
0.0
0.0
0.0
0.0
2850.0
3672.0
0.0
0.0
0.0
0.0
0.0
0.0
0.0
0.0
0.0
0.0
0.0
0.0
14793.0
0.0
0.0
0.0
0.0
0.0

1.0
1.0
1.0
1.0
1.0
1.0
1.0
1.0
1.0
1.0
1.0
1.0
1.0
1.0
1.0
1.0
1.0
1.0
1.0
1.0
1.0
1.0
1.0
1.0
1.0
1.0
1.0
1.0
1.0
1.0


574768.0
0.0
0.0
0.0
0.0
0.0
0.0
0.0
0.0
0.0
0.0
0.0
0.0
0.0
0.0
0.0
0.0
0.0
0.0
346576.0
228192.0
0.0
0.0
0.0
0.0
0.0
0.0
0.0
0.0
0.0

1.0
1.0
1.0
1.0
1.0
1.0
1.0
1.0
1.0
1.0
1.0
1.0
1.0
1.0
1.0
1.0
1.0
1.0
1.0
1.0
1.0
1.0
1.0
1.0
1.0
1.0
1.0
1.0
1.0
1.0


50102.0
629.0
328.0
3.0
0.0
0.0
0.0
5825.0
11183.0
11604.0
8353.0
5312.0
0.0
0.0
4828.0
0.0
0.0
35.0
926.0
0.0
0.0
0.0
0.0
0.0
14.0
0.0
446.0
0.0
616.0
0.0

0.99
0.99
0.99
0.99
0.99
0.99
0.99
0.99
0.99
0.99
0.99
0.99
0.99
0.99
0.99
0.99
0.99
0.99
0.99
0.99
0.99
0.99
0.99
0.99
0.99
0.99
0.99
0.99
0.99
0.99


14994.0
458.0
30.0
0.0
0.0
0.0
0.0
2513.0
1918.0
4979.0
3587.0
1509.0
0.0
0.0
0.0
0.0
0.0
0.0
0.0
0.0
0.0
0.0
0.0
0.0
0.0
0.0
0.0
0.0
0.0
0.0

0.93
0.93
0.93
0.93
0.93
0.93
0.93
0.93
0.93
0.93
0.93
0.93
0.93
0.93
0.93
0.93
0.93
0.93
0.93
0.93
0.93
0.93
0.93
0.93
0.93
0.93
0.93
0.93
0.93
0.93


11627.0
458.0
30.0
0.0
0.0
0.0
0.0
1474.0
1262.0
3359.0
3536.0
1508.0
0.0
0.0
0.0
0.0
0.0
0.0
0.0
0.0
0.0
0.0
0.0
0.0
0.0
0.0
0.0
0.0
0.0
0.0

1.0
1.0
1.0
1.0
1.0
1.0
1.0
1.0
1.0
1.0
1.0
1.0
1.0
1.0
1.0
1.0
1.0
1.0
1.0
1.0
1.0
1.0
1.0
1.0
1.0
1.0
1.0
1.0
1.0
1.0


3127.0
0.0
0.0
0.0
0.0
0.0
0.0
960.0
613.0
1502.0
51.0
1.0
0.0
0.0
0.0
0.0
0.0
0.0
0.0
0.0
0.0
0.0
0.0
0.0
0.0
0.0
0.0
0.0
0.0
0.0

1.0
1.0
1.0
1.0
1.0
1.0
1.0
1.0
1.0
1.0
1.0
1.0
1.0
1.0
1.0
1.0
1.0
1.0
1.0
1.0
1.0
1.0
1.0
1.0
1.0
1.0
1.0
1.0
1.0
1.0


52.0
0.0
0.0
0.0
0.0
0.0
0.0
9.0
22.0
21.0
0.0
0.0
0.0
0.0
0.0
0.0
0.0
0.0
0.0
0.0
0.0
0.0
0.0
0.0
0.0
0.0
0.0
0.0
0.0
0.0

1.0
1.0
1.0
1.0
1.0
1.0
1.0
1.0
1.0
1.0
1.0
1.0
1.0
1.0
1.0
1.0
1.0
1.0
1.0
1.0
1.0
1.0
1.0
1.0
1.0
1.0
1.0
1.0
1.0
1.0


18.0
0.0
0.0
0.0
0.0
0.0
0.0
0.0
0.0
18.0
0.0
0.0
0.0
0.0
0.0
0.0
0.0
0.0
0.0
0.0
0.0
0.0
0.0
0.0
0.0
0.0
0.0
0.0
0.0
0.0

1.0
1.0
1.0
1.0
1.0
1.0
1.0
1.0
1.0
1.0
1.0
1.0
1.0
1.0
1.0
1.0
1.0
1.0
1.0
1.0
1.0
1.0
1.0
1.0
1.0
1.0
1.0
1.0
1.0
1.0


8767.0
11.0
2.0
3.0
0.0
0.0
0.0
1191.0
3949.0
2638.0
816.0
157.0
0.0
0.0
0.0
0.0
0.0
0.0
0.0
0.0
0.0
0.0
0.0
0.0
0.0
0.0
0.0
0.0
0.0
0.0

0.96
0.96
0.96
0.96
0.96
0.96
0.96
0.96
0.96
0.96
0.96
0.96
0.96
0.96
0.96
0.96
0.96
0.96
0.96
0.96
0.96
0.96
0.96
0.96
0.96
0.96
0.96
0.96
0.96
0.96


6104.0
11.0
2.0
3.0
0.0
0.0
0.0
833.0
2903.0
1868.0
374.0
110.0
0.0
0.0
0.0
0.0
0.0
0.0
0.0
0.0
0.0
0.0
0.0
0.0
0.0
0.0
0.0
0.0
0.0
0.0

1.0
1.0
1.0
1.0
1.0
1.0
1.0
1.0
1.0
1.0
1.0
1.0
1.0
1.0
1.0
1.0
1.0
1.0
1.0
1.0
1.0
1.0
1.0
1.0
1.0
1.0
1.0
1.0
1.0
1.0


6327.0
0.0
0.0
0.0
0.0
0.0
0.0
444.0
2457.0
1088.0
1828.0
510.0
0.0
0.0
0.0
0.0
0.0
0.0
0.0
0.0
0.0
0.0
0.0
0.0
0.0
0.0
0.0
0.0
0.0
0.0


6327.0
0.0
0.0
0.0
0.0
0.0
0.0
444.0
2457.0
1088.0
1828.0
510.0
0.0
0.0
0.0
0.0
0.0
0.0
0.0
0.0
0.0
0.0
0.0
0.0
0.0
0.0
0.0
0.0
0.0
0.0

1.0
1.0
1.0
1.0
1.0
1.0
1.0
1.0
1.0
1.0
1.0
1.0
1.0
1.0
1.0
1.0
1.0
1.0
1.0
1.0
1.0
1.0
1.0
1.0
1.0
1.0
1.0
1.0
1.0
1.0


4106.0
0.0
0.0
0.0
0.0
0.0
0.0
931.0
1020.0
1717.0
403.0
35.0
0.0
0.0
0.0
0.0
0.0
0.0
0.0
0.0
0.0
0.0
0.0
0.0
0.0
0.0
0.0
0.0
0.0
0.0

1.0
1.0
1.0
1.0
1.0
1.0
1.0
1.0
1.0
1.0
1.0
1.0
1.0
1.0
1.0
1.0
1.0
1.0
1.0
1.0
1.0
1.0
1.0
1.0
1.0
1.0
1.0
1.0
1.0
1.0


1770.0
0.0
0.0
0.0
0.0
0.0
0.0
0.0
128.0
1226.0
389.0
27.0
0.0
0.0
0.0
0.0
0.0
0.0
0.0
0.0
0.0
0.0
0.0
0.0
0.0
0.0
0.0
0.0
0.0
0.0

1.0
1.0
1.0
1.0
1.0
1.0
1.0
1.0
1.0
1.0
1.0
1.0
1.0
1.0
1.0
1.0
1.0
1.0
1.0
1.0
1.0
1.0
1.0
1.0
1.0
1.0
1.0
1.0
1.0
1.0


497.0
0.0
0.0
0.0
0.0
0.0
0.0
259.0
79.0
137.0
14.0
8.0
0.0
0.0
0.0
0.0
0.0
0.0
0.0
0.0
0.0
0.0
0.0
0.0
0.0
0.0
0.0
0.0
0.0
0.0

1.0
1.0
1.0
1.0
1.0
1.0
1.0
1.0
1.0
1.0
1.0
1.0
1.0
1.0
1.0
1.0
1.0
1.0
1.0
1.0
1.0
1.0
1.0
1.0
1.0
1.0
1.0
1.0
1.0
1.0


380.0
0.0
0.0
0.0
0.0
0.0
0.0
0.0
195.0
160.0
25.0
0.0
0.0
0.0
0.0
0.0
0.0
0.0
0.0
0.0
0.0
0.0
0.0
0.0
0.0
0.0
0.0
0.0
0.0
0.0

0.93
0.93
0.93
0.93
0.93
0.93
0.93
0.93
0.93
0.93
0.93
0.93
0.93
0.93
0.93
0.93
0.93
0.93
0.93
0.93
0.93
0.93
0.93
0.93
0.93
0.93
0.93
0.93
0.93
0.93


983.0
0.0
0.0
0.0
0.0
0.0
0.0
91.0
223.0
0.0
0.0
0.0
0.0
0.0
0.0
0.0
0.0
0.0
669.0
0.0
0.0
0.0
0.0
0.0
0.0
0.0
0.0
0.0
0.0
0.0


302.0
0.0
0.0
0.0
0.0
0.0
0.0
91.0
211.0
0.0
0.0
0.0
0.0
0.0
0.0
0.0
0.0
0.0
0.0
0.0
0.0
0.0
0.0
0.0
0.0
0.0
0.0
0.0
0.0
0.0

1.0
1.0
1.0
1.0
1.0
1.0
1.0
1.0
1.0
1.0
1.0
1.0
1.0
1.0
1.0
1.0
1.0
1.0
1.0
1.0
1.0
1.0
1.0
1.0
1.0
1.0
1.0
1.0
1.0
1.0


681.0
0.0
0.0
0.0
0.0
0.0
0.0
0.0
12.0
0.0
0.0
0.0
0.0
0.0
0.0
0.0
0.0
0.0
669.0
0.0
0.0
0.0
0.0
0.0
0.0
0.0
0.0
0.0
0.0
0.0

0.97
0.97
0.97
0.97
0.97
0.97
0.97
0.97
0.97
0.97
0.97
0.97
0.97
0.97
0.97
0.97
0.97
0.97
0.97
0.97
0.97
0.97
0.97
0.97
0.97
0.97
0.97
0.97
0.97
0.97


427.0
0.0
0.0
0.0
0.0
0.0
0.0
0.0
0.0
268.0
143.0
16.0
0.0
0.0
0.0
0.0
0.0
0.0
0.0
0.0
0.0
0.0
0.0
0.0
0.0
0.0
0.0
0.0
0.0
0.0

0.75
0.75
0.75
0.75
0.75
0.75
0.75
0.75
0.75
0.75
0.75
0.75
0.75
0.75
0.75
0.75
0.75
0.75
0.75
0.75
0.75
0.75
0.75
0.75
0.75
0.75
0.75
0.75
0.75
0.75


872.0
0.0
0.0
0.0
0.0
0.0
0.0
153.0
479.0
116.0
107.0
17.0
0.0
0.0
0.0
0.0
0.0
0.0
0.0
0.0
0.0
0.0
0.0
0.0
0.0
0.0
0.0
0.0
0.0
0.0

1.0
1.0
1.0
1.0
1.0
1.0
1.0
1.0
1.0
1.0
1.0
1.0
1.0
1.0
1.0
1.0
1.0
1.0
1.0
1.0
1.0
1.0
1.0
1.0
1.0
1.0
1.0
1.0
1.0
1.0


324.0
0.0
0.0
0.0
0.0
0.0
0.0
0.0
212.0
39.0
73.0
0.0
0.0
0.0
0.0
0.0
0.0
0.0
0.0
0.0
0.0
0.0
0.0
0.0
0.0
0.0
0.0
0.0
0.0
0.0

1.0
1.0
1.0
1.0
1.0
1.0
1.0
1.0
1.0
1.0
1.0
1.0
1.0
1.0
1.0
1.0
1.0
1.0
1.0
1.0
1.0
1.0
1.0
1.0
1.0
1.0
1.0
1.0
1.0
1.0


59.0
0.0
0.0
0.0
0.0
0.0
0.0
0.0
35.0
24.0
0.0
0.0
0.0
0.0
0.0
0.0
0.0
0.0
0.0
0.0
0.0
0.0
0.0
0.0
0.0
0.0
0.0
0.0
0.0
0.0

1.0
1.0
1.0
1.0
1.0
1.0
1.0
1.0
1.0
1.0
1.0
1.0
1.0
1.0
1.0
1.0
1.0
1.0
1.0
1.0
1.0
1.0
1.0
1.0
1.0
1.0
1.0
1.0
1.0
1.0


536.0
0.0
0.0
0.0
0.0
0.0
0.0
0.0
213.0
0.0
238.0
85.0
0.0
0.0
0.0
0.0
0.0
0.0
0.0
0.0
0.0
0.0
0.0
0.0
0.0
0.0
0.0
0.0
0.0
0.0


536.0
0.0
0.0
0.0
0.0
0.0
0.0
0.0
213.0
0.0
238.0
85.0
0.0
0.0
0.0
0.0
0.0
0.0
0.0
0.0
0.0
0.0
0.0
0.0
0.0
0.0
0.0
0.0
0.0
0.0

1.0
1.0
1.0
1.0
1.0
1.0
1.0
1.0
1.0
1.0
1.0
1.0
1.0
1.0
1.0
1.0
1.0
1.0
1.0
1.0
1.0
1.0
1.0
1.0
1.0
1.0
1.0
1.0
1.0
1.0


376.0
0.0
0.0
0.0
0.0
0.0
0.0
231.0
52.0
90.0
3.0
0.0
0.0
0.0
0.0
0.0
0.0
0.0
0.0
0.0
0.0
0.0
0.0
0.0
0.0
0.0
0.0
0.0
0.0
0.0

1.0
1.0
1.0
1.0
1.0
1.0
1.0
1.0
1.0
1.0
1.0
1.0
1.0
1.0
1.0
1.0
1.0
1.0
1.0
1.0
1.0
1.0
1.0
1.0
1.0
1.0
1.0
1.0
1.0
1.0


179.0
1.0
0.0
0.0
0.0
0.0
0.0
0.0
91.0
0.0
79.0
8.0
0.0
0.0
0.0
0.0
0.0
0.0
0.0
0.0
0.0
0.0
0.0
0.0
0.0
0.0
0.0
0.0
0.0
0.0


159.0
1.0
0.0
0.0
0.0
0.0
0.0
0.0
91.0
0.0
59.0
8.0
0.0
0.0
0.0
0.0
0.0
0.0
0.0
0.0
0.0
0.0
0.0
0.0
0.0
0.0
0.0
0.0
0.0
0.0

1.0
1.0
1.0
1.0
1.0
1.0
1.0
1.0
1.0
1.0
1.0
1.0
1.0
1.0
1.0
1.0
1.0
1.0
1.0
1.0
1.0
1.0
1.0
1.0
1.0
1.0
1.0
1.0
1.0
1.0


20.0
0.0
0.0
0.0
0.0
0.0
0.0
0.0
0.0
0.0
20.0
0.0
0.0
0.0
0.0
0.0
0.0
0.0
0.0
0.0
0.0
0.0
0.0
0.0
0.0
0.0
0.0
0.0
0.0
0.0

0.99
0.99
0.99
0.99
0.99
0.99
0.99
0.99
0.99
0.99
0.99
0.99
0.99
0.99
0.99
0.99
0.99
0.99
0.99
0.99
0.99
0.99
0.99
0.99
0.99
0.99
0.99
0.99
0.99
0.99


78.0
0.0
0.0
0.0
0.0
0.0
0.0
0.0
0.0
78.0
0.0
0.0
0.0
0.0
0.0
0.0
0.0
0.0
0.0
0.0
0.0
0.0
0.0
0.0
0.0
0.0
0.0
0.0
0.0
0.0


78.0
0.0
0.0
0.0
0.0
0.0
0.0
0.0
0.0
78.0
0.0
0.0
0.0
0.0
0.0
0.0
0.0
0.0
0.0
0.0
0.0
0.0
0.0
0.0
0.0
0.0
0.0
0.0
0.0
0.0

1.0
1.0
1.0
1.0
1.0
1.0
1.0
1.0
1.0
1.0
1.0
1.0
1.0
1.0
1.0
1.0
1.0
1.0
1.0
1.0
1.0
1.0
1.0
1.0
1.0
1.0
1.0
1.0
1.0
1.0


55.0
0.0
0.0
0.0
0.0
0.0
0.0
55.0
0.0
0.0
0.0
0.0
0.0
0.0
0.0
0.0
0.0
0.0
0.0
0.0
0.0
0.0
0.0
0.0
0.0
0.0
0.0
0.0
0.0
0.0


55.0
0.0
0.0
0.0
0.0
0.0
0.0
55.0
0.0
0.0
0.0
0.0
0.0
0.0
0.0
0.0
0.0
0.0
0.0
0.0
0.0
0.0
0.0
0.0
0.0
0.0
0.0
0.0
0.0
0.0

1.0
1.0
1.0
1.0
1.0
1.0
1.0
1.0
1.0
1.0
1.0
1.0
1.0
1.0
1.0
1.0
1.0
1.0
1.0
1.0
1.0
1.0
1.0
1.0
1.0
1.0
1.0
1.0
1.0
1.0


40.0
0.0
0.0
0.0
0.0
0.0
0.0
0.0
0.0
25.0
15.0
0.0
0.0
0.0
0.0
0.0
0.0
0.0
0.0
0.0
0.0
0.0
0.0
0.0
0.0
0.0
0.0
0.0
0.0
0.0


40.0
0.0
0.0
0.0
0.0
0.0
0.0
0.0
0.0
25.0
15.0
0.0
0.0
0.0
0.0
0.0
0.0
0.0
0.0
0.0
0.0
0.0
0.0
0.0
0.0
0.0
0.0
0.0
0.0
0.0

1.0
1.0
1.0
1.0
1.0
1.0
1.0
1.0
1.0
1.0
1.0
1.0
1.0
1.0
1.0
1.0
1.0
1.0
1.0
1.0
1.0
1.0
1.0
1.0
1.0
1.0
1.0
1.0
1.0
1.0


11.0
0.0
0.0
0.0
0.0
0.0
0.0
0.0
0.0
0.0
11.0
0.0
0.0
0.0
0.0
0.0
0.0
0.0
0.0
0.0
0.0
0.0
0.0
0.0
0.0
0.0
0.0
0.0
0.0
0.0


11.0
0.0
0.0
0.0
0.0
0.0
0.0
0.0
0.0
0.0
11.0
0.0
0.0
0.0
0.0
0.0
0.0
0.0
0.0
0.0
0.0
0.0
0.0
0.0
0.0
0.0
0.0
0.0
0.0
0.0

0.89
0.89
0.89
0.89
0.89
0.89
0.89
0.89
0.89
0.89
0.89
0.89
0.89
0.89
0.89
0.89
0.89
0.89
0.89
0.89
0.89
0.89
0.89
0.89
0.89
0.89
0.89
0.89
0.89
0.89


4723.0
0.0
0.0
0.0
0.0
0.0
0.0
0.0
0.0
0.0
0.0
0.0
0.0
0.0
4723.0
0.0
0.0
0.0
0.0
0.0
0.0
0.0
0.0
0.0
0.0
0.0
0.0
0.0
0.0
0.0


4723.0
0.0
0.0
0.0
0.0
0.0
0.0
0.0
0.0
0.0
0.0
0.0
0.0
0.0
4723.0
0.0
0.0
0.0
0.0
0.0
0.0
0.0
0.0
0.0
0.0
0.0
0.0
0.0
0.0
0.0

1.0
1.0
1.0
1.0
1.0
1.0
1.0
1.0
1.0
1.0
1.0
1.0
1.0
1.0
1.0
1.0
1.0
1.0
1.0
1.0
1.0
1.0
1.0
1.0
1.0
1.0
1.0
1.0
1.0
1.0


1062.0
0.0
0.0
0.0
0.0
0.0
0.0
0.0
0.0
0.0
0.0
0.0
0.0
0.0
0.0
0.0
0.0
0.0
0.0
0.0
0.0
0.0
0.0
0.0
0.0
0.0
446.0
0.0
616.0
0.0


1062.0
0.0
0.0
0.0
0.0
0.0
0.0
0.0
0.0
0.0
0.0
0.0
0.0
0.0
0.0
0.0
0.0
0.0
0.0
0.0
0.0
0.0
0.0
0.0
0.0
0.0
446.0
0.0
616.0
0.0

1.0
1.0
1.0
1.0
1.0
1.0
1.0
1.0
1.0
1.0
1.0
1.0
1.0
1.0
1.0
1.0
1.0
1.0
1.0
1.0
1.0
1.0
1.0
1.0
1.0
1.0
1.0
1.0
1.0
1.0


14.0
0.0
0.0
0.0
0.0
0.0
0.0
0.0
0.0
0.0
0.0
0.0
0.0
0.0
0.0
0.0
0.0
0.0
0.0
0.0
0.0
0.0
0.0
0.0
14.0
0.0
0.0
0.0
0.0
0.0


14.0
0.0
0.0
0.0
0.0
0.0
0.0
0.0
0.0
0.0
0.0
0.0
0.0
0.0
0.0
0.0
0.0
0.0
0.0
0.0
0.0
0.0
0.0
0.0
14.0
0.0
0.0
0.0
0.0
0.0

1.0
1.0
1.0
1.0
1.0
1.0
1.0
1.0
1.0
1.0
1.0
1.0
1.0
1.0
1.0
1.0
1.0
1.0
1.0
1.0
1.0
1.0
1.0
1.0
1.0
1.0
1.0
1.0
1.0
1.0


2406733.0
4695.0
1667.0
1341.0
885.0
1734.0
3263.0
17084.0
10651.0
26255.0
20219.0
8284.0
71915.0
27852.0
3060.0
14592.0
3163.0
54605.0
75151.0
530921.0
189537.0
0.0
143213.0
71915.0
657232.0
129403.0
124862.0
43110.0
110859.0
59265.0

0.71
0.71
0.71
0.71
0.71
0.71
0.71
0.71
0.71
0.71
0.71
0.71
0.71
0.71
0.71
0.71
0.71
0.71
0.71
0.71
0.71
0.71
0.71
0.71
0.71
0.71
0.71
0.71
0.71
0.71


13316.0
0.0
0.0
0.0
0.0
0.0
0.0
9360.0
1372.0
1454.0
1130.0
0.0
0.0
0.0
0.0
0.0
0.0
0.0
0.0
0.0
0.0
0.0
0.0
0.0
0.0
0.0
0.0
0.0
0.0
0.0


13316.0
0.0
0.0
0.0
0.0
0.0
0.0
9360.0
1372.0
1454.0
1130.0
0.0
0.0
0.0
0.0
0.0
0.0
0.0
0.0
0.0
0.0
0.0
0.0
0.0
0.0
0.0
0.0
0.0
0.0
0.0

1.0
1.0
1.0
1.0
1.0
1.0
1.0
1.0
1.0
1.0
1.0
1.0
1.0
1.0
1.0
1.0
1.0
1.0
1.0
1.0
1.0
1.0
1.0
1.0
1.0
1.0
1.0
1.0
1.0
1.0


17605.0
1010.0
3.0
1.0
1.0
3.0
1.0
0.0
3903.0
9973.0
2709.0
1.0
0.0
0.0
0.0
0.0
0.0
0.0
0.0
0.0
0.0
0.0
0.0
0.0
0.0
0.0
0.0
0.0
0.0
0.0

1.0
1.0
1.0
1.0
1.0
1.0
1.0
1.0
1.0
1.0
1.0
1.0
1.0
1.0
1.0
1.0
1.0
1.0
1.0
1.0
1.0
1.0
1.0
1.0
1.0
1.0
1.0
1.0
1.0
1.0


352398.0
207.0
0.0
0.0
0.0
0.0
0.0
7724.0
1659.0
3014.0
2528.0
0.0
71915.0
0.0
0.0
0.0
0.0
0.0
0.0
20265.0
29958.0
0.0
143213.0
71915.0
0.0
0.0
0.0
0.0
0.0
0.0

1.0
1.0
1.0
1.0
1.0
1.0
1.0
1.0
1.0
1.0
1.0
1.0
1.0
1.0
1.0
1.0
1.0
1.0
1.0
1.0
1.0
1.0
1.0
1.0
1.0
1.0
1.0
1.0
1.0
1.0


352383.0
207.0
0.0
0.0
0.0
0.0
0.0
7724.0
1659.0
3014.0
2528.0
0.0
71915.0
0.0
0.0
0.0
0.0
0.0
0.0
20265.0
29943.0
0.0
143213.0
71915.0
0.0
0.0
0.0
0.0
0.0
0.0

1.0
1.0
1.0
1.0
1.0
1.0
1.0
1.0
1.0
1.0
1.0
1.0
1.0
1.0
1.0
1.0
1.0
1.0
1.0
1.0
1.0
1.0
1.0
1.0
1.0
1.0
1.0
1.0
1.0
1.0


1267181.0
795.0
1304.0
977.0
883.0
1730.0
2908.0
0.0
1502.0
2605.0
9600.0
2806.0
0.0
27852.0
837.0
14592.0
3163.0
25439.0
75151.0
510656.0
159579.0
0.0
0.0
0.0
10838.0
129403.0
123885.0
40620.0
60791.0
59265.0

1.0
1.0
1.0
1.0
1.0
1.0
1.0
1.0
1.0
1.0
1.0
1.0
1.0
1.0
1.0
1.0
1.0
1.0
1.0
1.0
1.0
1.0
1.0
1.0
1.0
1.0
1.0
1.0
1.0
1.0


14861.0
337.0
434.0
492.0
0.0
402.0
107.0
0.0
1502.0
1658.0
6439.0
1954.0
0.0
0.0
0.0
0.0
0.0
0.0
0.0
0.0
0.0
0.0
0.0
0.0
0.0
0.0
0.0
0.0
0.0
1536.0

1.0
1.0
1.0
1.0
1.0
1.0
1.0
1.0
1.0
1.0
1.0
1.0
1.0
1.0
1.0
1.0
1.0
1.0
1.0
1.0
1.0
1.0
1.0
1.0
1.0
1.0
1.0
1.0
1.0
1.0


1023190.0
187.0
537.0
178.0
531.0
1328.0
996.0
0.0
0.0
402.0
3161.0
576.0
0.0
24905.0
1.0
0.0
0.0
0.0
75151.0
510656.0
148770.0
0.0
0.0
0.0
0.0
11637.0
120441.0
19977.0
46027.0
57729.0

1.0
1.0
1.0
1.0
1.0
1.0
1.0
1.0
1.0
1.0
1.0
1.0
1.0
1.0
1.0
1.0
1.0
1.0
1.0
1.0
1.0
1.0
1.0
1.0
1.0
1.0
1.0
1.0
1.0
1.0


53204.0
271.0
249.0
28.0
0.0
0.0
809.0
0.0
0.0
0.0
0.0
276.0
0.0
0.0
0.0
0.0
0.0
0.0
0.0
0.0
0.0
0.0
0.0
0.0
0.0
16164.0
0.0
20643.0
14764.0
0.0

0.99
0.99
0.99
0.99
0.99
0.99
0.99
0.99
0.99
0.99
0.99
0.99
0.99
0.99
0.99
0.99
0.99
0.99
0.99
0.99
0.99
0.99
0.99
0.99
0.99
0.99
0.99
0.99
0.99
0.99


142106.0
0.0
84.0
279.0
352.0
0.0
705.0
0.0
0.0
0.0
0.0
0.0
0.0
2947.0
836.0
14592.0
3163.0
25439.0
0.0
0.0
10791.0
0.0
0.0
0.0
10838.0
68647.0
3433.0
0.0
0.0
0.0

1.0
1.0
1.0
1.0
1.0
1.0
1.0
1.0
1.0
1.0
1.0
1.0
1.0
1.0
1.0
1.0
1.0
1.0
1.0
1.0
1.0
1.0
1.0
1.0
1.0
1.0
1.0
1.0
1.0
1.0


545.0
0.0
0.0
0.0
0.0
0.0
0.0
0.0
0.0
545.0
0.0
0.0
0.0
0.0
0.0
0.0
0.0
0.0
0.0
0.0
0.0
0.0
0.0
0.0
0.0
0.0
0.0
0.0
0.0
0.0

1.0
1.0
1.0
1.0
1.0
1.0
1.0
1.0
1.0
1.0
1.0
1.0
1.0
1.0
1.0
1.0
1.0
1.0
1.0
1.0
1.0
1.0
1.0
1.0
1.0
1.0
1.0
1.0
1.0
1.0


32955.0
0.0
0.0
0.0
0.0
0.0
0.0
0.0
0.0
0.0
0.0
0.0
0.0
0.0
0.0
0.0
0.0
0.0
0.0
0.0
0.0
0.0
0.0
0.0
0.0
32955.0
0.0
0.0
0.0
0.0

0.97
0.97
0.97
0.97
0.97
0.97
0.97
0.97
0.97
0.97
0.97
0.97
0.97
0.97
0.97
0.97
0.97
0.97
0.97
0.97
0.97
0.97
0.97
0.97
0.97
0.97
0.97
0.97
0.97
0.97


571132.0
1931.0
262.0
0.0
1.0
0.0
0.0
0.0
0.0
5942.0
0.0
4493.0
0.0
0.0
0.0
0.0
0.0
29144.0
0.0
0.0
0.0
0.0
0.0
0.0
529359.0
0.0
0.0
0.0
0.0
0.0

1.0
1.0
1.0
1.0
1.0
1.0
1.0
1.0
1.0
1.0
1.0
1.0
1.0
1.0
1.0
1.0
1.0
1.0
1.0
1.0
1.0
1.0
1.0
1.0
1.0
1.0
1.0
1.0
1.0
1.0


7403.0
1459.0
1.0
0.0
1.0
0.0
0.0
0.0
0.0
5942.0
0.0
0.0
0.0
0.0
0.0
0.0
0.0
0.0
0.0
0.0
0.0
0.0
0.0
0.0
0.0
0.0
0.0
0.0
0.0
0.0

1.0
1.0
1.0
1.0
1.0
1.0
1.0
1.0
1.0
1.0
1.0
1.0
1.0
1.0
1.0
1.0
1.0
1.0
1.0
1.0
1.0
1.0
1.0
1.0
1.0
1.0
1.0
1.0
1.0
1.0


2037.0
0.0
0.0
0.0
0.0
0.0
0.0
0.0
0.0
0.0
0.0
2037.0
0.0
0.0
0.0
0.0
0.0
0.0
0.0
0.0
0.0
0.0
0.0
0.0
0.0
0.0
0.0
0.0
0.0
0.0

0.76
0.76
0.76
0.76
0.76
0.76
0.76
0.76
0.76
0.76
0.76
0.76
0.76
0.76
0.76
0.76
0.76
0.76
0.76
0.76
0.76
0.76
0.76
0.76
0.76
0.76
0.76
0.76
0.76
0.76


1172.0
0.0
0.0
0.0
0.0
0.0
0.0
0.0
0.0
0.0
0.0
1172.0
0.0
0.0
0.0
0.0
0.0
0.0
0.0
0.0
0.0
0.0
0.0
0.0
0.0
0.0
0.0
0.0
0.0
0.0

1.0
1.0
1.0
1.0
1.0
1.0
1.0
1.0
1.0
1.0
1.0
1.0
1.0
1.0
1.0
1.0
1.0
1.0
1.0
1.0
1.0
1.0
1.0
1.0
1.0
1.0
1.0
1.0
1.0
1.0


2003.0
458.0
261.0
0.0
0.0
0.0
0.0
0.0
0.0
0.0
0.0
1284.0
0.0
0.0
0.0
0.0
0.0
0.0
0.0
0.0
0.0
0.0
0.0
0.0
0.0
0.0
0.0
0.0
0.0
0.0

1.0
1.0
1.0
1.0
1.0
1.0
1.0
1.0
1.0
1.0
1.0
1.0
1.0
1.0
1.0
1.0
1.0
1.0
1.0
1.0
1.0
1.0
1.0
1.0
1.0
1.0
1.0
1.0
1.0
1.0


529248.0
0.0
0.0
0.0
0.0
0.0
0.0
0.0
0.0
0.0
0.0
0.0
0.0
0.0
0.0
0.0
0.0
0.0
0.0
0.0
0.0
0.0
0.0
0.0
529248.0
0.0
0.0
0.0
0.0
0.0

0.97
0.97
0.97
0.97
0.97
0.97
0.97
0.97
0.97
0.97
0.97
0.97
0.97
0.97
0.97
0.97
0.97
0.97
0.97
0.97
0.97
0.97
0.97
0.97
0.97
0.97
0.97
0.97
0.97
0.97


8149.0
453.0
0.0
0.0
0.0
1.0
0.0
0.0
394.0
2781.0
3862.0
658.0
0.0
0.0
0.0
0.0
0.0
0.0
0.0
0.0
0.0
0.0
0.0
0.0
0.0
0.0
0.0
0.0
0.0
0.0

1.0
1.0
1.0
1.0
1.0
1.0
1.0
1.0
1.0
1.0
1.0
1.0
1.0
1.0
1.0
1.0
1.0
1.0
1.0
1.0
1.0
1.0
1.0
1.0
1.0
1.0
1.0
1.0
1.0
1.0


965.0
0.0
0.0
0.0
0.0
0.0
0.0
0.0
394.0
571.0
0.0
0.0
0.0
0.0
0.0
0.0
0.0
0.0
0.0
0.0
0.0
0.0
0.0
0.0
0.0
0.0
0.0
0.0
0.0
0.0

0.84
0.84
0.84
0.84
0.84
0.84
0.84
0.84
0.84
0.84
0.84
0.84
0.84
0.84
0.84
0.84
0.84
0.84
0.84
0.84
0.84
0.84
0.84
0.84
0.84
0.84
0.84
0.84
0.84
0.84


1821.0
0.0
0.0
0.0
0.0
0.0
0.0
0.0
1821.0
0.0
0.0
0.0
0.0
0.0
0.0
0.0
0.0
0.0
0.0
0.0
0.0
0.0
0.0
0.0
0.0
0.0
0.0
0.0
0.0
0.0


1821.0
0.0
0.0
0.0
0.0
0.0
0.0
0.0
1821.0
0.0
0.0
0.0
0.0
0.0
0.0
0.0
0.0
0.0
0.0
0.0
0.0
0.0
0.0
0.0
0.0
0.0
0.0
0.0
0.0
0.0

1.0
1.0
1.0
1.0
1.0
1.0
1.0
1.0
1.0
1.0
1.0
1.0
1.0
1.0
1.0
1.0
1.0
1.0
1.0
1.0
1.0
1.0
1.0
1.0
1.0
1.0
1.0
1.0
1.0
1.0


174233.0
299.0
98.0
363.0
0.0
0.0
354.0
0.0
0.0
0.0
0.0
326.0
0.0
0.0
2223.0
0.0
0.0
0.0
0.0
0.0
0.0
0.0
0.0
0.0
117035.0
0.0
977.0
2490.0
50068.0
0.0

1.0
1.0
1.0
1.0
1.0
1.0
1.0
1.0
1.0
1.0
1.0
1.0
1.0
1.0
1.0
1.0
1.0
1.0
1.0
1.0
1.0
1.0
1.0
1.0
1.0
1.0
1.0
1.0
1.0
1.0


137804.0
299.0
0.0
325.0
0.0
0.0
0.0
0.0
0.0
0.0
0.0
0.0
0.0
0.0
0.0
0.0
0.0
0.0
0.0
0.0
0.0
0.0
0.0
0.0
87112.0
0.0
0.0
0.0
50068.0
0.0

0.96
0.96
0.96
0.96
0.96
0.96
0.96
0.96
0.96
0.96
0.96
0.96
0.96
0.96
0.96
0.96
0.96
0.96
0.96
0.96
0.96
0.96
0.96
0.96
0.96
0.96
0.96
0.96
0.96
0.96


185.0
0.0
51.0
0.0
0.0
0.0
0.0
0.0
0.0
0.0
0.0
134.0
0.0
0.0
0.0
0.0
0.0
0.0
0.0
0.0
0.0
0.0
0.0
0.0
0.0
0.0
0.0
0.0
0.0
0.0

1.0
1.0
1.0
1.0
1.0
1.0
1.0
1.0
1.0
1.0
1.0
1.0
1.0
1.0
1.0
1.0
1.0
1.0
1.0
1.0
1.0
1.0
1.0
1.0
1.0
1.0
1.0
1.0
1.0
1.0


30432.0
0.0
0.0
0.0
0.0
0.0
0.0
0.0
0.0
0.0
0.0
0.0
0.0
0.0
362.0
0.0
0.0
0.0
0.0
0.0
0.0
0.0
0.0
0.0
29918.0
0.0
0.0
152.0
0.0
0.0

0.91
0.91
0.91
0.91
0.91
0.91
0.91
0.91
0.91
0.91
0.91
0.91
0.91
0.91
0.91
0.91
0.91
0.91
0.91
0.91
0.91
0.91
0.91
0.91
0.91
0.91
0.91
0.91
0.91
0.91


977.0
0.0
0.0
0.0
0.0
0.0
0.0
0.0
0.0
0.0
0.0
0.0
0.0
0.0
0.0
0.0
0.0
0.0
0.0
0.0
0.0
0.0
0.0
0.0
0.0
0.0
977.0
0.0
0.0
0.0

0.77
0.77
0.77
0.77
0.77
0.77
0.77
0.77
0.77
0.77
0.77
0.77
0.77
0.77
0.77
0.77
0.77
0.77
0.77
0.77
0.77
0.77
0.77
0.77
0.77
0.77
0.77
0.77
0.77
0.77


486.0
0.0
0.0
0.0
0.0
0.0
0.0
0.0
0.0
486.0
0.0
0.0
0.0
0.0
0.0
0.0
0.0
0.0
0.0
0.0
0.0
0.0
0.0
0.0
0.0
0.0
0.0
0.0
0.0
0.0


486.0
0.0
0.0
0.0
0.0
0.0
0.0
0.0
0.0
486.0
0.0
0.0
0.0
0.0
0.0
0.0
0.0
0.0
0.0
0.0
0.0
0.0
0.0
0.0
0.0
0.0
0.0
0.0
0.0
0.0

0.99
0.99
0.99
0.99
0.99
0.99
0.99
0.99
0.99
0.99
0.99
0.99
0.99
0.99
0.99
0.99
0.99
0.99
0.99
0.99
0.99
0.99
0.99
0.99
0.99
0.99
0.99
0.99
0.99
0.99


390.0
0.0
0.0
0.0
0.0
0.0
0.0
0.0
0.0
0.0
390.0
0.0
0.0
0.0
0.0
0.0
0.0
0.0
0.0
0.0
0.0
0.0
0.0
0.0
0.0
0.0
0.0
0.0
0.0
0.0

1.0
1.0
1.0
1.0
1.0
1.0
1.0
1.0
1.0
1.0
1.0
1.0
1.0
1.0
1.0
1.0
1.0
1.0
1.0
1.0
1.0
1.0
1.0
1.0
1.0
1.0
1.0
1.0
1.0
1.0


13015.0
2084.0
451.0
0.0
0.0
0.0
0.0
0.0
0.0
0.0
5284.0
5196.0
0.0
0.0
0.0
0.0
0.0
0.0
0.0
0.0
0.0
0.0
0.0
0.0
0.0
0.0
0.0
0.0
0.0
0.0


13015.0
2084.0
451.0
0.0
0.0
0.0
0.0
0.0
0.0
0.0
5284.0
5196.0
0.0
0.0
0.0
0.0
0.0
0.0
0.0
0.0
0.0
0.0
0.0
0.0
0.0
0.0
0.0
0.0
0.0
0.0

0.93
0.93
0.93
0.93
0.93
0.93
0.93
0.93
0.93
0.93
0.93
0.93
0.93
0.93
0.93
0.93
0.93
0.93
0.93
0.93
0.93
0.93
0.93
0.93
0.93
0.93
0.93
0.93
0.93
0.93


291629.0
119.0
69.0
0.0
0.0
0.0
0.0
3199.0
3589.0
784.0
566.0
834.0
0.0
0.0
0.0
0.0
0.0
4.0
1.0
0.0
0.0
0.0
282458.0
0.0
0.0
0.0
6.0
0.0
0.0
0.0

0.76
0.76
0.76
0.76
0.76
0.76
0.76
0.76
0.76
0.76
0.76
0.76
0.76
0.76
0.76
0.76
0.76
0.76
0.76
0.76
0.76
0.76
0.76
0.76
0.76
0.76
0.76
0.76
0.76
0.76


289239.0
0.0
0.0
0.0
0.0
0.0
0.0
3199.0
3582.0
0.0
0.0
0.0
0.0
0.0
0.0
0.0
0.0
0.0
0.0
0.0
0.0
0.0
282458.0
0.0
0.0
0.0
0.0
0.0
0.0
0.0


289239.0
0.0
0.0
0.0
0.0
0.0
0.0
3199.0
3582.0
0.0
0.0
0.0
0.0
0.0
0.0
0.0
0.0
0.0
0.0
0.0
0.0
0.0
282458.0
0.0
0.0
0.0
0.0
0.0
0.0
0.0

0.99
0.99
0.99
0.99
0.99
0.99
0.99
0.99
0.99
0.99
0.99
0.99
0.99
0.99
0.99
0.99
0.99
0.99
0.99
0.99
0.99
0.99
0.99
0.99
0.99
0.99
0.99
0.99
0.99
0.99


724.0
0.0
0.0
0.0
0.0
0.0
0.0
0.0
0.0
0.0
0.0
724.0
0.0
0.0
0.0
0.0
0.0
0.0
0.0
0.0
0.0
0.0
0.0
0.0
0.0
0.0
0.0
0.0
0.0
0.0


724.0
0.0
0.0
0.0
0.0
0.0
0.0
0.0
0.0
0.0
0.0
724.0
0.0
0.0
0.0
0.0
0.0
0.0
0.0
0.0
0.0
0.0
0.0
0.0
0.0
0.0
0.0
0.0
0.0
0.0

1.0
1.0
1.0
1.0
1.0
1.0
1.0
1.0
1.0
1.0
1.0
1.0
1.0
1.0
1.0
1.0
1.0
1.0
1.0
1.0
1.0
1.0
1.0
1.0
1.0
1.0
1.0
1.0
1.0
1.0


797.0
0.0
0.0
0.0
0.0
0.0
0.0
0.0
7.0
784.0
0.0
0.0
0.0
0.0
0.0
0.0
0.0
0.0
0.0
0.0
0.0
0.0
0.0
0.0
0.0
0.0
6.0
0.0
0.0
0.0

0.95
0.95
0.95
0.95
0.95
0.95
0.95
0.95
0.95
0.95
0.95
0.95
0.95
0.95
0.95
0.95
0.95
0.95
0.95
0.95
0.95
0.95
0.95
0.95
0.95
0.95
0.95
0.95
0.95
0.95


52321.0
413.0
2.0
0.0
0.0
0.0
0.0
2247.0
1369.0
4216.0
1375.0
73.0
1051.0
0.0
247.0
0.0
0.0
15366.0
9893.0
0.0
0.0
0.0
3209.0
1051.0
0.0
7138.0
0.0
0.0
1041.0
3630.0

0.87
0.87
0.87
0.87
0.87
0.87
0.87
0.87
0.87
0.87
0.87
0.87
0.87
0.87
0.87
0.87
0.87
0.87
0.87
0.87
0.87
0.87
0.87
0.87
0.87
0.87
0.87
0.87
0.87
0.87


28394.0
66.0
0.0
0.0
0.0
0.0
0.0
1691.0
854.0
2589.0
687.0
0.0
1051.0
0.0
0.0
0.0
0.0
2333.0
9893.0
0.0
0.0
0.0
0.0
1051.0
0.0
7138.0
0.0
0.0
1041.0
0.0


28305.0
66.0
0.0
0.0
0.0
0.0
0.0
1691.0
854.0
2500.0
687.0
0.0
1051.0
0.0
0.0
0.0
0.0
2333.0
9893.0
0.0
0.0
0.0
0.0
1051.0
0.0
7138.0
0.0
0.0
1041.0
0.0

1.0
1.0
1.0
1.0
1.0
1.0
1.0
1.0
1.0
1.0
1.0
1.0
1.0
1.0
1.0
1.0
1.0
1.0
1.0
1.0
1.0
1.0
1.0
1.0
1.0
1.0
1.0
1.0
1.0
1.0


89.0
0.0
0.0
0.0
0.0
0.0
0.0
0.0
0.0
89.0
0.0
0.0
0.0
0.0
0.0
0.0
0.0
0.0
0.0
0.0
0.0
0.0
0.0
0.0
0.0
0.0
0.0
0.0
0.0
0.0

0.78
0.78
0.78
0.78
0.78
0.78
0.78
0.78
0.78
0.78
0.78
0.78
0.78
0.78
0.78
0.78
0.78
0.78
0.78
0.78
0.78
0.78
0.78
0.78
0.78
0.78
0.78
0.78
0.78
0.78


9781.0
224.0
1.0
0.0
0.0
0.0
0.0
110.0
493.0
1479.0
316.0
72.0
0.0
0.0
247.0
0.0
0.0
0.0
0.0
0.0
0.0
0.0
3209.0
0.0
0.0
0.0
0.0
0.0
0.0
3630.0

0.88
0.88
0.88
0.88
0.88
0.88
0.88
0.88
0.88
0.88
0.88
0.88
0.88
0.88
0.88
0.88
0.88
0.88
0.88
0.88
0.88
0.88
0.88
0.88
0.88
0.88
0.88
0.88
0.88
0.88


2003.0
215.0
1.0
0.0
0.0
0.0
0.0
0.0
476.0
1100.0
139.0
72.0
0.0
0.0
0.0
0.0
0.0
0.0
0.0
0.0
0.0
0.0
0.0
0.0
0.0
0.0
0.0
0.0
0.0
0.0

1.0
1.0
1.0
1.0
1.0
1.0
1.0
1.0
1.0
1.0
1.0
1.0
1.0
1.0
1.0
1.0
1.0
1.0
1.0
1.0
1.0
1.0
1.0
1.0
1.0
1.0
1.0
1.0
1.0
1.0


3869.0
9.0
0.0
0.0
0.0
0.0
0.0
110.0
17.0
347.0
177.0
0.0
0.0
0.0
0.0
0.0
0.0
0.0
0.0
0.0
0.0
0.0
3209.0
0.0
0.0
0.0
0.0
0.0
0.0
0.0

1.0
1.0
1.0
1.0
1.0
1.0
1.0
1.0
1.0
1.0
1.0
1.0
1.0
1.0
1.0
1.0
1.0
1.0
1.0
1.0
1.0
1.0
1.0
1.0
1.0
1.0
1.0
1.0
1.0
1.0


32.0
0.0
0.0
0.0
0.0
0.0
0.0
0.0
0.0
32.0
0.0
0.0
0.0
0.0
0.0
0.0
0.0
0.0
0.0
0.0
0.0
0.0
0.0
0.0
0.0
0.0
0.0
0.0
0.0
0.0

1.0
1.0
1.0
1.0
1.0
1.0
1.0
1.0
1.0
1.0
1.0
1.0
1.0
1.0
1.0
1.0
1.0
1.0
1.0
1.0
1.0
1.0
1.0
1.0
1.0
1.0
1.0
1.0
1.0
1.0


3630.0
0.0
0.0
0.0
0.0
0.0
0.0
0.0
0.0
0.0
0.0
0.0
0.0
0.0
0.0
0.0
0.0
0.0
0.0
0.0
0.0
0.0
0.0
0.0
0.0
0.0
0.0
0.0
0.0
3630.0

1.0
1.0
1.0
1.0
1.0
1.0
1.0
1.0
1.0
1.0
1.0
1.0
1.0
1.0
1.0
1.0
1.0
1.0
1.0
1.0
1.0
1.0
1.0
1.0
1.0
1.0
1.0
1.0
1.0
1.0


1041.0
113.0
0.0
0.0
0.0
0.0
0.0
446.0
0.0
118.0
364.0
0.0
0.0
0.0
0.0
0.0
0.0
0.0
0.0
0.0
0.0
0.0
0.0
0.0
0.0
0.0
0.0
0.0
0.0
0.0


1041.0
113.0
0.0
0.0
0.0
0.0
0.0
446.0
0.0
118.0
364.0
0.0
0.0
0.0
0.0
0.0
0.0
0.0
0.0
0.0
0.0
0.0
0.0
0.0
0.0
0.0
0.0
0.0
0.0
0.0

1.0
1.0
1.0
1.0
1.0
1.0
1.0
1.0
1.0
1.0
1.0
1.0
1.0
1.0
1.0
1.0
1.0
1.0
1.0
1.0
1.0
1.0
1.0
1.0
1.0
1.0
1.0
1.0
1.0
1.0


13033.0
0.0
0.0
0.0
0.0
0.0
0.0
0.0
0.0
0.0
0.0
0.0
0.0
0.0
0.0
0.0
0.0
13033.0
0.0
0.0
0.0
0.0
0.0
0.0
0.0
0.0
0.0
0.0
0.0
0.0

1.0
1.0
1.0
1.0
1.0
1.0
1.0
1.0
1.0
1.0
1.0
1.0
1.0
1.0
1.0
1.0
1.0
1.0
1.0
1.0
1.0
1.0
1.0
1.0
1.0
1.0
1.0
1.0
1.0
1.0


25575.0
0.0
0.0
0.0
0.0
0.0
0.0
2703.0
2490.0
6401.0
314.0
0.0
0.0
0.0
0.0
0.0
0.0
890.0
2962.0
4617.0
0.0
5198.0
0.0
0.0
0.0
0.0
0.0
0.0
0.0
0.0


25575.0
0.0
0.0
0.0
0.0
0.0
0.0
2703.0
2490.0
6401.0
314.0
0.0
0.0
0.0
0.0
0.0
0.0
890.0
2962.0
4617.0
0.0
5198.0
0.0
0.0
0.0
0.0
0.0
0.0
0.0
0.0

0.78
0.78
0.78
0.78
0.78
0.78
0.78
0.78
0.78
0.78
0.78
0.78
0.78
0.78
0.78
0.78
0.78
0.78
0.78
0.78
0.78
0.78
0.78
0.78
0.78
0.78
0.78
0.78
0.78
0.78


19136.0
0.0
0.0
0.0
0.0
0.0
0.0
659.0
696.0
5411.0
262.0
0.0
0.0
0.0
0.0
0.0
0.0
890.0
2955.0
4611.0
0.0
3652.0
0.0
0.0
0.0
0.0
0.0
0.0
0.0
0.0

1.0
1.0
1.0
1.0
1.0
1.0
1.0
1.0
1.0
1.0
1.0
1.0
1.0
1.0
1.0
1.0
1.0
1.0
1.0
1.0
1.0
1.0
1.0
1.0
1.0
1.0
1.0
1.0
1.0
1.0


6426.0
0.0
0.0
0.0
0.0
0.0
0.0
2044.0
1794.0
990.0
52.0
0.0
0.0
0.0
0.0
0.0
0.0
0.0
0.0
0.0
0.0
1546.0
0.0
0.0
0.0
0.0
0.0
0.0
0.0
0.0

1.0
1.0
1.0
1.0
1.0
1.0
1.0
1.0
1.0
1.0
1.0
1.0
1.0
1.0
1.0
1.0
1.0
1.0
1.0
1.0
1.0
1.0
1.0
1.0
1.0
1.0
1.0
1.0
1.0
1.0


719882.0
202.0
718.0
33.0
11.0
28.0
0.0
3496.0
3881.0
1728.0
516.0
874.0
43587.0
0.0
0.0
0.0
0.0
6188.0
162965.0
166016.0
931.0
64744.0
219685.0
43587.0
0.0
0.0
0.0
0.0
0.0
692.0

0.88
0.88
0.88
0.88
0.88
0.88
0.88
0.88
0.88
0.88
0.88
0.88
0.88
0.88
0.88
0.88
0.88
0.88
0.88
0.88
0.88
0.88
0.88
0.88
0.88
0.88
0.88
0.88
0.88
0.88


708504.0
0.0
0.0
0.0
0.0
0.0
0.0
3496.0
3197.0
1502.0
0.0
0.0
43587.0
0.0
0.0
0.0
0.0
0.0
162690.0
166016.0
0.0
64744.0
219685.0
43587.0
0.0
0.0
0.0
0.0
0.0
0.0


708504.0
0.0
0.0
0.0
0.0
0.0
0.0
3496.0
3197.0
1502.0
0.0
0.0
43587.0
0.0
0.0
0.0
0.0
0.0
162690.0
166016.0
0.0
64744.0
219685.0
43587.0
0.0
0.0
0.0
0.0
0.0
0.0

1.0
1.0
1.0
1.0
1.0
1.0
1.0
1.0
1.0
1.0
1.0
1.0
1.0
1.0
1.0
1.0
1.0
1.0
1.0
1.0
1.0
1.0
1.0
1.0
1.0
1.0
1.0
1.0
1.0
1.0


10518.0
202.0
718.0
33.0
0.0
0.0
0.0
0.0
0.0
89.0
516.0
874.0
0.0
0.0
0.0
0.0
0.0
6188.0
275.0
0.0
931.0
0.0
0.0
0.0
0.0
0.0
0.0
0.0
0.0
692.0


10518.0
202.0
718.0
33.0
0.0
0.0
0.0
0.0
0.0
89.0
516.0
874.0
0.0
0.0
0.0
0.0
0.0
6188.0
275.0
0.0
931.0
0.0
0.0
0.0
0.0
0.0
0.0
0.0
0.0
692.0

1.0
1.0
1.0
1.0
1.0
1.0
1.0
1.0
1.0
1.0
1.0
1.0
1.0
1.0
1.0
1.0
1.0
1.0
1.0
1.0
1.0
1.0
1.0
1.0
1.0
1.0
1.0
1.0
1.0
1.0


821.0
0.0
0.0
0.0
0.0
0.0
0.0
0.0
684.0
137.0
0.0
0.0
0.0
0.0
0.0
0.0
0.0
0.0
0.0
0.0
0.0
0.0
0.0
0.0
0.0
0.0
0.0
0.0
0.0
0.0


821.0
0.0
0.0
0.0
0.0
0.0
0.0
0.0
684.0
137.0
0.0
0.0
0.0
0.0
0.0
0.0
0.0
0.0
0.0
0.0
0.0
0.0
0.0
0.0
0.0
0.0
0.0
0.0
0.0
0.0

1.0
1.0
1.0
1.0
1.0
1.0
1.0
1.0
1.0
1.0
1.0
1.0
1.0
1.0
1.0
1.0
1.0
1.0
1.0
1.0
1.0
1.0
1.0
1.0
1.0
1.0
1.0
1.0
1.0
1.0


306591.0
48.0
11.0
0.0
12.0
0.0
0.0
1448.0
2487.0
1711.0
100.0
139.0
23.0
141.0
61916.0
910.0
5647.0
29866.0
85331.0
12355.0
1251.0
0.0
663.0
23.0
23289.0
8130.0
16209.0
5038.0
11131.0
38712.0

1.0
1.0
1.0
1.0
1.0
1.0
1.0
1.0
1.0
1.0
1.0
1.0
1.0
1.0
1.0
1.0
1.0
1.0
1.0
1.0
1.0
1.0
1.0
1.0
1.0
1.0
1.0
1.0
1.0
1.0


44609.0
28.0
0.0
0.0
0.0
0.0
0.0
655.0
677.0
726.0
11.0
50.0
18.0
0.0
0.0
0.0
0.0
25041.0
9966.0
277.0
0.0
0.0
0.0
18.0
0.0
289.0
0.0
115.0
2219.0
4519.0


44609.0
28.0
0.0
0.0
0.0
0.0
0.0
655.0
677.0
726.0
11.0
50.0
18.0
0.0
0.0
0.0
0.0
25041.0
9966.0
277.0
0.0
0.0
0.0
18.0
0.0
289.0
0.0
115.0
2219.0
4519.0

1.0
1.0
1.0
1.0
1.0
1.0
1.0
1.0
1.0
1.0
1.0
1.0
1.0
1.0
1.0
1.0
1.0
1.0
1.0
1.0
1.0
1.0
1.0
1.0
1.0
1.0
1.0
1.0
1.0
1.0


640.0
2.0
0.0
0.0
0.0
0.0
0.0
61.0
402.0
157.0
14.0
4.0
0.0
0.0
0.0
0.0
0.0
0.0
0.0
0.0
0.0
0.0
0.0
0.0
0.0
0.0
0.0
0.0
0.0
0.0

1.0
1.0
1.0
1.0
1.0
1.0
1.0
1.0
1.0
1.0
1.0
1.0
1.0
1.0
1.0
1.0
1.0
1.0
1.0
1.0
1.0
1.0
1.0
1.0
1.0
1.0
1.0
1.0
1.0
1.0


111.0
4.0
0.0
0.0
0.0
0.0
0.0
53.0
2.0
33.0
16.0
3.0
0.0
0.0
0.0
0.0
0.0
0.0
0.0
0.0
0.0
0.0
0.0
0.0
0.0
0.0
0.0
0.0
0.0
0.0

1.0
1.0
1.0
1.0
1.0
1.0
1.0
1.0
1.0
1.0
1.0
1.0
1.0
1.0
1.0
1.0
1.0
1.0
1.0
1.0
1.0
1.0
1.0
1.0
1.0
1.0
1.0
1.0
1.0
1.0


85.0
8.0
7.0
0.0
0.0
0.0
0.0
0.0
0.0
0.0
23.0
47.0
0.0
0.0
0.0
0.0
0.0
0.0
0.0
0.0
0.0
0.0
0.0
0.0
0.0
0.0
0.0
0.0
0.0
0.0


85.0
8.0
7.0
0.0
0.0
0.0
0.0
0.0
0.0
0.0
23.0
47.0
0.0
0.0
0.0
0.0
0.0
0.0
0.0
0.0
0.0
0.0
0.0
0.0
0.0
0.0
0.0
0.0
0.0
0.0

1.0
1.0
1.0
1.0
1.0
1.0
1.0
1.0
1.0
1.0
1.0
1.0
1.0
1.0
1.0
1.0
1.0
1.0
1.0
1.0
1.0
1.0
1.0
1.0
1.0
1.0
1.0
1.0
1.0
1.0


43.0
0.0
0.0
0.0
0.0
0.0
0.0
0.0
43.0
0.0
0.0
0.0
0.0
0.0
0.0
0.0
0.0
0.0
0.0
0.0
0.0
0.0
0.0
0.0
0.0
0.0
0.0
0.0
0.0
0.0


43.0
0.0
0.0
0.0
0.0
0.0
0.0
0.0
43.0
0.0
0.0
0.0
0.0
0.0
0.0
0.0
0.0
0.0
0.0
0.0
0.0
0.0
0.0
0.0
0.0
0.0
0.0
0.0
0.0
0.0

0.92
0.92
0.92
0.92
0.92
0.92
0.92
0.92
0.92
0.92
0.92
0.92
0.92
0.92
0.92
0.92
0.92
0.92
0.92
0.92
0.92
0.92
0.92
0.92
0.92
0.92
0.92
0.92
0.92
0.92


387.0
0.0
0.0
0.0
0.0
0.0
0.0
20.0
0.0
0.0
0.0
0.0
0.0
0.0
336.0
0.0
0.0
0.0
31.0
0.0
0.0
0.0
0.0
0.0
0.0
0.0
0.0
0.0
0.0
0.0


387.0
0.0
0.0
0.0
0.0
0.0
0.0
20.0
0.0
0.0
0.0
0.0
0.0
0.0
336.0
0.0
0.0
0.0
31.0
0.0
0.0
0.0
0.0
0.0
0.0
0.0
0.0
0.0
0.0
0.0

1.0
1.0
1.0
1.0
1.0
1.0
1.0
1.0
1.0
1.0
1.0
1.0
1.0
1.0
1.0
1.0
1.0
1.0
1.0
1.0
1.0
1.0
1.0
1.0
1.0
1.0
1.0
1.0
1.0
1.0


12.0
0.0
0.0
0.0
0.0
0.0
0.0
0.0
12.0
0.0
0.0
0.0
0.0
0.0
0.0
0.0
0.0
0.0
0.0
0.0
0.0
0.0
0.0
0.0
0.0
0.0
0.0
0.0
0.0
0.0

0.94
0.94
0.94
0.94
0.94
0.94
0.94
0.94
0.94
0.94
0.94
0.94
0.94
0.94
0.94
0.94
0.94
0.94
0.94
0.94
0.94
0.94
0.94
0.94
0.94
0.94
0.94
0.94
0.94
0.94


15532.0
0.0
0.0
0.0
0.0
0.0
0.0
0.0
0.0
0.0
0.0
0.0
0.0
0.0
0.0
0.0
0.0
0.0
15532.0
0.0
0.0
0.0
0.0
0.0
0.0
0.0
0.0
0.0
0.0
0.0

0.93
0.93
0.93
0.93
0.93
0.93
0.93
0.93
0.93
0.93
0.93
0.93
0.93
0.93
0.93
0.93
0.93
0.93
0.93
0.93
0.93
0.93
0.93
0.93
0.93
0.93
0.93
0.93
0.93
0.93


8941.0
0.0
0.0
0.0
0.0
0.0
0.0
0.0
0.0
0.0
0.0
0.0
0.0
0.0
0.0
0.0
0.0
51.0
0.0
8890.0
0.0
0.0
0.0
0.0
0.0
0.0
0.0
0.0
0.0
0.0

0.95
0.95
0.95
0.95
0.95
0.95
0.95
0.95
0.95
0.95
0.95
0.95
0.95
0.95
0.95
0.95
0.95
0.95
0.95
0.95
0.95
0.95
0.95
0.95
0.95
0.95
0.95
0.95
0.95
0.95


3800.0
0.0
0.0
0.0
0.0
0.0
0.0
0.0
0.0
0.0
0.0
0.0
0.0
0.0
0.0
572.0
0.0
0.0
0.0
0.0
0.0
0.0
0.0
0.0
0.0
0.0
0.0
0.0
1966.0
1262.0

0.97
0.97
0.97
0.97
0.97
0.97
0.97
0.97
0.97
0.97
0.97
0.97
0.97
0.97
0.97
0.97
0.97
0.97
0.97
0.97
0.97
0.97
0.97
0.97
0.97
0.97
0.97
0.97
0.97
0.97


1096.0
0.0
0.0
0.0
0.0
0.0
0.0
0.0
0.0
0.0
0.0
0.0
0.0
0.0
0.0
0.0
0.0
0.0
36.0
1060.0
0.0
0.0
0.0
0.0
0.0
0.0
0.0
0.0
0.0
0.0

0.91
0.91
0.91
0.91
0.91
0.91
0.91
0.91
0.91
0.91
0.91
0.91
0.91
0.91
0.91
0.91
0.91
0.91
0.91
0.91
0.91
0.91
0.91
0.91
0.91
0.91
0.91
0.91
0.91
0.91


780.0
0.0
0.0
0.0
0.0
0.0
0.0
0.0
0.0
0.0
0.0
0.0
0.0
0.0
0.0
0.0
0.0
686.0
0.0
0.0
0.0
0.0
0.0
0.0
0.0
17.0
0.0
0.0
0.0
77.0


780.0
0.0
0.0
0.0
0.0
0.0
0.0
0.0
0.0
0.0
0.0
0.0
0.0
0.0
0.0
0.0
0.0
686.0
0.0
0.0
0.0
0.0
0.0
0.0
0.0
17.0
0.0
0.0
0.0
77.0

0.98
0.98
0.98
0.98
0.98
0.98
0.98
0.98
0.98
0.98
0.98
0.98
0.98
0.98
0.98
0.98
0.98
0.98
0.98
0.98
0.98
0.98
0.98
0.98
0.98
0.98
0.98
0.98
0.98
0.98


875.0
0.0
0.0
0.0
0.0
0.0
0.0
0.0
0.0
0.0
0.0
0.0
0.0
25.0
0.0
0.0
0.0
0.0
0.0
0.0
0.0
0.0
0.0
0.0
103.0
366.0
0.0
93.0
0.0
288.0

1.0
1.0
1.0
1.0
1.0
1.0
1.0
1.0
1.0
1.0
1.0
1.0
1.0
1.0
1.0
1.0
1.0
1.0
1.0
1.0
1.0
1.0
1.0
1.0
1.0
1.0
1.0
1.0
1.0
1.0


629.0
0.0
0.0
0.0
0.0
0.0
0.0
0.0
0.0
0.0
0.0
0.0
0.0
0.0
0.0
8.0
0.0
0.0
101.0
0.0
0.0
0.0
0.0
0.0
0.0
453.0
18.0
46.0
3.0
0.0


629.0
0.0
0.0
0.0
0.0
0.0
0.0
0.0
0.0
0.0
0.0
0.0
0.0
0.0
0.0
8.0
0.0
0.0
101.0
0.0
0.0
0.0
0.0
0.0
0.0
453.0
18.0
46.0
3.0
0.0

1.0
1.0
1.0
1.0
1.0
1.0
1.0
1.0
1.0
1.0
1.0
1.0
1.0
1.0
1.0
1.0
1.0
1.0
1.0
1.0
1.0
1.0
1.0
1.0
1.0
1.0
1.0
1.0
1.0
1.0


504.0
0.0
0.0
0.0
0.0
0.0
0.0
0.0
0.0
0.0
0.0
0.0
0.0
0.0
0.0
0.0
0.0
0.0
0.0
0.0
0.0
0.0
0.0
0.0
0.0
0.0
0.0
0.0
0.0
504.0


504.0
0.0
0.0
0.0
0.0
0.0
0.0
0.0
0.0
0.0
0.0
0.0
0.0
0.0
0.0
0.0
0.0
0.0
0.0
0.0
0.0
0.0
0.0
0.0
0.0
0.0
0.0
0.0
0.0
504.0

0.99
0.99
0.99
0.99
0.99
0.99
0.99
0.99
0.99
0.99
0.99
0.99
0.99
0.99
0.99
0.99
0.99
0.99
0.99
0.99
0.99
0.99
0.99
0.99
0.99
0.99
0.99
0.99
0.99
0.99


431.0
0.0
0.0
0.0
0.0
0.0
0.0
0.0
0.0
0.0
0.0
0.0
0.0
0.0
0.0
0.0
0.0
0.0
0.0
0.0
0.0
0.0
0.0
0.0
0.0
5.0
76.0
12.0
0.0
338.0

1.0
1.0
1.0
1.0
1.0
1.0
1.0
1.0
1.0
1.0
1.0
1.0
1.0
1.0
1.0
1.0
1.0
1.0
1.0
1.0
1.0
1.0
1.0
1.0
1.0
1.0
1.0
1.0
1.0
1.0


326.0
0.0
0.0
0.0
0.0
0.0
0.0
0.0
0.0
0.0
0.0
0.0
0.0
0.0
0.0
0.0
0.0
18.0
308.0
0.0
0.0
0.0
0.0
0.0
0.0
0.0
0.0
0.0
0.0
0.0

0.75
0.75
0.75
0.75
0.75
0.75
0.75
0.75
0.75
0.75
0.75
0.75
0.75
0.75
0.75
0.75
0.75
0.75
0.75
0.75
0.75
0.75
0.75
0.75
0.75
0.75
0.75
0.75
0.75
0.75


301.0
0.0
0.0
0.0
0.0
0.0
0.0
0.0
0.0
0.0
0.0
0.0
0.0
0.0
0.0
0.0
0.0
301.0
0.0
0.0
0.0
0.0
0.0
0.0
0.0
0.0
0.0
0.0
0.0
0.0

0.71
0.71
0.71
0.71
0.71
0.71
0.71
0.71
0.71
0.71
0.71
0.71
0.71
0.71
0.71
0.71
0.71
0.71
0.71
0.71
0.71
0.71
0.71
0.71
0.71
0.71
0.71
0.71
0.71
0.71


231.0
0.0
0.0
0.0
0.0
0.0
0.0
0.0
0.0
0.0
0.0
0.0
0.0
0.0
0.0
0.0
0.0
0.0
0.0
221.0
0.0
0.0
0.0
0.0
0.0
0.0
0.0
0.0
0.0
10.0


221.0
0.0
0.0
0.0
0.0
0.0
0.0
0.0
0.0
0.0
0.0
0.0
0.0
0.0
0.0
0.0
0.0
0.0
0.0
221.0
0.0
0.0
0.0
0.0
0.0
0.0
0.0
0.0
0.0
0.0

1.0
1.0
1.0
1.0
1.0
1.0
1.0
1.0
1.0
1.0
1.0
1.0
1.0
1.0
1.0
1.0
1.0
1.0
1.0
1.0
1.0
1.0
1.0
1.0
1.0
1.0
1.0
1.0
1.0
1.0


10.0
0.0
0.0
0.0
0.0
0.0
0.0
0.0
0.0
0.0
0.0
0.0
0.0
0.0
0.0
0.0
0.0
0.0
0.0
0.0
0.0
0.0
0.0
0.0
0.0
0.0
0.0
0.0
0.0
10.0

1.0
1.0
1.0
1.0
1.0
1.0
1.0
1.0
1.0
1.0
1.0
1.0
1.0
1.0
1.0
1.0
1.0
1.0
1.0
1.0
1.0
1.0
1.0
1.0
1.0
1.0
1.0
1.0
1.0
1.0


200.0
0.0
0.0
0.0
0.0
0.0
0.0
0.0
0.0
0.0
0.0
0.0
0.0
41.0
0.0
0.0
0.0
148.0
0.0
0.0
0.0
0.0
0.0
0.0
0.0
0.0
0.0
6.0
0.0
5.0

0.74
0.74
0.74
0.74
0.74
0.74
0.74
0.74
0.74
0.74
0.74
0.74
0.74
0.74
0.74
0.74
0.74
0.74
0.74
0.74
0.74
0.74
0.74
0.74
0.74
0.74
0.74
0.74
0.74
0.74


50.0
0.0
0.0
0.0
0.0
0.0
0.0
0.0
0.0
0.0
0.0
0.0
0.0
0.0
0.0
0.0
25.0
0.0
0.0
0.0
0.0
0.0
0.0
0.0
0.0
0.0
0.0
0.0
0.0
25.0

0.74
0.74
0.74
0.74
0.74
0.74
0.74
0.74
0.74
0.74
0.74
0.74
0.74
0.74
0.74
0.74
0.74
0.74
0.74
0.74
0.74
0.74
0.74
0.74
0.74
0.74
0.74
0.74
0.74
0.74


28.0
0.0
0.0
0.0
0.0
0.0
0.0
0.0
0.0
0.0
0.0
0.0
0.0
0.0
0.0
0.0
0.0
0.0
0.0
0.0
0.0
0.0
0.0
0.0
0.0
0.0
0.0
0.0
0.0
28.0

0.7
0.7
0.7
0.7
0.7
0.7
0.7
0.7
0.7
0.7
0.7
0.7
0.7
0.7
0.7
0.7
0.7
0.7
0.7
0.7
0.7
0.7
0.7
0.7
0.7
0.7
0.7
0.7
0.7
0.7


13.0
0.0
0.0
0.0
0.0
0.0
0.0
0.0
0.0
0.0
0.0
0.0
0.0
0.0
0.0
0.0
0.0
0.0
0.0
0.0
0.0
0.0
0.0
0.0
0.0
0.0
0.0
0.0
0.0
13.0


13.0
0.0
0.0
0.0
0.0
0.0
0.0
0.0
0.0
0.0
0.0
0.0
0.0
0.0
0.0
0.0
0.0
0.0
0.0
0.0
0.0
0.0
0.0
0.0
0.0
0.0
0.0
0.0
0.0
13.0

1.0
1.0
1.0
1.0
1.0
1.0
1.0
1.0
1.0
1.0
1.0
1.0
1.0
1.0
1.0
1.0
1.0
1.0
1.0
1.0
1.0
1.0
1.0
1.0
1.0
1.0
1.0
1.0
1.0
1.0


4.0
0.0
0.0
0.0
0.0
0.0
0.0
0.0
0.0
0.0
0.0
0.0
0.0
0.0
0.0
0.0
0.0
4.0
0.0
0.0
0.0
0.0
0.0
0.0
0.0
0.0
0.0
0.0
0.0
0.0

0.93
0.93
0.93
0.93
0.93
0.93
0.93
0.93
0.93
0.93
0.93
0.93
0.93
0.93
0.93
0.93
0.93
0.93
0.93
0.93
0.93
0.93
0.93
0.93
0.93
0.93
0.93
0.93
0.93
0.93


1884.0
0.0
1884.0
0.0
0.0
0.0
0.0
0.0
0.0
0.0
0.0
0.0
0.0
0.0
0.0
0.0
0.0
0.0
0.0
0.0
0.0
0.0
0.0
0.0
0.0
0.0
0.0
0.0
0.0
0.0


1884.0
0.0
1884.0
0.0
0.0
0.0
0.0
0.0
0.0
0.0
0.0
0.0
0.0
0.0
0.0
0.0
0.0
0.0
0.0
0.0
0.0
0.0
0.0
0.0
0.0
0.0
0.0
0.0
0.0
0.0


1884.0
0.0
1884.0
0.0
0.0
0.0
0.0
0.0
0.0
0.0
0.0
0.0
0.0
0.0
0.0
0.0
0.0
0.0
0.0
0.0
0.0
0.0
0.0
0.0
0.0
0.0
0.0
0.0
0.0
0.0

1.0
1.0
1.0
1.0
1.0
1.0
1.0
1.0
1.0
1.0
1.0
1.0
1.0
1.0
1.0
1.0
1.0
1.0
1.0
1.0
1.0
1.0
1.0
1.0
1.0
1.0
1.0
1.0
1.0
1.0


54057.0
42.0
71.0
73.0
32.0
74.0
121.0
31.0
841.0
913.0
1388.0
428.0
0.0
0.0
22578.0
135.0
0.0
22.0
0.0
0.0
0.0
0.0
0.0
0.0
238.0
11468.0
5501.0
9462.0
52.0
587.0

0.93
0.93
0.93
0.93
0.93
0.93
0.93
0.93
0.93
0.93
0.93
0.93
0.93
0.93
0.93
0.93
0.93
0.93
0.93
0.93
0.93
0.93
0.93
0.93
0.93
0.93
0.93
0.93
0.93
0.93


4584.0
42.0
22.0
0.0
0.0
0.0
0.0
31.0
841.0
818.0
1302.0
178.0
0.0
0.0
1350.0
0.0
0.0
0.0
0.0
0.0
0.0
0.0
0.0
0.0
0.0
0.0
0.0
0.0
0.0
0.0


3421.0
38.0
22.0
0.0
0.0
0.0
0.0
31.0
791.0
772.0
1222.0
156.0
0.0
0.0
389.0
0.0
0.0
0.0
0.0
0.0
0.0
0.0
0.0
0.0
0.0
0.0
0.0
0.0
0.0
0.0

1.0
1.0
1.0
1.0
1.0
1.0
1.0
1.0
1.0
1.0
1.0
1.0
1.0
1.0
1.0
1.0
1.0
1.0
1.0
1.0
1.0
1.0
1.0
1.0
1.0
1.0
1.0
1.0
1.0
1.0


126.0
0.0
0.0
0.0
0.0
0.0
0.0
0.0
50.0
13.0
59.0
4.0
0.0
0.0
0.0
0.0
0.0
0.0
0.0
0.0
0.0
0.0
0.0
0.0
0.0
0.0
0.0
0.0
0.0
0.0

1.0
1.0
1.0
1.0
1.0
1.0
1.0
1.0
1.0
1.0
1.0
1.0
1.0
1.0
1.0
1.0
1.0
1.0
1.0
1.0
1.0
1.0
1.0
1.0
1.0
1.0
1.0
1.0
1.0
1.0


33.0
0.0
0.0
0.0
0.0
0.0
0.0
0.0
0.0
33.0
0.0
0.0
0.0
0.0
0.0
0.0
0.0
0.0
0.0
0.0
0.0
0.0
0.0
0.0
0.0
0.0
0.0
0.0
0.0
0.0

1.0
1.0
1.0
1.0
1.0
1.0
1.0
1.0
1.0
1.0
1.0
1.0
1.0
1.0
1.0
1.0
1.0
1.0
1.0
1.0
1.0
1.0
1.0
1.0
1.0
1.0
1.0
1.0
1.0
1.0


21.0
3.0
0.0
0.0
0.0
0.0
0.0
0.0
0.0
0.0
0.0
18.0
0.0
0.0
0.0
0.0
0.0
0.0
0.0
0.0
0.0
0.0
0.0
0.0
0.0
0.0
0.0
0.0
0.0
0.0

1.0
1.0
1.0
1.0
1.0
1.0
1.0
1.0
1.0
1.0
1.0
1.0
1.0
1.0
1.0
1.0
1.0
1.0
1.0
1.0
1.0
1.0
1.0
1.0
1.0
1.0
1.0
1.0
1.0
1.0


972.0
1.0
0.0
0.0
0.0
0.0
0.0
0.0
0.0
0.0
10.0
0.0
0.0
0.0
961.0
0.0
0.0
0.0
0.0
0.0
0.0
0.0
0.0
0.0
0.0
0.0
0.0
0.0
0.0
0.0

1.0
1.0
1.0
1.0
1.0
1.0
1.0
1.0
1.0
1.0
1.0
1.0
1.0
1.0
1.0
1.0
1.0
1.0
1.0
1.0
1.0
1.0
1.0
1.0
1.0
1.0
1.0
1.0
1.0
1.0


11.0
0.0
0.0
0.0
0.0
0.0
0.0
0.0
0.0
0.0
11.0
0.0
0.0
0.0
0.0
0.0
0.0
0.0
0.0
0.0
0.0
0.0
0.0
0.0
0.0
0.0
0.0
0.0
0.0
0.0

1.0
1.0
1.0
1.0
1.0
1.0
1.0
1.0
1.0
1.0
1.0
1.0
1.0
1.0
1.0
1.0
1.0
1.0
1.0
1.0
1.0
1.0
1.0
1.0
1.0
1.0
1.0
1.0
1.0
1.0


8093.0
0.0
48.0
43.0
11.0
8.0
93.0
0.0
0.0
0.0
0.0
76.0
0.0
0.0
0.0
135.0
0.0
22.0
0.0
0.0
0.0
0.0
0.0
0.0
0.0
5171.0
1207.0
640.0
52.0
587.0


8093.0
0.0
48.0
43.0
11.0
8.0
93.0
0.0
0.0
0.0
0.0
76.0
0.0
0.0
0.0
135.0
0.0
22.0
0.0
0.0
0.0
0.0
0.0
0.0
0.0
5171.0
1207.0
640.0
52.0
587.0

1.0
1.0
1.0
1.0
1.0
1.0
1.0
1.0
1.0
1.0
1.0
1.0
1.0
1.0
1.0
1.0
1.0
1.0
1.0
1.0
1.0
1.0
1.0
1.0
1.0
1.0
1.0
1.0
1.0
1.0


355.0
0.0
0.0
0.0
0.0
0.0
0.0
0.0
0.0
95.0
86.0
174.0
0.0
0.0
0.0
0.0
0.0
0.0
0.0
0.0
0.0
0.0
0.0
0.0
0.0
0.0
0.0
0.0
0.0
0.0


355.0
0.0
0.0
0.0
0.0
0.0
0.0
0.0
0.0
95.0
86.0
174.0
0.0
0.0
0.0
0.0
0.0
0.0
0.0
0.0
0.0
0.0
0.0
0.0
0.0
0.0
0.0
0.0
0.0
0.0

1.0
1.0
1.0
1.0
1.0
1.0
1.0
1.0
1.0
1.0
1.0
1.0
1.0
1.0
1.0
1.0
1.0
1.0
1.0
1.0
1.0
1.0
1.0
1.0
1.0
1.0
1.0
1.0
1.0
1.0


20922.0
0.0
1.0
30.0
21.0
66.0
28.0
0.0
0.0
0.0
0.0
0.0
0.0
0.0
1125.0
0.0
0.0
0.0
0.0
0.0
0.0
0.0
0.0
0.0
238.0
6297.0
4294.0
8822.0
0.0
0.0


8899.0
0.0
1.0
22.0
17.0
48.0
28.0
0.0
0.0
0.0
0.0
0.0
0.0
0.0
0.0
0.0
0.0
0.0
0.0
0.0
0.0
0.0
0.0
0.0
238.0
614.0
2628.0
5303.0
0.0
0.0

1.0
1.0
1.0
1.0
1.0
1.0
1.0
1.0
1.0
1.0
1.0
1.0
1.0
1.0
1.0
1.0
1.0
1.0
1.0
1.0
1.0
1.0
1.0
1.0
1.0
1.0
1.0
1.0
1.0
1.0


10998.0
0.0
0.0
8.0
4.0
18.0
0.0
0.0
0.0
0.0
0.0
0.0
0.0
0.0
1125.0
0.0
0.0
0.0
0.0
0.0
0.0
0.0
0.0
0.0
0.0
4748.0
1576.0
3519.0
0.0
0.0

1.0
1.0
1.0
1.0
1.0
1.0
1.0
1.0
1.0
1.0
1.0
1.0
1.0
1.0
1.0
1.0
1.0
1.0
1.0
1.0
1.0
1.0
1.0
1.0
1.0
1.0
1.0
1.0
1.0
1.0


1025.0
0.0
0.0
0.0
0.0
0.0
0.0
0.0
0.0
0.0
0.0
0.0
0.0
0.0
0.0
0.0
0.0
0.0
0.0
0.0
0.0
0.0
0.0
0.0
0.0
935.0
90.0
0.0
0.0
0.0

1.0
1.0
1.0
1.0
1.0
1.0
1.0
1.0
1.0
1.0
1.0
1.0
1.0
1.0
1.0
1.0
1.0
1.0
1.0
1.0
1.0
1.0
1.0
1.0
1.0
1.0
1.0
1.0
1.0
1.0


19565.0
0.0
0.0
0.0
0.0
0.0
0.0
0.0
0.0
0.0
0.0
0.0
0.0
0.0
19565.0
0.0
0.0
0.0
0.0
0.0
0.0
0.0
0.0
0.0
0.0
0.0
0.0
0.0
0.0
0.0


19565.0
0.0
0.0
0.0
0.0
0.0
0.0
0.0
0.0
0.0
0.0
0.0
0.0
0.0
19565.0
0.0
0.0
0.0
0.0
0.0
0.0
0.0
0.0
0.0
0.0
0.0
0.0
0.0
0.0
0.0

1.0
1.0
1.0
1.0
1.0
1.0
1.0
1.0
1.0
1.0
1.0
1.0
1.0
1.0
1.0
1.0
1.0
1.0
1.0
1.0
1.0
1.0
1.0
1.0
1.0
1.0
1.0
1.0
1.0
1.0


122786.0
153.0
313.0
292.0
327.0
324.0
138.0
0.0
0.0
88.0
1711.0
523.0
0.0
24884.0
21242.0
0.0
0.0
0.0
8234.0
0.0
13321.0
0.0
0.0
0.0
35118.0
0.0
11839.0
0.0
0.0
4279.0

1.0
1.0
1.0
1.0
1.0
1.0
1.0
1.0
1.0
1.0
1.0
1.0
1.0
1.0
1.0
1.0
1.0
1.0
1.0
1.0
1.0
1.0
1.0
1.0
1.0
1.0
1.0
1.0
1.0
1.0


74877.0
12.0
195.0
213.0
319.0
299.0
133.0
0.0
0.0
0.0
226.0
172.0
0.0
22409.0
14543.0
0.0
0.0
0.0
0.0
0.0
0.0
0.0
0.0
0.0
34935.0
0.0
0.0
0.0
0.0
1421.0

0.99
0.99
0.99
0.99
0.99
0.99
0.99
0.99
0.99
0.99
0.99
0.99
0.99
0.99
0.99
0.99
0.99
0.99
0.99
0.99
0.99
0.99
0.99
0.99
0.99
0.99
0.99
0.99
0.99
0.99


59295.0
12.0
195.0
213.0
319.0
299.0
133.0
0.0
0.0
0.0
226.0
172.0
0.0
22362.0
14543.0
0.0
0.0
0.0
0.0
0.0
0.0
0.0
0.0
0.0
20821.0
0.0
0.0
0.0
0.0
0.0

0.95
0.95
0.95
0.95
0.95
0.95
0.95
0.95
0.95
0.95
0.95
0.95
0.95
0.95
0.95
0.95
0.95
0.95
0.95
0.95
0.95
0.95
0.95
0.95
0.95
0.95
0.95
0.95
0.95
0.95


14114.0
0.0
0.0
0.0
0.0
0.0
0.0
0.0
0.0
0.0
0.0
0.0
0.0
0.0
0.0
0.0
0.0
0.0
0.0
0.0
0.0
0.0
0.0
0.0
14114.0
0.0
0.0
0.0
0.0
0.0

1.0
1.0
1.0
1.0
1.0
1.0
1.0
1.0
1.0
1.0
1.0
1.0
1.0
1.0
1.0
1.0
1.0
1.0
1.0
1.0
1.0
1.0
1.0
1.0
1.0
1.0
1.0
1.0
1.0
1.0


1300.0
0.0
0.0
0.0
0.0
0.0
0.0
0.0
0.0
0.0
0.0
0.0
0.0
0.0
0.0
0.0
0.0
0.0
0.0
0.0
0.0
0.0
0.0
0.0
0.0
0.0
0.0
0.0
0.0
1300.0

1.0
1.0
1.0
1.0
1.0
1.0
1.0
1.0
1.0
1.0
1.0
1.0
1.0
1.0
1.0
1.0
1.0
1.0
1.0
1.0
1.0
1.0
1.0
1.0
1.0
1.0
1.0
1.0
1.0
1.0


32898.0
141.0
118.0
79.0
8.0
25.0
5.0
0.0
0.0
88.0
1485.0
351.0
0.0
2397.0
6637.0
0.0
0.0
0.0
8223.0
0.0
13296.0
0.0
0.0
0.0
45.0
0.0
0.0
0.0
0.0
0.0


732.0
28.0
32.0
0.0
8.0
25.0
5.0
0.0
0.0
88.0
368.0
133.0
0.0
0.0
0.0
0.0
0.0
0.0
0.0
0.0
0.0
0.0
0.0
0.0
45.0
0.0
0.0
0.0
0.0
0.0

0.98
0.98
0.98
0.98
0.98
0.98
0.98
0.98
0.98
0.98
0.98
0.98
0.98
0.98
0.98
0.98
0.98
0.98
0.98
0.98
0.98
0.98
0.98
0.98
0.98
0.98
0.98
0.98
0.98
0.98


1134.0
76.0
48.0
25.0
0.0
0.0
0.0
0.0
0.0
0.0
862.0
123.0
0.0
0.0
0.0
0.0
0.0
0.0
0.0
0.0
0.0
0.0
0.0
0.0
0.0
0.0
0.0
0.0
0.0
0.0

1.0
1.0
1.0
1.0
1.0
1.0
1.0
1.0
1.0
1.0
1.0
1.0
1.0
1.0
1.0
1.0
1.0
1.0
1.0
1.0
1.0
1.0
1.0
1.0
1.0
1.0
1.0
1.0
1.0
1.0


29805.0
11.0
18.0
0.0
0.0
0.0
0.0
0.0
0.0
0.0
255.0
52.0
0.0
1313.0
6637.0
0.0
0.0
0.0
8223.0
0.0
13296.0
0.0
0.0
0.0
0.0
0.0
0.0
0.0
0.0
0.0

1.0
1.0
1.0
1.0
1.0
1.0
1.0
1.0
1.0
1.0
1.0
1.0
1.0
1.0
1.0
1.0
1.0
1.0
1.0
1.0
1.0
1.0
1.0
1.0
1.0
1.0
1.0
1.0
1.0
1.0


1227.0
26.0
20.0
54.0
0.0
0.0
0.0
0.0
0.0
0.0
0.0
43.0
0.0
1084.0
0.0
0.0
0.0
0.0
0.0
0.0
0.0
0.0
0.0
0.0
0.0
0.0
0.0
0.0
0.0
0.0

1.0
1.0
1.0
1.0
1.0
1.0
1.0
1.0
1.0
1.0
1.0
1.0
1.0
1.0
1.0
1.0
1.0
1.0
1.0
1.0
1.0
1.0
1.0
1.0
1.0
1.0
1.0
1.0
1.0
1.0


138.0
0.0
0.0
0.0
0.0
0.0
0.0
0.0
0.0
0.0
0.0
0.0
0.0
0.0
0.0
0.0
0.0
0.0
0.0
0.0
0.0
0.0
0.0
0.0
138.0
0.0
0.0
0.0
0.0
0.0


138.0
0.0
0.0
0.0
0.0
0.0
0.0
0.0
0.0
0.0
0.0
0.0
0.0
0.0
0.0
0.0
0.0
0.0
0.0
0.0
0.0
0.0
0.0
0.0
138.0
0.0
0.0
0.0
0.0
0.0

1.0
1.0
1.0
1.0
1.0
1.0
1.0
1.0
1.0
1.0
1.0
1.0
1.0
1.0
1.0
1.0
1.0
1.0
1.0
1.0
1.0
1.0
1.0
1.0
1.0
1.0
1.0
1.0
1.0
1.0


24.0
0.0
0.0
0.0
0.0
0.0
0.0
0.0
0.0
0.0
0.0
0.0
0.0
24.0
0.0
0.0
0.0
0.0
0.0
0.0
0.0
0.0
0.0
0.0
0.0
0.0
0.0
0.0
0.0
0.0

1.0
1.0
1.0
1.0
1.0
1.0
1.0
1.0
1.0
1.0
1.0
1.0
1.0
1.0
1.0
1.0
1.0
1.0
1.0
1.0
1.0
1.0
1.0
1.0
1.0
1.0
1.0
1.0
1.0
1.0


46717.0
92.0
2.0
0.0
0.0
0.0
0.0
1399.0
377.0
774.0
245.0
188.0
0.0
0.0
246.0
0.0
0.0
8062.0
4760.0
0.0
314.0
439.0
29066.0
0.0
0.0
0.0
505.0
0.0
0.0
248.0

0.92
0.92
0.92
0.92
0.92
0.92
0.92
0.92
0.92
0.92
0.92
0.92
0.92
0.92
0.92
0.92
0.92
0.92
0.92
0.92
0.92
0.92
0.92
0.92
0.92
0.92
0.92
0.92
0.92
0.92


4392.0
42.0
0.0
0.0
0.0
0.0
0.0
93.0
84.0
68.0
19.0
43.0
0.0
0.0
0.0
0.0
0.0
61.0
3982.0
0.0
0.0
0.0
0.0
0.0
0.0
0.0
0.0
0.0
0.0
0.0

0.99
0.99
0.99
0.99
0.99
0.99
0.99
0.99
0.99
0.99
0.99
0.99
0.99
0.99
0.99
0.99
0.99
0.99
0.99
0.99
0.99
0.99
0.99
0.99
0.99
0.99
0.99
0.99
0.99
0.99


4291.0
18.0
0.0
0.0
0.0
0.0
0.0
93.0
84.0
68.0
19.0
36.0
0.0
0.0
0.0
0.0
0.0
0.0
3973.0
0.0
0.0
0.0
0.0
0.0
0.0
0.0
0.0
0.0
0.0
0.0

1.0
1.0
1.0
1.0
1.0
1.0
1.0
1.0
1.0
1.0
1.0
1.0
1.0
1.0
1.0
1.0
1.0
1.0
1.0
1.0
1.0
1.0
1.0
1.0
1.0
1.0
1.0
1.0
1.0
1.0


31.0
24.0
0.0
0.0
0.0
0.0
0.0
0.0
0.0
0.0
0.0
7.0
0.0
0.0
0.0
0.0
0.0
0.0
0.0
0.0
0.0
0.0
0.0
0.0
0.0
0.0
0.0
0.0
0.0
0.0

1.0
1.0
1.0
1.0
1.0
1.0
1.0
1.0
1.0
1.0
1.0
1.0
1.0
1.0
1.0
1.0
1.0
1.0
1.0
1.0
1.0
1.0
1.0
1.0
1.0
1.0
1.0
1.0
1.0
1.0


712.0
24.0
2.0
0.0
0.0
0.0
0.0
39.0
160.0
357.0
91.0
39.0
0.0
0.0
0.0
0.0
0.0
0.0
0.0
0.0
0.0
0.0
0.0
0.0
0.0
0.0
0.0
0.0
0.0
0.0


712.0
24.0
2.0
0.0
0.0
0.0
0.0
39.0
160.0
357.0
91.0
39.0
0.0
0.0
0.0
0.0
0.0
0.0
0.0
0.0
0.0
0.0
0.0
0.0
0.0
0.0
0.0
0.0
0.0
0.0

1.0
1.0
1.0
1.0
1.0
1.0
1.0
1.0
1.0
1.0
1.0
1.0
1.0
1.0
1.0
1.0
1.0
1.0
1.0
1.0
1.0
1.0
1.0
1.0
1.0
1.0
1.0
1.0
1.0
1.0


3533.0
18.0
0.0
0.0
0.0
0.0
0.0
0.0
0.0
259.0
96.0
50.0
0.0
0.0
0.0
0.0
0.0
3110.0
0.0
0.0
0.0
0.0
0.0
0.0
0.0
0.0
0.0
0.0
0.0
0.0

1.0
1.0
1.0
1.0
1.0
1.0
1.0
1.0
1.0
1.0
1.0
1.0
1.0
1.0
1.0
1.0
1.0
1.0
1.0
1.0
1.0
1.0
1.0
1.0
1.0
1.0
1.0
1.0
1.0
1.0


108.0
1.0
0.0
0.0
0.0
0.0
0.0
0.0
0.0
96.0
0.0
11.0
0.0
0.0
0.0
0.0
0.0
0.0
0.0
0.0
0.0
0.0
0.0
0.0
0.0
0.0
0.0
0.0
0.0
0.0

1.0
1.0
1.0
1.0
1.0
1.0
1.0
1.0
1.0
1.0
1.0
1.0
1.0
1.0
1.0
1.0
1.0
1.0
1.0
1.0
1.0
1.0
1.0
1.0
1.0
1.0
1.0
1.0
1.0
1.0


25.0
0.0
0.0
0.0
0.0
0.0
0.0
0.0
0.0
9.0
16.0
0.0
0.0
0.0
0.0
0.0
0.0
0.0
0.0
0.0
0.0
0.0
0.0
0.0
0.0
0.0
0.0
0.0
0.0
0.0

1.0
1.0
1.0
1.0
1.0
1.0
1.0
1.0
1.0
1.0
1.0
1.0
1.0
1.0
1.0
1.0
1.0
1.0
1.0
1.0
1.0
1.0
1.0
1.0
1.0
1.0
1.0
1.0
1.0
1.0


126.0
0.0
0.0
0.0
0.0
0.0
0.0
59.0
44.0
0.0
23.0
0.0
0.0
0.0
0.0
0.0
0.0
0.0
0.0
0.0
0.0
0.0
0.0
0.0
0.0
0.0
0.0
0.0
0.0
0.0


126.0
0.0
0.0
0.0
0.0
0.0
0.0
59.0
44.0
0.0
23.0
0.0
0.0
0.0
0.0
0.0
0.0
0.0
0.0
0.0
0.0
0.0
0.0
0.0
0.0
0.0
0.0
0.0
0.0
0.0

1.0
1.0
1.0
1.0
1.0
1.0
1.0
1.0
1.0
1.0
1.0
1.0
1.0
1.0
1.0
1.0
1.0
1.0
1.0
1.0
1.0
1.0
1.0
1.0
1.0
1.0
1.0
1.0
1.0
1.0


96.0
0.0
0.0
0.0
0.0
0.0
0.0
0.0
0.0
0.0
0.0
52.0
0.0
0.0
0.0
0.0
0.0
0.0
0.0
0.0
0.0
0.0
0.0
0.0
0.0
0.0
0.0
0.0
0.0
44.0


52.0
0.0
0.0
0.0
0.0
0.0
0.0
0.0
0.0
0.0
0.0
52.0
0.0
0.0
0.0
0.0
0.0
0.0
0.0
0.0
0.0
0.0
0.0
0.0
0.0
0.0
0.0
0.0
0.0
0.0

1.0
1.0
1.0
1.0
1.0
1.0
1.0
1.0
1.0
1.0
1.0
1.0
1.0
1.0
1.0
1.0
1.0
1.0
1.0
1.0
1.0
1.0
1.0
1.0
1.0
1.0
1.0
1.0
1.0
1.0


44.0
0.0
0.0
0.0
0.0
0.0
0.0
0.0
0.0
0.0
0.0
0.0
0.0
0.0
0.0
0.0
0.0
0.0
0.0
0.0
0.0
0.0
0.0
0.0
0.0
0.0
0.0
0.0
0.0
44.0

1.0
1.0
1.0
1.0
1.0
1.0
1.0
1.0
1.0
1.0
1.0
1.0
1.0
1.0
1.0
1.0
1.0
1.0
1.0
1.0
1.0
1.0
1.0
1.0
1.0
1.0
1.0
1.0
1.0
1.0


36.0
0.0
0.0
0.0
0.0
0.0
0.0
0.0
0.0
36.0
0.0
0.0
0.0
0.0
0.0
0.0
0.0
0.0
0.0
0.0
0.0
0.0
0.0
0.0
0.0
0.0
0.0
0.0
0.0
0.0


36.0
0.0
0.0
0.0
0.0
0.0
0.0
0.0
0.0
36.0
0.0
0.0
0.0
0.0
0.0
0.0
0.0
0.0
0.0
0.0
0.0
0.0
0.0
0.0
0.0
0.0
0.0
0.0
0.0
0.0

1.0
1.0
1.0
1.0
1.0
1.0
1.0
1.0
1.0
1.0
1.0
1.0
1.0
1.0
1.0
1.0
1.0
1.0
1.0
1.0
1.0
1.0
1.0
1.0
1.0
1.0
1.0
1.0
1.0
1.0


34.0
0.0
0.0
0.0
0.0
0.0
0.0
0.0
0.0
34.0
0.0
0.0
0.0
0.0
0.0
0.0
0.0
0.0
0.0
0.0
0.0
0.0
0.0
0.0
0.0
0.0
0.0
0.0
0.0
0.0


34.0
0.0
0.0
0.0
0.0
0.0
0.0
0.0
0.0
34.0
0.0
0.0
0.0
0.0
0.0
0.0
0.0
0.0
0.0
0.0
0.0
0.0
0.0
0.0
0.0
0.0
0.0
0.0
0.0
0.0

1.0
1.0
1.0
1.0
1.0
1.0
1.0
1.0
1.0
1.0
1.0
1.0
1.0
1.0
1.0
1.0
1.0
1.0
1.0
1.0
1.0
1.0
1.0
1.0
1.0
1.0
1.0
1.0
1.0
1.0


217.0
8.0
0.0
0.0
0.0
0.0
0.0
0.0
0.0
0.0
16.0
4.0
0.0
0.0
189.0
0.0
0.0
0.0
0.0
0.0
0.0
0.0
0.0
0.0
0.0
0.0
0.0
0.0
0.0
0.0

0.97
0.97
0.97
0.97
0.97
0.97
0.97
0.97
0.97
0.97
0.97
0.97
0.97
0.97
0.97
0.97
0.97
0.97
0.97
0.97
0.97
0.97
0.97
0.97
0.97
0.97
0.97
0.97
0.97
0.97


4911.0
0.0
0.0
0.0
0.0
0.0
0.0
0.0
0.0
20.0
0.0
0.0
0.0
0.0
0.0
0.0
0.0
4891.0
0.0
0.0
0.0
0.0
0.0
0.0
0.0
0.0
0.0
0.0
0.0
0.0


4911.0
0.0
0.0
0.0
0.0
0.0
0.0
0.0
0.0
20.0
0.0
0.0
0.0
0.0
0.0
0.0
0.0
4891.0
0.0
0.0
0.0
0.0
0.0
0.0
0.0
0.0
0.0
0.0
0.0
0.0

1.0
1.0
1.0
1.0
1.0
1.0
1.0
1.0
1.0
1.0
1.0
1.0
1.0
1.0
1.0
1.0
1.0
1.0
1.0
1.0
1.0
1.0
1.0
1.0
1.0
1.0
1.0
1.0
1.0
1.0


710.0
0.0
0.0
0.0
0.0
0.0
0.0
0.0
12.0
0.0
0.0
0.0
0.0
0.0
0.0
0.0
0.0
0.0
0.0
0.0
0.0
0.0
0.0
0.0
0.0
0.0
505.0
0.0
0.0
193.0


710.0
0.0
0.0
0.0
0.0
0.0
0.0
0.0
12.0
0.0
0.0
0.0
0.0
0.0
0.0
0.0
0.0
0.0
0.0
0.0
0.0
0.0
0.0
0.0
0.0
0.0
505.0
0.0
0.0
193.0

1.0
1.0
1.0
1.0
1.0
1.0
1.0
1.0
1.0
1.0
1.0
1.0
1.0
1.0
1.0
1.0
1.0
1.0
1.0
1.0
1.0
1.0
1.0
1.0
1.0
1.0
1.0
1.0
1.0
1.0


29819.0
0.0
0.0
0.0
0.0
0.0
0.0
0.0
0.0
0.0
0.0
0.0
0.0
0.0
0.0
0.0
0.0
0.0
0.0
0.0
314.0
439.0
29066.0
0.0
0.0
0.0
0.0
0.0
0.0
0.0

0.7
0.7
0.7
0.7
0.7
0.7
0.7
0.7
0.7
0.7
0.7
0.7
0.7
0.7
0.7
0.7
0.7
0.7
0.7
0.7
0.7
0.7
0.7
0.7
0.7
0.7
0.7
0.7
0.7
0.7


17566.0
0.0
0.0
0.0
0.0
0.0
0.0
0.0
0.0
0.0
0.0
0.0
0.0
0.0
0.0
0.0
0.0
0.0
0.0
0.0
96.0
405.0
17065.0
0.0
0.0
0.0
0.0
0.0
0.0
0.0

0.86
0.86
0.86
0.86
0.86
0.86
0.86
0.86
0.86
0.86
0.86
0.86
0.86
0.86
0.86
0.86
0.86
0.86
0.86
0.86
0.86
0.86
0.86
0.86
0.86
0.86
0.86
0.86
0.86
0.86


778.0
0.0
0.0
0.0
0.0
0.0
0.0
0.0
0.0
0.0
0.0
0.0
0.0
0.0
0.0
0.0
0.0
0.0
778.0
0.0
0.0
0.0
0.0
0.0
0.0
0.0
0.0
0.0
0.0
0.0


778.0
0.0
0.0
0.0
0.0
0.0
0.0
0.0
0.0
0.0
0.0
0.0
0.0
0.0
0.0
0.0
0.0
0.0
778.0
0.0
0.0
0.0
0.0
0.0
0.0
0.0
0.0
0.0
0.0
0.0

1.0
1.0
1.0
1.0
1.0
1.0
1.0
1.0
1.0
1.0
1.0
1.0
1.0
1.0
1.0
1.0
1.0
1.0
1.0
1.0
1.0
1.0
1.0
1.0
1.0
1.0
1.0
1.0
1.0
1.0


1732.0
36.0
1.0
0.0
24.0
24.0
20.0
298.0
296.0
138.0
255.0
33.0
0.0
0.0
89.0
0.0
0.0
0.0
518.0
0.0
0.0
0.0
0.0
0.0
0.0
0.0
0.0
0.0
0.0
0.0


1387.0
36.0
1.0
0.0
0.0
0.0
0.0
231.0
296.0
48.0
242.0
15.0
0.0
0.0
0.0
0.0
0.0
0.0
518.0
0.0
0.0
0.0
0.0
0.0
0.0
0.0
0.0
0.0
0.0
0.0


1387.0
36.0
1.0
0.0
0.0
0.0
0.0
231.0
296.0
48.0
242.0
15.0
0.0
0.0
0.0
0.0
0.0
0.0
518.0
0.0
0.0
0.0
0.0
0.0
0.0
0.0
0.0
0.0
0.0
0.0

1.0
1.0
1.0
1.0
1.0
1.0
1.0
1.0
1.0
1.0
1.0
1.0
1.0
1.0
1.0
1.0
1.0
1.0
1.0
1.0
1.0
1.0
1.0
1.0
1.0
1.0
1.0
1.0
1.0
1.0


345.0
0.0
0.0
0.0
24.0
24.0
20.0
67.0
0.0
90.0
13.0
18.0
0.0
0.0
89.0
0.0
0.0
0.0
0.0
0.0
0.0
0.0
0.0
0.0
0.0
0.0
0.0
0.0
0.0
0.0


345.0
0.0
0.0
0.0
24.0
24.0
20.0
67.0
0.0
90.0
13.0
18.0
0.0
0.0
89.0
0.0
0.0
0.0
0.0
0.0
0.0
0.0
0.0
0.0
0.0
0.0
0.0
0.0
0.0
0.0

1.0
1.0
1.0
1.0
1.0
1.0
1.0
1.0
1.0
1.0
1.0
1.0
1.0
1.0
1.0
1.0
1.0
1.0
1.0
1.0
1.0
1.0
1.0
1.0
1.0
1.0
1.0
1.0
1.0
1.0


168302.0
49.0
214.0
25.0
122.0
88.0
0.0
0.0
116.0
252.0
1486.0
219.0
0.0
10230.0
2018.0
0.0
0.0
0.0
0.0
49.0
0.0
0.0
0.0
0.0
0.0
103770.0
0.0
0.0
0.0
49664.0

0.93
0.93
0.93
0.93
0.93
0.93
0.93
0.93
0.93
0.93
0.93
0.93
0.93
0.93
0.93
0.93
0.93
0.93
0.93
0.93
0.93
0.93
0.93
0.93
0.93
0.93
0.93
0.93
0.93
0.93


1074.0
0.0
0.0
0.0
0.0
0.0
0.0
0.0
0.0
0.0
1074.0
0.0
0.0
0.0
0.0
0.0
0.0
0.0
0.0
0.0
0.0
0.0
0.0
0.0
0.0
0.0
0.0
0.0
0.0
0.0


1074.0
0.0
0.0
0.0
0.0
0.0
0.0
0.0
0.0
0.0
1074.0
0.0
0.0
0.0
0.0
0.0
0.0
0.0
0.0
0.0
0.0
0.0
0.0
0.0
0.0
0.0
0.0
0.0
0.0
0.0

1.0
1.0
1.0
1.0
1.0
1.0
1.0
1.0
1.0
1.0
1.0
1.0
1.0
1.0
1.0
1.0
1.0
1.0
1.0
1.0
1.0
1.0
1.0
1.0
1.0
1.0
1.0
1.0
1.0
1.0


13691.0
49.0
214.0
25.0
122.0
88.0
0.0
0.0
0.0
0.0
214.0
219.0
0.0
10230.0
2018.0
0.0
0.0
0.0
0.0
49.0
0.0
0.0
0.0
0.0
0.0
463.0
0.0
0.0
0.0
0.0

1.0
1.0
1.0
1.0
1.0
1.0
1.0
1.0
1.0
1.0
1.0
1.0
1.0
1.0
1.0
1.0
1.0
1.0
1.0
1.0
1.0
1.0
1.0
1.0
1.0
1.0
1.0
1.0
1.0
1.0


186.0
0.0
0.0
0.0
0.0
0.0
0.0
0.0
0.0
0.0
186.0
0.0
0.0
0.0
0.0
0.0
0.0
0.0
0.0
0.0
0.0
0.0
0.0
0.0
0.0
0.0
0.0
0.0
0.0
0.0


186.0
0.0
0.0
0.0
0.0
0.0
0.0
0.0
0.0
0.0
186.0
0.0
0.0
0.0
0.0
0.0
0.0
0.0
0.0
0.0
0.0
0.0
0.0
0.0
0.0
0.0
0.0
0.0
0.0
0.0

0.93
0.93
0.93
0.93
0.93
0.93
0.93
0.93
0.93
0.93
0.93
0.93
0.93
0.93
0.93
0.93
0.93
0.93
0.93
0.93
0.93
0.93
0.93
0.93
0.93
0.93
0.93
0.93
0.93
0.93


177.0
0.0
0.0
0.0
0.0
0.0
0.0
0.0
0.0
177.0
0.0
0.0
0.0
0.0
0.0
0.0
0.0
0.0
0.0
0.0
0.0
0.0
0.0
0.0
0.0
0.0
0.0
0.0
0.0
0.0

1.0
1.0
1.0
1.0
1.0
1.0
1.0
1.0
1.0
1.0
1.0
1.0
1.0
1.0
1.0
1.0
1.0
1.0
1.0
1.0
1.0
1.0
1.0
1.0
1.0
1.0
1.0
1.0
1.0
1.0


76.0
0.0
0.0
0.0
0.0
0.0
0.0
0.0
0.0
64.0
12.0
0.0
0.0
0.0
0.0
0.0
0.0
0.0
0.0
0.0
0.0
0.0
0.0
0.0
0.0
0.0
0.0
0.0
0.0
0.0

0.99
0.99
0.99
0.99
0.99
0.99
0.99
0.99
0.99
0.99
0.99
0.99
0.99
0.99
0.99
0.99
0.99
0.99
0.99
0.99
0.99
0.99
0.99
0.99
0.99
0.99
0.99
0.99
0.99
0.99


11.0
0.0
0.0
0.0
0.0
0.0
0.0
0.0
0.0
11.0
0.0
0.0
0.0
0.0
0.0
0.0
0.0
0.0
0.0
0.0
0.0
0.0
0.0
0.0
0.0
0.0
0.0
0.0
0.0
0.0


11.0
0.0
0.0
0.0
0.0
0.0
0.0
0.0
0.0
11.0
0.0
0.0
0.0
0.0
0.0
0.0
0.0
0.0
0.0
0.0
0.0
0.0
0.0
0.0
0.0
0.0
0.0
0.0
0.0
0.0

1.0
1.0
1.0
1.0
1.0
1.0
1.0
1.0
1.0
1.0
1.0
1.0
1.0
1.0
1.0
1.0
1.0
1.0
1.0
1.0
1.0
1.0
1.0
1.0
1.0
1.0
1.0
1.0
1.0
1.0


612.0
0.0
0.0
0.0
0.0
0.0
0.0
0.0
0.0
0.0
0.0
0.0
0.0
0.0
0.0
0.0
0.0
0.0
0.0
0.0
0.0
0.0
0.0
0.0
0.0
612.0
0.0
0.0
0.0
0.0


612.0
0.0
0.0
0.0
0.0
0.0
0.0
0.0
0.0
0.0
0.0
0.0
0.0
0.0
0.0
0.0
0.0
0.0
0.0
0.0
0.0
0.0
0.0
0.0
0.0
612.0
0.0
0.0
0.0
0.0

0.97
0.97
0.97
0.97
0.97
0.97
0.97
0.97
0.97
0.97
0.97
0.97
0.97
0.97
0.97
0.97
0.97
0.97
0.97
0.97
0.97
0.97
0.97
0.97
0.97
0.97
0.97
0.97
0.97
0.97


28607.0
33.0
9.0
0.0
0.0
0.0
0.0
24.0
446.0
923.0
568.0
19.0
10440.0
0.0
703.0
0.0
0.0
270.0
183.0
2551.0
58.0
0.0
800.0
10440.0
0.0
0.0
1140.0
0.0
0.0
0.0

0.98
0.98
0.98
0.98
0.98
0.98
0.98
0.98
0.98
0.98
0.98
0.98
0.98
0.98
0.98
0.98
0.98
0.98
0.98
0.98
0.98
0.98
0.98
0.98
0.98
0.98
0.98
0.98
0.98
0.98


552.0
0.0
0.0
0.0
0.0
0.0
0.0
0.0
151.0
337.0
64.0
0.0
0.0
0.0
0.0
0.0
0.0
0.0
0.0
0.0
0.0
0.0
0.0
0.0
0.0
0.0
0.0
0.0
0.0
0.0


478.0
0.0
0.0
0.0
0.0
0.0
0.0
0.0
122.0
300.0
56.0
0.0
0.0
0.0
0.0
0.0
0.0
0.0
0.0
0.0
0.0
0.0
0.0
0.0
0.0
0.0
0.0
0.0
0.0
0.0

1.0
1.0
1.0
1.0
1.0
1.0
1.0
1.0
1.0
1.0
1.0
1.0
1.0
1.0
1.0
1.0
1.0
1.0
1.0
1.0
1.0
1.0
1.0
1.0
1.0
1.0
1.0
1.0
1.0
1.0


74.0
0.0
0.0
0.0
0.0
0.0
0.0
0.0
29.0
37.0
8.0
0.0
0.0
0.0
0.0
0.0
0.0
0.0
0.0
0.0
0.0
0.0
0.0
0.0
0.0
0.0
0.0
0.0
0.0
0.0

0.99
0.99
0.99
0.99
0.99
0.99
0.99
0.99
0.99
0.99
0.99
0.99
0.99
0.99
0.99
0.99
0.99
0.99
0.99
0.99
0.99
0.99
0.99
0.99
0.99
0.99
0.99
0.99
0.99
0.99


957.0
8.0
9.0
0.0
0.0
0.0
0.0
0.0
0.0
148.0
89.0
0.0
0.0
0.0
703.0
0.0
0.0
0.0
0.0
0.0
0.0
0.0
0.0
0.0
0.0
0.0
0.0
0.0
0.0
0.0


957.0
8.0
9.0
0.0
0.0
0.0
0.0
0.0
0.0
148.0
89.0
0.0
0.0
0.0
703.0
0.0
0.0
0.0
0.0
0.0
0.0
0.0
0.0
0.0
0.0
0.0
0.0
0.0
0.0
0.0

1.0
1.0
1.0
1.0
1.0
1.0
1.0
1.0
1.0
1.0
1.0
1.0
1.0
1.0
1.0
1.0
1.0
1.0
1.0
1.0
1.0
1.0
1.0
1.0
1.0
1.0
1.0
1.0
1.0
1.0


392.0
0.0
0.0
0.0
0.0
0.0
0.0
0.0
126.0
110.0
146.0
10.0
0.0
0.0
0.0
0.0
0.0
0.0
0.0
0.0
0.0
0.0
0.0
0.0
0.0
0.0
0.0
0.0
0.0
0.0


392.0
0.0
0.0
0.0
0.0
0.0
0.0
0.0
126.0
110.0
146.0
10.0
0.0
0.0
0.0
0.0
0.0
0.0
0.0
0.0
0.0
0.0
0.0
0.0
0.0
0.0
0.0
0.0
0.0
0.0

1.0
1.0
1.0
1.0
1.0
1.0
1.0
1.0
1.0
1.0
1.0
1.0
1.0
1.0
1.0
1.0
1.0
1.0
1.0
1.0
1.0
1.0
1.0
1.0
1.0
1.0
1.0
1.0
1.0
1.0


64.0
0.0
0.0
0.0
0.0
0.0
0.0
0.0
64.0
0.0
0.0
0.0
0.0
0.0
0.0
0.0
0.0
0.0
0.0
0.0
0.0
0.0
0.0
0.0
0.0
0.0
0.0
0.0
0.0
0.0


64.0
0.0
0.0
0.0
0.0
0.0
0.0
0.0
64.0
0.0
0.0
0.0
0.0
0.0
0.0
0.0
0.0
0.0
0.0
0.0
0.0
0.0
0.0
0.0
0.0
0.0
0.0
0.0
0.0
0.0

0.96
0.96
0.96
0.96
0.96
0.96
0.96
0.96
0.96
0.96
0.96
0.96
0.96
0.96
0.96
0.96
0.96
0.96
0.96
0.96
0.96
0.96
0.96
0.96
0.96
0.96
0.96
0.96
0.96
0.96


10.0
10.0
0.0
0.0
0.0
0.0
0.0
0.0
0.0
0.0
0.0
0.0
0.0
0.0
0.0
0.0
0.0
0.0
0.0
0.0
0.0
0.0
0.0
0.0
0.0
0.0
0.0
0.0
0.0
0.0


10.0
10.0
0.0
0.0
0.0
0.0
0.0
0.0
0.0
0.0
0.0
0.0
0.0
0.0
0.0
0.0
0.0
0.0
0.0
0.0
0.0
0.0
0.0
0.0
0.0
0.0
0.0
0.0
0.0
0.0

1.0
1.0
1.0
1.0
1.0
1.0
1.0
1.0
1.0
1.0
1.0
1.0
1.0
1.0
1.0
1.0
1.0
1.0
1.0
1.0
1.0
1.0
1.0
1.0
1.0
1.0
1.0
1.0
1.0
1.0


680.0
0.0
0.0
0.0
0.0
0.0
0.0
680.0
0.0
0.0
0.0
0.0
0.0
0.0
0.0
0.0
0.0
0.0
0.0
0.0
0.0
0.0
0.0
0.0
0.0
0.0
0.0
0.0
0.0
0.0


680.0
0.0
0.0
0.0
0.0
0.0
0.0
680.0
0.0
0.0
0.0
0.0
0.0
0.0
0.0
0.0
0.0
0.0
0.0
0.0
0.0
0.0
0.0
0.0
0.0
0.0
0.0
0.0
0.0
0.0

0.75
0.75
0.75
0.75
0.75
0.75
0.75
0.75
0.75
0.75
0.75
0.75
0.75
0.75
0.75
0.75
0.75
0.75
0.75
0.75
0.75
0.75
0.75
0.75
0.75
0.75
0.75
0.75
0.75
0.75


78785.0
28.0
146.0
56.0
112.0
537.0
491.0
180.0
197.0
633.0
470.0
131.0
0.0
0.0
5630.0
0.0
0.0
0.0
8622.0
0.0
477.0
0.0
0.0
0.0
3499.0
39806.0
5250.0
1297.0
2071.0
9152.0

0.85
0.85
0.85
0.85
0.85
0.85
0.85
0.85
0.85
0.85
0.85
0.85
0.85
0.85
0.85
0.85
0.85
0.85
0.85
0.85
0.85
0.85
0.85
0.85
0.85
0.85
0.85
0.85
0.85
0.85


17262.0
0.0
82.0
41.0
112.0
537.0
480.0
0.0
0.0
79.0
19.0
61.0
0.0
0.0
0.0
0.0
0.0
0.0
4628.0
0.0
0.0
0.0
0.0
0.0
343.0
0.0
159.0
208.0
1493.0
9020.0

0.73
0.73
0.73
0.73
0.73
0.73
0.73
0.73
0.73
0.73
0.73
0.73
0.73
0.73
0.73
0.73
0.73
0.73
0.73
0.73
0.73
0.73
0.73
0.73
0.73
0.73
0.73
0.73
0.73
0.73


17246.0
0.0
82.0
41.0
112.0
537.0
480.0
0.0
0.0
79.0
19.0
61.0
0.0
0.0
0.0
0.0
0.0
0.0
4628.0
0.0
0.0
0.0
0.0
0.0
343.0
0.0
159.0
206.0
1493.0
9006.0

1.0
1.0
1.0
1.0
1.0
1.0
1.0
1.0
1.0
1.0
1.0
1.0
1.0
1.0
1.0
1.0
1.0
1.0
1.0
1.0
1.0
1.0
1.0
1.0
1.0
1.0
1.0
1.0
1.0
1.0


1933.0
0.0
0.0
0.0
0.0
0.0
0.0
0.0
0.0
375.0
377.0
58.0
0.0
0.0
0.0
0.0
0.0
0.0
1123.0
0.0
0.0
0.0
0.0
0.0
0.0
0.0
0.0
0.0
0.0
0.0


1933.0
0.0
0.0
0.0
0.0
0.0
0.0
0.0
0.0
375.0
377.0
58.0
0.0
0.0
0.0
0.0
0.0
0.0
1123.0
0.0
0.0
0.0
0.0
0.0
0.0
0.0
0.0
0.0
0.0
0.0

1.0
1.0
1.0
1.0
1.0
1.0
1.0
1.0
1.0
1.0
1.0
1.0
1.0
1.0
1.0
1.0
1.0
1.0
1.0
1.0
1.0
1.0
1.0
1.0
1.0
1.0
1.0
1.0
1.0
1.0


10746.0
19.0
21.0
0.0
0.0
0.0
11.0
180.0
168.0
179.0
47.0
5.0
0.0
0.0
264.0
0.0
0.0
0.0
0.0
0.0
0.0
0.0
0.0
0.0
0.0
8992.0
860.0
0.0
0.0
0.0

0.93
0.93
0.93
0.93
0.93
0.93
0.93
0.93
0.93
0.93
0.93
0.93
0.93
0.93
0.93
0.93
0.93
0.93
0.93
0.93
0.93
0.93
0.93
0.93
0.93
0.93
0.93
0.93
0.93
0.93


10722.0
19.0
21.0
0.0
0.0
0.0
0.0
180.0
168.0
179.0
47.0
5.0
0.0
0.0
264.0
0.0
0.0
0.0
0.0
0.0
0.0
0.0
0.0
0.0
0.0
8979.0
860.0
0.0
0.0
0.0

1.0
1.0
1.0
1.0
1.0
1.0
1.0
1.0
1.0
1.0
1.0
1.0
1.0
1.0
1.0
1.0
1.0
1.0
1.0
1.0
1.0
1.0
1.0
1.0
1.0
1.0
1.0
1.0
1.0
1.0


69.0
4.0
43.0
15.0
0.0
0.0
0.0
0.0
0.0
0.0
0.0
7.0
0.0
0.0
0.0
0.0
0.0
0.0
0.0
0.0
0.0
0.0
0.0
0.0
0.0
0.0
0.0
0.0
0.0
0.0


69.0
4.0
43.0
15.0
0.0
0.0
0.0
0.0
0.0
0.0
0.0
7.0
0.0
0.0
0.0
0.0
0.0
0.0
0.0
0.0
0.0
0.0
0.0
0.0
0.0
0.0
0.0
0.0
0.0
0.0

1.0
1.0
1.0
1.0
1.0
1.0
1.0
1.0
1.0
1.0
1.0
1.0
1.0
1.0
1.0
1.0
1.0
1.0
1.0
1.0
1.0
1.0
1.0
1.0
1.0
1.0
1.0
1.0
1.0
1.0


34.0
5.0
0.0
0.0
0.0
0.0
0.0
0.0
29.0
0.0
0.0
0.0
0.0
0.0
0.0
0.0
0.0
0.0
0.0
0.0
0.0
0.0
0.0
0.0
0.0
0.0
0.0
0.0
0.0
0.0

1.0
1.0
1.0
1.0
1.0
1.0
1.0
1.0
1.0
1.0
1.0
1.0
1.0
1.0
1.0
1.0
1.0
1.0
1.0
1.0
1.0
1.0
1.0
1.0
1.0
1.0
1.0
1.0
1.0
1.0


11858.0
0.0
0.0
0.0
0.0
0.0
0.0
0.0
0.0
0.0
16.0
0.0
0.0
0.0
5349.0
0.0
0.0
0.0
2865.0
0.0
465.0
0.0
0.0
0.0
3134.0
0.0
0.0
0.0
0.0
29.0

0.99
0.99
0.99
0.99
0.99
0.99
0.99
0.99
0.99
0.99
0.99
0.99
0.99
0.99
0.99
0.99
0.99
0.99
0.99
0.99
0.99
0.99
0.99
0.99
0.99
0.99
0.99
0.99
0.99
0.99


2894.0
0.0
0.0
0.0
0.0
0.0
0.0
0.0
0.0
0.0
0.0
0.0
0.0
0.0
0.0
0.0
0.0
0.0
2865.0
0.0
0.0
0.0
0.0
0.0
0.0
0.0
0.0
0.0
0.0
29.0

1.0
1.0
1.0
1.0
1.0
1.0
1.0
1.0
1.0
1.0
1.0
1.0
1.0
1.0
1.0
1.0
1.0
1.0
1.0
1.0
1.0
1.0
1.0
1.0
1.0
1.0
1.0
1.0
1.0
1.0


96.0
0.0
0.0
0.0
0.0
0.0
0.0
0.0
0.0
0.0
0.0
0.0
0.0
0.0
0.0
0.0
0.0
0.0
0.0
0.0
0.0
0.0
0.0
0.0
0.0
69.0
27.0
0.0
0.0
0.0

0.93
0.93
0.93
0.93
0.93
0.93
0.93
0.93
0.93
0.93
0.93
0.93
0.93
0.93
0.93
0.93
0.93
0.93
0.93
0.93
0.93
0.93
0.93
0.93
0.93
0.93
0.93
0.93
0.93
0.93


3068.0
113.0
43.0
38.0
32.0
9.0
103.0
0.0
92.0
205.0
272.0
149.0
0.0
0.0
0.0
0.0
0.0
0.0
0.0
0.0
0.0
0.0
0.0
0.0
0.0
189.0
0.0
0.0
0.0
1823.0

0.9
0.9
0.9
0.9
0.9
0.9
0.9
0.9
0.9
0.9
0.9
0.9
0.9
0.9
0.9
0.9
0.9
0.9
0.9
0.9
0.9
0.9
0.9
0.9
0.9
0.9
0.9
0.9
0.9
0.9


526.0
89.0
43.0
38.0
17.0
0.0
103.0
0.0
0.0
11.0
128.0
97.0
0.0
0.0
0.0
0.0
0.0
0.0
0.0
0.0
0.0
0.0
0.0
0.0
0.0
0.0
0.0
0.0
0.0
0.0


526.0
89.0
43.0
38.0
17.0
0.0
103.0
0.0
0.0
11.0
128.0
97.0
0.0
0.0
0.0
0.0
0.0
0.0
0.0
0.0
0.0
0.0
0.0
0.0
0.0
0.0
0.0
0.0
0.0
0.0

1.0
1.0
1.0
1.0
1.0
1.0
1.0
1.0
1.0
1.0
1.0
1.0
1.0
1.0
1.0
1.0
1.0
1.0
1.0
1.0
1.0
1.0
1.0
1.0
1.0
1.0
1.0
1.0
1.0
1.0


162.0
0.0
0.0
0.0
0.0
0.0
0.0
0.0
39.0
57.0
66.0
0.0
0.0
0.0
0.0
0.0
0.0
0.0
0.0
0.0
0.0
0.0
0.0
0.0
0.0
0.0
0.0
0.0
0.0
0.0


162.0
0.0
0.0
0.0
0.0
0.0
0.0
0.0
39.0
57.0
66.0
0.0
0.0
0.0
0.0
0.0
0.0
0.0
0.0
0.0
0.0
0.0
0.0
0.0
0.0
0.0
0.0
0.0
0.0
0.0

1.0
1.0
1.0
1.0
1.0
1.0
1.0
1.0
1.0
1.0
1.0
1.0
1.0
1.0
1.0
1.0
1.0
1.0
1.0
1.0
1.0
1.0
1.0
1.0
1.0
1.0
1.0
1.0
1.0
1.0


175.0
24.0
0.0
0.0
0.0
0.0
0.0
0.0
53.0
54.0
44.0
0.0
0.0
0.0
0.0
0.0
0.0
0.0
0.0
0.0
0.0
0.0
0.0
0.0
0.0
0.0
0.0
0.0
0.0
0.0


175.0
24.0
0.0
0.0
0.0
0.0
0.0
0.0
53.0
54.0
44.0
0.0
0.0
0.0
0.0
0.0
0.0
0.0
0.0
0.0
0.0
0.0
0.0
0.0
0.0
0.0
0.0
0.0
0.0
0.0

1.0
1.0
1.0
1.0
1.0
1.0
1.0
1.0
1.0
1.0
1.0
1.0
1.0
1.0
1.0
1.0
1.0
1.0
1.0
1.0
1.0
1.0
1.0
1.0
1.0
1.0
1.0
1.0
1.0
1.0


65.0
0.0
0.0
0.0
0.0
0.0
0.0
0.0
0.0
65.0
0.0
0.0
0.0
0.0
0.0
0.0
0.0
0.0
0.0
0.0
0.0
0.0
0.0
0.0
0.0
0.0
0.0
0.0
0.0
0.0


65.0
0.0
0.0
0.0
0.0
0.0
0.0
0.0
0.0
65.0
0.0
0.0
0.0
0.0
0.0
0.0
0.0
0.0
0.0
0.0
0.0
0.0
0.0
0.0
0.0
0.0
0.0
0.0
0.0
0.0

1.0
1.0
1.0
1.0
1.0
1.0
1.0
1.0
1.0
1.0
1.0
1.0
1.0
1.0
1.0
1.0
1.0
1.0
1.0
1.0
1.0
1.0
1.0
1.0
1.0
1.0
1.0
1.0
1.0
1.0


49.0
0.0
0.0
0.0
0.0
0.0
0.0
0.0
0.0
4.0
0.0
45.0
0.0
0.0
0.0
0.0
0.0
0.0
0.0
0.0
0.0
0.0
0.0
0.0
0.0
0.0
0.0
0.0
0.0
0.0


49.0
0.0
0.0
0.0
0.0
0.0
0.0
0.0
0.0
4.0
0.0
45.0
0.0
0.0
0.0
0.0
0.0
0.0
0.0
0.0
0.0
0.0
0.0
0.0
0.0
0.0
0.0
0.0
0.0
0.0

1.0
1.0
1.0
1.0
1.0
1.0
1.0
1.0
1.0
1.0
1.0
1.0
1.0
1.0
1.0
1.0
1.0
1.0
1.0
1.0
1.0
1.0
1.0
1.0
1.0
1.0
1.0
1.0
1.0
1.0


36.0
0.0
0.0
0.0
0.0
0.0
0.0
0.0
0.0
14.0
15.0
7.0
0.0
0.0
0.0
0.0
0.0
0.0
0.0
0.0
0.0
0.0
0.0
0.0
0.0
0.0
0.0
0.0
0.0
0.0


36.0
0.0
0.0
0.0
0.0
0.0
0.0
0.0
0.0
14.0
15.0
7.0
0.0
0.0
0.0
0.0
0.0
0.0
0.0
0.0
0.0
0.0
0.0
0.0
0.0
0.0
0.0
0.0
0.0
0.0

1.0
1.0
1.0
1.0
1.0
1.0
1.0
1.0
1.0
1.0
1.0
1.0
1.0
1.0
1.0
1.0
1.0
1.0
1.0
1.0
1.0
1.0
1.0
1.0
1.0
1.0
1.0
1.0
1.0
1.0


34.0
0.0
0.0
0.0
15.0
9.0
0.0
0.0
0.0
0.0
10.0
0.0
0.0
0.0
0.0
0.0
0.0
0.0
0.0
0.0
0.0
0.0
0.0
0.0
0.0
0.0
0.0
0.0
0.0
0.0

1.0
1.0
1.0
1.0
1.0
1.0
1.0
1.0
1.0
1.0
1.0
1.0
1.0
1.0
1.0
1.0
1.0
1.0
1.0
1.0
1.0
1.0
1.0
1.0
1.0
1.0
1.0
1.0
1.0
1.0


9.0
0.0
0.0
0.0
0.0
0.0
0.0
0.0
0.0
0.0
9.0
0.0
0.0
0.0
0.0
0.0
0.0
0.0
0.0
0.0
0.0
0.0
0.0
0.0
0.0
0.0
0.0
0.0
0.0
0.0


9.0
0.0
0.0
0.0
0.0
0.0
0.0
0.0
0.0
0.0
9.0
0.0
0.0
0.0
0.0
0.0
0.0
0.0
0.0
0.0
0.0
0.0
0.0
0.0
0.0
0.0
0.0
0.0
0.0
0.0

1.0
1.0
1.0
1.0
1.0
1.0
1.0
1.0
1.0
1.0
1.0
1.0
1.0
1.0
1.0
1.0
1.0
1.0
1.0
1.0
1.0
1.0
1.0
1.0
1.0
1.0
1.0
1.0
1.0
1.0


1155.0
0.0
0.0
0.0
0.0
0.0
0.0
0.0
0.0
0.0
0.0
0.0
0.0
0.0
0.0
0.0
0.0
0.0
0.0
0.0
0.0
0.0
0.0
0.0
0.0
0.0
0.0
0.0
0.0
1155.0

0.77
0.77
0.77
0.77
0.77
0.77
0.77
0.77
0.77
0.77
0.77
0.77
0.77
0.77
0.77
0.77
0.77
0.77
0.77
0.77
0.77
0.77
0.77
0.77
0.77
0.77
0.77
0.77
0.77
0.77


112023.0
538.0
0.0
0.0
0.0
0.0
25.0
0.0
0.0
0.0
0.0
0.0
0.0
0.0
0.0
0.0
0.0
105330.0
0.0
0.0
0.0
0.0
0.0
0.0
0.0
0.0
0.0
4693.0
0.0
1437.0


105791.0
461.0
0.0
0.0
0.0
0.0
0.0
0.0
0.0
0.0
0.0
0.0
0.0
0.0
0.0
0.0
0.0
105330.0
0.0
0.0
0.0
0.0
0.0
0.0
0.0
0.0
0.0
0.0
0.0
0.0


105791.0
461.0
0.0
0.0
0.0
0.0
0.0
0.0
0.0
0.0
0.0
0.0
0.0
0.0
0.0
0.0
0.0
105330.0
0.0
0.0
0.0
0.0
0.0
0.0
0.0
0.0
0.0
0.0
0.0
0.0

1.0
1.0
1.0
1.0
1.0
1.0
1.0
1.0
1.0
1.0
1.0
1.0
1.0
1.0
1.0
1.0
1.0
1.0
1.0
1.0
1.0
1.0
1.0
1.0
1.0
1.0
1.0
1.0
1.0
1.0


6232.0
77.0
0.0
0.0
0.0
0.0
25.0
0.0
0.0
0.0
0.0
0.0
0.0
0.0
0.0
0.0
0.0
0.0
0.0
0.0
0.0
0.0
0.0
0.0
0.0
0.0
0.0
4693.0
0.0
1437.0


6232.0
77.0
0.0
0.0
0.0
0.0
25.0
0.0
0.0
0.0
0.0
0.0
0.0
0.0
0.0
0.0
0.0
0.0
0.0
0.0
0.0
0.0
0.0
0.0
0.0
0.0
0.0
4693.0
0.0
1437.0

1.0
1.0
1.0
1.0
1.0
1.0
1.0
1.0
1.0
1.0
1.0
1.0
1.0
1.0
1.0
1.0
1.0
1.0
1.0
1.0
1.0
1.0
1.0
1.0
1.0
1.0
1.0
1.0
1.0
1.0


419.0
0.0
0.0
0.0
0.0
0.0
0.0
0.0
419.0
0.0
0.0
0.0
0.0
0.0
0.0
0.0
0.0
0.0
0.0
0.0
0.0
0.0
0.0
0.0
0.0
0.0
0.0
0.0
0.0
0.0


419.0
0.0
0.0
0.0
0.0
0.0
0.0
0.0
419.0
0.0
0.0
0.0
0.0
0.0
0.0
0.0
0.0
0.0
0.0
0.0
0.0
0.0
0.0
0.0
0.0
0.0
0.0
0.0
0.0
0.0

0.99
0.99
0.99
0.99
0.99
0.99
0.99
0.99
0.99
0.99
0.99
0.99
0.99
0.99
0.99
0.99
0.99
0.99
0.99
0.99
0.99
0.99
0.99
0.99
0.99
0.99
0.99
0.99
0.99
0.99


13742.0
265.0
246.0
101.0
27.0
239.0
78.0
113.0
354.0
501.0
835.0
450.0
0.0
0.0
136.0
197.0
0.0
1498.0
4720.0
0.0
1295.0
0.0
0.0
0.0
1121.0
688.0
6.0
0.0
511.0
361.0

0.96
0.96
0.96
0.96
0.96
0.96
0.96
0.96
0.96
0.96
0.96
0.96
0.96
0.96
0.96
0.96
0.96
0.96
0.96
0.96
0.96
0.96
0.96
0.96
0.96
0.96
0.96
0.96
0.96
0.96


8962.0
37.0
62.0
56.0
27.0
71.0
78.0
0.0
0.0
149.0
295.0
81.0
0.0
0.0
136.0
0.0
0.0
1498.0
4624.0
0.0
1295.0
0.0
0.0
0.0
0.0
0.0
0.0
0.0
511.0
42.0


638.0
24.0
42.0
56.0
27.0
71.0
78.0
0.0
0.0
119.0
41.0
44.0
0.0
0.0
136.0
0.0
0.0
0.0
0.0
0.0
0.0
0.0
0.0
0.0
0.0
0.0
0.0
0.0
0.0
0.0

1.0
1.0
1.0
1.0
1.0
1.0
1.0
1.0
1.0
1.0
1.0
1.0
1.0
1.0
1.0
1.0
1.0
1.0
1.0
1.0
1.0
1.0
1.0
1.0
1.0
1.0
1.0
1.0
1.0
1.0


1565.0
13.0
20.0
0.0
0.0
0.0
0.0
0.0
0.0
30.0
170.0
37.0
0.0
0.0
0.0
0.0
0.0
0.0
0.0
0.0
1295.0
0.0
0.0
0.0
0.0
0.0
0.0
0.0
0.0
0.0

1.0
1.0
1.0
1.0
1.0
1.0
1.0
1.0
1.0
1.0
1.0
1.0
1.0
1.0
1.0
1.0
1.0
1.0
1.0
1.0
1.0
1.0
1.0
1.0
1.0
1.0
1.0
1.0
1.0
1.0


6023.0
0.0
0.0
0.0
0.0
0.0
0.0
0.0
0.0
0.0
84.0
0.0
0.0
0.0
0.0
0.0
0.0
1498.0
4441.0
0.0
0.0
0.0
0.0
0.0
0.0
0.0
0.0
0.0
0.0
0.0

1.0
1.0
1.0
1.0
1.0
1.0
1.0
1.0
1.0
1.0
1.0
1.0
1.0
1.0
1.0
1.0
1.0
1.0
1.0
1.0
1.0
1.0
1.0
1.0
1.0
1.0
1.0
1.0
1.0
1.0


736.0
0.0
0.0
0.0
0.0
0.0
0.0
0.0
0.0
0.0
0.0
0.0
0.0
0.0
0.0
0.0
0.0
0.0
183.0
0.0
0.0
0.0
0.0
0.0
0.0
0.0
0.0
0.0
511.0
42.0

1.0
1.0
1.0
1.0
1.0
1.0
1.0
1.0
1.0
1.0
1.0
1.0
1.0
1.0
1.0
1.0
1.0
1.0
1.0
1.0
1.0
1.0
1.0
1.0
1.0
1.0
1.0
1.0
1.0
1.0


1032.0
55.0
98.0
0.0
0.0
27.0
0.0
0.0
0.0
127.0
245.0
161.0
0.0
0.0
0.0
0.0
0.0
0.0
0.0
0.0
0.0
0.0
0.0
0.0
0.0
0.0
0.0
0.0
0.0
319.0

1.0
1.0
1.0
1.0
1.0
1.0
1.0
1.0
1.0
1.0
1.0
1.0
1.0
1.0
1.0
1.0
1.0
1.0
1.0
1.0
1.0
1.0
1.0
1.0
1.0
1.0
1.0
1.0
1.0
1.0


489.0
21.0
65.0
0.0
0.0
0.0
0.0
0.0
0.0
50.0
195.0
158.0
0.0
0.0
0.0
0.0
0.0
0.0
0.0
0.0
0.0
0.0
0.0
0.0
0.0
0.0
0.0
0.0
0.0
0.0

1.0
1.0
1.0
1.0
1.0
1.0
1.0
1.0
1.0
1.0
1.0
1.0
1.0
1.0
1.0
1.0
1.0
1.0
1.0
1.0
1.0
1.0
1.0
1.0
1.0
1.0
1.0
1.0
1.0
1.0


84.0
29.0
28.0
0.0
0.0
27.0
0.0
0.0
0.0
0.0
0.0
0.0
0.0
0.0
0.0
0.0
0.0
0.0
0.0
0.0
0.0
0.0
0.0
0.0
0.0
0.0
0.0
0.0
0.0
0.0

1.0
1.0
1.0
1.0
1.0
1.0
1.0
1.0
1.0
1.0
1.0
1.0
1.0
1.0
1.0
1.0
1.0
1.0
1.0
1.0
1.0
1.0
1.0
1.0
1.0
1.0
1.0
1.0
1.0
1.0


15.0
1.0
0.0
0.0
0.0
0.0
0.0
0.0
0.0
0.0
14.0
0.0
0.0
0.0
0.0
0.0
0.0
0.0
0.0
0.0
0.0
0.0
0.0
0.0
0.0
0.0
0.0
0.0
0.0
0.0

1.0
1.0
1.0
1.0
1.0
1.0
1.0
1.0
1.0
1.0
1.0
1.0
1.0
1.0
1.0
1.0
1.0
1.0
1.0
1.0
1.0
1.0
1.0
1.0
1.0
1.0
1.0
1.0
1.0
1.0


319.0
0.0
0.0
0.0
0.0
0.0
0.0
0.0
0.0
0.0
0.0
0.0
0.0
0.0
0.0
0.0
0.0
0.0
0.0
0.0
0.0
0.0
0.0
0.0
0.0
0.0
0.0
0.0
0.0
319.0

0.88
0.88
0.88
0.88
0.88
0.88
0.88
0.88
0.88
0.88
0.88
0.88
0.88
0.88
0.88
0.88
0.88
0.88
0.88
0.88
0.88
0.88
0.88
0.88
0.88
0.88
0.88
0.88
0.88
0.88


287.0
51.0
60.0
0.0
0.0
123.0
0.0
0.0
0.0
0.0
0.0
53.0
0.0
0.0
0.0
0.0
0.0
0.0
0.0
0.0
0.0
0.0
0.0
0.0
0.0
0.0
0.0
0.0
0.0
0.0

1.0
1.0
1.0
1.0
1.0
1.0
1.0
1.0
1.0
1.0
1.0
1.0
1.0
1.0
1.0
1.0
1.0
1.0
1.0
1.0
1.0
1.0
1.0
1.0
1.0
1.0
1.0
1.0
1.0
1.0


169.0
8.0
38.0
0.0
0.0
123.0
0.0
0.0
0.0
0.0
0.0
0.0
0.0
0.0
0.0
0.0
0.0
0.0
0.0
0.0
0.0
0.0
0.0
0.0
0.0
0.0
0.0
0.0
0.0
0.0

1.0
1.0
1.0
1.0
1.0
1.0
1.0
1.0
1.0
1.0
1.0
1.0
1.0
1.0
1.0
1.0
1.0
1.0
1.0
1.0
1.0
1.0
1.0
1.0
1.0
1.0
1.0
1.0
1.0
1.0


1261.0
90.0
7.0
0.0
0.0
8.0
0.0
113.0
249.0
218.0
287.0
92.0
0.0
0.0
0.0
197.0
0.0
0.0
0.0
0.0
0.0
0.0
0.0
0.0
0.0
0.0
0.0
0.0
0.0
0.0

0.99
0.99
0.99
0.99
0.99
0.99
0.99
0.99
0.99
0.99
0.99
0.99
0.99
0.99
0.99
0.99
0.99
0.99
0.99
0.99
0.99
0.99
0.99
0.99
0.99
0.99
0.99
0.99
0.99
0.99


179.0
0.0
0.0
0.0
0.0
0.0
0.0
70.0
34.0
75.0
0.0
0.0
0.0
0.0
0.0
0.0
0.0
0.0
0.0
0.0
0.0
0.0
0.0
0.0
0.0
0.0
0.0
0.0
0.0
0.0

1.0
1.0
1.0
1.0
1.0
1.0
1.0
1.0
1.0
1.0
1.0
1.0
1.0
1.0
1.0
1.0
1.0
1.0
1.0
1.0
1.0
1.0
1.0
1.0
1.0
1.0
1.0
1.0
1.0
1.0


237.0
0.0
0.0
0.0
0.0
0.0
0.0
0.0
187.0
37.0
0.0
13.0
0.0
0.0
0.0
0.0
0.0
0.0
0.0
0.0
0.0
0.0
0.0
0.0
0.0
0.0
0.0
0.0
0.0
0.0

0.92
0.92
0.92
0.92
0.92
0.92
0.92
0.92
0.92
0.92
0.92
0.92
0.92
0.92
0.92
0.92
0.92
0.92
0.92
0.92
0.92
0.92
0.92
0.92
0.92
0.92
0.92
0.92
0.92
0.92


89.0
0.0
3.0
0.0
0.0
0.0
0.0
0.0
0.0
20.0
66.0
0.0
0.0
0.0
0.0
0.0
0.0
0.0
0.0
0.0
0.0
0.0
0.0
0.0
0.0
0.0
0.0
0.0
0.0
0.0

1.0
1.0
1.0
1.0
1.0
1.0
1.0
1.0
1.0
1.0
1.0
1.0
1.0
1.0
1.0
1.0
1.0
1.0
1.0
1.0
1.0
1.0
1.0
1.0
1.0
1.0
1.0
1.0
1.0
1.0


47.0
0.0
0.0
0.0
0.0
0.0
0.0
0.0
0.0
12.0
0.0
35.0
0.0
0.0
0.0
0.0
0.0
0.0
0.0
0.0
0.0
0.0
0.0
0.0
0.0
0.0
0.0
0.0
0.0
0.0

1.0
1.0
1.0
1.0
1.0
1.0
1.0
1.0
1.0
1.0
1.0
1.0
1.0
1.0
1.0
1.0
1.0
1.0
1.0
1.0
1.0
1.0
1.0
1.0
1.0
1.0
1.0
1.0
1.0
1.0


29.0
20.0
4.0
0.0
0.0
0.0
0.0
0.0
0.0
0.0
0.0
5.0
0.0
0.0
0.0
0.0
0.0
0.0
0.0
0.0
0.0
0.0
0.0
0.0
0.0
0.0
0.0
0.0
0.0
0.0

1.0
1.0
1.0
1.0
1.0
1.0
1.0
1.0
1.0
1.0
1.0
1.0
1.0
1.0
1.0
1.0
1.0
1.0
1.0
1.0
1.0
1.0
1.0
1.0
1.0
1.0
1.0
1.0
1.0
1.0


8.0
0.0
0.0
0.0
0.0
8.0
0.0
0.0
0.0
0.0
0.0
0.0
0.0
0.0
0.0
0.0
0.0
0.0
0.0
0.0
0.0
0.0
0.0
0.0
0.0
0.0
0.0
0.0
0.0
0.0

1.0
1.0
1.0
1.0
1.0
1.0
1.0
1.0
1.0
1.0
1.0
1.0
1.0
1.0
1.0
1.0
1.0
1.0
1.0
1.0
1.0
1.0
1.0
1.0
1.0
1.0
1.0
1.0
1.0
1.0


80.0
0.0
0.0
0.0
0.0
0.0
0.0
0.0
80.0
0.0
0.0
0.0
0.0
0.0
0.0
0.0
0.0
0.0
0.0
0.0
0.0
0.0
0.0
0.0
0.0
0.0
0.0
0.0
0.0
0.0


80.0
0.0
0.0
0.0
0.0
0.0
0.0
0.0
80.0
0.0
0.0
0.0
0.0
0.0
0.0
0.0
0.0
0.0
0.0
0.0
0.0
0.0
0.0
0.0
0.0
0.0
0.0
0.0
0.0
0.0

1.0
1.0
1.0
1.0
1.0
1.0
1.0
1.0
1.0
1.0
1.0
1.0
1.0
1.0
1.0
1.0
1.0
1.0
1.0
1.0
1.0
1.0
1.0
1.0
1.0
1.0
1.0
1.0
1.0
1.0


77.0
10.0
6.0
0.0
0.0
0.0
0.0
0.0
0.0
0.0
0.0
61.0
0.0
0.0
0.0
0.0
0.0
0.0
0.0
0.0
0.0
0.0
0.0
0.0
0.0
0.0
0.0
0.0
0.0
0.0


77.0
10.0
6.0
0.0
0.0
0.0
0.0
0.0
0.0
0.0
0.0
61.0
0.0
0.0
0.0
0.0
0.0
0.0
0.0
0.0
0.0
0.0
0.0
0.0
0.0
0.0
0.0
0.0
0.0
0.0

1.0
1.0
1.0
1.0
1.0
1.0
1.0
1.0
1.0
1.0
1.0
1.0
1.0
1.0
1.0
1.0
1.0
1.0
1.0
1.0
1.0
1.0
1.0
1.0
1.0
1.0
1.0
1.0
1.0
1.0


119.0
20.0
0.0
0.0
0.0
0.0
0.0
0.0
25.0
7.0
8.0
2.0
0.0
0.0
0.0
0.0
0.0
0.0
57.0
0.0
0.0
0.0
0.0
0.0
0.0
0.0
0.0
0.0
0.0
0.0


43.0
11.0
0.0
0.0
0.0
0.0
0.0
0.0
25.0
7.0
0.0
0.0
0.0
0.0
0.0
0.0
0.0
0.0
0.0
0.0
0.0
0.0
0.0
0.0
0.0
0.0
0.0
0.0
0.0
0.0

1.0
1.0
1.0
1.0
1.0
1.0
1.0
1.0
1.0
1.0
1.0
1.0
1.0
1.0
1.0
1.0
1.0
1.0
1.0
1.0
1.0
1.0
1.0
1.0
1.0
1.0
1.0
1.0
1.0
1.0


76.0
9.0
0.0
0.0
0.0
0.0
0.0
0.0
0.0
0.0
8.0
2.0
0.0
0.0
0.0
0.0
0.0
0.0
57.0
0.0
0.0
0.0
0.0
0.0
0.0
0.0
0.0
0.0
0.0
0.0

1.0
1.0
1.0
1.0
1.0
1.0
1.0
1.0
1.0
1.0
1.0
1.0
1.0
1.0
1.0
1.0
1.0
1.0
1.0
1.0
1.0
1.0
1.0
1.0
1.0
1.0
1.0
1.0
1.0
1.0


94.0
2.0
13.0
45.0
0.0
10.0
0.0
0.0
0.0
0.0
0.0
0.0
0.0
0.0
0.0
0.0
0.0
0.0
18.0
0.0
0.0
0.0
0.0
0.0
0.0
0.0
6.0
0.0
0.0
0.0

0.91
0.91
0.91
0.91
0.91
0.91
0.91
0.91
0.91
0.91
0.91
0.91
0.91
0.91
0.91
0.91
0.91
0.91
0.91
0.91
0.91
0.91
0.91
0.91
0.91
0.91
0.91
0.91
0.91
0.91


31.0
2.0
13.0
16.0
0.0
0.0
0.0
0.0
0.0
0.0
0.0
0.0
0.0
0.0
0.0
0.0
0.0
0.0
0.0
0.0
0.0
0.0
0.0
0.0
0.0
0.0
0.0
0.0
0.0
0.0

1.0
1.0
1.0
1.0
1.0
1.0
1.0
1.0
1.0
1.0
1.0
1.0
1.0
1.0
1.0
1.0
1.0
1.0
1.0
1.0
1.0
1.0
1.0
1.0
1.0
1.0
1.0
1.0
1.0
1.0


203847.0
405.0
282.0
0.0
0.0
0.0
0.0
0.0
0.0
0.0
0.0
0.0
0.0
0.0
0.0
0.0
0.0
0.0
0.0
0.0
0.0
0.0
0.0
0.0
0.0
0.0
0.0
32639.0
0.0
170521.0


203847.0
405.0
282.0
0.0
0.0
0.0
0.0
0.0
0.0
0.0
0.0
0.0
0.0
0.0
0.0
0.0
0.0
0.0
0.0
0.0
0.0
0.0
0.0
0.0
0.0
0.0
0.0
32639.0
0.0
170521.0

1.0
1.0
1.0
1.0
1.0
1.0
1.0
1.0
1.0
1.0
1.0
1.0
1.0
1.0
1.0
1.0
1.0
1.0
1.0
1.0
1.0
1.0
1.0
1.0
1.0
1.0
1.0
1.0
1.0
1.0


623.0
405.0
218.0
0.0
0.0
0.0
0.0
0.0
0.0
0.0
0.0
0.0
0.0
0.0
0.0
0.0
0.0
0.0
0.0
0.0
0.0
0.0
0.0
0.0
0.0
0.0
0.0
0.0
0.0
0.0

1.0
1.0
1.0
1.0
1.0
1.0
1.0
1.0
1.0
1.0
1.0
1.0
1.0
1.0
1.0
1.0
1.0
1.0
1.0
1.0
1.0
1.0
1.0
1.0
1.0
1.0
1.0
1.0
1.0
1.0


54.0
0.0
54.0
0.0
0.0
0.0
0.0
0.0
0.0
0.0
0.0
0.0
0.0
0.0
0.0
0.0
0.0
0.0
0.0
0.0
0.0
0.0
0.0
0.0
0.0
0.0
0.0
0.0
0.0
0.0

1.0
1.0
1.0
1.0
1.0
1.0
1.0
1.0
1.0
1.0
1.0
1.0
1.0
1.0
1.0
1.0
1.0
1.0
1.0
1.0
1.0
1.0
1.0
1.0
1.0
1.0
1.0
1.0
1.0
1.0


5061.0
0.0
0.0
0.0
0.0
0.0
33.0
95.0
763.0
777.0
61.0
0.0
0.0
0.0
0.0
0.0
0.0
0.0
0.0
0.0
0.0
0.0
0.0
0.0
0.0
0.0
0.0
0.0
3332.0
0.0


1311.0
0.0
0.0
0.0
0.0
0.0
0.0
0.0
473.0
777.0
61.0
0.0
0.0
0.0
0.0
0.0
0.0
0.0
0.0
0.0
0.0
0.0
0.0
0.0
0.0
0.0
0.0
0.0
0.0
0.0


1311.0
0.0
0.0
0.0
0.0
0.0
0.0
0.0
473.0
777.0
61.0
0.0
0.0
0.0
0.0
0.0
0.0
0.0
0.0
0.0
0.0
0.0
0.0
0.0
0.0
0.0
0.0
0.0
0.0
0.0

1.0
1.0
1.0
1.0
1.0
1.0
1.0
1.0
1.0
1.0
1.0
1.0
1.0
1.0
1.0
1.0
1.0
1.0
1.0
1.0
1.0
1.0
1.0
1.0
1.0
1.0
1.0
1.0
1.0
1.0


290.0
0.0
0.0
0.0
0.0
0.0
0.0
0.0
290.0
0.0
0.0
0.0
0.0
0.0
0.0
0.0
0.0
0.0
0.0
0.0
0.0
0.0
0.0
0.0
0.0
0.0
0.0
0.0
0.0
0.0


290.0
0.0
0.0
0.0
0.0
0.0
0.0
0.0
290.0
0.0
0.0
0.0
0.0
0.0
0.0
0.0
0.0
0.0
0.0
0.0
0.0
0.0
0.0
0.0
0.0
0.0
0.0
0.0
0.0
0.0

1.0
1.0
1.0
1.0
1.0
1.0
1.0
1.0
1.0
1.0
1.0
1.0
1.0
1.0
1.0
1.0
1.0
1.0
1.0
1.0
1.0
1.0
1.0
1.0
1.0
1.0
1.0
1.0
1.0
1.0


3427.0
0.0
0.0
0.0
0.0
0.0
0.0
95.0
0.0
0.0
0.0
0.0
0.0
0.0
0.0
0.0
0.0
0.0
0.0
0.0
0.0
0.0
0.0
0.0
0.0
0.0
0.0
0.0
3332.0
0.0


3427.0
0.0
0.0
0.0
0.0
0.0
0.0
95.0
0.0
0.0
0.0
0.0
0.0
0.0
0.0
0.0
0.0
0.0
0.0
0.0
0.0
0.0
0.0
0.0
0.0
0.0
0.0
0.0
3332.0
0.0

1.0
1.0
1.0
1.0
1.0
1.0
1.0
1.0
1.0
1.0
1.0
1.0
1.0
1.0
1.0
1.0
1.0
1.0
1.0
1.0
1.0
1.0
1.0
1.0
1.0
1.0
1.0
1.0
1.0
1.0


33.0
0.0
0.0
0.0
0.0
0.0
33.0
0.0
0.0
0.0
0.0
0.0
0.0
0.0
0.0
0.0
0.0
0.0
0.0
0.0
0.0
0.0
0.0
0.0
0.0
0.0
0.0
0.0
0.0
0.0


33.0
0.0
0.0
0.0
0.0
0.0
33.0
0.0
0.0
0.0
0.0
0.0
0.0
0.0
0.0
0.0
0.0
0.0
0.0
0.0
0.0
0.0
0.0
0.0
0.0
0.0
0.0
0.0
0.0
0.0

1.0
1.0
1.0
1.0
1.0
1.0
1.0
1.0
1.0
1.0
1.0
1.0
1.0
1.0
1.0
1.0
1.0
1.0
1.0
1.0
1.0
1.0
1.0
1.0
1.0
1.0
1.0
1.0
1.0
1.0


287.0
3.0
0.0
0.0
0.0
0.0
0.0
0.0
148.0
75.0
61.0
0.0
0.0
0.0
0.0
0.0
0.0
0.0
0.0
0.0
0.0
0.0
0.0
0.0
0.0
0.0
0.0
0.0
0.0
0.0


287.0
3.0
0.0
0.0
0.0
0.0
0.0
0.0
148.0
75.0
61.0
0.0
0.0
0.0
0.0
0.0
0.0
0.0
0.0
0.0
0.0
0.0
0.0
0.0
0.0
0.0
0.0
0.0
0.0
0.0


279.0
3.0
0.0
0.0
0.0
0.0
0.0
0.0
148.0
67.0
61.0
0.0
0.0
0.0
0.0
0.0
0.0
0.0
0.0
0.0
0.0
0.0
0.0
0.0
0.0
0.0
0.0
0.0
0.0
0.0

1.0
1.0
1.0
1.0
1.0
1.0
1.0
1.0
1.0
1.0
1.0
1.0
1.0
1.0
1.0
1.0
1.0
1.0
1.0
1.0
1.0
1.0
1.0
1.0
1.0
1.0
1.0
1.0
1.0
1.0


8.0
0.0
0.0
0.0
0.0
0.0
0.0
0.0
0.0
8.0
0.0
0.0
0.0
0.0
0.0
0.0
0.0
0.0
0.0
0.0
0.0
0.0
0.0
0.0
0.0
0.0
0.0
0.0
0.0
0.0

0.99
0.99
0.99
0.99
0.99
0.99
0.99
0.99
0.99
0.99
0.99
0.99
0.99
0.99
0.99
0.99
0.99
0.99
0.99
0.99
0.99
0.99
0.99
0.99
0.99
0.99
0.99
0.99
0.99
0.99


472.0
0.0
0.0
0.0
0.0
0.0
0.0
0.0
192.0
125.0
155.0
0.0
0.0
0.0
0.0
0.0
0.0
0.0
0.0
0.0
0.0
0.0
0.0
0.0
0.0
0.0
0.0
0.0
0.0
0.0


472.0
0.0
0.0
0.0
0.0
0.0
0.0
0.0
192.0
125.0
155.0
0.0
0.0
0.0
0.0
0.0
0.0
0.0
0.0
0.0
0.0
0.0
0.0
0.0
0.0
0.0
0.0
0.0
0.0
0.0

1.0
1.0
1.0
1.0
1.0
1.0
1.0
1.0
1.0
1.0
1.0
1.0
1.0
1.0
1.0
1.0
1.0
1.0
1.0
1.0
1.0
1.0
1.0
1.0
1.0
1.0
1.0
1.0
1.0
1.0


192.0
0.0
0.0
0.0
0.0
0.0
0.0
0.0
192.0
0.0
0.0
0.0
0.0
0.0
0.0
0.0
0.0
0.0
0.0
0.0
0.0
0.0
0.0
0.0
0.0
0.0
0.0
0.0
0.0
0.0

0.88
0.88
0.88
0.88
0.88
0.88
0.88
0.88
0.88
0.88
0.88
0.88
0.88
0.88
0.88
0.88
0.88
0.88
0.88
0.88
0.88
0.88
0.88
0.88
0.88
0.88
0.88
0.88
0.88
0.88


318.0
39.0
0.0
0.0
0.0
0.0
0.0
0.0
0.0
101.0
121.0
57.0
0.0
0.0
0.0
0.0
0.0
0.0
0.0
0.0
0.0
0.0
0.0
0.0
0.0
0.0
0.0
0.0
0.0
0.0


318.0
39.0
0.0
0.0
0.0
0.0
0.0
0.0
0.0
101.0
121.0
57.0
0.0
0.0
0.0
0.0
0.0
0.0
0.0
0.0
0.0
0.0
0.0
0.0
0.0
0.0
0.0
0.0
0.0
0.0


318.0
39.0
0.0
0.0
0.0
0.0
0.0
0.0
0.0
101.0
121.0
57.0
0.0
0.0
0.0
0.0
0.0
0.0
0.0
0.0
0.0
0.0
0.0
0.0
0.0
0.0
0.0
0.0
0.0
0.0

1.0
1.0
1.0
1.0
1.0
1.0
1.0
1.0
1.0
1.0
1.0
1.0
1.0
1.0
1.0
1.0
1.0
1.0
1.0
1.0
1.0
1.0
1.0
1.0
1.0
1.0
1.0
1.0
1.0
1.0


851.0
0.0
0.0
0.0
0.0
0.0
0.0
0.0
0.0
93.0
0.0
0.0
0.0
0.0
0.0
0.0
0.0
0.0
0.0
0.0
0.0
0.0
0.0
0.0
0.0
758.0
0.0
0.0
0.0
0.0


851.0
0.0
0.0
0.0
0.0
0.0
0.0
0.0
0.0
93.0
0.0
0.0
0.0
0.0
0.0
0.0
0.0
0.0
0.0
0.0
0.0
0.0
0.0
0.0
0.0
758.0
0.0
0.0
0.0
0.0


851.0
0.0
0.0
0.0
0.0
0.0
0.0
0.0
0.0
93.0
0.0
0.0
0.0
0.0
0.0
0.0
0.0
0.0
0.0
0.0
0.0
0.0
0.0
0.0
0.0
758.0
0.0
0.0
0.0
0.0

1.0
1.0
1.0
1.0
1.0
1.0
1.0
1.0
1.0
1.0
1.0
1.0
1.0
1.0
1.0
1.0
1.0
1.0
1.0
1.0
1.0
1.0
1.0
1.0
1.0
1.0
1.0
1.0
1.0
1.0


6214.0
0.0
0.0
0.0
0.0
0.0
0.0
0.0
0.0
0.0
52.0
0.0
0.0
0.0
0.0
0.0
0.0
5707.0
0.0
0.0
0.0
0.0
0.0
0.0
455.0
0.0
0.0
0.0
0.0
0.0

0.81
0.81
0.81
0.81
0.81
0.81
0.81
0.81
0.81
0.81
0.81
0.81
0.81
0.81
0.81
0.81
0.81
0.81
0.81
0.81
0.81
0.81
0.81
0.81
0.81
0.81
0.81
0.81
0.81
0.81


5759.0
0.0
0.0
0.0
0.0
0.0
0.0
0.0
0.0
0.0
52.0
0.0
0.0
0.0
0.0
0.0
0.0
5707.0
0.0
0.0
0.0
0.0
0.0
0.0
0.0
0.0
0.0
0.0
0.0
0.0

1.0
1.0
1.0
1.0
1.0
1.0
1.0
1.0
1.0
1.0
1.0
1.0
1.0
1.0
1.0
1.0
1.0
1.0
1.0
1.0
1.0
1.0
1.0
1.0
1.0
1.0
1.0
1.0
1.0
1.0


5613.0
0.0
0.0
0.0
0.0
0.0
0.0
0.0
0.0
0.0
0.0
0.0
0.0
0.0
0.0
0.0
0.0
5613.0
0.0
0.0
0.0
0.0
0.0
0.0
0.0
0.0
0.0
0.0
0.0
0.0

1.0
1.0
1.0
1.0
1.0
1.0
1.0
1.0
1.0
1.0
1.0
1.0
1.0
1.0
1.0
1.0
1.0
1.0
1.0
1.0
1.0
1.0
1.0
1.0
1.0
1.0
1.0
1.0
1.0
1.0


29908.0
0.0
0.0
0.0
0.0
0.0
0.0
0.0
0.0
14.0
0.0
37.0
0.0
0.0
0.0
0.0
0.0
29857.0
0.0
0.0
0.0
0.0
0.0
0.0
0.0
0.0
0.0
0.0
0.0
0.0


29908.0
0.0
0.0
0.0
0.0
0.0
0.0
0.0
0.0
14.0
0.0
37.0
0.0
0.0
0.0
0.0
0.0
29857.0
0.0
0.0
0.0
0.0
0.0
0.0
0.0
0.0
0.0
0.0
0.0
0.0

1.0
1.0
1.0
1.0
1.0
1.0
1.0
1.0
1.0
1.0
1.0
1.0
1.0
1.0
1.0
1.0
1.0
1.0
1.0
1.0
1.0
1.0
1.0
1.0
1.0
1.0
1.0
1.0
1.0
1.0


29786.0
0.0
0.0
0.0
0.0
0.0
0.0
0.0
0.0
0.0
0.0
0.0
0.0
0.0
0.0
0.0
0.0
29786.0
0.0
0.0
0.0
0.0
0.0
0.0
0.0
0.0
0.0
0.0
0.0
0.0

1.0
1.0
1.0
1.0
1.0
1.0
1.0
1.0
1.0
1.0
1.0
1.0
1.0
1.0
1.0
1.0
1.0
1.0
1.0
1.0
1.0
1.0
1.0
1.0
1.0
1.0
1.0
1.0
1.0
1.0


8837.0
46.0
28.0
0.0
0.0
0.0
0.0
0.0
0.0
0.0
0.0
0.0
26.0
0.0
0.0
0.0
0.0
156.0
408.0
0.0
0.0
0.0
8147.0
26.0
0.0
0.0
0.0
0.0
0.0
0.0


8837.0
46.0
28.0
0.0
0.0
0.0
0.0
0.0
0.0
0.0
0.0
0.0
26.0
0.0
0.0
0.0
0.0
156.0
408.0
0.0
0.0
0.0
8147.0
26.0
0.0
0.0
0.0
0.0
0.0
0.0

0.73
0.73
0.73
0.73
0.73
0.73
0.73
0.73
0.73
0.73
0.73
0.73
0.73
0.73
0.73
0.73
0.73
0.73
0.73
0.73
0.73
0.73
0.73
0.73
0.73
0.73
0.73
0.73
0.73
0.73


46.0
46.0
0.0
0.0
0.0
0.0
0.0
0.0
0.0
0.0
0.0
0.0
0.0
0.0
0.0
0.0
0.0
0.0
0.0
0.0
0.0
0.0
0.0
0.0
0.0
0.0
0.0
0.0
0.0
0.0

1.0
1.0
1.0
1.0
1.0
1.0
1.0
1.0
1.0
1.0
1.0
1.0
1.0
1.0
1.0
1.0
1.0
1.0
1.0
1.0
1.0
1.0
1.0
1.0
1.0
1.0
1.0
1.0
1.0
1.0


28.0
0.0
28.0
0.0
0.0
0.0
0.0
0.0
0.0
0.0
0.0
0.0
0.0
0.0
0.0
0.0
0.0
0.0
0.0
0.0
0.0
0.0
0.0
0.0
0.0
0.0
0.0
0.0
0.0
0.0

1.0
1.0
1.0
1.0
1.0
1.0
1.0
1.0
1.0
1.0
1.0
1.0
1.0
1.0
1.0
1.0
1.0
1.0
1.0
1.0
1.0
1.0
1.0
1.0
1.0
1.0
1.0
1.0
1.0
1.0


23.0
0.0
0.0
0.0
0.0
0.0
0.0
0.0
0.0
23.0
0.0
0.0
0.0
0.0
0.0
0.0
0.0
0.0
0.0
0.0
0.0
0.0
0.0
0.0
0.0
0.0
0.0
0.0
0.0
0.0


23.0
0.0
0.0
0.0
0.0
0.0
0.0
0.0
0.0
23.0
0.0
0.0
0.0
0.0
0.0
0.0
0.0
0.0
0.0
0.0
0.0
0.0
0.0
0.0
0.0
0.0
0.0
0.0
0.0
0.0

0.99
0.99
0.99
0.99
0.99
0.99
0.99
0.99
0.99
0.99
0.99
0.99
0.99
0.99
0.99
0.99
0.99
0.99
0.99
0.99
0.99
0.99
0.99
0.99
0.99
0.99
0.99
0.99
0.99
0.99


19.0
0.0
0.0
0.0
0.0
0.0
0.0
0.0
0.0
0.0
19.0
0.0
0.0
0.0
0.0
0.0
0.0
0.0
0.0
0.0
0.0
0.0
0.0
0.0
0.0
0.0
0.0
0.0
0.0
0.0


19.0
0.0
0.0
0.0
0.0
0.0
0.0
0.0
0.0
0.0
19.0
0.0
0.0
0.0
0.0
0.0
0.0
0.0
0.0
0.0
0.0
0.0
0.0
0.0
0.0
0.0
0.0
0.0
0.0
0.0


19.0
0.0
0.0
0.0
0.0
0.0
0.0
0.0
0.0
0.0
19.0
0.0
0.0
0.0
0.0
0.0
0.0
0.0
0.0
0.0
0.0
0.0
0.0
0.0
0.0
0.0
0.0
0.0
0.0
0.0

1.0
1.0
1.0
1.0
1.0
1.0
1.0
1.0
1.0
1.0
1.0
1.0
1.0
1.0
1.0
1.0
1.0
1.0
1.0
1.0
1.0
1.0
1.0
1.0
1.0
1.0
1.0
1.0
1.0
1.0


27.0
0.0
0.0
8.0
10.0
0.0
9.0
0.0
0.0
0.0
0.0
0.0
0.0
0.0
0.0
0.0
0.0
0.0
0.0
0.0
0.0
0.0
0.0
0.0
0.0
0.0
0.0
0.0
0.0
0.0

0.73
0.73
0.73
0.73
0.73
0.73
0.73
0.73
0.73
0.73
0.73
0.73
0.73
0.73
0.73
0.73
0.73
0.73
0.73
0.73
0.73
0.73
0.73
0.73
0.73
0.73
0.73
0.73
0.73
0.73


36516.0
0.0
0.0
0.0
0.0
0.0
0.0
15.0
0.0
0.0
0.0
0.0
0.0
0.0
179.0
0.0
0.0
31426.0
3501.0
368.0
106.0
0.0
0.0
0.0
0.0
286.0
603.0
32.0
0.0
0.0


36516.0
0.0
0.0
0.0
0.0
0.0
0.0
15.0
0.0
0.0
0.0
0.0
0.0
0.0
179.0
0.0
0.0
31426.0
3501.0
368.0
106.0
0.0
0.0
0.0
0.0
286.0
603.0
32.0
0.0
0.0

0.97
0.97
0.97
0.97
0.97
0.97
0.97
0.97
0.97
0.97
0.97
0.97
0.97
0.97
0.97
0.97
0.97
0.97
0.97
0.97
0.97
0.97
0.97
0.97
0.97
0.97
0.97
0.97
0.97
0.97


35432.0
0.0
0.0
0.0
0.0
0.0
0.0
0.0
0.0
0.0
0.0
0.0
0.0
0.0
31.0
0.0
0.0
31426.0
3501.0
368.0
106.0
0.0
0.0
0.0
0.0
0.0
0.0
0.0
0.0
0.0

1.0
1.0
1.0
1.0
1.0
1.0
1.0
1.0
1.0
1.0
1.0
1.0
1.0
1.0
1.0
1.0
1.0
1.0
1.0
1.0
1.0
1.0
1.0
1.0
1.0
1.0
1.0
1.0
1.0
1.0


921.0
0.0
0.0
0.0
0.0
0.0
0.0
0.0
0.0
0.0
0.0
0.0
0.0
0.0
0.0
0.0
0.0
0.0
0.0
0.0
0.0
0.0
0.0
0.0
0.0
286.0
603.0
32.0
0.0
0.0

1.0
1.0
1.0
1.0
1.0
1.0
1.0
1.0
1.0
1.0
1.0
1.0
1.0
1.0
1.0
1.0
1.0
1.0
1.0
1.0
1.0
1.0
1.0
1.0
1.0
1.0
1.0
1.0
1.0
1.0


13.0
0.0
0.0
0.0
0.0
0.0
13.0
0.0
0.0
0.0
0.0
0.0
0.0
0.0
0.0
0.0
0.0
0.0
0.0
0.0
0.0
0.0
0.0
0.0
0.0
0.0
0.0
0.0
0.0
0.0


13.0
0.0
0.0
0.0
0.0
0.0
13.0
0.0
0.0
0.0
0.0
0.0
0.0
0.0
0.0
0.0
0.0
0.0
0.0
0.0
0.0
0.0
0.0
0.0
0.0
0.0
0.0
0.0
0.0
0.0


13.0
0.0
0.0
0.0
0.0
0.0
13.0
0.0
0.0
0.0
0.0
0.0
0.0
0.0
0.0
0.0
0.0
0.0
0.0
0.0
0.0
0.0
0.0
0.0
0.0
0.0
0.0
0.0
0.0
0.0

1.0
1.0
1.0
1.0
1.0
1.0
1.0
1.0
1.0
1.0
1.0
1.0
1.0
1.0
1.0
1.0
1.0
1.0
1.0
1.0
1.0
1.0
1.0
1.0
1.0
1.0
1.0
1.0
1.0
1.0


77294.0
0.0
0.0
0.0
0.0
0.0
0.0
0.0
0.0
0.0
0.0
0.0
0.0
0.0
77294.0
0.0
0.0
0.0
0.0
0.0
0.0
0.0
0.0
0.0
0.0
0.0
0.0
0.0
0.0
0.0

1.0
1.0
1.0
1.0
1.0
1.0
1.0
1.0
1.0
1.0
1.0
1.0
1.0
1.0
1.0
1.0
1.0
1.0
1.0
1.0
1.0
1.0
1.0
1.0
1.0
1.0
1.0
1.0
1.0
1.0


70745.0
0.0
0.0
0.0
0.0
0.0
0.0
0.0
0.0
0.0
0.0
0.0
0.0
0.0
70745.0
0.0
0.0
0.0
0.0
0.0
0.0
0.0
0.0
0.0
0.0
0.0
0.0
0.0
0.0
0.0

0.88
0.88
0.88
0.88
0.88
0.88
0.88
0.88
0.88
0.88
0.88
0.88
0.88
0.88
0.88
0.88
0.88
0.88
0.88
0.88
0.88
0.88
0.88
0.88
0.88
0.88
0.88
0.88
0.88
0.88


4.0
0.0
0.0
0.0
0.0
0.0
0.0
0.0
0.0
0.0
0.0
0.0
0.0
0.0
4.0
0.0
0.0
0.0
0.0
0.0
0.0
0.0
0.0
0.0
0.0
0.0
0.0
0.0
0.0
0.0

0.84
0.84
0.84
0.84
0.84
0.84
0.84
0.84
0.84
0.84
0.84
0.84
0.84
0.84
0.84
0.84
0.84
0.84
0.84
0.84
0.84
0.84
0.84
0.84
0.84
0.84
0.84
0.84
0.84
0.84


83958.0
0.0
0.0
0.0
0.0
0.0
0.0
0.0
0.0
0.0
0.0
0.0
0.0
0.0
0.0
1884.0
0.0
0.0
0.0
0.0
0.0
0.0
0.0
0.0
0.0
0.0
80897.0
0.0
1177.0
0.0


83958.0
0.0
0.0
0.0
0.0
0.0
0.0
0.0
0.0
0.0
0.0
0.0
0.0
0.0
0.0
1884.0
0.0
0.0
0.0
0.0
0.0
0.0
0.0
0.0
0.0
0.0
80897.0
0.0
1177.0
0.0


80897.0
0.0
0.0
0.0
0.0
0.0
0.0
0.0
0.0
0.0
0.0
0.0
0.0
0.0
0.0
0.0
0.0
0.0
0.0
0.0
0.0
0.0
0.0
0.0
0.0
0.0
80897.0
0.0
0.0
0.0

1.0
1.0
1.0
1.0
1.0
1.0
1.0
1.0
1.0
1.0
1.0
1.0
1.0
1.0
1.0
1.0
1.0
1.0
1.0
1.0
1.0
1.0
1.0
1.0
1.0
1.0
1.0
1.0
1.0
1.0


3061.0
0.0
0.0
0.0
0.0
0.0
0.0
0.0
0.0
0.0
0.0
0.0
0.0
0.0
0.0
1884.0
0.0
0.0
0.0
0.0
0.0
0.0
0.0
0.0
0.0
0.0
0.0
0.0
1177.0
0.0

1.0
1.0
1.0
1.0
1.0
1.0
1.0
1.0
1.0
1.0
1.0
1.0
1.0
1.0
1.0
1.0
1.0
1.0
1.0
1.0
1.0
1.0
1.0
1.0
1.0
1.0
1.0
1.0
1.0
1.0


25849.0
0.0
0.0
0.0
0.0
0.0
0.0
0.0
0.0
0.0
0.0
0.0
0.0
0.0
9430.0
0.0
0.0
4716.0
11703.0
0.0
0.0
0.0
0.0
0.0
0.0
0.0
0.0
0.0
0.0
0.0


16419.0
0.0
0.0
0.0
0.0
0.0
0.0
0.0
0.0
0.0
0.0
0.0
0.0
0.0
0.0
0.0
0.0
4716.0
11703.0
0.0
0.0
0.0
0.0
0.0
0.0
0.0
0.0
0.0
0.0
0.0

1.0
1.0
1.0
1.0
1.0
1.0
1.0
1.0
1.0
1.0
1.0
1.0
1.0
1.0
1.0
1.0
1.0
1.0
1.0
1.0
1.0
1.0
1.0
1.0
1.0
1.0
1.0
1.0
1.0
1.0


9430.0
0.0
0.0
0.0
0.0
0.0
0.0
0.0
0.0
0.0
0.0
0.0
0.0
0.0
9430.0
0.0
0.0
0.0
0.0
0.0
0.0
0.0
0.0
0.0
0.0
0.0
0.0
0.0
0.0
0.0


9430.0
0.0
0.0
0.0
0.0
0.0
0.0
0.0
0.0
0.0
0.0
0.0
0.0
0.0
9430.0
0.0
0.0
0.0
0.0
0.0
0.0
0.0
0.0
0.0
0.0
0.0
0.0
0.0
0.0
0.0

1.0
1.0
1.0
1.0
1.0
1.0
1.0
1.0
1.0
1.0
1.0
1.0
1.0
1.0
1.0
1.0
1.0
1.0
1.0
1.0
1.0
1.0
1.0
1.0
1.0
1.0
1.0
1.0
1.0
1.0


8820.0
0.0
0.0
0.0
0.0
0.0
0.0
0.0
0.0
0.0
0.0
0.0
0.0
0.0
0.0
0.0
0.0
8820.0
0.0
0.0
0.0
0.0
0.0
0.0
0.0
0.0
0.0
0.0
0.0
0.0


8820.0
0.0
0.0
0.0
0.0
0.0
0.0
0.0
0.0
0.0
0.0
0.0
0.0
0.0
0.0
0.0
0.0
8820.0
0.0
0.0
0.0
0.0
0.0
0.0
0.0
0.0
0.0
0.0
0.0
0.0


8820.0
0.0
0.0
0.0
0.0
0.0
0.0
0.0
0.0
0.0
0.0
0.0
0.0
0.0
0.0
0.0
0.0
8820.0
0.0
0.0
0.0
0.0
0.0
0.0
0.0
0.0
0.0
0.0
0.0
0.0

1.0
1.0
1.0
1.0
1.0
1.0
1.0
1.0
1.0
1.0
1.0
1.0
1.0
1.0
1.0
1.0
1.0
1.0
1.0
1.0
1.0
1.0
1.0
1.0
1.0
1.0
1.0
1.0
1.0
1.0


5894.0
0.0
0.0
0.0
0.0
0.0
0.0
0.0
0.0
0.0
0.0
0.0
0.0
2617.0
3277.0
0.0
0.0
0.0
0.0
0.0
0.0
0.0
0.0
0.0
0.0
0.0
0.0
0.0
0.0
0.0

0.71
0.71
0.71
0.71
0.71
0.71
0.71
0.71
0.71
0.71
0.71
0.71
0.71
0.71
0.71
0.71
0.71
0.71
0.71
0.71
0.71
0.71
0.71
0.71
0.71
0.71
0.71
0.71
0.71
0.71


5133.0
0.0
0.0
0.0
0.0
0.0
0.0
0.0
0.0
0.0
0.0
0.0
0.0
2263.0
2870.0
0.0
0.0
0.0
0.0
0.0
0.0
0.0
0.0
0.0
0.0
0.0
0.0
0.0
0.0
0.0

0.99
0.99
0.99
0.99
0.99
0.99
0.99
0.99
0.99
0.99
0.99
0.99
0.99
0.99
0.99
0.99
0.99
0.99
0.99
0.99
0.99
0.99
0.99
0.99
0.99
0.99
0.99
0.99
0.99
0.99


684.0
0.0
0.0
0.0
0.0
0.0
0.0
0.0
0.0
0.0
0.0
0.0
0.0
0.0
0.0
0.0
0.0
0.0
0.0
0.0
0.0
0.0
0.0
0.0
0.0
0.0
0.0
0.0
684.0
0.0


684.0
0.0
0.0
0.0
0.0
0.0
0.0
0.0
0.0
0.0
0.0
0.0
0.0
0.0
0.0
0.0
0.0
0.0
0.0
0.0
0.0
0.0
0.0
0.0
0.0
0.0
0.0
0.0
684.0
0.0


684.0
0.0
0.0
0.0
0.0
0.0
0.0
0.0
0.0
0.0
0.0
0.0
0.0
0.0
0.0
0.0
0.0
0.0
0.0
0.0
0.0
0.0
0.0
0.0
0.0
0.0
0.0
0.0
684.0
0.0

1.0
1.0
1.0
1.0
1.0
1.0
1.0
1.0
1.0
1.0
1.0
1.0
1.0
1.0
1.0
1.0
1.0
1.0
1.0
1.0
1.0
1.0
1.0
1.0
1.0
1.0
1.0
1.0
1.0
1.0


1802316.0
19218.0
9719.0
9622.0
20225.0
14106.0
24831.0
11243.0
4462.0
6406.0
13973.0
81774.0
47079.0
27093.0
136259.0
43356.0
232900.0
17538.0
31390.0
103124.0
75608.0
83849.0
100525.0
47079.0
248350.0
144029.0
108362.0
37165.0
13286.0
89745.0

1.0
1.0
1.0
1.0
1.0
1.0
1.0
1.0
1.0
1.0
1.0
1.0
1.0
1.0
1.0
1.0
1.0
1.0
1.0
1.0
1.0
1.0
1.0
1.0
1.0
1.0
1.0
1.0
1.0
1.0


639919.0
15295.0
6039.0
3783.0
6235.0
7895.0
20880.0
10591.0
2258.0
2214.0
7410.0
69972.0
7363.0
948.0
37955.0
548.0
687.0
15070.0
20168.0
48170.0
1825.0
27.0
6590.0
7363.0
94451.0
118592.0
94394.0
8043.0
5291.0
19862.0

1.0
1.0
1.0
1.0
1.0
1.0
1.0
1.0
1.0
1.0
1.0
1.0
1.0
1.0
1.0
1.0
1.0
1.0
1.0
1.0
1.0
1.0
1.0
1.0
1.0
1.0
1.0
1.0
1.0
1.0


86418.0
13919.0
2475.0
0.0
1226.0
0.0
0.0
111.0
806.0
91.0
0.0
67790.0
0.0
0.0
0.0
0.0
0.0
0.0
0.0
0.0
0.0
0.0
0.0
0.0
0.0
0.0
0.0
0.0
0.0
0.0


86418.0
13919.0
2475.0
0.0
1226.0
0.0
0.0
111.0
806.0
91.0
0.0
67790.0
0.0
0.0
0.0
0.0
0.0
0.0
0.0
0.0
0.0
0.0
0.0
0.0
0.0
0.0
0.0
0.0
0.0
0.0

1.0
1.0
1.0
1.0
1.0
1.0
1.0
1.0
1.0
1.0
1.0
1.0
1.0
1.0
1.0
1.0
1.0
1.0
1.0
1.0
1.0
1.0
1.0
1.0
1.0
1.0
1.0
1.0
1.0
1.0


16662.0
0.0
0.0
0.0
0.0
0.0
0.0
10341.0
1366.0
811.0
1779.0
336.0
2.0
0.0
0.0
0.0
0.0
0.0
0.0
173.0
0.0
27.0
1825.0
2.0
0.0
0.0
0.0
0.0
0.0
0.0


16662.0
0.0
0.0
0.0
0.0
0.0
0.0
10341.0
1366.0
811.0
1779.0
336.0
2.0
0.0
0.0
0.0
0.0
0.0
0.0
173.0
0.0
27.0
1825.0
2.0
0.0
0.0
0.0
0.0
0.0
0.0

1.0
1.0
1.0
1.0
1.0
1.0
1.0
1.0
1.0
1.0
1.0
1.0
1.0
1.0
1.0
1.0
1.0
1.0
1.0
1.0
1.0
1.0
1.0
1.0
1.0
1.0
1.0
1.0
1.0
1.0


186640.0
0.0
48.0
0.0
2052.0
3901.0
2284.0
0.0
0.0
0.0
0.0
0.0
0.0
0.0
0.0
0.0
0.0
0.0
0.0
0.0
0.0
0.0
0.0
0.0
57871.0
75274.0
45210.0
0.0
0.0
0.0

1.0
1.0
1.0
1.0
1.0
1.0
1.0
1.0
1.0
1.0
1.0
1.0
1.0
1.0
1.0
1.0
1.0
1.0
1.0
1.0
1.0
1.0
1.0
1.0
1.0
1.0
1.0
1.0
1.0
1.0


63753.0
0.0
48.0
0.0
2052.0
3891.0
88.0
0.0
0.0
0.0
0.0
0.0
0.0
0.0
0.0
0.0
0.0
0.0
0.0
0.0
0.0
0.0
0.0
0.0
57674.0
0.0
0.0
0.0
0.0
0.0

0.99
0.99
0.99
0.99
0.99
0.99
0.99
0.99
0.99
0.99
0.99
0.99
0.99
0.99
0.99
0.99
0.99
0.99
0.99
0.99
0.99
0.99
0.99
0.99
0.99
0.99
0.99
0.99
0.99
0.99


122680.0
0.0
0.0
0.0
0.0
0.0
2196.0
0.0
0.0
0.0
0.0
0.0
0.0
0.0
0.0
0.0
0.0
0.0
0.0
0.0
0.0
0.0
0.0
0.0
0.0
75274.0
45210.0
0.0
0.0
0.0

1.0
1.0
1.0
1.0
1.0
1.0
1.0
1.0
1.0
1.0
1.0
1.0
1.0
1.0
1.0
1.0
1.0
1.0
1.0
1.0
1.0
1.0
1.0
1.0
1.0
1.0
1.0
1.0
1.0
1.0


10.0
0.0
0.0
0.0
0.0
10.0
0.0
0.0
0.0
0.0
0.0
0.0
0.0
0.0
0.0
0.0
0.0
0.0
0.0
0.0
0.0
0.0
0.0
0.0
0.0
0.0
0.0
0.0
0.0
0.0

0.77
0.77
0.77
0.77
0.77
0.77
0.77
0.77
0.77
0.77
0.77
0.77
0.77
0.77
0.77
0.77
0.77
0.77
0.77
0.77
0.77
0.77
0.77
0.77
0.77
0.77
0.77
0.77
0.77
0.77


47259.0
0.0
568.0
0.0
275.0
575.0
14388.0
0.0
0.0
0.0
0.0
0.0
0.0
0.0
6968.0
0.0
0.0
2222.0
2933.0
0.0
620.0
0.0
0.0
0.0
0.0
6326.0
10802.0
0.0
1582.0
0.0

1.0
1.0
1.0
1.0
1.0
1.0
1.0
1.0
1.0
1.0
1.0
1.0
1.0
1.0
1.0
1.0
1.0
1.0
1.0
1.0
1.0
1.0
1.0
1.0
1.0
1.0
1.0
1.0
1.0
1.0


24277.0
0.0
335.0
0.0
0.0
34.0
10992.0
0.0
0.0
0.0
0.0
0.0
0.0
0.0
6661.0
0.0
0.0
0.0
0.0
0.0
0.0
0.0
0.0
0.0
0.0
6255.0
0.0
0.0
0.0
0.0

1.0
1.0
1.0
1.0
1.0
1.0
1.0
1.0
1.0
1.0
1.0
1.0
1.0
1.0
1.0
1.0
1.0
1.0
1.0
1.0
1.0
1.0
1.0
1.0
1.0
1.0
1.0
1.0
1.0
1.0


617.0
0.0
0.0
0.0
0.0
363.0
254.0
0.0
0.0
0.0
0.0
0.0
0.0
0.0
0.0
0.0
0.0
0.0
0.0
0.0
0.0
0.0
0.0
0.0
0.0
0.0
0.0
0.0
0.0
0.0

1.0
1.0
1.0
1.0
1.0
1.0
1.0
1.0
1.0
1.0
1.0
1.0
1.0
1.0
1.0
1.0
1.0
1.0
1.0
1.0
1.0
1.0
1.0
1.0
1.0
1.0
1.0
1.0
1.0
1.0


81.0
0.0
0.0
0.0
0.0
0.0
10.0
0.0
0.0
0.0
0.0
0.0
0.0
0.0
0.0
0.0
0.0
0.0
0.0
0.0
0.0
0.0
0.0
0.0
0.0
71.0
0.0
0.0
0.0
0.0

1.0
1.0
1.0
1.0
1.0
1.0
1.0
1.0
1.0
1.0
1.0
1.0
1.0
1.0
1.0
1.0
1.0
1.0
1.0
1.0
1.0
1.0
1.0
1.0
1.0
1.0
1.0
1.0
1.0
1.0


8054.0
528.0
755.0
1457.0
563.0
825.0
249.0
0.0
0.0
420.0
1501.0
657.0
0.0
0.0
0.0
0.0
0.0
0.0
0.0
0.0
0.0
0.0
0.0
0.0
1099.0
0.0
0.0
0.0
0.0
0.0


8054.0
528.0
755.0
1457.0
563.0
825.0
249.0
0.0
0.0
420.0
1501.0
657.0
0.0
0.0
0.0
0.0
0.0
0.0
0.0
0.0
0.0
0.0
0.0
0.0
1099.0
0.0
0.0
0.0
0.0
0.0

1.0
1.0
1.0
1.0
1.0
1.0
1.0
1.0
1.0
1.0
1.0
1.0
1.0
1.0
1.0
1.0
1.0
1.0
1.0
1.0
1.0
1.0
1.0
1.0
1.0
1.0
1.0
1.0
1.0
1.0


7857.0
48.0
209.0
427.0
280.0
645.0
925.0
0.0
0.0
84.0
2677.0
443.0
0.0
154.0
717.0
0.0
0.0
0.0
0.0
0.0
0.0
0.0
0.0
0.0
187.0
869.0
0.0
0.0
192.0
0.0


4993.0
48.0
209.0
427.0
280.0
645.0
925.0
0.0
0.0
84.0
148.0
108.0
0.0
154.0
717.0
0.0
0.0
0.0
0.0
0.0
0.0
0.0
0.0
0.0
187.0
869.0
0.0
0.0
192.0
0.0

0.96
0.96
0.96
0.96
0.96
0.96
0.96
0.96
0.96
0.96
0.96
0.96
0.96
0.96
0.96
0.96
0.96
0.96
0.96
0.96
0.96
0.96
0.96
0.96
0.96
0.96
0.96
0.96
0.96
0.96


2864.0
0.0
0.0
0.0
0.0
0.0
0.0
0.0
0.0
0.0
2529.0
335.0
0.0
0.0
0.0
0.0
0.0
0.0
0.0
0.0
0.0
0.0
0.0
0.0
0.0
0.0
0.0
0.0
0.0
0.0

1.0
1.0
1.0
1.0
1.0
1.0
1.0
1.0
1.0
1.0
1.0
1.0
1.0
1.0
1.0
1.0
1.0
1.0
1.0
1.0
1.0
1.0
1.0
1.0
1.0
1.0
1.0
1.0
1.0
1.0


2171.0
354.0
101.0
16.0
10.0
26.0
46.0
0.0
35.0
319.0
944.0
320.0
0.0
0.0
0.0
0.0
0.0
0.0
0.0
0.0
0.0
0.0
0.0
0.0
0.0
0.0
0.0
0.0
0.0
0.0

0.89
0.89
0.89
0.89
0.89
0.89
0.89
0.89
0.89
0.89
0.89
0.89
0.89
0.89
0.89
0.89
0.89
0.89
0.89
0.89
0.89
0.89
0.89
0.89
0.89
0.89
0.89
0.89
0.89
0.89


20.0
9.0
11.0
0.0
0.0
0.0
0.0
0.0
0.0
0.0
0.0
0.0
0.0
0.0
0.0
0.0
0.0
0.0
0.0
0.0
0.0
0.0
0.0
0.0
0.0
0.0
0.0
0.0
0.0
0.0

1.0
1.0
1.0
1.0
1.0
1.0
1.0
1.0
1.0
1.0
1.0
1.0
1.0
1.0
1.0
1.0
1.0
1.0
1.0
1.0
1.0
1.0
1.0
1.0
1.0
1.0
1.0
1.0
1.0
1.0


9017.0
0.0
0.0
0.0
0.0
0.0
786.0
0.0
0.0
0.0
0.0
0.0
0.0
0.0
0.0
0.0
0.0
0.0
4697.0
1552.0
0.0
0.0
0.0
0.0
0.0
0.0
1982.0
0.0
0.0
0.0


9017.0
0.0
0.0
0.0
0.0
0.0
786.0
0.0
0.0
0.0
0.0
0.0
0.0
0.0
0.0
0.0
0.0
0.0
4697.0
1552.0
0.0
0.0
0.0
0.0
0.0
0.0
1982.0
0.0
0.0
0.0

1.0
1.0
1.0
1.0
1.0
1.0
1.0
1.0
1.0
1.0
1.0
1.0
1.0
1.0
1.0
1.0
1.0
1.0
1.0
1.0
1.0
1.0
1.0
1.0
1.0
1.0
1.0
1.0
1.0
1.0


2986.0
0.0
0.0
0.0
467.0
0.0
80.0
0.0
0.0
0.0
0.0
0.0
0.0
0.0
0.0
0.0
0.0
0.0
0.0
0.0
0.0
0.0
0.0
0.0
2082.0
0.0
0.0
0.0
357.0
0.0


2986.0
0.0
0.0
0.0
467.0
0.0
80.0
0.0
0.0
0.0
0.0
0.0
0.0
0.0
0.0
0.0
0.0
0.0
0.0
0.0
0.0
0.0
0.0
0.0
2082.0
0.0
0.0
0.0
357.0
0.0

0.98
0.98
0.98
0.98
0.98
0.98
0.98
0.98
0.98
0.98
0.98
0.98
0.98
0.98
0.98
0.98
0.98
0.98
0.98
0.98
0.98
0.98
0.98
0.98
0.98
0.98
0.98
0.98
0.98
0.98


463.0
25.0
25.0
82.0
85.0
132.0
93.0
0.0
0.0
0.0
18.0
3.0
0.0
0.0
0.0
0.0
0.0
0.0
0.0
0.0
0.0
0.0
0.0
0.0
0.0
0.0
0.0
0.0
0.0
0.0


463.0
25.0
25.0
82.0
85.0
132.0
93.0
0.0
0.0
0.0
18.0
3.0
0.0
0.0
0.0
0.0
0.0
0.0
0.0
0.0
0.0
0.0
0.0
0.0
0.0
0.0
0.0
0.0
0.0
0.0

1.0
1.0
1.0
1.0
1.0
1.0
1.0
1.0
1.0
1.0
1.0
1.0
1.0
1.0
1.0
1.0
1.0
1.0
1.0
1.0
1.0
1.0
1.0
1.0
1.0
1.0
1.0
1.0
1.0
1.0


75911.0
119.0
414.0
0.0
4.0
0.0
0.0
0.0
0.0
0.0
27.0
4.0
7361.0
296.0
27721.0
0.0
0.0
3089.0
5425.0
18062.0
107.0
0.0
4765.0
7361.0
1156.0
0.0
0.0
0.0
0.0
0.0

1.0
1.0
1.0
1.0
1.0
1.0
1.0
1.0
1.0
1.0
1.0
1.0
1.0
1.0
1.0
1.0
1.0
1.0
1.0
1.0
1.0
1.0
1.0
1.0
1.0
1.0
1.0
1.0
1.0
1.0


38.0
11.0
0.0
0.0
0.0
0.0
0.0
0.0
0.0
0.0
27.0
0.0
0.0
0.0
0.0
0.0
0.0
0.0
0.0
0.0
0.0
0.0
0.0
0.0
0.0
0.0
0.0
0.0
0.0
0.0

0.81
0.81
0.81
0.81
0.81
0.81
0.81
0.81
0.81
0.81
0.81
0.81
0.81
0.81
0.81
0.81
0.81
0.81
0.81
0.81
0.81
0.81
0.81
0.81
0.81
0.81
0.81
0.81
0.81
0.81


20.0
20.0
0.0
0.0
0.0
0.0
0.0
0.0
0.0
0.0
0.0
0.0
0.0
0.0
0.0
0.0
0.0
0.0
0.0
0.0
0.0
0.0
0.0
0.0
0.0
0.0
0.0
0.0
0.0
0.0

1.0
1.0
1.0
1.0
1.0
1.0
1.0
1.0
1.0
1.0
1.0
1.0
1.0
1.0
1.0
1.0
1.0
1.0
1.0
1.0
1.0
1.0
1.0
1.0
1.0
1.0
1.0
1.0
1.0
1.0


2043.0
9.0
0.0
0.0
0.0
0.0
0.0
0.0
0.0
0.0
0.0
0.0
83.0
30.0
1256.0
0.0
0.0
17.0
103.0
258.0
13.0
0.0
184.0
83.0
7.0
0.0
0.0
0.0
0.0
0.0

1.0
1.0
1.0
1.0
1.0
1.0
1.0
1.0
1.0
1.0
1.0
1.0
1.0
1.0
1.0
1.0
1.0
1.0
1.0
1.0
1.0
1.0
1.0
1.0
1.0
1.0
1.0
1.0
1.0
1.0


25649.0
7.0
40.0
141.0
122.0
11.0
154.0
0.0
0.0
0.0
0.0
0.0
0.0
0.0
0.0
0.0
0.0
0.0
0.0
0.0
0.0
0.0
0.0
0.0
23430.0
0.0
0.0
1744.0
0.0
0.0


25649.0
7.0
40.0
141.0
122.0
11.0
154.0
0.0
0.0
0.0
0.0
0.0
0.0
0.0
0.0
0.0
0.0
0.0
0.0
0.0
0.0
0.0
0.0
0.0
23430.0
0.0
0.0
1744.0
0.0
0.0

1.0
1.0
1.0
1.0
1.0
1.0
1.0
1.0
1.0
1.0
1.0
1.0
1.0
1.0
1.0
1.0
1.0
1.0
1.0
1.0
1.0
1.0
1.0
1.0
1.0
1.0
1.0
1.0
1.0
1.0


510.0
0.0
86.0
4.0
3.0
325.0
39.0
0.0
0.0
3.0
0.0
0.0
0.0
0.0
33.0
0.0
0.0
0.0
0.0
0.0
0.0
0.0
0.0
0.0
8.0
9.0
0.0
0.0
0.0
0.0

0.87
0.87
0.87
0.87
0.87
0.87
0.87
0.87
0.87
0.87
0.87
0.87
0.87
0.87
0.87
0.87
0.87
0.87
0.87
0.87
0.87
0.87
0.87
0.87
0.87
0.87
0.87
0.87
0.87
0.87


280.0
0.0
67.0
0.0
0.0
213.0
0.0
0.0
0.0
0.0
0.0
0.0
0.0
0.0
0.0
0.0
0.0
0.0
0.0
0.0
0.0
0.0
0.0
0.0
0.0
0.0
0.0
0.0
0.0
0.0

1.0
1.0
1.0
1.0
1.0
1.0
1.0
1.0
1.0
1.0
1.0
1.0
1.0
1.0
1.0
1.0
1.0
1.0
1.0
1.0
1.0
1.0
1.0
1.0
1.0
1.0
1.0
1.0
1.0
1.0


81.0
0.0
11.0
0.0
0.0
70.0
0.0
0.0
0.0
0.0
0.0
0.0
0.0
0.0
0.0
0.0
0.0
0.0
0.0
0.0
0.0
0.0
0.0
0.0
0.0
0.0
0.0
0.0
0.0
0.0

0.83
0.83
0.83
0.83
0.83
0.83
0.83
0.83
0.83
0.83
0.83
0.83
0.83
0.83
0.83
0.83
0.83
0.83
0.83
0.83
0.83
0.83
0.83
0.83
0.83
0.83
0.83
0.83
0.83
0.83


819.0
56.0
68.0
24.0
2.0
15.0
2.0
0.0
39.0
11.0
16.0
87.0
0.0
0.0
0.0
0.0
0.0
0.0
0.0
0.0
0.0
0.0
0.0
0.0
499.0
0.0
0.0
0.0
0.0
0.0


736.0
52.0
44.0
24.0
2.0
15.0
2.0
0.0
0.0
11.0
0.0
87.0
0.0
0.0
0.0
0.0
0.0
0.0
0.0
0.0
0.0
0.0
0.0
0.0
499.0
0.0
0.0
0.0
0.0
0.0

1.0
1.0
1.0
1.0
1.0
1.0
1.0
1.0
1.0
1.0
1.0
1.0
1.0
1.0
1.0
1.0
1.0
1.0
1.0
1.0
1.0
1.0
1.0
1.0
1.0
1.0
1.0
1.0
1.0
1.0


83.0
4.0
24.0
0.0
0.0
0.0
0.0
0.0
39.0
0.0
16.0
0.0
0.0
0.0
0.0
0.0
0.0
0.0
0.0
0.0
0.0
0.0
0.0
0.0
0.0
0.0
0.0
0.0
0.0
0.0

1.0
1.0
1.0
1.0
1.0
1.0
1.0
1.0
1.0
1.0
1.0
1.0
1.0
1.0
1.0
1.0
1.0
1.0
1.0
1.0
1.0
1.0
1.0
1.0
1.0
1.0
1.0
1.0
1.0
1.0


344.0
63.0
0.0
0.0
0.0
0.0
0.0
0.0
0.0
229.0
52.0
0.0
0.0
0.0
0.0
0.0
0.0
0.0
0.0
0.0
0.0
0.0
0.0
0.0
0.0
0.0
0.0
0.0
0.0
0.0

1.0
1.0
1.0
1.0
1.0
1.0
1.0
1.0
1.0
1.0
1.0
1.0
1.0
1.0
1.0
1.0
1.0
1.0
1.0
1.0
1.0
1.0
1.0
1.0
1.0
1.0
1.0
1.0
1.0
1.0


193.0
53.0
0.0
0.0
0.0
0.0
0.0
0.0
0.0
88.0
52.0
0.0
0.0
0.0
0.0
0.0
0.0
0.0
0.0
0.0
0.0
0.0
0.0
0.0
0.0
0.0
0.0
0.0
0.0
0.0

1.0
1.0
1.0
1.0
1.0
1.0
1.0
1.0
1.0
1.0
1.0
1.0
1.0
1.0
1.0
1.0
1.0
1.0
1.0
1.0
1.0
1.0
1.0
1.0
1.0
1.0
1.0
1.0
1.0
1.0


6161.0
0.0
777.0
0.0
0.0
0.0
0.0
0.0
0.0
0.0
0.0
0.0
0.0
0.0
0.0
0.0
0.0
0.0
0.0
0.0
0.0
0.0
0.0
0.0
5384.0
0.0
0.0
0.0
0.0
0.0

1.0
1.0
1.0
1.0
1.0
1.0
1.0
1.0
1.0
1.0
1.0
1.0
1.0
1.0
1.0
1.0
1.0
1.0
1.0
1.0
1.0
1.0
1.0
1.0
1.0
1.0
1.0
1.0
1.0
1.0


109.0
0.0
15.0
0.0
0.0
0.0
0.0
94.0
0.0
0.0
0.0
0.0
0.0
0.0
0.0
0.0
0.0
0.0
0.0
0.0
0.0
0.0
0.0
0.0
0.0
0.0
0.0
0.0
0.0
0.0


109.0
0.0
15.0
0.0
0.0
0.0
0.0
94.0
0.0
0.0
0.0
0.0
0.0
0.0
0.0
0.0
0.0
0.0
0.0
0.0
0.0
0.0
0.0
0.0
0.0
0.0
0.0
0.0
0.0
0.0

1.0
1.0
1.0
1.0
1.0
1.0
1.0
1.0
1.0
1.0
1.0
1.0
1.0
1.0
1.0
1.0
1.0
1.0
1.0
1.0
1.0
1.0
1.0
1.0
1.0
1.0
1.0
1.0
1.0
1.0


106.0
0.0
0.0
0.0
1.0
0.0
0.0
0.0
0.0
0.0
0.0
105.0
0.0
0.0
0.0
0.0
0.0
0.0
0.0
0.0
0.0
0.0
0.0
0.0
0.0
0.0
0.0
0.0
0.0
0.0


106.0
0.0
0.0
0.0
1.0
0.0
0.0
0.0
0.0
0.0
0.0
105.0
0.0
0.0
0.0
0.0
0.0
0.0
0.0
0.0
0.0
0.0
0.0
0.0
0.0
0.0
0.0
0.0
0.0
0.0

1.0
1.0
1.0
1.0
1.0
1.0
1.0
1.0
1.0
1.0
1.0
1.0
1.0
1.0
1.0
1.0
1.0
1.0
1.0
1.0
1.0
1.0
1.0
1.0
1.0
1.0
1.0
1.0
1.0
1.0


435.0
4.0
0.0
0.0
0.0
0.0
0.0
0.0
0.0
98.0
43.0
1.0
0.0
0.0
24.0
0.0
0.0
0.0
209.0
56.0
0.0
0.0
0.0
0.0
0.0
0.0
0.0
0.0
0.0
0.0


435.0
4.0
0.0
0.0
0.0
0.0
0.0
0.0
0.0
98.0
43.0
1.0
0.0
0.0
24.0
0.0
0.0
0.0
209.0
56.0
0.0
0.0
0.0
0.0
0.0
0.0
0.0
0.0
0.0
0.0

1.0
1.0
1.0
1.0
1.0
1.0
1.0
1.0
1.0
1.0
1.0
1.0
1.0
1.0
1.0
1.0
1.0
1.0
1.0
1.0
1.0
1.0
1.0
1.0
1.0
1.0
1.0
1.0
1.0
1.0


759.0
7.0
6.0
54.0
26.0
70.0
0.0
0.0
0.0
0.0
0.0
35.0
0.0
0.0
0.0
0.0
0.0
0.0
0.0
0.0
0.0
0.0
0.0
0.0
0.0
0.0
0.0
0.0
0.0
561.0

1.0
1.0
1.0
1.0
1.0
1.0
1.0
1.0
1.0
1.0
1.0
1.0
1.0
1.0
1.0
1.0
1.0
1.0
1.0
1.0
1.0
1.0
1.0
1.0
1.0
1.0
1.0
1.0
1.0
1.0


131.0
7.0
6.0
0.0
26.0
70.0
0.0
0.0
0.0
0.0
0.0
22.0
0.0
0.0
0.0
0.0
0.0
0.0
0.0
0.0
0.0
0.0
0.0
0.0
0.0
0.0
0.0
0.0
0.0
0.0

1.0
1.0
1.0
1.0
1.0
1.0
1.0
1.0
1.0
1.0
1.0
1.0
1.0
1.0
1.0
1.0
1.0
1.0
1.0
1.0
1.0
1.0
1.0
1.0
1.0
1.0
1.0
1.0
1.0
1.0


67.0
0.0
0.0
54.0
0.0
0.0
0.0
0.0
0.0
0.0
0.0
13.0
0.0
0.0
0.0
0.0
0.0
0.0
0.0
0.0
0.0
0.0
0.0
0.0
0.0
0.0
0.0
0.0
0.0
0.0

0.9
0.9
0.9
0.9
0.9
0.9
0.9
0.9
0.9
0.9
0.9
0.9
0.9
0.9
0.9
0.9
0.9
0.9
0.9
0.9
0.9
0.9
0.9
0.9
0.9
0.9
0.9
0.9
0.9
0.9


106.0
106.0
0.0
0.0
0.0
0.0
0.0
0.0
0.0
0.0
0.0
0.0
0.0
0.0
0.0
0.0
0.0
0.0
0.0
0.0
0.0
0.0
0.0
0.0
0.0
0.0
0.0
0.0
0.0
0.0

1.0
1.0
1.0
1.0
1.0
1.0
1.0
1.0
1.0
1.0
1.0
1.0
1.0
1.0
1.0
1.0
1.0
1.0
1.0
1.0
1.0
1.0
1.0
1.0
1.0
1.0
1.0
1.0
1.0
1.0


107685.0
0.0
1.0
0.0
0.0
0.0
16.0
0.0
0.0
0.0
22.0
15.0
0.0
0.0
0.0
0.0
0.0
9629.0
2305.0
25422.0
0.0
0.0
0.0
0.0
0.0
24283.0
36400.0
5261.0
2009.0
2322.0

1.0
1.0
1.0
1.0
1.0
1.0
1.0
1.0
1.0
1.0
1.0
1.0
1.0
1.0
1.0
1.0
1.0
1.0
1.0
1.0
1.0
1.0
1.0
1.0
1.0
1.0
1.0
1.0
1.0
1.0


576.0
0.0
0.0
0.0
0.0
0.0
0.0
0.0
0.0
0.0
0.0
0.0
0.0
0.0
0.0
0.0
0.0
8.0
10.0
84.0
0.0
0.0
0.0
0.0
0.0
214.0
176.0
76.0
8.0
0.0

1.0
1.0
1.0
1.0
1.0
1.0
1.0
1.0
1.0
1.0
1.0
1.0
1.0
1.0
1.0
1.0
1.0
1.0
1.0
1.0
1.0
1.0
1.0
1.0
1.0
1.0
1.0
1.0
1.0
1.0


701.0
0.0
0.0
0.0
0.0
0.0
0.0
0.0
0.0
0.0
0.0
0.0
0.0
0.0
0.0
0.0
0.0
48.0
54.0
67.0
0.0
0.0
0.0
0.0
0.0
89.0
181.0
88.0
38.0
136.0

1.0
1.0
1.0
1.0
1.0
1.0
1.0
1.0
1.0
1.0
1.0
1.0
1.0
1.0
1.0
1.0
1.0
1.0
1.0
1.0
1.0
1.0
1.0
1.0
1.0
1.0
1.0
1.0
1.0
1.0


1016.0
0.0
0.0
0.0
0.0
0.0
45.0
0.0
0.0
0.0
0.0
0.0
0.0
0.0
0.0
0.0
0.0
0.0
971.0
0.0
0.0
0.0
0.0
0.0
0.0
0.0
0.0
0.0
0.0
0.0

1.0
1.0
1.0
1.0
1.0
1.0
1.0
1.0
1.0
1.0
1.0
1.0
1.0
1.0
1.0
1.0
1.0
1.0
1.0
1.0
1.0
1.0
1.0
1.0
1.0
1.0
1.0
1.0
1.0
1.0


1001.0
0.0
0.0
0.0
0.0
0.0
45.0
0.0
0.0
0.0
0.0
0.0
0.0
0.0
0.0
0.0
0.0
0.0
956.0
0.0
0.0
0.0
0.0
0.0
0.0
0.0
0.0
0.0
0.0
0.0

1.0
1.0
1.0
1.0
1.0
1.0
1.0
1.0
1.0
1.0
1.0
1.0
1.0
1.0
1.0
1.0
1.0
1.0
1.0
1.0
1.0
1.0
1.0
1.0
1.0
1.0
1.0
1.0
1.0
1.0


16963.0
0.0
44.0
0.0
0.0
0.0
0.0
0.0
0.0
0.0
0.0
0.0
0.0
0.0
0.0
0.0
0.0
0.0
0.0
0.0
0.0
0.0
0.0
0.0
0.0
0.0
0.0
0.0
0.0
16919.0


16963.0
0.0
44.0
0.0
0.0
0.0
0.0
0.0
0.0
0.0
0.0
0.0
0.0
0.0
0.0
0.0
0.0
0.0
0.0
0.0
0.0
0.0
0.0
0.0
0.0
0.0
0.0
0.0
0.0
16919.0

1.0
1.0
1.0
1.0
1.0
1.0
1.0
1.0
1.0
1.0
1.0
1.0
1.0
1.0
1.0
1.0
1.0
1.0
1.0
1.0
1.0
1.0
1.0
1.0
1.0
1.0
1.0
1.0
1.0
1.0


5199.0
0.0
0.0
0.0
0.0
41.0
0.0
0.0
0.0
0.0
0.0
0.0
0.0
0.0
0.0
0.0
0.0
0.0
3074.0
0.0
0.0
0.0
0.0
0.0
2084.0
0.0
0.0
0.0
0.0
0.0


5199.0
0.0
0.0
0.0
0.0
41.0
0.0
0.0
0.0
0.0
0.0
0.0
0.0
0.0
0.0
0.0
0.0
0.0
3074.0
0.0
0.0
0.0
0.0
0.0
2084.0
0.0
0.0
0.0
0.0
0.0

1.0
1.0
1.0
1.0
1.0
1.0
1.0
1.0
1.0
1.0
1.0
1.0
1.0
1.0
1.0
1.0
1.0
1.0
1.0
1.0
1.0
1.0
1.0
1.0
1.0
1.0
1.0
1.0
1.0
1.0


52.0
0.0
22.0
30.0
0.0
0.0
0.0
0.0
0.0
0.0
0.0
0.0
0.0
0.0
0.0
0.0
0.0
0.0
0.0
0.0
0.0
0.0
0.0
0.0
0.0
0.0
0.0
0.0
0.0
0.0

0.81
0.81
0.81
0.81
0.81
0.81
0.81
0.81
0.81
0.81
0.81
0.81
0.81
0.81
0.81
0.81
0.81
0.81
0.81
0.81
0.81
0.81
0.81
0.81
0.81
0.81
0.81
0.81
0.81
0.81


4302.0
0.0
3.0
14.0
0.0
1.0
6.0
0.0
0.0
6.0
0.0
0.0
0.0
0.0
0.0
0.0
0.0
130.0
0.0
2887.0
1098.0
0.0
0.0
0.0
157.0
0.0
0.0
0.0
0.0
0.0

1.0
1.0
1.0
1.0
1.0
1.0
1.0
1.0
1.0
1.0
1.0
1.0
1.0
1.0
1.0
1.0
1.0
1.0
1.0
1.0
1.0
1.0
1.0
1.0
1.0
1.0
1.0
1.0
1.0
1.0


30.0
0.0
3.0
14.0
0.0
1.0
6.0
0.0
0.0
6.0
0.0
0.0
0.0
0.0
0.0
0.0
0.0
0.0
0.0
0.0
0.0
0.0
0.0
0.0
0.0
0.0
0.0
0.0
0.0
0.0

1.0
1.0
1.0
1.0
1.0
1.0
1.0
1.0
1.0
1.0
1.0
1.0
1.0
1.0
1.0
1.0
1.0
1.0
1.0
1.0
1.0
1.0
1.0
1.0
1.0
1.0
1.0
1.0
1.0
1.0


11384.0
14.0
0.0
0.0
0.0
0.0
0.0
0.0
0.0
0.0
0.0
0.0
0.0
0.0
0.0
0.0
0.0
0.0
469.0
0.0
0.0
0.0
0.0
0.0
0.0
10901.0
0.0
0.0
0.0
0.0


11384.0
14.0
0.0
0.0
0.0
0.0
0.0
0.0
0.0
0.0
0.0
0.0
0.0
0.0
0.0
0.0
0.0
0.0
469.0
0.0
0.0
0.0
0.0
0.0
0.0
10901.0
0.0
0.0
0.0
0.0

1.0
1.0
1.0
1.0
1.0
1.0
1.0
1.0
1.0
1.0
1.0
1.0
1.0
1.0
1.0
1.0
1.0
1.0
1.0
1.0
1.0
1.0
1.0
1.0
1.0
1.0
1.0
1.0
1.0
1.0


493.0
0.0
4.0
6.0
0.0
0.0
0.0
0.0
0.0
0.0
3.0
0.0
0.0
113.0
367.0
0.0
0.0
0.0
0.0
0.0
0.0
0.0
0.0
0.0
0.0
0.0
0.0
0.0
0.0
0.0

0.97
0.97
0.97
0.97
0.97
0.97
0.97
0.97
0.97
0.97
0.97
0.97
0.97
0.97
0.97
0.97
0.97
0.97
0.97
0.97
0.97
0.97
0.97
0.97
0.97
0.97
0.97
0.97
0.97
0.97


94.0
0.0
0.0
0.0
0.0
0.0
0.0
0.0
0.0
0.0
0.0
0.0
0.0
18.0
76.0
0.0
0.0
0.0
0.0
0.0
0.0
0.0
0.0
0.0
0.0
0.0
0.0
0.0
0.0
0.0

0.95
0.95
0.95
0.95
0.95
0.95
0.95
0.95
0.95
0.95
0.95
0.95
0.95
0.95
0.95
0.95
0.95
0.95
0.95
0.95
0.95
0.95
0.95
0.95
0.95
0.95
0.95
0.95
0.95
0.95


3224.0
0.0
0.0
0.0
0.0
0.0
0.0
0.0
0.0
0.0
0.0
0.0
0.0
0.0
0.0
548.0
687.0
0.0
0.0
0.0
0.0
0.0
0.0
0.0
0.0
0.0
0.0
1038.0
951.0
0.0


3224.0
0.0
0.0
0.0
0.0
0.0
0.0
0.0
0.0
0.0
0.0
0.0
0.0
0.0
0.0
548.0
687.0
0.0
0.0
0.0
0.0
0.0
0.0
0.0
0.0
0.0
0.0
1038.0
951.0
0.0

1.0
1.0
1.0
1.0
1.0
1.0
1.0
1.0
1.0
1.0
1.0
1.0
1.0
1.0
1.0
1.0
1.0
1.0
1.0
1.0
1.0
1.0
1.0
1.0
1.0
1.0
1.0
1.0
1.0
1.0


2056.0
0.0
0.0
0.0
0.0
0.0
0.0
0.0
0.0
0.0
0.0
0.0
0.0
0.0
1619.0
0.0
0.0
0.0
0.0
0.0
0.0
0.0
0.0
0.0
437.0
0.0
0.0
0.0
0.0
0.0


2056.0
0.0
0.0
0.0
0.0
0.0
0.0
0.0
0.0
0.0
0.0
0.0
0.0
0.0
1619.0
0.0
0.0
0.0
0.0
0.0
0.0
0.0
0.0
0.0
437.0
0.0
0.0
0.0
0.0
0.0

1.0
1.0
1.0
1.0
1.0
1.0
1.0
1.0
1.0
1.0
1.0
1.0
1.0
1.0
1.0
1.0
1.0
1.0
1.0
1.0
1.0
1.0
1.0
1.0
1.0
1.0
1.0
1.0
1.0
1.0


18.0
0.0
0.0
0.0
0.0
0.0
0.0
0.0
0.0
0.0
0.0
0.0
0.0
0.0
0.0
0.0
0.0
0.0
0.0
18.0
0.0
0.0
0.0
0.0
0.0
0.0
0.0
0.0
0.0
0.0

0.83
0.83
0.83
0.83
0.83
0.83
0.83
0.83
0.83
0.83
0.83
0.83
0.83
0.83
0.83
0.83
0.83
0.83
0.83
0.83
0.83
0.83
0.83
0.83
0.83
0.83
0.83
0.83
0.83
0.83


14.0
0.0
0.0
0.0
0.0
0.0
0.0
0.0
0.0
0.0
0.0
0.0
0.0
14.0
0.0
0.0
0.0
0.0
0.0
0.0
0.0
0.0
0.0
0.0
0.0
0.0
0.0
0.0
0.0
0.0

0.86
0.86
0.86
0.86
0.86
0.86
0.86
0.86
0.86
0.86
0.86
0.86
0.86
0.86
0.86
0.86
0.86
0.86
0.86
0.86
0.86
0.86
0.86
0.86
0.86
0.86
0.86
0.86
0.86
0.86


19235.0
255.0
904.0
2034.0
10074.0
2100.0
497.0
0.0
0.0
42.0
292.0
1604.0
0.0
0.0
0.0
0.0
0.0
0.0
0.0
0.0
0.0
0.0
0.0
0.0
1183.0
0.0
0.0
0.0
250.0
0.0

1.0
1.0
1.0
1.0
1.0
1.0
1.0
1.0
1.0
1.0
1.0
1.0
1.0
1.0
1.0
1.0
1.0
1.0
1.0
1.0
1.0
1.0
1.0
1.0
1.0
1.0
1.0
1.0
1.0
1.0


6062.0
255.0
842.0
1740.0
329.0
0.0
0.0
0.0
0.0
0.0
99.0
1426.0
0.0
0.0
0.0
0.0
0.0
0.0
0.0
0.0
0.0
0.0
0.0
0.0
1121.0
0.0
0.0
0.0
250.0
0.0


6062.0
255.0
842.0
1740.0
329.0
0.0
0.0
0.0
0.0
0.0
99.0
1426.0
0.0
0.0
0.0
0.0
0.0
0.0
0.0
0.0
0.0
0.0
0.0
0.0
1121.0
0.0
0.0
0.0
250.0
0.0

1.0
1.0
1.0
1.0
1.0
1.0
1.0
1.0
1.0
1.0
1.0
1.0
1.0
1.0
1.0
1.0
1.0
1.0
1.0
1.0
1.0
1.0
1.0
1.0
1.0
1.0
1.0
1.0
1.0
1.0


477.0
0.0
0.0
0.0
0.0
0.0
477.0
0.0
0.0
0.0
0.0
0.0
0.0
0.0
0.0
0.0
0.0
0.0
0.0
0.0
0.0
0.0
0.0
0.0
0.0
0.0
0.0
0.0
0.0
0.0


477.0
0.0
0.0
0.0
0.0
0.0
477.0
0.0
0.0
0.0
0.0
0.0
0.0
0.0
0.0
0.0
0.0
0.0
0.0
0.0
0.0
0.0
0.0
0.0
0.0
0.0
0.0
0.0
0.0
0.0

1.0
1.0
1.0
1.0
1.0
1.0
1.0
1.0
1.0
1.0
1.0
1.0
1.0
1.0
1.0
1.0
1.0
1.0
1.0
1.0
1.0
1.0
1.0
1.0
1.0
1.0
1.0
1.0
1.0
1.0


258.0
0.0
0.0
0.0
0.0
65.0
0.0
0.0
0.0
0.0
193.0
0.0
0.0
0.0
0.0
0.0
0.0
0.0
0.0
0.0
0.0
0.0
0.0
0.0
0.0
0.0
0.0
0.0
0.0
0.0

1.0
1.0
1.0
1.0
1.0
1.0
1.0
1.0
1.0
1.0
1.0
1.0
1.0
1.0
1.0
1.0
1.0
1.0
1.0
1.0
1.0
1.0
1.0
1.0
1.0
1.0
1.0
1.0
1.0
1.0


138071.0
719.0
1809.0
2666.0
599.0
589.0
419.0
0.0
458.0
3529.0
4591.0
8170.0
629.0
5626.0
5599.0
31169.0
0.0
717.0
10950.0
13682.0
2320.0
0.0
1667.0
629.0
7949.0
0.0
429.0
3634.0
11.0
29511.0

0.99
0.99
0.99
0.99
0.99
0.99
0.99
0.99
0.99
0.99
0.99
0.99
0.99
0.99
0.99
0.99
0.99
0.99
0.99
0.99
0.99
0.99
0.99
0.99
0.99
0.99
0.99
0.99
0.99
0.99


8851.0
0.0
0.0
0.0
0.0
0.0
0.0
0.0
0.0
0.0
1649.0
7202.0
0.0
0.0
0.0
0.0
0.0
0.0
0.0
0.0
0.0
0.0
0.0
0.0
0.0
0.0
0.0
0.0
0.0
0.0


8851.0
0.0
0.0
0.0
0.0
0.0
0.0
0.0
0.0
0.0
1649.0
7202.0
0.0
0.0
0.0
0.0
0.0
0.0
0.0
0.0
0.0
0.0
0.0
0.0
0.0
0.0
0.0
0.0
0.0
0.0

1.0
1.0
1.0
1.0
1.0
1.0
1.0
1.0
1.0
1.0
1.0
1.0
1.0
1.0
1.0
1.0
1.0
1.0
1.0
1.0
1.0
1.0
1.0
1.0
1.0
1.0
1.0
1.0
1.0
1.0


3682.0
49.0
0.0
0.0
0.0
0.0
0.0
0.0
457.0
3033.0
143.0
0.0
0.0
0.0
0.0
0.0
0.0
0.0
0.0
0.0
0.0
0.0
0.0
0.0
0.0
0.0
0.0
0.0
0.0
0.0


3682.0
49.0
0.0
0.0
0.0
0.0
0.0
0.0
457.0
3033.0
143.0
0.0
0.0
0.0
0.0
0.0
0.0
0.0
0.0
0.0
0.0
0.0
0.0
0.0
0.0
0.0
0.0
0.0
0.0
0.0

1.0
1.0
1.0
1.0
1.0
1.0
1.0
1.0
1.0
1.0
1.0
1.0
1.0
1.0
1.0
1.0
1.0
1.0
1.0
1.0
1.0
1.0
1.0
1.0
1.0
1.0
1.0
1.0
1.0
1.0


5711.0
0.0
771.0
2387.0
380.0
0.0
13.0
0.0
0.0
0.0
0.0
0.0
0.0
2160.0
0.0
0.0
0.0
0.0
0.0
0.0
0.0
0.0
0.0
0.0
0.0
0.0
0.0
0.0
0.0
0.0

0.74
0.74
0.74
0.74
0.74
0.74
0.74
0.74
0.74
0.74
0.74
0.74
0.74
0.74
0.74
0.74
0.74
0.74
0.74
0.74
0.74
0.74
0.74
0.74
0.74
0.74
0.74
0.74
0.74
0.74


3348.0
0.0
731.0
2257.0
360.0
0.0
0.0
0.0
0.0
0.0
0.0
0.0
0.0
0.0
0.0
0.0
0.0
0.0
0.0
0.0
0.0
0.0
0.0
0.0
0.0
0.0
0.0
0.0
0.0
0.0

1.0
1.0
1.0
1.0
1.0
1.0
1.0
1.0
1.0
1.0
1.0
1.0
1.0
1.0
1.0
1.0
1.0
1.0
1.0
1.0
1.0
1.0
1.0
1.0
1.0
1.0
1.0
1.0
1.0
1.0


2306.0
0.0
40.0
130.0
20.0
0.0
0.0
0.0
0.0
0.0
0.0
0.0
0.0
2116.0
0.0
0.0
0.0
0.0
0.0
0.0
0.0
0.0
0.0
0.0
0.0
0.0
0.0
0.0
0.0
0.0

1.0
1.0
1.0
1.0
1.0
1.0
1.0
1.0
1.0
1.0
1.0
1.0
1.0
1.0
1.0
1.0
1.0
1.0
1.0
1.0
1.0
1.0
1.0
1.0
1.0
1.0
1.0
1.0
1.0
1.0


13.0
0.0
0.0
0.0
0.0
0.0
13.0
0.0
0.0
0.0
0.0
0.0
0.0
0.0
0.0
0.0
0.0
0.0
0.0
0.0
0.0
0.0
0.0
0.0
0.0
0.0
0.0
0.0
0.0
0.0

1.0
1.0
1.0
1.0
1.0
1.0
1.0
1.0
1.0
1.0
1.0
1.0
1.0
1.0
1.0
1.0
1.0
1.0
1.0
1.0
1.0
1.0
1.0
1.0
1.0
1.0
1.0
1.0
1.0
1.0


3009.0
281.0
404.0
0.0
0.0
0.0
0.0
0.0
0.0
112.0
1955.0
257.0
0.0
0.0
0.0
0.0
0.0
0.0
0.0
0.0
0.0
0.0
0.0
0.0
0.0
0.0
0.0
0.0
0.0
0.0


1761.0
74.0
15.0
0.0
0.0
0.0
0.0
0.0
0.0
0.0
1662.0
10.0
0.0
0.0
0.0
0.0
0.0
0.0
0.0
0.0
0.0
0.0
0.0
0.0
0.0
0.0
0.0
0.0
0.0
0.0

1.0
1.0
1.0
1.0
1.0
1.0
1.0
1.0
1.0
1.0
1.0
1.0
1.0
1.0
1.0
1.0
1.0
1.0
1.0
1.0
1.0
1.0
1.0
1.0
1.0
1.0
1.0
1.0
1.0
1.0


788.0
152.0
389.0
0.0
0.0
0.0
0.0
0.0
0.0
0.0
0.0
247.0
0.0
0.0
0.0
0.0
0.0
0.0
0.0
0.0
0.0
0.0
0.0
0.0
0.0
0.0
0.0
0.0
0.0
0.0

1.0
1.0
1.0
1.0
1.0
1.0
1.0
1.0
1.0
1.0
1.0
1.0
1.0
1.0
1.0
1.0
1.0
1.0
1.0
1.0
1.0
1.0
1.0
1.0
1.0
1.0
1.0
1.0
1.0
1.0


460.0
55.0
0.0
0.0
0.0
0.0
0.0
0.0
0.0
112.0
293.0
0.0
0.0
0.0
0.0
0.0
0.0
0.0
0.0
0.0
0.0
0.0
0.0
0.0
0.0
0.0
0.0
0.0
0.0
0.0

1.0
1.0
1.0
1.0
1.0
1.0
1.0
1.0
1.0
1.0
1.0
1.0
1.0
1.0
1.0
1.0
1.0
1.0
1.0
1.0
1.0
1.0
1.0
1.0
1.0
1.0
1.0
1.0
1.0
1.0


813.0
283.0
93.0
0.0
0.0
0.0
0.0
0.0
0.0
0.0
108.0
294.0
0.0
0.0
0.0
0.0
0.0
0.0
0.0
0.0
0.0
0.0
0.0
0.0
0.0
0.0
27.0
0.0
0.0
8.0

0.77
0.77
0.77
0.77
0.77
0.77
0.77
0.77
0.77
0.77
0.77
0.77
0.77
0.77
0.77
0.77
0.77
0.77
0.77
0.77
0.77
0.77
0.77
0.77
0.77
0.77
0.77
0.77
0.77
0.77


778.0
283.0
93.0
0.0
0.0
0.0
0.0
0.0
0.0
0.0
108.0
294.0
0.0
0.0
0.0
0.0
0.0
0.0
0.0
0.0
0.0
0.0
0.0
0.0
0.0
0.0
0.0
0.0
0.0
0.0

1.0
1.0
1.0
1.0
1.0
1.0
1.0
1.0
1.0
1.0
1.0
1.0
1.0
1.0
1.0
1.0
1.0
1.0
1.0
1.0
1.0
1.0
1.0
1.0
1.0
1.0
1.0
1.0
1.0
1.0


57896.0
0.0
0.0
31.0
182.0
476.0
308.0
0.0
1.0
0.0
0.0
0.0
0.0
0.0
5315.0
70.0
0.0
2.0
9770.0
10463.0
2320.0
0.0
0.0
0.0
0.0
0.0
0.0
0.0
0.0
28958.0

1.0
1.0
1.0
1.0
1.0
1.0
1.0
1.0
1.0
1.0
1.0
1.0
1.0
1.0
1.0
1.0
1.0
1.0
1.0
1.0
1.0
1.0
1.0
1.0
1.0
1.0
1.0
1.0
1.0
1.0


643.0
0.0
0.0
0.0
0.0
0.0
0.0
0.0
0.0
0.0
488.0
155.0
0.0
0.0
0.0
0.0
0.0
0.0
0.0
0.0
0.0
0.0
0.0
0.0
0.0
0.0
0.0
0.0
0.0
0.0

1.0
1.0
1.0
1.0
1.0
1.0
1.0
1.0
1.0
1.0
1.0
1.0
1.0
1.0
1.0
1.0
1.0
1.0
1.0
1.0
1.0
1.0
1.0
1.0
1.0
1.0
1.0
1.0
1.0
1.0


409.0
0.0
365.0
0.0
0.0
0.0
0.0
0.0
0.0
0.0
0.0
44.0
0.0
0.0
0.0
0.0
0.0
0.0
0.0
0.0
0.0
0.0
0.0
0.0
0.0
0.0
0.0
0.0
0.0
0.0


409.0
0.0
365.0
0.0
0.0
0.0
0.0
0.0
0.0
0.0
0.0
44.0
0.0
0.0
0.0
0.0
0.0
0.0
0.0
0.0
0.0
0.0
0.0
0.0
0.0
0.0
0.0
0.0
0.0
0.0

1.0
1.0
1.0
1.0
1.0
1.0
1.0
1.0
1.0
1.0
1.0
1.0
1.0
1.0
1.0
1.0
1.0
1.0
1.0
1.0
1.0
1.0
1.0
1.0
1.0
1.0
1.0
1.0
1.0
1.0


409.0
0.0
0.0
0.0
0.0
0.0
0.0
0.0
0.0
257.0
0.0
152.0
0.0
0.0
0.0
0.0
0.0
0.0
0.0
0.0
0.0
0.0
0.0
0.0
0.0
0.0
0.0
0.0
0.0
0.0


409.0
0.0
0.0
0.0
0.0
0.0
0.0
0.0
0.0
257.0
0.0
152.0
0.0
0.0
0.0
0.0
0.0
0.0
0.0
0.0
0.0
0.0
0.0
0.0
0.0
0.0
0.0
0.0
0.0
0.0

1.0
1.0
1.0
1.0
1.0
1.0
1.0
1.0
1.0
1.0
1.0
1.0
1.0
1.0
1.0
1.0
1.0
1.0
1.0
1.0
1.0
1.0
1.0
1.0
1.0
1.0
1.0
1.0
1.0
1.0


245.0
0.0
0.0
0.0
0.0
0.0
0.0
0.0
0.0
0.0
245.0
0.0
0.0
0.0
0.0
0.0
0.0
0.0
0.0
0.0
0.0
0.0
0.0
0.0
0.0
0.0
0.0
0.0
0.0
0.0


245.0
0.0
0.0
0.0
0.0
0.0
0.0
0.0
0.0
0.0
245.0
0.0
0.0
0.0
0.0
0.0
0.0
0.0
0.0
0.0
0.0
0.0
0.0
0.0
0.0
0.0
0.0
0.0
0.0
0.0

1.0
1.0
1.0
1.0
1.0
1.0
1.0
1.0
1.0
1.0
1.0
1.0
1.0
1.0
1.0
1.0
1.0
1.0
1.0
1.0
1.0
1.0
1.0
1.0
1.0
1.0
1.0
1.0
1.0
1.0


149.0
43.0
7.0
56.0
6.0
37.0
0.0
0.0
0.0
0.0
0.0
0.0
0.0
0.0
0.0
0.0
0.0
0.0
0.0
0.0
0.0
0.0
0.0
0.0
0.0
0.0
0.0
0.0
0.0
0.0

1.0
1.0
1.0
1.0
1.0
1.0
1.0
1.0
1.0
1.0
1.0
1.0
1.0
1.0
1.0
1.0
1.0
1.0
1.0
1.0
1.0
1.0
1.0
1.0
1.0
1.0
1.0
1.0
1.0
1.0


290.0
0.0
112.0
55.0
0.0
0.0
0.0
0.0
0.0
0.0
0.0
0.0
0.0
123.0
0.0
0.0
0.0
0.0
0.0
0.0
0.0
0.0
0.0
0.0
0.0
0.0
0.0
0.0
0.0
0.0


290.0
0.0
112.0
55.0
0.0
0.0
0.0
0.0
0.0
0.0
0.0
0.0
0.0
123.0
0.0
0.0
0.0
0.0
0.0
0.0
0.0
0.0
0.0
0.0
0.0
0.0
0.0
0.0
0.0
0.0

0.99
0.99
0.99
0.99
0.99
0.99
0.99
0.99
0.99
0.99
0.99
0.99
0.99
0.99
0.99
0.99
0.99
0.99
0.99
0.99
0.99
0.99
0.99
0.99
0.99
0.99
0.99
0.99
0.99
0.99


3121.0
9.0
51.0
108.0
0.0
0.0
0.0
0.0
0.0
0.0
0.0
0.0
0.0
2817.0
136.0
0.0
0.0
0.0
0.0
0.0
0.0
0.0
0.0
0.0
0.0
0.0
0.0
0.0
0.0
0.0


3121.0
9.0
51.0
108.0
0.0
0.0
0.0
0.0
0.0
0.0
0.0
0.0
0.0
2817.0
136.0
0.0
0.0
0.0
0.0
0.0
0.0
0.0
0.0
0.0
0.0
0.0
0.0
0.0
0.0
0.0

1.0
1.0
1.0
1.0
1.0
1.0
1.0
1.0
1.0
1.0
1.0
1.0
1.0
1.0
1.0
1.0
1.0
1.0
1.0
1.0
1.0
1.0
1.0
1.0
1.0
1.0
1.0
1.0
1.0
1.0


31.0
25.0
6.0
0.0
0.0
0.0
0.0
0.0
0.0
0.0
0.0
0.0
0.0
0.0
0.0
0.0
0.0
0.0
0.0
0.0
0.0
0.0
0.0
0.0
0.0
0.0
0.0
0.0
0.0
0.0

0.95
0.95
0.95
0.95
0.95
0.95
0.95
0.95
0.95
0.95
0.95
0.95
0.95
0.95
0.95
0.95
0.95
0.95
0.95
0.95
0.95
0.95
0.95
0.95
0.95
0.95
0.95
0.95
0.95
0.95


3030.0
0.0
0.0
18.0
0.0
0.0
25.0
0.0
0.0
0.0
0.0
0.0
0.0
0.0
0.0
0.0
0.0
0.0
0.0
0.0
0.0
0.0
0.0
0.0
0.0
0.0
0.0
2987.0
0.0
0.0


3030.0
0.0
0.0
18.0
0.0
0.0
25.0
0.0
0.0
0.0
0.0
0.0
0.0
0.0
0.0
0.0
0.0
0.0
0.0
0.0
0.0
0.0
0.0
0.0
0.0
0.0
0.0
2987.0
0.0
0.0

1.0
1.0
1.0
1.0
1.0
1.0
1.0
1.0
1.0
1.0
1.0
1.0
1.0
1.0
1.0
1.0
1.0
1.0
1.0
1.0
1.0
1.0
1.0
1.0
1.0
1.0
1.0
1.0
1.0
1.0


9.0
0.0
0.0
9.0
0.0
0.0
0.0
0.0
0.0
0.0
0.0
0.0
0.0
0.0
0.0
0.0
0.0
0.0
0.0
0.0
0.0
0.0
0.0
0.0
0.0
0.0
0.0
0.0
0.0
0.0


9.0
0.0
0.0
9.0
0.0
0.0
0.0
0.0
0.0
0.0
0.0
0.0
0.0
0.0
0.0
0.0
0.0
0.0
0.0
0.0
0.0
0.0
0.0
0.0
0.0
0.0
0.0
0.0
0.0
0.0

1.0
1.0
1.0
1.0
1.0
1.0
1.0
1.0
1.0
1.0
1.0
1.0
1.0
1.0
1.0
1.0
1.0
1.0
1.0
1.0
1.0
1.0
1.0
1.0
1.0
1.0
1.0
1.0
1.0
1.0


6144.0
0.0
0.0
0.0
0.0
0.0
0.0
0.0
0.0
0.0
0.0
0.0
629.0
0.0
0.0
0.0
0.0
0.0
0.0
3219.0
0.0
0.0
1667.0
629.0
0.0
0.0
0.0
0.0
0.0
0.0

0.93
0.93
0.93
0.93
0.93
0.93
0.93
0.93
0.93
0.93
0.93
0.93
0.93
0.93
0.93
0.93
0.93
0.93
0.93
0.93
0.93
0.93
0.93
0.93
0.93
0.93
0.93
0.93
0.93
0.93


1594.0
0.0
0.0
0.0
0.0
0.0
0.0
0.0
0.0
0.0
0.0
0.0
0.0
0.0
0.0
0.0
0.0
0.0
0.0
0.0
0.0
0.0
0.0
0.0
0.0
0.0
402.0
647.0
0.0
545.0


1594.0
0.0
0.0
0.0
0.0
0.0
0.0
0.0
0.0
0.0
0.0
0.0
0.0
0.0
0.0
0.0
0.0
0.0
0.0
0.0
0.0
0.0
0.0
0.0
0.0
0.0
402.0
647.0
0.0
545.0

1.0
1.0
1.0
1.0
1.0
1.0
1.0
1.0
1.0
1.0
1.0
1.0
1.0
1.0
1.0
1.0
1.0
1.0
1.0
1.0
1.0
1.0
1.0
1.0
1.0
1.0
1.0
1.0
1.0
1.0


441.0
0.0
0.0
0.0
0.0
0.0
0.0
0.0
0.0
0.0
0.0
0.0
0.0
350.0
80.0
0.0
0.0
0.0
0.0
0.0
0.0
0.0
0.0
0.0
0.0
0.0
0.0
0.0
11.0
0.0


430.0
0.0
0.0
0.0
0.0
0.0
0.0
0.0
0.0
0.0
0.0
0.0
0.0
350.0
80.0
0.0
0.0
0.0
0.0
0.0
0.0
0.0
0.0
0.0
0.0
0.0
0.0
0.0
0.0
0.0

0.99
0.99
0.99
0.99
0.99
0.99
0.99
0.99
0.99
0.99
0.99
0.99
0.99
0.99
0.99
0.99
0.99
0.99
0.99
0.99
0.99
0.99
0.99
0.99
0.99
0.99
0.99
0.99
0.99
0.99


11.0
0.0
0.0
0.0
0.0
0.0
0.0
0.0
0.0
0.0
0.0
0.0
0.0
0.0
0.0
0.0
0.0
0.0
0.0
0.0
0.0
0.0
0.0
0.0
0.0
0.0
0.0
0.0
11.0
0.0

1.0
1.0
1.0
1.0
1.0
1.0
1.0
1.0
1.0
1.0
1.0
1.0
1.0
1.0
1.0
1.0
1.0
1.0
1.0
1.0
1.0
1.0
1.0
1.0
1.0
1.0
1.0
1.0
1.0
1.0


8.0
0.0
0.0
0.0
0.0
0.0
0.0
0.0
0.0
0.0
0.0
0.0
0.0
6.0
0.0
0.0
0.0
0.0
0.0
0.0
0.0
0.0
0.0
0.0
2.0
0.0
0.0
0.0
0.0
0.0


8.0
0.0
0.0
0.0
0.0
0.0
0.0
0.0
0.0
0.0
0.0
0.0
0.0
6.0
0.0
0.0
0.0
0.0
0.0
0.0
0.0
0.0
0.0
0.0
2.0
0.0
0.0
0.0
0.0
0.0

1.0
1.0
1.0
1.0
1.0
1.0
1.0
1.0
1.0
1.0
1.0
1.0
1.0
1.0
1.0
1.0
1.0
1.0
1.0
1.0
1.0
1.0
1.0
1.0
1.0
1.0
1.0
1.0
1.0
1.0


57770.0
288.0
150.0
310.0
130.0
33.0
87.0
518.0
1266.0
0.0
198.0
443.0
329.0
685.0
97.0
3.0
653.0
1737.0
0.0
7158.0
1697.0
1676.0
3808.0
329.0
26700.0
0.0
3103.0
4384.0
470.0
1518.0


57770.0
288.0
150.0
310.0
130.0
33.0
87.0
518.0
1266.0
0.0
198.0
443.0
329.0
685.0
97.0
3.0
653.0
1737.0
0.0
7158.0
1697.0
1676.0
3808.0
329.0
26700.0
0.0
3103.0
4384.0
470.0
1518.0


57770.0
288.0
150.0
310.0
130.0
33.0
87.0
518.0
1266.0
0.0
198.0
443.0
329.0
685.0
97.0
3.0
653.0
1737.0
0.0
7158.0
1697.0
1676.0
3808.0
329.0
26700.0
0.0
3103.0
4384.0
470.0
1518.0

1.0
1.0
1.0
1.0
1.0
1.0
1.0
1.0
1.0
1.0
1.0
1.0
1.0
1.0
1.0
1.0
1.0
1.0
1.0
1.0
1.0
1.0
1.0
1.0
1.0
1.0
1.0
1.0
1.0
1.0


6385.0
555.0
68.0
10.0
0.0
672.0
0.0
86.0
0.0
0.0
0.0
14.0
0.0
430.0
259.0
0.0
0.0
0.0
0.0
1036.0
0.0
0.0
0.0
0.0
0.0
0.0
290.0
237.0
2362.0
366.0


1276.0
544.0
60.0
0.0
0.0
672.0
0.0
0.0
0.0
0.0
0.0
0.0
0.0
0.0
0.0
0.0
0.0
0.0
0.0
0.0
0.0
0.0
0.0
0.0
0.0
0.0
0.0
0.0
0.0
0.0


1276.0
544.0
60.0
0.0
0.0
672.0
0.0
0.0
0.0
0.0
0.0
0.0
0.0
0.0
0.0
0.0
0.0
0.0
0.0
0.0
0.0
0.0
0.0
0.0
0.0
0.0
0.0
0.0
0.0
0.0

1.0
1.0
1.0
1.0
1.0
1.0
1.0
1.0
1.0
1.0
1.0
1.0
1.0
1.0
1.0
1.0
1.0
1.0
1.0
1.0
1.0
1.0
1.0
1.0
1.0
1.0
1.0
1.0
1.0
1.0


5109.0
11.0
8.0
10.0
0.0
0.0
0.0
86.0
0.0
0.0
0.0
14.0
0.0
430.0
259.0
0.0
0.0
0.0
0.0
1036.0
0.0
0.0
0.0
0.0
0.0
0.0
290.0
237.0
2362.0
366.0


3783.0
11.0
8.0
10.0
0.0
0.0
0.0
86.0
0.0
0.0
0.0
14.0
0.0
430.0
259.0
0.0
0.0
0.0
0.0
0.0
0.0
0.0
0.0
0.0
0.0
0.0
0.0
237.0
2362.0
366.0

1.0
1.0
1.0
1.0
1.0
1.0
1.0
1.0
1.0
1.0
1.0
1.0
1.0
1.0
1.0
1.0
1.0
1.0
1.0
1.0
1.0
1.0
1.0
1.0
1.0
1.0
1.0
1.0
1.0
1.0


1326.0
0.0
0.0
0.0
0.0
0.0
0.0
0.0
0.0
0.0
0.0
0.0
0.0
0.0
0.0
0.0
0.0
0.0
0.0
1036.0
0.0
0.0
0.0
0.0
0.0
0.0
290.0
0.0
0.0
0.0

0.99
0.99
0.99
0.99
0.99
0.99
0.99
0.99
0.99
0.99
0.99
0.99
0.99
0.99
0.99
0.99
0.99
0.99
0.99
0.99
0.99
0.99
0.99
0.99
0.99
0.99
0.99
0.99
0.99
0.99


400061.0
1.0
93.0
193.0
772.0
637.0
1260.0
1.0
2.0
1.0
1.0
440.0
0.0
0.0
47489.0
578.0
70.0
0.0
0.0
32988.0
33607.0
79762.0
88460.0
0.0
113662.0
0.0
0.0
7.0
0.0
37.0

0.71
0.71
0.71
0.71
0.71
0.71
0.71
0.71
0.71
0.71
0.71
0.71
0.71
0.71
0.71
0.71
0.71
0.71
0.71
0.71
0.71
0.71
0.71
0.71
0.71
0.71
0.71
0.71
0.71
0.71


211042.0
1.0
8.0
0.0
0.0
637.0
1260.0
0.0
2.0
0.0
0.0
0.0
0.0
0.0
0.0
0.0
0.0
0.0
0.0
20261.0
29781.0
79762.0
79286.0
0.0
0.0
0.0
0.0
7.0
0.0
37.0

0.85
0.85
0.85
0.85
0.85
0.85
0.85
0.85
0.85
0.85
0.85
0.85
0.85
0.85
0.85
0.85
0.85
0.85
0.85
0.85
0.85
0.85
0.85
0.85
0.85
0.85
0.85
0.85
0.85
0.85


1269.0
1.0
8.0
0.0
0.0
0.0
1260.0
0.0
0.0
0.0
0.0
0.0
0.0
0.0
0.0
0.0
0.0
0.0
0.0
0.0
0.0
0.0
0.0
0.0
0.0
0.0
0.0
0.0
0.0
0.0

1.0
1.0
1.0
1.0
1.0
1.0
1.0
1.0
1.0
1.0
1.0
1.0
1.0
1.0
1.0
1.0
1.0
1.0
1.0
1.0
1.0
1.0
1.0
1.0
1.0
1.0
1.0
1.0
1.0
1.0


59453.0
0.0
0.0
0.0
0.0
637.0
0.0
0.0
2.0
0.0
0.0
0.0
0.0
0.0
0.0
0.0
0.0
0.0
0.0
604.0
3901.0
1285.0
53019.0
0.0
0.0
0.0
0.0
5.0
0.0
0.0

1.0
1.0
1.0
1.0
1.0
1.0
1.0
1.0
1.0
1.0
1.0
1.0
1.0
1.0
1.0
1.0
1.0
1.0
1.0
1.0
1.0
1.0
1.0
1.0
1.0
1.0
1.0
1.0
1.0
1.0


162081.0
0.0
85.0
77.0
772.0
0.0
0.0
1.0
0.0
1.0
1.0
0.0
0.0
0.0
47482.0
0.0
0.0
0.0
0.0
0.0
0.0
0.0
0.0
0.0
113662.0
0.0
0.0
0.0
0.0
0.0


162081.0
0.0
85.0
77.0
772.0
0.0
0.0
1.0
0.0
1.0
1.0
0.0
0.0
0.0
47482.0
0.0
0.0
0.0
0.0
0.0
0.0
0.0
0.0
0.0
113662.0
0.0
0.0
0.0
0.0
0.0

1.0
1.0
1.0
1.0
1.0
1.0
1.0
1.0
1.0
1.0
1.0
1.0
1.0
1.0
1.0
1.0
1.0
1.0
1.0
1.0
1.0
1.0
1.0
1.0
1.0
1.0
1.0
1.0
1.0
1.0


440.0
0.0
0.0
0.0
0.0
0.0
0.0
0.0
0.0
0.0
0.0
440.0
0.0
0.0
0.0
0.0
0.0
0.0
0.0
0.0
0.0
0.0
0.0
0.0
0.0
0.0
0.0
0.0
0.0
0.0


440.0
0.0
0.0
0.0
0.0
0.0
0.0
0.0
0.0
0.0
0.0
440.0
0.0
0.0
0.0
0.0
0.0
0.0
0.0
0.0
0.0
0.0
0.0
0.0
0.0
0.0
0.0
0.0
0.0
0.0

1.0
1.0
1.0
1.0
1.0
1.0
1.0
1.0
1.0
1.0
1.0
1.0
1.0
1.0
1.0
1.0
1.0
1.0
1.0
1.0
1.0
1.0
1.0
1.0
1.0
1.0
1.0
1.0
1.0
1.0


116.0
0.0
0.0
116.0
0.0
0.0
0.0
0.0
0.0
0.0
0.0
0.0
0.0
0.0
0.0
0.0
0.0
0.0
0.0
0.0
0.0
0.0
0.0
0.0
0.0
0.0
0.0
0.0
0.0
0.0


116.0
0.0
0.0
116.0
0.0
0.0
0.0
0.0
0.0
0.0
0.0
0.0
0.0
0.0
0.0
0.0
0.0
0.0
0.0
0.0
0.0
0.0
0.0
0.0
0.0
0.0
0.0
0.0
0.0
0.0

1.0
1.0
1.0
1.0
1.0
1.0
1.0
1.0
1.0
1.0
1.0
1.0
1.0
1.0
1.0
1.0
1.0
1.0
1.0
1.0
1.0
1.0
1.0
1.0
1.0
1.0
1.0
1.0
1.0
1.0


26375.0
0.0
0.0
0.0
0.0
0.0
0.0
0.0
0.0
0.0
0.0
0.0
0.0
0.0
0.0
578.0
70.0
0.0
0.0
12727.0
3826.0
0.0
9174.0
0.0
0.0
0.0
0.0
0.0
0.0
0.0


26375.0
0.0
0.0
0.0
0.0
0.0
0.0
0.0
0.0
0.0
0.0
0.0
0.0
0.0
0.0
578.0
70.0
0.0
0.0
12727.0
3826.0
0.0
9174.0
0.0
0.0
0.0
0.0
0.0
0.0
0.0

1.0
1.0
1.0
1.0
1.0
1.0
1.0
1.0
1.0
1.0
1.0
1.0
1.0
1.0
1.0
1.0
1.0
1.0
1.0
1.0
1.0
1.0
1.0
1.0
1.0
1.0
1.0
1.0
1.0
1.0


137956.0
1384.0
0.0
157.0
1249.0
423.0
544.0
0.0
0.0
0.0
0.0
0.0
38758.0
0.0
0.0
11001.0
0.0
0.0
0.0
0.0
36159.0
0.0
0.0
38758.0
0.0
0.0
9523.0
0.0
0.0
0.0

0.7
0.7
0.7
0.7
0.7
0.7
0.7
0.7
0.7
0.7
0.7
0.7
0.7
0.7
0.7
0.7
0.7
0.7
0.7
0.7
0.7
0.7
0.7
0.7
0.7
0.7
0.7
0.7
0.7
0.7


1510.0
0.0
0.0
1.0
965.0
0.0
544.0
0.0
0.0
0.0
0.0
0.0
0.0
0.0
0.0
0.0
0.0
0.0
0.0
0.0
0.0
0.0
0.0
0.0
0.0
0.0
0.0
0.0
0.0
0.0


965.0
0.0
0.0
0.0
965.0
0.0
0.0
0.0
0.0
0.0
0.0
0.0
0.0
0.0
0.0
0.0
0.0
0.0
0.0
0.0
0.0
0.0
0.0
0.0
0.0
0.0
0.0
0.0
0.0
0.0

0.99
0.99
0.99
0.99
0.99
0.99
0.99
0.99
0.99
0.99
0.99
0.99
0.99
0.99
0.99
0.99
0.99
0.99
0.99
0.99
0.99
0.99
0.99
0.99
0.99
0.99
0.99
0.99
0.99
0.99


545.0
0.0
0.0
1.0
0.0
0.0
544.0
0.0
0.0
0.0
0.0
0.0
0.0
0.0
0.0
0.0
0.0
0.0
0.0
0.0
0.0
0.0
0.0
0.0
0.0
0.0
0.0
0.0
0.0
0.0

1.0
1.0
1.0
1.0
1.0
1.0
1.0
1.0
1.0
1.0
1.0
1.0
1.0
1.0
1.0
1.0
1.0
1.0
1.0
1.0
1.0
1.0
1.0
1.0
1.0
1.0
1.0
1.0
1.0
1.0


78734.0
639.0
0.0
156.0
0.0
423.0
0.0
0.0
0.0
0.0
0.0
0.0
38758.0
0.0
0.0
0.0
0.0
0.0
0.0
0.0
0.0
0.0
0.0
38758.0
0.0
0.0
0.0
0.0
0.0
0.0

0.92
0.92
0.92
0.92
0.92
0.92
0.92
0.92
0.92
0.92
0.92
0.92
0.92
0.92
0.92
0.92
0.92
0.92
0.92
0.92
0.92
0.92
0.92
0.92
0.92
0.92
0.92
0.92
0.92
0.92


78524.0
639.0
0.0
156.0
0.0
423.0
0.0
0.0
0.0
0.0
0.0
0.0
38653.0
0.0
0.0
0.0
0.0
0.0
0.0
0.0
0.0
0.0
0.0
38653.0
0.0
0.0
0.0
0.0
0.0
0.0

1.0
1.0
1.0
1.0
1.0
1.0
1.0
1.0
1.0
1.0
1.0
1.0
1.0
1.0
1.0
1.0
1.0
1.0
1.0
1.0
1.0
1.0
1.0
1.0
1.0
1.0
1.0
1.0
1.0
1.0


679.0
679.0
0.0
0.0
0.0
0.0
0.0
0.0
0.0
0.0
0.0
0.0
0.0
0.0
0.0
0.0
0.0
0.0
0.0
0.0
0.0
0.0
0.0
0.0
0.0
0.0
0.0
0.0
0.0
0.0


679.0
679.0
0.0
0.0
0.0
0.0
0.0
0.0
0.0
0.0
0.0
0.0
0.0
0.0
0.0
0.0
0.0
0.0
0.0
0.0
0.0
0.0
0.0
0.0
0.0
0.0
0.0
0.0
0.0
0.0

1.0
1.0
1.0
1.0
1.0
1.0
1.0
1.0
1.0
1.0
1.0
1.0
1.0
1.0
1.0
1.0
1.0
1.0
1.0
1.0
1.0
1.0
1.0
1.0
1.0
1.0
1.0
1.0
1.0
1.0


36475.0
66.0
0.0
0.0
284.0
0.0
0.0
0.0
0.0
0.0
0.0
0.0
0.0
0.0
0.0
0.0
0.0
0.0
0.0
0.0
36125.0
0.0
0.0
0.0
0.0
0.0
0.0
0.0
0.0
0.0


36409.0
0.0
0.0
0.0
284.0
0.0
0.0
0.0
0.0
0.0
0.0
0.0
0.0
0.0
0.0
0.0
0.0
0.0
0.0
0.0
36125.0
0.0
0.0
0.0
0.0
0.0
0.0
0.0
0.0
0.0

1.0
1.0
1.0
1.0
1.0
1.0
1.0
1.0
1.0
1.0
1.0
1.0
1.0
1.0
1.0
1.0
1.0
1.0
1.0
1.0
1.0
1.0
1.0
1.0
1.0
1.0
1.0
1.0
1.0
1.0


66.0
66.0
0.0
0.0
0.0
0.0
0.0
0.0
0.0
0.0
0.0
0.0
0.0
0.0
0.0
0.0
0.0
0.0
0.0
0.0
0.0
0.0
0.0
0.0
0.0
0.0
0.0
0.0
0.0
0.0

1.0
1.0
1.0
1.0
1.0
1.0
1.0
1.0
1.0
1.0
1.0
1.0
1.0
1.0
1.0
1.0
1.0
1.0
1.0
1.0
1.0
1.0
1.0
1.0
1.0
1.0
1.0
1.0
1.0
1.0


11274.0
0.0
0.0
0.0
0.0
0.0
0.0
0.0
0.0
0.0
0.0
0.0
0.0
0.0
0.0
11001.0
0.0
0.0
0.0
0.0
0.0
0.0
0.0
0.0
0.0
0.0
273.0
0.0
0.0
0.0

1.0
1.0
1.0
1.0
1.0
1.0
1.0
1.0
1.0
1.0
1.0
1.0
1.0
1.0
1.0
1.0
1.0
1.0
1.0
1.0
1.0
1.0
1.0
1.0
1.0
1.0
1.0
1.0
1.0
1.0


9250.0
0.0
0.0
0.0
0.0
0.0
0.0
0.0
0.0
0.0
0.0
0.0
0.0
0.0
0.0
0.0
0.0
0.0
0.0
0.0
0.0
0.0
0.0
0.0
0.0
0.0
9250.0
0.0
0.0
0.0


9250.0
0.0
0.0
0.0
0.0
0.0
0.0
0.0
0.0
0.0
0.0
0.0
0.0
0.0
0.0
0.0
0.0
0.0
0.0
0.0
0.0
0.0
0.0
0.0
0.0
0.0
9250.0
0.0
0.0
0.0

1.0
1.0
1.0
1.0
1.0
1.0
1.0
1.0
1.0
1.0
1.0
1.0
1.0
1.0
1.0
1.0
1.0
1.0
1.0
1.0
1.0
1.0
1.0
1.0
1.0
1.0
1.0
1.0
1.0
1.0


960.0
0.0
0.0
0.0
0.0
960.0
0.0
0.0
0.0
0.0
0.0
0.0
0.0
0.0
0.0
0.0
0.0
0.0
0.0
0.0
0.0
0.0
0.0
0.0
0.0
0.0
0.0
0.0
0.0
0.0


960.0
0.0
0.0
0.0
0.0
960.0
0.0
0.0
0.0
0.0
0.0
0.0
0.0
0.0
0.0
0.0
0.0
0.0
0.0
0.0
0.0
0.0
0.0
0.0
0.0
0.0
0.0
0.0
0.0
0.0

1.0
1.0
1.0
1.0
1.0
1.0
1.0
1.0
1.0
1.0
1.0
1.0
1.0
1.0
1.0
1.0
1.0
1.0
1.0
1.0
1.0
1.0
1.0
1.0
1.0
1.0
1.0
1.0
1.0
1.0


635.0
0.0
0.0
0.0
0.0
0.0
0.0
0.0
0.0
0.0
635.0
0.0
0.0
0.0
0.0
0.0
0.0
0.0
0.0
0.0
0.0
0.0
0.0
0.0
0.0
0.0
0.0
0.0
0.0
0.0


635.0
0.0
0.0
0.0
0.0
0.0
0.0
0.0
0.0
0.0
635.0
0.0
0.0
0.0
0.0
0.0
0.0
0.0
0.0
0.0
0.0
0.0
0.0
0.0
0.0
0.0
0.0
0.0
0.0
0.0

1.0
1.0
1.0
1.0
1.0
1.0
1.0
1.0
1.0
1.0
1.0
1.0
1.0
1.0
1.0
1.0
1.0
1.0
1.0
1.0
1.0
1.0
1.0
1.0
1.0
1.0
1.0
1.0
1.0
1.0


2131.0
216.0
0.0
71.0
116.0
0.0
11.0
0.0
101.0
115.0
233.0
40.0
0.0
0.0
128.0
0.0
0.0
0.0
0.0
0.0
0.0
0.0
0.0
0.0
0.0
0.0
0.0
0.0
0.0
1100.0


2081.0
204.0
0.0
71.0
79.0
0.0
11.0
0.0
101.0
115.0
233.0
39.0
0.0
0.0
128.0
0.0
0.0
0.0
0.0
0.0
0.0
0.0
0.0
0.0
0.0
0.0
0.0
0.0
0.0
1100.0

0.98
0.98
0.98
0.98
0.98
0.98
0.98
0.98
0.98
0.98
0.98
0.98
0.98
0.98
0.98
0.98
0.98
0.98
0.98
0.98
0.98
0.98
0.98
0.98
0.98
0.98
0.98
0.98
0.98
0.98


1828.0
105.0
0.0
71.0
79.0
0.0
11.0
0.0
0.0
62.0
233.0
39.0
0.0
0.0
128.0
0.0
0.0
0.0
0.0
0.0
0.0
0.0
0.0
0.0
0.0
0.0
0.0
0.0
0.0
1100.0

1.0
1.0
1.0
1.0
1.0
1.0
1.0
1.0
1.0
1.0
1.0
1.0
1.0
1.0
1.0
1.0
1.0
1.0
1.0
1.0
1.0
1.0
1.0
1.0
1.0
1.0
1.0
1.0
1.0
1.0


99.0
99.0
0.0
0.0
0.0
0.0
0.0
0.0
0.0
0.0
0.0
0.0
0.0
0.0
0.0
0.0
0.0
0.0
0.0
0.0
0.0
0.0
0.0
0.0
0.0
0.0
0.0
0.0
0.0
0.0

1.0
1.0
1.0
1.0
1.0
1.0
1.0
1.0
1.0
1.0
1.0
1.0
1.0
1.0
1.0
1.0
1.0
1.0
1.0
1.0
1.0
1.0
1.0
1.0
1.0
1.0
1.0
1.0
1.0
1.0


37.0
0.0
0.0
0.0
37.0
0.0
0.0
0.0
0.0
0.0
0.0
0.0
0.0
0.0
0.0
0.0
0.0
0.0
0.0
0.0
0.0
0.0
0.0
0.0
0.0
0.0
0.0
0.0
0.0
0.0

1.0
1.0
1.0
1.0
1.0
1.0
1.0
1.0
1.0
1.0
1.0
1.0
1.0
1.0
1.0
1.0
1.0
1.0
1.0
1.0
1.0
1.0
1.0
1.0
1.0
1.0
1.0
1.0
1.0
1.0


13.0
12.0
0.0
0.0
0.0
0.0
0.0
0.0
0.0
0.0
0.0
1.0
0.0
0.0
0.0
0.0
0.0
0.0
0.0
0.0
0.0
0.0
0.0
0.0
0.0
0.0
0.0
0.0
0.0
0.0

0.77
0.77
0.77
0.77
0.77
0.77
0.77
0.77
0.77
0.77
0.77
0.77
0.77
0.77
0.77
0.77
0.77
0.77
0.77
0.77
0.77
0.77
0.77
0.77
0.77
0.77
0.77
0.77
0.77
0.77


559.0
0.0
254.0
305.0
0.0
0.0
0.0
0.0
0.0
0.0
0.0
0.0
0.0
0.0
0.0
0.0
0.0
0.0
0.0
0.0
0.0
0.0
0.0
0.0
0.0
0.0
0.0
0.0
0.0
0.0


559.0
0.0
254.0
305.0
0.0
0.0
0.0
0.0
0.0
0.0
0.0
0.0
0.0
0.0
0.0
0.0
0.0
0.0
0.0
0.0
0.0
0.0
0.0
0.0
0.0
0.0
0.0
0.0
0.0
0.0


559.0
0.0
254.0
305.0
0.0
0.0
0.0
0.0
0.0
0.0
0.0
0.0
0.0
0.0
0.0
0.0
0.0
0.0
0.0
0.0
0.0
0.0
0.0
0.0
0.0
0.0
0.0
0.0
0.0
0.0

1.0
1.0
1.0
1.0
1.0
1.0
1.0
1.0
1.0
1.0
1.0
1.0
1.0
1.0
1.0
1.0
1.0
1.0
1.0
1.0
1.0
1.0
1.0
1.0
1.0
1.0
1.0
1.0
1.0
1.0


27592.0
0.0
0.0
0.0
488.0
23.0
32.0
0.0
0.0
0.0
0.0
26.0
0.0
0.0
0.0
0.0
0.0
0.0
0.0
0.0
0.0
0.0
0.0
0.0
1204.0
22601.0
0.0
0.0
0.0
3218.0

0.79
0.79
0.79
0.79
0.79
0.79
0.79
0.79
0.79
0.79
0.79
0.79
0.79
0.79
0.79
0.79
0.79
0.79
0.79
0.79
0.79
0.79
0.79
0.79
0.79
0.79
0.79
0.79
0.79
0.79


26339.0
0.0
0.0
0.0
488.0
0.0
32.0
0.0
0.0
0.0
0.0
0.0
0.0
0.0
0.0
0.0
0.0
0.0
0.0
0.0
0.0
0.0
0.0
0.0
0.0
22601.0
0.0
0.0
0.0
3218.0


26339.0
0.0
0.0
0.0
488.0
0.0
32.0
0.0
0.0
0.0
0.0
0.0
0.0
0.0
0.0
0.0
0.0
0.0
0.0
0.0
0.0
0.0
0.0
0.0
0.0
22601.0
0.0
0.0
0.0
3218.0

1.0
1.0
1.0
1.0
1.0
1.0
1.0
1.0
1.0
1.0
1.0
1.0
1.0
1.0
1.0
1.0
1.0
1.0
1.0
1.0
1.0
1.0
1.0
1.0
1.0
1.0
1.0
1.0
1.0
1.0


23.0
0.0
0.0
0.0
0.0
23.0
0.0
0.0
0.0
0.0
0.0
0.0
0.0
0.0
0.0
0.0
0.0
0.0
0.0
0.0
0.0
0.0
0.0
0.0
0.0
0.0
0.0
0.0
0.0
0.0

0.97
0.97
0.97
0.97
0.97
0.97
0.97
0.97
0.97
0.97
0.97
0.97
0.97
0.97
0.97
0.97
0.97
0.97
0.97
0.97
0.97
0.97
0.97
0.97
0.97
0.97
0.97
0.97
0.97
0.97


1204.0
0.0
0.0
0.0
0.0
0.0
0.0
0.0
0.0
0.0
0.0
0.0
0.0
0.0
0.0
0.0
0.0
0.0
0.0
0.0
0.0
0.0
0.0
0.0
1204.0
0.0
0.0
0.0
0.0
0.0


1204.0
0.0
0.0
0.0
0.0
0.0
0.0
0.0
0.0
0.0
0.0
0.0
0.0
0.0
0.0
0.0
0.0
0.0
0.0
0.0
0.0
0.0
0.0
0.0
1204.0
0.0
0.0
0.0
0.0
0.0

1.0
1.0
1.0
1.0
1.0
1.0
1.0
1.0
1.0
1.0
1.0
1.0
1.0
1.0
1.0
1.0
1.0
1.0
1.0
1.0
1.0
1.0
1.0
1.0
1.0
1.0
1.0
1.0
1.0
1.0


942.0
97.0
126.0
0.0
0.0
0.0
429.0
0.0
0.0
0.0
0.0
290.0
0.0
0.0
0.0
0.0
0.0
0.0
0.0
0.0
0.0
0.0
0.0
0.0
0.0
0.0
0.0
0.0
0.0
0.0


513.0
97.0
126.0
0.0
0.0
0.0
0.0
0.0
0.0
0.0
0.0
290.0
0.0
0.0
0.0
0.0
0.0
0.0
0.0
0.0
0.0
0.0
0.0
0.0
0.0
0.0
0.0
0.0
0.0
0.0


513.0
97.0
126.0
0.0
0.0
0.0
0.0
0.0
0.0
0.0
0.0
290.0
0.0
0.0
0.0
0.0
0.0
0.0
0.0
0.0
0.0
0.0
0.0
0.0
0.0
0.0
0.0
0.0
0.0
0.0

1.0
1.0
1.0
1.0
1.0
1.0
1.0
1.0
1.0
1.0
1.0
1.0
1.0
1.0
1.0
1.0
1.0
1.0
1.0
1.0
1.0
1.0
1.0
1.0
1.0
1.0
1.0
1.0
1.0
1.0


429.0
0.0
0.0
0.0
0.0
0.0
429.0
0.0
0.0
0.0
0.0
0.0
0.0
0.0
0.0
0.0
0.0
0.0
0.0
0.0
0.0
0.0
0.0
0.0
0.0
0.0
0.0
0.0
0.0
0.0


429.0
0.0
0.0
0.0
0.0
0.0
429.0
0.0
0.0
0.0
0.0
0.0
0.0
0.0
0.0
0.0
0.0
0.0
0.0
0.0
0.0
0.0
0.0
0.0
0.0
0.0
0.0
0.0
0.0
0.0

1.0
1.0
1.0
1.0
1.0
1.0
1.0
1.0
1.0
1.0
1.0
1.0
1.0
1.0
1.0
1.0
1.0
1.0
1.0
1.0
1.0
1.0
1.0
1.0
1.0
1.0
1.0
1.0
1.0
1.0


1814.0
98.0
8.0
0.0
21.0
25.0
24.0
38.0
229.0
156.0
128.0
173.0
0.0
150.0
6.0
0.0
0.0
0.0
160.0
0.0
0.0
0.0
0.0
0.0
598.0
0.0
0.0
0.0
0.0
0.0


1370.0
77.0
8.0
0.0
21.0
25.0
24.0
0.0
41.0
59.0
42.0
159.0
0.0
150.0
6.0
0.0
0.0
0.0
160.0
0.0
0.0
0.0
0.0
0.0
598.0
0.0
0.0
0.0
0.0
0.0


1370.0
77.0
8.0
0.0
21.0
25.0
24.0
0.0
41.0
59.0
42.0
159.0
0.0
150.0
6.0
0.0
0.0
0.0
160.0
0.0
0.0
0.0
0.0
0.0
598.0
0.0
0.0
0.0
0.0
0.0

1.0
1.0
1.0
1.0
1.0
1.0
1.0
1.0
1.0
1.0
1.0
1.0
1.0
1.0
1.0
1.0
1.0
1.0
1.0
1.0
1.0
1.0
1.0
1.0
1.0
1.0
1.0
1.0
1.0
1.0


444.0
21.0
0.0
0.0
0.0
0.0
0.0
38.0
188.0
97.0
86.0
14.0
0.0
0.0
0.0
0.0
0.0
0.0
0.0
0.0
0.0
0.0
0.0
0.0
0.0
0.0
0.0
0.0
0.0
0.0


444.0
21.0
0.0
0.0
0.0
0.0
0.0
38.0
188.0
97.0
86.0
14.0
0.0
0.0
0.0
0.0
0.0
0.0
0.0
0.0
0.0
0.0
0.0
0.0
0.0
0.0
0.0
0.0
0.0
0.0

0.75
0.75
0.75
0.75
0.75
0.75
0.75
0.75
0.75
0.75
0.75
0.75
0.75
0.75
0.75
0.75
0.75
0.75
0.75
0.75
0.75
0.75
0.75
0.75
0.75
0.75
0.75
0.75
0.75
0.75


2647.0
120.0
146.0
7.0
123.0
503.0
164.0
0.0
6.0
68.0
293.0
86.0
0.0
0.0
44.0
57.0
0.0
0.0
0.0
83.0
0.0
0.0
0.0
0.0
0.0
129.0
173.0
18.0
213.0
414.0

0.97
0.97
0.97
0.97
0.97
0.97
0.97
0.97
0.97
0.97
0.97
0.97
0.97
0.97
0.97
0.97
0.97
0.97
0.97
0.97
0.97
0.97
0.97
0.97
0.97
0.97
0.97
0.97
0.97
0.97


657.0
11.0
6.0
1.0
0.0
0.0
0.0
0.0
6.0
68.0
266.0
66.0
0.0
0.0
44.0
57.0
0.0
0.0
0.0
83.0
0.0
0.0
0.0
0.0
0.0
0.0
0.0
0.0
0.0
49.0

1.0
1.0
1.0
1.0
1.0
1.0
1.0
1.0
1.0
1.0
1.0
1.0
1.0
1.0
1.0
1.0
1.0
1.0
1.0
1.0
1.0
1.0
1.0
1.0
1.0
1.0
1.0
1.0
1.0
1.0


106.0
0.0
0.0
0.0
0.0
0.0
0.0
0.0
0.0
0.0
0.0
0.0
0.0
0.0
0.0
57.0
0.0
0.0
0.0
0.0
0.0
0.0
0.0
0.0
0.0
0.0
0.0
0.0
0.0
49.0

1.0
1.0
1.0
1.0
1.0
1.0
1.0
1.0
1.0
1.0
1.0
1.0
1.0
1.0
1.0
1.0
1.0
1.0
1.0
1.0
1.0
1.0
1.0
1.0
1.0
1.0
1.0
1.0
1.0
1.0


44.0
0.0
0.0
0.0
0.0
0.0
0.0
0.0
0.0
0.0
0.0
0.0
0.0
0.0
44.0
0.0
0.0
0.0
0.0
0.0
0.0
0.0
0.0
0.0
0.0
0.0
0.0
0.0
0.0
0.0

0.9
0.9
0.9
0.9
0.9
0.9
0.9
0.9
0.9
0.9
0.9
0.9
0.9
0.9
0.9
0.9
0.9
0.9
0.9
0.9
0.9
0.9
0.9
0.9
0.9
0.9
0.9
0.9
0.9
0.9


1045.0
27.0
12.0
6.0
12.0
240.0
162.0
0.0
0.0
0.0
9.0
13.0
0.0
0.0
0.0
0.0
0.0
0.0
0.0
0.0
0.0
0.0
0.0
0.0
0.0
32.0
144.0
7.0
213.0
168.0


1045.0
27.0
12.0
6.0
12.0
240.0
162.0
0.0
0.0
0.0
9.0
13.0
0.0
0.0
0.0
0.0
0.0
0.0
0.0
0.0
0.0
0.0
0.0
0.0
0.0
32.0
144.0
7.0
213.0
168.0

1.0
1.0
1.0
1.0
1.0
1.0
1.0
1.0
1.0
1.0
1.0
1.0
1.0
1.0
1.0
1.0
1.0
1.0
1.0
1.0
1.0
1.0
1.0
1.0
1.0
1.0
1.0
1.0
1.0
1.0


120.0
0.0
0.0
0.0
0.0
0.0
0.0
0.0
0.0
0.0
0.0
0.0
0.0
0.0
0.0
0.0
0.0
0.0
0.0
0.0
0.0
0.0
0.0
0.0
0.0
97.0
0.0
0.0
0.0
23.0

0.77
0.77
0.77
0.77
0.77
0.77
0.77
0.77
0.77
0.77
0.77
0.77
0.77
0.77
0.77
0.77
0.77
0.77
0.77
0.77
0.77
0.77
0.77
0.77
0.77
0.77
0.77
0.77
0.77
0.77


214.0
0.0
0.0
0.0
0.0
0.0
0.0
0.0
0.0
0.0
0.0
0.0
0.0
0.0
0.0
0.0
0.0
0.0
0.0
0.0
0.0
0.0
0.0
0.0
0.0
0.0
29.0
11.0
0.0
174.0

1.0
1.0
1.0
1.0
1.0
1.0
1.0
1.0
1.0
1.0
1.0
1.0
1.0
1.0
1.0
1.0
1.0
1.0
1.0
1.0
1.0
1.0
1.0
1.0
1.0
1.0
1.0
1.0
1.0
1.0


715.0
0.0
0.0
0.0
381.0
108.0
151.0
0.0
0.0
0.0
0.0
0.0
0.0
0.0
0.0
0.0
0.0
0.0
0.0
0.0
0.0
0.0
0.0
0.0
0.0
0.0
0.0
75.0
0.0
0.0


715.0
0.0
0.0
0.0
381.0
108.0
151.0
0.0
0.0
0.0
0.0
0.0
0.0
0.0
0.0
0.0
0.0
0.0
0.0
0.0
0.0
0.0
0.0
0.0
0.0
0.0
0.0
75.0
0.0
0.0


715.0
0.0
0.0
0.0
381.0
108.0
151.0
0.0
0.0
0.0
0.0
0.0
0.0
0.0
0.0
0.0
0.0
0.0
0.0
0.0
0.0
0.0
0.0
0.0
0.0
0.0
0.0
75.0
0.0
0.0

1.0
1.0
1.0
1.0
1.0
1.0
1.0
1.0
1.0
1.0
1.0
1.0
1.0
1.0
1.0
1.0
1.0
1.0
1.0
1.0
1.0
1.0
1.0
1.0
1.0
1.0
1.0
1.0
1.0
1.0


165.0
0.0
0.0
0.0
26.0
105.0
0.0
0.0
0.0
23.0
0.0
11.0
0.0
0.0
0.0
0.0
0.0
0.0
0.0
0.0
0.0
0.0
0.0
0.0
0.0
0.0
0.0
0.0
0.0
0.0


131.0
0.0
0.0
0.0
26.0
105.0
0.0
0.0
0.0
0.0
0.0
0.0
0.0
0.0
0.0
0.0
0.0
0.0
0.0
0.0
0.0
0.0
0.0
0.0
0.0
0.0
0.0
0.0
0.0
0.0


131.0
0.0
0.0
0.0
26.0
105.0
0.0
0.0
0.0
0.0
0.0
0.0
0.0
0.0
0.0
0.0
0.0
0.0
0.0
0.0
0.0
0.0
0.0
0.0
0.0
0.0
0.0
0.0
0.0
0.0

1.0
1.0
1.0
1.0
1.0
1.0
1.0
1.0
1.0
1.0
1.0
1.0
1.0
1.0
1.0
1.0
1.0
1.0
1.0
1.0
1.0
1.0
1.0
1.0
1.0
1.0
1.0
1.0
1.0
1.0


34.0
0.0
0.0
0.0
0.0
0.0
0.0
0.0
0.0
23.0
0.0
11.0
0.0
0.0
0.0
0.0
0.0
0.0
0.0
0.0
0.0
0.0
0.0
0.0
0.0
0.0
0.0
0.0
0.0
0.0


20.0
0.0
0.0
0.0
0.0
0.0
0.0
0.0
0.0
19.0
0.0
1.0
0.0
0.0
0.0
0.0
0.0
0.0
0.0
0.0
0.0
0.0
0.0
0.0
0.0
0.0
0.0
0.0
0.0
0.0

1.0
1.0
1.0
1.0
1.0
1.0
1.0
1.0
1.0
1.0
1.0
1.0
1.0
1.0
1.0
1.0
1.0
1.0
1.0
1.0
1.0
1.0
1.0
1.0
1.0
1.0
1.0
1.0
1.0
1.0


14.0
0.0
0.0
0.0
0.0
0.0
0.0
0.0
0.0
4.0
0.0
10.0
0.0
0.0
0.0
0.0
0.0
0.0
0.0
0.0
0.0
0.0
0.0
0.0
0.0
0.0
0.0
0.0
0.0
0.0

1.0
1.0
1.0
1.0
1.0
1.0
1.0
1.0
1.0
1.0
1.0
1.0
1.0
1.0
1.0
1.0
1.0
1.0
1.0
1.0
1.0
1.0
1.0
1.0
1.0
1.0
1.0
1.0
1.0
1.0


103.0
21.0
0.0
0.0
0.0
0.0
0.0
0.0
0.0
82.0
0.0
0.0
0.0
0.0
0.0
0.0
0.0
0.0
0.0
0.0
0.0
0.0
0.0
0.0
0.0
0.0
0.0
0.0
0.0
0.0


103.0
21.0
0.0
0.0
0.0
0.0
0.0
0.0
0.0
82.0
0.0
0.0
0.0
0.0
0.0
0.0
0.0
0.0
0.0
0.0
0.0
0.0
0.0
0.0
0.0
0.0
0.0
0.0
0.0
0.0


103.0
21.0
0.0
0.0
0.0
0.0
0.0
0.0
0.0
82.0
0.0
0.0
0.0
0.0
0.0
0.0
0.0
0.0
0.0
0.0
0.0
0.0
0.0
0.0
0.0
0.0
0.0
0.0
0.0
0.0

1.0
1.0
1.0
1.0
1.0
1.0
1.0
1.0
1.0
1.0
1.0
1.0
1.0
1.0
1.0
1.0
1.0
1.0
1.0
1.0
1.0
1.0
1.0
1.0
1.0
1.0
1.0
1.0
1.0
1.0


6105.0
9.0
1.0
2.0
11.0
0.0
0.0
9.0
10.0
114.0
43.0
49.0
0.0
23.0
65.0
0.0
0.0
14.0
98.0
7.0
0.0
1392.0
0.0
0.0
2601.0
121.0
8.0
4.0
987.0
537.0

1.0
1.0
1.0
1.0
1.0
1.0
1.0
1.0
1.0
1.0
1.0
1.0
1.0
1.0
1.0
1.0
1.0
1.0
1.0
1.0
1.0
1.0
1.0
1.0
1.0
1.0
1.0
1.0
1.0
1.0


97.0
0.0
0.0
0.0
0.0
0.0
0.0
0.0
0.0
88.0
8.0
1.0
0.0
0.0
0.0
0.0
0.0
0.0
0.0
0.0
0.0
0.0
0.0
0.0
0.0
0.0
0.0
0.0
0.0
0.0


97.0
0.0
0.0
0.0
0.0
0.0
0.0
0.0
0.0
88.0
8.0
1.0
0.0
0.0
0.0
0.0
0.0
0.0
0.0
0.0
0.0
0.0
0.0
0.0
0.0
0.0
0.0
0.0
0.0
0.0

1.0
1.0
1.0
1.0
1.0
1.0
1.0
1.0
1.0
1.0
1.0
1.0
1.0
1.0
1.0
1.0
1.0
1.0
1.0
1.0
1.0
1.0
1.0
1.0
1.0
1.0
1.0
1.0
1.0
1.0


375.0
1.0
0.0
0.0
0.0
0.0
0.0
0.0
0.0
0.0
12.0
15.0
0.0
0.0
28.0
0.0
0.0
0.0
0.0
0.0
0.0
0.0
0.0
0.0
319.0
0.0
0.0
0.0
0.0
0.0


375.0
1.0
0.0
0.0
0.0
0.0
0.0
0.0
0.0
0.0
12.0
15.0
0.0
0.0
28.0
0.0
0.0
0.0
0.0
0.0
0.0
0.0
0.0
0.0
319.0
0.0
0.0
0.0
0.0
0.0

1.0
1.0
1.0
1.0
1.0
1.0
1.0
1.0
1.0
1.0
1.0
1.0
1.0
1.0
1.0
1.0
1.0
1.0
1.0
1.0
1.0
1.0
1.0
1.0
1.0
1.0
1.0
1.0
1.0
1.0


35.0
1.0
0.0
1.0
0.0
0.0
0.0
0.0
0.0
0.0
14.0
19.0
0.0
0.0
0.0
0.0
0.0
0.0
0.0
0.0
0.0
0.0
0.0
0.0
0.0
0.0
0.0
0.0
0.0
0.0


35.0
1.0
0.0
1.0
0.0
0.0
0.0
0.0
0.0
0.0
14.0
19.0
0.0
0.0
0.0
0.0
0.0
0.0
0.0
0.0
0.0
0.0
0.0
0.0
0.0
0.0
0.0
0.0
0.0
0.0

0.99
0.99
0.99
0.99
0.99
0.99
0.99
0.99
0.99
0.99
0.99
0.99
0.99
0.99
0.99
0.99
0.99
0.99
0.99
0.99
0.99
0.99
0.99
0.99
0.99
0.99
0.99
0.99
0.99
0.99


249.0
2.0
0.0
0.0
0.0
0.0
0.0
9.0
8.0
0.0
1.0
1.0
0.0
0.0
0.0
0.0
0.0
0.0
23.0
0.0
0.0
0.0
0.0
0.0
205.0
0.0
0.0
0.0
0.0
0.0


226.0
2.0
0.0
0.0
0.0
0.0
0.0
9.0
8.0
0.0
1.0
1.0
0.0
0.0
0.0
0.0
0.0
0.0
0.0
0.0
0.0
0.0
0.0
0.0
205.0
0.0
0.0
0.0
0.0
0.0

1.0
1.0
1.0
1.0
1.0
1.0
1.0
1.0
1.0
1.0
1.0
1.0
1.0
1.0
1.0
1.0
1.0
1.0
1.0
1.0
1.0
1.0
1.0
1.0
1.0
1.0
1.0
1.0
1.0
1.0


23.0
0.0
0.0
0.0
0.0
0.0
0.0
0.0
0.0
0.0
0.0
0.0
0.0
0.0
0.0
0.0
0.0
0.0
23.0
0.0
0.0
0.0
0.0
0.0
0.0
0.0
0.0
0.0
0.0
0.0

1.0
1.0
1.0
1.0
1.0
1.0
1.0
1.0
1.0
1.0
1.0
1.0
1.0
1.0
1.0
1.0
1.0
1.0
1.0
1.0
1.0
1.0
1.0
1.0
1.0
1.0
1.0
1.0
1.0
1.0


16.0
2.0
1.0
0.0
10.0
0.0
0.0
0.0
0.0
0.0
0.0
3.0
0.0
0.0
0.0
0.0
0.0
0.0
0.0
0.0
0.0
0.0
0.0
0.0
0.0
0.0
0.0
0.0
0.0
0.0


16.0
2.0
1.0
0.0
10.0
0.0
0.0
0.0
0.0
0.0
0.0
3.0
0.0
0.0
0.0
0.0
0.0
0.0
0.0
0.0
0.0
0.0
0.0
0.0
0.0
0.0
0.0
0.0
0.0
0.0

1.0
1.0
1.0
1.0
1.0
1.0
1.0
1.0
1.0
1.0
1.0
1.0
1.0
1.0
1.0
1.0
1.0
1.0
1.0
1.0
1.0
1.0
1.0
1.0
1.0
1.0
1.0
1.0
1.0
1.0


663.0
1.0
0.0
1.0
1.0
0.0
0.0
0.0
0.0
9.0
3.0
1.0
0.0
0.0
0.0
0.0
0.0
10.0
18.0
0.0
0.0
0.0
0.0
0.0
442.0
0.0
0.0
0.0
72.0
105.0

1.0
1.0
1.0
1.0
1.0
1.0
1.0
1.0
1.0
1.0
1.0
1.0
1.0
1.0
1.0
1.0
1.0
1.0
1.0
1.0
1.0
1.0
1.0
1.0
1.0
1.0
1.0
1.0
1.0
1.0


28.0
0.0
0.0
0.0
0.0
0.0
0.0
0.0
0.0
0.0
0.0
0.0
0.0
0.0
0.0
0.0
0.0
0.0
0.0
0.0
0.0
0.0
0.0
0.0
16.0
0.0
0.0
0.0
0.0
12.0

0.95
0.95
0.95
0.95
0.95
0.95
0.95
0.95
0.95
0.95
0.95
0.95
0.95
0.95
0.95
0.95
0.95
0.95
0.95
0.95
0.95
0.95
0.95
0.95
0.95
0.95
0.95
0.95
0.95
0.95


10.0
0.0
0.0
0.0
0.0
0.0
0.0
0.0
0.0
0.0
0.0
0.0
0.0
0.0
0.0
0.0
0.0
0.0
0.0
0.0
0.0
0.0
0.0
0.0
0.0
0.0
0.0
0.0
10.0
0.0

1.0
1.0
1.0
1.0
1.0
1.0
1.0
1.0
1.0
1.0
1.0
1.0
1.0
1.0
1.0
1.0
1.0
1.0
1.0
1.0
1.0
1.0
1.0
1.0
1.0
1.0
1.0
1.0
1.0
1.0


35.0
2.0
0.0
0.0
0.0
0.0
0.0
0.0
0.0
9.0
4.0
9.0
0.0
0.0
0.0
0.0
0.0
0.0
0.0
0.0
0.0
0.0
0.0
0.0
11.0
0.0
0.0
0.0
0.0
0.0

1.0
1.0
1.0
1.0
1.0
1.0
1.0
1.0
1.0
1.0
1.0
1.0
1.0
1.0
1.0
1.0
1.0
1.0
1.0
1.0
1.0
1.0
1.0
1.0
1.0
1.0
1.0
1.0
1.0
1.0


15.0
0.0
0.0
0.0
0.0
0.0
0.0
0.0
0.0
9.0
4.0
2.0
0.0
0.0
0.0
0.0
0.0
0.0
0.0
0.0
0.0
0.0
0.0
0.0
0.0
0.0
0.0
0.0
0.0
0.0

1.0
1.0
1.0
1.0
1.0
1.0
1.0
1.0
1.0
1.0
1.0
1.0
1.0
1.0
1.0
1.0
1.0
1.0
1.0
1.0
1.0
1.0
1.0
1.0
1.0
1.0
1.0
1.0
1.0
1.0


9.0
2.0
0.0
0.0
0.0
0.0
0.0
0.0
0.0
0.0
0.0
7.0
0.0
0.0
0.0
0.0
0.0
0.0
0.0
0.0
0.0
0.0
0.0
0.0
0.0
0.0
0.0
0.0
0.0
0.0

1.0
1.0
1.0
1.0
1.0
1.0
1.0
1.0
1.0
1.0
1.0
1.0
1.0
1.0
1.0
1.0
1.0
1.0
1.0
1.0
1.0
1.0
1.0
1.0
1.0
1.0
1.0
1.0
1.0
1.0


11.0
0.0
0.0
0.0
0.0
0.0
0.0
0.0
2.0
8.0
1.0
0.0
0.0
0.0
0.0
0.0
0.0
0.0
0.0
0.0
0.0
0.0
0.0
0.0
0.0
0.0
0.0
0.0
0.0
0.0


11.0
0.0
0.0
0.0
0.0
0.0
0.0
0.0
2.0
8.0
1.0
0.0
0.0
0.0
0.0
0.0
0.0
0.0
0.0
0.0
0.0
0.0
0.0
0.0
0.0
0.0
0.0
0.0
0.0
0.0

1.0
1.0
1.0
1.0
1.0
1.0
1.0
1.0
1.0
1.0
1.0
1.0
1.0
1.0
1.0
1.0
1.0
1.0
1.0
1.0
1.0
1.0
1.0
1.0
1.0
1.0
1.0
1.0
1.0
1.0


1459.0
0.0
0.0
0.0
0.0
0.0
0.0
0.0
0.0
0.0
0.0
0.0
0.0
0.0
0.0
0.0
0.0
0.0
0.0
0.0
0.0
0.0
0.0
0.0
1422.0
0.0
0.0
0.0
15.0
22.0


1435.0
0.0
0.0
0.0
0.0
0.0
0.0
0.0
0.0
0.0
0.0
0.0
0.0
0.0
0.0
0.0
0.0
0.0
0.0
0.0
0.0
0.0
0.0
0.0
1422.0
0.0
0.0
0.0
0.0
13.0

0.92
0.92
0.92
0.92
0.92
0.92
0.92
0.92
0.92
0.92
0.92
0.92
0.92
0.92
0.92
0.92
0.92
0.92
0.92
0.92
0.92
0.92
0.92
0.92
0.92
0.92
0.92
0.92
0.92
0.92


24.0
0.0
0.0
0.0
0.0
0.0
0.0
0.0
0.0
0.0
0.0
0.0
0.0
0.0
0.0
0.0
0.0
0.0
0.0
0.0
0.0
0.0
0.0
0.0
0.0
0.0
0.0
0.0
15.0
9.0

1.0
1.0
1.0
1.0
1.0
1.0
1.0
1.0
1.0
1.0
1.0
1.0
1.0
1.0
1.0
1.0
1.0
1.0
1.0
1.0
1.0
1.0
1.0
1.0
1.0
1.0
1.0
1.0
1.0
1.0


1392.0
0.0
0.0
0.0
0.0
0.0
0.0
0.0
0.0
0.0
0.0
0.0
0.0
0.0
0.0
0.0
0.0
0.0
0.0
0.0
0.0
1392.0
0.0
0.0
0.0
0.0
0.0
0.0
0.0
0.0


1392.0
0.0
0.0
0.0
0.0
0.0
0.0
0.0
0.0
0.0
0.0
0.0
0.0
0.0
0.0
0.0
0.0
0.0
0.0
0.0
0.0
1392.0
0.0
0.0
0.0
0.0
0.0
0.0
0.0
0.0

1.0
1.0
1.0
1.0
1.0
1.0
1.0
1.0
1.0
1.0
1.0
1.0
1.0
1.0
1.0
1.0
1.0
1.0
1.0
1.0
1.0
1.0
1.0
1.0
1.0
1.0
1.0
1.0
1.0
1.0


1310.0
0.0
0.0
0.0
0.0
0.0
0.0
0.0
0.0
0.0
0.0
0.0
0.0
0.0
0.0
0.0
0.0
0.0
0.0
0.0
0.0
0.0
0.0
0.0
0.0
0.0
0.0
0.0
900.0
410.0


1310.0
0.0
0.0
0.0
0.0
0.0
0.0
0.0
0.0
0.0
0.0
0.0
0.0
0.0
0.0
0.0
0.0
0.0
0.0
0.0
0.0
0.0
0.0
0.0
0.0
0.0
0.0
0.0
900.0
410.0

1.0
1.0
1.0
1.0
1.0
1.0
1.0
1.0
1.0
1.0
1.0
1.0
1.0
1.0
1.0
1.0
1.0
1.0
1.0
1.0
1.0
1.0
1.0
1.0
1.0
1.0
1.0
1.0
1.0
1.0


121.0
0.0
0.0
0.0
0.0
0.0
0.0
0.0
0.0
0.0
0.0
0.0
0.0
0.0
0.0
0.0
0.0
0.0
0.0
0.0
0.0
0.0
0.0
0.0
0.0
121.0
0.0
0.0
0.0
0.0

1.0
1.0
1.0
1.0
1.0
1.0
1.0
1.0
1.0
1.0
1.0
1.0
1.0
1.0
1.0
1.0
1.0
1.0
1.0
1.0
1.0
1.0
1.0
1.0
1.0
1.0
1.0
1.0
1.0
1.0


75.0
0.0
0.0
0.0
0.0
0.0
0.0
0.0
0.0
0.0
0.0
0.0
0.0
0.0
0.0
0.0
0.0
0.0
0.0
0.0
0.0
0.0
0.0
0.0
75.0
0.0
0.0
0.0
0.0
0.0


75.0
0.0
0.0
0.0
0.0
0.0
0.0
0.0
0.0
0.0
0.0
0.0
0.0
0.0
0.0
0.0
0.0
0.0
0.0
0.0
0.0
0.0
0.0
0.0
75.0
0.0
0.0
0.0
0.0
0.0

1.0
1.0
1.0
1.0
1.0
1.0
1.0
1.0
1.0
1.0
1.0
1.0
1.0
1.0
1.0
1.0
1.0
1.0
1.0
1.0
1.0
1.0
1.0
1.0
1.0
1.0
1.0
1.0
1.0
1.0


49.0
0.0
0.0
0.0
0.0
0.0
0.0
0.0
0.0
0.0
0.0
0.0
0.0
0.0
0.0
0.0
0.0
0.0
49.0
0.0
0.0
0.0
0.0
0.0
0.0
0.0
0.0
0.0
0.0
0.0


49.0
0.0
0.0
0.0
0.0
0.0
0.0
0.0
0.0
0.0
0.0
0.0
0.0
0.0
0.0
0.0
0.0
0.0
49.0
0.0
0.0
0.0
0.0
0.0
0.0
0.0
0.0
0.0
0.0
0.0

1.0
1.0
1.0
1.0
1.0
1.0
1.0
1.0
1.0
1.0
1.0
1.0
1.0
1.0
1.0
1.0
1.0
1.0
1.0
1.0
1.0
1.0
1.0
1.0
1.0
1.0
1.0
1.0
1.0
1.0


47.0
0.0
0.0
0.0
0.0
0.0
0.0
0.0
0.0
0.0
0.0
0.0
0.0
0.0
0.0
0.0
0.0
0.0
0.0
0.0
0.0
0.0
0.0
0.0
47.0
0.0
0.0
0.0
0.0
0.0


47.0
0.0
0.0
0.0
0.0
0.0
0.0
0.0
0.0
0.0
0.0
0.0
0.0
0.0
0.0
0.0
0.0
0.0
0.0
0.0
0.0
0.0
0.0
0.0
47.0
0.0
0.0
0.0
0.0
0.0

1.0
1.0
1.0
1.0
1.0
1.0
1.0
1.0
1.0
1.0
1.0
1.0
1.0
1.0
1.0
1.0
1.0
1.0
1.0
1.0
1.0
1.0
1.0
1.0
1.0
1.0
1.0
1.0
1.0
1.0


37.0
0.0
0.0
0.0
0.0
0.0
0.0
0.0
0.0
0.0
0.0
0.0
0.0
0.0
37.0
0.0
0.0
0.0
0.0
0.0
0.0
0.0
0.0
0.0
0.0
0.0
0.0
0.0
0.0
0.0


37.0
0.0
0.0
0.0
0.0
0.0
0.0
0.0
0.0
0.0
0.0
0.0
0.0
0.0
37.0
0.0
0.0
0.0
0.0
0.0
0.0
0.0
0.0
0.0
0.0
0.0
0.0
0.0
0.0
0.0

0.85
0.85
0.85
0.85
0.85
0.85
0.85
0.85
0.85
0.85
0.85
0.85
0.85
0.85
0.85
0.85
0.85
0.85
0.85
0.85
0.85
0.85
0.85
0.85
0.85
0.85
0.85
0.85
0.85
0.85


18.0
0.0
0.0
0.0
0.0
0.0
0.0
0.0
0.0
0.0
0.0
0.0
0.0
18.0
0.0
0.0
0.0
0.0
0.0
0.0
0.0
0.0
0.0
0.0
0.0
0.0
0.0
0.0
0.0
0.0

1.0
1.0
1.0
1.0
1.0
1.0
1.0
1.0
1.0
1.0
1.0
1.0
1.0
1.0
1.0
1.0
1.0
1.0
1.0
1.0
1.0
1.0
1.0
1.0
1.0
1.0
1.0
1.0
1.0
1.0


5.0
0.0
0.0
0.0
0.0
0.0
0.0
0.0
0.0
0.0
0.0
0.0
0.0
5.0
0.0
0.0
0.0
0.0
0.0
0.0
0.0
0.0
0.0
0.0
0.0
0.0
0.0
0.0
0.0
0.0

1.0
1.0
1.0
1.0
1.0
1.0
1.0
1.0
1.0
1.0
1.0
1.0
1.0
1.0
1.0
1.0
1.0
1.0
1.0
1.0
1.0
1.0
1.0
1.0
1.0
1.0
1.0
1.0
1.0
1.0


4.0
0.0
0.0
0.0
0.0
0.0
0.0
0.0
0.0
0.0
0.0
0.0
0.0
0.0
0.0
0.0
0.0
0.0
0.0
0.0
0.0
0.0
0.0
0.0
0.0
0.0
0.0
4.0
0.0
0.0


4.0
0.0
0.0
0.0
0.0
0.0
0.0
0.0
0.0
0.0
0.0
0.0
0.0
0.0
0.0
0.0
0.0
0.0
0.0
0.0
0.0
0.0
0.0
0.0
0.0
0.0
0.0
4.0
0.0
0.0

0.81
0.81
0.81
0.81
0.81
0.81
0.81
0.81
0.81
0.81
0.81
0.81
0.81
0.81
0.81
0.81
0.81
0.81
0.81
0.81
0.81
0.81
0.81
0.81
0.81
0.81
0.81
0.81
0.81
0.81


66.0
0.0
0.0
0.0
0.0
0.0
66.0
0.0
0.0
0.0
0.0
0.0
0.0
0.0
0.0
0.0
0.0
0.0
0.0
0.0
0.0
0.0
0.0
0.0
0.0
0.0
0.0
0.0
0.0
0.0


66.0
0.0
0.0
0.0
0.0
0.0
66.0
0.0
0.0
0.0
0.0
0.0
0.0
0.0
0.0
0.0
0.0
0.0
0.0
0.0
0.0
0.0
0.0
0.0
0.0
0.0
0.0
0.0
0.0
0.0


66.0
0.0
0.0
0.0
0.0
0.0
66.0
0.0
0.0
0.0
0.0
0.0
0.0
0.0
0.0
0.0
0.0
0.0
0.0
0.0
0.0
0.0
0.0
0.0
0.0
0.0
0.0
0.0
0.0
0.0

1.0
1.0
1.0
1.0
1.0
1.0
1.0
1.0
1.0
1.0
1.0
1.0
1.0
1.0
1.0
1.0
1.0
1.0
1.0
1.0
1.0
1.0
1.0
1.0
1.0
1.0
1.0
1.0
1.0
1.0


8919.0
0.0
65.0
34.0
0.0
0.0
0.0
0.0
0.0
0.0
0.0
0.0
0.0
8806.0
0.0
0.0
0.0
0.0
14.0
0.0
0.0
0.0
0.0
0.0
0.0
0.0
0.0
0.0
0.0
0.0


8885.0
0.0
65.0
0.0
0.0
0.0
0.0
0.0
0.0
0.0
0.0
0.0
0.0
8806.0
0.0
0.0
0.0
0.0
14.0
0.0
0.0
0.0
0.0
0.0
0.0
0.0
0.0
0.0
0.0
0.0

1.0
1.0
1.0
1.0
1.0
1.0
1.0
1.0
1.0
1.0
1.0
1.0
1.0
1.0
1.0
1.0
1.0
1.0
1.0
1.0
1.0
1.0
1.0
1.0
1.0
1.0
1.0
1.0
1.0
1.0


104.0
0.0
65.0
0.0
0.0
0.0
0.0
0.0
0.0
0.0
0.0
0.0
0.0
25.0
0.0
0.0
0.0
0.0
14.0
0.0
0.0
0.0
0.0
0.0
0.0
0.0
0.0
0.0
0.0
0.0

0.87
0.87
0.87
0.87
0.87
0.87
0.87
0.87
0.87
0.87
0.87
0.87
0.87
0.87
0.87
0.87
0.87
0.87
0.87
0.87
0.87
0.87
0.87
0.87
0.87
0.87
0.87
0.87
0.87
0.87


8579.0
0.0
0.0
0.0
0.0
0.0
0.0
0.0
0.0
0.0
0.0
0.0
0.0
8579.0
0.0
0.0
0.0
0.0
0.0
0.0
0.0
0.0
0.0
0.0
0.0
0.0
0.0
0.0
0.0
0.0

1.0
1.0
1.0
1.0
1.0
1.0
1.0
1.0
1.0
1.0
1.0
1.0
1.0
1.0
1.0
1.0
1.0
1.0
1.0
1.0
1.0
1.0
1.0
1.0
1.0
1.0
1.0
1.0
1.0
1.0


84.0
0.0
0.0
0.0
0.0
0.0
0.0
0.0
0.0
0.0
0.0
0.0
0.0
84.0
0.0
0.0
0.0
0.0
0.0
0.0
0.0
0.0
0.0
0.0
0.0
0.0
0.0
0.0
0.0
0.0

1.0
1.0
1.0
1.0
1.0
1.0
1.0
1.0
1.0
1.0
1.0
1.0
1.0
1.0
1.0
1.0
1.0
1.0
1.0
1.0
1.0
1.0
1.0
1.0
1.0
1.0
1.0
1.0
1.0
1.0


12.0
0.0
0.0
0.0
0.0
0.0
0.0
0.0
0.0
0.0
0.0
0.0
0.0
12.0
0.0
0.0
0.0
0.0
0.0
0.0
0.0
0.0
0.0
0.0
0.0
0.0
0.0
0.0
0.0
0.0

0.71
0.71
0.71
0.71
0.71
0.71
0.71
0.71
0.71
0.71
0.71
0.71
0.71
0.71
0.71
0.71
0.71
0.71
0.71
0.71
0.71
0.71
0.71
0.71
0.71
0.71
0.71
0.71
0.71
0.71


34.0
0.0
0.0
34.0
0.0
0.0
0.0
0.0
0.0
0.0
0.0
0.0
0.0
0.0
0.0
0.0
0.0
0.0
0.0
0.0
0.0
0.0
0.0
0.0
0.0
0.0
0.0
0.0
0.0
0.0


34.0
0.0
0.0
34.0
0.0
0.0
0.0
0.0
0.0
0.0
0.0
0.0
0.0
0.0
0.0
0.0
0.0
0.0
0.0
0.0
0.0
0.0
0.0
0.0
0.0
0.0
0.0
0.0
0.0
0.0

1.0
1.0
1.0
1.0
1.0
1.0
1.0
1.0
1.0
1.0
1.0
1.0
1.0
1.0
1.0
1.0
1.0
1.0
1.0
1.0
1.0
1.0
1.0
1.0
1.0
1.0
1.0
1.0
1.0
1.0


32771.0
0.0
0.0
0.0
0.0
0.0
52.0
0.0
0.0
0.0
0.0
0.0
0.0
0.0
0.0
0.0
0.0
0.0
0.0
0.0
0.0
0.0
0.0
0.0
0.0
24.0
0.0
0.0
0.0
32695.0


52.0
0.0
0.0
0.0
0.0
0.0
52.0
0.0
0.0
0.0
0.0
0.0
0.0
0.0
0.0
0.0
0.0
0.0
0.0
0.0
0.0
0.0
0.0
0.0
0.0
0.0
0.0
0.0
0.0
0.0


52.0
0.0
0.0
0.0
0.0
0.0
52.0
0.0
0.0
0.0
0.0
0.0
0.0
0.0
0.0
0.0
0.0
0.0
0.0
0.0
0.0
0.0
0.0
0.0
0.0
0.0
0.0
0.0
0.0
0.0

1.0
1.0
1.0
1.0
1.0
1.0
1.0
1.0
1.0
1.0
1.0
1.0
1.0
1.0
1.0
1.0
1.0
1.0
1.0
1.0
1.0
1.0
1.0
1.0
1.0
1.0
1.0
1.0
1.0
1.0


32719.0
0.0
0.0
0.0
0.0
0.0
0.0
0.0
0.0
0.0
0.0
0.0
0.0
0.0
0.0
0.0
0.0
0.0
0.0
0.0
0.0
0.0
0.0
0.0
0.0
24.0
0.0
0.0
0.0
32695.0


32719.0
0.0
0.0
0.0
0.0
0.0
0.0
0.0
0.0
0.0
0.0
0.0
0.0
0.0
0.0
0.0
0.0
0.0
0.0
0.0
0.0
0.0
0.0
0.0
0.0
24.0
0.0
0.0
0.0
32695.0

1.0
1.0
1.0
1.0
1.0
1.0
1.0
1.0
1.0
1.0
1.0
1.0
1.0
1.0
1.0
1.0
1.0
1.0
1.0
1.0
1.0
1.0
1.0
1.0
1.0
1.0
1.0
1.0
1.0
1.0


51012.0
0.0
0.0
50.0
0.0
0.0
0.0
0.0
0.0
0.0
0.0
0.0
0.0
6468.0
44494.0
0.0
0.0
0.0
0.0
0.0
0.0
0.0
0.0
0.0
0.0
0.0
0.0
0.0
0.0
0.0

1.0
1.0
1.0
1.0
1.0
1.0
1.0
1.0
1.0
1.0
1.0
1.0
1.0
1.0
1.0
1.0
1.0
1.0
1.0
1.0
1.0
1.0
1.0
1.0
1.0
1.0
1.0
1.0
1.0
1.0


1768.0
0.0
0.0
0.0
0.0
0.0
0.0
0.0
0.0
0.0
0.0
0.0
0.0
142.0
1626.0
0.0
0.0
0.0
0.0
0.0
0.0
0.0
0.0
0.0
0.0
0.0
0.0
0.0
0.0
0.0


1768.0
0.0
0.0
0.0
0.0
0.0
0.0
0.0
0.0
0.0
0.0
0.0
0.0
142.0
1626.0
0.0
0.0
0.0
0.0
0.0
0.0
0.0
0.0
0.0
0.0
0.0
0.0
0.0
0.0
0.0

1.0
1.0
1.0
1.0
1.0
1.0
1.0
1.0
1.0
1.0
1.0
1.0
1.0
1.0
1.0
1.0
1.0
1.0
1.0
1.0
1.0
1.0
1.0
1.0
1.0
1.0
1.0
1.0
1.0
1.0


935.0
45.0
0.0
0.0
0.0
0.0
0.0
0.0
0.0
0.0
0.0
0.0
0.0
0.0
0.0
0.0
0.0
0.0
0.0
0.0
0.0
890.0
0.0
0.0
0.0
0.0
0.0
0.0
0.0
0.0


935.0
45.0
0.0
0.0
0.0
0.0
0.0
0.0
0.0
0.0
0.0
0.0
0.0
0.0
0.0
0.0
0.0
0.0
0.0
0.0
0.0
890.0
0.0
0.0
0.0
0.0
0.0
0.0
0.0
0.0


935.0
45.0
0.0
0.0
0.0
0.0
0.0
0.0
0.0
0.0
0.0
0.0
0.0
0.0
0.0
0.0
0.0
0.0
0.0
0.0
0.0
890.0
0.0
0.0
0.0
0.0
0.0
0.0
0.0
0.0

1.0
1.0
1.0
1.0
1.0
1.0
1.0
1.0
1.0
1.0
1.0
1.0
1.0
1.0
1.0
1.0
1.0
1.0
1.0
1.0
1.0
1.0
1.0
1.0
1.0
1.0
1.0
1.0
1.0
1.0


36.0
0.0
36.0
0.0
0.0
0.0
0.0
0.0
0.0
0.0
0.0
0.0
0.0
0.0
0.0
0.0
0.0
0.0
0.0
0.0
0.0
0.0
0.0
0.0
0.0
0.0
0.0
0.0
0.0
0.0


36.0
0.0
36.0
0.0
0.0
0.0
0.0
0.0
0.0
0.0
0.0
0.0
0.0
0.0
0.0
0.0
0.0
0.0
0.0
0.0
0.0
0.0
0.0
0.0
0.0
0.0
0.0
0.0
0.0
0.0


36.0
0.0
36.0
0.0
0.0
0.0
0.0
0.0
0.0
0.0
0.0
0.0
0.0
0.0
0.0
0.0
0.0
0.0
0.0
0.0
0.0
0.0
0.0
0.0
0.0
0.0
0.0
0.0
0.0
0.0

1.0
1.0
1.0
1.0
1.0
1.0
1.0
1.0
1.0
1.0
1.0
1.0
1.0
1.0
1.0
1.0
1.0
1.0
1.0
1.0
1.0
1.0
1.0
1.0
1.0
1.0
1.0
1.0
1.0
1.0


33.0
0.0
0.0
0.0
0.0
33.0
0.0
0.0
0.0
0.0
0.0
0.0
0.0
0.0
0.0
0.0
0.0
0.0
0.0
0.0
0.0
0.0
0.0
0.0
0.0
0.0
0.0
0.0
0.0
0.0


33.0
0.0
0.0
0.0
0.0
33.0
0.0
0.0
0.0
0.0
0.0
0.0
0.0
0.0
0.0
0.0
0.0
0.0
0.0
0.0
0.0
0.0
0.0
0.0
0.0
0.0
0.0
0.0
0.0
0.0


33.0
0.0
0.0
0.0
0.0
33.0
0.0
0.0
0.0
0.0
0.0
0.0
0.0
0.0
0.0
0.0
0.0
0.0
0.0
0.0
0.0
0.0
0.0
0.0
0.0
0.0
0.0
0.0
0.0
0.0

0.71
0.71
0.71
0.71
0.71
0.71
0.71
0.71
0.71
0.71
0.71
0.71
0.71
0.71
0.71
0.71
0.71
0.71
0.71
0.71
0.71
0.71
0.71
0.71
0.71
0.71
0.71
0.71
0.71
0.71


2679.0
6.0
20.0
0.0
0.0
0.0
0.0
0.0
0.0
0.0
0.0
0.0
0.0
0.0
0.0
0.0
0.0
0.0
0.0
0.0
0.0
0.0
0.0
0.0
0.0
18.0
0.0
316.0
2113.0
206.0

1.0
1.0
1.0
1.0
1.0
1.0
1.0
1.0
1.0
1.0
1.0
1.0
1.0
1.0
1.0
1.0
1.0
1.0
1.0
1.0
1.0
1.0
1.0
1.0
1.0
1.0
1.0
1.0
1.0
1.0


2635.0
0.0
0.0
0.0
0.0
0.0
0.0
0.0
0.0
0.0
0.0
0.0
0.0
0.0
0.0
0.0
0.0
0.0
0.0
0.0
0.0
0.0
0.0
0.0
0.0
0.0
0.0
316.0
2113.0
206.0


2635.0
0.0
0.0
0.0
0.0
0.0
0.0
0.0
0.0
0.0
0.0
0.0
0.0
0.0
0.0
0.0
0.0
0.0
0.0
0.0
0.0
0.0
0.0
0.0
0.0
0.0
0.0
316.0
2113.0
206.0

0.79
0.79
0.79
0.79
0.79
0.79
0.79
0.79
0.79
0.79
0.79
0.79
0.79
0.79
0.79
0.79
0.79
0.79
0.79
0.79
0.79
0.79
0.79
0.79
0.79
0.79
0.79
0.79
0.79
0.79


18.0
0.0
0.0
0.0
0.0
0.0
0.0
0.0
0.0
0.0
0.0
0.0
0.0
0.0
0.0
0.0
0.0
0.0
0.0
0.0
0.0
0.0
0.0
0.0
0.0
18.0
0.0
0.0
0.0
0.0


18.0
0.0
0.0
0.0
0.0
0.0
0.0
0.0
0.0
0.0
0.0
0.0
0.0
0.0
0.0
0.0
0.0
0.0
0.0
0.0
0.0
0.0
0.0
0.0
0.0
18.0
0.0
0.0
0.0
0.0

1.0
1.0
1.0
1.0
1.0
1.0
1.0
1.0
1.0
1.0
1.0
1.0
1.0
1.0
1.0
1.0
1.0
1.0
1.0
1.0
1.0
1.0
1.0
1.0
1.0
1.0
1.0
1.0
1.0
1.0


17.0
0.0
0.0
0.0
0.0
0.0
0.0
0.0
14.0
3.0
0.0
0.0
0.0
0.0
0.0
0.0
0.0
0.0
0.0
0.0
0.0
0.0
0.0
0.0
0.0
0.0
0.0
0.0
0.0
0.0


17.0
0.0
0.0
0.0
0.0
0.0
0.0
0.0
14.0
3.0
0.0
0.0
0.0
0.0
0.0
0.0
0.0
0.0
0.0
0.0
0.0
0.0
0.0
0.0
0.0
0.0
0.0
0.0
0.0
0.0


17.0
0.0
0.0
0.0
0.0
0.0
0.0
0.0
14.0
3.0
0.0
0.0
0.0
0.0
0.0
0.0
0.0
0.0
0.0
0.0
0.0
0.0
0.0
0.0
0.0
0.0
0.0
0.0
0.0
0.0

0.89
0.89
0.89
0.89
0.89
0.89
0.89
0.89
0.89
0.89
0.89
0.89
0.89
0.89
0.89
0.89
0.89
0.89
0.89
0.89
0.89
0.89
0.89
0.89
0.89
0.89
0.89
0.89
0.89
0.89


225646.0
0.0
0.0
0.0
0.0
0.0
0.0
0.0
0.0
0.0
0.0
0.0
0.0
0.0
0.0
0.0
225646.0
0.0
0.0
0.0
0.0
0.0
0.0
0.0
0.0
0.0
0.0
0.0
0.0
0.0


225646.0
0.0
0.0
0.0
0.0
0.0
0.0
0.0
0.0
0.0
0.0
0.0
0.0
0.0
0.0
0.0
225646.0
0.0
0.0
0.0
0.0
0.0
0.0
0.0
0.0
0.0
0.0
0.0
0.0
0.0


225646.0
0.0
0.0
0.0
0.0
0.0
0.0
0.0
0.0
0.0
0.0
0.0
0.0
0.0
0.0
0.0
225646.0
0.0
0.0
0.0
0.0
0.0
0.0
0.0
0.0
0.0
0.0
0.0
0.0
0.0

0.96
0.96
0.96
0.96
0.96
0.96
0.96
0.96
0.96
0.96
0.96
0.96
0.96
0.96
0.96
0.96
0.96
0.96
0.96
0.96
0.96
0.96
0.96
0.96
0.96
0.96
0.96
0.96
0.96
0.96


4711.0
0.0
0.0
0.0
0.0
0.0
0.0
0.0
0.0
0.0
0.0
0.0
0.0
0.0
0.0
0.0
0.0
0.0
0.0
0.0
0.0
0.0
0.0
0.0
0.0
2544.0
442.0
8.0
1445.0
272.0


4711.0
0.0
0.0
0.0
0.0
0.0
0.0
0.0
0.0
0.0
0.0
0.0
0.0
0.0
0.0
0.0
0.0
0.0
0.0
0.0
0.0
0.0
0.0
0.0
0.0
2544.0
442.0
8.0
1445.0
272.0


4711.0
0.0
0.0
0.0
0.0
0.0
0.0
0.0
0.0
0.0
0.0
0.0
0.0
0.0
0.0
0.0
0.0
0.0
0.0
0.0
0.0
0.0
0.0
0.0
0.0
2544.0
442.0
8.0
1445.0
272.0

1.0
1.0
1.0
1.0
1.0
1.0
1.0
1.0
1.0
1.0
1.0
1.0
1.0
1.0
1.0
1.0
1.0
1.0
1.0
1.0
1.0
1.0
1.0
1.0
1.0
1.0
1.0
1.0
1.0
1.0


33.0
0.0
0.0
0.0
0.0
0.0
0.0
0.0
0.0
0.0
0.0
0.0
0.0
0.0
0.0
0.0
0.0
0.0
0.0
0.0
0.0
33.0
0.0
0.0
0.0
0.0
0.0
0.0
0.0
0.0


33.0
0.0
0.0
0.0
0.0
0.0
0.0
0.0
0.0
0.0
0.0
0.0
0.0
0.0
0.0
0.0
0.0
0.0
0.0
0.0
0.0
33.0
0.0
0.0
0.0
0.0
0.0
0.0
0.0
0.0


33.0
0.0
0.0
0.0
0.0
0.0
0.0
0.0
0.0
0.0
0.0
0.0
0.0
0.0
0.0
0.0
0.0
0.0
0.0
0.0
0.0
33.0
0.0
0.0
0.0
0.0
0.0
0.0
0.0
0.0

1.0
1.0
1.0
1.0
1.0
1.0
1.0
1.0
1.0
1.0
1.0
1.0
1.0
1.0
1.0
1.0
1.0
1.0
1.0
1.0
1.0
1.0
1.0
1.0
1.0
1.0
1.0
1.0
1.0
1.0


7.0
0.0
0.0
0.0
0.0
0.0
0.0
0.0
0.0
0.0
0.0
0.0
0.0
0.0
5.0
0.0
0.0
0.0
0.0
0.0
0.0
0.0
0.0
0.0
2.0
0.0
0.0
0.0
0.0
0.0


7.0
0.0
0.0
0.0
0.0
0.0
0.0
0.0
0.0
0.0
0.0
0.0
0.0
0.0
5.0
0.0
0.0
0.0
0.0
0.0
0.0
0.0
0.0
0.0
2.0
0.0
0.0
0.0
0.0
0.0


7.0
0.0
0.0
0.0
0.0
0.0
0.0
0.0
0.0
0.0
0.0
0.0
0.0
0.0
5.0
0.0
0.0
0.0
0.0
0.0
0.0
0.0
0.0
0.0
2.0
0.0
0.0
0.0
0.0
0.0

0.8
0.8
0.8
0.8
0.8
0.8
0.8
0.8
0.8
0.8
0.8
0.8
0.8
0.8
0.8
0.8
0.8
0.8
0.8
0.8
0.8
0.8
0.8
0.8
0.8
0.8
0.8
0.8
0.8
0.8


4.0
0.0
0.0
0.0
0.0
0.0
0.0
0.0
0.0
0.0
0.0
0.0
0.0
0.0
0.0
0.0
0.0
0.0
0.0
0.0
0.0
0.0
0.0
0.0
0.0
0.0
0.0
0.0
0.0
4.0


4.0
0.0
0.0
0.0
0.0
0.0
0.0
0.0
0.0
0.0
0.0
0.0
0.0
0.0
0.0
0.0
0.0
0.0
0.0
0.0
0.0
0.0
0.0
0.0
0.0
0.0
0.0
0.0
0.0
4.0


4.0
0.0
0.0
0.0
0.0
0.0
0.0
0.0
0.0
0.0
0.0
0.0
0.0
0.0
0.0
0.0
0.0
0.0
0.0
0.0
0.0
0.0
0.0
0.0
0.0
0.0
0.0
0.0
0.0
4.0

1.0
1.0
1.0
1.0
1.0
1.0
1.0
1.0
1.0
1.0
1.0
1.0
1.0
1.0
1.0
1.0
1.0
1.0
1.0
1.0
1.0
1.0
1.0
1.0
1.0
1.0
1.0
1.0
1.0
1.0


60437941.0
7673.0
5175.0
3360.0
14452.0
25136.0
10646.0
117348.0
21745.0
39468.0
11857.0
13965.0
3138145.0
5257153.0
598427.0
3927202.0
2423522.0
6018047.0
5844758.0
3756204.0
1569757.0
5860838.0
888336.0
3138145.0
568305.0
4077850.0
3834197.0
3853005.0
2763654.0
2649571.0

1.0
1.0
1.0
1.0
1.0
1.0
1.0
1.0
1.0
1.0
1.0
1.0
1.0
1.0
1.0
1.0
1.0
1.0
1.0
1.0
1.0
1.0
1.0
1.0
1.0
1.0
1.0
1.0
1.0
1.0


6574027.0
1768.0
278.0
919.0
238.0
1827.0
1246.0
51509.0
10903.0
4717.0
636.0
1827.0
34439.0
11141.0
109218.0
1412365.0
569144.0
296112.0
101853.0
139258.0
1.0
166679.0
485517.0
34439.0
0.0
958844.0
769660.0
436391.0
722702.0
250396.0

0.99
0.99
0.99
0.99
0.99
0.99
0.99
0.99
0.99
0.99
0.99
0.99
0.99
0.99
0.99
0.99
0.99
0.99
0.99
0.99
0.99
0.99
0.99
0.99
0.99
0.99
0.99
0.99
0.99
0.99


65902.0
0.0
0.0
0.0
0.0
0.0
0.0
51509.0
9883.0
4510.0
0.0
0.0
0.0
0.0
0.0
0.0
0.0
0.0
0.0
0.0
0.0
0.0
0.0
0.0
0.0
0.0
0.0
0.0
0.0
0.0


65902.0
0.0
0.0
0.0
0.0
0.0
0.0
51509.0
9883.0
4510.0
0.0
0.0
0.0
0.0
0.0
0.0
0.0
0.0
0.0
0.0
0.0
0.0
0.0
0.0
0.0
0.0
0.0
0.0
0.0
0.0

1.0
1.0
1.0
1.0
1.0
1.0
1.0
1.0
1.0
1.0
1.0
1.0
1.0
1.0
1.0
1.0
1.0
1.0
1.0
1.0
1.0
1.0
1.0
1.0
1.0
1.0
1.0
1.0
1.0
1.0


1657743.0
413.0
0.0
625.0
238.0
1692.0
338.0
0.0
0.0
0.0
0.0
983.0
1.0
0.0
0.0
0.0
0.0
0.0
0.0
0.0
0.0
0.0
0.0
1.0
0.0
821524.0
620147.0
210591.0
1190.0
0.0

1.0
1.0
1.0
1.0
1.0
1.0
1.0
1.0
1.0
1.0
1.0
1.0
1.0
1.0
1.0
1.0
1.0
1.0
1.0
1.0
1.0
1.0
1.0
1.0
1.0
1.0
1.0
1.0
1.0
1.0


16259.0
364.0
0.0
625.0
0.0
1692.0
189.0
0.0
0.0
0.0
0.0
983.0
0.0
0.0
0.0
0.0
0.0
0.0
0.0
0.0
0.0
0.0
0.0
0.0
0.0
0.0
6160.0
5056.0
1190.0
0.0

1.0
1.0
1.0
1.0
1.0
1.0
1.0
1.0
1.0
1.0
1.0
1.0
1.0
1.0
1.0
1.0
1.0
1.0
1.0
1.0
1.0
1.0
1.0
1.0
1.0
1.0
1.0
1.0
1.0
1.0


1637479.0
0.0
0.0
0.0
238.0
0.0
149.0
0.0
0.0
0.0
0.0
0.0
1.0
0.0
0.0
0.0
0.0
0.0
0.0
0.0
0.0
0.0
0.0
1.0
0.0
820954.0
610880.0
205256.0
0.0
0.0

1.0
1.0
1.0
1.0
1.0
1.0
1.0
1.0
1.0
1.0
1.0
1.0
1.0
1.0
1.0
1.0
1.0
1.0
1.0
1.0
1.0
1.0
1.0
1.0
1.0
1.0
1.0
1.0
1.0
1.0


49.0
49.0
0.0
0.0
0.0
0.0
0.0
0.0
0.0
0.0
0.0
0.0
0.0
0.0
0.0
0.0
0.0
0.0
0.0
0.0
0.0
0.0
0.0
0.0
0.0
0.0
0.0
0.0
0.0
0.0

1.0
1.0
1.0
1.0
1.0
1.0
1.0
1.0
1.0
1.0
1.0
1.0
1.0
1.0
1.0
1.0
1.0
1.0
1.0
1.0
1.0
1.0
1.0
1.0
1.0
1.0
1.0
1.0
1.0
1.0


126279.0
175.0
0.0
0.0
0.0
0.0
0.0
0.0
1011.0
0.0
518.0
439.0
0.0
0.0
0.0
0.0
0.0
124136.0
0.0
0.0
0.0
0.0
0.0
0.0
0.0
0.0
0.0
0.0
0.0
0.0


1529.0
0.0
0.0
0.0
0.0
0.0
0.0
0.0
1011.0
0.0
518.0
0.0
0.0
0.0
0.0
0.0
0.0
0.0
0.0
0.0
0.0
0.0
0.0
0.0
0.0
0.0
0.0
0.0
0.0
0.0

1.0
1.0
1.0
1.0
1.0
1.0
1.0
1.0
1.0
1.0
1.0
1.0
1.0
1.0
1.0
1.0
1.0
1.0
1.0
1.0
1.0
1.0
1.0
1.0
1.0
1.0
1.0
1.0
1.0
1.0


124750.0
175.0
0.0
0.0
0.0
0.0
0.0
0.0
0.0
0.0
0.0
439.0
0.0
0.0
0.0
0.0
0.0
124136.0
0.0
0.0
0.0
0.0
0.0
0.0
0.0
0.0
0.0
0.0
0.0
0.0

1.0
1.0
1.0
1.0
1.0
1.0
1.0
1.0
1.0
1.0
1.0
1.0
1.0
1.0
1.0
1.0
1.0
1.0
1.0
1.0
1.0
1.0
1.0
1.0
1.0
1.0
1.0
1.0
1.0
1.0


1471980.0
840.0
278.0
162.0
0.0
135.0
0.0
0.0
0.0
0.0
0.0
405.0
0.0
11140.0
26010.0
1125217.0
135831.0
171961.0
0.0
0.0
1.0
0.0
0.0
0.0
0.0
0.0
0.0
0.0
0.0
0.0


1471326.0
265.0
199.0
162.0
0.0
135.0
0.0
0.0
0.0
0.0
0.0
405.0
0.0
11140.0
26010.0
1125217.0
135831.0
171961.0
0.0
0.0
1.0
0.0
0.0
0.0
0.0
0.0
0.0
0.0
0.0
0.0

1.0
1.0
1.0
1.0
1.0
1.0
1.0
1.0
1.0
1.0
1.0
1.0
1.0
1.0
1.0
1.0
1.0
1.0
1.0
1.0
1.0
1.0
1.0
1.0
1.0
1.0
1.0
1.0
1.0
1.0


654.0
575.0
79.0
0.0
0.0
0.0
0.0
0.0
0.0
0.0
0.0
0.0
0.0
0.0
0.0
0.0
0.0
0.0
0.0
0.0
0.0
0.0
0.0
0.0
0.0
0.0
0.0
0.0
0.0
0.0

1.0
1.0
1.0
1.0
1.0
1.0
1.0
1.0
1.0
1.0
1.0
1.0
1.0
1.0
1.0
1.0
1.0
1.0
1.0
1.0
1.0
1.0
1.0
1.0
1.0
1.0
1.0
1.0
1.0
1.0


908.0
0.0
0.0
0.0
0.0
0.0
908.0
0.0
0.0
0.0
0.0
0.0
0.0
0.0
0.0
0.0
0.0
0.0
0.0
0.0
0.0
0.0
0.0
0.0
0.0
0.0
0.0
0.0
0.0
0.0


908.0
0.0
0.0
0.0
0.0
0.0
908.0
0.0
0.0
0.0
0.0
0.0
0.0
0.0
0.0
0.0
0.0
0.0
0.0
0.0
0.0
0.0
0.0
0.0
0.0
0.0
0.0
0.0
0.0
0.0

1.0
1.0
1.0
1.0
1.0
1.0
1.0
1.0
1.0
1.0
1.0
1.0
1.0
1.0
1.0
1.0
1.0
1.0
1.0
1.0
1.0
1.0
1.0
1.0
1.0
1.0
1.0
1.0
1.0
1.0


69210.0
0.0
0.0
0.0
0.0
0.0
0.0
0.0
9.0
207.0
118.0
0.0
34438.0
0.0
0.0
0.0
0.0
0.0
0.0
0.0
0.0
0.0
0.0
34438.0
0.0
0.0
0.0
0.0
0.0
0.0


69210.0
0.0
0.0
0.0
0.0
0.0
0.0
0.0
9.0
207.0
118.0
0.0
34438.0
0.0
0.0
0.0
0.0
0.0
0.0
0.0
0.0
0.0
0.0
34438.0
0.0
0.0
0.0
0.0
0.0
0.0

1.0
1.0
1.0
1.0
1.0
1.0
1.0
1.0
1.0
1.0
1.0
1.0
1.0
1.0
1.0
1.0
1.0
1.0
1.0
1.0
1.0
1.0
1.0
1.0
1.0
1.0
1.0
1.0
1.0
1.0


416772.0
340.0
0.0
0.0
0.0
0.0
0.0
0.0
0.0
0.0
0.0
0.0
0.0
0.0
0.0
0.0
0.0
0.0
0.0
139258.0
0.0
0.0
0.0
0.0
0.0
0.0
40292.0
174723.0
62159.0
0.0

1.0
1.0
1.0
1.0
1.0
1.0
1.0
1.0
1.0
1.0
1.0
1.0
1.0
1.0
1.0
1.0
1.0
1.0
1.0
1.0
1.0
1.0
1.0
1.0
1.0
1.0
1.0
1.0
1.0
1.0


132.0
0.0
0.0
132.0
0.0
0.0
0.0
0.0
0.0
0.0
0.0
0.0
0.0
0.0
0.0
0.0
0.0
0.0
0.0
0.0
0.0
0.0
0.0
0.0
0.0
0.0
0.0
0.0
0.0
0.0


132.0
0.0
0.0
132.0
0.0
0.0
0.0
0.0
0.0
0.0
0.0
0.0
0.0
0.0
0.0
0.0
0.0
0.0
0.0
0.0
0.0
0.0
0.0
0.0
0.0
0.0
0.0
0.0
0.0
0.0

1.0
1.0
1.0
1.0
1.0
1.0
1.0
1.0
1.0
1.0
1.0
1.0
1.0
1.0
1.0
1.0
1.0
1.0
1.0
1.0
1.0
1.0
1.0
1.0
1.0
1.0
1.0
1.0
1.0
1.0


1022133.0
0.0
0.0
0.0
0.0
0.0
0.0
0.0
0.0
0.0
0.0
0.0
0.0
0.0
0.0
0.0
0.0
0.0
0.0
0.0
0.0
166679.0
0.0
0.0
0.0
137314.0
102669.0
0.0
578945.0
36526.0

0.73
0.73
0.73
0.73
0.73
0.73
0.73
0.73
0.73
0.73
0.73
0.73
0.73
0.73
0.73
0.73
0.73
0.73
0.73
0.73
0.73
0.73
0.73
0.73
0.73
0.73
0.73
0.73
0.73
0.73


1022075.0
0.0
0.0
0.0
0.0
0.0
0.0
0.0
0.0
0.0
0.0
0.0
0.0
0.0
0.0
0.0
0.0
0.0
0.0
0.0
0.0
166679.0
0.0
0.0
0.0
137279.0
102646.0
0.0
578945.0
36526.0

1.0
1.0
1.0
1.0
1.0
1.0
1.0
1.0
1.0
1.0
1.0
1.0
1.0
1.0
1.0
1.0
1.0
1.0
1.0
1.0
1.0
1.0
1.0
1.0
1.0
1.0
1.0
1.0
1.0
1.0


941537.0
0.0
0.0
0.0
0.0
0.0
0.0
0.0
0.0
0.0
0.0
0.0
0.0
0.0
0.0
281838.0
428824.0
1.0
0.0
0.0
0.0
0.0
0.0
0.0
0.0
1.0
0.0
0.0
68157.0
162716.0


941537.0
0.0
0.0
0.0
0.0
0.0
0.0
0.0
0.0
0.0
0.0
0.0
0.0
0.0
0.0
281838.0
428824.0
1.0
0.0
0.0
0.0
0.0
0.0
0.0
0.0
1.0
0.0
0.0
68157.0
162716.0

1.0
1.0
1.0
1.0
1.0
1.0
1.0
1.0
1.0
1.0
1.0
1.0
1.0
1.0
1.0
1.0
1.0
1.0
1.0
1.0
1.0
1.0
1.0
1.0
1.0
1.0
1.0
1.0
1.0
1.0


568727.0
0.0
0.0
0.0
0.0
0.0
0.0
0.0
0.0
0.0
0.0
0.0
0.0
1.0
83208.0
0.0
0.0
0.0
1.0
0.0
0.0
0.0
485517.0
0.0
0.0
0.0
0.0
0.0
0.0
0.0

1.0
1.0
1.0
1.0
1.0
1.0
1.0
1.0
1.0
1.0
1.0
1.0
1.0
1.0
1.0
1.0
1.0
1.0
1.0
1.0
1.0
1.0
1.0
1.0
1.0
1.0
1.0
1.0
1.0
1.0


568671.0
0.0
0.0
0.0
0.0
0.0
0.0
0.0
0.0
0.0
0.0
0.0
0.0
1.0
83180.0
0.0
0.0
0.0
1.0
0.0
0.0
0.0
485489.0
0.0
0.0
0.0
0.0
0.0
0.0
0.0

0.9
0.9
0.9
0.9
0.9
0.9
0.9
0.9
0.9
0.9
0.9
0.9
0.9
0.9
0.9
0.9
0.9
0.9
0.9
0.9
0.9
0.9
0.9
0.9
0.9
0.9
0.9
0.9
0.9
0.9


101852.0
0.0
0.0
0.0
0.0
0.0
0.0
0.0
0.0
0.0
0.0
0.0
0.0
0.0
0.0
0.0
0.0
0.0
101852.0
0.0
0.0
0.0
0.0
0.0
0.0
0.0
0.0
0.0
0.0
0.0


101852.0
0.0
0.0
0.0
0.0
0.0
0.0
0.0
0.0
0.0
0.0
0.0
0.0
0.0
0.0
0.0
0.0
0.0
101852.0
0.0
0.0
0.0
0.0
0.0
0.0
0.0
0.0
0.0
0.0
0.0

1.0
1.0
1.0
1.0
1.0
1.0
1.0
1.0
1.0
1.0
1.0
1.0
1.0
1.0
1.0
1.0
1.0
1.0
1.0
1.0
1.0
1.0
1.0
1.0
1.0
1.0
1.0
1.0
1.0
1.0


110398.0
0.0
0.0
0.0
0.0
0.0
0.0
0.0
0.0
0.0
0.0
0.0
0.0
0.0
0.0
0.0
0.0
0.0
0.0
0.0
0.0
0.0
0.0
0.0
0.0
0.0
0.0
51046.0
12196.0
47156.0


110398.0
0.0
0.0
0.0
0.0
0.0
0.0
0.0
0.0
0.0
0.0
0.0
0.0
0.0
0.0
0.0
0.0
0.0
0.0
0.0
0.0
0.0
0.0
0.0
0.0
0.0
0.0
51046.0
12196.0
47156.0

1.0
1.0
1.0
1.0
1.0
1.0
1.0
1.0
1.0
1.0
1.0
1.0
1.0
1.0
1.0
1.0
1.0
1.0
1.0
1.0
1.0
1.0
1.0
1.0
1.0
1.0
1.0
1.0
1.0
1.0


8353.0
0.0
0.0
0.0
0.0
0.0
0.0
0.0
0.0
0.0
0.0
0.0
0.0
0.0
0.0
0.0
4355.0
0.0
0.0
0.0
0.0
0.0
0.0
0.0
0.0
0.0
0.0
0.0
0.0
3998.0


8353.0
0.0
0.0
0.0
0.0
0.0
0.0
0.0
0.0
0.0
0.0
0.0
0.0
0.0
0.0
0.0
4355.0
0.0
0.0
0.0
0.0
0.0
0.0
0.0
0.0
0.0
0.0
0.0
0.0
3998.0

1.0
1.0
1.0
1.0
1.0
1.0
1.0
1.0
1.0
1.0
1.0
1.0
1.0
1.0
1.0
1.0
1.0
1.0
1.0
1.0
1.0
1.0
1.0
1.0
1.0
1.0
1.0
1.0
1.0
1.0


6526.0
0.0
0.0
0.0
0.0
0.0
0.0
0.0
0.0
0.0
0.0
0.0
0.0
0.0
0.0
0.0
0.0
0.0
0.0
0.0
0.0
0.0
0.0
0.0
0.0
0.0
6526.0
0.0
0.0
0.0


6526.0
0.0
0.0
0.0
0.0
0.0
0.0
0.0
0.0
0.0
0.0
0.0
0.0
0.0
0.0
0.0
0.0
0.0
0.0
0.0
0.0
0.0
0.0
0.0
0.0
0.0
6526.0
0.0
0.0
0.0

1.0
1.0
1.0
1.0
1.0
1.0
1.0
1.0
1.0
1.0
1.0
1.0
1.0
1.0
1.0
1.0
1.0
1.0
1.0
1.0
1.0
1.0
1.0
1.0
1.0
1.0
1.0
1.0
1.0
1.0


31968171.0
928.0
1052.0
0.0
4187.0
0.0
1559.0
63278.0
8125.0
3862.0
6111.0
7387.0
3103701.0
3987655.0
0.0
887645.0
1249339.0
2254154.0
4707101.0
3310939.0
1430190.0
5672167.0
402785.0
3103701.0
0.0
582489.0
297182.0
195388.0
103879.0
583367.0

1.0
1.0
1.0
1.0
1.0
1.0
1.0
1.0
1.0
1.0
1.0
1.0
1.0
1.0
1.0
1.0
1.0
1.0
1.0
1.0
1.0
1.0
1.0
1.0
1.0
1.0
1.0
1.0
1.0
1.0


15972318.0
0.0
0.0
0.0
0.0
0.0
1559.0
32600.0
0.0
0.0
1.0
3.0
2419441.0
0.0
0.0
0.0
0.0
0.0
2702912.0
1888076.0
836023.0
5672166.0
96.0
2419441.0
0.0
0.0
0.0
0.0
0.0
0.0

0.78
0.78
0.78
0.78
0.78
0.78
0.78
0.78
0.78
0.78
0.78
0.78
0.78
0.78
0.78
0.78
0.78
0.78
0.78
0.78
0.78
0.78
0.78
0.78
0.78
0.78
0.78
0.78
0.78
0.78


15972308.0
0.0
0.0
0.0
0.0
0.0
1559.0
32600.0
0.0
0.0
1.0
3.0
2419436.0
0.0
0.0
0.0
0.0
0.0
2702912.0
1888076.0
836023.0
5672166.0
96.0
2419436.0
0.0
0.0
0.0
0.0
0.0
0.0

1.0
1.0
1.0
1.0
1.0
1.0
1.0
1.0
1.0
1.0
1.0
1.0
1.0
1.0
1.0
1.0
1.0
1.0
1.0
1.0
1.0
1.0
1.0
1.0
1.0
1.0
1.0
1.0
1.0
1.0


15287.0
0.0
0.0
0.0
0.0
0.0
0.0
15287.0
0.0
0.0
0.0
0.0
0.0
0.0
0.0
0.0
0.0
0.0
0.0
0.0
0.0
0.0
0.0
0.0
0.0
0.0
0.0
0.0
0.0
0.0


15287.0
0.0
0.0
0.0
0.0
0.0
0.0
15287.0
0.0
0.0
0.0
0.0
0.0
0.0
0.0
0.0
0.0
0.0
0.0
0.0
0.0
0.0
0.0
0.0
0.0
0.0
0.0
0.0
0.0
0.0

1.0
1.0
1.0
1.0
1.0
1.0
1.0
1.0
1.0
1.0
1.0
1.0
1.0
1.0
1.0
1.0
1.0
1.0
1.0
1.0
1.0
1.0
1.0
1.0
1.0
1.0
1.0
1.0
1.0
1.0


1614670.0
0.0
0.0
0.0
0.0
0.0
0.0
11736.0
0.0
3006.0
6110.0
0.0
684131.0
0.0
0.0
0.0
0.0
0.0
1.0
0.0
1.0
0.0
0.0
684131.0
0.0
0.0
225554.0
0.0
0.0
0.0


237290.0
0.0
0.0
0.0
0.0
0.0
0.0
11736.0
0.0
0.0
0.0
0.0
0.0
0.0
0.0
0.0
0.0
0.0
0.0
0.0
0.0
0.0
0.0
0.0
0.0
0.0
225554.0
0.0
0.0
0.0

1.0
1.0
1.0
1.0
1.0
1.0
1.0
1.0
1.0
1.0
1.0
1.0
1.0
1.0
1.0
1.0
1.0
1.0
1.0
1.0
1.0
1.0
1.0
1.0
1.0
1.0
1.0
1.0
1.0
1.0


6109.0
0.0
0.0
0.0
0.0
0.0
0.0
0.0
0.0
0.0
6109.0
0.0
0.0
0.0
0.0
0.0
0.0
0.0
0.0
0.0
0.0
0.0
0.0
0.0
0.0
0.0
0.0
0.0
0.0
0.0

0.83
0.83
0.83
0.83
0.83
0.83
0.83
0.83
0.83
0.83
0.83
0.83
0.83
0.83
0.83
0.83
0.83
0.83
0.83
0.83
0.83
0.83
0.83
0.83
0.83
0.83
0.83
0.83
0.83
0.83


1371271.0
0.0
0.0
0.0
0.0
0.0
0.0
0.0
0.0
3006.0
1.0
0.0
684131.0
0.0
0.0
0.0
0.0
0.0
1.0
0.0
1.0
0.0
0.0
684131.0
0.0
0.0
0.0
0.0
0.0
0.0

0.95
0.95
0.95
0.95
0.95
0.95
0.95
0.95
0.95
0.95
0.95
0.95
0.95
0.95
0.95
0.95
0.95
0.95
0.95
0.95
0.95
0.95
0.95
0.95
0.95
0.95
0.95
0.95
0.95
0.95


6187986.0
0.0
0.0
0.0
0.0
0.0
0.0
0.0
8125.0
856.0
0.0
0.0
0.0
3987623.0
0.0
657325.0
1249335.0
17975.0
84665.0
47471.0
2.0
1.0
1.0
0.0
0.0
0.0
0.0
134606.0
1.0
0.0

1.0
1.0
1.0
1.0
1.0
1.0
1.0
1.0
1.0
1.0
1.0
1.0
1.0
1.0
1.0
1.0
1.0
1.0
1.0
1.0
1.0
1.0
1.0
1.0
1.0
1.0
1.0
1.0
1.0
1.0


3995757.0
0.0
0.0
0.0
0.0
0.0
0.0
0.0
8125.0
0.0
0.0
0.0
0.0
3987623.0
0.0
0.0
1.0
1.0
2.0
0.0
2.0
1.0
1.0
0.0
0.0
0.0
0.0
0.0
1.0
0.0

0.97
0.97
0.97
0.97
0.97
0.97
0.97
0.97
0.97
0.97
0.97
0.97
0.97
0.97
0.97
0.97
0.97
0.97
0.97
0.97
0.97
0.97
0.97
0.97
0.97
0.97
0.97
0.97
0.97
0.97


856.0
0.0
0.0
0.0
0.0
0.0
0.0
0.0
0.0
856.0
0.0
0.0
0.0
0.0
0.0
0.0
0.0
0.0
0.0
0.0
0.0
0.0
0.0
0.0
0.0
0.0
0.0
0.0
0.0
0.0

0.96
0.96
0.96
0.96
0.96
0.96
0.96
0.96
0.96
0.96
0.96
0.96
0.96
0.96
0.96
0.96
0.96
0.96
0.96
0.96
0.96
0.96
0.96
0.96
0.96
0.96
0.96
0.96
0.96
0.96


1906623.0
0.0
0.0
0.0
0.0
0.0
0.0
0.0
0.0
0.0
0.0
0.0
0.0
0.0
0.0
657294.0
1249329.0
0.0
0.0
0.0
0.0
0.0
0.0
0.0
0.0
0.0
0.0
0.0
0.0
0.0

1.0
1.0
1.0
1.0
1.0
1.0
1.0
1.0
1.0
1.0
1.0
1.0
1.0
1.0
1.0
1.0
1.0
1.0
1.0
1.0
1.0
1.0
1.0
1.0
1.0
1.0
1.0
1.0
1.0
1.0


134606.0
0.0
0.0
0.0
0.0
0.0
0.0
0.0
0.0
0.0
0.0
0.0
0.0
0.0
0.0
0.0
0.0
0.0
0.0
0.0
0.0
0.0
0.0
0.0
0.0
0.0
0.0
134606.0
0.0
0.0

1.0
1.0
1.0
1.0
1.0
1.0
1.0
1.0
1.0
1.0
1.0
1.0
1.0
1.0
1.0
1.0
1.0
1.0
1.0
1.0
1.0
1.0
1.0
1.0
1.0
1.0
1.0
1.0
1.0
1.0


150108.0
0.0
0.0
0.0
0.0
0.0
0.0
0.0
0.0
0.0
0.0
0.0
0.0
0.0
0.0
0.0
0.0
17974.0
84663.0
47471.0
0.0
0.0
0.0
0.0
0.0
0.0
0.0
0.0
0.0
0.0

1.0
1.0
1.0
1.0
1.0
1.0
1.0
1.0
1.0
1.0
1.0
1.0
1.0
1.0
1.0
1.0
1.0
1.0
1.0
1.0
1.0
1.0
1.0
1.0
1.0
1.0
1.0
1.0
1.0
1.0


108194.0
0.0
133.0
0.0
4187.0
0.0
0.0
0.0
0.0
0.0
0.0
0.0
0.0
0.0
0.0
0.0
0.0
0.0
0.0
0.0
0.0
0.0
0.0
0.0
0.0
0.0
0.0
12.0
103862.0
0.0


108194.0
0.0
133.0
0.0
4187.0
0.0
0.0
0.0
0.0
0.0
0.0
0.0
0.0
0.0
0.0
0.0
0.0
0.0
0.0
0.0
0.0
0.0
0.0
0.0
0.0
0.0
0.0
12.0
103862.0
0.0

1.0
1.0
1.0
1.0
1.0
1.0
1.0
1.0
1.0
1.0
1.0
1.0
1.0
1.0
1.0
1.0
1.0
1.0
1.0
1.0
1.0
1.0
1.0
1.0
1.0
1.0
1.0
1.0
1.0
1.0


4208.0
0.0
0.0
0.0
0.0
0.0
0.0
0.0
0.0
0.0
0.0
4208.0
0.0
0.0
0.0
0.0
0.0
0.0
0.0
0.0
0.0
0.0
0.0
0.0
0.0
0.0
0.0
0.0
0.0
0.0


4208.0
0.0
0.0
0.0
0.0
0.0
0.0
0.0
0.0
0.0
0.0
4208.0
0.0
0.0
0.0
0.0
0.0
0.0
0.0
0.0
0.0
0.0
0.0
0.0
0.0
0.0
0.0
0.0
0.0
0.0

1.0
1.0
1.0
1.0
1.0
1.0
1.0
1.0
1.0
1.0
1.0
1.0
1.0
1.0
1.0
1.0
1.0
1.0
1.0
1.0
1.0
1.0
1.0
1.0
1.0
1.0
1.0
1.0
1.0
1.0


13872.0
0.0
919.0
0.0
0.0
0.0
0.0
0.0
0.0
0.0
0.0
3176.0
0.0
0.0
0.0
0.0
0.0
0.0
0.0
0.0
0.0
0.0
0.0
0.0
0.0
0.0
9777.0
0.0
0.0
0.0


13872.0
0.0
919.0
0.0
0.0
0.0
0.0
0.0
0.0
0.0
0.0
3176.0
0.0
0.0
0.0
0.0
0.0
0.0
0.0
0.0
0.0
0.0
0.0
0.0
0.0
0.0
9777.0
0.0
0.0
0.0

0.99
0.99
0.99
0.99
0.99
0.99
0.99
0.99
0.99
0.99
0.99
0.99
0.99
0.99
0.99
0.99
0.99
0.99
0.99
0.99
0.99
0.99
0.99
0.99
0.99
0.99
0.99
0.99
0.99
0.99


2777265.0
0.0
0.0
0.0
0.0
0.0
0.0
2592.0
0.0
0.0
0.0
0.0
0.0
1.0
0.0
1.0
0.0
0.0
536693.0
1375228.0
460061.0
0.0
402688.0
0.0
0.0
0.0
0.0
0.0
1.0
0.0

0.81
0.81
0.81
0.81
0.81
0.81
0.81
0.81
0.81
0.81
0.81
0.81
0.81
0.81
0.81
0.81
0.81
0.81
0.81
0.81
0.81
0.81
0.81
0.81
0.81
0.81
0.81
0.81
0.81
0.81


2774670.0
0.0
0.0
0.0
0.0
0.0
0.0
0.0
0.0
0.0
0.0
0.0
0.0
1.0
0.0
1.0
0.0
0.0
536693.0
1375226.0
460060.0
0.0
402688.0
0.0
0.0
0.0
0.0
0.0
1.0
0.0

0.74
0.74
0.74
0.74
0.74
0.74
0.74
0.74
0.74
0.74
0.74
0.74
0.74
0.74
0.74
0.74
0.74
0.74
0.74
0.74
0.74
0.74
0.74
0.74
0.74
0.74
0.74
0.74
0.74
0.74


481.0
481.0
0.0
0.0
0.0
0.0
0.0
0.0
0.0
0.0
0.0
0.0
0.0
0.0
0.0
0.0
0.0
0.0
0.0
0.0
0.0
0.0
0.0
0.0
0.0
0.0
0.0
0.0
0.0
0.0


481.0
481.0
0.0
0.0
0.0
0.0
0.0
0.0
0.0
0.0
0.0
0.0
0.0
0.0
0.0
0.0
0.0
0.0
0.0
0.0
0.0
0.0
0.0
0.0
0.0
0.0
0.0
0.0
0.0
0.0

1.0
1.0
1.0
1.0
1.0
1.0
1.0
1.0
1.0
1.0
1.0
1.0
1.0
1.0
1.0
1.0
1.0
1.0
1.0
1.0
1.0
1.0
1.0
1.0
1.0
1.0
1.0
1.0
1.0
1.0


341.0
341.0
0.0
0.0
0.0
0.0
0.0
0.0
0.0
0.0
0.0
0.0
0.0
0.0
0.0
0.0
0.0
0.0
0.0
0.0
0.0
0.0
0.0
0.0
0.0
0.0
0.0
0.0
0.0
0.0


341.0
341.0
0.0
0.0
0.0
0.0
0.0
0.0
0.0
0.0
0.0
0.0
0.0
0.0
0.0
0.0
0.0
0.0
0.0
0.0
0.0
0.0
0.0
0.0
0.0
0.0
0.0
0.0
0.0
0.0

1.0
1.0
1.0
1.0
1.0
1.0
1.0
1.0
1.0
1.0
1.0
1.0
1.0
1.0
1.0
1.0
1.0
1.0
1.0
1.0
1.0
1.0
1.0
1.0
1.0
1.0
1.0
1.0
1.0
1.0


106.0
106.0
0.0
0.0
0.0
0.0
0.0
0.0
0.0
0.0
0.0
0.0
0.0
0.0
0.0
0.0
0.0
0.0
0.0
0.0
0.0
0.0
0.0
0.0
0.0
0.0
0.0
0.0
0.0
0.0


106.0
106.0
0.0
0.0
0.0
0.0
0.0
0.0
0.0
0.0
0.0
0.0
0.0
0.0
0.0
0.0
0.0
0.0
0.0
0.0
0.0
0.0
0.0
0.0
0.0
0.0
0.0
0.0
0.0
0.0

0.95
0.95
0.95
0.95
0.95
0.95
0.95
0.95
0.95
0.95
0.95
0.95
0.95
0.95
0.95
0.95
0.95
0.95
0.95
0.95
0.95
0.95
0.95
0.95
0.95
0.95
0.95
0.95
0.95
0.95


3305852.0
0.0
0.0
0.0
0.0
0.0
0.0
0.0
0.0
0.0
0.0
0.0
0.0
0.0
0.0
0.0
0.0
2235860.0
1069989.0
2.0
0.0
0.0
0.0
0.0
0.0
0.0
1.0
0.0
0.0
0.0


3305852.0
0.0
0.0
0.0
0.0
0.0
0.0
0.0
0.0
0.0
0.0
0.0
0.0
0.0
0.0
0.0
0.0
2235860.0
1069989.0
2.0
0.0
0.0
0.0
0.0
0.0
0.0
1.0
0.0
0.0
0.0

0.99
0.99
0.99
0.99
0.99
0.99
0.99
0.99
0.99
0.99
0.99
0.99
0.99
0.99
0.99
0.99
0.99
0.99
0.99
0.99
0.99
0.99
0.99
0.99
0.99
0.99
0.99
0.99
0.99
0.99


487395.0
0.0
0.0
0.0
0.0
0.0
0.0
0.0
0.0
0.0
0.0
0.0
0.0
0.0
0.0
0.0
0.0
0.0
0.0
0.0
0.0
0.0
0.0
0.0
0.0
0.0
0.0
0.0
0.0
487395.0


487395.0
0.0
0.0
0.0
0.0
0.0
0.0
0.0
0.0
0.0
0.0
0.0
0.0
0.0
0.0
0.0
0.0
0.0
0.0
0.0
0.0
0.0
0.0
0.0
0.0
0.0
0.0
0.0
0.0
487395.0

0.99
0.99
0.99
0.99
0.99
0.99
0.99
0.99
0.99
0.99
0.99
0.99
0.99
0.99
0.99
0.99
0.99
0.99
0.99
0.99
0.99
0.99
0.99
0.99
0.99
0.99
0.99
0.99
0.99
0.99


446023.0
0.0
0.0
0.0
0.0
0.0
0.0
0.0
0.0
0.0
0.0
0.0
0.0
0.0
0.0
0.0
0.0
3.0
311923.0
0.0
134097.0
0.0
0.0
0.0
0.0
0.0
0.0
0.0
0.0
0.0


445436.0
0.0
0.0
0.0
0.0
0.0
0.0
0.0
0.0
0.0
0.0
0.0
0.0
0.0
0.0
0.0
0.0
3.0
311336.0
0.0
134097.0
0.0
0.0
0.0
0.0
0.0
0.0
0.0
0.0
0.0

1.0
1.0
1.0
1.0
1.0
1.0
1.0
1.0
1.0
1.0
1.0
1.0
1.0
1.0
1.0
1.0
1.0
1.0
1.0
1.0
1.0
1.0
1.0
1.0
1.0
1.0
1.0
1.0
1.0
1.0


587.0
0.0
0.0
0.0
0.0
0.0
0.0
0.0
0.0
0.0
0.0
0.0
0.0
0.0
0.0
0.0
0.0
0.0
587.0
0.0
0.0
0.0
0.0
0.0
0.0
0.0
0.0
0.0
0.0
0.0

0.98
0.98
0.98
0.98
0.98
0.98
0.98
0.98
0.98
0.98
0.98
0.98
0.98
0.98
0.98
0.98
0.98
0.98
0.98
0.98
0.98
0.98
0.98
0.98
0.98
0.98
0.98
0.98
0.98
0.98


246269.0
0.0
0.0
0.0
0.0
0.0
0.0
0.0
0.0
0.0
0.0
0.0
0.0
0.0
0.0
0.0
0.0
0.0
0.0
0.0
0.0
0.0
0.0
0.0
0.0
246269.0
0.0
0.0
0.0
0.0


246269.0
0.0
0.0
0.0
0.0
0.0
0.0
0.0
0.0
0.0
0.0
0.0
0.0
0.0
0.0
0.0
0.0
0.0
0.0
0.0
0.0
0.0
0.0
0.0
0.0
246269.0
0.0
0.0
0.0
0.0

1.0
1.0
1.0
1.0
1.0
1.0
1.0
1.0
1.0
1.0
1.0
1.0
1.0
1.0
1.0
1.0
1.0
1.0
1.0
1.0
1.0
1.0
1.0
1.0
1.0
1.0
1.0
1.0
1.0
1.0


230136.0
0.0
0.0
0.0
0.0
0.0
0.0
0.0
0.0
0.0
0.0
0.0
0.0
0.0
0.0
230136.0
0.0
0.0
0.0
0.0
0.0
0.0
0.0
0.0
0.0
0.0
0.0
0.0
0.0
0.0


230136.0
0.0
0.0
0.0
0.0
0.0
0.0
0.0
0.0
0.0
0.0
0.0
0.0
0.0
0.0
230136.0
0.0
0.0
0.0
0.0
0.0
0.0
0.0
0.0
0.0
0.0
0.0
0.0
0.0
0.0

1.0
1.0
1.0
1.0
1.0
1.0
1.0
1.0
1.0
1.0
1.0
1.0
1.0
1.0
1.0
1.0
1.0
1.0
1.0
1.0
1.0
1.0
1.0
1.0
1.0
1.0
1.0
1.0
1.0
1.0


150387.0
0.0
0.0
0.0
0.0
0.0
0.0
0.0
0.0
0.0
0.0
0.0
0.0
0.0
0.0
0.0
0.0
0.0
0.0
0.0
0.0
0.0
0.0
0.0
0.0
119377.0
0.0
31010.0
0.0
0.0

1.0
1.0
1.0
1.0
1.0
1.0
1.0
1.0
1.0
1.0
1.0
1.0
1.0
1.0
1.0
1.0
1.0
1.0
1.0
1.0
1.0
1.0
1.0
1.0
1.0
1.0
1.0
1.0
1.0
1.0


147389.0
0.0
0.0
0.0
0.0
0.0
0.0
0.0
0.0
0.0
0.0
0.0
0.0
0.0
0.0
0.0
0.0
0.0
0.0
0.0
0.0
0.0
0.0
0.0
0.0
85557.0
61832.0
0.0
0.0
0.0


147389.0
0.0
0.0
0.0
0.0
0.0
0.0
0.0
0.0
0.0
0.0
0.0
0.0
0.0
0.0
0.0
0.0
0.0
0.0
0.0
0.0
0.0
0.0
0.0
0.0
85557.0
61832.0
0.0
0.0
0.0

1.0
1.0
1.0
1.0
1.0
1.0
1.0
1.0
1.0
1.0
1.0
1.0
1.0
1.0
1.0
1.0
1.0
1.0
1.0
1.0
1.0
1.0
1.0
1.0
1.0
1.0
1.0
1.0
1.0
1.0


131279.0
0.0
0.0
0.0
0.0
0.0
0.0
0.0
0.0
0.0
0.0
0.0
0.0
0.0
0.0
0.0
0.0
0.0
0.0
0.0
0.0
0.0
0.0
0.0
0.0
131279.0
0.0
0.0
0.0
0.0


131279.0
0.0
0.0
0.0
0.0
0.0
0.0
0.0
0.0
0.0
0.0
0.0
0.0
0.0
0.0
0.0
0.0
0.0
0.0
0.0
0.0
0.0
0.0
0.0
0.0
131279.0
0.0
0.0
0.0
0.0

1.0
1.0
1.0
1.0
1.0
1.0
1.0
1.0
1.0
1.0
1.0
1.0
1.0
1.0
1.0
1.0
1.0
1.0
1.0
1.0
1.0
1.0
1.0
1.0
1.0
1.0
1.0
1.0
1.0
1.0


95010.0
0.0
0.0
0.0
0.0
0.0
0.0
0.0
0.0
0.0
0.0
0.0
0.0
0.0
0.0
0.0
0.0
0.0
0.0
0.0
0.0
0.0
0.0
0.0
0.0
0.0
0.0
0.0
0.0
95010.0

0.9
0.9
0.9
0.9
0.9
0.9
0.9
0.9
0.9
0.9
0.9
0.9
0.9
0.9
0.9
0.9
0.9
0.9
0.9
0.9
0.9
0.9
0.9
0.9
0.9
0.9
0.9
0.9
0.9
0.9


94757.0
0.0
0.0
0.0
0.0
0.0
0.0
0.0
0.0
0.0
0.0
0.0
0.0
0.0
0.0
0.0
0.0
0.0
0.0
0.0
0.0
0.0
0.0
0.0
0.0
0.0
0.0
0.0
0.0
94757.0

1.0
1.0
1.0
1.0
1.0
1.0
1.0
1.0
1.0
1.0
1.0
1.0
1.0
1.0
1.0
1.0
1.0
1.0
1.0
1.0
1.0
1.0
1.0
1.0
1.0
1.0
1.0
1.0
1.0
1.0


29738.0
0.0
0.0
0.0
0.0
0.0
0.0
0.0
0.0
0.0
0.0
0.0
0.0
0.0
0.0
0.0
0.0
0.0
0.0
0.0
0.0
0.0
0.0
0.0
0.0
0.0
0.0
29738.0
0.0
0.0


29738.0
0.0
0.0
0.0
0.0
0.0
0.0
0.0
0.0
0.0
0.0
0.0
0.0
0.0
0.0
0.0
0.0
0.0
0.0
0.0
0.0
0.0
0.0
0.0
0.0
0.0
0.0
29738.0
0.0
0.0

1.0
1.0
1.0
1.0
1.0
1.0
1.0
1.0
1.0
1.0
1.0
1.0
1.0
1.0
1.0
1.0
1.0
1.0
1.0
1.0
1.0
1.0
1.0
1.0
1.0
1.0
1.0
1.0
1.0
1.0


30.0
0.0
0.0
0.0
0.0
0.0
0.0
0.0
0.0
0.0
0.0
0.0
0.0
0.0
0.0
30.0
0.0
0.0
0.0
0.0
0.0
0.0
0.0
0.0
0.0
0.0
0.0
0.0
0.0
0.0

0.87
0.87
0.87
0.87
0.87
0.87
0.87
0.87
0.87
0.87
0.87
0.87
0.87
0.87
0.87
0.87
0.87
0.87
0.87
0.87
0.87
0.87
0.87
0.87
0.87
0.87
0.87
0.87
0.87
0.87


52.0
0.0
0.0
0.0
0.0
0.0
0.0
0.0
0.0
0.0
0.0
0.0
26.0
0.0
0.0
0.0
0.0
0.0
0.0
0.0
0.0
0.0
0.0
26.0
0.0
0.0
0.0
0.0
0.0
0.0

0.91
0.91
0.91
0.91
0.91
0.91
0.91
0.91
0.91
0.91
0.91
0.91
0.91
0.91
0.91
0.91
0.91
0.91
0.91
0.91
0.91
0.91
0.91
0.91
0.91
0.91
0.91
0.91
0.91
0.91


6.0
0.0
0.0
0.0
0.0
0.0
0.0
0.0
0.0
0.0
0.0
0.0
3.0
0.0
0.0
0.0
0.0
0.0
0.0
0.0
0.0
0.0
0.0
3.0
0.0
0.0
0.0
0.0
0.0
0.0

0.84
0.84
0.84
0.84
0.84
0.84
0.84
0.84
0.84
0.84
0.84
0.84
0.84
0.84
0.84
0.84
0.84
0.84
0.84
0.84
0.84
0.84
0.84
0.84
0.84
0.84
0.84
0.84
0.84
0.84


6.0
0.0
0.0
0.0
0.0
0.0
0.0
0.0
0.0
0.0
0.0
0.0
0.0
0.0
0.0
2.0
4.0
0.0
0.0
0.0
0.0
0.0
0.0
0.0
0.0
0.0
0.0
0.0
0.0
0.0

0.85
0.85
0.85
0.85
0.85
0.85
0.85
0.85
0.85
0.85
0.85
0.85
0.85
0.85
0.85
0.85
0.85
0.85
0.85
0.85
0.85
0.85
0.85
0.85
0.85
0.85
0.85
0.85
0.85
0.85


1135777.0
1530.0
1030.0
622.0
7445.0
20176.0
3954.0
896.0
1041.0
1104.0
2067.0
1366.0
0.0
54127.0
9132.0
981.0
14943.0
14668.0
2983.0
12548.0
49552.0
14183.0
0.0
0.0
27026.0
174713.0
209671.0
281886.0
136129.0
92004.0


198266.0
568.0
155.0
147.0
6235.0
19136.0
3150.0
896.0
1041.0
619.0
1656.0
334.0
0.0
0.0
1104.0
981.0
0.0
0.0
0.0
0.0
0.0
0.0
0.0
0.0
9029.0
20676.0
47779.0
57894.0
12856.0
14010.0

0.74
0.74
0.74
0.74
0.74
0.74
0.74
0.74
0.74
0.74
0.74
0.74
0.74
0.74
0.74
0.74
0.74
0.74
0.74
0.74
0.74
0.74
0.74
0.74
0.74
0.74
0.74
0.74
0.74
0.74


182841.0
568.0
155.0
147.0
6235.0
19105.0
3150.0
896.0
1041.0
619.0
1656.0
334.0
0.0
0.0
1104.0
981.0
0.0
0.0
0.0
0.0
0.0
0.0
0.0
0.0
9029.0
20676.0
47779.0
43887.0
12459.0
13020.0

1.0
1.0
1.0
1.0
1.0
1.0
1.0
1.0
1.0
1.0
1.0
1.0
1.0
1.0
1.0
1.0
1.0
1.0
1.0
1.0
1.0
1.0
1.0
1.0
1.0
1.0
1.0
1.0
1.0
1.0


14007.0
0.0
0.0
0.0
0.0
0.0
0.0
0.0
0.0
0.0
0.0
0.0
0.0
0.0
0.0
0.0
0.0
0.0
0.0
0.0
0.0
0.0
0.0
0.0
0.0
0.0
0.0
14007.0
0.0
0.0

1.0
1.0
1.0
1.0
1.0
1.0
1.0
1.0
1.0
1.0
1.0
1.0
1.0
1.0
1.0
1.0
1.0
1.0
1.0
1.0
1.0
1.0
1.0
1.0
1.0
1.0
1.0
1.0
1.0
1.0


1387.0
0.0
0.0
0.0
0.0
0.0
0.0
0.0
0.0
0.0
0.0
0.0
0.0
0.0
0.0
0.0
0.0
0.0
0.0
0.0
0.0
0.0
0.0
0.0
0.0
0.0
0.0
0.0
397.0
990.0

1.0
1.0
1.0
1.0
1.0
1.0
1.0
1.0
1.0
1.0
1.0
1.0
1.0
1.0
1.0
1.0
1.0
1.0
1.0
1.0
1.0
1.0
1.0
1.0
1.0
1.0
1.0
1.0
1.0
1.0


88380.0
368.0
445.0
336.0
1110.0
1015.0
489.0
0.0
0.0
452.0
306.0
91.0
0.0
0.0
1779.0
0.0
0.0
11597.0
0.0
0.0
0.0
0.0
0.0
0.0
3507.0
0.0
13402.0
5764.0
40474.0
7245.0


88380.0
368.0
445.0
336.0
1110.0
1015.0
489.0
0.0
0.0
452.0
306.0
91.0
0.0
0.0
1779.0
0.0
0.0
11597.0
0.0
0.0
0.0
0.0
0.0
0.0
3507.0
0.0
13402.0
5764.0
40474.0
7245.0

1.0
1.0
1.0
1.0
1.0
1.0
1.0
1.0
1.0
1.0
1.0
1.0
1.0
1.0
1.0
1.0
1.0
1.0
1.0
1.0
1.0
1.0
1.0
1.0
1.0
1.0
1.0
1.0
1.0
1.0


11615.0
591.0
397.0
0.0
0.0
0.0
0.0
0.0
0.0
0.0
0.0
896.0
0.0
0.0
0.0
0.0
0.0
0.0
2983.0
0.0
6748.0
0.0
0.0
0.0
0.0
0.0
0.0
0.0
0.0
0.0


1874.0
591.0
387.0
0.0
0.0
0.0
0.0
0.0
0.0
0.0
0.0
896.0
0.0
0.0
0.0
0.0
0.0
0.0
0.0
0.0
0.0
0.0
0.0
0.0
0.0
0.0
0.0
0.0
0.0
0.0

1.0
1.0
1.0
1.0
1.0
1.0
1.0
1.0
1.0
1.0
1.0
1.0
1.0
1.0
1.0
1.0
1.0
1.0
1.0
1.0
1.0
1.0
1.0
1.0
1.0
1.0
1.0
1.0
1.0
1.0


10.0
0.0
10.0
0.0
0.0
0.0
0.0
0.0
0.0
0.0
0.0
0.0
0.0
0.0
0.0
0.0
0.0
0.0
0.0
0.0
0.0
0.0
0.0
0.0
0.0
0.0
0.0
0.0
0.0
0.0

1.0
1.0
1.0
1.0
1.0
1.0
1.0
1.0
1.0
1.0
1.0
1.0
1.0
1.0
1.0
1.0
1.0
1.0
1.0
1.0
1.0
1.0
1.0
1.0
1.0
1.0
1.0
1.0
1.0
1.0


9731.0
0.0
0.0
0.0
0.0
0.0
0.0
0.0
0.0
0.0
0.0
0.0
0.0
0.0
0.0
0.0
0.0
0.0
2983.0
0.0
6748.0
0.0
0.0
0.0
0.0
0.0
0.0
0.0
0.0
0.0

1.0
1.0
1.0
1.0
1.0
1.0
1.0
1.0
1.0
1.0
1.0
1.0
1.0
1.0
1.0
1.0
1.0
1.0
1.0
1.0
1.0
1.0
1.0
1.0
1.0
1.0
1.0
1.0
1.0
1.0


118877.0
0.0
0.0
0.0
0.0
0.0
195.0
0.0
0.0
0.0
0.0
0.0
0.0
0.0
0.0
0.0
0.0
0.0
0.0
0.0
0.0
0.0
0.0
0.0
0.0
0.0
38981.0
18463.0
36884.0
24354.0


118877.0
0.0
0.0
0.0
0.0
0.0
195.0
0.0
0.0
0.0
0.0
0.0
0.0
0.0
0.0
0.0
0.0
0.0
0.0
0.0
0.0
0.0
0.0
0.0
0.0
0.0
38981.0
18463.0
36884.0
24354.0

1.0
1.0
1.0
1.0
1.0
1.0
1.0
1.0
1.0
1.0
1.0
1.0
1.0
1.0
1.0
1.0
1.0
1.0
1.0
1.0
1.0
1.0
1.0
1.0
1.0
1.0
1.0
1.0
1.0
1.0


132.0
0.0
0.0
0.0
100.0
0.0
32.0
0.0
0.0
0.0
0.0
0.0
0.0
0.0
0.0
0.0
0.0
0.0
0.0
0.0
0.0
0.0
0.0
0.0
0.0
0.0
0.0
0.0
0.0
0.0


132.0
0.0
0.0
0.0
100.0
0.0
32.0
0.0
0.0
0.0
0.0
0.0
0.0
0.0
0.0
0.0
0.0
0.0
0.0
0.0
0.0
0.0
0.0
0.0
0.0
0.0
0.0
0.0
0.0
0.0

1.0
1.0
1.0
1.0
1.0
1.0
1.0
1.0
1.0
1.0
1.0
1.0
1.0
1.0
1.0
1.0
1.0
1.0
1.0
1.0
1.0
1.0
1.0
1.0
1.0
1.0
1.0
1.0
1.0
1.0


13940.0
0.0
0.0
90.0
0.0
0.0
0.0
0.0
0.0
0.0
0.0
0.0
0.0
0.0
0.0
0.0
0.0
0.0
0.0
0.0
0.0
0.0
0.0
0.0
0.0
2658.0
11192.0
0.0
0.0
0.0


13940.0
0.0
0.0
90.0
0.0
0.0
0.0
0.0
0.0
0.0
0.0
0.0
0.0
0.0
0.0
0.0
0.0
0.0
0.0
0.0
0.0
0.0
0.0
0.0
0.0
2658.0
11192.0
0.0
0.0
0.0

1.0
1.0
1.0
1.0
1.0
1.0
1.0
1.0
1.0
1.0
1.0
1.0
1.0
1.0
1.0
1.0
1.0
1.0
1.0
1.0
1.0
1.0
1.0
1.0
1.0
1.0
1.0
1.0
1.0
1.0


3097.0
0.0
0.0
11.0
0.0
0.0
0.0
0.0
0.0
0.0
25.0
45.0
0.0
0.0
0.0
0.0
0.0
0.0
0.0
0.0
0.0
0.0
0.0
0.0
0.0
0.0
0.0
0.0
0.0
3016.0


3097.0
0.0
0.0
11.0
0.0
0.0
0.0
0.0
0.0
0.0
25.0
45.0
0.0
0.0
0.0
0.0
0.0
0.0
0.0
0.0
0.0
0.0
0.0
0.0
0.0
0.0
0.0
0.0
0.0
3016.0

1.0
1.0
1.0
1.0
1.0
1.0
1.0
1.0
1.0
1.0
1.0
1.0
1.0
1.0
1.0
1.0
1.0
1.0
1.0
1.0
1.0
1.0
1.0
1.0
1.0
1.0
1.0
1.0
1.0
1.0


149828.0
0.0
0.0
0.0
0.0
0.0
71.0
0.0
0.0
33.0
0.0
0.0
0.0
0.0
0.0
0.0
0.0
0.0
0.0
0.0
42804.0
14183.0
0.0
0.0
0.0
0.0
12553.0
64960.0
2955.0
12269.0

1.0
1.0
1.0
1.0
1.0
1.0
1.0
1.0
1.0
1.0
1.0
1.0
1.0
1.0
1.0
1.0
1.0
1.0
1.0
1.0
1.0
1.0
1.0
1.0
1.0
1.0
1.0
1.0
1.0
1.0


92612.0
0.0
0.0
0.0
0.0
0.0
71.0
0.0
0.0
0.0
0.0
0.0
0.0
0.0
0.0
0.0
0.0
0.0
0.0
0.0
0.0
0.0
0.0
0.0
0.0
0.0
12553.0
64764.0
2955.0
12269.0

1.0
1.0
1.0
1.0
1.0
1.0
1.0
1.0
1.0
1.0
1.0
1.0
1.0
1.0
1.0
1.0
1.0
1.0
1.0
1.0
1.0
1.0
1.0
1.0
1.0
1.0
1.0
1.0
1.0
1.0


57020.0
0.0
0.0
0.0
0.0
0.0
0.0
0.0
0.0
33.0
0.0
0.0
0.0
0.0
0.0
0.0
0.0
0.0
0.0
0.0
42804.0
14183.0
0.0
0.0
0.0
0.0
0.0
0.0
0.0
0.0

1.0
1.0
1.0
1.0
1.0
1.0
1.0
1.0
1.0
1.0
1.0
1.0
1.0
1.0
1.0
1.0
1.0
1.0
1.0
1.0
1.0
1.0
1.0
1.0
1.0
1.0
1.0
1.0
1.0
1.0


45.0
0.0
0.0
0.0
0.0
0.0
0.0
0.0
0.0
0.0
45.0
0.0
0.0
0.0
0.0
0.0
0.0
0.0
0.0
0.0
0.0
0.0
0.0
0.0
0.0
0.0
0.0
0.0
0.0
0.0

1.0
1.0
1.0
1.0
1.0
1.0
1.0
1.0
1.0
1.0
1.0
1.0
1.0
1.0
1.0
1.0
1.0
1.0
1.0
1.0
1.0
1.0
1.0
1.0
1.0
1.0
1.0
1.0
1.0
1.0


35.0
0.0
0.0
0.0
0.0
0.0
0.0
0.0
0.0
0.0
35.0
0.0
0.0
0.0
0.0
0.0
0.0
0.0
0.0
0.0
0.0
0.0
0.0
0.0
0.0
0.0
0.0
0.0
0.0
0.0


35.0
0.0
0.0
0.0
0.0
0.0
0.0
0.0
0.0
0.0
35.0
0.0
0.0
0.0
0.0
0.0
0.0
0.0
0.0
0.0
0.0
0.0
0.0
0.0
0.0
0.0
0.0
0.0
0.0
0.0

1.0
1.0
1.0
1.0
1.0
1.0
1.0
1.0
1.0
1.0
1.0
1.0
1.0
1.0
1.0
1.0
1.0
1.0
1.0
1.0
1.0
1.0
1.0
1.0
1.0
1.0
1.0
1.0
1.0
1.0


24244.0
0.0
3.0
18.0
0.0
8.0
0.0
0.0
0.0
0.0
0.0
0.0
0.0
0.0
0.0
0.0
8596.0
3071.0
0.0
12548.0
0.0
0.0
0.0
0.0
0.0
0.0
0.0
0.0
0.0
0.0


24244.0
0.0
3.0
18.0
0.0
8.0
0.0
0.0
0.0
0.0
0.0
0.0
0.0
0.0
0.0
0.0
8596.0
3071.0
0.0
12548.0
0.0
0.0
0.0
0.0
0.0
0.0
0.0
0.0
0.0
0.0

1.0
1.0
1.0
1.0
1.0
1.0
1.0
1.0
1.0
1.0
1.0
1.0
1.0
1.0
1.0
1.0
1.0
1.0
1.0
1.0
1.0
1.0
1.0
1.0
1.0
1.0
1.0
1.0
1.0
1.0


19.0
0.0
0.0
10.0
0.0
9.0
0.0
0.0
0.0
0.0
0.0
0.0
0.0
0.0
0.0
0.0
0.0
0.0
0.0
0.0
0.0
0.0
0.0
0.0
0.0
0.0
0.0
0.0
0.0
0.0


19.0
0.0
0.0
10.0
0.0
9.0
0.0
0.0
0.0
0.0
0.0
0.0
0.0
0.0
0.0
0.0
0.0
0.0
0.0
0.0
0.0
0.0
0.0
0.0
0.0
0.0
0.0
0.0
0.0
0.0

1.0
1.0
1.0
1.0
1.0
1.0
1.0
1.0
1.0
1.0
1.0
1.0
1.0
1.0
1.0
1.0
1.0
1.0
1.0
1.0
1.0
1.0
1.0
1.0
1.0
1.0
1.0
1.0
1.0
1.0


6247.0
0.0
4.0
0.0
0.0
8.0
6.0
0.0
0.0
0.0
0.0
0.0
0.0
0.0
0.0
0.0
0.0
0.0
0.0
0.0
0.0
0.0
0.0
0.0
6229.0
0.0
0.0
0.0
0.0
0.0


6247.0
0.0
4.0
0.0
0.0
8.0
6.0
0.0
0.0
0.0
0.0
0.0
0.0
0.0
0.0
0.0
0.0
0.0
0.0
0.0
0.0
0.0
0.0
0.0
6229.0
0.0
0.0
0.0
0.0
0.0

0.98
0.98
0.98
0.98
0.98
0.98
0.98
0.98
0.98
0.98
0.98
0.98
0.98
0.98
0.98
0.98
0.98
0.98
0.98
0.98
0.98
0.98
0.98
0.98
0.98
0.98
0.98
0.98
0.98
0.98


60403.0
3.0
14.0
10.0
0.0
0.0
0.0
0.0
0.0
0.0
0.0
0.0
0.0
54127.0
6249.0
0.0
0.0
0.0
0.0
0.0
0.0
0.0
0.0
0.0
0.0
0.0
0.0
0.0
0.0
0.0


60403.0
3.0
14.0
10.0
0.0
0.0
0.0
0.0
0.0
0.0
0.0
0.0
0.0
54127.0
6249.0
0.0
0.0
0.0
0.0
0.0
0.0
0.0
0.0
0.0
0.0
0.0
0.0
0.0
0.0
0.0

1.0
1.0
1.0
1.0
1.0
1.0
1.0
1.0
1.0
1.0
1.0
1.0
1.0
1.0
1.0
1.0
1.0
1.0
1.0
1.0
1.0
1.0
1.0
1.0
1.0
1.0
1.0
1.0
1.0
1.0


6076.0
0.0
12.0
0.0
0.0
0.0
0.0
0.0
0.0
0.0
0.0
0.0
0.0
0.0
0.0
0.0
0.0
0.0
0.0
0.0
0.0
0.0
0.0
0.0
6064.0
0.0
0.0
0.0
0.0
0.0


6076.0
0.0
12.0
0.0
0.0
0.0
0.0
0.0
0.0
0.0
0.0
0.0
0.0
0.0
0.0
0.0
0.0
0.0
0.0
0.0
0.0
0.0
0.0
0.0
6064.0
0.0
0.0
0.0
0.0
0.0

1.0
1.0
1.0
1.0
1.0
1.0
1.0
1.0
1.0
1.0
1.0
1.0
1.0
1.0
1.0
1.0
1.0
1.0
1.0
1.0
1.0
1.0
1.0
1.0
1.0
1.0
1.0
1.0
1.0
1.0


41434.0
0.0
0.0
0.0
0.0
0.0
11.0
0.0
0.0
0.0
0.0
0.0
0.0
0.0
0.0
0.0
0.0
0.0
0.0
0.0
0.0
0.0
0.0
0.0
0.0
34631.0
6792.0
0.0
0.0
0.0


41434.0
0.0
0.0
0.0
0.0
0.0
11.0
0.0
0.0
0.0
0.0
0.0
0.0
0.0
0.0
0.0
0.0
0.0
0.0
0.0
0.0
0.0
0.0
0.0
0.0
34631.0
6792.0
0.0
0.0
0.0

1.0
1.0
1.0
1.0
1.0
1.0
1.0
1.0
1.0
1.0
1.0
1.0
1.0
1.0
1.0
1.0
1.0
1.0
1.0
1.0
1.0
1.0
1.0
1.0
1.0
1.0
1.0
1.0
1.0
1.0


370522.0
0.0
0.0
0.0
0.0
0.0
0.0
0.0
0.0
0.0
0.0
0.0
0.0
0.0
0.0
0.0
6268.0
0.0
0.0
0.0
0.0
0.0
0.0
0.0
0.0
115410.0
78224.0
98112.0
41777.0
30731.0


370522.0
0.0
0.0
0.0
0.0
0.0
0.0
0.0
0.0
0.0
0.0
0.0
0.0
0.0
0.0
0.0
6268.0
0.0
0.0
0.0
0.0
0.0
0.0
0.0
0.0
115410.0
78224.0
98112.0
41777.0
30731.0

1.0
1.0
1.0
1.0
1.0
1.0
1.0
1.0
1.0
1.0
1.0
1.0
1.0
1.0
1.0
1.0
1.0
1.0
1.0
1.0
1.0
1.0
1.0
1.0
1.0
1.0
1.0
1.0
1.0
1.0


36257.0
0.0
0.0
0.0
0.0
0.0
0.0
0.0
0.0
0.0
0.0
0.0
0.0
0.0
0.0
0.0
0.0
0.0
0.0
0.0
0.0
0.0
0.0
0.0
0.0
0.0
0.0
36182.0
75.0
0.0


36257.0
0.0
0.0
0.0
0.0
0.0
0.0
0.0
0.0
0.0
0.0
0.0
0.0
0.0
0.0
0.0
0.0
0.0
0.0
0.0
0.0
0.0
0.0
0.0
0.0
0.0
0.0
36182.0
75.0
0.0

1.0
1.0
1.0
1.0
1.0
1.0
1.0
1.0
1.0
1.0
1.0
1.0
1.0
1.0
1.0
1.0
1.0
1.0
1.0
1.0
1.0
1.0
1.0
1.0
1.0
1.0
1.0
1.0
1.0
1.0


2536.0
0.0
0.0
0.0
0.0
0.0
0.0
0.0
0.0
0.0
0.0
0.0
0.0
0.0
0.0
0.0
0.0
0.0
0.0
0.0
0.0
0.0
0.0
0.0
2197.0
339.0
0.0
0.0
0.0
0.0


2197.0
0.0
0.0
0.0
0.0
0.0
0.0
0.0
0.0
0.0
0.0
0.0
0.0
0.0
0.0
0.0
0.0
0.0
0.0
0.0
0.0
0.0
0.0
0.0
2197.0
0.0
0.0
0.0
0.0
0.0

1.0
1.0
1.0
1.0
1.0
1.0
1.0
1.0
1.0
1.0
1.0
1.0
1.0
1.0
1.0
1.0
1.0
1.0
1.0
1.0
1.0
1.0
1.0
1.0
1.0
1.0
1.0
1.0
1.0
1.0


339.0
0.0
0.0
0.0
0.0
0.0
0.0
0.0
0.0
0.0
0.0
0.0
0.0
0.0
0.0
0.0
0.0
0.0
0.0
0.0
0.0
0.0
0.0
0.0
0.0
339.0
0.0
0.0
0.0
0.0

1.0
1.0
1.0
1.0
1.0
1.0
1.0
1.0
1.0
1.0
1.0
1.0
1.0
1.0
1.0
1.0
1.0
1.0
1.0
1.0
1.0
1.0
1.0
1.0
1.0
1.0
1.0
1.0
1.0
1.0


2959.0
0.0
0.0
0.0
0.0
0.0
0.0
0.0
0.0
0.0
0.0
0.0
0.0
0.0
0.0
0.0
79.0
0.0
0.0
0.0
0.0
0.0
0.0
0.0
0.0
999.0
748.0
511.0
243.0
379.0


2959.0
0.0
0.0
0.0
0.0
0.0
0.0
0.0
0.0
0.0
0.0
0.0
0.0
0.0
0.0
0.0
79.0
0.0
0.0
0.0
0.0
0.0
0.0
0.0
0.0
999.0
748.0
511.0
243.0
379.0

1.0
1.0
1.0
1.0
1.0
1.0
1.0
1.0
1.0
1.0
1.0
1.0
1.0
1.0
1.0
1.0
1.0
1.0
1.0
1.0
1.0
1.0
1.0
1.0
1.0
1.0
1.0
1.0
1.0
1.0


865.0
0.0
0.0
0.0
0.0
0.0
0.0
0.0
0.0
0.0
0.0
0.0
0.0
0.0
0.0
0.0
0.0
0.0
0.0
0.0
0.0
0.0
0.0
0.0
0.0
0.0
0.0
0.0
865.0
0.0


865.0
0.0
0.0
0.0
0.0
0.0
0.0
0.0
0.0
0.0
0.0
0.0
0.0
0.0
0.0
0.0
0.0
0.0
0.0
0.0
0.0
0.0
0.0
0.0
0.0
0.0
0.0
0.0
865.0
0.0

0.97
0.97
0.97
0.97
0.97
0.97
0.97
0.97
0.97
0.97
0.97
0.97
0.97
0.97
0.97
0.97
0.97
0.97
0.97
0.97
0.97
0.97
0.97
0.97
0.97
0.97
0.97
0.97
0.97
0.97


2863064.0
126.0
629.0
194.0
0.0
0.0
43.0
0.0
0.0
26234.0
355.0
2.0
0.0
251970.0
0.0
1.0
0.0
2537462.0
1.0
45816.0
2.0
2.0
0.0
0.0
1.0
0.0
1.0
225.0
0.0
0.0

0.7
0.7
0.7
0.7
0.7
0.7
0.7
0.7
0.7
0.7
0.7
0.7
0.7
0.7
0.7
0.7
0.7
0.7
0.7
0.7
0.7
0.7
0.7
0.7
0.7
0.7
0.7
0.7
0.7
0.7


2512843.0
0.0
0.0
0.0
0.0
0.0
0.0
0.0
0.0
26234.0
0.0
2.0
0.0
0.0
0.0
1.0
0.0
2486597.0
1.0
1.0
2.0
2.0
0.0
0.0
1.0
0.0
1.0
1.0
0.0
0.0


26236.0
0.0
0.0
0.0
0.0
0.0
0.0
0.0
0.0
26234.0
0.0
2.0
0.0
0.0
0.0
0.0
0.0
0.0
0.0
0.0
0.0
0.0
0.0
0.0
0.0
0.0
0.0
0.0
0.0
0.0

0.99
0.99
0.99
0.99
0.99
0.99
0.99
0.99
0.99
0.99
0.99
0.99
0.99
0.99
0.99
0.99
0.99
0.99
0.99
0.99
0.99
0.99
0.99
0.99
0.99
0.99
0.99
0.99
0.99
0.99


2486607.0
0.0
0.0
0.0
0.0
0.0
0.0
0.0
0.0
0.0
0.0
0.0
0.0
0.0
0.0
1.0
0.0
2486597.0
1.0
1.0
2.0
2.0
0.0
0.0
1.0
0.0
1.0
1.0
0.0
0.0

1.0
1.0
1.0
1.0
1.0
1.0
1.0
1.0
1.0
1.0
1.0
1.0
1.0
1.0
1.0
1.0
1.0
1.0
1.0
1.0
1.0
1.0
1.0
1.0
1.0
1.0
1.0
1.0
1.0
1.0


252793.0
0.0
629.0
194.0
0.0
0.0
0.0
0.0
0.0
0.0
0.0
0.0
0.0
251970.0
0.0
0.0
0.0
0.0
0.0
0.0
0.0
0.0
0.0
0.0
0.0
0.0
0.0
0.0
0.0
0.0


252793.0
0.0
629.0
194.0
0.0
0.0
0.0
0.0
0.0
0.0
0.0
0.0
0.0
251970.0
0.0
0.0
0.0
0.0
0.0
0.0
0.0
0.0
0.0
0.0
0.0
0.0
0.0
0.0
0.0
0.0

1.0
1.0
1.0
1.0
1.0
1.0
1.0
1.0
1.0
1.0
1.0
1.0
1.0
1.0
1.0
1.0
1.0
1.0
1.0
1.0
1.0
1.0
1.0
1.0
1.0
1.0
1.0
1.0
1.0
1.0


97199.0
126.0
0.0
0.0
0.0
0.0
43.0
0.0
0.0
0.0
355.0
0.0
0.0
0.0
0.0
0.0
0.0
50860.0
0.0
45815.0
0.0
0.0
0.0
0.0
0.0
0.0
0.0
0.0
0.0
0.0


97199.0
126.0
0.0
0.0
0.0
0.0
43.0
0.0
0.0
0.0
355.0
0.0
0.0
0.0
0.0
0.0
0.0
50860.0
0.0
45815.0
0.0
0.0
0.0
0.0
0.0
0.0
0.0
0.0
0.0
0.0

1.0
1.0
1.0
1.0
1.0
1.0
1.0
1.0
1.0
1.0
1.0
1.0
1.0
1.0
1.0
1.0
1.0
1.0
1.0
1.0
1.0
1.0
1.0
1.0
1.0
1.0
1.0
1.0
1.0
1.0


224.0
0.0
0.0
0.0
0.0
0.0
0.0
0.0
0.0
0.0
0.0
0.0
0.0
0.0
0.0
0.0
0.0
0.0
0.0
0.0
0.0
0.0
0.0
0.0
0.0
0.0
0.0
224.0
0.0
0.0

0.73
0.73
0.73
0.73
0.73
0.73
0.73
0.73
0.73
0.73
0.73
0.73
0.73
0.73
0.73
0.73
0.73
0.73
0.73
0.73
0.73
0.73
0.73
0.73
0.73
0.73
0.73
0.73
0.73
0.73


130566.0
1840.0
517.0
72.0
0.0
0.0
0.0
0.0
0.0
2676.0
759.0
671.0
0.0
21589.0
15213.0
0.0
0.0
0.0
87229.0
0.0
0.0
0.0
0.0
0.0
0.0
0.0
0.0
0.0
0.0
0.0


129567.0
1131.0
227.0
72.0
0.0
0.0
0.0
0.0
0.0
2676.0
759.0
671.0
0.0
21589.0
15213.0
0.0
0.0
0.0
87229.0
0.0
0.0
0.0
0.0
0.0
0.0
0.0
0.0
0.0
0.0
0.0


129567.0
1131.0
227.0
72.0
0.0
0.0
0.0
0.0
0.0
2676.0
759.0
671.0
0.0
21589.0
15213.0
0.0
0.0
0.0
87229.0
0.0
0.0
0.0
0.0
0.0
0.0
0.0
0.0
0.0
0.0
0.0

1.0
1.0
1.0
1.0
1.0
1.0
1.0
1.0
1.0
1.0
1.0
1.0
1.0
1.0
1.0
1.0
1.0
1.0
1.0
1.0
1.0
1.0
1.0
1.0
1.0
1.0
1.0
1.0
1.0
1.0


999.0
709.0
290.0
0.0
0.0
0.0
0.0
0.0
0.0
0.0
0.0
0.0
0.0
0.0
0.0
0.0
0.0
0.0
0.0
0.0
0.0
0.0
0.0
0.0
0.0
0.0
0.0
0.0
0.0
0.0

1.0
1.0
1.0
1.0
1.0
1.0
1.0
1.0
1.0
1.0
1.0
1.0
1.0
1.0
1.0
1.0
1.0
1.0
1.0
1.0
1.0
1.0
1.0
1.0
1.0
1.0
1.0
1.0
1.0
1.0


1237413.0
850.0
189.0
583.0
849.0
333.0
45.0
0.0
400.0
273.0
819.0
494.0
0.0
79012.0
0.0
231721.0
0.0
56690.0
41756.0
156693.0
35933.0
0.0
0.0
0.0
37550.0
115007.0
51311.0
151558.0
216287.0
59060.0

0.98
0.98
0.98
0.98
0.98
0.98
0.98
0.98
0.98
0.98
0.98
0.98
0.98
0.98
0.98
0.98
0.98
0.98
0.98
0.98
0.98
0.98
0.98
0.98
0.98
0.98
0.98
0.98
0.98
0.98


83419.0
666.0
82.0
86.0
0.0
167.0
30.0
0.0
0.0
0.0
325.0
446.0
0.0
0.0
0.0
0.0
0.0
0.0
0.0
0.0
35933.0
0.0
0.0
0.0
37550.0
0.0
0.0
0.0
0.0
8134.0


83419.0
666.0
82.0
86.0
0.0
167.0
30.0
0.0
0.0
0.0
325.0
446.0
0.0
0.0
0.0
0.0
0.0
0.0
0.0
0.0
35933.0
0.0
0.0
0.0
37550.0
0.0
0.0
0.0
0.0
8134.0

1.0
1.0
1.0
1.0
1.0
1.0
1.0
1.0
1.0
1.0
1.0
1.0
1.0
1.0
1.0
1.0
1.0
1.0
1.0
1.0
1.0
1.0
1.0
1.0
1.0
1.0
1.0
1.0
1.0
1.0


217684.0
161.0
76.0
0.0
498.0
166.0
0.0
0.0
0.0
23.0
473.0
48.0
0.0
0.0
0.0
42884.0
0.0
56689.0
41756.0
24309.0
0.0
0.0
0.0
0.0
0.0
30678.0
11731.0
0.0
3637.0
4555.0


217684.0
161.0
76.0
0.0
498.0
166.0
0.0
0.0
0.0
23.0
473.0
48.0
0.0
0.0
0.0
42884.0
0.0
56689.0
41756.0
24309.0
0.0
0.0
0.0
0.0
0.0
30678.0
11731.0
0.0
3637.0
4555.0

1.0
1.0
1.0
1.0
1.0
1.0
1.0
1.0
1.0
1.0
1.0
1.0
1.0
1.0
1.0
1.0
1.0
1.0
1.0
1.0
1.0
1.0
1.0
1.0
1.0
1.0
1.0
1.0
1.0
1.0


214607.0
0.0
0.0
471.0
318.0
0.0
0.0
0.0
0.0
0.0
0.0
0.0
0.0
79012.0
0.0
0.0
0.0
1.0
0.0
55786.0
0.0
0.0
0.0
0.0
0.0
23738.0
12822.0
26469.0
15990.0
0.0


53339.0
0.0
0.0
471.0
318.0
0.0
0.0
0.0
0.0
0.0
0.0
0.0
0.0
0.0
0.0
0.0
0.0
0.0
0.0
0.0
0.0
0.0
0.0
0.0
0.0
23738.0
12822.0
0.0
15990.0
0.0

1.0
1.0
1.0
1.0
1.0
1.0
1.0
1.0
1.0
1.0
1.0
1.0
1.0
1.0
1.0
1.0
1.0
1.0
1.0
1.0
1.0
1.0
1.0
1.0
1.0
1.0
1.0
1.0
1.0
1.0


134799.0
0.0
0.0
0.0
0.0
0.0
0.0
0.0
0.0
0.0
0.0
0.0
0.0
79012.0
0.0
0.0
0.0
1.0
0.0
55786.0
0.0
0.0
0.0
0.0
0.0
0.0
0.0
0.0
0.0
0.0

1.0
1.0
1.0
1.0
1.0
1.0
1.0
1.0
1.0
1.0
1.0
1.0
1.0
1.0
1.0
1.0
1.0
1.0
1.0
1.0
1.0
1.0
1.0
1.0
1.0
1.0
1.0
1.0
1.0
1.0


26469.0
0.0
0.0
0.0
0.0
0.0
0.0
0.0
0.0
0.0
0.0
0.0
0.0
0.0
0.0
0.0
0.0
0.0
0.0
0.0
0.0
0.0
0.0
0.0
0.0
0.0
0.0
26469.0
0.0
0.0

1.0
1.0
1.0
1.0
1.0
1.0
1.0
1.0
1.0
1.0
1.0
1.0
1.0
1.0
1.0
1.0
1.0
1.0
1.0
1.0
1.0
1.0
1.0
1.0
1.0
1.0
1.0
1.0
1.0
1.0


335.0
0.0
0.0
0.0
0.0
0.0
0.0
0.0
335.0
0.0
0.0
0.0
0.0
0.0
0.0
0.0
0.0
0.0
0.0
0.0
0.0
0.0
0.0
0.0
0.0
0.0
0.0
0.0
0.0
0.0


335.0
0.0
0.0
0.0
0.0
0.0
0.0
0.0
335.0
0.0
0.0
0.0
0.0
0.0
0.0
0.0
0.0
0.0
0.0
0.0
0.0
0.0
0.0
0.0
0.0
0.0
0.0
0.0
0.0
0.0

1.0
1.0
1.0
1.0
1.0
1.0
1.0
1.0
1.0
1.0
1.0
1.0
1.0
1.0
1.0
1.0
1.0
1.0
1.0
1.0
1.0
1.0
1.0
1.0
1.0
1.0
1.0
1.0
1.0
1.0


273.0
23.0
0.0
0.0
0.0
0.0
0.0
0.0
0.0
250.0
0.0
0.0
0.0
0.0
0.0
0.0
0.0
0.0
0.0
0.0
0.0
0.0
0.0
0.0
0.0
0.0
0.0
0.0
0.0
0.0


273.0
23.0
0.0
0.0
0.0
0.0
0.0
0.0
0.0
250.0
0.0
0.0
0.0
0.0
0.0
0.0
0.0
0.0
0.0
0.0
0.0
0.0
0.0
0.0
0.0
0.0
0.0
0.0
0.0
0.0

1.0
1.0
1.0
1.0
1.0
1.0
1.0
1.0
1.0
1.0
1.0
1.0
1.0
1.0
1.0
1.0
1.0
1.0
1.0
1.0
1.0
1.0
1.0
1.0
1.0
1.0
1.0
1.0
1.0
1.0


86.0
0.0
0.0
0.0
0.0
0.0
0.0
0.0
65.0
0.0
21.0
0.0
0.0
0.0
0.0
0.0
0.0
0.0
0.0
0.0
0.0
0.0
0.0
0.0
0.0
0.0
0.0
0.0
0.0
0.0


86.0
0.0
0.0
0.0
0.0
0.0
0.0
0.0
65.0
0.0
21.0
0.0
0.0
0.0
0.0
0.0
0.0
0.0
0.0
0.0
0.0
0.0
0.0
0.0
0.0
0.0
0.0
0.0
0.0
0.0

0.75
0.75
0.75
0.75
0.75
0.75
0.75
0.75
0.75
0.75
0.75
0.75
0.75
0.75
0.75
0.75
0.75
0.75
0.75
0.75
0.75
0.75
0.75
0.75
0.75
0.75
0.75
0.75
0.75
0.75


644365.0
0.0
0.0
26.0
33.0
0.0
0.0
0.0
0.0
0.0
0.0
0.0
0.0
0.0
0.0
188837.0
0.0
0.0
0.0
0.0
0.0
0.0
0.0
0.0
0.0
60591.0
26758.0
125089.0
196660.0
46371.0


644365.0
0.0
0.0
26.0
33.0
0.0
0.0
0.0
0.0
0.0
0.0
0.0
0.0
0.0
0.0
188837.0
0.0
0.0
0.0
0.0
0.0
0.0
0.0
0.0
0.0
60591.0
26758.0
125089.0
196660.0
46371.0

1.0
1.0
1.0
1.0
1.0
1.0
1.0
1.0
1.0
1.0
1.0
1.0
1.0
1.0
1.0
1.0
1.0
1.0
1.0
1.0
1.0
1.0
1.0
1.0
1.0
1.0
1.0
1.0
1.0
1.0


31.0
0.0
31.0
0.0
0.0
0.0
0.0
0.0
0.0
0.0
0.0
0.0
0.0
0.0
0.0
0.0
0.0
0.0
0.0
0.0
0.0
0.0
0.0
0.0
0.0
0.0
0.0
0.0
0.0
0.0


31.0
0.0
31.0
0.0
0.0
0.0
0.0
0.0
0.0
0.0
0.0
0.0
0.0
0.0
0.0
0.0
0.0
0.0
0.0
0.0
0.0
0.0
0.0
0.0
0.0
0.0
0.0
0.0
0.0
0.0

1.0
1.0
1.0
1.0
1.0
1.0
1.0
1.0
1.0
1.0
1.0
1.0
1.0
1.0
1.0
1.0
1.0
1.0
1.0
1.0
1.0
1.0
1.0
1.0
1.0
1.0
1.0
1.0
1.0
1.0


15.0
0.0
0.0
0.0
0.0
0.0
15.0
0.0
0.0
0.0
0.0
0.0
0.0
0.0
0.0
0.0
0.0
0.0
0.0
0.0
0.0
0.0
0.0
0.0
0.0
0.0
0.0
0.0
0.0
0.0


15.0
0.0
0.0
0.0
0.0
0.0
15.0
0.0
0.0
0.0
0.0
0.0
0.0
0.0
0.0
0.0
0.0
0.0
0.0
0.0
0.0
0.0
0.0
0.0
0.0
0.0
0.0
0.0
0.0
0.0

1.0
1.0
1.0
1.0
1.0
1.0
1.0
1.0
1.0
1.0
1.0
1.0
1.0
1.0
1.0
1.0
1.0
1.0
1.0
1.0
1.0
1.0
1.0
1.0
1.0
1.0
1.0
1.0
1.0
1.0


76577.0
0.0
0.0
0.0
0.0
0.0
0.0
0.0
0.0
0.0
0.0
0.0
0.0
0.0
0.0
0.0
0.0
0.0
0.0
76577.0
0.0
0.0
0.0
0.0
0.0
0.0
0.0
0.0
0.0
0.0


76577.0
0.0
0.0
0.0
0.0
0.0
0.0
0.0
0.0
0.0
0.0
0.0
0.0
0.0
0.0
0.0
0.0
0.0
0.0
76577.0
0.0
0.0
0.0
0.0
0.0
0.0
0.0
0.0
0.0
0.0

1.0
1.0
1.0
1.0
1.0
1.0
1.0
1.0
1.0
1.0
1.0
1.0
1.0
1.0
1.0
1.0
1.0
1.0
1.0
1.0
1.0
1.0
1.0
1.0
1.0
1.0
1.0
1.0
1.0
1.0


3905854.0
163.0
369.0
62.0
0.0
676.0
943.0
0.0
0.0
0.0
114.0
1470.0
0.0
52585.0
8786.0
254438.0
23551.0
0.0
0.0
0.0
0.0
0.0
10.0
0.0
154409.0
1513950.0
464562.0
1012615.0
327325.0
89826.0

0.98
0.98
0.98
0.98
0.98
0.98
0.98
0.98
0.98
0.98
0.98
0.98
0.98
0.98
0.98
0.98
0.98
0.98
0.98
0.98
0.98
0.98
0.98
0.98
0.98
0.98
0.98
0.98
0.98
0.98


84776.0
58.0
139.0
0.0
0.0
0.0
0.0
0.0
0.0
0.0
0.0
1053.0
0.0
0.0
0.0
0.0
0.0
0.0
0.0
0.0
0.0
0.0
0.0
0.0
83526.0
0.0
0.0
0.0
0.0
0.0


84776.0
58.0
139.0
0.0
0.0
0.0
0.0
0.0
0.0
0.0
0.0
1053.0
0.0
0.0
0.0
0.0
0.0
0.0
0.0
0.0
0.0
0.0
0.0
0.0
83526.0
0.0
0.0
0.0
0.0
0.0

1.0
1.0
1.0
1.0
1.0
1.0
1.0
1.0
1.0
1.0
1.0
1.0
1.0
1.0
1.0
1.0
1.0
1.0
1.0
1.0
1.0
1.0
1.0
1.0
1.0
1.0
1.0
1.0
1.0
1.0


1088848.0
89.0
222.0
19.0
0.0
56.0
916.0
0.0
0.0
0.0
0.0
352.0
0.0
52585.0
0.0
1.0
0.0
0.0
0.0
0.0
0.0
0.0
0.0
0.0
2.0
801650.0
130519.0
102437.0
0.0
0.0


1088848.0
89.0
222.0
19.0
0.0
56.0
916.0
0.0
0.0
0.0
0.0
352.0
0.0
52585.0
0.0
1.0
0.0
0.0
0.0
0.0
0.0
0.0
0.0
0.0
2.0
801650.0
130519.0
102437.0
0.0
0.0

1.0
1.0
1.0
1.0
1.0
1.0
1.0
1.0
1.0
1.0
1.0
1.0
1.0
1.0
1.0
1.0
1.0
1.0
1.0
1.0
1.0
1.0
1.0
1.0
1.0
1.0
1.0
1.0
1.0
1.0


11450.0
0.0
0.0
43.0
0.0
375.0
0.0
0.0
0.0
0.0
0.0
0.0
0.0
0.0
0.0
0.0
0.0
0.0
0.0
0.0
0.0
0.0
0.0
0.0
0.0
0.0
0.0
11032.0
0.0
0.0

1.0
1.0
1.0
1.0
1.0
1.0
1.0
1.0
1.0
1.0
1.0
1.0
1.0
1.0
1.0
1.0
1.0
1.0
1.0
1.0
1.0
1.0
1.0
1.0
1.0
1.0
1.0
1.0
1.0
1.0


324763.0
0.0
0.0
0.0
0.0
245.0
27.0
0.0
0.0
0.0
0.0
0.0
0.0
0.0
0.0
0.0
0.0
0.0
0.0
0.0
0.0
0.0
0.0
0.0
0.0
198175.0
18564.0
0.0
88811.0
18941.0


272.0
0.0
0.0
0.0
0.0
245.0
27.0
0.0
0.0
0.0
0.0
0.0
0.0
0.0
0.0
0.0
0.0
0.0
0.0
0.0
0.0
0.0
0.0
0.0
0.0
0.0
0.0
0.0
0.0
0.0

1.0
1.0
1.0
1.0
1.0
1.0
1.0
1.0
1.0
1.0
1.0
1.0
1.0
1.0
1.0
1.0
1.0
1.0
1.0
1.0
1.0
1.0
1.0
1.0
1.0
1.0
1.0
1.0
1.0
1.0


324491.0
0.0
0.0
0.0
0.0
0.0
0.0
0.0
0.0
0.0
0.0
0.0
0.0
0.0
0.0
0.0
0.0
0.0
0.0
0.0
0.0
0.0
0.0
0.0
0.0
198175.0
18564.0
0.0
88811.0
18941.0

1.0
1.0
1.0
1.0
1.0
1.0
1.0
1.0
1.0
1.0
1.0
1.0
1.0
1.0
1.0
1.0
1.0
1.0
1.0
1.0
1.0
1.0
1.0
1.0
1.0
1.0
1.0
1.0
1.0
1.0


407116.0
16.0
0.0
0.0
0.0
0.0
0.0
0.0
0.0
0.0
114.0
0.0
0.0
0.0
0.0
0.0
23551.0
0.0
0.0
0.0
0.0
0.0
0.0
0.0
0.0
307244.0
62329.0
1271.0
0.0
12591.0


383565.0
16.0
0.0
0.0
0.0
0.0
0.0
0.0
0.0
0.0
114.0
0.0
0.0
0.0
0.0
0.0
0.0
0.0
0.0
0.0
0.0
0.0
0.0
0.0
0.0
307244.0
62329.0
1271.0
0.0
12591.0

1.0
1.0
1.0
1.0
1.0
1.0
1.0
1.0
1.0
1.0
1.0
1.0
1.0
1.0
1.0
1.0
1.0
1.0
1.0
1.0
1.0
1.0
1.0
1.0
1.0
1.0
1.0
1.0
1.0
1.0


23551.0
0.0
0.0
0.0
0.0
0.0
0.0
0.0
0.0
0.0
0.0
0.0
0.0
0.0
0.0
0.0
23551.0
0.0
0.0
0.0
0.0
0.0
0.0
0.0
0.0
0.0
0.0
0.0
0.0
0.0

1.0
1.0
1.0
1.0
1.0
1.0
1.0
1.0
1.0
1.0
1.0
1.0
1.0
1.0
1.0
1.0
1.0
1.0
1.0
1.0
1.0
1.0
1.0
1.0
1.0
1.0
1.0
1.0
1.0
1.0


75373.0
0.0
8.0
0.0
0.0
0.0
0.0
0.0
0.0
0.0
0.0
65.0
0.0
0.0
0.0
75300.0
0.0
0.0
0.0
0.0
0.0
0.0
0.0
0.0
0.0
0.0
0.0
0.0
0.0
0.0


75373.0
0.0
8.0
0.0
0.0
0.0
0.0
0.0
0.0
0.0
0.0
65.0
0.0
0.0
0.0
75300.0
0.0
0.0
0.0
0.0
0.0
0.0
0.0
0.0
0.0
0.0
0.0
0.0
0.0
0.0

1.0
1.0
1.0
1.0
1.0
1.0
1.0
1.0
1.0
1.0
1.0
1.0
1.0
1.0
1.0
1.0
1.0
1.0
1.0
1.0
1.0
1.0
1.0
1.0
1.0
1.0
1.0
1.0
1.0
1.0


747058.0
0.0
0.0
0.0
0.0
0.0
0.0
0.0
0.0
0.0
0.0
0.0
0.0
0.0
0.0
25178.0
0.0
0.0
0.0
0.0
0.0
0.0
0.0
0.0
0.0
70851.0
80252.0
484420.0
86357.0
0.0


747058.0
0.0
0.0
0.0
0.0
0.0
0.0
0.0
0.0
0.0
0.0
0.0
0.0
0.0
0.0
25178.0
0.0
0.0
0.0
0.0
0.0
0.0
0.0
0.0
0.0
70851.0
80252.0
484420.0
86357.0
0.0

0.93
0.93
0.93
0.93
0.93
0.93
0.93
0.93
0.93
0.93
0.93
0.93
0.93
0.93
0.93
0.93
0.93
0.93
0.93
0.93
0.93
0.93
0.93
0.93
0.93
0.93
0.93
0.93
0.93
0.93


737401.0
0.0
0.0
0.0
0.0
0.0
0.0
0.0
0.0
0.0
0.0
0.0
0.0
0.0
0.0
0.0
0.0
0.0
0.0
0.0
0.0
0.0
0.0
0.0
0.0
136026.0
169973.0
346644.0
84758.0
0.0

1.0
1.0
1.0
1.0
1.0
1.0
1.0
1.0
1.0
1.0
1.0
1.0
1.0
1.0
1.0
1.0
1.0
1.0
1.0
1.0
1.0
1.0
1.0
1.0
1.0
1.0
1.0
1.0
1.0
1.0


217452.0
0.0
0.0
0.0
0.0
0.0
0.0
0.0
0.0
0.0
0.0
0.0
0.0
0.0
0.0
142817.0
0.0
0.0
0.0
0.0
0.0
0.0
10.0
0.0
0.0
0.0
0.0
66663.0
7962.0
0.0

1.0
1.0
1.0
1.0
1.0
1.0
1.0
1.0
1.0
1.0
1.0
1.0
1.0
1.0
1.0
1.0
1.0
1.0
1.0
1.0
1.0
1.0
1.0
1.0
1.0
1.0
1.0
1.0
1.0
1.0


166758.0
0.0
0.0
0.0
0.0
0.0
0.0
0.0
0.0
0.0
0.0
0.0
0.0
0.0
0.0
142278.0
0.0
0.0
0.0
0.0
0.0
0.0
10.0
0.0
0.0
0.0
0.0
16575.0
7895.0
0.0

1.0
1.0
1.0
1.0
1.0
1.0
1.0
1.0
1.0
1.0
1.0
1.0
1.0
1.0
1.0
1.0
1.0
1.0
1.0
1.0
1.0
1.0
1.0
1.0
1.0
1.0
1.0
1.0
1.0
1.0


204.0
0.0
0.0
0.0
0.0
0.0
0.0
0.0
0.0
0.0
0.0
0.0
0.0
0.0
0.0
190.0
0.0
0.0
0.0
0.0
0.0
0.0
0.0
0.0
0.0
0.0
0.0
5.0
9.0
0.0

0.75
0.75
0.75
0.75
0.75
0.75
0.75
0.75
0.75
0.75
0.75
0.75
0.75
0.75
0.75
0.75
0.75
0.75
0.75
0.75
0.75
0.75
0.75
0.75
0.75
0.75
0.75
0.75
0.75
0.75


70881.0
0.0
0.0
0.0
0.0
0.0
0.0
0.0
0.0
0.0
0.0
0.0
0.0
0.0
0.0
0.0
0.0
0.0
0.0
0.0
0.0
0.0
0.0
0.0
70881.0
0.0
0.0
0.0
0.0
0.0


70881.0
0.0
0.0
0.0
0.0
0.0
0.0
0.0
0.0
0.0
0.0
0.0
0.0
0.0
0.0
0.0
0.0
0.0
0.0
0.0
0.0
0.0
0.0
0.0
70881.0
0.0
0.0
0.0
0.0
0.0

1.0
1.0
1.0
1.0
1.0
1.0
1.0
1.0
1.0
1.0
1.0
1.0
1.0
1.0
1.0
1.0
1.0
1.0
1.0
1.0
1.0
1.0
1.0
1.0
1.0
1.0
1.0
1.0
1.0
1.0


59437.0
0.0
0.0
0.0
0.0
0.0
0.0
0.0
0.0
0.0
0.0
0.0
0.0
0.0
0.0
0.0
0.0
0.0
0.0
0.0
0.0
0.0
0.0
0.0
0.0
0.0
0.0
0.0
59437.0
0.0


59437.0
0.0
0.0
0.0
0.0
0.0
0.0
0.0
0.0
0.0
0.0
0.0
0.0
0.0
0.0
0.0
0.0
0.0
0.0
0.0
0.0
0.0
0.0
0.0
0.0
0.0
0.0
0.0
59437.0
0.0

1.0
1.0
1.0
1.0
1.0
1.0
1.0
1.0
1.0
1.0
1.0
1.0
1.0
1.0
1.0
1.0
1.0
1.0
1.0
1.0
1.0
1.0
1.0
1.0
1.0
1.0
1.0
1.0
1.0
1.0


70110.0
0.0
0.0
0.0
0.0
0.0
0.0
0.0
0.0
0.0
0.0
0.0
0.0
0.0
0.0
10601.0
0.0
0.0
0.0
0.0
0.0
0.0
0.0
0.0
0.0
0.0
2341.0
0.0
0.0
57168.0

0.84
0.84
0.84
0.84
0.84
0.84
0.84
0.84
0.84
0.84
0.84
0.84
0.84
0.84
0.84
0.84
0.84
0.84
0.84
0.84
0.84
0.84
0.84
0.84
0.84
0.84
0.84
0.84
0.84
0.84


69845.0
0.0
0.0
0.0
0.0
0.0
0.0
0.0
0.0
0.0
0.0
0.0
0.0
0.0
0.0
10578.0
0.0
0.0
0.0
0.0
0.0
0.0
0.0
0.0
0.0
0.0
2308.0
0.0
0.0
56959.0

1.0
1.0
1.0
1.0
1.0
1.0
1.0
1.0
1.0
1.0
1.0
1.0
1.0
1.0
1.0
1.0
1.0
1.0
1.0
1.0
1.0
1.0
1.0
1.0
1.0
1.0
1.0
1.0
1.0
1.0


163.0
0.0
0.0
0.0
0.0
0.0
0.0
0.0
0.0
0.0
0.0
0.0
0.0
0.0
0.0
23.0
0.0
0.0
0.0
0.0
0.0
0.0
0.0
0.0
0.0
0.0
0.0
0.0
0.0
140.0

0.8
0.8
0.8
0.8
0.8
0.8
0.8
0.8
0.8
0.8
0.8
0.8
0.8
0.8
0.8
0.8
0.8
0.8
0.8
0.8
0.8
0.8
0.8
0.8
0.8
0.8
0.8
0.8
0.8
0.8


8786.0
0.0
0.0
0.0
0.0
0.0
0.0
0.0
0.0
0.0
0.0
0.0
0.0
0.0
8786.0
0.0
0.0
0.0
0.0
0.0
0.0
0.0
0.0
0.0
0.0
0.0
0.0
0.0
0.0
0.0


8786.0
0.0
0.0
0.0
0.0
0.0
0.0
0.0
0.0
0.0
0.0
0.0
0.0
0.0
8786.0
0.0
0.0
0.0
0.0
0.0
0.0
0.0
0.0
0.0
0.0
0.0
0.0
0.0
0.0
0.0

1.0
1.0
1.0
1.0
1.0
1.0
1.0
1.0
1.0
1.0
1.0
1.0
1.0
1.0
1.0
1.0
1.0
1.0
1.0
1.0
1.0
1.0
1.0
1.0
1.0
1.0
1.0
1.0
1.0
1.0


1871469.0
40.0
568.0
118.0
499.0
0.0
0.0
0.0
0.0
0.0
0.0
0.0
0.0
220535.0
370227.0
284985.0
396497.0
239397.0
0.0
0.0
0.0
0.0
0.0
0.0
173598.0
0.0
0.0
0.0
132440.0
52565.0


765585.0
40.0
568.0
118.0
499.0
0.0
0.0
0.0
0.0
0.0
0.0
0.0
0.0
220535.0
370227.0
0.0
0.0
0.0
0.0
0.0
0.0
0.0
0.0
0.0
173598.0
0.0
0.0
0.0
0.0
0.0


765585.0
40.0
568.0
118.0
499.0
0.0
0.0
0.0
0.0
0.0
0.0
0.0
0.0
220535.0
370227.0
0.0
0.0
0.0
0.0
0.0
0.0
0.0
0.0
0.0
173598.0
0.0
0.0
0.0
0.0
0.0

0.99
0.99
0.99
0.99
0.99
0.99
0.99
0.99
0.99
0.99
0.99
0.99
0.99
0.99
0.99
0.99
0.99
0.99
0.99
0.99
0.99
0.99
0.99
0.99
0.99
0.99
0.99
0.99
0.99
0.99


1105884.0
0.0
0.0
0.0
0.0
0.0
0.0
0.0
0.0
0.0
0.0
0.0
0.0
0.0
0.0
284985.0
396497.0
239397.0
0.0
0.0
0.0
0.0
0.0
0.0
0.0
0.0
0.0
0.0
132440.0
52565.0

1.0
1.0
1.0
1.0
1.0
1.0
1.0
1.0
1.0
1.0
1.0
1.0
1.0
1.0
1.0
1.0
1.0
1.0
1.0
1.0
1.0
1.0
1.0
1.0
1.0
1.0
1.0
1.0
1.0
1.0


1088582.0
0.0
0.0
0.0
0.0
0.0
0.0
0.0
0.0
0.0
0.0
0.0
0.0
0.0
0.0
276835.0
392175.0
237213.0
0.0
0.0
0.0
0.0
0.0
0.0
0.0
0.0
0.0
0.0
130511.0
51848.0

1.0
1.0
1.0
1.0
1.0
1.0
1.0
1.0
1.0
1.0
1.0
1.0
1.0
1.0
1.0
1.0
1.0
1.0
1.0
1.0
1.0
1.0
1.0
1.0
1.0
1.0
1.0
1.0
1.0
1.0


6794.0
0.0
0.0
41.0
486.0
532.0
101.0
0.0
0.0
0.0
0.0
0.0
0.0
0.0
2155.0
0.0
0.0
0.0
0.0
0.0
0.0
0.0
0.0
0.0
3479.0
0.0
0.0
0.0
0.0
0.0


6794.0
0.0
0.0
41.0
486.0
532.0
101.0
0.0
0.0
0.0
0.0
0.0
0.0
0.0
2155.0
0.0
0.0
0.0
0.0
0.0
0.0
0.0
0.0
0.0
3479.0
0.0
0.0
0.0
0.0
0.0


6794.0
0.0
0.0
41.0
486.0
532.0
101.0
0.0
0.0
0.0
0.0
0.0
0.0
0.0
2155.0
0.0
0.0
0.0
0.0
0.0
0.0
0.0
0.0
0.0
3479.0
0.0
0.0
0.0
0.0
0.0

0.99
0.99
0.99
0.99
0.99
0.99
0.99
0.99
0.99
0.99
0.99
0.99
0.99
0.99
0.99
0.99
0.99
0.99
0.99
0.99
0.99
0.99
0.99
0.99
0.99
0.99
0.99
0.99
0.99
0.99


421597.0
0.0
97.0
0.0
25.0
48.0
697.0
0.0
227.0
90.0
34.0
47.0
0.0
0.0
113.0
275365.0
93203.0
36275.0
0.0
0.0
0.0
0.0
0.0
0.0
0.0
0.0
13983.0
1393.0
0.0
0.0


725.0
0.0
3.0
0.0
25.0
0.0
697.0
0.0
0.0
0.0
0.0
0.0
0.0
0.0
0.0
0.0
0.0
0.0
0.0
0.0
0.0
0.0
0.0
0.0
0.0
0.0
0.0
0.0
0.0
0.0


725.0
0.0
3.0
0.0
25.0
0.0
697.0
0.0
0.0
0.0
0.0
0.0
0.0
0.0
0.0
0.0
0.0
0.0
0.0
0.0
0.0
0.0
0.0
0.0
0.0
0.0
0.0
0.0
0.0
0.0

0.76
0.76
0.76
0.76
0.76
0.76
0.76
0.76
0.76
0.76
0.76
0.76
0.76
0.76
0.76
0.76
0.76
0.76
0.76
0.76
0.76
0.76
0.76
0.76
0.76
0.76
0.76
0.76
0.76
0.76


202.0
0.0
0.0
0.0
0.0
0.0
0.0
0.0
112.0
90.0
0.0
0.0
0.0
0.0
0.0
0.0
0.0
0.0
0.0
0.0
0.0
0.0
0.0
0.0
0.0
0.0
0.0
0.0
0.0
0.0


202.0
0.0
0.0
0.0
0.0
0.0
0.0
0.0
112.0
90.0
0.0
0.0
0.0
0.0
0.0
0.0
0.0
0.0
0.0
0.0
0.0
0.0
0.0
0.0
0.0
0.0
0.0
0.0
0.0
0.0

1.0
1.0
1.0
1.0
1.0
1.0
1.0
1.0
1.0
1.0
1.0
1.0
1.0
1.0
1.0
1.0
1.0
1.0
1.0
1.0
1.0
1.0
1.0
1.0
1.0
1.0
1.0
1.0
1.0
1.0


149.0
0.0
0.0
0.0
0.0
0.0
0.0
0.0
115.0
0.0
34.0
0.0
0.0
0.0
0.0
0.0
0.0
0.0
0.0
0.0
0.0
0.0
0.0
0.0
0.0
0.0
0.0
0.0
0.0
0.0


149.0
0.0
0.0
0.0
0.0
0.0
0.0
0.0
115.0
0.0
34.0
0.0
0.0
0.0
0.0
0.0
0.0
0.0
0.0
0.0
0.0
0.0
0.0
0.0
0.0
0.0
0.0
0.0
0.0
0.0

1.0
1.0
1.0
1.0
1.0
1.0
1.0
1.0
1.0
1.0
1.0
1.0
1.0
1.0
1.0
1.0
1.0
1.0
1.0
1.0
1.0
1.0
1.0
1.0
1.0
1.0
1.0
1.0
1.0
1.0


141.0
0.0
94.0
0.0
0.0
0.0
0.0
0.0
0.0
0.0
0.0
47.0
0.0
0.0
0.0
0.0
0.0
0.0
0.0
0.0
0.0
0.0
0.0
0.0
0.0
0.0
0.0
0.0
0.0
0.0


141.0
0.0
94.0
0.0
0.0
0.0
0.0
0.0
0.0
0.0
0.0
47.0
0.0
0.0
0.0
0.0
0.0
0.0
0.0
0.0
0.0
0.0
0.0
0.0
0.0
0.0
0.0
0.0
0.0
0.0

1.0
1.0
1.0
1.0
1.0
1.0
1.0
1.0
1.0
1.0
1.0
1.0
1.0
1.0
1.0
1.0
1.0
1.0
1.0
1.0
1.0
1.0
1.0
1.0
1.0
1.0
1.0
1.0
1.0
1.0


4037.0
0.0
0.0
0.0
0.0
48.0
0.0
0.0
0.0
0.0
0.0
0.0
0.0
0.0
0.0
2596.0
0.0
0.0
0.0
0.0
0.0
0.0
0.0
0.0
0.0
0.0
0.0
1393.0
0.0
0.0


4037.0
0.0
0.0
0.0
0.0
48.0
0.0
0.0
0.0
0.0
0.0
0.0
0.0
0.0
0.0
2596.0
0.0
0.0
0.0
0.0
0.0
0.0
0.0
0.0
0.0
0.0
0.0
1393.0
0.0
0.0

1.0
1.0
1.0
1.0
1.0
1.0
1.0
1.0
1.0
1.0
1.0
1.0
1.0
1.0
1.0
1.0
1.0
1.0
1.0
1.0
1.0
1.0
1.0
1.0
1.0
1.0
1.0
1.0
1.0
1.0


402247.0
0.0
0.0
0.0
0.0
0.0
0.0
0.0
0.0
0.0
0.0
0.0
0.0
0.0
0.0
272769.0
93203.0
36275.0
0.0
0.0
0.0
0.0
0.0
0.0
0.0
0.0
0.0
0.0
0.0
0.0


402247.0
0.0
0.0
0.0
0.0
0.0
0.0
0.0
0.0
0.0
0.0
0.0
0.0
0.0
0.0
272769.0
93203.0
36275.0
0.0
0.0
0.0
0.0
0.0
0.0
0.0
0.0
0.0
0.0
0.0
0.0

1.0
1.0
1.0
1.0
1.0
1.0
1.0
1.0
1.0
1.0
1.0
1.0
1.0
1.0
1.0
1.0
1.0
1.0
1.0
1.0
1.0
1.0
1.0
1.0
1.0
1.0
1.0
1.0
1.0
1.0


13983.0
0.0
0.0
0.0
0.0
0.0
0.0
0.0
0.0
0.0
0.0
0.0
0.0
0.0
0.0
0.0
0.0
0.0
0.0
0.0
0.0
0.0
0.0
0.0
0.0
0.0
13983.0
0.0
0.0
0.0


13983.0
0.0
0.0
0.0
0.0
0.0
0.0
0.0
0.0
0.0
0.0
0.0
0.0
0.0
0.0
0.0
0.0
0.0
0.0
0.0
0.0
0.0
0.0
0.0
0.0
0.0
13983.0
0.0
0.0
0.0

1.0
1.0
1.0
1.0
1.0
1.0
1.0
1.0
1.0
1.0
1.0
1.0
1.0
1.0
1.0
1.0
1.0
1.0
1.0
1.0
1.0
1.0
1.0
1.0
1.0
1.0
1.0
1.0
1.0
1.0


113.0
0.0
0.0
0.0
0.0
0.0
0.0
0.0
0.0
0.0
0.0
0.0
0.0
0.0
113.0
0.0
0.0
0.0
0.0
0.0
0.0
0.0
0.0
0.0
0.0
0.0
0.0
0.0
0.0
0.0

0.97
0.97
0.97
0.97
0.97
0.97
0.97
0.97
0.97
0.97
0.97
0.97
0.97
0.97
0.97
0.97
0.97
0.97
0.97
0.97
0.97
0.97
0.97
0.97
0.97
0.97
0.97
0.97
0.97
0.97


1349583.0
60.0
20.0
0.0
0.0
0.0
0.0
0.0
42.0
82.0
130.0
267.0
0.0
0.0
0.0
108947.0
15802.0
1.0
0.0
1.0
0.0
0.0
0.0
0.0
0.0
202788.0
493858.0
307778.0
118076.0
101731.0


1349583.0
60.0
20.0
0.0
0.0
0.0
0.0
0.0
42.0
82.0
130.0
267.0
0.0
0.0
0.0
108947.0
15802.0
1.0
0.0
1.0
0.0
0.0
0.0
0.0
0.0
202788.0
493858.0
307778.0
118076.0
101731.0

1.0
1.0
1.0
1.0
1.0
1.0
1.0
1.0
1.0
1.0
1.0
1.0
1.0
1.0
1.0
1.0
1.0
1.0
1.0
1.0
1.0
1.0
1.0
1.0
1.0
1.0
1.0
1.0
1.0
1.0


601.0
60.0
20.0
0.0
0.0
0.0
0.0
0.0
42.0
82.0
130.0
267.0
0.0
0.0
0.0
0.0
0.0
0.0
0.0
0.0
0.0
0.0
0.0
0.0
0.0
0.0
0.0
0.0
0.0
0.0

0.85
0.85
0.85
0.85
0.85
0.85
0.85
0.85
0.85
0.85
0.85
0.85
0.85
0.85
0.85
0.85
0.85
0.85
0.85
0.85
0.85
0.85
0.85
0.85
0.85
0.85
0.85
0.85
0.85
0.85


25530.0
51.0
25.0
84.0
52.0
18.0
128.0
0.0
0.0
0.0
0.0
21.0
0.0
0.0
0.0
0.0
0.0
11859.0
0.0
0.0
0.0
0.0
0.0
0.0
13292.0
0.0
0.0
0.0
0.0
0.0


25530.0
51.0
25.0
84.0
52.0
18.0
128.0
0.0
0.0
0.0
0.0
21.0
0.0
0.0
0.0
0.0
0.0
11859.0
0.0
0.0
0.0
0.0
0.0
0.0
13292.0
0.0
0.0
0.0
0.0
0.0


25530.0
51.0
25.0
84.0
52.0
18.0
128.0
0.0
0.0
0.0
0.0
21.0
0.0
0.0
0.0
0.0
0.0
11859.0
0.0
0.0
0.0
0.0
0.0
0.0
13292.0
0.0
0.0
0.0
0.0
0.0

1.0
1.0
1.0
1.0
1.0
1.0
1.0
1.0
1.0
1.0
1.0
1.0
1.0
1.0
1.0
1.0
1.0
1.0
1.0
1.0
1.0
1.0
1.0
1.0
1.0
1.0
1.0
1.0
1.0
1.0


931130.0
12.0
58.0
19.0
0.0
183.0
0.0
85.0
303.0
94.0
319.0
89.0
0.0
9322.0
6960.0
318512.0
2576.0
104270.0
64275.0
0.0
36686.0
7223.0
0.0
0.0
0.0
104540.0
118026.0
71965.0
82954.0
2659.0

0.71
0.71
0.71
0.71
0.71
0.71
0.71
0.71
0.71
0.71
0.71
0.71
0.71
0.71
0.71
0.71
0.71
0.71
0.71
0.71
0.71
0.71
0.71
0.71
0.71
0.71
0.71
0.71
0.71
0.71


96642.0
0.0
0.0
0.0
0.0
0.0
0.0
0.0
0.0
0.0
255.0
0.0
0.0
0.0
0.0
0.0
0.0
96387.0
0.0
0.0
0.0
0.0
0.0
0.0
0.0
0.0
0.0
0.0
0.0
0.0


96642.0
0.0
0.0
0.0
0.0
0.0
0.0
0.0
0.0
0.0
255.0
0.0
0.0
0.0
0.0
0.0
0.0
96387.0
0.0
0.0
0.0
0.0
0.0
0.0
0.0
0.0
0.0
0.0
0.0
0.0

0.99
0.99
0.99
0.99
0.99
0.99
0.99
0.99
0.99
0.99
0.99
0.99
0.99
0.99
0.99
0.99
0.99
0.99
0.99
0.99
0.99
0.99
0.99
0.99
0.99
0.99
0.99
0.99
0.99
0.99


207.0
0.0
0.0
0.0
0.0
0.0
0.0
0.0
121.0
86.0
0.0
0.0
0.0
0.0
0.0
0.0
0.0
0.0
0.0
0.0
0.0
0.0
0.0
0.0
0.0
0.0
0.0
0.0
0.0
0.0


207.0
0.0
0.0
0.0
0.0
0.0
0.0
0.0
121.0
86.0
0.0
0.0
0.0
0.0
0.0
0.0
0.0
0.0
0.0
0.0
0.0
0.0
0.0
0.0
0.0
0.0
0.0
0.0
0.0
0.0

1.0
1.0
1.0
1.0
1.0
1.0
1.0
1.0
1.0
1.0
1.0
1.0
1.0
1.0
1.0
1.0
1.0
1.0
1.0
1.0
1.0
1.0
1.0
1.0
1.0
1.0
1.0
1.0
1.0
1.0


180.0
0.0
0.0
0.0
0.0
0.0
0.0
0.0
180.0
0.0
0.0
0.0
0.0
0.0
0.0
0.0
0.0
0.0
0.0
0.0
0.0
0.0
0.0
0.0
0.0
0.0
0.0
0.0
0.0
0.0


180.0
0.0
0.0
0.0
0.0
0.0
0.0
0.0
180.0
0.0
0.0
0.0
0.0
0.0
0.0
0.0
0.0
0.0
0.0
0.0
0.0
0.0
0.0
0.0
0.0
0.0
0.0
0.0
0.0
0.0

1.0
1.0
1.0
1.0
1.0
1.0
1.0
1.0
1.0
1.0
1.0
1.0
1.0
1.0
1.0
1.0
1.0
1.0
1.0
1.0
1.0
1.0
1.0
1.0
1.0
1.0
1.0
1.0
1.0
1.0


280863.0
0.0
0.0
0.0
0.0
183.0
0.0
0.0
0.0
0.0
0.0
0.0
0.0
0.0
0.0
0.0
0.0
0.0
0.0
0.0
0.0
0.0
0.0
0.0
0.0
73150.0
66446.0
63824.0
74634.0
2626.0


280863.0
0.0
0.0
0.0
0.0
183.0
0.0
0.0
0.0
0.0
0.0
0.0
0.0
0.0
0.0
0.0
0.0
0.0
0.0
0.0
0.0
0.0
0.0
0.0
0.0
73150.0
66446.0
63824.0
74634.0
2626.0

1.0
1.0
1.0
1.0
1.0
1.0
1.0
1.0
1.0
1.0
1.0
1.0
1.0
1.0
1.0
1.0
1.0
1.0
1.0
1.0
1.0
1.0
1.0
1.0
1.0
1.0
1.0
1.0
1.0
1.0


30848.0
0.0
16.0
19.0
0.0
0.0
0.0
0.0
0.0
0.0
61.0
0.0
0.0
0.0
0.0
0.0
0.0
0.0
0.0
0.0
0.0
0.0
0.0
0.0
0.0
30752.0
0.0
0.0
0.0
0.0

1.0
1.0
1.0
1.0
1.0
1.0
1.0
1.0
1.0
1.0
1.0
1.0
1.0
1.0
1.0
1.0
1.0
1.0
1.0
1.0
1.0
1.0
1.0
1.0
1.0
1.0
1.0
1.0
1.0
1.0


122.0
0.0
28.0
0.0
0.0
0.0
0.0
85.0
0.0
0.0
0.0
9.0
0.0
0.0
0.0
0.0
0.0
0.0
0.0
0.0
0.0
0.0
0.0
0.0
0.0
0.0
0.0
0.0
0.0
0.0


122.0
0.0
28.0
0.0
0.0
0.0
0.0
85.0
0.0
0.0
0.0
9.0
0.0
0.0
0.0
0.0
0.0
0.0
0.0
0.0
0.0
0.0
0.0
0.0
0.0
0.0
0.0
0.0
0.0
0.0

1.0
1.0
1.0
1.0
1.0
1.0
1.0
1.0
1.0
1.0
1.0
1.0
1.0
1.0
1.0
1.0
1.0
1.0
1.0
1.0
1.0
1.0
1.0
1.0
1.0
1.0
1.0
1.0
1.0
1.0


176733.0
0.0
14.0
0.0
0.0
0.0
0.0
0.0
0.0
0.0
0.0
76.0
0.0
9322.0
6960.0
89544.0
2575.0
7883.0
0.0
0.0
0.0
0.0
0.0
0.0
0.0
638.0
51580.0
8141.0
0.0
0.0


47309.0
0.0
0.0
0.0
0.0
0.0
0.0
0.0
0.0
0.0
0.0
76.0
0.0
0.0
0.0
47233.0
0.0
0.0
0.0
0.0
0.0
0.0
0.0
0.0
0.0
0.0
0.0
0.0
0.0
0.0

1.0
1.0
1.0
1.0
1.0
1.0
1.0
1.0
1.0
1.0
1.0
1.0
1.0
1.0
1.0
1.0
1.0
1.0
1.0
1.0
1.0
1.0
1.0
1.0
1.0
1.0
1.0
1.0
1.0
1.0


2483.0
0.0
14.0
0.0
0.0
0.0
0.0
0.0
0.0
0.0
0.0
0.0
0.0
0.0
2469.0
0.0
0.0
0.0
0.0
0.0
0.0
0.0
0.0
0.0
0.0
0.0
0.0
0.0
0.0
0.0

1.0
1.0
1.0
1.0
1.0
1.0
1.0
1.0
1.0
1.0
1.0
1.0
1.0
1.0
1.0
1.0
1.0
1.0
1.0
1.0
1.0
1.0
1.0
1.0
1.0
1.0
1.0
1.0
1.0
1.0


91212.0
0.0
0.0
0.0
0.0
0.0
0.0
0.0
0.0
0.0
0.0
0.0
0.0
0.0
0.0
30853.0
0.0
0.0
0.0
0.0
0.0
0.0
0.0
0.0
0.0
638.0
51580.0
8141.0
0.0
0.0

1.0
1.0
1.0
1.0
1.0
1.0
1.0
1.0
1.0
1.0
1.0
1.0
1.0
1.0
1.0
1.0
1.0
1.0
1.0
1.0
1.0
1.0
1.0
1.0
1.0
1.0
1.0
1.0
1.0
1.0


35729.0
0.0
0.0
0.0
0.0
0.0
0.0
0.0
0.0
0.0
0.0
0.0
0.0
9322.0
4491.0
11458.0
2575.0
7883.0
0.0
0.0
0.0
0.0
0.0
0.0
0.0
0.0
0.0
0.0
0.0
0.0

1.0
1.0
1.0
1.0
1.0
1.0
1.0
1.0
1.0
1.0
1.0
1.0
1.0
1.0
1.0
1.0
1.0
1.0
1.0
1.0
1.0
1.0
1.0
1.0
1.0
1.0
1.0
1.0
1.0
1.0


12.0
12.0
0.0
0.0
0.0
0.0
0.0
0.0
0.0
0.0
0.0
0.0
0.0
0.0
0.0
0.0
0.0
0.0
0.0
0.0
0.0
0.0
0.0
0.0
0.0
0.0
0.0
0.0
0.0
0.0


12.0
12.0
0.0
0.0
0.0
0.0
0.0
0.0
0.0
0.0
0.0
0.0
0.0
0.0
0.0
0.0
0.0
0.0
0.0
0.0
0.0
0.0
0.0
0.0
0.0
0.0
0.0
0.0
0.0
0.0

1.0
1.0
1.0
1.0
1.0
1.0
1.0
1.0
1.0
1.0
1.0
1.0
1.0
1.0
1.0
1.0
1.0
1.0
1.0
1.0
1.0
1.0
1.0
1.0
1.0
1.0
1.0
1.0
1.0
1.0


228969.0
0.0
0.0
0.0
0.0
0.0
0.0
0.0
0.0
0.0
0.0
0.0
0.0
0.0
0.0
228968.0
1.0
0.0
0.0
0.0
0.0
0.0
0.0
0.0
0.0
0.0
0.0
0.0
0.0
0.0


228969.0
0.0
0.0
0.0
0.0
0.0
0.0
0.0
0.0
0.0
0.0
0.0
0.0
0.0
0.0
228968.0
1.0
0.0
0.0
0.0
0.0
0.0
0.0
0.0
0.0
0.0
0.0
0.0
0.0
0.0

1.0
1.0
1.0
1.0
1.0
1.0
1.0
1.0
1.0
1.0
1.0
1.0
1.0
1.0
1.0
1.0
1.0
1.0
1.0
1.0
1.0
1.0
1.0
1.0
1.0
1.0
1.0
1.0
1.0
1.0


108184.0
0.0
0.0
0.0
0.0
0.0
0.0
0.0
0.0
0.0
0.0
0.0
0.0
0.0
0.0
0.0
0.0
0.0
64275.0
0.0
36686.0
7223.0
0.0
0.0
0.0
0.0
0.0
0.0
0.0
0.0


64275.0
0.0
0.0
0.0
0.0
0.0
0.0
0.0
0.0
0.0
0.0
0.0
0.0
0.0
0.0
0.0
0.0
0.0
64275.0
0.0
0.0
0.0
0.0
0.0
0.0
0.0
0.0
0.0
0.0
0.0

1.0
1.0
1.0
1.0
1.0
1.0
1.0
1.0
1.0
1.0
1.0
1.0
1.0
1.0
1.0
1.0
1.0
1.0
1.0
1.0
1.0
1.0
1.0
1.0
1.0
1.0
1.0
1.0
1.0
1.0


43909.0
0.0
0.0
0.0
0.0
0.0
0.0
0.0
0.0
0.0
0.0
0.0
0.0
0.0
0.0
0.0
0.0
0.0
0.0
0.0
36686.0
7223.0
0.0
0.0
0.0
0.0
0.0
0.0
0.0
0.0

1.0
1.0
1.0
1.0
1.0
1.0
1.0
1.0
1.0
1.0
1.0
1.0
1.0
1.0
1.0
1.0
1.0
1.0
1.0
1.0
1.0
1.0
1.0
1.0
1.0
1.0
1.0
1.0
1.0
1.0


8320.0
0.0
0.0
0.0
0.0
0.0
0.0
0.0
0.0
0.0
0.0
0.0
0.0
0.0
0.0
0.0
0.0
0.0
0.0
0.0
0.0
0.0
0.0
0.0
0.0
0.0
0.0
0.0
8320.0
0.0


8320.0
0.0
0.0
0.0
0.0
0.0
0.0
0.0
0.0
0.0
0.0
0.0
0.0
0.0
0.0
0.0
0.0
0.0
0.0
0.0
0.0
0.0
0.0
0.0
0.0
0.0
0.0
0.0
8320.0
0.0

1.0
1.0
1.0
1.0
1.0
1.0
1.0
1.0
1.0
1.0
1.0
1.0
1.0
1.0
1.0
1.0
1.0
1.0
1.0
1.0
1.0
1.0
1.0
1.0
1.0
1.0
1.0
1.0
1.0
1.0


33.0
0.0
0.0
0.0
0.0
0.0
0.0
0.0
0.0
0.0
0.0
0.0
0.0
0.0
0.0
0.0
0.0
0.0
0.0
0.0
0.0
0.0
0.0
0.0
0.0
0.0
0.0
0.0
0.0
33.0


33.0
0.0
0.0
0.0
0.0
0.0
0.0
0.0
0.0
0.0
0.0
0.0
0.0
0.0
0.0
0.0
0.0
0.0
0.0
0.0
0.0
0.0
0.0
0.0
0.0
0.0
0.0
0.0
0.0
33.0

1.0
1.0
1.0
1.0
1.0
1.0
1.0
1.0
1.0
1.0
1.0
1.0
1.0
1.0
1.0
1.0
1.0
1.0
1.0
1.0
1.0
1.0
1.0
1.0
1.0
1.0
1.0
1.0
1.0
1.0


104125.0
32.0
11.0
20.0
0.0
0.0
4.0
124.0
112.0
135.0
134.0
31.0
0.0
0.0
0.0
4949.0
0.0
0.0
1414.0
7612.0
12632.0
0.0
0.0
0.0
5706.0
0.0
4257.0
28364.0
38588.0
0.0


104125.0
32.0
11.0
20.0
0.0
0.0
4.0
124.0
112.0
135.0
134.0
31.0
0.0
0.0
0.0
4949.0
0.0
0.0
1414.0
7612.0
12632.0
0.0
0.0
0.0
5706.0
0.0
4257.0
28364.0
38588.0
0.0


104125.0
32.0
11.0
20.0
0.0
0.0
4.0
124.0
112.0
135.0
134.0
31.0
0.0
0.0
0.0
4949.0
0.0
0.0
1414.0
7612.0
12632.0
0.0
0.0
0.0
5706.0
0.0
4257.0
28364.0
38588.0
0.0

1.0
1.0
1.0
1.0
1.0
1.0
1.0
1.0
1.0
1.0
1.0
1.0
1.0
1.0
1.0
1.0
1.0
1.0
1.0
1.0
1.0
1.0
1.0
1.0
1.0
1.0
1.0
1.0
1.0
1.0


40703.0
120.0
0.0
104.0
0.0
0.0
0.0
0.0
0.0
0.0
0.0
0.0
0.0
0.0
0.0
0.0
0.0
0.0
0.0
0.0
0.0
0.0
0.0
0.0
0.0
0.0
0.0
40479.0
0.0
0.0


224.0
120.0
0.0
104.0
0.0
0.0
0.0
0.0
0.0
0.0
0.0
0.0
0.0
0.0
0.0
0.0
0.0
0.0
0.0
0.0
0.0
0.0
0.0
0.0
0.0
0.0
0.0
0.0
0.0
0.0


224.0
120.0
0.0
104.0
0.0
0.0
0.0
0.0
0.0
0.0
0.0
0.0
0.0
0.0
0.0
0.0
0.0
0.0
0.0
0.0
0.0
0.0
0.0
0.0
0.0
0.0
0.0
0.0
0.0
0.0

1.0
1.0
1.0
1.0
1.0
1.0
1.0
1.0
1.0
1.0
1.0
1.0
1.0
1.0
1.0
1.0
1.0
1.0
1.0
1.0
1.0
1.0
1.0
1.0
1.0
1.0
1.0
1.0
1.0
1.0


40479.0
0.0
0.0
0.0
0.0
0.0
0.0
0.0
0.0
0.0
0.0
0.0
0.0
0.0
0.0
0.0
0.0
0.0
0.0
0.0
0.0
0.0
0.0
0.0
0.0
0.0
0.0
40479.0
0.0
0.0

1.0
1.0
1.0
1.0
1.0
1.0
1.0
1.0
1.0
1.0
1.0
1.0
1.0
1.0
1.0
1.0
1.0
1.0
1.0
1.0
1.0
1.0
1.0
1.0
1.0
1.0
1.0
1.0
1.0
1.0


40386.0
0.0
0.0
0.0
0.0
0.0
0.0
0.0
0.0
0.0
0.0
0.0
0.0
0.0
0.0
0.0
0.0
0.0
0.0
0.0
0.0
0.0
0.0
0.0
0.0
0.0
0.0
40386.0
0.0
0.0

1.0
1.0
1.0
1.0
1.0
1.0
1.0
1.0
1.0
1.0
1.0
1.0
1.0
1.0
1.0
1.0
1.0
1.0
1.0
1.0
1.0
1.0
1.0
1.0
1.0
1.0
1.0
1.0
1.0
1.0


14154.0
59.0
79.0
168.0
0.0
0.0
0.0
0.0
0.0
0.0
160.0
16.0
0.0
12097.0
1575.0
0.0
0.0
0.0
0.0
0.0
0.0
0.0
0.0
0.0
0.0
0.0
0.0
0.0
0.0
0.0


7562.0
59.0
0.0
0.0
0.0
0.0
0.0
0.0
0.0
0.0
160.0
0.0
0.0
7343.0
0.0
0.0
0.0
0.0
0.0
0.0
0.0
0.0
0.0
0.0
0.0
0.0
0.0
0.0
0.0
0.0


219.0
59.0
0.0
0.0
0.0
0.0
0.0
0.0
0.0
0.0
160.0
0.0
0.0
0.0
0.0
0.0
0.0
0.0
0.0
0.0
0.0
0.0
0.0
0.0
0.0
0.0
0.0
0.0
0.0
0.0

1.0
1.0
1.0
1.0
1.0
1.0
1.0
1.0
1.0
1.0
1.0
1.0
1.0
1.0
1.0
1.0
1.0
1.0
1.0
1.0
1.0
1.0
1.0
1.0
1.0
1.0
1.0
1.0
1.0
1.0


7343.0
0.0
0.0
0.0
0.0
0.0
0.0
0.0
0.0
0.0
0.0
0.0
0.0
7343.0
0.0
0.0
0.0
0.0
0.0
0.0
0.0
0.0
0.0
0.0
0.0
0.0
0.0
0.0
0.0
0.0

1.0
1.0
1.0
1.0
1.0
1.0
1.0
1.0
1.0
1.0
1.0
1.0
1.0
1.0
1.0
1.0
1.0
1.0
1.0
1.0
1.0
1.0
1.0
1.0
1.0
1.0
1.0
1.0
1.0
1.0


6592.0
0.0
79.0
168.0
0.0
0.0
0.0
0.0
0.0
0.0
0.0
16.0
0.0
4754.0
1575.0
0.0
0.0
0.0
0.0
0.0
0.0
0.0
0.0
0.0
0.0
0.0
0.0
0.0
0.0
0.0


6592.0
0.0
79.0
168.0
0.0
0.0
0.0
0.0
0.0
0.0
0.0
16.0
0.0
4754.0
1575.0
0.0
0.0
0.0
0.0
0.0
0.0
0.0
0.0
0.0
0.0
0.0
0.0
0.0
0.0
0.0

1.0
1.0
1.0
1.0
1.0
1.0
1.0
1.0
1.0
1.0
1.0
1.0
1.0
1.0
1.0
1.0
1.0
1.0
1.0
1.0
1.0
1.0
1.0
1.0
1.0
1.0
1.0
1.0
1.0
1.0


192.0
5.0
0.0
0.0
0.0
0.0
0.0
0.0
8.0
48.0
66.0
57.0
0.0
8.0
0.0
0.0
0.0
0.0
0.0
0.0
0.0
0.0
0.0
0.0
0.0
0.0
0.0
0.0
0.0
0.0


182.0
5.0
0.0
0.0
0.0
0.0
0.0
0.0
0.0
48.0
64.0
57.0
0.0
8.0
0.0
0.0
0.0
0.0
0.0
0.0
0.0
0.0
0.0
0.0
0.0
0.0
0.0
0.0
0.0
0.0


182.0
5.0
0.0
0.0
0.0
0.0
0.0
0.0
0.0
48.0
64.0
57.0
0.0
8.0
0.0
0.0
0.0
0.0
0.0
0.0
0.0
0.0
0.0
0.0
0.0
0.0
0.0
0.0
0.0
0.0

1.0
1.0
1.0
1.0
1.0
1.0
1.0
1.0
1.0
1.0
1.0
1.0
1.0
1.0
1.0
1.0
1.0
1.0
1.0
1.0
1.0
1.0
1.0
1.0
1.0
1.0
1.0
1.0
1.0
1.0


10.0
0.0
0.0
0.0
0.0
0.0
0.0
0.0
8.0
0.0
2.0
0.0
0.0
0.0
0.0
0.0
0.0
0.0
0.0
0.0
0.0
0.0
0.0
0.0
0.0
0.0
0.0
0.0
0.0
0.0


10.0
0.0
0.0
0.0
0.0
0.0
0.0
0.0
8.0
0.0
2.0
0.0
0.0
0.0
0.0
0.0
0.0
0.0
0.0
0.0
0.0
0.0
0.0
0.0
0.0
0.0
0.0
0.0
0.0
0.0

1.0
1.0
1.0
1.0
1.0
1.0
1.0
1.0
1.0
1.0
1.0
1.0
1.0
1.0
1.0
1.0
1.0
1.0
1.0
1.0
1.0
1.0
1.0
1.0
1.0
1.0
1.0
1.0
1.0
1.0


76248.0
0.0
78.0
73.0
0.0
0.0
51.0
8.0
0.0
0.0
0.0
0.0
0.0
0.0
0.0
0.0
0.0
0.0
0.0
0.0
0.0
0.0
0.0
0.0
57350.0
0.0
6531.0
0.0
0.0
12157.0


74098.0
0.0
66.0
53.0
0.0
0.0
51.0
8.0
0.0
0.0
0.0
0.0
0.0
0.0
0.0
0.0
0.0
0.0
0.0
0.0
0.0
0.0
0.0
0.0
57350.0
0.0
5641.0
0.0
0.0
10929.0


109.0
0.0
40.0
18.0
0.0
0.0
51.0
0.0
0.0
0.0
0.0
0.0
0.0
0.0
0.0
0.0
0.0
0.0
0.0
0.0
0.0
0.0
0.0
0.0
0.0
0.0
0.0
0.0
0.0
0.0

1.0
1.0
1.0
1.0
1.0
1.0
1.0
1.0
1.0
1.0
1.0
1.0
1.0
1.0
1.0
1.0
1.0
1.0
1.0
1.0
1.0
1.0
1.0
1.0
1.0
1.0
1.0
1.0
1.0
1.0


10963.0
0.0
0.0
26.0
0.0
0.0
0.0
8.0
0.0
0.0
0.0
0.0
0.0
0.0
0.0
0.0
0.0
0.0
0.0
0.0
0.0
0.0
0.0
0.0
0.0
0.0
0.0
0.0
0.0
10929.0

1.0
1.0
1.0
1.0
1.0
1.0
1.0
1.0
1.0
1.0
1.0
1.0
1.0
1.0
1.0
1.0
1.0
1.0
1.0
1.0
1.0
1.0
1.0
1.0
1.0
1.0
1.0
1.0
1.0
1.0


23.0
0.0
23.0
0.0
0.0
0.0
0.0
0.0
0.0
0.0
0.0
0.0
0.0
0.0
0.0
0.0
0.0
0.0
0.0
0.0
0.0
0.0
0.0
0.0
0.0
0.0
0.0
0.0
0.0
0.0

1.0
1.0
1.0
1.0
1.0
1.0
1.0
1.0
1.0
1.0
1.0
1.0
1.0
1.0
1.0
1.0
1.0
1.0
1.0
1.0
1.0
1.0
1.0
1.0
1.0
1.0
1.0
1.0
1.0
1.0


63003.0
0.0
3.0
9.0
0.0
0.0
0.0
0.0
0.0
0.0
0.0
0.0
0.0
0.0
0.0
0.0
0.0
0.0
0.0
0.0
0.0
0.0
0.0
0.0
57350.0
0.0
5641.0
0.0
0.0
0.0

0.99
0.99
0.99
0.99
0.99
0.99
0.99
0.99
0.99
0.99
0.99
0.99
0.99
0.99
0.99
0.99
0.99
0.99
0.99
0.99
0.99
0.99
0.99
0.99
0.99
0.99
0.99
0.99
0.99
0.99


32.0
0.0
12.0
20.0
0.0
0.0
0.0
0.0
0.0
0.0
0.0
0.0
0.0
0.0
0.0
0.0
0.0
0.0
0.0
0.0
0.0
0.0
0.0
0.0
0.0
0.0
0.0
0.0
0.0
0.0


32.0
0.0
12.0
20.0
0.0
0.0
0.0
0.0
0.0
0.0
0.0
0.0
0.0
0.0
0.0
0.0
0.0
0.0
0.0
0.0
0.0
0.0
0.0
0.0
0.0
0.0
0.0
0.0
0.0
0.0

1.0
1.0
1.0
1.0
1.0
1.0
1.0
1.0
1.0
1.0
1.0
1.0
1.0
1.0
1.0
1.0
1.0
1.0
1.0
1.0
1.0
1.0
1.0
1.0
1.0
1.0
1.0
1.0
1.0
1.0


2118.0
0.0
0.0
0.0
0.0
0.0
0.0
0.0
0.0
0.0
0.0
0.0
0.0
0.0
0.0
0.0
0.0
0.0
0.0
0.0
0.0
0.0
0.0
0.0
0.0
0.0
890.0
0.0
0.0
1228.0


2118.0
0.0
0.0
0.0
0.0
0.0
0.0
0.0
0.0
0.0
0.0
0.0
0.0
0.0
0.0
0.0
0.0
0.0
0.0
0.0
0.0
0.0
0.0
0.0
0.0
0.0
890.0
0.0
0.0
1228.0

1.0
1.0
1.0
1.0
1.0
1.0
1.0
1.0
1.0
1.0
1.0
1.0
1.0
1.0
1.0
1.0
1.0
1.0
1.0
1.0
1.0
1.0
1.0
1.0
1.0
1.0
1.0
1.0
1.0
1.0


1740723.0
0.0
109.0
0.0
0.0
0.0
0.0
0.0
0.0
0.0
0.0
0.0
0.0
0.0
0.0
53772.0
0.0
458329.0
818897.0
76876.0
0.0
0.0
0.0
0.0
87995.0
74556.0
0.0
0.0
0.0
170189.0

1.0
1.0
1.0
1.0
1.0
1.0
1.0
1.0
1.0
1.0
1.0
1.0
1.0
1.0
1.0
1.0
1.0
1.0
1.0
1.0
1.0
1.0
1.0
1.0
1.0
1.0
1.0
1.0
1.0
1.0


703183.0
0.0
109.0
0.0
0.0
0.0
0.0
0.0
0.0
0.0
0.0
0.0
0.0
0.0
0.0
0.0
0.0
458329.0
0.0
0.0
0.0
0.0
0.0
0.0
0.0
74556.0
0.0
0.0
0.0
170189.0

0.99
0.99
0.99
0.99
0.99
0.99
0.99
0.99
0.99
0.99
0.99
0.99
0.99
0.99
0.99
0.99
0.99
0.99
0.99
0.99
0.99
0.99
0.99
0.99
0.99
0.99
0.99
0.99
0.99
0.99


699858.0
0.0
109.0
0.0
0.0
0.0
0.0
0.0
0.0
0.0
0.0
0.0
0.0
0.0
0.0
0.0
0.0
458329.0
0.0
0.0
0.0
0.0
0.0
0.0
0.0
73424.0
0.0
0.0
0.0
167996.0

1.0
1.0
1.0
1.0
1.0
1.0
1.0
1.0
1.0
1.0
1.0
1.0
1.0
1.0
1.0
1.0
1.0
1.0
1.0
1.0
1.0
1.0
1.0
1.0
1.0
1.0
1.0
1.0
1.0
1.0


622114.0
0.0
0.0
0.0
0.0
0.0
0.0
0.0
0.0
0.0
0.0
0.0
0.0
0.0
0.0
0.0
0.0
0.0
534119.0
0.0
0.0
0.0
0.0
0.0
87995.0
0.0
0.0
0.0
0.0
0.0

0.99
0.99
0.99
0.99
0.99
0.99
0.99
0.99
0.99
0.99
0.99
0.99
0.99
0.99
0.99
0.99
0.99
0.99
0.99
0.99
0.99
0.99
0.99
0.99
0.99
0.99
0.99
0.99
0.99
0.99


622093.0
0.0
0.0
0.0
0.0
0.0
0.0
0.0
0.0
0.0
0.0
0.0
0.0
0.0
0.0
0.0
0.0
0.0
534098.0
0.0
0.0
0.0
0.0
0.0
87995.0
0.0
0.0
0.0
0.0
0.0

0.99
0.99
0.99
0.99
0.99
0.99
0.99
0.99
0.99
0.99
0.99
0.99
0.99
0.99
0.99
0.99
0.99
0.99
0.99
0.99
0.99
0.99
0.99
0.99
0.99
0.99
0.99
0.99
0.99
0.99


260570.0
0.0
0.0
0.0
0.0
0.0
0.0
0.0
0.0
0.0
0.0
0.0
0.0
0.0
0.0
0.0
0.0
0.0
260570.0
0.0
0.0
0.0
0.0
0.0
0.0
0.0
0.0
0.0
0.0
0.0


260570.0
0.0
0.0
0.0
0.0
0.0
0.0
0.0
0.0
0.0
0.0
0.0
0.0
0.0
0.0
0.0
0.0
0.0
260570.0
0.0
0.0
0.0
0.0
0.0
0.0
0.0
0.0
0.0
0.0
0.0

0.99
0.99
0.99
0.99
0.99
0.99
0.99
0.99
0.99
0.99
0.99
0.99
0.99
0.99
0.99
0.99
0.99
0.99
0.99
0.99
0.99
0.99
0.99
0.99
0.99
0.99
0.99
0.99
0.99
0.99


101073.0
0.0
0.0
0.0
0.0
0.0
0.0
0.0
0.0
0.0
0.0
0.0
0.0
0.0
0.0
0.0
0.0
0.0
24197.0
76876.0
0.0
0.0
0.0
0.0
0.0
0.0
0.0
0.0
0.0
0.0


101073.0
0.0
0.0
0.0
0.0
0.0
0.0
0.0
0.0
0.0
0.0
0.0
0.0
0.0
0.0
0.0
0.0
0.0
24197.0
76876.0
0.0
0.0
0.0
0.0
0.0
0.0
0.0
0.0
0.0
0.0

1.0
1.0
1.0
1.0
1.0
1.0
1.0
1.0
1.0
1.0
1.0
1.0
1.0
1.0
1.0
1.0
1.0
1.0
1.0
1.0
1.0
1.0
1.0
1.0
1.0
1.0
1.0
1.0
1.0
1.0


53772.0
0.0
0.0
0.0
0.0
0.0
0.0
0.0
0.0
0.0
0.0
0.0
0.0
0.0
0.0
53772.0
0.0
0.0
0.0
0.0
0.0
0.0
0.0
0.0
0.0
0.0
0.0
0.0
0.0
0.0


53772.0
0.0
0.0
0.0
0.0
0.0
0.0
0.0
0.0
0.0
0.0
0.0
0.0
0.0
0.0
53772.0
0.0
0.0
0.0
0.0
0.0
0.0
0.0
0.0
0.0
0.0
0.0
0.0
0.0
0.0

0.99
0.99
0.99
0.99
0.99
0.99
0.99
0.99
0.99
0.99
0.99
0.99
0.99
0.99
0.99
0.99
0.99
0.99
0.99
0.99
0.99
0.99
0.99
0.99
0.99
0.99
0.99
0.99
0.99
0.99


103.0
0.0
0.0
0.0
13.0
15.0
75.0
0.0
0.0
0.0
0.0
0.0
0.0
0.0
0.0
0.0
0.0
0.0
0.0
0.0
0.0
0.0
0.0
0.0
0.0
0.0
0.0
0.0
0.0
0.0


103.0
0.0
0.0
0.0
13.0
15.0
75.0
0.0
0.0
0.0
0.0
0.0
0.0
0.0
0.0
0.0
0.0
0.0
0.0
0.0
0.0
0.0
0.0
0.0
0.0
0.0
0.0
0.0
0.0
0.0


103.0
0.0
0.0
0.0
13.0
15.0
75.0
0.0
0.0
0.0
0.0
0.0
0.0
0.0
0.0
0.0
0.0
0.0
0.0
0.0
0.0
0.0
0.0
0.0
0.0
0.0
0.0
0.0
0.0
0.0

1.0
1.0
1.0
1.0
1.0
1.0
1.0
1.0
1.0
1.0
1.0
1.0
1.0
1.0
1.0
1.0
1.0
1.0
1.0
1.0
1.0
1.0
1.0
1.0
1.0
1.0
1.0
1.0
1.0
1.0


12060.0
0.0
0.0
0.0
0.0
0.0
0.0
34.0
37.0
7.0
16.0
16.0
0.0
0.0
0.0
0.0
0.0
0.0
9437.0
2198.0
0.0
0.0
0.0
0.0
315.0
0.0
0.0
0.0
0.0
0.0


12060.0
0.0
0.0
0.0
0.0
0.0
0.0
34.0
37.0
7.0
16.0
16.0
0.0
0.0
0.0
0.0
0.0
0.0
9437.0
2198.0
0.0
0.0
0.0
0.0
315.0
0.0
0.0
0.0
0.0
0.0

0.99
0.99
0.99
0.99
0.99
0.99
0.99
0.99
0.99
0.99
0.99
0.99
0.99
0.99
0.99
0.99
0.99
0.99
0.99
0.99
0.99
0.99
0.99
0.99
0.99
0.99
0.99
0.99
0.99
0.99


331.0
0.0
0.0
0.0
0.0
0.0
0.0
0.0
0.0
0.0
0.0
16.0
0.0
0.0
0.0
0.0
0.0
0.0
0.0
0.0
0.0
0.0
0.0
0.0
315.0
0.0
0.0
0.0
0.0
0.0

1.0
1.0
1.0
1.0
1.0
1.0
1.0
1.0
1.0
1.0
1.0
1.0
1.0
1.0
1.0
1.0
1.0
1.0
1.0
1.0
1.0
1.0
1.0
1.0
1.0
1.0
1.0
1.0
1.0
1.0


329885.0
1.0
5.0
14.0
175.0
34.0
46.0
0.0
13.0
0.0
0.0
78.0
0.0
8719.0
30705.0
28206.0
27293.0
0.0
0.0
0.0
0.0
0.0
0.0
0.0
2026.0
180720.0
4517.0
27683.0
2675.0
16975.0

1.0
1.0
1.0
1.0
1.0
1.0
1.0
1.0
1.0
1.0
1.0
1.0
1.0
1.0
1.0
1.0
1.0
1.0
1.0
1.0
1.0
1.0
1.0
1.0
1.0
1.0
1.0
1.0
1.0
1.0


84852.0
0.0
0.0
0.0
82.0
0.0
46.0
0.0
13.0
0.0
0.0
0.0
0.0
0.0
522.0
22417.0
4420.0
0.0
0.0
0.0
0.0
0.0
0.0
0.0
0.0
25152.0
4517.0
27683.0
0.0
0.0

0.99
0.99
0.99
0.99
0.99
0.99
0.99
0.99
0.99
0.99
0.99
0.99
0.99
0.99
0.99
0.99
0.99
0.99
0.99
0.99
0.99
0.99
0.99
0.99
0.99
0.99
0.99
0.99
0.99
0.99


604.0
0.0
0.0
0.0
82.0
0.0
0.0
0.0
0.0
0.0
0.0
0.0
0.0
0.0
522.0
0.0
0.0
0.0
0.0
0.0
0.0
0.0
0.0
0.0
0.0
0.0
0.0
0.0
0.0
0.0

1.0
1.0
1.0
1.0
1.0
1.0
1.0
1.0
1.0
1.0
1.0
1.0
1.0
1.0
1.0
1.0
1.0
1.0
1.0
1.0
1.0
1.0
1.0
1.0
1.0
1.0
1.0
1.0
1.0
1.0


4576.0
0.0
0.0
0.0
0.0
0.0
46.0
0.0
13.0
0.0
0.0
0.0
0.0
0.0
0.0
0.0
0.0
0.0
0.0
0.0
0.0
0.0
0.0
0.0
0.0
0.0
4517.0
0.0
0.0
0.0

0.86
0.86
0.86
0.86
0.86
0.86
0.86
0.86
0.86
0.86
0.86
0.86
0.86
0.86
0.86
0.86
0.86
0.86
0.86
0.86
0.86
0.86
0.86
0.86
0.86
0.86
0.86
0.86
0.86
0.86


20212.0
0.0
0.0
0.0
0.0
0.0
0.0
0.0
0.0
0.0
0.0
0.0
0.0
0.0
0.0
0.0
0.0
0.0
0.0
0.0
0.0
0.0
0.0
0.0
0.0
0.0
0.0
20212.0
0.0
0.0

1.0
1.0
1.0
1.0
1.0
1.0
1.0
1.0
1.0
1.0
1.0
1.0
1.0
1.0
1.0
1.0
1.0
1.0
1.0
1.0
1.0
1.0
1.0
1.0
1.0
1.0
1.0
1.0
1.0
1.0


45920.0
0.0
1.0
14.0
93.0
0.0
0.0
0.0
0.0
0.0
0.0
0.0
0.0
0.0
0.0
5774.0
22392.0
0.0
0.0
0.0
0.0
0.0
0.0
0.0
1781.0
0.0
0.0
0.0
32.0
15833.0

0.86
0.86
0.86
0.86
0.86
0.86
0.86
0.86
0.86
0.86
0.86
0.86
0.86
0.86
0.86
0.86
0.86
0.86
0.86
0.86
0.86
0.86
0.86
0.86
0.86
0.86
0.86
0.86
0.86
0.86


185394.0
0.0
0.0
0.0
0.0
34.0
0.0
0.0
0.0
0.0
0.0
0.0
0.0
0.0
29792.0
0.0
0.0
0.0
0.0
0.0
0.0
0.0
0.0
0.0
0.0
155568.0
0.0
0.0
0.0
0.0


185394.0
0.0
0.0
0.0
0.0
34.0
0.0
0.0
0.0
0.0
0.0
0.0
0.0
0.0
29792.0
0.0
0.0
0.0
0.0
0.0
0.0
0.0
0.0
0.0
0.0
155568.0
0.0
0.0
0.0
0.0

1.0
1.0
1.0
1.0
1.0
1.0
1.0
1.0
1.0
1.0
1.0
1.0
1.0
1.0
1.0
1.0
1.0
1.0
1.0
1.0
1.0
1.0
1.0
1.0
1.0
1.0
1.0
1.0
1.0
1.0


41852.0
0.0
3.0
0.0
17.0
67.0
32.0
69.0
0.0
0.0
11.0
4.0
0.0
0.0
0.0
0.0
0.0
0.0
0.0
34.0
0.0
0.0
0.0
0.0
0.0
13152.0
26812.0
1333.0
84.0
234.0


31369.0
0.0
3.0
0.0
0.0
0.0
1.0
69.0
0.0
0.0
11.0
0.0
0.0
0.0
0.0
0.0
0.0
0.0
0.0
0.0
0.0
0.0
0.0
0.0
0.0
12208.0
19077.0
0.0
0.0
0.0


31369.0
0.0
3.0
0.0
0.0
0.0
1.0
69.0
0.0
0.0
11.0
0.0
0.0
0.0
0.0
0.0
0.0
0.0
0.0
0.0
0.0
0.0
0.0
0.0
0.0
12208.0
19077.0
0.0
0.0
0.0

1.0
1.0
1.0
1.0
1.0
1.0
1.0
1.0
1.0
1.0
1.0
1.0
1.0
1.0
1.0
1.0
1.0
1.0
1.0
1.0
1.0
1.0
1.0
1.0
1.0
1.0
1.0
1.0
1.0
1.0


7841.0
0.0
0.0
0.0
17.0
57.0
28.0
0.0
0.0
0.0
0.0
4.0
0.0
0.0
0.0
0.0
0.0
0.0
0.0
0.0
0.0
0.0
0.0
0.0
0.0
0.0
7735.0
0.0
0.0
0.0


7841.0
0.0
0.0
0.0
17.0
57.0
28.0
0.0
0.0
0.0
0.0
4.0
0.0
0.0
0.0
0.0
0.0
0.0
0.0
0.0
0.0
0.0
0.0
0.0
0.0
0.0
7735.0
0.0
0.0
0.0

0.98
0.98
0.98
0.98
0.98
0.98
0.98
0.98
0.98
0.98
0.98
0.98
0.98
0.98
0.98
0.98
0.98
0.98
0.98
0.98
0.98
0.98
0.98
0.98
0.98
0.98
0.98
0.98
0.98
0.98


2389.0
0.0
0.0
0.0
0.0
10.0
3.0
0.0
0.0
0.0
0.0
0.0
0.0
0.0
0.0
0.0
0.0
0.0
0.0
34.0
0.0
0.0
0.0
0.0
0.0
944.0
0.0
1333.0
0.0
65.0


1148.0
0.0
0.0
0.0
0.0
10.0
3.0
0.0
0.0
0.0
0.0
0.0
0.0
0.0
0.0
0.0
0.0
0.0
0.0
0.0
0.0
0.0
0.0
0.0
0.0
0.0
0.0
1135.0
0.0
0.0

1.0
1.0
1.0
1.0
1.0
1.0
1.0
1.0
1.0
1.0
1.0
1.0
1.0
1.0
1.0
1.0
1.0
1.0
1.0
1.0
1.0
1.0
1.0
1.0
1.0
1.0
1.0
1.0
1.0
1.0


1207.0
0.0
0.0
0.0
0.0
0.0
0.0
0.0
0.0
0.0
0.0
0.0
0.0
0.0
0.0
0.0
0.0
0.0
0.0
0.0
0.0
0.0
0.0
0.0
0.0
944.0
0.0
198.0
0.0
65.0

1.0
1.0
1.0
1.0
1.0
1.0
1.0
1.0
1.0
1.0
1.0
1.0
1.0
1.0
1.0
1.0
1.0
1.0
1.0
1.0
1.0
1.0
1.0
1.0
1.0
1.0
1.0
1.0
1.0
1.0


34.0
0.0
0.0
0.0
0.0
0.0
0.0
0.0
0.0
0.0
0.0
0.0
0.0
0.0
0.0
0.0
0.0
0.0
0.0
34.0
0.0
0.0
0.0
0.0
0.0
0.0
0.0
0.0
0.0
0.0

1.0
1.0
1.0
1.0
1.0
1.0
1.0
1.0
1.0
1.0
1.0
1.0
1.0
1.0
1.0
1.0
1.0
1.0
1.0
1.0
1.0
1.0
1.0
1.0
1.0
1.0
1.0
1.0
1.0
1.0


253.0
0.0
0.0
0.0
0.0
0.0
0.0
0.0
0.0
0.0
0.0
0.0
0.0
0.0
0.0
0.0
0.0
0.0
0.0
0.0
0.0
0.0
0.0
0.0
0.0
0.0
0.0
0.0
84.0
169.0


253.0
0.0
0.0
0.0
0.0
0.0
0.0
0.0
0.0
0.0
0.0
0.0
0.0
0.0
0.0
0.0
0.0
0.0
0.0
0.0
0.0
0.0
0.0
0.0
0.0
0.0
0.0
0.0
84.0
169.0

1.0
1.0
1.0
1.0
1.0
1.0
1.0
1.0
1.0
1.0
1.0
1.0
1.0
1.0
1.0
1.0
1.0
1.0
1.0
1.0
1.0
1.0
1.0
1.0
1.0
1.0
1.0
1.0
1.0
1.0


3966.0
0.0
36.0
48.0
0.0
0.0
0.0
0.0
0.0
0.0
0.0
0.0
0.0
3845.0
37.0
0.0
0.0
0.0
0.0
0.0
0.0
0.0
0.0
0.0
0.0
0.0
0.0
0.0
0.0
0.0


104.0
0.0
19.0
48.0
0.0
0.0
0.0
0.0
0.0
0.0
0.0
0.0
0.0
0.0
37.0
0.0
0.0
0.0
0.0
0.0
0.0
0.0
0.0
0.0
0.0
0.0
0.0
0.0
0.0
0.0


104.0
0.0
19.0
48.0
0.0
0.0
0.0
0.0
0.0
0.0
0.0
0.0
0.0
0.0
37.0
0.0
0.0
0.0
0.0
0.0
0.0
0.0
0.0
0.0
0.0
0.0
0.0
0.0
0.0
0.0

1.0
1.0
1.0
1.0
1.0
1.0
1.0
1.0
1.0
1.0
1.0
1.0
1.0
1.0
1.0
1.0
1.0
1.0
1.0
1.0
1.0
1.0
1.0
1.0
1.0
1.0
1.0
1.0
1.0
1.0


3862.0
0.0
17.0
0.0
0.0
0.0
0.0
0.0
0.0
0.0
0.0
0.0
0.0
3845.0
0.0
0.0
0.0
0.0
0.0
0.0
0.0
0.0
0.0
0.0
0.0
0.0
0.0
0.0
0.0
0.0


3862.0
0.0
17.0
0.0
0.0
0.0
0.0
0.0
0.0
0.0
0.0
0.0
0.0
3845.0
0.0
0.0
0.0
0.0
0.0
0.0
0.0
0.0
0.0
0.0
0.0
0.0
0.0
0.0
0.0
0.0

1.0
1.0
1.0
1.0
1.0
1.0
1.0
1.0
1.0
1.0
1.0
1.0
1.0
1.0
1.0
1.0
1.0
1.0
1.0
1.0
1.0
1.0
1.0
1.0
1.0
1.0
1.0
1.0
1.0
1.0


83.0
36.0
0.0
0.0
0.0
0.0
0.0
0.0
0.0
22.0
0.0
25.0
0.0
0.0
0.0
0.0
0.0
0.0
0.0
0.0
0.0
0.0
0.0
0.0
0.0
0.0
0.0
0.0
0.0
0.0


83.0
36.0
0.0
0.0
0.0
0.0
0.0
0.0
0.0
22.0
0.0
25.0
0.0
0.0
0.0
0.0
0.0
0.0
0.0
0.0
0.0
0.0
0.0
0.0
0.0
0.0
0.0
0.0
0.0
0.0


83.0
36.0
0.0
0.0
0.0
0.0
0.0
0.0
0.0
22.0
0.0
25.0
0.0
0.0
0.0
0.0
0.0
0.0
0.0
0.0
0.0
0.0
0.0
0.0
0.0
0.0
0.0
0.0
0.0
0.0

1.0
1.0
1.0
1.0
1.0
1.0
1.0
1.0
1.0
1.0
1.0
1.0
1.0
1.0
1.0
1.0
1.0
1.0
1.0
1.0
1.0
1.0
1.0
1.0
1.0
1.0
1.0
1.0
1.0
1.0


19994.0
0.0
0.0
0.0
0.0
0.0
0.0
55.0
0.0
0.0
0.0
0.0
0.0
0.0
0.0
0.0
0.0
8428.0
0.0
0.0
2351.0
0.0
0.0
0.0
0.0
0.0
0.0
0.0
9160.0
0.0


19994.0
0.0
0.0
0.0
0.0
0.0
0.0
55.0
0.0
0.0
0.0
0.0
0.0
0.0
0.0
0.0
0.0
8428.0
0.0
0.0
2351.0
0.0
0.0
0.0
0.0
0.0
0.0
0.0
9160.0
0.0


19994.0
0.0
0.0
0.0
0.0
0.0
0.0
55.0
0.0
0.0
0.0
0.0
0.0
0.0
0.0
0.0
0.0
8428.0
0.0
0.0
2351.0
0.0
0.0
0.0
0.0
0.0
0.0
0.0
9160.0
0.0

1.0
1.0
1.0
1.0
1.0
1.0
1.0
1.0
1.0
1.0
1.0
1.0
1.0
1.0
1.0
1.0
1.0
1.0
1.0
1.0
1.0
1.0
1.0
1.0
1.0
1.0
1.0
1.0
1.0
1.0


6262.0
4.0
5.0
0.0
0.0
0.0
0.0
0.0
33.0
0.0
0.0
0.0
0.0
0.0
0.0
2070.0
0.0
0.0
2926.0
0.0
0.0
0.0
0.0
0.0
39.0
0.0
0.0
0.0
0.0
1185.0


6262.0
4.0
5.0
0.0
0.0
0.0
0.0
0.0
33.0
0.0
0.0
0.0
0.0
0.0
0.0
2070.0
0.0
0.0
2926.0
0.0
0.0
0.0
0.0
0.0
39.0
0.0
0.0
0.0
0.0
1185.0


6262.0
4.0
5.0
0.0
0.0
0.0
0.0
0.0
33.0
0.0
0.0
0.0
0.0
0.0
0.0
2070.0
0.0
0.0
2926.0
0.0
0.0
0.0
0.0
0.0
39.0
0.0
0.0
0.0
0.0
1185.0

1.0
1.0
1.0
1.0
1.0
1.0
1.0
1.0
1.0
1.0
1.0
1.0
1.0
1.0
1.0
1.0
1.0
1.0
1.0
1.0
1.0
1.0
1.0
1.0
1.0
1.0
1.0
1.0
1.0
1.0


24.0
0.0
0.0
0.0
0.0
0.0
0.0
0.0
0.0
0.0
24.0
0.0
0.0
0.0
0.0
0.0
0.0
0.0
0.0
0.0
0.0
0.0
0.0
0.0
0.0
0.0
0.0
0.0
0.0
0.0


24.0
0.0
0.0
0.0
0.0
0.0
0.0
0.0
0.0
0.0
24.0
0.0
0.0
0.0
0.0
0.0
0.0
0.0
0.0
0.0
0.0
0.0
0.0
0.0
0.0
0.0
0.0
0.0
0.0
0.0


24.0
0.0
0.0
0.0
0.0
0.0
0.0
0.0
0.0
0.0
24.0
0.0
0.0
0.0
0.0
0.0
0.0
0.0
0.0
0.0
0.0
0.0
0.0
0.0
0.0
0.0
0.0
0.0
0.0
0.0

1.0
1.0
1.0
1.0
1.0
1.0
1.0
1.0
1.0
1.0
1.0
1.0
1.0
1.0
1.0
1.0
1.0
1.0
1.0
1.0
1.0
1.0
1.0
1.0
1.0
1.0
1.0
1.0
1.0
1.0


1751.0
0.0
0.0
0.0
22.0
0.0
0.0
0.0
0.0
0.0
16.0
0.0
0.0
0.0
0.0
798.0
0.0
0.0
0.0
0.0
0.0
0.0
0.0
0.0
0.0
0.0
915.0
0.0
0.0
0.0


1751.0
0.0
0.0
0.0
22.0
0.0
0.0
0.0
0.0
0.0
16.0
0.0
0.0
0.0
0.0
798.0
0.0
0.0
0.0
0.0
0.0
0.0
0.0
0.0
0.0
0.0
915.0
0.0
0.0
0.0


1751.0
0.0
0.0
0.0
22.0
0.0
0.0
0.0
0.0
0.0
16.0
0.0
0.0
0.0
0.0
798.0
0.0
0.0
0.0
0.0
0.0
0.0
0.0
0.0
0.0
0.0
915.0
0.0
0.0
0.0

1.0
1.0
1.0
1.0
1.0
1.0
1.0
1.0
1.0
1.0
1.0
1.0
1.0
1.0
1.0
1.0
1.0
1.0
1.0
1.0
1.0
1.0
1.0
1.0
1.0
1.0
1.0
1.0
1.0
1.0


30035.0
25.0
7.0
2.0
0.0
0.0
9.0
0.0
0.0
0.0
0.0
12.0
0.0
0.0
0.0
0.0
0.0
0.0
0.0
0.0
0.0
0.0
0.0
0.0
0.0
0.0
0.0
0.0
29980.0
0.0


21.0
0.0
7.0
2.0
0.0
0.0
0.0
0.0
0.0
0.0
0.0
12.0
0.0
0.0
0.0
0.0
0.0
0.0
0.0
0.0
0.0
0.0
0.0
0.0
0.0
0.0
0.0
0.0
0.0
0.0


21.0
0.0
7.0
2.0
0.0
0.0
0.0
0.0
0.0
0.0
0.0
12.0
0.0
0.0
0.0
0.0
0.0
0.0
0.0
0.0
0.0
0.0
0.0
0.0
0.0
0.0
0.0
0.0
0.0
0.0

1.0
1.0
1.0
1.0
1.0
1.0
1.0
1.0
1.0
1.0
1.0
1.0
1.0
1.0
1.0
1.0
1.0
1.0
1.0
1.0
1.0
1.0
1.0
1.0
1.0
1.0
1.0
1.0
1.0
1.0


16.0
16.0
0.0
0.0
0.0
0.0
0.0
0.0
0.0
0.0
0.0
0.0
0.0
0.0
0.0
0.0
0.0
0.0
0.0
0.0
0.0
0.0
0.0
0.0
0.0
0.0
0.0
0.0
0.0
0.0


16.0
16.0
0.0
0.0
0.0
0.0
0.0
0.0
0.0
0.0
0.0
0.0
0.0
0.0
0.0
0.0
0.0
0.0
0.0
0.0
0.0
0.0
0.0
0.0
0.0
0.0
0.0
0.0
0.0
0.0

1.0
1.0
1.0
1.0
1.0
1.0
1.0
1.0
1.0
1.0
1.0
1.0
1.0
1.0
1.0
1.0
1.0
1.0
1.0
1.0
1.0
1.0
1.0
1.0
1.0
1.0
1.0
1.0
1.0
1.0


9.0
9.0
0.0
0.0
0.0
0.0
0.0
0.0
0.0
0.0
0.0
0.0
0.0
0.0
0.0
0.0
0.0
0.0
0.0
0.0
0.0
0.0
0.0
0.0
0.0
0.0
0.0
0.0
0.0
0.0


9.0
9.0
0.0
0.0
0.0
0.0
0.0
0.0
0.0
0.0
0.0
0.0
0.0
0.0
0.0
0.0
0.0
0.0
0.0
0.0
0.0
0.0
0.0
0.0
0.0
0.0
0.0
0.0
0.0
0.0

1.0
1.0
1.0
1.0
1.0
1.0
1.0
1.0
1.0
1.0
1.0
1.0
1.0
1.0
1.0
1.0
1.0
1.0
1.0
1.0
1.0
1.0
1.0
1.0
1.0
1.0
1.0
1.0
1.0
1.0


29989.0
0.0
0.0
0.0
0.0
0.0
9.0
0.0
0.0
0.0
0.0
0.0
0.0
0.0
0.0
0.0
0.0
0.0
0.0
0.0
0.0
0.0
0.0
0.0
0.0
0.0
0.0
0.0
29980.0
0.0


29989.0
0.0
0.0
0.0
0.0
0.0
9.0
0.0
0.0
0.0
0.0
0.0
0.0
0.0
0.0
0.0
0.0
0.0
0.0
0.0
0.0
0.0
0.0
0.0
0.0
0.0
0.0
0.0
29980.0
0.0

1.0
1.0
1.0
1.0
1.0
1.0
1.0
1.0
1.0
1.0
1.0
1.0
1.0
1.0
1.0
1.0
1.0
1.0
1.0
1.0
1.0
1.0
1.0
1.0
1.0
1.0
1.0
1.0
1.0
1.0


17350.0
0.0
0.0
1.0
0.0
2.0
11.0
0.0
0.0
0.0
0.0
0.0
0.0
0.0
0.0
0.0
53.0
0.0
0.0
0.0
0.0
0.0
0.0
0.0
540.0
554.0
8773.0
6348.0
438.0
630.0


15481.0
0.0
0.0
1.0
0.0
2.0
11.0
0.0
0.0
0.0
0.0
0.0
0.0
0.0
0.0
0.0
0.0
0.0
0.0
0.0
0.0
0.0
0.0
0.0
0.0
347.0
8633.0
5988.0
256.0
243.0


15481.0
0.0
0.0
1.0
0.0
2.0
11.0
0.0
0.0
0.0
0.0
0.0
0.0
0.0
0.0
0.0
0.0
0.0
0.0
0.0
0.0
0.0
0.0
0.0
0.0
347.0
8633.0
5988.0
256.0
243.0

1.0
1.0
1.0
1.0
1.0
1.0
1.0
1.0
1.0
1.0
1.0
1.0
1.0
1.0
1.0
1.0
1.0
1.0
1.0
1.0
1.0
1.0
1.0
1.0
1.0
1.0
1.0
1.0
1.0
1.0


1869.0
0.0
0.0
0.0
0.0
0.0
0.0
0.0
0.0
0.0
0.0
0.0
0.0
0.0
0.0
0.0
53.0
0.0
0.0
0.0
0.0
0.0
0.0
0.0
540.0
207.0
140.0
360.0
182.0
387.0

0.79
0.79
0.79
0.79
0.79
0.79
0.79
0.79
0.79
0.79
0.79
0.79
0.79
0.79
0.79
0.79
0.79
0.79
0.79
0.79
0.79
0.79
0.79
0.79
0.79
0.79
0.79
0.79
0.79
0.79


781.0
0.0
0.0
0.0
0.0
0.0
0.0
0.0
0.0
0.0
0.0
0.0
0.0
0.0
0.0
0.0
0.0
0.0
0.0
0.0
0.0
0.0
0.0
0.0
0.0
77.0
42.0
335.0
35.0
292.0

1.0
1.0
1.0
1.0
1.0
1.0
1.0
1.0
1.0
1.0
1.0
1.0
1.0
1.0
1.0
1.0
1.0
1.0
1.0
1.0
1.0
1.0
1.0
1.0
1.0
1.0
1.0
1.0
1.0
1.0


200.0
0.0
0.0
0.0
0.0
0.0
0.0
0.0
0.0
0.0
0.0
0.0
0.0
0.0
0.0
0.0
0.0
0.0
0.0
0.0
0.0
0.0
0.0
0.0
104.0
9.0
0.0
0.0
87.0
0.0

1.0
1.0
1.0
1.0
1.0
1.0
1.0
1.0
1.0
1.0
1.0
1.0
1.0
1.0
1.0
1.0
1.0
1.0
1.0
1.0
1.0
1.0
1.0
1.0
1.0
1.0
1.0
1.0
1.0
1.0


134.0
0.0
0.0
0.0
0.0
0.0
0.0
0.0
0.0
0.0
0.0
0.0
0.0
0.0
0.0
0.0
0.0
0.0
0.0
0.0
0.0
0.0
0.0
0.0
0.0
0.0
98.0
20.0
16.0
0.0

1.0
1.0
1.0
1.0
1.0
1.0
1.0
1.0
1.0
1.0
1.0
1.0
1.0
1.0
1.0
1.0
1.0
1.0
1.0
1.0
1.0
1.0
1.0
1.0
1.0
1.0
1.0
1.0
1.0
1.0


66.0
0.0
0.0
0.0
0.0
0.0
0.0
0.0
0.0
0.0
0.0
0.0
0.0
0.0
0.0
0.0
0.0
0.0
0.0
0.0
0.0
0.0
0.0
0.0
0.0
0.0
0.0
0.0
0.0
66.0

1.0
1.0
1.0
1.0
1.0
1.0
1.0
1.0
1.0
1.0
1.0
1.0
1.0
1.0
1.0
1.0
1.0
1.0
1.0
1.0
1.0
1.0
1.0
1.0
1.0
1.0
1.0
1.0
1.0
1.0


53.0
0.0
0.0
0.0
0.0
0.0
0.0
0.0
0.0
0.0
0.0
0.0
0.0
0.0
0.0
0.0
53.0
0.0
0.0
0.0
0.0
0.0
0.0
0.0
0.0
0.0
0.0
0.0
0.0
0.0

0.79
0.79
0.79
0.79
0.79
0.79
0.79
0.79
0.79
0.79
0.79
0.79
0.79
0.79
0.79
0.79
0.79
0.79
0.79
0.79
0.79
0.79
0.79
0.79
0.79
0.79
0.79
0.79
0.79
0.79


18.0
0.0
0.0
0.0
0.0
0.0
0.0
0.0
0.0
0.0
0.0
0.0
0.0
0.0
0.0
0.0
0.0
0.0
0.0
0.0
0.0
0.0
0.0
0.0
0.0
0.0
0.0
0.0
18.0
0.0

1.0
1.0
1.0
1.0
1.0
1.0
1.0
1.0
1.0
1.0
1.0
1.0
1.0
1.0
1.0
1.0
1.0
1.0
1.0
1.0
1.0
1.0
1.0
1.0
1.0
1.0
1.0
1.0
1.0
1.0


13.0
13.0
0.0
0.0
0.0
0.0
0.0
0.0
0.0
0.0
0.0
0.0
0.0
0.0
0.0
0.0
0.0
0.0
0.0
0.0
0.0
0.0
0.0
0.0
0.0
0.0
0.0
0.0
0.0
0.0


13.0
13.0
0.0
0.0
0.0
0.0
0.0
0.0
0.0
0.0
0.0
0.0
0.0
0.0
0.0
0.0
0.0
0.0
0.0
0.0
0.0
0.0
0.0
0.0
0.0
0.0
0.0
0.0
0.0
0.0


13.0
13.0
0.0
0.0
0.0
0.0
0.0
0.0
0.0
0.0
0.0
0.0
0.0
0.0
0.0
0.0
0.0
0.0
0.0
0.0
0.0
0.0
0.0
0.0
0.0
0.0
0.0
0.0
0.0
0.0

1.0
1.0
1.0
1.0
1.0
1.0
1.0
1.0
1.0
1.0
1.0
1.0
1.0
1.0
1.0
1.0
1.0
1.0
1.0
1.0
1.0
1.0
1.0
1.0
1.0
1.0
1.0
1.0
1.0
1.0


2455.0
0.0
0.0
0.0
0.0
0.0
0.0
0.0
0.0
0.0
13.0
0.0
0.0
0.0
0.0
0.0
0.0
0.0
0.0
0.0
0.0
0.0
0.0
0.0
0.0
0.0
0.0
0.0
2442.0
0.0


2455.0
0.0
0.0
0.0
0.0
0.0
0.0
0.0
0.0
0.0
13.0
0.0
0.0
0.0
0.0
0.0
0.0
0.0
0.0
0.0
0.0
0.0
0.0
0.0
0.0
0.0
0.0
0.0
2442.0
0.0


13.0
0.0
0.0
0.0
0.0
0.0
0.0
0.0
0.0
0.0
13.0
0.0
0.0
0.0
0.0
0.0
0.0
0.0
0.0
0.0
0.0
0.0
0.0
0.0
0.0
0.0
0.0
0.0
0.0
0.0

1.0
1.0
1.0
1.0
1.0
1.0
1.0
1.0
1.0
1.0
1.0
1.0
1.0
1.0
1.0
1.0
1.0
1.0
1.0
1.0
1.0
1.0
1.0
1.0
1.0
1.0
1.0
1.0
1.0
1.0


2442.0
0.0
0.0
0.0
0.0
0.0
0.0
0.0
0.0
0.0
0.0
0.0
0.0
0.0
0.0
0.0
0.0
0.0
0.0
0.0
0.0
0.0
0.0
0.0
0.0
0.0
0.0
0.0
2442.0
0.0

1.0
1.0
1.0
1.0
1.0
1.0
1.0
1.0
1.0
1.0
1.0
1.0
1.0
1.0
1.0
1.0
1.0
1.0
1.0
1.0
1.0
1.0
1.0
1.0
1.0
1.0
1.0
1.0
1.0
1.0


4536823.0
0.0
0.0
9.0
0.0
0.0
0.0
0.0
0.0
0.0
0.0
0.0
0.0
0.0
2.0
45304.0
12041.0
1.0
13.0
2.0
0.0
0.0
3.0
0.0
0.0
1.0
1258484.0
1200161.0
829272.0
1191530.0

1.0
1.0
1.0
1.0
1.0
1.0
1.0
1.0
1.0
1.0
1.0
1.0
1.0
1.0
1.0
1.0
1.0
1.0
1.0
1.0
1.0
1.0
1.0
1.0
1.0
1.0
1.0
1.0
1.0
1.0


12050.0
0.0
0.0
9.0
0.0
0.0
0.0
0.0
0.0
0.0
0.0
0.0
0.0
0.0
0.0
0.0
12041.0
0.0
0.0
0.0
0.0
0.0
0.0
0.0
0.0
0.0
0.0
0.0
0.0
0.0


12050.0
0.0
0.0
9.0
0.0
0.0
0.0
0.0
0.0
0.0
0.0
0.0
0.0
0.0
0.0
0.0
12041.0
0.0
0.0
0.0
0.0
0.0
0.0
0.0
0.0
0.0
0.0
0.0
0.0
0.0

1.0
1.0
1.0
1.0
1.0
1.0
1.0
1.0
1.0
1.0
1.0
1.0
1.0
1.0
1.0
1.0
1.0
1.0
1.0
1.0
1.0
1.0
1.0
1.0
1.0
1.0
1.0
1.0
1.0
1.0


3348237.0
0.0
0.0
0.0
0.0
0.0
0.0
0.0
0.0
0.0
0.0
0.0
0.0
0.0
2.0
2.0
0.0
1.0
0.0
2.0
0.0
0.0
2.0
0.0
0.0
1.0
430181.0
1200152.0
829256.0
888638.0

0.73
0.73
0.73
0.73
0.73
0.73
0.73
0.73
0.73
0.73
0.73
0.73
0.73
0.73
0.73
0.73
0.73
0.73
0.73
0.73
0.73
0.73
0.73
0.73
0.73
0.73
0.73
0.73
0.73
0.73


3348227.0
0.0
0.0
0.0
0.0
0.0
0.0
0.0
0.0
0.0
0.0
0.0
0.0
0.0
2.0
2.0
0.0
1.0
0.0
2.0
0.0
0.0
2.0
0.0
0.0
1.0
430181.0
1200152.0
829251.0
888633.0

1.0
1.0
1.0
1.0
1.0
1.0
1.0
1.0
1.0
1.0
1.0
1.0
1.0
1.0
1.0
1.0
1.0
1.0
1.0
1.0
1.0
1.0
1.0
1.0
1.0
1.0
1.0
1.0
1.0
1.0


759463.0
0.0
0.0
0.0
0.0
0.0
0.0
0.0
0.0
0.0
0.0
0.0
0.0
0.0
0.0
0.0
0.0
0.0
0.0
0.0
0.0
0.0
1.0
0.0
0.0
0.0
759462.0
0.0
0.0
0.0


759463.0
0.0
0.0
0.0
0.0
0.0
0.0
0.0
0.0
0.0
0.0
0.0
0.0
0.0
0.0
0.0
0.0
0.0
0.0
0.0
0.0
0.0
1.0
0.0
0.0
0.0
759462.0
0.0
0.0
0.0

1.0
1.0
1.0
1.0
1.0
1.0
1.0
1.0
1.0
1.0
1.0
1.0
1.0
1.0
1.0
1.0
1.0
1.0
1.0
1.0
1.0
1.0
1.0
1.0
1.0
1.0
1.0
1.0
1.0
1.0


289667.0
0.0
0.0
0.0
0.0
0.0
0.0
0.0
0.0
0.0
0.0
0.0
0.0
0.0
0.0
0.0
0.0
0.0
0.0
0.0
0.0
0.0
0.0
0.0
0.0
0.0
0.0
0.0
0.0
289667.0


289667.0
0.0
0.0
0.0
0.0
0.0
0.0
0.0
0.0
0.0
0.0
0.0
0.0
0.0
0.0
0.0
0.0
0.0
0.0
0.0
0.0
0.0
0.0
0.0
0.0
0.0
0.0
0.0
0.0
289667.0

0.99
0.99
0.99
0.99
0.99
0.99
0.99
0.99
0.99
0.99
0.99
0.99
0.99
0.99
0.99
0.99
0.99
0.99
0.99
0.99
0.99
0.99
0.99
0.99
0.99
0.99
0.99
0.99
0.99
0.99


47971.0
0.0
0.0
0.0
0.0
0.0
0.0
0.0
0.0
0.0
0.0
0.0
0.0
0.0
0.0
0.0
0.0
0.0
0.0
0.0
0.0
0.0
0.0
0.0
0.0
0.0
47971.0
0.0
0.0
0.0


47971.0
0.0
0.0
0.0
0.0
0.0
0.0
0.0
0.0
0.0
0.0
0.0
0.0
0.0
0.0
0.0
0.0
0.0
0.0
0.0
0.0
0.0
0.0
0.0
0.0
0.0
47971.0
0.0
0.0
0.0

1.0
1.0
1.0
1.0
1.0
1.0
1.0
1.0
1.0
1.0
1.0
1.0
1.0
1.0
1.0
1.0
1.0
1.0
1.0
1.0
1.0
1.0
1.0
1.0
1.0
1.0
1.0
1.0
1.0
1.0


45302.0
0.0
0.0
0.0
0.0
0.0
0.0
0.0
0.0
0.0
0.0
0.0
0.0
0.0
0.0
45302.0
0.0
0.0
0.0
0.0
0.0
0.0
0.0
0.0
0.0
0.0
0.0
0.0
0.0
0.0

1.0
1.0
1.0
1.0
1.0
1.0
1.0
1.0
1.0
1.0
1.0
1.0
1.0
1.0
1.0
1.0
1.0
1.0
1.0
1.0
1.0
1.0
1.0
1.0
1.0
1.0
1.0
1.0
1.0
1.0


17.0
0.0
0.0
0.0
0.0
0.0
0.0
0.0
0.0
0.0
0.0
0.0
0.0
0.0
0.0
17.0
0.0
0.0
0.0
0.0
0.0
0.0
0.0
0.0
0.0
0.0
0.0
0.0
0.0
0.0

0.71
0.71
0.71
0.71
0.71
0.71
0.71
0.71
0.71
0.71
0.71
0.71
0.71
0.71
0.71
0.71
0.71
0.71
0.71
0.71
0.71
0.71
0.71
0.71
0.71
0.71
0.71
0.71
0.71
0.71


31016.0
0.0
0.0
0.0
0.0
0.0
0.0
0.0
0.0
0.0
0.0
0.0
0.0
0.0
0.0
0.0
0.0
0.0
0.0
0.0
0.0
0.0
0.0
0.0
0.0
0.0
17802.0
0.0
0.0
13214.0


16448.0
0.0
0.0
0.0
0.0
0.0
0.0
0.0
0.0
0.0
0.0
0.0
0.0
0.0
0.0
0.0
0.0
0.0
0.0
0.0
0.0
0.0
0.0
0.0
0.0
0.0
3234.0
0.0
0.0
13214.0

0.98
0.98
0.98
0.98
0.98
0.98
0.98
0.98
0.98
0.98
0.98
0.98
0.98
0.98
0.98
0.98
0.98
0.98
0.98
0.98
0.98
0.98
0.98
0.98
0.98
0.98
0.98
0.98
0.98
0.98


14568.0
0.0
0.0
0.0
0.0
0.0
0.0
0.0
0.0
0.0
0.0
0.0
0.0
0.0
0.0
0.0
0.0
0.0
0.0
0.0
0.0
0.0
0.0
0.0
0.0
0.0
14568.0
0.0
0.0
0.0

0.95
0.95
0.95
0.95
0.95
0.95
0.95
0.95
0.95
0.95
0.95
0.95
0.95
0.95
0.95
0.95
0.95
0.95
0.95
0.95
0.95
0.95
0.95
0.95
0.95
0.95
0.95
0.95
0.95
0.95


542081.0
0.0
0.0
0.0
0.0
0.0
0.0
0.0
0.0
0.0
0.0
0.0
0.0
542079.0
0.0
0.0
0.0
0.0
0.0
0.0
0.0
0.0
0.0
0.0
0.0
0.0
0.0
0.0
2.0
0.0


542081.0
0.0
0.0
0.0
0.0
0.0
0.0
0.0
0.0
0.0
0.0
0.0
0.0
542079.0
0.0
0.0
0.0
0.0
0.0
0.0
0.0
0.0
0.0
0.0
0.0
0.0
0.0
0.0
2.0
0.0

0.84
0.84
0.84
0.84
0.84
0.84
0.84
0.84
0.84
0.84
0.84
0.84
0.84
0.84
0.84
0.84
0.84
0.84
0.84
0.84
0.84
0.84
0.84
0.84
0.84
0.84
0.84
0.84
0.84
0.84


42727.0
0.0
0.0
0.0
0.0
0.0
0.0
0.0
0.0
0.0
0.0
0.0
0.0
0.0
0.0
0.0
0.0
0.0
0.0
0.0
0.0
0.0
0.0
0.0
0.0
27251.0
15476.0
0.0
0.0
0.0


42727.0
0.0
0.0
0.0
0.0
0.0
0.0
0.0
0.0
0.0
0.0
0.0
0.0
0.0
0.0
0.0
0.0
0.0
0.0
0.0
0.0
0.0
0.0
0.0
0.0
27251.0
15476.0
0.0
0.0
0.0


42727.0
0.0
0.0
0.0
0.0
0.0
0.0
0.0
0.0
0.0
0.0
0.0
0.0
0.0
0.0
0.0
0.0
0.0
0.0
0.0
0.0
0.0
0.0
0.0
0.0
27251.0
15476.0
0.0
0.0
0.0

1.0
1.0
1.0
1.0
1.0
1.0
1.0
1.0
1.0
1.0
1.0
1.0
1.0
1.0
1.0
1.0
1.0
1.0
1.0
1.0
1.0
1.0
1.0
1.0
1.0
1.0
1.0
1.0
1.0
1.0


67861.0
0.0
0.0
0.0
0.0
0.0
0.0
0.0
0.0
0.0
0.0
0.0
0.0
0.0
0.0
0.0
0.0
0.0
0.0
0.0
0.0
0.0
0.0
0.0
0.0
0.0
67861.0
0.0
0.0
0.0


67861.0
0.0
0.0
0.0
0.0
0.0
0.0
0.0
0.0
0.0
0.0
0.0
0.0
0.0
0.0
0.0
0.0
0.0
0.0
0.0
0.0
0.0
0.0
0.0
0.0
0.0
67861.0
0.0
0.0
0.0

1.0
1.0
1.0
1.0
1.0
1.0
1.0
1.0
1.0
1.0
1.0
1.0
1.0
1.0
1.0
1.0
1.0
1.0
1.0
1.0
1.0
1.0
1.0
1.0
1.0
1.0
1.0
1.0
1.0
1.0


23774.0
0.0
0.0
0.0
0.0
0.0
0.0
0.0
0.0
0.0
0.0
0.0
0.0
0.0
0.0
0.0
0.0
0.0
0.0
0.0
0.0
0.0
0.0
0.0
0.0
17353.0
0.0
6421.0
0.0
0.0


23774.0
0.0
0.0
0.0
0.0
0.0
0.0
0.0
0.0
0.0
0.0
0.0
0.0
0.0
0.0
0.0
0.0
0.0
0.0
0.0
0.0
0.0
0.0
0.0
0.0
17353.0
0.0
6421.0
0.0
0.0


23774.0
0.0
0.0
0.0
0.0
0.0
0.0
0.0
0.0
0.0
0.0
0.0
0.0
0.0
0.0
0.0
0.0
0.0
0.0
0.0
0.0
0.0
0.0
0.0
0.0
17353.0
0.0
6421.0
0.0
0.0

1.0
1.0
1.0
1.0
1.0
1.0
1.0
1.0
1.0
1.0
1.0
1.0
1.0
1.0
1.0
1.0
1.0
1.0
1.0
1.0
1.0
1.0
1.0
1.0
1.0
1.0
1.0
1.0
1.0
1.0


15528.0
0.0
0.0
0.0
0.0
0.0
0.0
0.0
0.0
0.0
0.0
0.0
0.0
1701.0
2064.0
0.0
0.0
0.0
0.0
1484.0
2403.0
584.0
0.0
0.0
643.0
0.0
0.0
2822.0
1214.0
2613.0


15528.0
0.0
0.0
0.0
0.0
0.0
0.0
0.0
0.0
0.0
0.0
0.0
0.0
1701.0
2064.0
0.0
0.0
0.0
0.0
1484.0
2403.0
584.0
0.0
0.0
643.0
0.0
0.0
2822.0
1214.0
2613.0


15528.0
0.0
0.0
0.0
0.0
0.0
0.0
0.0
0.0
0.0
0.0
0.0
0.0
1701.0
2064.0
0.0
0.0
0.0
0.0
1484.0
2403.0
584.0
0.0
0.0
643.0
0.0
0.0
2822.0
1214.0
2613.0

1.0
1.0
1.0
1.0
1.0
1.0
1.0
1.0
1.0
1.0
1.0
1.0
1.0
1.0
1.0
1.0
1.0
1.0
1.0
1.0
1.0
1.0
1.0
1.0
1.0
1.0
1.0
1.0
1.0
1.0


137.0
0.0
0.0
0.0
0.0
0.0
0.0
0.0
0.0
0.0
0.0
0.0
0.0
0.0
0.0
0.0
0.0
0.0
0.0
0.0
0.0
0.0
0.0
0.0
0.0
9.0
116.0
12.0
0.0
0.0

0.73
0.73
0.73
0.73
0.73
0.73
0.73
0.73
0.73
0.73
0.73
0.73
0.73
0.73
0.73
0.73
0.73
0.73
0.73
0.73
0.73
0.73
0.73
0.73
0.73
0.73
0.73
0.73
0.73
0.73


31.0
0.0
0.0
0.0
0.0
0.0
0.0
0.0
0.0
0.0
0.0
0.0
0.0
0.0
0.0
0.0
0.0
0.0
0.0
0.0
0.0
0.0
0.0
0.0
0.0
0.0
31.0
0.0
0.0
0.0


31.0
0.0
0.0
0.0
0.0
0.0
0.0
0.0
0.0
0.0
0.0
0.0
0.0
0.0
0.0
0.0
0.0
0.0
0.0
0.0
0.0
0.0
0.0
0.0
0.0
0.0
31.0
0.0
0.0
0.0


31.0
0.0
0.0
0.0
0.0
0.0
0.0
0.0
0.0
0.0
0.0
0.0
0.0
0.0
0.0
0.0
0.0
0.0
0.0
0.0
0.0
0.0
0.0
0.0
0.0
0.0
31.0
0.0
0.0
0.0

1.0
1.0
1.0
1.0
1.0
1.0
1.0
1.0
1.0
1.0
1.0
1.0
1.0
1.0
1.0
1.0
1.0
1.0
1.0
1.0
1.0
1.0
1.0
1.0
1.0
1.0
1.0
1.0
1.0
1.0


2131252.0
15940.0
70949.0
68190.0
22388.0
33923.0
49672.0
1652.0
4918.0
25013.0
56940.0
37266.0
1493.0
198339.0
90264.0
0.0
58186.0
134159.0
47351.0
77707.0
328970.0
28198.0
26328.0
1493.0
567472.0
18142.0
74010.0
27746.0
39395.0
25148.0

0.96
0.96
0.96
0.96
0.96
0.96
0.96
0.96
0.96
0.96
0.96
0.96
0.96
0.96
0.96
0.96
0.96
0.96
0.96
0.96
0.96
0.96
0.96
0.96
0.96
0.96
0.96
0.96
0.96
0.96


193541.0
7527.0
15675.0
1779.0
6018.0
0.0
357.0
0.0
0.0
1.0
36206.0
14982.0
0.0
45103.0
32030.0
0.0
0.0
0.0
0.0
0.0
0.0
0.0
0.0
0.0
5.0
0.0
26164.0
4755.0
1991.0
948.0

1.0
1.0
1.0
1.0
1.0
1.0
1.0
1.0
1.0
1.0
1.0
1.0
1.0
1.0
1.0
1.0
1.0
1.0
1.0
1.0
1.0
1.0
1.0
1.0
1.0
1.0
1.0
1.0
1.0
1.0


151864.0
6831.0
15135.0
1576.0
0.0
0.0
0.0
0.0
0.0
1.0
36206.0
14982.0
0.0
45103.0
32030.0
0.0
0.0
0.0
0.0
0.0
0.0
0.0
0.0
0.0
0.0
0.0
0.0
0.0
0.0
0.0


151864.0
6831.0
15135.0
1576.0
0.0
0.0
0.0
0.0
0.0
1.0
36206.0
14982.0
0.0
45103.0
32030.0
0.0
0.0
0.0
0.0
0.0
0.0
0.0
0.0
0.0
0.0
0.0
0.0
0.0
0.0
0.0

1.0
1.0
1.0
1.0
1.0
1.0
1.0
1.0
1.0
1.0
1.0
1.0
1.0
1.0
1.0
1.0
1.0
1.0
1.0
1.0
1.0
1.0
1.0
1.0
1.0
1.0
1.0
1.0
1.0
1.0


5592.0
0.0
0.0
0.0
5592.0
0.0
0.0
0.0
0.0
0.0
0.0
0.0
0.0
0.0
0.0
0.0
0.0
0.0
0.0
0.0
0.0
0.0
0.0
0.0
0.0
0.0
0.0
0.0
0.0
0.0


5592.0
0.0
0.0
0.0
5592.0
0.0
0.0
0.0
0.0
0.0
0.0
0.0
0.0
0.0
0.0
0.0
0.0
0.0
0.0
0.0
0.0
0.0
0.0
0.0
0.0
0.0
0.0
0.0
0.0
0.0

0.99
0.99
0.99
0.99
0.99
0.99
0.99
0.99
0.99
0.99
0.99
0.99
0.99
0.99
0.99
0.99
0.99
0.99
0.99
0.99
0.99
0.99
0.99
0.99
0.99
0.99
0.99
0.99
0.99
0.99


35875.0
696.0
423.0
203.0
338.0
0.0
357.0
0.0
0.0
0.0
0.0
0.0
0.0
0.0
0.0
0.0
0.0
0.0
0.0
0.0
0.0
0.0
0.0
0.0
0.0
0.0
26164.0
4755.0
1991.0
948.0


35875.0
696.0
423.0
203.0
338.0
0.0
357.0
0.0
0.0
0.0
0.0
0.0
0.0
0.0
0.0
0.0
0.0
0.0
0.0
0.0
0.0
0.0
0.0
0.0
0.0
0.0
26164.0
4755.0
1991.0
948.0

1.0
1.0
1.0
1.0
1.0
1.0
1.0
1.0
1.0
1.0
1.0
1.0
1.0
1.0
1.0
1.0
1.0
1.0
1.0
1.0
1.0
1.0
1.0
1.0
1.0
1.0
1.0
1.0
1.0
1.0


117.0
0.0
117.0
0.0
0.0
0.0
0.0
0.0
0.0
0.0
0.0
0.0
0.0
0.0
0.0
0.0
0.0
0.0
0.0
0.0
0.0
0.0
0.0
0.0
0.0
0.0
0.0
0.0
0.0
0.0


117.0
0.0
117.0
0.0
0.0
0.0
0.0
0.0
0.0
0.0
0.0
0.0
0.0
0.0
0.0
0.0
0.0
0.0
0.0
0.0
0.0
0.0
0.0
0.0
0.0
0.0
0.0
0.0
0.0
0.0

1.0
1.0
1.0
1.0
1.0
1.0
1.0
1.0
1.0
1.0
1.0
1.0
1.0
1.0
1.0
1.0
1.0
1.0
1.0
1.0
1.0
1.0
1.0
1.0
1.0
1.0
1.0
1.0
1.0
1.0


65858.0
1.0
7236.0
10655.0
13822.0
17031.0
5966.0
0.0
0.0
0.0
0.0
55.0
0.0
867.0
7430.0
0.0
0.0
0.0
0.0
0.0
0.0
0.0
0.0
0.0
2795.0
0.0
0.0
0.0
0.0
0.0

1.0
1.0
1.0
1.0
1.0
1.0
1.0
1.0
1.0
1.0
1.0
1.0
1.0
1.0
1.0
1.0
1.0
1.0
1.0
1.0
1.0
1.0
1.0
1.0
1.0
1.0
1.0
1.0
1.0
1.0


60697.0
1.0
7236.0
10655.0
13822.0
17031.0
805.0
0.0
0.0
0.0
0.0
55.0
0.0
867.0
7430.0
0.0
0.0
0.0
0.0
0.0
0.0
0.0
0.0
0.0
2795.0
0.0
0.0
0.0
0.0
0.0

1.0
1.0
1.0
1.0
1.0
1.0
1.0
1.0
1.0
1.0
1.0
1.0
1.0
1.0
1.0
1.0
1.0
1.0
1.0
1.0
1.0
1.0
1.0
1.0
1.0
1.0
1.0
1.0
1.0
1.0


31086.0
1.0
6463.0
10026.0
8146.0
6450.0
0.0
0.0
0.0
0.0
0.0
0.0
0.0
0.0
0.0
0.0
0.0
0.0
0.0
0.0
0.0
0.0
0.0
0.0
0.0
0.0
0.0
0.0
0.0
0.0

1.0
1.0
1.0
1.0
1.0
1.0
1.0
1.0
1.0
1.0
1.0
1.0
1.0
1.0
1.0
1.0
1.0
1.0
1.0
1.0
1.0
1.0
1.0
1.0
1.0
1.0
1.0
1.0
1.0
1.0


15755.0
0.0
79.0
266.0
3942.0
1864.0
146.0
0.0
0.0
0.0
0.0
0.0
0.0
867.0
5796.0
0.0
0.0
0.0
0.0
0.0
0.0
0.0
0.0
0.0
2795.0
0.0
0.0
0.0
0.0
0.0

0.96
0.96
0.96
0.96
0.96
0.96
0.96
0.96
0.96
0.96
0.96
0.96
0.96
0.96
0.96
0.96
0.96
0.96
0.96
0.96
0.96
0.96
0.96
0.96
0.96
0.96
0.96
0.96
0.96
0.96


11449.0
0.0
540.0
363.0
1659.0
8228.0
659.0
0.0
0.0
0.0
0.0
0.0
0.0
0.0
0.0
0.0
0.0
0.0
0.0
0.0
0.0
0.0
0.0
0.0
0.0
0.0
0.0
0.0
0.0
0.0

1.0
1.0
1.0
1.0
1.0
1.0
1.0
1.0
1.0
1.0
1.0
1.0
1.0
1.0
1.0
1.0
1.0
1.0
1.0
1.0
1.0
1.0
1.0
1.0
1.0
1.0
1.0
1.0
1.0
1.0


736.0
0.0
97.0
0.0
67.0
361.0
0.0
0.0
0.0
0.0
0.0
0.0
0.0
0.0
211.0
0.0
0.0
0.0
0.0
0.0
0.0
0.0
0.0
0.0
0.0
0.0
0.0
0.0
0.0
0.0

1.0
1.0
1.0
1.0
1.0
1.0
1.0
1.0
1.0
1.0
1.0
1.0
1.0
1.0
1.0
1.0
1.0
1.0
1.0
1.0
1.0
1.0
1.0
1.0
1.0
1.0
1.0
1.0
1.0
1.0


185.0
0.0
57.0
0.0
0.0
128.0
0.0
0.0
0.0
0.0
0.0
0.0
0.0
0.0
0.0
0.0
0.0
0.0
0.0
0.0
0.0
0.0
0.0
0.0
0.0
0.0
0.0
0.0
0.0
0.0

1.0
1.0
1.0
1.0
1.0
1.0
1.0
1.0
1.0
1.0
1.0
1.0
1.0
1.0
1.0
1.0
1.0
1.0
1.0
1.0
1.0
1.0
1.0
1.0
1.0
1.0
1.0
1.0
1.0
1.0


55.0
0.0
0.0
0.0
0.0
0.0
0.0
0.0
0.0
0.0
0.0
55.0
0.0
0.0
0.0
0.0
0.0
0.0
0.0
0.0
0.0
0.0
0.0
0.0
0.0
0.0
0.0
0.0
0.0
0.0

1.0
1.0
1.0
1.0
1.0
1.0
1.0
1.0
1.0
1.0
1.0
1.0
1.0
1.0
1.0
1.0
1.0
1.0
1.0
1.0
1.0
1.0
1.0
1.0
1.0
1.0
1.0
1.0
1.0
1.0


1423.0
0.0
0.0
0.0
0.0
0.0
0.0
0.0
0.0
0.0
0.0
0.0
0.0
0.0
1423.0
0.0
0.0
0.0
0.0
0.0
0.0
0.0
0.0
0.0
0.0
0.0
0.0
0.0
0.0
0.0

1.0
1.0
1.0
1.0
1.0
1.0
1.0
1.0
1.0
1.0
1.0
1.0
1.0
1.0
1.0
1.0
1.0
1.0
1.0
1.0
1.0
1.0
1.0
1.0
1.0
1.0
1.0
1.0
1.0
1.0


1627.0
0.0
0.0
0.0
0.0
0.0
1627.0
0.0
0.0
0.0
0.0
0.0
0.0
0.0
0.0
0.0
0.0
0.0
0.0
0.0
0.0
0.0
0.0
0.0
0.0
0.0
0.0
0.0
0.0
0.0

0.91
0.91
0.91
0.91
0.91
0.91
0.91
0.91
0.91
0.91
0.91
0.91
0.91
0.91
0.91
0.91
0.91
0.91
0.91
0.91
0.91
0.91
0.91
0.91
0.91
0.91
0.91
0.91
0.91
0.91


50085.0
0.0
0.0
0.0
0.0
13518.0
36567.0
0.0
0.0
0.0
0.0
0.0
0.0
0.0
0.0
0.0
0.0
0.0
0.0
0.0
0.0
0.0
0.0
0.0
0.0
0.0
0.0
0.0
0.0
0.0

0.97
0.97
0.97
0.97
0.97
0.97
0.97
0.97
0.97
0.97
0.97
0.97
0.97
0.97
0.97
0.97
0.97
0.97
0.97
0.97
0.97
0.97
0.97
0.97
0.97
0.97
0.97
0.97
0.97
0.97


49986.0
0.0
0.0
0.0
0.0
13467.0
36519.0
0.0
0.0
0.0
0.0
0.0
0.0
0.0
0.0
0.0
0.0
0.0
0.0
0.0
0.0
0.0
0.0
0.0
0.0
0.0
0.0
0.0
0.0
0.0

0.95
0.95
0.95
0.95
0.95
0.95
0.95
0.95
0.95
0.95
0.95
0.95
0.95
0.95
0.95
0.95
0.95
0.95
0.95
0.95
0.95
0.95
0.95
0.95
0.95
0.95
0.95
0.95
0.95
0.95


73743.0
2.0
30635.0
36065.0
11.0
2.0
0.0
0.0
0.0
0.0
0.0
7011.0
0.0
0.0
0.0
0.0
0.0
17.0
0.0
0.0
0.0
0.0
0.0
0.0
0.0
0.0
0.0
0.0
0.0
0.0


73726.0
2.0
30635.0
36065.0
11.0
2.0
0.0
0.0
0.0
0.0
0.0
7011.0
0.0
0.0
0.0
0.0
0.0
0.0
0.0
0.0
0.0
0.0
0.0
0.0
0.0
0.0
0.0
0.0
0.0
0.0

1.0
1.0
1.0
1.0
1.0
1.0
1.0
1.0
1.0
1.0
1.0
1.0
1.0
1.0
1.0
1.0
1.0
1.0
1.0
1.0
1.0
1.0
1.0
1.0
1.0
1.0
1.0
1.0
1.0
1.0


73713.0
2.0
30623.0
36064.0
11.0
2.0
0.0
0.0
0.0
0.0
0.0
7011.0
0.0
0.0
0.0
0.0
0.0
0.0
0.0
0.0
0.0
0.0
0.0
0.0
0.0
0.0
0.0
0.0
0.0
0.0

1.0
1.0
1.0
1.0
1.0
1.0
1.0
1.0
1.0
1.0
1.0
1.0
1.0
1.0
1.0
1.0
1.0
1.0
1.0
1.0
1.0
1.0
1.0
1.0
1.0
1.0
1.0
1.0
1.0
1.0


17.0
0.0
0.0
0.0
0.0
0.0
0.0
0.0
0.0
0.0
0.0
0.0
0.0
0.0
0.0
0.0
0.0
17.0
0.0
0.0
0.0
0.0
0.0
0.0
0.0
0.0
0.0
0.0
0.0
0.0


17.0
0.0
0.0
0.0
0.0
0.0
0.0
0.0
0.0
0.0
0.0
0.0
0.0
0.0
0.0
0.0
0.0
17.0
0.0
0.0
0.0
0.0
0.0
0.0
0.0
0.0
0.0
0.0
0.0
0.0

1.0
1.0
1.0
1.0
1.0
1.0
1.0
1.0
1.0
1.0
1.0
1.0
1.0
1.0
1.0
1.0
1.0
1.0
1.0
1.0
1.0
1.0
1.0
1.0
1.0
1.0
1.0
1.0
1.0
1.0


17270.0
0.0
2470.0
597.0
0.0
0.0
0.0
0.0
0.0
0.0
8850.0
5162.0
0.0
191.0
0.0
0.0
0.0
0.0
0.0
0.0
0.0
0.0
0.0
0.0
0.0
0.0
0.0
0.0
0.0
0.0


16632.0
0.0
2198.0
386.0
0.0
0.0
0.0
0.0
0.0
0.0
8850.0
5007.0
0.0
191.0
0.0
0.0
0.0
0.0
0.0
0.0
0.0
0.0
0.0
0.0
0.0
0.0
0.0
0.0
0.0
0.0

0.99
0.99
0.99
0.99
0.99
0.99
0.99
0.99
0.99
0.99
0.99
0.99
0.99
0.99
0.99
0.99
0.99
0.99
0.99
0.99
0.99
0.99
0.99
0.99
0.99
0.99
0.99
0.99
0.99
0.99


619.0
0.0
272.0
211.0
0.0
0.0
0.0
0.0
0.0
0.0
0.0
136.0
0.0
0.0
0.0
0.0
0.0
0.0
0.0
0.0
0.0
0.0
0.0
0.0
0.0
0.0
0.0
0.0
0.0
0.0


619.0
0.0
272.0
211.0
0.0
0.0
0.0
0.0
0.0
0.0
0.0
136.0
0.0
0.0
0.0
0.0
0.0
0.0
0.0
0.0
0.0
0.0
0.0
0.0
0.0
0.0
0.0
0.0
0.0
0.0

1.0
1.0
1.0
1.0
1.0
1.0
1.0
1.0
1.0
1.0
1.0
1.0
1.0
1.0
1.0
1.0
1.0
1.0
1.0
1.0
1.0
1.0
1.0
1.0
1.0
1.0
1.0
1.0
1.0
1.0


19.0
0.0
0.0
0.0
0.0
0.0
0.0
0.0
0.0
0.0
0.0
19.0
0.0
0.0
0.0
0.0
0.0
0.0
0.0
0.0
0.0
0.0
0.0
0.0
0.0
0.0
0.0
0.0
0.0
0.0


19.0
0.0
0.0
0.0
0.0
0.0
0.0
0.0
0.0
0.0
0.0
19.0
0.0
0.0
0.0
0.0
0.0
0.0
0.0
0.0
0.0
0.0
0.0
0.0
0.0
0.0
0.0
0.0
0.0
0.0

1.0
1.0
1.0
1.0
1.0
1.0
1.0
1.0
1.0
1.0
1.0
1.0
1.0
1.0
1.0
1.0
1.0
1.0
1.0
1.0
1.0
1.0
1.0
1.0
1.0
1.0
1.0
1.0
1.0
1.0


20164.0
1060.0
346.0
0.0
0.0
0.0
0.0
267.0
607.0
10016.0
4149.0
1129.0
0.0
0.0
97.0
0.0
0.0
2493.0
0.0
0.0
0.0
0.0
0.0
0.0
0.0
0.0
0.0
0.0
0.0
0.0

0.99
0.99
0.99
0.99
0.99
0.99
0.99
0.99
0.99
0.99
0.99
0.99
0.99
0.99
0.99
0.99
0.99
0.99
0.99
0.99
0.99
0.99
0.99
0.99
0.99
0.99
0.99
0.99
0.99
0.99


16356.0
937.0
125.0
0.0
0.0
0.0
0.0
267.0
603.0
9279.0
4028.0
1117.0
0.0
0.0
0.0
0.0
0.0
0.0
0.0
0.0
0.0
0.0
0.0
0.0
0.0
0.0
0.0
0.0
0.0
0.0

0.96
0.96
0.96
0.96
0.96
0.96
0.96
0.96
0.96
0.96
0.96
0.96
0.96
0.96
0.96
0.96
0.96
0.96
0.96
0.96
0.96
0.96
0.96
0.96
0.96
0.96
0.96
0.96
0.96
0.96


582.0
0.0
61.0
0.0
0.0
0.0
0.0
0.0
0.0
521.0
0.0
0.0
0.0
0.0
0.0
0.0
0.0
0.0
0.0
0.0
0.0
0.0
0.0
0.0
0.0
0.0
0.0
0.0
0.0
0.0

0.94
0.94
0.94
0.94
0.94
0.94
0.94
0.94
0.94
0.94
0.94
0.94
0.94
0.94
0.94
0.94
0.94
0.94
0.94
0.94
0.94
0.94
0.94
0.94
0.94
0.94
0.94
0.94
0.94
0.94


135.0
42.0
0.0
0.0
0.0
0.0
0.0
0.0
0.0
0.0
93.0
0.0
0.0
0.0
0.0
0.0
0.0
0.0
0.0
0.0
0.0
0.0
0.0
0.0
0.0
0.0
0.0
0.0
0.0
0.0


135.0
42.0
0.0
0.0
0.0
0.0
0.0
0.0
0.0
0.0
93.0
0.0
0.0
0.0
0.0
0.0
0.0
0.0
0.0
0.0
0.0
0.0
0.0
0.0
0.0
0.0
0.0
0.0
0.0
0.0

0.94
0.94
0.94
0.94
0.94
0.94
0.94
0.94
0.94
0.94
0.94
0.94
0.94
0.94
0.94
0.94
0.94
0.94
0.94
0.94
0.94
0.94
0.94
0.94
0.94
0.94
0.94
0.94
0.94
0.94


55.0
8.0
4.0
0.0
0.0
0.0
0.0
0.0
0.0
9.0
23.0
11.0
0.0
0.0
0.0
0.0
0.0
0.0
0.0
0.0
0.0
0.0
0.0
0.0
0.0
0.0
0.0
0.0
0.0
0.0

1.0
1.0
1.0
1.0
1.0
1.0
1.0
1.0
1.0
1.0
1.0
1.0
1.0
1.0
1.0
1.0
1.0
1.0
1.0
1.0
1.0
1.0
1.0
1.0
1.0
1.0
1.0
1.0
1.0
1.0


46949.0
185.0
1001.0
3947.0
804.0
449.0
125.0
0.0
0.0
298.0
595.0
473.0
0.0
5109.0
3196.0
0.0
0.0
0.0
0.0
0.0
0.0
0.0
0.0
0.0
30767.0
0.0
0.0
0.0
0.0
0.0

0.93
0.93
0.93
0.93
0.93
0.93
0.93
0.93
0.93
0.93
0.93
0.93
0.93
0.93
0.93
0.93
0.93
0.93
0.93
0.93
0.93
0.93
0.93
0.93
0.93
0.93
0.93
0.93
0.93
0.93


2114.0
51.0
312.0
856.0
149.0
86.0
66.0
0.0
0.0
298.0
284.0
12.0
0.0
0.0
0.0
0.0
0.0
0.0
0.0
0.0
0.0
0.0
0.0
0.0
0.0
0.0
0.0
0.0
0.0
0.0

0.8
0.8
0.8
0.8
0.8
0.8
0.8
0.8
0.8
0.8
0.8
0.8
0.8
0.8
0.8
0.8
0.8
0.8
0.8
0.8
0.8
0.8
0.8
0.8
0.8
0.8
0.8
0.8
0.8
0.8


36016.0
0.0
0.0
0.0
0.0
0.0
0.0
0.0
0.0
0.0
0.0
0.0
0.0
2053.0
3196.0
0.0
0.0
0.0
0.0
0.0
0.0
0.0
0.0
0.0
30767.0
0.0
0.0
0.0
0.0
0.0

0.99
0.99
0.99
0.99
0.99
0.99
0.99
0.99
0.99
0.99
0.99
0.99
0.99
0.99
0.99
0.99
0.99
0.99
0.99
0.99
0.99
0.99
0.99
0.99
0.99
0.99
0.99
0.99
0.99
0.99


261804.0
425.0
5299.0
1086.0
604.0
97.0
647.0
312.0
522.0
37.0
256.0
558.0
1493.0
95445.0
300.0
0.0
57112.0
592.0
4272.0
8004.0
0.0
0.0
5716.0
1493.0
0.0
6071.0
11529.0
4751.0
37388.0
17795.0

1.0
1.0
1.0
1.0
1.0
1.0
1.0
1.0
1.0
1.0
1.0
1.0
1.0
1.0
1.0
1.0
1.0
1.0
1.0
1.0
1.0
1.0
1.0
1.0
1.0
1.0
1.0
1.0
1.0
1.0


4813.0
0.0
4813.0
0.0
0.0
0.0
0.0
0.0
0.0
0.0
0.0
0.0
0.0
0.0
0.0
0.0
0.0
0.0
0.0
0.0
0.0
0.0
0.0
0.0
0.0
0.0
0.0
0.0
0.0
0.0

1.0
1.0
1.0
1.0
1.0
1.0
1.0
1.0
1.0
1.0
1.0
1.0
1.0
1.0
1.0
1.0
1.0
1.0
1.0
1.0
1.0
1.0
1.0
1.0
1.0
1.0
1.0
1.0
1.0
1.0


9585.0
0.0
13.0
37.0
46.0
0.0
0.0
312.0
493.0
0.0
0.0
0.0
1484.0
0.0
0.0
0.0
0.0
0.0
0.0
0.0
0.0
0.0
5716.0
1484.0
0.0
0.0
0.0
0.0
0.0
0.0


9489.0
0.0
0.0
0.0
0.0
0.0
0.0
312.0
493.0
0.0
0.0
0.0
1484.0
0.0
0.0
0.0
0.0
0.0
0.0
0.0
0.0
0.0
5716.0
1484.0
0.0
0.0
0.0
0.0
0.0
0.0

1.0
1.0
1.0
1.0
1.0
1.0
1.0
1.0
1.0
1.0
1.0
1.0
1.0
1.0
1.0
1.0
1.0
1.0
1.0
1.0
1.0
1.0
1.0
1.0
1.0
1.0
1.0
1.0
1.0
1.0


96.0
0.0
13.0
37.0
46.0
0.0
0.0
0.0
0.0
0.0
0.0
0.0
0.0
0.0
0.0
0.0
0.0
0.0
0.0
0.0
0.0
0.0
0.0
0.0
0.0
0.0
0.0
0.0
0.0
0.0

1.0
1.0
1.0
1.0
1.0
1.0
1.0
1.0
1.0
1.0
1.0
1.0
1.0
1.0
1.0
1.0
1.0
1.0
1.0
1.0
1.0
1.0
1.0
1.0
1.0
1.0
1.0
1.0
1.0
1.0


4084.0
0.0
0.0
0.0
552.0
0.0
0.0
0.0
0.0
0.0
155.0
196.0
0.0
0.0
0.0
0.0
0.0
0.0
0.0
0.0
0.0
0.0
0.0
0.0
0.0
0.0
0.0
0.0
3181.0
0.0

1.0
1.0
1.0
1.0
1.0
1.0
1.0
1.0
1.0
1.0
1.0
1.0
1.0
1.0
1.0
1.0
1.0
1.0
1.0
1.0
1.0
1.0
1.0
1.0
1.0
1.0
1.0
1.0
1.0
1.0


351.0
0.0
0.0
0.0
0.0
0.0
0.0
0.0
0.0
0.0
155.0
196.0
0.0
0.0
0.0
0.0
0.0
0.0
0.0
0.0
0.0
0.0
0.0
0.0
0.0
0.0
0.0
0.0
0.0
0.0

1.0
1.0
1.0
1.0
1.0
1.0
1.0
1.0
1.0
1.0
1.0
1.0
1.0
1.0
1.0
1.0
1.0
1.0
1.0
1.0
1.0
1.0
1.0
1.0
1.0
1.0
1.0
1.0
1.0
1.0


52813.0
16.0
47.0
0.0
0.0
0.0
626.0
0.0
0.0
0.0
0.0
0.0
0.0
0.0
0.0
0.0
21493.0
0.0
0.0
0.0
0.0
0.0
0.0
0.0
0.0
0.0
739.0
1083.0
26106.0
2703.0


52725.0
0.0
0.0
0.0
0.0
0.0
601.0
0.0
0.0
0.0
0.0
0.0
0.0
0.0
0.0
0.0
21493.0
0.0
0.0
0.0
0.0
0.0
0.0
0.0
0.0
0.0
739.0
1083.0
26106.0
2703.0

1.0
1.0
1.0
1.0
1.0
1.0
1.0
1.0
1.0
1.0
1.0
1.0
1.0
1.0
1.0
1.0
1.0
1.0
1.0
1.0
1.0
1.0
1.0
1.0
1.0
1.0
1.0
1.0
1.0
1.0


47.0
0.0
47.0
0.0
0.0
0.0
0.0
0.0
0.0
0.0
0.0
0.0
0.0
0.0
0.0
0.0
0.0
0.0
0.0
0.0
0.0
0.0
0.0
0.0
0.0
0.0
0.0
0.0
0.0
0.0

1.0
1.0
1.0
1.0
1.0
1.0
1.0
1.0
1.0
1.0
1.0
1.0
1.0
1.0
1.0
1.0
1.0
1.0
1.0
1.0
1.0
1.0
1.0
1.0
1.0
1.0
1.0
1.0
1.0
1.0


25.0
0.0
0.0
0.0
0.0
0.0
25.0
0.0
0.0
0.0
0.0
0.0
0.0
0.0
0.0
0.0
0.0
0.0
0.0
0.0
0.0
0.0
0.0
0.0
0.0
0.0
0.0
0.0
0.0
0.0

1.0
1.0
1.0
1.0
1.0
1.0
1.0
1.0
1.0
1.0
1.0
1.0
1.0
1.0
1.0
1.0
1.0
1.0
1.0
1.0
1.0
1.0
1.0
1.0
1.0
1.0
1.0
1.0
1.0
1.0


16.0
16.0
0.0
0.0
0.0
0.0
0.0
0.0
0.0
0.0
0.0
0.0
0.0
0.0
0.0
0.0
0.0
0.0
0.0
0.0
0.0
0.0
0.0
0.0
0.0
0.0
0.0
0.0
0.0
0.0

1.0
1.0
1.0
1.0
1.0
1.0
1.0
1.0
1.0
1.0
1.0
1.0
1.0
1.0
1.0
1.0
1.0
1.0
1.0
1.0
1.0
1.0
1.0
1.0
1.0
1.0
1.0
1.0
1.0
1.0


211.0
0.0
211.0
0.0
0.0
0.0
0.0
0.0
0.0
0.0
0.0
0.0
0.0
0.0
0.0
0.0
0.0
0.0
0.0
0.0
0.0
0.0
0.0
0.0
0.0
0.0
0.0
0.0
0.0
0.0


211.0
0.0
211.0
0.0
0.0
0.0
0.0
0.0
0.0
0.0
0.0
0.0
0.0
0.0
0.0
0.0
0.0
0.0
0.0
0.0
0.0
0.0
0.0
0.0
0.0
0.0
0.0
0.0
0.0
0.0

1.0
1.0
1.0
1.0
1.0
1.0
1.0
1.0
1.0
1.0
1.0
1.0
1.0
1.0
1.0
1.0
1.0
1.0
1.0
1.0
1.0
1.0
1.0
1.0
1.0
1.0
1.0
1.0
1.0
1.0


689.0
208.0
42.0
23.0
0.0
0.0
0.0
0.0
0.0
0.0
101.0
315.0
0.0
0.0
0.0
0.0
0.0
0.0
0.0
0.0
0.0
0.0
0.0
0.0
0.0
0.0
0.0
0.0
0.0
0.0

1.0
1.0
1.0
1.0
1.0
1.0
1.0
1.0
1.0
1.0
1.0
1.0
1.0
1.0
1.0
1.0
1.0
1.0
1.0
1.0
1.0
1.0
1.0
1.0
1.0
1.0
1.0
1.0
1.0
1.0


482.0
182.0
18.0
0.0
0.0
0.0
0.0
0.0
0.0
0.0
101.0
181.0
0.0
0.0
0.0
0.0
0.0
0.0
0.0
0.0
0.0
0.0
0.0
0.0
0.0
0.0
0.0
0.0
0.0
0.0

1.0
1.0
1.0
1.0
1.0
1.0
1.0
1.0
1.0
1.0
1.0
1.0
1.0
1.0
1.0
1.0
1.0
1.0
1.0
1.0
1.0
1.0
1.0
1.0
1.0
1.0
1.0
1.0
1.0
1.0


23.0
0.0
0.0
23.0
0.0
0.0
0.0
0.0
0.0
0.0
0.0
0.0
0.0
0.0
0.0
0.0
0.0
0.0
0.0
0.0
0.0
0.0
0.0
0.0
0.0
0.0
0.0
0.0
0.0
0.0

1.0
1.0
1.0
1.0
1.0
1.0
1.0
1.0
1.0
1.0
1.0
1.0
1.0
1.0
1.0
1.0
1.0
1.0
1.0
1.0
1.0
1.0
1.0
1.0
1.0
1.0
1.0
1.0
1.0
1.0


231.0
87.0
0.0
0.0
0.0
97.0
0.0
0.0
0.0
0.0
0.0
47.0
0.0
0.0
0.0
0.0
0.0
0.0
0.0
0.0
0.0
0.0
0.0
0.0
0.0
0.0
0.0
0.0
0.0
0.0

1.0
1.0
1.0
1.0
1.0
1.0
1.0
1.0
1.0
1.0
1.0
1.0
1.0
1.0
1.0
1.0
1.0
1.0
1.0
1.0
1.0
1.0
1.0
1.0
1.0
1.0
1.0
1.0
1.0
1.0


106.0
106.0
0.0
0.0
0.0
0.0
0.0
0.0
0.0
0.0
0.0
0.0
0.0
0.0
0.0
0.0
0.0
0.0
0.0
0.0
0.0
0.0
0.0
0.0
0.0
0.0
0.0
0.0
0.0
0.0


106.0
106.0
0.0
0.0
0.0
0.0
0.0
0.0
0.0
0.0
0.0
0.0
0.0
0.0
0.0
0.0
0.0
0.0
0.0
0.0
0.0
0.0
0.0
0.0
0.0
0.0
0.0
0.0
0.0
0.0

0.85
0.85
0.85
0.85
0.85
0.85
0.85
0.85
0.85
0.85
0.85
0.85
0.85
0.85
0.85
0.85
0.85
0.85
0.85
0.85
0.85
0.85
0.85
0.85
0.85
0.85
0.85
0.85
0.85
0.85


47990.0
0.0
133.0
0.0
0.0
0.0
0.0
0.0
0.0
0.0
0.0
0.0
0.0
0.0
0.0
0.0
35619.0
0.0
4272.0
7966.0
0.0
0.0
0.0
0.0
0.0
0.0
0.0
0.0
0.0
0.0


47990.0
0.0
133.0
0.0
0.0
0.0
0.0
0.0
0.0
0.0
0.0
0.0
0.0
0.0
0.0
0.0
35619.0
0.0
4272.0
7966.0
0.0
0.0
0.0
0.0
0.0
0.0
0.0
0.0
0.0
0.0

1.0
1.0
1.0
1.0
1.0
1.0
1.0
1.0
1.0
1.0
1.0
1.0
1.0
1.0
1.0
1.0
1.0
1.0
1.0
1.0
1.0
1.0
1.0
1.0
1.0
1.0
1.0
1.0
1.0
1.0


35386.0
8.0
39.0
93.0
6.0
0.0
21.0
0.0
29.0
37.0
0.0
0.0
0.0
2373.0
300.0
0.0
0.0
592.0
0.0
0.0
0.0
0.0
0.0
0.0
0.0
6071.0
0.0
3668.0
7123.0
15026.0

1.0
1.0
1.0
1.0
1.0
1.0
1.0
1.0
1.0
1.0
1.0
1.0
1.0
1.0
1.0
1.0
1.0
1.0
1.0
1.0
1.0
1.0
1.0
1.0
1.0
1.0
1.0
1.0
1.0
1.0


2788.0
8.0
13.0
59.0
6.0
0.0
0.0
0.0
29.0
0.0
0.0
0.0
0.0
2373.0
300.0
0.0
0.0
0.0
0.0
0.0
0.0
0.0
0.0
0.0
0.0
0.0
0.0
0.0
0.0
0.0

1.0
1.0
1.0
1.0
1.0
1.0
1.0
1.0
1.0
1.0
1.0
1.0
1.0
1.0
1.0
1.0
1.0
1.0
1.0
1.0
1.0
1.0
1.0
1.0
1.0
1.0
1.0
1.0
1.0
1.0


31946.0
0.0
0.0
0.0
0.0
0.0
21.0
0.0
0.0
37.0
0.0
0.0
0.0
0.0
0.0
0.0
0.0
0.0
0.0
0.0
0.0
0.0
0.0
0.0
0.0
6071.0
0.0
3668.0
7123.0
15026.0

1.0
1.0
1.0
1.0
1.0
1.0
1.0
1.0
1.0
1.0
1.0
1.0
1.0
1.0
1.0
1.0
1.0
1.0
1.0
1.0
1.0
1.0
1.0
1.0
1.0
1.0
1.0
1.0
1.0
1.0


34.0
0.0
0.0
34.0
0.0
0.0
0.0
0.0
0.0
0.0
0.0
0.0
0.0
0.0
0.0
0.0
0.0
0.0
0.0
0.0
0.0
0.0
0.0
0.0
0.0
0.0
0.0
0.0
0.0
0.0

1.0
1.0
1.0
1.0
1.0
1.0
1.0
1.0
1.0
1.0
1.0
1.0
1.0
1.0
1.0
1.0
1.0
1.0
1.0
1.0
1.0
1.0
1.0
1.0
1.0
1.0
1.0
1.0
1.0
1.0


9.0
0.0
9.0
0.0
0.0
0.0
0.0
0.0
0.0
0.0
0.0
0.0
0.0
0.0
0.0
0.0
0.0
0.0
0.0
0.0
0.0
0.0
0.0
0.0
0.0
0.0
0.0
0.0
0.0
0.0

0.94
0.94
0.94
0.94
0.94
0.94
0.94
0.94
0.94
0.94
0.94
0.94
0.94
0.94
0.94
0.94
0.94
0.94
0.94
0.94
0.94
0.94
0.94
0.94
0.94
0.94
0.94
0.94
0.94
0.94


592.0
0.0
0.0
0.0
0.0
0.0
0.0
0.0
0.0
0.0
0.0
0.0
0.0
0.0
0.0
0.0
0.0
592.0
0.0
0.0
0.0
0.0
0.0
0.0
0.0
0.0
0.0
0.0
0.0
0.0

1.0
1.0
1.0
1.0
1.0
1.0
1.0
1.0
1.0
1.0
1.0
1.0
1.0
1.0
1.0
1.0
1.0
1.0
1.0
1.0
1.0
1.0
1.0
1.0
1.0
1.0
1.0
1.0
1.0
1.0


93072.0
0.0
0.0
0.0
0.0
0.0
0.0
0.0
0.0
0.0
0.0
0.0
0.0
93072.0
0.0
0.0
0.0
0.0
0.0
0.0
0.0
0.0
0.0
0.0
0.0
0.0
0.0
0.0
0.0
0.0


93072.0
0.0
0.0
0.0
0.0
0.0
0.0
0.0
0.0
0.0
0.0
0.0
0.0
93072.0
0.0
0.0
0.0
0.0
0.0
0.0
0.0
0.0
0.0
0.0
0.0
0.0
0.0
0.0
0.0
0.0

0.84
0.84
0.84
0.84
0.84
0.84
0.84
0.84
0.84
0.84
0.84
0.84
0.84
0.84
0.84
0.84
0.84
0.84
0.84
0.84
0.84
0.84
0.84
0.84
0.84
0.84
0.84
0.84
0.84
0.84


11834.0
0.0
0.0
0.0
0.0
0.0
0.0
0.0
0.0
0.0
0.0
0.0
0.0
0.0
0.0
0.0
0.0
0.0
0.0
0.0
0.0
0.0
0.0
0.0
0.0
0.0
10790.0
0.0
978.0
66.0

0.77
0.77
0.77
0.77
0.77
0.77
0.77
0.77
0.77
0.77
0.77
0.77
0.77
0.77
0.77
0.77
0.77
0.77
0.77
0.77
0.77
0.77
0.77
0.77
0.77
0.77
0.77
0.77
0.77
0.77


56.0
0.0
0.0
0.0
0.0
0.0
0.0
0.0
0.0
0.0
0.0
0.0
9.0
0.0
0.0
0.0
0.0
0.0
0.0
38.0
0.0
0.0
0.0
9.0
0.0
0.0
0.0
0.0
0.0
0.0


56.0
0.0
0.0
0.0
0.0
0.0
0.0
0.0
0.0
0.0
0.0
0.0
9.0
0.0
0.0
0.0
0.0
0.0
0.0
38.0
0.0
0.0
0.0
9.0
0.0
0.0
0.0
0.0
0.0
0.0

1.0
1.0
1.0
1.0
1.0
1.0
1.0
1.0
1.0
1.0
1.0
1.0
1.0
1.0
1.0
1.0
1.0
1.0
1.0
1.0
1.0
1.0
1.0
1.0
1.0
1.0
1.0
1.0
1.0
1.0


63991.0
1039.0
35.0
86.0
107.0
46.0
0.0
433.0
562.0
675.0
3858.0
277.0
0.0
0.0
3361.0
0.0
0.0
3033.0
849.0
0.0
0.0
28198.0
20590.0
0.0
797.0
45.0
0.0
0.0
0.0
0.0

0.98
0.98
0.98
0.98
0.98
0.98
0.98
0.98
0.98
0.98
0.98
0.98
0.98
0.98
0.98
0.98
0.98
0.98
0.98
0.98
0.98
0.98
0.98
0.98
0.98
0.98
0.98
0.98
0.98
0.98


2695.0
0.0
0.0
0.0
0.0
0.0
0.0
0.0
0.0
0.0
2695.0
0.0
0.0
0.0
0.0
0.0
0.0
0.0
0.0
0.0
0.0
0.0
0.0
0.0
0.0
0.0
0.0
0.0
0.0
0.0

0.97
0.97
0.97
0.97
0.97
0.97
0.97
0.97
0.97
0.97
0.97
0.97
0.97
0.97
0.97
0.97
0.97
0.97
0.97
0.97
0.97
0.97
0.97
0.97
0.97
0.97
0.97
0.97
0.97
0.97


49439.0
691.0
22.0
0.0
0.0
0.0
0.0
344.0
185.0
126.0
367.0
0.0
0.0
0.0
0.0
0.0
0.0
0.0
500.0
0.0
0.0
26859.0
20345.0
0.0
0.0
0.0
0.0
0.0
0.0
0.0

1.0
1.0
1.0
1.0
1.0
1.0
1.0
1.0
1.0
1.0
1.0
1.0
1.0
1.0
1.0
1.0
1.0
1.0
1.0
1.0
1.0
1.0
1.0
1.0
1.0
1.0
1.0
1.0
1.0
1.0


193.0
0.0
0.0
0.0
0.0
0.0
0.0
0.0
0.0
0.0
193.0
0.0
0.0
0.0
0.0
0.0
0.0
0.0
0.0
0.0
0.0
0.0
0.0
0.0
0.0
0.0
0.0
0.0
0.0
0.0

1.0
1.0
1.0
1.0
1.0
1.0
1.0
1.0
1.0
1.0
1.0
1.0
1.0
1.0
1.0
1.0
1.0
1.0
1.0
1.0
1.0
1.0
1.0
1.0
1.0
1.0
1.0
1.0
1.0
1.0


6300.0
13.0
0.0
0.0
0.0
0.0
0.0
0.0
0.0
0.0
0.0
0.0
0.0
0.0
0.0
0.0
0.0
0.0
0.0
0.0
0.0
0.0
6287.0
0.0
0.0
0.0
0.0
0.0
0.0
0.0

1.0
1.0
1.0
1.0
1.0
1.0
1.0
1.0
1.0
1.0
1.0
1.0
1.0
1.0
1.0
1.0
1.0
1.0
1.0
1.0
1.0
1.0
1.0
1.0
1.0
1.0
1.0
1.0
1.0
1.0


1674.0
1.0
0.0
0.0
0.0
0.0
0.0
43.0
126.0
273.0
155.0
22.0
0.0
0.0
0.0
0.0
0.0
1054.0
0.0
0.0
0.0
0.0
0.0
0.0
0.0
0.0
0.0
0.0
0.0
0.0

0.98
0.98
0.98
0.98
0.98
0.98
0.98
0.98
0.98
0.98
0.98
0.98
0.98
0.98
0.98
0.98
0.98
0.98
0.98
0.98
0.98
0.98
0.98
0.98
0.98
0.98
0.98
0.98
0.98
0.98


357.0
48.0
2.0
0.0
0.0
0.0
0.0
0.0
99.0
12.0
143.0
40.0
0.0
0.0
0.0
0.0
0.0
0.0
0.0
0.0
0.0
0.0
0.0
0.0
13.0
0.0
0.0
0.0
0.0
0.0


357.0
48.0
2.0
0.0
0.0
0.0
0.0
0.0
99.0
12.0
143.0
40.0
0.0
0.0
0.0
0.0
0.0
0.0
0.0
0.0
0.0
0.0
0.0
0.0
13.0
0.0
0.0
0.0
0.0
0.0

1.0
1.0
1.0
1.0
1.0
1.0
1.0
1.0
1.0
1.0
1.0
1.0
1.0
1.0
1.0
1.0
1.0
1.0
1.0
1.0
1.0
1.0
1.0
1.0
1.0
1.0
1.0
1.0
1.0
1.0


3767.0
159.0
0.0
0.0
0.0
0.0
0.0
0.0
0.0
0.0
57.0
0.0
0.0
0.0
2284.0
0.0
0.0
1267.0
0.0
0.0
0.0
0.0
0.0
0.0
0.0
0.0
0.0
0.0
0.0
0.0


3767.0
159.0
0.0
0.0
0.0
0.0
0.0
0.0
0.0
0.0
57.0
0.0
0.0
0.0
2284.0
0.0
0.0
1267.0
0.0
0.0
0.0
0.0
0.0
0.0
0.0
0.0
0.0
0.0
0.0
0.0

0.98
0.98
0.98
0.98
0.98
0.98
0.98
0.98
0.98
0.98
0.98
0.98
0.98
0.98
0.98
0.98
0.98
0.98
0.98
0.98
0.98
0.98
0.98
0.98
0.98
0.98
0.98
0.98
0.98
0.98


998.0
7.0
9.0
45.0
107.0
46.0
0.0
0.0
0.0
0.0
0.0
0.0
0.0
0.0
0.0
0.0
0.0
0.0
0.0
0.0
0.0
0.0
0.0
0.0
784.0
0.0
0.0
0.0
0.0
0.0


998.0
7.0
9.0
45.0
107.0
46.0
0.0
0.0
0.0
0.0
0.0
0.0
0.0
0.0
0.0
0.0
0.0
0.0
0.0
0.0
0.0
0.0
0.0
0.0
784.0
0.0
0.0
0.0
0.0
0.0

0.95
0.95
0.95
0.95
0.95
0.95
0.95
0.95
0.95
0.95
0.95
0.95
0.95
0.95
0.95
0.95
0.95
0.95
0.95
0.95
0.95
0.95
0.95
0.95
0.95
0.95
0.95
0.95
0.95
0.95


115.0
0.0
0.0
0.0
0.0
0.0
0.0
0.0
0.0
57.0
58.0
0.0
0.0
0.0
0.0
0.0
0.0
0.0
0.0
0.0
0.0
0.0
0.0
0.0
0.0
0.0
0.0
0.0
0.0
0.0

1.0
1.0
1.0
1.0
1.0
1.0
1.0
1.0
1.0
1.0
1.0
1.0
1.0
1.0
1.0
1.0
1.0
1.0
1.0
1.0
1.0
1.0
1.0
1.0
1.0
1.0
1.0
1.0
1.0
1.0


85.0
85.0
0.0
0.0
0.0
0.0
0.0
0.0
0.0
0.0
0.0
0.0
0.0
0.0
0.0
0.0
0.0
0.0
0.0
0.0
0.0
0.0
0.0
0.0
0.0
0.0
0.0
0.0
0.0
0.0

1.0
1.0
1.0
1.0
1.0
1.0
1.0
1.0
1.0
1.0
1.0
1.0
1.0
1.0
1.0
1.0
1.0
1.0
1.0
1.0
1.0
1.0
1.0
1.0
1.0
1.0
1.0
1.0
1.0
1.0


85.0
0.0
0.0
0.0
0.0
0.0
0.0
0.0
0.0
32.0
0.0
53.0
0.0
0.0
0.0
0.0
0.0
0.0
0.0
0.0
0.0
0.0
0.0
0.0
0.0
0.0
0.0
0.0
0.0
0.0


53.0
0.0
0.0
0.0
0.0
0.0
0.0
0.0
0.0
0.0
0.0
53.0
0.0
0.0
0.0
0.0
0.0
0.0
0.0
0.0
0.0
0.0
0.0
0.0
0.0
0.0
0.0
0.0
0.0
0.0

0.82
0.82
0.82
0.82
0.82
0.82
0.82
0.82
0.82
0.82
0.82
0.82
0.82
0.82
0.82
0.82
0.82
0.82
0.82
0.82
0.82
0.82
0.82
0.82
0.82
0.82
0.82
0.82
0.82
0.82


32.0
0.0
0.0
0.0
0.0
0.0
0.0
0.0
0.0
32.0
0.0
0.0
0.0
0.0
0.0
0.0
0.0
0.0
0.0
0.0
0.0
0.0
0.0
0.0
0.0
0.0
0.0
0.0
0.0
0.0

0.93
0.93
0.93
0.93
0.93
0.93
0.93
0.93
0.93
0.93
0.93
0.93
0.93
0.93
0.93
0.93
0.93
0.93
0.93
0.93
0.93
0.93
0.93
0.93
0.93
0.93
0.93
0.93
0.93
0.93


120.0
0.0
0.0
41.0
0.0
0.0
0.0
0.0
0.0
0.0
0.0
0.0
0.0
0.0
60.0
0.0
0.0
0.0
19.0
0.0
0.0
0.0
0.0
0.0
0.0
0.0
0.0
0.0
0.0
0.0

1.0
1.0
1.0
1.0
1.0
1.0
1.0
1.0
1.0
1.0
1.0
1.0
1.0
1.0
1.0
1.0
1.0
1.0
1.0
1.0
1.0
1.0
1.0
1.0
1.0
1.0
1.0
1.0
1.0
1.0


36.0
0.0
0.0
0.0
0.0
0.0
0.0
0.0
36.0
0.0
0.0
0.0
0.0
0.0
0.0
0.0
0.0
0.0
0.0
0.0
0.0
0.0
0.0
0.0
0.0
0.0
0.0
0.0
0.0
0.0


36.0
0.0
0.0
0.0
0.0
0.0
0.0
0.0
36.0
0.0
0.0
0.0
0.0
0.0
0.0
0.0
0.0
0.0
0.0
0.0
0.0
0.0
0.0
0.0
0.0
0.0
0.0
0.0
0.0
0.0

1.0
1.0
1.0
1.0
1.0
1.0
1.0
1.0
1.0
1.0
1.0
1.0
1.0
1.0
1.0
1.0
1.0
1.0
1.0
1.0
1.0
1.0
1.0
1.0
1.0
1.0
1.0
1.0
1.0
1.0


847.0
0.0
0.0
0.0
0.0
0.0
0.0
0.0
0.0
15.0
0.0
17.0
0.0
0.0
0.0
0.0
0.0
515.0
300.0
0.0
0.0
0.0
0.0
0.0
0.0
0.0
0.0
0.0
0.0
0.0


832.0
0.0
0.0
0.0
0.0
0.0
0.0
0.0
0.0
0.0
0.0
17.0
0.0
0.0
0.0
0.0
0.0
515.0
300.0
0.0
0.0
0.0
0.0
0.0
0.0
0.0
0.0
0.0
0.0
0.0

1.0
1.0
1.0
1.0
1.0
1.0
1.0
1.0
1.0
1.0
1.0
1.0
1.0
1.0
1.0
1.0
1.0
1.0
1.0
1.0
1.0
1.0
1.0
1.0
1.0
1.0
1.0
1.0
1.0
1.0


15.0
0.0
0.0
0.0
0.0
0.0
0.0
0.0
0.0
15.0
0.0
0.0
0.0
0.0
0.0
0.0
0.0
0.0
0.0
0.0
0.0
0.0
0.0
0.0
0.0
0.0
0.0
0.0
0.0
0.0

1.0
1.0
1.0
1.0
1.0
1.0
1.0
1.0
1.0
1.0
1.0
1.0
1.0
1.0
1.0
1.0
1.0
1.0
1.0
1.0
1.0
1.0
1.0
1.0
1.0
1.0
1.0
1.0
1.0
1.0


15.0
15.0
0.0
0.0
0.0
0.0
0.0
0.0
0.0
0.0
0.0
0.0
0.0
0.0
0.0
0.0
0.0
0.0
0.0
0.0
0.0
0.0
0.0
0.0
0.0
0.0
0.0
0.0
0.0
0.0

0.99
0.99
0.99
0.99
0.99
0.99
0.99
0.99
0.99
0.99
0.99
0.99
0.99
0.99
0.99
0.99
0.99
0.99
0.99
0.99
0.99
0.99
0.99
0.99
0.99
0.99
0.99
0.99
0.99
0.99


1017.0
0.0
0.0
0.0
0.0
0.0
0.0
0.0
0.0
0.0
0.0
0.0
0.0
0.0
1017.0
0.0
0.0
0.0
0.0
0.0
0.0
0.0
0.0
0.0
0.0
0.0
0.0
0.0
0.0
0.0


1017.0
0.0
0.0
0.0
0.0
0.0
0.0
0.0
0.0
0.0
0.0
0.0
0.0
0.0
1017.0
0.0
0.0
0.0
0.0
0.0
0.0
0.0
0.0
0.0
0.0
0.0
0.0
0.0
0.0
0.0

1.0
1.0
1.0
1.0
1.0
1.0
1.0
1.0
1.0
1.0
1.0
1.0
1.0
1.0
1.0
1.0
1.0
1.0
1.0
1.0
1.0
1.0
1.0
1.0
1.0
1.0
1.0
1.0
1.0
1.0


959.0
0.0
0.0
0.0
0.0
0.0
0.0
0.0
0.0
0.0
0.0
0.0
0.0
0.0
0.0
0.0
0.0
0.0
0.0
0.0
0.0
959.0
0.0
0.0
0.0
0.0
0.0
0.0
0.0
0.0

0.89
0.89
0.89
0.89
0.89
0.89
0.89
0.89
0.89
0.89
0.89
0.89
0.89
0.89
0.89
0.89
0.89
0.89
0.89
0.89
0.89
0.89
0.89
0.89
0.89
0.89
0.89
0.89
0.89
0.89


45.0
0.0
0.0
0.0
0.0
0.0
0.0
0.0
0.0
0.0
0.0
0.0
0.0
0.0
0.0
0.0
0.0
0.0
0.0
0.0
0.0
0.0
0.0
0.0
0.0
45.0
0.0
0.0
0.0
0.0


45.0
0.0
0.0
0.0
0.0
0.0
0.0
0.0
0.0
0.0
0.0
0.0
0.0
0.0
0.0
0.0
0.0
0.0
0.0
0.0
0.0
0.0
0.0
0.0
0.0
45.0
0.0
0.0
0.0
0.0

0.83
0.83
0.83
0.83
0.83
0.83
0.83
0.83
0.83
0.83
0.83
0.83
0.83
0.83
0.83
0.83
0.83
0.83
0.83
0.83
0.83
0.83
0.83
0.83
0.83
0.83
0.83
0.83
0.83
0.83


5824.0
1854.0
1351.0
197.0
0.0
0.0
0.0
0.0
0.0
0.0
780.0
1642.0
0.0
0.0
0.0
0.0
0.0
0.0
0.0
0.0
0.0
0.0
0.0
0.0
0.0
0.0
0.0
0.0
0.0
0.0


5824.0
1854.0
1351.0
197.0
0.0
0.0
0.0
0.0
0.0
0.0
780.0
1642.0
0.0
0.0
0.0
0.0
0.0
0.0
0.0
0.0
0.0
0.0
0.0
0.0
0.0
0.0
0.0
0.0
0.0
0.0


4376.0
1296.0
1087.0
196.0
0.0
0.0
0.0
0.0
0.0
0.0
648.0
1149.0
0.0
0.0
0.0
0.0
0.0
0.0
0.0
0.0
0.0
0.0
0.0
0.0
0.0
0.0
0.0
0.0
0.0
0.0

1.0
1.0
1.0
1.0
1.0
1.0
1.0
1.0
1.0
1.0
1.0
1.0
1.0
1.0
1.0
1.0
1.0
1.0
1.0
1.0
1.0
1.0
1.0
1.0
1.0
1.0
1.0
1.0
1.0
1.0


1448.0
558.0
264.0
1.0
0.0
0.0
0.0
0.0
0.0
0.0
132.0
493.0
0.0
0.0
0.0
0.0
0.0
0.0
0.0
0.0
0.0
0.0
0.0
0.0
0.0
0.0
0.0
0.0
0.0
0.0

1.0
1.0
1.0
1.0
1.0
1.0
1.0
1.0
1.0
1.0
1.0
1.0
1.0
1.0
1.0
1.0
1.0
1.0
1.0
1.0
1.0
1.0
1.0
1.0
1.0
1.0
1.0
1.0
1.0
1.0


48713.0
0.0
407.0
29.0
0.0
1223.0
3205.0
0.0
0.0
0.0
0.0
0.0
0.0
0.0
43849.0
0.0
0.0
0.0
0.0
0.0
0.0
0.0
0.0
0.0
0.0
0.0
0.0
0.0
0.0
0.0


4428.0
0.0
0.0
0.0
0.0
1223.0
3205.0
0.0
0.0
0.0
0.0
0.0
0.0
0.0
0.0
0.0
0.0
0.0
0.0
0.0
0.0
0.0
0.0
0.0
0.0
0.0
0.0
0.0
0.0
0.0

1.0
1.0
1.0
1.0
1.0
1.0
1.0
1.0
1.0
1.0
1.0
1.0
1.0
1.0
1.0
1.0
1.0
1.0
1.0
1.0
1.0
1.0
1.0
1.0
1.0
1.0
1.0
1.0
1.0
1.0


284.0
0.0
255.0
29.0
0.0
0.0
0.0
0.0
0.0
0.0
0.0
0.0
0.0
0.0
0.0
0.0
0.0
0.0
0.0
0.0
0.0
0.0
0.0
0.0
0.0
0.0
0.0
0.0
0.0
0.0


284.0
0.0
255.0
29.0
0.0
0.0
0.0
0.0
0.0
0.0
0.0
0.0
0.0
0.0
0.0
0.0
0.0
0.0
0.0
0.0
0.0
0.0
0.0
0.0
0.0
0.0
0.0
0.0
0.0
0.0

1.0
1.0
1.0
1.0
1.0
1.0
1.0
1.0
1.0
1.0
1.0
1.0
1.0
1.0
1.0
1.0
1.0
1.0
1.0
1.0
1.0
1.0
1.0
1.0
1.0
1.0
1.0
1.0
1.0
1.0


44001.0
0.0
152.0
0.0
0.0
0.0
0.0
0.0
0.0
0.0
0.0
0.0
0.0
0.0
43849.0
0.0
0.0
0.0
0.0
0.0
0.0
0.0
0.0
0.0
0.0
0.0
0.0
0.0
0.0
0.0


44001.0
0.0
152.0
0.0
0.0
0.0
0.0
0.0
0.0
0.0
0.0
0.0
0.0
0.0
43849.0
0.0
0.0
0.0
0.0
0.0
0.0
0.0
0.0
0.0
0.0
0.0
0.0
0.0
0.0
0.0

1.0
1.0
1.0
1.0
1.0
1.0
1.0
1.0
1.0
1.0
1.0
1.0
1.0
1.0
1.0
1.0
1.0
1.0
1.0
1.0
1.0
1.0
1.0
1.0
1.0
1.0
1.0
1.0
1.0
1.0


3534.0
708.0
77.0
0.0
0.0
0.0
1179.0
0.0
0.0
187.0
1.0
1368.0
0.0
0.0
0.0
0.0
0.0
14.0
0.0
0.0
0.0
0.0
0.0
0.0
0.0
0.0
0.0
0.0
0.0
0.0


2154.0
708.0
77.0
0.0
0.0
0.0
0.0
0.0
0.0
0.0
1.0
1368.0
0.0
0.0
0.0
0.0
0.0
0.0
0.0
0.0
0.0
0.0
0.0
0.0
0.0
0.0
0.0
0.0
0.0
0.0


2154.0
708.0
77.0
0.0
0.0
0.0
0.0
0.0
0.0
0.0
1.0
1368.0
0.0
0.0
0.0
0.0
0.0
0.0
0.0
0.0
0.0
0.0
0.0
0.0
0.0
0.0
0.0
0.0
0.0
0.0

0.93
0.93
0.93
0.93
0.93
0.93
0.93
0.93
0.93
0.93
0.93
0.93
0.93
0.93
0.93
0.93
0.93
0.93
0.93
0.93
0.93
0.93
0.93
0.93
0.93
0.93
0.93
0.93
0.93
0.93


1179.0
0.0
0.0
0.0
0.0
0.0
1179.0
0.0
0.0
0.0
0.0
0.0
0.0
0.0
0.0
0.0
0.0
0.0
0.0
0.0
0.0
0.0
0.0
0.0
0.0
0.0
0.0
0.0
0.0
0.0

1.0
1.0
1.0
1.0
1.0
1.0
1.0
1.0
1.0
1.0
1.0
1.0
1.0
1.0
1.0
1.0
1.0
1.0
1.0
1.0
1.0
1.0
1.0
1.0
1.0
1.0
1.0
1.0
1.0
1.0


201.0
0.0
0.0
0.0
0.0
0.0
0.0
0.0
0.0
187.0
0.0
0.0
0.0
0.0
0.0
0.0
0.0
14.0
0.0
0.0
0.0
0.0
0.0
0.0
0.0
0.0
0.0
0.0
0.0
0.0


187.0
0.0
0.0
0.0
0.0
0.0
0.0
0.0
0.0
187.0
0.0
0.0
0.0
0.0
0.0
0.0
0.0
0.0
0.0
0.0
0.0
0.0
0.0
0.0
0.0
0.0
0.0
0.0
0.0
0.0

1.0
1.0
1.0
1.0
1.0
1.0
1.0
1.0
1.0
1.0
1.0
1.0
1.0
1.0
1.0
1.0
1.0
1.0
1.0
1.0
1.0
1.0
1.0
1.0
1.0
1.0
1.0
1.0
1.0
1.0


14.0
0.0
0.0
0.0
0.0
0.0
0.0
0.0
0.0
0.0
0.0
0.0
0.0
0.0
0.0
0.0
0.0
14.0
0.0
0.0
0.0
0.0
0.0
0.0
0.0
0.0
0.0
0.0
0.0
0.0

1.0
1.0
1.0
1.0
1.0
1.0
1.0
1.0
1.0
1.0
1.0
1.0
1.0
1.0
1.0
1.0
1.0
1.0
1.0
1.0
1.0
1.0
1.0
1.0
1.0
1.0
1.0
1.0
1.0
1.0


2683.0
589.0
404.0
1690.0
0.0
0.0
0.0
0.0
0.0
0.0
0.0
0.0
0.0
0.0
0.0
0.0
0.0
0.0
0.0
0.0
0.0
0.0
0.0
0.0
0.0
0.0
0.0
0.0
0.0
0.0


2683.0
589.0
404.0
1690.0
0.0
0.0
0.0
0.0
0.0
0.0
0.0
0.0
0.0
0.0
0.0
0.0
0.0
0.0
0.0
0.0
0.0
0.0
0.0
0.0
0.0
0.0
0.0
0.0
0.0
0.0

1.0
1.0
1.0
1.0
1.0
1.0
1.0
1.0
1.0
1.0
1.0
1.0
1.0
1.0
1.0
1.0
1.0
1.0
1.0
1.0
1.0
1.0
1.0
1.0
1.0
1.0
1.0
1.0
1.0
1.0


1499.0
489.0
262.0
748.0
0.0
0.0
0.0
0.0
0.0
0.0
0.0
0.0
0.0
0.0
0.0
0.0
0.0
0.0
0.0
0.0
0.0
0.0
0.0
0.0
0.0
0.0
0.0
0.0
0.0
0.0

1.0
1.0
1.0
1.0
1.0
1.0
1.0
1.0
1.0
1.0
1.0
1.0
1.0
1.0
1.0
1.0
1.0
1.0
1.0
1.0
1.0
1.0
1.0
1.0
1.0
1.0
1.0
1.0
1.0
1.0


307.0
0.0
0.0
307.0
0.0
0.0
0.0
0.0
0.0
0.0
0.0
0.0
0.0
0.0
0.0
0.0
0.0
0.0
0.0
0.0
0.0
0.0
0.0
0.0
0.0
0.0
0.0
0.0
0.0
0.0

1.0
1.0
1.0
1.0
1.0
1.0
1.0
1.0
1.0
1.0
1.0
1.0
1.0
1.0
1.0
1.0
1.0
1.0
1.0
1.0
1.0
1.0
1.0
1.0
1.0
1.0
1.0
1.0
1.0
1.0


4815.0
202.0
337.0
8.0
94.0
42.0
605.0
69.0
183.0
462.0
249.0
1140.0
0.0
0.0
0.0
0.0
0.0
0.0
0.0
83.0
0.0
0.0
0.0
0.0
1341.0
0.0
0.0
0.0
0.0
0.0

0.76
0.76
0.76
0.76
0.76
0.76
0.76
0.76
0.76
0.76
0.76
0.76
0.76
0.76
0.76
0.76
0.76
0.76
0.76
0.76
0.76
0.76
0.76
0.76
0.76
0.76
0.76
0.76
0.76
0.76


1376.0
38.0
2.0
0.0
0.0
0.0
0.0
14.0
163.0
347.0
223.0
589.0
0.0
0.0
0.0
0.0
0.0
0.0
0.0
0.0
0.0
0.0
0.0
0.0
0.0
0.0
0.0
0.0
0.0
0.0


1376.0
38.0
2.0
0.0
0.0
0.0
0.0
14.0
163.0
347.0
223.0
589.0
0.0
0.0
0.0
0.0
0.0
0.0
0.0
0.0
0.0
0.0
0.0
0.0
0.0
0.0
0.0
0.0
0.0
0.0

1.0
1.0
1.0
1.0
1.0
1.0
1.0
1.0
1.0
1.0
1.0
1.0
1.0
1.0
1.0
1.0
1.0
1.0
1.0
1.0
1.0
1.0
1.0
1.0
1.0
1.0
1.0
1.0
1.0
1.0


1054.0
74.0
329.0
0.0
0.0
0.0
74.0
0.0
0.0
0.0
0.0
551.0
0.0
0.0
0.0
0.0
0.0
0.0
0.0
0.0
0.0
0.0
0.0
0.0
26.0
0.0
0.0
0.0
0.0
0.0


1054.0
74.0
329.0
0.0
0.0
0.0
74.0
0.0
0.0
0.0
0.0
551.0
0.0
0.0
0.0
0.0
0.0
0.0
0.0
0.0
0.0
0.0
0.0
0.0
26.0
0.0
0.0
0.0
0.0
0.0

1.0
1.0
1.0
1.0
1.0
1.0
1.0
1.0
1.0
1.0
1.0
1.0
1.0
1.0
1.0
1.0
1.0
1.0
1.0
1.0
1.0
1.0
1.0
1.0
1.0
1.0
1.0
1.0
1.0
1.0


1772.0
4.0
6.0
0.0
94.0
42.0
177.0
55.0
2.0
77.0
0.0
0.0
0.0
0.0
0.0
0.0
0.0
0.0
0.0
0.0
0.0
0.0
0.0
0.0
1315.0
0.0
0.0
0.0
0.0
0.0


1772.0
4.0
6.0
0.0
94.0
42.0
177.0
55.0
2.0
77.0
0.0
0.0
0.0
0.0
0.0
0.0
0.0
0.0
0.0
0.0
0.0
0.0
0.0
0.0
1315.0
0.0
0.0
0.0
0.0
0.0

1.0
1.0
1.0
1.0
1.0
1.0
1.0
1.0
1.0
1.0
1.0
1.0
1.0
1.0
1.0
1.0
1.0
1.0
1.0
1.0
1.0
1.0
1.0
1.0
1.0
1.0
1.0
1.0
1.0
1.0


338.0
0.0
0.0
0.0
0.0
0.0
338.0
0.0
0.0
0.0
0.0
0.0
0.0
0.0
0.0
0.0
0.0
0.0
0.0
0.0
0.0
0.0
0.0
0.0
0.0
0.0
0.0
0.0
0.0
0.0


338.0
0.0
0.0
0.0
0.0
0.0
338.0
0.0
0.0
0.0
0.0
0.0
0.0
0.0
0.0
0.0
0.0
0.0
0.0
0.0
0.0
0.0
0.0
0.0
0.0
0.0
0.0
0.0
0.0
0.0

1.0
1.0
1.0
1.0
1.0
1.0
1.0
1.0
1.0
1.0
1.0
1.0
1.0
1.0
1.0
1.0
1.0
1.0
1.0
1.0
1.0
1.0
1.0
1.0
1.0
1.0
1.0
1.0
1.0
1.0


34.0
34.0
0.0
0.0
0.0
0.0
0.0
0.0
0.0
0.0
0.0
0.0
0.0
0.0
0.0
0.0
0.0
0.0
0.0
0.0
0.0
0.0
0.0
0.0
0.0
0.0
0.0
0.0
0.0
0.0

1.0
1.0
1.0
1.0
1.0
1.0
1.0
1.0
1.0
1.0
1.0
1.0
1.0
1.0
1.0
1.0
1.0
1.0
1.0
1.0
1.0
1.0
1.0
1.0
1.0
1.0
1.0
1.0
1.0
1.0


48.0
48.0
0.0
0.0
0.0
0.0
0.0
0.0
0.0
0.0
0.0
0.0
0.0
0.0
0.0
0.0
0.0
0.0
0.0
0.0
0.0
0.0
0.0
0.0
0.0
0.0
0.0
0.0
0.0
0.0


48.0
48.0
0.0
0.0
0.0
0.0
0.0
0.0
0.0
0.0
0.0
0.0
0.0
0.0
0.0
0.0
0.0
0.0
0.0
0.0
0.0
0.0
0.0
0.0
0.0
0.0
0.0
0.0
0.0
0.0

1.0
1.0
1.0
1.0
1.0
1.0
1.0
1.0
1.0
1.0
1.0
1.0
1.0
1.0
1.0
1.0
1.0
1.0
1.0
1.0
1.0
1.0
1.0
1.0
1.0
1.0
1.0
1.0
1.0
1.0


53.0
0.0
0.0
0.0
0.0
0.0
0.0
0.0
0.0
27.0
26.0
0.0
0.0
0.0
0.0
0.0
0.0
0.0
0.0
0.0
0.0
0.0
0.0
0.0
0.0
0.0
0.0
0.0
0.0
0.0


53.0
0.0
0.0
0.0
0.0
0.0
0.0
0.0
0.0
27.0
26.0
0.0
0.0
0.0
0.0
0.0
0.0
0.0
0.0
0.0
0.0
0.0
0.0
0.0
0.0
0.0
0.0
0.0
0.0
0.0

0.9
0.9
0.9
0.9
0.9
0.9
0.9
0.9
0.9
0.9
0.9
0.9
0.9
0.9
0.9
0.9
0.9
0.9
0.9
0.9
0.9
0.9
0.9
0.9
0.9
0.9
0.9
0.9
0.9
0.9


15.0
4.0
0.0
0.0
0.0
0.0
0.0
0.0
0.0
11.0
0.0
0.0
0.0
0.0
0.0
0.0
0.0
0.0
0.0
0.0
0.0
0.0
0.0
0.0
0.0
0.0
0.0
0.0
0.0
0.0


15.0
4.0
0.0
0.0
0.0
0.0
0.0
0.0
0.0
11.0
0.0
0.0
0.0
0.0
0.0
0.0
0.0
0.0
0.0
0.0
0.0
0.0
0.0
0.0
0.0
0.0
0.0
0.0
0.0
0.0

0.98
0.98
0.98
0.98
0.98
0.98
0.98
0.98
0.98
0.98
0.98
0.98
0.98
0.98
0.98
0.98
0.98
0.98
0.98
0.98
0.98
0.98
0.98
0.98
0.98
0.98
0.98
0.98
0.98
0.98


18.0
0.0
0.0
0.0
0.0
0.0
0.0
0.0
18.0
0.0
0.0
0.0
0.0
0.0
0.0
0.0
0.0
0.0
0.0
0.0
0.0
0.0
0.0
0.0
0.0
0.0
0.0
0.0
0.0
0.0

1.0
1.0
1.0
1.0
1.0
1.0
1.0
1.0
1.0
1.0
1.0
1.0
1.0
1.0
1.0
1.0
1.0
1.0
1.0
1.0
1.0
1.0
1.0
1.0
1.0
1.0
1.0
1.0
1.0
1.0


8.0
0.0
0.0
8.0
0.0
0.0
0.0
0.0
0.0
0.0
0.0
0.0
0.0
0.0
0.0
0.0
0.0
0.0
0.0
0.0
0.0
0.0
0.0
0.0
0.0
0.0
0.0
0.0
0.0
0.0


8.0
0.0
0.0
8.0
0.0
0.0
0.0
0.0
0.0
0.0
0.0
0.0
0.0
0.0
0.0
0.0
0.0
0.0
0.0
0.0
0.0
0.0
0.0
0.0
0.0
0.0
0.0
0.0
0.0
0.0

0.85
0.85
0.85
0.85
0.85
0.85
0.85
0.85
0.85
0.85
0.85
0.85
0.85
0.85
0.85
0.85
0.85
0.85
0.85
0.85
0.85
0.85
0.85
0.85
0.85
0.85
0.85
0.85
0.85
0.85


83.0
0.0
0.0
0.0
0.0
0.0
0.0
0.0
0.0
0.0
0.0
0.0
0.0
0.0
0.0
0.0
0.0
0.0
0.0
83.0
0.0
0.0
0.0
0.0
0.0
0.0
0.0
0.0
0.0
0.0


83.0
0.0
0.0
0.0
0.0
0.0
0.0
0.0
0.0
0.0
0.0
0.0
0.0
0.0
0.0
0.0
0.0
0.0
0.0
83.0
0.0
0.0
0.0
0.0
0.0
0.0
0.0
0.0
0.0
0.0

0.97
0.97
0.97
0.97
0.97
0.97
0.97
0.97
0.97
0.97
0.97
0.97
0.97
0.97
0.97
0.97
0.97
0.97
0.97
0.97
0.97
0.97
0.97
0.97
0.97
0.97
0.97
0.97
0.97
0.97


179933.0
364.0
608.0
953.0
0.0
0.0
0.0
0.0
0.0
0.0
909.0
708.0
0.0
26091.0
0.0
0.0
0.0
0.0
0.0
0.0
0.0
0.0
0.0
0.0
150300.0
0.0
0.0
0.0
0.0
0.0

0.81
0.81
0.81
0.81
0.81
0.81
0.81
0.81
0.81
0.81
0.81
0.81
0.81
0.81
0.81
0.81
0.81
0.81
0.81
0.81
0.81
0.81
0.81
0.81
0.81
0.81
0.81
0.81
0.81
0.81


1112.0
364.0
562.0
0.0
0.0
0.0
0.0
0.0
0.0
0.0
0.0
186.0
0.0
0.0
0.0
0.0
0.0
0.0
0.0
0.0
0.0
0.0
0.0
0.0
0.0
0.0
0.0
0.0
0.0
0.0

1.0
1.0
1.0
1.0
1.0
1.0
1.0
1.0
1.0
1.0
1.0
1.0
1.0
1.0
1.0
1.0
1.0
1.0
1.0
1.0
1.0
1.0
1.0
1.0
1.0
1.0
1.0
1.0
1.0
1.0


11.0
7.0
4.0
0.0
0.0
0.0
0.0
0.0
0.0
0.0
0.0
0.0
0.0
0.0
0.0
0.0
0.0
0.0
0.0
0.0
0.0
0.0
0.0
0.0
0.0
0.0
0.0
0.0
0.0
0.0

1.0
1.0
1.0
1.0
1.0
1.0
1.0
1.0
1.0
1.0
1.0
1.0
1.0
1.0
1.0
1.0
1.0
1.0
1.0
1.0
1.0
1.0
1.0
1.0
1.0
1.0
1.0
1.0
1.0
1.0


909.0
0.0
0.0
0.0
0.0
0.0
0.0
0.0
0.0
0.0
909.0
0.0
0.0
0.0
0.0
0.0
0.0
0.0
0.0
0.0
0.0
0.0
0.0
0.0
0.0
0.0
0.0
0.0
0.0
0.0


909.0
0.0
0.0
0.0
0.0
0.0
0.0
0.0
0.0
0.0
909.0
0.0
0.0
0.0
0.0
0.0
0.0
0.0
0.0
0.0
0.0
0.0
0.0
0.0
0.0
0.0
0.0
0.0
0.0
0.0

1.0
1.0
1.0
1.0
1.0
1.0
1.0
1.0
1.0
1.0
1.0
1.0
1.0
1.0
1.0
1.0
1.0
1.0
1.0
1.0
1.0
1.0
1.0
1.0
1.0
1.0
1.0
1.0
1.0
1.0


119233.0
0.0
0.0
641.0
0.0
0.0
0.0
0.0
0.0
0.0
0.0
0.0
0.0
0.0
0.0
0.0
0.0
0.0
0.0
0.0
0.0
0.0
0.0
0.0
118592.0
0.0
0.0
0.0
0.0
0.0


119233.0
0.0
0.0
641.0
0.0
0.0
0.0
0.0
0.0
0.0
0.0
0.0
0.0
0.0
0.0
0.0
0.0
0.0
0.0
0.0
0.0
0.0
0.0
0.0
118592.0
0.0
0.0
0.0
0.0
0.0

1.0
1.0
1.0
1.0
1.0
1.0
1.0
1.0
1.0
1.0
1.0
1.0
1.0
1.0
1.0
1.0
1.0
1.0
1.0
1.0
1.0
1.0
1.0
1.0
1.0
1.0
1.0
1.0
1.0
1.0


26659.0
0.0
46.0
0.0
0.0
0.0
0.0
0.0
0.0
0.0
0.0
522.0
0.0
26091.0
0.0
0.0
0.0
0.0
0.0
0.0
0.0
0.0
0.0
0.0
0.0
0.0
0.0
0.0
0.0
0.0


26659.0
0.0
46.0
0.0
0.0
0.0
0.0
0.0
0.0
0.0
0.0
522.0
0.0
26091.0
0.0
0.0
0.0
0.0
0.0
0.0
0.0
0.0
0.0
0.0
0.0
0.0
0.0
0.0
0.0
0.0

1.0
1.0
1.0
1.0
1.0
1.0
1.0
1.0
1.0
1.0
1.0
1.0
1.0
1.0
1.0
1.0
1.0
1.0
1.0
1.0
1.0
1.0
1.0
1.0
1.0
1.0
1.0
1.0
1.0
1.0


30834.0
0.0
0.0
289.0
0.0
0.0
0.0
0.0
0.0
0.0
0.0
0.0
0.0
0.0
0.0
0.0
0.0
0.0
0.0
0.0
0.0
0.0
0.0
0.0
30545.0
0.0
0.0
0.0
0.0
0.0


30834.0
0.0
0.0
289.0
0.0
0.0
0.0
0.0
0.0
0.0
0.0
0.0
0.0
0.0
0.0
0.0
0.0
0.0
0.0
0.0
0.0
0.0
0.0
0.0
30545.0
0.0
0.0
0.0
0.0
0.0

1.0
1.0
1.0
1.0
1.0
1.0
1.0
1.0
1.0
1.0
1.0
1.0
1.0
1.0
1.0
1.0
1.0
1.0
1.0
1.0
1.0
1.0
1.0
1.0
1.0
1.0
1.0
1.0
1.0
1.0


1574.0
858.0
16.0
0.0
0.0
0.0
0.0
0.0
0.0
0.0
0.0
0.0
0.0
0.0
0.0
0.0
700.0
0.0
0.0
0.0
0.0
0.0
0.0
0.0
0.0
0.0
0.0
0.0
0.0
0.0


858.0
858.0
0.0
0.0
0.0
0.0
0.0
0.0
0.0
0.0
0.0
0.0
0.0
0.0
0.0
0.0
0.0
0.0
0.0
0.0
0.0
0.0
0.0
0.0
0.0
0.0
0.0
0.0
0.0
0.0


858.0
858.0
0.0
0.0
0.0
0.0
0.0
0.0
0.0
0.0
0.0
0.0
0.0
0.0
0.0
0.0
0.0
0.0
0.0
0.0
0.0
0.0
0.0
0.0
0.0
0.0
0.0
0.0
0.0
0.0

1.0
1.0
1.0
1.0
1.0
1.0
1.0
1.0
1.0
1.0
1.0
1.0
1.0
1.0
1.0
1.0
1.0
1.0
1.0
1.0
1.0
1.0
1.0
1.0
1.0
1.0
1.0
1.0
1.0
1.0


16.0
0.0
16.0
0.0
0.0
0.0
0.0
0.0
0.0
0.0
0.0
0.0
0.0
0.0
0.0
0.0
0.0
0.0
0.0
0.0
0.0
0.0
0.0
0.0
0.0
0.0
0.0
0.0
0.0
0.0


16.0
0.0
16.0
0.0
0.0
0.0
0.0
0.0
0.0
0.0
0.0
0.0
0.0
0.0
0.0
0.0
0.0
0.0
0.0
0.0
0.0
0.0
0.0
0.0
0.0
0.0
0.0
0.0
0.0
0.0

1.0
1.0
1.0
1.0
1.0
1.0
1.0
1.0
1.0
1.0
1.0
1.0
1.0
1.0
1.0
1.0
1.0
1.0
1.0
1.0
1.0
1.0
1.0
1.0
1.0
1.0
1.0
1.0
1.0
1.0


700.0
0.0
0.0
0.0
0.0
0.0
0.0
0.0
0.0
0.0
0.0
0.0
0.0
0.0
0.0
0.0
700.0
0.0
0.0
0.0
0.0
0.0
0.0
0.0
0.0
0.0
0.0
0.0
0.0
0.0


700.0
0.0
0.0
0.0
0.0
0.0
0.0
0.0
0.0
0.0
0.0
0.0
0.0
0.0
0.0
0.0
700.0
0.0
0.0
0.0
0.0
0.0
0.0
0.0
0.0
0.0
0.0
0.0
0.0
0.0

1.0
1.0
1.0
1.0
1.0
1.0
1.0
1.0
1.0
1.0
1.0
1.0
1.0
1.0
1.0
1.0
1.0
1.0
1.0
1.0
1.0
1.0
1.0
1.0
1.0
1.0
1.0
1.0
1.0
1.0


536286.0
77.0
139.0
93.0
282.0
0.0
0.0
119.0
246.0
361.0
67.0
0.0
0.0
0.0
1.0
0.0
0.0
127910.0
0.0
69620.0
325476.0
0.0
0.0
0.0
11895.0
0.0
0.0
0.0
0.0
0.0

1.0
1.0
1.0
1.0
1.0
1.0
1.0
1.0
1.0
1.0
1.0
1.0
1.0
1.0
1.0
1.0
1.0
1.0
1.0
1.0
1.0
1.0
1.0
1.0
1.0
1.0
1.0
1.0
1.0
1.0


549.0
0.0
0.0
0.0
0.0
0.0
0.0
0.0
246.0
303.0
0.0
0.0
0.0
0.0
0.0
0.0
0.0
0.0
0.0
0.0
0.0
0.0
0.0
0.0
0.0
0.0
0.0
0.0
0.0
0.0

0.77
0.77
0.77
0.77
0.77
0.77
0.77
0.77
0.77
0.77
0.77
0.77
0.77
0.77
0.77
0.77
0.77
0.77
0.77
0.77
0.77
0.77
0.77
0.77
0.77
0.77
0.77
0.77
0.77
0.77


699.0
19.0
128.0
75.0
245.0
0.0
0.0
119.0
0.0
46.0
67.0
0.0
0.0
0.0
0.0
0.0
0.0
0.0
0.0
0.0
0.0
0.0
0.0
0.0
0.0
0.0
0.0
0.0
0.0
0.0


699.0
19.0
128.0
75.0
245.0
0.0
0.0
119.0
0.0
46.0
67.0
0.0
0.0
0.0
0.0
0.0
0.0
0.0
0.0
0.0
0.0
0.0
0.0
0.0
0.0
0.0
0.0
0.0
0.0
0.0

1.0
1.0
1.0
1.0
1.0
1.0
1.0
1.0
1.0
1.0
1.0
1.0
1.0
1.0
1.0
1.0
1.0
1.0
1.0
1.0
1.0
1.0
1.0
1.0
1.0
1.0
1.0
1.0
1.0
1.0


52.0
46.0
0.0
0.0
0.0
0.0
0.0
0.0
0.0
0.0
0.0
0.0
0.0
0.0
0.0
0.0
0.0
6.0
0.0
0.0
0.0
0.0
0.0
0.0
0.0
0.0
0.0
0.0
0.0
0.0

0.75
0.75
0.75
0.75
0.75
0.75
0.75
0.75
0.75
0.75
0.75
0.75
0.75
0.75
0.75
0.75
0.75
0.75
0.75
0.75
0.75
0.75
0.75
0.75
0.75
0.75
0.75
0.75
0.75
0.75


46.0
46.0
0.0
0.0
0.0
0.0
0.0
0.0
0.0
0.0
0.0
0.0
0.0
0.0
0.0
0.0
0.0
0.0
0.0
0.0
0.0
0.0
0.0
0.0
0.0
0.0
0.0
0.0
0.0
0.0

0.99
0.99
0.99
0.99
0.99
0.99
0.99
0.99
0.99
0.99
0.99
0.99
0.99
0.99
0.99
0.99
0.99
0.99
0.99
0.99
0.99
0.99
0.99
0.99
0.99
0.99
0.99
0.99
0.99
0.99


37.0
0.0
0.0
0.0
37.0
0.0
0.0
0.0
0.0
0.0
0.0
0.0
0.0
0.0
0.0
0.0
0.0
0.0
0.0
0.0
0.0
0.0
0.0
0.0
0.0
0.0
0.0
0.0
0.0
0.0

1.0
1.0
1.0
1.0
1.0
1.0
1.0
1.0
1.0
1.0
1.0
1.0
1.0
1.0
1.0
1.0
1.0
1.0
1.0
1.0
1.0
1.0
1.0
1.0
1.0
1.0
1.0
1.0
1.0
1.0


11913.0
0.0
0.0
18.0
0.0
0.0
0.0
0.0
0.0
0.0
0.0
0.0
0.0
0.0
0.0
0.0
0.0
0.0
0.0
0.0
0.0
0.0
0.0
0.0
11895.0
0.0
0.0
0.0
0.0
0.0

1.0
1.0
1.0
1.0
1.0
1.0
1.0
1.0
1.0
1.0
1.0
1.0
1.0
1.0
1.0
1.0
1.0
1.0
1.0
1.0
1.0
1.0
1.0
1.0
1.0
1.0
1.0
1.0
1.0
1.0


15.0
0.0
3.0
0.0
0.0
0.0
0.0
0.0
0.0
12.0
0.0
0.0
0.0
0.0
0.0
0.0
0.0
0.0
0.0
0.0
0.0
0.0
0.0
0.0
0.0
0.0
0.0
0.0
0.0
0.0


15.0
0.0
3.0
0.0
0.0
0.0
0.0
0.0
0.0
12.0
0.0
0.0
0.0
0.0
0.0
0.0
0.0
0.0
0.0
0.0
0.0
0.0
0.0
0.0
0.0
0.0
0.0
0.0
0.0
0.0

0.98
0.98
0.98
0.98
0.98
0.98
0.98
0.98
0.98
0.98
0.98
0.98
0.98
0.98
0.98
0.98
0.98
0.98
0.98
0.98
0.98
0.98
0.98
0.98
0.98
0.98
0.98
0.98
0.98
0.98


12.0
12.0
0.0
0.0
0.0
0.0
0.0
0.0
0.0
0.0
0.0
0.0
0.0
0.0
0.0
0.0
0.0
0.0
0.0
0.0
0.0
0.0
0.0
0.0
0.0
0.0
0.0
0.0
0.0
0.0

0.71
0.71
0.71
0.71
0.71
0.71
0.71
0.71
0.71
0.71
0.71
0.71
0.71
0.71
0.71
0.71
0.71
0.71
0.71
0.71
0.71
0.71
0.71
0.71
0.71
0.71
0.71
0.71
0.71
0.71


325477.0
0.0
0.0
0.0
0.0
0.0
0.0
0.0
0.0
0.0
0.0
0.0
0.0
0.0
1.0
0.0
0.0
0.0
0.0
0.0
325476.0
0.0
0.0
0.0
0.0
0.0
0.0
0.0
0.0
0.0


325477.0
0.0
0.0
0.0
0.0
0.0
0.0
0.0
0.0
0.0
0.0
0.0
0.0
0.0
1.0
0.0
0.0
0.0
0.0
0.0
325476.0
0.0
0.0
0.0
0.0
0.0
0.0
0.0
0.0
0.0

1.0
1.0
1.0
1.0
1.0
1.0
1.0
1.0
1.0
1.0
1.0
1.0
1.0
1.0
1.0
1.0
1.0
1.0
1.0
1.0
1.0
1.0
1.0
1.0
1.0
1.0
1.0
1.0
1.0
1.0


127082.0
0.0
0.0
0.0
0.0
0.0
0.0
0.0
0.0
0.0
0.0
0.0
0.0
0.0
0.0
0.0
0.0
127082.0
0.0
0.0
0.0
0.0
0.0
0.0
0.0
0.0
0.0
0.0
0.0
0.0


127082.0
0.0
0.0
0.0
0.0
0.0
0.0
0.0
0.0
0.0
0.0
0.0
0.0
0.0
0.0
0.0
0.0
127082.0
0.0
0.0
0.0
0.0
0.0
0.0
0.0
0.0
0.0
0.0
0.0
0.0

1.0
1.0
1.0
1.0
1.0
1.0
1.0
1.0
1.0
1.0
1.0
1.0
1.0
1.0
1.0
1.0
1.0
1.0
1.0
1.0
1.0
1.0
1.0
1.0
1.0
1.0
1.0
1.0
1.0
1.0


69620.0
0.0
0.0
0.0
0.0
0.0
0.0
0.0
0.0
0.0
0.0
0.0
0.0
0.0
0.0
0.0
0.0
0.0
0.0
69620.0
0.0
0.0
0.0
0.0
0.0
0.0
0.0
0.0
0.0
0.0


69620.0
0.0
0.0
0.0
0.0
0.0
0.0
0.0
0.0
0.0
0.0
0.0
0.0
0.0
0.0
0.0
0.0
0.0
0.0
69620.0
0.0
0.0
0.0
0.0
0.0
0.0
0.0
0.0
0.0
0.0

1.0
1.0
1.0
1.0
1.0
1.0
1.0
1.0
1.0
1.0
1.0
1.0
1.0
1.0
1.0
1.0
1.0
1.0
1.0
1.0
1.0
1.0
1.0
1.0
1.0
1.0
1.0
1.0
1.0
1.0


154053.0
137.0
95.0
1.0
0.0
188.0
637.0
0.0
0.0
20.0
45.0
42.0
0.0
0.0
0.0
0.0
374.0
0.0
0.0
0.0
0.0
0.0
0.0
0.0
152514.0
0.0
0.0
0.0
0.0
0.0

1.0
1.0
1.0
1.0
1.0
1.0
1.0
1.0
1.0
1.0
1.0
1.0
1.0
1.0
1.0
1.0
1.0
1.0
1.0
1.0
1.0
1.0
1.0
1.0
1.0
1.0
1.0
1.0
1.0
1.0


153107.0
9.0
0.0
1.0
0.0
150.0
384.0
0.0
0.0
7.0
0.0
42.0
0.0
0.0
0.0
0.0
0.0
0.0
0.0
0.0
0.0
0.0
0.0
0.0
152514.0
0.0
0.0
0.0
0.0
0.0


384.0
0.0
0.0
0.0
0.0
0.0
384.0
0.0
0.0
0.0
0.0
0.0
0.0
0.0
0.0
0.0
0.0
0.0
0.0
0.0
0.0
0.0
0.0
0.0
0.0
0.0
0.0
0.0
0.0
0.0

1.0
1.0
1.0
1.0
1.0
1.0
1.0
1.0
1.0
1.0
1.0
1.0
1.0
1.0
1.0
1.0
1.0
1.0
1.0
1.0
1.0
1.0
1.0
1.0
1.0
1.0
1.0
1.0
1.0
1.0


3374.0
0.0
0.0
0.0
0.0
150.0
0.0
0.0
0.0
0.0
0.0
0.0
0.0
0.0
0.0
0.0
0.0
0.0
0.0
0.0
0.0
0.0
0.0
0.0
3224.0
0.0
0.0
0.0
0.0
0.0

1.0
1.0
1.0
1.0
1.0
1.0
1.0
1.0
1.0
1.0
1.0
1.0
1.0
1.0
1.0
1.0
1.0
1.0
1.0
1.0
1.0
1.0
1.0
1.0
1.0
1.0
1.0
1.0
1.0
1.0


149349.0
9.0
0.0
1.0
0.0
0.0
0.0
0.0
0.0
7.0
0.0
42.0
0.0
0.0
0.0
0.0
0.0
0.0
0.0
0.0
0.0
0.0
0.0
0.0
149290.0
0.0
0.0
0.0
0.0
0.0

1.0
1.0
1.0
1.0
1.0
1.0
1.0
1.0
1.0
1.0
1.0
1.0
1.0
1.0
1.0
1.0
1.0
1.0
1.0
1.0
1.0
1.0
1.0
1.0
1.0
1.0
1.0
1.0
1.0
1.0


176.0
0.0
7.0
0.0
0.0
0.0
169.0
0.0
0.0
0.0
0.0
0.0
0.0
0.0
0.0
0.0
0.0
0.0
0.0
0.0
0.0
0.0
0.0
0.0
0.0
0.0
0.0
0.0
0.0
0.0


176.0
0.0
7.0
0.0
0.0
0.0
169.0
0.0
0.0
0.0
0.0
0.0
0.0
0.0
0.0
0.0
0.0
0.0
0.0
0.0
0.0
0.0
0.0
0.0
0.0
0.0
0.0
0.0
0.0
0.0

1.0
1.0
1.0
1.0
1.0
1.0
1.0
1.0
1.0
1.0
1.0
1.0
1.0
1.0
1.0
1.0
1.0
1.0
1.0
1.0
1.0
1.0
1.0
1.0
1.0
1.0
1.0
1.0
1.0
1.0


219.0
128.0
46.0
0.0
0.0
0.0
0.0
0.0
0.0
0.0
45.0
0.0
0.0
0.0
0.0
0.0
0.0
0.0
0.0
0.0
0.0
0.0
0.0
0.0
0.0
0.0
0.0
0.0
0.0
0.0


219.0
128.0
46.0
0.0
0.0
0.0
0.0
0.0
0.0
0.0
45.0
0.0
0.0
0.0
0.0
0.0
0.0
0.0
0.0
0.0
0.0
0.0
0.0
0.0
0.0
0.0
0.0
0.0
0.0
0.0

0.98
0.98
0.98
0.98
0.98
0.98
0.98
0.98
0.98
0.98
0.98
0.98
0.98
0.98
0.98
0.98
0.98
0.98
0.98
0.98
0.98
0.98
0.98
0.98
0.98
0.98
0.98
0.98
0.98
0.98


452.0
0.0
0.0
0.0
0.0
0.0
84.0
0.0
0.0
0.0
0.0
0.0
0.0
0.0
0.0
0.0
368.0
0.0
0.0
0.0
0.0
0.0
0.0
0.0
0.0
0.0
0.0
0.0
0.0
0.0

1.0
1.0
1.0
1.0
1.0
1.0
1.0
1.0
1.0
1.0
1.0
1.0
1.0
1.0
1.0
1.0
1.0
1.0
1.0
1.0
1.0
1.0
1.0
1.0
1.0
1.0
1.0
1.0
1.0
1.0


266.0
0.0
0.0
0.0
0.0
0.0
84.0
0.0
0.0
0.0
0.0
0.0
0.0
0.0
0.0
0.0
182.0
0.0
0.0
0.0
0.0
0.0
0.0
0.0
0.0
0.0
0.0
0.0
0.0
0.0

1.0
1.0
1.0
1.0
1.0
1.0
1.0
1.0
1.0
1.0
1.0
1.0
1.0
1.0
1.0
1.0
1.0
1.0
1.0
1.0
1.0
1.0
1.0
1.0
1.0
1.0
1.0
1.0
1.0
1.0


38.0
0.0
0.0
0.0
0.0
38.0
0.0
0.0
0.0
0.0
0.0
0.0
0.0
0.0
0.0
0.0
0.0
0.0
0.0
0.0
0.0
0.0
0.0
0.0
0.0
0.0
0.0
0.0
0.0
0.0


38.0
0.0
0.0
0.0
0.0
38.0
0.0
0.0
0.0
0.0
0.0
0.0
0.0
0.0
0.0
0.0
0.0
0.0
0.0
0.0
0.0
0.0
0.0
0.0
0.0
0.0
0.0
0.0
0.0
0.0

1.0
1.0
1.0
1.0
1.0
1.0
1.0
1.0
1.0
1.0
1.0
1.0
1.0
1.0
1.0
1.0
1.0
1.0
1.0
1.0
1.0
1.0
1.0
1.0
1.0
1.0
1.0
1.0
1.0
1.0


30.0
0.0
30.0
0.0
0.0
0.0
0.0
0.0
0.0
0.0
0.0
0.0
0.0
0.0
0.0
0.0
0.0
0.0
0.0
0.0
0.0
0.0
0.0
0.0
0.0
0.0
0.0
0.0
0.0
0.0


30.0
0.0
30.0
0.0
0.0
0.0
0.0
0.0
0.0
0.0
0.0
0.0
0.0
0.0
0.0
0.0
0.0
0.0
0.0
0.0
0.0
0.0
0.0
0.0
0.0
0.0
0.0
0.0
0.0
0.0

1.0
1.0
1.0
1.0
1.0
1.0
1.0
1.0
1.0
1.0
1.0
1.0
1.0
1.0
1.0
1.0
1.0
1.0
1.0
1.0
1.0
1.0
1.0
1.0
1.0
1.0
1.0
1.0
1.0
1.0


25.0
0.0
12.0
0.0
0.0
0.0
0.0
0.0
0.0
13.0
0.0
0.0
0.0
0.0
0.0
0.0
0.0
0.0
0.0
0.0
0.0
0.0
0.0
0.0
0.0
0.0
0.0
0.0
0.0
0.0

0.99
0.99
0.99
0.99
0.99
0.99
0.99
0.99
0.99
0.99
0.99
0.99
0.99
0.99
0.99
0.99
0.99
0.99
0.99
0.99
0.99
0.99
0.99
0.99
0.99
0.99
0.99
0.99
0.99
0.99


12.0
0.0
12.0
0.0
0.0
0.0
0.0
0.0
0.0
0.0
0.0
0.0
0.0
0.0
0.0
0.0
0.0
0.0
0.0
0.0
0.0
0.0
0.0
0.0
0.0
0.0
0.0
0.0
0.0
0.0

1.0
1.0
1.0
1.0
1.0
1.0
1.0
1.0
1.0
1.0
1.0
1.0
1.0
1.0
1.0
1.0
1.0
1.0
1.0
1.0
1.0
1.0
1.0
1.0
1.0
1.0
1.0
1.0
1.0
1.0


229478.0
12.0
14.0
19.0
0.0
0.0
36.0
76.0
137.0
28.0
0.0
0.0
0.0
24953.0
0.0
0.0
0.0
0.0
0.0
0.0
0.0
0.0
0.0
0.0
204203.0
0.0
0.0
0.0
0.0
0.0

0.81
0.81
0.81
0.81
0.81
0.81
0.81
0.81
0.81
0.81
0.81
0.81
0.81
0.81
0.81
0.81
0.81
0.81
0.81
0.81
0.81
0.81
0.81
0.81
0.81
0.81
0.81
0.81
0.81
0.81


289.0
12.0
0.0
0.0
0.0
0.0
36.0
76.0
137.0
28.0
0.0
0.0
0.0
0.0
0.0
0.0
0.0
0.0
0.0
0.0
0.0
0.0
0.0
0.0
0.0
0.0
0.0
0.0
0.0
0.0


289.0
12.0
0.0
0.0
0.0
0.0
36.0
76.0
137.0
28.0
0.0
0.0
0.0
0.0
0.0
0.0
0.0
0.0
0.0
0.0
0.0
0.0
0.0
0.0
0.0
0.0
0.0
0.0
0.0
0.0

0.97
0.97
0.97
0.97
0.97
0.97
0.97
0.97
0.97
0.97
0.97
0.97
0.97
0.97
0.97
0.97
0.97
0.97
0.97
0.97
0.97
0.97
0.97
0.97
0.97
0.97
0.97
0.97
0.97
0.97


33.0
0.0
14.0
19.0
0.0
0.0
0.0
0.0
0.0
0.0
0.0
0.0
0.0
0.0
0.0
0.0
0.0
0.0
0.0
0.0
0.0
0.0
0.0
0.0
0.0
0.0
0.0
0.0
0.0
0.0


33.0
0.0
14.0
19.0
0.0
0.0
0.0
0.0
0.0
0.0
0.0
0.0
0.0
0.0
0.0
0.0
0.0
0.0
0.0
0.0
0.0
0.0
0.0
0.0
0.0
0.0
0.0
0.0
0.0
0.0

1.0
1.0
1.0
1.0
1.0
1.0
1.0
1.0
1.0
1.0
1.0
1.0
1.0
1.0
1.0
1.0
1.0
1.0
1.0
1.0
1.0
1.0
1.0
1.0
1.0
1.0
1.0
1.0
1.0
1.0


1432.0
31.0
22.0
26.0
44.0
145.0
96.0
0.0
0.0
0.0
0.0
21.0
0.0
0.0
0.0
0.0
0.0
0.0
0.0
0.0
0.0
0.0
0.0
0.0
879.0
29.0
139.0
0.0
0.0
0.0


1432.0
31.0
22.0
26.0
44.0
145.0
96.0
0.0
0.0
0.0
0.0
21.0
0.0
0.0
0.0
0.0
0.0
0.0
0.0
0.0
0.0
0.0
0.0
0.0
879.0
29.0
139.0
0.0
0.0
0.0


1432.0
31.0
22.0
26.0
44.0
145.0
96.0
0.0
0.0
0.0
0.0
21.0
0.0
0.0
0.0
0.0
0.0
0.0
0.0
0.0
0.0
0.0
0.0
0.0
879.0
29.0
139.0
0.0
0.0
0.0

1.0
1.0
1.0
1.0
1.0
1.0
1.0
1.0
1.0
1.0
1.0
1.0
1.0
1.0
1.0
1.0
1.0
1.0
1.0
1.0
1.0
1.0
1.0
1.0
1.0
1.0
1.0
1.0
1.0
1.0


28530.0
0.0
0.0
0.0
0.0
0.0
0.0
255.0
0.0
28.0
0.0
127.0
0.0
0.0
0.0
0.0
0.0
0.0
28120.0
0.0
0.0
0.0
0.0
0.0
0.0
0.0
0.0
0.0
0.0
0.0


255.0
0.0
0.0
0.0
0.0
0.0
0.0
255.0
0.0
0.0
0.0
0.0
0.0
0.0
0.0
0.0
0.0
0.0
0.0
0.0
0.0
0.0
0.0
0.0
0.0
0.0
0.0
0.0
0.0
0.0

1.0
1.0
1.0
1.0
1.0
1.0
1.0
1.0
1.0
1.0
1.0
1.0
1.0
1.0
1.0
1.0
1.0
1.0
1.0
1.0
1.0
1.0
1.0
1.0
1.0
1.0
1.0
1.0
1.0
1.0


127.0
0.0
0.0
0.0
0.0
0.0
0.0
0.0
0.0
0.0
0.0
127.0
0.0
0.0
0.0
0.0
0.0
0.0
0.0
0.0
0.0
0.0
0.0
0.0
0.0
0.0
0.0
0.0
0.0
0.0


127.0
0.0
0.0
0.0
0.0
0.0
0.0
0.0
0.0
0.0
0.0
127.0
0.0
0.0
0.0
0.0
0.0
0.0
0.0
0.0
0.0
0.0
0.0
0.0
0.0
0.0
0.0
0.0
0.0
0.0

1.0
1.0
1.0
1.0
1.0
1.0
1.0
1.0
1.0
1.0
1.0
1.0
1.0
1.0
1.0
1.0
1.0
1.0
1.0
1.0
1.0
1.0
1.0
1.0
1.0
1.0
1.0
1.0
1.0
1.0


28.0
0.0
0.0
0.0
0.0
0.0
0.0
0.0
0.0
28.0
0.0
0.0
0.0
0.0
0.0
0.0
0.0
0.0
0.0
0.0
0.0
0.0
0.0
0.0
0.0
0.0
0.0
0.0
0.0
0.0

0.83
0.83
0.83
0.83
0.83
0.83
0.83
0.83
0.83
0.83
0.83
0.83
0.83
0.83
0.83
0.83
0.83
0.83
0.83
0.83
0.83
0.83
0.83
0.83
0.83
0.83
0.83
0.83
0.83
0.83


28120.0
0.0
0.0
0.0
0.0
0.0
0.0
0.0
0.0
0.0
0.0
0.0
0.0
0.0
0.0
0.0
0.0
0.0
28120.0
0.0
0.0
0.0
0.0
0.0
0.0
0.0
0.0
0.0
0.0
0.0


28120.0
0.0
0.0
0.0
0.0
0.0
0.0
0.0
0.0
0.0
0.0
0.0
0.0
0.0
0.0
0.0
0.0
0.0
28120.0
0.0
0.0
0.0
0.0
0.0
0.0
0.0
0.0
0.0
0.0
0.0

1.0
1.0
1.0
1.0
1.0
1.0
1.0
1.0
1.0
1.0
1.0
1.0
1.0
1.0
1.0
1.0
1.0
1.0
1.0
1.0
1.0
1.0
1.0
1.0
1.0
1.0
1.0
1.0
1.0
1.0


431.0
49.0
1.0
0.0
0.0
0.0
0.0
121.0
67.0
27.0
148.0
18.0
0.0
0.0
0.0
0.0
0.0
0.0
0.0
0.0
0.0
0.0
0.0
0.0
0.0
0.0
0.0
0.0
0.0
0.0


302.0
49.0
1.0
0.0
0.0
0.0
0.0
121.0
67.0
8.0
38.0
18.0
0.0
0.0
0.0
0.0
0.0
0.0
0.0
0.0
0.0
0.0
0.0
0.0
0.0
0.0
0.0
0.0
0.0
0.0


302.0
49.0
1.0
0.0
0.0
0.0
0.0
121.0
67.0
8.0
38.0
18.0
0.0
0.0
0.0
0.0
0.0
0.0
0.0
0.0
0.0
0.0
0.0
0.0
0.0
0.0
0.0
0.0
0.0
0.0

1.0
1.0
1.0
1.0
1.0
1.0
1.0
1.0
1.0
1.0
1.0
1.0
1.0
1.0
1.0
1.0
1.0
1.0
1.0
1.0
1.0
1.0
1.0
1.0
1.0
1.0
1.0
1.0
1.0
1.0


129.0
0.0
0.0
0.0
0.0
0.0
0.0
0.0
0.0
19.0
110.0
0.0
0.0
0.0
0.0
0.0
0.0
0.0
0.0
0.0
0.0
0.0
0.0
0.0
0.0
0.0
0.0
0.0
0.0
0.0

0.86
0.86
0.86
0.86
0.86
0.86
0.86
0.86
0.86
0.86
0.86
0.86
0.86
0.86
0.86
0.86
0.86
0.86
0.86
0.86
0.86
0.86
0.86
0.86
0.86
0.86
0.86
0.86
0.86
0.86


8074.0
0.0
0.0
0.0
0.0
0.0
227.0
0.0
0.0
0.0
0.0
0.0
0.0
0.0
0.0
0.0
0.0
0.0
0.0
0.0
0.0
0.0
0.0
0.0
0.0
0.0
0.0
7847.0
0.0
0.0


8074.0
0.0
0.0
0.0
0.0
0.0
227.0
0.0
0.0
0.0
0.0
0.0
0.0
0.0
0.0
0.0
0.0
0.0
0.0
0.0
0.0
0.0
0.0
0.0
0.0
0.0
0.0
7847.0
0.0
0.0


8074.0
0.0
0.0
0.0
0.0
0.0
227.0
0.0
0.0
0.0
0.0
0.0
0.0
0.0
0.0
0.0
0.0
0.0
0.0
0.0
0.0
0.0
0.0
0.0
0.0
0.0
0.0
7847.0
0.0
0.0

1.0
1.0
1.0
1.0
1.0
1.0
1.0
1.0
1.0
1.0
1.0
1.0
1.0
1.0
1.0
1.0
1.0
1.0
1.0
1.0
1.0
1.0
1.0
1.0
1.0
1.0
1.0
1.0
1.0
1.0


287.0
9.0
14.0
135.0
0.0
15.0
0.0
0.0
0.0
0.0
0.0
0.0
0.0
0.0
0.0
0.0
0.0
0.0
0.0
0.0
0.0
0.0
0.0
0.0
114.0
0.0
0.0
0.0
0.0
0.0


263.0
9.0
14.0
111.0
0.0
15.0
0.0
0.0
0.0
0.0
0.0
0.0
0.0
0.0
0.0
0.0
0.0
0.0
0.0
0.0
0.0
0.0
0.0
0.0
114.0
0.0
0.0
0.0
0.0
0.0

1.0
1.0
1.0
1.0
1.0
1.0
1.0
1.0
1.0
1.0
1.0
1.0
1.0
1.0
1.0
1.0
1.0
1.0
1.0
1.0
1.0
1.0
1.0
1.0
1.0
1.0
1.0
1.0
1.0
1.0


244.0
9.0
14.0
111.0
0.0
0.0
0.0
0.0
0.0
0.0
0.0
0.0
0.0
0.0
0.0
0.0
0.0
0.0
0.0
0.0
0.0
0.0
0.0
0.0
110.0
0.0
0.0
0.0
0.0
0.0

1.0
1.0
1.0
1.0
1.0
1.0
1.0
1.0
1.0
1.0
1.0
1.0
1.0
1.0
1.0
1.0
1.0
1.0
1.0
1.0
1.0
1.0
1.0
1.0
1.0
1.0
1.0
1.0
1.0
1.0


15.0
0.0
0.0
0.0
0.0
15.0
0.0
0.0
0.0
0.0
0.0
0.0
0.0
0.0
0.0
0.0
0.0
0.0
0.0
0.0
0.0
0.0
0.0
0.0
0.0
0.0
0.0
0.0
0.0
0.0

0.82
0.82
0.82
0.82
0.82
0.82
0.82
0.82
0.82
0.82
0.82
0.82
0.82
0.82
0.82
0.82
0.82
0.82
0.82
0.82
0.82
0.82
0.82
0.82
0.82
0.82
0.82
0.82
0.82
0.82


24.0
0.0
0.0
24.0
0.0
0.0
0.0
0.0
0.0
0.0
0.0
0.0
0.0
0.0
0.0
0.0
0.0
0.0
0.0
0.0
0.0
0.0
0.0
0.0
0.0
0.0
0.0
0.0
0.0
0.0

1.0
1.0
1.0
1.0
1.0
1.0
1.0
1.0
1.0
1.0
1.0
1.0
1.0
1.0
1.0
1.0
1.0
1.0
1.0
1.0
1.0
1.0
1.0
1.0
1.0
1.0
1.0
1.0
1.0
1.0


3578.0
16.0
25.0
1.0
1.0
14.0
10.0
0.0
0.0
0.0
0.0
17.0
0.0
0.0
0.0
0.0
0.0
0.0
0.0
0.0
3494.0
0.0
0.0
0.0
0.0
0.0
0.0
0.0
0.0
0.0


40.0
16.0
18.0
0.0
0.0
0.0
0.0
0.0
0.0
0.0
0.0
6.0
0.0
0.0
0.0
0.0
0.0
0.0
0.0
0.0
0.0
0.0
0.0
0.0
0.0
0.0
0.0
0.0
0.0
0.0


40.0
16.0
18.0
0.0
0.0
0.0
0.0
0.0
0.0
0.0
0.0
6.0
0.0
0.0
0.0
0.0
0.0
0.0
0.0
0.0
0.0
0.0
0.0
0.0
0.0
0.0
0.0
0.0
0.0
0.0

0.92
0.92
0.92
0.92
0.92
0.92
0.92
0.92
0.92
0.92
0.92
0.92
0.92
0.92
0.92
0.92
0.92
0.92
0.92
0.92
0.92
0.92
0.92
0.92
0.92
0.92
0.92
0.92
0.92
0.92


33.0
0.0
7.0
1.0
1.0
14.0
10.0
0.0
0.0
0.0
0.0
0.0
0.0
0.0
0.0
0.0
0.0
0.0
0.0
0.0
0.0
0.0
0.0
0.0
0.0
0.0
0.0
0.0
0.0
0.0


33.0
0.0
7.0
1.0
1.0
14.0
10.0
0.0
0.0
0.0
0.0
0.0
0.0
0.0
0.0
0.0
0.0
0.0
0.0
0.0
0.0
0.0
0.0
0.0
0.0
0.0
0.0
0.0
0.0
0.0

0.89
0.89
0.89
0.89
0.89
0.89
0.89
0.89
0.89
0.89
0.89
0.89
0.89
0.89
0.89
0.89
0.89
0.89
0.89
0.89
0.89
0.89
0.89
0.89
0.89
0.89
0.89
0.89
0.89
0.89


11.0
0.0
0.0
0.0
0.0
0.0
0.0
0.0
0.0
0.0
0.0
11.0
0.0
0.0
0.0
0.0
0.0
0.0
0.0
0.0
0.0
0.0
0.0
0.0
0.0
0.0
0.0
0.0
0.0
0.0


11.0
0.0
0.0
0.0
0.0
0.0
0.0
0.0
0.0
0.0
0.0
11.0
0.0
0.0
0.0
0.0
0.0
0.0
0.0
0.0
0.0
0.0
0.0
0.0
0.0
0.0
0.0
0.0
0.0
0.0

1.0
1.0
1.0
1.0
1.0
1.0
1.0
1.0
1.0
1.0
1.0
1.0
1.0
1.0
1.0
1.0
1.0
1.0
1.0
1.0
1.0
1.0
1.0
1.0
1.0
1.0
1.0
1.0
1.0
1.0


3494.0
0.0
0.0
0.0
0.0
0.0
0.0
0.0
0.0
0.0
0.0
0.0
0.0
0.0
0.0
0.0
0.0
0.0
0.0
0.0
3494.0
0.0
0.0
0.0
0.0
0.0
0.0
0.0
0.0
0.0


3494.0
0.0
0.0
0.0
0.0
0.0
0.0
0.0
0.0
0.0
0.0
0.0
0.0
0.0
0.0
0.0
0.0
0.0
0.0
0.0
3494.0
0.0
0.0
0.0
0.0
0.0
0.0
0.0
0.0
0.0

1.0
1.0
1.0
1.0
1.0
1.0
1.0
1.0
1.0
1.0
1.0
1.0
1.0
1.0
1.0
1.0
1.0
1.0
1.0
1.0
1.0
1.0
1.0
1.0
1.0
1.0
1.0
1.0
1.0
1.0


551.0
3.0
2.0
6.0
0.0
0.0
0.0
0.0
0.0
0.0
0.0
0.0
0.0
0.0
0.0
0.0
0.0
0.0
0.0
0.0
0.0
0.0
0.0
0.0
0.0
0.0
540.0
0.0
0.0
0.0


551.0
3.0
2.0
6.0
0.0
0.0
0.0
0.0
0.0
0.0
0.0
0.0
0.0
0.0
0.0
0.0
0.0
0.0
0.0
0.0
0.0
0.0
0.0
0.0
0.0
0.0
540.0
0.0
0.0
0.0


551.0
3.0
2.0
6.0
0.0
0.0
0.0
0.0
0.0
0.0
0.0
0.0
0.0
0.0
0.0
0.0
0.0
0.0
0.0
0.0
0.0
0.0
0.0
0.0
0.0
0.0
540.0
0.0
0.0
0.0

1.0
1.0
1.0
1.0
1.0
1.0
1.0
1.0
1.0
1.0
1.0
1.0
1.0
1.0
1.0
1.0
1.0
1.0
1.0
1.0
1.0
1.0
1.0
1.0
1.0
1.0
1.0
1.0
1.0
1.0


2764.0
0.0
0.0
0.0
0.0
0.0
0.0
0.0
0.0
0.0
0.0
0.0
0.0
0.0
0.0
0.0
0.0
0.0
1121.0
0.0
0.0
0.0
0.0
0.0
0.0
0.0
0.0
0.0
0.0
1643.0


2764.0
0.0
0.0
0.0
0.0
0.0
0.0
0.0
0.0
0.0
0.0
0.0
0.0
0.0
0.0
0.0
0.0
0.0
1121.0
0.0
0.0
0.0
0.0
0.0
0.0
0.0
0.0
0.0
0.0
1643.0

1.0
1.0
1.0
1.0
1.0
1.0
1.0
1.0
1.0
1.0
1.0
1.0
1.0
1.0
1.0
1.0
1.0
1.0
1.0
1.0
1.0
1.0
1.0
1.0
1.0
1.0
1.0
1.0
1.0
1.0


586.0
0.0
0.0
0.0
0.0
0.0
0.0
0.0
0.0
0.0
0.0
0.0
0.0
0.0
0.0
0.0
0.0
0.0
0.0
0.0
0.0
0.0
0.0
0.0
432.0
0.0
145.0
0.0
0.0
9.0

0.97
0.97
0.97
0.97
0.97
0.97
0.97
0.97
0.97
0.97
0.97
0.97
0.97
0.97
0.97
0.97
0.97
0.97
0.97
0.97
0.97
0.97
0.97
0.97
0.97
0.97
0.97
0.97
0.97
0.97


500.0
0.0
0.0
0.0
0.0
0.0
0.0
0.0
0.0
0.0
0.0
0.0
0.0
0.0
0.0
0.0
0.0
0.0
0.0
0.0
0.0
0.0
0.0
0.0
397.0
0.0
100.0
0.0
0.0
3.0

1.0
1.0
1.0
1.0
1.0
1.0
1.0
1.0
1.0
1.0
1.0
1.0
1.0
1.0
1.0
1.0
1.0
1.0
1.0
1.0
1.0
1.0
1.0
1.0
1.0
1.0
1.0
1.0
1.0
1.0


59.0
0.0
0.0
0.0
0.0
0.0
0.0
0.0
0.0
0.0
0.0
0.0
0.0
0.0
0.0
0.0
0.0
0.0
0.0
0.0
0.0
0.0
0.0
0.0
59.0
0.0
0.0
0.0
0.0
0.0


59.0
0.0
0.0
0.0
0.0
0.0
0.0
0.0
0.0
0.0
0.0
0.0
0.0
0.0
0.0
0.0
0.0
0.0
0.0
0.0
0.0
0.0
0.0
0.0
59.0
0.0
0.0
0.0
0.0
0.0


59.0
0.0
0.0
0.0
0.0
0.0
0.0
0.0
0.0
0.0
0.0
0.0
0.0
0.0
0.0
0.0
0.0
0.0
0.0
0.0
0.0
0.0
0.0
0.0
59.0
0.0
0.0
0.0
0.0
0.0

1.0
1.0
1.0
1.0
1.0
1.0
1.0
1.0
1.0
1.0
1.0
1.0
1.0
1.0
1.0
1.0
1.0
1.0
1.0
1.0
1.0
1.0
1.0
1.0
1.0
1.0
1.0
1.0
1.0
1.0


9.0
0.0
0.0
0.0
0.0
0.0
0.0
0.0
0.0
0.0
0.0
0.0
0.0
0.0
0.0
0.0
0.0
0.0
0.0
0.0
0.0
0.0
0.0
0.0
0.0
0.0
0.0
0.0
9.0
0.0

0.88
0.88
0.88
0.88
0.88
0.88
0.88
0.88
0.88
0.88
0.88
0.88
0.88
0.88
0.88
0.88
0.88
0.88
0.88
0.88
0.88
0.88
0.88
0.88
0.88
0.88
0.88
0.88
0.88
0.88


14673988.0
9629.0
5702.0
33641.0
6699.0
3744.0
5632.0
39205.0
30081.0
79643.0
47225.0
44395.0
35188.0
185331.0
530052.0
3544517.0
1501676.0
756405.0
413942.0
564802.0
333611.0
567634.0
489868.0
35188.0
1524527.0
1200211.0
797446.0
505125.0
1117824.0
265045.0

1.0
1.0
1.0
1.0
1.0
1.0
1.0
1.0
1.0
1.0
1.0
1.0
1.0
1.0
1.0
1.0
1.0
1.0
1.0
1.0
1.0
1.0
1.0
1.0
1.0
1.0
1.0
1.0
1.0
1.0


154641.0
372.0
0.0
21.0
0.0
51.0
19.0
0.0
0.0
38612.0
15965.0
717.0
0.0
0.0
0.0
0.0
0.0
1839.0
23331.0
1586.0
631.0
5114.0
0.0
0.0
0.0
24825.0
2882.0
6845.0
30513.0
1318.0

1.0
1.0
1.0
1.0
1.0
1.0
1.0
1.0
1.0
1.0
1.0
1.0
1.0
1.0
1.0
1.0
1.0
1.0
1.0
1.0
1.0
1.0
1.0
1.0
1.0
1.0
1.0
1.0
1.0
1.0


114745.0
372.0
0.0
21.0
0.0
10.0
19.0
0.0
0.0
38612.0
15965.0
717.0
0.0
0.0
0.0
0.0
0.0
1611.0
701.0
1586.0
631.0
5114.0
0.0
0.0
0.0
21089.0
2882.0
4296.0
19801.0
1318.0

1.0
1.0
1.0
1.0
1.0
1.0
1.0
1.0
1.0
1.0
1.0
1.0
1.0
1.0
1.0
1.0
1.0
1.0
1.0
1.0
1.0
1.0
1.0
1.0
1.0
1.0
1.0
1.0
1.0
1.0


62390.0
367.0
0.0
0.0
0.0
0.0
0.0
0.0
0.0
38379.0
15824.0
716.0
0.0
0.0
0.0
0.0
0.0
0.0
0.0
1511.0
576.0
5017.0
0.0
0.0
0.0
0.0
0.0
0.0
0.0
0.0

0.99
0.99
0.99
0.99
0.99
0.99
0.99
0.99
0.99
0.99
0.99
0.99
0.99
0.99
0.99
0.99
0.99
0.99
0.99
0.99
0.99
0.99
0.99
0.99
0.99
0.99
0.99
0.99
0.99
0.99


70.0
0.0
0.0
21.0
0.0
0.0
0.0
0.0
0.0
0.0
49.0
0.0
0.0
0.0
0.0
0.0
0.0
0.0
0.0
0.0
0.0
0.0
0.0
0.0
0.0
0.0
0.0
0.0
0.0
0.0

1.0
1.0
1.0
1.0
1.0
1.0
1.0
1.0
1.0
1.0
1.0
1.0
1.0
1.0
1.0
1.0
1.0
1.0
1.0
1.0
1.0
1.0
1.0
1.0
1.0
1.0
1.0
1.0
1.0
1.0


42.0
0.0
0.0
0.0
0.0
0.0
0.0
0.0
0.0
42.0
0.0
0.0
0.0
0.0
0.0
0.0
0.0
0.0
0.0
0.0
0.0
0.0
0.0
0.0
0.0
0.0
0.0
0.0
0.0
0.0

0.98
0.98
0.98
0.98
0.98
0.98
0.98
0.98
0.98
0.98
0.98
0.98
0.98
0.98
0.98
0.98
0.98
0.98
0.98
0.98
0.98
0.98
0.98
0.98
0.98
0.98
0.98
0.98
0.98
0.98


2431.0
0.0
0.0
0.0
0.0
0.0
0.0
0.0
0.0
9.0
22.0
0.0
0.0
0.0
0.0
0.0
0.0
1611.0
0.0
0.0
0.0
0.0
0.0
0.0
0.0
0.0
0.0
0.0
789.0
0.0

0.9
0.9
0.9
0.9
0.9
0.9
0.9
0.9
0.9
0.9
0.9
0.9
0.9
0.9
0.9
0.9
0.9
0.9
0.9
0.9
0.9
0.9
0.9
0.9
0.9
0.9
0.9
0.9
0.9
0.9


19554.0
0.0
0.0
0.0
0.0
10.0
19.0
0.0
0.0
0.0
0.0
0.0
0.0
0.0
0.0
0.0
0.0
0.0
0.0
0.0
0.0
0.0
0.0
0.0
0.0
14283.0
1868.0
3230.0
0.0
144.0

1.0
1.0
1.0
1.0
1.0
1.0
1.0
1.0
1.0
1.0
1.0
1.0
1.0
1.0
1.0
1.0
1.0
1.0
1.0
1.0
1.0
1.0
1.0
1.0
1.0
1.0
1.0
1.0
1.0
1.0


18.0
0.0
0.0
0.0
0.0
0.0
0.0
0.0
0.0
18.0
0.0
0.0
0.0
0.0
0.0
0.0
0.0
0.0
0.0
0.0
0.0
0.0
0.0
0.0
0.0
0.0
0.0
0.0
0.0
0.0

1.0
1.0
1.0
1.0
1.0
1.0
1.0
1.0
1.0
1.0
1.0
1.0
1.0
1.0
1.0
1.0
1.0
1.0
1.0
1.0
1.0
1.0
1.0
1.0
1.0
1.0
1.0
1.0
1.0
1.0


18584.0
0.0
0.0
0.0
0.0
0.0
0.0
0.0
0.0
0.0
0.0
0.0
0.0
0.0
0.0
0.0
0.0
0.0
0.0
0.0
0.0
0.0
0.0
0.0
0.0
0.0
0.0
0.0
18584.0
0.0

1.0
1.0
1.0
1.0
1.0
1.0
1.0
1.0
1.0
1.0
1.0
1.0
1.0
1.0
1.0
1.0
1.0
1.0
1.0
1.0
1.0
1.0
1.0
1.0
1.0
1.0
1.0
1.0
1.0
1.0


1397.0
0.0
0.0
0.0
0.0
0.0
0.0
0.0
0.0
0.0
0.0
0.0
0.0
0.0
0.0
0.0
0.0
0.0
0.0
0.0
0.0
0.0
0.0
0.0
0.0
1364.0
33.0
0.0
0.0
0.0

0.95
0.95
0.95
0.95
0.95
0.95
0.95
0.95
0.95
0.95
0.95
0.95
0.95
0.95
0.95
0.95
0.95
0.95
0.95
0.95
0.95
0.95
0.95
0.95
0.95
0.95
0.95
0.95
0.95
0.95


697.0
0.0
0.0
0.0
0.0
0.0
0.0
0.0
0.0
0.0
0.0
0.0
0.0
0.0
0.0
0.0
0.0
0.0
697.0
0.0
0.0
0.0
0.0
0.0
0.0
0.0
0.0
0.0
0.0
0.0

1.0
1.0
1.0
1.0
1.0
1.0
1.0
1.0
1.0
1.0
1.0
1.0
1.0
1.0
1.0
1.0
1.0
1.0
1.0
1.0
1.0
1.0
1.0
1.0
1.0
1.0
1.0
1.0
1.0
1.0


126.0
0.0
0.0
0.0
0.0
0.0
0.0
0.0
0.0
0.0
0.0
0.0
0.0
0.0
0.0
0.0
0.0
0.0
0.0
0.0
0.0
0.0
0.0
0.0
0.0
126.0
0.0
0.0
0.0
0.0

0.88
0.88
0.88
0.88
0.88
0.88
0.88
0.88
0.88
0.88
0.88
0.88
0.88
0.88
0.88
0.88
0.88
0.88
0.88
0.88
0.88
0.88
0.88
0.88
0.88
0.88
0.88
0.88
0.88
0.88


250.0
0.0
0.0
0.0
0.0
0.0
0.0
0.0
0.0
0.0
0.0
0.0
0.0
0.0
0.0
0.0
0.0
0.0
0.0
0.0
0.0
0.0
0.0
0.0
0.0
0.0
0.0
0.0
250.0
0.0

1.0
1.0
1.0
1.0
1.0
1.0
1.0
1.0
1.0
1.0
1.0
1.0
1.0
1.0
1.0
1.0
1.0
1.0
1.0
1.0
1.0
1.0
1.0
1.0
1.0
1.0
1.0
1.0
1.0
1.0


13530.0
0.0
0.0
0.0
0.0
41.0
0.0
0.0
0.0
0.0
0.0
0.0
0.0
0.0
0.0
0.0
0.0
228.0
0.0
0.0
0.0
0.0
0.0
0.0
0.0
0.0
0.0
2549.0
10712.0
0.0

1.0
1.0
1.0
1.0
1.0
1.0
1.0
1.0
1.0
1.0
1.0
1.0
1.0
1.0
1.0
1.0
1.0
1.0
1.0
1.0
1.0
1.0
1.0
1.0
1.0
1.0
1.0
1.0
1.0
1.0


9314.0
0.0
0.0
0.0
0.0
41.0
0.0
0.0
0.0
0.0
0.0
0.0
0.0
0.0
0.0
0.0
0.0
0.0
0.0
0.0
0.0
0.0
0.0
0.0
0.0
0.0
0.0
2486.0
6787.0
0.0

1.0
1.0
1.0
1.0
1.0
1.0
1.0
1.0
1.0
1.0
1.0
1.0
1.0
1.0
1.0
1.0
1.0
1.0
1.0
1.0
1.0
1.0
1.0
1.0
1.0
1.0
1.0
1.0
1.0
1.0


195.0
0.0
0.0
0.0
0.0
0.0
0.0
0.0
0.0
0.0
0.0
0.0
0.0
0.0
0.0
0.0
0.0
195.0
0.0
0.0
0.0
0.0
0.0
0.0
0.0
0.0
0.0
0.0
0.0
0.0

1.0
1.0
1.0
1.0
1.0
1.0
1.0
1.0
1.0
1.0
1.0
1.0
1.0
1.0
1.0
1.0
1.0
1.0
1.0
1.0
1.0
1.0
1.0
1.0
1.0
1.0
1.0
1.0
1.0
1.0


26186.0
0.0
0.0
0.0
0.0
0.0
0.0
0.0
0.0
0.0
0.0
0.0
0.0
0.0
0.0
0.0
0.0
0.0
22450.0
0.0
0.0
0.0
0.0
0.0
0.0
3736.0
0.0
0.0
0.0
0.0


22450.0
0.0
0.0
0.0
0.0
0.0
0.0
0.0
0.0
0.0
0.0
0.0
0.0
0.0
0.0
0.0
0.0
0.0
22450.0
0.0
0.0
0.0
0.0
0.0
0.0
0.0
0.0
0.0
0.0
0.0

1.0
1.0
1.0
1.0
1.0
1.0
1.0
1.0
1.0
1.0
1.0
1.0
1.0
1.0
1.0
1.0
1.0
1.0
1.0
1.0
1.0
1.0
1.0
1.0
1.0
1.0
1.0
1.0
1.0
1.0


3736.0
0.0
0.0
0.0
0.0
0.0
0.0
0.0
0.0
0.0
0.0
0.0
0.0
0.0
0.0
0.0
0.0
0.0
0.0
0.0
0.0
0.0
0.0
0.0
0.0
3736.0
0.0
0.0
0.0
0.0

1.0
1.0
1.0
1.0
1.0
1.0
1.0
1.0
1.0
1.0
1.0
1.0
1.0
1.0
1.0
1.0
1.0
1.0
1.0
1.0
1.0
1.0
1.0
1.0
1.0
1.0
1.0
1.0
1.0
1.0


4772407.0
3500.0
23.0
0.0
0.0
1.0
1665.0
0.0
90.0
24693.0
15810.0
3790.0
0.0
0.0
0.0
3195435.0
1178261.0
15875.0
0.0
0.0
35307.0
0.0
0.0
0.0
24137.0
107548.0
34338.0
26003.0
75933.0
29998.0


504663.0
3500.0
0.0
0.0
0.0
1.0
1665.0
0.0
0.0
24451.0
15809.0
3790.0
0.0
0.0
0.0
113567.0
46280.0
15875.0
0.0
0.0
35307.0
0.0
0.0
0.0
0.0
84733.0
31577.0
23526.0
75933.0
28649.0


504663.0
3500.0
0.0
0.0
0.0
1.0
1665.0
0.0
0.0
24451.0
15809.0
3790.0
0.0
0.0
0.0
113567.0
46280.0
15875.0
0.0
0.0
35307.0
0.0
0.0
0.0
0.0
84733.0
31577.0
23526.0
75933.0
28649.0

1.0
1.0
1.0
1.0
1.0
1.0
1.0
1.0
1.0
1.0
1.0
1.0
1.0
1.0
1.0
1.0
1.0
1.0
1.0
1.0
1.0
1.0
1.0
1.0
1.0
1.0
1.0
1.0
1.0
1.0


333.0
0.0
0.0
0.0
0.0
0.0
0.0
0.0
90.0
242.0
1.0
0.0
0.0
0.0
0.0
0.0
0.0
0.0
0.0
0.0
0.0
0.0
0.0
0.0
0.0
0.0
0.0
0.0
0.0
0.0


243.0
0.0
0.0
0.0
0.0
0.0
0.0
0.0
0.0
242.0
1.0
0.0
0.0
0.0
0.0
0.0
0.0
0.0
0.0
0.0
0.0
0.0
0.0
0.0
0.0
0.0
0.0
0.0
0.0
0.0

0.95
0.95
0.95
0.95
0.95
0.95
0.95
0.95
0.95
0.95
0.95
0.95
0.95
0.95
0.95
0.95
0.95
0.95
0.95
0.95
0.95
0.95
0.95
0.95
0.95
0.95
0.95
0.95
0.95
0.95


90.0
0.0
0.0
0.0
0.0
0.0
0.0
0.0
90.0
0.0
0.0
0.0
0.0
0.0
0.0
0.0
0.0
0.0
0.0
0.0
0.0
0.0
0.0
0.0
0.0
0.0
0.0
0.0
0.0
0.0

1.0
1.0
1.0
1.0
1.0
1.0
1.0
1.0
1.0
1.0
1.0
1.0
1.0
1.0
1.0
1.0
1.0
1.0
1.0
1.0
1.0
1.0
1.0
1.0
1.0
1.0
1.0
1.0
1.0
1.0


15438.0
0.0
23.0
0.0
0.0
0.0
0.0
0.0
0.0
0.0
0.0
0.0
0.0
0.0
0.0
0.0
0.0
0.0
0.0
0.0
0.0
0.0
0.0
0.0
0.0
8829.0
2760.0
2477.0
0.0
1349.0


11161.0
0.0
23.0
0.0
0.0
0.0
0.0
0.0
0.0
0.0
0.0
0.0
0.0
0.0
0.0
0.0
0.0
0.0
0.0
0.0
0.0
0.0
0.0
0.0
0.0
7452.0
0.0
2420.0
0.0
1266.0

1.0
1.0
1.0
1.0
1.0
1.0
1.0
1.0
1.0
1.0
1.0
1.0
1.0
1.0
1.0
1.0
1.0
1.0
1.0
1.0
1.0
1.0
1.0
1.0
1.0
1.0
1.0
1.0
1.0
1.0


3972.0
0.0
0.0
0.0
0.0
0.0
0.0
0.0
0.0
0.0
0.0
0.0
0.0
0.0
0.0
0.0
0.0
0.0
0.0
0.0
0.0
0.0
0.0
0.0
0.0
1177.0
2760.0
0.0
0.0
35.0

0.98
0.98
0.98
0.98
0.98
0.98
0.98
0.98
0.98
0.98
0.98
0.98
0.98
0.98
0.98
0.98
0.98
0.98
0.98
0.98
0.98
0.98
0.98
0.98
0.98
0.98
0.98
0.98
0.98
0.98


305.0
0.0
0.0
0.0
0.0
0.0
0.0
0.0
0.0
0.0
0.0
0.0
0.0
0.0
0.0
0.0
0.0
0.0
0.0
0.0
0.0
0.0
0.0
0.0
0.0
200.0
0.0
57.0
0.0
48.0

0.99
0.99
0.99
0.99
0.99
0.99
0.99
0.99
0.99
0.99
0.99
0.99
0.99
0.99
0.99
0.99
0.99
0.99
0.99
0.99
0.99
0.99
0.99
0.99
0.99
0.99
0.99
0.99
0.99
0.99


4213850.0
0.0
0.0
0.0
0.0
0.0
0.0
0.0
0.0
0.0
0.0
0.0
0.0
0.0
0.0
3081868.0
1131981.0
0.0
0.0
0.0
0.0
0.0
0.0
0.0
0.0
0.0
1.0
0.0
0.0
0.0


4213850.0
0.0
0.0
0.0
0.0
0.0
0.0
0.0
0.0
0.0
0.0
0.0
0.0
0.0
0.0
3081868.0
1131981.0
0.0
0.0
0.0
0.0
0.0
0.0
0.0
0.0
0.0
1.0
0.0
0.0
0.0

0.99
0.99
0.99
0.99
0.99
0.99
0.99
0.99
0.99
0.99
0.99
0.99
0.99
0.99
0.99
0.99
0.99
0.99
0.99
0.99
0.99
0.99
0.99
0.99
0.99
0.99
0.99
0.99
0.99
0.99


24137.0
0.0
0.0
0.0
0.0
0.0
0.0
0.0
0.0
0.0
0.0
0.0
0.0
0.0
0.0
0.0
0.0
0.0
0.0
0.0
0.0
0.0
0.0
0.0
24137.0
0.0
0.0
0.0
0.0
0.0


24137.0
0.0
0.0
0.0
0.0
0.0
0.0
0.0
0.0
0.0
0.0
0.0
0.0
0.0
0.0
0.0
0.0
0.0
0.0
0.0
0.0
0.0
0.0
0.0
24137.0
0.0
0.0
0.0
0.0
0.0

1.0
1.0
1.0
1.0
1.0
1.0
1.0
1.0
1.0
1.0
1.0
1.0
1.0
1.0
1.0
1.0
1.0
1.0
1.0
1.0
1.0
1.0
1.0
1.0
1.0
1.0
1.0
1.0
1.0
1.0


13986.0
0.0
0.0
0.0
0.0
0.0
0.0
0.0
0.0
0.0
0.0
0.0
0.0
0.0
0.0
0.0
0.0
0.0
0.0
0.0
0.0
0.0
0.0
0.0
0.0
13986.0
0.0
0.0
0.0
0.0


13986.0
0.0
0.0
0.0
0.0
0.0
0.0
0.0
0.0
0.0
0.0
0.0
0.0
0.0
0.0
0.0
0.0
0.0
0.0
0.0
0.0
0.0
0.0
0.0
0.0
13986.0
0.0
0.0
0.0
0.0

1.0
1.0
1.0
1.0
1.0
1.0
1.0
1.0
1.0
1.0
1.0
1.0
1.0
1.0
1.0
1.0
1.0
1.0
1.0
1.0
1.0
1.0
1.0
1.0
1.0
1.0
1.0
1.0
1.0
1.0


2565604.0
2806.0
6.0
0.0
1838.0
228.0
719.0
29770.0
23608.0
4054.0
3847.0
2192.0
0.0
0.0
0.0
1.0
0.0
1.0
2.0
7955.0
141263.0
1.0
0.0
0.0
26011.0
781991.0
628844.0
249237.0
588027.0
73203.0


2502814.0
2794.0
0.0
0.0
1838.0
228.0
710.0
28201.0
23519.0
3941.0
3816.0
1802.0
0.0
0.0
0.0
1.0
0.0
1.0
2.0
0.0
141263.0
1.0
0.0
0.0
0.0
772510.0
628844.0
238420.0
588027.0
66896.0


2502814.0
2794.0
0.0
0.0
1838.0
228.0
710.0
28201.0
23519.0
3941.0
3816.0
1802.0
0.0
0.0
0.0
1.0
0.0
1.0
2.0
0.0
141263.0
1.0
0.0
0.0
0.0
772510.0
628844.0
238420.0
588027.0
66896.0

1.0
1.0
1.0
1.0
1.0
1.0
1.0
1.0
1.0
1.0
1.0
1.0
1.0
1.0
1.0
1.0
1.0
1.0
1.0
1.0
1.0
1.0
1.0
1.0
1.0
1.0
1.0
1.0
1.0
1.0


1772.0
1.0
0.0
0.0
0.0
0.0
0.0
1547.0
0.0
96.0
0.0
128.0
0.0
0.0
0.0
0.0
0.0
0.0
0.0
0.0
0.0
0.0
0.0
0.0
0.0
0.0
0.0
0.0
0.0
0.0

1.0
1.0
1.0
1.0
1.0
1.0
1.0
1.0
1.0
1.0
1.0
1.0
1.0
1.0
1.0
1.0
1.0
1.0
1.0
1.0
1.0
1.0
1.0
1.0
1.0
1.0
1.0
1.0
1.0
1.0


75.0
1.0
0.0
0.0
0.0
0.0
0.0
0.0
0.0
74.0
0.0
0.0
0.0
0.0
0.0
0.0
0.0
0.0
0.0
0.0
0.0
0.0
0.0
0.0
0.0
0.0
0.0
0.0
0.0
0.0

0.93
0.93
0.93
0.93
0.93
0.93
0.93
0.93
0.93
0.93
0.93
0.93
0.93
0.93
0.93
0.93
0.93
0.93
0.93
0.93
0.93
0.93
0.93
0.93
0.93
0.93
0.93
0.93
0.93
0.93


432.0
11.0
0.0
0.0
0.0
0.0
0.0
22.0
89.0
17.0
31.0
262.0
0.0
0.0
0.0
0.0
0.0
0.0
0.0
0.0
0.0
0.0
0.0
0.0
0.0
0.0
0.0
0.0
0.0
0.0

1.0
1.0
1.0
1.0
1.0
1.0
1.0
1.0
1.0
1.0
1.0
1.0
1.0
1.0
1.0
1.0
1.0
1.0
1.0
1.0
1.0
1.0
1.0
1.0
1.0
1.0
1.0
1.0
1.0
1.0


42.0
0.0
0.0
0.0
0.0
0.0
0.0
0.0
23.0
13.0
6.0
0.0
0.0
0.0
0.0
0.0
0.0
0.0
0.0
0.0
0.0
0.0
0.0
0.0
0.0
0.0
0.0
0.0
0.0
0.0

1.0
1.0
1.0
1.0
1.0
1.0
1.0
1.0
1.0
1.0
1.0
1.0
1.0
1.0
1.0
1.0
1.0
1.0
1.0
1.0
1.0
1.0
1.0
1.0
1.0
1.0
1.0
1.0
1.0
1.0


12.0
0.0
0.0
0.0
0.0
0.0
0.0
0.0
0.0
4.0
1.0
7.0
0.0
0.0
0.0
0.0
0.0
0.0
0.0
0.0
0.0
0.0
0.0
0.0
0.0
0.0
0.0
0.0
0.0
0.0

1.0
1.0
1.0
1.0
1.0
1.0
1.0
1.0
1.0
1.0
1.0
1.0
1.0
1.0
1.0
1.0
1.0
1.0
1.0
1.0
1.0
1.0
1.0
1.0
1.0
1.0
1.0
1.0
1.0
1.0


10.0
0.0
0.0
0.0
0.0
0.0
0.0
0.0
10.0
0.0
0.0
0.0
0.0
0.0
0.0
0.0
0.0
0.0
0.0
0.0
0.0
0.0
0.0
0.0
0.0
0.0
0.0
0.0
0.0
0.0

1.0
1.0
1.0
1.0
1.0
1.0
1.0
1.0
1.0
1.0
1.0
1.0
1.0
1.0
1.0
1.0
1.0
1.0
1.0
1.0
1.0
1.0
1.0
1.0
1.0
1.0
1.0
1.0
1.0
1.0


52631.0
0.0
6.0
0.0
0.0
0.0
9.0
0.0
0.0
0.0
0.0
0.0
0.0
0.0
0.0
0.0
0.0
0.0
0.0
0.0
0.0
0.0
0.0
0.0
26011.0
9481.0
0.0
10817.0
0.0
6307.0

1.0
1.0
1.0
1.0
1.0
1.0
1.0
1.0
1.0
1.0
1.0
1.0
1.0
1.0
1.0
1.0
1.0
1.0
1.0
1.0
1.0
1.0
1.0
1.0
1.0
1.0
1.0
1.0
1.0
1.0


2998.0
0.0
6.0
0.0
0.0
0.0
9.0
0.0
0.0
0.0
0.0
0.0
0.0
0.0
0.0
0.0
0.0
0.0
0.0
0.0
0.0
0.0
0.0
0.0
0.0
360.0
0.0
2623.0
0.0
0.0

1.0
1.0
1.0
1.0
1.0
1.0
1.0
1.0
1.0
1.0
1.0
1.0
1.0
1.0
1.0
1.0
1.0
1.0
1.0
1.0
1.0
1.0
1.0
1.0
1.0
1.0
1.0
1.0
1.0
1.0


7955.0
0.0
0.0
0.0
0.0
0.0
0.0
0.0
0.0
0.0
0.0
0.0
0.0
0.0
0.0
0.0
0.0
0.0
0.0
7955.0
0.0
0.0
0.0
0.0
0.0
0.0
0.0
0.0
0.0
0.0

1.0
1.0
1.0
1.0
1.0
1.0
1.0
1.0
1.0
1.0
1.0
1.0
1.0
1.0
1.0
1.0
1.0
1.0
1.0
1.0
1.0
1.0
1.0
1.0
1.0
1.0
1.0
1.0
1.0
1.0


1714323.0
515.0
4086.0
32802.0
3455.0
1542.0
1017.0
318.0
433.0
2716.0
2018.0
818.0
2849.0
24671.0
34996.0
1.0
7018.0
6215.0
11909.0
5181.0
31095.0
53549.0
73754.0
2849.0
590742.0
148580.0
94074.0
100327.0
381531.0
95262.0

1.0
1.0
1.0
1.0
1.0
1.0
1.0
1.0
1.0
1.0
1.0
1.0
1.0
1.0
1.0
1.0
1.0
1.0
1.0
1.0
1.0
1.0
1.0
1.0
1.0
1.0
1.0
1.0
1.0
1.0


41712.0
33.0
3767.0
32047.0
3149.0
467.0
0.0
0.0
12.0
9.0
113.0
53.0
0.0
0.0
0.0
0.0
0.0
0.0
0.0
0.0
0.0
0.0
2.0
0.0
1974.0
10.0
0.0
46.0
0.0
30.0

0.89
0.89
0.89
0.89
0.89
0.89
0.89
0.89
0.89
0.89
0.89
0.89
0.89
0.89
0.89
0.89
0.89
0.89
0.89
0.89
0.89
0.89
0.89
0.89
0.89
0.89
0.89
0.89
0.89
0.89


39650.0
33.0
3767.0
32047.0
3149.0
467.0
0.0
0.0
12.0
9.0
113.0
53.0
0.0
0.0
0.0
0.0
0.0
0.0
0.0
0.0
0.0
0.0
0.0
0.0
0.0
0.0
0.0
0.0
0.0
0.0

1.0
1.0
1.0
1.0
1.0
1.0
1.0
1.0
1.0
1.0
1.0
1.0
1.0
1.0
1.0
1.0
1.0
1.0
1.0
1.0
1.0
1.0
1.0
1.0
1.0
1.0
1.0
1.0
1.0
1.0


10.0
0.0
0.0
0.0
0.0
0.0
0.0
0.0
0.0
0.0
0.0
0.0
0.0
0.0
0.0
0.0
0.0
0.0
0.0
0.0
0.0
0.0
0.0
0.0
0.0
10.0
0.0
0.0
0.0
0.0

1.0
1.0
1.0
1.0
1.0
1.0
1.0
1.0
1.0
1.0
1.0
1.0
1.0
1.0
1.0
1.0
1.0
1.0
1.0
1.0
1.0
1.0
1.0
1.0
1.0
1.0
1.0
1.0
1.0
1.0


1309896.0
286.0
136.0
257.0
268.0
198.0
462.0
318.0
384.0
1543.0
696.0
162.0
2849.0
0.0
30707.0
1.0
5291.0
6188.0
11909.0
794.0
25817.0
6105.0
61789.0
2849.0
435074.0
101633.0
69071.0
81655.0
375331.0
88123.0


1309896.0
286.0
136.0
257.0
268.0
198.0
462.0
318.0
384.0
1543.0
696.0
162.0
2849.0
0.0
30707.0
1.0
5291.0
6188.0
11909.0
794.0
25817.0
6105.0
61789.0
2849.0
435074.0
101633.0
69071.0
81655.0
375331.0
88123.0

1.0
1.0
1.0
1.0
1.0
1.0
1.0
1.0
1.0
1.0
1.0
1.0
1.0
1.0
1.0
1.0
1.0
1.0
1.0
1.0
1.0
1.0
1.0
1.0
1.0
1.0
1.0
1.0
1.0
1.0


70424.0
0.0
141.0
288.0
0.0
302.0
314.0
0.0
0.0
1090.0
967.0
230.0
0.0
0.0
7.0
0.0
0.0
0.0
0.0
0.0
5268.0
27479.0
11177.0
0.0
3007.0
15495.0
0.0
4659.0
0.0
0.0


65637.0
0.0
141.0
288.0
0.0
302.0
314.0
0.0
0.0
0.0
514.0
0.0
0.0
0.0
0.0
0.0
0.0
0.0
0.0
0.0
5268.0
27479.0
11177.0
0.0
0.0
15495.0
0.0
4659.0
0.0
0.0

1.0
1.0
1.0
1.0
1.0
1.0
1.0
1.0
1.0
1.0
1.0
1.0
1.0
1.0
1.0
1.0
1.0
1.0
1.0
1.0
1.0
1.0
1.0
1.0
1.0
1.0
1.0
1.0
1.0
1.0


4733.0
0.0
0.0
0.0
0.0
0.0
0.0
0.0
0.0
1090.0
453.0
176.0
0.0
0.0
7.0
0.0
0.0
0.0
0.0
0.0
0.0
0.0
0.0
0.0
3007.0
0.0
0.0
0.0
0.0
0.0

1.0
1.0
1.0
1.0
1.0
1.0
1.0
1.0
1.0
1.0
1.0
1.0
1.0
1.0
1.0
1.0
1.0
1.0
1.0
1.0
1.0
1.0
1.0
1.0
1.0
1.0
1.0
1.0
1.0
1.0


54.0
0.0
0.0
0.0
0.0
0.0
0.0
0.0
0.0
0.0
0.0
54.0
0.0
0.0
0.0
0.0
0.0
0.0
0.0
0.0
0.0
0.0
0.0
0.0
0.0
0.0
0.0
0.0
0.0
0.0

1.0
1.0
1.0
1.0
1.0
1.0
1.0
1.0
1.0
1.0
1.0
1.0
1.0
1.0
1.0
1.0
1.0
1.0
1.0
1.0
1.0
1.0
1.0
1.0
1.0
1.0
1.0
1.0
1.0
1.0


45183.0
0.0
0.0
15.0
17.0
78.0
198.0
0.0
0.0
0.0
3.0
7.0
0.0
0.0
0.0
0.0
0.0
0.0
0.0
1842.0
0.0
19965.0
0.0
0.0
1653.0
6120.0
4403.0
6166.0
2678.0
2038.0

1.0
1.0
1.0
1.0
1.0
1.0
1.0
1.0
1.0
1.0
1.0
1.0
1.0
1.0
1.0
1.0
1.0
1.0
1.0
1.0
1.0
1.0
1.0
1.0
1.0
1.0
1.0
1.0
1.0
1.0


3109.0
32.0
27.0
0.0
0.0
32.0
0.0
0.0
37.0
44.0
133.0
46.0
0.0
0.0
0.0
0.0
0.0
0.0
0.0
0.0
0.0
0.0
0.0
0.0
2559.0
0.0
0.0
0.0
199.0
0.0

1.0
1.0
1.0
1.0
1.0
1.0
1.0
1.0
1.0
1.0
1.0
1.0
1.0
1.0
1.0
1.0
1.0
1.0
1.0
1.0
1.0
1.0
1.0
1.0
1.0
1.0
1.0
1.0
1.0
1.0


379.0
0.0
0.0
0.0
0.0
0.0
0.0
0.0
0.0
0.0
0.0
0.0
0.0
0.0
0.0
0.0
0.0
0.0
0.0
0.0
0.0
0.0
0.0
0.0
379.0
0.0
0.0
0.0
0.0
0.0

0.93
0.93
0.93
0.93
0.93
0.93
0.93
0.93
0.93
0.93
0.93
0.93
0.93
0.93
0.93
0.93
0.93
0.93
0.93
0.93
0.93
0.93
0.93
0.93
0.93
0.93
0.93
0.93
0.93
0.93


44931.0
138.0
0.0
3.0
2.0
6.0
13.0
0.0
0.0
30.0
106.0
269.0
0.0
24671.0
4200.0
0.0
0.0
0.0
0.0
2424.0
0.0
0.0
786.0
0.0
9614.0
0.0
1985.0
0.0
684.0
0.0

1.0
1.0
1.0
1.0
1.0
1.0
1.0
1.0
1.0
1.0
1.0
1.0
1.0
1.0
1.0
1.0
1.0
1.0
1.0
1.0
1.0
1.0
1.0
1.0
1.0
1.0
1.0
1.0
1.0
1.0


4032.0
25.0
0.0
0.0
0.0
0.0
0.0
0.0
0.0
27.0
73.0
17.0
0.0
3292.0
251.0
0.0
0.0
0.0
0.0
292.0
0.0
0.0
55.0
0.0
0.0
0.0
0.0
0.0
0.0
0.0

1.0
1.0
1.0
1.0
1.0
1.0
1.0
1.0
1.0
1.0
1.0
1.0
1.0
1.0
1.0
1.0
1.0
1.0
1.0
1.0
1.0
1.0
1.0
1.0
1.0
1.0
1.0
1.0
1.0
1.0


12283.0
0.0
0.0
0.0
0.0
0.0
0.0
0.0
0.0
0.0
0.0
0.0
0.0
0.0
0.0
0.0
0.0
0.0
0.0
0.0
0.0
0.0
0.0
0.0
9614.0
0.0
1985.0
0.0
684.0
0.0

1.0
1.0
1.0
1.0
1.0
1.0
1.0
1.0
1.0
1.0
1.0
1.0
1.0
1.0
1.0
1.0
1.0
1.0
1.0
1.0
1.0
1.0
1.0
1.0
1.0
1.0
1.0
1.0
1.0
1.0


358.0
26.0
0.0
1.0
0.0
160.0
5.0
0.0
0.0
0.0
0.0
0.0
0.0
0.0
82.0
0.0
0.0
23.0
0.0
0.0
0.0
0.0
0.0
0.0
23.0
0.0
0.0
3.0
35.0
0.0

0.8
0.8
0.8
0.8
0.8
0.8
0.8
0.8
0.8
0.8
0.8
0.8
0.8
0.8
0.8
0.8
0.8
0.8
0.8
0.8
0.8
0.8
0.8
0.8
0.8
0.8
0.8
0.8
0.8
0.8


174.0
2.0
0.0
1.0
0.0
0.0
5.0
0.0
0.0
0.0
0.0
0.0
0.0
0.0
82.0
0.0
0.0
23.0
0.0
0.0
0.0
0.0
0.0
0.0
23.0
0.0
0.0
3.0
35.0
0.0

0.97
0.97
0.97
0.97
0.97
0.97
0.97
0.97
0.97
0.97
0.97
0.97
0.97
0.97
0.97
0.97
0.97
0.97
0.97
0.97
0.97
0.97
0.97
0.97
0.97
0.97
0.97
0.97
0.97
0.97


79.0
0.0
13.0
15.0
0.0
0.0
0.0
0.0
0.0
0.0
0.0
51.0
0.0
0.0
0.0
0.0
0.0
0.0
0.0
0.0
0.0
0.0
0.0
0.0
0.0
0.0
0.0
0.0
0.0
0.0

0.97
0.97
0.97
0.97
0.97
0.97
0.97
0.97
0.97
0.97
0.97
0.97
0.97
0.97
0.97
0.97
0.97
0.97
0.97
0.97
0.97
0.97
0.97
0.97
0.97
0.97
0.97
0.97
0.97
0.97


4660.0
0.0
0.0
47.0
0.0
0.0
0.0
0.0
0.0
0.0
0.0
0.0
0.0
0.0
0.0
0.0
0.0
0.0
0.0
0.0
0.0
0.0
0.0
0.0
0.0
0.0
0.0
0.0
1326.0
3287.0


4660.0
0.0
0.0
47.0
0.0
0.0
0.0
0.0
0.0
0.0
0.0
0.0
0.0
0.0
0.0
0.0
0.0
0.0
0.0
0.0
0.0
0.0
0.0
0.0
0.0
0.0
0.0
0.0
1326.0
3287.0

1.0
1.0
1.0
1.0
1.0
1.0
1.0
1.0
1.0
1.0
1.0
1.0
1.0
1.0
1.0
1.0
1.0
1.0
1.0
1.0
1.0
1.0
1.0
1.0
1.0
1.0
1.0
1.0
1.0
1.0


8739.0
0.0
1.0
1.0
9.0
5.0
25.0
0.0
0.0
0.0
0.0
0.0
0.0
0.0
0.0
0.0
1727.0
0.0
0.0
0.0
0.0
0.0
0.0
0.0
79.0
2122.0
3140.0
343.0
530.0
757.0


8739.0
0.0
1.0
1.0
9.0
5.0
25.0
0.0
0.0
0.0
0.0
0.0
0.0
0.0
0.0
0.0
1727.0
0.0
0.0
0.0
0.0
0.0
0.0
0.0
79.0
2122.0
3140.0
343.0
530.0
757.0

1.0
1.0
1.0
1.0
1.0
1.0
1.0
1.0
1.0
1.0
1.0
1.0
1.0
1.0
1.0
1.0
1.0
1.0
1.0
1.0
1.0
1.0
1.0
1.0
1.0
1.0
1.0
1.0
1.0
1.0


35.0
0.0
0.0
2.0
9.0
20.0
0.0
0.0
0.0
0.0
0.0
0.0
0.0
0.0
0.0
0.0
0.0
4.0
0.0
0.0
0.0
0.0
0.0
0.0
0.0
0.0
0.0
0.0
0.0
0.0

1.0
1.0
1.0
1.0
1.0
1.0
1.0
1.0
1.0
1.0
1.0
1.0
1.0
1.0
1.0
1.0
1.0
1.0
1.0
1.0
1.0
1.0
1.0
1.0
1.0
1.0
1.0
1.0
1.0
1.0


14694.0
0.0
0.0
0.0
0.0
0.0
0.0
0.0
0.0
0.0
0.0
0.0
0.0
0.0
0.0
0.0
0.0
0.0
0.0
0.0
0.0
0.0
0.0
0.0
0.0
4337.0
9791.0
566.0
0.0
0.0


14694.0
0.0
0.0
0.0
0.0
0.0
0.0
0.0
0.0
0.0
0.0
0.0
0.0
0.0
0.0
0.0
0.0
0.0
0.0
0.0
0.0
0.0
0.0
0.0
0.0
4337.0
9791.0
566.0
0.0
0.0

0.95
0.95
0.95
0.95
0.95
0.95
0.95
0.95
0.95
0.95
0.95
0.95
0.95
0.95
0.95
0.95
0.95
0.95
0.95
0.95
0.95
0.95
0.95
0.95
0.95
0.95
0.95
0.95
0.95
0.95


25383.0
0.0
0.0
0.0
0.0
0.0
0.0
0.0
0.0
0.0
0.0
0.0
0.0
0.0
0.0
0.0
0.0
0.0
0.0
0.0
0.0
0.0
0.0
0.0
0.0
18863.0
5684.0
88.0
748.0
0.0

0.95
0.95
0.95
0.95
0.95
0.95
0.95
0.95
0.95
0.95
0.95
0.95
0.95
0.95
0.95
0.95
0.95
0.95
0.95
0.95
0.95
0.95
0.95
0.95
0.95
0.95
0.95
0.95
0.95
0.95


2791908.0
2011.0
551.0
260.0
122.0
144.0
2.0
7200.0
4251.0
7150.0
8495.0
34560.0
32334.0
0.0
1942.0
32519.0
52413.0
627197.0
274045.0
461328.0
111112.0
508831.0
415540.0
32334.0
167592.0
0.0
1.0
1180.0
5749.0
3045.0

1.0
1.0
1.0
1.0
1.0
1.0
1.0
1.0
1.0
1.0
1.0
1.0
1.0
1.0
1.0
1.0
1.0
1.0
1.0
1.0
1.0
1.0
1.0
1.0
1.0
1.0
1.0
1.0
1.0
1.0


34032.0
521.0
30.0
0.0
0.0
0.0
0.0
0.0
441.0
689.0
3379.0
28972.0
0.0
0.0
0.0
0.0
0.0
0.0
0.0
0.0
0.0
0.0
0.0
0.0
0.0
0.0
0.0
0.0
0.0
0.0


34032.0
521.0
30.0
0.0
0.0
0.0
0.0
0.0
441.0
689.0
3379.0
28972.0
0.0
0.0
0.0
0.0
0.0
0.0
0.0
0.0
0.0
0.0
0.0
0.0
0.0
0.0
0.0
0.0
0.0
0.0

1.0
1.0
1.0
1.0
1.0
1.0
1.0
1.0
1.0
1.0
1.0
1.0
1.0
1.0
1.0
1.0
1.0
1.0
1.0
1.0
1.0
1.0
1.0
1.0
1.0
1.0
1.0
1.0
1.0
1.0


6136.0
873.0
0.0
0.0
0.0
0.0
0.0
0.0
0.0
2619.0
1348.0
1296.0
0.0
0.0
0.0
0.0
0.0
0.0
0.0
0.0
0.0
0.0
0.0
0.0
0.0
0.0
0.0
0.0
0.0
0.0


6136.0
873.0
0.0
0.0
0.0
0.0
0.0
0.0
0.0
2619.0
1348.0
1296.0
0.0
0.0
0.0
0.0
0.0
0.0
0.0
0.0
0.0
0.0
0.0
0.0
0.0
0.0
0.0
0.0
0.0
0.0

1.0
1.0
1.0
1.0
1.0
1.0
1.0
1.0
1.0
1.0
1.0
1.0
1.0
1.0
1.0
1.0
1.0
1.0
1.0
1.0
1.0
1.0
1.0
1.0
1.0
1.0
1.0
1.0
1.0
1.0


13202.0
179.0
19.0
14.0
0.0
0.0
0.0
5921.0
2092.0
425.0
1389.0
856.0
0.0
0.0
0.0
0.0
0.0
0.0
0.0
0.0
0.0
0.0
0.0
0.0
2307.0
0.0
0.0
0.0
0.0
0.0


9525.0
101.0
0.0
0.0
0.0
0.0
0.0
5921.0
2092.0
0.0
1032.0
379.0
0.0
0.0
0.0
0.0
0.0
0.0
0.0
0.0
0.0
0.0
0.0
0.0
0.0
0.0
0.0
0.0
0.0
0.0

1.0
1.0
1.0
1.0
1.0
1.0
1.0
1.0
1.0
1.0
1.0
1.0
1.0
1.0
1.0
1.0
1.0
1.0
1.0
1.0
1.0
1.0
1.0
1.0
1.0
1.0
1.0
1.0
1.0
1.0


3677.0
78.0
19.0
14.0
0.0
0.0
0.0
0.0
0.0
425.0
357.0
477.0
0.0
0.0
0.0
0.0
0.0
0.0
0.0
0.0
0.0
0.0
0.0
0.0
2307.0
0.0
0.0
0.0
0.0
0.0

1.0
1.0
1.0
1.0
1.0
1.0
1.0
1.0
1.0
1.0
1.0
1.0
1.0
1.0
1.0
1.0
1.0
1.0
1.0
1.0
1.0
1.0
1.0
1.0
1.0
1.0
1.0
1.0
1.0
1.0


3736.0
135.0
44.0
11.0
0.0
0.0
0.0
0.0
257.0
1183.0
826.0
1280.0
0.0
0.0
0.0
0.0
0.0
0.0
0.0
0.0
0.0
0.0
0.0
0.0
0.0
0.0
0.0
0.0
0.0
0.0

1.0
1.0
1.0
1.0
1.0
1.0
1.0
1.0
1.0
1.0
1.0
1.0
1.0
1.0
1.0
1.0
1.0
1.0
1.0
1.0
1.0
1.0
1.0
1.0
1.0
1.0
1.0
1.0
1.0
1.0


90253.0
0.0
0.0
0.0
0.0
0.0
0.0
0.0
171.0
527.0
0.0
366.0
0.0
0.0
0.0
0.0
0.0
69473.0
130.0
8012.0
11574.0
0.0
0.0
0.0
0.0
0.0
0.0
0.0
0.0
0.0

0.9
0.9
0.9
0.9
0.9
0.9
0.9
0.9
0.9
0.9
0.9
0.9
0.9
0.9
0.9
0.9
0.9
0.9
0.9
0.9
0.9
0.9
0.9
0.9
0.9
0.9
0.9
0.9
0.9
0.9


88938.0
0.0
0.0
0.0
0.0
0.0
0.0
0.0
171.0
527.0
0.0
366.0
0.0
0.0
0.0
0.0
0.0
68840.0
121.0
7660.0
11253.0
0.0
0.0
0.0
0.0
0.0
0.0
0.0
0.0
0.0

1.0
1.0
1.0
1.0
1.0
1.0
1.0
1.0
1.0
1.0
1.0
1.0
1.0
1.0
1.0
1.0
1.0
1.0
1.0
1.0
1.0
1.0
1.0
1.0
1.0
1.0
1.0
1.0
1.0
1.0


2727.0
0.0
17.0
0.0
0.0
0.0
0.0
0.0
0.0
340.0
0.0
364.0
0.0
0.0
0.0
0.0
0.0
927.0
0.0
0.0
1079.0
0.0
0.0
0.0
0.0
0.0
0.0
0.0
0.0
0.0


2727.0
0.0
17.0
0.0
0.0
0.0
0.0
0.0
0.0
340.0
0.0
364.0
0.0
0.0
0.0
0.0
0.0
927.0
0.0
0.0
1079.0
0.0
0.0
0.0
0.0
0.0
0.0
0.0
0.0
0.0

1.0
1.0
1.0
1.0
1.0
1.0
1.0
1.0
1.0
1.0
1.0
1.0
1.0
1.0
1.0
1.0
1.0
1.0
1.0
1.0
1.0
1.0
1.0
1.0
1.0
1.0
1.0
1.0
1.0
1.0


577.0
0.0
0.0
0.0
0.0
0.0
0.0
0.0
153.0
0.0
269.0
155.0
0.0
0.0
0.0
0.0
0.0
0.0
0.0
0.0
0.0
0.0
0.0
0.0
0.0
0.0
0.0
0.0
0.0
0.0


577.0
0.0
0.0
0.0
0.0
0.0
0.0
0.0
153.0
0.0
269.0
155.0
0.0
0.0
0.0
0.0
0.0
0.0
0.0
0.0
0.0
0.0
0.0
0.0
0.0
0.0
0.0
0.0
0.0
0.0

1.0
1.0
1.0
1.0
1.0
1.0
1.0
1.0
1.0
1.0
1.0
1.0
1.0
1.0
1.0
1.0
1.0
1.0
1.0
1.0
1.0
1.0
1.0
1.0
1.0
1.0
1.0
1.0
1.0
1.0


93705.0
29.0
59.0
125.0
0.0
70.0
0.0
0.0
0.0
0.0
0.0
291.0
0.0
0.0
0.0
0.0
9262.0
3309.0
59447.0
0.0
3189.0
0.0
7455.0
0.0
10469.0
0.0
0.0
0.0
0.0
0.0


93705.0
29.0
59.0
125.0
0.0
70.0
0.0
0.0
0.0
0.0
0.0
291.0
0.0
0.0
0.0
0.0
9262.0
3309.0
59447.0
0.0
3189.0
0.0
7455.0
0.0
10469.0
0.0
0.0
0.0
0.0
0.0

0.97
0.97
0.97
0.97
0.97
0.97
0.97
0.97
0.97
0.97
0.97
0.97
0.97
0.97
0.97
0.97
0.97
0.97
0.97
0.97
0.97
0.97
0.97
0.97
0.97
0.97
0.97
0.97
0.97
0.97


599.0
29.0
2.0
0.0
0.0
0.0
0.0
0.0
0.0
128.0
139.0
64.0
0.0
0.0
0.0
0.0
0.0
0.0
237.0
0.0
0.0
0.0
0.0
0.0
0.0
0.0
0.0
0.0
0.0
0.0


599.0
29.0
2.0
0.0
0.0
0.0
0.0
0.0
0.0
128.0
139.0
64.0
0.0
0.0
0.0
0.0
0.0
0.0
237.0
0.0
0.0
0.0
0.0
0.0
0.0
0.0
0.0
0.0
0.0
0.0

1.0
1.0
1.0
1.0
1.0
1.0
1.0
1.0
1.0
1.0
1.0
1.0
1.0
1.0
1.0
1.0
1.0
1.0
1.0
1.0
1.0
1.0
1.0
1.0
1.0
1.0
1.0
1.0
1.0
1.0


343.0
0.0
5.0
0.0
0.0
0.0
0.0
78.0
46.0
76.0
126.0
12.0
0.0
0.0
0.0
0.0
0.0
0.0
0.0
0.0
0.0
0.0
0.0
0.0
0.0
0.0
0.0
0.0
0.0
0.0

1.0
1.0
1.0
1.0
1.0
1.0
1.0
1.0
1.0
1.0
1.0
1.0
1.0
1.0
1.0
1.0
1.0
1.0
1.0
1.0
1.0
1.0
1.0
1.0
1.0
1.0
1.0
1.0
1.0
1.0


133.0
17.0
0.0
0.0
0.0
0.0
0.0
0.0
0.0
116.0
0.0
0.0
0.0
0.0
0.0
0.0
0.0
0.0
0.0
0.0
0.0
0.0
0.0
0.0
0.0
0.0
0.0
0.0
0.0
0.0


133.0
17.0
0.0
0.0
0.0
0.0
0.0
0.0
0.0
116.0
0.0
0.0
0.0
0.0
0.0
0.0
0.0
0.0
0.0
0.0
0.0
0.0
0.0
0.0
0.0
0.0
0.0
0.0
0.0
0.0

1.0
1.0
1.0
1.0
1.0
1.0
1.0
1.0
1.0
1.0
1.0
1.0
1.0
1.0
1.0
1.0
1.0
1.0
1.0
1.0
1.0
1.0
1.0
1.0
1.0
1.0
1.0
1.0
1.0
1.0


511390.0
57.0
0.0
0.0
0.0
0.0
0.0
0.0
0.0
0.0
0.0
53.0
0.0
0.0
0.0
0.0
0.0
177394.0
102664.0
191934.0
39288.0
0.0
0.0
0.0
0.0
0.0
0.0
0.0
0.0
0.0


511390.0
57.0
0.0
0.0
0.0
0.0
0.0
0.0
0.0
0.0
0.0
53.0
0.0
0.0
0.0
0.0
0.0
177394.0
102664.0
191934.0
39288.0
0.0
0.0
0.0
0.0
0.0
0.0
0.0
0.0
0.0

1.0
1.0
1.0
1.0
1.0
1.0
1.0
1.0
1.0
1.0
1.0
1.0
1.0
1.0
1.0
1.0
1.0
1.0
1.0
1.0
1.0
1.0
1.0
1.0
1.0
1.0
1.0
1.0
1.0
1.0


188.0
6.0
13.0
2.0
0.0
0.0
0.0
0.0
17.0
68.0
27.0
55.0
0.0
0.0
0.0
0.0
0.0
0.0
0.0
0.0
0.0
0.0
0.0
0.0
0.0
0.0
0.0
0.0
0.0
0.0


188.0
6.0
13.0
2.0
0.0
0.0
0.0
0.0
17.0
68.0
27.0
55.0
0.0
0.0
0.0
0.0
0.0
0.0
0.0
0.0
0.0
0.0
0.0
0.0
0.0
0.0
0.0
0.0
0.0
0.0

0.86
0.86
0.86
0.86
0.86
0.86
0.86
0.86
0.86
0.86
0.86
0.86
0.86
0.86
0.86
0.86
0.86
0.86
0.86
0.86
0.86
0.86
0.86
0.86
0.86
0.86
0.86
0.86
0.86
0.86


446713.0
9.0
37.0
0.0
0.0
0.0
0.0
0.0
0.0
0.0
0.0
138.0
0.0
0.0
0.0
0.0
0.0
143644.0
46049.0
223952.0
10096.0
17716.0
2.0
0.0
5068.0
0.0
1.0
0.0
1.0
0.0

1.0
1.0
1.0
1.0
1.0
1.0
1.0
1.0
1.0
1.0
1.0
1.0
1.0
1.0
1.0
1.0
1.0
1.0
1.0
1.0
1.0
1.0
1.0
1.0
1.0
1.0
1.0
1.0
1.0
1.0


9.0
9.0
0.0
0.0
0.0
0.0
0.0
0.0
0.0
0.0
0.0
0.0
0.0
0.0
0.0
0.0
0.0
0.0
0.0
0.0
0.0
0.0
0.0
0.0
0.0
0.0
0.0
0.0
0.0
0.0

1.0
1.0
1.0
1.0
1.0
1.0
1.0
1.0
1.0
1.0
1.0
1.0
1.0
1.0
1.0
1.0
1.0
1.0
1.0
1.0
1.0
1.0
1.0
1.0
1.0
1.0
1.0
1.0
1.0
1.0


1114.0
4.0
0.0
11.0
38.0
0.0
0.0
0.0
0.0
0.0
36.0
0.0
0.0
0.0
0.0
0.0
0.0
1025.0
0.0
0.0
0.0
0.0
0.0
0.0
0.0
0.0
0.0
0.0
0.0
0.0


1114.0
4.0
0.0
11.0
38.0
0.0
0.0
0.0
0.0
0.0
36.0
0.0
0.0
0.0
0.0
0.0
0.0
1025.0
0.0
0.0
0.0
0.0
0.0
0.0
0.0
0.0
0.0
0.0
0.0
0.0

1.0
1.0
1.0
1.0
1.0
1.0
1.0
1.0
1.0
1.0
1.0
1.0
1.0
1.0
1.0
1.0
1.0
1.0
1.0
1.0
1.0
1.0
1.0
1.0
1.0
1.0
1.0
1.0
1.0
1.0


79.0
6.0
0.0
0.0
0.0
0.0
0.0
0.0
18.0
26.0
29.0
0.0
0.0
0.0
0.0
0.0
0.0
0.0
0.0
0.0
0.0
0.0
0.0
0.0
0.0
0.0
0.0
0.0
0.0
0.0


79.0
6.0
0.0
0.0
0.0
0.0
0.0
0.0
18.0
26.0
29.0
0.0
0.0
0.0
0.0
0.0
0.0
0.0
0.0
0.0
0.0
0.0
0.0
0.0
0.0
0.0
0.0
0.0
0.0
0.0

1.0
1.0
1.0
1.0
1.0
1.0
1.0
1.0
1.0
1.0
1.0
1.0
1.0
1.0
1.0
1.0
1.0
1.0
1.0
1.0
1.0
1.0
1.0
1.0
1.0
1.0
1.0
1.0
1.0
1.0


249553.0
2.0
0.0
1.0
12.0
0.0
0.0
0.0
0.0
0.0
0.0
48.0
0.0
0.0
0.0
22991.0
29976.0
105787.0
38426.0
0.0
0.0
0.0
0.0
0.0
52310.0
0.0
0.0
0.0
0.0
0.0


249553.0
2.0
0.0
1.0
12.0
0.0
0.0
0.0
0.0
0.0
0.0
48.0
0.0
0.0
0.0
22991.0
29976.0
105787.0
38426.0
0.0
0.0
0.0
0.0
0.0
52310.0
0.0
0.0
0.0
0.0
0.0

1.0
1.0
1.0
1.0
1.0
1.0
1.0
1.0
1.0
1.0
1.0
1.0
1.0
1.0
1.0
1.0
1.0
1.0
1.0
1.0
1.0
1.0
1.0
1.0
1.0
1.0
1.0
1.0
1.0
1.0


158922.0
7.0
1.0
1.0
0.0
0.0
2.0
0.0
0.0
20.0
30.0
14.0
28953.0
0.0
0.0
4056.0
10067.0
19644.0
26199.0
13561.0
13050.0
5712.0
0.0
28953.0
0.0
0.0
0.0
0.0
5748.0
2904.0

0.99
0.99
0.99
0.99
0.99
0.99
0.99
0.99
0.99
0.99
0.99
0.99
0.99
0.99
0.99
0.99
0.99
0.99
0.99
0.99
0.99
0.99
0.99
0.99
0.99
0.99
0.99
0.99
0.99
0.99


116724.0
7.0
1.0
1.0
0.0
0.0
2.0
0.0
0.0
8.0
30.0
14.0
14639.0
0.0
0.0
4056.0
10067.0
15500.0
26032.0
10147.0
13023.0
5712.0
0.0
14639.0
0.0
0.0
0.0
0.0
0.0
2846.0

1.0
1.0
1.0
1.0
1.0
1.0
1.0
1.0
1.0
1.0
1.0
1.0
1.0
1.0
1.0
1.0
1.0
1.0
1.0
1.0
1.0
1.0
1.0
1.0
1.0
1.0
1.0
1.0
1.0
1.0


31264.0
0.0
0.0
0.0
0.0
0.0
0.0
0.0
0.0
0.0
0.0
0.0
11780.0
0.0
0.0
0.0
0.0
4144.0
167.0
3308.0
27.0
0.0
0.0
11780.0
0.0
0.0
0.0
0.0
0.0
58.0

1.0
1.0
1.0
1.0
1.0
1.0
1.0
1.0
1.0
1.0
1.0
1.0
1.0
1.0
1.0
1.0
1.0
1.0
1.0
1.0
1.0
1.0
1.0
1.0
1.0
1.0
1.0
1.0
1.0
1.0


5068.0
0.0
0.0
0.0
0.0
0.0
0.0
0.0
0.0
0.0
0.0
0.0
2534.0
0.0
0.0
0.0
0.0
0.0
0.0
0.0
0.0
0.0
0.0
2534.0
0.0
0.0
0.0
0.0
0.0
0.0

1.0
1.0
1.0
1.0
1.0
1.0
1.0
1.0
1.0
1.0
1.0
1.0
1.0
1.0
1.0
1.0
1.0
1.0
1.0
1.0
1.0
1.0
1.0
1.0
1.0
1.0
1.0
1.0
1.0
1.0


106.0
0.0
0.0
0.0
0.0
0.0
0.0
0.0
0.0
0.0
0.0
0.0
0.0
0.0
0.0
0.0
0.0
0.0
0.0
106.0
0.0
0.0
0.0
0.0
0.0
0.0
0.0
0.0
0.0
0.0

0.97
0.97
0.97
0.97
0.97
0.97
0.97
0.97
0.97
0.97
0.97
0.97
0.97
0.97
0.97
0.97
0.97
0.97
0.97
0.97
0.97
0.97
0.97
0.97
0.97
0.97
0.97
0.97
0.97
0.97


62.0
6.0
0.0
0.0
0.0
0.0
0.0
0.0
0.0
18.0
38.0
0.0
0.0
0.0
0.0
0.0
0.0
0.0
0.0
0.0
0.0
0.0
0.0
0.0
0.0
0.0
0.0
0.0
0.0
0.0


62.0
6.0
0.0
0.0
0.0
0.0
0.0
0.0
0.0
18.0
38.0
0.0
0.0
0.0
0.0
0.0
0.0
0.0
0.0
0.0
0.0
0.0
0.0
0.0
0.0
0.0
0.0
0.0
0.0
0.0

1.0
1.0
1.0
1.0
1.0
1.0
1.0
1.0
1.0
1.0
1.0
1.0
1.0
1.0
1.0
1.0
1.0
1.0
1.0
1.0
1.0
1.0
1.0
1.0
1.0
1.0
1.0
1.0
1.0
1.0


54.0
0.0
0.0
0.0
0.0
0.0
0.0
0.0
0.0
0.0
54.0
0.0
0.0
0.0
0.0
0.0
0.0
0.0
0.0
0.0
0.0
0.0
0.0
0.0
0.0
0.0
0.0
0.0
0.0
0.0

1.0
1.0
1.0
1.0
1.0
1.0
1.0
1.0
1.0
1.0
1.0
1.0
1.0
1.0
1.0
1.0
1.0
1.0
1.0
1.0
1.0
1.0
1.0
1.0
1.0
1.0
1.0
1.0
1.0
1.0


39.0
0.0
0.0
0.0
0.0
39.0
0.0
0.0
0.0
0.0
0.0
0.0
0.0
0.0
0.0
0.0
0.0
0.0
0.0
0.0
0.0
0.0
0.0
0.0
0.0
0.0
0.0
0.0
0.0
0.0


39.0
0.0
0.0
0.0
0.0
39.0
0.0
0.0
0.0
0.0
0.0
0.0
0.0
0.0
0.0
0.0
0.0
0.0
0.0
0.0
0.0
0.0
0.0
0.0
0.0
0.0
0.0
0.0
0.0
0.0

1.0
1.0
1.0
1.0
1.0
1.0
1.0
1.0
1.0
1.0
1.0
1.0
1.0
1.0
1.0
1.0
1.0
1.0
1.0
1.0
1.0
1.0
1.0
1.0
1.0
1.0
1.0
1.0
1.0
1.0


30.0
0.0
0.0
0.0
0.0
30.0
0.0
0.0
0.0
0.0
0.0
0.0
0.0
0.0
0.0
0.0
0.0
0.0
0.0
0.0
0.0
0.0
0.0
0.0
0.0
0.0
0.0
0.0
0.0
0.0

0.74
0.74
0.74
0.74
0.74
0.74
0.74
0.74
0.74
0.74
0.74
0.74
0.74
0.74
0.74
0.74
0.74
0.74
0.74
0.74
0.74
0.74
0.74
0.74
0.74
0.74
0.74
0.74
0.74
0.74


54.0
0.0
0.0
0.0
0.0
0.0
0.0
0.0
0.0
0.0
51.0
3.0
0.0
0.0
0.0
0.0
0.0
0.0
0.0
0.0
0.0
0.0
0.0
0.0
0.0
0.0
0.0
0.0
0.0
0.0


54.0
0.0
0.0
0.0
0.0
0.0
0.0
0.0
0.0
0.0
51.0
3.0
0.0
0.0
0.0
0.0
0.0
0.0
0.0
0.0
0.0
0.0
0.0
0.0
0.0
0.0
0.0
0.0
0.0
0.0

0.97
0.97
0.97
0.97
0.97
0.97
0.97
0.97
0.97
0.97
0.97
0.97
0.97
0.97
0.97
0.97
0.97
0.97
0.97
0.97
0.97
0.97
0.97
0.97
0.97
0.97
0.97
0.97
0.97
0.97


23.0
23.0
0.0
0.0
0.0
0.0
0.0
0.0
0.0
0.0
0.0
0.0
0.0
0.0
0.0
0.0
0.0
0.0
0.0
0.0
0.0
0.0
0.0
0.0
0.0
0.0
0.0
0.0
0.0
0.0


23.0
23.0
0.0
0.0
0.0
0.0
0.0
0.0
0.0
0.0
0.0
0.0
0.0
0.0
0.0
0.0
0.0
0.0
0.0
0.0
0.0
0.0
0.0
0.0
0.0
0.0
0.0
0.0
0.0
0.0

1.0
1.0
1.0
1.0
1.0
1.0
1.0
1.0
1.0
1.0
1.0
1.0
1.0
1.0
1.0
1.0
1.0
1.0
1.0
1.0
1.0
1.0
1.0
1.0
1.0
1.0
1.0
1.0
1.0
1.0


21.0
0.0
21.0
0.0
0.0
0.0
0.0
0.0
0.0
0.0
0.0
0.0
0.0
0.0
0.0
0.0
0.0
0.0
0.0
0.0
0.0
0.0
0.0
0.0
0.0
0.0
0.0
0.0
0.0
0.0


21.0
0.0
21.0
0.0
0.0
0.0
0.0
0.0
0.0
0.0
0.0
0.0
0.0
0.0
0.0
0.0
0.0
0.0
0.0
0.0
0.0
0.0
0.0
0.0
0.0
0.0
0.0
0.0
0.0
0.0

1.0
1.0
1.0
1.0
1.0
1.0
1.0
1.0
1.0
1.0
1.0
1.0
1.0
1.0
1.0
1.0
1.0
1.0
1.0
1.0
1.0
1.0
1.0
1.0
1.0
1.0
1.0
1.0
1.0
1.0


18.0
0.0
0.0
0.0
0.0
0.0
0.0
0.0
0.0
0.0
0.0
18.0
0.0
0.0
0.0
0.0
0.0
0.0
0.0
0.0
0.0
0.0
0.0
0.0
0.0
0.0
0.0
0.0
0.0
0.0


18.0
0.0
0.0
0.0
0.0
0.0
0.0
0.0
0.0
0.0
0.0
18.0
0.0
0.0
0.0
0.0
0.0
0.0
0.0
0.0
0.0
0.0
0.0
0.0
0.0
0.0
0.0
0.0
0.0
0.0

1.0
1.0
1.0
1.0
1.0
1.0
1.0
1.0
1.0
1.0
1.0
1.0
1.0
1.0
1.0
1.0
1.0
1.0
1.0
1.0
1.0
1.0
1.0
1.0
1.0
1.0
1.0
1.0
1.0
1.0


14.0
0.0
1.0
0.0
0.0
5.0
0.0
0.0
0.0
0.0
8.0
0.0
0.0
0.0
0.0
0.0
0.0
0.0
0.0
0.0
0.0
0.0
0.0
0.0
0.0
0.0
0.0
0.0
0.0
0.0

1.0
1.0
1.0
1.0
1.0
1.0
1.0
1.0
1.0
1.0
1.0
1.0
1.0
1.0
1.0
1.0
1.0
1.0
1.0
1.0
1.0
1.0
1.0
1.0
1.0
1.0
1.0
1.0
1.0
1.0


14.0
0.0
0.0
0.0
0.0
0.0
0.0
0.0
0.0
11.0
0.0
3.0
0.0
0.0
0.0
0.0
0.0
0.0
0.0
0.0
0.0
0.0
0.0
0.0
0.0
0.0
0.0
0.0
0.0
0.0


14.0
0.0
0.0
0.0
0.0
0.0
0.0
0.0
0.0
11.0
0.0
3.0
0.0
0.0
0.0
0.0
0.0
0.0
0.0
0.0
0.0
0.0
0.0
0.0
0.0
0.0
0.0
0.0
0.0
0.0

1.0
1.0
1.0
1.0
1.0
1.0
1.0
1.0
1.0
1.0
1.0
1.0
1.0
1.0
1.0
1.0
1.0
1.0
1.0
1.0
1.0
1.0
1.0
1.0
1.0
1.0
1.0
1.0
1.0
1.0


13.0
1.0
4.0
0.0
0.0
0.0
0.0
0.0
0.0
8.0
0.0
0.0
0.0
0.0
0.0
0.0
0.0
0.0
0.0
0.0
0.0
0.0
0.0
0.0
0.0
0.0
0.0
0.0
0.0
0.0


13.0
1.0
4.0
0.0
0.0
0.0
0.0
0.0
0.0
8.0
0.0
0.0
0.0
0.0
0.0
0.0
0.0
0.0
0.0
0.0
0.0
0.0
0.0
0.0
0.0
0.0
0.0
0.0
0.0
0.0

1.0
1.0
1.0
1.0
1.0
1.0
1.0
1.0
1.0
1.0
1.0
1.0
1.0
1.0
1.0
1.0
1.0
1.0
1.0
1.0
1.0
1.0
1.0
1.0
1.0
1.0
1.0
1.0
1.0
1.0


11.0
0.0
0.0
0.0
0.0
0.0
0.0
0.0
0.0
6.0
5.0
0.0
0.0
0.0
0.0
0.0
0.0
0.0
0.0
0.0
0.0
0.0
0.0
0.0
0.0
0.0
0.0
0.0
0.0
0.0


11.0
0.0
0.0
0.0
0.0
0.0
0.0
0.0
0.0
6.0
5.0
0.0
0.0
0.0
0.0
0.0
0.0
0.0
0.0
0.0
0.0
0.0
0.0
0.0
0.0
0.0
0.0
0.0
0.0
0.0

1.0
1.0
1.0
1.0
1.0
1.0
1.0
1.0
1.0
1.0
1.0
1.0
1.0
1.0
1.0
1.0
1.0
1.0
1.0
1.0
1.0
1.0
1.0
1.0
1.0
1.0
1.0
1.0
1.0
1.0


2598.0
0.0
0.0
0.0
0.0
0.0
0.0
0.0
0.0
0.0
0.0
8.0
0.0
0.0
0.0
537.0
592.0
0.0
0.0
1186.0
0.0
0.0
0.0
0.0
134.0
0.0
0.0
0.0
0.0
141.0

1.0
1.0
1.0
1.0
1.0
1.0
1.0
1.0
1.0
1.0
1.0
1.0
1.0
1.0
1.0
1.0
1.0
1.0
1.0
1.0
1.0
1.0
1.0
1.0
1.0
1.0
1.0
1.0
1.0
1.0


721044.0
0.0
0.0
0.0
0.0
0.0
0.0
0.0
0.0
0.0
0.0
0.0
1.0
0.0
0.0
0.0
0.0
1.0
1.0
0.0
0.0
426913.0
294127.0
1.0
0.0
0.0
0.0
0.0
0.0
0.0

1.0
1.0
1.0
1.0
1.0
1.0
1.0
1.0
1.0
1.0
1.0
1.0
1.0
1.0
1.0
1.0
1.0
1.0
1.0
1.0
1.0
1.0
1.0
1.0
1.0
1.0
1.0
1.0
1.0
1.0


165264.0
0.0
0.0
0.0
0.0
0.0
0.0
0.0
0.0
0.0
0.0
0.0
0.0
0.0
0.0
0.0
0.0
0.0
0.0
0.0
3430.0
55996.0
105838.0
0.0
0.0
0.0
0.0
0.0
0.0
0.0


165264.0
0.0
0.0
0.0
0.0
0.0
0.0
0.0
0.0
0.0
0.0
0.0
0.0
0.0
0.0
0.0
0.0
0.0
0.0
0.0
3430.0
55996.0
105838.0
0.0
0.0
0.0
0.0
0.0
0.0
0.0

1.0
1.0
1.0
1.0
1.0
1.0
1.0
1.0
1.0
1.0
1.0
1.0
1.0
1.0
1.0
1.0
1.0
1.0
1.0
1.0
1.0
1.0
1.0
1.0
1.0
1.0
1.0
1.0
1.0
1.0


107392.0
0.0
0.0
0.0
0.0
0.0
0.0
0.0
0.0
0.0
0.0
0.0
0.0
0.0
0.0
0.0
0.0
99207.0
0.0
0.0
8055.0
0.0
0.0
0.0
130.0
0.0
0.0
0.0
0.0
0.0

1.0
1.0
1.0
1.0
1.0
1.0
1.0
1.0
1.0
1.0
1.0
1.0
1.0
1.0
1.0
1.0
1.0
1.0
1.0
1.0
1.0
1.0
1.0
1.0
1.0
1.0
1.0
1.0
1.0
1.0


22597.0
0.0
0.0
0.0
0.0
0.0
0.0
0.0
0.0
0.0
0.0
0.0
1221.0
0.0
201.0
0.0
0.0
2515.0
0.0
13517.0
3922.0
0.0
0.0
1221.0
0.0
0.0
0.0
0.0
0.0
0.0

1.0
1.0
1.0
1.0
1.0
1.0
1.0
1.0
1.0
1.0
1.0
1.0
1.0
1.0
1.0
1.0
1.0
1.0
1.0
1.0
1.0
1.0
1.0
1.0
1.0
1.0
1.0
1.0
1.0
1.0


201.0
0.0
0.0
0.0
0.0
0.0
0.0
0.0
0.0
0.0
0.0
0.0
0.0
0.0
201.0
0.0
0.0
0.0
0.0
0.0
0.0
0.0
0.0
0.0
0.0
0.0
0.0
0.0
0.0
0.0

0.96
0.96
0.96
0.96
0.96
0.96
0.96
0.96
0.96
0.96
0.96
0.96
0.96
0.96
0.96
0.96
0.96
0.96
0.96
0.96
0.96
0.96
0.96
0.96
0.96
0.96
0.96
0.96
0.96
0.96


149.0
0.0
0.0
0.0
0.0
0.0
0.0
0.0
0.0
0.0
0.0
0.0
0.0
0.0
0.0
0.0
0.0
0.0
0.0
149.0
0.0
0.0
0.0
0.0
0.0
0.0
0.0
0.0
0.0
0.0

1.0
1.0
1.0
1.0
1.0
1.0
1.0
1.0
1.0
1.0
1.0
1.0
1.0
1.0
1.0
1.0
1.0
1.0
1.0
1.0
1.0
1.0
1.0
1.0
1.0
1.0
1.0
1.0
1.0
1.0


15851.0
0.0
0.0
0.0
0.0
0.0
0.0
0.0
0.0
0.0
0.0
0.0
0.0
0.0
0.0
0.0
0.0
0.0
0.0
0.0
15851.0
0.0
0.0
0.0
0.0
0.0
0.0
0.0
0.0
0.0

1.0
1.0
1.0
1.0
1.0
1.0
1.0
1.0
1.0
1.0
1.0
1.0
1.0
1.0
1.0
1.0
1.0
1.0
1.0
1.0
1.0
1.0
1.0
1.0
1.0
1.0
1.0
1.0
1.0
1.0


9951.0
0.0
0.0
0.0
0.0
0.0
0.0
0.0
0.0
0.0
0.0
0.0
0.0
0.0
0.0
4650.0
1952.0
1319.0
347.0
503.0
0.0
0.0
0.0
0.0
0.0
0.0
0.0
1180.0
0.0
0.0


9951.0
0.0
0.0
0.0
0.0
0.0
0.0
0.0
0.0
0.0
0.0
0.0
0.0
0.0
0.0
4650.0
1952.0
1319.0
347.0
503.0
0.0
0.0
0.0
0.0
0.0
0.0
0.0
1180.0
0.0
0.0

0.92
0.92
0.92
0.92
0.92
0.92
0.92
0.92
0.92
0.92
0.92
0.92
0.92
0.92
0.92
0.92
0.92
0.92
0.92
0.92
0.92
0.92
0.92
0.92
0.92
0.92
0.92
0.92
0.92
0.92


10210.0
0.0
0.0
0.0
0.0
0.0
0.0
0.0
0.0
0.0
0.0
0.0
0.0
0.0
0.0
0.0
0.0
0.0
0.0
0.0
0.0
2092.0
8118.0
0.0
0.0
0.0
0.0
0.0
0.0
0.0

0.87
0.87
0.87
0.87
0.87
0.87
0.87
0.87
0.87
0.87
0.87
0.87
0.87
0.87
0.87
0.87
0.87
0.87
0.87
0.87
0.87
0.87
0.87
0.87
0.87
0.87
0.87
0.87
0.87
0.87


1638.0
0.0
0.0
0.0
0.0
0.0
0.0
0.0
0.0
0.0
0.0
0.0
0.0
0.0
0.0
0.0
0.0
0.0
0.0
657.0
579.0
402.0
0.0
0.0
0.0
0.0
0.0
0.0
0.0
0.0


1638.0
0.0
0.0
0.0
0.0
0.0
0.0
0.0
0.0
0.0
0.0
0.0
0.0
0.0
0.0
0.0
0.0
0.0
0.0
657.0
579.0
402.0
0.0
0.0
0.0
0.0
0.0
0.0
0.0
0.0

1.0
1.0
1.0
1.0
1.0
1.0
1.0
1.0
1.0
1.0
1.0
1.0
1.0
1.0
1.0
1.0
1.0
1.0
1.0
1.0
1.0
1.0
1.0
1.0
1.0
1.0
1.0
1.0
1.0
1.0


1126.0
0.0
0.0
0.0
0.0
0.0
0.0
0.0
0.0
0.0
0.0
0.0
563.0
0.0
0.0
0.0
0.0
0.0
0.0
0.0
0.0
0.0
0.0
563.0
0.0
0.0
0.0
0.0
0.0
0.0

1.0
1.0
1.0
1.0
1.0
1.0
1.0
1.0
1.0
1.0
1.0
1.0
1.0
1.0
1.0
1.0
1.0
1.0
1.0
1.0
1.0
1.0
1.0
1.0
1.0
1.0
1.0
1.0
1.0
1.0


750.0
0.0
0.0
0.0
0.0
0.0
0.0
0.0
0.0
0.0
0.0
0.0
0.0
0.0
0.0
0.0
0.0
242.0
0.0
316.0
0.0
0.0
0.0
0.0
192.0
0.0
0.0
0.0
0.0
0.0


750.0
0.0
0.0
0.0
0.0
0.0
0.0
0.0
0.0
0.0
0.0
0.0
0.0
0.0
0.0
0.0
0.0
242.0
0.0
316.0
0.0
0.0
0.0
0.0
192.0
0.0
0.0
0.0
0.0
0.0

1.0
1.0
1.0
1.0
1.0
1.0
1.0
1.0
1.0
1.0
1.0
1.0
1.0
1.0
1.0
1.0
1.0
1.0
1.0
1.0
1.0
1.0
1.0
1.0
1.0
1.0
1.0
1.0
1.0
1.0


415.0
0.0
0.0
0.0
0.0
0.0
0.0
0.0
0.0
0.0
0.0
0.0
0.0
0.0
0.0
0.0
0.0
415.0
0.0
0.0
0.0
0.0
0.0
0.0
0.0
0.0
0.0
0.0
0.0
0.0

1.0
1.0
1.0
1.0
1.0
1.0
1.0
1.0
1.0
1.0
1.0
1.0
1.0
1.0
1.0
1.0
1.0
1.0
1.0
1.0
1.0
1.0
1.0
1.0
1.0
1.0
1.0
1.0
1.0
1.0


5164.0
61.0
277.0
0.0
0.0
0.0
0.0
1577.0
1153.0
1667.0
345.0
84.0
0.0
0.0
0.0
0.0
0.0
0.0
0.0
0.0
0.0
0.0
0.0
0.0
0.0
0.0
0.0
0.0
0.0
0.0


5164.0
61.0
277.0
0.0
0.0
0.0
0.0
1577.0
1153.0
1667.0
345.0
84.0
0.0
0.0
0.0
0.0
0.0
0.0
0.0
0.0
0.0
0.0
0.0
0.0
0.0
0.0
0.0
0.0
0.0
0.0


1685.0
0.0
0.0
0.0
0.0
0.0
0.0
0.0
921.0
764.0
0.0
0.0
0.0
0.0
0.0
0.0
0.0
0.0
0.0
0.0
0.0
0.0
0.0
0.0
0.0
0.0
0.0
0.0
0.0
0.0

1.0
1.0
1.0
1.0
1.0
1.0
1.0
1.0
1.0
1.0
1.0
1.0
1.0
1.0
1.0
1.0
1.0
1.0
1.0
1.0
1.0
1.0
1.0
1.0
1.0
1.0
1.0
1.0
1.0
1.0


1577.0
0.0
0.0
0.0
0.0
0.0
0.0
1577.0
0.0
0.0
0.0
0.0
0.0
0.0
0.0
0.0
0.0
0.0
0.0
0.0
0.0
0.0
0.0
0.0
0.0
0.0
0.0
0.0
0.0
0.0

1.0
1.0
1.0
1.0
1.0
1.0
1.0
1.0
1.0
1.0
1.0
1.0
1.0
1.0
1.0
1.0
1.0
1.0
1.0
1.0
1.0
1.0
1.0
1.0
1.0
1.0
1.0
1.0
1.0
1.0


1598.0
54.0
3.0
0.0
0.0
0.0
0.0
0.0
232.0
903.0
322.0
84.0
0.0
0.0
0.0
0.0
0.0
0.0
0.0
0.0
0.0
0.0
0.0
0.0
0.0
0.0
0.0
0.0
0.0
0.0

1.0
1.0
1.0
1.0
1.0
1.0
1.0
1.0
1.0
1.0
1.0
1.0
1.0
1.0
1.0
1.0
1.0
1.0
1.0
1.0
1.0
1.0
1.0
1.0
1.0
1.0
1.0
1.0
1.0
1.0


274.0
0.0
274.0
0.0
0.0
0.0
0.0
0.0
0.0
0.0
0.0
0.0
0.0
0.0
0.0
0.0
0.0
0.0
0.0
0.0
0.0
0.0
0.0
0.0
0.0
0.0
0.0
0.0
0.0
0.0

1.0
1.0
1.0
1.0
1.0
1.0
1.0
1.0
1.0
1.0
1.0
1.0
1.0
1.0
1.0
1.0
1.0
1.0
1.0
1.0
1.0
1.0
1.0
1.0
1.0
1.0
1.0
1.0
1.0
1.0


30.0
7.0
0.0
0.0
0.0
0.0
0.0
0.0
0.0
0.0
23.0
0.0
0.0
0.0
0.0
0.0
0.0
0.0
0.0
0.0
0.0
0.0
0.0
0.0
0.0
0.0
0.0
0.0
0.0
0.0

1.0
1.0
1.0
1.0
1.0
1.0
1.0
1.0
1.0
1.0
1.0
1.0
1.0
1.0
1.0
1.0
1.0
1.0
1.0
1.0
1.0
1.0
1.0
1.0
1.0
1.0
1.0
1.0
1.0
1.0


339339.0
0.0
185.0
42.0
988.0
723.0
1693.0
0.0
0.0
0.0
139.0
1458.0
0.0
0.0
0.0
21342.0
4039.0
24375.0
29917.0
46242.0
8694.0
0.0
0.0
0.0
73978.0
95490.0
1358.0
16383.0
8575.0
3718.0

0.98
0.98
0.98
0.98
0.98
0.98
0.98
0.98
0.98
0.98
0.98
0.98
0.98
0.98
0.98
0.98
0.98
0.98
0.98
0.98
0.98
0.98
0.98
0.98
0.98
0.98
0.98
0.98
0.98
0.98


1738.0
0.0
9.0
0.0
122.0
408.0
679.0
0.0
0.0
0.0
0.0
66.0
0.0
0.0
0.0
277.0
0.0
0.0
0.0
0.0
0.0
0.0
0.0
0.0
0.0
0.0
0.0
177.0
0.0
0.0

1.0
1.0
1.0
1.0
1.0
1.0
1.0
1.0
1.0
1.0
1.0
1.0
1.0
1.0
1.0
1.0
1.0
1.0
1.0
1.0
1.0
1.0
1.0
1.0
1.0
1.0
1.0
1.0
1.0
1.0


1284.0
0.0
9.0
0.0
122.0
408.0
679.0
0.0
0.0
0.0
0.0
66.0
0.0
0.0
0.0
0.0
0.0
0.0
0.0
0.0
0.0
0.0
0.0
0.0
0.0
0.0
0.0
0.0
0.0
0.0

0.72
0.72
0.72
0.72
0.72
0.72
0.72
0.72
0.72
0.72
0.72
0.72
0.72
0.72
0.72
0.72
0.72
0.72
0.72
0.72
0.72
0.72
0.72
0.72
0.72
0.72
0.72
0.72
0.72
0.72


1087.0
0.0
122.0
37.0
612.0
0.0
165.0
0.0
0.0
0.0
0.0
151.0
0.0
0.0
0.0
0.0
0.0
0.0
0.0
0.0
0.0
0.0
0.0
0.0
0.0
0.0
0.0
0.0
0.0
0.0

0.71
0.71
0.71
0.71
0.71
0.71
0.71
0.71
0.71
0.71
0.71
0.71
0.71
0.71
0.71
0.71
0.71
0.71
0.71
0.71
0.71
0.71
0.71
0.71
0.71
0.71
0.71
0.71
0.71
0.71


1010.0
0.0
0.0
0.0
0.0
0.0
0.0
0.0
0.0
0.0
0.0
1010.0
0.0
0.0
0.0
0.0
0.0
0.0
0.0
0.0
0.0
0.0
0.0
0.0
0.0
0.0
0.0
0.0
0.0
0.0


1010.0
0.0
0.0
0.0
0.0
0.0
0.0
0.0
0.0
0.0
0.0
1010.0
0.0
0.0
0.0
0.0
0.0
0.0
0.0
0.0
0.0
0.0
0.0
0.0
0.0
0.0
0.0
0.0
0.0
0.0

1.0
1.0
1.0
1.0
1.0
1.0
1.0
1.0
1.0
1.0
1.0
1.0
1.0
1.0
1.0
1.0
1.0
1.0
1.0
1.0
1.0
1.0
1.0
1.0
1.0
1.0
1.0
1.0
1.0
1.0


60535.0
0.0
20.0
0.0
86.0
37.0
445.0
0.0
0.0
0.0
0.0
21.0
0.0
0.0
0.0
19993.0
4039.0
9462.0
9167.0
11244.0
0.0
0.0
0.0
0.0
0.0
6021.0
0.0
0.0
0.0
0.0


60411.0
0.0
20.0
0.0
86.0
37.0
321.0
0.0
0.0
0.0
0.0
21.0
0.0
0.0
0.0
19993.0
4039.0
9462.0
9167.0
11244.0
0.0
0.0
0.0
0.0
0.0
6021.0
0.0
0.0
0.0
0.0

1.0
1.0
1.0
1.0
1.0
1.0
1.0
1.0
1.0
1.0
1.0
1.0
1.0
1.0
1.0
1.0
1.0
1.0
1.0
1.0
1.0
1.0
1.0
1.0
1.0
1.0
1.0
1.0
1.0
1.0


124.0
0.0
0.0
0.0
0.0
0.0
124.0
0.0
0.0
0.0
0.0
0.0
0.0
0.0
0.0
0.0
0.0
0.0
0.0
0.0
0.0
0.0
0.0
0.0
0.0
0.0
0.0
0.0
0.0
0.0

1.0
1.0
1.0
1.0
1.0
1.0
1.0
1.0
1.0
1.0
1.0
1.0
1.0
1.0
1.0
1.0
1.0
1.0
1.0
1.0
1.0
1.0
1.0
1.0
1.0
1.0
1.0
1.0
1.0
1.0


45988.0
0.0
0.0
0.0
0.0
0.0
202.0
0.0
0.0
0.0
0.0
0.0
0.0
0.0
0.0
0.0
0.0
0.0
20750.0
25036.0
0.0
0.0
0.0
0.0
0.0
0.0
0.0
0.0
0.0
0.0


45988.0
0.0
0.0
0.0
0.0
0.0
202.0
0.0
0.0
0.0
0.0
0.0
0.0
0.0
0.0
0.0
0.0
0.0
20750.0
25036.0
0.0
0.0
0.0
0.0
0.0
0.0
0.0
0.0
0.0
0.0

1.0
1.0
1.0
1.0
1.0
1.0
1.0
1.0
1.0
1.0
1.0
1.0
1.0
1.0
1.0
1.0
1.0
1.0
1.0
1.0
1.0
1.0
1.0
1.0
1.0
1.0
1.0
1.0
1.0
1.0


186.0
0.0
0.0
0.0
0.0
26.0
160.0
0.0
0.0
0.0
0.0
0.0
0.0
0.0
0.0
0.0
0.0
0.0
0.0
0.0
0.0
0.0
0.0
0.0
0.0
0.0
0.0
0.0
0.0
0.0


186.0
0.0
0.0
0.0
0.0
26.0
160.0
0.0
0.0
0.0
0.0
0.0
0.0
0.0
0.0
0.0
0.0
0.0
0.0
0.0
0.0
0.0
0.0
0.0
0.0
0.0
0.0
0.0
0.0
0.0

1.0
1.0
1.0
1.0
1.0
1.0
1.0
1.0
1.0
1.0
1.0
1.0
1.0
1.0
1.0
1.0
1.0
1.0
1.0
1.0
1.0
1.0
1.0
1.0
1.0
1.0
1.0
1.0
1.0
1.0


188.0
0.0
0.0
0.0
0.0
188.0
0.0
0.0
0.0
0.0
0.0
0.0
0.0
0.0
0.0
0.0
0.0
0.0
0.0
0.0
0.0
0.0
0.0
0.0
0.0
0.0
0.0
0.0
0.0
0.0


188.0
0.0
0.0
0.0
0.0
188.0
0.0
0.0
0.0
0.0
0.0
0.0
0.0
0.0
0.0
0.0
0.0
0.0
0.0
0.0
0.0
0.0
0.0
0.0
0.0
0.0
0.0
0.0
0.0
0.0

1.0
1.0
1.0
1.0
1.0
1.0
1.0
1.0
1.0
1.0
1.0
1.0
1.0
1.0
1.0
1.0
1.0
1.0
1.0
1.0
1.0
1.0
1.0
1.0
1.0
1.0
1.0
1.0
1.0
1.0


216.0
0.0
0.0
0.0
0.0
0.0
0.0
0.0
0.0
0.0
139.0
77.0
0.0
0.0
0.0
0.0
0.0
0.0
0.0
0.0
0.0
0.0
0.0
0.0
0.0
0.0
0.0
0.0
0.0
0.0


216.0
0.0
0.0
0.0
0.0
0.0
0.0
0.0
0.0
0.0
139.0
77.0
0.0
0.0
0.0
0.0
0.0
0.0
0.0
0.0
0.0
0.0
0.0
0.0
0.0
0.0
0.0
0.0
0.0
0.0

1.0
1.0
1.0
1.0
1.0
1.0
1.0
1.0
1.0
1.0
1.0
1.0
1.0
1.0
1.0
1.0
1.0
1.0
1.0
1.0
1.0
1.0
1.0
1.0
1.0
1.0
1.0
1.0
1.0
1.0


74088.0
0.0
0.0
5.0
0.0
0.0
0.0
0.0
0.0
0.0
0.0
105.0
0.0
0.0
0.0
0.0
0.0
0.0
0.0
0.0
0.0
0.0
0.0
0.0
73978.0
0.0
0.0
0.0
0.0
0.0

1.0
1.0
1.0
1.0
1.0
1.0
1.0
1.0
1.0
1.0
1.0
1.0
1.0
1.0
1.0
1.0
1.0
1.0
1.0
1.0
1.0
1.0
1.0
1.0
1.0
1.0
1.0
1.0
1.0
1.0


110.0
0.0
0.0
5.0
0.0
0.0
0.0
0.0
0.0
0.0
0.0
105.0
0.0
0.0
0.0
0.0
0.0
0.0
0.0
0.0
0.0
0.0
0.0
0.0
0.0
0.0
0.0
0.0
0.0
0.0

1.0
1.0
1.0
1.0
1.0
1.0
1.0
1.0
1.0
1.0
1.0
1.0
1.0
1.0
1.0
1.0
1.0
1.0
1.0
1.0
1.0
1.0
1.0
1.0
1.0
1.0
1.0
1.0
1.0
1.0


64313.0
0.0
0.0
0.0
0.0
0.0
0.0
0.0
0.0
0.0
0.0
0.0
0.0
0.0
0.0
0.0
0.0
0.0
0.0
0.0
0.0
0.0
0.0
0.0
64313.0
0.0
0.0
0.0
0.0
0.0

1.0
1.0
1.0
1.0
1.0
1.0
1.0
1.0
1.0
1.0
1.0
1.0
1.0
1.0
1.0
1.0
1.0
1.0
1.0
1.0
1.0
1.0
1.0
1.0
1.0
1.0
1.0
1.0
1.0
1.0


1502.0
0.0
34.0
0.0
0.0
64.0
39.0
0.0
0.0
0.0
0.0
0.0
0.0
0.0
0.0
0.0
0.0
0.0
0.0
0.0
0.0
0.0
0.0
0.0
0.0
1365.0
0.0
0.0
0.0
0.0


1438.0
0.0
34.0
0.0
0.0
0.0
39.0
0.0
0.0
0.0
0.0
0.0
0.0
0.0
0.0
0.0
0.0
0.0
0.0
0.0
0.0
0.0
0.0
0.0
0.0
1365.0
0.0
0.0
0.0
0.0

1.0
1.0
1.0
1.0
1.0
1.0
1.0
1.0
1.0
1.0
1.0
1.0
1.0
1.0
1.0
1.0
1.0
1.0
1.0
1.0
1.0
1.0
1.0
1.0
1.0
1.0
1.0
1.0
1.0
1.0


64.0
0.0
0.0
0.0
0.0
64.0
0.0
0.0
0.0
0.0
0.0
0.0
0.0
0.0
0.0
0.0
0.0
0.0
0.0
0.0
0.0
0.0
0.0
0.0
0.0
0.0
0.0
0.0
0.0
0.0

1.0
1.0
1.0
1.0
1.0
1.0
1.0
1.0
1.0
1.0
1.0
1.0
1.0
1.0
1.0
1.0
1.0
1.0
1.0
1.0
1.0
1.0
1.0
1.0
1.0
1.0
1.0
1.0
1.0
1.0


28.0
0.0
0.0
0.0
0.0
0.0
0.0
0.0
0.0
0.0
0.0
28.0
0.0
0.0
0.0
0.0
0.0
0.0
0.0
0.0
0.0
0.0
0.0
0.0
0.0
0.0
0.0
0.0
0.0
0.0


28.0
0.0
0.0
0.0
0.0
0.0
0.0
0.0
0.0
0.0
0.0
28.0
0.0
0.0
0.0
0.0
0.0
0.0
0.0
0.0
0.0
0.0
0.0
0.0
0.0
0.0
0.0
0.0
0.0
0.0

0.98
0.98
0.98
0.98
0.98
0.98
0.98
0.98
0.98
0.98
0.98
0.98
0.98
0.98
0.98
0.98
0.98
0.98
0.98
0.98
0.98
0.98
0.98
0.98
0.98
0.98
0.98
0.98
0.98
0.98


14913.0
0.0
0.0
0.0
0.0
0.0
0.0
0.0
0.0
0.0
0.0
0.0
0.0
0.0
0.0
0.0
0.0
14913.0
0.0
0.0
0.0
0.0
0.0
0.0
0.0
0.0
0.0
0.0
0.0
0.0

1.0
1.0
1.0
1.0
1.0
1.0
1.0
1.0
1.0
1.0
1.0
1.0
1.0
1.0
1.0
1.0
1.0
1.0
1.0
1.0
1.0
1.0
1.0
1.0
1.0
1.0
1.0
1.0
1.0
1.0


9962.0
0.0
0.0
0.0
0.0
0.0
0.0
0.0
0.0
0.0
0.0
0.0
0.0
0.0
0.0
0.0
0.0
0.0
0.0
9962.0
0.0
0.0
0.0
0.0
0.0
0.0
0.0
0.0
0.0
0.0

1.0
1.0
1.0
1.0
1.0
1.0
1.0
1.0
1.0
1.0
1.0
1.0
1.0
1.0
1.0
1.0
1.0
1.0
1.0
1.0
1.0
1.0
1.0
1.0
1.0
1.0
1.0
1.0
1.0
1.0


9954.0
0.0
0.0
0.0
0.0
0.0
0.0
0.0
0.0
0.0
0.0
0.0
0.0
0.0
0.0
0.0
0.0
0.0
0.0
9954.0
0.0
0.0
0.0
0.0
0.0
0.0
0.0
0.0
0.0
0.0

1.0
1.0
1.0
1.0
1.0
1.0
1.0
1.0
1.0
1.0
1.0
1.0
1.0
1.0
1.0
1.0
1.0
1.0
1.0
1.0
1.0
1.0
1.0
1.0
1.0
1.0
1.0
1.0
1.0
1.0


8694.0
0.0
0.0
0.0
0.0
0.0
0.0
0.0
0.0
0.0
0.0
0.0
0.0
0.0
0.0
0.0
0.0
0.0
0.0
0.0
8694.0
0.0
0.0
0.0
0.0
0.0
0.0
0.0
0.0
0.0


8694.0
0.0
0.0
0.0
0.0
0.0
0.0
0.0
0.0
0.0
0.0
0.0
0.0
0.0
0.0
0.0
0.0
0.0
0.0
0.0
8694.0
0.0
0.0
0.0
0.0
0.0
0.0
0.0
0.0
0.0

1.0
1.0
1.0
1.0
1.0
1.0
1.0
1.0
1.0
1.0
1.0
1.0
1.0
1.0
1.0
1.0
1.0
1.0
1.0
1.0
1.0
1.0
1.0
1.0
1.0
1.0
1.0
1.0
1.0
1.0


411.0
0.0
0.0
0.0
0.0
0.0
0.0
0.0
0.0
0.0
0.0
0.0
0.0
0.0
0.0
411.0
0.0
0.0
0.0
0.0
0.0
0.0
0.0
0.0
0.0
0.0
0.0
0.0
0.0
0.0


411.0
0.0
0.0
0.0
0.0
0.0
0.0
0.0
0.0
0.0
0.0
0.0
0.0
0.0
0.0
411.0
0.0
0.0
0.0
0.0
0.0
0.0
0.0
0.0
0.0
0.0
0.0
0.0
0.0
0.0

1.0
1.0
1.0
1.0
1.0
1.0
1.0
1.0
1.0
1.0
1.0
1.0
1.0
1.0
1.0
1.0
1.0
1.0
1.0
1.0
1.0
1.0
1.0
1.0
1.0
1.0
1.0
1.0
1.0
1.0


1002820.0
173.0
8.0
20.0
6.0
0.0
0.0
0.0
301.0
538.0
0.0
0.0
0.0
159495.0
492075.0
0.0
0.0
0.0
0.0
0.0
0.0
0.0
0.0
0.0
350199.0
5.0
0.0
0.0
0.0
0.0

1.0
1.0
1.0
1.0
1.0
1.0
1.0
1.0
1.0
1.0
1.0
1.0
1.0
1.0
1.0
1.0
1.0
1.0
1.0
1.0
1.0
1.0
1.0
1.0
1.0
1.0
1.0
1.0
1.0
1.0


936.0
172.0
0.0
0.0
0.0
0.0
0.0
0.0
265.0
499.0
0.0
0.0
0.0
0.0
0.0
0.0
0.0
0.0
0.0
0.0
0.0
0.0
0.0
0.0
0.0
0.0
0.0
0.0
0.0
0.0


936.0
172.0
0.0
0.0
0.0
0.0
0.0
0.0
265.0
499.0
0.0
0.0
0.0
0.0
0.0
0.0
0.0
0.0
0.0
0.0
0.0
0.0
0.0
0.0
0.0
0.0
0.0
0.0
0.0
0.0

1.0
1.0
1.0
1.0
1.0
1.0
1.0
1.0
1.0
1.0
1.0
1.0
1.0
1.0
1.0
1.0
1.0
1.0
1.0
1.0
1.0
1.0
1.0
1.0
1.0
1.0
1.0
1.0
1.0
1.0


40.0
1.0
0.0
0.0
0.0
0.0
0.0
0.0
0.0
39.0
0.0
0.0
0.0
0.0
0.0
0.0
0.0
0.0
0.0
0.0
0.0
0.0
0.0
0.0
0.0
0.0
0.0
0.0
0.0
0.0


40.0
1.0
0.0
0.0
0.0
0.0
0.0
0.0
0.0
39.0
0.0
0.0
0.0
0.0
0.0
0.0
0.0
0.0
0.0
0.0
0.0
0.0
0.0
0.0
0.0
0.0
0.0
0.0
0.0
0.0

1.0
1.0
1.0
1.0
1.0
1.0
1.0
1.0
1.0
1.0
1.0
1.0
1.0
1.0
1.0
1.0
1.0
1.0
1.0
1.0
1.0
1.0
1.0
1.0
1.0
1.0
1.0
1.0
1.0
1.0


36.0
0.0
0.0
0.0
0.0
0.0
0.0
0.0
36.0
0.0
0.0
0.0
0.0
0.0
0.0
0.0
0.0
0.0
0.0
0.0
0.0
0.0
0.0
0.0
0.0
0.0
0.0
0.0
0.0
0.0


36.0
0.0
0.0
0.0
0.0
0.0
0.0
0.0
36.0
0.0
0.0
0.0
0.0
0.0
0.0
0.0
0.0
0.0
0.0
0.0
0.0
0.0
0.0
0.0
0.0
0.0
0.0
0.0
0.0
0.0

1.0
1.0
1.0
1.0
1.0
1.0
1.0
1.0
1.0
1.0
1.0
1.0
1.0
1.0
1.0
1.0
1.0
1.0
1.0
1.0
1.0
1.0
1.0
1.0
1.0
1.0
1.0
1.0
1.0
1.0


935107.0
0.0
0.0
0.0
0.0
0.0
0.0
0.0
0.0
0.0
0.0
0.0
0.0
148401.0
448514.0
0.0
0.0
0.0
0.0
0.0
0.0
0.0
0.0
0.0
338187.0
5.0
0.0
0.0
0.0
0.0


935107.0
0.0
0.0
0.0
0.0
0.0
0.0
0.0
0.0
0.0
0.0
0.0
0.0
148401.0
448514.0
0.0
0.0
0.0
0.0
0.0
0.0
0.0
0.0
0.0
338187.0
5.0
0.0
0.0
0.0
0.0

1.0
1.0
1.0
1.0
1.0
1.0
1.0
1.0
1.0
1.0
1.0
1.0
1.0
1.0
1.0
1.0
1.0
1.0
1.0
1.0
1.0
1.0
1.0
1.0
1.0
1.0
1.0
1.0
1.0
1.0


112549.0
0.0
95.0
379.0
0.0
358.0
0.0
0.0
0.0
0.0
14.0
0.0
0.0
0.0
0.0
5965.0
0.0
22886.0
0.0
0.0
0.0
0.0
0.0
0.0
69803.0
0.0
0.0
0.0
13049.0
0.0

0.83
0.83
0.83
0.83
0.83
0.83
0.83
0.83
0.83
0.83
0.83
0.83
0.83
0.83
0.83
0.83
0.83
0.83
0.83
0.83
0.83
0.83
0.83
0.83
0.83
0.83
0.83
0.83
0.83
0.83


112535.0
0.0
95.0
379.0
0.0
358.0
0.0
0.0
0.0
0.0
0.0
0.0
0.0
0.0
0.0
5965.0
0.0
22886.0
0.0
0.0
0.0
0.0
0.0
0.0
69803.0
0.0
0.0
0.0
13049.0
0.0


112535.0
0.0
95.0
379.0
0.0
358.0
0.0
0.0
0.0
0.0
0.0
0.0
0.0
0.0
0.0
5965.0
0.0
22886.0
0.0
0.0
0.0
0.0
0.0
0.0
69803.0
0.0
0.0
0.0
13049.0
0.0

1.0
1.0
1.0
1.0
1.0
1.0
1.0
1.0
1.0
1.0
1.0
1.0
1.0
1.0
1.0
1.0
1.0
1.0
1.0
1.0
1.0
1.0
1.0
1.0
1.0
1.0
1.0
1.0
1.0
1.0


657.0
0.0
56.0
0.0
120.0
480.0
1.0
0.0
0.0
0.0
0.0
0.0
0.0
0.0
0.0
0.0
0.0
0.0
0.0
0.0
0.0
0.0
0.0
0.0
0.0
0.0
0.0
0.0
0.0
0.0


657.0
0.0
56.0
0.0
120.0
480.0
1.0
0.0
0.0
0.0
0.0
0.0
0.0
0.0
0.0
0.0
0.0
0.0
0.0
0.0
0.0
0.0
0.0
0.0
0.0
0.0
0.0
0.0
0.0
0.0


657.0
0.0
56.0
0.0
120.0
480.0
1.0
0.0
0.0
0.0
0.0
0.0
0.0
0.0
0.0
0.0
0.0
0.0
0.0
0.0
0.0
0.0
0.0
0.0
0.0
0.0
0.0
0.0
0.0
0.0

0.92
0.92
0.92
0.92
0.92
0.92
0.92
0.92
0.92
0.92
0.92
0.92
0.92
0.92
0.92
0.92
0.92
0.92
0.92
0.92
0.92
0.92
0.92
0.92
0.92
0.92
0.92
0.92
0.92
0.92


213525.0
2.0
282.0
0.0
0.0
0.0
0.0
0.0
0.0
10.0
316.0
366.0
0.0
0.0
0.0
12505.0
0.0
0.0
0.0
0.0
0.0
0.0
0.0
0.0
49896.0
0.0
21323.0
87262.0
0.0
41563.0

0.74
0.74
0.74
0.74
0.74
0.74
0.74
0.74
0.74
0.74
0.74
0.74
0.74
0.74
0.74
0.74
0.74
0.74
0.74
0.74
0.74
0.74
0.74
0.74
0.74
0.74
0.74
0.74
0.74
0.74


50784.0
2.0
211.0
0.0
0.0
0.0
0.0
0.0
0.0
0.0
309.0
366.0
0.0
0.0
0.0
0.0
0.0
0.0
0.0
0.0
0.0
0.0
0.0
0.0
49896.0
0.0
0.0
0.0
0.0
0.0

0.87
0.87
0.87
0.87
0.87
0.87
0.87
0.87
0.87
0.87
0.87
0.87
0.87
0.87
0.87
0.87
0.87
0.87
0.87
0.87
0.87
0.87
0.87
0.87
0.87
0.87
0.87
0.87
0.87
0.87


48076.0
2.0
211.0
0.0
0.0
0.0
0.0
0.0
0.0
0.0
309.0
366.0
0.0
0.0
0.0
0.0
0.0
0.0
0.0
0.0
0.0
0.0
0.0
0.0
47188.0
0.0
0.0
0.0
0.0
0.0

1.0
1.0
1.0
1.0
1.0
1.0
1.0
1.0
1.0
1.0
1.0
1.0
1.0
1.0
1.0
1.0
1.0
1.0
1.0
1.0
1.0
1.0
1.0
1.0
1.0
1.0
1.0
1.0
1.0
1.0


88.0
0.0
71.0
0.0
0.0
0.0
0.0
0.0
0.0
10.0
7.0
0.0
0.0
0.0
0.0
0.0
0.0
0.0
0.0
0.0
0.0
0.0
0.0
0.0
0.0
0.0
0.0
0.0
0.0
0.0


71.0
0.0
71.0
0.0
0.0
0.0
0.0
0.0
0.0
0.0
0.0
0.0
0.0
0.0
0.0
0.0
0.0
0.0
0.0
0.0
0.0
0.0
0.0
0.0
0.0
0.0
0.0
0.0
0.0
0.0

0.83
0.83
0.83
0.83
0.83
0.83
0.83
0.83
0.83
0.83
0.83
0.83
0.83
0.83
0.83
0.83
0.83
0.83
0.83
0.83
0.83
0.83
0.83
0.83
0.83
0.83
0.83
0.83
0.83
0.83


17.0
0.0
0.0
0.0
0.0
0.0
0.0
0.0
0.0
10.0
7.0
0.0
0.0
0.0
0.0
0.0
0.0
0.0
0.0
0.0
0.0
0.0
0.0
0.0
0.0
0.0
0.0
0.0
0.0
0.0

1.0
1.0
1.0
1.0
1.0
1.0
1.0
1.0
1.0
1.0
1.0
1.0
1.0
1.0
1.0
1.0
1.0
1.0
1.0
1.0
1.0
1.0
1.0
1.0
1.0
1.0
1.0
1.0
1.0
1.0


142110.0
0.0
0.0
0.0
0.0
0.0
0.0
0.0
0.0
0.0
0.0
0.0
0.0
0.0
0.0
12505.0
0.0
0.0
0.0
0.0
0.0
0.0
0.0
0.0
0.0
0.0
20915.0
87262.0
0.0
21428.0

1.0
1.0
1.0
1.0
1.0
1.0
1.0
1.0
1.0
1.0
1.0
1.0
1.0
1.0
1.0
1.0
1.0
1.0
1.0
1.0
1.0
1.0
1.0
1.0
1.0
1.0
1.0
1.0
1.0
1.0


106365.0
0.0
0.0
0.0
0.0
0.0
0.0
0.0
0.0
0.0
0.0
0.0
0.0
0.0
0.0
0.0
0.0
0.0
0.0
0.0
0.0
0.0
0.0
0.0
0.0
0.0
0.0
85678.0
0.0
20687.0

0.84
0.84
0.84
0.84
0.84
0.84
0.84
0.84
0.84
0.84
0.84
0.84
0.84
0.84
0.84
0.84
0.84
0.84
0.84
0.84
0.84
0.84
0.84
0.84
0.84
0.84
0.84
0.84
0.84
0.84


16538.0
0.0
0.0
0.0
0.0
0.0
0.0
0.0
0.0
0.0
0.0
0.0
0.0
0.0
0.0
0.0
0.0
0.0
0.0
0.0
0.0
0.0
0.0
0.0
0.0
0.0
16538.0
0.0
0.0
0.0

0.98
0.98
0.98
0.98
0.98
0.98
0.98
0.98
0.98
0.98
0.98
0.98
0.98
0.98
0.98
0.98
0.98
0.98
0.98
0.98
0.98
0.98
0.98
0.98
0.98
0.98
0.98
0.98
0.98
0.98


12505.0
0.0
0.0
0.0
0.0
0.0
0.0
0.0
0.0
0.0
0.0
0.0
0.0
0.0
0.0
12505.0
0.0
0.0
0.0
0.0
0.0
0.0
0.0
0.0
0.0
0.0
0.0
0.0
0.0
0.0

1.0
1.0
1.0
1.0
1.0
1.0
1.0
1.0
1.0
1.0
1.0
1.0
1.0
1.0
1.0
1.0
1.0
1.0
1.0
1.0
1.0
1.0
1.0
1.0
1.0
1.0
1.0
1.0
1.0
1.0


14955.0
0.0
0.0
0.0
0.0
0.0
0.0
0.0
0.0
0.0
0.0
0.0
0.0
0.0
0.0
0.0
0.0
0.0
0.0
0.0
0.0
0.0
0.0
0.0
0.0
0.0
0.0
0.0
0.0
14955.0


14955.0
0.0
0.0
0.0
0.0
0.0
0.0
0.0
0.0
0.0
0.0
0.0
0.0
0.0
0.0
0.0
0.0
0.0
0.0
0.0
0.0
0.0
0.0
0.0
0.0
0.0
0.0
0.0
0.0
14955.0

1.0
1.0
1.0
1.0
1.0
1.0
1.0
1.0
1.0
1.0
1.0
1.0
1.0
1.0
1.0
1.0
1.0
1.0
1.0
1.0
1.0
1.0
1.0
1.0
1.0
1.0
1.0
1.0
1.0
1.0


5180.0
0.0
0.0
0.0
0.0
0.0
0.0
0.0
0.0
0.0
0.0
0.0
0.0
0.0
0.0
0.0
0.0
0.0
0.0
0.0
0.0
0.0
0.0
0.0
0.0
0.0
0.0
0.0
0.0
5180.0


5180.0
0.0
0.0
0.0
0.0
0.0
0.0
0.0
0.0
0.0
0.0
0.0
0.0
0.0
0.0
0.0
0.0
0.0
0.0
0.0
0.0
0.0
0.0
0.0
0.0
0.0
0.0
0.0
0.0
5180.0

0.97
0.97
0.97
0.97
0.97
0.97
0.97
0.97
0.97
0.97
0.97
0.97
0.97
0.97
0.97
0.97
0.97
0.97
0.97
0.97
0.97
0.97
0.97
0.97
0.97
0.97
0.97
0.97
0.97
0.97


536135.0
142.0
49.0
31.0
14.0
13.0
19.0
193.0
176.0
160.0
239.0
293.0
0.0
203.0
147.0
258051.0
191183.0
27595.0
37672.0
6312.0
4791.0
0.0
51.0
0.0
1338.0
517.0
112.0
0.0
2342.0
4492.0

1.0
1.0
1.0
1.0
1.0
1.0
1.0
1.0
1.0
1.0
1.0
1.0
1.0
1.0
1.0
1.0
1.0
1.0
1.0
1.0
1.0
1.0
1.0
1.0
1.0
1.0
1.0
1.0
1.0
1.0


320.0
5.0
2.0
6.0
0.0
2.0
0.0
125.0
117.0
31.0
31.0
1.0
0.0
0.0
0.0
0.0
0.0
0.0
0.0
0.0
0.0
0.0
0.0
0.0
0.0
0.0
0.0
0.0
0.0
0.0


320.0
5.0
2.0
6.0
0.0
2.0
0.0
125.0
117.0
31.0
31.0
1.0
0.0
0.0
0.0
0.0
0.0
0.0
0.0
0.0
0.0
0.0
0.0
0.0
0.0
0.0
0.0
0.0
0.0
0.0

1.0
1.0
1.0
1.0
1.0
1.0
1.0
1.0
1.0
1.0
1.0
1.0
1.0
1.0
1.0
1.0
1.0
1.0
1.0
1.0
1.0
1.0
1.0
1.0
1.0
1.0
1.0
1.0
1.0
1.0


30410.0
1.0
6.0
19.0
3.0
8.0
0.0
0.0
0.0
40.0
53.0
109.0
0.0
0.0
0.0
0.0
0.0
152.0
19605.0
5636.0
4778.0
0.0
0.0
0.0
0.0
0.0
0.0
0.0
0.0
0.0

1.0
1.0
1.0
1.0
1.0
1.0
1.0
1.0
1.0
1.0
1.0
1.0
1.0
1.0
1.0
1.0
1.0
1.0
1.0
1.0
1.0
1.0
1.0
1.0
1.0
1.0
1.0
1.0
1.0
1.0


1414.0
1.0
5.0
19.0
3.0
8.0
0.0
0.0
0.0
19.0
40.0
95.0
0.0
0.0
0.0
0.0
0.0
152.0
0.0
0.0
1072.0
0.0
0.0
0.0
0.0
0.0
0.0
0.0
0.0
0.0

1.0
1.0
1.0
1.0
1.0
1.0
1.0
1.0
1.0
1.0
1.0
1.0
1.0
1.0
1.0
1.0
1.0
1.0
1.0
1.0
1.0
1.0
1.0
1.0
1.0
1.0
1.0
1.0
1.0
1.0


211.0
9.0
0.0
0.0
2.0
0.0
0.0
44.0
26.0
34.0
75.0
21.0
0.0
0.0
0.0
0.0
0.0
0.0
0.0
0.0
0.0
0.0
0.0
0.0
0.0
0.0
0.0
0.0
0.0
0.0

1.0
1.0
1.0
1.0
1.0
1.0
1.0
1.0
1.0
1.0
1.0
1.0
1.0
1.0
1.0
1.0
1.0
1.0
1.0
1.0
1.0
1.0
1.0
1.0
1.0
1.0
1.0
1.0
1.0
1.0


83.0
4.0
0.0
0.0
0.0
0.0
0.0
2.0
7.0
22.0
31.0
17.0
0.0
0.0
0.0
0.0
0.0
0.0
0.0
0.0
0.0
0.0
0.0
0.0
0.0
0.0
0.0
0.0
0.0
0.0

0.98
0.98
0.98
0.98
0.98
0.98
0.98
0.98
0.98
0.98
0.98
0.98
0.98
0.98
0.98
0.98
0.98
0.98
0.98
0.98
0.98
0.98
0.98
0.98
0.98
0.98
0.98
0.98
0.98
0.98


16.0
4.0
0.0
0.0
0.0
0.0
0.0
3.0
0.0
1.0
8.0
0.0
0.0
0.0
0.0
0.0
0.0
0.0
0.0
0.0
0.0
0.0
0.0
0.0
0.0
0.0
0.0
0.0
0.0
0.0

1.0
1.0
1.0
1.0
1.0
1.0
1.0
1.0
1.0
1.0
1.0
1.0
1.0
1.0
1.0
1.0
1.0
1.0
1.0
1.0
1.0
1.0
1.0
1.0
1.0
1.0
1.0
1.0
1.0
1.0


8.0
0.0
0.0
0.0
0.0
0.0
0.0
0.0
0.0
0.0
8.0
0.0
0.0
0.0
0.0
0.0
0.0
0.0
0.0
0.0
0.0
0.0
0.0
0.0
0.0
0.0
0.0
0.0
0.0
0.0

0.97
0.97
0.97
0.97
0.97
0.97
0.97
0.97
0.97
0.97
0.97
0.97
0.97
0.97
0.97
0.97
0.97
0.97
0.97
0.97
0.97
0.97
0.97
0.97
0.97
0.97
0.97
0.97
0.97
0.97


83.0
2.0
0.0
0.0
0.0
0.0
0.0
5.0
15.0
31.0
29.0
1.0
0.0
0.0
0.0
0.0
0.0
0.0
0.0
0.0
0.0
0.0
0.0
0.0
0.0
0.0
0.0
0.0
0.0
0.0


83.0
2.0
0.0
0.0
0.0
0.0
0.0
5.0
15.0
31.0
29.0
1.0
0.0
0.0
0.0
0.0
0.0
0.0
0.0
0.0
0.0
0.0
0.0
0.0
0.0
0.0
0.0
0.0
0.0
0.0

1.0
1.0
1.0
1.0
1.0
1.0
1.0
1.0
1.0
1.0
1.0
1.0
1.0
1.0
1.0
1.0
1.0
1.0
1.0
1.0
1.0
1.0
1.0
1.0
1.0
1.0
1.0
1.0
1.0
1.0


62.0
39.0
4.0
0.0
0.0
0.0
0.0
0.0
0.0
0.0
0.0
19.0
0.0
0.0
0.0
0.0
0.0
0.0
0.0
0.0
0.0
0.0
0.0
0.0
0.0
0.0
0.0
0.0
0.0
0.0


62.0
39.0
4.0
0.0
0.0
0.0
0.0
0.0
0.0
0.0
0.0
19.0
0.0
0.0
0.0
0.0
0.0
0.0
0.0
0.0
0.0
0.0
0.0
0.0
0.0
0.0
0.0
0.0
0.0
0.0

1.0
1.0
1.0
1.0
1.0
1.0
1.0
1.0
1.0
1.0
1.0
1.0
1.0
1.0
1.0
1.0
1.0
1.0
1.0
1.0
1.0
1.0
1.0
1.0
1.0
1.0
1.0
1.0
1.0
1.0


491876.0
0.0
21.0
1.0
0.0
0.0
19.0
0.0
0.0
0.0
0.0
24.0
0.0
49.0
85.0
256919.0
190979.0
24083.0
13179.0
0.0
0.0
0.0
0.0
0.0
665.0
0.0
0.0
0.0
1784.0
4068.0

1.0
1.0
1.0
1.0
1.0
1.0
1.0
1.0
1.0
1.0
1.0
1.0
1.0
1.0
1.0
1.0
1.0
1.0
1.0
1.0
1.0
1.0
1.0
1.0
1.0
1.0
1.0
1.0
1.0
1.0


2130.0
0.0
19.0
0.0
0.0
0.0
0.0
0.0
0.0
0.0
0.0
0.0
0.0
0.0
0.0
0.0
0.0
2111.0
0.0
0.0
0.0
0.0
0.0
0.0
0.0
0.0
0.0
0.0
0.0
0.0

0.99
0.99
0.99
0.99
0.99
0.99
0.99
0.99
0.99
0.99
0.99
0.99
0.99
0.99
0.99
0.99
0.99
0.99
0.99
0.99
0.99
0.99
0.99
0.99
0.99
0.99
0.99
0.99
0.99
0.99


3249.0
0.0
0.0
0.0
0.0
0.0
0.0
0.0
0.0
0.0
0.0
0.0
0.0
0.0
0.0
0.0
885.0
2364.0
0.0
0.0
0.0
0.0
0.0
0.0
0.0
0.0
0.0
0.0
0.0
0.0

0.98
0.98
0.98
0.98
0.98
0.98
0.98
0.98
0.98
0.98
0.98
0.98
0.98
0.98
0.98
0.98
0.98
0.98
0.98
0.98
0.98
0.98
0.98
0.98
0.98
0.98
0.98
0.98
0.98
0.98


242.0
8.0
2.0
0.0
0.0
0.0
0.0
0.0
1.0
0.0
8.0
4.0
0.0
0.0
0.0
0.0
0.0
0.0
0.0
0.0
0.0
0.0
0.0
0.0
219.0
0.0
0.0
0.0
0.0
0.0

1.0
1.0
1.0
1.0
1.0
1.0
1.0
1.0
1.0
1.0
1.0
1.0
1.0
1.0
1.0
1.0
1.0
1.0
1.0
1.0
1.0
1.0
1.0
1.0
1.0
1.0
1.0
1.0
1.0
1.0


23.0
8.0
2.0
0.0
0.0
0.0
0.0
0.0
1.0
0.0
8.0
4.0
0.0
0.0
0.0
0.0
0.0
0.0
0.0
0.0
0.0
0.0
0.0
0.0
0.0
0.0
0.0
0.0
0.0
0.0

1.0
1.0
1.0
1.0
1.0
1.0
1.0
1.0
1.0
1.0
1.0
1.0
1.0
1.0
1.0
1.0
1.0
1.0
1.0
1.0
1.0
1.0
1.0
1.0
1.0
1.0
1.0
1.0
1.0
1.0


23.0
0.0
0.0
0.0
0.0
0.0
0.0
7.0
7.0
3.0
6.0
0.0
0.0
0.0
0.0
0.0
0.0
0.0
0.0
0.0
0.0
0.0
0.0
0.0
0.0
0.0
0.0
0.0
0.0
0.0

0.89
0.89
0.89
0.89
0.89
0.89
0.89
0.89
0.89
0.89
0.89
0.89
0.89
0.89
0.89
0.89
0.89
0.89
0.89
0.89
0.89
0.89
0.89
0.89
0.89
0.89
0.89
0.89
0.89
0.89


17.0
0.0
0.0
0.0
0.0
0.0
0.0
0.0
0.0
4.0
11.0
2.0
0.0
0.0
0.0
0.0
0.0
0.0
0.0
0.0
0.0
0.0
0.0
0.0
0.0
0.0
0.0
0.0
0.0
0.0


17.0
0.0
0.0
0.0
0.0
0.0
0.0
0.0
0.0
4.0
11.0
2.0
0.0
0.0
0.0
0.0
0.0
0.0
0.0
0.0
0.0
0.0
0.0
0.0
0.0
0.0
0.0
0.0
0.0
0.0

1.0
1.0
1.0
1.0
1.0
1.0
1.0
1.0
1.0
1.0
1.0
1.0
1.0
1.0
1.0
1.0
1.0
1.0
1.0
1.0
1.0
1.0
1.0
1.0
1.0
1.0
1.0
1.0
1.0
1.0


16.0
12.0
4.0
0.0
0.0
0.0
0.0
0.0
0.0
0.0
0.0
0.0
0.0
0.0
0.0
0.0
0.0
0.0
0.0
0.0
0.0
0.0
0.0
0.0
0.0
0.0
0.0
0.0
0.0
0.0


16.0
12.0
4.0
0.0
0.0
0.0
0.0
0.0
0.0
0.0
0.0
0.0
0.0
0.0
0.0
0.0
0.0
0.0
0.0
0.0
0.0
0.0
0.0
0.0
0.0
0.0
0.0
0.0
0.0
0.0

1.0
1.0
1.0
1.0
1.0
1.0
1.0
1.0
1.0
1.0
1.0
1.0
1.0
1.0
1.0
1.0
1.0
1.0
1.0
1.0
1.0
1.0
1.0
1.0
1.0
1.0
1.0
1.0
1.0
1.0


13.0
12.0
1.0
0.0
0.0
0.0
0.0
0.0
0.0
0.0
0.0
0.0
0.0
0.0
0.0
0.0
0.0
0.0
0.0
0.0
0.0
0.0
0.0
0.0
0.0
0.0
0.0
0.0
0.0
0.0


13.0
12.0
1.0
0.0
0.0
0.0
0.0
0.0
0.0
0.0
0.0
0.0
0.0
0.0
0.0
0.0
0.0
0.0
0.0
0.0
0.0
0.0
0.0
0.0
0.0
0.0
0.0
0.0
0.0
0.0

1.0
1.0
1.0
1.0
1.0
1.0
1.0
1.0
1.0
1.0
1.0
1.0
1.0
1.0
1.0
1.0
1.0
1.0
1.0
1.0
1.0
1.0
1.0
1.0
1.0
1.0
1.0
1.0
1.0
1.0


9.0
0.0
0.0
0.0
0.0
0.0
0.0
0.0
0.0
1.0
0.0
8.0
0.0
0.0
0.0
0.0
0.0
0.0
0.0
0.0
0.0
0.0
0.0
0.0
0.0
0.0
0.0
0.0
0.0
0.0


9.0
0.0
0.0
0.0
0.0
0.0
0.0
0.0
0.0
1.0
0.0
8.0
0.0
0.0
0.0
0.0
0.0
0.0
0.0
0.0
0.0
0.0
0.0
0.0
0.0
0.0
0.0
0.0
0.0
0.0

1.0
1.0
1.0
1.0
1.0
1.0
1.0
1.0
1.0
1.0
1.0
1.0
1.0
1.0
1.0
1.0
1.0
1.0
1.0
1.0
1.0
1.0
1.0
1.0
1.0
1.0
1.0
1.0
1.0
1.0


5230.0
0.0
0.0
0.0
0.0
0.0
0.0
0.0
0.0
0.0
0.0
0.0
0.0
138.0
0.0
0.0
204.0
0.0
4888.0
0.0
0.0
0.0
0.0
0.0
0.0
0.0
0.0
0.0
0.0
0.0

1.0
1.0
1.0
1.0
1.0
1.0
1.0
1.0
1.0
1.0
1.0
1.0
1.0
1.0
1.0
1.0
1.0
1.0
1.0
1.0
1.0
1.0
1.0
1.0
1.0
1.0
1.0
1.0
1.0
1.0


4336.0
0.0
0.0
0.0
0.0
0.0
0.0
0.0
0.0
0.0
0.0
0.0
0.0
0.0
0.0
1062.0
0.0
2896.0
0.0
378.0
0.0
0.0
0.0
0.0
0.0
0.0
0.0
0.0
0.0
0.0

0.71
0.71
0.71
0.71
0.71
0.71
0.71
0.71
0.71
0.71
0.71
0.71
0.71
0.71
0.71
0.71
0.71
0.71
0.71
0.71
0.71
0.71
0.71
0.71
0.71
0.71
0.71
0.71
0.71
0.71


2184.0
0.0
0.0
0.0
0.0
0.0
0.0
0.0
0.0
0.0
0.0
0.0
0.0
0.0
0.0
435.0
0.0
1749.0
0.0
0.0
0.0
0.0
0.0
0.0
0.0
0.0
0.0
0.0
0.0
0.0

0.96
0.96
0.96
0.96
0.96
0.96
0.96
0.96
0.96
0.96
0.96
0.96
0.96
0.96
0.96
0.96
0.96
0.96
0.96
0.96
0.96
0.96
0.96
0.96
0.96
0.96
0.96
0.96
0.96
0.96


1147.0
0.0
0.0
0.0
0.0
0.0
0.0
0.0
0.0
0.0
0.0
0.0
0.0
0.0
0.0
0.0
0.0
1147.0
0.0
0.0
0.0
0.0
0.0
0.0
0.0
0.0
0.0
0.0
0.0
0.0

0.91
0.91
0.91
0.91
0.91
0.91
0.91
0.91
0.91
0.91
0.91
0.91
0.91
0.91
0.91
0.91
0.91
0.91
0.91
0.91
0.91
0.91
0.91
0.91
0.91
0.91
0.91
0.91
0.91
0.91


378.0
0.0
0.0
0.0
0.0
0.0
0.0
0.0
0.0
0.0
0.0
0.0
0.0
0.0
0.0
0.0
0.0
0.0
0.0
378.0
0.0
0.0
0.0
0.0
0.0
0.0
0.0
0.0
0.0
0.0

0.96
0.96
0.96
0.96
0.96
0.96
0.96
0.96
0.96
0.96
0.96
0.96
0.96
0.96
0.96
0.96
0.96
0.96
0.96
0.96
0.96
0.96
0.96
0.96
0.96
0.96
0.96
0.96
0.96
0.96


362.0
0.0
0.0
0.0
0.0
0.0
0.0
0.0
0.0
0.0
0.0
0.0
0.0
0.0
0.0
0.0
0.0
362.0
0.0
0.0
0.0
0.0
0.0
0.0
0.0
0.0
0.0
0.0
0.0
0.0


362.0
0.0
0.0
0.0
0.0
0.0
0.0
0.0
0.0
0.0
0.0
0.0
0.0
0.0
0.0
0.0
0.0
362.0
0.0
0.0
0.0
0.0
0.0
0.0
0.0
0.0
0.0
0.0
0.0
0.0

1.0
1.0
1.0
1.0
1.0
1.0
1.0
1.0
1.0
1.0
1.0
1.0
1.0
1.0
1.0
1.0
1.0
1.0
1.0
1.0
1.0
1.0
1.0
1.0
1.0
1.0
1.0
1.0
1.0
1.0


427.0
0.0
0.0
0.0
0.0
0.0
0.0
0.0
0.0
0.0
0.0
0.0
0.0
0.0
0.0
0.0
0.0
102.0
0.0
0.0
0.0
0.0
0.0
0.0
0.0
200.0
0.0
0.0
0.0
125.0

1.0
1.0
1.0
1.0
1.0
1.0
1.0
1.0
1.0
1.0
1.0
1.0
1.0
1.0
1.0
1.0
1.0
1.0
1.0
1.0
1.0
1.0
1.0
1.0
1.0
1.0
1.0
1.0
1.0
1.0


296.0
0.0
0.0
0.0
0.0
0.0
0.0
0.0
0.0
0.0
0.0
0.0
0.0
16.0
0.0
0.0
0.0
0.0
0.0
267.0
13.0
0.0
0.0
0.0
0.0
0.0
0.0
0.0
0.0
0.0

1.0
1.0
1.0
1.0
1.0
1.0
1.0
1.0
1.0
1.0
1.0
1.0
1.0
1.0
1.0
1.0
1.0
1.0
1.0
1.0
1.0
1.0
1.0
1.0
1.0
1.0
1.0
1.0
1.0
1.0


245.0
0.0
0.0
0.0
0.0
0.0
0.0
0.0
0.0
0.0
0.0
0.0
0.0
0.0
0.0
0.0
0.0
0.0
0.0
0.0
0.0
0.0
0.0
0.0
0.0
10.0
0.0
0.0
0.0
235.0

1.0
1.0
1.0
1.0
1.0
1.0
1.0
1.0
1.0
1.0
1.0
1.0
1.0
1.0
1.0
1.0
1.0
1.0
1.0
1.0
1.0
1.0
1.0
1.0
1.0
1.0
1.0
1.0
1.0
1.0


172.0
0.0
0.0
0.0
0.0
0.0
0.0
0.0
0.0
0.0
0.0
0.0
0.0
0.0
0.0
0.0
0.0
0.0
0.0
0.0
0.0
0.0
0.0
0.0
0.0
172.0
0.0
0.0
0.0
0.0

1.0
1.0
1.0
1.0
1.0
1.0
1.0
1.0
1.0
1.0
1.0
1.0
1.0
1.0
1.0
1.0
1.0
1.0
1.0
1.0
1.0
1.0
1.0
1.0
1.0
1.0
1.0
1.0
1.0
1.0


141.0
0.0
0.0
0.0
0.0
0.0
0.0
0.0
0.0
0.0
0.0
0.0
0.0
0.0
0.0
0.0
0.0
0.0
0.0
0.0
0.0
0.0
0.0
0.0
141.0
0.0
0.0
0.0
0.0
0.0


141.0
0.0
0.0
0.0
0.0
0.0
0.0
0.0
0.0
0.0
0.0
0.0
0.0
0.0
0.0
0.0
0.0
0.0
0.0
0.0
0.0
0.0
0.0
0.0
141.0
0.0
0.0
0.0
0.0
0.0

1.0
1.0
1.0
1.0
1.0
1.0
1.0
1.0
1.0
1.0
1.0
1.0
1.0
1.0
1.0
1.0
1.0
1.0
1.0
1.0
1.0
1.0
1.0
1.0
1.0
1.0
1.0
1.0
1.0
1.0


112.0
0.0
0.0
0.0
0.0
0.0
0.0
0.0
0.0
0.0
0.0
0.0
0.0
0.0
0.0
70.0
0.0
0.0
0.0
0.0
0.0
0.0
0.0
0.0
42.0
0.0
0.0
0.0
0.0
0.0


70.0
0.0
0.0
0.0
0.0
0.0
0.0
0.0
0.0
0.0
0.0
0.0
0.0
0.0
0.0
70.0
0.0
0.0
0.0
0.0
0.0
0.0
0.0
0.0
0.0
0.0
0.0
0.0
0.0
0.0

0.99
0.99
0.99
0.99
0.99
0.99
0.99
0.99
0.99
0.99
0.99
0.99
0.99
0.99
0.99
0.99
0.99
0.99
0.99
0.99
0.99
0.99
0.99
0.99
0.99
0.99
0.99
0.99
0.99
0.99


42.0
0.0
0.0
0.0
0.0
0.0
0.0
0.0
0.0
0.0
0.0
0.0
0.0
0.0
0.0
0.0
0.0
0.0
0.0
0.0
0.0
0.0
0.0
0.0
42.0
0.0
0.0
0.0
0.0
0.0

1.0
1.0
1.0
1.0
1.0
1.0
1.0
1.0
1.0
1.0
1.0
1.0
1.0
1.0
1.0
1.0
1.0
1.0
1.0
1.0
1.0
1.0
1.0
1.0
1.0
1.0
1.0
1.0
1.0
1.0


62.0
0.0
0.0
0.0
0.0
0.0
0.0
0.0
0.0
0.0
0.0
0.0
0.0
0.0
62.0
0.0
0.0
0.0
0.0
0.0
0.0
0.0
0.0
0.0
0.0
0.0
0.0
0.0
0.0
0.0

0.97
0.97
0.97
0.97
0.97
0.97
0.97
0.97
0.97
0.97
0.97
0.97
0.97
0.97
0.97
0.97
0.97
0.97
0.97
0.97
0.97
0.97
0.97
0.97
0.97
0.97
0.97
0.97
0.97
0.97


62.0
0.0
0.0
0.0
0.0
0.0
0.0
0.0
0.0
0.0
0.0
0.0
0.0
0.0
0.0
0.0
0.0
0.0
0.0
0.0
0.0
0.0
0.0
0.0
0.0
0.0
43.0
0.0
0.0
19.0

1.0
1.0
1.0
1.0
1.0
1.0
1.0
1.0
1.0
1.0
1.0
1.0
1.0
1.0
1.0
1.0
1.0
1.0
1.0
1.0
1.0
1.0
1.0
1.0
1.0
1.0
1.0
1.0
1.0
1.0


51.0
0.0
0.0
0.0
0.0
0.0
0.0
0.0
0.0
0.0
0.0
0.0
0.0
0.0
0.0
0.0
0.0
0.0
0.0
0.0
0.0
0.0
51.0
0.0
0.0
0.0
0.0
0.0
0.0
0.0


51.0
0.0
0.0
0.0
0.0
0.0
0.0
0.0
0.0
0.0
0.0
0.0
0.0
0.0
0.0
0.0
0.0
0.0
0.0
0.0
0.0
0.0
51.0
0.0
0.0
0.0
0.0
0.0
0.0
0.0

1.0
1.0
1.0
1.0
1.0
1.0
1.0
1.0
1.0
1.0
1.0
1.0
1.0
1.0
1.0
1.0
1.0
1.0
1.0
1.0
1.0
1.0
1.0
1.0
1.0
1.0
1.0
1.0
1.0
1.0


35.0
0.0
0.0
0.0
0.0
0.0
0.0
0.0
0.0
0.0
0.0
0.0
0.0
0.0
0.0
0.0
0.0
0.0
0.0
0.0
0.0
0.0
0.0
0.0
35.0
0.0
0.0
0.0
0.0
0.0


35.0
0.0
0.0
0.0
0.0
0.0
0.0
0.0
0.0
0.0
0.0
0.0
0.0
0.0
0.0
0.0
0.0
0.0
0.0
0.0
0.0
0.0
0.0
0.0
35.0
0.0
0.0
0.0
0.0
0.0

1.0
1.0
1.0
1.0
1.0
1.0
1.0
1.0
1.0
1.0
1.0
1.0
1.0
1.0
1.0
1.0
1.0
1.0
1.0
1.0
1.0
1.0
1.0
1.0
1.0
1.0
1.0
1.0
1.0
1.0


31.0
0.0
0.0
0.0
0.0
0.0
0.0
0.0
0.0
0.0
0.0
0.0
0.0
0.0
0.0
0.0
0.0
0.0
0.0
31.0
0.0
0.0
0.0
0.0
0.0
0.0
0.0
0.0
0.0
0.0


31.0
0.0
0.0
0.0
0.0
0.0
0.0
0.0
0.0
0.0
0.0
0.0
0.0
0.0
0.0
0.0
0.0
0.0
0.0
31.0
0.0
0.0
0.0
0.0
0.0
0.0
0.0
0.0
0.0
0.0

0.99
0.99
0.99
0.99
0.99
0.99
0.99
0.99
0.99
0.99
0.99
0.99
0.99
0.99
0.99
0.99
0.99
0.99
0.99
0.99
0.99
0.99
0.99
0.99
0.99
0.99
0.99
0.99
0.99
0.99


9.0
0.0
0.0
0.0
0.0
0.0
0.0
0.0
0.0
0.0
0.0
0.0
0.0
0.0
0.0
0.0
0.0
0.0
0.0
0.0
0.0
0.0
0.0
0.0
0.0
2.0
7.0
0.0
0.0
0.0

0.99
0.99
0.99
0.99
0.99
0.99
0.99
0.99
0.99
0.99
0.99
0.99
0.99
0.99
0.99
0.99
0.99
0.99
0.99
0.99
0.99
0.99
0.99
0.99
0.99
0.99
0.99
0.99
0.99
0.99


17779.0
28.0
7.0
0.0
0.0
0.0
216.0
0.0
0.0
0.0
0.0
0.0
0.0
0.0
0.0
0.0
0.0
17528.0
0.0
0.0
0.0
0.0
0.0
0.0
0.0
0.0
0.0
0.0
0.0
0.0


251.0
28.0
7.0
0.0
0.0
0.0
216.0
0.0
0.0
0.0
0.0
0.0
0.0
0.0
0.0
0.0
0.0
0.0
0.0
0.0
0.0
0.0
0.0
0.0
0.0
0.0
0.0
0.0
0.0
0.0

1.0
1.0
1.0
1.0
1.0
1.0
1.0
1.0
1.0
1.0
1.0
1.0
1.0
1.0
1.0
1.0
1.0
1.0
1.0
1.0
1.0
1.0
1.0
1.0
1.0
1.0
1.0
1.0
1.0
1.0


216.0
0.0
0.0
0.0
0.0
0.0
216.0
0.0
0.0
0.0
0.0
0.0
0.0
0.0
0.0
0.0
0.0
0.0
0.0
0.0
0.0
0.0
0.0
0.0
0.0
0.0
0.0
0.0
0.0
0.0

1.0
1.0
1.0
1.0
1.0
1.0
1.0
1.0
1.0
1.0
1.0
1.0
1.0
1.0
1.0
1.0
1.0
1.0
1.0
1.0
1.0
1.0
1.0
1.0
1.0
1.0
1.0
1.0
1.0
1.0


10.0
3.0
7.0
0.0
0.0
0.0
0.0
0.0
0.0
0.0
0.0
0.0
0.0
0.0
0.0
0.0
0.0
0.0
0.0
0.0
0.0
0.0
0.0
0.0
0.0
0.0
0.0
0.0
0.0
0.0

0.98
0.98
0.98
0.98
0.98
0.98
0.98
0.98
0.98
0.98
0.98
0.98
0.98
0.98
0.98
0.98
0.98
0.98
0.98
0.98
0.98
0.98
0.98
0.98
0.98
0.98
0.98
0.98
0.98
0.98


17528.0
0.0
0.0
0.0
0.0
0.0
0.0
0.0
0.0
0.0
0.0
0.0
0.0
0.0
0.0
0.0
0.0
17528.0
0.0
0.0
0.0
0.0
0.0
0.0
0.0
0.0
0.0
0.0
0.0
0.0


17528.0
0.0
0.0
0.0
0.0
0.0
0.0
0.0
0.0
0.0
0.0
0.0
0.0
0.0
0.0
0.0
0.0
17528.0
0.0
0.0
0.0
0.0
0.0
0.0
0.0
0.0
0.0
0.0
0.0
0.0

1.0
1.0
1.0
1.0
1.0
1.0
1.0
1.0
1.0
1.0
1.0
1.0
1.0
1.0
1.0
1.0
1.0
1.0
1.0
1.0
1.0
1.0
1.0
1.0
1.0
1.0
1.0
1.0
1.0
1.0


510.0
0.0
0.0
0.0
92.0
80.0
38.0
0.0
0.0
0.0
0.0
0.0
0.0
0.0
0.0
0.0
235.0
0.0
0.0
0.0
0.0
0.0
0.0
0.0
14.0
0.0
8.0
15.0
28.0
0.0


151.0
0.0
0.0
0.0
92.0
21.0
38.0
0.0
0.0
0.0
0.0
0.0
0.0
0.0
0.0
0.0
0.0
0.0
0.0
0.0
0.0
0.0
0.0
0.0
0.0
0.0
0.0
0.0
0.0
0.0

1.0
1.0
1.0
1.0
1.0
1.0
1.0
1.0
1.0
1.0
1.0
1.0
1.0
1.0
1.0
1.0
1.0
1.0
1.0
1.0
1.0
1.0
1.0
1.0
1.0
1.0
1.0
1.0
1.0
1.0


130.0
0.0
0.0
0.0
92.0
0.0
38.0
0.0
0.0
0.0
0.0
0.0
0.0
0.0
0.0
0.0
0.0
0.0
0.0
0.0
0.0
0.0
0.0
0.0
0.0
0.0
0.0
0.0
0.0
0.0

1.0
1.0
1.0
1.0
1.0
1.0
1.0
1.0
1.0
1.0
1.0
1.0
1.0
1.0
1.0
1.0
1.0
1.0
1.0
1.0
1.0
1.0
1.0
1.0
1.0
1.0
1.0
1.0
1.0
1.0


59.0
0.0
0.0
0.0
0.0
59.0
0.0
0.0
0.0
0.0
0.0
0.0
0.0
0.0
0.0
0.0
0.0
0.0
0.0
0.0
0.0
0.0
0.0
0.0
0.0
0.0
0.0
0.0
0.0
0.0


32.0
0.0
0.0
0.0
0.0
32.0
0.0
0.0
0.0
0.0
0.0
0.0
0.0
0.0
0.0
0.0
0.0
0.0
0.0
0.0
0.0
0.0
0.0
0.0
0.0
0.0
0.0
0.0
0.0
0.0

0.72
0.72
0.72
0.72
0.72
0.72
0.72
0.72
0.72
0.72
0.72
0.72
0.72
0.72
0.72
0.72
0.72
0.72
0.72
0.72
0.72
0.72
0.72
0.72
0.72
0.72
0.72
0.72
0.72
0.72


27.0
0.0
0.0
0.0
0.0
27.0
0.0
0.0
0.0
0.0
0.0
0.0
0.0
0.0
0.0
0.0
0.0
0.0
0.0
0.0
0.0
0.0
0.0
0.0
0.0
0.0
0.0
0.0
0.0
0.0

1.0
1.0
1.0
1.0
1.0
1.0
1.0
1.0
1.0
1.0
1.0
1.0
1.0
1.0
1.0
1.0
1.0
1.0
1.0
1.0
1.0
1.0
1.0
1.0
1.0
1.0
1.0
1.0
1.0
1.0


162.0
0.0
0.0
0.0
0.0
0.0
0.0
0.0
0.0
0.0
0.0
0.0
0.0
0.0
0.0
0.0
147.0
0.0
0.0
0.0
0.0
0.0
0.0
0.0
0.0
0.0
0.0
15.0
0.0
0.0

1.0
1.0
1.0
1.0
1.0
1.0
1.0
1.0
1.0
1.0
1.0
1.0
1.0
1.0
1.0
1.0
1.0
1.0
1.0
1.0
1.0
1.0
1.0
1.0
1.0
1.0
1.0
1.0
1.0
1.0


102.0
0.0
0.0
0.0
0.0
0.0
0.0
0.0
0.0
0.0
0.0
0.0
0.0
0.0
0.0
0.0
88.0
0.0
0.0
0.0
0.0
0.0
0.0
0.0
14.0
0.0
0.0
0.0
0.0
0.0


102.0
0.0
0.0
0.0
0.0
0.0
0.0
0.0
0.0
0.0
0.0
0.0
0.0
0.0
0.0
0.0
88.0
0.0
0.0
0.0
0.0
0.0
0.0
0.0
14.0
0.0
0.0
0.0
0.0
0.0

1.0
1.0
1.0
1.0
1.0
1.0
1.0
1.0
1.0
1.0
1.0
1.0
1.0
1.0
1.0
1.0
1.0
1.0
1.0
1.0
1.0
1.0
1.0
1.0
1.0
1.0
1.0
1.0
1.0
1.0


36.0
0.0
0.0
0.0
0.0
0.0
0.0
0.0
0.0
0.0
0.0
0.0
0.0
0.0
0.0
0.0
0.0
0.0
0.0
0.0
0.0
0.0
0.0
0.0
0.0
0.0
8.0
0.0
28.0
0.0


36.0
0.0
0.0
0.0
0.0
0.0
0.0
0.0
0.0
0.0
0.0
0.0
0.0
0.0
0.0
0.0
0.0
0.0
0.0
0.0
0.0
0.0
0.0
0.0
0.0
0.0
8.0
0.0
28.0
0.0

1.0
1.0
1.0
1.0
1.0
1.0
1.0
1.0
1.0
1.0
1.0
1.0
1.0
1.0
1.0
1.0
1.0
1.0
1.0
1.0
1.0
1.0
1.0
1.0
1.0
1.0
1.0
1.0
1.0
1.0


8985.0
0.0
39.0
16.0
0.0
0.0
99.0
0.0
0.0
0.0
0.0
10.0
0.0
0.0
0.0
0.0
2273.0
0.0
3920.0
0.0
0.0
0.0
0.0
0.0
0.0
197.0
0.0
1607.0
0.0
824.0


76.0
0.0
0.0
0.0
0.0
0.0
76.0
0.0
0.0
0.0
0.0
0.0
0.0
0.0
0.0
0.0
0.0
0.0
0.0
0.0
0.0
0.0
0.0
0.0
0.0
0.0
0.0
0.0
0.0
0.0

1.0
1.0
1.0
1.0
1.0
1.0
1.0
1.0
1.0
1.0
1.0
1.0
1.0
1.0
1.0
1.0
1.0
1.0
1.0
1.0
1.0
1.0
1.0
1.0
1.0
1.0
1.0
1.0
1.0
1.0


8909.0
0.0
39.0
16.0
0.0
0.0
23.0
0.0
0.0
0.0
0.0
10.0
0.0
0.0
0.0
0.0
2273.0
0.0
3920.0
0.0
0.0
0.0
0.0
0.0
0.0
197.0
0.0
1607.0
0.0
824.0

1.0
1.0
1.0
1.0
1.0
1.0
1.0
1.0
1.0
1.0
1.0
1.0
1.0
1.0
1.0
1.0
1.0
1.0
1.0
1.0
1.0
1.0
1.0
1.0
1.0
1.0
1.0
1.0
1.0
1.0


31.0
0.0
31.0
0.0
0.0
0.0
0.0
0.0
0.0
0.0
0.0
0.0
0.0
0.0
0.0
0.0
0.0
0.0
0.0
0.0
0.0
0.0
0.0
0.0
0.0
0.0
0.0
0.0
0.0
0.0

1.0
1.0
1.0
1.0
1.0
1.0
1.0
1.0
1.0
1.0
1.0
1.0
1.0
1.0
1.0
1.0
1.0
1.0
1.0
1.0
1.0
1.0
1.0
1.0
1.0
1.0
1.0
1.0
1.0
1.0


23.0
0.0
0.0
0.0
0.0
0.0
23.0
0.0
0.0
0.0
0.0
0.0
0.0
0.0
0.0
0.0
0.0
0.0
0.0
0.0
0.0
0.0
0.0
0.0
0.0
0.0
0.0
0.0
0.0
0.0

1.0
1.0
1.0
1.0
1.0
1.0
1.0
1.0
1.0
1.0
1.0
1.0
1.0
1.0
1.0
1.0
1.0
1.0
1.0
1.0
1.0
1.0
1.0
1.0
1.0
1.0
1.0
1.0
1.0
1.0


789.0
0.0
0.0
16.0
0.0
0.0
0.0
0.0
0.0
0.0
0.0
0.0
0.0
0.0
0.0
0.0
0.0
0.0
0.0
0.0
0.0
0.0
0.0
0.0
0.0
0.0
0.0
0.0
0.0
773.0

1.0
1.0
1.0
1.0
1.0
1.0
1.0
1.0
1.0
1.0
1.0
1.0
1.0
1.0
1.0
1.0
1.0
1.0
1.0
1.0
1.0
1.0
1.0
1.0
1.0
1.0
1.0
1.0
1.0
1.0


7976.0
0.0
8.0
0.0
0.0
0.0
0.0
0.0
0.0
0.0
0.0
0.0
0.0
0.0
0.0
0.0
2273.0
0.0
3920.0
0.0
0.0
0.0
0.0
0.0
0.0
197.0
0.0
1578.0
0.0
0.0

1.0
1.0
1.0
1.0
1.0
1.0
1.0
1.0
1.0
1.0
1.0
1.0
1.0
1.0
1.0
1.0
1.0
1.0
1.0
1.0
1.0
1.0
1.0
1.0
1.0
1.0
1.0
1.0
1.0
1.0


5572.0
13.0
9.0
50.0
32.0
46.0
25.0
0.0
0.0
2.0
11.0
5.0
0.0
0.0
27.0
62.0
34.0
0.0
14.0
0.0
0.0
0.0
0.0
0.0
945.0
695.0
134.0
2625.0
464.0
379.0

1.0
1.0
1.0
1.0
1.0
1.0
1.0
1.0
1.0
1.0
1.0
1.0
1.0
1.0
1.0
1.0
1.0
1.0
1.0
1.0
1.0
1.0
1.0
1.0
1.0
1.0
1.0
1.0
1.0
1.0


580.0
7.0
1.0
32.0
21.0
24.0
9.0
0.0
0.0
0.0
0.0
0.0
0.0
0.0
27.0
0.0
0.0
0.0
0.0
0.0
0.0
0.0
0.0
0.0
410.0
0.0
45.0
0.0
4.0
0.0

1.0
1.0
1.0
1.0
1.0
1.0
1.0
1.0
1.0
1.0
1.0
1.0
1.0
1.0
1.0
1.0
1.0
1.0
1.0
1.0
1.0
1.0
1.0
1.0
1.0
1.0
1.0
1.0
1.0
1.0


36.0
5.0
0.0
4.0
0.0
0.0
0.0
0.0
0.0
0.0
0.0
0.0
0.0
0.0
27.0
0.0
0.0
0.0
0.0
0.0
0.0
0.0
0.0
0.0
0.0
0.0
0.0
0.0
0.0
0.0

1.0
1.0
1.0
1.0
1.0
1.0
1.0
1.0
1.0
1.0
1.0
1.0
1.0
1.0
1.0
1.0
1.0
1.0
1.0
1.0
1.0
1.0
1.0
1.0
1.0
1.0
1.0
1.0
1.0
1.0


575.0
6.0
8.0
3.0
0.0
17.0
9.0
0.0
0.0
2.0
11.0
5.0
0.0
0.0
0.0
62.0
34.0
0.0
14.0
0.0
0.0
0.0
0.0
0.0
215.0
0.0
5.0
24.0
46.0
114.0

1.0
1.0
1.0
1.0
1.0
1.0
1.0
1.0
1.0
1.0
1.0
1.0
1.0
1.0
1.0
1.0
1.0
1.0
1.0
1.0
1.0
1.0
1.0
1.0
1.0
1.0
1.0
1.0
1.0
1.0


3938.0
0.0
0.0
0.0
2.0
2.0
7.0
0.0
0.0
0.0
0.0
0.0
0.0
0.0
0.0
0.0
0.0
0.0
0.0
0.0
0.0
0.0
0.0
0.0
316.0
633.0
55.0
2324.0
391.0
208.0

1.0
1.0
1.0
1.0
1.0
1.0
1.0
1.0
1.0
1.0
1.0
1.0
1.0
1.0
1.0
1.0
1.0
1.0
1.0
1.0
1.0
1.0
1.0
1.0
1.0
1.0
1.0
1.0
1.0
1.0


129.0
0.0
0.0
0.0
0.0
0.0
0.0
0.0
0.0
0.0
0.0
0.0
0.0
0.0
0.0
0.0
0.0
0.0
0.0
0.0
0.0
0.0
0.0
0.0
0.0
129.0
0.0
0.0
0.0
0.0

1.0
1.0
1.0
1.0
1.0
1.0
1.0
1.0
1.0
1.0
1.0
1.0
1.0
1.0
1.0
1.0
1.0
1.0
1.0
1.0
1.0
1.0
1.0
1.0
1.0
1.0
1.0
1.0
1.0
1.0


364.0
0.0
0.0
0.0
0.0
0.0
0.0
0.0
0.0
0.0
0.0
0.0
0.0
0.0
0.0
0.0
0.0
0.0
0.0
0.0
0.0
0.0
0.0
0.0
0.0
33.0
0.0
264.0
12.0
55.0

0.84
0.84
0.84
0.84
0.84
0.84
0.84
0.84
0.84
0.84
0.84
0.84
0.84
0.84
0.84
0.84
0.84
0.84
0.84
0.84
0.84
0.84
0.84
0.84
0.84
0.84
0.84
0.84
0.84
0.84


31.0
0.0
0.0
0.0
0.0
0.0
0.0
0.0
0.0
0.0
0.0
0.0
0.0
0.0
0.0
0.0
0.0
0.0
0.0
0.0
0.0
0.0
0.0
0.0
0.0
29.0
0.0
0.0
0.0
2.0


31.0
0.0
0.0
0.0
0.0
0.0
0.0
0.0
0.0
0.0
0.0
0.0
0.0
0.0
0.0
0.0
0.0
0.0
0.0
0.0
0.0
0.0
0.0
0.0
0.0
29.0
0.0
0.0
0.0
2.0

1.0
1.0
1.0
1.0
1.0
1.0
1.0
1.0
1.0
1.0
1.0
1.0
1.0
1.0
1.0
1.0
1.0
1.0
1.0
1.0
1.0
1.0
1.0
1.0
1.0
1.0
1.0
1.0
1.0
1.0


56.0
0.0
0.0
0.0
0.0
0.0
0.0
0.0
0.0
0.0
0.0
56.0
0.0
0.0
0.0
0.0
0.0
0.0
0.0
0.0
0.0
0.0
0.0
0.0
0.0
0.0
0.0
0.0
0.0
0.0


56.0
0.0
0.0
0.0
0.0
0.0
0.0
0.0
0.0
0.0
0.0
56.0
0.0
0.0
0.0
0.0
0.0
0.0
0.0
0.0
0.0
0.0
0.0
0.0
0.0
0.0
0.0
0.0
0.0
0.0


56.0
0.0
0.0
0.0
0.0
0.0
0.0
0.0
0.0
0.0
0.0
56.0
0.0
0.0
0.0
0.0
0.0
0.0
0.0
0.0
0.0
0.0
0.0
0.0
0.0
0.0
0.0
0.0
0.0
0.0

1.0
1.0
1.0
1.0
1.0
1.0
1.0
1.0
1.0
1.0
1.0
1.0
1.0
1.0
1.0
1.0
1.0
1.0
1.0
1.0
1.0
1.0
1.0
1.0
1.0
1.0
1.0
1.0
1.0
1.0


48.0
0.0
2.0
0.0
18.0
0.0
8.0
0.0
0.0
0.0
0.0
20.0
0.0
0.0
0.0
0.0
0.0
0.0
0.0
0.0
0.0
0.0
0.0
0.0
0.0
0.0
0.0
0.0
0.0
0.0


48.0
0.0
2.0
0.0
18.0
0.0
8.0
0.0
0.0
0.0
0.0
20.0
0.0
0.0
0.0
0.0
0.0
0.0
0.0
0.0
0.0
0.0
0.0
0.0
0.0
0.0
0.0
0.0
0.0
0.0


48.0
0.0
2.0
0.0
18.0
0.0
8.0
0.0
0.0
0.0
0.0
20.0
0.0
0.0
0.0
0.0
0.0
0.0
0.0
0.0
0.0
0.0
0.0
0.0
0.0
0.0
0.0
0.0
0.0
0.0

1.0
1.0
1.0
1.0
1.0
1.0
1.0
1.0
1.0
1.0
1.0
1.0
1.0
1.0
1.0
1.0
1.0
1.0
1.0
1.0
1.0
1.0
1.0
1.0
1.0
1.0
1.0
1.0
1.0
1.0


58.0
1.0
1.0
3.0
0.0
0.0
0.0
17.0
9.0
11.0
12.0
4.0
0.0
0.0
0.0
0.0
0.0
0.0
0.0
0.0
0.0
0.0
0.0
0.0
0.0
0.0
0.0
0.0
0.0
0.0

0.96
0.96
0.96
0.96
0.96
0.96
0.96
0.96
0.96
0.96
0.96
0.96
0.96
0.96
0.96
0.96
0.96
0.96
0.96
0.96
0.96
0.96
0.96
0.96
0.96
0.96
0.96
0.96
0.96
0.96


41.0
1.0
1.0
3.0
0.0
0.0
0.0
0.0
9.0
11.0
12.0
4.0
0.0
0.0
0.0
0.0
0.0
0.0
0.0
0.0
0.0
0.0
0.0
0.0
0.0
0.0
0.0
0.0
0.0
0.0

1.0
1.0
1.0
1.0
1.0
1.0
1.0
1.0
1.0
1.0
1.0
1.0
1.0
1.0
1.0
1.0
1.0
1.0
1.0
1.0
1.0
1.0
1.0
1.0
1.0
1.0
1.0
1.0
1.0
1.0


2480.0
0.0
2.0
0.0
0.0
37.0
39.0
0.0
0.0
0.0
0.0
0.0
0.0
0.0
0.0
383.0
0.0
63.0
451.0
789.0
0.0
0.0
0.0
0.0
5.0
276.0
264.0
34.0
6.0
131.0

0.98
0.98
0.98
0.98
0.98
0.98
0.98
0.98
0.98
0.98
0.98
0.98
0.98
0.98
0.98
0.98
0.98
0.98
0.98
0.98
0.98
0.98
0.98
0.98
0.98
0.98
0.98
0.98
0.98
0.98


318.0
0.0
0.0
0.0
0.0
25.0
39.0
0.0
0.0
0.0
0.0
0.0
0.0
0.0
0.0
37.0
0.0
0.0
0.0
217.0
0.0
0.0
0.0
0.0
0.0
0.0
0.0
0.0
0.0
0.0

1.0
1.0
1.0
1.0
1.0
1.0
1.0
1.0
1.0
1.0
1.0
1.0
1.0
1.0
1.0
1.0
1.0
1.0
1.0
1.0
1.0
1.0
1.0
1.0
1.0
1.0
1.0
1.0
1.0
1.0


183.0
0.0
2.0
0.0
0.0
12.0
0.0
0.0
0.0
0.0
0.0
0.0
0.0
0.0
0.0
169.0
0.0
0.0
0.0
0.0
0.0
0.0
0.0
0.0
0.0
0.0
0.0
0.0
0.0
0.0

1.0
1.0
1.0
1.0
1.0
1.0
1.0
1.0
1.0
1.0
1.0
1.0
1.0
1.0
1.0
1.0
1.0
1.0
1.0
1.0
1.0
1.0
1.0
1.0
1.0
1.0
1.0
1.0
1.0
1.0


14.0
0.0
2.0
0.0
0.0
12.0
0.0
0.0
0.0
0.0
0.0
0.0
0.0
0.0
0.0
0.0
0.0
0.0
0.0
0.0
0.0
0.0
0.0
0.0
0.0
0.0
0.0
0.0
0.0
0.0

1.0
1.0
1.0
1.0
1.0
1.0
1.0
1.0
1.0
1.0
1.0
1.0
1.0
1.0
1.0
1.0
1.0
1.0
1.0
1.0
1.0
1.0
1.0
1.0
1.0
1.0
1.0
1.0
1.0
1.0


134.0
0.0
0.0
0.0
0.0
0.0
0.0
0.0
0.0
0.0
0.0
0.0
0.0
0.0
0.0
0.0
0.0
0.0
0.0
130.0
0.0
0.0
0.0
0.0
0.0
0.0
0.0
0.0
0.0
4.0

0.8
0.8
0.8
0.8
0.8
0.8
0.8
0.8
0.8
0.8
0.8
0.8
0.8
0.8
0.8
0.8
0.8
0.8
0.8
0.8
0.8
0.8
0.8
0.8
0.8
0.8
0.8
0.8
0.8
0.8


4.0
0.0
0.0
0.0
0.0
0.0
0.0
0.0
0.0
0.0
0.0
0.0
0.0
0.0
0.0
0.0
0.0
0.0
0.0
0.0
0.0
0.0
0.0
0.0
0.0
0.0
0.0
0.0
0.0
4.0

1.0
1.0
1.0
1.0
1.0
1.0
1.0
1.0
1.0
1.0
1.0
1.0
1.0
1.0
1.0
1.0
1.0
1.0
1.0
1.0
1.0
1.0
1.0
1.0
1.0
1.0
1.0
1.0
1.0
1.0


50.0
0.0
0.0
0.0
0.0
0.0
0.0
20.0
4.0
8.0
14.0
4.0
0.0
0.0
0.0
0.0
0.0
0.0
0.0
0.0
0.0
0.0
0.0
0.0
0.0
0.0
0.0
0.0
0.0
0.0


21.0
0.0
0.0
0.0
0.0
0.0
0.0
20.0
0.0
0.0
1.0
0.0
0.0
0.0
0.0
0.0
0.0
0.0
0.0
0.0
0.0
0.0
0.0
0.0
0.0
0.0
0.0
0.0
0.0
0.0

0.98
0.98
0.98
0.98
0.98
0.98
0.98
0.98
0.98
0.98
0.98
0.98
0.98
0.98
0.98
0.98
0.98
0.98
0.98
0.98
0.98
0.98
0.98
0.98
0.98
0.98
0.98
0.98
0.98
0.98


29.0
0.0
0.0
0.0
0.0
0.0
0.0
0.0
4.0
8.0
13.0
4.0
0.0
0.0
0.0
0.0
0.0
0.0
0.0
0.0
0.0
0.0
0.0
0.0
0.0
0.0
0.0
0.0
0.0
0.0


29.0
0.0
0.0
0.0
0.0
0.0
0.0
0.0
4.0
8.0
13.0
4.0
0.0
0.0
0.0
0.0
0.0
0.0
0.0
0.0
0.0
0.0
0.0
0.0
0.0
0.0
0.0
0.0
0.0
0.0

1.0
1.0
1.0
1.0
1.0
1.0
1.0
1.0
1.0
1.0
1.0
1.0
1.0
1.0
1.0
1.0
1.0
1.0
1.0
1.0
1.0
1.0
1.0
1.0
1.0
1.0
1.0
1.0
1.0
1.0


138994.0
0.0
0.0
17.0
0.0
0.0
0.0
0.0
0.0
0.0
0.0
0.0
0.0
0.0
0.0
0.0
0.0
0.0
0.0
0.0
0.0
0.0
0.0
0.0
138977.0
0.0
0.0
0.0
0.0
0.0

0.72
0.72
0.72
0.72
0.72
0.72
0.72
0.72
0.72
0.72
0.72
0.72
0.72
0.72
0.72
0.72
0.72
0.72
0.72
0.72
0.72
0.72
0.72
0.72
0.72
0.72
0.72
0.72
0.72
0.72


124878.0
0.0
0.0
17.0
0.0
0.0
0.0
0.0
0.0
0.0
0.0
0.0
0.0
0.0
0.0
0.0
0.0
0.0
0.0
0.0
0.0
0.0
0.0
0.0
124861.0
0.0
0.0
0.0
0.0
0.0


124878.0
0.0
0.0
17.0
0.0
0.0
0.0
0.0
0.0
0.0
0.0
0.0
0.0
0.0
0.0
0.0
0.0
0.0
0.0
0.0
0.0
0.0
0.0
0.0
124861.0
0.0
0.0
0.0
0.0
0.0

1.0
1.0
1.0
1.0
1.0
1.0
1.0
1.0
1.0
1.0
1.0
1.0
1.0
1.0
1.0
1.0
1.0
1.0
1.0
1.0
1.0
1.0
1.0
1.0
1.0
1.0
1.0
1.0
1.0
1.0


13851.0
0.0
0.0
0.0
0.0
0.0
0.0
0.0
0.0
0.0
0.0
0.0
0.0
0.0
0.0
0.0
0.0
0.0
0.0
0.0
0.0
0.0
0.0
0.0
13851.0
0.0
0.0
0.0
0.0
0.0


13851.0
0.0
0.0
0.0
0.0
0.0
0.0
0.0
0.0
0.0
0.0
0.0
0.0
0.0
0.0
0.0
0.0
0.0
0.0
0.0
0.0
0.0
0.0
0.0
13851.0
0.0
0.0
0.0
0.0
0.0

1.0
1.0
1.0
1.0
1.0
1.0
1.0
1.0
1.0
1.0
1.0
1.0
1.0
1.0
1.0
1.0
1.0
1.0
1.0
1.0
1.0
1.0
1.0
1.0
1.0
1.0
1.0
1.0
1.0
1.0


5063.0
0.0
0.0
0.0
0.0
0.0
11.0
0.0
0.0
0.0
0.0
0.0
0.0
0.0
0.0
0.0
0.0
0.0
0.0
0.0
0.0
0.0
0.0
0.0
1016.0
498.0
127.0
2226.0
900.0
285.0

0.83
0.83
0.83
0.83
0.83
0.83
0.83
0.83
0.83
0.83
0.83
0.83
0.83
0.83
0.83
0.83
0.83
0.83
0.83
0.83
0.83
0.83
0.83
0.83
0.83
0.83
0.83
0.83
0.83
0.83


3883.0
0.0
0.0
0.0
0.0
0.0
0.0
0.0
0.0
0.0
0.0
0.0
0.0
0.0
0.0
0.0
0.0
0.0
0.0
0.0
0.0
0.0
0.0
0.0
0.0
498.0
127.0
2226.0
747.0
285.0


3883.0
0.0
0.0
0.0
0.0
0.0
0.0
0.0
0.0
0.0
0.0
0.0
0.0
0.0
0.0
0.0
0.0
0.0
0.0
0.0
0.0
0.0
0.0
0.0
0.0
498.0
127.0
2226.0
747.0
285.0

1.0
1.0
1.0
1.0
1.0
1.0
1.0
1.0
1.0
1.0
1.0
1.0
1.0
1.0
1.0
1.0
1.0
1.0
1.0
1.0
1.0
1.0
1.0
1.0
1.0
1.0
1.0
1.0
1.0
1.0


11.0
0.0
0.0
0.0
0.0
0.0
0.0
0.0
0.0
0.0
0.0
0.0
0.0
0.0
0.0
0.0
0.0
0.0
0.0
0.0
0.0
0.0
0.0
0.0
11.0
0.0
0.0
0.0
0.0
0.0

0.99
0.99
0.99
0.99
0.99
0.99
0.99
0.99
0.99
0.99
0.99
0.99
0.99
0.99
0.99
0.99
0.99
0.99
0.99
0.99
0.99
0.99
0.99
0.99
0.99
0.99
0.99
0.99
0.99
0.99


3336.0
5.0
0.0
0.0
0.0
10.0
0.0
2.0
1.0
8.0
0.0
10.0
0.0
0.0
0.0
61.0
0.0
0.0
6.0
0.0
0.0
0.0
0.0
0.0
567.0
23.0
232.0
0.0
655.0
1756.0

1.0
1.0
1.0
1.0
1.0
1.0
1.0
1.0
1.0
1.0
1.0
1.0
1.0
1.0
1.0
1.0
1.0
1.0
1.0
1.0
1.0
1.0
1.0
1.0
1.0
1.0
1.0
1.0
1.0
1.0


1138.0
5.0
0.0
0.0
0.0
10.0
0.0
2.0
1.0
0.0
0.0
10.0
0.0
0.0
0.0
0.0
0.0
0.0
0.0
0.0
0.0
0.0
0.0
0.0
0.0
0.0
150.0
0.0
539.0
421.0

1.0
1.0
1.0
1.0
1.0
1.0
1.0
1.0
1.0
1.0
1.0
1.0
1.0
1.0
1.0
1.0
1.0
1.0
1.0
1.0
1.0
1.0
1.0
1.0
1.0
1.0
1.0
1.0
1.0
1.0


8.0
5.0
0.0
0.0
0.0
0.0
0.0
2.0
1.0
0.0
0.0
0.0
0.0
0.0
0.0
0.0
0.0
0.0
0.0
0.0
0.0
0.0
0.0
0.0
0.0
0.0
0.0
0.0
0.0
0.0

1.0
1.0
1.0
1.0
1.0
1.0
1.0
1.0
1.0
1.0
1.0
1.0
1.0
1.0
1.0
1.0
1.0
1.0
1.0
1.0
1.0
1.0
1.0
1.0
1.0
1.0
1.0
1.0
1.0
1.0


104.0
0.0
0.0
0.0
0.0
0.0
0.0
0.0
0.0
0.0
0.0
0.0
0.0
0.0
0.0
0.0
0.0
0.0
0.0
0.0
0.0
0.0
0.0
0.0
0.0
0.0
92.0
0.0
12.0
0.0

0.8
0.8
0.8
0.8
0.8
0.8
0.8
0.8
0.8
0.8
0.8
0.8
0.8
0.8
0.8
0.8
0.8
0.8
0.8
0.8
0.8
0.8
0.8
0.8
0.8
0.8
0.8
0.8
0.8
0.8


369.0
0.0
0.0
0.0
0.0
0.0
0.0
0.0
0.0
0.0
0.0
0.0
0.0
0.0
0.0
61.0
0.0
0.0
0.0
0.0
0.0
0.0
0.0
0.0
0.0
0.0
82.0
0.0
94.0
132.0


369.0
0.0
0.0
0.0
0.0
0.0
0.0
0.0
0.0
0.0
0.0
0.0
0.0
0.0
0.0
61.0
0.0
0.0
0.0
0.0
0.0
0.0
0.0
0.0
0.0
0.0
82.0
0.0
94.0
132.0

1.0
1.0
1.0
1.0
1.0
1.0
1.0
1.0
1.0
1.0
1.0
1.0
1.0
1.0
1.0
1.0
1.0
1.0
1.0
1.0
1.0
1.0
1.0
1.0
1.0
1.0
1.0
1.0
1.0
1.0


474.0
0.0
0.0
0.0
0.0
0.0
0.0
0.0
0.0
0.0
0.0
0.0
0.0
0.0
0.0
0.0
0.0
0.0
0.0
0.0
0.0
0.0
0.0
0.0
474.0
0.0
0.0
0.0
0.0
0.0


474.0
0.0
0.0
0.0
0.0
0.0
0.0
0.0
0.0
0.0
0.0
0.0
0.0
0.0
0.0
0.0
0.0
0.0
0.0
0.0
0.0
0.0
0.0
0.0
474.0
0.0
0.0
0.0
0.0
0.0

1.0
1.0
1.0
1.0
1.0
1.0
1.0
1.0
1.0
1.0
1.0
1.0
1.0
1.0
1.0
1.0
1.0
1.0
1.0
1.0
1.0
1.0
1.0
1.0
1.0
1.0
1.0
1.0
1.0
1.0


42.0
0.0
0.0
0.0
0.0
0.0
0.0
0.0
0.0
0.0
0.0
0.0
0.0
0.0
0.0
0.0
0.0
0.0
0.0
0.0
0.0
0.0
0.0
0.0
42.0
0.0
0.0
0.0
0.0
0.0


42.0
0.0
0.0
0.0
0.0
0.0
0.0
0.0
0.0
0.0
0.0
0.0
0.0
0.0
0.0
0.0
0.0
0.0
0.0
0.0
0.0
0.0
0.0
0.0
42.0
0.0
0.0
0.0
0.0
0.0

0.86
0.86
0.86
0.86
0.86
0.86
0.86
0.86
0.86
0.86
0.86
0.86
0.86
0.86
0.86
0.86
0.86
0.86
0.86
0.86
0.86
0.86
0.86
0.86
0.86
0.86
0.86
0.86
0.86
0.86


4177.0
0.0
0.0
0.0
0.0
0.0
10.0
0.0
0.0
0.0
0.0
0.0
0.0
0.0
0.0
0.0
0.0
0.0
71.0
2953.0
0.0
0.0
0.0
0.0
81.0
664.0
18.0
57.0
0.0
323.0

0.71
0.71
0.71
0.71
0.71
0.71
0.71
0.71
0.71
0.71
0.71
0.71
0.71
0.71
0.71
0.71
0.71
0.71
0.71
0.71
0.71
0.71
0.71
0.71
0.71
0.71
0.71
0.71
0.71
0.71


22.0
0.0
0.0
0.0
0.0
0.0
10.0
0.0
0.0
0.0
0.0
0.0
0.0
0.0
0.0
0.0
0.0
0.0
0.0
0.0
0.0
0.0
0.0
0.0
0.0
0.0
0.0
12.0
0.0
0.0


10.0
0.0
0.0
0.0
0.0
0.0
10.0
0.0
0.0
0.0
0.0
0.0
0.0
0.0
0.0
0.0
0.0
0.0
0.0
0.0
0.0
0.0
0.0
0.0
0.0
0.0
0.0
0.0
0.0
0.0

0.92
0.92
0.92
0.92
0.92
0.92
0.92
0.92
0.92
0.92
0.92
0.92
0.92
0.92
0.92
0.92
0.92
0.92
0.92
0.92
0.92
0.92
0.92
0.92
0.92
0.92
0.92
0.92
0.92
0.92


12.0
0.0
0.0
0.0
0.0
0.0
0.0
0.0
0.0
0.0
0.0
0.0
0.0
0.0
0.0
0.0
0.0
0.0
0.0
0.0
0.0
0.0
0.0
0.0
0.0
0.0
0.0
12.0
0.0
0.0

0.85
0.85
0.85
0.85
0.85
0.85
0.85
0.85
0.85
0.85
0.85
0.85
0.85
0.85
0.85
0.85
0.85
0.85
0.85
0.85
0.85
0.85
0.85
0.85
0.85
0.85
0.85
0.85
0.85
0.85


3896.0
0.0
0.0
0.0
0.0
0.0
0.0
0.0
0.0
0.0
0.0
0.0
0.0
0.0
0.0
0.0
0.0
0.0
0.0
2953.0
0.0
0.0
0.0
0.0
81.0
625.0
0.0
45.0
0.0
192.0

0.94
0.94
0.94
0.94
0.94
0.94
0.94
0.94
0.94
0.94
0.94
0.94
0.94
0.94
0.94
0.94
0.94
0.94
0.94
0.94
0.94
0.94
0.94
0.94
0.94
0.94
0.94
0.94
0.94
0.94


81.0
0.0
0.0
0.0
0.0
0.0
0.0
0.0
0.0
0.0
0.0
0.0
0.0
0.0
0.0
0.0
0.0
0.0
0.0
0.0
0.0
0.0
0.0
0.0
81.0
0.0
0.0
0.0
0.0
0.0

0.97
0.97
0.97
0.97
0.97
0.97
0.97
0.97
0.97
0.97
0.97
0.97
0.97
0.97
0.97
0.97
0.97
0.97
0.97
0.97
0.97
0.97
0.97
0.97
0.97
0.97
0.97
0.97
0.97
0.97


220.0
0.0
0.0
0.0
0.0
0.0
0.0
0.0
0.0
0.0
0.0
0.0
0.0
0.0
0.0
0.0
0.0
0.0
71.0
0.0
0.0
0.0
0.0
0.0
0.0
0.0
18.0
0.0
0.0
131.0


220.0
0.0
0.0
0.0
0.0
0.0
0.0
0.0
0.0
0.0
0.0
0.0
0.0
0.0
0.0
0.0
0.0
0.0
71.0
0.0
0.0
0.0
0.0
0.0
0.0
0.0
18.0
0.0
0.0
131.0

1.0
1.0
1.0
1.0
1.0
1.0
1.0
1.0
1.0
1.0
1.0
1.0
1.0
1.0
1.0
1.0
1.0
1.0
1.0
1.0
1.0
1.0
1.0
1.0
1.0
1.0
1.0
1.0
1.0
1.0


5657.0
0.0
0.0
0.0
2.0
6.0
9.0
0.0
0.0
0.0
0.0
0.0
0.0
0.0
148.0
3.0
34.0
175.0
13.0
4.0
0.0
0.0
0.0
0.0
2417.0
268.0
319.0
141.0
1884.0
234.0

1.0
1.0
1.0
1.0
1.0
1.0
1.0
1.0
1.0
1.0
1.0
1.0
1.0
1.0
1.0
1.0
1.0
1.0
1.0
1.0
1.0
1.0
1.0
1.0
1.0
1.0
1.0
1.0
1.0
1.0


1419.0
0.0
0.0
0.0
2.0
6.0
9.0
0.0
0.0
0.0
0.0
0.0
0.0
0.0
0.0
0.0
0.0
0.0
0.0
0.0
0.0
0.0
0.0
0.0
7.0
0.0
0.0
0.0
1387.0
8.0

1.0
1.0
1.0
1.0
1.0
1.0
1.0
1.0
1.0
1.0
1.0
1.0
1.0
1.0
1.0
1.0
1.0
1.0
1.0
1.0
1.0
1.0
1.0
1.0
1.0
1.0
1.0
1.0
1.0
1.0


1234.0
0.0
0.0
0.0
0.0
0.0
0.0
0.0
0.0
0.0
0.0
0.0
0.0
0.0
0.0
0.0
4.0
0.0
0.0
0.0
0.0
0.0
0.0
0.0
815.0
0.0
0.0
24.0
391.0
0.0

0.99
0.99
0.99
0.99
0.99
0.99
0.99
0.99
0.99
0.99
0.99
0.99
0.99
0.99
0.99
0.99
0.99
0.99
0.99
0.99
0.99
0.99
0.99
0.99
0.99
0.99
0.99
0.99
0.99
0.99


233.0
0.0
0.0
0.0
0.0
0.0
0.0
0.0
0.0
0.0
0.0
0.0
0.0
0.0
0.0
0.0
0.0
0.0
0.0
0.0
0.0
0.0
0.0
0.0
0.0
25.0
208.0
0.0
0.0
0.0

1.0
1.0
1.0
1.0
1.0
1.0
1.0
1.0
1.0
1.0
1.0
1.0
1.0
1.0
1.0
1.0
1.0
1.0
1.0
1.0
1.0
1.0
1.0
1.0
1.0
1.0
1.0
1.0
1.0
1.0


175.0
0.0
0.0
0.0
0.0
0.0
0.0
0.0
0.0
0.0
0.0
0.0
0.0
0.0
0.0
0.0
0.0
175.0
0.0
0.0
0.0
0.0
0.0
0.0
0.0
0.0
0.0
0.0
0.0
0.0


175.0
0.0
0.0
0.0
0.0
0.0
0.0
0.0
0.0
0.0
0.0
0.0
0.0
0.0
0.0
0.0
0.0
175.0
0.0
0.0
0.0
0.0
0.0
0.0
0.0
0.0
0.0
0.0
0.0
0.0

1.0
1.0
1.0
1.0
1.0
1.0
1.0
1.0
1.0
1.0
1.0
1.0
1.0
1.0
1.0
1.0
1.0
1.0
1.0
1.0
1.0
1.0
1.0
1.0
1.0
1.0
1.0
1.0
1.0
1.0


148.0
0.0
0.0
0.0
0.0
0.0
0.0
0.0
0.0
0.0
0.0
0.0
0.0
0.0
148.0
0.0
0.0
0.0
0.0
0.0
0.0
0.0
0.0
0.0
0.0
0.0
0.0
0.0
0.0
0.0


148.0
0.0
0.0
0.0
0.0
0.0
0.0
0.0
0.0
0.0
0.0
0.0
0.0
0.0
148.0
0.0
0.0
0.0
0.0
0.0
0.0
0.0
0.0
0.0
0.0
0.0
0.0
0.0
0.0
0.0

1.0
1.0
1.0
1.0
1.0
1.0
1.0
1.0
1.0
1.0
1.0
1.0
1.0
1.0
1.0
1.0
1.0
1.0
1.0
1.0
1.0
1.0
1.0
1.0
1.0
1.0
1.0
1.0
1.0
1.0


54.0
0.0
0.0
0.0
0.0
0.0
0.0
0.0
0.0
0.0
0.0
0.0
0.0
0.0
0.0
0.0
0.0
0.0
0.0
0.0
0.0
0.0
0.0
0.0
23.0
0.0
4.0
0.0
0.0
27.0

0.79
0.79
0.79
0.79
0.79
0.79
0.79
0.79
0.79
0.79
0.79
0.79
0.79
0.79
0.79
0.79
0.79
0.79
0.79
0.79
0.79
0.79
0.79
0.79
0.79
0.79
0.79
0.79
0.79
0.79


32.0
0.0
0.0
0.0
0.0
0.0
0.0
0.0
0.0
0.0
0.0
0.0
0.0
0.0
0.0
0.0
0.0
0.0
0.0
0.0
0.0
0.0
0.0
0.0
0.0
0.0
0.0
0.0
0.0
32.0

0.71
0.71
0.71
0.71
0.71
0.71
0.71
0.71
0.71
0.71
0.71
0.71
0.71
0.71
0.71
0.71
0.71
0.71
0.71
0.71
0.71
0.71
0.71
0.71
0.71
0.71
0.71
0.71
0.71
0.71


14.0
0.0
0.0
0.0
0.0
0.0
0.0
0.0
0.0
0.0
0.0
0.0
0.0
0.0
0.0
0.0
0.0
0.0
0.0
0.0
0.0
0.0
0.0
0.0
0.0
14.0
0.0
0.0
0.0
0.0


14.0
0.0
0.0
0.0
0.0
0.0
0.0
0.0
0.0
0.0
0.0
0.0
0.0
0.0
0.0
0.0
0.0
0.0
0.0
0.0
0.0
0.0
0.0
0.0
0.0
14.0
0.0
0.0
0.0
0.0

1.0
1.0
1.0
1.0
1.0
1.0
1.0
1.0
1.0
1.0
1.0
1.0
1.0
1.0
1.0
1.0
1.0
1.0
1.0
1.0
1.0
1.0
1.0
1.0
1.0
1.0
1.0
1.0
1.0
1.0


8.0
0.0
0.0
0.0
0.0
0.0
0.0
0.0
0.0
0.0
0.0
0.0
0.0
0.0
0.0
0.0
0.0
0.0
0.0
0.0
0.0
0.0
0.0
0.0
8.0
0.0
0.0
0.0
0.0
0.0


8.0
0.0
0.0
0.0
0.0
0.0
0.0
0.0
0.0
0.0
0.0
0.0
0.0
0.0
0.0
0.0
0.0
0.0
0.0
0.0
0.0
0.0
0.0
0.0
8.0
0.0
0.0
0.0
0.0
0.0

1.0
1.0
1.0
1.0
1.0
1.0
1.0
1.0
1.0
1.0
1.0
1.0
1.0
1.0
1.0
1.0
1.0
1.0
1.0
1.0
1.0
1.0
1.0
1.0
1.0
1.0
1.0
1.0
1.0
1.0


37177.0
0.0
0.0
0.0
0.0
0.0
8.0
0.0
0.0
0.0
0.0
0.0
0.0
80.0
0.0
825.0
335.0
0.0
8674.0
25779.0
0.0
128.0
171.0
0.0
253.0
156.0
547.0
9.0
168.0
44.0


35270.0
0.0
0.0
0.0
0.0
0.0
8.0
0.0
0.0
0.0
0.0
0.0
0.0
80.0
0.0
0.0
0.0
0.0
8674.0
25779.0
0.0
128.0
171.0
0.0
193.0
4.0
29.0
9.0
156.0
39.0

1.0
1.0
1.0
1.0
1.0
1.0
1.0
1.0
1.0
1.0
1.0
1.0
1.0
1.0
1.0
1.0
1.0
1.0
1.0
1.0
1.0
1.0
1.0
1.0
1.0
1.0
1.0
1.0
1.0
1.0


239.0
0.0
0.0
0.0
0.0
0.0
8.0
0.0
0.0
0.0
0.0
0.0
0.0
0.0
0.0
0.0
0.0
0.0
0.0
0.0
0.0
0.0
0.0
0.0
147.0
4.0
29.0
9.0
3.0
39.0

0.97
0.97
0.97
0.97
0.97
0.97
0.97
0.97
0.97
0.97
0.97
0.97
0.97
0.97
0.97
0.97
0.97
0.97
0.97
0.97
0.97
0.97
0.97
0.97
0.97
0.97
0.97
0.97
0.97
0.97


30458.0
0.0
0.0
0.0
0.0
0.0
0.0
0.0
0.0
0.0
0.0
0.0
0.0
0.0
0.0
0.0
0.0
0.0
6803.0
23655.0
0.0
0.0
0.0
0.0
0.0
0.0
0.0
0.0
0.0
0.0

1.0
1.0
1.0
1.0
1.0
1.0
1.0
1.0
1.0
1.0
1.0
1.0
1.0
1.0
1.0
1.0
1.0
1.0
1.0
1.0
1.0
1.0
1.0
1.0
1.0
1.0
1.0
1.0
1.0
1.0


538.0
0.0
0.0
0.0
0.0
0.0
0.0
0.0
0.0
0.0
0.0
0.0
0.0
0.0
0.0
0.0
0.0
0.0
0.0
193.0
0.0
128.0
171.0
0.0
46.0
0.0
0.0
0.0
0.0
0.0

1.0
1.0
1.0
1.0
1.0
1.0
1.0
1.0
1.0
1.0
1.0
1.0
1.0
1.0
1.0
1.0
1.0
1.0
1.0
1.0
1.0
1.0
1.0
1.0
1.0
1.0
1.0
1.0
1.0
1.0


233.0
0.0
0.0
0.0
0.0
0.0
0.0
0.0
0.0
0.0
0.0
0.0
0.0
80.0
0.0
0.0
0.0
0.0
0.0
0.0
0.0
0.0
0.0
0.0
0.0
0.0
0.0
0.0
153.0
0.0

1.0
1.0
1.0
1.0
1.0
1.0
1.0
1.0
1.0
1.0
1.0
1.0
1.0
1.0
1.0
1.0
1.0
1.0
1.0
1.0
1.0
1.0
1.0
1.0
1.0
1.0
1.0
1.0
1.0
1.0


1907.0
0.0
0.0
0.0
0.0
0.0
0.0
0.0
0.0
0.0
0.0
0.0
0.0
0.0
0.0
825.0
335.0
0.0
0.0
0.0
0.0
0.0
0.0
0.0
60.0
152.0
518.0
0.0
12.0
5.0

0.99
0.99
0.99
0.99
0.99
0.99
0.99
0.99
0.99
0.99
0.99
0.99
0.99
0.99
0.99
0.99
0.99
0.99
0.99
0.99
0.99
0.99
0.99
0.99
0.99
0.99
0.99
0.99
0.99
0.99


13207.0
0.0
0.0
0.0
0.0
0.0
0.0
0.0
0.0
0.0
0.0
0.0
0.0
0.0
0.0
0.0
0.0
0.0
0.0
243.0
0.0
0.0
11.0
0.0
1850.0
365.0
1687.0
4043.0
4817.0
191.0


13207.0
0.0
0.0
0.0
0.0
0.0
0.0
0.0
0.0
0.0
0.0
0.0
0.0
0.0
0.0
0.0
0.0
0.0
0.0
243.0
0.0
0.0
11.0
0.0
1850.0
365.0
1687.0
4043.0
4817.0
191.0

1.0
1.0
1.0
1.0
1.0
1.0
1.0
1.0
1.0
1.0
1.0
1.0
1.0
1.0
1.0
1.0
1.0
1.0
1.0
1.0
1.0
1.0
1.0
1.0
1.0
1.0
1.0
1.0
1.0
1.0


12453.0
0.0
0.0
0.0
0.0
0.0
0.0
0.0
0.0
0.0
0.0
0.0
0.0
0.0
0.0
0.0
0.0
0.0
0.0
0.0
0.0
0.0
0.0
0.0
1454.0
365.0
1660.0
4038.0
4745.0
191.0

1.0
1.0
1.0
1.0
1.0
1.0
1.0
1.0
1.0
1.0
1.0
1.0
1.0
1.0
1.0
1.0
1.0
1.0
1.0
1.0
1.0
1.0
1.0
1.0
1.0
1.0
1.0
1.0
1.0
1.0


573.0
0.0
0.0
0.0
0.0
0.0
0.0
0.0
0.0
0.0
0.0
0.0
0.0
0.0
0.0
0.0
0.0
0.0
0.0
227.0
0.0
0.0
11.0
0.0
335.0
0.0
0.0
0.0
0.0
0.0

0.8
0.8
0.8
0.8
0.8
0.8
0.8
0.8
0.8
0.8
0.8
0.8
0.8
0.8
0.8
0.8
0.8
0.8
0.8
0.8
0.8
0.8
0.8
0.8
0.8
0.8
0.8
0.8
0.8
0.8


3050.0
0.0
0.0
0.0
0.0
0.0
0.0
0.0
0.0
0.0
0.0
0.0
0.0
0.0
0.0
1130.0
1920.0
0.0
0.0
0.0
0.0
0.0
0.0
0.0
0.0
0.0
0.0
0.0
0.0
0.0


3050.0
0.0
0.0
0.0
0.0
0.0
0.0
0.0
0.0
0.0
0.0
0.0
0.0
0.0
0.0
1130.0
1920.0
0.0
0.0
0.0
0.0
0.0
0.0
0.0
0.0
0.0
0.0
0.0
0.0
0.0


3050.0
0.0
0.0
0.0
0.0
0.0
0.0
0.0
0.0
0.0
0.0
0.0
0.0
0.0
0.0
1130.0
1920.0
0.0
0.0
0.0
0.0
0.0
0.0
0.0
0.0
0.0
0.0
0.0
0.0
0.0

1.0
1.0
1.0
1.0
1.0
1.0
1.0
1.0
1.0
1.0
1.0
1.0
1.0
1.0
1.0
1.0
1.0
1.0
1.0
1.0
1.0
1.0
1.0
1.0
1.0
1.0
1.0
1.0
1.0
1.0


2852.0
0.0
0.0
0.0
0.0
0.0
0.0
0.0
0.0
0.0
0.0
0.0
0.0
0.0
0.0
0.0
0.0
0.0
0.0
0.0
0.0
0.0
0.0
0.0
112.0
6.0
648.0
0.0
0.0
2086.0

0.97
0.97
0.97
0.97
0.97
0.97
0.97
0.97
0.97
0.97
0.97
0.97
0.97
0.97
0.97
0.97
0.97
0.97
0.97
0.97
0.97
0.97
0.97
0.97
0.97
0.97
0.97
0.97
0.97
0.97


1935.0
0.0
0.0
0.0
0.0
0.0
0.0
0.0
0.0
0.0
0.0
0.0
0.0
0.0
0.0
0.0
0.0
0.0
0.0
0.0
0.0
0.0
0.0
0.0
0.0
0.0
0.0
0.0
0.0
1935.0

0.99
0.99
0.99
0.99
0.99
0.99
0.99
0.99
0.99
0.99
0.99
0.99
0.99
0.99
0.99
0.99
0.99
0.99
0.99
0.99
0.99
0.99
0.99
0.99
0.99
0.99
0.99
0.99
0.99
0.99


4153.0
0.0
0.0
0.0
0.0
0.0
0.0
0.0
0.0
0.0
0.0
0.0
0.0
0.0
0.0
0.0
0.0
0.0
0.0
0.0
0.0
0.0
0.0
0.0
1899.0
0.0
2254.0
0.0
0.0
0.0


1899.0
0.0
0.0
0.0
0.0
0.0
0.0
0.0
0.0
0.0
0.0
0.0
0.0
0.0
0.0
0.0
0.0
0.0
0.0
0.0
0.0
0.0
0.0
0.0
1899.0
0.0
0.0
0.0
0.0
0.0


1899.0
0.0
0.0
0.0
0.0
0.0
0.0
0.0
0.0
0.0
0.0
0.0
0.0
0.0
0.0
0.0
0.0
0.0
0.0
0.0
0.0
0.0
0.0
0.0
1899.0
0.0
0.0
0.0
0.0
0.0

0.98
0.98
0.98
0.98
0.98
0.98
0.98
0.98
0.98
0.98
0.98
0.98
0.98
0.98
0.98
0.98
0.98
0.98
0.98
0.98
0.98
0.98
0.98
0.98
0.98
0.98
0.98
0.98
0.98
0.98


2254.0
0.0
0.0
0.0
0.0
0.0
0.0
0.0
0.0
0.0
0.0
0.0
0.0
0.0
0.0
0.0
0.0
0.0
0.0
0.0
0.0
0.0
0.0
0.0
0.0
0.0
2254.0
0.0
0.0
0.0

0.96
0.96
0.96
0.96
0.96
0.96
0.96
0.96
0.96
0.96
0.96
0.96
0.96
0.96
0.96
0.96
0.96
0.96
0.96
0.96
0.96
0.96
0.96
0.96
0.96
0.96
0.96
0.96
0.96
0.96


2147.0
0.0
0.0
0.0
0.0
0.0
0.0
0.0
0.0
0.0
0.0
0.0
0.0
0.0
0.0
0.0
0.0
0.0
0.0
0.0
0.0
0.0
0.0
0.0
2095.0
0.0
0.0
0.0
52.0
0.0


2095.0
0.0
0.0
0.0
0.0
0.0
0.0
0.0
0.0
0.0
0.0
0.0
0.0
0.0
0.0
0.0
0.0
0.0
0.0
0.0
0.0
0.0
0.0
0.0
2095.0
0.0
0.0
0.0
0.0
0.0

0.94
0.94
0.94
0.94
0.94
0.94
0.94
0.94
0.94
0.94
0.94
0.94
0.94
0.94
0.94
0.94
0.94
0.94
0.94
0.94
0.94
0.94
0.94
0.94
0.94
0.94
0.94
0.94
0.94
0.94


52.0
0.0
0.0
0.0
0.0
0.0
0.0
0.0
0.0
0.0
0.0
0.0
0.0
0.0
0.0
0.0
0.0
0.0
0.0
0.0
0.0
0.0
0.0
0.0
0.0
0.0
0.0
0.0
52.0
0.0

0.89
0.89
0.89
0.89
0.89
0.89
0.89
0.89
0.89
0.89
0.89
0.89
0.89
0.89
0.89
0.89
0.89
0.89
0.89
0.89
0.89
0.89
0.89
0.89
0.89
0.89
0.89
0.89
0.89
0.89


1592.0
0.0
0.0
0.0
0.0
0.0
0.0
0.0
0.0
0.0
0.0
0.0
0.0
0.0
0.0
0.0
1592.0
0.0
0.0
0.0
0.0
0.0
0.0
0.0
0.0
0.0
0.0
0.0
0.0
0.0


1592.0
0.0
0.0
0.0
0.0
0.0
0.0
0.0
0.0
0.0
0.0
0.0
0.0
0.0
0.0
0.0
1592.0
0.0
0.0
0.0
0.0
0.0
0.0
0.0
0.0
0.0
0.0
0.0
0.0
0.0

1.0
1.0
1.0
1.0
1.0
1.0
1.0
1.0
1.0
1.0
1.0
1.0
1.0
1.0
1.0
1.0
1.0
1.0
1.0
1.0
1.0
1.0
1.0
1.0
1.0
1.0
1.0
1.0
1.0
1.0


1224.0
0.0
0.0
0.0
0.0
0.0
0.0
0.0
0.0
0.0
0.0
0.0
0.0
0.0
0.0
6.0
0.0
34.0
764.0
5.0
68.0
0.0
0.0
0.0
215.0
50.0
21.0
0.0
33.0
28.0

1.0
1.0
1.0
1.0
1.0
1.0
1.0
1.0
1.0
1.0
1.0
1.0
1.0
1.0
1.0
1.0
1.0
1.0
1.0
1.0
1.0
1.0
1.0
1.0
1.0
1.0
1.0
1.0
1.0
1.0


207.0
0.0
0.0
0.0
0.0
0.0
0.0
0.0
0.0
0.0
0.0
0.0
0.0
0.0
0.0
0.0
0.0
0.0
0.0
0.0
45.0
0.0
0.0
0.0
162.0
0.0
0.0
0.0
0.0
0.0

1.0
1.0
1.0
1.0
1.0
1.0
1.0
1.0
1.0
1.0
1.0
1.0
1.0
1.0
1.0
1.0
1.0
1.0
1.0
1.0
1.0
1.0
1.0
1.0
1.0
1.0
1.0
1.0
1.0
1.0


75.0
0.0
0.0
0.0
0.0
0.0
0.0
0.0
0.0
0.0
0.0
0.0
0.0
0.0
0.0
0.0
0.0
0.0
0.0
0.0
0.0
0.0
0.0
0.0
75.0
0.0
0.0
0.0
0.0
0.0

1.0
1.0
1.0
1.0
1.0
1.0
1.0
1.0
1.0
1.0
1.0
1.0
1.0
1.0
1.0
1.0
1.0
1.0
1.0
1.0
1.0
1.0
1.0
1.0
1.0
1.0
1.0
1.0
1.0
1.0


81.0
0.0
0.0
0.0
0.0
0.0
0.0
0.0
0.0
0.0
0.0
0.0
0.0
0.0
0.0
0.0
0.0
27.0
31.0
0.0
23.0
0.0
0.0
0.0
0.0
0.0
0.0
0.0
0.0
0.0

1.0
1.0
1.0
1.0
1.0
1.0
1.0
1.0
1.0
1.0
1.0
1.0
1.0
1.0
1.0
1.0
1.0
1.0
1.0
1.0
1.0
1.0
1.0
1.0
1.0
1.0
1.0
1.0
1.0
1.0


26.0
0.0
0.0
0.0
0.0
0.0
0.0
0.0
0.0
0.0
0.0
0.0
0.0
0.0
0.0
0.0
0.0
0.0
0.0
0.0
0.0
0.0
0.0
0.0
0.0
0.0
6.0
0.0
0.0
20.0


26.0
0.0
0.0
0.0
0.0
0.0
0.0
0.0
0.0
0.0
0.0
0.0
0.0
0.0
0.0
0.0
0.0
0.0
0.0
0.0
0.0
0.0
0.0
0.0
0.0
0.0
6.0
0.0
0.0
20.0

0.96
0.96
0.96
0.96
0.96
0.96
0.96
0.96
0.96
0.96
0.96
0.96
0.96
0.96
0.96
0.96
0.96
0.96
0.96
0.96
0.96
0.96
0.96
0.96
0.96
0.96
0.96
0.96
0.96
0.96


12.0
0.0
0.0
0.0
0.0
0.0
0.0
0.0
0.0
0.0
0.0
0.0
0.0
0.0
0.0
0.0
0.0
0.0
0.0
0.0
0.0
0.0
0.0
0.0
0.0
7.0
0.0
0.0
5.0
0.0

0.72
0.72
0.72
0.72
0.72
0.72
0.72
0.72
0.72
0.72
0.72
0.72
0.72
0.72
0.72
0.72
0.72
0.72
0.72
0.72
0.72
0.72
0.72
0.72
0.72
0.72
0.72
0.72
0.72
0.72


1062.0
0.0
0.0
0.0
0.0
0.0
0.0
0.0
0.0
0.0
0.0
0.0
0.0
0.0
587.0
0.0
0.0
11.0
0.0
58.0
0.0
0.0
0.0
0.0
41.0
341.0
18.0
5.0
1.0
0.0


722.0
0.0
0.0
0.0
0.0
0.0
0.0
0.0
0.0
0.0
0.0
0.0
0.0
0.0
587.0
0.0
0.0
0.0
0.0
58.0
0.0
0.0
0.0
0.0
28.0
48.0
0.0
0.0
1.0
0.0

0.92
0.92
0.92
0.92
0.92
0.92
0.92
0.92
0.92
0.92
0.92
0.92
0.92
0.92
0.92
0.92
0.92
0.92
0.92
0.92
0.92
0.92
0.92
0.92
0.92
0.92
0.92
0.92
0.92
0.92


28.0
0.0
0.0
0.0
0.0
0.0
0.0
0.0
0.0
0.0
0.0
0.0
0.0
0.0
0.0
0.0
0.0
0.0
0.0
0.0
0.0
0.0
0.0
0.0
28.0
0.0
0.0
0.0
0.0
0.0

1.0
1.0
1.0
1.0
1.0
1.0
1.0
1.0
1.0
1.0
1.0
1.0
1.0
1.0
1.0
1.0
1.0
1.0
1.0
1.0
1.0
1.0
1.0
1.0
1.0
1.0
1.0
1.0
1.0
1.0


241.0
0.0
0.0
0.0
0.0
0.0
0.0
0.0
0.0
0.0
0.0
0.0
0.0
0.0
0.0
0.0
0.0
0.0
0.0
0.0
0.0
0.0
0.0
0.0
0.0
229.0
12.0
0.0
0.0
0.0

1.0
1.0
1.0
1.0
1.0
1.0
1.0
1.0
1.0
1.0
1.0
1.0
1.0
1.0
1.0
1.0
1.0
1.0
1.0
1.0
1.0
1.0
1.0
1.0
1.0
1.0
1.0
1.0
1.0
1.0


78.0
0.0
0.0
0.0
0.0
0.0
0.0
0.0
0.0
0.0
0.0
0.0
0.0
0.0
0.0
0.0
0.0
5.0
0.0
0.0
0.0
0.0
0.0
0.0
4.0
64.0
0.0
5.0
0.0
0.0

0.74
0.74
0.74
0.74
0.74
0.74
0.74
0.74
0.74
0.74
0.74
0.74
0.74
0.74
0.74
0.74
0.74
0.74
0.74
0.74
0.74
0.74
0.74
0.74
0.74
0.74
0.74
0.74
0.74
0.74


14.0
0.0
0.0
0.0
0.0
0.0
0.0
0.0
0.0
0.0
0.0
0.0
0.0
0.0
0.0
0.0
0.0
5.0
0.0
0.0
0.0
0.0
0.0
0.0
4.0
0.0
0.0
5.0
0.0
0.0

1.0
1.0
1.0
1.0
1.0
1.0
1.0
1.0
1.0
1.0
1.0
1.0
1.0
1.0
1.0
1.0
1.0
1.0
1.0
1.0
1.0
1.0
1.0
1.0
1.0
1.0
1.0
1.0
1.0
1.0


9.0
0.0
0.0
0.0
0.0
0.0
0.0
0.0
0.0
0.0
0.0
0.0
0.0
0.0
0.0
0.0
0.0
0.0
0.0
0.0
0.0
0.0
0.0
0.0
9.0
0.0
0.0
0.0
0.0
0.0

0.87
0.87
0.87
0.87
0.87
0.87
0.87
0.87
0.87
0.87
0.87
0.87
0.87
0.87
0.87
0.87
0.87
0.87
0.87
0.87
0.87
0.87
0.87
0.87
0.87
0.87
0.87
0.87
0.87
0.87


12.0
0.0
0.0
0.0
0.0
0.0
0.0
0.0
0.0
0.0
0.0
0.0
0.0
0.0
0.0
0.0
0.0
6.0
0.0
0.0
0.0
0.0
0.0
0.0
0.0
0.0
6.0
0.0
0.0
0.0

1.0
1.0
1.0
1.0
1.0
1.0
1.0
1.0
1.0
1.0
1.0
1.0
1.0
1.0
1.0
1.0
1.0
1.0
1.0
1.0
1.0
1.0
1.0
1.0
1.0
1.0
1.0
1.0
1.0
1.0


3147.0
0.0
0.0
0.0
0.0
0.0
0.0
0.0
0.0
0.0
0.0
0.0
0.0
0.0
0.0
442.0
0.0
112.0
12.0
0.0
414.0
0.0
226.0
0.0
73.0
34.0
161.0
967.0
568.0
138.0

0.94
0.94
0.94
0.94
0.94
0.94
0.94
0.94
0.94
0.94
0.94
0.94
0.94
0.94
0.94
0.94
0.94
0.94
0.94
0.94
0.94
0.94
0.94
0.94
0.94
0.94
0.94
0.94
0.94
0.94


654.0
0.0
0.0
0.0
0.0
0.0
0.0
0.0
0.0
0.0
0.0
0.0
0.0
0.0
0.0
0.0
0.0
0.0
0.0
0.0
0.0
0.0
0.0
0.0
4.0
0.0
89.0
520.0
16.0
25.0

0.97
0.97
0.97
0.97
0.97
0.97
0.97
0.97
0.97
0.97
0.97
0.97
0.97
0.97
0.97
0.97
0.97
0.97
0.97
0.97
0.97
0.97
0.97
0.97
0.97
0.97
0.97
0.97
0.97
0.97


875.0
0.0
0.0
0.0
0.0
0.0
0.0
0.0
0.0
0.0
0.0
0.0
0.0
0.0
0.0
442.0
0.0
8.0
5.0
0.0
0.0
0.0
0.0
0.0
60.0
22.0
0.0
337.0
1.0
0.0


875.0
0.0
0.0
0.0
0.0
0.0
0.0
0.0
0.0
0.0
0.0
0.0
0.0
0.0
0.0
442.0
0.0
8.0
5.0
0.0
0.0
0.0
0.0
0.0
60.0
22.0
0.0
337.0
1.0
0.0

1.0
1.0
1.0
1.0
1.0
1.0
1.0
1.0
1.0
1.0
1.0
1.0
1.0
1.0
1.0
1.0
1.0
1.0
1.0
1.0
1.0
1.0
1.0
1.0
1.0
1.0
1.0
1.0
1.0
1.0


1194.0
0.0
0.0
0.0
0.0
0.0
0.0
0.0
0.0
0.0
0.0
0.0
0.0
0.0
0.0
0.0
0.0
0.0
0.0
0.0
414.0
0.0
0.0
0.0
6.0
12.0
72.0
33.0
544.0
113.0

1.0
1.0
1.0
1.0
1.0
1.0
1.0
1.0
1.0
1.0
1.0
1.0
1.0
1.0
1.0
1.0
1.0
1.0
1.0
1.0
1.0
1.0
1.0
1.0
1.0
1.0
1.0
1.0
1.0
1.0


320.0
0.0
0.0
0.0
0.0
0.0
0.0
0.0
0.0
0.0
0.0
0.0
0.0
0.0
0.0
0.0
0.0
0.0
0.0
0.0
311.0
0.0
0.0
0.0
0.0
0.0
0.0
0.0
1.0
8.0

1.0
1.0
1.0
1.0
1.0
1.0
1.0
1.0
1.0
1.0
1.0
1.0
1.0
1.0
1.0
1.0
1.0
1.0
1.0
1.0
1.0
1.0
1.0
1.0
1.0
1.0
1.0
1.0
1.0
1.0


104.0
0.0
0.0
0.0
0.0
0.0
0.0
0.0
0.0
0.0
0.0
0.0
0.0
0.0
0.0
0.0
0.0
104.0
0.0
0.0
0.0
0.0
0.0
0.0
0.0
0.0
0.0
0.0
0.0
0.0

1.0
1.0
1.0
1.0
1.0
1.0
1.0
1.0
1.0
1.0
1.0
1.0
1.0
1.0
1.0
1.0
1.0
1.0
1.0
1.0
1.0
1.0
1.0
1.0
1.0
1.0
1.0
1.0
1.0
1.0


99.0
0.0
0.0
0.0
0.0
0.0
0.0
0.0
0.0
0.0
0.0
0.0
0.0
0.0
0.0
0.0
0.0
99.0
0.0
0.0
0.0
0.0
0.0
0.0
0.0
0.0
0.0
0.0
0.0
0.0

1.0
1.0
1.0
1.0
1.0
1.0
1.0
1.0
1.0
1.0
1.0
1.0
1.0
1.0
1.0
1.0
1.0
1.0
1.0
1.0
1.0
1.0
1.0
1.0
1.0
1.0
1.0
1.0
1.0
1.0


236.0
0.0
0.0
0.0
0.0
0.0
0.0
0.0
0.0
0.0
0.0
0.0
0.0
0.0
0.0
0.0
0.0
0.0
7.0
0.0
0.0
0.0
226.0
0.0
3.0
0.0
0.0
0.0
0.0
0.0

1.0
1.0
1.0
1.0
1.0
1.0
1.0
1.0
1.0
1.0
1.0
1.0
1.0
1.0
1.0
1.0
1.0
1.0
1.0
1.0
1.0
1.0
1.0
1.0
1.0
1.0
1.0
1.0
1.0
1.0


74.0
0.0
0.0
0.0
0.0
0.0
0.0
0.0
0.0
0.0
0.0
0.0
0.0
0.0
0.0
0.0
0.0
0.0
0.0
0.0
0.0
0.0
74.0
0.0
0.0
0.0
0.0
0.0
0.0
0.0

0.9
0.9
0.9
0.9
0.9
0.9
0.9
0.9
0.9
0.9
0.9
0.9
0.9
0.9
0.9
0.9
0.9
0.9
0.9
0.9
0.9
0.9
0.9
0.9
0.9
0.9
0.9
0.9
0.9
0.9


54.0
0.0
0.0
0.0
0.0
0.0
0.0
0.0
0.0
0.0
0.0
0.0
0.0
0.0
0.0
0.0
0.0
0.0
0.0
0.0
0.0
0.0
54.0
0.0
0.0
0.0
0.0
0.0
0.0
0.0

1.0
1.0
1.0
1.0
1.0
1.0
1.0
1.0
1.0
1.0
1.0
1.0
1.0
1.0
1.0
1.0
1.0
1.0
1.0
1.0
1.0
1.0
1.0
1.0
1.0
1.0
1.0
1.0
1.0
1.0


819.0
0.0
0.0
0.0
0.0
0.0
0.0
0.0
0.0
0.0
0.0
0.0
0.0
0.0
0.0
634.0
0.0
0.0
0.0
0.0
0.0
0.0
0.0
0.0
30.0
155.0
0.0
0.0
0.0
0.0


819.0
0.0
0.0
0.0
0.0
0.0
0.0
0.0
0.0
0.0
0.0
0.0
0.0
0.0
0.0
634.0
0.0
0.0
0.0
0.0
0.0
0.0
0.0
0.0
30.0
155.0
0.0
0.0
0.0
0.0

0.97
0.97
0.97
0.97
0.97
0.97
0.97
0.97
0.97
0.97
0.97
0.97
0.97
0.97
0.97
0.97
0.97
0.97
0.97
0.97
0.97
0.97
0.97
0.97
0.97
0.97
0.97
0.97
0.97
0.97


173.0
0.0
0.0
0.0
0.0
0.0
0.0
0.0
0.0
0.0
0.0
0.0
0.0
0.0
0.0
0.0
0.0
0.0
153.0
0.0
0.0
0.0
0.0
0.0
0.0
20.0
0.0
0.0
0.0
0.0


173.0
0.0
0.0
0.0
0.0
0.0
0.0
0.0
0.0
0.0
0.0
0.0
0.0
0.0
0.0
0.0
0.0
0.0
153.0
0.0
0.0
0.0
0.0
0.0
0.0
20.0
0.0
0.0
0.0
0.0

1.0
1.0
1.0
1.0
1.0
1.0
1.0
1.0
1.0
1.0
1.0
1.0
1.0
1.0
1.0
1.0
1.0
1.0
1.0
1.0
1.0
1.0
1.0
1.0
1.0
1.0
1.0
1.0
1.0
1.0


178.0
0.0
0.0
0.0
0.0
0.0
0.0
0.0
0.0
0.0
0.0
0.0
0.0
0.0
0.0
0.0
0.0
0.0
0.0
0.0
0.0
0.0
0.0
0.0
178.0
0.0
0.0
0.0
0.0
0.0


178.0
0.0
0.0
0.0
0.0
0.0
0.0
0.0
0.0
0.0
0.0
0.0
0.0
0.0
0.0
0.0
0.0
0.0
0.0
0.0
0.0
0.0
0.0
0.0
178.0
0.0
0.0
0.0
0.0
0.0


178.0
0.0
0.0
0.0
0.0
0.0
0.0
0.0
0.0
0.0
0.0
0.0
0.0
0.0
0.0
0.0
0.0
0.0
0.0
0.0
0.0
0.0
0.0
0.0
178.0
0.0
0.0
0.0
0.0
0.0

1.0
1.0
1.0
1.0
1.0
1.0
1.0
1.0
1.0
1.0
1.0
1.0
1.0
1.0
1.0
1.0
1.0
1.0
1.0
1.0
1.0
1.0
1.0
1.0
1.0
1.0
1.0
1.0
1.0
1.0


168.0
0.0
0.0
0.0
0.0
0.0
0.0
0.0
0.0
0.0
0.0
0.0
5.0
0.0
0.0
0.0
0.0
0.0
16.0
0.0
0.0
11.0
115.0
5.0
0.0
16.0
0.0
0.0
0.0
0.0

0.7
0.7
0.7
0.7
0.7
0.7
0.7
0.7
0.7
0.7
0.7
0.7
0.7
0.7
0.7
0.7
0.7
0.7
0.7
0.7
0.7
0.7
0.7
0.7
0.7
0.7
0.7
0.7
0.7
0.7


115.0
0.0
0.0
0.0
0.0
0.0
0.0
0.0
0.0
0.0
0.0
0.0
0.0
0.0
0.0
0.0
0.0
0.0
0.0
0.0
0.0
0.0
115.0
0.0
0.0
0.0
0.0
0.0
0.0
0.0

0.98
0.98
0.98
0.98
0.98
0.98
0.98
0.98
0.98
0.98
0.98
0.98
0.98
0.98
0.98
0.98
0.98
0.98
0.98
0.98
0.98
0.98
0.98
0.98
0.98
0.98
0.98
0.98
0.98
0.98


21.0
0.0
0.0
0.0
0.0
0.0
0.0
0.0
0.0
0.0
0.0
0.0
5.0
0.0
0.0
0.0
0.0
0.0
0.0
0.0
0.0
11.0
0.0
5.0
0.0
0.0
0.0
0.0
0.0
0.0

1.0
1.0
1.0
1.0
1.0
1.0
1.0
1.0
1.0
1.0
1.0
1.0
1.0
1.0
1.0
1.0
1.0
1.0
1.0
1.0
1.0
1.0
1.0
1.0
1.0
1.0
1.0
1.0
1.0
1.0


16.0
0.0
0.0
0.0
0.0
0.0
0.0
0.0
0.0
0.0
0.0
0.0
0.0
0.0
0.0
0.0
0.0
0.0
16.0
0.0
0.0
0.0
0.0
0.0
0.0
0.0
0.0
0.0
0.0
0.0

1.0
1.0
1.0
1.0
1.0
1.0
1.0
1.0
1.0
1.0
1.0
1.0
1.0
1.0
1.0
1.0
1.0
1.0
1.0
1.0
1.0
1.0
1.0
1.0
1.0
1.0
1.0
1.0
1.0
1.0


120.0
0.0
0.0
0.0
0.0
0.0
0.0
0.0
0.0
0.0
0.0
0.0
0.0
0.0
0.0
0.0
25.0
0.0
0.0
46.0
0.0
0.0
0.0
0.0
49.0
0.0
0.0
0.0
0.0
0.0

0.94
0.94
0.94
0.94
0.94
0.94
0.94
0.94
0.94
0.94
0.94
0.94
0.94
0.94
0.94
0.94
0.94
0.94
0.94
0.94
0.94
0.94
0.94
0.94
0.94
0.94
0.94
0.94
0.94
0.94


64.0
0.0
0.0
0.0
0.0
0.0
0.0
0.0
0.0
0.0
0.0
0.0
0.0
0.0
0.0
0.0
20.0
0.0
0.0
0.0
0.0
0.0
0.0
0.0
44.0
0.0
0.0
0.0
0.0
0.0

1.0
1.0
1.0
1.0
1.0
1.0
1.0
1.0
1.0
1.0
1.0
1.0
1.0
1.0
1.0
1.0
1.0
1.0
1.0
1.0
1.0
1.0
1.0
1.0
1.0
1.0
1.0
1.0
1.0
1.0


46.0
0.0
0.0
0.0
0.0
0.0
0.0
0.0
0.0
0.0
0.0
0.0
0.0
0.0
0.0
0.0
0.0
0.0
0.0
46.0
0.0
0.0
0.0
0.0
0.0
0.0
0.0
0.0
0.0
0.0

0.94
0.94
0.94
0.94
0.94
0.94
0.94
0.94
0.94
0.94
0.94
0.94
0.94
0.94
0.94
0.94
0.94
0.94
0.94
0.94
0.94
0.94
0.94
0.94
0.94
0.94
0.94
0.94
0.94
0.94


28.0
0.0
0.0
0.0
0.0
0.0
0.0
0.0
0.0
0.0
0.0
0.0
0.0
0.0
0.0
0.0
0.0
0.0
0.0
0.0
0.0
0.0
0.0
0.0
0.0
0.0
0.0
0.0
28.0
0.0

0.92
0.92
0.92
0.92
0.92
0.92
0.92
0.92
0.92
0.92
0.92
0.92
0.92
0.92
0.92
0.92
0.92
0.92
0.92
0.92
0.92
0.92
0.92
0.92
0.92
0.92
0.92
0.92
0.92
0.92


85.0
0.0
0.0
0.0
0.0
0.0
0.0
0.0
0.0
0.0
0.0
0.0
0.0
0.0
0.0
0.0
0.0
0.0
0.0
0.0
0.0
0.0
0.0
0.0
0.0
0.0
85.0
0.0
0.0
0.0


85.0
0.0
0.0
0.0
0.0
0.0
0.0
0.0
0.0
0.0
0.0
0.0
0.0
0.0
0.0
0.0
0.0
0.0
0.0
0.0
0.0
0.0
0.0
0.0
0.0
0.0
85.0
0.0
0.0
0.0


85.0
0.0
0.0
0.0
0.0
0.0
0.0
0.0
0.0
0.0
0.0
0.0
0.0
0.0
0.0
0.0
0.0
0.0
0.0
0.0
0.0
0.0
0.0
0.0
0.0
0.0
85.0
0.0
0.0
0.0

1.0
1.0
1.0
1.0
1.0
1.0
1.0
1.0
1.0
1.0
1.0
1.0
1.0
1.0
1.0
1.0
1.0
1.0
1.0
1.0
1.0
1.0
1.0
1.0
1.0
1.0
1.0
1.0
1.0
1.0


6029250.0
239.0
13618.0
12311.0
38982.0
44768.0
20722.0
0.0
0.0
2233.0
15680.0
6681.0
0.0
1321660.0
0.0
3.0
0.0
148383.0
266964.0
0.0
0.0
0.0
2340720.0
0.0
1796136.0
0.0
62.0
0.0
88.0
0.0

0.73
0.73
0.73
0.73
0.73
0.73
0.73
0.73
0.73
0.73
0.73
0.73
0.73
0.73
0.73
0.73
0.73
0.73
0.73
0.73
0.73
0.73
0.73
0.73
0.73
0.73
0.73
0.73
0.73
0.73


4519849.0
0.0
10518.0
11688.0
35749.0
22826.0
14822.0
0.0
0.0
600.0
13467.0
6205.0
0.0
1.0
0.0
3.0
0.0
0.0
266964.0
0.0
0.0
0.0
2340720.0
0.0
1796136.0
0.0
62.0
0.0
88.0
0.0

1.0
1.0
1.0
1.0
1.0
1.0
1.0
1.0
1.0
1.0
1.0
1.0
1.0
1.0
1.0
1.0
1.0
1.0
1.0
1.0
1.0
1.0
1.0
1.0
1.0
1.0
1.0
1.0
1.0
1.0


3153025.0
0.0
3899.0
7822.0
34635.0
0.0
8661.0
0.0
0.0
599.0
987.0
6087.0
0.0
1.0
0.0
1.0
0.0
0.0
1.0
0.0
0.0
0.0
2326246.0
0.0
764086.0
0.0
0.0
0.0
0.0
0.0

1.0
1.0
1.0
1.0
1.0
1.0
1.0
1.0
1.0
1.0
1.0
1.0
1.0
1.0
1.0
1.0
1.0
1.0
1.0
1.0
1.0
1.0
1.0
1.0
1.0
1.0
1.0
1.0
1.0
1.0


3077631.0
0.0
3869.0
7797.0
34561.0
0.0
8649.0
0.0
0.0
585.0
979.0
6038.0
0.0
1.0
0.0
1.0
0.0
0.0
1.0
0.0
0.0
0.0
2277479.0
0.0
737671.0
0.0
0.0
0.0
0.0
0.0

1.0
1.0
1.0
1.0
1.0
1.0
1.0
1.0
1.0
1.0
1.0
1.0
1.0
1.0
1.0
1.0
1.0
1.0
1.0
1.0
1.0
1.0
1.0
1.0
1.0
1.0
1.0
1.0
1.0
1.0


20.0
0.0
0.0
0.0
0.0
0.0
0.0
0.0
0.0
0.0
0.0
0.0
0.0
0.0
0.0
0.0
0.0
0.0
0.0
0.0
0.0
0.0
17.0
0.0
3.0
0.0
0.0
0.0
0.0
0.0

0.95
0.95
0.95
0.95
0.95
0.95
0.95
0.95
0.95
0.95
0.95
0.95
0.95
0.95
0.95
0.95
0.95
0.95
0.95
0.95
0.95
0.95
0.95
0.95
0.95
0.95
0.95
0.95
0.95
0.95


1038272.0
0.0
6195.0
3321.0
2.0
0.0
0.0
0.0
0.0
1.0
12226.0
0.0
0.0
0.0
0.0
0.0
0.0
0.0
219598.0
0.0
0.0
0.0
0.0
0.0
796929.0
0.0
0.0
0.0
0.0
0.0

1.0
1.0
1.0
1.0
1.0
1.0
1.0
1.0
1.0
1.0
1.0
1.0
1.0
1.0
1.0
1.0
1.0
1.0
1.0
1.0
1.0
1.0
1.0
1.0
1.0
1.0
1.0
1.0
1.0
1.0


50104.0
0.0
24.0
5.0
0.0
0.0
0.0
0.0
0.0
0.0
2.0
0.0
0.0
0.0
0.0
0.0
0.0
0.0
11818.0
0.0
0.0
0.0
0.0
0.0
38255.0
0.0
0.0
0.0
0.0
0.0

1.0
1.0
1.0
1.0
1.0
1.0
1.0
1.0
1.0
1.0
1.0
1.0
1.0
1.0
1.0
1.0
1.0
1.0
1.0
1.0
1.0
1.0
1.0
1.0
1.0
1.0
1.0
1.0
1.0
1.0


12081.0
0.0
0.0
0.0
0.0
0.0
0.0
0.0
0.0
0.0
0.0
0.0
0.0
0.0
0.0
0.0
0.0
0.0
1433.0
0.0
0.0
0.0
0.0
0.0
10648.0
0.0
0.0
0.0
0.0
0.0

0.97
0.97
0.97
0.97
0.97
0.97
0.97
0.97
0.97
0.97
0.97
0.97
0.97
0.97
0.97
0.97
0.97
0.97
0.97
0.97
0.97
0.97
0.97
0.97
0.97
0.97
0.97
0.97
0.97
0.97


3923.0
0.0
0.0
0.0
0.0
0.0
0.0
0.0
0.0
0.0
0.0
0.0
0.0
0.0
0.0
0.0
0.0
0.0
3923.0
0.0
0.0
0.0
0.0
0.0
0.0
0.0
0.0
0.0
0.0
0.0

0.99
0.99
0.99
0.99
0.99
0.99
0.99
0.99
0.99
0.99
0.99
0.99
0.99
0.99
0.99
0.99
0.99
0.99
0.99
0.99
0.99
0.99
0.99
0.99
0.99
0.99
0.99
0.99
0.99
0.99


71010.0
0.0
0.0
293.0
1052.0
21643.0
3754.0
0.0
0.0
0.0
0.0
0.0
0.0
0.0
0.0
2.0
0.0
0.0
0.0
0.0
0.0
0.0
0.0
0.0
44116.0
0.0
62.0
0.0
88.0
0.0

1.0
1.0
1.0
1.0
1.0
1.0
1.0
1.0
1.0
1.0
1.0
1.0
1.0
1.0
1.0
1.0
1.0
1.0
1.0
1.0
1.0
1.0
1.0
1.0
1.0
1.0
1.0
1.0
1.0
1.0


70577.0
0.0
0.0
286.0
1040.0
21424.0
3720.0
0.0
0.0
0.0
0.0
0.0
0.0
0.0
0.0
2.0
0.0
0.0
0.0
0.0
0.0
0.0
0.0
0.0
43955.0
0.0
62.0
0.0
88.0
0.0

1.0
1.0
1.0
1.0
1.0
1.0
1.0
1.0
1.0
1.0
1.0
1.0
1.0
1.0
1.0
1.0
1.0
1.0
1.0
1.0
1.0
1.0
1.0
1.0
1.0
1.0
1.0
1.0
1.0
1.0


2193.0
0.0
0.0
0.0
0.0
0.0
2193.0
0.0
0.0
0.0
0.0
0.0
0.0
0.0
0.0
0.0
0.0
0.0
0.0
0.0
0.0
0.0
0.0
0.0
0.0
0.0
0.0
0.0
0.0
0.0


2193.0
0.0
0.0
0.0
0.0
0.0
2193.0
0.0
0.0
0.0
0.0
0.0
0.0
0.0
0.0
0.0
0.0
0.0
0.0
0.0
0.0
0.0
0.0
0.0
0.0
0.0
0.0
0.0
0.0
0.0

0.94
0.94
0.94
0.94
0.94
0.94
0.94
0.94
0.94
0.94
0.94
0.94
0.94
0.94
0.94
0.94
0.94
0.94
0.94
0.94
0.94
0.94
0.94
0.94
0.94
0.94
0.94
0.94
0.94
0.94


486.0
0.0
185.0
183.0
0.0
0.0
0.0
0.0
0.0
0.0
0.0
118.0
0.0
0.0
0.0
0.0
0.0
0.0
0.0
0.0
0.0
0.0
0.0
0.0
0.0
0.0
0.0
0.0
0.0
0.0


486.0
0.0
185.0
183.0
0.0
0.0
0.0
0.0
0.0
0.0
0.0
118.0
0.0
0.0
0.0
0.0
0.0
0.0
0.0
0.0
0.0
0.0
0.0
0.0
0.0
0.0
0.0
0.0
0.0
0.0

1.0
1.0
1.0
1.0
1.0
1.0
1.0
1.0
1.0
1.0
1.0
1.0
1.0
1.0
1.0
1.0
1.0
1.0
1.0
1.0
1.0
1.0
1.0
1.0
1.0
1.0
1.0
1.0
1.0
1.0


344.0
0.0
0.0
2.0
19.0
266.0
57.0
0.0
0.0
0.0
0.0
0.0
0.0
0.0
0.0
0.0
0.0
0.0
0.0
0.0
0.0
0.0
0.0
0.0
0.0
0.0
0.0
0.0
0.0
0.0

0.97
0.97
0.97
0.97
0.97
0.97
0.97
0.97
0.97
0.97
0.97
0.97
0.97
0.97
0.97
0.97
0.97
0.97
0.97
0.97
0.97
0.97
0.97
0.97
0.97
0.97
0.97
0.97
0.97
0.97


2804.0
0.0
0.0
0.0
0.0
0.0
0.0
0.0
0.0
0.0
0.0
0.0
0.0
0.0
0.0
0.0
0.0
0.0
1900.0
0.0
0.0
0.0
0.0
0.0
904.0
0.0
0.0
0.0
0.0
0.0


2804.0
0.0
0.0
0.0
0.0
0.0
0.0
0.0
0.0
0.0
0.0
0.0
0.0
0.0
0.0
0.0
0.0
0.0
1900.0
0.0
0.0
0.0
0.0
0.0
904.0
0.0
0.0
0.0
0.0
0.0

1.0
1.0
1.0
1.0
1.0
1.0
1.0
1.0
1.0
1.0
1.0
1.0
1.0
1.0
1.0
1.0
1.0
1.0
1.0
1.0
1.0
1.0
1.0
1.0
1.0
1.0
1.0
1.0
1.0
1.0


3724.0
0.0
0.0
0.0
0.0
0.0
0.0
0.0
0.0
0.0
0.0
0.0
0.0
0.0
0.0
0.0
0.0
0.0
1140.0
0.0
0.0
0.0
0.0
0.0
2584.0
0.0
0.0
0.0
0.0
0.0

0.87
0.87
0.87
0.87
0.87
0.87
0.87
0.87
0.87
0.87
0.87
0.87
0.87
0.87
0.87
0.87
0.87
0.87
0.87
0.87
0.87
0.87
0.87
0.87
0.87
0.87
0.87
0.87
0.87
0.87


20435.0
0.0
0.0
0.0
0.0
0.0
0.0
0.0
0.0
0.0
0.0
0.0
0.0
0.0
0.0
0.0
0.0
0.0
4938.0
0.0
0.0
0.0
0.0
0.0
15497.0
0.0
0.0
0.0
0.0
0.0

0.97
0.97
0.97
0.97
0.97
0.97
0.97
0.97
0.97
0.97
0.97
0.97
0.97
0.97
0.97
0.97
0.97
0.97
0.97
0.97
0.97
0.97
0.97
0.97
0.97
0.97
0.97
0.97
0.97
0.97


1200.0
0.0
0.0
0.0
0.0
0.0
0.0
0.0
0.0
0.0
0.0
0.0
0.0
0.0
0.0
0.0
0.0
0.0
0.0
0.0
0.0
0.0
0.0
0.0
1200.0
0.0
0.0
0.0
0.0
0.0

0.77
0.77
0.77
0.77
0.77
0.77
0.77
0.77
0.77
0.77
0.77
0.77
0.77
0.77
0.77
0.77
0.77
0.77
0.77
0.77
0.77
0.77
0.77
0.77
0.77
0.77
0.77
0.77
0.77
0.77


3205.0
0.0
0.0
0.0
0.0
0.0
0.0
0.0
0.0
0.0
0.0
0.0
0.0
0.0
0.0
0.0
0.0
0.0
567.0
0.0
0.0
0.0
0.0
0.0
2638.0
0.0
0.0
0.0
0.0
0.0

0.89
0.89
0.89
0.89
0.89
0.89
0.89
0.89
0.89
0.89
0.89
0.89
0.89
0.89
0.89
0.89
0.89
0.89
0.89
0.89
0.89
0.89
0.89
0.89
0.89
0.89
0.89
0.89
0.89
0.89


839.0
0.0
0.0
0.0
0.0
0.0
0.0
0.0
0.0
0.0
0.0
0.0
0.0
0.0
0.0
0.0
0.0
0.0
0.0
0.0
0.0
0.0
0.0
0.0
839.0
0.0
0.0
0.0
0.0
0.0

0.89
0.89
0.89
0.89
0.89
0.89
0.89
0.89
0.89
0.89
0.89
0.89
0.89
0.89
0.89
0.89
0.89
0.89
0.89
0.89
0.89
0.89
0.89
0.89
0.89
0.89
0.89
0.89
0.89
0.89


1348351.0
239.0
3100.0
623.0
3202.0
21942.0
5312.0
0.0
0.0
1633.0
2213.0
476.0
0.0
1309611.0
0.0
0.0
0.0
0.0
0.0
0.0
0.0
0.0
0.0
0.0
0.0
0.0
0.0
0.0
0.0
0.0

1.0
1.0
1.0
1.0
1.0
1.0
1.0
1.0
1.0
1.0
1.0
1.0
1.0
1.0
1.0
1.0
1.0
1.0
1.0
1.0
1.0
1.0
1.0
1.0
1.0
1.0
1.0
1.0
1.0
1.0


18496.0
0.0
0.0
0.0
0.0
13184.0
5312.0
0.0
0.0
0.0
0.0
0.0
0.0
0.0
0.0
0.0
0.0
0.0
0.0
0.0
0.0
0.0
0.0
0.0
0.0
0.0
0.0
0.0
0.0
0.0


18496.0
0.0
0.0
0.0
0.0
13184.0
5312.0
0.0
0.0
0.0
0.0
0.0
0.0
0.0
0.0
0.0
0.0
0.0
0.0
0.0
0.0
0.0
0.0
0.0
0.0
0.0
0.0
0.0
0.0
0.0

1.0
1.0
1.0
1.0
1.0
1.0
1.0
1.0
1.0
1.0
1.0
1.0
1.0
1.0
1.0
1.0
1.0
1.0
1.0
1.0
1.0
1.0
1.0
1.0
1.0
1.0
1.0
1.0
1.0
1.0


2963.0
0.0
2963.0
0.0
0.0
0.0
0.0
0.0
0.0
0.0
0.0
0.0
0.0
0.0
0.0
0.0
0.0
0.0
0.0
0.0
0.0
0.0
0.0
0.0
0.0
0.0
0.0
0.0
0.0
0.0


2963.0
0.0
2963.0
0.0
0.0
0.0
0.0
0.0
0.0
0.0
0.0
0.0
0.0
0.0
0.0
0.0
0.0
0.0
0.0
0.0
0.0
0.0
0.0
0.0
0.0
0.0
0.0
0.0
0.0
0.0

1.0
1.0
1.0
1.0
1.0
1.0
1.0
1.0
1.0
1.0
1.0
1.0
1.0
1.0
1.0
1.0
1.0
1.0
1.0
1.0
1.0
1.0
1.0
1.0
1.0
1.0
1.0
1.0
1.0
1.0


2689.0
0.0
0.0
0.0
0.0
0.0
0.0
0.0
0.0
0.0
2213.0
476.0
0.0
0.0
0.0
0.0
0.0
0.0
0.0
0.0
0.0
0.0
0.0
0.0
0.0
0.0
0.0
0.0
0.0
0.0

1.0
1.0
1.0
1.0
1.0
1.0
1.0
1.0
1.0
1.0
1.0
1.0
1.0
1.0
1.0
1.0
1.0
1.0
1.0
1.0
1.0
1.0
1.0
1.0
1.0
1.0
1.0
1.0
1.0
1.0


2658.0
0.0
0.0
0.0
0.0
0.0
0.0
0.0
0.0
0.0
2182.0
476.0
0.0
0.0
0.0
0.0
0.0
0.0
0.0
0.0
0.0
0.0
0.0
0.0
0.0
0.0
0.0
0.0
0.0
0.0

1.0
1.0
1.0
1.0
1.0
1.0
1.0
1.0
1.0
1.0
1.0
1.0
1.0
1.0
1.0
1.0
1.0
1.0
1.0
1.0
1.0
1.0
1.0
1.0
1.0
1.0
1.0
1.0
1.0
1.0


1615.0
0.0
0.0
0.0
0.0
0.0
0.0
0.0
0.0
1615.0
0.0
0.0
0.0
0.0
0.0
0.0
0.0
0.0
0.0
0.0
0.0
0.0
0.0
0.0
0.0
0.0
0.0
0.0
0.0
0.0


1615.0
0.0
0.0
0.0
0.0
0.0
0.0
0.0
0.0
1615.0
0.0
0.0
0.0
0.0
0.0
0.0
0.0
0.0
0.0
0.0
0.0
0.0
0.0
0.0
0.0
0.0
0.0
0.0
0.0
0.0

1.0
1.0
1.0
1.0
1.0
1.0
1.0
1.0
1.0
1.0
1.0
1.0
1.0
1.0
1.0
1.0
1.0
1.0
1.0
1.0
1.0
1.0
1.0
1.0
1.0
1.0
1.0
1.0
1.0
1.0


137.0
0.0
137.0
0.0
0.0
0.0
0.0
0.0
0.0
0.0
0.0
0.0
0.0
0.0
0.0
0.0
0.0
0.0
0.0
0.0
0.0
0.0
0.0
0.0
0.0
0.0
0.0
0.0
0.0
0.0


137.0
0.0
137.0
0.0
0.0
0.0
0.0
0.0
0.0
0.0
0.0
0.0
0.0
0.0
0.0
0.0
0.0
0.0
0.0
0.0
0.0
0.0
0.0
0.0
0.0
0.0
0.0
0.0
0.0
0.0

0.72
0.72
0.72
0.72
0.72
0.72
0.72
0.72
0.72
0.72
0.72
0.72
0.72
0.72
0.72
0.72
0.72
0.72
0.72
0.72
0.72
0.72
0.72
0.72
0.72
0.72
0.72
0.72
0.72
0.72


90.0
0.0
0.0
0.0
3.0
87.0
0.0
0.0
0.0
0.0
0.0
0.0
0.0
0.0
0.0
0.0
0.0
0.0
0.0
0.0
0.0
0.0
0.0
0.0
0.0
0.0
0.0
0.0
0.0
0.0

0.92
0.92
0.92
0.92
0.92
0.92
0.92
0.92
0.92
0.92
0.92
0.92
0.92
0.92
0.92
0.92
0.92
0.92
0.92
0.92
0.92
0.92
0.92
0.92
0.92
0.92
0.92
0.92
0.92
0.92


44.0
0.0
0.0
0.0
0.0
44.0
0.0
0.0
0.0
0.0
0.0
0.0
0.0
0.0
0.0
0.0
0.0
0.0
0.0
0.0
0.0
0.0
0.0
0.0
0.0
0.0
0.0
0.0
0.0
0.0

1.0
1.0
1.0
1.0
1.0
1.0
1.0
1.0
1.0
1.0
1.0
1.0
1.0
1.0
1.0
1.0
1.0
1.0
1.0
1.0
1.0
1.0
1.0
1.0
1.0
1.0
1.0
1.0
1.0
1.0


299.0
15.0
0.0
1.0
0.0
0.0
0.0
0.0
0.0
0.0
0.0
0.0
0.0
283.0
0.0
0.0
0.0
0.0
0.0
0.0
0.0
0.0
0.0
0.0
0.0
0.0
0.0
0.0
0.0
0.0

0.99
0.99
0.99
0.99
0.99
0.99
0.99
0.99
0.99
0.99
0.99
0.99
0.99
0.99
0.99
0.99
0.99
0.99
0.99
0.99
0.99
0.99
0.99
0.99
0.99
0.99
0.99
0.99
0.99
0.99


1016486.0
16.0
0.0
4.0
0.0
0.0
0.0
0.0
0.0
0.0
0.0
0.0
0.0
1016466.0
0.0
0.0
0.0
0.0
0.0
0.0
0.0
0.0
0.0
0.0
0.0
0.0
0.0
0.0
0.0
0.0

1.0
1.0
1.0
1.0
1.0
1.0
1.0
1.0
1.0
1.0
1.0
1.0
1.0
1.0
1.0
1.0
1.0
1.0
1.0
1.0
1.0
1.0
1.0
1.0
1.0
1.0
1.0
1.0
1.0
1.0


619.0
0.0
0.0
0.0
31.0
0.0
588.0
0.0
0.0
0.0
0.0
0.0
0.0
0.0
0.0
0.0
0.0
0.0
0.0
0.0
0.0
0.0
0.0
0.0
0.0
0.0
0.0
0.0
0.0
0.0


619.0
0.0
0.0
0.0
31.0
0.0
588.0
0.0
0.0
0.0
0.0
0.0
0.0
0.0
0.0
0.0
0.0
0.0
0.0
0.0
0.0
0.0
0.0
0.0
0.0
0.0
0.0
0.0
0.0
0.0


619.0
0.0
0.0
0.0
31.0
0.0
588.0
0.0
0.0
0.0
0.0
0.0
0.0
0.0
0.0
0.0
0.0
0.0
0.0
0.0
0.0
0.0
0.0
0.0
0.0
0.0
0.0
0.0
0.0
0.0

1.0
1.0
1.0
1.0
1.0
1.0
1.0
1.0
1.0
1.0
1.0
1.0
1.0
1.0
1.0
1.0
1.0
1.0
1.0
1.0
1.0
1.0
1.0
1.0
1.0
1.0
1.0
1.0
1.0
1.0


148383.0
0.0
0.0
0.0
0.0
0.0
0.0
0.0
0.0
0.0
0.0
0.0
0.0
0.0
0.0
0.0
0.0
148383.0
0.0
0.0
0.0
0.0
0.0
0.0
0.0
0.0
0.0
0.0
0.0
0.0


148383.0
0.0
0.0
0.0
0.0
0.0
0.0
0.0
0.0
0.0
0.0
0.0
0.0
0.0
0.0
0.0
0.0
148383.0
0.0
0.0
0.0
0.0
0.0
0.0
0.0
0.0
0.0
0.0
0.0
0.0

1.0
1.0
1.0
1.0
1.0
1.0
1.0
1.0
1.0
1.0
1.0
1.0
1.0
1.0
1.0
1.0
1.0
1.0
1.0
1.0
1.0
1.0
1.0
1.0
1.0
1.0
1.0
1.0
1.0
1.0


148239.0
0.0
0.0
0.0
0.0
0.0
0.0
0.0
0.0
0.0
0.0
0.0
0.0
0.0
0.0
0.0
0.0
148239.0
0.0
0.0
0.0
0.0
0.0
0.0
0.0
0.0
0.0
0.0
0.0
0.0

1.0
1.0
1.0
1.0
1.0
1.0
1.0
1.0
1.0
1.0
1.0
1.0
1.0
1.0
1.0
1.0
1.0
1.0
1.0
1.0
1.0
1.0
1.0
1.0
1.0
1.0
1.0
1.0
1.0
1.0


3475415.0
45.0
1.0
296.0
2054.0
1674.0
10996.0
0.0
0.0
106.0
109.0
32.0
0.0
0.0
867485.0
378011.0
872336.0
0.0
0.0
0.0
0.0
1.0
0.0
0.0
195914.0
986002.0
63612.0
90335.0
5240.0
1166.0

0.89
0.89
0.89
0.89
0.89
0.89
0.89
0.89
0.89
0.89
0.89
0.89
0.89
0.89
0.89
0.89
0.89
0.89
0.89
0.89
0.89
0.89
0.89
0.89
0.89
0.89
0.89
0.89
0.89
0.89


2805891.0
0.0
0.0
296.0
1926.0
1451.0
10786.0
0.0
0.0
0.0
0.0
0.0
0.0
0.0
865034.0
376520.0
871804.0
0.0
0.0
0.0
0.0
1.0
0.0
0.0
194305.0
483766.0
2.0
0.0
0.0
0.0


10461.0
0.0
0.0
0.0
1.0
0.0
10460.0
0.0
0.0
0.0
0.0
0.0
0.0
0.0
0.0
0.0
0.0
0.0
0.0
0.0
0.0
0.0
0.0
0.0
0.0
0.0
0.0
0.0
0.0
0.0


10461.0
0.0
0.0
0.0
1.0
0.0
10460.0
0.0
0.0
0.0
0.0
0.0
0.0
0.0
0.0
0.0
0.0
0.0
0.0
0.0
0.0
0.0
0.0
0.0
0.0
0.0
0.0
0.0
0.0
0.0

1.0
1.0
1.0
1.0
1.0
1.0
1.0
1.0
1.0
1.0
1.0
1.0
1.0
1.0
1.0
1.0
1.0
1.0
1.0
1.0
1.0
1.0
1.0
1.0
1.0
1.0
1.0
1.0
1.0
1.0


1930396.0
0.0
0.0
296.0
1925.0
1451.0
326.0
0.0
0.0
0.0
0.0
0.0
0.0
0.0
0.0
376520.0
871804.0
0.0
0.0
0.0
0.0
1.0
0.0
0.0
194305.0
483766.0
2.0
0.0
0.0
0.0

0.99
0.99
0.99
0.99
0.99
0.99
0.99
0.99
0.99
0.99
0.99
0.99
0.99
0.99
0.99
0.99
0.99
0.99
0.99
0.99
0.99
0.99
0.99
0.99
0.99
0.99
0.99
0.99
0.99
0.99


2938.0
0.0
0.0
0.0
0.0
0.0
0.0
0.0
0.0
0.0
0.0
0.0
0.0
0.0
0.0
0.0
0.0
0.0
0.0
0.0
0.0
0.0
0.0
0.0
769.0
2169.0
0.0
0.0
0.0
0.0

0.94
0.94
0.94
0.94
0.94
0.94
0.94
0.94
0.94
0.94
0.94
0.94
0.94
0.94
0.94
0.94
0.94
0.94
0.94
0.94
0.94
0.94
0.94
0.94
0.94
0.94
0.94
0.94
0.94
0.94


865034.0
0.0
0.0
0.0
0.0
0.0
0.0
0.0
0.0
0.0
0.0
0.0
0.0
0.0
865034.0
0.0
0.0
0.0
0.0
0.0
0.0
0.0
0.0
0.0
0.0
0.0
0.0
0.0
0.0
0.0


865034.0
0.0
0.0
0.0
0.0
0.0
0.0
0.0
0.0
0.0
0.0
0.0
0.0
0.0
865034.0
0.0
0.0
0.0
0.0
0.0
0.0
0.0
0.0
0.0
0.0
0.0
0.0
0.0
0.0
0.0

1.0
1.0
1.0
1.0
1.0
1.0
1.0
1.0
1.0
1.0
1.0
1.0
1.0
1.0
1.0
1.0
1.0
1.0
1.0
1.0
1.0
1.0
1.0
1.0
1.0
1.0
1.0
1.0
1.0
1.0


527.0
0.0
0.0
0.0
120.0
197.0
210.0
0.0
0.0
0.0
0.0
0.0
0.0
0.0
0.0
0.0
0.0
0.0
0.0
0.0
0.0
0.0
0.0
0.0
0.0
0.0
0.0
0.0
0.0
0.0


527.0
0.0
0.0
0.0
120.0
197.0
210.0
0.0
0.0
0.0
0.0
0.0
0.0
0.0
0.0
0.0
0.0
0.0
0.0
0.0
0.0
0.0
0.0
0.0
0.0
0.0
0.0
0.0
0.0
0.0


527.0
0.0
0.0
0.0
120.0
197.0
210.0
0.0
0.0
0.0
0.0
0.0
0.0
0.0
0.0
0.0
0.0
0.0
0.0
0.0
0.0
0.0
0.0
0.0
0.0
0.0
0.0
0.0
0.0
0.0

1.0
1.0
1.0
1.0
1.0
1.0
1.0
1.0
1.0
1.0
1.0
1.0
1.0
1.0
1.0
1.0
1.0
1.0
1.0
1.0
1.0
1.0
1.0
1.0
1.0
1.0
1.0
1.0
1.0
1.0


312259.0
45.0
0.0
0.0
0.0
20.0
0.0
0.0
0.0
106.0
109.0
32.0
0.0
0.0
2451.0
0.0
0.0
0.0
0.0
0.0
0.0
0.0
0.0
0.0
0.0
149336.0
63575.0
90179.0
5240.0
1166.0

0.99
0.99
0.99
0.99
0.99
0.99
0.99
0.99
0.99
0.99
0.99
0.99
0.99
0.99
0.99
0.99
0.99
0.99
0.99
0.99
0.99
0.99
0.99
0.99
0.99
0.99
0.99
0.99
0.99
0.99


240240.0
45.0
0.0
0.0
0.0
9.0
0.0
0.0
0.0
106.0
109.0
32.0
0.0
0.0
2451.0
0.0
0.0
0.0
0.0
0.0
0.0
0.0
0.0
0.0
0.0
133340.0
34612.0
64296.0
5240.0
0.0

1.0
1.0
1.0
1.0
1.0
1.0
1.0
1.0
1.0
1.0
1.0
1.0
1.0
1.0
1.0
1.0
1.0
1.0
1.0
1.0
1.0
1.0
1.0
1.0
1.0
1.0
1.0
1.0
1.0
1.0


189952.0
0.0
0.0
0.0
0.0
9.0
0.0
0.0
0.0
0.0
0.0
0.0
0.0
0.0
0.0
0.0
0.0
0.0
0.0
0.0
0.0
0.0
0.0
0.0
0.0
133340.0
34612.0
21991.0
0.0
0.0

1.0
1.0
1.0
1.0
1.0
1.0
1.0
1.0
1.0
1.0
1.0
1.0
1.0
1.0
1.0
1.0
1.0
1.0
1.0
1.0
1.0
1.0
1.0
1.0
1.0
1.0
1.0
1.0
1.0
1.0


72015.0
0.0
0.0
0.0
0.0
11.0
0.0
0.0
0.0
0.0
0.0
0.0
0.0
0.0
0.0
0.0
0.0
0.0
0.0
0.0
0.0
0.0
0.0
0.0
0.0
15996.0
28963.0
25879.0
0.0
1166.0

1.0
1.0
1.0
1.0
1.0
1.0
1.0
1.0
1.0
1.0
1.0
1.0
1.0
1.0
1.0
1.0
1.0
1.0
1.0
1.0
1.0
1.0
1.0
1.0
1.0
1.0
1.0
1.0
1.0
1.0


11.0
0.0
0.0
0.0
0.0
11.0
0.0
0.0
0.0
0.0
0.0
0.0
0.0
0.0
0.0
0.0
0.0
0.0
0.0
0.0
0.0
0.0
0.0
0.0
0.0
0.0
0.0
0.0
0.0
0.0

0.95
0.95
0.95
0.95
0.95
0.95
0.95
0.95
0.95
0.95
0.95
0.95
0.95
0.95
0.95
0.95
0.95
0.95
0.95
0.95
0.95
0.95
0.95
0.95
0.95
0.95
0.95
0.95
0.95
0.95


47348.0
0.0
0.0
0.0
0.0
0.0
0.0
0.0
0.0
0.0
0.0
0.0
0.0
0.0
0.0
0.0
0.0
0.0
0.0
0.0
0.0
0.0
0.0
0.0
0.0
13439.0
8030.0
25879.0
0.0
0.0

1.0
1.0
1.0
1.0
1.0
1.0
1.0
1.0
1.0
1.0
1.0
1.0
1.0
1.0
1.0
1.0
1.0
1.0
1.0
1.0
1.0
1.0
1.0
1.0
1.0
1.0
1.0
1.0
1.0
1.0


3883.0
0.0
0.0
0.0
0.0
0.0
0.0
0.0
0.0
0.0
0.0
0.0
0.0
0.0
0.0
0.0
0.0
0.0
0.0
0.0
0.0
0.0
0.0
0.0
0.0
0.0
2717.0
0.0
0.0
1166.0

0.95
0.95
0.95
0.95
0.95
0.95
0.95
0.95
0.95
0.95
0.95
0.95
0.95
0.95
0.95
0.95
0.95
0.95
0.95
0.95
0.95
0.95
0.95
0.95
0.95
0.95
0.95
0.95
0.95
0.95


2031.0
0.0
0.0
0.0
0.0
0.0
0.0
0.0
0.0
0.0
0.0
0.0
0.0
0.0
0.0
0.0
0.0
0.0
0.0
0.0
0.0
0.0
0.0
0.0
0.0
0.0
2031.0
0.0
0.0
0.0

0.99
0.99
0.99
0.99
0.99
0.99
0.99
0.99
0.99
0.99
0.99
0.99
0.99
0.99
0.99
0.99
0.99
0.99
0.99
0.99
0.99
0.99
0.99
0.99
0.99
0.99
0.99
0.99
0.99
0.99


337.0
0.0
1.0
0.0
8.0
6.0
0.0
0.0
0.0
0.0
0.0
0.0
0.0
0.0
0.0
131.0
0.0
0.0
0.0
0.0
0.0
0.0
0.0
0.0
0.0
0.0
35.0
156.0
0.0
0.0


337.0
0.0
1.0
0.0
8.0
6.0
0.0
0.0
0.0
0.0
0.0
0.0
0.0
0.0
0.0
131.0
0.0
0.0
0.0
0.0
0.0
0.0
0.0
0.0
0.0
0.0
35.0
156.0
0.0
0.0


337.0
0.0
1.0
0.0
8.0
6.0
0.0
0.0
0.0
0.0
0.0
0.0
0.0
0.0
0.0
131.0
0.0
0.0
0.0
0.0
0.0
0.0
0.0
0.0
0.0
0.0
35.0
156.0
0.0
0.0

1.0
1.0
1.0
1.0
1.0
1.0
1.0
1.0
1.0
1.0
1.0
1.0
1.0
1.0
1.0
1.0
1.0
1.0
1.0
1.0
1.0
1.0
1.0
1.0
1.0
1.0
1.0
1.0
1.0
1.0


16469.0
714.0
0.0
0.0
0.0
0.0
0.0
0.0
0.0
0.0
1203.0
366.0
0.0
0.0
0.0
0.0
0.0
14021.0
0.0
165.0
0.0
0.0
0.0
0.0
0.0
0.0
0.0
0.0
0.0
0.0


16068.0
714.0
0.0
0.0
0.0
0.0
0.0
0.0
0.0
0.0
1203.0
366.0
0.0
0.0
0.0
0.0
0.0
13785.0
0.0
0.0
0.0
0.0
0.0
0.0
0.0
0.0
0.0
0.0
0.0
0.0

1.0
1.0
1.0
1.0
1.0
1.0
1.0
1.0
1.0
1.0
1.0
1.0
1.0
1.0
1.0
1.0
1.0
1.0
1.0
1.0
1.0
1.0
1.0
1.0
1.0
1.0
1.0
1.0
1.0
1.0


2283.0
714.0
0.0
0.0
0.0
0.0
0.0
0.0
0.0
0.0
1203.0
366.0
0.0
0.0
0.0
0.0
0.0
0.0
0.0
0.0
0.0
0.0
0.0
0.0
0.0
0.0
0.0
0.0
0.0
0.0

1.0
1.0
1.0
1.0
1.0
1.0
1.0
1.0
1.0
1.0
1.0
1.0
1.0
1.0
1.0
1.0
1.0
1.0
1.0
1.0
1.0
1.0
1.0
1.0
1.0
1.0
1.0
1.0
1.0
1.0


165.0
0.0
0.0
0.0
0.0
0.0
0.0
0.0
0.0
0.0
0.0
0.0
0.0
0.0
0.0
0.0
0.0
0.0
0.0
165.0
0.0
0.0
0.0
0.0
0.0
0.0
0.0
0.0
0.0
0.0

0.78
0.78
0.78
0.78
0.78
0.78
0.78
0.78
0.78
0.78
0.78
0.78
0.78
0.78
0.78
0.78
0.78
0.78
0.78
0.78
0.78
0.78
0.78
0.78
0.78
0.78
0.78
0.78
0.78
0.78


40.0
0.0
0.0
0.0
0.0
0.0
0.0
0.0
0.0
0.0
0.0
0.0
0.0
0.0
0.0
0.0
0.0
0.0
0.0
40.0
0.0
0.0
0.0
0.0
0.0
0.0
0.0
0.0
0.0
0.0

0.74
0.74
0.74
0.74
0.74
0.74
0.74
0.74
0.74
0.74
0.74
0.74
0.74
0.74
0.74
0.74
0.74
0.74
0.74
0.74
0.74
0.74
0.74
0.74
0.74
0.74
0.74
0.74
0.74
0.74


236.0
0.0
0.0
0.0
0.0
0.0
0.0
0.0
0.0
0.0
0.0
0.0
0.0
0.0
0.0
0.0
0.0
236.0
0.0
0.0
0.0
0.0
0.0
0.0
0.0
0.0
0.0
0.0
0.0
0.0


236.0
0.0
0.0
0.0
0.0
0.0
0.0
0.0
0.0
0.0
0.0
0.0
0.0
0.0
0.0
0.0
0.0
236.0
0.0
0.0
0.0
0.0
0.0
0.0
0.0
0.0
0.0
0.0
0.0
0.0


236.0
0.0
0.0
0.0
0.0
0.0
0.0
0.0
0.0
0.0
0.0
0.0
0.0
0.0
0.0
0.0
0.0
236.0
0.0
0.0
0.0
0.0
0.0
0.0
0.0
0.0
0.0
0.0
0.0
0.0

0.96
0.96
0.96
0.96
0.96
0.96
0.96
0.96
0.96
0.96
0.96
0.96
0.96
0.96
0.96
0.96
0.96
0.96
0.96
0.96
0.96
0.96
0.96
0.96
0.96
0.96
0.96
0.96
0.96
0.96


958863.0
219.0
77.0
91.0
0.0
0.0
33.0
0.0
0.0
0.0
0.0
544.0
0.0
0.0
0.0
0.0
644022.0
0.0
0.0
274635.0
39242.0
0.0
0.0
0.0
0.0
0.0
0.0
0.0
0.0
0.0

0.94
0.94
0.94
0.94
0.94
0.94
0.94
0.94
0.94
0.94
0.94
0.94
0.94
0.94
0.94
0.94
0.94
0.94
0.94
0.94
0.94
0.94
0.94
0.94
0.94
0.94
0.94
0.94
0.94
0.94


314841.0
219.0
77.0
91.0
0.0
0.0
33.0
0.0
0.0
0.0
0.0
544.0
0.0
0.0
0.0
0.0
0.0
0.0
0.0
274635.0
39242.0
0.0
0.0
0.0
0.0
0.0
0.0
0.0
0.0
0.0


314841.0
219.0
77.0
91.0
0.0
0.0
33.0
0.0
0.0
0.0
0.0
544.0
0.0
0.0
0.0
0.0
0.0
0.0
0.0
274635.0
39242.0
0.0
0.0
0.0
0.0
0.0
0.0
0.0
0.0
0.0

1.0
1.0
1.0
1.0
1.0
1.0
1.0
1.0
1.0
1.0
1.0
1.0
1.0
1.0
1.0
1.0
1.0
1.0
1.0
1.0
1.0
1.0
1.0
1.0
1.0
1.0
1.0
1.0
1.0
1.0


271269.0
219.0
77.0
0.0
0.0
0.0
33.0
0.0
0.0
0.0
0.0
544.0
0.0
0.0
0.0
0.0
0.0
0.0
0.0
270396.0
0.0
0.0
0.0
0.0
0.0
0.0
0.0
0.0
0.0
0.0

1.0
1.0
1.0
1.0
1.0
1.0
1.0
1.0
1.0
1.0
1.0
1.0
1.0
1.0
1.0
1.0
1.0
1.0
1.0
1.0
1.0
1.0
1.0
1.0
1.0
1.0
1.0
1.0
1.0
1.0


39290.0
0.0
0.0
91.0
0.0
0.0
0.0
0.0
0.0
0.0
0.0
0.0
0.0
0.0
0.0
0.0
0.0
0.0
0.0
0.0
39199.0
0.0
0.0
0.0
0.0
0.0
0.0
0.0
0.0
0.0

1.0
1.0
1.0
1.0
1.0
1.0
1.0
1.0
1.0
1.0
1.0
1.0
1.0
1.0
1.0
1.0
1.0
1.0
1.0
1.0
1.0
1.0
1.0
1.0
1.0
1.0
1.0
1.0
1.0
1.0


643134.0
0.0
0.0
0.0
0.0
0.0
0.0
0.0
0.0
0.0
0.0
0.0
0.0
0.0
0.0
0.0
643134.0
0.0
0.0
0.0
0.0
0.0
0.0
0.0
0.0
0.0
0.0
0.0
0.0
0.0


643134.0
0.0
0.0
0.0
0.0
0.0
0.0
0.0
0.0
0.0
0.0
0.0
0.0
0.0
0.0
0.0
643134.0
0.0
0.0
0.0
0.0
0.0
0.0
0.0
0.0
0.0
0.0
0.0
0.0
0.0


643134.0
0.0
0.0
0.0
0.0
0.0
0.0
0.0
0.0
0.0
0.0
0.0
0.0
0.0
0.0
0.0
643134.0
0.0
0.0
0.0
0.0
0.0
0.0
0.0
0.0
0.0
0.0
0.0
0.0
0.0

1.0
1.0
1.0
1.0
1.0
1.0
1.0
1.0
1.0
1.0
1.0
1.0
1.0
1.0
1.0
1.0
1.0
1.0
1.0
1.0
1.0
1.0
1.0
1.0
1.0
1.0
1.0
1.0
1.0
1.0


11723.0
305.0
12.0
22.0
39.0
0.0
13.0
614.0
0.0
439.0
30.0
162.0
0.0
0.0
52.0
0.0
0.0
0.0
474.0
0.0
6945.0
751.0
646.0
0.0
1212.0
7.0
0.0
0.0
0.0
0.0


2687.0
0.0
0.0
0.0
0.0
0.0
0.0
560.0
0.0
51.0
0.0
0.0
0.0
0.0
0.0
0.0
0.0
0.0
474.0
0.0
205.0
751.0
646.0
0.0
0.0
0.0
0.0
0.0
0.0
0.0


2526.0
0.0
0.0
0.0
0.0
0.0
0.0
399.0
0.0
51.0
0.0
0.0
0.0
0.0
0.0
0.0
0.0
0.0
474.0
0.0
205.0
751.0
646.0
0.0
0.0
0.0
0.0
0.0
0.0
0.0


2526.0
0.0
0.0
0.0
0.0
0.0
0.0
399.0
0.0
51.0
0.0
0.0
0.0
0.0
0.0
0.0
0.0
0.0
474.0
0.0
205.0
751.0
646.0
0.0
0.0
0.0
0.0
0.0
0.0
0.0

1.0
1.0
1.0
1.0
1.0
1.0
1.0
1.0
1.0
1.0
1.0
1.0
1.0
1.0
1.0
1.0
1.0
1.0
1.0
1.0
1.0
1.0
1.0
1.0
1.0
1.0
1.0
1.0
1.0
1.0


161.0
0.0
0.0
0.0
0.0
0.0
0.0
161.0
0.0
0.0
0.0
0.0
0.0
0.0
0.0
0.0
0.0
0.0
0.0
0.0
0.0
0.0
0.0
0.0
0.0
0.0
0.0
0.0
0.0
0.0


161.0
0.0
0.0
0.0
0.0
0.0
0.0
161.0
0.0
0.0
0.0
0.0
0.0
0.0
0.0
0.0
0.0
0.0
0.0
0.0
0.0
0.0
0.0
0.0
0.0
0.0
0.0
0.0
0.0
0.0

1.0
1.0
1.0
1.0
1.0
1.0
1.0
1.0
1.0
1.0
1.0
1.0
1.0
1.0
1.0
1.0
1.0
1.0
1.0
1.0
1.0
1.0
1.0
1.0
1.0
1.0
1.0
1.0
1.0
1.0


7195.0
44.0
0.0
0.0
0.0
0.0
0.0
0.0
0.0
275.0
0.0
136.0
0.0
0.0
0.0
0.0
0.0
0.0
0.0
0.0
6740.0
0.0
0.0
0.0
0.0
0.0
0.0
0.0
0.0
0.0


4632.0
0.0
0.0
0.0
0.0
0.0
0.0
0.0
0.0
275.0
0.0
102.0
0.0
0.0
0.0
0.0
0.0
0.0
0.0
0.0
4255.0
0.0
0.0
0.0
0.0
0.0
0.0
0.0
0.0
0.0

1.0
1.0
1.0
1.0
1.0
1.0
1.0
1.0
1.0
1.0
1.0
1.0
1.0
1.0
1.0
1.0
1.0
1.0
1.0
1.0
1.0
1.0
1.0
1.0
1.0
1.0
1.0
1.0
1.0
1.0


4624.0
0.0
0.0
0.0
0.0
0.0
0.0
0.0
0.0
275.0
0.0
102.0
0.0
0.0
0.0
0.0
0.0
0.0
0.0
0.0
4247.0
0.0
0.0
0.0
0.0
0.0
0.0
0.0
0.0
0.0

1.0
1.0
1.0
1.0
1.0
1.0
1.0
1.0
1.0
1.0
1.0
1.0
1.0
1.0
1.0
1.0
1.0
1.0
1.0
1.0
1.0
1.0
1.0
1.0
1.0
1.0
1.0
1.0
1.0
1.0


2563.0
44.0
0.0
0.0
0.0
0.0
0.0
0.0
0.0
0.0
0.0
34.0
0.0
0.0
0.0
0.0
0.0
0.0
0.0
0.0
2485.0
0.0
0.0
0.0
0.0
0.0
0.0
0.0
0.0
0.0


2563.0
44.0
0.0
0.0
0.0
0.0
0.0
0.0
0.0
0.0
0.0
34.0
0.0
0.0
0.0
0.0
0.0
0.0
0.0
0.0
2485.0
0.0
0.0
0.0
0.0
0.0
0.0
0.0
0.0
0.0

1.0
1.0
1.0
1.0
1.0
1.0
1.0
1.0
1.0
1.0
1.0
1.0
1.0
1.0
1.0
1.0
1.0
1.0
1.0
1.0
1.0
1.0
1.0
1.0
1.0
1.0
1.0
1.0
1.0
1.0


1585.0
261.0
12.0
22.0
39.0
0.0
13.0
54.0
0.0
113.0
30.0
26.0
0.0
0.0
52.0
0.0
0.0
0.0
0.0
0.0
0.0
0.0
0.0
0.0
963.0
0.0
0.0
0.0
0.0
0.0


1585.0
261.0
12.0
22.0
39.0
0.0
13.0
54.0
0.0
113.0
30.0
26.0
0.0
0.0
52.0
0.0
0.0
0.0
0.0
0.0
0.0
0.0
0.0
0.0
963.0
0.0
0.0
0.0
0.0
0.0


610.0
261.0
12.0
22.0
39.0
0.0
13.0
54.0
0.0
113.0
30.0
14.0
0.0
0.0
52.0
0.0
0.0
0.0
0.0
0.0
0.0
0.0
0.0
0.0
0.0
0.0
0.0
0.0
0.0
0.0

1.0
1.0
1.0
1.0
1.0
1.0
1.0
1.0
1.0
1.0
1.0
1.0
1.0
1.0
1.0
1.0
1.0
1.0
1.0
1.0
1.0
1.0
1.0
1.0
1.0
1.0
1.0
1.0
1.0
1.0


975.0
0.0
0.0
0.0
0.0
0.0
0.0
0.0
0.0
0.0
0.0
12.0
0.0
0.0
0.0
0.0
0.0
0.0
0.0
0.0
0.0
0.0
0.0
0.0
963.0
0.0
0.0
0.0
0.0
0.0

1.0
1.0
1.0
1.0
1.0
1.0
1.0
1.0
1.0
1.0
1.0
1.0
1.0
1.0
1.0
1.0
1.0
1.0
1.0
1.0
1.0
1.0
1.0
1.0
1.0
1.0
1.0
1.0
1.0
1.0


216.0
0.0
0.0
0.0
0.0
0.0
0.0
0.0
0.0
0.0
0.0
0.0
0.0
0.0
0.0
0.0
0.0
0.0
0.0
0.0
0.0
0.0
0.0
0.0
216.0
0.0
0.0
0.0
0.0
0.0


216.0
0.0
0.0
0.0
0.0
0.0
0.0
0.0
0.0
0.0
0.0
0.0
0.0
0.0
0.0
0.0
0.0
0.0
0.0
0.0
0.0
0.0
0.0
0.0
216.0
0.0
0.0
0.0
0.0
0.0


216.0
0.0
0.0
0.0
0.0
0.0
0.0
0.0
0.0
0.0
0.0
0.0
0.0
0.0
0.0
0.0
0.0
0.0
0.0
0.0
0.0
0.0
0.0
0.0
216.0
0.0
0.0
0.0
0.0
0.0

1.0
1.0
1.0
1.0
1.0
1.0
1.0
1.0
1.0
1.0
1.0
1.0
1.0
1.0
1.0
1.0
1.0
1.0
1.0
1.0
1.0
1.0
1.0
1.0
1.0
1.0
1.0
1.0
1.0
1.0


33.0
0.0
0.0
0.0
0.0
0.0
0.0
0.0
0.0
0.0
0.0
0.0
0.0
0.0
0.0
0.0
0.0
0.0
0.0
0.0
0.0
0.0
0.0
0.0
33.0
0.0
0.0
0.0
0.0
0.0


33.0
0.0
0.0
0.0
0.0
0.0
0.0
0.0
0.0
0.0
0.0
0.0
0.0
0.0
0.0
0.0
0.0
0.0
0.0
0.0
0.0
0.0
0.0
0.0
33.0
0.0
0.0
0.0
0.0
0.0


33.0
0.0
0.0
0.0
0.0
0.0
0.0
0.0
0.0
0.0
0.0
0.0
0.0
0.0
0.0
0.0
0.0
0.0
0.0
0.0
0.0
0.0
0.0
0.0
33.0
0.0
0.0
0.0
0.0
0.0

1.0
1.0
1.0
1.0
1.0
1.0
1.0
1.0
1.0
1.0
1.0
1.0
1.0
1.0
1.0
1.0
1.0
1.0
1.0
1.0
1.0
1.0
1.0
1.0
1.0
1.0
1.0
1.0
1.0
1.0


7.0
0.0
0.0
0.0
0.0
0.0
0.0
0.0
0.0
0.0
0.0
0.0
0.0
0.0
0.0
0.0
0.0
0.0
0.0
0.0
0.0
0.0
0.0
0.0
0.0
7.0
0.0
0.0
0.0
0.0


7.0
0.0
0.0
0.0
0.0
0.0
0.0
0.0
0.0
0.0
0.0
0.0
0.0
0.0
0.0
0.0
0.0
0.0
0.0
0.0
0.0
0.0
0.0
0.0
0.0
7.0
0.0
0.0
0.0
0.0


7.0
0.0
0.0
0.0
0.0
0.0
0.0
0.0
0.0
0.0
0.0
0.0
0.0
0.0
0.0
0.0
0.0
0.0
0.0
0.0
0.0
0.0
0.0
0.0
0.0
7.0
0.0
0.0
0.0
0.0

1.0
1.0
1.0
1.0
1.0
1.0
1.0
1.0
1.0
1.0
1.0
1.0
1.0
1.0
1.0
1.0
1.0
1.0
1.0
1.0
1.0
1.0
1.0
1.0
1.0
1.0
1.0
1.0
1.0
1.0


1016.0
14.0
2.0
0.0
0.0
0.0
0.0
4.0
14.0
100.0
241.0
15.0
14.0
0.0
0.0
0.0
0.0
0.0
0.0
0.0
0.0
44.0
0.0
14.0
0.0
5.0
105.0
59.0
254.0
131.0


309.0
7.0
0.0
0.0
0.0
0.0
0.0
0.0
7.0
61.0
162.0
0.0
14.0
0.0
0.0
0.0
0.0
0.0
0.0
0.0
0.0
44.0
0.0
14.0
0.0
0.0
0.0
0.0
0.0
0.0

0.81
0.81
0.81
0.81
0.81
0.81
0.81
0.81
0.81
0.81
0.81
0.81
0.81
0.81
0.81
0.81
0.81
0.81
0.81
0.81
0.81
0.81
0.81
0.81
0.81
0.81
0.81
0.81
0.81
0.81


219.0
6.0
0.0
0.0
0.0
0.0
0.0
0.0
6.0
49.0
158.0
0.0
0.0
0.0
0.0
0.0
0.0
0.0
0.0
0.0
0.0
0.0
0.0
0.0
0.0
0.0
0.0
0.0
0.0
0.0

0.99
0.99
0.99
0.99
0.99
0.99
0.99
0.99
0.99
0.99
0.99
0.99
0.99
0.99
0.99
0.99
0.99
0.99
0.99
0.99
0.99
0.99
0.99
0.99
0.99
0.99
0.99
0.99
0.99
0.99


18.0
1.0
0.0
0.0
0.0
0.0
0.0
0.0
1.0
12.0
4.0
0.0
0.0
0.0
0.0
0.0
0.0
0.0
0.0
0.0
0.0
0.0
0.0
0.0
0.0
0.0
0.0
0.0
0.0
0.0


10.0
0.0
0.0
0.0
0.0
0.0
0.0
0.0
0.0
7.0
3.0
0.0
0.0
0.0
0.0
0.0
0.0
0.0
0.0
0.0
0.0
0.0
0.0
0.0
0.0
0.0
0.0
0.0
0.0
0.0

1.0
1.0
1.0
1.0
1.0
1.0
1.0
1.0
1.0
1.0
1.0
1.0
1.0
1.0
1.0
1.0
1.0
1.0
1.0
1.0
1.0
1.0
1.0
1.0
1.0
1.0
1.0
1.0
1.0
1.0


8.0
1.0
0.0
0.0
0.0
0.0
0.0
0.0
1.0
5.0
1.0
0.0
0.0
0.0
0.0
0.0
0.0
0.0
0.0
0.0
0.0
0.0
0.0
0.0
0.0
0.0
0.0
0.0
0.0
0.0

1.0
1.0
1.0
1.0
1.0
1.0
1.0
1.0
1.0
1.0
1.0
1.0
1.0
1.0
1.0
1.0
1.0
1.0
1.0
1.0
1.0
1.0
1.0
1.0
1.0
1.0
1.0
1.0
1.0
1.0


145.0
1.0
0.0
0.0
0.0
0.0
0.0
4.0
7.0
39.0
79.0
15.0
0.0
0.0
0.0
0.0
0.0
0.0
0.0
0.0
0.0
0.0
0.0
0.0
0.0
0.0
0.0
0.0
0.0
0.0


145.0
1.0
0.0
0.0
0.0
0.0
0.0
4.0
7.0
39.0
79.0
15.0
0.0
0.0
0.0
0.0
0.0
0.0
0.0
0.0
0.0
0.0
0.0
0.0
0.0
0.0
0.0
0.0
0.0
0.0


145.0
1.0
0.0
0.0
0.0
0.0
0.0
4.0
7.0
39.0
79.0
15.0
0.0
0.0
0.0
0.0
0.0
0.0
0.0
0.0
0.0
0.0
0.0
0.0
0.0
0.0
0.0
0.0
0.0
0.0

1.0
1.0
1.0
1.0
1.0
1.0
1.0
1.0
1.0
1.0
1.0
1.0
1.0
1.0
1.0
1.0
1.0
1.0
1.0
1.0
1.0
1.0
1.0
1.0
1.0
1.0
1.0
1.0
1.0
1.0


8.0
6.0
2.0
0.0
0.0
0.0
0.0
0.0
0.0
0.0
0.0
0.0
0.0
0.0
0.0
0.0
0.0
0.0
0.0
0.0
0.0
0.0
0.0
0.0
0.0
0.0
0.0
0.0
0.0
0.0


8.0
6.0
2.0
0.0
0.0
0.0
0.0
0.0
0.0
0.0
0.0
0.0
0.0
0.0
0.0
0.0
0.0
0.0
0.0
0.0
0.0
0.0
0.0
0.0
0.0
0.0
0.0
0.0
0.0
0.0


8.0
6.0
2.0
0.0
0.0
0.0
0.0
0.0
0.0
0.0
0.0
0.0
0.0
0.0
0.0
0.0
0.0
0.0
0.0
0.0
0.0
0.0
0.0
0.0
0.0
0.0
0.0
0.0
0.0
0.0

1.0
1.0
1.0
1.0
1.0
1.0
1.0
1.0
1.0
1.0
1.0
1.0
1.0
1.0
1.0
1.0
1.0
1.0
1.0
1.0
1.0
1.0
1.0
1.0
1.0
1.0
1.0
1.0
1.0
1.0


554.0
0.0
0.0
0.0
0.0
0.0
0.0
0.0
0.0
0.0
0.0
0.0
0.0
0.0
0.0
0.0
0.0
0.0
0.0
0.0
0.0
0.0
0.0
0.0
0.0
5.0
105.0
59.0
254.0
131.0


410.0
0.0
0.0
0.0
0.0
0.0
0.0
0.0
0.0
0.0
0.0
0.0
0.0
0.0
0.0
0.0
0.0
0.0
0.0
0.0
0.0
0.0
0.0
0.0
0.0
5.0
81.0
33.0
179.0
112.0

0.97
0.97
0.97
0.97
0.97
0.97
0.97
0.97
0.97
0.97
0.97
0.97
0.97
0.97
0.97
0.97
0.97
0.97
0.97
0.97
0.97
0.97
0.97
0.97
0.97
0.97
0.97
0.97
0.97
0.97


144.0
0.0
0.0
0.0
0.0
0.0
0.0
0.0
0.0
0.0
0.0
0.0
0.0
0.0
0.0
0.0
0.0
0.0
0.0
0.0
0.0
0.0
0.0
0.0
0.0
0.0
24.0
26.0
75.0
19.0


144.0
0.0
0.0
0.0
0.0
0.0
0.0
0.0
0.0
0.0
0.0
0.0
0.0
0.0
0.0
0.0
0.0
0.0
0.0
0.0
0.0
0.0
0.0
0.0
0.0
0.0
24.0
26.0
75.0
19.0

1.0
1.0
1.0
1.0
1.0
1.0
1.0
1.0
1.0
1.0
1.0
1.0
1.0
1.0
1.0
1.0
1.0
1.0
1.0
1.0
1.0
1.0
1.0
1.0
1.0
1.0
1.0
1.0
1.0
1.0


198.0
198.0
0.0
0.0
0.0
0.0
0.0
0.0
0.0
0.0
0.0
0.0
0.0
0.0
0.0
0.0
0.0
0.0
0.0
0.0
0.0
0.0
0.0
0.0
0.0
0.0
0.0
0.0
0.0
0.0


198.0
198.0
0.0
0.0
0.0
0.0
0.0
0.0
0.0
0.0
0.0
0.0
0.0
0.0
0.0
0.0
0.0
0.0
0.0
0.0
0.0
0.0
0.0
0.0
0.0
0.0
0.0
0.0
0.0
0.0


198.0
198.0
0.0
0.0
0.0
0.0
0.0
0.0
0.0
0.0
0.0
0.0
0.0
0.0
0.0
0.0
0.0
0.0
0.0
0.0
0.0
0.0
0.0
0.0
0.0
0.0
0.0
0.0
0.0
0.0


198.0
198.0
0.0
0.0
0.0
0.0
0.0
0.0
0.0
0.0
0.0
0.0
0.0
0.0
0.0
0.0
0.0
0.0
0.0
0.0
0.0
0.0
0.0
0.0
0.0
0.0
0.0
0.0
0.0
0.0

0.95
0.95
0.95
0.95
0.95
0.95
0.95
0.95
0.95
0.95
0.95
0.95
0.95
0.95
0.95
0.95
0.95
0.95
0.95
0.95
0.95
0.95
0.95
0.95
0.95
0.95
0.95
0.95
0.95
0.95


145.0
0.0
0.0
0.0
0.0
0.0
0.0
0.0
0.0
0.0
145.0
0.0
0.0
0.0
0.0
0.0
0.0
0.0
0.0
0.0
0.0
0.0
0.0
0.0
0.0
0.0
0.0
0.0
0.0
0.0


145.0
0.0
0.0
0.0
0.0
0.0
0.0
0.0
0.0
0.0
145.0
0.0
0.0
0.0
0.0
0.0
0.0
0.0
0.0
0.0
0.0
0.0
0.0
0.0
0.0
0.0
0.0
0.0
0.0
0.0


145.0
0.0
0.0
0.0
0.0
0.0
0.0
0.0
0.0
0.0
145.0
0.0
0.0
0.0
0.0
0.0
0.0
0.0
0.0
0.0
0.0
0.0
0.0
0.0
0.0
0.0
0.0
0.0
0.0
0.0


145.0
0.0
0.0
0.0
0.0
0.0
0.0
0.0
0.0
0.0
145.0
0.0
0.0
0.0
0.0
0.0
0.0
0.0
0.0
0.0
0.0
0.0
0.0
0.0
0.0
0.0
0.0
0.0
0.0
0.0

1.0
1.0
1.0
1.0
1.0
1.0
1.0
1.0
1.0
1.0
1.0
1.0
1.0
1.0
1.0
1.0
1.0
1.0
1.0
1.0
1.0
1.0
1.0
1.0
1.0
1.0
1.0
1.0
1.0
1.0


22.0
0.0
0.0
0.0
0.0
0.0
0.0
0.0
0.0
22.0
0.0
0.0
0.0
0.0
0.0
0.0
0.0
0.0
0.0
0.0
0.0
0.0
0.0
0.0
0.0
0.0
0.0
0.0
0.0
0.0


22.0
0.0
0.0
0.0
0.0
0.0
0.0
0.0
0.0
22.0
0.0
0.0
0.0
0.0
0.0
0.0
0.0
0.0
0.0
0.0
0.0
0.0
0.0
0.0
0.0
0.0
0.0
0.0
0.0
0.0


22.0
0.0
0.0
0.0
0.0
0.0
0.0
0.0
0.0
22.0
0.0
0.0
0.0
0.0
0.0
0.0
0.0
0.0
0.0
0.0
0.0
0.0
0.0
0.0
0.0
0.0
0.0
0.0
0.0
0.0

0.93
0.93
0.93
0.93
0.93
0.93
0.93
0.93
0.93
0.93
0.93
0.93
0.93
0.93
0.93
0.93
0.93
0.93
0.93
0.93
0.93
0.93
0.93
0.93
0.93
0.93
0.93
0.93
0.93
0.93


10031.0
291.0
57.0
24.0
7.0
0.0
0.0
83.0
39.0
82.0
458.0
334.0
166.0
762.0
0.0
9.0
0.0
0.0
0.0
643.0
407.0
46.0
632.0
166.0
939.0
0.0
2653.0
678.0
9.0
1546.0

0.74
0.74
0.74
0.74
0.74
0.74
0.74
0.74
0.74
0.74
0.74
0.74
0.74
0.74
0.74
0.74
0.74
0.74
0.74
0.74
0.74
0.74
0.74
0.74
0.74
0.74
0.74
0.74
0.74
0.74


9540.0
291.0
57.0
24.0
7.0
0.0
0.0
83.0
39.0
82.0
458.0
334.0
68.0
762.0
0.0
9.0
0.0
0.0
0.0
643.0
407.0
46.0
632.0
68.0
939.0
0.0
2642.0
497.0
9.0
1443.0

0.9
0.9
0.9
0.9
0.9
0.9
0.9
0.9
0.9
0.9
0.9
0.9
0.9
0.9
0.9
0.9
0.9
0.9
0.9
0.9
0.9
0.9
0.9
0.9
0.9
0.9
0.9
0.9
0.9
0.9


9351.0
288.0
55.0
24.0
7.0
0.0
0.0
71.0
39.0
46.0
458.0
333.0
16.0
762.0
0.0
9.0
0.0
0.0
0.0
643.0
407.0
46.0
601.0
16.0
939.0
0.0
2642.0
497.0
9.0
1443.0

0.79
0.79
0.79
0.79
0.79
0.79
0.79
0.79
0.79
0.79
0.79
0.79
0.79
0.79
0.79
0.79
0.79
0.79
0.79
0.79
0.79
0.79
0.79
0.79
0.79
0.79
0.79
0.79
0.79
0.79


9343.0
288.0
55.0
24.0
7.0
0.0
0.0
71.0
39.0
46.0
458.0
333.0
16.0
762.0
0.0
9.0
0.0
0.0
0.0
643.0
407.0
46.0
601.0
16.0
931.0
0.0
2642.0
497.0
9.0
1443.0

0.95
0.95
0.95
0.95
0.95
0.95
0.95
0.95
0.95
0.95
0.95
0.95
0.95
0.95
0.95
0.95
0.95
0.95
0.95
0.95
0.95
0.95
0.95
0.95
0.95
0.95
0.95
0.95
0.95
0.95


3605.0
288.0
55.0
24.0
7.0
0.0
0.0
71.0
39.0
37.0
458.0
333.0
14.0
0.0
0.0
0.0
0.0
0.0
0.0
406.0
379.0
46.0
503.0
14.0
931.0
0.0
0.0
0.0
0.0
0.0

1.0
1.0
1.0
1.0
1.0
1.0
1.0
1.0
1.0
1.0
1.0
1.0
1.0
1.0
1.0
1.0
1.0
1.0
1.0
1.0
1.0
1.0
1.0
1.0
1.0
1.0
1.0
1.0
1.0
1.0


4591.0
0.0
0.0
0.0
0.0
0.0
0.0
0.0
0.0
0.0
0.0
0.0
0.0
0.0
0.0
9.0
0.0
0.0
0.0
0.0
0.0
0.0
0.0
0.0
0.0
0.0
2642.0
488.0
9.0
1443.0

0.94
0.94
0.94
0.94
0.94
0.94
0.94
0.94
0.94
0.94
0.94
0.94
0.94
0.94
0.94
0.94
0.94
0.94
0.94
0.94
0.94
0.94
0.94
0.94
0.94
0.94
0.94
0.94
0.94
0.94


762.0
0.0
0.0
0.0
0.0
0.0
0.0
0.0
0.0
0.0
0.0
0.0
0.0
762.0
0.0
0.0
0.0
0.0
0.0
0.0
0.0
0.0
0.0
0.0
0.0
0.0
0.0
0.0
0.0
0.0

0.86
0.86
0.86
0.86
0.86
0.86
0.86
0.86
0.86
0.86
0.86
0.86
0.86
0.86
0.86
0.86
0.86
0.86
0.86
0.86
0.86
0.86
0.86
0.86
0.86
0.86
0.86
0.86
0.86
0.86


24.0
0.0
0.0
0.0
0.0
0.0
0.0
0.0
0.0
0.0
0.0
0.0
0.0
0.0
0.0
0.0
0.0
0.0
0.0
0.0
24.0
0.0
0.0
0.0
0.0
0.0
0.0
0.0
0.0
0.0

0.72
0.72
0.72
0.72
0.72
0.72
0.72
0.72
0.72
0.72
0.72
0.72
0.72
0.72
0.72
0.72
0.72
0.72
0.72
0.72
0.72
0.72
0.72
0.72
0.72
0.72
0.72
0.72
0.72
0.72


5.0
0.0
0.0
0.0
0.0
0.0
0.0
0.0
0.0
0.0
0.0
0.0
0.0
0.0
0.0
0.0
0.0
0.0
0.0
0.0
0.0
0.0
5.0
0.0
0.0
0.0
0.0
0.0
0.0
0.0

0.82
0.82
0.82
0.82
0.82
0.82
0.82
0.82
0.82
0.82
0.82
0.82
0.82
0.82
0.82
0.82
0.82
0.82
0.82
0.82
0.82
0.82
0.82
0.82
0.82
0.82
0.82
0.82
0.82
0.82


54.0
3.0
2.0
0.0
0.0
0.0
0.0
12.0
0.0
36.0
0.0
1.0
0.0
0.0
0.0
0.0
0.0
0.0
0.0
0.0
0.0
0.0
0.0
0.0
0.0
0.0
0.0
0.0
0.0
0.0


54.0
3.0
2.0
0.0
0.0
0.0
0.0
12.0
0.0
36.0
0.0
1.0
0.0
0.0
0.0
0.0
0.0
0.0
0.0
0.0
0.0
0.0
0.0
0.0
0.0
0.0
0.0
0.0
0.0
0.0


54.0
3.0
2.0
0.0
0.0
0.0
0.0
12.0
0.0
36.0
0.0
1.0
0.0
0.0
0.0
0.0
0.0
0.0
0.0
0.0
0.0
0.0
0.0
0.0
0.0
0.0
0.0
0.0
0.0
0.0

0.82
0.82
0.82
0.82
0.82
0.82
0.82
0.82
0.82
0.82
0.82
0.82
0.82
0.82
0.82
0.82
0.82
0.82
0.82
0.82
0.82
0.82
0.82
0.82
0.82
0.82
0.82
0.82
0.82
0.82


132263.0
95.0
201.0
151.0
200.0
12.0
88.0
661.0
153.0
82.0
331.0
275.0
6898.0
8333.0
16399.0
29518.0
98.0
3834.0
572.0
144.0
2260.0
11.0
647.0
6898.0
37438.0
3040.0
6625.0
4249.0
2857.0
193.0

0.72
0.72
0.72
0.72
0.72
0.72
0.72
0.72
0.72
0.72
0.72
0.72
0.72
0.72
0.72
0.72
0.72
0.72
0.72
0.72
0.72
0.72
0.72
0.72
0.72
0.72
0.72
0.72
0.72
0.72


131623.0
83.0
201.0
151.0
200.0
12.0
88.0
640.0
153.0
82.0
256.0
268.0
6898.0
8327.0
16399.0
29518.0
98.0
3834.0
264.0
5.0
2238.0
6.0
610.0
6898.0
37434.0
3040.0
6625.0
4249.0
2853.0
193.0

1.0
1.0
1.0
1.0
1.0
1.0
1.0
1.0
1.0
1.0
1.0
1.0
1.0
1.0
1.0
1.0
1.0
1.0
1.0
1.0
1.0
1.0
1.0
1.0
1.0
1.0
1.0
1.0
1.0
1.0


32325.0
55.0
54.0
49.0
24.0
1.0
0.0
181.0
0.0
11.0
122.0
22.0
3821.0
343.0
86.0
0.0
0.0
597.0
0.0
0.0
1629.0
0.0
0.0
3821.0
21509.0
0.0
0.0
0.0
0.0
0.0

0.72
0.72
0.72
0.72
0.72
0.72
0.72
0.72
0.72
0.72
0.72
0.72
0.72
0.72
0.72
0.72
0.72
0.72
0.72
0.72
0.72
0.72
0.72
0.72
0.72
0.72
0.72
0.72
0.72
0.72


19243.0
55.0
54.0
49.0
24.0
1.0
0.0
181.0
0.0
11.0
122.0
22.0
0.0
343.0
0.0
0.0
0.0
0.0
0.0
0.0
0.0
0.0
0.0
0.0
18381.0
0.0
0.0
0.0
0.0
0.0

0.8
0.8
0.8
0.8
0.8
0.8
0.8
0.8
0.8
0.8
0.8
0.8
0.8
0.8
0.8
0.8
0.8
0.8
0.8
0.8
0.8
0.8
0.8
0.8
0.8
0.8
0.8
0.8
0.8
0.8


18771.0
27.0
3.0
11.0
0.0
0.0
0.0
0.0
0.0
0.0
0.0
6.0
0.0
343.0
0.0
0.0
0.0
0.0
0.0
0.0
0.0
0.0
0.0
0.0
18381.0
0.0
0.0
0.0
0.0
0.0

1.0
1.0
1.0
1.0
1.0
1.0
1.0
1.0
1.0
1.0
1.0
1.0
1.0
1.0
1.0
1.0
1.0
1.0
1.0
1.0
1.0
1.0
1.0
1.0
1.0
1.0
1.0
1.0
1.0
1.0


27408.0
4.0
132.0
10.0
84.0
2.0
18.0
28.0
17.0
29.0
27.0
68.0
0.0
0.0
11567.0
0.0
0.0
0.0
0.0
0.0
0.0
0.0
0.0
0.0
15258.0
0.0
0.0
0.0
164.0
0.0


24290.0
4.0
121.0
5.0
84.0
2.0
18.0
28.0
17.0
29.0
27.0
67.0
0.0
0.0
8466.0
0.0
0.0
0.0
0.0
0.0
0.0
0.0
0.0
0.0
15258.0
0.0
0.0
0.0
164.0
0.0


24252.0
4.0
121.0
5.0
84.0
2.0
18.0
28.0
17.0
29.0
27.0
67.0
0.0
0.0
8466.0
0.0
0.0
0.0
0.0
0.0
0.0
0.0
0.0
0.0
15221.0
0.0
0.0
0.0
163.0
0.0

1.0
1.0
1.0
1.0
1.0
1.0
1.0
1.0
1.0
1.0
1.0
1.0
1.0
1.0
1.0
1.0
1.0
1.0
1.0
1.0
1.0
1.0
1.0
1.0
1.0
1.0
1.0
1.0
1.0
1.0


38.0
0.0
0.0
0.0
0.0
0.0
0.0
0.0
0.0
0.0
0.0
0.0
0.0
0.0
0.0
0.0
0.0
0.0
0.0
0.0
0.0
0.0
0.0
0.0
37.0
0.0
0.0
0.0
1.0
0.0

0.9
0.9
0.9
0.9
0.9
0.9
0.9
0.9
0.9
0.9
0.9
0.9
0.9
0.9
0.9
0.9
0.9
0.9
0.9
0.9
0.9
0.9
0.9
0.9
0.9
0.9
0.9
0.9
0.9
0.9


3118.0
0.0
11.0
5.0
0.0
0.0
0.0
0.0
0.0
0.0
0.0
1.0
0.0
0.0
3101.0
0.0
0.0
0.0
0.0
0.0
0.0
0.0
0.0
0.0
0.0
0.0
0.0
0.0
0.0
0.0


3118.0
0.0
11.0
5.0
0.0
0.0
0.0
0.0
0.0
0.0
0.0
1.0
0.0
0.0
3101.0
0.0
0.0
0.0
0.0
0.0
0.0
0.0
0.0
0.0
0.0
0.0
0.0
0.0
0.0
0.0

1.0
1.0
1.0
1.0
1.0
1.0
1.0
1.0
1.0
1.0
1.0
1.0
1.0
1.0
1.0
1.0
1.0
1.0
1.0
1.0
1.0
1.0
1.0
1.0
1.0
1.0
1.0
1.0
1.0
1.0


13656.0
9.0
0.0
6.0
59.0
1.0
60.0
83.0
120.0
18.0
0.0
85.0
0.0
6505.0
4598.0
0.0
0.0
268.0
0.0
0.0
145.0
0.0
0.0
0.0
0.0
850.0
0.0
758.0
0.0
91.0


4979.0
9.0
0.0
0.0
0.0
0.0
0.0
0.0
117.0
18.0
0.0
54.0
0.0
0.0
4513.0
0.0
0.0
268.0
0.0
0.0
0.0
0.0
0.0
0.0
0.0
0.0
0.0
0.0
0.0
0.0


4979.0
9.0
0.0
0.0
0.0
0.0
0.0
0.0
117.0
18.0
0.0
54.0
0.0
0.0
4513.0
0.0
0.0
268.0
0.0
0.0
0.0
0.0
0.0
0.0
0.0
0.0
0.0
0.0
0.0
0.0

0.93
0.93
0.93
0.93
0.93
0.93
0.93
0.93
0.93
0.93
0.93
0.93
0.93
0.93
0.93
0.93
0.93
0.93
0.93
0.93
0.93
0.93
0.93
0.93
0.93
0.93
0.93
0.93
0.93
0.93


983.0
0.0
0.0
0.0
2.0
1.0
47.0
83.0
0.0
0.0
0.0
0.0
0.0
0.0
0.0
0.0
0.0
0.0
0.0
0.0
0.0
0.0
0.0
0.0
0.0
850.0
0.0
0.0
0.0
0.0


983.0
0.0
0.0
0.0
2.0
1.0
47.0
83.0
0.0
0.0
0.0
0.0
0.0
0.0
0.0
0.0
0.0
0.0
0.0
0.0
0.0
0.0
0.0
0.0
0.0
850.0
0.0
0.0
0.0
0.0

1.0
1.0
1.0
1.0
1.0
1.0
1.0
1.0
1.0
1.0
1.0
1.0
1.0
1.0
1.0
1.0
1.0
1.0
1.0
1.0
1.0
1.0
1.0
1.0
1.0
1.0
1.0
1.0
1.0
1.0


6792.0
0.0
0.0
0.0
57.0
0.0
0.0
0.0
0.0
0.0
0.0
0.0
0.0
6505.0
85.0
0.0
0.0
0.0
0.0
0.0
145.0
0.0
0.0
0.0
0.0
0.0
0.0
0.0
0.0
0.0

0.82
0.82
0.82
0.82
0.82
0.82
0.82
0.82
0.82
0.82
0.82
0.82
0.82
0.82
0.82
0.82
0.82
0.82
0.82
0.82
0.82
0.82
0.82
0.82
0.82
0.82
0.82
0.82
0.82
0.82


6650.0
0.0
0.0
0.0
0.0
0.0
0.0
0.0
0.0
0.0
0.0
0.0
0.0
6505.0
0.0
0.0
0.0
0.0
0.0
0.0
145.0
0.0
0.0
0.0
0.0
0.0
0.0
0.0
0.0
0.0

0.95
0.95
0.95
0.95
0.95
0.95
0.95
0.95
0.95
0.95
0.95
0.95
0.95
0.95
0.95
0.95
0.95
0.95
0.95
0.95
0.95
0.95
0.95
0.95
0.95
0.95
0.95
0.95
0.95
0.95


133.0
0.0
0.0
6.0
0.0
0.0
13.0
0.0
3.0
0.0
0.0
20.0
0.0
0.0
0.0
0.0
0.0
0.0
0.0
0.0
0.0
0.0
0.0
0.0
0.0
0.0
0.0
0.0
0.0
91.0

0.94
0.94
0.94
0.94
0.94
0.94
0.94
0.94
0.94
0.94
0.94
0.94
0.94
0.94
0.94
0.94
0.94
0.94
0.94
0.94
0.94
0.94
0.94
0.94
0.94
0.94
0.94
0.94
0.94
0.94


91.0
0.0
0.0
0.0
0.0
0.0
0.0
0.0
0.0
0.0
0.0
0.0
0.0
0.0
0.0
0.0
0.0
0.0
0.0
0.0
0.0
0.0
0.0
0.0
0.0
0.0
0.0
0.0
0.0
91.0

0.93
0.93
0.93
0.93
0.93
0.93
0.93
0.93
0.93
0.93
0.93
0.93
0.93
0.93
0.93
0.93
0.93
0.93
0.93
0.93
0.93
0.93
0.93
0.93
0.93
0.93
0.93
0.93
0.93
0.93


11.0
0.0
0.0
0.0
0.0
0.0
0.0
0.0
0.0
0.0
0.0
11.0
0.0
0.0
0.0
0.0
0.0
0.0
0.0
0.0
0.0
0.0
0.0
0.0
0.0
0.0
0.0
0.0
0.0
0.0


11.0
0.0
0.0
0.0
0.0
0.0
0.0
0.0
0.0
0.0
0.0
11.0
0.0
0.0
0.0
0.0
0.0
0.0
0.0
0.0
0.0
0.0
0.0
0.0
0.0
0.0
0.0
0.0
0.0
0.0

1.0
1.0
1.0
1.0
1.0
1.0
1.0
1.0
1.0
1.0
1.0
1.0
1.0
1.0
1.0
1.0
1.0
1.0
1.0
1.0
1.0
1.0
1.0
1.0
1.0
1.0
1.0
1.0
1.0
1.0


758.0
0.0
0.0
0.0
0.0
0.0
0.0
0.0
0.0
0.0
0.0
0.0
0.0
0.0
0.0
0.0
0.0
0.0
0.0
0.0
0.0
0.0
0.0
0.0
0.0
0.0
0.0
758.0
0.0
0.0

1.0
1.0
1.0
1.0
1.0
1.0
1.0
1.0
1.0
1.0
1.0
1.0
1.0
1.0
1.0
1.0
1.0
1.0
1.0
1.0
1.0
1.0
1.0
1.0
1.0
1.0
1.0
1.0
1.0
1.0


6830.0
0.0
0.0
0.0
5.0
7.0
0.0
152.0
1.0
9.0
32.0
2.0
2674.0
0.0
0.0
0.0
0.0
0.0
0.0
0.0
0.0
0.0
0.0
2674.0
0.0
1031.0
243.0
0.0
0.0
0.0

0.85
0.85
0.85
0.85
0.85
0.85
0.85
0.85
0.85
0.85
0.85
0.85
0.85
0.85
0.85
0.85
0.85
0.85
0.85
0.85
0.85
0.85
0.85
0.85
0.85
0.85
0.85
0.85
0.85
0.85


5494.0
0.0
0.0
0.0
0.0
0.0
0.0
129.0
0.0
0.0
15.0
2.0
2674.0
0.0
0.0
0.0
0.0
0.0
0.0
0.0
0.0
0.0
0.0
2674.0
0.0
0.0
0.0
0.0
0.0
0.0


5494.0
0.0
0.0
0.0
0.0
0.0
0.0
129.0
0.0
0.0
15.0
2.0
2674.0
0.0
0.0
0.0
0.0
0.0
0.0
0.0
0.0
0.0
0.0
2674.0
0.0
0.0
0.0
0.0
0.0
0.0

1.0
1.0
1.0
1.0
1.0
1.0
1.0
1.0
1.0
1.0
1.0
1.0
1.0
1.0
1.0
1.0
1.0
1.0
1.0
1.0
1.0
1.0
1.0
1.0
1.0
1.0
1.0
1.0
1.0
1.0


261.0
0.0
0.0
0.0
5.0
7.0
0.0
0.0
0.0
0.0
6.0
0.0
0.0
0.0
0.0
0.0
0.0
0.0
0.0
0.0
0.0
0.0
0.0
0.0
0.0
0.0
243.0
0.0
0.0
0.0


261.0
0.0
0.0
0.0
5.0
7.0
0.0
0.0
0.0
0.0
6.0
0.0
0.0
0.0
0.0
0.0
0.0
0.0
0.0
0.0
0.0
0.0
0.0
0.0
0.0
0.0
243.0
0.0
0.0
0.0

1.0
1.0
1.0
1.0
1.0
1.0
1.0
1.0
1.0
1.0
1.0
1.0
1.0
1.0
1.0
1.0
1.0
1.0
1.0
1.0
1.0
1.0
1.0
1.0
1.0
1.0
1.0
1.0
1.0
1.0


1031.0
0.0
0.0
0.0
0.0
0.0
0.0
0.0
0.0
0.0
0.0
0.0
0.0
0.0
0.0
0.0
0.0
0.0
0.0
0.0
0.0
0.0
0.0
0.0
0.0
1031.0
0.0
0.0
0.0
0.0


1031.0
0.0
0.0
0.0
0.0
0.0
0.0
0.0
0.0
0.0
0.0
0.0
0.0
0.0
0.0
0.0
0.0
0.0
0.0
0.0
0.0
0.0
0.0
0.0
0.0
1031.0
0.0
0.0
0.0
0.0

1.0
1.0
1.0
1.0
1.0
1.0
1.0
1.0
1.0
1.0
1.0
1.0
1.0
1.0
1.0
1.0
1.0
1.0
1.0
1.0
1.0
1.0
1.0
1.0
1.0
1.0
1.0
1.0
1.0
1.0


2234.0
10.0
0.0
33.0
0.0
0.0
0.0
130.0
0.0
0.0
0.0
0.0
0.0
0.0
93.0
0.0
0.0
1711.0
197.0
0.0
0.0
0.0
0.0
0.0
60.0
0.0
0.0
0.0
0.0
0.0

0.98
0.98
0.98
0.98
0.98
0.98
0.98
0.98
0.98
0.98
0.98
0.98
0.98
0.98
0.98
0.98
0.98
0.98
0.98
0.98
0.98
0.98
0.98
0.98
0.98
0.98
0.98
0.98
0.98
0.98


220.0
10.0
0.0
0.0
0.0
0.0
0.0
117.0
0.0
0.0
0.0
0.0
0.0
0.0
93.0
0.0
0.0
0.0
0.0
0.0
0.0
0.0
0.0
0.0
0.0
0.0
0.0
0.0
0.0
0.0


220.0
10.0
0.0
0.0
0.0
0.0
0.0
117.0
0.0
0.0
0.0
0.0
0.0
0.0
93.0
0.0
0.0
0.0
0.0
0.0
0.0
0.0
0.0
0.0
0.0
0.0
0.0
0.0
0.0
0.0

1.0
1.0
1.0
1.0
1.0
1.0
1.0
1.0
1.0
1.0
1.0
1.0
1.0
1.0
1.0
1.0
1.0
1.0
1.0
1.0
1.0
1.0
1.0
1.0
1.0
1.0
1.0
1.0
1.0
1.0


22.0
0.0
0.0
22.0
0.0
0.0
0.0
0.0
0.0
0.0
0.0
0.0
0.0
0.0
0.0
0.0
0.0
0.0
0.0
0.0
0.0
0.0
0.0
0.0
0.0
0.0
0.0
0.0
0.0
0.0


22.0
0.0
0.0
22.0
0.0
0.0
0.0
0.0
0.0
0.0
0.0
0.0
0.0
0.0
0.0
0.0
0.0
0.0
0.0
0.0
0.0
0.0
0.0
0.0
0.0
0.0
0.0
0.0
0.0
0.0

1.0
1.0
1.0
1.0
1.0
1.0
1.0
1.0
1.0
1.0
1.0
1.0
1.0
1.0
1.0
1.0
1.0
1.0
1.0
1.0
1.0
1.0
1.0
1.0
1.0
1.0
1.0
1.0
1.0
1.0


268.0
0.0
0.0
11.0
0.0
0.0
0.0
0.0
0.0
0.0
0.0
0.0
0.0
0.0
0.0
0.0
0.0
0.0
197.0
0.0
0.0
0.0
0.0
0.0
60.0
0.0
0.0
0.0
0.0
0.0


71.0
0.0
0.0
11.0
0.0
0.0
0.0
0.0
0.0
0.0
0.0
0.0
0.0
0.0
0.0
0.0
0.0
0.0
0.0
0.0
0.0
0.0
0.0
0.0
60.0
0.0
0.0
0.0
0.0
0.0

1.0
1.0
1.0
1.0
1.0
1.0
1.0
1.0
1.0
1.0
1.0
1.0
1.0
1.0
1.0
1.0
1.0
1.0
1.0
1.0
1.0
1.0
1.0
1.0
1.0
1.0
1.0
1.0
1.0
1.0


197.0
0.0
0.0
0.0
0.0
0.0
0.0
0.0
0.0
0.0
0.0
0.0
0.0
0.0
0.0
0.0
0.0
0.0
197.0
0.0
0.0
0.0
0.0
0.0
0.0
0.0
0.0
0.0
0.0
0.0

1.0
1.0
1.0
1.0
1.0
1.0
1.0
1.0
1.0
1.0
1.0
1.0
1.0
1.0
1.0
1.0
1.0
1.0
1.0
1.0
1.0
1.0
1.0
1.0
1.0
1.0
1.0
1.0
1.0
1.0


1711.0
0.0
0.0
0.0
0.0
0.0
0.0
0.0
0.0
0.0
0.0
0.0
0.0
0.0
0.0
0.0
0.0
1711.0
0.0
0.0
0.0
0.0
0.0
0.0
0.0
0.0
0.0
0.0
0.0
0.0


1711.0
0.0
0.0
0.0
0.0
0.0
0.0
0.0
0.0
0.0
0.0
0.0
0.0
0.0
0.0
0.0
0.0
1711.0
0.0
0.0
0.0
0.0
0.0
0.0
0.0
0.0
0.0
0.0
0.0
0.0

1.0
1.0
1.0
1.0
1.0
1.0
1.0
1.0
1.0
1.0
1.0
1.0
1.0
1.0
1.0
1.0
1.0
1.0
1.0
1.0
1.0
1.0
1.0
1.0
1.0
1.0
1.0
1.0
1.0
1.0


1357.0
0.0
0.0
0.0
0.0
0.0
0.0
66.0
0.0
0.0
0.0
0.0
4.0
0.0
15.0
0.0
0.0
1142.0
60.0
5.0
6.0
0.0
0.0
4.0
55.0
0.0
0.0
0.0
0.0
0.0

1.0
1.0
1.0
1.0
1.0
1.0
1.0
1.0
1.0
1.0
1.0
1.0
1.0
1.0
1.0
1.0
1.0
1.0
1.0
1.0
1.0
1.0
1.0
1.0
1.0
1.0
1.0
1.0
1.0
1.0


1263.0
0.0
0.0
0.0
0.0
0.0
0.0
66.0
0.0
0.0
0.0
0.0
0.0
0.0
0.0
0.0
0.0
1142.0
0.0
0.0
0.0
0.0
0.0
0.0
55.0
0.0
0.0
0.0
0.0
0.0


66.0
0.0
0.0
0.0
0.0
0.0
0.0
66.0
0.0
0.0
0.0
0.0
0.0
0.0
0.0
0.0
0.0
0.0
0.0
0.0
0.0
0.0
0.0
0.0
0.0
0.0
0.0
0.0
0.0
0.0

1.0
1.0
1.0
1.0
1.0
1.0
1.0
1.0
1.0
1.0
1.0
1.0
1.0
1.0
1.0
1.0
1.0
1.0
1.0
1.0
1.0
1.0
1.0
1.0
1.0
1.0
1.0
1.0
1.0
1.0


1197.0
0.0
0.0
0.0
0.0
0.0
0.0
0.0
0.0
0.0
0.0
0.0
0.0
0.0
0.0
0.0
0.0
1142.0
0.0
0.0
0.0
0.0
0.0
0.0
55.0
0.0
0.0
0.0
0.0
0.0

0.79
0.79
0.79
0.79
0.79
0.79
0.79
0.79
0.79
0.79
0.79
0.79
0.79
0.79
0.79
0.79
0.79
0.79
0.79
0.79
0.79
0.79
0.79
0.79
0.79
0.79
0.79
0.79
0.79
0.79


60.0
0.0
0.0
0.0
0.0
0.0
0.0
0.0
0.0
0.0
0.0
0.0
0.0
0.0
0.0
0.0
0.0
0.0
60.0
0.0
0.0
0.0
0.0
0.0
0.0
0.0
0.0
0.0
0.0
0.0


60.0
0.0
0.0
0.0
0.0
0.0
0.0
0.0
0.0
0.0
0.0
0.0
0.0
0.0
0.0
0.0
0.0
0.0
60.0
0.0
0.0
0.0
0.0
0.0
0.0
0.0
0.0
0.0
0.0
0.0

1.0
1.0
1.0
1.0
1.0
1.0
1.0
1.0
1.0
1.0
1.0
1.0
1.0
1.0
1.0
1.0
1.0
1.0
1.0
1.0
1.0
1.0
1.0
1.0
1.0
1.0
1.0
1.0
1.0
1.0


56.0
0.0
0.0
6.0
0.0
0.0
0.0
0.0
0.0
0.0
50.0
0.0
0.0
0.0
0.0
0.0
0.0
0.0
0.0
0.0
0.0
0.0
0.0
0.0
0.0
0.0
0.0
0.0
0.0
0.0


56.0
0.0
0.0
6.0
0.0
0.0
0.0
0.0
0.0
0.0
50.0
0.0
0.0
0.0
0.0
0.0
0.0
0.0
0.0
0.0
0.0
0.0
0.0
0.0
0.0
0.0
0.0
0.0
0.0
0.0


56.0
0.0
0.0
6.0
0.0
0.0
0.0
0.0
0.0
0.0
50.0
0.0
0.0
0.0
0.0
0.0
0.0
0.0
0.0
0.0
0.0
0.0
0.0
0.0
0.0
0.0
0.0
0.0
0.0
0.0

1.0
1.0
1.0
1.0
1.0
1.0
1.0
1.0
1.0
1.0
1.0
1.0
1.0
1.0
1.0
1.0
1.0
1.0
1.0
1.0
1.0
1.0
1.0
1.0
1.0
1.0
1.0
1.0
1.0
1.0


1583.0
0.0
0.0
0.0
0.0
0.0
0.0
0.0
4.0
0.0
10.0
26.0
0.0
747.0
0.0
0.0
0.0
0.0
0.0
0.0
0.0
0.0
504.0
0.0
0.0
0.0
0.0
0.0
292.0
0.0

0.7
0.7
0.7
0.7
0.7
0.7
0.7
0.7
0.7
0.7
0.7
0.7
0.7
0.7
0.7
0.7
0.7
0.7
0.7
0.7
0.7
0.7
0.7
0.7
0.7
0.7
0.7
0.7
0.7
0.7


769.0
0.0
0.0
0.0
0.0
0.0
0.0
0.0
0.0
0.0
0.0
22.0
0.0
747.0
0.0
0.0
0.0
0.0
0.0
0.0
0.0
0.0
0.0
0.0
0.0
0.0
0.0
0.0
0.0
0.0


769.0
0.0
0.0
0.0
0.0
0.0
0.0
0.0
0.0
0.0
0.0
22.0
0.0
747.0
0.0
0.0
0.0
0.0
0.0
0.0
0.0
0.0
0.0
0.0
0.0
0.0
0.0
0.0
0.0
0.0

0.95
0.95
0.95
0.95
0.95
0.95
0.95
0.95
0.95
0.95
0.95
0.95
0.95
0.95
0.95
0.95
0.95
0.95
0.95
0.95
0.95
0.95
0.95
0.95
0.95
0.95
0.95
0.95
0.95
0.95


10.0
0.0
0.0
0.0
0.0
0.0
0.0
0.0
0.0
0.0
10.0
0.0
0.0
0.0
0.0
0.0
0.0
0.0
0.0
0.0
0.0
0.0
0.0
0.0
0.0
0.0
0.0
0.0
0.0
0.0

0.99
0.99
0.99
0.99
0.99
0.99
0.99
0.99
0.99
0.99
0.99
0.99
0.99
0.99
0.99
0.99
0.99
0.99
0.99
0.99
0.99
0.99
0.99
0.99
0.99
0.99
0.99
0.99
0.99
0.99


504.0
0.0
0.0
0.0
0.0
0.0
0.0
0.0
0.0
0.0
0.0
0.0
0.0
0.0
0.0
0.0
0.0
0.0
0.0
0.0
0.0
0.0
504.0
0.0
0.0
0.0
0.0
0.0
0.0
0.0


504.0
0.0
0.0
0.0
0.0
0.0
0.0
0.0
0.0
0.0
0.0
0.0
0.0
0.0
0.0
0.0
0.0
0.0
0.0
0.0
0.0
0.0
504.0
0.0
0.0
0.0
0.0
0.0
0.0
0.0

1.0
1.0
1.0
1.0
1.0
1.0
1.0
1.0
1.0
1.0
1.0
1.0
1.0
1.0
1.0
1.0
1.0
1.0
1.0
1.0
1.0
1.0
1.0
1.0
1.0
1.0
1.0
1.0
1.0
1.0


292.0
0.0
0.0
0.0
0.0
0.0
0.0
0.0
0.0
0.0
0.0
0.0
0.0
0.0
0.0
0.0
0.0
0.0
0.0
0.0
0.0
0.0
0.0
0.0
0.0
0.0
0.0
0.0
292.0
0.0


292.0
0.0
0.0
0.0
0.0
0.0
0.0
0.0
0.0
0.0
0.0
0.0
0.0
0.0
0.0
0.0
0.0
0.0
0.0
0.0
0.0
0.0
0.0
0.0
0.0
0.0
0.0
0.0
292.0
0.0

1.0
1.0
1.0
1.0
1.0
1.0
1.0
1.0
1.0
1.0
1.0
1.0
1.0
1.0
1.0
1.0
1.0
1.0
1.0
1.0
1.0
1.0
1.0
1.0
1.0
1.0
1.0
1.0
1.0
1.0


8572.0
0.0
0.0
0.0
0.0
0.0
0.0
0.0
0.0
15.0
0.0
0.0
0.0
732.0
0.0
0.0
0.0
0.0
0.0
0.0
0.0
0.0
0.0
0.0
0.0
0.0
5968.0
2.0
1855.0
0.0


15.0
0.0
0.0
0.0
0.0
0.0
0.0
0.0
0.0
15.0
0.0
0.0
0.0
0.0
0.0
0.0
0.0
0.0
0.0
0.0
0.0
0.0
0.0
0.0
0.0
0.0
0.0
0.0
0.0
0.0


15.0
0.0
0.0
0.0
0.0
0.0
0.0
0.0
0.0
15.0
0.0
0.0
0.0
0.0
0.0
0.0
0.0
0.0
0.0
0.0
0.0
0.0
0.0
0.0
0.0
0.0
0.0
0.0
0.0
0.0

1.0
1.0
1.0
1.0
1.0
1.0
1.0
1.0
1.0
1.0
1.0
1.0
1.0
1.0
1.0
1.0
1.0
1.0
1.0
1.0
1.0
1.0
1.0
1.0
1.0
1.0
1.0
1.0
1.0
1.0


5242.0
0.0
0.0
0.0
0.0
0.0
0.0
0.0
0.0
0.0
0.0
0.0
0.0
0.0
0.0
0.0
0.0
0.0
0.0
0.0
0.0
0.0
0.0
0.0
0.0
0.0
5240.0
2.0
0.0
0.0


5242.0
0.0
0.0
0.0
0.0
0.0
0.0
0.0
0.0
0.0
0.0
0.0
0.0
0.0
0.0
0.0
0.0
0.0
0.0
0.0
0.0
0.0
0.0
0.0
0.0
0.0
5240.0
2.0
0.0
0.0

0.73
0.73
0.73
0.73
0.73
0.73
0.73
0.73
0.73
0.73
0.73
0.73
0.73
0.73
0.73
0.73
0.73
0.73
0.73
0.73
0.73
0.73
0.73
0.73
0.73
0.73
0.73
0.73
0.73
0.73


2583.0
0.0
0.0
0.0
0.0
0.0
0.0
0.0
0.0
0.0
0.0
0.0
0.0
0.0
0.0
0.0
0.0
0.0
0.0
0.0
0.0
0.0
0.0
0.0
0.0
0.0
728.0
0.0
1855.0
0.0


2583.0
0.0
0.0
0.0
0.0
0.0
0.0
0.0
0.0
0.0
0.0
0.0
0.0
0.0
0.0
0.0
0.0
0.0
0.0
0.0
0.0
0.0
0.0
0.0
0.0
0.0
728.0
0.0
1855.0
0.0

1.0
1.0
1.0
1.0
1.0
1.0
1.0
1.0
1.0
1.0
1.0
1.0
1.0
1.0
1.0
1.0
1.0
1.0
1.0
1.0
1.0
1.0
1.0
1.0
1.0
1.0
1.0
1.0
1.0
1.0


732.0
0.0
0.0
0.0
0.0
0.0
0.0
0.0
0.0
0.0
0.0
0.0
0.0
732.0
0.0
0.0
0.0
0.0
0.0
0.0
0.0
0.0
0.0
0.0
0.0
0.0
0.0
0.0
0.0
0.0


732.0
0.0
0.0
0.0
0.0
0.0
0.0
0.0
0.0
0.0
0.0
0.0
0.0
732.0
0.0
0.0
0.0
0.0
0.0
0.0
0.0
0.0
0.0
0.0
0.0
0.0
0.0
0.0
0.0
0.0

0.87
0.87
0.87
0.87
0.87
0.87
0.87
0.87
0.87
0.87
0.87
0.87
0.87
0.87
0.87
0.87
0.87
0.87
0.87
0.87
0.87
0.87
0.87
0.87
0.87
0.87
0.87
0.87
0.87
0.87


12.0
0.0
0.0
0.0
0.0
0.0
0.0
0.0
0.0
0.0
12.0
0.0
0.0
0.0
0.0
0.0
0.0
0.0
0.0
0.0
0.0
0.0
0.0
0.0
0.0
0.0
0.0
0.0
0.0
0.0


12.0
0.0
0.0
0.0
0.0
0.0
0.0
0.0
0.0
0.0
12.0
0.0
0.0
0.0
0.0
0.0
0.0
0.0
0.0
0.0
0.0
0.0
0.0
0.0
0.0
0.0
0.0
0.0
0.0
0.0


12.0
0.0
0.0
0.0
0.0
0.0
0.0
0.0
0.0
0.0
12.0
0.0
0.0
0.0
0.0
0.0
0.0
0.0
0.0
0.0
0.0
0.0
0.0
0.0
0.0
0.0
0.0
0.0
0.0
0.0

0.95
0.95
0.95
0.95
0.95
0.95
0.95
0.95
0.95
0.95
0.95
0.95
0.95
0.95
0.95
0.95
0.95
0.95
0.95
0.95
0.95
0.95
0.95
0.95
0.95
0.95
0.95
0.95
0.95
0.95


24234.0
0.0
0.0
0.0
0.0
0.0
0.0
0.0
0.0
0.0
0.0
0.0
0.0
0.0
0.0
24234.0
0.0
0.0
0.0
0.0
0.0
0.0
0.0
0.0
0.0
0.0
0.0
0.0
0.0
0.0


24234.0
0.0
0.0
0.0
0.0
0.0
0.0
0.0
0.0
0.0
0.0
0.0
0.0
0.0
0.0
24234.0
0.0
0.0
0.0
0.0
0.0
0.0
0.0
0.0
0.0
0.0
0.0
0.0
0.0
0.0


24234.0
0.0
0.0
0.0
0.0
0.0
0.0
0.0
0.0
0.0
0.0
0.0
0.0
0.0
0.0
24234.0
0.0
0.0
0.0
0.0
0.0
0.0
0.0
0.0
0.0
0.0
0.0
0.0
0.0
0.0

1.0
1.0
1.0
1.0
1.0
1.0
1.0
1.0
1.0
1.0
1.0
1.0
1.0
1.0
1.0
1.0
1.0
1.0
1.0
1.0
1.0
1.0
1.0
1.0
1.0
1.0
1.0
1.0
1.0
1.0


430.0
0.0
0.0
0.0
0.0
0.0
0.0
0.0
0.0
0.0
0.0
0.0
0.0
0.0
0.0
0.0
0.0
0.0
0.0
0.0
131.0
0.0
0.0
0.0
0.0
299.0
0.0
0.0
0.0
0.0


430.0
0.0
0.0
0.0
0.0
0.0
0.0
0.0
0.0
0.0
0.0
0.0
0.0
0.0
0.0
0.0
0.0
0.0
0.0
0.0
131.0
0.0
0.0
0.0
0.0
299.0
0.0
0.0
0.0
0.0


430.0
0.0
0.0
0.0
0.0
0.0
0.0
0.0
0.0
0.0
0.0
0.0
0.0
0.0
0.0
0.0
0.0
0.0
0.0
0.0
131.0
0.0
0.0
0.0
0.0
299.0
0.0
0.0
0.0
0.0

1.0
1.0
1.0
1.0
1.0
1.0
1.0
1.0
1.0
1.0
1.0
1.0
1.0
1.0
1.0
1.0
1.0
1.0
1.0
1.0
1.0
1.0
1.0
1.0
1.0
1.0
1.0
1.0
1.0
1.0


86.0
0.0
0.0
0.0
0.0
0.0
0.0
0.0
0.0
0.0
0.0
0.0
0.0
0.0
0.0
0.0
0.0
0.0
7.0
0.0
0.0
0.0
0.0
0.0
0.0
0.0
0.0
79.0
0.0
0.0


86.0
0.0
0.0
0.0
0.0
0.0
0.0
0.0
0.0
0.0
0.0
0.0
0.0
0.0
0.0
0.0
0.0
0.0
7.0
0.0
0.0
0.0
0.0
0.0
0.0
0.0
0.0
79.0
0.0
0.0


86.0
0.0
0.0
0.0
0.0
0.0
0.0
0.0
0.0
0.0
0.0
0.0
0.0
0.0
0.0
0.0
0.0
0.0
7.0
0.0
0.0
0.0
0.0
0.0
0.0
0.0
0.0
79.0
0.0
0.0

0.84
0.84
0.84
0.84
0.84
0.84
0.84
0.84
0.84
0.84
0.84
0.84
0.84
0.84
0.84
0.84
0.84
0.84
0.84
0.84
0.84
0.84
0.84
0.84
0.84
0.84
0.84
0.84
0.84
0.84


125.0
12.0
0.0
0.0
0.0
0.0
0.0
21.0
0.0
0.0
75.0
7.0
0.0
6.0
0.0
0.0
0.0
0.0
0.0
0.0
0.0
0.0
0.0
0.0
4.0
0.0
0.0
0.0
0.0
0.0


115.0
12.0
0.0
0.0
0.0
0.0
0.0
21.0
0.0
0.0
75.0
7.0
0.0
0.0
0.0
0.0
0.0
0.0
0.0
0.0
0.0
0.0
0.0
0.0
0.0
0.0
0.0
0.0
0.0
0.0


115.0
12.0
0.0
0.0
0.0
0.0
0.0
21.0
0.0
0.0
75.0
7.0
0.0
0.0
0.0
0.0
0.0
0.0
0.0
0.0
0.0
0.0
0.0
0.0
0.0
0.0
0.0
0.0
0.0
0.0


115.0
12.0
0.0
0.0
0.0
0.0
0.0
21.0
0.0
0.0
75.0
7.0
0.0
0.0
0.0
0.0
0.0
0.0
0.0
0.0
0.0
0.0
0.0
0.0
0.0
0.0
0.0
0.0
0.0
0.0

1.0
1.0
1.0
1.0
1.0
1.0
1.0
1.0
1.0
1.0
1.0
1.0
1.0
1.0
1.0
1.0
1.0
1.0
1.0
1.0
1.0
1.0
1.0
1.0
1.0
1.0
1.0
1.0
1.0
1.0


10.0
0.0
0.0
0.0
0.0
0.0
0.0
0.0
0.0
0.0
0.0
0.0
0.0
6.0
0.0
0.0
0.0
0.0
0.0
0.0
0.0
0.0
0.0
0.0
4.0
0.0
0.0
0.0
0.0
0.0

0.74
0.74
0.74
0.74
0.74
0.74
0.74
0.74
0.74
0.74
0.74
0.74
0.74
0.74
0.74
0.74
0.74
0.74
0.74
0.74
0.74
0.74
0.74
0.74
0.74
0.74
0.74
0.74
0.74
0.74


4.0
0.0
0.0
0.0
0.0
0.0
0.0
0.0
0.0
0.0
0.0
0.0
0.0
0.0
0.0
0.0
0.0
0.0
0.0
0.0
0.0
0.0
0.0
0.0
4.0
0.0
0.0
0.0
0.0
0.0


4.0
0.0
0.0
0.0
0.0
0.0
0.0
0.0
0.0
0.0
0.0
0.0
0.0
0.0
0.0
0.0
0.0
0.0
0.0
0.0
0.0
0.0
0.0
0.0
4.0
0.0
0.0
0.0
0.0
0.0

0.96
0.96
0.96
0.96
0.96
0.96
0.96
0.96
0.96
0.96
0.96
0.96
0.96
0.96
0.96
0.96
0.96
0.96
0.96
0.96
0.96
0.96
0.96
0.96
0.96
0.96
0.96
0.96
0.96
0.96


473.0
0.0
0.0
0.0
0.0
0.0
0.0
0.0
0.0
0.0
0.0
0.0
0.0
0.0
0.0
0.0
0.0
0.0
308.0
139.0
22.0
0.0
0.0
0.0
0.0
0.0
0.0
0.0
4.0
0.0


447.0
0.0
0.0
0.0
0.0
0.0
0.0
0.0
0.0
0.0
0.0
0.0
0.0
0.0
0.0
0.0
0.0
0.0
308.0
139.0
0.0
0.0
0.0
0.0
0.0
0.0
0.0
0.0
0.0
0.0


447.0
0.0
0.0
0.0
0.0
0.0
0.0
0.0
0.0
0.0
0.0
0.0
0.0
0.0
0.0
0.0
0.0
0.0
308.0
139.0
0.0
0.0
0.0
0.0
0.0
0.0
0.0
0.0
0.0
0.0


447.0
0.0
0.0
0.0
0.0
0.0
0.0
0.0
0.0
0.0
0.0
0.0
0.0
0.0
0.0
0.0
0.0
0.0
308.0
139.0
0.0
0.0
0.0
0.0
0.0
0.0
0.0
0.0
0.0
0.0

1.0
1.0
1.0
1.0
1.0
1.0
1.0
1.0
1.0
1.0
1.0
1.0
1.0
1.0
1.0
1.0
1.0
1.0
1.0
1.0
1.0
1.0
1.0
1.0
1.0
1.0
1.0
1.0
1.0
1.0


22.0
0.0
0.0
0.0
0.0
0.0
0.0
0.0
0.0
0.0
0.0
0.0
0.0
0.0
0.0
0.0
0.0
0.0
0.0
0.0
22.0
0.0
0.0
0.0
0.0
0.0
0.0
0.0
0.0
0.0


22.0
0.0
0.0
0.0
0.0
0.0
0.0
0.0
0.0
0.0
0.0
0.0
0.0
0.0
0.0
0.0
0.0
0.0
0.0
0.0
22.0
0.0
0.0
0.0
0.0
0.0
0.0
0.0
0.0
0.0


22.0
0.0
0.0
0.0
0.0
0.0
0.0
0.0
0.0
0.0
0.0
0.0
0.0
0.0
0.0
0.0
0.0
0.0
0.0
0.0
22.0
0.0
0.0
0.0
0.0
0.0
0.0
0.0
0.0
0.0

0.77
0.77
0.77
0.77
0.77
0.77
0.77
0.77
0.77
0.77
0.77
0.77
0.77
0.77
0.77
0.77
0.77
0.77
0.77
0.77
0.77
0.77
0.77
0.77
0.77
0.77
0.77
0.77
0.77
0.77


4.0
0.0
0.0
0.0
0.0
0.0
0.0
0.0
0.0
0.0
0.0
0.0
0.0
0.0
0.0
0.0
0.0
0.0
0.0
0.0
0.0
0.0
0.0
0.0
0.0
0.0
0.0
0.0
4.0
0.0


4.0
0.0
0.0
0.0
0.0
0.0
0.0
0.0
0.0
0.0
0.0
0.0
0.0
0.0
0.0
0.0
0.0
0.0
0.0
0.0
0.0
0.0
0.0
0.0
0.0
0.0
0.0
0.0
4.0
0.0


4.0
0.0
0.0
0.0
0.0
0.0
0.0
0.0
0.0
0.0
0.0
0.0
0.0
0.0
0.0
0.0
0.0
0.0
0.0
0.0
0.0
0.0
0.0
0.0
0.0
0.0
0.0
0.0
4.0
0.0

1.0
1.0
1.0
1.0
1.0
1.0
1.0
1.0
1.0
1.0
1.0
1.0
1.0
1.0
1.0
1.0
1.0
1.0
1.0
1.0
1.0
1.0
1.0
1.0
1.0
1.0
1.0
1.0
1.0
1.0


5937.0
0.0
0.0
0.0
0.0
0.0
0.0
0.0
0.0
0.0
0.0
0.0
0.0
0.0
0.0
0.0
5937.0
0.0
0.0
0.0
0.0
0.0
0.0
0.0
0.0
0.0
0.0
0.0
0.0
0.0


5937.0
0.0
0.0
0.0
0.0
0.0
0.0
0.0
0.0
0.0
0.0
0.0
0.0
0.0
0.0
0.0
5937.0
0.0
0.0
0.0
0.0
0.0
0.0
0.0
0.0
0.0
0.0
0.0
0.0
0.0


5937.0
0.0
0.0
0.0
0.0
0.0
0.0
0.0
0.0
0.0
0.0
0.0
0.0
0.0
0.0
0.0
5937.0
0.0
0.0
0.0
0.0
0.0
0.0
0.0
0.0
0.0
0.0
0.0
0.0
0.0

0.73
0.73
0.73
0.73
0.73
0.73
0.73
0.73
0.73
0.73
0.73
0.73
0.73
0.73
0.73
0.73
0.73
0.73
0.73
0.73
0.73
0.73
0.73
0.73
0.73
0.73
0.73
0.73
0.73
0.73


5845.0
0.0
0.0
0.0
0.0
0.0
0.0
0.0
0.0
0.0
0.0
0.0
0.0
0.0
0.0
0.0
5845.0
0.0
0.0
0.0
0.0
0.0
0.0
0.0
0.0
0.0
0.0
0.0
0.0
0.0


5845.0
0.0
0.0
0.0
0.0
0.0
0.0
0.0
0.0
0.0
0.0
0.0
0.0
0.0
0.0
0.0
5845.0
0.0
0.0
0.0
0.0
0.0
0.0
0.0
0.0
0.0
0.0
0.0
0.0
0.0

1.0
1.0
1.0
1.0
1.0
1.0
1.0
1.0
1.0
1.0
1.0
1.0
1.0
1.0
1.0
1.0
1.0
1.0
1.0
1.0
1.0
1.0
1.0
1.0
1.0
1.0
1.0
1.0
1.0
1.0


385643.0
32.0
2644.0
71.0
1119.0
0.0
0.0
3315.0
1.0
0.0
757.0
25249.0
0.0
0.0
0.0
0.0
0.0
0.0
0.0
0.0
199008.0
0.0
0.0
0.0
153443.0
0.0
0.0
0.0
4.0
0.0


385643.0
32.0
2644.0
71.0
1119.0
0.0
0.0
3315.0
1.0
0.0
757.0
25249.0
0.0
0.0
0.0
0.0
0.0
0.0
0.0
0.0
199008.0
0.0
0.0
0.0
153443.0
0.0
0.0
0.0
4.0
0.0


385643.0
32.0
2644.0
71.0
1119.0
0.0
0.0
3315.0
1.0
0.0
757.0
25249.0
0.0
0.0
0.0
0.0
0.0
0.0
0.0
0.0
199008.0
0.0
0.0
0.0
153443.0
0.0
0.0
0.0
4.0
0.0


384473.0
1.0
2644.0
71.0
1119.0
0.0
0.0
3315.0
1.0
0.0
6.0
25249.0
0.0
0.0
0.0
0.0
0.0
0.0
0.0
0.0
198620.0
0.0
0.0
0.0
153443.0
0.0
0.0
0.0
4.0
0.0


345553.0
0.0
2497.0
0.0
0.0
0.0
0.0
4.0
1.0
0.0
6.0
19444.0
0.0
0.0
0.0
0.0
0.0
0.0
0.0
0.0
198620.0
0.0
0.0
0.0
124977.0
0.0
0.0
0.0
4.0
0.0


345553.0
0.0
2497.0
0.0
0.0
0.0
0.0
4.0
1.0
0.0
6.0
19444.0
0.0
0.0
0.0
0.0
0.0
0.0
0.0
0.0
198620.0
0.0
0.0
0.0
124977.0
0.0
0.0
0.0
4.0
0.0

1.0
1.0
1.0
1.0
1.0
1.0
1.0
1.0
1.0
1.0
1.0
1.0
1.0
1.0
1.0
1.0
1.0
1.0
1.0
1.0
1.0
1.0
1.0
1.0
1.0
1.0
1.0
1.0
1.0
1.0


6801.0
0.0
135.0
0.0
0.0
0.0
0.0
3311.0
0.0
0.0
0.0
3049.0
0.0
0.0
0.0
0.0
0.0
0.0
0.0
0.0
0.0
0.0
0.0
0.0
306.0
0.0
0.0
0.0
0.0
0.0


6801.0
0.0
135.0
0.0
0.0
0.0
0.0
3311.0
0.0
0.0
0.0
3049.0
0.0
0.0
0.0
0.0
0.0
0.0
0.0
0.0
0.0
0.0
0.0
0.0
306.0
0.0
0.0
0.0
0.0
0.0

1.0
1.0
1.0
1.0
1.0
1.0
1.0
1.0
1.0
1.0
1.0
1.0
1.0
1.0
1.0
1.0
1.0
1.0
1.0
1.0
1.0
1.0
1.0
1.0
1.0
1.0
1.0
1.0
1.0
1.0


30208.0
1.0
0.0
71.0
1119.0
0.0
0.0
0.0
0.0
0.0
0.0
857.0
0.0
0.0
0.0
0.0
0.0
0.0
0.0
0.0
0.0
0.0
0.0
0.0
28160.0
0.0
0.0
0.0
0.0
0.0

1.0
1.0
1.0
1.0
1.0
1.0
1.0
1.0
1.0
1.0
1.0
1.0
1.0
1.0
1.0
1.0
1.0
1.0
1.0
1.0
1.0
1.0
1.0
1.0
1.0
1.0
1.0
1.0
1.0
1.0


1909.0
1.0
0.0
65.0
1039.0
0.0
0.0
0.0
0.0
0.0
0.0
804.0
0.0
0.0
0.0
0.0
0.0
0.0
0.0
0.0
0.0
0.0
0.0
0.0
0.0
0.0
0.0
0.0
0.0
0.0

0.7
0.7
0.7
0.7
0.7
0.7
0.7
0.7
0.7
0.7
0.7
0.7
0.7
0.7
0.7
0.7
0.7
0.7
0.7
0.7
0.7
0.7
0.7
0.7
0.7
0.7
0.7
0.7
0.7
0.7


28236.0
0.0
0.0
4.0
26.0
0.0
0.0
0.0
0.0
0.0
0.0
46.0
0.0
0.0
0.0
0.0
0.0
0.0
0.0
0.0
0.0
0.0
0.0
0.0
28160.0
0.0
0.0
0.0
0.0
0.0

1.0
1.0
1.0
1.0
1.0
1.0
1.0
1.0
1.0
1.0
1.0
1.0
1.0
1.0
1.0
1.0
1.0
1.0
1.0
1.0
1.0
1.0
1.0
1.0
1.0
1.0
1.0
1.0
1.0
1.0


1911.0
0.0
12.0
0.0
0.0
0.0
0.0
0.0
0.0
0.0
0.0
1899.0
0.0
0.0
0.0
0.0
0.0
0.0
0.0
0.0
0.0
0.0
0.0
0.0
0.0
0.0
0.0
0.0
0.0
0.0

0.87
0.87
0.87
0.87
0.87
0.87
0.87
0.87
0.87
0.87
0.87
0.87
0.87
0.87
0.87
0.87
0.87
0.87
0.87
0.87
0.87
0.87
0.87
0.87
0.87
0.87
0.87
0.87
0.87
0.87


1170.0
31.0
0.0
0.0
0.0
0.0
0.0
0.0
0.0
0.0
751.0
0.0
0.0
0.0
0.0
0.0
0.0
0.0
0.0
0.0
388.0
0.0
0.0
0.0
0.0
0.0
0.0
0.0
0.0
0.0


1170.0
31.0
0.0
0.0
0.0
0.0
0.0
0.0
0.0
0.0
751.0
0.0
0.0
0.0
0.0
0.0
0.0
0.0
0.0
0.0
388.0
0.0
0.0
0.0
0.0
0.0
0.0
0.0
0.0
0.0


1146.0
31.0
0.0
0.0
0.0
0.0
0.0
0.0
0.0
0.0
727.0
0.0
0.0
0.0
0.0
0.0
0.0
0.0
0.0
0.0
388.0
0.0
0.0
0.0
0.0
0.0
0.0
0.0
0.0
0.0

1.0
1.0
1.0
1.0
1.0
1.0
1.0
1.0
1.0
1.0
1.0
1.0
1.0
1.0
1.0
1.0
1.0
1.0
1.0
1.0
1.0
1.0
1.0
1.0
1.0
1.0
1.0
1.0
1.0
1.0


24.0
0.0
0.0
0.0
0.0
0.0
0.0
0.0
0.0
0.0
24.0
0.0
0.0
0.0
0.0
0.0
0.0
0.0
0.0
0.0
0.0
0.0
0.0
0.0
0.0
0.0
0.0
0.0
0.0
0.0

0.92
0.92
0.92
0.92
0.92
0.92
0.92
0.92
0.92
0.92
0.92
0.92
0.92
0.92
0.92
0.92
0.92
0.92
0.92
0.92
0.92
0.92
0.92
0.92
0.92
0.92
0.92
0.92
0.92
0.92


1061.0
0.0
18.0
0.0
8.0
0.0
9.0
20.0
53.0
28.0
15.0
3.0
0.0
0.0
5.0
0.0
0.0
57.0
0.0
0.0
0.0
357.0
78.0
0.0
0.0
339.0
43.0
28.0
0.0
0.0


1061.0
0.0
18.0
0.0
8.0
0.0
9.0
20.0
53.0
28.0
15.0
3.0
0.0
0.0
5.0
0.0
0.0
57.0
0.0
0.0
0.0
357.0
78.0
0.0
0.0
339.0
43.0
28.0
0.0
0.0


1061.0
0.0
18.0
0.0
8.0
0.0
9.0
20.0
53.0
28.0
15.0
3.0
0.0
0.0
5.0
0.0
0.0
57.0
0.0
0.0
0.0
357.0
78.0
0.0
0.0
339.0
43.0
28.0
0.0
0.0

1.0
1.0
1.0
1.0
1.0
1.0
1.0
1.0
1.0
1.0
1.0
1.0
1.0
1.0
1.0
1.0
1.0
1.0
1.0
1.0
1.0
1.0
1.0
1.0
1.0
1.0
1.0
1.0
1.0
1.0


375.0
0.0
0.0
0.0
0.0
0.0
0.0
18.0
0.0
0.0
0.0
0.0
0.0
0.0
0.0
0.0
0.0
0.0
0.0
0.0
0.0
357.0
0.0
0.0
0.0
0.0
0.0
0.0
0.0
0.0


375.0
0.0
0.0
0.0
0.0
0.0
0.0
18.0
0.0
0.0
0.0
0.0
0.0
0.0
0.0
0.0
0.0
0.0
0.0
0.0
0.0
357.0
0.0
0.0
0.0
0.0
0.0
0.0
0.0
0.0


375.0
0.0
0.0
0.0
0.0
0.0
0.0
18.0
0.0
0.0
0.0
0.0
0.0
0.0
0.0
0.0
0.0
0.0
0.0
0.0
0.0
357.0
0.0
0.0
0.0
0.0
0.0
0.0
0.0
0.0

0.93
0.93
0.93
0.93
0.93
0.93
0.93
0.93
0.93
0.93
0.93
0.93
0.93
0.93
0.93
0.93
0.93
0.93
0.93
0.93
0.93
0.93
0.93
0.93
0.93
0.93
0.93
0.93
0.93
0.93


78.0
0.0
0.0
0.0
0.0
0.0
0.0
0.0
0.0
0.0
0.0
0.0
0.0
0.0
0.0
0.0
0.0
0.0
0.0
0.0
0.0
0.0
78.0
0.0
0.0
0.0
0.0
0.0
0.0
0.0


78.0
0.0
0.0
0.0
0.0
0.0
0.0
0.0
0.0
0.0
0.0
0.0
0.0
0.0
0.0
0.0
0.0
0.0
0.0
0.0
0.0
0.0
78.0
0.0
0.0
0.0
0.0
0.0
0.0
0.0


78.0
0.0
0.0
0.0
0.0
0.0
0.0
0.0
0.0
0.0
0.0
0.0
0.0
0.0
0.0
0.0
0.0
0.0
0.0
0.0
0.0
0.0
78.0
0.0
0.0
0.0
0.0
0.0
0.0
0.0

0.76
0.76
0.76
0.76
0.76
0.76
0.76
0.76
0.76
0.76
0.76
0.76
0.76
0.76
0.76
0.76
0.76
0.76
0.76
0.76
0.76
0.76
0.76
0.76
0.76
0.76
0.76
0.76
0.76
0.76


57.0
0.0
0.0
0.0
0.0
0.0
0.0
0.0
0.0
0.0
0.0
0.0
0.0
0.0
0.0
0.0
0.0
57.0
0.0
0.0
0.0
0.0
0.0
0.0
0.0
0.0
0.0
0.0
0.0
0.0


57.0
0.0
0.0
0.0
0.0
0.0
0.0
0.0
0.0
0.0
0.0
0.0
0.0
0.0
0.0
0.0
0.0
57.0
0.0
0.0
0.0
0.0
0.0
0.0
0.0
0.0
0.0
0.0
0.0
0.0

0.88
0.88
0.88
0.88
0.88
0.88
0.88
0.88
0.88
0.88
0.88
0.88
0.88
0.88
0.88
0.88
0.88
0.88
0.88
0.88
0.88
0.88
0.88
0.88
0.88
0.88
0.88
0.88
0.88
0.88


20.0
0.0
0.0
0.0
0.0
0.0
0.0
0.0
0.0
0.0
0.0
0.0
0.0
0.0
0.0
0.0
0.0
0.0
0.0
20.0
0.0
0.0
0.0
0.0
0.0
0.0
0.0
0.0
0.0
0.0


20.0
0.0
0.0
0.0
0.0
0.0
0.0
0.0
0.0
0.0
0.0
0.0
0.0
0.0
0.0
0.0
0.0
0.0
0.0
20.0
0.0
0.0
0.0
0.0
0.0
0.0
0.0
0.0
0.0
0.0


20.0
0.0
0.0
0.0
0.0
0.0
0.0
0.0
0.0
0.0
0.0
0.0
0.0
0.0
0.0
0.0
0.0
0.0
0.0
20.0
0.0
0.0
0.0
0.0
0.0
0.0
0.0
0.0
0.0
0.0


20.0
0.0
0.0
0.0
0.0
0.0
0.0
0.0
0.0
0.0
0.0
0.0
0.0
0.0
0.0
0.0
0.0
0.0
0.0
20.0
0.0
0.0
0.0
0.0
0.0
0.0
0.0
0.0
0.0
0.0


20.0
0.0
0.0
0.0
0.0
0.0
0.0
0.0
0.0
0.0
0.0
0.0
0.0
0.0
0.0
0.0
0.0
0.0
0.0
20.0
0.0
0.0
0.0
0.0
0.0
0.0
0.0
0.0
0.0
0.0


20.0
0.0
0.0
0.0
0.0
0.0
0.0
0.0
0.0
0.0
0.0
0.0
0.0
0.0
0.0
0.0
0.0
0.0
0.0
20.0
0.0
0.0
0.0
0.0
0.0
0.0
0.0
0.0
0.0
0.0

1.0
1.0
1.0
1.0
1.0
1.0
1.0
1.0
1.0
1.0
1.0
1.0
1.0
1.0
1.0
1.0
1.0
1.0
1.0
1.0
1.0
1.0
1.0
1.0
1.0
1.0
1.0
1.0
1.0
1.0


565.0
0.0
0.0
0.0
0.0
0.0
0.0
47.0
448.0
70.0
0.0
0.0
0.0
0.0
0.0
0.0
0.0
0.0
0.0
0.0
0.0
0.0
0.0
0.0
0.0
0.0
0.0
0.0
0.0
0.0


565.0
0.0
0.0
0.0
0.0
0.0
0.0
47.0
448.0
70.0
0.0
0.0
0.0
0.0
0.0
0.0
0.0
0.0
0.0
0.0
0.0
0.0
0.0
0.0
0.0
0.0
0.0
0.0
0.0
0.0


565.0
0.0
0.0
0.0
0.0
0.0
0.0
47.0
448.0
70.0
0.0
0.0
0.0
0.0
0.0
0.0
0.0
0.0
0.0
0.0
0.0
0.0
0.0
0.0
0.0
0.0
0.0
0.0
0.0
0.0

0.73
0.73
0.73
0.73
0.73
0.73
0.73
0.73
0.73
0.73
0.73
0.73
0.73
0.73
0.73
0.73
0.73
0.73
0.73
0.73
0.73
0.73
0.73
0.73
0.73
0.73
0.73
0.73
0.73
0.73


2216.0
169.0
100.0
71.0
103.0
254.0
450.0
132.0
64.0
417.0
215.0
110.0
11.0
0.0
0.0
0.0
0.0
13.0
12.0
20.0
9.0
4.0
10.0
11.0
35.0
0.0
0.0
5.0
0.0
1.0


2216.0
169.0
100.0
71.0
103.0
254.0
450.0
132.0
64.0
417.0
215.0
110.0
11.0
0.0
0.0
0.0
0.0
13.0
12.0
20.0
9.0
4.0
10.0
11.0
35.0
0.0
0.0
5.0
0.0
1.0


2216.0
169.0
100.0
71.0
103.0
254.0
450.0
132.0
64.0
417.0
215.0
110.0
11.0
0.0
0.0
0.0
0.0
13.0
12.0
20.0
9.0
4.0
10.0
11.0
35.0
0.0
0.0
5.0
0.0
1.0


2216.0
169.0
100.0
71.0
103.0
254.0
450.0
132.0
64.0
417.0
215.0
110.0
11.0
0.0
0.0
0.0
0.0
13.0
12.0
20.0
9.0
4.0
10.0
11.0
35.0
0.0
0.0
5.0
0.0
1.0


2216.0
169.0
100.0
71.0
103.0
254.0
450.0
132.0
64.0
417.0
215.0
110.0
11.0
0.0
0.0
0.0
0.0
13.0
12.0
20.0
9.0
4.0
10.0
11.0
35.0
0.0
0.0
5.0
0.0
1.0


2216.0
169.0
100.0
71.0
103.0
254.0
450.0
132.0
64.0
417.0
215.0
110.0
11.0
0.0
0.0
0.0
0.0
13.0
12.0
20.0
9.0
4.0
10.0
11.0
35.0
0.0
0.0
5.0
0.0
1.0

1.0
1.0
1.0
1.0
1.0
1.0
1.0
1.0
1.0
1.0
1.0
1.0
1.0
1.0
1.0
1.0
1.0
1.0
1.0
1.0
1.0
1.0
1.0
1.0
1.0
1.0
1.0
1.0
1.0
1.0


1856.0
134.0
89.0
67.0
78.0
212.0
447.0
110.0
40.0
382.0
192.0
64.0
11.0
0.0
0.0
0.0
0.0
0.0
1.0
0.0
3.0
4.0
10.0
11.0
0.0
0.0
0.0
0.0
0.0
1.0

1.0
1.0
1.0
1.0
1.0
1.0
1.0
1.0
1.0
1.0
1.0
1.0
1.0
1.0
1.0
1.0
1.0
1.0
1.0
1.0
1.0
1.0
1.0
1.0
1.0
1.0
1.0
1.0
1.0
1.0


283.0
10.0
1.0
0.0
1.0
9.0
14.0
33.0
10.0
9.0
12.0
11.0
0.0
0.0
0.0
0.0
3.0
16.0
12.0
1.0
0.0
0.0
4.0
0.0
36.0
24.0
41.0
0.0
9.0
27.0


283.0
10.0
1.0
0.0
1.0
9.0
14.0
33.0
10.0
9.0
12.0
11.0
0.0
0.0
0.0
0.0
3.0
16.0
12.0
1.0
0.0
0.0
4.0
0.0
36.0
24.0
41.0
0.0
9.0
27.0

0.99
0.99
0.99
0.99
0.99
0.99
0.99
0.99
0.99
0.99
0.99
0.99
0.99
0.99
0.99
0.99
0.99
0.99
0.99
0.99
0.99
0.99
0.99
0.99
0.99
0.99
0.99
0.99
0.99
0.99


224.0
8.0
0.0
0.0
1.0
9.0
14.0
33.0
10.0
9.0
12.0
4.0
0.0
0.0
0.0
0.0
3.0
16.0
12.0
1.0
0.0
0.0
4.0
0.0
27.0
19.0
27.0
0.0
9.0
6.0


212.0
8.0
0.0
0.0
1.0
9.0
14.0
33.0
10.0
9.0
12.0
4.0
0.0
0.0
0.0
0.0
3.0
16.0
12.0
1.0
0.0
0.0
4.0
0.0
27.0
7.0
27.0
0.0
9.0
6.0


212.0
8.0
0.0
0.0
1.0
9.0
14.0
33.0
10.0
9.0
12.0
4.0
0.0
0.0
0.0
0.0
3.0
16.0
12.0
1.0
0.0
0.0
4.0
0.0
27.0
7.0
27.0
0.0
9.0
6.0


212.0
8.0
0.0
0.0
1.0
9.0
14.0
33.0
10.0
9.0
12.0
4.0
0.0
0.0
0.0
0.0
3.0
16.0
12.0
1.0
0.0
0.0
4.0
0.0
27.0
7.0
27.0
0.0
9.0
6.0

0.84
0.84
0.84
0.84
0.84
0.84
0.84
0.84
0.84
0.84
0.84
0.84
0.84
0.84
0.84
0.84
0.84
0.84
0.84
0.84
0.84
0.84
0.84
0.84
0.84
0.84
0.84
0.84
0.84
0.84


47.0
4.0
0.0
0.0
1.0
5.0
7.0
11.0
5.0
4.0
7.0
3.0
0.0
0.0
0.0
0.0
0.0
0.0
0.0
0.0
0.0
0.0
0.0
0.0
0.0
0.0
0.0
0.0
0.0
0.0

1.0
1.0
1.0
1.0
1.0
1.0
1.0
1.0
1.0
1.0
1.0
1.0
1.0
1.0
1.0
1.0
1.0
1.0
1.0
1.0
1.0
1.0
1.0
1.0
1.0
1.0
1.0
1.0
1.0
1.0


57.0
0.0
0.0
0.0
0.0
0.0
0.0
0.0
0.0
0.0
0.0
0.0
0.0
0.0
0.0
0.0
3.0
9.0
0.0
0.0
0.0
0.0
4.0
0.0
3.0
5.0
27.0
0.0
0.0
6.0

0.83
0.83
0.83
0.83
0.83
0.83
0.83
0.83
0.83
0.83
0.83
0.83
0.83
0.83
0.83
0.83
0.83
0.83
0.83
0.83
0.83
0.83
0.83
0.83
0.83
0.83
0.83
0.83
0.83
0.83


48.0
0.0
0.0
0.0
0.0
0.0
0.0
0.0
0.0
0.0
0.0
0.0
0.0
0.0
0.0
0.0
0.0
7.0
12.0
1.0
0.0
0.0
0.0
0.0
24.0
2.0
0.0
0.0
2.0
0.0

0.8
0.8
0.8
0.8
0.8
0.8
0.8
0.8
0.8
0.8
0.8
0.8
0.8
0.8
0.8
0.8
0.8
0.8
0.8
0.8
0.8
0.8
0.8
0.8
0.8
0.8
0.8
0.8
0.8
0.8


7.0
0.0
0.0
0.0
0.0
0.0
0.0
0.0
0.0
0.0
0.0
0.0
0.0
0.0
0.0
0.0
0.0
0.0
0.0
0.0
0.0
0.0
0.0
0.0
0.0
0.0
0.0
0.0
7.0
0.0

0.92
0.92
0.92
0.92
0.92
0.92
0.92
0.92
0.92
0.92
0.92
0.92
0.92
0.92
0.92
0.92
0.92
0.92
0.92
0.92
0.92
0.92
0.92
0.92
0.92
0.92
0.92
0.92
0.92
0.92


12.0
0.0
0.0
0.0
0.0
0.0
0.0
0.0
0.0
0.0
0.0
0.0
0.0
0.0
0.0
0.0
0.0
0.0
0.0
0.0
0.0
0.0
0.0
0.0
0.0
12.0
0.0
0.0
0.0
0.0


12.0
0.0
0.0
0.0
0.0
0.0
0.0
0.0
0.0
0.0
0.0
0.0
0.0
0.0
0.0
0.0
0.0
0.0
0.0
0.0
0.0
0.0
0.0
0.0
0.0
12.0
0.0
0.0
0.0
0.0


12.0
0.0
0.0
0.0
0.0
0.0
0.0
0.0
0.0
0.0
0.0
0.0
0.0
0.0
0.0
0.0
0.0
0.0
0.0
0.0
0.0
0.0
0.0
0.0
0.0
12.0
0.0
0.0
0.0
0.0


12.0
0.0
0.0
0.0
0.0
0.0
0.0
0.0
0.0
0.0
0.0
0.0
0.0
0.0
0.0
0.0
0.0
0.0
0.0
0.0
0.0
0.0
0.0
0.0
0.0
12.0
0.0
0.0
0.0
0.0

0.87
0.87
0.87
0.87
0.87
0.87
0.87
0.87
0.87
0.87
0.87
0.87
0.87
0.87
0.87
0.87
0.87
0.87
0.87
0.87
0.87
0.87
0.87
0.87
0.87
0.87
0.87
0.87
0.87
0.87


19.0
2.0
1.0
0.0
0.0
0.0
0.0
0.0
0.0
0.0
0.0
7.0
0.0
0.0
0.0
0.0
0.0
0.0
0.0
0.0
0.0
0.0
0.0
0.0
9.0
0.0
0.0
0.0
0.0
0.0


19.0
2.0
1.0
0.0
0.0
0.0
0.0
0.0
0.0
0.0
0.0
7.0
0.0
0.0
0.0
0.0
0.0
0.0
0.0
0.0
0.0
0.0
0.0
0.0
9.0
0.0
0.0
0.0
0.0
0.0


19.0
2.0
1.0
0.0
0.0
0.0
0.0
0.0
0.0
0.0
0.0
7.0
0.0
0.0
0.0
0.0
0.0
0.0
0.0
0.0
0.0
0.0
0.0
0.0
9.0
0.0
0.0
0.0
0.0
0.0

0.9
0.9
0.9
0.9
0.9
0.9
0.9
0.9
0.9
0.9
0.9
0.9
0.9
0.9
0.9
0.9
0.9
0.9
0.9
0.9
0.9
0.9
0.9
0.9
0.9
0.9
0.9
0.9
0.9
0.9


9.0
0.0
0.0
0.0
0.0
0.0
0.0
0.0
0.0
0.0
0.0
0.0
0.0
0.0
0.0
0.0
0.0
0.0
0.0
0.0
0.0
0.0
0.0
0.0
9.0
0.0
0.0
0.0
0.0
0.0

0.84
0.84
0.84
0.84
0.84
0.84
0.84
0.84
0.84
0.84
0.84
0.84
0.84
0.84
0.84
0.84
0.84
0.84
0.84
0.84
0.84
0.84
0.84
0.84
0.84
0.84
0.84
0.84
0.84
0.84


35.0
0.0
0.0
0.0
0.0
0.0
0.0
0.0
0.0
0.0
0.0
0.0
0.0
0.0
0.0
0.0
0.0
0.0
0.0
0.0
0.0
0.0
0.0
0.0
0.0
0.0
14.0
0.0
0.0
21.0


21.0
0.0
0.0
0.0
0.0
0.0
0.0
0.0
0.0
0.0
0.0
0.0
0.0
0.0
0.0
0.0
0.0
0.0
0.0
0.0
0.0
0.0
0.0
0.0
0.0
0.0
0.0
0.0
0.0
21.0


21.0
0.0
0.0
0.0
0.0
0.0
0.0
0.0
0.0
0.0
0.0
0.0
0.0
0.0
0.0
0.0
0.0
0.0
0.0
0.0
0.0
0.0
0.0
0.0
0.0
0.0
0.0
0.0
0.0
21.0


21.0
0.0
0.0
0.0
0.0
0.0
0.0
0.0
0.0
0.0
0.0
0.0
0.0
0.0
0.0
0.0
0.0
0.0
0.0
0.0
0.0
0.0
0.0
0.0
0.0
0.0
0.0
0.0
0.0
21.0

0.76
0.76
0.76
0.76
0.76
0.76
0.76
0.76
0.76
0.76
0.76
0.76
0.76
0.76
0.76
0.76
0.76
0.76
0.76
0.76
0.76
0.76
0.76
0.76
0.76
0.76
0.76
0.76
0.76
0.76


14.0
0.0
0.0
0.0
0.0
0.0
0.0
0.0
0.0
0.0
0.0
0.0
0.0
0.0
0.0
0.0
0.0
0.0
0.0
0.0
0.0
0.0
0.0
0.0
0.0
0.0
14.0
0.0
0.0
0.0


14.0
0.0
0.0
0.0
0.0
0.0
0.0
0.0
0.0
0.0
0.0
0.0
0.0
0.0
0.0
0.0
0.0
0.0
0.0
0.0
0.0
0.0
0.0
0.0
0.0
0.0
14.0
0.0
0.0
0.0


14.0
0.0
0.0
0.0
0.0
0.0
0.0
0.0
0.0
0.0
0.0
0.0
0.0
0.0
0.0
0.0
0.0
0.0
0.0
0.0
0.0
0.0
0.0
0.0
0.0
0.0
14.0
0.0
0.0
0.0


14.0
0.0
0.0
0.0
0.0
0.0
0.0
0.0
0.0
0.0
0.0
0.0
0.0
0.0
0.0
0.0
0.0
0.0
0.0
0.0
0.0
0.0
0.0
0.0
0.0
0.0
14.0
0.0
0.0
0.0

1.0
1.0
1.0
1.0
1.0
1.0
1.0
1.0
1.0
1.0
1.0
1.0
1.0
1.0
1.0
1.0
1.0
1.0
1.0
1.0
1.0
1.0
1.0
1.0
1.0
1.0
1.0
1.0
1.0
1.0
