## Supplementary material for "Charting the Insect Biodiversity of Crete: Insights from a Pilot Metabarcoding Survey": Krona_esvs.html

Javascript must be enabled to view this page.

xml version="1.0" ?


esvs
conf

All
2021-04-25\_T1
2021-05-08\_T1
2021-05-22\_T1
2021-06-05\_T1
2021-06-19\_T1
2021-07-03\_T1
2022-03-12\_T1
2022-03-28\_T2
2022-04-09\_T1
2022-04-23\_T2
2022-05-07\_T2
2022-05-21\_T1
2022-05-21\_T3
2022-05-28\_T1
2022-09-17\_T3
2022-09-24\_T3
2022-10-22\_T3
2022-10-29\_T3
2022-11-05\_T3
2022-11-18\_T3
2022-11-26\_T3
2022-12-04\_T3
2022-12-31\_T3
2023-06-04\_T2
2023-08-21\_T2
2023-08-28\_T2
2023-09-04\_T2
2023-09-11\_T2
2023-09-18\_T2


20290
906
910
588
508
671
671
613
774
1140
1101
895
276
452
747
534
349
813
941
615
431
219
354
276
1814
759
800
660
624
849


20031
878
888
576
490
651
646
598
758
1120
1079
870
275
452
747
534
348
808
937
612
429
218
352
275
1810
755
798
659
622
846

1.0
1.0
1.0
1.0
1.0
1.0
1.0
1.0
1.0
1.0
1.0
1.0
1.0
1.0
1.0
1.0
1.0
1.0
1.0
1.0
1.0
1.0
1.0
1.0
1.0
1.0
1.0
1.0
1.0
1.0


19955
872
880
572
485
650
644
593
755
1116
1071
860
275
451
745
534
348
807
937
611
426
216
351
275
1805
754
797
658
621
846

1.0
1.0
1.0
1.0
1.0
1.0
1.0
1.0
1.0
1.0
1.0
1.0
1.0
1.0
1.0
1.0
1.0
1.0
1.0
1.0
1.0
1.0
1.0
1.0
1.0
1.0
1.0
1.0
1.0
1.0


19563
853
867
557
475
643
637
569
744
1100
1043
831
258
444
733
520
337
802
931
603
414
210
328
258
1782
747
785
645
607
840

1.0
1.0
1.0
1.0
1.0
1.0
1.0
1.0
1.0
1.0
1.0
1.0
1.0
1.0
1.0
1.0
1.0
1.0
1.0
1.0
1.0
1.0
1.0
1.0
1.0
1.0
1.0
1.0
1.0
1.0


8359
426
357
159
159
178
162
409
541
732
593
407
177
95
444
249
127
393
299
250
210
113
150
177
233
269
264
175
204
407

1.0
1.0
1.0
1.0
1.0
1.0
1.0
1.0
1.0
1.0
1.0
1.0
1.0
1.0
1.0
1.0
1.0
1.0
1.0
1.0
1.0
1.0
1.0
1.0
1.0
1.0
1.0
1.0
1.0
1.0


351
14
10
4
5
5
4
46
48
51
35
11
6
1
9
1
1
8
19
18
14
5
3
6
1
8
1
7
9
1

1.0
1.0
1.0
1.0
1.0
1.0
1.0
1.0
1.0
1.0
1.0
1.0
1.0
1.0
1.0
1.0
1.0
1.0
1.0
1.0
1.0
1.0
1.0
1.0
1.0
1.0
1.0
1.0
1.0
1.0


154
8
5
1
1
2
0
24
25
22
17
6
4
0
2
1
0
4
8
7
5
3
0
4
0
0
0
5
0
0

1.0
1.0
1.0
1.0
1.0
1.0
1.0
1.0
1.0
1.0
1.0
1.0
1.0
1.0
1.0
1.0
1.0
1.0
1.0
1.0
1.0
1.0
1.0
1.0
1.0
1.0
1.0
1.0
1.0
1.0


100
4
2
0
0
1
0
18
19
14
11
3
4
0
0
0
0
3
6
5
5
1
0
4
0
0
0
0
0
0

1.0
1.0
1.0
1.0
1.0
1.0
1.0
1.0
1.0
1.0
1.0
1.0
1.0
1.0
1.0
1.0
1.0
1.0
1.0
1.0
1.0
1.0
1.0
1.0
1.0
1.0
1.0
1.0
1.0
1.0


11
1
0
0
0
0
0
3
2
2
2
1
0
0
0
0
0
0
0
0
0
0
0
0
0
0
0
0
0
0

1.0
1.0
1.0
1.0
1.0
1.0
1.0
1.0
1.0
1.0
1.0
1.0
1.0
1.0
1.0
1.0
1.0
1.0
1.0
1.0
1.0
1.0
1.0
1.0
1.0
1.0
1.0
1.0
1.0
1.0


9
0
1
0
0
0
0
0
0
1
1
1
0
0
0
0
0
0
0
0
0
0
0
0
0
0
0
5
0
0

1.0
1.0
1.0
1.0
1.0
1.0
1.0
1.0
1.0
1.0
1.0
1.0
1.0
1.0
1.0
1.0
1.0
1.0
1.0
1.0
1.0
1.0
1.0
1.0
1.0
1.0
1.0
1.0
1.0
1.0


3
1
0
0
0
0
0
1
0
0
1
0
0
0
0
0
0
0
0
0
0
0
0
0
0
0
0
0
0
0

0.97
0.97
0.97
0.97
0.97
0.97
0.97
0.97
0.97
0.97
0.97
0.97
0.97
0.97
0.97
0.97
0.97
0.97
0.97
0.97
0.97
0.97
0.97
0.97
0.97
0.97
0.97
0.97
0.97
0.97


7
1
1
0
0
0
0
1
1
1
0
0
0
0
2
0
0
0
0
0
0
0
0
0
0
0
0
0
0
0

1.0
1.0
1.0
1.0
1.0
1.0
1.0
1.0
1.0
1.0
1.0
1.0
1.0
1.0
1.0
1.0
1.0
1.0
1.0
1.0
1.0
1.0
1.0
1.0
1.0
1.0
1.0
1.0
1.0
1.0


12
2
1
1
0
0
0
1
1
2
2
1
0
0
1
0
0
0
0
0
0
0
0
0
0
0
0
0
0
0


12
2
1
1
0
0
0
1
1
2
2
1
0
0
1
0
0
0
0
0
0
0
0
0
0
0
0
0
0
0

1.0
1.0
1.0
1.0
1.0
1.0
1.0
1.0
1.0
1.0
1.0
1.0
1.0
1.0
1.0
1.0
1.0
1.0
1.0
1.0
1.0
1.0
1.0
1.0
1.0
1.0
1.0
1.0
1.0
1.0


11
1
1
1
1
1
1
0
0
0
0
0
0
0
0
0
1
1
0
1
1
1
0
0
0
0
0
0
0
0

1.0
1.0
1.0
1.0
1.0
1.0
1.0
1.0
1.0
1.0
1.0
1.0
1.0
1.0
1.0
1.0
1.0
1.0
1.0
1.0
1.0
1.0
1.0
1.0
1.0
1.0
1.0
1.0
1.0
1.0


27
0
1
0
0
1
1
5
6
6
4
1
0
0
0
0
0
0
0
1
1
0
0
0
0
0
0
0
0
0

0.99
0.99
0.99
0.99
0.99
0.99
0.99
0.99
0.99
0.99
0.99
0.99
0.99
0.99
0.99
0.99
0.99
0.99
0.99
0.99
0.99
0.99
0.99
0.99
0.99
0.99
0.99
0.99
0.99
0.99


5
0
0
0
0
1
1
1
0
1
0
1
0
0
0
0
0
0
0
0
0
0
0
0
0
0
0
0
0
0

1.0
1.0
1.0
1.0
1.0
1.0
1.0
1.0
1.0
1.0
1.0
1.0
1.0
1.0
1.0
1.0
1.0
1.0
1.0
1.0
1.0
1.0
1.0
1.0
1.0
1.0
1.0
1.0
1.0
1.0


30
2
0
0
2
0
0
2
2
4
3
1
0
0
6
0
0
2
1
1
2
0
2
0
0
0
0
0
0
0

0.93
0.93
0.93
0.93
0.93
0.93
0.93
0.93
0.93
0.93
0.93
0.93
0.93
0.93
0.93
0.93
0.93
0.93
0.93
0.93
0.93
0.93
0.93
0.93
0.93
0.93
0.93
0.93
0.93
0.93


5
0
0
0
0
0
0
2
1
1
1
0
0
0
0
0
0
0
0
0
0
0
0
0
0
0
0
0
0
0

1.0
1.0
1.0
1.0
1.0
1.0
1.0
1.0
1.0
1.0
1.0
1.0
1.0
1.0
1.0
1.0
1.0
1.0
1.0
1.0
1.0
1.0
1.0
1.0
1.0
1.0
1.0
1.0
1.0
1.0


14
1
0
0
1
0
0
0
0
2
1
0
0
0
6
0
0
2
1
0
0
0
0
0
0
0
0
0
0
0

1.0
1.0
1.0
1.0
1.0
1.0
1.0
1.0
1.0
1.0
1.0
1.0
1.0
1.0
1.0
1.0
1.0
1.0
1.0
1.0
1.0
1.0
1.0
1.0
1.0
1.0
1.0
1.0
1.0
1.0


7
1
0
0
1
0
0
0
1
1
1
1
0
0
0
0
0
0
0
1
0
0
0
0
0
0
0
0
0
0

1.0
1.0
1.0
1.0
1.0
1.0
1.0
1.0
1.0
1.0
1.0
1.0
1.0
1.0
1.0
1.0
1.0
1.0
1.0
1.0
1.0
1.0
1.0
1.0
1.0
1.0
1.0
1.0
1.0
1.0


8
1
1
0
0
0
0
0
0
0
2
1
0
0
0
0
0
0
1
1
1
0
0
0
0
0
0
0
0
0

1.0
1.0
1.0
1.0
1.0
1.0
1.0
1.0
1.0
1.0
1.0
1.0
1.0
1.0
1.0
1.0
1.0
1.0
1.0
1.0
1.0
1.0
1.0
1.0
1.0
1.0
1.0
1.0
1.0
1.0


46
5
4
1
1
0
6
1
6
7
3
4
0
0
0
0
0
0
0
0
0
0
0
0
2
0
3
3
0
0


31
5
3
1
1
0
0
1
6
7
3
4
0
0
0
0
0
0
0
0
0
0
0
0
0
0
0
0
0
0

0.75
0.75
0.75
0.75
0.75
0.75
0.75
0.75
0.75
0.75
0.75
0.75
0.75
0.75
0.75
0.75
0.75
0.75
0.75
0.75
0.75
0.75
0.75
0.75
0.75
0.75
0.75
0.75
0.75
0.75


16
5
3
1
1
0
0
0
0
1
2
3
0
0
0
0
0
0
0
0
0
0
0
0
0
0
0
0
0
0

1.0
1.0
1.0
1.0
1.0
1.0
1.0
1.0
1.0
1.0
1.0
1.0
1.0
1.0
1.0
1.0
1.0
1.0
1.0
1.0
1.0
1.0
1.0
1.0
1.0
1.0
1.0
1.0
1.0
1.0


10
0
0
0
0
0
0
0
4
4
1
1
0
0
0
0
0
0
0
0
0
0
0
0
0
0
0
0
0
0

0.99
0.99
0.99
0.99
0.99
0.99
0.99
0.99
0.99
0.99
0.99
0.99
0.99
0.99
0.99
0.99
0.99
0.99
0.99
0.99
0.99
0.99
0.99
0.99
0.99
0.99
0.99
0.99
0.99
0.99


12
0
0
0
0
0
6
0
0
0
0
0
0
0
0
0
0
0
0
0
0
0
0
0
0
0
3
3
0
0


12
0
0
0
0
0
6
0
0
0
0
0
0
0
0
0
0
0
0
0
0
0
0
0
0
0
3
3
0
0

1.0
1.0
1.0
1.0
1.0
1.0
1.0
1.0
1.0
1.0
1.0
1.0
1.0
1.0
1.0
1.0
1.0
1.0
1.0
1.0
1.0
1.0
1.0
1.0
1.0
1.0
1.0
1.0
1.0
1.0


3
0
1
0
0
0
0
0
0
0
0
0
0
0
0
0
0
0
0
0
0
0
0
0
2
0
0
0
0
0


1
0
1
0
0
0
0
0
0
0
0
0
0
0
0
0
0
0
0
0
0
0
0
0
0
0
0
0
0
0

0.99
0.99
0.99
0.99
0.99
0.99
0.99
0.99
0.99
0.99
0.99
0.99
0.99
0.99
0.99
0.99
0.99
0.99
0.99
0.99
0.99
0.99
0.99
0.99
0.99
0.99
0.99
0.99
0.99
0.99


2
0
0
0
0
0
0
0
0
0
0
0
0
0
0
0
0
0
0
0
0
0
0
0
2
0
0
0
0
0

0.84
0.84
0.84
0.84
0.84
0.84
0.84
0.84
0.84
0.84
0.84
0.84
0.84
0.84
0.84
0.84
0.84
0.84
0.84
0.84
0.84
0.84
0.84
0.84
0.84
0.84
0.84
0.84
0.84
0.84


530
11
9
4
1
1
1
30
19
24
23
8
25
0
4
74
36
34
6
23
22
7
6
25
0
15
37
29
33
23

0.93
0.93
0.93
0.93
0.93
0.93
0.93
0.93
0.93
0.93
0.93
0.93
0.93
0.93
0.93
0.93
0.93
0.93
0.93
0.93
0.93
0.93
0.93
0.93
0.93
0.93
0.93
0.93
0.93
0.93


303
3
3
1
0
0
0
10
9
6
6
1
4
0
1
29
32
22
3
23
17
5
5
4
0
13
35
23
30
18

1.0
1.0
1.0
1.0
1.0
1.0
1.0
1.0
1.0
1.0
1.0
1.0
1.0
1.0
1.0
1.0
1.0
1.0
1.0
1.0
1.0
1.0
1.0
1.0
1.0
1.0
1.0
1.0
1.0
1.0


120
2
1
0
0
0
0
9
8
4
4
1
4
0
1
3
20
9
2
4
5
2
4
4
0
7
10
7
4
5

1.0
1.0
1.0
1.0
1.0
1.0
1.0
1.0
1.0
1.0
1.0
1.0
1.0
1.0
1.0
1.0
1.0
1.0
1.0
1.0
1.0
1.0
1.0
1.0
1.0
1.0
1.0
1.0
1.0
1.0


37
1
1
1
0
0
0
1
1
2
1
0
0
0
0
4
4
3
0
0
0
3
0
0
0
0
4
3
4
4

1.0
1.0
1.0
1.0
1.0
1.0
1.0
1.0
1.0
1.0
1.0
1.0
1.0
1.0
1.0
1.0
1.0
1.0
1.0
1.0
1.0
1.0
1.0
1.0
1.0
1.0
1.0
1.0
1.0
1.0


106
0
1
0
0
0
0
0
0
0
1
0
0
0
0
17
6
6
1
15
10
0
1
0
0
4
14
10
15
5

1.0
1.0
1.0
1.0
1.0
1.0
1.0
1.0
1.0
1.0
1.0
1.0
1.0
1.0
1.0
1.0
1.0
1.0
1.0
1.0
1.0
1.0
1.0
1.0
1.0
1.0
1.0
1.0
1.0
1.0


68
1
1
1
1
1
1
17
6
9
8
2
4
0
0
0
0
0
3
0
5
2
1
4
0
0
0
0
0
1

1.0
1.0
1.0
1.0
1.0
1.0
1.0
1.0
1.0
1.0
1.0
1.0
1.0
1.0
1.0
1.0
1.0
1.0
1.0
1.0
1.0
1.0
1.0
1.0
1.0
1.0
1.0
1.0
1.0
1.0


45
1
1
1
1
1
1
12
4
5
6
1
4
0
0
0
0
0
1
0
0
0
1
4
0
0
0
0
0
1

1.0
1.0
1.0
1.0
1.0
1.0
1.0
1.0
1.0
1.0
1.0
1.0
1.0
1.0
1.0
1.0
1.0
1.0
1.0
1.0
1.0
1.0
1.0
1.0
1.0
1.0
1.0
1.0
1.0
1.0


106
2
2
0
0
0
0
0
0
3
3
2
13
0
0
43
4
11
0
0
0
0
0
13
0
0
0
4
3
3


106
2
2
0
0
0
0
0
0
3
3
2
13
0
0
43
4
11
0
0
0
0
0
13
0
0
0
4
3
3

1.0
1.0
1.0
1.0
1.0
1.0
1.0
1.0
1.0
1.0
1.0
1.0
1.0
1.0
1.0
1.0
1.0
1.0
1.0
1.0
1.0
1.0
1.0
1.0
1.0
1.0
1.0
1.0
1.0
1.0


11
0
0
1
0
0
0
2
4
2
2
0
0
0
0
0
0
0
0
0
0
0
0
0
0
0
0
0
0
0


11
0
0
1
0
0
0
2
4
2
2
0
0
0
0
0
0
0
0
0
0
0
0
0
0
0
0
0
0
0

1.0
1.0
1.0
1.0
1.0
1.0
1.0
1.0
1.0
1.0
1.0
1.0
1.0
1.0
1.0
1.0
1.0
1.0
1.0
1.0
1.0
1.0
1.0
1.0
1.0
1.0
1.0
1.0
1.0
1.0


2
1
0
0
0
0
0
1
0
0
0
0
0
0
0
0
0
0
0
0
0
0
0
0
0
0
0
0
0
0

0.93
0.93
0.93
0.93
0.93
0.93
0.93
0.93
0.93
0.93
0.93
0.93
0.93
0.93
0.93
0.93
0.93
0.93
0.93
0.93
0.93
0.93
0.93
0.93
0.93
0.93
0.93
0.93
0.93
0.93


1
1
0
0
0
0
0
0
0
0
0
0
0
0
0
0
0
0
0
0
0
0
0
0
0
0
0
0
0
0

0.91
0.91
0.91
0.91
0.91
0.91
0.91
0.91
0.91
0.91
0.91
0.91
0.91
0.91
0.91
0.91
0.91
0.91
0.91
0.91
0.91
0.91
0.91
0.91
0.91
0.91
0.91
0.91
0.91
0.91


1
0
0
0
0
0
0
0
0
1
0
0
0
0
0
0
0
0
0
0
0
0
0
0
0
0
0
0
0
0


1
0
0
0
0
0
0
0
0
1
0
0
0
0
0
0
0
0
0
0
0
0
0
0
0
0
0
0
0
0

1.0
1.0
1.0
1.0
1.0
1.0
1.0
1.0
1.0
1.0
1.0
1.0
1.0
1.0
1.0
1.0
1.0
1.0
1.0
1.0
1.0
1.0
1.0
1.0
1.0
1.0
1.0
1.0
1.0
1.0


4
0
0
0
0
0
0
0
0
0
0
0
2
0
0
0
0
0
0
0
0
0
0
2
0
0
0
0
0
0


4
0
0
0
0
0
0
0
0
0
0
0
2
0
0
0
0
0
0
0
0
0
0
2
0
0
0
0
0
0

0.88
0.88
0.88
0.88
0.88
0.88
0.88
0.88
0.88
0.88
0.88
0.88
0.88
0.88
0.88
0.88
0.88
0.88
0.88
0.88
0.88
0.88
0.88
0.88
0.88
0.88
0.88
0.88
0.88
0.88


672
31
10
1
1
3
3
16
36
52
54
24
55
7
37
11
6
36
12
13
9
0
32
55
10
46
24
14
21
53

0.98
0.98
0.98
0.98
0.98
0.98
0.98
0.98
0.98
0.98
0.98
0.98
0.98
0.98
0.98
0.98
0.98
0.98
0.98
0.98
0.98
0.98
0.98
0.98
0.98
0.98
0.98
0.98
0.98
0.98


32
3
0
0
0
0
0
8
10
5
3
3
0
0
0
0
0
0
0
0
0
0
0
0
0
0
0
0
0
0


24
0
0
0
0
0
0
5
8
5
3
3
0
0
0
0
0
0
0
0
0
0
0
0
0
0
0
0
0
0

1.0
1.0
1.0
1.0
1.0
1.0
1.0
1.0
1.0
1.0
1.0
1.0
1.0
1.0
1.0
1.0
1.0
1.0
1.0
1.0
1.0
1.0
1.0
1.0
1.0
1.0
1.0
1.0
1.0
1.0


6
1
0
0
0
0
0
3
2
0
0
0
0
0
0
0
0
0
0
0
0
0
0
0
0
0
0
0
0
0

1.0
1.0
1.0
1.0
1.0
1.0
1.0
1.0
1.0
1.0
1.0
1.0
1.0
1.0
1.0
1.0
1.0
1.0
1.0
1.0
1.0
1.0
1.0
1.0
1.0
1.0
1.0
1.0
1.0
1.0


2
2
0
0
0
0
0
0
0
0
0
0
0
0
0
0
0
0
0
0
0
0
0
0
0
0
0
0
0
0

1.0
1.0
1.0
1.0
1.0
1.0
1.0
1.0
1.0
1.0
1.0
1.0
1.0
1.0
1.0
1.0
1.0
1.0
1.0
1.0
1.0
1.0
1.0
1.0
1.0
1.0
1.0
1.0
1.0
1.0


166
6
0
0
0
0
0
4
6
4
0
4
43
0
0
1
0
9
0
0
0
0
32
43
0
9
5
0
0
0

0.99
0.99
0.99
0.99
0.99
0.99
0.99
0.99
0.99
0.99
0.99
0.99
0.99
0.99
0.99
0.99
0.99
0.99
0.99
0.99
0.99
0.99
0.99
0.99
0.99
0.99
0.99
0.99
0.99
0.99


59
2
0
0
0
0
0
4
6
4
0
4
4
0
0
1
0
9
0
0
0
0
7
4
0
9
5
0
0
0

1.0
1.0
1.0
1.0
1.0
1.0
1.0
1.0
1.0
1.0
1.0
1.0
1.0
1.0
1.0
1.0
1.0
1.0
1.0
1.0
1.0
1.0
1.0
1.0
1.0
1.0
1.0
1.0
1.0
1.0


44
4
0
0
0
0
0
0
0
0
0
0
15
0
0
0
0
0
0
0
0
0
10
15
0
0
0
0
0
0

1.0
1.0
1.0
1.0
1.0
1.0
1.0
1.0
1.0
1.0
1.0
1.0
1.0
1.0
1.0
1.0
1.0
1.0
1.0
1.0
1.0
1.0
1.0
1.0
1.0
1.0
1.0
1.0
1.0
1.0


100
14
3
0
0
0
0
0
11
24
32
7
1
0
5
0
0
0
0
0
0
0
0
1
0
2
0
0
0
0

0.98
0.98
0.98
0.98
0.98
0.98
0.98
0.98
0.98
0.98
0.98
0.98
0.98
0.98
0.98
0.98
0.98
0.98
0.98
0.98
0.98
0.98
0.98
0.98
0.98
0.98
0.98
0.98
0.98
0.98


73
1
0
1
0
2
1
0
3
8
5
4
0
7
27
0
0
0
0
0
0
0
0
0
10
0
0
0
4
0

0.7
0.7
0.7
0.7
0.7
0.7
0.7
0.7
0.7
0.7
0.7
0.7
0.7
0.7
0.7
0.7
0.7
0.7
0.7
0.7
0.7
0.7
0.7
0.7
0.7
0.7
0.7
0.7
0.7
0.7


7
0
0
0
0
0
1
0
0
1
1
1
0
0
1
0
0
0
0
0
0
0
0
0
2
0
0
0
0
0

1.0
1.0
1.0
1.0
1.0
1.0
1.0
1.0
1.0
1.0
1.0
1.0
1.0
1.0
1.0
1.0
1.0
1.0
1.0
1.0
1.0
1.0
1.0
1.0
1.0
1.0
1.0
1.0
1.0
1.0


41
1
0
0
0
1
0
0
1
2
2
2
0
5
19
0
0
0
0
0
0
0
0
0
8
0
0
0
0
0

1.0
1.0
1.0
1.0
1.0
1.0
1.0
1.0
1.0
1.0
1.0
1.0
1.0
1.0
1.0
1.0
1.0
1.0
1.0
1.0
1.0
1.0
1.0
1.0
1.0
1.0
1.0
1.0
1.0
1.0


8
0
0
0
0
0
0
0
2
4
2
0
0
0
0
0
0
0
0
0
0
0
0
0
0
0
0
0
0
0

1.0
1.0
1.0
1.0
1.0
1.0
1.0
1.0
1.0
1.0
1.0
1.0
1.0
1.0
1.0
1.0
1.0
1.0
1.0
1.0
1.0
1.0
1.0
1.0
1.0
1.0
1.0
1.0
1.0
1.0


16
0
0
1
0
1
0
0
0
1
0
1
0
2
6
0
0
0
0
0
0
0
0
0
0
0
0
0
4
0

1.0
1.0
1.0
1.0
1.0
1.0
1.0
1.0
1.0
1.0
1.0
1.0
1.0
1.0
1.0
1.0
1.0
1.0
1.0
1.0
1.0
1.0
1.0
1.0
1.0
1.0
1.0
1.0
1.0
1.0


14
1
1
0
0
0
0
2
1
2
1
0
0
0
0
0
0
1
3
2
0
0
0
0
0
0
0
0
0
0

0.94
0.94
0.94
0.94
0.94
0.94
0.94
0.94
0.94
0.94
0.94
0.94
0.94
0.94
0.94
0.94
0.94
0.94
0.94
0.94
0.94
0.94
0.94
0.94
0.94
0.94
0.94
0.94
0.94
0.94


11
1
1
0
0
0
0
2
1
2
1
0
0
0
0
0
0
1
2
0
0
0
0
0
0
0
0
0
0
0

1.0
1.0
1.0
1.0
1.0
1.0
1.0
1.0
1.0
1.0
1.0
1.0
1.0
1.0
1.0
1.0
1.0
1.0
1.0
1.0
1.0
1.0
1.0
1.0
1.0
1.0
1.0
1.0
1.0
1.0


21
1
1
0
0
0
0
1
0
1
1
0
4
0
0
0
0
0
0
0
8
0
0
4
0
0
0
0
0
0


21
1
1
0
0
0
0
1
0
1
1
0
4
0
0
0
0
0
0
0
8
0
0
4
0
0
0
0
0
0

1.0
1.0
1.0
1.0
1.0
1.0
1.0
1.0
1.0
1.0
1.0
1.0
1.0
1.0
1.0
1.0
1.0
1.0
1.0
1.0
1.0
1.0
1.0
1.0
1.0
1.0
1.0
1.0
1.0
1.0


45
0
0
0
0
0
0
0
0
0
2
0
3
0
0
1
1
0
1
1
1
0
0
3
0
6
6
3
4
13


44
0
0
0
0
0
0
0
0
0
1
0
3
0
0
1
1
0
1
1
1
0
0
3
0
6
6
3
4
13

1.0
1.0
1.0
1.0
1.0
1.0
1.0
1.0
1.0
1.0
1.0
1.0
1.0
1.0
1.0
1.0
1.0
1.0
1.0
1.0
1.0
1.0
1.0
1.0
1.0
1.0
1.0
1.0
1.0
1.0


1
0
0
0
0
0
0
0
0
0
1
0
0
0
0
0
0
0
0
0
0
0
0
0
0
0
0
0
0
0

1.0
1.0
1.0
1.0
1.0
1.0
1.0
1.0
1.0
1.0
1.0
1.0
1.0
1.0
1.0
1.0
1.0
1.0
1.0
1.0
1.0
1.0
1.0
1.0
1.0
1.0
1.0
1.0
1.0
1.0


23
1
0
0
0
0
0
0
2
0
3
4
0
0
0
6
0
5
0
0
0
0
0
0
0
1
0
0
1
0

1.0
1.0
1.0
1.0
1.0
1.0
1.0
1.0
1.0
1.0
1.0
1.0
1.0
1.0
1.0
1.0
1.0
1.0
1.0
1.0
1.0
1.0
1.0
1.0
1.0
1.0
1.0
1.0
1.0
1.0


5
1
0
0
0
0
0
0
2
0
0
0
0
0
0
0
0
0
0
0
0
0
0
0
0
1
0
0
1
0

1.0
1.0
1.0
1.0
1.0
1.0
1.0
1.0
1.0
1.0
1.0
1.0
1.0
1.0
1.0
1.0
1.0
1.0
1.0
1.0
1.0
1.0
1.0
1.0
1.0
1.0
1.0
1.0
1.0
1.0


5
0
0
0
0
0
0
0
0
0
2
3
0
0
0
0
0
0
0
0
0
0
0
0
0
0
0
0
0
0

1.0
1.0
1.0
1.0
1.0
1.0
1.0
1.0
1.0
1.0
1.0
1.0
1.0
1.0
1.0
1.0
1.0
1.0
1.0
1.0
1.0
1.0
1.0
1.0
1.0
1.0
1.0
1.0
1.0
1.0


11
0
0
0
0
0
0
0
0
0
0
0
0
0
0
6
0
5
0
0
0
0
0
0
0
0
0
0
0
0

0.94
0.94
0.94
0.94
0.94
0.94
0.94
0.94
0.94
0.94
0.94
0.94
0.94
0.94
0.94
0.94
0.94
0.94
0.94
0.94
0.94
0.94
0.94
0.94
0.94
0.94
0.94
0.94
0.94
0.94


52
1
4
0
0
0
1
0
2
5
4
1
0
0
0
0
0
16
5
8
0
0
0
0
0
0
0
0
0
5

1.0
1.0
1.0
1.0
1.0
1.0
1.0
1.0
1.0
1.0
1.0
1.0
1.0
1.0
1.0
1.0
1.0
1.0
1.0
1.0
1.0
1.0
1.0
1.0
1.0
1.0
1.0
1.0
1.0
1.0


28
1
2
0
0
0
0
0
1
5
3
0
0
0
0
0
0
10
0
1
0
0
0
0
0
0
0
0
0
5

1.0
1.0
1.0
1.0
1.0
1.0
1.0
1.0
1.0
1.0
1.0
1.0
1.0
1.0
1.0
1.0
1.0
1.0
1.0
1.0
1.0
1.0
1.0
1.0
1.0
1.0
1.0
1.0
1.0
1.0


5
0
1
0
0
0
1
0
1
0
1
0
0
0
0
0
0
0
1
0
0
0
0
0
0
0
0
0
0
0

0.97
0.97
0.97
0.97
0.97
0.97
0.97
0.97
0.97
0.97
0.97
0.97
0.97
0.97
0.97
0.97
0.97
0.97
0.97
0.97
0.97
0.97
0.97
0.97
0.97
0.97
0.97
0.97
0.97
0.97


33
0
0
0
0
0
0
0
0
1
1
1
0
0
0
0
0
0
0
0
0
0
0
0
0
6
3
8
3
10

0.99
0.99
0.99
0.99
0.99
0.99
0.99
0.99
0.99
0.99
0.99
0.99
0.99
0.99
0.99
0.99
0.99
0.99
0.99
0.99
0.99
0.99
0.99
0.99
0.99
0.99
0.99
0.99
0.99
0.99


5
0
0
0
0
0
0
0
0
1
1
1
0
0
0
0
0
0
0
0
0
0
0
0
0
0
2
0
0
0

1.0
1.0
1.0
1.0
1.0
1.0
1.0
1.0
1.0
1.0
1.0
1.0
1.0
1.0
1.0
1.0
1.0
1.0
1.0
1.0
1.0
1.0
1.0
1.0
1.0
1.0
1.0
1.0
1.0
1.0


21
0
0
0
0
0
0
0
0
0
0
0
0
0
0
0
0
0
0
0
0
0
0
0
0
1
0
7
3
10

1.0
1.0
1.0
1.0
1.0
1.0
1.0
1.0
1.0
1.0
1.0
1.0
1.0
1.0
1.0
1.0
1.0
1.0
1.0
1.0
1.0
1.0
1.0
1.0
1.0
1.0
1.0
1.0
1.0
1.0


1
0
0
0
0
0
0
0
0
0
0
0
0
0
0
0
0
0
0
0
0
0
0
0
0
0
0
1
0
0

0.97
0.97
0.97
0.97
0.97
0.97
0.97
0.97
0.97
0.97
0.97
0.97
0.97
0.97
0.97
0.97
0.97
0.97
0.97
0.97
0.97
0.97
0.97
0.97
0.97
0.97
0.97
0.97
0.97
0.97


23
1
1
0
1
0
0
0
0
0
0
0
0
0
0
0
0
0
3
0
0
0
0
0
0
6
0
1
3
7

1.0
1.0
1.0
1.0
1.0
1.0
1.0
1.0
1.0
1.0
1.0
1.0
1.0
1.0
1.0
1.0
1.0
1.0
1.0
1.0
1.0
1.0
1.0
1.0
1.0
1.0
1.0
1.0
1.0
1.0


18
0
0
0
0
0
0
0
0
0
0
0
0
0
0
0
0
0
3
0
0
0
0
0
0
6
0
0
3
6

0.86
0.86
0.86
0.86
0.86
0.86
0.86
0.86
0.86
0.86
0.86
0.86
0.86
0.86
0.86
0.86
0.86
0.86
0.86
0.86
0.86
0.86
0.86
0.86
0.86
0.86
0.86
0.86
0.86
0.86


1
0
0
0
0
0
0
0
0
0
1
0
0
0
0
0
0
0
0
0
0
0
0
0
0
0
0
0
0
0


1
0
0
0
0
0
0
0
0
0
1
0
0
0
0
0
0
0
0
0
0
0
0
0
0
0
0
0
0
0

1.0
1.0
1.0
1.0
1.0
1.0
1.0
1.0
1.0
1.0
1.0
1.0
1.0
1.0
1.0
1.0
1.0
1.0
1.0
1.0
1.0
1.0
1.0
1.0
1.0
1.0
1.0
1.0
1.0
1.0


4
0
0
0
0
0
0
0
0
0
0
0
0
0
2
0
0
2
0
0
0
0
0
0
0
0
0
0
0
0

0.8
0.8
0.8
0.8
0.8
0.8
0.8
0.8
0.8
0.8
0.8
0.8
0.8
0.8
0.8
0.8
0.8
0.8
0.8
0.8
0.8
0.8
0.8
0.8
0.8
0.8
0.8
0.8
0.8
0.8


5
0
0
0
0
0
0
0
0
0
0
0
0
0
0
0
1
0
0
0
0
0
0
0
0
0
1
1
2
0

0.89
0.89
0.89
0.89
0.89
0.89
0.89
0.89
0.89
0.89
0.89
0.89
0.89
0.89
0.89
0.89
0.89
0.89
0.89
0.89
0.89
0.89
0.89
0.89
0.89
0.89
0.89
0.89
0.89
0.89


2
0
0
0
0
0
0
0
0
0
0
0
1
0
0
0
0
0
0
0
0
0
0
1
0
0
0
0
0
0

0.8
0.8
0.8
0.8
0.8
0.8
0.8
0.8
0.8
0.8
0.8
0.8
0.8
0.8
0.8
0.8
0.8
0.8
0.8
0.8
0.8
0.8
0.8
0.8
0.8
0.8
0.8
0.8
0.8
0.8


4
0
0
0
0
0
0
0
0
0
0
0
0
0
0
0
0
0
0
0
0
0
0
0
0
0
4
0
0
0

1.0
1.0
1.0
1.0
1.0
1.0
1.0
1.0
1.0
1.0
1.0
1.0
1.0
1.0
1.0
1.0
1.0
1.0
1.0
1.0
1.0
1.0
1.0
1.0
1.0
1.0
1.0
1.0
1.0
1.0


399
27
37
12
15
6
6
2
17
29
19
18
4
5
21
5
6
55
12
16
14
8
0
4
11
15
12
5
12
6

1.0
1.0
1.0
1.0
1.0
1.0
1.0
1.0
1.0
1.0
1.0
1.0
1.0
1.0
1.0
1.0
1.0
1.0
1.0
1.0
1.0
1.0
1.0
1.0
1.0
1.0
1.0
1.0
1.0
1.0


47
6
1
0
0
0
0
0
6
7
4
5
0
0
0
0
0
11
0
0
7
0
0
0
0
0
0
0
0
0

1.0
1.0
1.0
1.0
1.0
1.0
1.0
1.0
1.0
1.0
1.0
1.0
1.0
1.0
1.0
1.0
1.0
1.0
1.0
1.0
1.0
1.0
1.0
1.0
1.0
1.0
1.0
1.0
1.0
1.0


5
0
0
0
0
0
0
0
0
2
0
3
0
0
0
0
0
0
0
0
0
0
0
0
0
0
0
0
0
0

1.0
1.0
1.0
1.0
1.0
1.0
1.0
1.0
1.0
1.0
1.0
1.0
1.0
1.0
1.0
1.0
1.0
1.0
1.0
1.0
1.0
1.0
1.0
1.0
1.0
1.0
1.0
1.0
1.0
1.0


34
2
6
0
0
0
0
2
4
6
3
2
0
0
0
0
0
0
3
3
0
3
0
0
0
0
0
0
0
0

1.0
1.0
1.0
1.0
1.0
1.0
1.0
1.0
1.0
1.0
1.0
1.0
1.0
1.0
1.0
1.0
1.0
1.0
1.0
1.0
1.0
1.0
1.0
1.0
1.0
1.0
1.0
1.0
1.0
1.0


3
0
0
0
0
0
0
0
1
2
0
0
0
0
0
0
0
0
0
0
0
0
0
0
0
0
0
0
0
0


3
0
0
0
0
0
0
0
1
2
0
0
0
0
0
0
0
0
0
0
0
0
0
0
0
0
0
0
0
0

1.0
1.0
1.0
1.0
1.0
1.0
1.0
1.0
1.0
1.0
1.0
1.0
1.0
1.0
1.0
1.0
1.0
1.0
1.0
1.0
1.0
1.0
1.0
1.0
1.0
1.0
1.0
1.0
1.0
1.0


18
0
2
1
0
2
0
0
0
0
0
1
0
5
0
0
5
0
0
0
0
0
0
0
0
0
0
0
0
2


9
0
1
1
0
2
0
0
0
0
0
0
0
5
0
0
0
0
0
0
0
0
0
0
0
0
0
0
0
0

0.98
0.98
0.98
0.98
0.98
0.98
0.98
0.98
0.98
0.98
0.98
0.98
0.98
0.98
0.98
0.98
0.98
0.98
0.98
0.98
0.98
0.98
0.98
0.98
0.98
0.98
0.98
0.98
0.98
0.98


2
0
1
0
0
0
0
0
0
0
0
1
0
0
0
0
0
0
0
0
0
0
0
0
0
0
0
0
0
0

1.0
1.0
1.0
1.0
1.0
1.0
1.0
1.0
1.0
1.0
1.0
1.0
1.0
1.0
1.0
1.0
1.0
1.0
1.0
1.0
1.0
1.0
1.0
1.0
1.0
1.0
1.0
1.0
1.0
1.0


7
0
0
0
0
0
0
0
0
0
0
0
0
0
0
0
5
0
0
0
0
0
0
0
0
0
0
0
0
2

1.0
1.0
1.0
1.0
1.0
1.0
1.0
1.0
1.0
1.0
1.0
1.0
1.0
1.0
1.0
1.0
1.0
1.0
1.0
1.0
1.0
1.0
1.0
1.0
1.0
1.0
1.0
1.0
1.0
1.0


11
2
0
0
0
0
0
0
4
2
3
0
0
0
0
0
0
0
0
0
0
0
0
0
0
0
0
0
0
0


11
2
0
0
0
0
0
0
4
2
3
0
0
0
0
0
0
0
0
0
0
0
0
0
0
0
0
0
0
0

1.0
1.0
1.0
1.0
1.0
1.0
1.0
1.0
1.0
1.0
1.0
1.0
1.0
1.0
1.0
1.0
1.0
1.0
1.0
1.0
1.0
1.0
1.0
1.0
1.0
1.0
1.0
1.0
1.0
1.0


7
0
1
2
4
0
0
0
0
0
0
0
0
0
0
0
0
0
0
0
0
0
0
0
0
0
0
0
0
0


7
0
1
2
4
0
0
0
0
0
0
0
0
0
0
0
0
0
0
0
0
0
0
0
0
0
0
0
0
0

1.0
1.0
1.0
1.0
1.0
1.0
1.0
1.0
1.0
1.0
1.0
1.0
1.0
1.0
1.0
1.0
1.0
1.0
1.0
1.0
1.0
1.0
1.0
1.0
1.0
1.0
1.0
1.0
1.0
1.0


3
0
1
0
0
0
0
0
1
1
0
0
0
0
0
0
0
0
0
0
0
0
0
0
0
0
0
0
0
0

1.0
1.0
1.0
1.0
1.0
1.0
1.0
1.0
1.0
1.0
1.0
1.0
1.0
1.0
1.0
1.0
1.0
1.0
1.0
1.0
1.0
1.0
1.0
1.0
1.0
1.0
1.0
1.0
1.0
1.0


5
2
1
0
0
0
0
0
0
1
1
0
0
0
0
0
0
0
0
0
0
0
0
0
0
0
0
0
0
0

1.0
1.0
1.0
1.0
1.0
1.0
1.0
1.0
1.0
1.0
1.0
1.0
1.0
1.0
1.0
1.0
1.0
1.0
1.0
1.0
1.0
1.0
1.0
1.0
1.0
1.0
1.0
1.0
1.0
1.0


7
2
1
0
0
0
0
0
0
3
1
0
0
0
0
0
0
0
0
0
0
0
0
0
0
0
0
0
0
0


7
2
1
0
0
0
0
0
0
3
1
0
0
0
0
0
0
0
0
0
0
0
0
0
0
0
0
0
0
0

1.0
1.0
1.0
1.0
1.0
1.0
1.0
1.0
1.0
1.0
1.0
1.0
1.0
1.0
1.0
1.0
1.0
1.0
1.0
1.0
1.0
1.0
1.0
1.0
1.0
1.0
1.0
1.0
1.0
1.0


34
0
0
0
3
3
3
0
0
0
3
1
0
0
0
0
0
0
0
0
0
0
0
0
0
12
9
0
0
0

1.0
1.0
1.0
1.0
1.0
1.0
1.0
1.0
1.0
1.0
1.0
1.0
1.0
1.0
1.0
1.0
1.0
1.0
1.0
1.0
1.0
1.0
1.0
1.0
1.0
1.0
1.0
1.0
1.0
1.0


4
0
1
0
0
0
0
0
0
1
0
1
0
0
0
0
0
0
0
0
0
0
0
0
1
0
0
0
0
0


4
0
1
0
0
0
0
0
0
1
0
1
0
0
0
0
0
0
0
0
0
0
0
0
1
0
0
0
0
0

1.0
1.0
1.0
1.0
1.0
1.0
1.0
1.0
1.0
1.0
1.0
1.0
1.0
1.0
1.0
1.0
1.0
1.0
1.0
1.0
1.0
1.0
1.0
1.0
1.0
1.0
1.0
1.0
1.0
1.0


8
2
0
0
0
0
0
0
0
3
0
3
0
0
0
0
0
0
0
0
0
0
0
0
0
0
0
0
0
0


8
2
0
0
0
0
0
0
0
3
0
3
0
0
0
0
0
0
0
0
0
0
0
0
0
0
0
0
0
0

1.0
1.0
1.0
1.0
1.0
1.0
1.0
1.0
1.0
1.0
1.0
1.0
1.0
1.0
1.0
1.0
1.0
1.0
1.0
1.0
1.0
1.0
1.0
1.0
1.0
1.0
1.0
1.0
1.0
1.0


35
2
8
5
1
1
1
0
0
0
0
1
4
0
6
2
0
0
0
0
0
0
0
4
0
0
0
0
0
0

1.0
1.0
1.0
1.0
1.0
1.0
1.0
1.0
1.0
1.0
1.0
1.0
1.0
1.0
1.0
1.0
1.0
1.0
1.0
1.0
1.0
1.0
1.0
1.0
1.0
1.0
1.0
1.0
1.0
1.0


7
0
4
2
0
0
0
0
0
0
0
1
0
0
0
0
0
0
0
0
0
0
0
0
0
0
0
0
0
0

0.93
0.93
0.93
0.93
0.93
0.93
0.93
0.93
0.93
0.93
0.93
0.93
0.93
0.93
0.93
0.93
0.93
0.93
0.93
0.93
0.93
0.93
0.93
0.93
0.93
0.93
0.93
0.93
0.93
0.93


2
2
0
0
0
0
0
0
0
0
0
0
0
0
0
0
0
0
0
0
0
0
0
0
0
0
0
0
0
0


2
2
0
0
0
0
0
0
0
0
0
0
0
0
0
0
0
0
0
0
0
0
0
0
0
0
0
0
0
0

1.0
1.0
1.0
1.0
1.0
1.0
1.0
1.0
1.0
1.0
1.0
1.0
1.0
1.0
1.0
1.0
1.0
1.0
1.0
1.0
1.0
1.0
1.0
1.0
1.0
1.0
1.0
1.0
1.0
1.0


20
0
1
0
2
0
2
0
0
0
0
0
0
0
0
1
0
13
0
0
0
0
0
0
0
1
0
0
0
0

1.0
1.0
1.0
1.0
1.0
1.0
1.0
1.0
1.0
1.0
1.0
1.0
1.0
1.0
1.0
1.0
1.0
1.0
1.0
1.0
1.0
1.0
1.0
1.0
1.0
1.0
1.0
1.0
1.0
1.0


14
0
1
0
0
0
0
0
0
0
0
0
0
0
0
0
0
13
0
0
0
0
0
0
0
0
0
0
0
0

1.0
1.0
1.0
1.0
1.0
1.0
1.0
1.0
1.0
1.0
1.0
1.0
1.0
1.0
1.0
1.0
1.0
1.0
1.0
1.0
1.0
1.0
1.0
1.0
1.0
1.0
1.0
1.0
1.0
1.0


34
2
1
0
1
0
0
0
0
2
2
0
0
0
0
0
0
6
0
0
0
0
0
0
7
0
0
0
10
3

1.0
1.0
1.0
1.0
1.0
1.0
1.0
1.0
1.0
1.0
1.0
1.0
1.0
1.0
1.0
1.0
1.0
1.0
1.0
1.0
1.0
1.0
1.0
1.0
1.0
1.0
1.0
1.0
1.0
1.0


7
2
1
0
0
0
0
0
0
2
2
0
0
0
0
0
0
0
0
0
0
0
0
0
0
0
0
0
0
0

0.99
0.99
0.99
0.99
0.99
0.99
0.99
0.99
0.99
0.99
0.99
0.99
0.99
0.99
0.99
0.99
0.99
0.99
0.99
0.99
0.99
0.99
0.99
0.99
0.99
0.99
0.99
0.99
0.99
0.99


17
0
0
0
1
0
0
0
0
0
0
0
0
0
0
0
0
6
0
0
0
0
0
0
3
0
0
0
5
2

0.99
0.99
0.99
0.99
0.99
0.99
0.99
0.99
0.99
0.99
0.99
0.99
0.99
0.99
0.99
0.99
0.99
0.99
0.99
0.99
0.99
0.99
0.99
0.99
0.99
0.99
0.99
0.99
0.99
0.99


19
0
2
0
4
0
0
0
0
0
0
0
0
0
0
0
1
6
1
5
0
0
0
0
0
0
0
0
0
0


19
0
2
0
4
0
0
0
0
0
0
0
0
0
0
0
1
6
1
5
0
0
0
0
0
0
0
0
0
0

1.0
1.0
1.0
1.0
1.0
1.0
1.0
1.0
1.0
1.0
1.0
1.0
1.0
1.0
1.0
1.0
1.0
1.0
1.0
1.0
1.0
1.0
1.0
1.0
1.0
1.0
1.0
1.0
1.0
1.0


10
2
2
2
0
0
0
0
0
0
0
2
0
0
0
0
0
0
0
0
0
0
0
0
2
0
0
0
0
0


10
2
2
2
0
0
0
0
0
0
0
2
0
0
0
0
0
0
0
0
0
0
0
0
2
0
0
0
0
0

1.0
1.0
1.0
1.0
1.0
1.0
1.0
1.0
1.0
1.0
1.0
1.0
1.0
1.0
1.0
1.0
1.0
1.0
1.0
1.0
1.0
1.0
1.0
1.0
1.0
1.0
1.0
1.0
1.0
1.0


22
0
1
0
0
0
0
0
0
0
1
0
0
0
0
0
0
0
5
5
5
5
0
0
0
0
0
0
0
0

1.0
1.0
1.0
1.0
1.0
1.0
1.0
1.0
1.0
1.0
1.0
1.0
1.0
1.0
1.0
1.0
1.0
1.0
1.0
1.0
1.0
1.0
1.0
1.0
1.0
1.0
1.0
1.0
1.0
1.0


10
0
1
0
0
0
0
0
0
0
1
0
0
0
0
0
0
0
2
2
2
2
0
0
0
0
0
0
0
0

1.0
1.0
1.0
1.0
1.0
1.0
1.0
1.0
1.0
1.0
1.0
1.0
1.0
1.0
1.0
1.0
1.0
1.0
1.0
1.0
1.0
1.0
1.0
1.0
1.0
1.0
1.0
1.0
1.0
1.0


1
0
0
0
0
0
0
0
0
1
0
0
0
0
0
0
0
0
0
0
0
0
0
0
0
0
0
0
0
0


1
0
0
0
0
0
0
0
0
1
0
0
0
0
0
0
0
0
0
0
0
0
0
0
0
0
0
0
0
0

0.99
0.99
0.99
0.99
0.99
0.99
0.99
0.99
0.99
0.99
0.99
0.99
0.99
0.99
0.99
0.99
0.99
0.99
0.99
0.99
0.99
0.99
0.99
0.99
0.99
0.99
0.99
0.99
0.99
0.99


1
0
0
0
0
0
0
0
0
0
1
0
0
0
0
0
0
0
0
0
0
0
0
0
0
0
0
0
0
0


1
0
0
0
0
0
0
0
0
0
1
0
0
0
0
0
0
0
0
0
0
0
0
0
0
0
0
0
0
0

1.0
1.0
1.0
1.0
1.0
1.0
1.0
1.0
1.0
1.0
1.0
1.0
1.0
1.0
1.0
1.0
1.0
1.0
1.0
1.0
1.0
1.0
1.0
1.0
1.0
1.0
1.0
1.0
1.0
1.0


2
1
1
0
0
0
0
0
0
0
0
0
0
0
0
0
0
0
0
0
0
0
0
0
0
0
0
0
0
0

0.95
0.95
0.95
0.95
0.95
0.95
0.95
0.95
0.95
0.95
0.95
0.95
0.95
0.95
0.95
0.95
0.95
0.95
0.95
0.95
0.95
0.95
0.95
0.95
0.95
0.95
0.95
0.95
0.95
0.95


1
0
1
0
0
0
0
0
0
0
0
0
0
0
0
0
0
0
0
0
0
0
0
0
0
0
0
0
0
0

0.89
0.89
0.89
0.89
0.89
0.89
0.89
0.89
0.89
0.89
0.89
0.89
0.89
0.89
0.89
0.89
0.89
0.89
0.89
0.89
0.89
0.89
0.89
0.89
0.89
0.89
0.89
0.89
0.89
0.89


3
1
2
0
0
0
0
0
0
0
0
0
0
0
0
0
0
0
0
0
0
0
0
0
0
0
0
0
0
0


2
1
1
0
0
0
0
0
0
0
0
0
0
0
0
0
0
0
0
0
0
0
0
0
0
0
0
0
0
0

1.0
1.0
1.0
1.0
1.0
1.0
1.0
1.0
1.0
1.0
1.0
1.0
1.0
1.0
1.0
1.0
1.0
1.0
1.0
1.0
1.0
1.0
1.0
1.0
1.0
1.0
1.0
1.0
1.0
1.0


1
0
1
0
0
0
0
0
0
0
0
0
0
0
0
0
0
0
0
0
0
0
0
0
0
0
0
0
0
0

1.0
1.0
1.0
1.0
1.0
1.0
1.0
1.0
1.0
1.0
1.0
1.0
1.0
1.0
1.0
1.0
1.0
1.0
1.0
1.0
1.0
1.0
1.0
1.0
1.0
1.0
1.0
1.0
1.0
1.0


2
0
2
0
0
0
0
0
0
0
0
0
0
0
0
0
0
0
0
0
0
0
0
0
0
0
0
0
0
0

1.0
1.0
1.0
1.0
1.0
1.0
1.0
1.0
1.0
1.0
1.0
1.0
1.0
1.0
1.0
1.0
1.0
1.0
1.0
1.0
1.0
1.0
1.0
1.0
1.0
1.0
1.0
1.0
1.0
1.0


17
0
1
1
0
0
0
0
1
0
0
0
0
0
14
0
0
0
0
0
0
0
0
0
0
0
0
0
0
0


17
0
1
1
0
0
0
0
1
0
0
0
0
0
14
0
0
0
0
0
0
0
0
0
0
0
0
0
0
0

1.0
1.0
1.0
1.0
1.0
1.0
1.0
1.0
1.0
1.0
1.0
1.0
1.0
1.0
1.0
1.0
1.0
1.0
1.0
1.0
1.0
1.0
1.0
1.0
1.0
1.0
1.0
1.0
1.0
1.0


7
0
0
0
0
0
0
0
0
0
0
0
0
0
0
1
0
1
1
1
0
0
0
0
0
0
1
0
1
1

0.75
0.75
0.75
0.75
0.75
0.75
0.75
0.75
0.75
0.75
0.75
0.75
0.75
0.75
0.75
0.75
0.75
0.75
0.75
0.75
0.75
0.75
0.75
0.75
0.75
0.75
0.75
0.75
0.75
0.75


6
0
0
0
0
0
0
0
0
0
0
0
0
0
0
0
0
6
0
0
0
0
0
0
0
0
0
0
0
0

0.96
0.96
0.96
0.96
0.96
0.96
0.96
0.96
0.96
0.96
0.96
0.96
0.96
0.96
0.96
0.96
0.96
0.96
0.96
0.96
0.96
0.96
0.96
0.96
0.96
0.96
0.96
0.96
0.96
0.96


5
0
0
0
0
0
0
0
0
0
0
0
0
0
0
0
0
5
0
0
0
0
0
0
0
0
0
0
0
0

1.0
1.0
1.0
1.0
1.0
1.0
1.0
1.0
1.0
1.0
1.0
1.0
1.0
1.0
1.0
1.0
1.0
1.0
1.0
1.0
1.0
1.0
1.0
1.0
1.0
1.0
1.0
1.0
1.0
1.0


5
0
0
0
0
0
0
0
0
0
0
0
0
0
0
0
0
0
0
0
0
0
0
0
0
0
0
5
0
0


5
0
0
0
0
0
0
0
0
0
0
0
0
0
0
0
0
0
0
0
0
0
0
0
0
0
0
5
0
0

1.0
1.0
1.0
1.0
1.0
1.0
1.0
1.0
1.0
1.0
1.0
1.0
1.0
1.0
1.0
1.0
1.0
1.0
1.0
1.0
1.0
1.0
1.0
1.0
1.0
1.0
1.0
1.0
1.0
1.0


2
0
0
0
0
0
0
0
0
0
0
0
0
0
0
0
0
1
0
0
1
0
0
0
0
0
0
0
0
0


2
0
0
0
0
0
0
0
0
0
0
0
0
0
0
0
0
1
0
0
1
0
0
0
0
0
0
0
0
0

1.0
1.0
1.0
1.0
1.0
1.0
1.0
1.0
1.0
1.0
1.0
1.0
1.0
1.0
1.0
1.0
1.0
1.0
1.0
1.0
1.0
1.0
1.0
1.0
1.0
1.0
1.0
1.0
1.0
1.0


1
0
0
0
0
0
0
0
0
0
0
0
0
0
1
0
0
0
0
0
0
0
0
0
0
0
0
0
0
0

0.7
0.7
0.7
0.7
0.7
0.7
0.7
0.7
0.7
0.7
0.7
0.7
0.7
0.7
0.7
0.7
0.7
0.7
0.7
0.7
0.7
0.7
0.7
0.7
0.7
0.7
0.7
0.7
0.7
0.7


434
6
33
31
58
46
44
0
1
15
2
1
0
0
4
43
19
12
0
13
6
3
8
0
11
15
41
12
9
1

0.92
0.92
0.92
0.92
0.92
0.92
0.92
0.92
0.92
0.92
0.92
0.92
0.92
0.92
0.92
0.92
0.92
0.92
0.92
0.92
0.92
0.92
0.92
0.92
0.92
0.92
0.92
0.92
0.92
0.92


365
4
32
31
58
44
41
0
1
14
1
1
0
0
1
31
17
12
0
13
6
3
8
0
11
11
13
10
2
0

1.0
1.0
1.0
1.0
1.0
1.0
1.0
1.0
1.0
1.0
1.0
1.0
1.0
1.0
1.0
1.0
1.0
1.0
1.0
1.0
1.0
1.0
1.0
1.0
1.0
1.0
1.0
1.0
1.0
1.0


152
0
15
15
36
27
21
0
0
4
0
1
0
0
0
15
8
0
0
0
0
0
0
0
1
1
7
1
0
0

1.0
1.0
1.0
1.0
1.0
1.0
1.0
1.0
1.0
1.0
1.0
1.0
1.0
1.0
1.0
1.0
1.0
1.0
1.0
1.0
1.0
1.0
1.0
1.0
1.0
1.0
1.0
1.0
1.0
1.0


37
1
5
3
5
5
4
0
0
0
0
0
0
0
0
1
1
0
0
0
5
0
0
0
5
1
0
1
0
0

1.0
1.0
1.0
1.0
1.0
1.0
1.0
1.0
1.0
1.0
1.0
1.0
1.0
1.0
1.0
1.0
1.0
1.0
1.0
1.0
1.0
1.0
1.0
1.0
1.0
1.0
1.0
1.0
1.0
1.0


15
0
3
3
3
3
3
0
0
0
0
0
0
0
0
0
0
0
0
0
0
0
0
0
0
0
0
0
0
0

0.94
0.94
0.94
0.94
0.94
0.94
0.94
0.94
0.94
0.94
0.94
0.94
0.94
0.94
0.94
0.94
0.94
0.94
0.94
0.94
0.94
0.94
0.94
0.94
0.94
0.94
0.94
0.94
0.94
0.94


5
1
0
1
1
0
0
0
1
0
1
0
0
0
0
0
0
0
0
0
0
0
0
0
0
0
0
0
0
0

1.0
1.0
1.0
1.0
1.0
1.0
1.0
1.0
1.0
1.0
1.0
1.0
1.0
1.0
1.0
1.0
1.0
1.0
1.0
1.0
1.0
1.0
1.0
1.0
1.0
1.0
1.0
1.0
1.0
1.0


15
0
2
3
3
2
3
0
0
2
0
0
0
0
0
0
0
0
0
0
0
0
0
0
0
0
0
0
0
0

1.0
1.0
1.0
1.0
1.0
1.0
1.0
1.0
1.0
1.0
1.0
1.0
1.0
1.0
1.0
1.0
1.0
1.0
1.0
1.0
1.0
1.0
1.0
1.0
1.0
1.0
1.0
1.0
1.0
1.0


2
1
0
0
0
0
0
0
0
1
0
0
0
0
0
0
0
0
0
0
0
0
0
0
0
0
0
0
0
0

1.0
1.0
1.0
1.0
1.0
1.0
1.0
1.0
1.0
1.0
1.0
1.0
1.0
1.0
1.0
1.0
1.0
1.0
1.0
1.0
1.0
1.0
1.0
1.0
1.0
1.0
1.0
1.0
1.0
1.0


6
0
0
0
0
0
0
0
0
0
0
0
0
0
0
0
0
0
0
0
0
0
0
0
0
2
2
1
1
0

1.0
1.0
1.0
1.0
1.0
1.0
1.0
1.0
1.0
1.0
1.0
1.0
1.0
1.0
1.0
1.0
1.0
1.0
1.0
1.0
1.0
1.0
1.0
1.0
1.0
1.0
1.0
1.0
1.0
1.0


2
0
0
0
0
0
0
0
0
0
0
0
0
0
0
0
0
0
0
2
0
0
0
0
0
0
0
0
0
0

0.89
0.89
0.89
0.89
0.89
0.89
0.89
0.89
0.89
0.89
0.89
0.89
0.89
0.89
0.89
0.89
0.89
0.89
0.89
0.89
0.89
0.89
0.89
0.89
0.89
0.89
0.89
0.89
0.89
0.89


4
0
0
0
0
0
0
0
0
0
0
0
0
0
0
0
0
1
0
0
0
1
2
0
0
0
0
0
0
0

0.97
0.97
0.97
0.97
0.97
0.97
0.97
0.97
0.97
0.97
0.97
0.97
0.97
0.97
0.97
0.97
0.97
0.97
0.97
0.97
0.97
0.97
0.97
0.97
0.97
0.97
0.97
0.97
0.97
0.97


29
1
0
0
0
0
1
0
0
1
1
0
0
0
0
8
1
0
0
0
0
0
0
0
0
4
5
0
7
0

1.0
1.0
1.0
1.0
1.0
1.0
1.0
1.0
1.0
1.0
1.0
1.0
1.0
1.0
1.0
1.0
1.0
1.0
1.0
1.0
1.0
1.0
1.0
1.0
1.0
1.0
1.0
1.0
1.0
1.0


3
0
0
0
0
2
0
0
0
0
0
0
0
0
0
0
0
0
0
0
0
0
0
0
0
0
0
1
0
0


3
0
0
0
0
2
0
0
0
0
0
0
0
0
0
0
0
0
0
0
0
0
0
0
0
0
0
1
0
0

1.0
1.0
1.0
1.0
1.0
1.0
1.0
1.0
1.0
1.0
1.0
1.0
1.0
1.0
1.0
1.0
1.0
1.0
1.0
1.0
1.0
1.0
1.0
1.0
1.0
1.0
1.0
1.0
1.0
1.0


7
1
1
0
0
0
1
0
0
0
0
0
0
0
3
0
0
0
0
0
0
0
0
0
0
0
0
0
0
1


7
1
1
0
0
0
1
0
0
0
0
0
0
0
3
0
0
0
0
0
0
0
0
0
0
0
0
0
0
1

1.0
1.0
1.0
1.0
1.0
1.0
1.0
1.0
1.0
1.0
1.0
1.0
1.0
1.0
1.0
1.0
1.0
1.0
1.0
1.0
1.0
1.0
1.0
1.0
1.0
1.0
1.0
1.0
1.0
1.0


2
0
0
0
0
0
1
0
0
0
0
0
0
0
0
0
0
0
0
0
0
0
0
0
0
0
0
1
0
0

0.82
0.82
0.82
0.82
0.82
0.82
0.82
0.82
0.82
0.82
0.82
0.82
0.82
0.82
0.82
0.82
0.82
0.82
0.82
0.82
0.82
0.82
0.82
0.82
0.82
0.82
0.82
0.82
0.82
0.82


1
0
0
0
0
0
1
0
0
0
0
0
0
0
0
0
0
0
0
0
0
0
0
0
0
0
0
0
0
0

1.0
1.0
1.0
1.0
1.0
1.0
1.0
1.0
1.0
1.0
1.0
1.0
1.0
1.0
1.0
1.0
1.0
1.0
1.0
1.0
1.0
1.0
1.0
1.0
1.0
1.0
1.0
1.0
1.0
1.0


23
0
0
0
0
0
0
0
0
0
0
0
0
0
0
0
0
0
0
0
0
0
0
0
0
0
23
0
0
0

1.0
1.0
1.0
1.0
1.0
1.0
1.0
1.0
1.0
1.0
1.0
1.0
1.0
1.0
1.0
1.0
1.0
1.0
1.0
1.0
1.0
1.0
1.0
1.0
1.0
1.0
1.0
1.0
1.0
1.0


8
0
0
0
0
0
0
0
0
0
0
0
0
0
0
0
0
0
0
0
0
0
0
0
0
0
8
0
0
0

1.0
1.0
1.0
1.0
1.0
1.0
1.0
1.0
1.0
1.0
1.0
1.0
1.0
1.0
1.0
1.0
1.0
1.0
1.0
1.0
1.0
1.0
1.0
1.0
1.0
1.0
1.0
1.0
1.0
1.0


125
9
12
0
0
6
1
7
6
11
8
7
5
0
9
0
0
0
1
1
0
0
0
5
0
8
9
0
6
14

1.0
1.0
1.0
1.0
1.0
1.0
1.0
1.0
1.0
1.0
1.0
1.0
1.0
1.0
1.0
1.0
1.0
1.0
1.0
1.0
1.0
1.0
1.0
1.0
1.0
1.0
1.0
1.0
1.0
1.0


89
4
7
0
0
5
1
7
5
7
7
5
5
0
0
0
0
0
0
0
0
0
0
5
0
8
9
0
6
8

1.0
1.0
1.0
1.0
1.0
1.0
1.0
1.0
1.0
1.0
1.0
1.0
1.0
1.0
1.0
1.0
1.0
1.0
1.0
1.0
1.0
1.0
1.0
1.0
1.0
1.0
1.0
1.0
1.0
1.0


55
2
4
0
0
3
1
4
3
4
4
3
3
0
0
0
0
0
0
0
0
0
0
3
0
5
6
0
4
6

1.0
1.0
1.0
1.0
1.0
1.0
1.0
1.0
1.0
1.0
1.0
1.0
1.0
1.0
1.0
1.0
1.0
1.0
1.0
1.0
1.0
1.0
1.0
1.0
1.0
1.0
1.0
1.0
1.0
1.0


8
1
1
0
0
1
0
1
1
1
1
1
0
0
0
0
0
0
0
0
0
0
0
0
0
0
0
0
0
0

0.71
0.71
0.71
0.71
0.71
0.71
0.71
0.71
0.71
0.71
0.71
0.71
0.71
0.71
0.71
0.71
0.71
0.71
0.71
0.71
0.71
0.71
0.71
0.71
0.71
0.71
0.71
0.71
0.71
0.71


6
1
1
0
0
0
0
0
0
1
0
0
0
0
3
0
0
0
0
0
0
0
0
0
0
0
0
0
0
0


6
1
1
0
0
0
0
0
0
1
0
0
0
0
3
0
0
0
0
0
0
0
0
0
0
0
0
0
0
0

0.99
0.99
0.99
0.99
0.99
0.99
0.99
0.99
0.99
0.99
0.99
0.99
0.99
0.99
0.99
0.99
0.99
0.99
0.99
0.99
0.99
0.99
0.99
0.99
0.99
0.99
0.99
0.99
0.99
0.99


6
1
0
0
0
0
0
0
1
2
1
0
0
0
0
0
0
0
1
0
0
0
0
0
0
0
0
0
0
0

0.8
0.8
0.8
0.8
0.8
0.8
0.8
0.8
0.8
0.8
0.8
0.8
0.8
0.8
0.8
0.8
0.8
0.8
0.8
0.8
0.8
0.8
0.8
0.8
0.8
0.8
0.8
0.8
0.8
0.8


1
1
0
0
0
0
0
0
0
0
0
0
0
0
0
0
0
0
0
0
0
0
0
0
0
0
0
0
0
0

1.0
1.0
1.0
1.0
1.0
1.0
1.0
1.0
1.0
1.0
1.0
1.0
1.0
1.0
1.0
1.0
1.0
1.0
1.0
1.0
1.0
1.0
1.0
1.0
1.0
1.0
1.0
1.0
1.0
1.0


1
0
0
0
0
0
0
0
0
1
0
0
0
0
0
0
0
0
0
0
0
0
0
0
0
0
0
0
0
0

1.0
1.0
1.0
1.0
1.0
1.0
1.0
1.0
1.0
1.0
1.0
1.0
1.0
1.0
1.0
1.0
1.0
1.0
1.0
1.0
1.0
1.0
1.0
1.0
1.0
1.0
1.0
1.0
1.0
1.0


2
1
1
0
0
0
0
0
0
0
0
0
0
0
0
0
0
0
0
0
0
0
0
0
0
0
0
0
0
0


2
1
1
0
0
0
0
0
0
0
0
0
0
0
0
0
0
0
0
0
0
0
0
0
0
0
0
0
0
0

0.89
0.89
0.89
0.89
0.89
0.89
0.89
0.89
0.89
0.89
0.89
0.89
0.89
0.89
0.89
0.89
0.89
0.89
0.89
0.89
0.89
0.89
0.89
0.89
0.89
0.89
0.89
0.89
0.89
0.89


9
1
1
0
0
1
0
0
0
0
0
0
0
0
0
0
0
0
0
0
0
0
0
0
0
0
0
0
0
6

1.0
1.0
1.0
1.0
1.0
1.0
1.0
1.0
1.0
1.0
1.0
1.0
1.0
1.0
1.0
1.0
1.0
1.0
1.0
1.0
1.0
1.0
1.0
1.0
1.0
1.0
1.0
1.0
1.0
1.0


4
1
1
0
0
1
0
0
0
0
0
0
0
0
0
0
0
0
0
0
0
0
0
0
0
0
0
0
0
1

1.0
1.0
1.0
1.0
1.0
1.0
1.0
1.0
1.0
1.0
1.0
1.0
1.0
1.0
1.0
1.0
1.0
1.0
1.0
1.0
1.0
1.0
1.0
1.0
1.0
1.0
1.0
1.0
1.0
1.0


2
0
0
0
0
0
0
0
0
0
0
0
0
0
0
0
0
0
0
0
0
0
0
0
0
0
0
0
0
2

0.98
0.98
0.98
0.98
0.98
0.98
0.98
0.98
0.98
0.98
0.98
0.98
0.98
0.98
0.98
0.98
0.98
0.98
0.98
0.98
0.98
0.98
0.98
0.98
0.98
0.98
0.98
0.98
0.98
0.98


3
0
0
0
0
0
0
0
0
1
0
2
0
0
0
0
0
0
0
0
0
0
0
0
0
0
0
0
0
0


3
0
0
0
0
0
0
0
0
1
0
2
0
0
0
0
0
0
0
0
0
0
0
0
0
0
0
0
0
0

0.92
0.92
0.92
0.92
0.92
0.92
0.92
0.92
0.92
0.92
0.92
0.92
0.92
0.92
0.92
0.92
0.92
0.92
0.92
0.92
0.92
0.92
0.92
0.92
0.92
0.92
0.92
0.92
0.92
0.92


1
1
0
0
0
0
0
0
0
0
0
0
0
0
0
0
0
0
0
0
0
0
0
0
0
0
0
0
0
0


1
1
0
0
0
0
0
0
0
0
0
0
0
0
0
0
0
0
0
0
0
0
0
0
0
0
0
0
0
0

0.96
0.96
0.96
0.96
0.96
0.96
0.96
0.96
0.96
0.96
0.96
0.96
0.96
0.96
0.96
0.96
0.96
0.96
0.96
0.96
0.96
0.96
0.96
0.96
0.96
0.96
0.96
0.96
0.96
0.96


1
0
1
0
0
0
0
0
0
0
0
0
0
0
0
0
0
0
0
0
0
0
0
0
0
0
0
0
0
0

1.0
1.0
1.0
1.0
1.0
1.0
1.0
1.0
1.0
1.0
1.0
1.0
1.0
1.0
1.0
1.0
1.0
1.0
1.0
1.0
1.0
1.0
1.0
1.0
1.0
1.0
1.0
1.0
1.0
1.0


2
0
1
0
0
0
0
0
0
0
0
0
0
0
0
0
0
0
0
1
0
0
0
0
0
0
0
0
0
0


2
0
1
0
0
0
0
0
0
0
0
0
0
0
0
0
0
0
0
1
0
0
0
0
0
0
0
0
0
0

1.0
1.0
1.0
1.0
1.0
1.0
1.0
1.0
1.0
1.0
1.0
1.0
1.0
1.0
1.0
1.0
1.0
1.0
1.0
1.0
1.0
1.0
1.0
1.0
1.0
1.0
1.0
1.0
1.0
1.0


1
0
0
0
0
0
0
0
0
0
0
0
0
0
1
0
0
0
0
0
0
0
0
0
0
0
0
0
0
0


1
0
0
0
0
0
0
0
0
0
0
0
0
0
1
0
0
0
0
0
0
0
0
0
0
0
0
0
0
0

1.0
1.0
1.0
1.0
1.0
1.0
1.0
1.0
1.0
1.0
1.0
1.0
1.0
1.0
1.0
1.0
1.0
1.0
1.0
1.0
1.0
1.0
1.0
1.0
1.0
1.0
1.0
1.0
1.0
1.0


5
1
1
1
0
0
0
0
0
0
1
1
0
0
0
0
0
0
0
0
0
0
0
0
0
0
0
0
0
0

0.78
0.78
0.78
0.78
0.78
0.78
0.78
0.78
0.78
0.78
0.78
0.78
0.78
0.78
0.78
0.78
0.78
0.78
0.78
0.78
0.78
0.78
0.78
0.78
0.78
0.78
0.78
0.78
0.78
0.78


102
3
1
0
0
1
1
10
15
10
9
2
1
0
6
0
2
4
4
1
0
0
0
1
2
9
8
3
1
8

0.91
0.91
0.91
0.91
0.91
0.91
0.91
0.91
0.91
0.91
0.91
0.91
0.91
0.91
0.91
0.91
0.91
0.91
0.91
0.91
0.91
0.91
0.91
0.91
0.91
0.91
0.91
0.91
0.91
0.91


52
2
1
0
0
1
1
4
6
3
3
2
1
0
5
0
2
4
4
1
0
0
0
1
2
2
3
1
0
3

1.0
1.0
1.0
1.0
1.0
1.0
1.0
1.0
1.0
1.0
1.0
1.0
1.0
1.0
1.0
1.0
1.0
1.0
1.0
1.0
1.0
1.0
1.0
1.0
1.0
1.0
1.0
1.0
1.0
1.0


20
2
0
0
0
1
1
4
4
3
3
2
0
0
0
0
0
0
0
0
0
0
0
0
0
0
0
0
0
0

0.99
0.99
0.99
0.99
0.99
0.99
0.99
0.99
0.99
0.99
0.99
0.99
0.99
0.99
0.99
0.99
0.99
0.99
0.99
0.99
0.99
0.99
0.99
0.99
0.99
0.99
0.99
0.99
0.99
0.99


2
0
1
0
0
0
0
0
0
0
0
0
0
0
1
0
0
0
0
0
0
0
0
0
0
0
0
0
0
0

1.0
1.0
1.0
1.0
1.0
1.0
1.0
1.0
1.0
1.0
1.0
1.0
1.0
1.0
1.0
1.0
1.0
1.0
1.0
1.0
1.0
1.0
1.0
1.0
1.0
1.0
1.0
1.0
1.0
1.0


3
0
0
0
0
0
0
0
1
0
0
0
0
0
0
0
1
0
1
0
0
0
0
0
0
0
0
0
0
0

1.0
1.0
1.0
1.0
1.0
1.0
1.0
1.0
1.0
1.0
1.0
1.0
1.0
1.0
1.0
1.0
1.0
1.0
1.0
1.0
1.0
1.0
1.0
1.0
1.0
1.0
1.0
1.0
1.0
1.0


1
0
0
0
0
0
0
0
1
0
0
0
0
0
0
0
0
0
0
0
0
0
0
0
0
0
0
0
0
0

1.0
1.0
1.0
1.0
1.0
1.0
1.0
1.0
1.0
1.0
1.0
1.0
1.0
1.0
1.0
1.0
1.0
1.0
1.0
1.0
1.0
1.0
1.0
1.0
1.0
1.0
1.0
1.0
1.0
1.0


4
0
0
0
0
0
0
0
0
0
0
0
0
0
0
0
0
0
0
0
0
0
0
0
2
0
0
0
0
2

1.0
1.0
1.0
1.0
1.0
1.0
1.0
1.0
1.0
1.0
1.0
1.0
1.0
1.0
1.0
1.0
1.0
1.0
1.0
1.0
1.0
1.0
1.0
1.0
1.0
1.0
1.0
1.0
1.0
1.0


28
0
0
0
0
0
0
1
4
3
3
0
0
0
1
0
0
0
0
0
0
0
0
0
0
5
4
2
0
5

1.0
1.0
1.0
1.0
1.0
1.0
1.0
1.0
1.0
1.0
1.0
1.0
1.0
1.0
1.0
1.0
1.0
1.0
1.0
1.0
1.0
1.0
1.0
1.0
1.0
1.0
1.0
1.0
1.0
1.0


11
0
0
0
0
0
0
0
1
1
1
0
0
0
0
0
0
0
0
0
0
0
0
0
0
3
2
0
0
3

1.0
1.0
1.0
1.0
1.0
1.0
1.0
1.0
1.0
1.0
1.0
1.0
1.0
1.0
1.0
1.0
1.0
1.0
1.0
1.0
1.0
1.0
1.0
1.0
1.0
1.0
1.0
1.0
1.0
1.0


8
0
0
0
0
0
0
0
0
0
0
0
0
0
0
0
0
0
0
0
0
0
0
0
0
2
2
2
0
2

1.0
1.0
1.0
1.0
1.0
1.0
1.0
1.0
1.0
1.0
1.0
1.0
1.0
1.0
1.0
1.0
1.0
1.0
1.0
1.0
1.0
1.0
1.0
1.0
1.0
1.0
1.0
1.0
1.0
1.0


1
0
0
0
0
0
0
0
0
0
0
0
0
0
1
0
0
0
0
0
0
0
0
0
0
0
0
0
0
0

0.96
0.96
0.96
0.96
0.96
0.96
0.96
0.96
0.96
0.96
0.96
0.96
0.96
0.96
0.96
0.96
0.96
0.96
0.96
0.96
0.96
0.96
0.96
0.96
0.96
0.96
0.96
0.96
0.96
0.96


4
0
0
0
0
0
0
2
2
0
0
0
0
0
0
0
0
0
0
0
0
0
0
0
0
0
0
0
0
0

1.0
1.0
1.0
1.0
1.0
1.0
1.0
1.0
1.0
1.0
1.0
1.0
1.0
1.0
1.0
1.0
1.0
1.0
1.0
1.0
1.0
1.0
1.0
1.0
1.0
1.0
1.0
1.0
1.0
1.0


1
0
0
0
0
0
0
0
1
0
0
0
0
0
0
0
0
0
0
0
0
0
0
0
0
0
0
0
0
0

1.0
1.0
1.0
1.0
1.0
1.0
1.0
1.0
1.0
1.0
1.0
1.0
1.0
1.0
1.0
1.0
1.0
1.0
1.0
1.0
1.0
1.0
1.0
1.0
1.0
1.0
1.0
1.0
1.0
1.0


6
0
0
0
0
0
0
0
0
1
1
0
0
0
0
0
0
0
0
0
0
0
0
0
0
2
1
0
1
0

1.0
1.0
1.0
1.0
1.0
1.0
1.0
1.0
1.0
1.0
1.0
1.0
1.0
1.0
1.0
1.0
1.0
1.0
1.0
1.0
1.0
1.0
1.0
1.0
1.0
1.0
1.0
1.0
1.0
1.0


2
0
0
0
0
0
0
0
0
1
1
0
0
0
0
0
0
0
0
0
0
0
0
0
0
0
0
0
0
0

0.98
0.98
0.98
0.98
0.98
0.98
0.98
0.98
0.98
0.98
0.98
0.98
0.98
0.98
0.98
0.98
0.98
0.98
0.98
0.98
0.98
0.98
0.98
0.98
0.98
0.98
0.98
0.98
0.98
0.98


2
0
0
0
0
0
0
0
0
0
0
0
0
0
0
0
0
0
0
0
0
0
0
0
0
1
1
0
0
0

0.96
0.96
0.96
0.96
0.96
0.96
0.96
0.96
0.96
0.96
0.96
0.96
0.96
0.96
0.96
0.96
0.96
0.96
0.96
0.96
0.96
0.96
0.96
0.96
0.96
0.96
0.96
0.96
0.96
0.96


1
0
0
0
0
0
0
0
0
0
0
0
0
0
0
0
0
0
0
0
0
0
0
0
0
0
0
0
1
0

0.9
0.9
0.9
0.9
0.9
0.9
0.9
0.9
0.9
0.9
0.9
0.9
0.9
0.9
0.9
0.9
0.9
0.9
0.9
0.9
0.9
0.9
0.9
0.9
0.9
0.9
0.9
0.9
0.9
0.9


34
4
2
0
0
0
0
2
3
10
8
3
1
0
0
0
0
0
0
0
0
0
0
1
0
0
0
0
0
0


26
3
1
0
0
0
0
2
2
8
7
3
0
0
0
0
0
0
0
0
0
0
0
0
0
0
0
0
0
0

1.0
1.0
1.0
1.0
1.0
1.0
1.0
1.0
1.0
1.0
1.0
1.0
1.0
1.0
1.0
1.0
1.0
1.0
1.0
1.0
1.0
1.0
1.0
1.0
1.0
1.0
1.0
1.0
1.0
1.0


4
0
0
0
0
0
0
0
0
3
1
0
0
0
0
0
0
0
0
0
0
0
0
0
0
0
0
0
0
0

1.0
1.0
1.0
1.0
1.0
1.0
1.0
1.0
1.0
1.0
1.0
1.0
1.0
1.0
1.0
1.0
1.0
1.0
1.0
1.0
1.0
1.0
1.0
1.0
1.0
1.0
1.0
1.0
1.0
1.0


2
0
0
0
0
0
0
0
0
0
1
1
0
0
0
0
0
0
0
0
0
0
0
0
0
0
0
0
0
0

1.0
1.0
1.0
1.0
1.0
1.0
1.0
1.0
1.0
1.0
1.0
1.0
1.0
1.0
1.0
1.0
1.0
1.0
1.0
1.0
1.0
1.0
1.0
1.0
1.0
1.0
1.0
1.0
1.0
1.0


2
1
0
0
0
0
0
0
0
1
0
0
0
0
0
0
0
0
0
0
0
0
0
0
0
0
0
0
0
0

1.0
1.0
1.0
1.0
1.0
1.0
1.0
1.0
1.0
1.0
1.0
1.0
1.0
1.0
1.0
1.0
1.0
1.0
1.0
1.0
1.0
1.0
1.0
1.0
1.0
1.0
1.0
1.0
1.0
1.0


4
0
0
0
0
0
0
0
1
2
1
0
0
0
0
0
0
0
0
0
0
0
0
0
0
0
0
0
0
0

1.0
1.0
1.0
1.0
1.0
1.0
1.0
1.0
1.0
1.0
1.0
1.0
1.0
1.0
1.0
1.0
1.0
1.0
1.0
1.0
1.0
1.0
1.0
1.0
1.0
1.0
1.0
1.0
1.0
1.0


1
0
0
0
0
0
0
0
0
1
0
0
0
0
0
0
0
0
0
0
0
0
0
0
0
0
0
0
0
0

0.82
0.82
0.82
0.82
0.82
0.82
0.82
0.82
0.82
0.82
0.82
0.82
0.82
0.82
0.82
0.82
0.82
0.82
0.82
0.82
0.82
0.82
0.82
0.82
0.82
0.82
0.82
0.82
0.82
0.82


4
1
1
0
0
0
0
0
0
0
0
0
1
0
0
0
0
0
0
0
0
0
0
1
0
0
0
0
0
0


4
1
1
0
0
0
0
0
0
0
0
0
1
0
0
0
0
0
0
0
0
0
0
1
0
0
0
0
0
0

1.0
1.0
1.0
1.0
1.0
1.0
1.0
1.0
1.0
1.0
1.0
1.0
1.0
1.0
1.0
1.0
1.0
1.0
1.0
1.0
1.0
1.0
1.0
1.0
1.0
1.0
1.0
1.0
1.0
1.0


90
9
0
0
0
0
0
10
0
32
28
11
0
0
0
0
0
0
0
0
0
0
0
0
0
0
0
0
0
0


90
9
0
0
0
0
0
10
0
32
28
11
0
0
0
0
0
0
0
0
0
0
0
0
0
0
0
0
0
0

1.0
1.0
1.0
1.0
1.0
1.0
1.0
1.0
1.0
1.0
1.0
1.0
1.0
1.0
1.0
1.0
1.0
1.0
1.0
1.0
1.0
1.0
1.0
1.0
1.0
1.0
1.0
1.0
1.0
1.0


1165
84
57
34
27
35
24
53
77
93
77
68
25
26
34
15
14
80
78
47
39
37
24
25
20
10
11
12
21
18

1.0
1.0
1.0
1.0
1.0
1.0
1.0
1.0
1.0
1.0
1.0
1.0
1.0
1.0
1.0
1.0
1.0
1.0
1.0
1.0
1.0
1.0
1.0
1.0
1.0
1.0
1.0
1.0
1.0
1.0


813
57
33
15
13
16
11
47
63
71
54
42
25
6
22
8
8
58
66
40
32
23
24
25
15
7
6
6
11
9

1.0
1.0
1.0
1.0
1.0
1.0
1.0
1.0
1.0
1.0
1.0
1.0
1.0
1.0
1.0
1.0
1.0
1.0
1.0
1.0
1.0
1.0
1.0
1.0
1.0
1.0
1.0
1.0
1.0
1.0


40
3
2
1
0
0
0
2
4
4
4
1
2
0
0
1
0
2
2
0
0
0
4
2
4
0
0
0
2
0

1.0
1.0
1.0
1.0
1.0
1.0
1.0
1.0
1.0
1.0
1.0
1.0
1.0
1.0
1.0
1.0
1.0
1.0
1.0
1.0
1.0
1.0
1.0
1.0
1.0
1.0
1.0
1.0
1.0
1.0


28
2
2
1
1
0
0
1
1
2
2
2
0
0
8
0
0
2
4
0
0
0
0
0
0
0
0
0
0
0

1.0
1.0
1.0
1.0
1.0
1.0
1.0
1.0
1.0
1.0
1.0
1.0
1.0
1.0
1.0
1.0
1.0
1.0
1.0
1.0
1.0
1.0
1.0
1.0
1.0
1.0
1.0
1.0
1.0
1.0


33
2
0
0
0
1
0
1
3
4
5
1
1
0
0
0
0
8
1
1
1
1
2
1
0
0
0
0
0
0

1.0
1.0
1.0
1.0
1.0
1.0
1.0
1.0
1.0
1.0
1.0
1.0
1.0
1.0
1.0
1.0
1.0
1.0
1.0
1.0
1.0
1.0
1.0
1.0
1.0
1.0
1.0
1.0
1.0
1.0


8
1
0
0
0
0
0
1
1
1
1
0
1
0
0
0
0
0
0
0
0
0
1
1
0
0
0
0
0
0

1.0
1.0
1.0
1.0
1.0
1.0
1.0
1.0
1.0
1.0
1.0
1.0
1.0
1.0
1.0
1.0
1.0
1.0
1.0
1.0
1.0
1.0
1.0
1.0
1.0
1.0
1.0
1.0
1.0
1.0


18
1
0
0
0
0
0
2
3
4
3
1
0
0
0
0
0
2
2
0
0
0
0
0
0
0
0
0
0
0

1.0
1.0
1.0
1.0
1.0
1.0
1.0
1.0
1.0
1.0
1.0
1.0
1.0
1.0
1.0
1.0
1.0
1.0
1.0
1.0
1.0
1.0
1.0
1.0
1.0
1.0
1.0
1.0
1.0
1.0


25
3
1
0
0
0
0
0
3
3
3
3
1
0
0
0
0
1
1
2
1
1
1
1
0
0
0
0
0
0

1.0
1.0
1.0
1.0
1.0
1.0
1.0
1.0
1.0
1.0
1.0
1.0
1.0
1.0
1.0
1.0
1.0
1.0
1.0
1.0
1.0
1.0
1.0
1.0
1.0
1.0
1.0
1.0
1.0
1.0


10
2
1
0
0
0
0
1
0
1
1
1
0
0
0
0
0
1
0
0
1
0
0
0
1
0
0
0
0
0

1.0
1.0
1.0
1.0
1.0
1.0
1.0
1.0
1.0
1.0
1.0
1.0
1.0
1.0
1.0
1.0
1.0
1.0
1.0
1.0
1.0
1.0
1.0
1.0
1.0
1.0
1.0
1.0
1.0
1.0


9
1
0
0
0
0
0
1
1
1
0
0
0
1
0
0
0
0
0
2
1
0
0
0
0
0
0
0
0
1

1.0
1.0
1.0
1.0
1.0
1.0
1.0
1.0
1.0
1.0
1.0
1.0
1.0
1.0
1.0
1.0
1.0
1.0
1.0
1.0
1.0
1.0
1.0
1.0
1.0
1.0
1.0
1.0
1.0
1.0


17
1
1
0
0
0
0
1
1
1
1
1
0
0
0
0
0
4
2
0
1
1
2
0
0
0
0
0
0
0

1.0
1.0
1.0
1.0
1.0
1.0
1.0
1.0
1.0
1.0
1.0
1.0
1.0
1.0
1.0
1.0
1.0
1.0
1.0
1.0
1.0
1.0
1.0
1.0
1.0
1.0
1.0
1.0
1.0
1.0


33
4
1
0
0
0
0
0
3
5
2
9
4
1
0
0
0
0
0
0
0
0
0
4
0
0
0
0
0
0

1.0
1.0
1.0
1.0
1.0
1.0
1.0
1.0
1.0
1.0
1.0
1.0
1.0
1.0
1.0
1.0
1.0
1.0
1.0
1.0
1.0
1.0
1.0
1.0
1.0
1.0
1.0
1.0
1.0
1.0


7
1
1
0
1
0
0
0
1
1
1
1
0
0
0
0
0
0
0
0
0
0
0
0
0
0
0
0
0
0

0.93
0.93
0.93
0.93
0.93
0.93
0.93
0.93
0.93
0.93
0.93
0.93
0.93
0.93
0.93
0.93
0.93
0.93
0.93
0.93
0.93
0.93
0.93
0.93
0.93
0.93
0.93
0.93
0.93
0.93


15
0
1
1
1
1
1
0
0
0
0
0
0
1
1
0
1
1
1
0
1
0
0
0
1
1
0
0
1
1

1.0
1.0
1.0
1.0
1.0
1.0
1.0
1.0
1.0
1.0
1.0
1.0
1.0
1.0
1.0
1.0
1.0
1.0
1.0
1.0
1.0
1.0
1.0
1.0
1.0
1.0
1.0
1.0
1.0
1.0


7
1
1
0
0
0
0
0
1
1
0
1
0
0
0
0
0
0
1
0
0
0
0
0
1
0
0
0
0
0

1.0
1.0
1.0
1.0
1.0
1.0
1.0
1.0
1.0
1.0
1.0
1.0
1.0
1.0
1.0
1.0
1.0
1.0
1.0
1.0
1.0
1.0
1.0
1.0
1.0
1.0
1.0
1.0
1.0
1.0


16
0
1
0
1
2
1
0
0
1
0
0
0
0
1
1
1
1
1
0
0
0
0
0
1
1
1
0
1
1

1.0
1.0
1.0
1.0
1.0
1.0
1.0
1.0
1.0
1.0
1.0
1.0
1.0
1.0
1.0
1.0
1.0
1.0
1.0
1.0
1.0
1.0
1.0
1.0
1.0
1.0
1.0
1.0
1.0
1.0


6
1
1
1
0
0
0
0
1
1
1
0
0
0
0
0
0
0
0
0
0
0
0
0
0
0
0
0
0
0

0.71
0.71
0.71
0.71
0.71
0.71
0.71
0.71
0.71
0.71
0.71
0.71
0.71
0.71
0.71
0.71
0.71
0.71
0.71
0.71
0.71
0.71
0.71
0.71
0.71
0.71
0.71
0.71
0.71
0.71


5
1
0
0
0
0
0
0
0
1
1
0
0
0
0
0
0
1
1
0
0
0
0
0
0
0
0
0
0
0

1.0
1.0
1.0
1.0
1.0
1.0
1.0
1.0
1.0
1.0
1.0
1.0
1.0
1.0
1.0
1.0
1.0
1.0
1.0
1.0
1.0
1.0
1.0
1.0
1.0
1.0
1.0
1.0
1.0
1.0


1
0
0
0
0
0
0
1
0
0
0
0
0
0
0
0
0
0
0
0
0
0
0
0
0
0
0
0
0
0

1.0
1.0
1.0
1.0
1.0
1.0
1.0
1.0
1.0
1.0
1.0
1.0
1.0
1.0
1.0
1.0
1.0
1.0
1.0
1.0
1.0
1.0
1.0
1.0
1.0
1.0
1.0
1.0
1.0
1.0


2
1
1
0
0
0
0
0
0
0
0
0
0
0
0
0
0
0
0
0
0
0
0
0
0
0
0
0
0
0

1.0
1.0
1.0
1.0
1.0
1.0
1.0
1.0
1.0
1.0
1.0
1.0
1.0
1.0
1.0
1.0
1.0
1.0
1.0
1.0
1.0
1.0
1.0
1.0
1.0
1.0
1.0
1.0
1.0
1.0


28
0
0
0
0
0
0
0
0
0
0
0
3
0
0
0
0
2
5
7
6
2
0
3
0
0
0
0
0
0

1.0
1.0
1.0
1.0
1.0
1.0
1.0
1.0
1.0
1.0
1.0
1.0
1.0
1.0
1.0
1.0
1.0
1.0
1.0
1.0
1.0
1.0
1.0
1.0
1.0
1.0
1.0
1.0
1.0
1.0


2
0
0
0
0
0
0
0
0
0
0
0
0
0
0
0
0
0
2
0
0
0
0
0
0
0
0
0
0
0

1.0
1.0
1.0
1.0
1.0
1.0
1.0
1.0
1.0
1.0
1.0
1.0
1.0
1.0
1.0
1.0
1.0
1.0
1.0
1.0
1.0
1.0
1.0
1.0
1.0
1.0
1.0
1.0
1.0
1.0


5
0
0
0
0
0
0
0
0
0
0
0
0
0
0
0
0
0
1
2
1
1
0
0
0
0
0
0
0
0

1.0
1.0
1.0
1.0
1.0
1.0
1.0
1.0
1.0
1.0
1.0
1.0
1.0
1.0
1.0
1.0
1.0
1.0
1.0
1.0
1.0
1.0
1.0
1.0
1.0
1.0
1.0
1.0
1.0
1.0


3
0
0
0
0
0
0
0
0
0
0
0
0
0
0
0
0
0
3
0
0
0
0
0
0
0
0
0
0
0

1.0
1.0
1.0
1.0
1.0
1.0
1.0
1.0
1.0
1.0
1.0
1.0
1.0
1.0
1.0
1.0
1.0
1.0
1.0
1.0
1.0
1.0
1.0
1.0
1.0
1.0
1.0
1.0
1.0
1.0


1
0
0
0
0
0
0
0
0
0
0
0
0
0
0
0
0
0
1
0
0
0
0
0
0
0
0
0
0
0

0.71
0.71
0.71
0.71
0.71
0.71
0.71
0.71
0.71
0.71
0.71
0.71
0.71
0.71
0.71
0.71
0.71
0.71
0.71
0.71
0.71
0.71
0.71
0.71
0.71
0.71
0.71
0.71
0.71
0.71


153
16
14
12
6
9
5
0
6
6
14
15
0
3
4
3
3
4
5
3
0
8
0
0
3
3
2
3
3
3

1.0
1.0
1.0
1.0
1.0
1.0
1.0
1.0
1.0
1.0
1.0
1.0
1.0
1.0
1.0
1.0
1.0
1.0
1.0
1.0
1.0
1.0
1.0
1.0
1.0
1.0
1.0
1.0
1.0
1.0


36
6
5
3
0
0
0
0
6
3
7
6
0
0
0
0
0
0
0
0
0
0
0
0
0
0
0
0
0
0

1.0
1.0
1.0
1.0
1.0
1.0
1.0
1.0
1.0
1.0
1.0
1.0
1.0
1.0
1.0
1.0
1.0
1.0
1.0
1.0
1.0
1.0
1.0
1.0
1.0
1.0
1.0
1.0
1.0
1.0


21
1
1
1
1
1
0
1
2
3
0
1
0
9
0
0
0
0
0
0
0
0
0
0
0
0
0
0
0
0

0.7
0.7
0.7
0.7
0.7
0.7
0.7
0.7
0.7
0.7
0.7
0.7
0.7
0.7
0.7
0.7
0.7
0.7
0.7
0.7
0.7
0.7
0.7
0.7
0.7
0.7
0.7
0.7
0.7
0.7


19
1
1
1
1
1
0
0
2
2
0
1
0
9
0
0
0
0
0
0
0
0
0
0
0
0
0
0
0
0

1.0
1.0
1.0
1.0
1.0
1.0
1.0
1.0
1.0
1.0
1.0
1.0
1.0
1.0
1.0
1.0
1.0
1.0
1.0
1.0
1.0
1.0
1.0
1.0
1.0
1.0
1.0
1.0
1.0
1.0


12
1
0
0
0
0
0
2
0
4
4
0
0
0
0
0
0
0
0
0
0
1
0
0
0
0
0
0
0
0

1.0
1.0
1.0
1.0
1.0
1.0
1.0
1.0
1.0
1.0
1.0
1.0
1.0
1.0
1.0
1.0
1.0
1.0
1.0
1.0
1.0
1.0
1.0
1.0
1.0
1.0
1.0
1.0
1.0
1.0


69
3
3
1
2
3
2
0
0
2
2
3
0
3
3
4
3
5
7
4
3
2
0
0
2
0
3
3
3
3


69
3
3
1
2
3
2
0
0
2
2
3
0
3
3
4
3
5
7
4
3
2
0
0
2
0
3
3
3
3

1.0
1.0
1.0
1.0
1.0
1.0
1.0
1.0
1.0
1.0
1.0
1.0
1.0
1.0
1.0
1.0
1.0
1.0
1.0
1.0
1.0
1.0
1.0
1.0
1.0
1.0
1.0
1.0
1.0
1.0


54
2
4
3
1
4
3
0
2
3
0
3
0
5
5
0
0
13
0
0
0
0
0
0
0
0
0
0
3
3


54
2
4
3
1
4
3
0
2
3
0
3
0
5
5
0
0
13
0
0
0
0
0
0
0
0
0
0
3
3

1.0
1.0
1.0
1.0
1.0
1.0
1.0
1.0
1.0
1.0
1.0
1.0
1.0
1.0
1.0
1.0
1.0
1.0
1.0
1.0
1.0
1.0
1.0
1.0
1.0
1.0
1.0
1.0
1.0
1.0


1
0
0
0
0
0
0
0
0
0
0
1
0
0
0
0
0
0
0
0
0
0
0
0
0
0
0
0
0
0

1.0
1.0
1.0
1.0
1.0
1.0
1.0
1.0
1.0
1.0
1.0
1.0
1.0
1.0
1.0
1.0
1.0
1.0
1.0
1.0
1.0
1.0
1.0
1.0
1.0
1.0
1.0
1.0
1.0
1.0


78
3
1
0
0
0
0
4
5
3
5
2
9
0
0
0
0
0
0
5
7
5
8
9
2
0
0
0
0
10


78
3
1
0
0
0
0
4
5
3
5
2
9
0
0
0
0
0
0
5
7
5
8
9
2
0
0
0
0
10

1.0
1.0
1.0
1.0
1.0
1.0
1.0
1.0
1.0
1.0
1.0
1.0
1.0
1.0
1.0
1.0
1.0
1.0
1.0
1.0
1.0
1.0
1.0
1.0
1.0
1.0
1.0
1.0
1.0
1.0


49
2
0
0
0
0
0
4
5
2
3
0
8
0
0
0
0
0
0
0
0
0
7
8
0
0
0
0
0
10

1.0
1.0
1.0
1.0
1.0
1.0
1.0
1.0
1.0
1.0
1.0
1.0
1.0
1.0
1.0
1.0
1.0
1.0
1.0
1.0
1.0
1.0
1.0
1.0
1.0
1.0
1.0
1.0
1.0
1.0


4
0
0
0
0
0
0
0
0
0
1
2
0
0
0
0
0
0
0
0
0
0
0
0
1
0
0
0
0
0

1.0
1.0
1.0
1.0
1.0
1.0
1.0
1.0
1.0
1.0
1.0
1.0
1.0
1.0
1.0
1.0
1.0
1.0
1.0
1.0
1.0
1.0
1.0
1.0
1.0
1.0
1.0
1.0
1.0
1.0


12
0
0
0
0
0
0
0
0
0
0
0
0
0
0
0
0
0
0
5
7
0
0
0
0
0
0
0
0
0

1.0
1.0
1.0
1.0
1.0
1.0
1.0
1.0
1.0
1.0
1.0
1.0
1.0
1.0
1.0
1.0
1.0
1.0
1.0
1.0
1.0
1.0
1.0
1.0
1.0
1.0
1.0
1.0
1.0
1.0


319
6
4
1
0
0
0
62
85
75
49
29
0
0
2
0
0
1
2
0
0
0
0
0
1
0
1
0
1
0

0.99
0.99
0.99
0.99
0.99
0.99
0.99
0.99
0.99
0.99
0.99
0.99
0.99
0.99
0.99
0.99
0.99
0.99
0.99
0.99
0.99
0.99
0.99
0.99
0.99
0.99
0.99
0.99
0.99
0.99


58
1
1
0
0
0
0
27
16
9
2
2
0
0
0
0
0
0
0
0
0
0
0
0
0
0
0
0
0
0

0.93
0.93
0.93
0.93
0.93
0.93
0.93
0.93
0.93
0.93
0.93
0.93
0.93
0.93
0.93
0.93
0.93
0.93
0.93
0.93
0.93
0.93
0.93
0.93
0.93
0.93
0.93
0.93
0.93
0.93


40
1
1
0
0
0
0
22
11
3
1
1
0
0
0
0
0
0
0
0
0
0
0
0
0
0
0
0
0
0

1.0
1.0
1.0
1.0
1.0
1.0
1.0
1.0
1.0
1.0
1.0
1.0
1.0
1.0
1.0
1.0
1.0
1.0
1.0
1.0
1.0
1.0
1.0
1.0
1.0
1.0
1.0
1.0
1.0
1.0


8
0
0
0
0
0
0
2
2
2
1
1
0
0
0
0
0
0
0
0
0
0
0
0
0
0
0
0
0
0

1.0
1.0
1.0
1.0
1.0
1.0
1.0
1.0
1.0
1.0
1.0
1.0
1.0
1.0
1.0
1.0
1.0
1.0
1.0
1.0
1.0
1.0
1.0
1.0
1.0
1.0
1.0
1.0
1.0
1.0


5
0
0
0
0
0
0
1
2
2
0
0
0
0
0
0
0
0
0
0
0
0
0
0
0
0
0
0
0
0

1.0
1.0
1.0
1.0
1.0
1.0
1.0
1.0
1.0
1.0
1.0
1.0
1.0
1.0
1.0
1.0
1.0
1.0
1.0
1.0
1.0
1.0
1.0
1.0
1.0
1.0
1.0
1.0
1.0
1.0


1
0
0
0
0
0
0
0
0
1
0
0
0
0
0
0
0
0
0
0
0
0
0
0
0
0
0
0
0
0

1.0
1.0
1.0
1.0
1.0
1.0
1.0
1.0
1.0
1.0
1.0
1.0
1.0
1.0
1.0
1.0
1.0
1.0
1.0
1.0
1.0
1.0
1.0
1.0
1.0
1.0
1.0
1.0
1.0
1.0


103
2
1
1
0
0
0
13
28
27
19
12
0
0
0
0
0
0
0
0
0
0
0
0
0
0
0
0
0
0

0.96
0.96
0.96
0.96
0.96
0.96
0.96
0.96
0.96
0.96
0.96
0.96
0.96
0.96
0.96
0.96
0.96
0.96
0.96
0.96
0.96
0.96
0.96
0.96
0.96
0.96
0.96
0.96
0.96
0.96


29
2
1
1
0
0
0
5
6
6
3
5
0
0
0
0
0
0
0
0
0
0
0
0
0
0
0
0
0
0

1.0
1.0
1.0
1.0
1.0
1.0
1.0
1.0
1.0
1.0
1.0
1.0
1.0
1.0
1.0
1.0
1.0
1.0
1.0
1.0
1.0
1.0
1.0
1.0
1.0
1.0
1.0
1.0
1.0
1.0


23
0
0
0
0
0
0
5
5
6
4
3
0
0
0
0
0
0
0
0
0
0
0
0
0
0
0
0
0
0


23
0
0
0
0
0
0
5
5
6
4
3
0
0
0
0
0
0
0
0
0
0
0
0
0
0
0
0
0
0

1.0
1.0
1.0
1.0
1.0
1.0
1.0
1.0
1.0
1.0
1.0
1.0
1.0
1.0
1.0
1.0
1.0
1.0
1.0
1.0
1.0
1.0
1.0
1.0
1.0
1.0
1.0
1.0
1.0
1.0


40
0
0
0
0
0
0
7
11
13
5
4
0
0
0
0
0
0
0
0
0
0
0
0
0
0
0
0
0
0

1.0
1.0
1.0
1.0
1.0
1.0
1.0
1.0
1.0
1.0
1.0
1.0
1.0
1.0
1.0
1.0
1.0
1.0
1.0
1.0
1.0
1.0
1.0
1.0
1.0
1.0
1.0
1.0
1.0
1.0


11
0
0
0
0
0
0
0
3
3
3
2
0
0
0
0
0
0
0
0
0
0
0
0
0
0
0
0
0
0

1.0
1.0
1.0
1.0
1.0
1.0
1.0
1.0
1.0
1.0
1.0
1.0
1.0
1.0
1.0
1.0
1.0
1.0
1.0
1.0
1.0
1.0
1.0
1.0
1.0
1.0
1.0
1.0
1.0
1.0


14
0
0
0
0
0
0
4
2
4
2
2
0
0
0
0
0
0
0
0
0
0
0
0
0
0
0
0
0
0

1.0
1.0
1.0
1.0
1.0
1.0
1.0
1.0
1.0
1.0
1.0
1.0
1.0
1.0
1.0
1.0
1.0
1.0
1.0
1.0
1.0
1.0
1.0
1.0
1.0
1.0
1.0
1.0
1.0
1.0


9
0
0
0
0
0
0
0
4
3
2
0
0
0
0
0
0
0
0
0
0
0
0
0
0
0
0
0
0
0

0.93
0.93
0.93
0.93
0.93
0.93
0.93
0.93
0.93
0.93
0.93
0.93
0.93
0.93
0.93
0.93
0.93
0.93
0.93
0.93
0.93
0.93
0.93
0.93
0.93
0.93
0.93
0.93
0.93
0.93


4
0
0
0
0
0
0
1
2
0
0
0
0
0
0
0
0
0
1
0
0
0
0
0
0
0
0
0
0
0


2
0
0
0
0
0
0
1
1
0
0
0
0
0
0
0
0
0
0
0
0
0
0
0
0
0
0
0
0
0

1.0
1.0
1.0
1.0
1.0
1.0
1.0
1.0
1.0
1.0
1.0
1.0
1.0
1.0
1.0
1.0
1.0
1.0
1.0
1.0
1.0
1.0
1.0
1.0
1.0
1.0
1.0
1.0
1.0
1.0


2
0
0
0
0
0
0
0
1
0
0
0
0
0
0
0
0
0
1
0
0
0
0
0
0
0
0
0
0
0

0.97
0.97
0.97
0.97
0.97
0.97
0.97
0.97
0.97
0.97
0.97
0.97
0.97
0.97
0.97
0.97
0.97
0.97
0.97
0.97
0.97
0.97
0.97
0.97
0.97
0.97
0.97
0.97
0.97
0.97


4
0
0
0
0
0
0
0
0
1
2
1
0
0
0
0
0
0
0
0
0
0
0
0
0
0
0
0
0
0

0.75
0.75
0.75
0.75
0.75
0.75
0.75
0.75
0.75
0.75
0.75
0.75
0.75
0.75
0.75
0.75
0.75
0.75
0.75
0.75
0.75
0.75
0.75
0.75
0.75
0.75
0.75
0.75
0.75
0.75


21
0
0
0
0
0
0
1
7
6
5
2
0
0
0
0
0
0
0
0
0
0
0
0
0
0
0
0
0
0

1.0
1.0
1.0
1.0
1.0
1.0
1.0
1.0
1.0
1.0
1.0
1.0
1.0
1.0
1.0
1.0
1.0
1.0
1.0
1.0
1.0
1.0
1.0
1.0
1.0
1.0
1.0
1.0
1.0
1.0


8
0
0
0
0
0
0
0
3
2
3
0
0
0
0
0
0
0
0
0
0
0
0
0
0
0
0
0
0
0

1.0
1.0
1.0
1.0
1.0
1.0
1.0
1.0
1.0
1.0
1.0
1.0
1.0
1.0
1.0
1.0
1.0
1.0
1.0
1.0
1.0
1.0
1.0
1.0
1.0
1.0
1.0
1.0
1.0
1.0


2
0
0
0
0
0
0
0
1
1
0
0
0
0
0
0
0
0
0
0
0
0
0
0
0
0
0
0
0
0

1.0
1.0
1.0
1.0
1.0
1.0
1.0
1.0
1.0
1.0
1.0
1.0
1.0
1.0
1.0
1.0
1.0
1.0
1.0
1.0
1.0
1.0
1.0
1.0
1.0
1.0
1.0
1.0
1.0
1.0


4
0
0
0
0
0
0
0
2
0
1
1
0
0
0
0
0
0
0
0
0
0
0
0
0
0
0
0
0
0


4
0
0
0
0
0
0
0
2
0
1
1
0
0
0
0
0
0
0
0
0
0
0
0
0
0
0
0
0
0

1.0
1.0
1.0
1.0
1.0
1.0
1.0
1.0
1.0
1.0
1.0
1.0
1.0
1.0
1.0
1.0
1.0
1.0
1.0
1.0
1.0
1.0
1.0
1.0
1.0
1.0
1.0
1.0
1.0
1.0


11
0
0
0
0
0
0
4
3
3
1
0
0
0
0
0
0
0
0
0
0
0
0
0
0
0
0
0
0
0

1.0
1.0
1.0
1.0
1.0
1.0
1.0
1.0
1.0
1.0
1.0
1.0
1.0
1.0
1.0
1.0
1.0
1.0
1.0
1.0
1.0
1.0
1.0
1.0
1.0
1.0
1.0
1.0
1.0
1.0


7
1
0
0
0
0
0
0
2
0
2
2
0
0
0
0
0
0
0
0
0
0
0
0
0
0
0
0
0
0


6
1
0
0
0
0
0
0
2
0
1
2
0
0
0
0
0
0
0
0
0
0
0
0
0
0
0
0
0
0

1.0
1.0
1.0
1.0
1.0
1.0
1.0
1.0
1.0
1.0
1.0
1.0
1.0
1.0
1.0
1.0
1.0
1.0
1.0
1.0
1.0
1.0
1.0
1.0
1.0
1.0
1.0
1.0
1.0
1.0


1
0
0
0
0
0
0
0
0
0
1
0
0
0
0
0
0
0
0
0
0
0
0
0
0
0
0
0
0
0

0.99
0.99
0.99
0.99
0.99
0.99
0.99
0.99
0.99
0.99
0.99
0.99
0.99
0.99
0.99
0.99
0.99
0.99
0.99
0.99
0.99
0.99
0.99
0.99
0.99
0.99
0.99
0.99
0.99
0.99


1
0
0
0
0
0
0
0
0
1
0
0
0
0
0
0
0
0
0
0
0
0
0
0
0
0
0
0
0
0


1
0
0
0
0
0
0
0
0
1
0
0
0
0
0
0
0
0
0
0
0
0
0
0
0
0
0
0
0
0

1.0
1.0
1.0
1.0
1.0
1.0
1.0
1.0
1.0
1.0
1.0
1.0
1.0
1.0
1.0
1.0
1.0
1.0
1.0
1.0
1.0
1.0
1.0
1.0
1.0
1.0
1.0
1.0
1.0
1.0


1
0
0
0
0
0
0
1
0
0
0
0
0
0
0
0
0
0
0
0
0
0
0
0
0
0
0
0
0
0


1
0
0
0
0
0
0
1
0
0
0
0
0
0
0
0
0
0
0
0
0
0
0
0
0
0
0
0
0
0

1.0
1.0
1.0
1.0
1.0
1.0
1.0
1.0
1.0
1.0
1.0
1.0
1.0
1.0
1.0
1.0
1.0
1.0
1.0
1.0
1.0
1.0
1.0
1.0
1.0
1.0
1.0
1.0
1.0
1.0


2
0
0
0
0
0
0
0
0
1
1
0
0
0
0
0
0
0
0
0
0
0
0
0
0
0
0
0
0
0


2
0
0
0
0
0
0
0
0
1
1
0
0
0
0
0
0
0
0
0
0
0
0
0
0
0
0
0
0
0

1.0
1.0
1.0
1.0
1.0
1.0
1.0
1.0
1.0
1.0
1.0
1.0
1.0
1.0
1.0
1.0
1.0
1.0
1.0
1.0
1.0
1.0
1.0
1.0
1.0
1.0
1.0
1.0
1.0
1.0


1
0
0
0
0
0
0
0
0
0
1
0
0
0
0
0
0
0
0
0
0
0
0
0
0
0
0
0
0
0


1
0
0
0
0
0
0
0
0
0
1
0
0
0
0
0
0
0
0
0
0
0
0
0
0
0
0
0
0
0

0.89
0.89
0.89
0.89
0.89
0.89
0.89
0.89
0.89
0.89
0.89
0.89
0.89
0.89
0.89
0.89
0.89
0.89
0.89
0.89
0.89
0.89
0.89
0.89
0.89
0.89
0.89
0.89
0.89
0.89


1
0
0
0
0
0
0
0
0
0
0
0
0
0
1
0
0
0
0
0
0
0
0
0
0
0
0
0
0
0


1
0
0
0
0
0
0
0
0
0
0
0
0
0
1
0
0
0
0
0
0
0
0
0
0
0
0
0
0
0

1.0
1.0
1.0
1.0
1.0
1.0
1.0
1.0
1.0
1.0
1.0
1.0
1.0
1.0
1.0
1.0
1.0
1.0
1.0
1.0
1.0
1.0
1.0
1.0
1.0
1.0
1.0
1.0
1.0
1.0


2
0
0
0
0
0
0
0
0
0
0
0
0
0
0
0
0
0
0
0
0
0
0
0
0
0
1
0
1
0


2
0
0
0
0
0
0
0
0
0
0
0
0
0
0
0
0
0
0
0
0
0
0
0
0
0
1
0
1
0

1.0
1.0
1.0
1.0
1.0
1.0
1.0
1.0
1.0
1.0
1.0
1.0
1.0
1.0
1.0
1.0
1.0
1.0
1.0
1.0
1.0
1.0
1.0
1.0
1.0
1.0
1.0
1.0
1.0
1.0


1
0
0
0
0
0
0
0
0
0
0
0
0
0
0
0
0
0
0
0
0
0
0
0
1
0
0
0
0
0


1
0
0
0
0
0
0
0
0
0
0
0
0
0
0
0
0
0
0
0
0
0
0
0
1
0
0
0
0
0

1.0
1.0
1.0
1.0
1.0
1.0
1.0
1.0
1.0
1.0
1.0
1.0
1.0
1.0
1.0
1.0
1.0
1.0
1.0
1.0
1.0
1.0
1.0
1.0
1.0
1.0
1.0
1.0
1.0
1.0


377
23
14
9
8
10
14
9
17
33
33
20
1
8
7
2
2
9
7
16
39
0
3
1
19
13
11
11
13
25

0.71
0.71
0.71
0.71
0.71
0.71
0.71
0.71
0.71
0.71
0.71
0.71
0.71
0.71
0.71
0.71
0.71
0.71
0.71
0.71
0.71
0.71
0.71
0.71
0.71
0.71
0.71
0.71
0.71
0.71


13
0
0
0
0
0
0
5
2
3
3
0
0
0
0
0
0
0
0
0
0
0
0
0
0
0
0
0
0
0


13
0
0
0
0
0
0
5
2
3
3
0
0
0
0
0
0
0
0
0
0
0
0
0
0
0
0
0
0
0

1.0
1.0
1.0
1.0
1.0
1.0
1.0
1.0
1.0
1.0
1.0
1.0
1.0
1.0
1.0
1.0
1.0
1.0
1.0
1.0
1.0
1.0
1.0
1.0
1.0
1.0
1.0
1.0
1.0
1.0


27
4
2
1
1
3
1
0
5
5
4
1
0
0
0
0
0
0
0
0
0
0
0
0
0
0
0
0
0
0

1.0
1.0
1.0
1.0
1.0
1.0
1.0
1.0
1.0
1.0
1.0
1.0
1.0
1.0
1.0
1.0
1.0
1.0
1.0
1.0
1.0
1.0
1.0
1.0
1.0
1.0
1.0
1.0
1.0
1.0


60
2
0
0
0
0
0
4
6
7
7
0
1
0
0
0
0
0
0
2
27
0
3
1
0
0
0
0
0
0

1.0
1.0
1.0
1.0
1.0
1.0
1.0
1.0
1.0
1.0
1.0
1.0
1.0
1.0
1.0
1.0
1.0
1.0
1.0
1.0
1.0
1.0
1.0
1.0
1.0
1.0
1.0
1.0
1.0
1.0


57
2
0
0
0
0
0
4
6
7
7
0
1
0
0
0
0
0
0
2
24
0
3
1
0
0
0
0
0
0

1.0
1.0
1.0
1.0
1.0
1.0
1.0
1.0
1.0
1.0
1.0
1.0
1.0
1.0
1.0
1.0
1.0
1.0
1.0
1.0
1.0
1.0
1.0
1.0
1.0
1.0
1.0
1.0
1.0
1.0


195
7
8
6
6
6
12
0
2
8
13
9
0
8
3
2
2
2
7
14
12
0
0
0
2
13
10
8
10
25

1.0
1.0
1.0
1.0
1.0
1.0
1.0
1.0
1.0
1.0
1.0
1.0
1.0
1.0
1.0
1.0
1.0
1.0
1.0
1.0
1.0
1.0
1.0
1.0
1.0
1.0
1.0
1.0
1.0
1.0


20
2
3
1
0
1
1
0
2
3
3
3
0
0
0
0
0
0
0
0
0
0
0
0
0
0
0
0
0
1

1.0
1.0
1.0
1.0
1.0
1.0
1.0
1.0
1.0
1.0
1.0
1.0
1.0
1.0
1.0
1.0
1.0
1.0
1.0
1.0
1.0
1.0
1.0
1.0
1.0
1.0
1.0
1.0
1.0
1.0


131
3
3
3
5
5
7
0
0
3
10
5
0
6
1
0
0
0
7
14
9
0
0
0
0
5
7
6
8
24

1.0
1.0
1.0
1.0
1.0
1.0
1.0
1.0
1.0
1.0
1.0
1.0
1.0
1.0
1.0
1.0
1.0
1.0
1.0
1.0
1.0
1.0
1.0
1.0
1.0
1.0
1.0
1.0
1.0
1.0


14
2
1
1
0
0
2
0
0
0
0
1
0
0
0
0
0
0
0
0
0
0
0
0
0
3
0
2
2
0

0.99
0.99
0.99
0.99
0.99
0.99
0.99
0.99
0.99
0.99
0.99
0.99
0.99
0.99
0.99
0.99
0.99
0.99
0.99
0.99
0.99
0.99
0.99
0.99
0.99
0.99
0.99
0.99
0.99
0.99


23
0
1
1
1
0
1
0
0
0
0
0
0
2
2
2
2
2
0
0
2
0
0
0
2
3
2
0
0
0

1.0
1.0
1.0
1.0
1.0
1.0
1.0
1.0
1.0
1.0
1.0
1.0
1.0
1.0
1.0
1.0
1.0
1.0
1.0
1.0
1.0
1.0
1.0
1.0
1.0
1.0
1.0
1.0
1.0
1.0


2
0
0
0
0
0
0
0
0
2
0
0
0
0
0
0
0
0
0
0
0
0
0
0
0
0
0
0
0
0

1.0
1.0
1.0
1.0
1.0
1.0
1.0
1.0
1.0
1.0
1.0
1.0
1.0
1.0
1.0
1.0
1.0
1.0
1.0
1.0
1.0
1.0
1.0
1.0
1.0
1.0
1.0
1.0
1.0
1.0


2
0
0
0
0
0
0
0
0
0
0
0
0
0
0
0
0
0
0
0
0
0
0
0
0
2
0
0
0
0

0.97
0.97
0.97
0.97
0.97
0.97
0.97
0.97
0.97
0.97
0.97
0.97
0.97
0.97
0.97
0.97
0.97
0.97
0.97
0.97
0.97
0.97
0.97
0.97
0.97
0.97
0.97
0.97
0.97
0.97


30
4
2
0
1
0
0
0
0
4
0
4
0
0
0
0
0
6
0
0
0
0
0
0
9
0
0
0
0
0

1.0
1.0
1.0
1.0
1.0
1.0
1.0
1.0
1.0
1.0
1.0
1.0
1.0
1.0
1.0
1.0
1.0
1.0
1.0
1.0
1.0
1.0
1.0
1.0
1.0
1.0
1.0
1.0
1.0
1.0


8
2
1
0
1
0
0
0
0
4
0
0
0
0
0
0
0
0
0
0
0
0
0
0
0
0
0
0
0
0

1.0
1.0
1.0
1.0
1.0
1.0
1.0
1.0
1.0
1.0
1.0
1.0
1.0
1.0
1.0
1.0
1.0
1.0
1.0
1.0
1.0
1.0
1.0
1.0
1.0
1.0
1.0
1.0
1.0
1.0


1
0
0
0
0
0
0
0
0
0
0
1
0
0
0
0
0
0
0
0
0
0
0
0
0
0
0
0
0
0

0.76
0.76
0.76
0.76
0.76
0.76
0.76
0.76
0.76
0.76
0.76
0.76
0.76
0.76
0.76
0.76
0.76
0.76
0.76
0.76
0.76
0.76
0.76
0.76
0.76
0.76
0.76
0.76
0.76
0.76


1
0
0
0
0
0
0
0
0
0
0
1
0
0
0
0
0
0
0
0
0
0
0
0
0
0
0
0
0
0

1.0
1.0
1.0
1.0
1.0
1.0
1.0
1.0
1.0
1.0
1.0
1.0
1.0
1.0
1.0
1.0
1.0
1.0
1.0
1.0
1.0
1.0
1.0
1.0
1.0
1.0
1.0
1.0
1.0
1.0


4
1
1
0
0
0
0
0
0
0
0
2
0
0
0
0
0
0
0
0
0
0
0
0
0
0
0
0
0
0

1.0
1.0
1.0
1.0
1.0
1.0
1.0
1.0
1.0
1.0
1.0
1.0
1.0
1.0
1.0
1.0
1.0
1.0
1.0
1.0
1.0
1.0
1.0
1.0
1.0
1.0
1.0
1.0
1.0
1.0


6
0
0
0
0
0
0
0
0
0
0
0
0
0
0
0
0
0
0
0
0
0
0
0
6
0
0
0
0
0

0.97
0.97
0.97
0.97
0.97
0.97
0.97
0.97
0.97
0.97
0.97
0.97
0.97
0.97
0.97
0.97
0.97
0.97
0.97
0.97
0.97
0.97
0.97
0.97
0.97
0.97
0.97
0.97
0.97
0.97


21
5
0
0
0
1
0
0
1
5
5
4
0
0
0
0
0
0
0
0
0
0
0
0
0
0
0
0
0
0

1.0
1.0
1.0
1.0
1.0
1.0
1.0
1.0
1.0
1.0
1.0
1.0
1.0
1.0
1.0
1.0
1.0
1.0
1.0
1.0
1.0
1.0
1.0
1.0
1.0
1.0
1.0
1.0
1.0
1.0


2
0
0
0
0
0
0
0
1
1
0
0
0
0
0
0
0
0
0
0
0
0
0
0
0
0
0
0
0
0

0.84
0.84
0.84
0.84
0.84
0.84
0.84
0.84
0.84
0.84
0.84
0.84
0.84
0.84
0.84
0.84
0.84
0.84
0.84
0.84
0.84
0.84
0.84
0.84
0.84
0.84
0.84
0.84
0.84
0.84


1
0
0
0
0
0
0
0
1
0
0
0
0
0
0
0
0
0
0
0
0
0
0
0
0
0
0
0
0
0


1
0
0
0
0
0
0
0
1
0
0
0
0
0
0
0
0
0
0
0
0
0
0
0
0
0
0
0
0
0

1.0
1.0
1.0
1.0
1.0
1.0
1.0
1.0
1.0
1.0
1.0
1.0
1.0
1.0
1.0
1.0
1.0
1.0
1.0
1.0
1.0
1.0
1.0
1.0
1.0
1.0
1.0
1.0
1.0
1.0


27
1
2
2
0
0
1
0
0
0
0
2
0
0
4
0
0
0
0
0
0
0
0
0
8
0
1
3
3
0

1.0
1.0
1.0
1.0
1.0
1.0
1.0
1.0
1.0
1.0
1.0
1.0
1.0
1.0
1.0
1.0
1.0
1.0
1.0
1.0
1.0
1.0
1.0
1.0
1.0
1.0
1.0
1.0
1.0
1.0


9
1
0
1
0
0
0
0
0
0
0
0
0
0
0
0
0
0
0
0
0
0
0
0
4
0
0
0
3
0

0.96
0.96
0.96
0.96
0.96
0.96
0.96
0.96
0.96
0.96
0.96
0.96
0.96
0.96
0.96
0.96
0.96
0.96
0.96
0.96
0.96
0.96
0.96
0.96
0.96
0.96
0.96
0.96
0.96
0.96


2
0
1
0
0
0
0
0
0
0
0
1
0
0
0
0
0
0
0
0
0
0
0
0
0
0
0
0
0
0

1.0
1.0
1.0
1.0
1.0
1.0
1.0
1.0
1.0
1.0
1.0
1.0
1.0
1.0
1.0
1.0
1.0
1.0
1.0
1.0
1.0
1.0
1.0
1.0
1.0
1.0
1.0
1.0
1.0
1.0


8
0
0
0
0
0
0
0
0
0
0
0
0
0
3
0
0
0
0
0
0
0
0
0
3
0
0
2
0
0

0.91
0.91
0.91
0.91
0.91
0.91
0.91
0.91
0.91
0.91
0.91
0.91
0.91
0.91
0.91
0.91
0.91
0.91
0.91
0.91
0.91
0.91
0.91
0.91
0.91
0.91
0.91
0.91
0.91
0.91


1
0
0
0
0
0
0
0
0
0
0
0
0
0
0
0
0
0
0
0
0
0
0
0
0
0
1
0
0
0

0.77
0.77
0.77
0.77
0.77
0.77
0.77
0.77
0.77
0.77
0.77
0.77
0.77
0.77
0.77
0.77
0.77
0.77
0.77
0.77
0.77
0.77
0.77
0.77
0.77
0.77
0.77
0.77
0.77
0.77


1
0
0
0
0
0
0
0
0
1
0
0
0
0
0
0
0
0
0
0
0
0
0
0
0
0
0
0
0
0


1
0
0
0
0
0
0
0
0
1
0
0
0
0
0
0
0
0
0
0
0
0
0
0
0
0
0
0
0
0

0.99
0.99
0.99
0.99
0.99
0.99
0.99
0.99
0.99
0.99
0.99
0.99
0.99
0.99
0.99
0.99
0.99
0.99
0.99
0.99
0.99
0.99
0.99
0.99
0.99
0.99
0.99
0.99
0.99
0.99


1
0
0
0
0
0
0
0
0
0
1
0
0
0
0
0
0
0
0
0
0
0
0
0
0
0
0
0
0
0

1.0
1.0
1.0
1.0
1.0
1.0
1.0
1.0
1.0
1.0
1.0
1.0
1.0
1.0
1.0
1.0
1.0
1.0
1.0
1.0
1.0
1.0
1.0
1.0
1.0
1.0
1.0
1.0
1.0
1.0


17
5
4
0
0
0
0
0
0
0
4
4
0
0
0
0
0
0
0
0
0
0
0
0
0
0
0
0
0
0


17
5
4
0
0
0
0
0
0
0
4
4
0
0
0
0
0
0
0
0
0
0
0
0
0
0
0
0
0
0

0.93
0.93
0.93
0.93
0.93
0.93
0.93
0.93
0.93
0.93
0.93
0.93
0.93
0.93
0.93
0.93
0.93
0.93
0.93
0.93
0.93
0.93
0.93
0.93
0.93
0.93
0.93
0.93
0.93
0.93


20
1
1
0
0
0
0
2
3
3
1
2
0
0
0
0
0
1
1
0
0
0
4
0
0
0
1
0
0
0

0.76
0.76
0.76
0.76
0.76
0.76
0.76
0.76
0.76
0.76
0.76
0.76
0.76
0.76
0.76
0.76
0.76
0.76
0.76
0.76
0.76
0.76
0.76
0.76
0.76
0.76
0.76
0.76
0.76
0.76


8
0
0
0
0
0
0
2
2
0
0
0
0
0
0
0
0
0
0
0
0
0
4
0
0
0
0
0
0
0


8
0
0
0
0
0
0
2
2
0
0
0
0
0
0
0
0
0
0
0
0
0
4
0
0
0
0
0
0
0

0.99
0.99
0.99
0.99
0.99
0.99
0.99
0.99
0.99
0.99
0.99
0.99
0.99
0.99
0.99
0.99
0.99
0.99
0.99
0.99
0.99
0.99
0.99
0.99
0.99
0.99
0.99
0.99
0.99
0.99


1
0
0
0
0
0
0
0
0
0
0
1
0
0
0
0
0
0
0
0
0
0
0
0
0
0
0
0
0
0


1
0
0
0
0
0
0
0
0
0
0
1
0
0
0
0
0
0
0
0
0
0
0
0
0
0
0
0
0
0

1.0
1.0
1.0
1.0
1.0
1.0
1.0
1.0
1.0
1.0
1.0
1.0
1.0
1.0
1.0
1.0
1.0
1.0
1.0
1.0
1.0
1.0
1.0
1.0
1.0
1.0
1.0
1.0
1.0
1.0


5
0
0
0
0
0
0
0
1
3
0
0
0
0
0
0
0
0
0
0
0
0
0
0
0
0
1
0
0
0

0.95
0.95
0.95
0.95
0.95
0.95
0.95
0.95
0.95
0.95
0.95
0.95
0.95
0.95
0.95
0.95
0.95
0.95
0.95
0.95
0.95
0.95
0.95
0.95
0.95
0.95
0.95
0.95
0.95
0.95


95
13
2
0
0
0
0
8
10
22
16
3
1
0
2
0
0
4
3
0
0
0
1
1
0
3
0
0
3
3

0.87
0.87
0.87
0.87
0.87
0.87
0.87
0.87
0.87
0.87
0.87
0.87
0.87
0.87
0.87
0.87
0.87
0.87
0.87
0.87
0.87
0.87
0.87
0.87
0.87
0.87
0.87
0.87
0.87
0.87


32
2
0
0
0
0
0
3
3
8
2
0
1
0
0
0
0
3
3
0
0
0
0
1
0
3
0
0
3
0


31
2
0
0
0
0
0
3
3
7
2
0
1
0
0
0
0
3
3
0
0
0
0
1
0
3
0
0
3
0

1.0
1.0
1.0
1.0
1.0
1.0
1.0
1.0
1.0
1.0
1.0
1.0
1.0
1.0
1.0
1.0
1.0
1.0
1.0
1.0
1.0
1.0
1.0
1.0
1.0
1.0
1.0
1.0
1.0
1.0


1
0
0
0
0
0
0
0
0
1
0
0
0
0
0
0
0
0
0
0
0
0
0
0
0
0
0
0
0
0

0.78
0.78
0.78
0.78
0.78
0.78
0.78
0.78
0.78
0.78
0.78
0.78
0.78
0.78
0.78
0.78
0.78
0.78
0.78
0.78
0.78
0.78
0.78
0.78
0.78
0.78
0.78
0.78
0.78
0.78


39
5
1
0
0
0
0
2
5
10
8
2
0
0
2
0
0
0
0
0
0
0
1
0
0
0
0
0
0
3

0.88
0.88
0.88
0.88
0.88
0.88
0.88
0.88
0.88
0.88
0.88
0.88
0.88
0.88
0.88
0.88
0.88
0.88
0.88
0.88
0.88
0.88
0.88
0.88
0.88
0.88
0.88
0.88
0.88
0.88


21
4
1
0
0
0
0
0
4
5
5
2
0
0
0
0
0
0
0
0
0
0
0
0
0
0
0
0
0
0

1.0
1.0
1.0
1.0
1.0
1.0
1.0
1.0
1.0
1.0
1.0
1.0
1.0
1.0
1.0
1.0
1.0
1.0
1.0
1.0
1.0
1.0
1.0
1.0
1.0
1.0
1.0
1.0
1.0
1.0


12
1
0
0
0
0
0
2
1
4
3
0
0
0
0
0
0
0
0
0
0
0
1
0
0
0
0
0
0
0

1.0
1.0
1.0
1.0
1.0
1.0
1.0
1.0
1.0
1.0
1.0
1.0
1.0
1.0
1.0
1.0
1.0
1.0
1.0
1.0
1.0
1.0
1.0
1.0
1.0
1.0
1.0
1.0
1.0
1.0


1
0
0
0
0
0
0
0
0
1
0
0
0
0
0
0
0
0
0
0
0
0
0
0
0
0
0
0
0
0

1.0
1.0
1.0
1.0
1.0
1.0
1.0
1.0
1.0
1.0
1.0
1.0
1.0
1.0
1.0
1.0
1.0
1.0
1.0
1.0
1.0
1.0
1.0
1.0
1.0
1.0
1.0
1.0
1.0
1.0


3
0
0
0
0
0
0
0
0
0
0
0
0
0
0
0
0
0
0
0
0
0
0
0
0
0
0
0
0
3

1.0
1.0
1.0
1.0
1.0
1.0
1.0
1.0
1.0
1.0
1.0
1.0
1.0
1.0
1.0
1.0
1.0
1.0
1.0
1.0
1.0
1.0
1.0
1.0
1.0
1.0
1.0
1.0
1.0
1.0


13
4
0
0
0
0
0
3
0
2
4
0
0
0
0
0
0
0
0
0
0
0
0
0
0
0
0
0
0
0


13
4
0
0
0
0
0
3
0
2
4
0
0
0
0
0
0
0
0
0
0
0
0
0
0
0
0
0
0
0

1.0
1.0
1.0
1.0
1.0
1.0
1.0
1.0
1.0
1.0
1.0
1.0
1.0
1.0
1.0
1.0
1.0
1.0
1.0
1.0
1.0
1.0
1.0
1.0
1.0
1.0
1.0
1.0
1.0
1.0


1
0
0
0
0
0
0
0
0
0
0
0
0
0
0
0
0
1
0
0
0
0
0
0
0
0
0
0
0
0

1.0
1.0
1.0
1.0
1.0
1.0
1.0
1.0
1.0
1.0
1.0
1.0
1.0
1.0
1.0
1.0
1.0
1.0
1.0
1.0
1.0
1.0
1.0
1.0
1.0
1.0
1.0
1.0
1.0
1.0


116
0
0
0
0
0
0
13
15
18
9
0
0
0
0
0
0
5
13
18
0
25
0
0
0
0
0
0
0
0


116
0
0
0
0
0
0
13
15
18
9
0
0
0
0
0
0
5
13
18
0
25
0
0
0
0
0
0
0
0

0.78
0.78
0.78
0.78
0.78
0.78
0.78
0.78
0.78
0.78
0.78
0.78
0.78
0.78
0.78
0.78
0.78
0.78
0.78
0.78
0.78
0.78
0.78
0.78
0.78
0.78
0.78
0.78
0.78
0.78


80
0
0
0
0
0
0
6
7
9
7
0
0
0
0
0
0
5
12
17
0
17
0
0
0
0
0
0
0
0

1.0
1.0
1.0
1.0
1.0
1.0
1.0
1.0
1.0
1.0
1.0
1.0
1.0
1.0
1.0
1.0
1.0
1.0
1.0
1.0
1.0
1.0
1.0
1.0
1.0
1.0
1.0
1.0
1.0
1.0


34
0
0
0
0
0
0
7
8
9
2
0
0
0
0
0
0
0
0
0
0
8
0
0
0
0
0
0
0
0

1.0
1.0
1.0
1.0
1.0
1.0
1.0
1.0
1.0
1.0
1.0
1.0
1.0
1.0
1.0
1.0
1.0
1.0
1.0
1.0
1.0
1.0
1.0
1.0
1.0
1.0
1.0
1.0
1.0
1.0


113
6
6
1
1
1
0
4
7
8
4
8
6
0
0
0
0
9
22
6
4
5
7
6
0
0
0
0
0
2

0.88
0.88
0.88
0.88
0.88
0.88
0.88
0.88
0.88
0.88
0.88
0.88
0.88
0.88
0.88
0.88
0.88
0.88
0.88
0.88
0.88
0.88
0.88
0.88
0.88
0.88
0.88
0.88
0.88
0.88


63
0
0
0
0
0
0
4
5
5
0
0
6
0
0
0
0
0
19
6
0
5
7
6
0
0
0
0
0
0


63
0
0
0
0
0
0
4
5
5
0
0
6
0
0
0
0
0
19
6
0
5
7
6
0
0
0
0
0
0

1.0
1.0
1.0
1.0
1.0
1.0
1.0
1.0
1.0
1.0
1.0
1.0
1.0
1.0
1.0
1.0
1.0
1.0
1.0
1.0
1.0
1.0
1.0
1.0
1.0
1.0
1.0
1.0
1.0
1.0


44
6
6
1
0
0
0
0
0
1
4
8
0
0
0
0
0
9
3
0
4
0
0
0
0
0
0
0
0
2


44
6
6
1
0
0
0
0
0
1
4
8
0
0
0
0
0
9
3
0
4
0
0
0
0
0
0
0
0
2

1.0
1.0
1.0
1.0
1.0
1.0
1.0
1.0
1.0
1.0
1.0
1.0
1.0
1.0
1.0
1.0
1.0
1.0
1.0
1.0
1.0
1.0
1.0
1.0
1.0
1.0
1.0
1.0
1.0
1.0


4
0
0
0
0
0
0
0
2
2
0
0
0
0
0
0
0
0
0
0
0
0
0
0
0
0
0
0
0
0


4
0
0
0
0
0
0
0
2
2
0
0
0
0
0
0
0
0
0
0
0
0
0
0
0
0
0
0
0
0

1.0
1.0
1.0
1.0
1.0
1.0
1.0
1.0
1.0
1.0
1.0
1.0
1.0
1.0
1.0
1.0
1.0
1.0
1.0
1.0
1.0
1.0
1.0
1.0
1.0
1.0
1.0
1.0
1.0
1.0


400
9
3
0
1
0
0
12
14
11
8
10
2
16
9
10
8
18
28
20
8
0
1
2
78
10
16
23
17
66

1.0
1.0
1.0
1.0
1.0
1.0
1.0
1.0
1.0
1.0
1.0
1.0
1.0
1.0
1.0
1.0
1.0
1.0
1.0
1.0
1.0
1.0
1.0
1.0
1.0
1.0
1.0
1.0
1.0
1.0


29
3
0
0
0
0
0
2
3
3
1
2
1
0
0
0
0
3
5
1
0
0
0
1
0
1
0
1
1
1


29
3
0
0
0
0
0
2
3
3
1
2
1
0
0
0
0
3
5
1
0
0
0
1
0
1
0
1
1
1

1.0
1.0
1.0
1.0
1.0
1.0
1.0
1.0
1.0
1.0
1.0
1.0
1.0
1.0
1.0
1.0
1.0
1.0
1.0
1.0
1.0
1.0
1.0
1.0
1.0
1.0
1.0
1.0
1.0
1.0


10
1
0
0
0
0
0
2
2
2
1
2
0
0
0
0
0
0
0
0
0
0
0
0
0
0
0
0
0
0

1.0
1.0
1.0
1.0
1.0
1.0
1.0
1.0
1.0
1.0
1.0
1.0
1.0
1.0
1.0
1.0
1.0
1.0
1.0
1.0
1.0
1.0
1.0
1.0
1.0
1.0
1.0
1.0
1.0
1.0


6
1
0
0
0
0
0
1
1
1
1
1
0
0
0
0
0
0
0
0
0
0
0
0
0
0
0
0
0
0

1.0
1.0
1.0
1.0
1.0
1.0
1.0
1.0
1.0
1.0
1.0
1.0
1.0
1.0
1.0
1.0
1.0
1.0
1.0
1.0
1.0
1.0
1.0
1.0
1.0
1.0
1.0
1.0
1.0
1.0


4
1
1
0
0
0
0
0
0
0
1
1
0
0
0
0
0
0
0
0
0
0
0
0
0
0
0
0
0
0


4
1
1
0
0
0
0
0
0
0
1
1
0
0
0
0
0
0
0
0
0
0
0
0
0
0
0
0
0
0

1.0
1.0
1.0
1.0
1.0
1.0
1.0
1.0
1.0
1.0
1.0
1.0
1.0
1.0
1.0
1.0
1.0
1.0
1.0
1.0
1.0
1.0
1.0
1.0
1.0
1.0
1.0
1.0
1.0
1.0


1
0
0
0
0
0
0
0
1
0
0
0
0
0
0
0
0
0
0
0
0
0
0
0
0
0
0
0
0
0


1
0
0
0
0
0
0
0
1
0
0
0
0
0
0
0
0
0
0
0
0
0
0
0
0
0
0
0
0
0

0.92
0.92
0.92
0.92
0.92
0.92
0.92
0.92
0.92
0.92
0.92
0.92
0.92
0.92
0.92
0.92
0.92
0.92
0.92
0.92
0.92
0.92
0.92
0.92
0.92
0.92
0.92
0.92
0.92
0.92


4
0
0
0
0
0
0
2
0
0
0
0
0
0
1
0
0
0
1
0
0
0
0
0
0
0
0
0
0
0


4
0
0
0
0
0
0
2
0
0
0
0
0
0
1
0
0
0
1
0
0
0
0
0
0
0
0
0
0
0

1.0
1.0
1.0
1.0
1.0
1.0
1.0
1.0
1.0
1.0
1.0
1.0
1.0
1.0
1.0
1.0
1.0
1.0
1.0
1.0
1.0
1.0
1.0
1.0
1.0
1.0
1.0
1.0
1.0
1.0


1
0
0
0
0
0
0
0
1
0
0
0
0
0
0
0
0
0
0
0
0
0
0
0
0
0
0
0
0
0

0.94
0.94
0.94
0.94
0.94
0.94
0.94
0.94
0.94
0.94
0.94
0.94
0.94
0.94
0.94
0.94
0.94
0.94
0.94
0.94
0.94
0.94
0.94
0.94
0.94
0.94
0.94
0.94
0.94
0.94


2
0
0
0
0
0
0
0
0
0
0
0
0
0
0
0
0
0
2
0
0
0
0
0
0
0
0
0
0
0

0.93
0.93
0.93
0.93
0.93
0.93
0.93
0.93
0.93
0.93
0.93
0.93
0.93
0.93
0.93
0.93
0.93
0.93
0.93
0.93
0.93
0.93
0.93
0.93
0.93
0.93
0.93
0.93
0.93
0.93


2
0
0
0
0
0
0
0
0
0
0
0
0
0
0
0
0
1
0
1
0
0
0
0
0
0
0
0
0
0

0.95
0.95
0.95
0.95
0.95
0.95
0.95
0.95
0.95
0.95
0.95
0.95
0.95
0.95
0.95
0.95
0.95
0.95
0.95
0.95
0.95
0.95
0.95
0.95
0.95
0.95
0.95
0.95
0.95
0.95


7
0
0
0
0
0
0
0
0
0
0
0
0
0
0
1
0
0
0
0
0
0
0
0
0
0
0
0
1
5

0.97
0.97
0.97
0.97
0.97
0.97
0.97
0.97
0.97
0.97
0.97
0.97
0.97
0.97
0.97
0.97
0.97
0.97
0.97
0.97
0.97
0.97
0.97
0.97
0.97
0.97
0.97
0.97
0.97
0.97


2
0
0
0
0
0
0
0
0
0
0
0
0
0
0
0
0
0
1
1
0
0
0
0
0
0
0
0
0
0

0.91
0.91
0.91
0.91
0.91
0.91
0.91
0.91
0.91
0.91
0.91
0.91
0.91
0.91
0.91
0.91
0.91
0.91
0.91
0.91
0.91
0.91
0.91
0.91
0.91
0.91
0.91
0.91
0.91
0.91


3
0
0
0
0
0
0
0
0
0
0
0
0
0
0
0
0
1
0
0
0
0
0
0
0
1
0
0
0
1


3
0
0
0
0
0
0
0
0
0
0
0
0
0
0
0
0
1
0
0
0
0
0
0
0
1
0
0
0
1

0.98
0.98
0.98
0.98
0.98
0.98
0.98
0.98
0.98
0.98
0.98
0.98
0.98
0.98
0.98
0.98
0.98
0.98
0.98
0.98
0.98
0.98
0.98
0.98
0.98
0.98
0.98
0.98
0.98
0.98


8
0
0
0
0
0
0
0
0
0
0
0
0
1
0
0
0
0
0
0
0
0
0
0
3
1
0
1
0
2

1.0
1.0
1.0
1.0
1.0
1.0
1.0
1.0
1.0
1.0
1.0
1.0
1.0
1.0
1.0
1.0
1.0
1.0
1.0
1.0
1.0
1.0
1.0
1.0
1.0
1.0
1.0
1.0
1.0
1.0


6
0
0
0
0
0
0
0
0
0
0
0
0
0
0
1
0
0
1
0
0
0
0
0
0
1
1
1
1
0


6
0
0
0
0
0
0
0
0
0
0
0
0
0
0
1
0
0
1
0
0
0
0
0
0
1
1
1
1
0

1.0
1.0
1.0
1.0
1.0
1.0
1.0
1.0
1.0
1.0
1.0
1.0
1.0
1.0
1.0
1.0
1.0
1.0
1.0
1.0
1.0
1.0
1.0
1.0
1.0
1.0
1.0
1.0
1.0
1.0


1
0
0
0
0
0
0
0
0
0
0
0
0
0
0
0
0
0
0
0
0
0
0
0
0
0
0
0
0
1


1
0
0
0
0
0
0
0
0
0
0
0
0
0
0
0
0
0
0
0
0
0
0
0
0
0
0
0
0
1

0.99
0.99
0.99
0.99
0.99
0.99
0.99
0.99
0.99
0.99
0.99
0.99
0.99
0.99
0.99
0.99
0.99
0.99
0.99
0.99
0.99
0.99
0.99
0.99
0.99
0.99
0.99
0.99
0.99
0.99


4
0
0
0
0
0
0
0
0
0
0
0
0
0
0
0
0
0
0
0
0
0
0
0
0
1
1
1
0
1

1.0
1.0
1.0
1.0
1.0
1.0
1.0
1.0
1.0
1.0
1.0
1.0
1.0
1.0
1.0
1.0
1.0
1.0
1.0
1.0
1.0
1.0
1.0
1.0
1.0
1.0
1.0
1.0
1.0
1.0


2
0
0
0
0
0
0
0
0
0
0
0
0
0
0
0
0
1
1
0
0
0
0
0
0
0
0
0
0
0

0.75
0.75
0.75
0.75
0.75
0.75
0.75
0.75
0.75
0.75
0.75
0.75
0.75
0.75
0.75
0.75
0.75
0.75
0.75
0.75
0.75
0.75
0.75
0.75
0.75
0.75
0.75
0.75
0.75
0.75


1
0
0
0
0
0
0
0
0
0
0
0
0
0
0
0
0
1
0
0
0
0
0
0
0
0
0
0
0
0

0.71
0.71
0.71
0.71
0.71
0.71
0.71
0.71
0.71
0.71
0.71
0.71
0.71
0.71
0.71
0.71
0.71
0.71
0.71
0.71
0.71
0.71
0.71
0.71
0.71
0.71
0.71
0.71
0.71
0.71


2
0
0
0
0
0
0
0
0
0
0
0
0
0
0
0
0
0
0
1
0
0
0
0
0
0
0
0
0
1


1
0
0
0
0
0
0
0
0
0
0
0
0
0
0
0
0
0
0
1
0
0
0
0
0
0
0
0
0
0

1.0
1.0
1.0
1.0
1.0
1.0
1.0
1.0
1.0
1.0
1.0
1.0
1.0
1.0
1.0
1.0
1.0
1.0
1.0
1.0
1.0
1.0
1.0
1.0
1.0
1.0
1.0
1.0
1.0
1.0


1
0
0
0
0
0
0
0
0
0
0
0
0
0
0
0
0
0
0
0
0
0
0
0
0
0
0
0
0
1

1.0
1.0
1.0
1.0
1.0
1.0
1.0
1.0
1.0
1.0
1.0
1.0
1.0
1.0
1.0
1.0
1.0
1.0
1.0
1.0
1.0
1.0
1.0
1.0
1.0
1.0
1.0
1.0
1.0
1.0


4
0
0
0
0
0
0
0
0
0
0
0
0
1
0
0
0
1
0
0
0
0
0
0
0
0
0
1
0
1

0.74
0.74
0.74
0.74
0.74
0.74
0.74
0.74
0.74
0.74
0.74
0.74
0.74
0.74
0.74
0.74
0.74
0.74
0.74
0.74
0.74
0.74
0.74
0.74
0.74
0.74
0.74
0.74
0.74
0.74


2
0
0
0
0
0
0
0
0
0
0
0
0
0
0
0
1
0
0
0
0
0
0
0
0
0
0
0
0
1

0.74
0.74
0.74
0.74
0.74
0.74
0.74
0.74
0.74
0.74
0.74
0.74
0.74
0.74
0.74
0.74
0.74
0.74
0.74
0.74
0.74
0.74
0.74
0.74
0.74
0.74
0.74
0.74
0.74
0.74


1
0
0
0
0
0
0
0
0
0
0
0
0
0
0
0
0
0
0
0
0
0
0
0
0
0
0
0
0
1

0.7
0.7
0.7
0.7
0.7
0.7
0.7
0.7
0.7
0.7
0.7
0.7
0.7
0.7
0.7
0.7
0.7
0.7
0.7
0.7
0.7
0.7
0.7
0.7
0.7
0.7
0.7
0.7
0.7
0.7


1
0
0
0
0
0
0
0
0
0
0
0
0
0
0
0
0
0
0
0
0
0
0
0
0
0
0
0
0
1


1
0
0
0
0
0
0
0
0
0
0
0
0
0
0
0
0
0
0
0
0
0
0
0
0
0
0
0
0
1

1.0
1.0
1.0
1.0
1.0
1.0
1.0
1.0
1.0
1.0
1.0
1.0
1.0
1.0
1.0
1.0
1.0
1.0
1.0
1.0
1.0
1.0
1.0
1.0
1.0
1.0
1.0
1.0
1.0
1.0


1
0
0
0
0
0
0
0
0
0
0
0
0
0
0
0
0
1
0
0
0
0
0
0
0
0
0
0
0
0

0.93
0.93
0.93
0.93
0.93
0.93
0.93
0.93
0.93
0.93
0.93
0.93
0.93
0.93
0.93
0.93
0.93
0.93
0.93
0.93
0.93
0.93
0.93
0.93
0.93
0.93
0.93
0.93
0.93
0.93


1
0
1
0
0
0
0
0
0
0
0
0
0
0
0
0
0
0
0
0
0
0
0
0
0
0
0
0
0
0


1
0
1
0
0
0
0
0
0
0
0
0
0
0
0
0
0
0
0
0
0
0
0
0
0
0
0
0
0
0


1
0
1
0
0
0
0
0
0
0
0
0
0
0
0
0
0
0
0
0
0
0
0
0
0
0
0
0
0
0

1.0
1.0
1.0
1.0
1.0
1.0
1.0
1.0
1.0
1.0
1.0
1.0
1.0
1.0
1.0
1.0
1.0
1.0
1.0
1.0
1.0
1.0
1.0
1.0
1.0
1.0
1.0
1.0
1.0
1.0


113
4
6
4
4
4
6
1
4
8
10
15
0
0
19
2
0
1
0
0
0
0
0
0
1
8
6
5
2
3

0.93
0.93
0.93
0.93
0.93
0.93
0.93
0.93
0.93
0.93
0.93
0.93
0.93
0.93
0.93
0.93
0.93
0.93
0.93
0.93
0.93
0.93
0.93
0.93
0.93
0.93
0.93
0.93
0.93
0.93


31
4
1
0
0
0
0
1
4
5
7
6
0
0
3
0
0
0
0
0
0
0
0
0
0
0
0
0
0
0


14
2
1
0
0
0
0
1
2
2
2
2
0
0
2
0
0
0
0
0
0
0
0
0
0
0
0
0
0
0

1.0
1.0
1.0
1.0
1.0
1.0
1.0
1.0
1.0
1.0
1.0
1.0
1.0
1.0
1.0
1.0
1.0
1.0
1.0
1.0
1.0
1.0
1.0
1.0
1.0
1.0
1.0
1.0
1.0
1.0


10
0
0
0
0
0
0
0
2
2
3
3
0
0
0
0
0
0
0
0
0
0
0
0
0
0
0
0
0
0

1.0
1.0
1.0
1.0
1.0
1.0
1.0
1.0
1.0
1.0
1.0
1.0
1.0
1.0
1.0
1.0
1.0
1.0
1.0
1.0
1.0
1.0
1.0
1.0
1.0
1.0
1.0
1.0
1.0
1.0


1
0
0
0
0
0
0
0
0
1
0
0
0
0
0
0
0
0
0
0
0
0
0
0
0
0
0
0
0
0

1.0
1.0
1.0
1.0
1.0
1.0
1.0
1.0
1.0
1.0
1.0
1.0
1.0
1.0
1.0
1.0
1.0
1.0
1.0
1.0
1.0
1.0
1.0
1.0
1.0
1.0
1.0
1.0
1.0
1.0


2
1
0
0
0
0
0
0
0
0
0
1
0
0
0
0
0
0
0
0
0
0
0
0
0
0
0
0
0
0

1.0
1.0
1.0
1.0
1.0
1.0
1.0
1.0
1.0
1.0
1.0
1.0
1.0
1.0
1.0
1.0
1.0
1.0
1.0
1.0
1.0
1.0
1.0
1.0
1.0
1.0
1.0
1.0
1.0
1.0


3
1
0
0
0
0
0
0
0
0
1
0
0
0
1
0
0
0
0
0
0
0
0
0
0
0
0
0
0
0

1.0
1.0
1.0
1.0
1.0
1.0
1.0
1.0
1.0
1.0
1.0
1.0
1.0
1.0
1.0
1.0
1.0
1.0
1.0
1.0
1.0
1.0
1.0
1.0
1.0
1.0
1.0
1.0
1.0
1.0


1
0
0
0
0
0
0
0
0
0
1
0
0
0
0
0
0
0
0
0
0
0
0
0
0
0
0
0
0
0

1.0
1.0
1.0
1.0
1.0
1.0
1.0
1.0
1.0
1.0
1.0
1.0
1.0
1.0
1.0
1.0
1.0
1.0
1.0
1.0
1.0
1.0
1.0
1.0
1.0
1.0
1.0
1.0
1.0
1.0


34
0
4
2
2
2
5
0
0
0
0
3
0
0
0
2
0
1
0
0
0
0
0
0
0
2
3
3
2
3


34
0
4
2
2
2
5
0
0
0
0
3
0
0
0
2
0
1
0
0
0
0
0
0
0
2
3
3
2
3

1.0
1.0
1.0
1.0
1.0
1.0
1.0
1.0
1.0
1.0
1.0
1.0
1.0
1.0
1.0
1.0
1.0
1.0
1.0
1.0
1.0
1.0
1.0
1.0
1.0
1.0
1.0
1.0
1.0
1.0


12
0
0
0
0
0
0
0
0
3
3
6
0
0
0
0
0
0
0
0
0
0
0
0
0
0
0
0
0
0


12
0
0
0
0
0
0
0
0
3
3
6
0
0
0
0
0
0
0
0
0
0
0
0
0
0
0
0
0
0

1.0
1.0
1.0
1.0
1.0
1.0
1.0
1.0
1.0
1.0
1.0
1.0
1.0
1.0
1.0
1.0
1.0
1.0
1.0
1.0
1.0
1.0
1.0
1.0
1.0
1.0
1.0
1.0
1.0
1.0


21
0
1
2
2
2
1
0
0
0
0
0
0
0
1
0
0
0
0
0
0
0
0
0
1
6
3
2
0
0


9
0
1
1
1
1
1
0
0
0
0
0
0
0
0
0
0
0
0
0
0
0
0
0
1
1
1
1
0
0

1.0
1.0
1.0
1.0
1.0
1.0
1.0
1.0
1.0
1.0
1.0
1.0
1.0
1.0
1.0
1.0
1.0
1.0
1.0
1.0
1.0
1.0
1.0
1.0
1.0
1.0
1.0
1.0
1.0
1.0


7
0
0
1
1
1
0
0
0
0
0
0
0
0
1
0
0
0
0
0
0
0
0
0
0
1
1
1
0
0

1.0
1.0
1.0
1.0
1.0
1.0
1.0
1.0
1.0
1.0
1.0
1.0
1.0
1.0
1.0
1.0
1.0
1.0
1.0
1.0
1.0
1.0
1.0
1.0
1.0
1.0
1.0
1.0
1.0
1.0


5
0
0
0
0
0
0
0
0
0
0
0
0
0
0
0
0
0
0
0
0
0
0
0
0
4
1
0
0
0

1.0
1.0
1.0
1.0
1.0
1.0
1.0
1.0
1.0
1.0
1.0
1.0
1.0
1.0
1.0
1.0
1.0
1.0
1.0
1.0
1.0
1.0
1.0
1.0
1.0
1.0
1.0
1.0
1.0
1.0


14
0
0
0
0
0
0
0
0
0
0
0
0
0
14
0
0
0
0
0
0
0
0
0
0
0
0
0
0
0


14
0
0
0
0
0
0
0
0
0
0
0
0
0
14
0
0
0
0
0
0
0
0
0
0
0
0
0
0
0

1.0
1.0
1.0
1.0
1.0
1.0
1.0
1.0
1.0
1.0
1.0
1.0
1.0
1.0
1.0
1.0
1.0
1.0
1.0
1.0
1.0
1.0
1.0
1.0
1.0
1.0
1.0
1.0
1.0
1.0


92
7
6
4
2
2
2
0
0
1
9
8
0
8
10
0
0
0
3
0
5
0
0
0
9
0
1
0
0
15

1.0
1.0
1.0
1.0
1.0
1.0
1.0
1.0
1.0
1.0
1.0
1.0
1.0
1.0
1.0
1.0
1.0
1.0
1.0
1.0
1.0
1.0
1.0
1.0
1.0
1.0
1.0
1.0
1.0
1.0


36
1
1
1
1
1
1
0
0
0
1
1
0
3
5
0
0
0
0
0
0
0
0
0
6
0
0
0
0
14

0.99
0.99
0.99
0.99
0.99
0.99
0.99
0.99
0.99
0.99
0.99
0.99
0.99
0.99
0.99
0.99
0.99
0.99
0.99
0.99
0.99
0.99
0.99
0.99
0.99
0.99
0.99
0.99
0.99
0.99


19
1
1
1
1
1
1
0
0
0
1
1
0
2
5
0
0
0
0
0
0
0
0
0
4
0
0
0
0
0

0.95
0.95
0.95
0.95
0.95
0.95
0.95
0.95
0.95
0.95
0.95
0.95
0.95
0.95
0.95
0.95
0.95
0.95
0.95
0.95
0.95
0.95
0.95
0.95
0.95
0.95
0.95
0.95
0.95
0.95


2
0
0
0
0
0
0
0
0
0
0
0
0
0
0
0
0
0
0
0
0
0
0
0
2
0
0
0
0
0

1.0
1.0
1.0
1.0
1.0
1.0
1.0
1.0
1.0
1.0
1.0
1.0
1.0
1.0
1.0
1.0
1.0
1.0
1.0
1.0
1.0
1.0
1.0
1.0
1.0
1.0
1.0
1.0
1.0
1.0


13
0
0
0
0
0
0
0
0
0
0
0
0
0
0
0
0
0
0
0
0
0
0
0
0
0
0
0
0
13

1.0
1.0
1.0
1.0
1.0
1.0
1.0
1.0
1.0
1.0
1.0
1.0
1.0
1.0
1.0
1.0
1.0
1.0
1.0
1.0
1.0
1.0
1.0
1.0
1.0
1.0
1.0
1.0
1.0
1.0


43
6
5
3
1
1
1
0
0
1
8
7
0
3
2
0
0
0
2
0
2
0
0
0
1
0
0
0
0
0


9
1
1
0
1
1
1
0
0
1
1
1
0
0
0
0
0
0
0
0
0
0
0
0
1
0
0
0
0
0

0.98
0.98
0.98
0.98
0.98
0.98
0.98
0.98
0.98
0.98
0.98
0.98
0.98
0.98
0.98
0.98
0.98
0.98
0.98
0.98
0.98
0.98
0.98
0.98
0.98
0.98
0.98
0.98
0.98
0.98


13
3
1
2
0
0
0
0
0
0
4
3
0
0
0
0
0
0
0
0
0
0
0
0
0
0
0
0
0
0

1.0
1.0
1.0
1.0
1.0
1.0
1.0
1.0
1.0
1.0
1.0
1.0
1.0
1.0
1.0
1.0
1.0
1.0
1.0
1.0
1.0
1.0
1.0
1.0
1.0
1.0
1.0
1.0
1.0
1.0


15
1
2
0
0
0
0
0
0
0
3
2
0
1
2
0
0
0
2
0
2
0
0
0
0
0
0
0
0
0

1.0
1.0
1.0
1.0
1.0
1.0
1.0
1.0
1.0
1.0
1.0
1.0
1.0
1.0
1.0
1.0
1.0
1.0
1.0
1.0
1.0
1.0
1.0
1.0
1.0
1.0
1.0
1.0
1.0
1.0


6
1
1
1
0
0
0
0
0
0
0
1
0
2
0
0
0
0
0
0
0
0
0
0
0
0
0
0
0
0

1.0
1.0
1.0
1.0
1.0
1.0
1.0
1.0
1.0
1.0
1.0
1.0
1.0
1.0
1.0
1.0
1.0
1.0
1.0
1.0
1.0
1.0
1.0
1.0
1.0
1.0
1.0
1.0
1.0
1.0


2
0
0
0
0
0
0
0
0
0
0
0
0
0
0
0
0
0
0
0
0
0
0
0
2
0
0
0
0
0


2
0
0
0
0
0
0
0
0
0
0
0
0
0
0
0
0
0
0
0
0
0
0
0
2
0
0
0
0
0

1.0
1.0
1.0
1.0
1.0
1.0
1.0
1.0
1.0
1.0
1.0
1.0
1.0
1.0
1.0
1.0
1.0
1.0
1.0
1.0
1.0
1.0
1.0
1.0
1.0
1.0
1.0
1.0
1.0
1.0


1
0
0
0
0
0
0
0
0
0
0
0
0
1
0
0
0
0
0
0
0
0
0
0
0
0
0
0
0
0

1.0
1.0
1.0
1.0
1.0
1.0
1.0
1.0
1.0
1.0
1.0
1.0
1.0
1.0
1.0
1.0
1.0
1.0
1.0
1.0
1.0
1.0
1.0
1.0
1.0
1.0
1.0
1.0
1.0
1.0


94
10
1
0
0
0
0
8
10
18
13
10
0
0
2
0
0
9
3
0
2
2
2
0
0
0
1
0
0
3

0.92
0.92
0.92
0.92
0.92
0.92
0.92
0.92
0.92
0.92
0.92
0.92
0.92
0.92
0.92
0.92
0.92
0.92
0.92
0.92
0.92
0.92
0.92
0.92
0.92
0.92
0.92
0.92
0.92
0.92


14
2
0
0
0
0
0
1
1
1
1
2
0
0
0
0
0
4
2
0
0
0
0
0
0
0
0
0
0
0

0.99
0.99
0.99
0.99
0.99
0.99
0.99
0.99
0.99
0.99
0.99
0.99
0.99
0.99
0.99
0.99
0.99
0.99
0.99
0.99
0.99
0.99
0.99
0.99
0.99
0.99
0.99
0.99
0.99
0.99


7
1
0
0
0
0
0
1
1
1
1
1
0
0
0
0
0
0
1
0
0
0
0
0
0
0
0
0
0
0

1.0
1.0
1.0
1.0
1.0
1.0
1.0
1.0
1.0
1.0
1.0
1.0
1.0
1.0
1.0
1.0
1.0
1.0
1.0
1.0
1.0
1.0
1.0
1.0
1.0
1.0
1.0
1.0
1.0
1.0


2
1
0
0
0
0
0
0
0
0
0
1
0
0
0
0
0
0
0
0
0
0
0
0
0
0
0
0
0
0

1.0
1.0
1.0
1.0
1.0
1.0
1.0
1.0
1.0
1.0
1.0
1.0
1.0
1.0
1.0
1.0
1.0
1.0
1.0
1.0
1.0
1.0
1.0
1.0
1.0
1.0
1.0
1.0
1.0
1.0


31
4
1
0
0
0
0
2
5
8
7
4
0
0
0
0
0
0
0
0
0
0
0
0
0
0
0
0
0
0


31
4
1
0
0
0
0
2
5
8
7
4
0
0
0
0
0
0
0
0
0
0
0
0
0
0
0
0
0
0

1.0
1.0
1.0
1.0
1.0
1.0
1.0
1.0
1.0
1.0
1.0
1.0
1.0
1.0
1.0
1.0
1.0
1.0
1.0
1.0
1.0
1.0
1.0
1.0
1.0
1.0
1.0
1.0
1.0
1.0


17
3
0
0
0
0
0
0
0
6
3
2
0
0
0
0
0
3
0
0
0
0
0
0
0
0
0
0
0
0

1.0
1.0
1.0
1.0
1.0
1.0
1.0
1.0
1.0
1.0
1.0
1.0
1.0
1.0
1.0
1.0
1.0
1.0
1.0
1.0
1.0
1.0
1.0
1.0
1.0
1.0
1.0
1.0
1.0
1.0


5
1
0
0
0
0
0
0
0
3
0
1
0
0
0
0
0
0
0
0
0
0
0
0
0
0
0
0
0
0

1.0
1.0
1.0
1.0
1.0
1.0
1.0
1.0
1.0
1.0
1.0
1.0
1.0
1.0
1.0
1.0
1.0
1.0
1.0
1.0
1.0
1.0
1.0
1.0
1.0
1.0
1.0
1.0
1.0
1.0


2
0
0
0
0
0
0
0
0
1
1
0
0
0
0
0
0
0
0
0
0
0
0
0
0
0
0
0
0
0

1.0
1.0
1.0
1.0
1.0
1.0
1.0
1.0
1.0
1.0
1.0
1.0
1.0
1.0
1.0
1.0
1.0
1.0
1.0
1.0
1.0
1.0
1.0
1.0
1.0
1.0
1.0
1.0
1.0
1.0


3
0
0
0
0
0
0
1
1
0
1
0
0
0
0
0
0
0
0
0
0
0
0
0
0
0
0
0
0
0


3
0
0
0
0
0
0
1
1
0
1
0
0
0
0
0
0
0
0
0
0
0
0
0
0
0
0
0
0
0

1.0
1.0
1.0
1.0
1.0
1.0
1.0
1.0
1.0
1.0
1.0
1.0
1.0
1.0
1.0
1.0
1.0
1.0
1.0
1.0
1.0
1.0
1.0
1.0
1.0
1.0
1.0
1.0
1.0
1.0


2
0
0
0
0
0
0
0
0
0
0
1
0
0
0
0
0
0
0
0
0
0
0
0
0
0
0
0
0
1


1
0
0
0
0
0
0
0
0
0
0
1
0
0
0
0
0
0
0
0
0
0
0
0
0
0
0
0
0
0

1.0
1.0
1.0
1.0
1.0
1.0
1.0
1.0
1.0
1.0
1.0
1.0
1.0
1.0
1.0
1.0
1.0
1.0
1.0
1.0
1.0
1.0
1.0
1.0
1.0
1.0
1.0
1.0
1.0
1.0


1
0
0
0
0
0
0
0
0
0
0
0
0
0
0
0
0
0
0
0
0
0
0
0
0
0
0
0
0
1

1.0
1.0
1.0
1.0
1.0
1.0
1.0
1.0
1.0
1.0
1.0
1.0
1.0
1.0
1.0
1.0
1.0
1.0
1.0
1.0
1.0
1.0
1.0
1.0
1.0
1.0
1.0
1.0
1.0
1.0


1
0
0
0
0
0
0
0
0
1
0
0
0
0
0
0
0
0
0
0
0
0
0
0
0
0
0
0
0
0


1
0
0
0
0
0
0
0
0
1
0
0
0
0
0
0
0
0
0
0
0
0
0
0
0
0
0
0
0
0

1.0
1.0
1.0
1.0
1.0
1.0
1.0
1.0
1.0
1.0
1.0
1.0
1.0
1.0
1.0
1.0
1.0
1.0
1.0
1.0
1.0
1.0
1.0
1.0
1.0
1.0
1.0
1.0
1.0
1.0


1
0
0
0
0
0
0
0
0
1
0
0
0
0
0
0
0
0
0
0
0
0
0
0
0
0
0
0
0
0


1
0
0
0
0
0
0
0
0
1
0
0
0
0
0
0
0
0
0
0
0
0
0
0
0
0
0
0
0
0

1.0
1.0
1.0
1.0
1.0
1.0
1.0
1.0
1.0
1.0
1.0
1.0
1.0
1.0
1.0
1.0
1.0
1.0
1.0
1.0
1.0
1.0
1.0
1.0
1.0
1.0
1.0
1.0
1.0
1.0


4
1
0
0
0
0
0
0
0
0
1
1
0
0
1
0
0
0
0
0
0
0
0
0
0
0
0
0
0
0

0.97
0.97
0.97
0.97
0.97
0.97
0.97
0.97
0.97
0.97
0.97
0.97
0.97
0.97
0.97
0.97
0.97
0.97
0.97
0.97
0.97
0.97
0.97
0.97
0.97
0.97
0.97
0.97
0.97
0.97


3
0
0
0
0
0
0
0
0
1
0
0
0
0
0
0
0
2
0
0
0
0
0
0
0
0
0
0
0
0


3
0
0
0
0
0
0
0
0
1
0
0
0
0
0
0
0
2
0
0
0
0
0
0
0
0
0
0
0
0

1.0
1.0
1.0
1.0
1.0
1.0
1.0
1.0
1.0
1.0
1.0
1.0
1.0
1.0
1.0
1.0
1.0
1.0
1.0
1.0
1.0
1.0
1.0
1.0
1.0
1.0
1.0
1.0
1.0
1.0


3
0
0
0
0
0
0
0
1
0
0
0
0
0
0
0
0
0
0
0
0
0
0
0
0
0
1
0
0
1


3
0
0
0
0
0
0
0
1
0
0
0
0
0
0
0
0
0
0
0
0
0
0
0
0
0
1
0
0
1

1.0
1.0
1.0
1.0
1.0
1.0
1.0
1.0
1.0
1.0
1.0
1.0
1.0
1.0
1.0
1.0
1.0
1.0
1.0
1.0
1.0
1.0
1.0
1.0
1.0
1.0
1.0
1.0
1.0
1.0


6
0
0
0
0
0
0
0
0
0
0
0
0
0
0
0
0
0
0
0
2
2
2
0
0
0
0
0
0
0

0.7
0.7
0.7
0.7
0.7
0.7
0.7
0.7
0.7
0.7
0.7
0.7
0.7
0.7
0.7
0.7
0.7
0.7
0.7
0.7
0.7
0.7
0.7
0.7
0.7
0.7
0.7
0.7
0.7
0.7


3
0
0
0
0
0
0
0
0
0
0
0
0
0
0
0
0
0
0
0
1
1
1
0
0
0
0
0
0
0

0.86
0.86
0.86
0.86
0.86
0.86
0.86
0.86
0.86
0.86
0.86
0.86
0.86
0.86
0.86
0.86
0.86
0.86
0.86
0.86
0.86
0.86
0.86
0.86
0.86
0.86
0.86
0.86
0.86
0.86


1
0
0
0
0
0
0
0
0
0
0
0
0
0
0
0
0
0
1
0
0
0
0
0
0
0
0
0
0
0


1
0
0
0
0
0
0
0
0
0
0
0
0
0
0
0
0
0
1
0
0
0
0
0
0
0
0
0
0
0

1.0
1.0
1.0
1.0
1.0
1.0
1.0
1.0
1.0
1.0
1.0
1.0
1.0
1.0
1.0
1.0
1.0
1.0
1.0
1.0
1.0
1.0
1.0
1.0
1.0
1.0
1.0
1.0
1.0
1.0


24
1
1
0
1
2
1
4
1
5
3
3
0
0
1
0
0
0
1
0
0
0
0
0
0
0
0
0
0
0


9
1
1
0
0
0
0
2
1
1
1
1
0
0
0
0
0
0
1
0
0
0
0
0
0
0
0
0
0
0


9
1
1
0
0
0
0
2
1
1
1
1
0
0
0
0
0
0
1
0
0
0
0
0
0
0
0
0
0
0

1.0
1.0
1.0
1.0
1.0
1.0
1.0
1.0
1.0
1.0
1.0
1.0
1.0
1.0
1.0
1.0
1.0
1.0
1.0
1.0
1.0
1.0
1.0
1.0
1.0
1.0
1.0
1.0
1.0
1.0


15
0
0
0
1
2
1
2
0
4
2
2
0
0
1
0
0
0
0
0
0
0
0
0
0
0
0
0
0
0


15
0
0
0
1
2
1
2
0
4
2
2
0
0
1
0
0
0
0
0
0
0
0
0
0
0
0
0
0
0

1.0
1.0
1.0
1.0
1.0
1.0
1.0
1.0
1.0
1.0
1.0
1.0
1.0
1.0
1.0
1.0
1.0
1.0
1.0
1.0
1.0
1.0
1.0
1.0
1.0
1.0
1.0
1.0
1.0
1.0


43
3
6
1
3
1
0
0
1
3
6
5
0
2
2
0
0
0
0
1
0
0
0
0
0
6
0
0
0
3

0.93
0.93
0.93
0.93
0.93
0.93
0.93
0.93
0.93
0.93
0.93
0.93
0.93
0.93
0.93
0.93
0.93
0.93
0.93
0.93
0.93
0.93
0.93
0.93
0.93
0.93
0.93
0.93
0.93
0.93


2
0
0
0
0
0
0
0
0
0
2
0
0
0
0
0
0
0
0
0
0
0
0
0
0
0
0
0
0
0


2
0
0
0
0
0
0
0
0
0
2
0
0
0
0
0
0
0
0
0
0
0
0
0
0
0
0
0
0
0

1.0
1.0
1.0
1.0
1.0
1.0
1.0
1.0
1.0
1.0
1.0
1.0
1.0
1.0
1.0
1.0
1.0
1.0
1.0
1.0
1.0
1.0
1.0
1.0
1.0
1.0
1.0
1.0
1.0
1.0


27
3
6
1
3
1
0
0
0
0
2
5
0
2
2
0
0
0
0
1
0
0
0
0
0
1
0
0
0
0

1.0
1.0
1.0
1.0
1.0
1.0
1.0
1.0
1.0
1.0
1.0
1.0
1.0
1.0
1.0
1.0
1.0
1.0
1.0
1.0
1.0
1.0
1.0
1.0
1.0
1.0
1.0
1.0
1.0
1.0


1
0
0
0
0
0
0
0
0
0
1
0
0
0
0
0
0
0
0
0
0
0
0
0
0
0
0
0
0
0


1
0
0
0
0
0
0
0
0
0
1
0
0
0
0
0
0
0
0
0
0
0
0
0
0
0
0
0
0
0

0.93
0.93
0.93
0.93
0.93
0.93
0.93
0.93
0.93
0.93
0.93
0.93
0.93
0.93
0.93
0.93
0.93
0.93
0.93
0.93
0.93
0.93
0.93
0.93
0.93
0.93
0.93
0.93
0.93
0.93


1
0
0
0
0
0
0
0
0
1
0
0
0
0
0
0
0
0
0
0
0
0
0
0
0
0
0
0
0
0

1.0
1.0
1.0
1.0
1.0
1.0
1.0
1.0
1.0
1.0
1.0
1.0
1.0
1.0
1.0
1.0
1.0
1.0
1.0
1.0
1.0
1.0
1.0
1.0
1.0
1.0
1.0
1.0
1.0
1.0


2
0
0
0
0
0
0
0
0
1
1
0
0
0
0
0
0
0
0
0
0
0
0
0
0
0
0
0
0
0

0.99
0.99
0.99
0.99
0.99
0.99
0.99
0.99
0.99
0.99
0.99
0.99
0.99
0.99
0.99
0.99
0.99
0.99
0.99
0.99
0.99
0.99
0.99
0.99
0.99
0.99
0.99
0.99
0.99
0.99


1
0
0
0
0
0
0
0
0
1
0
0
0
0
0
0
0
0
0
0
0
0
0
0
0
0
0
0
0
0


1
0
0
0
0
0
0
0
0
1
0
0
0
0
0
0
0
0
0
0
0
0
0
0
0
0
0
0
0
0

1.0
1.0
1.0
1.0
1.0
1.0
1.0
1.0
1.0
1.0
1.0
1.0
1.0
1.0
1.0
1.0
1.0
1.0
1.0
1.0
1.0
1.0
1.0
1.0
1.0
1.0
1.0
1.0
1.0
1.0


1
0
0
0
0
0
0
0
0
0
0
0
0
0
0
0
0
0
0
0
0
0
0
0
0
1
0
0
0
0


1
0
0
0
0
0
0
0
0
0
0
0
0
0
0
0
0
0
0
0
0
0
0
0
0
1
0
0
0
0

0.97
0.97
0.97
0.97
0.97
0.97
0.97
0.97
0.97
0.97
0.97
0.97
0.97
0.97
0.97
0.97
0.97
0.97
0.97
0.97
0.97
0.97
0.97
0.97
0.97
0.97
0.97
0.97
0.97
0.97


53
3
1
0
0
0
0
1
7
12
12
2
2
0
4
0
0
1
1
1
2
0
1
2
0
0
1
0
0
0

0.98
0.98
0.98
0.98
0.98
0.98
0.98
0.98
0.98
0.98
0.98
0.98
0.98
0.98
0.98
0.98
0.98
0.98
0.98
0.98
0.98
0.98
0.98
0.98
0.98
0.98
0.98
0.98
0.98
0.98


11
0
0
0
0
0
0
0
2
5
4
0
0
0
0
0
0
0
0
0
0
0
0
0
0
0
0
0
0
0


7
0
0
0
0
0
0
0
1
3
3
0
0
0
0
0
0
0
0
0
0
0
0
0
0
0
0
0
0
0

1.0
1.0
1.0
1.0
1.0
1.0
1.0
1.0
1.0
1.0
1.0
1.0
1.0
1.0
1.0
1.0
1.0
1.0
1.0
1.0
1.0
1.0
1.0
1.0
1.0
1.0
1.0
1.0
1.0
1.0


4
0
0
0
0
0
0
0
1
2
1
0
0
0
0
0
0
0
0
0
0
0
0
0
0
0
0
0
0
0

0.99
0.99
0.99
0.99
0.99
0.99
0.99
0.99
0.99
0.99
0.99
0.99
0.99
0.99
0.99
0.99
0.99
0.99
0.99
0.99
0.99
0.99
0.99
0.99
0.99
0.99
0.99
0.99
0.99
0.99


10
1
1
0
0
0
0
0
0
3
1
0
0
0
4
0
0
0
0
0
0
0
0
0
0
0
0
0
0
0


10
1
1
0
0
0
0
0
0
3
1
0
0
0
4
0
0
0
0
0
0
0
0
0
0
0
0
0
0
0

1.0
1.0
1.0
1.0
1.0
1.0
1.0
1.0
1.0
1.0
1.0
1.0
1.0
1.0
1.0
1.0
1.0
1.0
1.0
1.0
1.0
1.0
1.0
1.0
1.0
1.0
1.0
1.0
1.0
1.0


13
0
0
0
0
0
0
0
3
3
6
1
0
0
0
0
0
0
0
0
0
0
0
0
0
0
0
0
0
0


13
0
0
0
0
0
0
0
3
3
6
1
0
0
0
0
0
0
0
0
0
0
0
0
0
0
0
0
0
0

1.0
1.0
1.0
1.0
1.0
1.0
1.0
1.0
1.0
1.0
1.0
1.0
1.0
1.0
1.0
1.0
1.0
1.0
1.0
1.0
1.0
1.0
1.0
1.0
1.0
1.0
1.0
1.0
1.0
1.0


1
0
0
0
0
0
0
0
1
0
0
0
0
0
0
0
0
0
0
0
0
0
0
0
0
0
0
0
0
0


1
0
0
0
0
0
0
0
1
0
0
0
0
0
0
0
0
0
0
0
0
0
0
0
0
0
0
0
0
0

0.96
0.96
0.96
0.96
0.96
0.96
0.96
0.96
0.96
0.96
0.96
0.96
0.96
0.96
0.96
0.96
0.96
0.96
0.96
0.96
0.96
0.96
0.96
0.96
0.96
0.96
0.96
0.96
0.96
0.96


1
1
0
0
0
0
0
0
0
0
0
0
0
0
0
0
0
0
0
0
0
0
0
0
0
0
0
0
0
0


1
1
0
0
0
0
0
0
0
0
0
0
0
0
0
0
0
0
0
0
0
0
0
0
0
0
0
0
0
0

1.0
1.0
1.0
1.0
1.0
1.0
1.0
1.0
1.0
1.0
1.0
1.0
1.0
1.0
1.0
1.0
1.0
1.0
1.0
1.0
1.0
1.0
1.0
1.0
1.0
1.0
1.0
1.0
1.0
1.0


1
0
0
0
0
0
0
1
0
0
0
0
0
0
0
0
0
0
0
0
0
0
0
0
0
0
0
0
0
0


1
0
0
0
0
0
0
1
0
0
0
0
0
0
0
0
0
0
0
0
0
0
0
0
0
0
0
0
0
0

0.75
0.75
0.75
0.75
0.75
0.75
0.75
0.75
0.75
0.75
0.75
0.75
0.75
0.75
0.75
0.75
0.75
0.75
0.75
0.75
0.75
0.75
0.75
0.75
0.75
0.75
0.75
0.75
0.75
0.75


154
6
7
5
3
6
7
4
4
10
8
6
0
0
5
0
0
0
25
0
6
0
0
0
4
14
8
9
8
9

0.85
0.85
0.85
0.85
0.85
0.85
0.85
0.85
0.85
0.85
0.85
0.85
0.85
0.85
0.85
0.85
0.85
0.85
0.85
0.85
0.85
0.85
0.85
0.85
0.85
0.85
0.85
0.85
0.85
0.85


35
0
3
3
3
6
6
0
0
1
1
2
0
0
0
0
0
0
2
0
0
0
0
0
1
0
1
2
1
3

0.73
0.73
0.73
0.73
0.73
0.73
0.73
0.73
0.73
0.73
0.73
0.73
0.73
0.73
0.73
0.73
0.73
0.73
0.73
0.73
0.73
0.73
0.73
0.73
0.73
0.73
0.73
0.73
0.73
0.73


33
0
3
3
3
6
6
0
0
1
1
2
0
0
0
0
0
0
2
0
0
0
0
0
1
0
1
1
1
2

1.0
1.0
1.0
1.0
1.0
1.0
1.0
1.0
1.0
1.0
1.0
1.0
1.0
1.0
1.0
1.0
1.0
1.0
1.0
1.0
1.0
1.0
1.0
1.0
1.0
1.0
1.0
1.0
1.0
1.0


16
0
0
0
0
0
0
0
0
4
2
2
0
0
0
0
0
0
8
0
0
0
0
0
0
0
0
0
0
0


16
0
0
0
0
0
0
0
0
4
2
2
0
0
0
0
0
0
8
0
0
0
0
0
0
0
0
0
0
0

1.0
1.0
1.0
1.0
1.0
1.0
1.0
1.0
1.0
1.0
1.0
1.0
1.0
1.0
1.0
1.0
1.0
1.0
1.0
1.0
1.0
1.0
1.0
1.0
1.0
1.0
1.0
1.0
1.0
1.0


29
4
1
0
0
0
1
4
3
5
3
1
0
0
2
0
0
0
0
0
0
0
0
0
0
3
2
0
0
0

0.93
0.93
0.93
0.93
0.93
0.93
0.93
0.93
0.93
0.93
0.93
0.93
0.93
0.93
0.93
0.93
0.93
0.93
0.93
0.93
0.93
0.93
0.93
0.93
0.93
0.93
0.93
0.93
0.93
0.93


27
4
1
0
0
0
0
4
3
5
3
1
0
0
2
0
0
0
0
0
0
0
0
0
0
2
2
0
0
0

1.0
1.0
1.0
1.0
1.0
1.0
1.0
1.0
1.0
1.0
1.0
1.0
1.0
1.0
1.0
1.0
1.0
1.0
1.0
1.0
1.0
1.0
1.0
1.0
1.0
1.0
1.0
1.0
1.0
1.0


7
1
3
2
0
0
0
0
0
0
0
1
0
0
0
0
0
0
0
0
0
0
0
0
0
0
0
0
0
0


7
1
3
2
0
0
0
0
0
0
0
1
0
0
0
0
0
0
0
0
0
0
0
0
0
0
0
0
0
0

1.0
1.0
1.0
1.0
1.0
1.0
1.0
1.0
1.0
1.0
1.0
1.0
1.0
1.0
1.0
1.0
1.0
1.0
1.0
1.0
1.0
1.0
1.0
1.0
1.0
1.0
1.0
1.0
1.0
1.0


2
1
0
0
0
0
0
0
1
0
0
0
0
0
0
0
0
0
0
0
0
0
0
0
0
0
0
0
0
0

1.0
1.0
1.0
1.0
1.0
1.0
1.0
1.0
1.0
1.0
1.0
1.0
1.0
1.0
1.0
1.0
1.0
1.0
1.0
1.0
1.0
1.0
1.0
1.0
1.0
1.0
1.0
1.0
1.0
1.0


22
0
0
0
0
0
0
0
0
0
1
0
0
0
1
0
0
0
14
0
4
0
0
0
1
0
0
0
0
1

0.99
0.99
0.99
0.99
0.99
0.99
0.99
0.99
0.99
0.99
0.99
0.99
0.99
0.99
0.99
0.99
0.99
0.99
0.99
0.99
0.99
0.99
0.99
0.99
0.99
0.99
0.99
0.99
0.99
0.99


15
0
0
0
0
0
0
0
0
0
0
0
0
0
0
0
0
0
14
0
0
0
0
0
0
0
0
0
0
1

1.0
1.0
1.0
1.0
1.0
1.0
1.0
1.0
1.0
1.0
1.0
1.0
1.0
1.0
1.0
1.0
1.0
1.0
1.0
1.0
1.0
1.0
1.0
1.0
1.0
1.0
1.0
1.0
1.0
1.0


4
0
0
0
0
0
0
0
0
0
0
0
0
0
0
0
0
0
0
0
0
0
0
0
0
3
1
0
0
0

0.93
0.93
0.93
0.93
0.93
0.93
0.93
0.93
0.93
0.93
0.93
0.93
0.93
0.93
0.93
0.93
0.93
0.93
0.93
0.93
0.93
0.93
0.93
0.93
0.93
0.93
0.93
0.93
0.93
0.93


33
2
1
1
2
1
1
0
2
7
7
3
0
0
0
0
0
0
0
0
0
0
0
0
0
2
0
0
0
4

0.9
0.9
0.9
0.9
0.9
0.9
0.9
0.9
0.9
0.9
0.9
0.9
0.9
0.9
0.9
0.9
0.9
0.9
0.9
0.9
0.9
0.9
0.9
0.9
0.9
0.9
0.9
0.9
0.9
0.9


8
1
1
1
1
0
1
0
0
1
1
1
0
0
0
0
0
0
0
0
0
0
0
0
0
0
0
0
0
0


8
1
1
1
1
0
1
0
0
1
1
1
0
0
0
0
0
0
0
0
0
0
0
0
0
0
0
0
0
0

1.0
1.0
1.0
1.0
1.0
1.0
1.0
1.0
1.0
1.0
1.0
1.0
1.0
1.0
1.0
1.0
1.0
1.0
1.0
1.0
1.0
1.0
1.0
1.0
1.0
1.0
1.0
1.0
1.0
1.0


3
0
0
0
0
0
0
0
1
1
1
0
0
0
0
0
0
0
0
0
0
0
0
0
0
0
0
0
0
0


3
0
0
0
0
0
0
0
1
1
1
0
0
0
0
0
0
0
0
0
0
0
0
0
0
0
0
0
0
0

1.0
1.0
1.0
1.0
1.0
1.0
1.0
1.0
1.0
1.0
1.0
1.0
1.0
1.0
1.0
1.0
1.0
1.0
1.0
1.0
1.0
1.0
1.0
1.0
1.0
1.0
1.0
1.0
1.0
1.0


6
1
0
0
0
0
0
0
1
2
2
0
0
0
0
0
0
0
0
0
0
0
0
0
0
0
0
0
0
0


6
1
0
0
0
0
0
0
1
2
2
0
0
0
0
0
0
0
0
0
0
0
0
0
0
0
0
0
0
0

1.0
1.0
1.0
1.0
1.0
1.0
1.0
1.0
1.0
1.0
1.0
1.0
1.0
1.0
1.0
1.0
1.0
1.0
1.0
1.0
1.0
1.0
1.0
1.0
1.0
1.0
1.0
1.0
1.0
1.0


1
0
0
0
0
0
0
0
0
1
0
0
0
0
0
0
0
0
0
0
0
0
0
0
0
0
0
0
0
0


1
0
0
0
0
0
0
0
0
1
0
0
0
0
0
0
0
0
0
0
0
0
0
0
0
0
0
0
0
0

1.0
1.0
1.0
1.0
1.0
1.0
1.0
1.0
1.0
1.0
1.0
1.0
1.0
1.0
1.0
1.0
1.0
1.0
1.0
1.0
1.0
1.0
1.0
1.0
1.0
1.0
1.0
1.0
1.0
1.0


2
0
0
0
0
0
0
0
0
1
0
1
0
0
0
0
0
0
0
0
0
0
0
0
0
0
0
0
0
0


2
0
0
0
0
0
0
0
0
1
0
1
0
0
0
0
0
0
0
0
0
0
0
0
0
0
0
0
0
0

1.0
1.0
1.0
1.0
1.0
1.0
1.0
1.0
1.0
1.0
1.0
1.0
1.0
1.0
1.0
1.0
1.0
1.0
1.0
1.0
1.0
1.0
1.0
1.0
1.0
1.0
1.0
1.0
1.0
1.0


3
0
0
0
0
0
0
0
0
1
1
1
0
0
0
0
0
0
0
0
0
0
0
0
0
0
0
0
0
0


3
0
0
0
0
0
0
0
0
1
1
1
0
0
0
0
0
0
0
0
0
0
0
0
0
0
0
0
0
0

1.0
1.0
1.0
1.0
1.0
1.0
1.0
1.0
1.0
1.0
1.0
1.0
1.0
1.0
1.0
1.0
1.0
1.0
1.0
1.0
1.0
1.0
1.0
1.0
1.0
1.0
1.0
1.0
1.0
1.0


3
0
0
0
1
1
0
0
0
0
1
0
0
0
0
0
0
0
0
0
0
0
0
0
0
0
0
0
0
0

1.0
1.0
1.0
1.0
1.0
1.0
1.0
1.0
1.0
1.0
1.0
1.0
1.0
1.0
1.0
1.0
1.0
1.0
1.0
1.0
1.0
1.0
1.0
1.0
1.0
1.0
1.0
1.0
1.0
1.0


1
0
0
0
0
0
0
0
0
0
1
0
0
0
0
0
0
0
0
0
0
0
0
0
0
0
0
0
0
0


1
0
0
0
0
0
0
0
0
0
1
0
0
0
0
0
0
0
0
0
0
0
0
0
0
0
0
0
0
0

1.0
1.0
1.0
1.0
1.0
1.0
1.0
1.0
1.0
1.0
1.0
1.0
1.0
1.0
1.0
1.0
1.0
1.0
1.0
1.0
1.0
1.0
1.0
1.0
1.0
1.0
1.0
1.0
1.0
1.0


1
0
0
0
0
0
0
0
0
0
0
0
0
0
0
0
0
0
0
0
0
0
0
0
0
0
0
0
0
1

0.77
0.77
0.77
0.77
0.77
0.77
0.77
0.77
0.77
0.77
0.77
0.77
0.77
0.77
0.77
0.77
0.77
0.77
0.77
0.77
0.77
0.77
0.77
0.77
0.77
0.77
0.77
0.77
0.77
0.77


27
3
0
0
0
0
1
0
0
0
0
0
0
0
0
0
0
20
0
0
0
0
0
0
0
0
0
1
0
2


22
2
0
0
0
0
0
0
0
0
0
0
0
0
0
0
0
20
0
0
0
0
0
0
0
0
0
0
0
0


22
2
0
0
0
0
0
0
0
0
0
0
0
0
0
0
0
20
0
0
0
0
0
0
0
0
0
0
0
0

1.0
1.0
1.0
1.0
1.0
1.0
1.0
1.0
1.0
1.0
1.0
1.0
1.0
1.0
1.0
1.0
1.0
1.0
1.0
1.0
1.0
1.0
1.0
1.0
1.0
1.0
1.0
1.0
1.0
1.0


5
1
0
0
0
0
1
0
0
0
0
0
0
0
0
0
0
0
0
0
0
0
0
0
0
0
0
1
0
2


5
1
0
0
0
0
1
0
0
0
0
0
0
0
0
0
0
0
0
0
0
0
0
0
0
0
0
1
0
2

1.0
1.0
1.0
1.0
1.0
1.0
1.0
1.0
1.0
1.0
1.0
1.0
1.0
1.0
1.0
1.0
1.0
1.0
1.0
1.0
1.0
1.0
1.0
1.0
1.0
1.0
1.0
1.0
1.0
1.0


2
0
0
0
0
0
0
0
2
0
0
0
0
0
0
0
0
0
0
0
0
0
0
0
0
0
0
0
0
0


2
0
0
0
0
0
0
0
2
0
0
0
0
0
0
0
0
0
0
0
0
0
0
0
0
0
0
0
0
0

0.99
0.99
0.99
0.99
0.99
0.99
0.99
0.99
0.99
0.99
0.99
0.99
0.99
0.99
0.99
0.99
0.99
0.99
0.99
0.99
0.99
0.99
0.99
0.99
0.99
0.99
0.99
0.99
0.99
0.99


150
22
18
6
1
5
1
4
11
18
23
19
0
0
1
1
0
1
9
0
1
0
0
0
1
1
1
0
1
5

0.96
0.96
0.96
0.96
0.96
0.96
0.96
0.96
0.96
0.96
0.96
0.96
0.96
0.96
0.96
0.96
0.96
0.96
0.96
0.96
0.96
0.96
0.96
0.96
0.96
0.96
0.96
0.96
0.96
0.96


29
3
4
2
1
1
1
0
0
2
3
2
0
0
1
0
0
1
2
0
1
0
0
0
0
0
0
0
1
4


14
2
3
2
1
1
1
0
0
1
1
1
0
0
1
0
0
0
0
0
0
0
0
0
0
0
0
0
0
0

1.0
1.0
1.0
1.0
1.0
1.0
1.0
1.0
1.0
1.0
1.0
1.0
1.0
1.0
1.0
1.0
1.0
1.0
1.0
1.0
1.0
1.0
1.0
1.0
1.0
1.0
1.0
1.0
1.0
1.0


6
1
1
0
0
0
0
0
0
1
1
1
0
0
0
0
0
0
0
0
1
0
0
0
0
0
0
0
0
0

1.0
1.0
1.0
1.0
1.0
1.0
1.0
1.0
1.0
1.0
1.0
1.0
1.0
1.0
1.0
1.0
1.0
1.0
1.0
1.0
1.0
1.0
1.0
1.0
1.0
1.0
1.0
1.0
1.0
1.0


3
0
0
0
0
0
0
0
0
0
1
0
0
0
0
0
0
1
1
0
0
0
0
0
0
0
0
0
0
0

1.0
1.0
1.0
1.0
1.0
1.0
1.0
1.0
1.0
1.0
1.0
1.0
1.0
1.0
1.0
1.0
1.0
1.0
1.0
1.0
1.0
1.0
1.0
1.0
1.0
1.0
1.0
1.0
1.0
1.0


6
0
0
0
0
0
0
0
0
0
0
0
0
0
0
0
0
0
1
0
0
0
0
0
0
0
0
0
1
4

1.0
1.0
1.0
1.0
1.0
1.0
1.0
1.0
1.0
1.0
1.0
1.0
1.0
1.0
1.0
1.0
1.0
1.0
1.0
1.0
1.0
1.0
1.0
1.0
1.0
1.0
1.0
1.0
1.0
1.0


33
6
7
0
0
1
0
0
0
4
7
7
0
0
0
0
0
0
0
0
0
0
0
0
0
0
0
0
0
1

1.0
1.0
1.0
1.0
1.0
1.0
1.0
1.0
1.0
1.0
1.0
1.0
1.0
1.0
1.0
1.0
1.0
1.0
1.0
1.0
1.0
1.0
1.0
1.0
1.0
1.0
1.0
1.0
1.0
1.0


21
3
5
0
0
0
0
0
0
3
4
6
0
0
0
0
0
0
0
0
0
0
0
0
0
0
0
0
0
0

1.0
1.0
1.0
1.0
1.0
1.0
1.0
1.0
1.0
1.0
1.0
1.0
1.0
1.0
1.0
1.0
1.0
1.0
1.0
1.0
1.0
1.0
1.0
1.0
1.0
1.0
1.0
1.0
1.0
1.0


3
1
1
0
0
1
0
0
0
0
0
0
0
0
0
0
0
0
0
0
0
0
0
0
0
0
0
0
0
0

1.0
1.0
1.0
1.0
1.0
1.0
1.0
1.0
1.0
1.0
1.0
1.0
1.0
1.0
1.0
1.0
1.0
1.0
1.0
1.0
1.0
1.0
1.0
1.0
1.0
1.0
1.0
1.0
1.0
1.0


2
1
0
0
0
0
0
0
0
0
1
0
0
0
0
0
0
0
0
0
0
0
0
0
0
0
0
0
0
0

1.0
1.0
1.0
1.0
1.0
1.0
1.0
1.0
1.0
1.0
1.0
1.0
1.0
1.0
1.0
1.0
1.0
1.0
1.0
1.0
1.0
1.0
1.0
1.0
1.0
1.0
1.0
1.0
1.0
1.0


1
0
0
0
0
0
0
0
0
0
0
0
0
0
0
0
0
0
0
0
0
0
0
0
0
0
0
0
0
1

0.88
0.88
0.88
0.88
0.88
0.88
0.88
0.88
0.88
0.88
0.88
0.88
0.88
0.88
0.88
0.88
0.88
0.88
0.88
0.88
0.88
0.88
0.88
0.88
0.88
0.88
0.88
0.88
0.88
0.88


10
4
2
0
0
1
0
0
0
0
0
3
0
0
0
0
0
0
0
0
0
0
0
0
0
0
0
0
0
0

1.0
1.0
1.0
1.0
1.0
1.0
1.0
1.0
1.0
1.0
1.0
1.0
1.0
1.0
1.0
1.0
1.0
1.0
1.0
1.0
1.0
1.0
1.0
1.0
1.0
1.0
1.0
1.0
1.0
1.0


3
1
1
0
0
1
0
0
0
0
0
0
0
0
0
0
0
0
0
0
0
0
0
0
0
0
0
0
0
0

1.0
1.0
1.0
1.0
1.0
1.0
1.0
1.0
1.0
1.0
1.0
1.0
1.0
1.0
1.0
1.0
1.0
1.0
1.0
1.0
1.0
1.0
1.0
1.0
1.0
1.0
1.0
1.0
1.0
1.0


50
5
2
0
0
1
0
4
9
11
12
5
0
0
0
1
0
0
0
0
0
0
0
0
0
0
0
0
0
0

0.99
0.99
0.99
0.99
0.99
0.99
0.99
0.99
0.99
0.99
0.99
0.99
0.99
0.99
0.99
0.99
0.99
0.99
0.99
0.99
0.99
0.99
0.99
0.99
0.99
0.99
0.99
0.99
0.99
0.99


5
0
0
0
0
0
0
1
2
2
0
0
0
0
0
0
0
0
0
0
0
0
0
0
0
0
0
0
0
0

1.0
1.0
1.0
1.0
1.0
1.0
1.0
1.0
1.0
1.0
1.0
1.0
1.0
1.0
1.0
1.0
1.0
1.0
1.0
1.0
1.0
1.0
1.0
1.0
1.0
1.0
1.0
1.0
1.0
1.0


8
0
0
0
0
0
0
0
5
2
0
1
0
0
0
0
0
0
0
0
0
0
0
0
0
0
0
0
0
0

0.92
0.92
0.92
0.92
0.92
0.92
0.92
0.92
0.92
0.92
0.92
0.92
0.92
0.92
0.92
0.92
0.92
0.92
0.92
0.92
0.92
0.92
0.92
0.92
0.92
0.92
0.92
0.92
0.92
0.92


3
0
1
0
0
0
0
0
0
1
1
0
0
0
0
0
0
0
0
0
0
0
0
0
0
0
0
0
0
0

1.0
1.0
1.0
1.0
1.0
1.0
1.0
1.0
1.0
1.0
1.0
1.0
1.0
1.0
1.0
1.0
1.0
1.0
1.0
1.0
1.0
1.0
1.0
1.0
1.0
1.0
1.0
1.0
1.0
1.0


2
0
0
0
0
0
0
0
0
1
0
1
0
0
0
0
0
0
0
0
0
0
0
0
0
0
0
0
0
0

1.0
1.0
1.0
1.0
1.0
1.0
1.0
1.0
1.0
1.0
1.0
1.0
1.0
1.0
1.0
1.0
1.0
1.0
1.0
1.0
1.0
1.0
1.0
1.0
1.0
1.0
1.0
1.0
1.0
1.0


4
2
1
0
0
0
0
0
0
0
0
1
0
0
0
0
0
0
0
0
0
0
0
0
0
0
0
0
0
0

1.0
1.0
1.0
1.0
1.0
1.0
1.0
1.0
1.0
1.0
1.0
1.0
1.0
1.0
1.0
1.0
1.0
1.0
1.0
1.0
1.0
1.0
1.0
1.0
1.0
1.0
1.0
1.0
1.0
1.0


1
0
0
0
0
1
0
0
0
0
0
0
0
0
0
0
0
0
0
0
0
0
0
0
0
0
0
0
0
0

1.0
1.0
1.0
1.0
1.0
1.0
1.0
1.0
1.0
1.0
1.0
1.0
1.0
1.0
1.0
1.0
1.0
1.0
1.0
1.0
1.0
1.0
1.0
1.0
1.0
1.0
1.0
1.0
1.0
1.0


1
0
0
0
0
0
0
0
1
0
0
0
0
0
0
0
0
0
0
0
0
0
0
0
0
0
0
0
0
0


1
0
0
0
0
0
0
0
1
0
0
0
0
0
0
0
0
0
0
0
0
0
0
0
0
0
0
0
0
0

1.0
1.0
1.0
1.0
1.0
1.0
1.0
1.0
1.0
1.0
1.0
1.0
1.0
1.0
1.0
1.0
1.0
1.0
1.0
1.0
1.0
1.0
1.0
1.0
1.0
1.0
1.0
1.0
1.0
1.0


3
1
1
0
0
0
0
0
0
0
0
1
0
0
0
0
0
0
0
0
0
0
0
0
0
0
0
0
0
0


3
1
1
0
0
0
0
0
0
0
0
1
0
0
0
0
0
0
0
0
0
0
0
0
0
0
0
0
0
0

1.0
1.0
1.0
1.0
1.0
1.0
1.0
1.0
1.0
1.0
1.0
1.0
1.0
1.0
1.0
1.0
1.0
1.0
1.0
1.0
1.0
1.0
1.0
1.0
1.0
1.0
1.0
1.0
1.0
1.0


10
2
0
0
0
0
0
0
1
1
1
1
0
0
0
0
0
0
4
0
0
0
0
0
0
0
0
0
0
0


3
1
0
0
0
0
0
0
1
1
0
0
0
0
0
0
0
0
0
0
0
0
0
0
0
0
0
0
0
0

1.0
1.0
1.0
1.0
1.0
1.0
1.0
1.0
1.0
1.0
1.0
1.0
1.0
1.0
1.0
1.0
1.0
1.0
1.0
1.0
1.0
1.0
1.0
1.0
1.0
1.0
1.0
1.0
1.0
1.0


7
1
0
0
0
0
0
0
0
0
1
1
0
0
0
0
0
0
4
0
0
0
0
0
0
0
0
0
0
0

1.0
1.0
1.0
1.0
1.0
1.0
1.0
1.0
1.0
1.0
1.0
1.0
1.0
1.0
1.0
1.0
1.0
1.0
1.0
1.0
1.0
1.0
1.0
1.0
1.0
1.0
1.0
1.0
1.0
1.0


11
1
2
4
0
1
0
0
0
0
0
0
0
0
0
0
0
0
2
0
0
0
0
0
0
0
1
0
0
0

0.91
0.91
0.91
0.91
0.91
0.91
0.91
0.91
0.91
0.91
0.91
0.91
0.91
0.91
0.91
0.91
0.91
0.91
0.91
0.91
0.91
0.91
0.91
0.91
0.91
0.91
0.91
0.91
0.91
0.91


5
1
2
2
0
0
0
0
0
0
0
0
0
0
0
0
0
0
0
0
0
0
0
0
0
0
0
0
0
0

1.0
1.0
1.0
1.0
1.0
1.0
1.0
1.0
1.0
1.0
1.0
1.0
1.0
1.0
1.0
1.0
1.0
1.0
1.0
1.0
1.0
1.0
1.0
1.0
1.0
1.0
1.0
1.0
1.0
1.0


21
3
4
0
0
0
0
0
0
0
0
0
0
0
0
0
0
0
0
0
0
0
0
0
0
0
0
4
0
10


21
3
4
0
0
0
0
0
0
0
0
0
0
0
0
0
0
0
0
0
0
0
0
0
0
0
0
4
0
10

1.0
1.0
1.0
1.0
1.0
1.0
1.0
1.0
1.0
1.0
1.0
1.0
1.0
1.0
1.0
1.0
1.0
1.0
1.0
1.0
1.0
1.0
1.0
1.0
1.0
1.0
1.0
1.0
1.0
1.0


5
3
2
0
0
0
0
0
0
0
0
0
0
0
0
0
0
0
0
0
0
0
0
0
0
0
0
0
0
0

1.0
1.0
1.0
1.0
1.0
1.0
1.0
1.0
1.0
1.0
1.0
1.0
1.0
1.0
1.0
1.0
1.0
1.0
1.0
1.0
1.0
1.0
1.0
1.0
1.0
1.0
1.0
1.0
1.0
1.0


1
0
1
0
0
0
0
0
0
0
0
0
0
0
0
0
0
0
0
0
0
0
0
0
0
0
0
0
0
0

1.0
1.0
1.0
1.0
1.0
1.0
1.0
1.0
1.0
1.0
1.0
1.0
1.0
1.0
1.0
1.0
1.0
1.0
1.0
1.0
1.0
1.0
1.0
1.0
1.0
1.0
1.0
1.0
1.0
1.0


14
0
0
0
0
0
1
2
4
5
1
0
0
0
0
0
0
0
0
0
0
0
0
0
0
0
0
0
1
0


9
0
0
0
0
0
0
0
3
5
1
0
0
0
0
0
0
0
0
0
0
0
0
0
0
0
0
0
0
0


9
0
0
0
0
0
0
0
3
5
1
0
0
0
0
0
0
0
0
0
0
0
0
0
0
0
0
0
0
0

1.0
1.0
1.0
1.0
1.0
1.0
1.0
1.0
1.0
1.0
1.0
1.0
1.0
1.0
1.0
1.0
1.0
1.0
1.0
1.0
1.0
1.0
1.0
1.0
1.0
1.0
1.0
1.0
1.0
1.0


1
0
0
0
0
0
0
0
1
0
0
0
0
0
0
0
0
0
0
0
0
0
0
0
0
0
0
0
0
0


1
0
0
0
0
0
0
0
1
0
0
0
0
0
0
0
0
0
0
0
0
0
0
0
0
0
0
0
0
0

1.0
1.0
1.0
1.0
1.0
1.0
1.0
1.0
1.0
1.0
1.0
1.0
1.0
1.0
1.0
1.0
1.0
1.0
1.0
1.0
1.0
1.0
1.0
1.0
1.0
1.0
1.0
1.0
1.0
1.0


3
0
0
0
0
0
0
2
0
0
0
0
0
0
0
0
0
0
0
0
0
0
0
0
0
0
0
0
1
0


3
0
0
0
0
0
0
2
0
0
0
0
0
0
0
0
0
0
0
0
0
0
0
0
0
0
0
0
1
0

1.0
1.0
1.0
1.0
1.0
1.0
1.0
1.0
1.0
1.0
1.0
1.0
1.0
1.0
1.0
1.0
1.0
1.0
1.0
1.0
1.0
1.0
1.0
1.0
1.0
1.0
1.0
1.0
1.0
1.0


1
0
0
0
0
0
1
0
0
0
0
0
0
0
0
0
0
0
0
0
0
0
0
0
0
0
0
0
0
0


1
0
0
0
0
0
1
0
0
0
0
0
0
0
0
0
0
0
0
0
0
0
0
0
0
0
0
0
0
0

1.0
1.0
1.0
1.0
1.0
1.0
1.0
1.0
1.0
1.0
1.0
1.0
1.0
1.0
1.0
1.0
1.0
1.0
1.0
1.0
1.0
1.0
1.0
1.0
1.0
1.0
1.0
1.0
1.0
1.0


5
1
0
0
0
0
0
0
1
2
1
0
0
0
0
0
0
0
0
0
0
0
0
0
0
0
0
0
0
0


5
1
0
0
0
0
0
0
1
2
1
0
0
0
0
0
0
0
0
0
0
0
0
0
0
0
0
0
0
0


4
1
0
0
0
0
0
0
1
1
1
0
0
0
0
0
0
0
0
0
0
0
0
0
0
0
0
0
0
0

1.0
1.0
1.0
1.0
1.0
1.0
1.0
1.0
1.0
1.0
1.0
1.0
1.0
1.0
1.0
1.0
1.0
1.0
1.0
1.0
1.0
1.0
1.0
1.0
1.0
1.0
1.0
1.0
1.0
1.0


1
0
0
0
0
0
0
0
0
1
0
0
0
0
0
0
0
0
0
0
0
0
0
0
0
0
0
0
0
0

0.99
0.99
0.99
0.99
0.99
0.99
0.99
0.99
0.99
0.99
0.99
0.99
0.99
0.99
0.99
0.99
0.99
0.99
0.99
0.99
0.99
0.99
0.99
0.99
0.99
0.99
0.99
0.99
0.99
0.99


3
0
0
0
0
0
0
0
1
1
1
0
0
0
0
0
0
0
0
0
0
0
0
0
0
0
0
0
0
0


3
0
0
0
0
0
0
0
1
1
1
0
0
0
0
0
0
0
0
0
0
0
0
0
0
0
0
0
0
0

1.0
1.0
1.0
1.0
1.0
1.0
1.0
1.0
1.0
1.0
1.0
1.0
1.0
1.0
1.0
1.0
1.0
1.0
1.0
1.0
1.0
1.0
1.0
1.0
1.0
1.0
1.0
1.0
1.0
1.0


1
0
0
0
0
0
0
0
1
0
0
0
0
0
0
0
0
0
0
0
0
0
0
0
0
0
0
0
0
0

0.88
0.88
0.88
0.88
0.88
0.88
0.88
0.88
0.88
0.88
0.88
0.88
0.88
0.88
0.88
0.88
0.88
0.88
0.88
0.88
0.88
0.88
0.88
0.88
0.88
0.88
0.88
0.88
0.88
0.88


16
3
0
0
0
0
0
0
0
5
4
4
0
0
0
0
0
0
0
0
0
0
0
0
0
0
0
0
0
0


16
3
0
0
0
0
0
0
0
5
4
4
0
0
0
0
0
0
0
0
0
0
0
0
0
0
0
0
0
0


16
3
0
0
0
0
0
0
0
5
4
4
0
0
0
0
0
0
0
0
0
0
0
0
0
0
0
0
0
0

1.0
1.0
1.0
1.0
1.0
1.0
1.0
1.0
1.0
1.0
1.0
1.0
1.0
1.0
1.0
1.0
1.0
1.0
1.0
1.0
1.0
1.0
1.0
1.0
1.0
1.0
1.0
1.0
1.0
1.0


3
0
0
0
0
0
0
0
0
1
0
0
0
0
0
0
0
0
0
0
0
0
0
0
0
2
0
0
0
0


3
0
0
0
0
0
0
0
0
1
0
0
0
0
0
0
0
0
0
0
0
0
0
0
0
2
0
0
0
0


3
0
0
0
0
0
0
0
0
1
0
0
0
0
0
0
0
0
0
0
0
0
0
0
0
2
0
0
0
0

1.0
1.0
1.0
1.0
1.0
1.0
1.0
1.0
1.0
1.0
1.0
1.0
1.0
1.0
1.0
1.0
1.0
1.0
1.0
1.0
1.0
1.0
1.0
1.0
1.0
1.0
1.0
1.0
1.0
1.0


7
0
0
0
0
0
0
0
0
0
1
0
0
0
0
0
0
5
0
0
0
0
0
0
1
0
0
0
0
0

0.81
0.81
0.81
0.81
0.81
0.81
0.81
0.81
0.81
0.81
0.81
0.81
0.81
0.81
0.81
0.81
0.81
0.81
0.81
0.81
0.81
0.81
0.81
0.81
0.81
0.81
0.81
0.81
0.81
0.81


6
0
0
0
0
0
0
0
0
0
1
0
0
0
0
0
0
5
0
0
0
0
0
0
0
0
0
0
0
0

1.0
1.0
1.0
1.0
1.0
1.0
1.0
1.0
1.0
1.0
1.0
1.0
1.0
1.0
1.0
1.0
1.0
1.0
1.0
1.0
1.0
1.0
1.0
1.0
1.0
1.0
1.0
1.0
1.0
1.0


4
0
0
0
0
0
0
0
0
0
0
0
0
0
0
0
0
4
0
0
0
0
0
0
0
0
0
0
0
0

1.0
1.0
1.0
1.0
1.0
1.0
1.0
1.0
1.0
1.0
1.0
1.0
1.0
1.0
1.0
1.0
1.0
1.0
1.0
1.0
1.0
1.0
1.0
1.0
1.0
1.0
1.0
1.0
1.0
1.0


8
0
0
0
0
0
0
0
0
1
0
1
0
0
0
0
0
6
0
0
0
0
0
0
0
0
0
0
0
0


8
0
0
0
0
0
0
0
0
1
0
1
0
0
0
0
0
6
0
0
0
0
0
0
0
0
0
0
0
0

1.0
1.0
1.0
1.0
1.0
1.0
1.0
1.0
1.0
1.0
1.0
1.0
1.0
1.0
1.0
1.0
1.0
1.0
1.0
1.0
1.0
1.0
1.0
1.0
1.0
1.0
1.0
1.0
1.0
1.0


4
0
0
0
0
0
0
0
0
0
0
0
0
0
0
0
0
4
0
0
0
0
0
0
0
0
0
0
0
0

1.0
1.0
1.0
1.0
1.0
1.0
1.0
1.0
1.0
1.0
1.0
1.0
1.0
1.0
1.0
1.0
1.0
1.0
1.0
1.0
1.0
1.0
1.0
1.0
1.0
1.0
1.0
1.0
1.0
1.0


7
1
1
0
0
0
0
0
0
0
0
0
1
0
0
0
0
1
1
0
0
0
1
1
0
0
0
0
0
0


7
1
1
0
0
0
0
0
0
0
0
0
1
0
0
0
0
1
1
0
0
0
1
1
0
0
0
0
0
0

0.73
0.73
0.73
0.73
0.73
0.73
0.73
0.73
0.73
0.73
0.73
0.73
0.73
0.73
0.73
0.73
0.73
0.73
0.73
0.73
0.73
0.73
0.73
0.73
0.73
0.73
0.73
0.73
0.73
0.73


1
1
0
0
0
0
0
0
0
0
0
0
0
0
0
0
0
0
0
0
0
0
0
0
0
0
0
0
0
0

1.0
1.0
1.0
1.0
1.0
1.0
1.0
1.0
1.0
1.0
1.0
1.0
1.0
1.0
1.0
1.0
1.0
1.0
1.0
1.0
1.0
1.0
1.0
1.0
1.0
1.0
1.0
1.0
1.0
1.0


1
0
1
0
0
0
0
0
0
0
0
0
0
0
0
0
0
0
0
0
0
0
0
0
0
0
0
0
0
0

1.0
1.0
1.0
1.0
1.0
1.0
1.0
1.0
1.0
1.0
1.0
1.0
1.0
1.0
1.0
1.0
1.0
1.0
1.0
1.0
1.0
1.0
1.0
1.0
1.0
1.0
1.0
1.0
1.0
1.0


1
0
0
0
0
0
0
0
0
1
0
0
0
0
0
0
0
0
0
0
0
0
0
0
0
0
0
0
0
0


1
0
0
0
0
0
0
0
0
1
0
0
0
0
0
0
0
0
0
0
0
0
0
0
0
0
0
0
0
0

0.99
0.99
0.99
0.99
0.99
0.99
0.99
0.99
0.99
0.99
0.99
0.99
0.99
0.99
0.99
0.99
0.99
0.99
0.99
0.99
0.99
0.99
0.99
0.99
0.99
0.99
0.99
0.99
0.99
0.99


1
0
0
0
0
0
0
0
0
0
1
0
0
0
0
0
0
0
0
0
0
0
0
0
0
0
0
0
0
0


1
0
0
0
0
0
0
0
0
0
1
0
0
0
0
0
0
0
0
0
0
0
0
0
0
0
0
0
0
0


1
0
0
0
0
0
0
0
0
0
1
0
0
0
0
0
0
0
0
0
0
0
0
0
0
0
0
0
0
0

1.0
1.0
1.0
1.0
1.0
1.0
1.0
1.0
1.0
1.0
1.0
1.0
1.0
1.0
1.0
1.0
1.0
1.0
1.0
1.0
1.0
1.0
1.0
1.0
1.0
1.0
1.0
1.0
1.0
1.0


3
0
0
1
1
0
1
0
0
0
0
0
0
0
0
0
0
0
0
0
0
0
0
0
0
0
0
0
0
0

0.73
0.73
0.73
0.73
0.73
0.73
0.73
0.73
0.73
0.73
0.73
0.73
0.73
0.73
0.73
0.73
0.73
0.73
0.73
0.73
0.73
0.73
0.73
0.73
0.73
0.73
0.73
0.73
0.73
0.73


24
0
0
0
0
0
0
1
0
0
0
0
0
0
5
0
0
3
3
3
1
0
0
0
0
1
6
1
0
0


24
0
0
0
0
0
0
1
0
0
0
0
0
0
5
0
0
3
3
3
1
0
0
0
0
1
6
1
0
0

0.97
0.97
0.97
0.97
0.97
0.97
0.97
0.97
0.97
0.97
0.97
0.97
0.97
0.97
0.97
0.97
0.97
0.97
0.97
0.97
0.97
0.97
0.97
0.97
0.97
0.97
0.97
0.97
0.97
0.97


11
0
0
0
0
0
0
0
0
0
0
0
0
0
1
0
0
3
3
3
1
0
0
0
0
0
0
0
0
0

1.0
1.0
1.0
1.0
1.0
1.0
1.0
1.0
1.0
1.0
1.0
1.0
1.0
1.0
1.0
1.0
1.0
1.0
1.0
1.0
1.0
1.0
1.0
1.0
1.0
1.0
1.0
1.0
1.0
1.0


8
0
0
0
0
0
0
0
0
0
0
0
0
0
0
0
0
0
0
0
0
0
0
0
0
1
6
1
0
0

1.0
1.0
1.0
1.0
1.0
1.0
1.0
1.0
1.0
1.0
1.0
1.0
1.0
1.0
1.0
1.0
1.0
1.0
1.0
1.0
1.0
1.0
1.0
1.0
1.0
1.0
1.0
1.0
1.0
1.0


1
0
0
0
0
0
1
0
0
0
0
0
0
0
0
0
0
0
0
0
0
0
0
0
0
0
0
0
0
0


1
0
0
0
0
0
1
0
0
0
0
0
0
0
0
0
0
0
0
0
0
0
0
0
0
0
0
0
0
0


1
0
0
0
0
0
1
0
0
0
0
0
0
0
0
0
0
0
0
0
0
0
0
0
0
0
0
0
0
0

1.0
1.0
1.0
1.0
1.0
1.0
1.0
1.0
1.0
1.0
1.0
1.0
1.0
1.0
1.0
1.0
1.0
1.0
1.0
1.0
1.0
1.0
1.0
1.0
1.0
1.0
1.0
1.0
1.0
1.0


15
0
0
0
0
0
0
0
0
0
0
0
0
0
15
0
0
0
0
0
0
0
0
0
0
0
0
0
0
0

1.0
1.0
1.0
1.0
1.0
1.0
1.0
1.0
1.0
1.0
1.0
1.0
1.0
1.0
1.0
1.0
1.0
1.0
1.0
1.0
1.0
1.0
1.0
1.0
1.0
1.0
1.0
1.0
1.0
1.0


7
0
0
0
0
0
0
0
0
0
0
0
0
0
7
0
0
0
0
0
0
0
0
0
0
0
0
0
0
0

0.88
0.88
0.88
0.88
0.88
0.88
0.88
0.88
0.88
0.88
0.88
0.88
0.88
0.88
0.88
0.88
0.88
0.88
0.88
0.88
0.88
0.88
0.88
0.88
0.88
0.88
0.88
0.88
0.88
0.88


1
0
0
0
0
0
0
0
0
0
0
0
0
0
1
0
0
0
0
0
0
0
0
0
0
0
0
0
0
0

0.84
0.84
0.84
0.84
0.84
0.84
0.84
0.84
0.84
0.84
0.84
0.84
0.84
0.84
0.84
0.84
0.84
0.84
0.84
0.84
0.84
0.84
0.84
0.84
0.84
0.84
0.84
0.84
0.84
0.84


10
0
0
0
0
0
0
0
0
0
0
0
0
0
0
1
0
0
0
0
0
0
0
0
0
0
8
0
1
0


10
0
0
0
0
0
0
0
0
0
0
0
0
0
0
1
0
0
0
0
0
0
0
0
0
0
8
0
1
0


8
0
0
0
0
0
0
0
0
0
0
0
0
0
0
0
0
0
0
0
0
0
0
0
0
0
8
0
0
0

1.0
1.0
1.0
1.0
1.0
1.0
1.0
1.0
1.0
1.0
1.0
1.0
1.0
1.0
1.0
1.0
1.0
1.0
1.0
1.0
1.0
1.0
1.0
1.0
1.0
1.0
1.0
1.0
1.0
1.0


2
0
0
0
0
0
0
0
0
0
0
0
0
0
0
1
0
0
0
0
0
0
0
0
0
0
0
0
1
0

1.0
1.0
1.0
1.0
1.0
1.0
1.0
1.0
1.0
1.0
1.0
1.0
1.0
1.0
1.0
1.0
1.0
1.0
1.0
1.0
1.0
1.0
1.0
1.0
1.0
1.0
1.0
1.0
1.0
1.0


3
0
0
0
0
0
0
0
0
0
0
0
0
0
1
0
0
1
1
0
0
0
0
0
0
0
0
0
0
0


2
0
0
0
0
0
0
0
0
0
0
0
0
0
0
0
0
1
1
0
0
0
0
0
0
0
0
0
0
0

1.0
1.0
1.0
1.0
1.0
1.0
1.0
1.0
1.0
1.0
1.0
1.0
1.0
1.0
1.0
1.0
1.0
1.0
1.0
1.0
1.0
1.0
1.0
1.0
1.0
1.0
1.0
1.0
1.0
1.0


1
0
0
0
0
0
0
0
0
0
0
0
0
0
1
0
0
0
0
0
0
0
0
0
0
0
0
0
0
0


1
0
0
0
0
0
0
0
0
0
0
0
0
0
1
0
0
0
0
0
0
0
0
0
0
0
0
0
0
0

1.0
1.0
1.0
1.0
1.0
1.0
1.0
1.0
1.0
1.0
1.0
1.0
1.0
1.0
1.0
1.0
1.0
1.0
1.0
1.0
1.0
1.0
1.0
1.0
1.0
1.0
1.0
1.0
1.0
1.0


2
0
0
0
0
0
0
0
0
0
0
0
0
0
0
0
0
2
0
0
0
0
0
0
0
0
0
0
0
0


2
0
0
0
0
0
0
0
0
0
0
0
0
0
0
0
0
2
0
0
0
0
0
0
0
0
0
0
0
0


2
0
0
0
0
0
0
0
0
0
0
0
0
0
0
0
0
2
0
0
0
0
0
0
0
0
0
0
0
0

1.0
1.0
1.0
1.0
1.0
1.0
1.0
1.0
1.0
1.0
1.0
1.0
1.0
1.0
1.0
1.0
1.0
1.0
1.0
1.0
1.0
1.0
1.0
1.0
1.0
1.0
1.0
1.0
1.0
1.0


5
0
0
0
0
0
0
0
0
0
0
0
0
2
3
0
0
0
0
0
0
0
0
0
0
0
0
0
0
0

0.71
0.71
0.71
0.71
0.71
0.71
0.71
0.71
0.71
0.71
0.71
0.71
0.71
0.71
0.71
0.71
0.71
0.71
0.71
0.71
0.71
0.71
0.71
0.71
0.71
0.71
0.71
0.71
0.71
0.71


3
0
0
0
0
0
0
0
0
0
0
0
0
1
2
0
0
0
0
0
0
0
0
0
0
0
0
0
0
0

0.99
0.99
0.99
0.99
0.99
0.99
0.99
0.99
0.99
0.99
0.99
0.99
0.99
0.99
0.99
0.99
0.99
0.99
0.99
0.99
0.99
0.99
0.99
0.99
0.99
0.99
0.99
0.99
0.99
0.99


1
0
0
0
0
0
0
0
0
0
0
0
0
0
0
0
0
0
0
0
0
0
0
0
0
0
0
0
1
0


1
0
0
0
0
0
0
0
0
0
0
0
0
0
0
0
0
0
0
0
0
0
0
0
0
0
0
0
1
0


1
0
0
0
0
0
0
0
0
0
0
0
0
0
0
0
0
0
0
0
0
0
0
0
0
0
0
0
1
0

1.0
1.0
1.0
1.0
1.0
1.0
1.0
1.0
1.0
1.0
1.0
1.0
1.0
1.0
1.0
1.0
1.0
1.0
1.0
1.0
1.0
1.0
1.0
1.0
1.0
1.0
1.0
1.0
1.0
1.0


1553
99
112
60
66
64
100
14
37
59
73
91
12
128
126
18
13
26
46
45
26
16
26
12
63
47
40
39
44
51

1.0
1.0
1.0
1.0
1.0
1.0
1.0
1.0
1.0
1.0
1.0
1.0
1.0
1.0
1.0
1.0
1.0
1.0
1.0
1.0
1.0
1.0
1.0
1.0
1.0
1.0
1.0
1.0
1.0
1.0


752
48
71
23
32
36
63
8
17
24
26
40
5
30
51
6
7
20
35
27
7
1
8
5
35
38
25
24
22
18

1.0
1.0
1.0
1.0
1.0
1.0
1.0
1.0
1.0
1.0
1.0
1.0
1.0
1.0
1.0
1.0
1.0
1.0
1.0
1.0
1.0
1.0
1.0
1.0
1.0
1.0
1.0
1.0
1.0
1.0


68
18
10
0
8
0
0
2
8
4
0
18
0
0
0
0
0
0
0
0
0
0
0
0
0
0
0
0
0
0


68
18
10
0
8
0
0
2
8
4
0
18
0
0
0
0
0
0
0
0
0
0
0
0
0
0
0
0
0
0

1.0
1.0
1.0
1.0
1.0
1.0
1.0
1.0
1.0
1.0
1.0
1.0
1.0
1.0
1.0
1.0
1.0
1.0
1.0
1.0
1.0
1.0
1.0
1.0
1.0
1.0
1.0
1.0
1.0
1.0


17
0
0
0
0
0
0
2
2
1
2
1
1
0
0
0
0
0
0
2
0
1
4
1
0
0
0
0
0
0


17
0
0
0
0
0
0
2
2
1
2
1
1
0
0
0
0
0
0
2
0
1
4
1
0
0
0
0
0
0

1.0
1.0
1.0
1.0
1.0
1.0
1.0
1.0
1.0
1.0
1.0
1.0
1.0
1.0
1.0
1.0
1.0
1.0
1.0
1.0
1.0
1.0
1.0
1.0
1.0
1.0
1.0
1.0
1.0
1.0


26
0
1
0
3
5
8
0
0
0
0
0
0
0
0
0
0
0
0
0
0
0
0
0
4
3
2
0
0
0

1.0
1.0
1.0
1.0
1.0
1.0
1.0
1.0
1.0
1.0
1.0
1.0
1.0
1.0
1.0
1.0
1.0
1.0
1.0
1.0
1.0
1.0
1.0
1.0
1.0
1.0
1.0
1.0
1.0
1.0


13
0
1
0
3
4
3
0
0
0
0
0
0
0
0
0
0
0
0
0
0
0
0
0
2
0
0
0
0
0

0.99
0.99
0.99
0.99
0.99
0.99
0.99
0.99
0.99
0.99
0.99
0.99
0.99
0.99
0.99
0.99
0.99
0.99
0.99
0.99
0.99
0.99
0.99
0.99
0.99
0.99
0.99
0.99
0.99
0.99


10
0
0
0
0
0
5
0
0
0
0
0
0
0
0
0
0
0
0
0
0
0
0
0
0
3
2
0
0
0

1.0
1.0
1.0
1.0
1.0
1.0
1.0
1.0
1.0
1.0
1.0
1.0
1.0
1.0
1.0
1.0
1.0
1.0
1.0
1.0
1.0
1.0
1.0
1.0
1.0
1.0
1.0
1.0
1.0
1.0


1
0
0
0
0
1
0
0
0
0
0
0
0
0
0
0
0
0
0
0
0
0
0
0
0
0
0
0
0
0

0.77
0.77
0.77
0.77
0.77
0.77
0.77
0.77
0.77
0.77
0.77
0.77
0.77
0.77
0.77
0.77
0.77
0.77
0.77
0.77
0.77
0.77
0.77
0.77
0.77
0.77
0.77
0.77
0.77
0.77


71
0
11
0
1
7
30
0
0
0
0
0
0
0
7
0
0
2
3
0
2
0
0
0
0
3
3
0
2
0

1.0
1.0
1.0
1.0
1.0
1.0
1.0
1.0
1.0
1.0
1.0
1.0
1.0
1.0
1.0
1.0
1.0
1.0
1.0
1.0
1.0
1.0
1.0
1.0
1.0
1.0
1.0
1.0
1.0
1.0


33
0
7
0
0
3
18
0
0
0
0
0
0
0
3
0
0
0
0
0
0
0
0
0
0
2
0
0
0
0

1.0
1.0
1.0
1.0
1.0
1.0
1.0
1.0
1.0
1.0
1.0
1.0
1.0
1.0
1.0
1.0
1.0
1.0
1.0
1.0
1.0
1.0
1.0
1.0
1.0
1.0
1.0
1.0
1.0
1.0


2
0
0
0
0
1
1
0
0
0
0
0
0
0
0
0
0
0
0
0
0
0
0
0
0
0
0
0
0
0

1.0
1.0
1.0
1.0
1.0
1.0
1.0
1.0
1.0
1.0
1.0
1.0
1.0
1.0
1.0
1.0
1.0
1.0
1.0
1.0
1.0
1.0
1.0
1.0
1.0
1.0
1.0
1.0
1.0
1.0


2
0
0
0
0
0
1
0
0
0
0
0
0
0
0
0
0
0
0
0
0
0
0
0
0
1
0
0
0
0

1.0
1.0
1.0
1.0
1.0
1.0
1.0
1.0
1.0
1.0
1.0
1.0
1.0
1.0
1.0
1.0
1.0
1.0
1.0
1.0
1.0
1.0
1.0
1.0
1.0
1.0
1.0
1.0
1.0
1.0


21
2
2
3
1
3
2
0
0
1
2
2
0
0
0
0
0
0
0
0
0
0
0
0
3
0
0
0
0
0


21
2
2
3
1
3
2
0
0
1
2
2
0
0
0
0
0
0
0
0
0
0
0
0
3
0
0
0
0
0

1.0
1.0
1.0
1.0
1.0
1.0
1.0
1.0
1.0
1.0
1.0
1.0
1.0
1.0
1.0
1.0
1.0
1.0
1.0
1.0
1.0
1.0
1.0
1.0
1.0
1.0
1.0
1.0
1.0
1.0


19
1
2
2
1
1
2
0
0
1
2
2
0
1
1
0
0
0
0
0
0
0
0
0
1
1
0
0
1
0


17
1
2
2
1
1
2
0
0
1
1
1
0
1
1
0
0
0
0
0
0
0
0
0
1
1
0
0
1
0

0.96
0.96
0.96
0.96
0.96
0.96
0.96
0.96
0.96
0.96
0.96
0.96
0.96
0.96
0.96
0.96
0.96
0.96
0.96
0.96
0.96
0.96
0.96
0.96
0.96
0.96
0.96
0.96
0.96
0.96


2
0
0
0
0
0
0
0
0
0
1
1
0
0
0
0
0
0
0
0
0
0
0
0
0
0
0
0
0
0

1.0
1.0
1.0
1.0
1.0
1.0
1.0
1.0
1.0
1.0
1.0
1.0
1.0
1.0
1.0
1.0
1.0
1.0
1.0
1.0
1.0
1.0
1.0
1.0
1.0
1.0
1.0
1.0
1.0
1.0


27
4
2
2
1
1
2
0
3
4
4
4
0
0
0
0
0
0
0
0
0
0
0
0
0
0
0
0
0
0

0.89
0.89
0.89
0.89
0.89
0.89
0.89
0.89
0.89
0.89
0.89
0.89
0.89
0.89
0.89
0.89
0.89
0.89
0.89
0.89
0.89
0.89
0.89
0.89
0.89
0.89
0.89
0.89
0.89
0.89


2
1
1
0
0
0
0
0
0
0
0
0
0
0
0
0
0
0
0
0
0
0
0
0
0
0
0
0
0
0

1.0
1.0
1.0
1.0
1.0
1.0
1.0
1.0
1.0
1.0
1.0
1.0
1.0
1.0
1.0
1.0
1.0
1.0
1.0
1.0
1.0
1.0
1.0
1.0
1.0
1.0
1.0
1.0
1.0
1.0


13
0
0
0
0
0
2
0
0
0
0
0
0
0
0
0
0
0
5
4
0
0
0
0
0
0
2
0
0
0


13
0
0
0
0
0
2
0
0
0
0
0
0
0
0
0
0
0
5
4
0
0
0
0
0
0
2
0
0
0

1.0
1.0
1.0
1.0
1.0
1.0
1.0
1.0
1.0
1.0
1.0
1.0
1.0
1.0
1.0
1.0
1.0
1.0
1.0
1.0
1.0
1.0
1.0
1.0
1.0
1.0
1.0
1.0
1.0
1.0


5
0
0
0
1
0
1
0
0
0
0
0
0
0
0
0
0
0
0
0
0
0
0
0
2
0
0
0
1
0


5
0
0
0
1
0
1
0
0
0
0
0
0
0
0
0
0
0
0
0
0
0
0
0
2
0
0
0
1
0

0.98
0.98
0.98
0.98
0.98
0.98
0.98
0.98
0.98
0.98
0.98
0.98
0.98
0.98
0.98
0.98
0.98
0.98
0.98
0.98
0.98
0.98
0.98
0.98
0.98
0.98
0.98
0.98
0.98
0.98


12
2
1
1
2
2
2
0
0
0
1
1
0
0
0
0
0
0
0
0
0
0
0
0
0
0
0
0
0
0


12
2
1
1
2
2
2
0
0
0
1
1
0
0
0
0
0
0
0
0
0
0
0
0
0
0
0
0
0
0

1.0
1.0
1.0
1.0
1.0
1.0
1.0
1.0
1.0
1.0
1.0
1.0
1.0
1.0
1.0
1.0
1.0
1.0
1.0
1.0
1.0
1.0
1.0
1.0
1.0
1.0
1.0
1.0
1.0
1.0


52
6
4
0
1
0
0
0
0
0
1
1
4
2
7
0
0
4
5
4
3
0
4
4
2
0
0
0
0
0

1.0
1.0
1.0
1.0
1.0
1.0
1.0
1.0
1.0
1.0
1.0
1.0
1.0
1.0
1.0
1.0
1.0
1.0
1.0
1.0
1.0
1.0
1.0
1.0
1.0
1.0
1.0
1.0
1.0
1.0


2
1
0
0
0
0
0
0
0
0
1
0
0
0
0
0
0
0
0
0
0
0
0
0
0
0
0
0
0
0

0.81
0.81
0.81
0.81
0.81
0.81
0.81
0.81
0.81
0.81
0.81
0.81
0.81
0.81
0.81
0.81
0.81
0.81
0.81
0.81
0.81
0.81
0.81
0.81
0.81
0.81
0.81
0.81
0.81
0.81


1
1
0
0
0
0
0
0
0
0
0
0
0
0
0
0
0
0
0
0
0
0
0
0
0
0
0
0
0
0

1.0
1.0
1.0
1.0
1.0
1.0
1.0
1.0
1.0
1.0
1.0
1.0
1.0
1.0
1.0
1.0
1.0
1.0
1.0
1.0
1.0
1.0
1.0
1.0
1.0
1.0
1.0
1.0
1.0
1.0


18
1
0
0
0
0
0
0
0
0
0
0
2
1
3
0
0
1
2
2
1
0
2
2
1
0
0
0
0
0

1.0
1.0
1.0
1.0
1.0
1.0
1.0
1.0
1.0
1.0
1.0
1.0
1.0
1.0
1.0
1.0
1.0
1.0
1.0
1.0
1.0
1.0
1.0
1.0
1.0
1.0
1.0
1.0
1.0
1.0


15
1
3
3
2
1
3
0
0
0
0
0
0
0
0
0
0
0
0
0
0
0
0
0
1
0
0
1
0
0


15
1
3
3
2
1
3
0
0
0
0
0
0
0
0
0
0
0
0
0
0
0
0
0
1
0
0
1
0
0

1.0
1.0
1.0
1.0
1.0
1.0
1.0
1.0
1.0
1.0
1.0
1.0
1.0
1.0
1.0
1.0
1.0
1.0
1.0
1.0
1.0
1.0
1.0
1.0
1.0
1.0
1.0
1.0
1.0
1.0


22
0
5
2
1
7
2
0
0
1
0
0
0
0
2
0
0
0
0
0
0
0
0
0
1
1
0
0
0
0

0.87
0.87
0.87
0.87
0.87
0.87
0.87
0.87
0.87
0.87
0.87
0.87
0.87
0.87
0.87
0.87
0.87
0.87
0.87
0.87
0.87
0.87
0.87
0.87
0.87
0.87
0.87
0.87
0.87
0.87


2
0
1
0
0
1
0
0
0
0
0
0
0
0
0
0
0
0
0
0
0
0
0
0
0
0
0
0
0
0

1.0
1.0
1.0
1.0
1.0
1.0
1.0
1.0
1.0
1.0
1.0
1.0
1.0
1.0
1.0
1.0
1.0
1.0
1.0
1.0
1.0
1.0
1.0
1.0
1.0
1.0
1.0
1.0
1.0
1.0


6
0
2
0
0
4
0
0
0
0
0
0
0
0
0
0
0
0
0
0
0
0
0
0
0
0
0
0
0
0

0.83
0.83
0.83
0.83
0.83
0.83
0.83
0.83
0.83
0.83
0.83
0.83
0.83
0.83
0.83
0.83
0.83
0.83
0.83
0.83
0.83
0.83
0.83
0.83
0.83
0.83
0.83
0.83
0.83
0.83


17
3
4
1
1
1
1
0
1
2
1
1
0
0
0
0
0
0
0
0
0
0
0
0
1
0
0
0
0
0


12
2
2
1
1
1
1
0
0
2
0
1
0
0
0
0
0
0
0
0
0
0
0
0
1
0
0
0
0
0

1.0
1.0
1.0
1.0
1.0
1.0
1.0
1.0
1.0
1.0
1.0
1.0
1.0
1.0
1.0
1.0
1.0
1.0
1.0
1.0
1.0
1.0
1.0
1.0
1.0
1.0
1.0
1.0
1.0
1.0


5
1
2
0
0
0
0
0
1
0
1
0
0
0
0
0
0
0
0
0
0
0
0
0
0
0
0
0
0
0

1.0
1.0
1.0
1.0
1.0
1.0
1.0
1.0
1.0
1.0
1.0
1.0
1.0
1.0
1.0
1.0
1.0
1.0
1.0
1.0
1.0
1.0
1.0
1.0
1.0
1.0
1.0
1.0
1.0
1.0


6
2
0
0
0
0
0
0
0
2
2
0
0
0
0
0
0
0
0
0
0
0
0
0
0
0
0
0
0
0

1.0
1.0
1.0
1.0
1.0
1.0
1.0
1.0
1.0
1.0
1.0
1.0
1.0
1.0
1.0
1.0
1.0
1.0
1.0
1.0
1.0
1.0
1.0
1.0
1.0
1.0
1.0
1.0
1.0
1.0


4
1
0
0
0
0
0
0
0
1
2
0
0
0
0
0
0
0
0
0
0
0
0
0
0
0
0
0
0
0

1.0
1.0
1.0
1.0
1.0
1.0
1.0
1.0
1.0
1.0
1.0
1.0
1.0
1.0
1.0
1.0
1.0
1.0
1.0
1.0
1.0
1.0
1.0
1.0
1.0
1.0
1.0
1.0
1.0
1.0


17
0
12
0
0
0
0
0
0
0
0
0
0
0
0
0
0
0
0
0
0
0
0
0
5
0
0
0
0
0

1.0
1.0
1.0
1.0
1.0
1.0
1.0
1.0
1.0
1.0
1.0
1.0
1.0
1.0
1.0
1.0
1.0
1.0
1.0
1.0
1.0
1.0
1.0
1.0
1.0
1.0
1.0
1.0
1.0
1.0


2
0
1
0
0
0
0
1
0
0
0
0
0
0
0
0
0
0
0
0
0
0
0
0
0
0
0
0
0
0


2
0
1
0
0
0
0
1
0
0
0
0
0
0
0
0
0
0
0
0
0
0
0
0
0
0
0
0
0
0

1.0
1.0
1.0
1.0
1.0
1.0
1.0
1.0
1.0
1.0
1.0
1.0
1.0
1.0
1.0
1.0
1.0
1.0
1.0
1.0
1.0
1.0
1.0
1.0
1.0
1.0
1.0
1.0
1.0
1.0


2
0
0
0
1
0
0
0
0
0
0
1
0
0
0
0
0
0
0
0
0
0
0
0
0
0
0
0
0
0


2
0
0
0
1
0
0
0
0
0
0
1
0
0
0
0
0
0
0
0
0
0
0
0
0
0
0
0
0
0

1.0
1.0
1.0
1.0
1.0
1.0
1.0
1.0
1.0
1.0
1.0
1.0
1.0
1.0
1.0
1.0
1.0
1.0
1.0
1.0
1.0
1.0
1.0
1.0
1.0
1.0
1.0
1.0
1.0
1.0


11
2
0
0
0
0
0
0
0
2
3
1
0
0
1
0
0
0
1
1
0
0
0
0
0
0
0
0
0
0


11
2
0
0
0
0
0
0
0
2
3
1
0
0
1
0
0
0
1
1
0
0
0
0
0
0
0
0
0
0

1.0
1.0
1.0
1.0
1.0
1.0
1.0
1.0
1.0
1.0
1.0
1.0
1.0
1.0
1.0
1.0
1.0
1.0
1.0
1.0
1.0
1.0
1.0
1.0
1.0
1.0
1.0
1.0
1.0
1.0


9
1
1
1
1
1
0
0
0
0
0
2
0
0
0
0
0
0
0
0
0
0
0
0
0
0
0
0
0
2

1.0
1.0
1.0
1.0
1.0
1.0
1.0
1.0
1.0
1.0
1.0
1.0
1.0
1.0
1.0
1.0
1.0
1.0
1.0
1.0
1.0
1.0
1.0
1.0
1.0
1.0
1.0
1.0
1.0
1.0


5
1
1
0
1
1
0
0
0
0
0
1
0
0
0
0
0
0
0
0
0
0
0
0
0
0
0
0
0
0

1.0
1.0
1.0
1.0
1.0
1.0
1.0
1.0
1.0
1.0
1.0
1.0
1.0
1.0
1.0
1.0
1.0
1.0
1.0
1.0
1.0
1.0
1.0
1.0
1.0
1.0
1.0
1.0
1.0
1.0


2
0
0
1
0
0
0
0
0
0
0
1
0
0
0
0
0
0
0
0
0
0
0
0
0
0
0
0
0
0

0.9
0.9
0.9
0.9
0.9
0.9
0.9
0.9
0.9
0.9
0.9
0.9
0.9
0.9
0.9
0.9
0.9
0.9
0.9
0.9
0.9
0.9
0.9
0.9
0.9
0.9
0.9
0.9
0.9
0.9


2
2
0
0
0
0
0
0
0
0
0
0
0
0
0
0
0
0
0
0
0
0
0
0
0
0
0
0
0
0

1.0
1.0
1.0
1.0
1.0
1.0
1.0
1.0
1.0
1.0
1.0
1.0
1.0
1.0
1.0
1.0
1.0
1.0
1.0
1.0
1.0
1.0
1.0
1.0
1.0
1.0
1.0
1.0
1.0
1.0


109
0
1
0
0
0
1
0
0
0
1
1
0
0
0
0
0
13
10
10
0
0
0
0
0
18
18
14
10
12

1.0
1.0
1.0
1.0
1.0
1.0
1.0
1.0
1.0
1.0
1.0
1.0
1.0
1.0
1.0
1.0
1.0
1.0
1.0
1.0
1.0
1.0
1.0
1.0
1.0
1.0
1.0
1.0
1.0
1.0


13
0
0
0
0
0
0
0
0
0
0
0
0
0
0
0
0
1
1
2
0
0
0
0
0
3
3
2
1
0

1.0
1.0
1.0
1.0
1.0
1.0
1.0
1.0
1.0
1.0
1.0
1.0
1.0
1.0
1.0
1.0
1.0
1.0
1.0
1.0
1.0
1.0
1.0
1.0
1.0
1.0
1.0
1.0
1.0
1.0


18
0
0
0
0
0
0
0
0
0
0
0
0
0
0
0
0
3
1
2
0
0
0
0
0
1
3
2
2
4

1.0
1.0
1.0
1.0
1.0
1.0
1.0
1.0
1.0
1.0
1.0
1.0
1.0
1.0
1.0
1.0
1.0
1.0
1.0
1.0
1.0
1.0
1.0
1.0
1.0
1.0
1.0
1.0
1.0
1.0


4
0
0
0
0
0
1
0
0
0
0
0
0
0
0
0
0
0
3
0
0
0
0
0
0
0
0
0
0
0

1.0
1.0
1.0
1.0
1.0
1.0
1.0
1.0
1.0
1.0
1.0
1.0
1.0
1.0
1.0
1.0
1.0
1.0
1.0
1.0
1.0
1.0
1.0
1.0
1.0
1.0
1.0
1.0
1.0
1.0


3
0
0
0
0
0
1
0
0
0
0
0
0
0
0
0
0
0
2
0
0
0
0
0
0
0
0
0
0
0

1.0
1.0
1.0
1.0
1.0
1.0
1.0
1.0
1.0
1.0
1.0
1.0
1.0
1.0
1.0
1.0
1.0
1.0
1.0
1.0
1.0
1.0
1.0
1.0
1.0
1.0
1.0
1.0
1.0
1.0


4
0
1
0
0
0
0
0
0
0
0
0
0
0
0
0
0
0
0
0
0
0
0
0
0
0
0
0
0
3


4
0
1
0
0
0
0
0
0
0
0
0
0
0
0
0
0
0
0
0
0
0
0
0
0
0
0
0
0
3

1.0
1.0
1.0
1.0
1.0
1.0
1.0
1.0
1.0
1.0
1.0
1.0
1.0
1.0
1.0
1.0
1.0
1.0
1.0
1.0
1.0
1.0
1.0
1.0
1.0
1.0
1.0
1.0
1.0
1.0


8
0
0
0
0
1
0
0
0
0
0
0
0
0
0
0
0
0
4
0
0
0
0
0
3
0
0
0
0
0


8
0
0
0
0
1
0
0
0
0
0
0
0
0
0
0
0
0
4
0
0
0
0
0
3
0
0
0
0
0

1.0
1.0
1.0
1.0
1.0
1.0
1.0
1.0
1.0
1.0
1.0
1.0
1.0
1.0
1.0
1.0
1.0
1.0
1.0
1.0
1.0
1.0
1.0
1.0
1.0
1.0
1.0
1.0
1.0
1.0


4
0
2
2
0
0
0
0
0
0
0
0
0
0
0
0
0
0
0
0
0
0
0
0
0
0
0
0
0
0

0.81
0.81
0.81
0.81
0.81
0.81
0.81
0.81
0.81
0.81
0.81
0.81
0.81
0.81
0.81
0.81
0.81
0.81
0.81
0.81
0.81
0.81
0.81
0.81
0.81
0.81
0.81
0.81
0.81
0.81


13
0
1
1
0
1
1
0
0
1
0
0
0
0
0
0
0
1
0
4
2
0
0
0
1
0
0
0
0
0

1.0
1.0
1.0
1.0
1.0
1.0
1.0
1.0
1.0
1.0
1.0
1.0
1.0
1.0
1.0
1.0
1.0
1.0
1.0
1.0
1.0
1.0
1.0
1.0
1.0
1.0
1.0
1.0
1.0
1.0


5
0
1
1
0
1
1
0
0
1
0
0
0
0
0
0
0
0
0
0
0
0
0
0
0
0
0
0
0
0

1.0
1.0
1.0
1.0
1.0
1.0
1.0
1.0
1.0
1.0
1.0
1.0
1.0
1.0
1.0
1.0
1.0
1.0
1.0
1.0
1.0
1.0
1.0
1.0
1.0
1.0
1.0
1.0
1.0
1.0


13
1
0
0
0
0
0
0
0
0
0
0
0
0
0
0
0
0
3
0
0
0
0
0
0
9
0
0
0
0


13
1
0
0
0
0
0
0
0
0
0
0
0
0
0
0
0
0
3
0
0
0
0
0
0
9
0
0
0
0

1.0
1.0
1.0
1.0
1.0
1.0
1.0
1.0
1.0
1.0
1.0
1.0
1.0
1.0
1.0
1.0
1.0
1.0
1.0
1.0
1.0
1.0
1.0
1.0
1.0
1.0
1.0
1.0
1.0
1.0


29
0
1
1
0
0
0
0
0
0
1
0
0
13
13
0
0
0
0
0
0
0
0
0
0
0
0
0
0
0

0.97
0.97
0.97
0.97
0.97
0.97
0.97
0.97
0.97
0.97
0.97
0.97
0.97
0.97
0.97
0.97
0.97
0.97
0.97
0.97
0.97
0.97
0.97
0.97
0.97
0.97
0.97
0.97
0.97
0.97


3
0
0
0
0
0
0
0
0
0
0
0
0
2
1
0
0
0
0
0
0
0
0
0
0
0
0
0
0
0

0.95
0.95
0.95
0.95
0.95
0.95
0.95
0.95
0.95
0.95
0.95
0.95
0.95
0.95
0.95
0.95
0.95
0.95
0.95
0.95
0.95
0.95
0.95
0.95
0.95
0.95
0.95
0.95
0.95
0.95


28
0
0
0
0
0
0
0
0
0
0
0
0
0
0
6
7
0
0
0
0
0
0
0
0
0
0
9
6
0


28
0
0
0
0
0
0
0
0
0
0
0
0
0
0
6
7
0
0
0
0
0
0
0
0
0
0
9
6
0

1.0
1.0
1.0
1.0
1.0
1.0
1.0
1.0
1.0
1.0
1.0
1.0
1.0
1.0
1.0
1.0
1.0
1.0
1.0
1.0
1.0
1.0
1.0
1.0
1.0
1.0
1.0
1.0
1.0
1.0


13
0
0
0
0
0
0
0
0
0
0
0
0
0
7
0
0
0
0
0
0
0
0
0
6
0
0
0
0
0


13
0
0
0
0
0
0
0
0
0
0
0
0
0
7
0
0
0
0
0
0
0
0
0
6
0
0
0
0
0

1.0
1.0
1.0
1.0
1.0
1.0
1.0
1.0
1.0
1.0
1.0
1.0
1.0
1.0
1.0
1.0
1.0
1.0
1.0
1.0
1.0
1.0
1.0
1.0
1.0
1.0
1.0
1.0
1.0
1.0


2
0
0
0
0
0
0
0
0
0
0
0
0
0
0
0
0
0
0
2
0
0
0
0
0
0
0
0
0
0

0.83
0.83
0.83
0.83
0.83
0.83
0.83
0.83
0.83
0.83
0.83
0.83
0.83
0.83
0.83
0.83
0.83
0.83
0.83
0.83
0.83
0.83
0.83
0.83
0.83
0.83
0.83
0.83
0.83
0.83


2
0
0
0
0
0
0
0
0
0
0
0
0
2
0
0
0
0
0
0
0
0
0
0
0
0
0
0
0
0

0.86
0.86
0.86
0.86
0.86
0.86
0.86
0.86
0.86
0.86
0.86
0.86
0.86
0.86
0.86
0.86
0.86
0.86
0.86
0.86
0.86
0.86
0.86
0.86
0.86
0.86
0.86
0.86
0.86
0.86


35
1
4
5
6
3
3
0
0
1
4
3
0
0
0
0
0
0
0
0
0
0
0
0
4
0
0
0
1
0

1.0
1.0
1.0
1.0
1.0
1.0
1.0
1.0
1.0
1.0
1.0
1.0
1.0
1.0
1.0
1.0
1.0
1.0
1.0
1.0
1.0
1.0
1.0
1.0
1.0
1.0
1.0
1.0
1.0
1.0


16
1
3
3
2
0
0
0
0
0
2
2
0
0
0
0
0
0
0
0
0
0
0
0
2
0
0
0
1
0


16
1
3
3
2
0
0
0
0
0
2
2
0
0
0
0
0
0
0
0
0
0
0
0
2
0
0
0
1
0

1.0
1.0
1.0
1.0
1.0
1.0
1.0
1.0
1.0
1.0
1.0
1.0
1.0
1.0
1.0
1.0
1.0
1.0
1.0
1.0
1.0
1.0
1.0
1.0
1.0
1.0
1.0
1.0
1.0
1.0


1
0
0
0
0
0
1
0
0
0
0
0
0
0
0
0
0
0
0
0
0
0
0
0
0
0
0
0
0
0


1
0
0
0
0
0
1
0
0
0
0
0
0
0
0
0
0
0
0
0
0
0
0
0
0
0
0
0
0
0

1.0
1.0
1.0
1.0
1.0
1.0
1.0
1.0
1.0
1.0
1.0
1.0
1.0
1.0
1.0
1.0
1.0
1.0
1.0
1.0
1.0
1.0
1.0
1.0
1.0
1.0
1.0
1.0
1.0
1.0


3
0
0
0
0
1
0
0
0
0
2
0
0
0
0
0
0
0
0
0
0
0
0
0
0
0
0
0
0
0

1.0
1.0
1.0
1.0
1.0
1.0
1.0
1.0
1.0
1.0
1.0
1.0
1.0
1.0
1.0
1.0
1.0
1.0
1.0
1.0
1.0
1.0
1.0
1.0
1.0
1.0
1.0
1.0
1.0
1.0


176
12
15
14
8
4
6
0
7
14
18
16
1
20
7
6
0
2
5
2
2
0
1
1
3
0
2
3
2
5

0.99
0.99
0.99
0.99
0.99
0.99
0.99
0.99
0.99
0.99
0.99
0.99
0.99
0.99
0.99
0.99
0.99
0.99
0.99
0.99
0.99
0.99
0.99
0.99
0.99
0.99
0.99
0.99
0.99
0.99


13
0
0
0
0
0
0
0
0
0
6
7
0
0
0
0
0
0
0
0
0
0
0
0
0
0
0
0
0
0


13
0
0
0
0
0
0
0
0
0
6
7
0
0
0
0
0
0
0
0
0
0
0
0
0
0
0
0
0
0

1.0
1.0
1.0
1.0
1.0
1.0
1.0
1.0
1.0
1.0
1.0
1.0
1.0
1.0
1.0
1.0
1.0
1.0
1.0
1.0
1.0
1.0
1.0
1.0
1.0
1.0
1.0
1.0
1.0
1.0


19
2
0
0
0
0
0
0
6
10
1
0
0
0
0
0
0
0
0
0
0
0
0
0
0
0
0
0
0
0


19
2
0
0
0
0
0
0
6
10
1
0
0
0
0
0
0
0
0
0
0
0
0
0
0
0
0
0
0
0

1.0
1.0
1.0
1.0
1.0
1.0
1.0
1.0
1.0
1.0
1.0
1.0
1.0
1.0
1.0
1.0
1.0
1.0
1.0
1.0
1.0
1.0
1.0
1.0
1.0
1.0
1.0
1.0
1.0
1.0


27
0
6
6
5
0
1
0
0
0
0
0
0
9
0
0
0
0
0
0
0
0
0
0
0
0
0
0
0
0

0.74
0.74
0.74
0.74
0.74
0.74
0.74
0.74
0.74
0.74
0.74
0.74
0.74
0.74
0.74
0.74
0.74
0.74
0.74
0.74
0.74
0.74
0.74
0.74
0.74
0.74
0.74
0.74
0.74
0.74


11
0
4
4
3
0
0
0
0
0
0
0
0
0
0
0
0
0
0
0
0
0
0
0
0
0
0
0
0
0

1.0
1.0
1.0
1.0
1.0
1.0
1.0
1.0
1.0
1.0
1.0
1.0
1.0
1.0
1.0
1.0
1.0
1.0
1.0
1.0
1.0
1.0
1.0
1.0
1.0
1.0
1.0
1.0
1.0
1.0


13
0
2
2
2
0
0
0
0
0
0
0
0
7
0
0
0
0
0
0
0
0
0
0
0
0
0
0
0
0

1.0
1.0
1.0
1.0
1.0
1.0
1.0
1.0
1.0
1.0
1.0
1.0
1.0
1.0
1.0
1.0
1.0
1.0
1.0
1.0
1.0
1.0
1.0
1.0
1.0
1.0
1.0
1.0
1.0
1.0


1
0
0
0
0
0
1
0
0
0
0
0
0
0
0
0
0
0
0
0
0
0
0
0
0
0
0
0
0
0

1.0
1.0
1.0
1.0
1.0
1.0
1.0
1.0
1.0
1.0
1.0
1.0
1.0
1.0
1.0
1.0
1.0
1.0
1.0
1.0
1.0
1.0
1.0
1.0
1.0
1.0
1.0
1.0
1.0
1.0


18
5
2
0
0
0
0
0
0
2
7
2
0
0
0
0
0
0
0
0
0
0
0
0
0
0
0
0
0
0


9
3
1
0
0
0
0
0
0
0
4
1
0
0
0
0
0
0
0
0
0
0
0
0
0
0
0
0
0
0

1.0
1.0
1.0
1.0
1.0
1.0
1.0
1.0
1.0
1.0
1.0
1.0
1.0
1.0
1.0
1.0
1.0
1.0
1.0
1.0
1.0
1.0
1.0
1.0
1.0
1.0
1.0
1.0
1.0
1.0


3
1
1
0
0
0
0
0
0
0
0
1
0
0
0
0
0
0
0
0
0
0
0
0
0
0
0
0
0
0

1.0
1.0
1.0
1.0
1.0
1.0
1.0
1.0
1.0
1.0
1.0
1.0
1.0
1.0
1.0
1.0
1.0
1.0
1.0
1.0
1.0
1.0
1.0
1.0
1.0
1.0
1.0
1.0
1.0
1.0


6
1
0
0
0
0
0
0
0
2
3
0
0
0
0
0
0
0
0
0
0
0
0
0
0
0
0
0
0
0

1.0
1.0
1.0
1.0
1.0
1.0
1.0
1.0
1.0
1.0
1.0
1.0
1.0
1.0
1.0
1.0
1.0
1.0
1.0
1.0
1.0
1.0
1.0
1.0
1.0
1.0
1.0
1.0
1.0
1.0


6
1
1
0
0
0
0
0
0
0
1
1
0
0
0
0
0
0
0
0
0
0
0
0
0
0
1
0
0
1

0.77
0.77
0.77
0.77
0.77
0.77
0.77
0.77
0.77
0.77
0.77
0.77
0.77
0.77
0.77
0.77
0.77
0.77
0.77
0.77
0.77
0.77
0.77
0.77
0.77
0.77
0.77
0.77
0.77
0.77


4
1
1
0
0
0
0
0
0
0
1
1
0
0
0
0
0
0
0
0
0
0
0
0
0
0
0
0
0
0

1.0
1.0
1.0
1.0
1.0
1.0
1.0
1.0
1.0
1.0
1.0
1.0
1.0
1.0
1.0
1.0
1.0
1.0
1.0
1.0
1.0
1.0
1.0
1.0
1.0
1.0
1.0
1.0
1.0
1.0


23
0
0
1
1
2
2
0
1
0
0
0
0
0
1
4
0
1
4
1
2
0
0
0
0
0
0
0
0
3

1.0
1.0
1.0
1.0
1.0
1.0
1.0
1.0
1.0
1.0
1.0
1.0
1.0
1.0
1.0
1.0
1.0
1.0
1.0
1.0
1.0
1.0
1.0
1.0
1.0
1.0
1.0
1.0
1.0
1.0


2
0
0
0
0
0
0
0
0
0
1
1
0
0
0
0
0
0
0
0
0
0
0
0
0
0
0
0
0
0

1.0
1.0
1.0
1.0
1.0
1.0
1.0
1.0
1.0
1.0
1.0
1.0
1.0
1.0
1.0
1.0
1.0
1.0
1.0
1.0
1.0
1.0
1.0
1.0
1.0
1.0
1.0
1.0
1.0
1.0


2
0
1
0
0
0
0
0
0
0
0
1
0
0
0
0
0
0
0
0
0
0
0
0
0
0
0
0
0
0


2
0
1
0
0
0
0
0
0
0
0
1
0
0
0
0
0
0
0
0
0
0
0
0
0
0
0
0
0
0

1.0
1.0
1.0
1.0
1.0
1.0
1.0
1.0
1.0
1.0
1.0
1.0
1.0
1.0
1.0
1.0
1.0
1.0
1.0
1.0
1.0
1.0
1.0
1.0
1.0
1.0
1.0
1.0
1.0
1.0


2
0
0
0
0
0
0
0
0
1
0
1
0
0
0
0
0
0
0
0
0
0
0
0
0
0
0
0
0
0


2
0
0
0
0
0
0
0
0
1
0
1
0
0
0
0
0
0
0
0
0
0
0
0
0
0
0
0
0
0

1.0
1.0
1.0
1.0
1.0
1.0
1.0
1.0
1.0
1.0
1.0
1.0
1.0
1.0
1.0
1.0
1.0
1.0
1.0
1.0
1.0
1.0
1.0
1.0
1.0
1.0
1.0
1.0
1.0
1.0


1
0
0
0
0
0
0
0
0
0
1
0
0
0
0
0
0
0
0
0
0
0
0
0
0
0
0
0
0
0


1
0
0
0
0
0
0
0
0
0
1
0
0
0
0
0
0
0
0
0
0
0
0
0
0
0
0
0
0
0

1.0
1.0
1.0
1.0
1.0
1.0
1.0
1.0
1.0
1.0
1.0
1.0
1.0
1.0
1.0
1.0
1.0
1.0
1.0
1.0
1.0
1.0
1.0
1.0
1.0
1.0
1.0
1.0
1.0
1.0


5
1
1
1
1
1
0
0
0
0
0
0
0
0
0
0
0
0
0
0
0
0
0
0
0
0
0
0
0
0

1.0
1.0
1.0
1.0
1.0
1.0
1.0
1.0
1.0
1.0
1.0
1.0
1.0
1.0
1.0
1.0
1.0
1.0
1.0
1.0
1.0
1.0
1.0
1.0
1.0
1.0
1.0
1.0
1.0
1.0


3
0
1
1
0
0
0
0
0
0
0
0
0
1
0
0
0
0
0
0
0
0
0
0
0
0
0
0
0
0


3
0
1
1
0
0
0
0
0
0
0
0
0
1
0
0
0
0
0
0
0
0
0
0
0
0
0
0
0
0

0.99
0.99
0.99
0.99
0.99
0.99
0.99
0.99
0.99
0.99
0.99
0.99
0.99
0.99
0.99
0.99
0.99
0.99
0.99
0.99
0.99
0.99
0.99
0.99
0.99
0.99
0.99
0.99
0.99
0.99


9
1
2
2
0
0
0
0
0
0
0
0
0
3
1
0
0
0
0
0
0
0
0
0
0
0
0
0
0
0


9
1
2
2
0
0
0
0
0
0
0
0
0
3
1
0
0
0
0
0
0
0
0
0
0
0
0
0
0
0

1.0
1.0
1.0
1.0
1.0
1.0
1.0
1.0
1.0
1.0
1.0
1.0
1.0
1.0
1.0
1.0
1.0
1.0
1.0
1.0
1.0
1.0
1.0
1.0
1.0
1.0
1.0
1.0
1.0
1.0


2
1
1
0
0
0
0
0
0
0
0
0
0
0
0
0
0
0
0
0
0
0
0
0
0
0
0
0
0
0

0.95
0.95
0.95
0.95
0.95
0.95
0.95
0.95
0.95
0.95
0.95
0.95
0.95
0.95
0.95
0.95
0.95
0.95
0.95
0.95
0.95
0.95
0.95
0.95
0.95
0.95
0.95
0.95
0.95
0.95


4
0
0
1
0
0
1
0
0
0
0
0
0
0
0
0
0
0
0
0
0
0
0
0
0
0
0
2
0
0


4
0
0
1
0
0
1
0
0
0
0
0
0
0
0
0
0
0
0
0
0
0
0
0
0
0
0
2
0
0

1.0
1.0
1.0
1.0
1.0
1.0
1.0
1.0
1.0
1.0
1.0
1.0
1.0
1.0
1.0
1.0
1.0
1.0
1.0
1.0
1.0
1.0
1.0
1.0
1.0
1.0
1.0
1.0
1.0
1.0


1
0
0
1
0
0
0
0
0
0
0
0
0
0
0
0
0
0
0
0
0
0
0
0
0
0
0
0
0
0


1
0
0
1
0
0
0
0
0
0
0
0
0
0
0
0
0
0
0
0
0
0
0
0
0
0
0
0
0
0

1.0
1.0
1.0
1.0
1.0
1.0
1.0
1.0
1.0
1.0
1.0
1.0
1.0
1.0
1.0
1.0
1.0
1.0
1.0
1.0
1.0
1.0
1.0
1.0
1.0
1.0
1.0
1.0
1.0
1.0


4
0
0
0
0
0
0
0
0
0
0
0
1
0
0
0
0
0
0
1
0
0
1
1
0
0
0
0
0
0

0.93
0.93
0.93
0.93
0.93
0.93
0.93
0.93
0.93
0.93
0.93
0.93
0.93
0.93
0.93
0.93
0.93
0.93
0.93
0.93
0.93
0.93
0.93
0.93
0.93
0.93
0.93
0.93
0.93
0.93


3
0
0
0
0
0
0
0
0
0
0
0
0
0
0
0
0
0
0
0
0
0
0
0
0
0
1
1
0
1


3
0
0
0
0
0
0
0
0
0
0
0
0
0
0
0
0
0
0
0
0
0
0
0
0
0
1
1
0
1

1.0
1.0
1.0
1.0
1.0
1.0
1.0
1.0
1.0
1.0
1.0
1.0
1.0
1.0
1.0
1.0
1.0
1.0
1.0
1.0
1.0
1.0
1.0
1.0
1.0
1.0
1.0
1.0
1.0
1.0


7
0
0
0
0
0
0
0
0
0
0
0
0
3
2
0
0
0
0
0
0
0
0
0
0
0
0
0
2
0


5
0
0
0
0
0
0
0
0
0
0
0
0
3
2
0
0
0
0
0
0
0
0
0
0
0
0
0
0
0

0.99
0.99
0.99
0.99
0.99
0.99
0.99
0.99
0.99
0.99
0.99
0.99
0.99
0.99
0.99
0.99
0.99
0.99
0.99
0.99
0.99
0.99
0.99
0.99
0.99
0.99
0.99
0.99
0.99
0.99


2
0
0
0
0
0
0
0
0
0
0
0
0
0
0
0
0
0
0
0
0
0
0
0
0
0
0
0
2
0

1.0
1.0
1.0
1.0
1.0
1.0
1.0
1.0
1.0
1.0
1.0
1.0
1.0
1.0
1.0
1.0
1.0
1.0
1.0
1.0
1.0
1.0
1.0
1.0
1.0
1.0
1.0
1.0
1.0
1.0


2
0
0
0
0
0
0
0
0
0
0
0
0
1
0
0
0
0
0
0
0
0
0
0
1
0
0
0
0
0


2
0
0
0
0
0
0
0
0
0
0
0
0
1
0
0
0
0
0
0
0
0
0
0
1
0
0
0
0
0

1.0
1.0
1.0
1.0
1.0
1.0
1.0
1.0
1.0
1.0
1.0
1.0
1.0
1.0
1.0
1.0
1.0
1.0
1.0
1.0
1.0
1.0
1.0
1.0
1.0
1.0
1.0
1.0
1.0
1.0


28
1
1
1
1
1
1
2
2
0
1
1
1
1
1
1
1
1
0
1
1
1
1
1
1
0
1
1
1
1


28
1
1
1
1
1
1
2
2
0
1
1
1
1
1
1
1
1
0
1
1
1
1
1
1
0
1
1
1
1


28
1
1
1
1
1
1
2
2
0
1
1
1
1
1
1
1
1
0
1
1
1
1
1
1
0
1
1
1
1

1.0
1.0
1.0
1.0
1.0
1.0
1.0
1.0
1.0
1.0
1.0
1.0
1.0
1.0
1.0
1.0
1.0
1.0
1.0
1.0
1.0
1.0
1.0
1.0
1.0
1.0
1.0
1.0
1.0
1.0


18
2
2
1
0
1
0
1
0
0
0
1
0
1
1
0
0
0
0
2
0
0
0
0
0
0
1
1
3
1


3
1
1
0
0
1
0
0
0
0
0
0
0
0
0
0
0
0
0
0
0
0
0
0
0
0
0
0
0
0


3
1
1
0
0
1
0
0
0
0
0
0
0
0
0
0
0
0
0
0
0
0
0
0
0
0
0
0
0
0

1.0
1.0
1.0
1.0
1.0
1.0
1.0
1.0
1.0
1.0
1.0
1.0
1.0
1.0
1.0
1.0
1.0
1.0
1.0
1.0
1.0
1.0
1.0
1.0
1.0
1.0
1.0
1.0
1.0
1.0


15
1
1
1
0
0
0
1
0
0
0
1
0
1
1
0
0
0
0
2
0
0
0
0
0
0
1
1
3
1


12
1
1
1
0
0
0
1
0
0
0
1
0
1
1
0
0
0
0
0
0
0
0
0
0
0
0
1
3
1

1.0
1.0
1.0
1.0
1.0
1.0
1.0
1.0
1.0
1.0
1.0
1.0
1.0
1.0
1.0
1.0
1.0
1.0
1.0
1.0
1.0
1.0
1.0
1.0
1.0
1.0
1.0
1.0
1.0
1.0


3
0
0
0
0
0
0
0
0
0
0
0
0
0
0
0
0
0
0
2
0
0
0
0
0
0
1
0
0
0

0.99
0.99
0.99
0.99
0.99
0.99
0.99
0.99
0.99
0.99
0.99
0.99
0.99
0.99
0.99
0.99
0.99
0.99
0.99
0.99
0.99
0.99
0.99
0.99
0.99
0.99
0.99
0.99
0.99
0.99


90
1
3
3
1
1
3
1
1
1
1
1
0
0
8
3
3
0
0
11
13
9
16
0
6
0
0
2
0
2

0.71
0.71
0.71
0.71
0.71
0.71
0.71
0.71
0.71
0.71
0.71
0.71
0.71
0.71
0.71
0.71
0.71
0.71
0.71
0.71
0.71
0.71
0.71
0.71
0.71
0.71
0.71
0.71
0.71
0.71


50
1
2
0
0
1
3
0
1
0
0
0
0
0
0
0
0
0
0
8
9
9
12
0
0
0
0
2
0
2

0.85
0.85
0.85
0.85
0.85
0.85
0.85
0.85
0.85
0.85
0.85
0.85
0.85
0.85
0.85
0.85
0.85
0.85
0.85
0.85
0.85
0.85
0.85
0.85
0.85
0.85
0.85
0.85
0.85
0.85


6
1
2
0
0
0
3
0
0
0
0
0
0
0
0
0
0
0
0
0
0
0
0
0
0
0
0
0
0
0

1.0
1.0
1.0
1.0
1.0
1.0
1.0
1.0
1.0
1.0
1.0
1.0
1.0
1.0
1.0
1.0
1.0
1.0
1.0
1.0
1.0
1.0
1.0
1.0
1.0
1.0
1.0
1.0
1.0
1.0


19
0
0
0
0
1
0
0
1
0
0
0
0
0
0
0
0
0
0
4
4
4
4
0
0
0
0
1
0
0

1.0
1.0
1.0
1.0
1.0
1.0
1.0
1.0
1.0
1.0
1.0
1.0
1.0
1.0
1.0
1.0
1.0
1.0
1.0
1.0
1.0
1.0
1.0
1.0
1.0
1.0
1.0
1.0
1.0
1.0


19
0
1
1
1
0
0
1
0
1
1
0
0
0
7
0
0
0
0
0
0
0
0
0
6
0
0
0
0
0


19
0
1
1
1
0
0
1
0
1
1
0
0
0
7
0
0
0
0
0
0
0
0
0
6
0
0
0
0
0

1.0
1.0
1.0
1.0
1.0
1.0
1.0
1.0
1.0
1.0
1.0
1.0
1.0
1.0
1.0
1.0
1.0
1.0
1.0
1.0
1.0
1.0
1.0
1.0
1.0
1.0
1.0
1.0
1.0
1.0


1
0
0
0
0
0
0
0
0
0
0
1
0
0
0
0
0
0
0
0
0
0
0
0
0
0
0
0
0
0


1
0
0
0
0
0
0
0
0
0
0
1
0
0
0
0
0
0
0
0
0
0
0
0
0
0
0
0
0
0

1.0
1.0
1.0
1.0
1.0
1.0
1.0
1.0
1.0
1.0
1.0
1.0
1.0
1.0
1.0
1.0
1.0
1.0
1.0
1.0
1.0
1.0
1.0
1.0
1.0
1.0
1.0
1.0
1.0
1.0


2
0
0
2
0
0
0
0
0
0
0
0
0
0
0
0
0
0
0
0
0
0
0
0
0
0
0
0
0
0


2
0
0
2
0
0
0
0
0
0
0
0
0
0
0
0
0
0
0
0
0
0
0
0
0
0
0
0
0
0

1.0
1.0
1.0
1.0
1.0
1.0
1.0
1.0
1.0
1.0
1.0
1.0
1.0
1.0
1.0
1.0
1.0
1.0
1.0
1.0
1.0
1.0
1.0
1.0
1.0
1.0
1.0
1.0
1.0
1.0


17
0
0
0
0
0
0
0
0
0
0
0
0
0
0
3
3
0
0
3
4
0
4
0
0
0
0
0
0
0


17
0
0
0
0
0
0
0
0
0
0
0
0
0
0
3
3
0
0
3
4
0
4
0
0
0
0
0
0
0

1.0
1.0
1.0
1.0
1.0
1.0
1.0
1.0
1.0
1.0
1.0
1.0
1.0
1.0
1.0
1.0
1.0
1.0
1.0
1.0
1.0
1.0
1.0
1.0
1.0
1.0
1.0
1.0
1.0
1.0


33
6
0
4
2
2
3
0
0
0
0
0
5
0
0
1
0
0
0
0
3
0
0
5
0
0
2
0
0
0

0.7
0.7
0.7
0.7
0.7
0.7
0.7
0.7
0.7
0.7
0.7
0.7
0.7
0.7
0.7
0.7
0.7
0.7
0.7
0.7
0.7
0.7
0.7
0.7
0.7
0.7
0.7
0.7
0.7
0.7


5
0
0
1
1
0
3
0
0
0
0
0
0
0
0
0
0
0
0
0
0
0
0
0
0
0
0
0
0
0


1
0
0
0
1
0
0
0
0
0
0
0
0
0
0
0
0
0
0
0
0
0
0
0
0
0
0
0
0
0

0.99
0.99
0.99
0.99
0.99
0.99
0.99
0.99
0.99
0.99
0.99
0.99
0.99
0.99
0.99
0.99
0.99
0.99
0.99
0.99
0.99
0.99
0.99
0.99
0.99
0.99
0.99
0.99
0.99
0.99


4
0
0
1
0
0
3
0
0
0
0
0
0
0
0
0
0
0
0
0
0
0
0
0
0
0
0
0
0
0

1.0
1.0
1.0
1.0
1.0
1.0
1.0
1.0
1.0
1.0
1.0
1.0
1.0
1.0
1.0
1.0
1.0
1.0
1.0
1.0
1.0
1.0
1.0
1.0
1.0
1.0
1.0
1.0
1.0
1.0


19
4
0
3
0
2
0
0
0
0
0
0
5
0
0
0
0
0
0
0
0
0
0
5
0
0
0
0
0
0

0.92
0.92
0.92
0.92
0.92
0.92
0.92
0.92
0.92
0.92
0.92
0.92
0.92
0.92
0.92
0.92
0.92
0.92
0.92
0.92
0.92
0.92
0.92
0.92
0.92
0.92
0.92
0.92
0.92
0.92


15
4
0
3
0
2
0
0
0
0
0
0
3
0
0
0
0
0
0
0
0
0
0
3
0
0
0
0
0
0

1.0
1.0
1.0
1.0
1.0
1.0
1.0
1.0
1.0
1.0
1.0
1.0
1.0
1.0
1.0
1.0
1.0
1.0
1.0
1.0
1.0
1.0
1.0
1.0
1.0
1.0
1.0
1.0
1.0
1.0


1
1
0
0
0
0
0
0
0
0
0
0
0
0
0
0
0
0
0
0
0
0
0
0
0
0
0
0
0
0


1
1
0
0
0
0
0
0
0
0
0
0
0
0
0
0
0
0
0
0
0
0
0
0
0
0
0
0
0
0

1.0
1.0
1.0
1.0
1.0
1.0
1.0
1.0
1.0
1.0
1.0
1.0
1.0
1.0
1.0
1.0
1.0
1.0
1.0
1.0
1.0
1.0
1.0
1.0
1.0
1.0
1.0
1.0
1.0
1.0


4
1
0
0
1
0
0
0
0
0
0
0
0
0
0
0
0
0
0
0
2
0
0
0
0
0
0
0
0
0


3
0
0
0
1
0
0
0
0
0
0
0
0
0
0
0
0
0
0
0
2
0
0
0
0
0
0
0
0
0

1.0
1.0
1.0
1.0
1.0
1.0
1.0
1.0
1.0
1.0
1.0
1.0
1.0
1.0
1.0
1.0
1.0
1.0
1.0
1.0
1.0
1.0
1.0
1.0
1.0
1.0
1.0
1.0
1.0
1.0


1
1
0
0
0
0
0
0
0
0
0
0
0
0
0
0
0
0
0
0
0
0
0
0
0
0
0
0
0
0

1.0
1.0
1.0
1.0
1.0
1.0
1.0
1.0
1.0
1.0
1.0
1.0
1.0
1.0
1.0
1.0
1.0
1.0
1.0
1.0
1.0
1.0
1.0
1.0
1.0
1.0
1.0
1.0
1.0
1.0


2
0
0
0
0
0
0
0
0
0
0
0
0
0
0
1
0
0
0
0
0
0
0
0
0
0
1
0
0
0

1.0
1.0
1.0
1.0
1.0
1.0
1.0
1.0
1.0
1.0
1.0
1.0
1.0
1.0
1.0
1.0
1.0
1.0
1.0
1.0
1.0
1.0
1.0
1.0
1.0
1.0
1.0
1.0
1.0
1.0


1
0
0
0
0
0
0
0
0
0
0
0
0
0
0
0
0
0
0
0
0
0
0
0
0
0
1
0
0
0


1
0
0
0
0
0
0
0
0
0
0
0
0
0
0
0
0
0
0
0
0
0
0
0
0
0
1
0
0
0

1.0
1.0
1.0
1.0
1.0
1.0
1.0
1.0
1.0
1.0
1.0
1.0
1.0
1.0
1.0
1.0
1.0
1.0
1.0
1.0
1.0
1.0
1.0
1.0
1.0
1.0
1.0
1.0
1.0
1.0


3
0
0
0
0
3
0
0
0
0
0
0
0
0
0
0
0
0
0
0
0
0
0
0
0
0
0
0
0
0


3
0
0
0
0
3
0
0
0
0
0
0
0
0
0
0
0
0
0
0
0
0
0
0
0
0
0
0
0
0

1.0
1.0
1.0
1.0
1.0
1.0
1.0
1.0
1.0
1.0
1.0
1.0
1.0
1.0
1.0
1.0
1.0
1.0
1.0
1.0
1.0
1.0
1.0
1.0
1.0
1.0
1.0
1.0
1.0
1.0


1
0
0
0
0
0
0
0
0
0
1
0
0
0
0
0
0
0
0
0
0
0
0
0
0
0
0
0
0
0


1
0
0
0
0
0
0
0
0
0
1
0
0
0
0
0
0
0
0
0
0
0
0
0
0
0
0
0
0
0

1.0
1.0
1.0
1.0
1.0
1.0
1.0
1.0
1.0
1.0
1.0
1.0
1.0
1.0
1.0
1.0
1.0
1.0
1.0
1.0
1.0
1.0
1.0
1.0
1.0
1.0
1.0
1.0
1.0
1.0


21
4
0
2
2
0
1
0
1
2
2
2
0
0
2
0
0
0
0
0
0
0
0
0
0
0
0
0
0
3


18
3
0
2
1
0
1
0
1
2
2
1
0
0
2
0
0
0
0
0
0
0
0
0
0
0
0
0
0
3

0.98
0.98
0.98
0.98
0.98
0.98
0.98
0.98
0.98
0.98
0.98
0.98
0.98
0.98
0.98
0.98
0.98
0.98
0.98
0.98
0.98
0.98
0.98
0.98
0.98
0.98
0.98
0.98
0.98
0.98


15
2
0
2
1
0
1
0
0
1
2
1
0
0
2
0
0
0
0
0
0
0
0
0
0
0
0
0
0
3

1.0
1.0
1.0
1.0
1.0
1.0
1.0
1.0
1.0
1.0
1.0
1.0
1.0
1.0
1.0
1.0
1.0
1.0
1.0
1.0
1.0
1.0
1.0
1.0
1.0
1.0
1.0
1.0
1.0
1.0


1
1
0
0
0
0
0
0
0
0
0
0
0
0
0
0
0
0
0
0
0
0
0
0
0
0
0
0
0
0

1.0
1.0
1.0
1.0
1.0
1.0
1.0
1.0
1.0
1.0
1.0
1.0
1.0
1.0
1.0
1.0
1.0
1.0
1.0
1.0
1.0
1.0
1.0
1.0
1.0
1.0
1.0
1.0
1.0
1.0


1
0
0
0
1
0
0
0
0
0
0
0
0
0
0
0
0
0
0
0
0
0
0
0
0
0
0
0
0
0

1.0
1.0
1.0
1.0
1.0
1.0
1.0
1.0
1.0
1.0
1.0
1.0
1.0
1.0
1.0
1.0
1.0
1.0
1.0
1.0
1.0
1.0
1.0
1.0
1.0
1.0
1.0
1.0
1.0
1.0


2
1
0
0
0
0
0
0
0
0
0
1
0
0
0
0
0
0
0
0
0
0
0
0
0
0
0
0
0
0

0.77
0.77
0.77
0.77
0.77
0.77
0.77
0.77
0.77
0.77
0.77
0.77
0.77
0.77
0.77
0.77
0.77
0.77
0.77
0.77
0.77
0.77
0.77
0.77
0.77
0.77
0.77
0.77
0.77
0.77


2
0
1
1
0
0
0
0
0
0
0
0
0
0
0
0
0
0
0
0
0
0
0
0
0
0
0
0
0
0


2
0
1
1
0
0
0
0
0
0
0
0
0
0
0
0
0
0
0
0
0
0
0
0
0
0
0
0
0
0


2
0
1
1
0
0
0
0
0
0
0
0
0
0
0
0
0
0
0
0
0
0
0
0
0
0
0
0
0
0

1.0
1.0
1.0
1.0
1.0
1.0
1.0
1.0
1.0
1.0
1.0
1.0
1.0
1.0
1.0
1.0
1.0
1.0
1.0
1.0
1.0
1.0
1.0
1.0
1.0
1.0
1.0
1.0
1.0
1.0


7
0
0
0
1
1
1
0
0
0
0
1
0
0
0
0
0
0
0
0
0
0
0
0
1
1
0
0
0
1

0.79
0.79
0.79
0.79
0.79
0.79
0.79
0.79
0.79
0.79
0.79
0.79
0.79
0.79
0.79
0.79
0.79
0.79
0.79
0.79
0.79
0.79
0.79
0.79
0.79
0.79
0.79
0.79
0.79
0.79


4
0
0
0
1
0
1
0
0
0
0
0
0
0
0
0
0
0
0
0
0
0
0
0
0
1
0
0
0
1


4
0
0
0
1
0
1
0
0
0
0
0
0
0
0
0
0
0
0
0
0
0
0
0
0
1
0
0
0
1

1.0
1.0
1.0
1.0
1.0
1.0
1.0
1.0
1.0
1.0
1.0
1.0
1.0
1.0
1.0
1.0
1.0
1.0
1.0
1.0
1.0
1.0
1.0
1.0
1.0
1.0
1.0
1.0
1.0
1.0


1
0
0
0
0
1
0
0
0
0
0
0
0
0
0
0
0
0
0
0
0
0
0
0
0
0
0
0
0
0

0.97
0.97
0.97
0.97
0.97
0.97
0.97
0.97
0.97
0.97
0.97
0.97
0.97
0.97
0.97
0.97
0.97
0.97
0.97
0.97
0.97
0.97
0.97
0.97
0.97
0.97
0.97
0.97
0.97
0.97


1
0
0
0
0
0
0
0
0
0
0
0
0
0
0
0
0
0
0
0
0
0
0
0
1
0
0
0
0
0


1
0
0
0
0
0
0
0
0
0
0
0
0
0
0
0
0
0
0
0
0
0
0
0
1
0
0
0
0
0

1.0
1.0
1.0
1.0
1.0
1.0
1.0
1.0
1.0
1.0
1.0
1.0
1.0
1.0
1.0
1.0
1.0
1.0
1.0
1.0
1.0
1.0
1.0
1.0
1.0
1.0
1.0
1.0
1.0
1.0


6
1
1
0
0
0
3
0
0
0
0
1
0
0
0
0
0
0
0
0
0
0
0
0
0
0
0
0
0
0


3
1
1
0
0
0
0
0
0
0
0
1
0
0
0
0
0
0
0
0
0
0
0
0
0
0
0
0
0
0


3
1
1
0
0
0
0
0
0
0
0
1
0
0
0
0
0
0
0
0
0
0
0
0
0
0
0
0
0
0

1.0
1.0
1.0
1.0
1.0
1.0
1.0
1.0
1.0
1.0
1.0
1.0
1.0
1.0
1.0
1.0
1.0
1.0
1.0
1.0
1.0
1.0
1.0
1.0
1.0
1.0
1.0
1.0
1.0
1.0


3
0
0
0
0
0
3
0
0
0
0
0
0
0
0
0
0
0
0
0
0
0
0
0
0
0
0
0
0
0


3
0
0
0
0
0
3
0
0
0
0
0
0
0
0
0
0
0
0
0
0
0
0
0
0
0
0
0
0
0

1.0
1.0
1.0
1.0
1.0
1.0
1.0
1.0
1.0
1.0
1.0
1.0
1.0
1.0
1.0
1.0
1.0
1.0
1.0
1.0
1.0
1.0
1.0
1.0
1.0
1.0
1.0
1.0
1.0
1.0


19
2
1
0
1
1
1
1
2
2
2
2
0
1
1
0
0
0
1
0
0
0
0
0
1
0
0
0
0
0


13
1
1
0
1
1
1
0
1
1
1
1
0
1
1
0
0
0
1
0
0
0
0
0
1
0
0
0
0
0


13
1
1
0
1
1
1
0
1
1
1
1
0
1
1
0
0
0
1
0
0
0
0
0
1
0
0
0
0
0

1.0
1.0
1.0
1.0
1.0
1.0
1.0
1.0
1.0
1.0
1.0
1.0
1.0
1.0
1.0
1.0
1.0
1.0
1.0
1.0
1.0
1.0
1.0
1.0
1.0
1.0
1.0
1.0
1.0
1.0


6
1
0
0
0
0
0
1
1
1
1
1
0
0
0
0
0
0
0
0
0
0
0
0
0
0
0
0
0
0


6
1
0
0
0
0
0
1
1
1
1
1
0
0
0
0
0
0
0
0
0
0
0
0
0
0
0
0
0
0

0.75
0.75
0.75
0.75
0.75
0.75
0.75
0.75
0.75
0.75
0.75
0.75
0.75
0.75
0.75
0.75
0.75
0.75
0.75
0.75
0.75
0.75
0.75
0.75
0.75
0.75
0.75
0.75
0.75
0.75


80
7
9
2
6
8
7
0
1
1
5
6
0
0
1
1
0
0
0
1
0
0
0
0
0
3
6
2
5
9

0.97
0.97
0.97
0.97
0.97
0.97
0.97
0.97
0.97
0.97
0.97
0.97
0.97
0.97
0.97
0.97
0.97
0.97
0.97
0.97
0.97
0.97
0.97
0.97
0.97
0.97
0.97
0.97
0.97
0.97


11
1
1
1
0
0
0
0
1
1
1
1
0
0
1
1
0
0
0
1
0
0
0
0
0
0
0
0
0
1

1.0
1.0
1.0
1.0
1.0
1.0
1.0
1.0
1.0
1.0
1.0
1.0
1.0
1.0
1.0
1.0
1.0
1.0
1.0
1.0
1.0
1.0
1.0
1.0
1.0
1.0
1.0
1.0
1.0
1.0


2
0
0
0
0
0
0
0
0
0
0
0
0
0
0
1
0
0
0
0
0
0
0
0
0
0
0
0
0
1

1.0
1.0
1.0
1.0
1.0
1.0
1.0
1.0
1.0
1.0
1.0
1.0
1.0
1.0
1.0
1.0
1.0
1.0
1.0
1.0
1.0
1.0
1.0
1.0
1.0
1.0
1.0
1.0
1.0
1.0


1
0
0
0
0
0
0
0
0
0
0
0
0
0
1
0
0
0
0
0
0
0
0
0
0
0
0
0
0
0

0.9
0.9
0.9
0.9
0.9
0.9
0.9
0.9
0.9
0.9
0.9
0.9
0.9
0.9
0.9
0.9
0.9
0.9
0.9
0.9
0.9
0.9
0.9
0.9
0.9
0.9
0.9
0.9
0.9
0.9


39
2
3
1
2
4
5
0
0
0
2
3
0
0
0
0
0
0
0
0
0
0
0
0
0
2
4
1
5
5


39
2
3
1
2
4
5
0
0
0
2
3
0
0
0
0
0
0
0
0
0
0
0
0
0
2
4
1
5
5

1.0
1.0
1.0
1.0
1.0
1.0
1.0
1.0
1.0
1.0
1.0
1.0
1.0
1.0
1.0
1.0
1.0
1.0
1.0
1.0
1.0
1.0
1.0
1.0
1.0
1.0
1.0
1.0
1.0
1.0


2
0
0
0
0
0
0
0
0
0
0
0
0
0
0
0
0
0
0
0
0
0
0
0
0
1
0
0
0
1

0.77
0.77
0.77
0.77
0.77
0.77
0.77
0.77
0.77
0.77
0.77
0.77
0.77
0.77
0.77
0.77
0.77
0.77
0.77
0.77
0.77
0.77
0.77
0.77
0.77
0.77
0.77
0.77
0.77
0.77


5
0
0
0
0
0
0
0
0
0
0
0
0
0
0
0
0
0
0
0
0
0
0
0
0
0
2
1
0
2

1.0
1.0
1.0
1.0
1.0
1.0
1.0
1.0
1.0
1.0
1.0
1.0
1.0
1.0
1.0
1.0
1.0
1.0
1.0
1.0
1.0
1.0
1.0
1.0
1.0
1.0
1.0
1.0
1.0
1.0


6
0
0
0
3
1
1
0
0
0
0
0
0
0
0
0
0
0
0
0
0
0
0
0
0
0
0
1
0
0


6
0
0
0
3
1
1
0
0
0
0
0
0
0
0
0
0
0
0
0
0
0
0
0
0
0
0
1
0
0


6
0
0
0
3
1
1
0
0
0
0
0
0
0
0
0
0
0
0
0
0
0
0
0
0
0
0
1
0
0

1.0
1.0
1.0
1.0
1.0
1.0
1.0
1.0
1.0
1.0
1.0
1.0
1.0
1.0
1.0
1.0
1.0
1.0
1.0
1.0
1.0
1.0
1.0
1.0
1.0
1.0
1.0
1.0
1.0
1.0


6
0
0
0
1
1
0
0
0
2
0
2
0
0
0
0
0
0
0
0
0
0
0
0
0
0
0
0
0
0


2
0
0
0
1
1
0
0
0
0
0
0
0
0
0
0
0
0
0
0
0
0
0
0
0
0
0
0
0
0


2
0
0
0
1
1
0
0
0
0
0
0
0
0
0
0
0
0
0
0
0
0
0
0
0
0
0
0
0
0

1.0
1.0
1.0
1.0
1.0
1.0
1.0
1.0
1.0
1.0
1.0
1.0
1.0
1.0
1.0
1.0
1.0
1.0
1.0
1.0
1.0
1.0
1.0
1.0
1.0
1.0
1.0
1.0
1.0
1.0


4
0
0
0
0
0
0
0
0
2
0
2
0
0
0
0
0
0
0
0
0
0
0
0
0
0
0
0
0
0


2
0
0
0
0
0
0
0
0
1
0
1
0
0
0
0
0
0
0
0
0
0
0
0
0
0
0
0
0
0

1.0
1.0
1.0
1.0
1.0
1.0
1.0
1.0
1.0
1.0
1.0
1.0
1.0
1.0
1.0
1.0
1.0
1.0
1.0
1.0
1.0
1.0
1.0
1.0
1.0
1.0
1.0
1.0
1.0
1.0


2
0
0
0
0
0
0
0
0
1
0
1
0
0
0
0
0
0
0
0
0
0
0
0
0
0
0
0
0
0

1.0
1.0
1.0
1.0
1.0
1.0
1.0
1.0
1.0
1.0
1.0
1.0
1.0
1.0
1.0
1.0
1.0
1.0
1.0
1.0
1.0
1.0
1.0
1.0
1.0
1.0
1.0
1.0
1.0
1.0


2
1
0
0
0
0
0
0
0
1
0
0
0
0
0
0
0
0
0
0
0
0
0
0
0
0
0
0
0
0


2
1
0
0
0
0
0
0
0
1
0
0
0
0
0
0
0
0
0
0
0
0
0
0
0
0
0
0
0
0


2
1
0
0
0
0
0
0
0
1
0
0
0
0
0
0
0
0
0
0
0
0
0
0
0
0
0
0
0
0

1.0
1.0
1.0
1.0
1.0
1.0
1.0
1.0
1.0
1.0
1.0
1.0
1.0
1.0
1.0
1.0
1.0
1.0
1.0
1.0
1.0
1.0
1.0
1.0
1.0
1.0
1.0
1.0
1.0
1.0


78
6
1
2
2
0
0
1
2
7
10
9
0
2
2
0
0
3
4
1
0
1
0
0
11
1
1
1
5
6

1.0
1.0
1.0
1.0
1.0
1.0
1.0
1.0
1.0
1.0
1.0
1.0
1.0
1.0
1.0
1.0
1.0
1.0
1.0
1.0
1.0
1.0
1.0
1.0
1.0
1.0
1.0
1.0
1.0
1.0


8
0
0
0
0
0
0
0
0
4
3
1
0
0
0
0
0
0
0
0
0
0
0
0
0
0
0
0
0
0


8
0
0
0
0
0
0
0
0
4
3
1
0
0
0
0
0
0
0
0
0
0
0
0
0
0
0
0
0
0

1.0
1.0
1.0
1.0
1.0
1.0
1.0
1.0
1.0
1.0
1.0
1.0
1.0
1.0
1.0
1.0
1.0
1.0
1.0
1.0
1.0
1.0
1.0
1.0
1.0
1.0
1.0
1.0
1.0
1.0


5
1
0
0
0
0
0
0
0
0
1
1
0
0
1
0
0
0
0
0
0
0
0
0
1
0
0
0
0
0


5
1
0
0
0
0
0
0
0
0
1
1
0
0
1
0
0
0
0
0
0
0
0
0
1
0
0
0
0
0

1.0
1.0
1.0
1.0
1.0
1.0
1.0
1.0
1.0
1.0
1.0
1.0
1.0
1.0
1.0
1.0
1.0
1.0
1.0
1.0
1.0
1.0
1.0
1.0
1.0
1.0
1.0
1.0
1.0
1.0


6
1
0
1
0
0
0
0
0
0
2
2
0
0
0
0
0
0
0
0
0
0
0
0
0
0
0
0
0
0


6
1
0
1
0
0
0
0
0
0
2
2
0
0
0
0
0
0
0
0
0
0
0
0
0
0
0
0
0
0

0.99
0.99
0.99
0.99
0.99
0.99
0.99
0.99
0.99
0.99
0.99
0.99
0.99
0.99
0.99
0.99
0.99
0.99
0.99
0.99
0.99
0.99
0.99
0.99
0.99
0.99
0.99
0.99
0.99
0.99


7
1
0
0
0
0
0
1
1
0
1
1
0
0
0
0
0
0
1
0
0
0
0
0
1
0
0
0
0
0


6
1
0
0
0
0
0
1
1
0
1
1
0
0
0
0
0
0
0
0
0
0
0
0
1
0
0
0
0
0

1.0
1.0
1.0
1.0
1.0
1.0
1.0
1.0
1.0
1.0
1.0
1.0
1.0
1.0
1.0
1.0
1.0
1.0
1.0
1.0
1.0
1.0
1.0
1.0
1.0
1.0
1.0
1.0
1.0
1.0


1
0
0
0
0
0
0
0
0
0
0
0
0
0
0
0
0
0
1
0
0
0
0
0
0
0
0
0
0
0

1.0
1.0
1.0
1.0
1.0
1.0
1.0
1.0
1.0
1.0
1.0
1.0
1.0
1.0
1.0
1.0
1.0
1.0
1.0
1.0
1.0
1.0
1.0
1.0
1.0
1.0
1.0
1.0
1.0
1.0


4
1
1
0
1
0
0
0
0
0
0
1
0
0
0
0
0
0
0
0
0
0
0
0
0
0
0
0
0
0


4
1
1
0
1
0
0
0
0
0
0
1
0
0
0
0
0
0
0
0
0
0
0
0
0
0
0
0
0
0

1.0
1.0
1.0
1.0
1.0
1.0
1.0
1.0
1.0
1.0
1.0
1.0
1.0
1.0
1.0
1.0
1.0
1.0
1.0
1.0
1.0
1.0
1.0
1.0
1.0
1.0
1.0
1.0
1.0
1.0


15
1
0
1
1
0
0
0
0
1
1
1
0
0
0
0
0
2
1
0
0
0
0
0
2
0
0
0
2
2

1.0
1.0
1.0
1.0
1.0
1.0
1.0
1.0
1.0
1.0
1.0
1.0
1.0
1.0
1.0
1.0
1.0
1.0
1.0
1.0
1.0
1.0
1.0
1.0
1.0
1.0
1.0
1.0
1.0
1.0


2
0
0
0
0
0
0
0
0
0
0
0
0
0
0
0
0
0
0
0
0
0
0
0
1
0
0
0
0
1

0.95
0.95
0.95
0.95
0.95
0.95
0.95
0.95
0.95
0.95
0.95
0.95
0.95
0.95
0.95
0.95
0.95
0.95
0.95
0.95
0.95
0.95
0.95
0.95
0.95
0.95
0.95
0.95
0.95
0.95


1
0
0
0
0
0
0
0
0
0
0
0
0
0
0
0
0
0
0
0
0
0
0
0
0
0
0
0
1
0

1.0
1.0
1.0
1.0
1.0
1.0
1.0
1.0
1.0
1.0
1.0
1.0
1.0
1.0
1.0
1.0
1.0
1.0
1.0
1.0
1.0
1.0
1.0
1.0
1.0
1.0
1.0
1.0
1.0
1.0


7
1
0
0
0
0
0
0
0
1
1
2
0
0
0
0
0
0
0
0
0
0
0
0
2
0
0
0
0
0

1.0
1.0
1.0
1.0
1.0
1.0
1.0
1.0
1.0
1.0
1.0
1.0
1.0
1.0
1.0
1.0
1.0
1.0
1.0
1.0
1.0
1.0
1.0
1.0
1.0
1.0
1.0
1.0
1.0
1.0


3
0
0
0
0
0
0
0
0
1
1
1
0
0
0
0
0
0
0
0
0
0
0
0
0
0
0
0
0
0

1.0
1.0
1.0
1.0
1.0
1.0
1.0
1.0
1.0
1.0
1.0
1.0
1.0
1.0
1.0
1.0
1.0
1.0
1.0
1.0
1.0
1.0
1.0
1.0
1.0
1.0
1.0
1.0
1.0
1.0


2
1
0
0
0
0
0
0
0
0
0
1
0
0
0
0
0
0
0
0
0
0
0
0
0
0
0
0
0
0

1.0
1.0
1.0
1.0
1.0
1.0
1.0
1.0
1.0
1.0
1.0
1.0
1.0
1.0
1.0
1.0
1.0
1.0
1.0
1.0
1.0
1.0
1.0
1.0
1.0
1.0
1.0
1.0
1.0
1.0


3
0
0
0
0
0
0
0
1
1
1
0
0
0
0
0
0
0
0
0
0
0
0
0
0
0
0
0
0
0


3
0
0
0
0
0
0
0
1
1
1
0
0
0
0
0
0
0
0
0
0
0
0
0
0
0
0
0
0
0

1.0
1.0
1.0
1.0
1.0
1.0
1.0
1.0
1.0
1.0
1.0
1.0
1.0
1.0
1.0
1.0
1.0
1.0
1.0
1.0
1.0
1.0
1.0
1.0
1.0
1.0
1.0
1.0
1.0
1.0


4
0
0
0
0
0
0
0
0
0
0
0
0
0
0
0
0
0
0
0
0
0
0
0
1
0
0
0
1
2


2
0
0
0
0
0
0
0
0
0
0
0
0
0
0
0
0
0
0
0
0
0
0
0
1
0
0
0
0
1

0.92
0.92
0.92
0.92
0.92
0.92
0.92
0.92
0.92
0.92
0.92
0.92
0.92
0.92
0.92
0.92
0.92
0.92
0.92
0.92
0.92
0.92
0.92
0.92
0.92
0.92
0.92
0.92
0.92
0.92


2
0
0
0
0
0
0
0
0
0
0
0
0
0
0
0
0
0
0
0
0
0
0
0
0
0
0
0
1
1

1.0
1.0
1.0
1.0
1.0
1.0
1.0
1.0
1.0
1.0
1.0
1.0
1.0
1.0
1.0
1.0
1.0
1.0
1.0
1.0
1.0
1.0
1.0
1.0
1.0
1.0
1.0
1.0
1.0
1.0


1
0
0
0
0
0
0
0
0
0
0
0
0
0
0
0
0
0
0
0
0
1
0
0
0
0
0
0
0
0


1
0
0
0
0
0
0
0
0
0
0
0
0
0
0
0
0
0
0
0
0
1
0
0
0
0
0
0
0
0

1.0
1.0
1.0
1.0
1.0
1.0
1.0
1.0
1.0
1.0
1.0
1.0
1.0
1.0
1.0
1.0
1.0
1.0
1.0
1.0
1.0
1.0
1.0
1.0
1.0
1.0
1.0
1.0
1.0
1.0


4
0
0
0
0
0
0
0
0
0
0
0
0
0
0
0
0
0
0
0
0
0
0
0
0
0
0
0
2
2


4
0
0
0
0
0
0
0
0
0
0
0
0
0
0
0
0
0
0
0
0
0
0
0
0
0
0
0
2
2

1.0
1.0
1.0
1.0
1.0
1.0
1.0
1.0
1.0
1.0
1.0
1.0
1.0
1.0
1.0
1.0
1.0
1.0
1.0
1.0
1.0
1.0
1.0
1.0
1.0
1.0
1.0
1.0
1.0
1.0


1
0
0
0
0
0
0
0
0
0
0
0
0
0
0
0
0
0
0
0
0
0
0
0
0
1
0
0
0
0

1.0
1.0
1.0
1.0
1.0
1.0
1.0
1.0
1.0
1.0
1.0
1.0
1.0
1.0
1.0
1.0
1.0
1.0
1.0
1.0
1.0
1.0
1.0
1.0
1.0
1.0
1.0
1.0
1.0
1.0


1
0
0
0
0
0
0
0
0
0
0
0
0
0
0
0
0
0
0
0
0
0
0
0
1
0
0
0
0
0


1
0
0
0
0
0
0
0
0
0
0
0
0
0
0
0
0
0
0
0
0
0
0
0
1
0
0
0
0
0

1.0
1.0
1.0
1.0
1.0
1.0
1.0
1.0
1.0
1.0
1.0
1.0
1.0
1.0
1.0
1.0
1.0
1.0
1.0
1.0
1.0
1.0
1.0
1.0
1.0
1.0
1.0
1.0
1.0
1.0


1
0
0
0
0
0
0
0
0
0
0
0
0
0
0
0
0
0
1
0
0
0
0
0
0
0
0
0
0
0


1
0
0
0
0
0
0
0
0
0
0
0
0
0
0
0
0
0
1
0
0
0
0
0
0
0
0
0
0
0

1.0
1.0
1.0
1.0
1.0
1.0
1.0
1.0
1.0
1.0
1.0
1.0
1.0
1.0
1.0
1.0
1.0
1.0
1.0
1.0
1.0
1.0
1.0
1.0
1.0
1.0
1.0
1.0
1.0
1.0


1
0
0
0
0
0
0
0
0
0
0
0
0
0
0
0
0
0
0
0
0
0
0
0
1
0
0
0
0
0


1
0
0
0
0
0
0
0
0
0
0
0
0
0
0
0
0
0
0
0
0
0
0
0
1
0
0
0
0
0

1.0
1.0
1.0
1.0
1.0
1.0
1.0
1.0
1.0
1.0
1.0
1.0
1.0
1.0
1.0
1.0
1.0
1.0
1.0
1.0
1.0
1.0
1.0
1.0
1.0
1.0
1.0
1.0
1.0
1.0


1
0
0
0
0
0
0
0
0
0
0
0
0
0
1
0
0
0
0
0
0
0
0
0
0
0
0
0
0
0


1
0
0
0
0
0
0
0
0
0
0
0
0
0
1
0
0
0
0
0
0
0
0
0
0
0
0
0
0
0

0.85
0.85
0.85
0.85
0.85
0.85
0.85
0.85
0.85
0.85
0.85
0.85
0.85
0.85
0.85
0.85
0.85
0.85
0.85
0.85
0.85
0.85
0.85
0.85
0.85
0.85
0.85
0.85
0.85
0.85


1
0
0
0
0
0
0
0
0
0
0
0
0
1
0
0
0
0
0
0
0
0
0
0
0
0
0
0
0
0

1.0
1.0
1.0
1.0
1.0
1.0
1.0
1.0
1.0
1.0
1.0
1.0
1.0
1.0
1.0
1.0
1.0
1.0
1.0
1.0
1.0
1.0
1.0
1.0
1.0
1.0
1.0
1.0
1.0
1.0


1
0
0
0
0
0
0
0
0
0
0
0
0
1
0
0
0
0
0
0
0
0
0
0
0
0
0
0
0
0

1.0
1.0
1.0
1.0
1.0
1.0
1.0
1.0
1.0
1.0
1.0
1.0
1.0
1.0
1.0
1.0
1.0
1.0
1.0
1.0
1.0
1.0
1.0
1.0
1.0
1.0
1.0
1.0
1.0
1.0


1
0
0
0
0
0
0
0
0
0
0
0
0
0
0
0
0
0
0
0
0
0
0
0
0
0
0
1
0
0


1
0
0
0
0
0
0
0
0
0
0
0
0
0
0
0
0
0
0
0
0
0
0
0
0
0
0
1
0
0

0.81
0.81
0.81
0.81
0.81
0.81
0.81
0.81
0.81
0.81
0.81
0.81
0.81
0.81
0.81
0.81
0.81
0.81
0.81
0.81
0.81
0.81
0.81
0.81
0.81
0.81
0.81
0.81
0.81
0.81


1
0
0
0
0
0
1
0
0
0
0
0
0
0
0
0
0
0
0
0
0
0
0
0
0
0
0
0
0
0


1
0
0
0
0
0
1
0
0
0
0
0
0
0
0
0
0
0
0
0
0
0
0
0
0
0
0
0
0
0


1
0
0
0
0
0
1
0
0
0
0
0
0
0
0
0
0
0
0
0
0
0
0
0
0
0
0
0
0
0

1.0
1.0
1.0
1.0
1.0
1.0
1.0
1.0
1.0
1.0
1.0
1.0
1.0
1.0
1.0
1.0
1.0
1.0
1.0
1.0
1.0
1.0
1.0
1.0
1.0
1.0
1.0
1.0
1.0
1.0


55
0
1
1
0
0
0
0
0
0
0
0
0
52
0
0
0
0
1
0
0
0
0
0
0
0
0
0
0
0


54
0
1
0
0
0
0
0
0
0
0
0
0
52
0
0
0
0
1
0
0
0
0
0
0
0
0
0
0
0

1.0
1.0
1.0
1.0
1.0
1.0
1.0
1.0
1.0
1.0
1.0
1.0
1.0
1.0
1.0
1.0
1.0
1.0
1.0
1.0
1.0
1.0
1.0
1.0
1.0
1.0
1.0
1.0
1.0
1.0


3
0
1
0
0
0
0
0
0
0
0
0
0
1
0
0
0
0
1
0
0
0
0
0
0
0
0
0
0
0

0.87
0.87
0.87
0.87
0.87
0.87
0.87
0.87
0.87
0.87
0.87
0.87
0.87
0.87
0.87
0.87
0.87
0.87
0.87
0.87
0.87
0.87
0.87
0.87
0.87
0.87
0.87
0.87
0.87
0.87


43
0
0
0
0
0
0
0
0
0
0
0
0
43
0
0
0
0
0
0
0
0
0
0
0
0
0
0
0
0

1.0
1.0
1.0
1.0
1.0
1.0
1.0
1.0
1.0
1.0
1.0
1.0
1.0
1.0
1.0
1.0
1.0
1.0
1.0
1.0
1.0
1.0
1.0
1.0
1.0
1.0
1.0
1.0
1.0
1.0


3
0
0
0
0
0
0
0
0
0
0
0
0
3
0
0
0
0
0
0
0
0
0
0
0
0
0
0
0
0

1.0
1.0
1.0
1.0
1.0
1.0
1.0
1.0
1.0
1.0
1.0
1.0
1.0
1.0
1.0
1.0
1.0
1.0
1.0
1.0
1.0
1.0
1.0
1.0
1.0
1.0
1.0
1.0
1.0
1.0


1
0
0
0
0
0
0
0
0
0
0
0
0
1
0
0
0
0
0
0
0
0
0
0
0
0
0
0
0
0

0.71
0.71
0.71
0.71
0.71
0.71
0.71
0.71
0.71
0.71
0.71
0.71
0.71
0.71
0.71
0.71
0.71
0.71
0.71
0.71
0.71
0.71
0.71
0.71
0.71
0.71
0.71
0.71
0.71
0.71


1
0
0
1
0
0
0
0
0
0
0
0
0
0
0
0
0
0
0
0
0
0
0
0
0
0
0
0
0
0


1
0
0
1
0
0
0
0
0
0
0
0
0
0
0
0
0
0
0
0
0
0
0
0
0
0
0
0
0
0

1.0
1.0
1.0
1.0
1.0
1.0
1.0
1.0
1.0
1.0
1.0
1.0
1.0
1.0
1.0
1.0
1.0
1.0
1.0
1.0
1.0
1.0
1.0
1.0
1.0
1.0
1.0
1.0
1.0
1.0


3
0
0
0
0
0
1
0
0
0
0
0
0
0
0
0
0
0
0
0
0
0
0
0
0
1
0
0
0
1


1
0
0
0
0
0
1
0
0
0
0
0
0
0
0
0
0
0
0
0
0
0
0
0
0
0
0
0
0
0


1
0
0
0
0
0
1
0
0
0
0
0
0
0
0
0
0
0
0
0
0
0
0
0
0
0
0
0
0
0

1.0
1.0
1.0
1.0
1.0
1.0
1.0
1.0
1.0
1.0
1.0
1.0
1.0
1.0
1.0
1.0
1.0
1.0
1.0
1.0
1.0
1.0
1.0
1.0
1.0
1.0
1.0
1.0
1.0
1.0


2
0
0
0
0
0
0
0
0
0
0
0
0
0
0
0
0
0
0
0
0
0
0
0
0
1
0
0
0
1


2
0
0
0
0
0
0
0
0
0
0
0
0
0
0
0
0
0
0
0
0
0
0
0
0
1
0
0
0
1

1.0
1.0
1.0
1.0
1.0
1.0
1.0
1.0
1.0
1.0
1.0
1.0
1.0
1.0
1.0
1.0
1.0
1.0
1.0
1.0
1.0
1.0
1.0
1.0
1.0
1.0
1.0
1.0
1.0
1.0


71
0
0
1
0
0
0
0
0
0
0
0
0
20
50
0
0
0
0
0
0
0
0
0
0
0
0
0
0
0

1.0
1.0
1.0
1.0
1.0
1.0
1.0
1.0
1.0
1.0
1.0
1.0
1.0
1.0
1.0
1.0
1.0
1.0
1.0
1.0
1.0
1.0
1.0
1.0
1.0
1.0
1.0
1.0
1.0
1.0


8
0
0
0
0
0
0
0
0
0
0
0
0
2
6
0
0
0
0
0
0
0
0
0
0
0
0
0
0
0


8
0
0
0
0
0
0
0
0
0
0
0
0
2
6
0
0
0
0
0
0
0
0
0
0
0
0
0
0
0

1.0
1.0
1.0
1.0
1.0
1.0
1.0
1.0
1.0
1.0
1.0
1.0
1.0
1.0
1.0
1.0
1.0
1.0
1.0
1.0
1.0
1.0
1.0
1.0
1.0
1.0
1.0
1.0
1.0
1.0


2
1
0
0
0
0
0
0
0
0
0
0
0
0
0
0
0
0
0
0
0
1
0
0
0
0
0
0
0
0


2
1
0
0
0
0
0
0
0
0
0
0
0
0
0
0
0
0
0
0
0
1
0
0
0
0
0
0
0
0


2
1
0
0
0
0
0
0
0
0
0
0
0
0
0
0
0
0
0
0
0
1
0
0
0
0
0
0
0
0

1.0
1.0
1.0
1.0
1.0
1.0
1.0
1.0
1.0
1.0
1.0
1.0
1.0
1.0
1.0
1.0
1.0
1.0
1.0
1.0
1.0
1.0
1.0
1.0
1.0
1.0
1.0
1.0
1.0
1.0


1
0
1
0
0
0
0
0
0
0
0
0
0
0
0
0
0
0
0
0
0
0
0
0
0
0
0
0
0
0


1
0
1
0
0
0
0
0
0
0
0
0
0
0
0
0
0
0
0
0
0
0
0
0
0
0
0
0
0
0


1
0
1
0
0
0
0
0
0
0
0
0
0
0
0
0
0
0
0
0
0
0
0
0
0
0
0
0
0
0

1.0
1.0
1.0
1.0
1.0
1.0
1.0
1.0
1.0
1.0
1.0
1.0
1.0
1.0
1.0
1.0
1.0
1.0
1.0
1.0
1.0
1.0
1.0
1.0
1.0
1.0
1.0
1.0
1.0
1.0


1
0
0
0
0
1
0
0
0
0
0
0
0
0
0
0
0
0
0
0
0
0
0
0
0
0
0
0
0
0


1
0
0
0
0
1
0
0
0
0
0
0
0
0
0
0
0
0
0
0
0
0
0
0
0
0
0
0
0
0


1
0
0
0
0
1
0
0
0
0
0
0
0
0
0
0
0
0
0
0
0
0
0
0
0
0
0
0
0
0

0.71
0.71
0.71
0.71
0.71
0.71
0.71
0.71
0.71
0.71
0.71
0.71
0.71
0.71
0.71
0.71
0.71
0.71
0.71
0.71
0.71
0.71
0.71
0.71
0.71
0.71
0.71
0.71
0.71
0.71


6
1
1
0
0
0
0
0
0
0
0
0
0
0
0
0
0
0
0
0
0
0
0
0
0
1
0
1
1
1

1.0
1.0
1.0
1.0
1.0
1.0
1.0
1.0
1.0
1.0
1.0
1.0
1.0
1.0
1.0
1.0
1.0
1.0
1.0
1.0
1.0
1.0
1.0
1.0
1.0
1.0
1.0
1.0
1.0
1.0


3
0
0
0
0
0
0
0
0
0
0
0
0
0
0
0
0
0
0
0
0
0
0
0
0
0
0
1
1
1


3
0
0
0
0
0
0
0
0
0
0
0
0
0
0
0
0
0
0
0
0
0
0
0
0
0
0
1
1
1

0.79
0.79
0.79
0.79
0.79
0.79
0.79
0.79
0.79
0.79
0.79
0.79
0.79
0.79
0.79
0.79
0.79
0.79
0.79
0.79
0.79
0.79
0.79
0.79
0.79
0.79
0.79
0.79
0.79
0.79


1
0
0
0
0
0
0
0
0
0
0
0
0
0
0
0
0
0
0
0
0
0
0
0
0
1
0
0
0
0


1
0
0
0
0
0
0
0
0
0
0
0
0
0
0
0
0
0
0
0
0
0
0
0
0
1
0
0
0
0

1.0
1.0
1.0
1.0
1.0
1.0
1.0
1.0
1.0
1.0
1.0
1.0
1.0
1.0
1.0
1.0
1.0
1.0
1.0
1.0
1.0
1.0
1.0
1.0
1.0
1.0
1.0
1.0
1.0
1.0


2
0
0
0
0
0
0
0
1
1
0
0
0
0
0
0
0
0
0
0
0
0
0
0
0
0
0
0
0
0


2
0
0
0
0
0
0
0
1
1
0
0
0
0
0
0
0
0
0
0
0
0
0
0
0
0
0
0
0
0


2
0
0
0
0
0
0
0
1
1
0
0
0
0
0
0
0
0
0
0
0
0
0
0
0
0
0
0
0
0

0.89
0.89
0.89
0.89
0.89
0.89
0.89
0.89
0.89
0.89
0.89
0.89
0.89
0.89
0.89
0.89
0.89
0.89
0.89
0.89
0.89
0.89
0.89
0.89
0.89
0.89
0.89
0.89
0.89
0.89


1
0
0
0
0
0
0
0
0
0
0
0
0
0
0
0
1
0
0
0
0
0
0
0
0
0
0
0
0
0


1
0
0
0
0
0
0
0
0
0
0
0
0
0
0
0
1
0
0
0
0
0
0
0
0
0
0
0
0
0


1
0
0
0
0
0
0
0
0
0
0
0
0
0
0
0
1
0
0
0
0
0
0
0
0
0
0
0
0
0

0.96
0.96
0.96
0.96
0.96
0.96
0.96
0.96
0.96
0.96
0.96
0.96
0.96
0.96
0.96
0.96
0.96
0.96
0.96
0.96
0.96
0.96
0.96
0.96
0.96
0.96
0.96
0.96
0.96
0.96


8
0
0
0
0
0
0
0
0
0
0
0
0
0
0
0
0
0
0
0
0
0
0
0
0
2
2
1
2
1


8
0
0
0
0
0
0
0
0
0
0
0
0
0
0
0
0
0
0
0
0
0
0
0
0
2
2
1
2
1


8
0
0
0
0
0
0
0
0
0
0
0
0
0
0
0
0
0
0
0
0
0
0
0
0
2
2
1
2
1

1.0
1.0
1.0
1.0
1.0
1.0
1.0
1.0
1.0
1.0
1.0
1.0
1.0
1.0
1.0
1.0
1.0
1.0
1.0
1.0
1.0
1.0
1.0
1.0
1.0
1.0
1.0
1.0
1.0
1.0


1
0
0
0
0
0
0
0
0
0
0
0
0
0
0
0
0
0
0
0
0
1
0
0
0
0
0
0
0
0


1
0
0
0
0
0
0
0
0
0
0
0
0
0
0
0
0
0
0
0
0
1
0
0
0
0
0
0
0
0


1
0
0
0
0
0
0
0
0
0
0
0
0
0
0
0
0
0
0
0
0
1
0
0
0
0
0
0
0
0

1.0
1.0
1.0
1.0
1.0
1.0
1.0
1.0
1.0
1.0
1.0
1.0
1.0
1.0
1.0
1.0
1.0
1.0
1.0
1.0
1.0
1.0
1.0
1.0
1.0
1.0
1.0
1.0
1.0
1.0


2
0
0
0
0
0
0
0
0
0
0
0
0
0
1
0
0
0
0
0
0
0
0
0
1
0
0
0
0
0


2
0
0
0
0
0
0
0
0
0
0
0
0
0
1
0
0
0
0
0
0
0
0
0
1
0
0
0
0
0


2
0
0
0
0
0
0
0
0
0
0
0
0
0
1
0
0
0
0
0
0
0
0
0
1
0
0
0
0
0

0.8
0.8
0.8
0.8
0.8
0.8
0.8
0.8
0.8
0.8
0.8
0.8
0.8
0.8
0.8
0.8
0.8
0.8
0.8
0.8
0.8
0.8
0.8
0.8
0.8
0.8
0.8
0.8
0.8
0.8


1
0
0
0
0
0
0
0
0
0
0
0
0
0
0
0
0
0
0
0
0
0
0
0
0
0
0
0
0
1


1
0
0
0
0
0
0
0
0
0
0
0
0
0
0
0
0
0
0
0
0
0
0
0
0
0
0
0
0
1


1
0
0
0
0
0
0
0
0
0
0
0
0
0
0
0
0
0
0
0
0
0
0
0
0
0
0
0
0
1

1.0
1.0
1.0
1.0
1.0
1.0
1.0
1.0
1.0
1.0
1.0
1.0
1.0
1.0
1.0
1.0
1.0
1.0
1.0
1.0
1.0
1.0
1.0
1.0
1.0
1.0
1.0
1.0
1.0
1.0


2104
59
61
47
31
49
55
38
27
31
44
51
38
50
46
123
62
140
98
65
50
12
16
38
34
159
215
186
129
150

1.0
1.0
1.0
1.0
1.0
1.0
1.0
1.0
1.0
1.0
1.0
1.0
1.0
1.0
1.0
1.0
1.0
1.0
1.0
1.0
1.0
1.0
1.0
1.0
1.0
1.0
1.0
1.0
1.0
1.0


231
9
3
5
1
6
8
5
4
3
2
6
3
2
8
20
9
10
5
3
1
3
7
3
0
23
36
20
14
12

0.99
0.99
0.99
0.99
0.99
0.99
0.99
0.99
0.99
0.99
0.99
0.99
0.99
0.99
0.99
0.99
0.99
0.99
0.99
0.99
0.99
0.99
0.99
0.99
0.99
0.99
0.99
0.99
0.99
0.99


7
0
0
0
0
0
0
5
1
1
0
0
0
0
0
0
0
0
0
0
0
0
0
0
0
0
0
0
0
0


7
0
0
0
0
0
0
5
1
1
0
0
0
0
0
0
0
0
0
0
0
0
0
0
0
0
0
0
0
0

1.0
1.0
1.0
1.0
1.0
1.0
1.0
1.0
1.0
1.0
1.0
1.0
1.0
1.0
1.0
1.0
1.0
1.0
1.0
1.0
1.0
1.0
1.0
1.0
1.0
1.0
1.0
1.0
1.0
1.0


72
3
0
2
1
5
3
0
0
0
0
4
1
0
0
0
0
0
0
0
0
0
0
1
0
16
22
12
2
0

1.0
1.0
1.0
1.0
1.0
1.0
1.0
1.0
1.0
1.0
1.0
1.0
1.0
1.0
1.0
1.0
1.0
1.0
1.0
1.0
1.0
1.0
1.0
1.0
1.0
1.0
1.0
1.0
1.0
1.0


25
2
0
2
0
5
2
0
0
0
0
4
0
0
0
0
0
0
0
0
0
0
0
0
0
0
4
4
2
0

1.0
1.0
1.0
1.0
1.0
1.0
1.0
1.0
1.0
1.0
1.0
1.0
1.0
1.0
1.0
1.0
1.0
1.0
1.0
1.0
1.0
1.0
1.0
1.0
1.0
1.0
1.0
1.0
1.0
1.0


10
0
0
0
1
0
1
0
0
0
0
0
1
0
0
0
0
0
0
0
0
0
0
1
0
2
2
2
0
0

1.0
1.0
1.0
1.0
1.0
1.0
1.0
1.0
1.0
1.0
1.0
1.0
1.0
1.0
1.0
1.0
1.0
1.0
1.0
1.0
1.0
1.0
1.0
1.0
1.0
1.0
1.0
1.0
1.0
1.0


1
1
0
0
0
0
0
0
0
0
0
0
0
0
0
0
0
0
0
0
0
0
0
0
0
0
0
0
0
0

1.0
1.0
1.0
1.0
1.0
1.0
1.0
1.0
1.0
1.0
1.0
1.0
1.0
1.0
1.0
1.0
1.0
1.0
1.0
1.0
1.0
1.0
1.0
1.0
1.0
1.0
1.0
1.0
1.0
1.0


5
1
0
0
0
0
0
0
1
0
1
1
0
0
0
0
0
1
0
0
0
0
0
0
0
0
0
0
0
0


2
0
0
0
0
0
0
0
1
0
1
0
0
0
0
0
0
0
0
0
0
0
0
0
0
0
0
0
0
0

1.0
1.0
1.0
1.0
1.0
1.0
1.0
1.0
1.0
1.0
1.0
1.0
1.0
1.0
1.0
1.0
1.0
1.0
1.0
1.0
1.0
1.0
1.0
1.0
1.0
1.0
1.0
1.0
1.0
1.0


3
1
0
0
0
0
0
0
0
0
0
1
0
0
0
0
0
1
0
0
0
0
0
0
0
0
0
0
0
0

1.0
1.0
1.0
1.0
1.0
1.0
1.0
1.0
1.0
1.0
1.0
1.0
1.0
1.0
1.0
1.0
1.0
1.0
1.0
1.0
1.0
1.0
1.0
1.0
1.0
1.0
1.0
1.0
1.0
1.0


37
3
3
1
0
1
0
0
0
0
0
1
0
1
3
14
3
6
0
0
1
0
0
0
0
0
0
0
0
0


33
1
1
1
0
1
0
0
0
0
0
1
0
1
3
14
3
6
0
0
1
0
0
0
0
0
0
0
0
0

1.0
1.0
1.0
1.0
1.0
1.0
1.0
1.0
1.0
1.0
1.0
1.0
1.0
1.0
1.0
1.0
1.0
1.0
1.0
1.0
1.0
1.0
1.0
1.0
1.0
1.0
1.0
1.0
1.0
1.0


4
2
2
0
0
0
0
0
0
0
0
0
0
0
0
0
0
0
0
0
0
0
0
0
0
0
0
0
0
0

1.0
1.0
1.0
1.0
1.0
1.0
1.0
1.0
1.0
1.0
1.0
1.0
1.0
1.0
1.0
1.0
1.0
1.0
1.0
1.0
1.0
1.0
1.0
1.0
1.0
1.0
1.0
1.0
1.0
1.0


5
0
0
0
0
0
5
0
0
0
0
0
0
0
0
0
0
0
0
0
0
0
0
0
0
0
0
0
0
0


5
0
0
0
0
0
5
0
0
0
0
0
0
0
0
0
0
0
0
0
0
0
0
0
0
0
0
0
0
0

1.0
1.0
1.0
1.0
1.0
1.0
1.0
1.0
1.0
1.0
1.0
1.0
1.0
1.0
1.0
1.0
1.0
1.0
1.0
1.0
1.0
1.0
1.0
1.0
1.0
1.0
1.0
1.0
1.0
1.0


9
0
0
0
0
0
0
0
2
2
1
0
2
0
0
0
0
0
0
0
0
0
0
2
0
0
0
0
0
0


9
0
0
0
0
0
0
0
2
2
1
0
2
0
0
0
0
0
0
0
0
0
0
2
0
0
0
0
0
0

1.0
1.0
1.0
1.0
1.0
1.0
1.0
1.0
1.0
1.0
1.0
1.0
1.0
1.0
1.0
1.0
1.0
1.0
1.0
1.0
1.0
1.0
1.0
1.0
1.0
1.0
1.0
1.0
1.0
1.0


13
2
0
0
0
0
0
0
0
0
0
0
0
0
0
0
0
0
0
3
0
0
0
0
0
0
4
3
1
0

1.0
1.0
1.0
1.0
1.0
1.0
1.0
1.0
1.0
1.0
1.0
1.0
1.0
1.0
1.0
1.0
1.0
1.0
1.0
1.0
1.0
1.0
1.0
1.0
1.0
1.0
1.0
1.0
1.0
1.0


2
0
0
2
0
0
0
0
0
0
0
0
0
0
0
0
0
0
0
0
0
0
0
0
0
0
0
0
0
0


2
0
0
2
0
0
0
0
0
0
0
0
0
0
0
0
0
0
0
0
0
0
0
0
0
0
0
0
0
0

1.0
1.0
1.0
1.0
1.0
1.0
1.0
1.0
1.0
1.0
1.0
1.0
1.0
1.0
1.0
1.0
1.0
1.0
1.0
1.0
1.0
1.0
1.0
1.0
1.0
1.0
1.0
1.0
1.0
1.0


19
0
0
0
0
0
0
0
0
0
0
0
0
0
0
0
0
0
0
0
0
3
0
0
0
5
5
0
4
2

0.73
0.73
0.73
0.73
0.73
0.73
0.73
0.73
0.73
0.73
0.73
0.73
0.73
0.73
0.73
0.73
0.73
0.73
0.73
0.73
0.73
0.73
0.73
0.73
0.73
0.73
0.73
0.73
0.73
0.73


16
0
0
0
0
0
0
0
0
0
0
0
0
0
0
0
0
0
0
0
0
3
0
0
0
4
3
0
4
2

1.0
1.0
1.0
1.0
1.0
1.0
1.0
1.0
1.0
1.0
1.0
1.0
1.0
1.0
1.0
1.0
1.0
1.0
1.0
1.0
1.0
1.0
1.0
1.0
1.0
1.0
1.0
1.0
1.0
1.0


11
0
0
0
0
0
0
0
0
0
0
0
0
0
0
3
2
1
0
0
0
0
0
0
0
1
0
0
2
2


11
0
0
0
0
0
0
0
0
0
0
0
0
0
0
3
2
1
0
0
0
0
0
0
0
1
0
0
2
2

1.0
1.0
1.0
1.0
1.0
1.0
1.0
1.0
1.0
1.0
1.0
1.0
1.0
1.0
1.0
1.0
1.0
1.0
1.0
1.0
1.0
1.0
1.0
1.0
1.0
1.0
1.0
1.0
1.0
1.0


14
0
0
0
0
0
0
0
0
0
0
0
0
1
5
0
0
0
1
0
0
0
7
0
0
0
0
0
0
0

1.0
1.0
1.0
1.0
1.0
1.0
1.0
1.0
1.0
1.0
1.0
1.0
1.0
1.0
1.0
1.0
1.0
1.0
1.0
1.0
1.0
1.0
1.0
1.0
1.0
1.0
1.0
1.0
1.0
1.0


10
0
0
0
0
0
0
0
0
0
0
0
0
1
3
0
0
0
1
0
0
0
5
0
0
0
0
0
0
0

0.9
0.9
0.9
0.9
0.9
0.9
0.9
0.9
0.9
0.9
0.9
0.9
0.9
0.9
0.9
0.9
0.9
0.9
0.9
0.9
0.9
0.9
0.9
0.9
0.9
0.9
0.9
0.9
0.9
0.9


4
0
0
0
0
0
0
0
0
0
0
0
0
0
0
0
0
0
4
0
0
0
0
0
0
0
0
0
0
0


4
0
0
0
0
0
0
0
0
0
0
0
0
0
0
0
0
0
4
0
0
0
0
0
0
0
0
0
0
0

1.0
1.0
1.0
1.0
1.0
1.0
1.0
1.0
1.0
1.0
1.0
1.0
1.0
1.0
1.0
1.0
1.0
1.0
1.0
1.0
1.0
1.0
1.0
1.0
1.0
1.0
1.0
1.0
1.0
1.0


14
0
0
0
0
0
0
0
0
0
0
0
0
0
0
0
0
0
0
0
0
0
0
0
0
0
0
4
4
6


14
0
0
0
0
0
0
0
0
0
0
0
0
0
0
0
0
0
0
0
0
0
0
0
0
0
0
4
4
6

1.0
1.0
1.0
1.0
1.0
1.0
1.0
1.0
1.0
1.0
1.0
1.0
1.0
1.0
1.0
1.0
1.0
1.0
1.0
1.0
1.0
1.0
1.0
1.0
1.0
1.0
1.0
1.0
1.0
1.0


4
0
0
0
0
0
0
0
0
0
0
0
0
0
0
0
2
0
0
0
0
0
0
0
0
0
0
0
0
2


4
0
0
0
0
0
0
0
0
0
0
0
0
0
0
0
2
0
0
0
0
0
0
0
0
0
0
0
0
2

1.0
1.0
1.0
1.0
1.0
1.0
1.0
1.0
1.0
1.0
1.0
1.0
1.0
1.0
1.0
1.0
1.0
1.0
1.0
1.0
1.0
1.0
1.0
1.0
1.0
1.0
1.0
1.0
1.0
1.0


1
0
0
0
0
0
0
0
0
0
0
0
0
0
0
0
0
0
0
0
0
0
0
0
0
0
1
0
0
0


1
0
0
0
0
0
0
0
0
0
0
0
0
0
0
0
0
0
0
0
0
0
0
0
0
0
1
0
0
0

1.0
1.0
1.0
1.0
1.0
1.0
1.0
1.0
1.0
1.0
1.0
1.0
1.0
1.0
1.0
1.0
1.0
1.0
1.0
1.0
1.0
1.0
1.0
1.0
1.0
1.0
1.0
1.0
1.0
1.0


330
3
2
0
1
0
1
15
1
3
3
3
33
9
0
17
8
30
41
39
27
3
4
33
0
14
14
7
5
14

1.0
1.0
1.0
1.0
1.0
1.0
1.0
1.0
1.0
1.0
1.0
1.0
1.0
1.0
1.0
1.0
1.0
1.0
1.0
1.0
1.0
1.0
1.0
1.0
1.0
1.0
1.0
1.0
1.0
1.0


89
0
0
0
0
0
1
5
0
0
1
1
8
0
0
0
0
0
17
28
17
2
1
8
0
0
0
0
0
0

0.78
0.78
0.78
0.78
0.78
0.78
0.78
0.78
0.78
0.78
0.78
0.78
0.78
0.78
0.78
0.78
0.78
0.78
0.78
0.78
0.78
0.78
0.78
0.78
0.78
0.78
0.78
0.78
0.78
0.78


85
0
0
0
0
0
1
5
0
0
1
1
6
0
0
0
0
0
17
28
17
2
1
6
0
0
0
0
0
0

1.0
1.0
1.0
1.0
1.0
1.0
1.0
1.0
1.0
1.0
1.0
1.0
1.0
1.0
1.0
1.0
1.0
1.0
1.0
1.0
1.0
1.0
1.0
1.0
1.0
1.0
1.0
1.0
1.0
1.0


3
0
0
0
0
0
0
3
0
0
0
0
0
0
0
0
0
0
0
0
0
0
0
0
0
0
0
0
0
0


3
0
0
0
0
0
0
3
0
0
0
0
0
0
0
0
0
0
0
0
0
0
0
0
0
0
0
0
0
0

1.0
1.0
1.0
1.0
1.0
1.0
1.0
1.0
1.0
1.0
1.0
1.0
1.0
1.0
1.0
1.0
1.0
1.0
1.0
1.0
1.0
1.0
1.0
1.0
1.0
1.0
1.0
1.0
1.0
1.0


18
0
0
0
0
0
0
4
0
1
2
0
2
0
0
0
0
0
1
0
1
0
0
2
0
0
5
0
0
0


9
0
0
0
0
0
0
4
0
0
0
0
0
0
0
0
0
0
0
0
0
0
0
0
0
0
5
0
0
0

1.0
1.0
1.0
1.0
1.0
1.0
1.0
1.0
1.0
1.0
1.0
1.0
1.0
1.0
1.0
1.0
1.0
1.0
1.0
1.0
1.0
1.0
1.0
1.0
1.0
1.0
1.0
1.0
1.0
1.0


1
0
0
0
0
0
0
0
0
0
1
0
0
0
0
0
0
0
0
0
0
0
0
0
0
0
0
0
0
0

0.83
0.83
0.83
0.83
0.83
0.83
0.83
0.83
0.83
0.83
0.83
0.83
0.83
0.83
0.83
0.83
0.83
0.83
0.83
0.83
0.83
0.83
0.83
0.83
0.83
0.83
0.83
0.83
0.83
0.83


8
0
0
0
0
0
0
0
0
1
1
0
2
0
0
0
0
0
1
0
1
0
0
2
0
0
0
0
0
0

0.95
0.95
0.95
0.95
0.95
0.95
0.95
0.95
0.95
0.95
0.95
0.95
0.95
0.95
0.95
0.95
0.95
0.95
0.95
0.95
0.95
0.95
0.95
0.95
0.95
0.95
0.95
0.95
0.95
0.95


40
0
0
0
0
0
0
0
1
2
0
0
0
7
0
6
7
5
5
2
1
1
1
0
0
0
0
1
1
0

1.0
1.0
1.0
1.0
1.0
1.0
1.0
1.0
1.0
1.0
1.0
1.0
1.0
1.0
1.0
1.0
1.0
1.0
1.0
1.0
1.0
1.0
1.0
1.0
1.0
1.0
1.0
1.0
1.0
1.0


15
0
0
0
0
0
0
0
1
0
0
0
0
7
0
0
1
1
1
0
1
1
1
0
0
0
0
0
1
0

0.97
0.97
0.97
0.97
0.97
0.97
0.97
0.97
0.97
0.97
0.97
0.97
0.97
0.97
0.97
0.97
0.97
0.97
0.97
0.97
0.97
0.97
0.97
0.97
0.97
0.97
0.97
0.97
0.97
0.97


2
0
0
0
0
0
0
0
0
2
0
0
0
0
0
0
0
0
0
0
0
0
0
0
0
0
0
0
0
0

0.96
0.96
0.96
0.96
0.96
0.96
0.96
0.96
0.96
0.96
0.96
0.96
0.96
0.96
0.96
0.96
0.96
0.96
0.96
0.96
0.96
0.96
0.96
0.96
0.96
0.96
0.96
0.96
0.96
0.96


10
0
0
0
0
0
0
0
0
0
0
0
0
0
0
5
5
0
0
0
0
0
0
0
0
0
0
0
0
0

1.0
1.0
1.0
1.0
1.0
1.0
1.0
1.0
1.0
1.0
1.0
1.0
1.0
1.0
1.0
1.0
1.0
1.0
1.0
1.0
1.0
1.0
1.0
1.0
1.0
1.0
1.0
1.0
1.0
1.0


1
0
0
0
0
0
0
0
0
0
0
0
0
0
0
0
0
0
0
0
0
0
0
0
0
0
0
1
0
0

1.0
1.0
1.0
1.0
1.0
1.0
1.0
1.0
1.0
1.0
1.0
1.0
1.0
1.0
1.0
1.0
1.0
1.0
1.0
1.0
1.0
1.0
1.0
1.0
1.0
1.0
1.0
1.0
1.0
1.0


10
0
0
0
0
0
0
0
0
0
0
0
0
0
0
0
0
4
4
2
0
0
0
0
0
0
0
0
0
0

1.0
1.0
1.0
1.0
1.0
1.0
1.0
1.0
1.0
1.0
1.0
1.0
1.0
1.0
1.0
1.0
1.0
1.0
1.0
1.0
1.0
1.0
1.0
1.0
1.0
1.0
1.0
1.0
1.0
1.0


4
0
1
0
1
0
0
0
0
0
0
0
0
0
0
0
0
0
0
0
0
0
0
0
0
0
0
1
1
0


4
0
1
0
1
0
0
0
0
0
0
0
0
0
0
0
0
0
0
0
0
0
0
0
0
0
0
1
1
0

1.0
1.0
1.0
1.0
1.0
1.0
1.0
1.0
1.0
1.0
1.0
1.0
1.0
1.0
1.0
1.0
1.0
1.0
1.0
1.0
1.0
1.0
1.0
1.0
1.0
1.0
1.0
1.0
1.0
1.0


1
0
0
0
0
0
0
0
0
0
0
1
0
0
0
0
0
0
0
0
0
0
0
0
0
0
0
0
0
0


1
0
0
0
0
0
0
0
0
0
0
1
0
0
0
0
0
0
0
0
0
0
0
0
0
0
0
0
0
0

1.0
1.0
1.0
1.0
1.0
1.0
1.0
1.0
1.0
1.0
1.0
1.0
1.0
1.0
1.0
1.0
1.0
1.0
1.0
1.0
1.0
1.0
1.0
1.0
1.0
1.0
1.0
1.0
1.0
1.0


4
0
1
0
0
0
0
0
0
0
0
1
0
0
0
0
0
0
0
0
0
0
0
0
0
0
2
0
0
0


4
0
1
0
0
0
0
0
0
0
0
1
0
0
0
0
0
0
0
0
0
0
0
0
0
0
2
0
0
0

0.99
0.99
0.99
0.99
0.99
0.99
0.99
0.99
0.99
0.99
0.99
0.99
0.99
0.99
0.99
0.99
0.99
0.99
0.99
0.99
0.99
0.99
0.99
0.99
0.99
0.99
0.99
0.99
0.99
0.99


16
0
0
0
0
0
0
1
0
0
0
0
0
1
0
1
0
0
2
4
4
0
2
0
0
0
0
0
1
0

0.81
0.81
0.81
0.81
0.81
0.81
0.81
0.81
0.81
0.81
0.81
0.81
0.81
0.81
0.81
0.81
0.81
0.81
0.81
0.81
0.81
0.81
0.81
0.81
0.81
0.81
0.81
0.81
0.81
0.81


13
0
0
0
0
0
0
0
0
0
0
0
0
1
0
1
0
0
2
3
3
0
2
0
0
0
0
0
1
0

0.74
0.74
0.74
0.74
0.74
0.74
0.74
0.74
0.74
0.74
0.74
0.74
0.74
0.74
0.74
0.74
0.74
0.74
0.74
0.74
0.74
0.74
0.74
0.74
0.74
0.74
0.74
0.74
0.74
0.74


1
1
0
0
0
0
0
0
0
0
0
0
0
0
0
0
0
0
0
0
0
0
0
0
0
0
0
0
0
0


1
1
0
0
0
0
0
0
0
0
0
0
0
0
0
0
0
0
0
0
0
0
0
0
0
0
0
0
0
0

1.0
1.0
1.0
1.0
1.0
1.0
1.0
1.0
1.0
1.0
1.0
1.0
1.0
1.0
1.0
1.0
1.0
1.0
1.0
1.0
1.0
1.0
1.0
1.0
1.0
1.0
1.0
1.0
1.0
1.0


1
1
0
0
0
0
0
0
0
0
0
0
0
0
0
0
0
0
0
0
0
0
0
0
0
0
0
0
0
0


1
1
0
0
0
0
0
0
0
0
0
0
0
0
0
0
0
0
0
0
0
0
0
0
0
0
0
0
0
0

1.0
1.0
1.0
1.0
1.0
1.0
1.0
1.0
1.0
1.0
1.0
1.0
1.0
1.0
1.0
1.0
1.0
1.0
1.0
1.0
1.0
1.0
1.0
1.0
1.0
1.0
1.0
1.0
1.0
1.0


1
1
0
0
0
0
0
0
0
0
0
0
0
0
0
0
0
0
0
0
0
0
0
0
0
0
0
0
0
0


1
1
0
0
0
0
0
0
0
0
0
0
0
0
0
0
0
0
0
0
0
0
0
0
0
0
0
0
0
0

0.95
0.95
0.95
0.95
0.95
0.95
0.95
0.95
0.95
0.95
0.95
0.95
0.95
0.95
0.95
0.95
0.95
0.95
0.95
0.95
0.95
0.95
0.95
0.95
0.95
0.95
0.95
0.95
0.95
0.95


12
0
0
0
0
0
0
0
0
0
0
0
0
0
0
0
0
6
4
1
0
0
0
0
0
0
1
0
0
0


12
0
0
0
0
0
0
0
0
0
0
0
0
0
0
0
0
6
4
1
0
0
0
0
0
0
1
0
0
0

0.99
0.99
0.99
0.99
0.99
0.99
0.99
0.99
0.99
0.99
0.99
0.99
0.99
0.99
0.99
0.99
0.99
0.99
0.99
0.99
0.99
0.99
0.99
0.99
0.99
0.99
0.99
0.99
0.99
0.99


9
0
0
0
0
0
0
0
0
0
0
0
0
0
0
0
0
0
0
0
0
0
0
0
0
0
0
0
0
9


9
0
0
0
0
0
0
0
0
0
0
0
0
0
0
0
0
0
0
0
0
0
0
0
0
0
0
0
0
9

0.99
0.99
0.99
0.99
0.99
0.99
0.99
0.99
0.99
0.99
0.99
0.99
0.99
0.99
0.99
0.99
0.99
0.99
0.99
0.99
0.99
0.99
0.99
0.99
0.99
0.99
0.99
0.99
0.99
0.99


5
0
0
0
0
0
0
0
0
0
0
0
0
0
0
0
0
1
2
0
2
0
0
0
0
0
0
0
0
0


4
0
0
0
0
0
0
0
0
0
0
0
0
0
0
0
0
1
1
0
2
0
0
0
0
0
0
0
0
0

1.0
1.0
1.0
1.0
1.0
1.0
1.0
1.0
1.0
1.0
1.0
1.0
1.0
1.0
1.0
1.0
1.0
1.0
1.0
1.0
1.0
1.0
1.0
1.0
1.0
1.0
1.0
1.0
1.0
1.0


1
0
0
0
0
0
0
0
0
0
0
0
0
0
0
0
0
0
1
0
0
0
0
0
0
0
0
0
0
0

0.98
0.98
0.98
0.98
0.98
0.98
0.98
0.98
0.98
0.98
0.98
0.98
0.98
0.98
0.98
0.98
0.98
0.98
0.98
0.98
0.98
0.98
0.98
0.98
0.98
0.98
0.98
0.98
0.98
0.98


4
0
0
0
0
0
0
0
0
0
0
0
0
0
0
0
0
0
0
0
0
0
0
0
0
4
0
0
0
0


4
0
0
0
0
0
0
0
0
0
0
0
0
0
0
0
0
0
0
0
0
0
0
0
0
4
0
0
0
0

1.0
1.0
1.0
1.0
1.0
1.0
1.0
1.0
1.0
1.0
1.0
1.0
1.0
1.0
1.0
1.0
1.0
1.0
1.0
1.0
1.0
1.0
1.0
1.0
1.0
1.0
1.0
1.0
1.0
1.0


1
0
0
0
0
0
0
0
0
0
0
0
0
0
0
1
0
0
0
0
0
0
0
0
0
0
0
0
0
0


1
0
0
0
0
0
0
0
0
0
0
0
0
0
0
1
0
0
0
0
0
0
0
0
0
0
0
0
0
0

1.0
1.0
1.0
1.0
1.0
1.0
1.0
1.0
1.0
1.0
1.0
1.0
1.0
1.0
1.0
1.0
1.0
1.0
1.0
1.0
1.0
1.0
1.0
1.0
1.0
1.0
1.0
1.0
1.0
1.0


2
0
0
0
0
0
0
0
0
0
0
0
0
0
0
0
0
0
0
0
0
0
0
0
0
1
0
1
0
0

1.0
1.0
1.0
1.0
1.0
1.0
1.0
1.0
1.0
1.0
1.0
1.0
1.0
1.0
1.0
1.0
1.0
1.0
1.0
1.0
1.0
1.0
1.0
1.0
1.0
1.0
1.0
1.0
1.0
1.0


9
0
0
0
0
0
0
0
0
0
0
0
0
0
0
0
0
0
0
0
0
0
0
0
0
6
3
0
0
0


9
0
0
0
0
0
0
0
0
0
0
0
0
0
0
0
0
0
0
0
0
0
0
0
0
6
3
0
0
0

1.0
1.0
1.0
1.0
1.0
1.0
1.0
1.0
1.0
1.0
1.0
1.0
1.0
1.0
1.0
1.0
1.0
1.0
1.0
1.0
1.0
1.0
1.0
1.0
1.0
1.0
1.0
1.0
1.0
1.0


1
0
0
0
0
0
0
0
0
0
0
0
0
0
0
0
0
0
0
0
0
0
0
0
0
1
0
0
0
0


1
0
0
0
0
0
0
0
0
0
0
0
0
0
0
0
0
0
0
0
0
0
0
0
0
1
0
0
0
0

1.0
1.0
1.0
1.0
1.0
1.0
1.0
1.0
1.0
1.0
1.0
1.0
1.0
1.0
1.0
1.0
1.0
1.0
1.0
1.0
1.0
1.0
1.0
1.0
1.0
1.0
1.0
1.0
1.0
1.0


2
0
0
0
0
0
0
0
0
0
0
0
0
0
0
0
0
0
0
0
0
0
0
0
0
0
0
0
0
2

0.9
0.9
0.9
0.9
0.9
0.9
0.9
0.9
0.9
0.9
0.9
0.9
0.9
0.9
0.9
0.9
0.9
0.9
0.9
0.9
0.9
0.9
0.9
0.9
0.9
0.9
0.9
0.9
0.9
0.9


1
0
0
0
0
0
0
0
0
0
0
0
0
0
0
0
0
0
0
0
0
0
0
0
0
0
0
0
0
1

1.0
1.0
1.0
1.0
1.0
1.0
1.0
1.0
1.0
1.0
1.0
1.0
1.0
1.0
1.0
1.0
1.0
1.0
1.0
1.0
1.0
1.0
1.0
1.0
1.0
1.0
1.0
1.0
1.0
1.0


1
0
0
0
0
0
0
0
0
0
0
0
0
0
0
0
0
0
0
0
0
0
0
0
0
0
0
1
0
0


1
0
0
0
0
0
0
0
0
0
0
0
0
0
0
0
0
0
0
0
0
0
0
0
0
0
0
1
0
0

1.0
1.0
1.0
1.0
1.0
1.0
1.0
1.0
1.0
1.0
1.0
1.0
1.0
1.0
1.0
1.0
1.0
1.0
1.0
1.0
1.0
1.0
1.0
1.0
1.0
1.0
1.0
1.0
1.0
1.0


2
0
0
0
0
0
0
0
0
0
0
0
0
0
0
2
0
0
0
0
0
0
0
0
0
0
0
0
0
0

0.87
0.87
0.87
0.87
0.87
0.87
0.87
0.87
0.87
0.87
0.87
0.87
0.87
0.87
0.87
0.87
0.87
0.87
0.87
0.87
0.87
0.87
0.87
0.87
0.87
0.87
0.87
0.87
0.87
0.87


14
0
0
0
0
0
0
0
0
0
0
0
7
0
0
0
0
0
0
0
0
0
0
7
0
0
0
0
0
0

0.91
0.91
0.91
0.91
0.91
0.91
0.91
0.91
0.91
0.91
0.91
0.91
0.91
0.91
0.91
0.91
0.91
0.91
0.91
0.91
0.91
0.91
0.91
0.91
0.91
0.91
0.91
0.91
0.91
0.91


2
0
0
0
0
0
0
0
0
0
0
0
1
0
0
0
0
0
0
0
0
0
0
1
0
0
0
0
0
0

0.84
0.84
0.84
0.84
0.84
0.84
0.84
0.84
0.84
0.84
0.84
0.84
0.84
0.84
0.84
0.84
0.84
0.84
0.84
0.84
0.84
0.84
0.84
0.84
0.84
0.84
0.84
0.84
0.84
0.84


2
0
0
0
0
0
0
0
0
0
0
0
0
0
0
1
1
0
0
0
0
0
0
0
0
0
0
0
0
0

0.85
0.85
0.85
0.85
0.85
0.85
0.85
0.85
0.85
0.85
0.85
0.85
0.85
0.85
0.85
0.85
0.85
0.85
0.85
0.85
0.85
0.85
0.85
0.85
0.85
0.85
0.85
0.85
0.85
0.85


255
9
11
8
7
12
11
1
1
3
5
7
0
9
5
2
6
3
5
3
10
3
0
0
6
22
33
27
24
22


55
3
1
1
4
8
4
1
1
1
1
1
0
0
2
2
0
0
0
0
0
0
0
0
2
4
6
5
4
4

0.74
0.74
0.74
0.74
0.74
0.74
0.74
0.74
0.74
0.74
0.74
0.74
0.74
0.74
0.74
0.74
0.74
0.74
0.74
0.74
0.74
0.74
0.74
0.74
0.74
0.74
0.74
0.74
0.74
0.74


50
3
1
1
4
7
4
1
1
1
1
1
0
0
2
2
0
0
0
0
0
0
0
0
2
4
6
4
3
2

1.0
1.0
1.0
1.0
1.0
1.0
1.0
1.0
1.0
1.0
1.0
1.0
1.0
1.0
1.0
1.0
1.0
1.0
1.0
1.0
1.0
1.0
1.0
1.0
1.0
1.0
1.0
1.0
1.0
1.0


1
0
0
0
0
0
0
0
0
0
0
0
0
0
0
0
0
0
0
0
0
0
0
0
0
0
0
1
0
0

1.0
1.0
1.0
1.0
1.0
1.0
1.0
1.0
1.0
1.0
1.0
1.0
1.0
1.0
1.0
1.0
1.0
1.0
1.0
1.0
1.0
1.0
1.0
1.0
1.0
1.0
1.0
1.0
1.0
1.0


3
0
0
0
0
0
0
0
0
0
0
0
0
0
0
0
0
0
0
0
0
0
0
0
0
0
0
0
1
2

1.0
1.0
1.0
1.0
1.0
1.0
1.0
1.0
1.0
1.0
1.0
1.0
1.0
1.0
1.0
1.0
1.0
1.0
1.0
1.0
1.0
1.0
1.0
1.0
1.0
1.0
1.0
1.0
1.0
1.0


24
1
1
1
1
1
1
0
0
1
1
1
0
0
1
0
0
1
0
0
0
0
0
0
1
0
4
1
5
2


24
1
1
1
1
1
1
0
0
1
1
1
0
0
1
0
0
1
0
0
0
0
0
0
1
0
4
1
5
2

1.0
1.0
1.0
1.0
1.0
1.0
1.0
1.0
1.0
1.0
1.0
1.0
1.0
1.0
1.0
1.0
1.0
1.0
1.0
1.0
1.0
1.0
1.0
1.0
1.0
1.0
1.0
1.0
1.0
1.0


21
4
4
0
0
0
0
0
0
0
0
4
0
0
0
0
0
0
5
0
4
0
0
0
0
0
0
0
0
0


11
4
3
0
0
0
0
0
0
0
0
4
0
0
0
0
0
0
0
0
0
0
0
0
0
0
0
0
0
0

1.0
1.0
1.0
1.0
1.0
1.0
1.0
1.0
1.0
1.0
1.0
1.0
1.0
1.0
1.0
1.0
1.0
1.0
1.0
1.0
1.0
1.0
1.0
1.0
1.0
1.0
1.0
1.0
1.0
1.0


1
0
1
0
0
0
0
0
0
0
0
0
0
0
0
0
0
0
0
0
0
0
0
0
0
0
0
0
0
0

1.0
1.0
1.0
1.0
1.0
1.0
1.0
1.0
1.0
1.0
1.0
1.0
1.0
1.0
1.0
1.0
1.0
1.0
1.0
1.0
1.0
1.0
1.0
1.0
1.0
1.0
1.0
1.0
1.0
1.0


9
0
0
0
0
0
0
0
0
0
0
0
0
0
0
0
0
0
5
0
4
0
0
0
0
0
0
0
0
0

1.0
1.0
1.0
1.0
1.0
1.0
1.0
1.0
1.0
1.0
1.0
1.0
1.0
1.0
1.0
1.0
1.0
1.0
1.0
1.0
1.0
1.0
1.0
1.0
1.0
1.0
1.0
1.0
1.0
1.0


17
0
0
0
0
0
1
0
0
0
0
0
0
0
0
0
0
0
0
0
0
0
0
0
0
0
4
3
5
4


17
0
0
0
0
0
1
0
0
0
0
0
0
0
0
0
0
0
0
0
0
0
0
0
0
0
4
3
5
4

1.0
1.0
1.0
1.0
1.0
1.0
1.0
1.0
1.0
1.0
1.0
1.0
1.0
1.0
1.0
1.0
1.0
1.0
1.0
1.0
1.0
1.0
1.0
1.0
1.0
1.0
1.0
1.0
1.0
1.0


4
0
0
0
2
0
2
0
0
0
0
0
0
0
0
0
0
0
0
0
0
0
0
0
0
0
0
0
0
0


4
0
0
0
2
0
2
0
0
0
0
0
0
0
0
0
0
0
0
0
0
0
0
0
0
0
0
0
0
0

1.0
1.0
1.0
1.0
1.0
1.0
1.0
1.0
1.0
1.0
1.0
1.0
1.0
1.0
1.0
1.0
1.0
1.0
1.0
1.0
1.0
1.0
1.0
1.0
1.0
1.0
1.0
1.0
1.0
1.0


13
0
0
1
0
0
0
0
0
0
0
0
0
0
0
0
0
0
0
0
0
0
0
0
0
6
6
0
0
0


13
0
0
1
0
0
0
0
0
0
0
0
0
0
0
0
0
0
0
0
0
0
0
0
0
6
6
0
0
0

1.0
1.0
1.0
1.0
1.0
1.0
1.0
1.0
1.0
1.0
1.0
1.0
1.0
1.0
1.0
1.0
1.0
1.0
1.0
1.0
1.0
1.0
1.0
1.0
1.0
1.0
1.0
1.0
1.0
1.0


4
0
0
1
0
0
0
0
0
0
1
1
0
0
0
0
0
0
0
0
0
0
0
0
0
0
0
0
0
1


4
0
0
1
0
0
0
0
0
0
1
1
0
0
0
0
0
0
0
0
0
0
0
0
0
0
0
0
0
1

1.0
1.0
1.0
1.0
1.0
1.0
1.0
1.0
1.0
1.0
1.0
1.0
1.0
1.0
1.0
1.0
1.0
1.0
1.0
1.0
1.0
1.0
1.0
1.0
1.0
1.0
1.0
1.0
1.0
1.0


37
0
0
0
0
0
1
0
0
1
0
0
0
0
0
0
0
0
0
0
6
3
0
0
0
0
6
11
4
5

1.0
1.0
1.0
1.0
1.0
1.0
1.0
1.0
1.0
1.0
1.0
1.0
1.0
1.0
1.0
1.0
1.0
1.0
1.0
1.0
1.0
1.0
1.0
1.0
1.0
1.0
1.0
1.0
1.0
1.0


26
0
0
0
0
0
1
0
0
0
0
0
0
0
0
0
0
0
0
0
0
0
0
0
0
0
6
10
4
5

1.0
1.0
1.0
1.0
1.0
1.0
1.0
1.0
1.0
1.0
1.0
1.0
1.0
1.0
1.0
1.0
1.0
1.0
1.0
1.0
1.0
1.0
1.0
1.0
1.0
1.0
1.0
1.0
1.0
1.0


10
0
0
0
0
0
0
0
0
1
0
0
0
0
0
0
0
0
0
0
6
3
0
0
0
0
0
0
0
0

1.0
1.0
1.0
1.0
1.0
1.0
1.0
1.0
1.0
1.0
1.0
1.0
1.0
1.0
1.0
1.0
1.0
1.0
1.0
1.0
1.0
1.0
1.0
1.0
1.0
1.0
1.0
1.0
1.0
1.0


1
0
0
0
0
0
0
0
0
0
1
0
0
0
0
0
0
0
0
0
0
0
0
0
0
0
0
0
0
0

1.0
1.0
1.0
1.0
1.0
1.0
1.0
1.0
1.0
1.0
1.0
1.0
1.0
1.0
1.0
1.0
1.0
1.0
1.0
1.0
1.0
1.0
1.0
1.0
1.0
1.0
1.0
1.0
1.0
1.0


1
0
0
0
0
0
0
0
0
0
1
0
0
0
0
0
0
0
0
0
0
0
0
0
0
0
0
0
0
0


1
0
0
0
0
0
0
0
0
0
1
0
0
0
0
0
0
0
0
0
0
0
0
0
0
0
0
0
0
0

1.0
1.0
1.0
1.0
1.0
1.0
1.0
1.0
1.0
1.0
1.0
1.0
1.0
1.0
1.0
1.0
1.0
1.0
1.0
1.0
1.0
1.0
1.0
1.0
1.0
1.0
1.0
1.0
1.0
1.0


11
0
1
2
0
1
0
0
0
0
0
0
0
0
0
0
2
2
0
3
0
0
0
0
0
0
0
0
0
0


11
0
1
2
0
1
0
0
0
0
0
0
0
0
0
0
2
2
0
3
0
0
0
0
0
0
0
0
0
0

1.0
1.0
1.0
1.0
1.0
1.0
1.0
1.0
1.0
1.0
1.0
1.0
1.0
1.0
1.0
1.0
1.0
1.0
1.0
1.0
1.0
1.0
1.0
1.0
1.0
1.0
1.0
1.0
1.0
1.0


2
0
0
1
0
1
0
0
0
0
0
0
0
0
0
0
0
0
0
0
0
0
0
0
0
0
0
0
0
0


2
0
0
1
0
1
0
0
0
0
0
0
0
0
0
0
0
0
0
0
0
0
0
0
0
0
0
0
0
0

1.0
1.0
1.0
1.0
1.0
1.0
1.0
1.0
1.0
1.0
1.0
1.0
1.0
1.0
1.0
1.0
1.0
1.0
1.0
1.0
1.0
1.0
1.0
1.0
1.0
1.0
1.0
1.0
1.0
1.0


4
0
1
0
0
1
1
0
0
0
0
0
0
0
0
0
0
0
0
0
0
0
0
0
1
0
0
0
0
0


4
0
1
0
0
1
1
0
0
0
0
0
0
0
0
0
0
0
0
0
0
0
0
0
1
0
0
0
0
0

0.98
0.98
0.98
0.98
0.98
0.98
0.98
0.98
0.98
0.98
0.98
0.98
0.98
0.98
0.98
0.98
0.98
0.98
0.98
0.98
0.98
0.98
0.98
0.98
0.98
0.98
0.98
0.98
0.98
0.98


15
1
2
1
0
0
0
0
0
0
0
0
0
9
2
0
0
0
0
0
0
0
0
0
0
0
0
0
0
0


15
1
2
1
0
0
0
0
0
0
0
0
0
9
2
0
0
0
0
0
0
0
0
0
0
0
0
0
0
0

1.0
1.0
1.0
1.0
1.0
1.0
1.0
1.0
1.0
1.0
1.0
1.0
1.0
1.0
1.0
1.0
1.0
1.0
1.0
1.0
1.0
1.0
1.0
1.0
1.0
1.0
1.0
1.0
1.0
1.0


2
0
1
0
0
0
0
0
0
0
0
0
0
0
0
0
0
0
0
0
0
0
0
0
1
0
0
0
0
0


2
0
1
0
0
0
0
0
0
0
0
0
0
0
0
0
0
0
0
0
0
0
0
0
1
0
0
0
0
0

1.0
1.0
1.0
1.0
1.0
1.0
1.0
1.0
1.0
1.0
1.0
1.0
1.0
1.0
1.0
1.0
1.0
1.0
1.0
1.0
1.0
1.0
1.0
1.0
1.0
1.0
1.0
1.0
1.0
1.0


3
0
0
0
0
0
1
0
0
0
0
0
0
0
0
0
0
0
0
0
0
0
0
0
0
1
1
0
0
0


3
0
0
0
0
0
1
0
0
0
0
0
0
0
0
0
0
0
0
0
0
0
0
0
0
1
1
0
0
0

1.0
1.0
1.0
1.0
1.0
1.0
1.0
1.0
1.0
1.0
1.0
1.0
1.0
1.0
1.0
1.0
1.0
1.0
1.0
1.0
1.0
1.0
1.0
1.0
1.0
1.0
1.0
1.0
1.0
1.0


6
0
0
0
0
0
0
0
0
0
0
0
0
0
0
0
1
0
0
0
0
0
0
0
0
1
1
1
1
1


6
0
0
0
0
0
0
0
0
0
0
0
0
0
0
0
1
0
0
0
0
0
0
0
0
1
1
1
1
1

1.0
1.0
1.0
1.0
1.0
1.0
1.0
1.0
1.0
1.0
1.0
1.0
1.0
1.0
1.0
1.0
1.0
1.0
1.0
1.0
1.0
1.0
1.0
1.0
1.0
1.0
1.0
1.0
1.0
1.0


2
0
0
0
0
0
0
0
0
0
0
0
0
0
0
0
0
0
0
0
0
0
0
0
0
0
0
1
1
0


2
0
0
0
0
0
0
0
0
0
0
0
0
0
0
0
0
0
0
0
0
0
0
0
0
0
0
1
1
0

1.0
1.0
1.0
1.0
1.0
1.0
1.0
1.0
1.0
1.0
1.0
1.0
1.0
1.0
1.0
1.0
1.0
1.0
1.0
1.0
1.0
1.0
1.0
1.0
1.0
1.0
1.0
1.0
1.0
1.0


3
0
0
0
0
0
0
0
0
0
0
0
0
0
0
0
0
0
0
0
0
0
0
0
1
2
0
0
0
0


1
0
0
0
0
0
0
0
0
0
0
0
0
0
0
0
0
0
0
0
0
0
0
0
1
0
0
0
0
0

1.0
1.0
1.0
1.0
1.0
1.0
1.0
1.0
1.0
1.0
1.0
1.0
1.0
1.0
1.0
1.0
1.0
1.0
1.0
1.0
1.0
1.0
1.0
1.0
1.0
1.0
1.0
1.0
1.0
1.0


2
0
0
0
0
0
0
0
0
0
0
0
0
0
0
0
0
0
0
0
0
0
0
0
0
2
0
0
0
0

1.0
1.0
1.0
1.0
1.0
1.0
1.0
1.0
1.0
1.0
1.0
1.0
1.0
1.0
1.0
1.0
1.0
1.0
1.0
1.0
1.0
1.0
1.0
1.0
1.0
1.0
1.0
1.0
1.0
1.0


29
0
0
0
0
0
0
0
0
0
0
0
0
0
0
0
3
0
0
0
0
0
0
0
0
8
5
5
3
5


29
0
0
0
0
0
0
0
0
0
0
0
0
0
0
0
3
0
0
0
0
0
0
0
0
8
5
5
3
5

1.0
1.0
1.0
1.0
1.0
1.0
1.0
1.0
1.0
1.0
1.0
1.0
1.0
1.0
1.0
1.0
1.0
1.0
1.0
1.0
1.0
1.0
1.0
1.0
1.0
1.0
1.0
1.0
1.0
1.0


1
0
0
0
0
0
0
0
0
0
0
0
0
0
0
0
0
0
0
0
0
0
0
0
0
0
0
0
1
0


1
0
0
0
0
0
0
0
0
0
0
0
0
0
0
0
0
0
0
0
0
0
0
0
0
0
0
0
1
0

0.97
0.97
0.97
0.97
0.97
0.97
0.97
0.97
0.97
0.97
0.97
0.97
0.97
0.97
0.97
0.97
0.97
0.97
0.97
0.97
0.97
0.97
0.97
0.97
0.97
0.97
0.97
0.97
0.97
0.97


37
1
2
2
0
0
1
0
0
1
1
1
0
1
0
1
0
16
1
3
1
1
0
0
1
0
1
2
0
0

0.7
0.7
0.7
0.7
0.7
0.7
0.7
0.7
0.7
0.7
0.7
0.7
0.7
0.7
0.7
0.7
0.7
0.7
0.7
0.7
0.7
0.7
0.7
0.7
0.7
0.7
0.7
0.7
0.7
0.7


24
0
0
0
0
0
0
0
0
1
0
1
0
0
0
1
0
14
1
1
1
1
0
0
1
0
1
1
0
0


2
0
0
0
0
0
0
0
0
1
0
1
0
0
0
0
0
0
0
0
0
0
0
0
0
0
0
0
0
0

0.99
0.99
0.99
0.99
0.99
0.99
0.99
0.99
0.99
0.99
0.99
0.99
0.99
0.99
0.99
0.99
0.99
0.99
0.99
0.99
0.99
0.99
0.99
0.99
0.99
0.99
0.99
0.99
0.99
0.99


22
0
0
0
0
0
0
0
0
0
0
0
0
0
0
1
0
14
1
1
1
1
0
0
1
0
1
1
0
0

1.0
1.0
1.0
1.0
1.0
1.0
1.0
1.0
1.0
1.0
1.0
1.0
1.0
1.0
1.0
1.0
1.0
1.0
1.0
1.0
1.0
1.0
1.0
1.0
1.0
1.0
1.0
1.0
1.0
1.0


5
0
2
2
0
0
0
0
0
0
0
0
0
1
0
0
0
0
0
0
0
0
0
0
0
0
0
0
0
0


5
0
2
2
0
0
0
0
0
0
0
0
0
1
0
0
0
0
0
0
0
0
0
0
0
0
0
0
0
0

1.0
1.0
1.0
1.0
1.0
1.0
1.0
1.0
1.0
1.0
1.0
1.0
1.0
1.0
1.0
1.0
1.0
1.0
1.0
1.0
1.0
1.0
1.0
1.0
1.0
1.0
1.0
1.0
1.0
1.0


6
1
0
0
0
0
1
0
0
0
1
0
0
0
0
0
0
1
0
2
0
0
0
0
0
0
0
0
0
0


6
1
0
0
0
0
1
0
0
0
1
0
0
0
0
0
0
1
0
2
0
0
0
0
0
0
0
0
0
0

1.0
1.0
1.0
1.0
1.0
1.0
1.0
1.0
1.0
1.0
1.0
1.0
1.0
1.0
1.0
1.0
1.0
1.0
1.0
1.0
1.0
1.0
1.0
1.0
1.0
1.0
1.0
1.0
1.0
1.0


1
0
0
0
0
0
0
0
0
0
0
0
0
0
0
0
0
0
0
0
0
0
0
0
0
0
0
1
0
0

0.73
0.73
0.73
0.73
0.73
0.73
0.73
0.73
0.73
0.73
0.73
0.73
0.73
0.73
0.73
0.73
0.73
0.73
0.73
0.73
0.73
0.73
0.73
0.73
0.73
0.73
0.73
0.73
0.73
0.73


28
8
4
1
0
0
0
0
0
4
3
1
0
1
2
0
0
0
4
0
0
0
0
0
0
0
0
0
0
0


24
5
3
1
0
0
0
0
0
4
3
1
0
1
2
0
0
0
4
0
0
0
0
0
0
0
0
0
0
0


24
5
3
1
0
0
0
0
0
4
3
1
0
1
2
0
0
0
4
0
0
0
0
0
0
0
0
0
0
0

1.0
1.0
1.0
1.0
1.0
1.0
1.0
1.0
1.0
1.0
1.0
1.0
1.0
1.0
1.0
1.0
1.0
1.0
1.0
1.0
1.0
1.0
1.0
1.0
1.0
1.0
1.0
1.0
1.0
1.0


4
3
1
0
0
0
0
0
0
0
0
0
0
0
0
0
0
0
0
0
0
0
0
0
0
0
0
0
0
0

1.0
1.0
1.0
1.0
1.0
1.0
1.0
1.0
1.0
1.0
1.0
1.0
1.0
1.0
1.0
1.0
1.0
1.0
1.0
1.0
1.0
1.0
1.0
1.0
1.0
1.0
1.0
1.0
1.0
1.0


91
5
3
5
3
4
2
0
3
4
3
3
0
1
0
9
0
3
2
7
1
0
0
0
1
7
6
6
6
7

0.98
0.98
0.98
0.98
0.98
0.98
0.98
0.98
0.98
0.98
0.98
0.98
0.98
0.98
0.98
0.98
0.98
0.98
0.98
0.98
0.98
0.98
0.98
0.98
0.98
0.98
0.98
0.98
0.98
0.98


14
2
1
2
0
2
1
0
0
0
1
2
0
0
0
0
0
0
0
0
1
0
0
0
1
0
0
0
0
1


14
2
1
2
0
2
1
0
0
0
1
2
0
0
0
0
0
0
0
0
1
0
0
0
1
0
0
0
0
1

1.0
1.0
1.0
1.0
1.0
1.0
1.0
1.0
1.0
1.0
1.0
1.0
1.0
1.0
1.0
1.0
1.0
1.0
1.0
1.0
1.0
1.0
1.0
1.0
1.0
1.0
1.0
1.0
1.0
1.0


21
2
1
0
1
2
0
0
0
1
1
1
0
0
0
1
0
2
2
2
0
0
0
0
0
1
2
0
1
1


21
2
1
0
1
2
0
0
0
1
1
1
0
0
0
1
0
2
2
2
0
0
0
0
0
1
2
0
1
1

1.0
1.0
1.0
1.0
1.0
1.0
1.0
1.0
1.0
1.0
1.0
1.0
1.0
1.0
1.0
1.0
1.0
1.0
1.0
1.0
1.0
1.0
1.0
1.0
1.0
1.0
1.0
1.0
1.0
1.0


12
0
0
2
1
0
0
0
0
0
0
0
0
1
0
0
0
1
0
1
0
0
0
0
0
1
2
2
1
0


7
0
0
2
1
0
0
0
0
0
0
0
0
0
0
0
0
0
0
0
0
0
0
0
0
1
2
0
1
0

1.0
1.0
1.0
1.0
1.0
1.0
1.0
1.0
1.0
1.0
1.0
1.0
1.0
1.0
1.0
1.0
1.0
1.0
1.0
1.0
1.0
1.0
1.0
1.0
1.0
1.0
1.0
1.0
1.0
1.0


3
0
0
0
0
0
0
0
0
0
0
0
0
1
0
0
0
1
0
1
0
0
0
0
0
0
0
0
0
0

1.0
1.0
1.0
1.0
1.0
1.0
1.0
1.0
1.0
1.0
1.0
1.0
1.0
1.0
1.0
1.0
1.0
1.0
1.0
1.0
1.0
1.0
1.0
1.0
1.0
1.0
1.0
1.0
1.0
1.0


2
0
0
0
0
0
0
0
0
0
0
0
0
0
0
0
0
0
0
0
0
0
0
0
0
0
0
2
0
0

1.0
1.0
1.0
1.0
1.0
1.0
1.0
1.0
1.0
1.0
1.0
1.0
1.0
1.0
1.0
1.0
1.0
1.0
1.0
1.0
1.0
1.0
1.0
1.0
1.0
1.0
1.0
1.0
1.0
1.0


2
0
0
0
0
0
0
0
2
0
0
0
0
0
0
0
0
0
0
0
0
0
0
0
0
0
0
0
0
0


2
0
0
0
0
0
0
0
2
0
0
0
0
0
0
0
0
0
0
0
0
0
0
0
0
0
0
0
0
0

1.0
1.0
1.0
1.0
1.0
1.0
1.0
1.0
1.0
1.0
1.0
1.0
1.0
1.0
1.0
1.0
1.0
1.0
1.0
1.0
1.0
1.0
1.0
1.0
1.0
1.0
1.0
1.0
1.0
1.0


4
1
0
0
0
0
0
0
0
3
0
0
0
0
0
0
0
0
0
0
0
0
0
0
0
0
0
0
0
0


4
1
0
0
0
0
0
0
0
3
0
0
0
0
0
0
0
0
0
0
0
0
0
0
0
0
0
0
0
0

1.0
1.0
1.0
1.0
1.0
1.0
1.0
1.0
1.0
1.0
1.0
1.0
1.0
1.0
1.0
1.0
1.0
1.0
1.0
1.0
1.0
1.0
1.0
1.0
1.0
1.0
1.0
1.0
1.0
1.0


2
0
0
0
0
0
0
0
1
0
1
0
0
0
0
0
0
0
0
0
0
0
0
0
0
0
0
0
0
0


2
0
0
0
0
0
0
0
1
0
1
0
0
0
0
0
0
0
0
0
0
0
0
0
0
0
0
0
0
0

0.75
0.75
0.75
0.75
0.75
0.75
0.75
0.75
0.75
0.75
0.75
0.75
0.75
0.75
0.75
0.75
0.75
0.75
0.75
0.75
0.75
0.75
0.75
0.75
0.75
0.75
0.75
0.75
0.75
0.75


30
0
0
1
1
0
0
0
0
0
0
0
0
0
0
8
0
0
0
0
0
0
0
0
0
5
2
4
4
5


30
0
0
1
1
0
0
0
0
0
0
0
0
0
0
8
0
0
0
0
0
0
0
0
0
5
2
4
4
5

1.0
1.0
1.0
1.0
1.0
1.0
1.0
1.0
1.0
1.0
1.0
1.0
1.0
1.0
1.0
1.0
1.0
1.0
1.0
1.0
1.0
1.0
1.0
1.0
1.0
1.0
1.0
1.0
1.0
1.0


1
0
1
0
0
0
0
0
0
0
0
0
0
0
0
0
0
0
0
0
0
0
0
0
0
0
0
0
0
0


1
0
1
0
0
0
0
0
0
0
0
0
0
0
0
0
0
0
0
0
0
0
0
0
0
0
0
0
0
0

1.0
1.0
1.0
1.0
1.0
1.0
1.0
1.0
1.0
1.0
1.0
1.0
1.0
1.0
1.0
1.0
1.0
1.0
1.0
1.0
1.0
1.0
1.0
1.0
1.0
1.0
1.0
1.0
1.0
1.0


1
0
0
0
0
0
1
0
0
0
0
0
0
0
0
0
0
0
0
0
0
0
0
0
0
0
0
0
0
0


1
0
0
0
0
0
1
0
0
0
0
0
0
0
0
0
0
0
0
0
0
0
0
0
0
0
0
0
0
0

1.0
1.0
1.0
1.0
1.0
1.0
1.0
1.0
1.0
1.0
1.0
1.0
1.0
1.0
1.0
1.0
1.0
1.0
1.0
1.0
1.0
1.0
1.0
1.0
1.0
1.0
1.0
1.0
1.0
1.0


2
0
0
0
0
0
0
0
0
0
0
0
0
0
0
0
0
0
0
2
0
0
0
0
0
0
0
0
0
0


2
0
0
0
0
0
0
0
0
0
0
0
0
0
0
0
0
0
0
2
0
0
0
0
0
0
0
0
0
0

1.0
1.0
1.0
1.0
1.0
1.0
1.0
1.0
1.0
1.0
1.0
1.0
1.0
1.0
1.0
1.0
1.0
1.0
1.0
1.0
1.0
1.0
1.0
1.0
1.0
1.0
1.0
1.0
1.0
1.0


179
4
6
2
0
4
6
0
0
0
1
5
0
5
1
21
1
0
0
0
0
0
1
0
3
34
27
26
8
24

0.98
0.98
0.98
0.98
0.98
0.98
0.98
0.98
0.98
0.98
0.98
0.98
0.98
0.98
0.98
0.98
0.98
0.98
0.98
0.98
0.98
0.98
0.98
0.98
0.98
0.98
0.98
0.98
0.98
0.98


4
1
1
0
0
0
0
0
0
0
0
1
0
0
0
0
0
0
0
0
0
0
0
0
1
0
0
0
0
0


4
1
1
0
0
0
0
0
0
0
0
1
0
0
0
0
0
0
0
0
0
0
0
0
1
0
0
0
0
0

1.0
1.0
1.0
1.0
1.0
1.0
1.0
1.0
1.0
1.0
1.0
1.0
1.0
1.0
1.0
1.0
1.0
1.0
1.0
1.0
1.0
1.0
1.0
1.0
1.0
1.0
1.0
1.0
1.0
1.0


74
2
4
1
0
1
5
0
0
0
0
3
0
5
0
1
0
0
0
0
0
0
0
0
1
26
13
12
0
0


74
2
4
1
0
1
5
0
0
0
0
3
0
5
0
1
0
0
0
0
0
0
0
0
1
26
13
12
0
0

1.0
1.0
1.0
1.0
1.0
1.0
1.0
1.0
1.0
1.0
1.0
1.0
1.0
1.0
1.0
1.0
1.0
1.0
1.0
1.0
1.0
1.0
1.0
1.0
1.0
1.0
1.0
1.0
1.0
1.0


4
0
0
1
0
2
0
0
0
0
0
0
0
0
0
0
0
0
0
0
0
0
0
0
0
0
0
1
0
0

1.0
1.0
1.0
1.0
1.0
1.0
1.0
1.0
1.0
1.0
1.0
1.0
1.0
1.0
1.0
1.0
1.0
1.0
1.0
1.0
1.0
1.0
1.0
1.0
1.0
1.0
1.0
1.0
1.0
1.0


13
0
0
0
0
1
1
0
0
0
0
0
0
0
0
0
0
0
0
0
0
0
0
0
0
3
1
0
1
6


2
0
0
0
0
1
1
0
0
0
0
0
0
0
0
0
0
0
0
0
0
0
0
0
0
0
0
0
0
0

1.0
1.0
1.0
1.0
1.0
1.0
1.0
1.0
1.0
1.0
1.0
1.0
1.0
1.0
1.0
1.0
1.0
1.0
1.0
1.0
1.0
1.0
1.0
1.0
1.0
1.0
1.0
1.0
1.0
1.0


11
0
0
0
0
0
0
0
0
0
0
0
0
0
0
0
0
0
0
0
0
0
0
0
0
3
1
0
1
6

1.0
1.0
1.0
1.0
1.0
1.0
1.0
1.0
1.0
1.0
1.0
1.0
1.0
1.0
1.0
1.0
1.0
1.0
1.0
1.0
1.0
1.0
1.0
1.0
1.0
1.0
1.0
1.0
1.0
1.0


8
1
0
0
0
0
0
0
0
0
1
0
0
0
0
0
1
0
0
0
0
0
0
0
0
2
1
1
0
1


7
1
0
0
0
0
0
0
0
0
1
0
0
0
0
0
0
0
0
0
0
0
0
0
0
2
1
1
0
1

1.0
1.0
1.0
1.0
1.0
1.0
1.0
1.0
1.0
1.0
1.0
1.0
1.0
1.0
1.0
1.0
1.0
1.0
1.0
1.0
1.0
1.0
1.0
1.0
1.0
1.0
1.0
1.0
1.0
1.0


1
0
0
0
0
0
0
0
0
0
0
0
0
0
0
0
1
0
0
0
0
0
0
0
0
0
0
0
0
0

1.0
1.0
1.0
1.0
1.0
1.0
1.0
1.0
1.0
1.0
1.0
1.0
1.0
1.0
1.0
1.0
1.0
1.0
1.0
1.0
1.0
1.0
1.0
1.0
1.0
1.0
1.0
1.0
1.0
1.0


4
0
1
0
0
0
0
0
0
0
0
1
0
0
0
2
0
0
0
0
0
0
0
0
0
0
0
0
0
0


4
0
1
0
0
0
0
0
0
0
0
1
0
0
0
2
0
0
0
0
0
0
0
0
0
0
0
0
0
0

1.0
1.0
1.0
1.0
1.0
1.0
1.0
1.0
1.0
1.0
1.0
1.0
1.0
1.0
1.0
1.0
1.0
1.0
1.0
1.0
1.0
1.0
1.0
1.0
1.0
1.0
1.0
1.0
1.0
1.0


5
0
0
0
0
0
0
0
0
0
0
0
0
0
0
1
0
0
0
0
0
0
0
0
0
1
1
1
1
0


5
0
0
0
0
0
0
0
0
0
0
0
0
0
0
1
0
0
0
0
0
0
0
0
0
1
1
1
1
0

0.93
0.93
0.93
0.93
0.93
0.93
0.93
0.93
0.93
0.93
0.93
0.93
0.93
0.93
0.93
0.93
0.93
0.93
0.93
0.93
0.93
0.93
0.93
0.93
0.93
0.93
0.93
0.93
0.93
0.93


5
0
0
0
0
0
0
0
0
0
0
0
0
0
0
0
0
0
0
0
0
0
0
0
0
1
2
1
1
0

1.0
1.0
1.0
1.0
1.0
1.0
1.0
1.0
1.0
1.0
1.0
1.0
1.0
1.0
1.0
1.0
1.0
1.0
1.0
1.0
1.0
1.0
1.0
1.0
1.0
1.0
1.0
1.0
1.0
1.0


14
0
0
0
0
0
0
0
0
0
0
0
0
0
0
5
0
0
0
0
0
0
1
0
0
0
0
5
3
0

1.0
1.0
1.0
1.0
1.0
1.0
1.0
1.0
1.0
1.0
1.0
1.0
1.0
1.0
1.0
1.0
1.0
1.0
1.0
1.0
1.0
1.0
1.0
1.0
1.0
1.0
1.0
1.0
1.0
1.0


6
0
0
0
0
0
0
0
0
0
0
0
0
0
0
3
0
0
0
0
0
0
1
0
0
0
0
1
1
0

1.0
1.0
1.0
1.0
1.0
1.0
1.0
1.0
1.0
1.0
1.0
1.0
1.0
1.0
1.0
1.0
1.0
1.0
1.0
1.0
1.0
1.0
1.0
1.0
1.0
1.0
1.0
1.0
1.0
1.0


3
0
0
0
0
0
0
0
0
0
0
0
0
0
0
1
0
0
0
0
0
0
0
0
0
0
0
1
1
0

0.75
0.75
0.75
0.75
0.75
0.75
0.75
0.75
0.75
0.75
0.75
0.75
0.75
0.75
0.75
0.75
0.75
0.75
0.75
0.75
0.75
0.75
0.75
0.75
0.75
0.75
0.75
0.75
0.75
0.75


1
0
0
0
0
0
0
0
0
0
0
0
0
0
0
0
0
0
0
0
0
0
0
0
1
0
0
0
0
0


1
0
0
0
0
0
0
0
0
0
0
0
0
0
0
0
0
0
0
0
0
0
0
0
1
0
0
0
0
0

1.0
1.0
1.0
1.0
1.0
1.0
1.0
1.0
1.0
1.0
1.0
1.0
1.0
1.0
1.0
1.0
1.0
1.0
1.0
1.0
1.0
1.0
1.0
1.0
1.0
1.0
1.0
1.0
1.0
1.0


2
0
0
0
0
0
0
0
0
0
0
0
0
0
0
0
0
0
0
0
0
0
0
0
0
0
0
0
2
0


2
0
0
0
0
0
0
0
0
0
0
0
0
0
0
0
0
0
0
0
0
0
0
0
0
0
0
0
2
0

1.0
1.0
1.0
1.0
1.0
1.0
1.0
1.0
1.0
1.0
1.0
1.0
1.0
1.0
1.0
1.0
1.0
1.0
1.0
1.0
1.0
1.0
1.0
1.0
1.0
1.0
1.0
1.0
1.0
1.0


15
0
0
0
0
0
0
0
0
0
0
0
0
0
0
5
0
0
0
0
0
0
0
0
0
0
3
0
0
7

0.84
0.84
0.84
0.84
0.84
0.84
0.84
0.84
0.84
0.84
0.84
0.84
0.84
0.84
0.84
0.84
0.84
0.84
0.84
0.84
0.84
0.84
0.84
0.84
0.84
0.84
0.84
0.84
0.84
0.84


10
0
0
0
0
0
0
0
0
0
0
0
0
0
0
4
0
0
0
0
0
0
0
0
0
0
2
0
0
4

1.0
1.0
1.0
1.0
1.0
1.0
1.0
1.0
1.0
1.0
1.0
1.0
1.0
1.0
1.0
1.0
1.0
1.0
1.0
1.0
1.0
1.0
1.0
1.0
1.0
1.0
1.0
1.0
1.0
1.0


3
0
0
0
0
0
0
0
0
0
0
0
0
0
0
1
0
0
0
0
0
0
0
0
0
0
0
0
0
2

0.8
0.8
0.8
0.8
0.8
0.8
0.8
0.8
0.8
0.8
0.8
0.8
0.8
0.8
0.8
0.8
0.8
0.8
0.8
0.8
0.8
0.8
0.8
0.8
0.8
0.8
0.8
0.8
0.8
0.8


1
0
0
0
0
0
0
0
0
0
0
0
0
0
1
0
0
0
0
0
0
0
0
0
0
0
0
0
0
0


1
0
0
0
0
0
0
0
0
0
0
0
0
0
1
0
0
0
0
0
0
0
0
0
0
0
0
0
0
0

1.0
1.0
1.0
1.0
1.0
1.0
1.0
1.0
1.0
1.0
1.0
1.0
1.0
1.0
1.0
1.0
1.0
1.0
1.0
1.0
1.0
1.0
1.0
1.0
1.0
1.0
1.0
1.0
1.0
1.0


48
1
1
1
2
0
0
0
0
0
0
0
0
1
5
4
7
12
0
0
0
0
0
0
1
0
0
0
5
8


12
1
1
1
2
0
0
0
0
0
0
0
0
1
5
0
0
0
0
0
0
0
0
0
1
0
0
0
0
0


12
1
1
1
2
0
0
0
0
0
0
0
0
1
5
0
0
0
0
0
0
0
0
0
1
0
0
0
0
0

0.99
0.99
0.99
0.99
0.99
0.99
0.99
0.99
0.99
0.99
0.99
0.99
0.99
0.99
0.99
0.99
0.99
0.99
0.99
0.99
0.99
0.99
0.99
0.99
0.99
0.99
0.99
0.99
0.99
0.99


36
0
0
0
0
0
0
0
0
0
0
0
0
0
0
4
7
12
0
0
0
0
0
0
0
0
0
0
5
8

1.0
1.0
1.0
1.0
1.0
1.0
1.0
1.0
1.0
1.0
1.0
1.0
1.0
1.0
1.0
1.0
1.0
1.0
1.0
1.0
1.0
1.0
1.0
1.0
1.0
1.0
1.0
1.0
1.0
1.0


25
0
0
0
0
0
0
0
0
0
0
0
0
0
0
2
4
10
0
0
0
0
0
0
0
0
0
0
3
6

1.0
1.0
1.0
1.0
1.0
1.0
1.0
1.0
1.0
1.0
1.0
1.0
1.0
1.0
1.0
1.0
1.0
1.0
1.0
1.0
1.0
1.0
1.0
1.0
1.0
1.0
1.0
1.0
1.0
1.0


6
0
0
1
1
1
1
0
0
0
0
0
0
0
1
0
0
0
0
0
0
0
0
0
1
0
0
0
0
0


6
0
0
1
1
1
1
0
0
0
0
0
0
0
1
0
0
0
0
0
0
0
0
0
1
0
0
0
0
0


6
0
0
1
1
1
1
0
0
0
0
0
0
0
1
0
0
0
0
0
0
0
0
0
1
0
0
0
0
0

0.99
0.99
0.99
0.99
0.99
0.99
0.99
0.99
0.99
0.99
0.99
0.99
0.99
0.99
0.99
0.99
0.99
0.99
0.99
0.99
0.99
0.99
0.99
0.99
0.99
0.99
0.99
0.99
0.99
0.99


27
0
2
0
1
2
1
0
2
1
1
1
0
0
4
5
2
2
0
0
0
0
0
0
0
0
1
2
0
0


3
0
1
0
1
0
1
0
0
0
0
0
0
0
0
0
0
0
0
0
0
0
0
0
0
0
0
0
0
0


3
0
1
0
1
0
1
0
0
0
0
0
0
0
0
0
0
0
0
0
0
0
0
0
0
0
0
0
0
0

0.76
0.76
0.76
0.76
0.76
0.76
0.76
0.76
0.76
0.76
0.76
0.76
0.76
0.76
0.76
0.76
0.76
0.76
0.76
0.76
0.76
0.76
0.76
0.76
0.76
0.76
0.76
0.76
0.76
0.76


2
0
0
0
0
0
0
0
1
1
0
0
0
0
0
0
0
0
0
0
0
0
0
0
0
0
0
0
0
0


2
0
0
0
0
0
0
0
1
1
0
0
0
0
0
0
0
0
0
0
0
0
0
0
0
0
0
0
0
0

1.0
1.0
1.0
1.0
1.0
1.0
1.0
1.0
1.0
1.0
1.0
1.0
1.0
1.0
1.0
1.0
1.0
1.0
1.0
1.0
1.0
1.0
1.0
1.0
1.0
1.0
1.0
1.0
1.0
1.0


2
0
0
0
0
0
0
0
1
0
1
0
0
0
0
0
0
0
0
0
0
0
0
0
0
0
0
0
0
0


2
0
0
0
0
0
0
0
1
0
1
0
0
0
0
0
0
0
0
0
0
0
0
0
0
0
0
0
0
0

1.0
1.0
1.0
1.0
1.0
1.0
1.0
1.0
1.0
1.0
1.0
1.0
1.0
1.0
1.0
1.0
1.0
1.0
1.0
1.0
1.0
1.0
1.0
1.0
1.0
1.0
1.0
1.0
1.0
1.0


2
0
1
0
0
0
0
0
0
0
0
1
0
0
0
0
0
0
0
0
0
0
0
0
0
0
0
0
0
0


2
0
1
0
0
0
0
0
0
0
0
1
0
0
0
0
0
0
0
0
0
0
0
0
0
0
0
0
0
0

1.0
1.0
1.0
1.0
1.0
1.0
1.0
1.0
1.0
1.0
1.0
1.0
1.0
1.0
1.0
1.0
1.0
1.0
1.0
1.0
1.0
1.0
1.0
1.0
1.0
1.0
1.0
1.0
1.0
1.0


5
0
0
0
0
2
0
0
0
0
0
0
0
0
0
1
0
0
0
0
0
0
0
0
0
0
0
2
0
0


5
0
0
0
0
2
0
0
0
0
0
0
0
0
0
1
0
0
0
0
0
0
0
0
0
0
0
2
0
0

1.0
1.0
1.0
1.0
1.0
1.0
1.0
1.0
1.0
1.0
1.0
1.0
1.0
1.0
1.0
1.0
1.0
1.0
1.0
1.0
1.0
1.0
1.0
1.0
1.0
1.0
1.0
1.0
1.0
1.0


8
0
0
0
0
0
0
0
0
0
0
0
0
0
0
4
2
2
0
0
0
0
0
0
0
0
0
0
0
0


8
0
0
0
0
0
0
0
0
0
0
0
0
0
0
4
2
2
0
0
0
0
0
0
0
0
0
0
0
0

1.0
1.0
1.0
1.0
1.0
1.0
1.0
1.0
1.0
1.0
1.0
1.0
1.0
1.0
1.0
1.0
1.0
1.0
1.0
1.0
1.0
1.0
1.0
1.0
1.0
1.0
1.0
1.0
1.0
1.0


1
0
0
0
0
0
0
0
0
0
0
0
0
0
0
0
0
0
0
0
0
0
0
0
0
0
1
0
0
0


1
0
0
0
0
0
0
0
0
0
0
0
0
0
0
0
0
0
0
0
0
0
0
0
0
0
1
0
0
0

1.0
1.0
1.0
1.0
1.0
1.0
1.0
1.0
1.0
1.0
1.0
1.0
1.0
1.0
1.0
1.0
1.0
1.0
1.0
1.0
1.0
1.0
1.0
1.0
1.0
1.0
1.0
1.0
1.0
1.0


4
0
0
0
0
0
0
0
0
0
0
0
0
0
4
0
0
0
0
0
0
0
0
0
0
0
0
0
0
0

0.97
0.97
0.97
0.97
0.97
0.97
0.97
0.97
0.97
0.97
0.97
0.97
0.97
0.97
0.97
0.97
0.97
0.97
0.97
0.97
0.97
0.97
0.97
0.97
0.97
0.97
0.97
0.97
0.97
0.97


52
1
1
0
0
0
0
0
1
1
1
1
0
0
0
5
4
1
0
1
0
0
0
0
0
6
8
8
7
6


52
1
1
0
0
0
0
0
1
1
1
1
0
0
0
5
4
1
0
1
0
0
0
0
0
6
8
8
7
6

1.0
1.0
1.0
1.0
1.0
1.0
1.0
1.0
1.0
1.0
1.0
1.0
1.0
1.0
1.0
1.0
1.0
1.0
1.0
1.0
1.0
1.0
1.0
1.0
1.0
1.0
1.0
1.0
1.0
1.0


6
1
1
0
0
0
0
0
1
1
1
1
0
0
0
0
0
0
0
0
0
0
0
0
0
0
0
0
0
0

0.85
0.85
0.85
0.85
0.85
0.85
0.85
0.85
0.85
0.85
0.85
0.85
0.85
0.85
0.85
0.85
0.85
0.85
0.85
0.85
0.85
0.85
0.85
0.85
0.85
0.85
0.85
0.85
0.85
0.85


13
1
1
1
1
1
3
0
0
0
0
1
0
0
0
0
0
2
0
0
0
0
0
0
2
0
0
0
0
0


13
1
1
1
1
1
3
0
0
0
0
1
0
0
0
0
0
2
0
0
0
0
0
0
2
0
0
0
0
0


13
1
1
1
1
1
3
0
0
0
0
1
0
0
0
0
0
2
0
0
0
0
0
0
2
0
0
0
0
0

1.0
1.0
1.0
1.0
1.0
1.0
1.0
1.0
1.0
1.0
1.0
1.0
1.0
1.0
1.0
1.0
1.0
1.0
1.0
1.0
1.0
1.0
1.0
1.0
1.0
1.0
1.0
1.0
1.0
1.0


81
1
4
1
0
4
0
1
3
2
3
4
0
1
3
10
2
3
1
0
1
1
0
0
0
8
6
5
14
3

0.71
0.71
0.71
0.71
0.71
0.71
0.71
0.71
0.71
0.71
0.71
0.71
0.71
0.71
0.71
0.71
0.71
0.71
0.71
0.71
0.71
0.71
0.71
0.71
0.71
0.71
0.71
0.71
0.71
0.71


2
0
0
0
0
0
0
0
0
0
1
0
0
0
0
0
0
1
0
0
0
0
0
0
0
0
0
0
0
0


2
0
0
0
0
0
0
0
0
0
1
0
0
0
0
0
0
1
0
0
0
0
0
0
0
0
0
0
0
0

0.99
0.99
0.99
0.99
0.99
0.99
0.99
0.99
0.99
0.99
0.99
0.99
0.99
0.99
0.99
0.99
0.99
0.99
0.99
0.99
0.99
0.99
0.99
0.99
0.99
0.99
0.99
0.99
0.99
0.99


2
0
0
0
0
0
0
0
1
1
0
0
0
0
0
0
0
0
0
0
0
0
0
0
0
0
0
0
0
0


2
0
0
0
0
0
0
0
1
1
0
0
0
0
0
0
0
0
0
0
0
0
0
0
0
0
0
0
0
0

1.0
1.0
1.0
1.0
1.0
1.0
1.0
1.0
1.0
1.0
1.0
1.0
1.0
1.0
1.0
1.0
1.0
1.0
1.0
1.0
1.0
1.0
1.0
1.0
1.0
1.0
1.0
1.0
1.0
1.0


1
0
0
0
0
0
0
0
1
0
0
0
0
0
0
0
0
0
0
0
0
0
0
0
0
0
0
0
0
0


1
0
0
0
0
0
0
0
1
0
0
0
0
0
0
0
0
0
0
0
0
0
0
0
0
0
0
0
0
0

1.0
1.0
1.0
1.0
1.0
1.0
1.0
1.0
1.0
1.0
1.0
1.0
1.0
1.0
1.0
1.0
1.0
1.0
1.0
1.0
1.0
1.0
1.0
1.0
1.0
1.0
1.0
1.0
1.0
1.0


22
0
0
0
0
4
0
0
0
0
0
0
0
0
0
0
0
0
0
0
0
0
0
0
0
5
4
4
3
2


22
0
0
0
0
4
0
0
0
0
0
0
0
0
0
0
0
0
0
0
0
0
0
0
0
5
4
4
3
2

1.0
1.0
1.0
1.0
1.0
1.0
1.0
1.0
1.0
1.0
1.0
1.0
1.0
1.0
1.0
1.0
1.0
1.0
1.0
1.0
1.0
1.0
1.0
1.0
1.0
1.0
1.0
1.0
1.0
1.0


5
0
1
1
0
0
0
0
0
0
1
0
0
0
0
0
0
0
0
0
0
0
0
0
0
2
0
0
0
0

1.0
1.0
1.0
1.0
1.0
1.0
1.0
1.0
1.0
1.0
1.0
1.0
1.0
1.0
1.0
1.0
1.0
1.0
1.0
1.0
1.0
1.0
1.0
1.0
1.0
1.0
1.0
1.0
1.0
1.0


4
0
2
0
0
0
0
1
0
0
0
1
0
0
0
0
0
0
0
0
0
0
0
0
0
0
0
0
0
0


4
0
2
0
0
0
0
1
0
0
0
1
0
0
0
0
0
0
0
0
0
0
0
0
0
0
0
0
0
0

1.0
1.0
1.0
1.0
1.0
1.0
1.0
1.0
1.0
1.0
1.0
1.0
1.0
1.0
1.0
1.0
1.0
1.0
1.0
1.0
1.0
1.0
1.0
1.0
1.0
1.0
1.0
1.0
1.0
1.0


20
0
1
0
0
0
0
0
0
0
0
2
0
1
3
6
1
2
0
0
0
0
0
0
0
1
2
1
0
0


5
0
0
0
0
0
0
0
0
0
0
2
0
0
0
3
0
0
0
0
0
0
0
0
0
0
0
0
0
0

1.0
1.0
1.0
1.0
1.0
1.0
1.0
1.0
1.0
1.0
1.0
1.0
1.0
1.0
1.0
1.0
1.0
1.0
1.0
1.0
1.0
1.0
1.0
1.0
1.0
1.0
1.0
1.0
1.0
1.0


3
0
1
0
0
0
0
0
0
0
0
0
0
0
2
0
0
0
0
0
0
0
0
0
0
0
0
0
0
0

1.0
1.0
1.0
1.0
1.0
1.0
1.0
1.0
1.0
1.0
1.0
1.0
1.0
1.0
1.0
1.0
1.0
1.0
1.0
1.0
1.0
1.0
1.0
1.0
1.0
1.0
1.0
1.0
1.0
1.0


6
0
0
0
0
0
0
0
0
0
0
0
0
0
0
2
0
0
0
0
0
0
0
0
0
1
2
1
0
0

1.0
1.0
1.0
1.0
1.0
1.0
1.0
1.0
1.0
1.0
1.0
1.0
1.0
1.0
1.0
1.0
1.0
1.0
1.0
1.0
1.0
1.0
1.0
1.0
1.0
1.0
1.0
1.0
1.0
1.0


6
0
0
0
0
0
0
0
0
0
0
0
0
1
1
1
1
2
0
0
0
0
0
0
0
0
0
0
0
0

1.0
1.0
1.0
1.0
1.0
1.0
1.0
1.0
1.0
1.0
1.0
1.0
1.0
1.0
1.0
1.0
1.0
1.0
1.0
1.0
1.0
1.0
1.0
1.0
1.0
1.0
1.0
1.0
1.0
1.0


1
1
0
0
0
0
0
0
0
0
0
0
0
0
0
0
0
0
0
0
0
0
0
0
0
0
0
0
0
0


1
1
0
0
0
0
0
0
0
0
0
0
0
0
0
0
0
0
0
0
0
0
0
0
0
0
0
0
0
0

1.0
1.0
1.0
1.0
1.0
1.0
1.0
1.0
1.0
1.0
1.0
1.0
1.0
1.0
1.0
1.0
1.0
1.0
1.0
1.0
1.0
1.0
1.0
1.0
1.0
1.0
1.0
1.0
1.0
1.0


5
0
0
0
0
0
0
0
0
0
0
0
0
0
0
4
1
0
0
0
0
0
0
0
0
0
0
0
0
0


5
0
0
0
0
0
0
0
0
0
0
0
0
0
0
4
1
0
0
0
0
0
0
0
0
0
0
0
0
0

1.0
1.0
1.0
1.0
1.0
1.0
1.0
1.0
1.0
1.0
1.0
1.0
1.0
1.0
1.0
1.0
1.0
1.0
1.0
1.0
1.0
1.0
1.0
1.0
1.0
1.0
1.0
1.0
1.0
1.0


3
0
0
0
0
0
0
0
0
0
0
0
0
0
0
0
0
0
1
0
1
1
0
0
0
0
0
0
0
0


1
0
0
0
0
0
0
0
0
0
0
0
0
0
0
0
0
0
1
0
0
0
0
0
0
0
0
0
0
0

1.0
1.0
1.0
1.0
1.0
1.0
1.0
1.0
1.0
1.0
1.0
1.0
1.0
1.0
1.0
1.0
1.0
1.0
1.0
1.0
1.0
1.0
1.0
1.0
1.0
1.0
1.0
1.0
1.0
1.0


2
0
0
0
0
0
0
0
0
0
0
0
0
0
0
0
0
0
0
0
1
1
0
0
0
0
0
0
0
0

1.0
1.0
1.0
1.0
1.0
1.0
1.0
1.0
1.0
1.0
1.0
1.0
1.0
1.0
1.0
1.0
1.0
1.0
1.0
1.0
1.0
1.0
1.0
1.0
1.0
1.0
1.0
1.0
1.0
1.0


11
0
0
0
0
0
0
0
0
0
0
0
0
0
0
0
0
0
0
0
0
0
0
0
0
0
0
0
11
0


11
0
0
0
0
0
0
0
0
0
0
0
0
0
0
0
0
0
0
0
0
0
0
0
0
0
0
0
11
0

1.0
1.0
1.0
1.0
1.0
1.0
1.0
1.0
1.0
1.0
1.0
1.0
1.0
1.0
1.0
1.0
1.0
1.0
1.0
1.0
1.0
1.0
1.0
1.0
1.0
1.0
1.0
1.0
1.0
1.0


1
0
0
0
0
0
0
0
0
0
0
0
0
0
0
0
0
0
0
0
0
0
0
0
0
0
0
0
0
1


1
0
0
0
0
0
0
0
0
0
0
0
0
0
0
0
0
0
0
0
0
0
0
0
0
0
0
0
0
1

1.0
1.0
1.0
1.0
1.0
1.0
1.0
1.0
1.0
1.0
1.0
1.0
1.0
1.0
1.0
1.0
1.0
1.0
1.0
1.0
1.0
1.0
1.0
1.0
1.0
1.0
1.0
1.0
1.0
1.0


66
3
3
2
0
0
2
4
4
4
6
5
0
0
0
3
0
0
3
3
5
0
0
0
3
0
3
6
7
0


66
3
3
2
0
0
2
4
4
4
6
5
0
0
0
3
0
0
3
3
5
0
0
0
3
0
3
6
7
0


66
3
3
2
0
0
2
4
4
4
6
5
0
0
0
3
0
0
3
3
5
0
0
0
3
0
3
6
7
0

1.0
1.0
1.0
1.0
1.0
1.0
1.0
1.0
1.0
1.0
1.0
1.0
1.0
1.0
1.0
1.0
1.0
1.0
1.0
1.0
1.0
1.0
1.0
1.0
1.0
1.0
1.0
1.0
1.0
1.0


4
1
0
1
0
0
0
0
0
0
0
0
0
0
0
0
0
0
0
0
0
0
0
0
0
0
0
2
0
0


2
1
0
1
0
0
0
0
0
0
0
0
0
0
0
0
0
0
0
0
0
0
0
0
0
0
0
0
0
0


2
1
0
1
0
0
0
0
0
0
0
0
0
0
0
0
0
0
0
0
0
0
0
0
0
0
0
0
0
0

1.0
1.0
1.0
1.0
1.0
1.0
1.0
1.0
1.0
1.0
1.0
1.0
1.0
1.0
1.0
1.0
1.0
1.0
1.0
1.0
1.0
1.0
1.0
1.0
1.0
1.0
1.0
1.0
1.0
1.0


2
0
0
0
0
0
0
0
0
0
0
0
0
0
0
0
0
0
0
0
0
0
0
0
0
0
0
2
0
0

1.0
1.0
1.0
1.0
1.0
1.0
1.0
1.0
1.0
1.0
1.0
1.0
1.0
1.0
1.0
1.0
1.0
1.0
1.0
1.0
1.0
1.0
1.0
1.0
1.0
1.0
1.0
1.0
1.0
1.0


1
0
0
0
0
0
0
0
0
0
0
0
0
0
0
0
0
0
0
0
0
0
0
0
0
0
0
1
0
0

1.0
1.0
1.0
1.0
1.0
1.0
1.0
1.0
1.0
1.0
1.0
1.0
1.0
1.0
1.0
1.0
1.0
1.0
1.0
1.0
1.0
1.0
1.0
1.0
1.0
1.0
1.0
1.0
1.0
1.0


15
1
4
5
0
0
0
0
0
0
1
1
0
2
1
0
0
0
0
0
0
0
0
0
0
0
0
0
0
0


3
1
0
0
0
0
0
0
0
0
1
0
0
1
0
0
0
0
0
0
0
0
0
0
0
0
0
0
0
0


2
1
0
0
0
0
0
0
0
0
1
0
0
0
0
0
0
0
0
0
0
0
0
0
0
0
0
0
0
0

1.0
1.0
1.0
1.0
1.0
1.0
1.0
1.0
1.0
1.0
1.0
1.0
1.0
1.0
1.0
1.0
1.0
1.0
1.0
1.0
1.0
1.0
1.0
1.0
1.0
1.0
1.0
1.0
1.0
1.0


1
0
0
0
0
0
0
0
0
0
0
0
0
1
0
0
0
0
0
0
0
0
0
0
0
0
0
0
0
0

1.0
1.0
1.0
1.0
1.0
1.0
1.0
1.0
1.0
1.0
1.0
1.0
1.0
1.0
1.0
1.0
1.0
1.0
1.0
1.0
1.0
1.0
1.0
1.0
1.0
1.0
1.0
1.0
1.0
1.0


12
0
4
5
0
0
0
0
0
0
0
1
0
1
1
0
0
0
0
0
0
0
0
0
0
0
0
0
0
0


12
0
4
5
0
0
0
0
0
0
0
1
0
1
1
0
0
0
0
0
0
0
0
0
0
0
0
0
0
0

1.0
1.0
1.0
1.0
1.0
1.0
1.0
1.0
1.0
1.0
1.0
1.0
1.0
1.0
1.0
1.0
1.0
1.0
1.0
1.0
1.0
1.0
1.0
1.0
1.0
1.0
1.0
1.0
1.0
1.0


8
1
0
0
0
0
0
0
1
1
3
1
0
1
0
0
0
0
0
0
0
0
0
0
0
0
0
0
0
0


6
1
0
0
0
0
0
0
0
1
2
1
0
1
0
0
0
0
0
0
0
0
0
0
0
0
0
0
0
0


6
1
0
0
0
0
0
0
0
1
2
1
0
1
0
0
0
0
0
0
0
0
0
0
0
0
0
0
0
0

1.0
1.0
1.0
1.0
1.0
1.0
1.0
1.0
1.0
1.0
1.0
1.0
1.0
1.0
1.0
1.0
1.0
1.0
1.0
1.0
1.0
1.0
1.0
1.0
1.0
1.0
1.0
1.0
1.0
1.0


2
0
0
0
0
0
0
0
1
0
1
0
0
0
0
0
0
0
0
0
0
0
0
0
0
0
0
0
0
0


2
0
0
0
0
0
0
0
1
0
1
0
0
0
0
0
0
0
0
0
0
0
0
0
0
0
0
0
0
0

1.0
1.0
1.0
1.0
1.0
1.0
1.0
1.0
1.0
1.0
1.0
1.0
1.0
1.0
1.0
1.0
1.0
1.0
1.0
1.0
1.0
1.0
1.0
1.0
1.0
1.0
1.0
1.0
1.0
1.0


15
0
4
4
0
0
1
1
0
0
0
0
0
0
0
0
0
0
0
0
0
0
0
0
1
0
2
0
0
2


11
0
3
3
0
0
1
1
0
0
0
0
0
0
0
0
0
0
0
0
0
0
0
0
1
0
1
0
0
1


3
0
1
1
0
0
1
0
0
0
0
0
0
0
0
0
0
0
0
0
0
0
0
0
0
0
0
0
0
0

1.0
1.0
1.0
1.0
1.0
1.0
1.0
1.0
1.0
1.0
1.0
1.0
1.0
1.0
1.0
1.0
1.0
1.0
1.0
1.0
1.0
1.0
1.0
1.0
1.0
1.0
1.0
1.0
1.0
1.0


3
0
0
1
0
0
0
1
0
0
0
0
0
0
0
0
0
0
0
0
0
0
0
0
0
0
0
0
0
1

1.0
1.0
1.0
1.0
1.0
1.0
1.0
1.0
1.0
1.0
1.0
1.0
1.0
1.0
1.0
1.0
1.0
1.0
1.0
1.0
1.0
1.0
1.0
1.0
1.0
1.0
1.0
1.0
1.0
1.0


1
0
1
0
0
0
0
0
0
0
0
0
0
0
0
0
0
0
0
0
0
0
0
0
0
0
0
0
0
0

1.0
1.0
1.0
1.0
1.0
1.0
1.0
1.0
1.0
1.0
1.0
1.0
1.0
1.0
1.0
1.0
1.0
1.0
1.0
1.0
1.0
1.0
1.0
1.0
1.0
1.0
1.0
1.0
1.0
1.0


4
0
1
1
0
0
0
0
0
0
0
0
0
0
0
0
0
0
0
0
0
0
0
0
1
0
1
0
0
0

0.99
0.99
0.99
0.99
0.99
0.99
0.99
0.99
0.99
0.99
0.99
0.99
0.99
0.99
0.99
0.99
0.99
0.99
0.99
0.99
0.99
0.99
0.99
0.99
0.99
0.99
0.99
0.99
0.99
0.99


2
0
1
1
0
0
0
0
0
0
0
0
0
0
0
0
0
0
0
0
0
0
0
0
0
0
0
0
0
0


2
0
1
1
0
0
0
0
0
0
0
0
0
0
0
0
0
0
0
0
0
0
0
0
0
0
0
0
0
0

1.0
1.0
1.0
1.0
1.0
1.0
1.0
1.0
1.0
1.0
1.0
1.0
1.0
1.0
1.0
1.0
1.0
1.0
1.0
1.0
1.0
1.0
1.0
1.0
1.0
1.0
1.0
1.0
1.0
1.0


2
0
0
0
0
0
0
0
0
0
0
0
0
0
0
0
0
0
0
0
0
0
0
0
0
0
1
0
0
1


2
0
0
0
0
0
0
0
0
0
0
0
0
0
0
0
0
0
0
0
0
0
0
0
0
0
1
0
0
1

1.0
1.0
1.0
1.0
1.0
1.0
1.0
1.0
1.0
1.0
1.0
1.0
1.0
1.0
1.0
1.0
1.0
1.0
1.0
1.0
1.0
1.0
1.0
1.0
1.0
1.0
1.0
1.0
1.0
1.0


67
0
1
0
0
0
0
0
0
0
0
0
0
0
0
1
0
32
17
1
0
0
0
0
1
5
0
0
0
9

1.0
1.0
1.0
1.0
1.0
1.0
1.0
1.0
1.0
1.0
1.0
1.0
1.0
1.0
1.0
1.0
1.0
1.0
1.0
1.0
1.0
1.0
1.0
1.0
1.0
1.0
1.0
1.0
1.0
1.0


47
0
1
0
0
0
0
0
0
0
0
0
0
0
0
0
0
32
0
0
0
0
0
0
0
5
0
0
0
9

0.99
0.99
0.99
0.99
0.99
0.99
0.99
0.99
0.99
0.99
0.99
0.99
0.99
0.99
0.99
0.99
0.99
0.99
0.99
0.99
0.99
0.99
0.99
0.99
0.99
0.99
0.99
0.99
0.99
0.99


45
0
1
0
0
0
0
0
0
0
0
0
0
0
0
0
0
32
0
0
0
0
0
0
0
4
0
0
0
8

1.0
1.0
1.0
1.0
1.0
1.0
1.0
1.0
1.0
1.0
1.0
1.0
1.0
1.0
1.0
1.0
1.0
1.0
1.0
1.0
1.0
1.0
1.0
1.0
1.0
1.0
1.0
1.0
1.0
1.0


7
0
0
0
0
0
0
0
0
0
0
0
0
0
0
0
0
0
6
0
0
0
0
0
1
0
0
0
0
0

0.99
0.99
0.99
0.99
0.99
0.99
0.99
0.99
0.99
0.99
0.99
0.99
0.99
0.99
0.99
0.99
0.99
0.99
0.99
0.99
0.99
0.99
0.99
0.99
0.99
0.99
0.99
0.99
0.99
0.99


5
0
0
0
0
0
0
0
0
0
0
0
0
0
0
0
0
0
4
0
0
0
0
0
1
0
0
0
0
0

0.99
0.99
0.99
0.99
0.99
0.99
0.99
0.99
0.99
0.99
0.99
0.99
0.99
0.99
0.99
0.99
0.99
0.99
0.99
0.99
0.99
0.99
0.99
0.99
0.99
0.99
0.99
0.99
0.99
0.99


8
0
0
0
0
0
0
0
0
0
0
0
0
0
0
0
0
0
8
0
0
0
0
0
0
0
0
0
0
0


8
0
0
0
0
0
0
0
0
0
0
0
0
0
0
0
0
0
8
0
0
0
0
0
0
0
0
0
0
0

0.99
0.99
0.99
0.99
0.99
0.99
0.99
0.99
0.99
0.99
0.99
0.99
0.99
0.99
0.99
0.99
0.99
0.99
0.99
0.99
0.99
0.99
0.99
0.99
0.99
0.99
0.99
0.99
0.99
0.99


2
0
0
0
0
0
0
0
0
0
0
0
0
0
0
0
0
0
1
1
0
0
0
0
0
0
0
0
0
0


2
0
0
0
0
0
0
0
0
0
0
0
0
0
0
0
0
0
1
1
0
0
0
0
0
0
0
0
0
0

1.0
1.0
1.0
1.0
1.0
1.0
1.0
1.0
1.0
1.0
1.0
1.0
1.0
1.0
1.0
1.0
1.0
1.0
1.0
1.0
1.0
1.0
1.0
1.0
1.0
1.0
1.0
1.0
1.0
1.0


1
0
0
0
0
0
0
0
0
0
0
0
0
0
0
1
0
0
0
0
0
0
0
0
0
0
0
0
0
0


1
0
0
0
0
0
0
0
0
0
0
0
0
0
0
1
0
0
0
0
0
0
0
0
0
0
0
0
0
0

0.99
0.99
0.99
0.99
0.99
0.99
0.99
0.99
0.99
0.99
0.99
0.99
0.99
0.99
0.99
0.99
0.99
0.99
0.99
0.99
0.99
0.99
0.99
0.99
0.99
0.99
0.99
0.99
0.99
0.99


3
0
0
0
1
1
1
0
0
0
0
0
0
0
0
0
0
0
0
0
0
0
0
0
0
0
0
0
0
0


3
0
0
0
1
1
1
0
0
0
0
0
0
0
0
0
0
0
0
0
0
0
0
0
0
0
0
0
0
0


3
0
0
0
1
1
1
0
0
0
0
0
0
0
0
0
0
0
0
0
0
0
0
0
0
0
0
0
0
0

1.0
1.0
1.0
1.0
1.0
1.0
1.0
1.0
1.0
1.0
1.0
1.0
1.0
1.0
1.0
1.0
1.0
1.0
1.0
1.0
1.0
1.0
1.0
1.0
1.0
1.0
1.0
1.0
1.0
1.0


8
0
0
0
0
0
0
1
1
1
1
1
0
0
0
0
0
0
1
1
0
0
0
0
1
0
0
0
0
0


8
0
0
0
0
0
0
1
1
1
1
1
0
0
0
0
0
0
1
1
0
0
0
0
1
0
0
0
0
0

0.99
0.99
0.99
0.99
0.99
0.99
0.99
0.99
0.99
0.99
0.99
0.99
0.99
0.99
0.99
0.99
0.99
0.99
0.99
0.99
0.99
0.99
0.99
0.99
0.99
0.99
0.99
0.99
0.99
0.99


2
0
0
0
0
0
0
0
0
0
0
1
0
0
0
0
0
0
0
0
0
0
0
0
1
0
0
0
0
0

1.0
1.0
1.0
1.0
1.0
1.0
1.0
1.0
1.0
1.0
1.0
1.0
1.0
1.0
1.0
1.0
1.0
1.0
1.0
1.0
1.0
1.0
1.0
1.0
1.0
1.0
1.0
1.0
1.0
1.0


51
1
2
1
3
1
2
0
1
0
0
1
0
1
3
8
7
0
0
0
0
0
0
0
3
5
1
3
3
5

1.0
1.0
1.0
1.0
1.0
1.0
1.0
1.0
1.0
1.0
1.0
1.0
1.0
1.0
1.0
1.0
1.0
1.0
1.0
1.0
1.0
1.0
1.0
1.0
1.0
1.0
1.0
1.0
1.0
1.0


17
0
0
0
1
0
2
0
1
0
0
0
0
0
1
3
2
0
0
0
0
0
0
0
0
3
1
3
0
0

0.99
0.99
0.99
0.99
0.99
0.99
0.99
0.99
0.99
0.99
0.99
0.99
0.99
0.99
0.99
0.99
0.99
0.99
0.99
0.99
0.99
0.99
0.99
0.99
0.99
0.99
0.99
0.99
0.99
0.99


2
0
0
0
1
0
0
0
0
0
0
0
0
0
1
0
0
0
0
0
0
0
0
0
0
0
0
0
0
0

1.0
1.0
1.0
1.0
1.0
1.0
1.0
1.0
1.0
1.0
1.0
1.0
1.0
1.0
1.0
1.0
1.0
1.0
1.0
1.0
1.0
1.0
1.0
1.0
1.0
1.0
1.0
1.0
1.0
1.0


4
0
0
0
0
0
2
0
1
0
0
0
0
0
0
0
0
0
0
0
0
0
0
0
0
0
1
0
0
0

0.86
0.86
0.86
0.86
0.86
0.86
0.86
0.86
0.86
0.86
0.86
0.86
0.86
0.86
0.86
0.86
0.86
0.86
0.86
0.86
0.86
0.86
0.86
0.86
0.86
0.86
0.86
0.86
0.86
0.86


1
0
0
0
0
0
0
0
0
0
0
0
0
0
0
0
0
0
0
0
0
0
0
0
0
0
0
1
0
0

1.0
1.0
1.0
1.0
1.0
1.0
1.0
1.0
1.0
1.0
1.0
1.0
1.0
1.0
1.0
1.0
1.0
1.0
1.0
1.0
1.0
1.0
1.0
1.0
1.0
1.0
1.0
1.0
1.0
1.0


20
0
1
1
2
0
0
0
0
0
0
0
0
0
0
4
4
0
0
0
0
0
0
0
2
0
0
0
2
4

0.86
0.86
0.86
0.86
0.86
0.86
0.86
0.86
0.86
0.86
0.86
0.86
0.86
0.86
0.86
0.86
0.86
0.86
0.86
0.86
0.86
0.86
0.86
0.86
0.86
0.86
0.86
0.86
0.86
0.86


4
0
0
0
0
1
0
0
0
0
0
0
0
0
1
0
0
0
0
0
0
0
0
0
0
2
0
0
0
0


4
0
0
0
0
1
0
0
0
0
0
0
0
0
1
0
0
0
0
0
0
0
0
0
0
2
0
0
0
0

1.0
1.0
1.0
1.0
1.0
1.0
1.0
1.0
1.0
1.0
1.0
1.0
1.0
1.0
1.0
1.0
1.0
1.0
1.0
1.0
1.0
1.0
1.0
1.0
1.0
1.0
1.0
1.0
1.0
1.0


23
0
1
0
1
2
4
1
0
0
2
1
0
0
0
0
0
0
0
1
0
0
0
0
0
2
2
2
1
3


7
0
1
0
0
0
1
1
0
0
2
0
0
0
0
0
0
0
0
0
0
0
0
0
0
1
1
0
0
0


7
0
1
0
0
0
1
1
0
0
2
0
0
0
0
0
0
0
0
0
0
0
0
0
0
1
1
0
0
0

1.0
1.0
1.0
1.0
1.0
1.0
1.0
1.0
1.0
1.0
1.0
1.0
1.0
1.0
1.0
1.0
1.0
1.0
1.0
1.0
1.0
1.0
1.0
1.0
1.0
1.0
1.0
1.0
1.0
1.0


6
0
0
0
1
1
2
0
0
0
0
1
0
0
0
0
0
0
0
0
0
0
0
0
0
0
1
0
0
0


6
0
0
0
1
1
2
0
0
0
0
1
0
0
0
0
0
0
0
0
0
0
0
0
0
0
1
0
0
0

0.98
0.98
0.98
0.98
0.98
0.98
0.98
0.98
0.98
0.98
0.98
0.98
0.98
0.98
0.98
0.98
0.98
0.98
0.98
0.98
0.98
0.98
0.98
0.98
0.98
0.98
0.98
0.98
0.98
0.98


7
0
0
0
0
1
1
0
0
0
0
0
0
0
0
0
0
0
0
1
0
0
0
0
0
1
0
2
0
1


3
0
0
0
0
1
1
0
0
0
0
0
0
0
0
0
0
0
0
0
0
0
0
0
0
0
0
1
0
0

1.0
1.0
1.0
1.0
1.0
1.0
1.0
1.0
1.0
1.0
1.0
1.0
1.0
1.0
1.0
1.0
1.0
1.0
1.0
1.0
1.0
1.0
1.0
1.0
1.0
1.0
1.0
1.0
1.0
1.0


3
0
0
0
0
0
0
0
0
0
0
0
0
0
0
0
0
0
0
0
0
0
0
0
0
1
0
1
0
1

1.0
1.0
1.0
1.0
1.0
1.0
1.0
1.0
1.0
1.0
1.0
1.0
1.0
1.0
1.0
1.0
1.0
1.0
1.0
1.0
1.0
1.0
1.0
1.0
1.0
1.0
1.0
1.0
1.0
1.0


1
0
0
0
0
0
0
0
0
0
0
0
0
0
0
0
0
0
0
1
0
0
0
0
0
0
0
0
0
0

1.0
1.0
1.0
1.0
1.0
1.0
1.0
1.0
1.0
1.0
1.0
1.0
1.0
1.0
1.0
1.0
1.0
1.0
1.0
1.0
1.0
1.0
1.0
1.0
1.0
1.0
1.0
1.0
1.0
1.0


3
0
0
0
0
0
0
0
0
0
0
0
0
0
0
0
0
0
0
0
0
0
0
0
0
0
0
0
1
2


3
0
0
0
0
0
0
0
0
0
0
0
0
0
0
0
0
0
0
0
0
0
0
0
0
0
0
0
1
2

1.0
1.0
1.0
1.0
1.0
1.0
1.0
1.0
1.0
1.0
1.0
1.0
1.0
1.0
1.0
1.0
1.0
1.0
1.0
1.0
1.0
1.0
1.0
1.0
1.0
1.0
1.0
1.0
1.0
1.0


5
0
2
1
0
0
0
0
0
0
0
0
0
1
1
0
0
0
0
0
0
0
0
0
0
0
0
0
0
0


3
0
1
1
0
0
0
0
0
0
0
0
0
0
1
0
0
0
0
0
0
0
0
0
0
0
0
0
0
0


3
0
1
1
0
0
0
0
0
0
0
0
0
0
1
0
0
0
0
0
0
0
0
0
0
0
0
0
0
0

1.0
1.0
1.0
1.0
1.0
1.0
1.0
1.0
1.0
1.0
1.0
1.0
1.0
1.0
1.0
1.0
1.0
1.0
1.0
1.0
1.0
1.0
1.0
1.0
1.0
1.0
1.0
1.0
1.0
1.0


2
0
1
0
0
0
0
0
0
0
0
0
0
1
0
0
0
0
0
0
0
0
0
0
0
0
0
0
0
0


2
0
1
0
0
0
0
0
0
0
0
0
0
1
0
0
0
0
0
0
0
0
0
0
0
0
0
0
0
0

1.0
1.0
1.0
1.0
1.0
1.0
1.0
1.0
1.0
1.0
1.0
1.0
1.0
1.0
1.0
1.0
1.0
1.0
1.0
1.0
1.0
1.0
1.0
1.0
1.0
1.0
1.0
1.0
1.0
1.0


5
2
0
0
0
0
0
0
0
1
0
2
0
0
0
0
0
0
0
0
0
0
0
0
0
0
0
0
0
0


5
2
0
0
0
0
0
0
0
1
0
2
0
0
0
0
0
0
0
0
0
0
0
0
0
0
0
0
0
0


5
2
0
0
0
0
0
0
0
1
0
2
0
0
0
0
0
0
0
0
0
0
0
0
0
0
0
0
0
0

1.0
1.0
1.0
1.0
1.0
1.0
1.0
1.0
1.0
1.0
1.0
1.0
1.0
1.0
1.0
1.0
1.0
1.0
1.0
1.0
1.0
1.0
1.0
1.0
1.0
1.0
1.0
1.0
1.0
1.0


8
0
0
0
0
0
0
1
0
0
0
0
0
0
0
0
0
2
0
0
1
0
0
0
0
0
0
0
4
0


8
0
0
0
0
0
0
1
0
0
0
0
0
0
0
0
0
2
0
0
1
0
0
0
0
0
0
0
4
0


8
0
0
0
0
0
0
1
0
0
0
0
0
0
0
0
0
2
0
0
1
0
0
0
0
0
0
0
4
0

1.0
1.0
1.0
1.0
1.0
1.0
1.0
1.0
1.0
1.0
1.0
1.0
1.0
1.0
1.0
1.0
1.0
1.0
1.0
1.0
1.0
1.0
1.0
1.0
1.0
1.0
1.0
1.0
1.0
1.0


7
1
1
0
0
0
0
0
1
0
0
0
0
0
0
1
0
0
1
0
0
0
0
0
1
0
0
0
0
1


7
1
1
0
0
0
0
0
1
0
0
0
0
0
0
1
0
0
1
0
0
0
0
0
1
0
0
0
0
1


7
1
1
0
0
0
0
0
1
0
0
0
0
0
0
1
0
0
1
0
0
0
0
0
1
0
0
0
0
1

1.0
1.0
1.0
1.0
1.0
1.0
1.0
1.0
1.0
1.0
1.0
1.0
1.0
1.0
1.0
1.0
1.0
1.0
1.0
1.0
1.0
1.0
1.0
1.0
1.0
1.0
1.0
1.0
1.0
1.0


1
0
0
0
0
0
0
0
0
0
1
0
0
0
0
0
0
0
0
0
0
0
0
0
0
0
0
0
0
0


1
0
0
0
0
0
0
0
0
0
1
0
0
0
0
0
0
0
0
0
0
0
0
0
0
0
0
0
0
0


1
0
0
0
0
0
0
0
0
0
1
0
0
0
0
0
0
0
0
0
0
0
0
0
0
0
0
0
0
0

1.0
1.0
1.0
1.0
1.0
1.0
1.0
1.0
1.0
1.0
1.0
1.0
1.0
1.0
1.0
1.0
1.0
1.0
1.0
1.0
1.0
1.0
1.0
1.0
1.0
1.0
1.0
1.0
1.0
1.0


4
0
0
0
1
0
0
0
0
0
1
0
0
0
0
1
0
0
0
0
0
0
0
0
0
0
1
0
0
0


4
0
0
0
1
0
0
0
0
0
1
0
0
0
0
1
0
0
0
0
0
0
0
0
0
0
1
0
0
0


4
0
0
0
1
0
0
0
0
0
1
0
0
0
0
1
0
0
0
0
0
0
0
0
0
0
1
0
0
0

1.0
1.0
1.0
1.0
1.0
1.0
1.0
1.0
1.0
1.0
1.0
1.0
1.0
1.0
1.0
1.0
1.0
1.0
1.0
1.0
1.0
1.0
1.0
1.0
1.0
1.0
1.0
1.0
1.0
1.0


7
2
1
1
0
0
1
0
0
0
0
1
0
0
0
0
0
0
0
0
0
0
0
0
0
0
0
0
1
0


3
0
1
1
0
0
0
0
0
0
0
1
0
0
0
0
0
0
0
0
0
0
0
0
0
0
0
0
0
0


3
0
1
1
0
0
0
0
0
0
0
1
0
0
0
0
0
0
0
0
0
0
0
0
0
0
0
0
0
0

1.0
1.0
1.0
1.0
1.0
1.0
1.0
1.0
1.0
1.0
1.0
1.0
1.0
1.0
1.0
1.0
1.0
1.0
1.0
1.0
1.0
1.0
1.0
1.0
1.0
1.0
1.0
1.0
1.0
1.0


1
1
0
0
0
0
0
0
0
0
0
0
0
0
0
0
0
0
0
0
0
0
0
0
0
0
0
0
0
0


1
1
0
0
0
0
0
0
0
0
0
0
0
0
0
0
0
0
0
0
0
0
0
0
0
0
0
0
0
0

1.0
1.0
1.0
1.0
1.0
1.0
1.0
1.0
1.0
1.0
1.0
1.0
1.0
1.0
1.0
1.0
1.0
1.0
1.0
1.0
1.0
1.0
1.0
1.0
1.0
1.0
1.0
1.0
1.0
1.0


1
1
0
0
0
0
0
0
0
0
0
0
0
0
0
0
0
0
0
0
0
0
0
0
0
0
0
0
0
0


1
1
0
0
0
0
0
0
0
0
0
0
0
0
0
0
0
0
0
0
0
0
0
0
0
0
0
0
0
0

1.0
1.0
1.0
1.0
1.0
1.0
1.0
1.0
1.0
1.0
1.0
1.0
1.0
1.0
1.0
1.0
1.0
1.0
1.0
1.0
1.0
1.0
1.0
1.0
1.0
1.0
1.0
1.0
1.0
1.0


2
0
0
0
0
0
1
0
0
0
0
0
0
0
0
0
0
0
0
0
0
0
0
0
0
0
0
0
1
0


2
0
0
0
0
0
1
0
0
0
0
0
0
0
0
0
0
0
0
0
0
0
0
0
0
0
0
0
1
0

1.0
1.0
1.0
1.0
1.0
1.0
1.0
1.0
1.0
1.0
1.0
1.0
1.0
1.0
1.0
1.0
1.0
1.0
1.0
1.0
1.0
1.0
1.0
1.0
1.0
1.0
1.0
1.0
1.0
1.0


34
0
0
1
0
1
1
0
0
0
0
0
0
0
0
0
1
0
0
0
0
0
0
0
2
4
7
6
6
5


13
0
0
1
0
1
1
0
0
0
0
0
0
0
0
0
0
0
0
0
0
0
0
0
0
1
4
2
1
2


13
0
0
1
0
1
1
0
0
0
0
0
0
0
0
0
0
0
0
0
0
0
0
0
0
1
4
2
1
2

1.0
1.0
1.0
1.0
1.0
1.0
1.0
1.0
1.0
1.0
1.0
1.0
1.0
1.0
1.0
1.0
1.0
1.0
1.0
1.0
1.0
1.0
1.0
1.0
1.0
1.0
1.0
1.0
1.0
1.0


21
0
0
0
0
0
0
0
0
0
0
0
0
0
0
0
1
0
0
0
0
0
0
0
2
3
3
4
5
3

0.79
0.79
0.79
0.79
0.79
0.79
0.79
0.79
0.79
0.79
0.79
0.79
0.79
0.79
0.79
0.79
0.79
0.79
0.79
0.79
0.79
0.79
0.79
0.79
0.79
0.79
0.79
0.79
0.79
0.79


6
0
0
0
0
0
0
0
0
0
0
0
0
0
0
0
0
0
0
0
0
0
0
0
0
1
1
2
1
1

1.0
1.0
1.0
1.0
1.0
1.0
1.0
1.0
1.0
1.0
1.0
1.0
1.0
1.0
1.0
1.0
1.0
1.0
1.0
1.0
1.0
1.0
1.0
1.0
1.0
1.0
1.0
1.0
1.0
1.0


3
0
0
0
0
0
0
0
0
0
0
0
0
0
0
0
0
0
0
0
0
0
0
0
1
1
0
0
1
0

1.0
1.0
1.0
1.0
1.0
1.0
1.0
1.0
1.0
1.0
1.0
1.0
1.0
1.0
1.0
1.0
1.0
1.0
1.0
1.0
1.0
1.0
1.0
1.0
1.0
1.0
1.0
1.0
1.0
1.0


4
0
0
0
0
0
0
0
0
0
0
0
0
0
0
0
0
0
0
0
0
0
0
0
0
0
2
1
1
0

1.0
1.0
1.0
1.0
1.0
1.0
1.0
1.0
1.0
1.0
1.0
1.0
1.0
1.0
1.0
1.0
1.0
1.0
1.0
1.0
1.0
1.0
1.0
1.0
1.0
1.0
1.0
1.0
1.0
1.0


1
0
0
0
0
0
0
0
0
0
0
0
0
0
0
0
0
0
0
0
0
0
0
0
0
0
0
0
0
1

1.0
1.0
1.0
1.0
1.0
1.0
1.0
1.0
1.0
1.0
1.0
1.0
1.0
1.0
1.0
1.0
1.0
1.0
1.0
1.0
1.0
1.0
1.0
1.0
1.0
1.0
1.0
1.0
1.0
1.0


1
0
0
0
0
0
0
0
0
0
0
0
0
0
0
0
1
0
0
0
0
0
0
0
0
0
0
0
0
0

0.79
0.79
0.79
0.79
0.79
0.79
0.79
0.79
0.79
0.79
0.79
0.79
0.79
0.79
0.79
0.79
0.79
0.79
0.79
0.79
0.79
0.79
0.79
0.79
0.79
0.79
0.79
0.79
0.79
0.79


1
0
0
0
0
0
0
0
0
0
0
0
0
0
0
0
0
0
0
0
0
0
0
0
0
0
0
0
1
0

1.0
1.0
1.0
1.0
1.0
1.0
1.0
1.0
1.0
1.0
1.0
1.0
1.0
1.0
1.0
1.0
1.0
1.0
1.0
1.0
1.0
1.0
1.0
1.0
1.0
1.0
1.0
1.0
1.0
1.0


1
1
0
0
0
0
0
0
0
0
0
0
0
0
0
0
0
0
0
0
0
0
0
0
0
0
0
0
0
0


1
1
0
0
0
0
0
0
0
0
0
0
0
0
0
0
0
0
0
0
0
0
0
0
0
0
0
0
0
0


1
1
0
0
0
0
0
0
0
0
0
0
0
0
0
0
0
0
0
0
0
0
0
0
0
0
0
0
0
0

1.0
1.0
1.0
1.0
1.0
1.0
1.0
1.0
1.0
1.0
1.0
1.0
1.0
1.0
1.0
1.0
1.0
1.0
1.0
1.0
1.0
1.0
1.0
1.0
1.0
1.0
1.0
1.0
1.0
1.0


2
0
0
0
0
0
0
0
0
0
1
0
0
0
0
0
0
0
0
0
0
0
0
0
0
0
0
0
1
0


2
0
0
0
0
0
0
0
0
0
1
0
0
0
0
0
0
0
0
0
0
0
0
0
0
0
0
0
1
0


1
0
0
0
0
0
0
0
0
0
1
0
0
0
0
0
0
0
0
0
0
0
0
0
0
0
0
0
0
0

1.0
1.0
1.0
1.0
1.0
1.0
1.0
1.0
1.0
1.0
1.0
1.0
1.0
1.0
1.0
1.0
1.0
1.0
1.0
1.0
1.0
1.0
1.0
1.0
1.0
1.0
1.0
1.0
1.0
1.0


1
0
0
0
0
0
0
0
0
0
0
0
0
0
0
0
0
0
0
0
0
0
0
0
0
0
0
0
1
0

1.0
1.0
1.0
1.0
1.0
1.0
1.0
1.0
1.0
1.0
1.0
1.0
1.0
1.0
1.0
1.0
1.0
1.0
1.0
1.0
1.0
1.0
1.0
1.0
1.0
1.0
1.0
1.0
1.0
1.0


69
0
0
1
0
0
0
0
0
0
0
0
0
0
1
3
1
1
1
1
0
0
2
0
0
1
23
13
9
12

1.0
1.0
1.0
1.0
1.0
1.0
1.0
1.0
1.0
1.0
1.0
1.0
1.0
1.0
1.0
1.0
1.0
1.0
1.0
1.0
1.0
1.0
1.0
1.0
1.0
1.0
1.0
1.0
1.0
1.0


2
0
0
1
0
0
0
0
0
0
0
0
0
0
0
0
1
0
0
0
0
0
0
0
0
0
0
0
0
0


2
0
0
1
0
0
0
0
0
0
0
0
0
0
0
0
1
0
0
0
0
0
0
0
0
0
0
0
0
0

1.0
1.0
1.0
1.0
1.0
1.0
1.0
1.0
1.0
1.0
1.0
1.0
1.0
1.0
1.0
1.0
1.0
1.0
1.0
1.0
1.0
1.0
1.0
1.0
1.0
1.0
1.0
1.0
1.0
1.0


39
0
0
0
0
0
0
0
0
0
0
0
0
0
1
1
0
1
0
1
0
0
1
0
0
1
7
12
7
7

0.73
0.73
0.73
0.73
0.73
0.73
0.73
0.73
0.73
0.73
0.73
0.73
0.73
0.73
0.73
0.73
0.73
0.73
0.73
0.73
0.73
0.73
0.73
0.73
0.73
0.73
0.73
0.73
0.73
0.73


37
0
0
0
0
0
0
0
0
0
0
0
0
0
1
1
0
1
0
1
0
0
1
0
0
1
7
12
6
6

1.0
1.0
1.0
1.0
1.0
1.0
1.0
1.0
1.0
1.0
1.0
1.0
1.0
1.0
1.0
1.0
1.0
1.0
1.0
1.0
1.0
1.0
1.0
1.0
1.0
1.0
1.0
1.0
1.0
1.0


7
0
0
0
0
0
0
0
0
0
0
0
0
0
0
0
0
0
0
0
0
0
1
0
0
0
6
0
0
0


7
0
0
0
0
0
0
0
0
0
0
0
0
0
0
0
0
0
0
0
0
0
1
0
0
0
6
0
0
0

1.0
1.0
1.0
1.0
1.0
1.0
1.0
1.0
1.0
1.0
1.0
1.0
1.0
1.0
1.0
1.0
1.0
1.0
1.0
1.0
1.0
1.0
1.0
1.0
1.0
1.0
1.0
1.0
1.0
1.0


2
0
0
0
0
0
0
0
0
0
0
0
0
0
0
0
0
0
0
0
0
0
0
0
0
0
0
0
0
2


2
0
0
0
0
0
0
0
0
0
0
0
0
0
0
0
0
0
0
0
0
0
0
0
0
0
0
0
0
2

0.99
0.99
0.99
0.99
0.99
0.99
0.99
0.99
0.99
0.99
0.99
0.99
0.99
0.99
0.99
0.99
0.99
0.99
0.99
0.99
0.99
0.99
0.99
0.99
0.99
0.99
0.99
0.99
0.99
0.99


3
0
0
0
0
0
0
0
0
0
0
0
0
0
0
0
0
0
0
0
0
0
0
0
0
0
3
0
0
0


3
0
0
0
0
0
0
0
0
0
0
0
0
0
0
0
0
0
0
0
0
0
0
0
0
0
3
0
0
0

1.0
1.0
1.0
1.0
1.0
1.0
1.0
1.0
1.0
1.0
1.0
1.0
1.0
1.0
1.0
1.0
1.0
1.0
1.0
1.0
1.0
1.0
1.0
1.0
1.0
1.0
1.0
1.0
1.0
1.0


2
0
0
0
0
0
0
0
0
0
0
0
0
0
0
2
0
0
0
0
0
0
0
0
0
0
0
0
0
0

1.0
1.0
1.0
1.0
1.0
1.0
1.0
1.0
1.0
1.0
1.0
1.0
1.0
1.0
1.0
1.0
1.0
1.0
1.0
1.0
1.0
1.0
1.0
1.0
1.0
1.0
1.0
1.0
1.0
1.0


1
0
0
0
0
0
0
0
0
0
0
0
0
0
0
1
0
0
0
0
0
0
0
0
0
0
0
0
0
0

0.71
0.71
0.71
0.71
0.71
0.71
0.71
0.71
0.71
0.71
0.71
0.71
0.71
0.71
0.71
0.71
0.71
0.71
0.71
0.71
0.71
0.71
0.71
0.71
0.71
0.71
0.71
0.71
0.71
0.71


5
0
0
0
0
0
0
0
0
0
0
0
0
0
0
0
0
0
0
0
0
0
0
0
0
0
3
0
0
2


3
0
0
0
0
0
0
0
0
0
0
0
0
0
0
0
0
0
0
0
0
0
0
0
0
0
1
0
0
2

0.98
0.98
0.98
0.98
0.98
0.98
0.98
0.98
0.98
0.98
0.98
0.98
0.98
0.98
0.98
0.98
0.98
0.98
0.98
0.98
0.98
0.98
0.98
0.98
0.98
0.98
0.98
0.98
0.98
0.98


2
0
0
0
0
0
0
0
0
0
0
0
0
0
0
0
0
0
0
0
0
0
0
0
0
0
2
0
0
0

0.95
0.95
0.95
0.95
0.95
0.95
0.95
0.95
0.95
0.95
0.95
0.95
0.95
0.95
0.95
0.95
0.95
0.95
0.95
0.95
0.95
0.95
0.95
0.95
0.95
0.95
0.95
0.95
0.95
0.95


4
0
0
0
0
0
0
0
0
0
0
0
0
3
0
0
0
0
0
0
0
0
0
0
0
0
0
0
1
0


4
0
0
0
0
0
0
0
0
0
0
0
0
3
0
0
0
0
0
0
0
0
0
0
0
0
0
0
1
0

0.84
0.84
0.84
0.84
0.84
0.84
0.84
0.84
0.84
0.84
0.84
0.84
0.84
0.84
0.84
0.84
0.84
0.84
0.84
0.84
0.84
0.84
0.84
0.84
0.84
0.84
0.84
0.84
0.84
0.84


4
0
0
0
0
0
0
0
0
0
0
0
0
0
0
0
0
0
0
0
0
0
0
0
0
2
2
0
0
0


4
0
0
0
0
0
0
0
0
0
0
0
0
0
0
0
0
0
0
0
0
0
0
0
0
2
2
0
0
0


4
0
0
0
0
0
0
0
0
0
0
0
0
0
0
0
0
0
0
0
0
0
0
0
0
2
2
0
0
0

1.0
1.0
1.0
1.0
1.0
1.0
1.0
1.0
1.0
1.0
1.0
1.0
1.0
1.0
1.0
1.0
1.0
1.0
1.0
1.0
1.0
1.0
1.0
1.0
1.0
1.0
1.0
1.0
1.0
1.0


2
0
0
0
0
0
0
0
0
0
0
0
0
0
0
0
0
0
0
0
0
0
0
0
0
0
2
0
0
0


2
0
0
0
0
0
0
0
0
0
0
0
0
0
0
0
0
0
0
0
0
0
0
0
0
0
2
0
0
0

1.0
1.0
1.0
1.0
1.0
1.0
1.0
1.0
1.0
1.0
1.0
1.0
1.0
1.0
1.0
1.0
1.0
1.0
1.0
1.0
1.0
1.0
1.0
1.0
1.0
1.0
1.0
1.0
1.0
1.0


5
0
0
0
0
0
0
0
0
0
0
0
0
0
0
0
0
0
0
0
0
0
0
0
0
3
0
2
0
0


5
0
0
0
0
0
0
0
0
0
0
0
0
0
0
0
0
0
0
0
0
0
0
0
0
3
0
2
0
0


5
0
0
0
0
0
0
0
0
0
0
0
0
0
0
0
0
0
0
0
0
0
0
0
0
3
0
2
0
0

1.0
1.0
1.0
1.0
1.0
1.0
1.0
1.0
1.0
1.0
1.0
1.0
1.0
1.0
1.0
1.0
1.0
1.0
1.0
1.0
1.0
1.0
1.0
1.0
1.0
1.0
1.0
1.0
1.0
1.0


9
0
0
0
0
0
0
0
0
0
0
0
0
1
1
0
0
0
0
1
1
1
0
0
1
0
0
1
1
1


9
0
0
0
0
0
0
0
0
0
0
0
0
1
1
0
0
0
0
1
1
1
0
0
1
0
0
1
1
1


9
0
0
0
0
0
0
0
0
0
0
0
0
1
1
0
0
0
0
1
1
1
0
0
1
0
0
1
1
1

1.0
1.0
1.0
1.0
1.0
1.0
1.0
1.0
1.0
1.0
1.0
1.0
1.0
1.0
1.0
1.0
1.0
1.0
1.0
1.0
1.0
1.0
1.0
1.0
1.0
1.0
1.0
1.0
1.0
1.0


6
0
0
0
0
0
0
0
0
0
0
0
0
0
0
0
0
0
0
0
0
0
0
0
0
1
3
2
0
0

0.73
0.73
0.73
0.73
0.73
0.73
0.73
0.73
0.73
0.73
0.73
0.73
0.73
0.73
0.73
0.73
0.73
0.73
0.73
0.73
0.73
0.73
0.73
0.73
0.73
0.73
0.73
0.73
0.73
0.73


1
0
0
0
0
0
0
0
0
0
0
0
0
0
0
0
0
0
0
0
0
0
0
0
0
0
1
0
0
0


1
0
0
0
0
0
0
0
0
0
0
0
0
0
0
0
0
0
0
0
0
0
0
0
0
0
1
0
0
0


1
0
0
0
0
0
0
0
0
0
0
0
0
0
0
0
0
0
0
0
0
0
0
0
0
0
1
0
0
0

1.0
1.0
1.0
1.0
1.0
1.0
1.0
1.0
1.0
1.0
1.0
1.0
1.0
1.0
1.0
1.0
1.0
1.0
1.0
1.0
1.0
1.0
1.0
1.0
1.0
1.0
1.0
1.0
1.0
1.0


1078
90
136
126
39
46
45
14
23
45
66
71
3
20
23
0
29
91
14
8
5
6
8
3
56
11
23
13
31
33

0.96
0.96
0.96
0.96
0.96
0.96
0.96
0.96
0.96
0.96
0.96
0.96
0.96
0.96
0.96
0.96
0.96
0.96
0.96
0.96
0.96
0.96
0.96
0.96
0.96
0.96
0.96
0.96
0.96
0.96


51
7
9
4
5
0
2
0
0
1
6
4
0
3
4
0
0
0
0
0
0
0
0
0
1
0
2
1
1
1

1.0
1.0
1.0
1.0
1.0
1.0
1.0
1.0
1.0
1.0
1.0
1.0
1.0
1.0
1.0
1.0
1.0
1.0
1.0
1.0
1.0
1.0
1.0
1.0
1.0
1.0
1.0
1.0
1.0
1.0


33
6
6
3
0
0
0
0
0
1
6
4
0
3
4
0
0
0
0
0
0
0
0
0
0
0
0
0
0
0


33
6
6
3
0
0
0
0
0
1
6
4
0
3
4
0
0
0
0
0
0
0
0
0
0
0
0
0
0
0

1.0
1.0
1.0
1.0
1.0
1.0
1.0
1.0
1.0
1.0
1.0
1.0
1.0
1.0
1.0
1.0
1.0
1.0
1.0
1.0
1.0
1.0
1.0
1.0
1.0
1.0
1.0
1.0
1.0
1.0


2
0
0
0
2
0
0
0
0
0
0
0
0
0
0
0
0
0
0
0
0
0
0
0
0
0
0
0
0
0


2
0
0
0
2
0
0
0
0
0
0
0
0
0
0
0
0
0
0
0
0
0
0
0
0
0
0
0
0
0

0.99
0.99
0.99
0.99
0.99
0.99
0.99
0.99
0.99
0.99
0.99
0.99
0.99
0.99
0.99
0.99
0.99
0.99
0.99
0.99
0.99
0.99
0.99
0.99
0.99
0.99
0.99
0.99
0.99
0.99


12
1
2
1
1
0
2
0
0
0
0
0
0
0
0
0
0
0
0
0
0
0
0
0
0
0
2
1
1
1


12
1
2
1
1
0
2
0
0
0
0
0
0
0
0
0
0
0
0
0
0
0
0
0
0
0
2
1
1
1

1.0
1.0
1.0
1.0
1.0
1.0
1.0
1.0
1.0
1.0
1.0
1.0
1.0
1.0
1.0
1.0
1.0
1.0
1.0
1.0
1.0
1.0
1.0
1.0
1.0
1.0
1.0
1.0
1.0
1.0


1
0
1
0
0
0
0
0
0
0
0
0
0
0
0
0
0
0
0
0
0
0
0
0
0
0
0
0
0
0


1
0
1
0
0
0
0
0
0
0
0
0
0
0
0
0
0
0
0
0
0
0
0
0
0
0
0
0
0
0

1.0
1.0
1.0
1.0
1.0
1.0
1.0
1.0
1.0
1.0
1.0
1.0
1.0
1.0
1.0
1.0
1.0
1.0
1.0
1.0
1.0
1.0
1.0
1.0
1.0
1.0
1.0
1.0
1.0
1.0


61
1
9
6
12
14
9
0
0
0
0
1
0
2
6
0
0
0
0
0
0
0
0
0
1
0
0
0
0
0

1.0
1.0
1.0
1.0
1.0
1.0
1.0
1.0
1.0
1.0
1.0
1.0
1.0
1.0
1.0
1.0
1.0
1.0
1.0
1.0
1.0
1.0
1.0
1.0
1.0
1.0
1.0
1.0
1.0
1.0


57
1
9
6
12
14
5
0
0
0
0
1
0
2
6
0
0
0
0
0
0
0
0
0
1
0
0
0
0
0

1.0
1.0
1.0
1.0
1.0
1.0
1.0
1.0
1.0
1.0
1.0
1.0
1.0
1.0
1.0
1.0
1.0
1.0
1.0
1.0
1.0
1.0
1.0
1.0
1.0
1.0
1.0
1.0
1.0
1.0


14
1
3
4
3
3
0
0
0
0
0
0
0
0
0
0
0
0
0
0
0
0
0
0
0
0
0
0
0
0

1.0
1.0
1.0
1.0
1.0
1.0
1.0
1.0
1.0
1.0
1.0
1.0
1.0
1.0
1.0
1.0
1.0
1.0
1.0
1.0
1.0
1.0
1.0
1.0
1.0
1.0
1.0
1.0
1.0
1.0


10
0
1
1
1
1
1
0
0
0
0
0
0
2
2
0
0
0
0
0
0
0
0
0
1
0
0
0
0
0

0.96
0.96
0.96
0.96
0.96
0.96
0.96
0.96
0.96
0.96
0.96
0.96
0.96
0.96
0.96
0.96
0.96
0.96
0.96
0.96
0.96
0.96
0.96
0.96
0.96
0.96
0.96
0.96
0.96
0.96


22
0
3
1
6
8
4
0
0
0
0
0
0
0
0
0
0
0
0
0
0
0
0
0
0
0
0
0
0
0

1.0
1.0
1.0
1.0
1.0
1.0
1.0
1.0
1.0
1.0
1.0
1.0
1.0
1.0
1.0
1.0
1.0
1.0
1.0
1.0
1.0
1.0
1.0
1.0
1.0
1.0
1.0
1.0
1.0
1.0


6
0
1
0
1
1
0
0
0
0
0
0
0
0
3
0
0
0
0
0
0
0
0
0
0
0
0
0
0
0

1.0
1.0
1.0
1.0
1.0
1.0
1.0
1.0
1.0
1.0
1.0
1.0
1.0
1.0
1.0
1.0
1.0
1.0
1.0
1.0
1.0
1.0
1.0
1.0
1.0
1.0
1.0
1.0
1.0
1.0


2
0
1
0
0
1
0
0
0
0
0
0
0
0
0
0
0
0
0
0
0
0
0
0
0
0
0
0
0
0

1.0
1.0
1.0
1.0
1.0
1.0
1.0
1.0
1.0
1.0
1.0
1.0
1.0
1.0
1.0
1.0
1.0
1.0
1.0
1.0
1.0
1.0
1.0
1.0
1.0
1.0
1.0
1.0
1.0
1.0


1
0
0
0
0
0
0
0
0
0
0
1
0
0
0
0
0
0
0
0
0
0
0
0
0
0
0
0
0
0

1.0
1.0
1.0
1.0
1.0
1.0
1.0
1.0
1.0
1.0
1.0
1.0
1.0
1.0
1.0
1.0
1.0
1.0
1.0
1.0
1.0
1.0
1.0
1.0
1.0
1.0
1.0
1.0
1.0
1.0


1
0
0
0
0
0
0
0
0
0
0
0
0
0
1
0
0
0
0
0
0
0
0
0
0
0
0
0
0
0

1.0
1.0
1.0
1.0
1.0
1.0
1.0
1.0
1.0
1.0
1.0
1.0
1.0
1.0
1.0
1.0
1.0
1.0
1.0
1.0
1.0
1.0
1.0
1.0
1.0
1.0
1.0
1.0
1.0
1.0


2
0
0
0
0
0
2
0
0
0
0
0
0
0
0
0
0
0
0
0
0
0
0
0
0
0
0
0
0
0

0.91
0.91
0.91
0.91
0.91
0.91
0.91
0.91
0.91
0.91
0.91
0.91
0.91
0.91
0.91
0.91
0.91
0.91
0.91
0.91
0.91
0.91
0.91
0.91
0.91
0.91
0.91
0.91
0.91
0.91


14
0
0
0
0
7
7
0
0
0
0
0
0
0
0
0
0
0
0
0
0
0
0
0
0
0
0
0
0
0

0.97
0.97
0.97
0.97
0.97
0.97
0.97
0.97
0.97
0.97
0.97
0.97
0.97
0.97
0.97
0.97
0.97
0.97
0.97
0.97
0.97
0.97
0.97
0.97
0.97
0.97
0.97
0.97
0.97
0.97


6
0
0
0
0
3
3
0
0
0
0
0
0
0
0
0
0
0
0
0
0
0
0
0
0
0
0
0
0
0

0.95
0.95
0.95
0.95
0.95
0.95
0.95
0.95
0.95
0.95
0.95
0.95
0.95
0.95
0.95
0.95
0.95
0.95
0.95
0.95
0.95
0.95
0.95
0.95
0.95
0.95
0.95
0.95
0.95
0.95


115
1
37
67
4
1
0
0
0
0
0
4
0
0
0
0
0
1
0
0
0
0
0
0
0
0
0
0
0
0


114
1
37
67
4
1
0
0
0
0
0
4
0
0
0
0
0
0
0
0
0
0
0
0
0
0
0
0
0
0

1.0
1.0
1.0
1.0
1.0
1.0
1.0
1.0
1.0
1.0
1.0
1.0
1.0
1.0
1.0
1.0
1.0
1.0
1.0
1.0
1.0
1.0
1.0
1.0
1.0
1.0
1.0
1.0
1.0
1.0


112
1
36
66
4
1
0
0
0
0
0
4
0
0
0
0
0
0
0
0
0
0
0
0
0
0
0
0
0
0

1.0
1.0
1.0
1.0
1.0
1.0
1.0
1.0
1.0
1.0
1.0
1.0
1.0
1.0
1.0
1.0
1.0
1.0
1.0
1.0
1.0
1.0
1.0
1.0
1.0
1.0
1.0
1.0
1.0
1.0


1
0
0
0
0
0
0
0
0
0
0
0
0
0
0
0
0
1
0
0
0
0
0
0
0
0
0
0
0
0


1
0
0
0
0
0
0
0
0
0
0
0
0
0
0
0
0
1
0
0
0
0
0
0
0
0
0
0
0
0

1.0
1.0
1.0
1.0
1.0
1.0
1.0
1.0
1.0
1.0
1.0
1.0
1.0
1.0
1.0
1.0
1.0
1.0
1.0
1.0
1.0
1.0
1.0
1.0
1.0
1.0
1.0
1.0
1.0
1.0


35
0
8
4
0
0
0
0
0
0
8
14
0
1
0
0
0
0
0
0
0
0
0
0
0
0
0
0
0
0


28
0
6
3
0
0
0
0
0
0
8
10
0
1
0
0
0
0
0
0
0
0
0
0
0
0
0
0
0
0

0.99
0.99
0.99
0.99
0.99
0.99
0.99
0.99
0.99
0.99
0.99
0.99
0.99
0.99
0.99
0.99
0.99
0.99
0.99
0.99
0.99
0.99
0.99
0.99
0.99
0.99
0.99
0.99
0.99
0.99


6
0
2
1
0
0
0
0
0
0
0
3
0
0
0
0
0
0
0
0
0
0
0
0
0
0
0
0
0
0


6
0
2
1
0
0
0
0
0
0
0
3
0
0
0
0
0
0
0
0
0
0
0
0
0
0
0
0
0
0

1.0
1.0
1.0
1.0
1.0
1.0
1.0
1.0
1.0
1.0
1.0
1.0
1.0
1.0
1.0
1.0
1.0
1.0
1.0
1.0
1.0
1.0
1.0
1.0
1.0
1.0
1.0
1.0
1.0
1.0


1
0
0
0
0
0
0
0
0
0
0
1
0
0
0
0
0
0
0
0
0
0
0
0
0
0
0
0
0
0


1
0
0
0
0
0
0
0
0
0
0
1
0
0
0
0
0
0
0
0
0
0
0
0
0
0
0
0
0
0

1.0
1.0
1.0
1.0
1.0
1.0
1.0
1.0
1.0
1.0
1.0
1.0
1.0
1.0
1.0
1.0
1.0
1.0
1.0
1.0
1.0
1.0
1.0
1.0
1.0
1.0
1.0
1.0
1.0
1.0


55
11
8
0
0
0
0
1
5
12
11
5
0
0
1
0
0
1
0
0
0
0
0
0
0
0
0
0
0
0

0.99
0.99
0.99
0.99
0.99
0.99
0.99
0.99
0.99
0.99
0.99
0.99
0.99
0.99
0.99
0.99
0.99
0.99
0.99
0.99
0.99
0.99
0.99
0.99
0.99
0.99
0.99
0.99
0.99
0.99


32
6
4
0
0
0
0
1
4
7
7
3
0
0
0
0
0
0
0
0
0
0
0
0
0
0
0
0
0
0

0.96
0.96
0.96
0.96
0.96
0.96
0.96
0.96
0.96
0.96
0.96
0.96
0.96
0.96
0.96
0.96
0.96
0.96
0.96
0.96
0.96
0.96
0.96
0.96
0.96
0.96
0.96
0.96
0.96
0.96


3
0
1
0
0
0
0
0
0
2
0
0
0
0
0
0
0
0
0
0
0
0
0
0
0
0
0
0
0
0

0.94
0.94
0.94
0.94
0.94
0.94
0.94
0.94
0.94
0.94
0.94
0.94
0.94
0.94
0.94
0.94
0.94
0.94
0.94
0.94
0.94
0.94
0.94
0.94
0.94
0.94
0.94
0.94
0.94
0.94


2
1
0
0
0
0
0
0
0
0
1
0
0
0
0
0
0
0
0
0
0
0
0
0
0
0
0
0
0
0


2
1
0
0
0
0
0
0
0
0
1
0
0
0
0
0
0
0
0
0
0
0
0
0
0
0
0
0
0
0

0.94
0.94
0.94
0.94
0.94
0.94
0.94
0.94
0.94
0.94
0.94
0.94
0.94
0.94
0.94
0.94
0.94
0.94
0.94
0.94
0.94
0.94
0.94
0.94
0.94
0.94
0.94
0.94
0.94
0.94


5
1
1
0
0
0
0
0
0
1
1
1
0
0
0
0
0
0
0
0
0
0
0
0
0
0
0
0
0
0

1.0
1.0
1.0
1.0
1.0
1.0
1.0
1.0
1.0
1.0
1.0
1.0
1.0
1.0
1.0
1.0
1.0
1.0
1.0
1.0
1.0
1.0
1.0
1.0
1.0
1.0
1.0
1.0
1.0
1.0


35
4
3
3
3
3
3
0
0
1
2
3
0
3
2
0
0
0
0
0
0
0
0
0
5
0
0
0
0
0

0.93
0.93
0.93
0.93
0.93
0.93
0.93
0.93
0.93
0.93
0.93
0.93
0.93
0.93
0.93
0.93
0.93
0.93
0.93
0.93
0.93
0.93
0.93
0.93
0.93
0.93
0.93
0.93
0.93
0.93


16
2
2
2
2
2
2
0
0
1
1
2
0
0
0
0
0
0
0
0
0
0
0
0
0
0
0
0
0
0

0.8
0.8
0.8
0.8
0.8
0.8
0.8
0.8
0.8
0.8
0.8
0.8
0.8
0.8
0.8
0.8
0.8
0.8
0.8
0.8
0.8
0.8
0.8
0.8
0.8
0.8
0.8
0.8
0.8
0.8


9
0
0
0
0
0
0
0
0
0
0
0
0
2
2
0
0
0
0
0
0
0
0
0
5
0
0
0
0
0

0.99
0.99
0.99
0.99
0.99
0.99
0.99
0.99
0.99
0.99
0.99
0.99
0.99
0.99
0.99
0.99
0.99
0.99
0.99
0.99
0.99
0.99
0.99
0.99
0.99
0.99
0.99
0.99
0.99
0.99


156
11
18
6
4
1
4
2
2
1
4
8
3
8
2
0
11
3
2
3
0
0
2
3
0
3
3
7
27
18

1.0
1.0
1.0
1.0
1.0
1.0
1.0
1.0
1.0
1.0
1.0
1.0
1.0
1.0
1.0
1.0
1.0
1.0
1.0
1.0
1.0
1.0
1.0
1.0
1.0
1.0
1.0
1.0
1.0
1.0


4
0
4
0
0
0
0
0
0
0
0
0
0
0
0
0
0
0
0
0
0
0
0
0
0
0
0
0
0
0

1.0
1.0
1.0
1.0
1.0
1.0
1.0
1.0
1.0
1.0
1.0
1.0
1.0
1.0
1.0
1.0
1.0
1.0
1.0
1.0
1.0
1.0
1.0
1.0
1.0
1.0
1.0
1.0
1.0
1.0


13
0
1
1
2
0
0
2
1
0
0
0
2
0
0
0
0
0
0
0
0
0
2
2
0
0
0
0
0
0


9
0
0
0
0
0
0
2
1
0
0
0
2
0
0
0
0
0
0
0
0
0
2
2
0
0
0
0
0
0

1.0
1.0
1.0
1.0
1.0
1.0
1.0
1.0
1.0
1.0
1.0
1.0
1.0
1.0
1.0
1.0
1.0
1.0
1.0
1.0
1.0
1.0
1.0
1.0
1.0
1.0
1.0
1.0
1.0
1.0


4
0
1
1
2
0
0
0
0
0
0
0
0
0
0
0
0
0
0
0
0
0
0
0
0
0
0
0
0
0

1.0
1.0
1.0
1.0
1.0
1.0
1.0
1.0
1.0
1.0
1.0
1.0
1.0
1.0
1.0
1.0
1.0
1.0
1.0
1.0
1.0
1.0
1.0
1.0
1.0
1.0
1.0
1.0
1.0
1.0


6
0
0
0
1
0
0
0
0
0
1
1
0
0
0
0
0
0
0
0
0
0
0
0
0
0
0
0
3
0

1.0
1.0
1.0
1.0
1.0
1.0
1.0
1.0
1.0
1.0
1.0
1.0
1.0
1.0
1.0
1.0
1.0
1.0
1.0
1.0
1.0
1.0
1.0
1.0
1.0
1.0
1.0
1.0
1.0
1.0


2
0
0
0
0
0
0
0
0
0
1
1
0
0
0
0
0
0
0
0
0
0
0
0
0
0
0
0
0
0

1.0
1.0
1.0
1.0
1.0
1.0
1.0
1.0
1.0
1.0
1.0
1.0
1.0
1.0
1.0
1.0
1.0
1.0
1.0
1.0
1.0
1.0
1.0
1.0
1.0
1.0
1.0
1.0
1.0
1.0


46
1
1
0
0
0
3
0
0
0
0
0
0
0
0
0
7
0
0
0
0
0
0
0
0
0
2
4
18
10


43
0
0
0
0
0
2
0
0
0
0
0
0
0
0
0
7
0
0
0
0
0
0
0
0
0
2
4
18
10

1.0
1.0
1.0
1.0
1.0
1.0
1.0
1.0
1.0
1.0
1.0
1.0
1.0
1.0
1.0
1.0
1.0
1.0
1.0
1.0
1.0
1.0
1.0
1.0
1.0
1.0
1.0
1.0
1.0
1.0


1
0
1
0
0
0
0
0
0
0
0
0
0
0
0
0
0
0
0
0
0
0
0
0
0
0
0
0
0
0

1.0
1.0
1.0
1.0
1.0
1.0
1.0
1.0
1.0
1.0
1.0
1.0
1.0
1.0
1.0
1.0
1.0
1.0
1.0
1.0
1.0
1.0
1.0
1.0
1.0
1.0
1.0
1.0
1.0
1.0


1
0
0
0
0
0
1
0
0
0
0
0
0
0
0
0
0
0
0
0
0
0
0
0
0
0
0
0
0
0

1.0
1.0
1.0
1.0
1.0
1.0
1.0
1.0
1.0
1.0
1.0
1.0
1.0
1.0
1.0
1.0
1.0
1.0
1.0
1.0
1.0
1.0
1.0
1.0
1.0
1.0
1.0
1.0
1.0
1.0


1
1
0
0
0
0
0
0
0
0
0
0
0
0
0
0
0
0
0
0
0
0
0
0
0
0
0
0
0
0

1.0
1.0
1.0
1.0
1.0
1.0
1.0
1.0
1.0
1.0
1.0
1.0
1.0
1.0
1.0
1.0
1.0
1.0
1.0
1.0
1.0
1.0
1.0
1.0
1.0
1.0
1.0
1.0
1.0
1.0


1
0
1
0
0
0
0
0
0
0
0
0
0
0
0
0
0
0
0
0
0
0
0
0
0
0
0
0
0
0


1
0
1
0
0
0
0
0
0
0
0
0
0
0
0
0
0
0
0
0
0
0
0
0
0
0
0
0
0
0

1.0
1.0
1.0
1.0
1.0
1.0
1.0
1.0
1.0
1.0
1.0
1.0
1.0
1.0
1.0
1.0
1.0
1.0
1.0
1.0
1.0
1.0
1.0
1.0
1.0
1.0
1.0
1.0
1.0
1.0


18
5
4
1
0
0
0
0
0
0
3
5
0
0
0
0
0
0
0
0
0
0
0
0
0
0
0
0
0
0

1.0
1.0
1.0
1.0
1.0
1.0
1.0
1.0
1.0
1.0
1.0
1.0
1.0
1.0
1.0
1.0
1.0
1.0
1.0
1.0
1.0
1.0
1.0
1.0
1.0
1.0
1.0
1.0
1.0
1.0


13
4
3
0
0
0
0
0
0
0
3
3
0
0
0
0
0
0
0
0
0
0
0
0
0
0
0
0
0
0

1.0
1.0
1.0
1.0
1.0
1.0
1.0
1.0
1.0
1.0
1.0
1.0
1.0
1.0
1.0
1.0
1.0
1.0
1.0
1.0
1.0
1.0
1.0
1.0
1.0
1.0
1.0
1.0
1.0
1.0


1
0
0
1
0
0
0
0
0
0
0
0
0
0
0
0
0
0
0
0
0
0
0
0
0
0
0
0
0
0

1.0
1.0
1.0
1.0
1.0
1.0
1.0
1.0
1.0
1.0
1.0
1.0
1.0
1.0
1.0
1.0
1.0
1.0
1.0
1.0
1.0
1.0
1.0
1.0
1.0
1.0
1.0
1.0
1.0
1.0


6
3
0
0
0
1
0
0
0
0
0
2
0
0
0
0
0
0
0
0
0
0
0
0
0
0
0
0
0
0

1.0
1.0
1.0
1.0
1.0
1.0
1.0
1.0
1.0
1.0
1.0
1.0
1.0
1.0
1.0
1.0
1.0
1.0
1.0
1.0
1.0
1.0
1.0
1.0
1.0
1.0
1.0
1.0
1.0
1.0


1
1
0
0
0
0
0
0
0
0
0
0
0
0
0
0
0
0
0
0
0
0
0
0
0
0
0
0
0
0


1
1
0
0
0
0
0
0
0
0
0
0
0
0
0
0
0
0
0
0
0
0
0
0
0
0
0
0
0
0

0.85
0.85
0.85
0.85
0.85
0.85
0.85
0.85
0.85
0.85
0.85
0.85
0.85
0.85
0.85
0.85
0.85
0.85
0.85
0.85
0.85
0.85
0.85
0.85
0.85
0.85
0.85
0.85
0.85
0.85


10
0
2
0
0
0
0
0
0
0
0
0
0
0
0
0
4
0
2
2
0
0
0
0
0
0
0
0
0
0


10
0
2
0
0
0
0
0
0
0
0
0
0
0
0
0
4
0
2
2
0
0
0
0
0
0
0
0
0
0

1.0
1.0
1.0
1.0
1.0
1.0
1.0
1.0
1.0
1.0
1.0
1.0
1.0
1.0
1.0
1.0
1.0
1.0
1.0
1.0
1.0
1.0
1.0
1.0
1.0
1.0
1.0
1.0
1.0
1.0


38
1
4
3
1
0
1
0
1
1
0
0
0
4
2
0
0
3
0
0
0
0
0
0
0
3
0
3
5
6

1.0
1.0
1.0
1.0
1.0
1.0
1.0
1.0
1.0
1.0
1.0
1.0
1.0
1.0
1.0
1.0
1.0
1.0
1.0
1.0
1.0
1.0
1.0
1.0
1.0
1.0
1.0
1.0
1.0
1.0


13
1
2
2
1
0
0
0
1
0
0
0
0
4
2
0
0
0
0
0
0
0
0
0
0
0
0
0
0
0

1.0
1.0
1.0
1.0
1.0
1.0
1.0
1.0
1.0
1.0
1.0
1.0
1.0
1.0
1.0
1.0
1.0
1.0
1.0
1.0
1.0
1.0
1.0
1.0
1.0
1.0
1.0
1.0
1.0
1.0


19
0
0
0
0
0
1
0
0
1
0
0
0
0
0
0
0
0
0
0
0
0
0
0
0
3
0
3
5
6

1.0
1.0
1.0
1.0
1.0
1.0
1.0
1.0
1.0
1.0
1.0
1.0
1.0
1.0
1.0
1.0
1.0
1.0
1.0
1.0
1.0
1.0
1.0
1.0
1.0
1.0
1.0
1.0
1.0
1.0


1
0
0
1
0
0
0
0
0
0
0
0
0
0
0
0
0
0
0
0
0
0
0
0
0
0
0
0
0
0

1.0
1.0
1.0
1.0
1.0
1.0
1.0
1.0
1.0
1.0
1.0
1.0
1.0
1.0
1.0
1.0
1.0
1.0
1.0
1.0
1.0
1.0
1.0
1.0
1.0
1.0
1.0
1.0
1.0
1.0


1
0
1
0
0
0
0
0
0
0
0
0
0
0
0
0
0
0
0
0
0
0
0
0
0
0
0
0
0
0

0.94
0.94
0.94
0.94
0.94
0.94
0.94
0.94
0.94
0.94
0.94
0.94
0.94
0.94
0.94
0.94
0.94
0.94
0.94
0.94
0.94
0.94
0.94
0.94
0.94
0.94
0.94
0.94
0.94
0.94


3
0
0
0
0
0
0
0
0
0
0
0
0
0
0
0
0
3
0
0
0
0
0
0
0
0
0
0
0
0

1.0
1.0
1.0
1.0
1.0
1.0
1.0
1.0
1.0
1.0
1.0
1.0
1.0
1.0
1.0
1.0
1.0
1.0
1.0
1.0
1.0
1.0
1.0
1.0
1.0
1.0
1.0
1.0
1.0
1.0


4
0
0
0
0
0
0
0
0
0
0
0
0
4
0
0
0
0
0
0
0
0
0
0
0
0
0
0
0
0


4
0
0
0
0
0
0
0
0
0
0
0
0
4
0
0
0
0
0
0
0
0
0
0
0
0
0
0
0
0

0.84
0.84
0.84
0.84
0.84
0.84
0.84
0.84
0.84
0.84
0.84
0.84
0.84
0.84
0.84
0.84
0.84
0.84
0.84
0.84
0.84
0.84
0.84
0.84
0.84
0.84
0.84
0.84
0.84
0.84


4
0
0
0
0
0
0
0
0
0
0
0
0
0
0
0
0
0
0
0
0
0
0
0
0
0
1
0
1
2

0.77
0.77
0.77
0.77
0.77
0.77
0.77
0.77
0.77
0.77
0.77
0.77
0.77
0.77
0.77
0.77
0.77
0.77
0.77
0.77
0.77
0.77
0.77
0.77
0.77
0.77
0.77
0.77
0.77
0.77


3
0
0
0
0
0
0
0
0
0
0
0
1
0
0
0
0
0
0
1
0
0
0
1
0
0
0
0
0
0


3
0
0
0
0
0
0
0
0
0
0
0
1
0
0
0
0
0
0
1
0
0
0
1
0
0
0
0
0
0

1.0
1.0
1.0
1.0
1.0
1.0
1.0
1.0
1.0
1.0
1.0
1.0
1.0
1.0
1.0
1.0
1.0
1.0
1.0
1.0
1.0
1.0
1.0
1.0
1.0
1.0
1.0
1.0
1.0
1.0


141
13
4
4
2
1
0
3
6
9
14
6
0
0
4
0
0
55
6
0
0
6
5
0
2
1
0
0
0
0

0.98
0.98
0.98
0.98
0.98
0.98
0.98
0.98
0.98
0.98
0.98
0.98
0.98
0.98
0.98
0.98
0.98
0.98
0.98
0.98
0.98
0.98
0.98
0.98
0.98
0.98
0.98
0.98
0.98
0.98


4
0
0
0
0
0
0
0
0
0
4
0
0
0
0
0
0
0
0
0
0
0
0
0
0
0
0
0
0
0

0.97
0.97
0.97
0.97
0.97
0.97
0.97
0.97
0.97
0.97
0.97
0.97
0.97
0.97
0.97
0.97
0.97
0.97
0.97
0.97
0.97
0.97
0.97
0.97
0.97
0.97
0.97
0.97
0.97
0.97


19
5
1
0
0
0
0
1
1
1
2
0
0
0
0
0
0
0
1
0
0
3
4
0
0
0
0
0
0
0

1.0
1.0
1.0
1.0
1.0
1.0
1.0
1.0
1.0
1.0
1.0
1.0
1.0
1.0
1.0
1.0
1.0
1.0
1.0
1.0
1.0
1.0
1.0
1.0
1.0
1.0
1.0
1.0
1.0
1.0


1
0
0
0
0
0
0
0
0
0
1
0
0
0
0
0
0
0
0
0
0
0
0
0
0
0
0
0
0
0

1.0
1.0
1.0
1.0
1.0
1.0
1.0
1.0
1.0
1.0
1.0
1.0
1.0
1.0
1.0
1.0
1.0
1.0
1.0
1.0
1.0
1.0
1.0
1.0
1.0
1.0
1.0
1.0
1.0
1.0


2
1
0
0
0
0
0
0
0
0
0
0
0
0
0
0
0
0
0
0
0
0
1
0
0
0
0
0
0
0

1.0
1.0
1.0
1.0
1.0
1.0
1.0
1.0
1.0
1.0
1.0
1.0
1.0
1.0
1.0
1.0
1.0
1.0
1.0
1.0
1.0
1.0
1.0
1.0
1.0
1.0
1.0
1.0
1.0
1.0


51
1
0
0
0
0
0
1
1
1
1
1
0
0
0
0
0
45
0
0
0
0
0
0
0
0
0
0
0
0

0.98
0.98
0.98
0.98
0.98
0.98
0.98
0.98
0.98
0.98
0.98
0.98
0.98
0.98
0.98
0.98
0.98
0.98
0.98
0.98
0.98
0.98
0.98
0.98
0.98
0.98
0.98
0.98
0.98
0.98


7
1
1
0
0
0
0
0
1
1
1
1
0
0
0
0
0
0
0
0
0
0
0
0
1
0
0
0
0
0


7
1
1
0
0
0
0
0
1
1
1
1
0
0
0
0
0
0
0
0
0
0
0
0
1
0
0
0
0
0

1.0
1.0
1.0
1.0
1.0
1.0
1.0
1.0
1.0
1.0
1.0
1.0
1.0
1.0
1.0
1.0
1.0
1.0
1.0
1.0
1.0
1.0
1.0
1.0
1.0
1.0
1.0
1.0
1.0
1.0


4
1
0
0
0
0
0
0
0
0
1
0
0
0
1
0
0
1
0
0
0
0
0
0
0
0
0
0
0
0


4
1
0
0
0
0
0
0
0
0
1
0
0
0
1
0
0
1
0
0
0
0
0
0
0
0
0
0
0
0

0.98
0.98
0.98
0.98
0.98
0.98
0.98
0.98
0.98
0.98
0.98
0.98
0.98
0.98
0.98
0.98
0.98
0.98
0.98
0.98
0.98
0.98
0.98
0.98
0.98
0.98
0.98
0.98
0.98
0.98


9
1
1
3
2
1
0
0
0
0
0
0
0
0
0
0
0
0
0
0
0
0
0
0
1
0
0
0
0
0


9
1
1
3
2
1
0
0
0
0
0
0
0
0
0
0
0
0
0
0
0
0
0
0
1
0
0
0
0
0

0.95
0.95
0.95
0.95
0.95
0.95
0.95
0.95
0.95
0.95
0.95
0.95
0.95
0.95
0.95
0.95
0.95
0.95
0.95
0.95
0.95
0.95
0.95
0.95
0.95
0.95
0.95
0.95
0.95
0.95


2
0
0
0
0
0
0
0
0
1
1
0
0
0
0
0
0
0
0
0
0
0
0
0
0
0
0
0
0
0

1.0
1.0
1.0
1.0
1.0
1.0
1.0
1.0
1.0
1.0
1.0
1.0
1.0
1.0
1.0
1.0
1.0
1.0
1.0
1.0
1.0
1.0
1.0
1.0
1.0
1.0
1.0
1.0
1.0
1.0


2
2
0
0
0
0
0
0
0
0
0
0
0
0
0
0
0
0
0
0
0
0
0
0
0
0
0
0
0
0

1.0
1.0
1.0
1.0
1.0
1.0
1.0
1.0
1.0
1.0
1.0
1.0
1.0
1.0
1.0
1.0
1.0
1.0
1.0
1.0
1.0
1.0
1.0
1.0
1.0
1.0
1.0
1.0
1.0
1.0


3
0
0
0
0
0
0
0
0
2
0
1
0
0
0
0
0
0
0
0
0
0
0
0
0
0
0
0
0
0


1
0
0
0
0
0
0
0
0
0
0
1
0
0
0
0
0
0
0
0
0
0
0
0
0
0
0
0
0
0

0.82
0.82
0.82
0.82
0.82
0.82
0.82
0.82
0.82
0.82
0.82
0.82
0.82
0.82
0.82
0.82
0.82
0.82
0.82
0.82
0.82
0.82
0.82
0.82
0.82
0.82
0.82
0.82
0.82
0.82


2
0
0
0
0
0
0
0
0
2
0
0
0
0
0
0
0
0
0
0
0
0
0
0
0
0
0
0
0
0

0.93
0.93
0.93
0.93
0.93
0.93
0.93
0.93
0.93
0.93
0.93
0.93
0.93
0.93
0.93
0.93
0.93
0.93
0.93
0.93
0.93
0.93
0.93
0.93
0.93
0.93
0.93
0.93
0.93
0.93


5
0
0
1
0
0
0
0
0
0
0
0
0
0
2
0
0
0
2
0
0
0
0
0
0
0
0
0
0
0

1.0
1.0
1.0
1.0
1.0
1.0
1.0
1.0
1.0
1.0
1.0
1.0
1.0
1.0
1.0
1.0
1.0
1.0
1.0
1.0
1.0
1.0
1.0
1.0
1.0
1.0
1.0
1.0
1.0
1.0


1
0
0
0
0
0
0
0
1
0
0
0
0
0
0
0
0
0
0
0
0
0
0
0
0
0
0
0
0
0


1
0
0
0
0
0
0
0
1
0
0
0
0
0
0
0
0
0
0
0
0
0
0
0
0
0
0
0
0
0

1.0
1.0
1.0
1.0
1.0
1.0
1.0
1.0
1.0
1.0
1.0
1.0
1.0
1.0
1.0
1.0
1.0
1.0
1.0
1.0
1.0
1.0
1.0
1.0
1.0
1.0
1.0
1.0
1.0
1.0


5
0
0
0
0
0
0
0
0
1
0
1
0
0
0
0
0
1
2
0
0
0
0
0
0
0
0
0
0
0


4
0
0
0
0
0
0
0
0
0
0
1
0
0
0
0
0
1
2
0
0
0
0
0
0
0
0
0
0
0

1.0
1.0
1.0
1.0
1.0
1.0
1.0
1.0
1.0
1.0
1.0
1.0
1.0
1.0
1.0
1.0
1.0
1.0
1.0
1.0
1.0
1.0
1.0
1.0
1.0
1.0
1.0
1.0
1.0
1.0


1
0
0
0
0
0
0
0
0
1
0
0
0
0
0
0
0
0
0
0
0
0
0
0
0
0
0
0
0
0

1.0
1.0
1.0
1.0
1.0
1.0
1.0
1.0
1.0
1.0
1.0
1.0
1.0
1.0
1.0
1.0
1.0
1.0
1.0
1.0
1.0
1.0
1.0
1.0
1.0
1.0
1.0
1.0
1.0
1.0


1
1
0
0
0
0
0
0
0
0
0
0
0
0
0
0
0
0
0
0
0
0
0
0
0
0
0
0
0
0

0.99
0.99
0.99
0.99
0.99
0.99
0.99
0.99
0.99
0.99
0.99
0.99
0.99
0.99
0.99
0.99
0.99
0.99
0.99
0.99
0.99
0.99
0.99
0.99
0.99
0.99
0.99
0.99
0.99
0.99


1
0
0
0
0
0
0
0
0
0
0
0
0
0
1
0
0
0
0
0
0
0
0
0
0
0
0
0
0
0


1
0
0
0
0
0
0
0
0
0
0
0
0
0
1
0
0
0
0
0
0
0
0
0
0
0
0
0
0
0

1.0
1.0
1.0
1.0
1.0
1.0
1.0
1.0
1.0
1.0
1.0
1.0
1.0
1.0
1.0
1.0
1.0
1.0
1.0
1.0
1.0
1.0
1.0
1.0
1.0
1.0
1.0
1.0
1.0
1.0


1
0
0
0
0
0
0
0
0
0
0
0
0
0
0
0
0
0
0
0
0
1
0
0
0
0
0
0
0
0

0.89
0.89
0.89
0.89
0.89
0.89
0.89
0.89
0.89
0.89
0.89
0.89
0.89
0.89
0.89
0.89
0.89
0.89
0.89
0.89
0.89
0.89
0.89
0.89
0.89
0.89
0.89
0.89
0.89
0.89


1
0
0
0
0
0
0
0
0
0
0
0
0
0
0
0
0
0
0
0
0
0
0
0
0
1
0
0
0
0


1
0
0
0
0
0
0
0
0
0
0
0
0
0
0
0
0
0
0
0
0
0
0
0
0
1
0
0
0
0

0.83
0.83
0.83
0.83
0.83
0.83
0.83
0.83
0.83
0.83
0.83
0.83
0.83
0.83
0.83
0.83
0.83
0.83
0.83
0.83
0.83
0.83
0.83
0.83
0.83
0.83
0.83
0.83
0.83
0.83


32
8
6
6
0
0
0
0
0
0
5
7
0
0
0
0
0
0
0
0
0
0
0
0
0
0
0
0
0
0


32
8
6
6
0
0
0
0
0
0
5
7
0
0
0
0
0
0
0
0
0
0
0
0
0
0
0
0
0
0


27
7
5
5
0
0
0
0
0
0
4
6
0
0
0
0
0
0
0
0
0
0
0
0
0
0
0
0
0
0

1.0
1.0
1.0
1.0
1.0
1.0
1.0
1.0
1.0
1.0
1.0
1.0
1.0
1.0
1.0
1.0
1.0
1.0
1.0
1.0
1.0
1.0
1.0
1.0
1.0
1.0
1.0
1.0
1.0
1.0


5
1
1
1
0
0
0
0
0
0
1
1
0
0
0
0
0
0
0
0
0
0
0
0
0
0
0
0
0
0

1.0
1.0
1.0
1.0
1.0
1.0
1.0
1.0
1.0
1.0
1.0
1.0
1.0
1.0
1.0
1.0
1.0
1.0
1.0
1.0
1.0
1.0
1.0
1.0
1.0
1.0
1.0
1.0
1.0
1.0


14
0
2
1
0
4
4
0
0
0
0
0
0
0
3
0
0
0
0
0
0
0
0
0
0
0
0
0
0
0


8
0
0
0
0
4
4
0
0
0
0
0
0
0
0
0
0
0
0
0
0
0
0
0
0
0
0
0
0
0

1.0
1.0
1.0
1.0
1.0
1.0
1.0
1.0
1.0
1.0
1.0
1.0
1.0
1.0
1.0
1.0
1.0
1.0
1.0
1.0
1.0
1.0
1.0
1.0
1.0
1.0
1.0
1.0
1.0
1.0


2
0
1
1
0
0
0
0
0
0
0
0
0
0
0
0
0
0
0
0
0
0
0
0
0
0
0
0
0
0


2
0
1
1
0
0
0
0
0
0
0
0
0
0
0
0
0
0
0
0
0
0
0
0
0
0
0
0
0
0

1.0
1.0
1.0
1.0
1.0
1.0
1.0
1.0
1.0
1.0
1.0
1.0
1.0
1.0
1.0
1.0
1.0
1.0
1.0
1.0
1.0
1.0
1.0
1.0
1.0
1.0
1.0
1.0
1.0
1.0


4
0
1
0
0
0
0
0
0
0
0
0
0
0
3
0
0
0
0
0
0
0
0
0
0
0
0
0
0
0


4
0
1
0
0
0
0
0
0
0
0
0
0
0
3
0
0
0
0
0
0
0
0
0
0
0
0
0
0
0

1.0
1.0
1.0
1.0
1.0
1.0
1.0
1.0
1.0
1.0
1.0
1.0
1.0
1.0
1.0
1.0
1.0
1.0
1.0
1.0
1.0
1.0
1.0
1.0
1.0
1.0
1.0
1.0
1.0
1.0


11
3
1
0
0
0
1
0
0
1
1
3
0
0
0
0
0
1
0
0
0
0
0
0
0
0
0
0
0
0


8
3
1
0
0
0
0
0
0
0
1
3
0
0
0
0
0
0
0
0
0
0
0
0
0
0
0
0
0
0


8
3
1
0
0
0
0
0
0
0
1
3
0
0
0
0
0
0
0
0
0
0
0
0
0
0
0
0
0
0

0.93
0.93
0.93
0.93
0.93
0.93
0.93
0.93
0.93
0.93
0.93
0.93
0.93
0.93
0.93
0.93
0.93
0.93
0.93
0.93
0.93
0.93
0.93
0.93
0.93
0.93
0.93
0.93
0.93
0.93


1
0
0
0
0
0
1
0
0
0
0
0
0
0
0
0
0
0
0
0
0
0
0
0
0
0
0
0
0
0

1.0
1.0
1.0
1.0
1.0
1.0
1.0
1.0
1.0
1.0
1.0
1.0
1.0
1.0
1.0
1.0
1.0
1.0
1.0
1.0
1.0
1.0
1.0
1.0
1.0
1.0
1.0
1.0
1.0
1.0


2
0
0
0
0
0
0
0
0
1
0
0
0
0
0
0
0
1
0
0
0
0
0
0
0
0
0
0
0
0


1
0
0
0
0
0
0
0
0
1
0
0
0
0
0
0
0
0
0
0
0
0
0
0
0
0
0
0
0
0

1.0
1.0
1.0
1.0
1.0
1.0
1.0
1.0
1.0
1.0
1.0
1.0
1.0
1.0
1.0
1.0
1.0
1.0
1.0
1.0
1.0
1.0
1.0
1.0
1.0
1.0
1.0
1.0
1.0
1.0


1
0
0
0
0
0
0
0
0
0
0
0
0
0
0
0
0
1
0
0
0
0
0
0
0
0
0
0
0
0

1.0
1.0
1.0
1.0
1.0
1.0
1.0
1.0
1.0
1.0
1.0
1.0
1.0
1.0
1.0
1.0
1.0
1.0
1.0
1.0
1.0
1.0
1.0
1.0
1.0
1.0
1.0
1.0
1.0
1.0


8
2
2
4
0
0
0
0
0
0
0
0
0
0
0
0
0
0
0
0
0
0
0
0
0
0
0
0
0
0


8
2
2
4
0
0
0
0
0
0
0
0
0
0
0
0
0
0
0
0
0
0
0
0
0
0
0
0
0
0

1.0
1.0
1.0
1.0
1.0
1.0
1.0
1.0
1.0
1.0
1.0
1.0
1.0
1.0
1.0
1.0
1.0
1.0
1.0
1.0
1.0
1.0
1.0
1.0
1.0
1.0
1.0
1.0
1.0
1.0


3
1
1
1
0
0
0
0
0
0
0
0
0
0
0
0
0
0
0
0
0
0
0
0
0
0
0
0
0
0

1.0
1.0
1.0
1.0
1.0
1.0
1.0
1.0
1.0
1.0
1.0
1.0
1.0
1.0
1.0
1.0
1.0
1.0
1.0
1.0
1.0
1.0
1.0
1.0
1.0
1.0
1.0
1.0
1.0
1.0


2
0
0
2
0
0
0
0
0
0
0
0
0
0
0
0
0
0
0
0
0
0
0
0
0
0
0
0
0
0

1.0
1.0
1.0
1.0
1.0
1.0
1.0
1.0
1.0
1.0
1.0
1.0
1.0
1.0
1.0
1.0
1.0
1.0
1.0
1.0
1.0
1.0
1.0
1.0
1.0
1.0
1.0
1.0
1.0
1.0


37
7
4
1
1
1
4
2
4
4
2
3
0
0
0
0
0
0
0
2
0
0
0
0
2
0
0
0
0
0

0.76
0.76
0.76
0.76
0.76
0.76
0.76
0.76
0.76
0.76
0.76
0.76
0.76
0.76
0.76
0.76
0.76
0.76
0.76
0.76
0.76
0.76
0.76
0.76
0.76
0.76
0.76
0.76
0.76
0.76


7
1
1
0
0
0
0
1
1
1
1
1
0
0
0
0
0
0
0
0
0
0
0
0
0
0
0
0
0
0


7
1
1
0
0
0
0
1
1
1
1
1
0
0
0
0
0
0
0
0
0
0
0
0
0
0
0
0
0
0

1.0
1.0
1.0
1.0
1.0
1.0
1.0
1.0
1.0
1.0
1.0
1.0
1.0
1.0
1.0
1.0
1.0
1.0
1.0
1.0
1.0
1.0
1.0
1.0
1.0
1.0
1.0
1.0
1.0
1.0


7
1
2
0
0
0
1
0
0
0
0
2
0
0
0
0
0
0
0
0
0
0
0
0
1
0
0
0
0
0


7
1
2
0
0
0
1
0
0
0
0
2
0
0
0
0
0
0
0
0
0
0
0
0
1
0
0
0
0
0

1.0
1.0
1.0
1.0
1.0
1.0
1.0
1.0
1.0
1.0
1.0
1.0
1.0
1.0
1.0
1.0
1.0
1.0
1.0
1.0
1.0
1.0
1.0
1.0
1.0
1.0
1.0
1.0
1.0
1.0


9
1
1
0
1
1
1
1
1
1
0
0
0
0
0
0
0
0
0
0
0
0
0
0
1
0
0
0
0
0


9
1
1
0
1
1
1
1
1
1
0
0
0
0
0
0
0
0
0
0
0
0
0
0
1
0
0
0
0
0

1.0
1.0
1.0
1.0
1.0
1.0
1.0
1.0
1.0
1.0
1.0
1.0
1.0
1.0
1.0
1.0
1.0
1.0
1.0
1.0
1.0
1.0
1.0
1.0
1.0
1.0
1.0
1.0
1.0
1.0


1
0
0
0
0
0
1
0
0
0
0
0
0
0
0
0
0
0
0
0
0
0
0
0
0
0
0
0
0
0


1
0
0
0
0
0
1
0
0
0
0
0
0
0
0
0
0
0
0
0
0
0
0
0
0
0
0
0
0
0

1.0
1.0
1.0
1.0
1.0
1.0
1.0
1.0
1.0
1.0
1.0
1.0
1.0
1.0
1.0
1.0
1.0
1.0
1.0
1.0
1.0
1.0
1.0
1.0
1.0
1.0
1.0
1.0
1.0
1.0


1
1
0
0
0
0
0
0
0
0
0
0
0
0
0
0
0
0
0
0
0
0
0
0
0
0
0
0
0
0

1.0
1.0
1.0
1.0
1.0
1.0
1.0
1.0
1.0
1.0
1.0
1.0
1.0
1.0
1.0
1.0
1.0
1.0
1.0
1.0
1.0
1.0
1.0
1.0
1.0
1.0
1.0
1.0
1.0
1.0


2
2
0
0
0
0
0
0
0
0
0
0
0
0
0
0
0
0
0
0
0
0
0
0
0
0
0
0
0
0


2
2
0
0
0
0
0
0
0
0
0
0
0
0
0
0
0
0
0
0
0
0
0
0
0
0
0
0
0
0

1.0
1.0
1.0
1.0
1.0
1.0
1.0
1.0
1.0
1.0
1.0
1.0
1.0
1.0
1.0
1.0
1.0
1.0
1.0
1.0
1.0
1.0
1.0
1.0
1.0
1.0
1.0
1.0
1.0
1.0


2
0
0
0
0
0
0
0
0
1
1
0
0
0
0
0
0
0
0
0
0
0
0
0
0
0
0
0
0
0


2
0
0
0
0
0
0
0
0
1
1
0
0
0
0
0
0
0
0
0
0
0
0
0
0
0
0
0
0
0

0.9
0.9
0.9
0.9
0.9
0.9
0.9
0.9
0.9
0.9
0.9
0.9
0.9
0.9
0.9
0.9
0.9
0.9
0.9
0.9
0.9
0.9
0.9
0.9
0.9
0.9
0.9
0.9
0.9
0.9


2
1
0
0
0
0
0
0
0
1
0
0
0
0
0
0
0
0
0
0
0
0
0
0
0
0
0
0
0
0


2
1
0
0
0
0
0
0
0
1
0
0
0
0
0
0
0
0
0
0
0
0
0
0
0
0
0
0
0
0

0.98
0.98
0.98
0.98
0.98
0.98
0.98
0.98
0.98
0.98
0.98
0.98
0.98
0.98
0.98
0.98
0.98
0.98
0.98
0.98
0.98
0.98
0.98
0.98
0.98
0.98
0.98
0.98
0.98
0.98


2
0
0
0
0
0
0
0
2
0
0
0
0
0
0
0
0
0
0
0
0
0
0
0
0
0
0
0
0
0

1.0
1.0
1.0
1.0
1.0
1.0
1.0
1.0
1.0
1.0
1.0
1.0
1.0
1.0
1.0
1.0
1.0
1.0
1.0
1.0
1.0
1.0
1.0
1.0
1.0
1.0
1.0
1.0
1.0
1.0


1
0
0
1
0
0
0
0
0
0
0
0
0
0
0
0
0
0
0
0
0
0
0
0
0
0
0
0
0
0


1
0
0
1
0
0
0
0
0
0
0
0
0
0
0
0
0
0
0
0
0
0
0
0
0
0
0
0
0
0

0.85
0.85
0.85
0.85
0.85
0.85
0.85
0.85
0.85
0.85
0.85
0.85
0.85
0.85
0.85
0.85
0.85
0.85
0.85
0.85
0.85
0.85
0.85
0.85
0.85
0.85
0.85
0.85
0.85
0.85


2
0
0
0
0
0
0
0
0
0
0
0
0
0
0
0
0
0
0
2
0
0
0
0
0
0
0
0
0
0


2
0
0
0
0
0
0
0
0
0
0
0
0
0
0
0
0
0
0
2
0
0
0
0
0
0
0
0
0
0

0.97
0.97
0.97
0.97
0.97
0.97
0.97
0.97
0.97
0.97
0.97
0.97
0.97
0.97
0.97
0.97
0.97
0.97
0.97
0.97
0.97
0.97
0.97
0.97
0.97
0.97
0.97
0.97
0.97
0.97


18
2
3
4
0
0
0
0
0
0
1
2
0
1
0
0
0
0
0
0
0
0
0
0
5
0
0
0
0
0

0.81
0.81
0.81
0.81
0.81
0.81
0.81
0.81
0.81
0.81
0.81
0.81
0.81
0.81
0.81
0.81
0.81
0.81
0.81
0.81
0.81
0.81
0.81
0.81
0.81
0.81
0.81
0.81
0.81
0.81


5
2
2
0
0
0
0
0
0
0
0
1
0
0
0
0
0
0
0
0
0
0
0
0
0
0
0
0
0
0

1.0
1.0
1.0
1.0
1.0
1.0
1.0
1.0
1.0
1.0
1.0
1.0
1.0
1.0
1.0
1.0
1.0
1.0
1.0
1.0
1.0
1.0
1.0
1.0
1.0
1.0
1.0
1.0
1.0
1.0


2
1
1
0
0
0
0
0
0
0
0
0
0
0
0
0
0
0
0
0
0
0
0
0
0
0
0
0
0
0

1.0
1.0
1.0
1.0
1.0
1.0
1.0
1.0
1.0
1.0
1.0
1.0
1.0
1.0
1.0
1.0
1.0
1.0
1.0
1.0
1.0
1.0
1.0
1.0
1.0
1.0
1.0
1.0
1.0
1.0


1
0
0
0
0
0
0
0
0
0
1
0
0
0
0
0
0
0
0
0
0
0
0
0
0
0
0
0
0
0


1
0
0
0
0
0
0
0
0
0
1
0
0
0
0
0
0
0
0
0
0
0
0
0
0
0
0
0
0
0

1.0
1.0
1.0
1.0
1.0
1.0
1.0
1.0
1.0
1.0
1.0
1.0
1.0
1.0
1.0
1.0
1.0
1.0
1.0
1.0
1.0
1.0
1.0
1.0
1.0
1.0
1.0
1.0
1.0
1.0


3
0
0
1
0
0
0
0
0
0
0
0
0
0
0
0
0
0
0
0
0
0
0
0
2
0
0
0
0
0


3
0
0
1
0
0
0
0
0
0
0
0
0
0
0
0
0
0
0
0
0
0
0
0
2
0
0
0
0
0

1.0
1.0
1.0
1.0
1.0
1.0
1.0
1.0
1.0
1.0
1.0
1.0
1.0
1.0
1.0
1.0
1.0
1.0
1.0
1.0
1.0
1.0
1.0
1.0
1.0
1.0
1.0
1.0
1.0
1.0


3
0
1
0
0
0
0
0
0
0
0
1
0
1
0
0
0
0
0
0
0
0
0
0
0
0
0
0
0
0


3
0
1
0
0
0
0
0
0
0
0
1
0
1
0
0
0
0
0
0
0
0
0
0
0
0
0
0
0
0

1.0
1.0
1.0
1.0
1.0
1.0
1.0
1.0
1.0
1.0
1.0
1.0
1.0
1.0
1.0
1.0
1.0
1.0
1.0
1.0
1.0
1.0
1.0
1.0
1.0
1.0
1.0
1.0
1.0
1.0


4
0
0
2
0
0
0
0
0
0
0
0
0
0
0
0
0
0
0
0
0
0
0
0
2
0
0
0
0
0


4
0
0
2
0
0
0
0
0
0
0
0
0
0
0
0
0
0
0
0
0
0
0
0
2
0
0
0
0
0

1.0
1.0
1.0
1.0
1.0
1.0
1.0
1.0
1.0
1.0
1.0
1.0
1.0
1.0
1.0
1.0
1.0
1.0
1.0
1.0
1.0
1.0
1.0
1.0
1.0
1.0
1.0
1.0
1.0
1.0


4
2
1
0
0
0
0
0
0
0
0
0
0
0
0
0
1
0
0
0
0
0
0
0
0
0
0
0
0
0


2
2
0
0
0
0
0
0
0
0
0
0
0
0
0
0
0
0
0
0
0
0
0
0
0
0
0
0
0
0


2
2
0
0
0
0
0
0
0
0
0
0
0
0
0
0
0
0
0
0
0
0
0
0
0
0
0
0
0
0

1.0
1.0
1.0
1.0
1.0
1.0
1.0
1.0
1.0
1.0
1.0
1.0
1.0
1.0
1.0
1.0
1.0
1.0
1.0
1.0
1.0
1.0
1.0
1.0
1.0
1.0
1.0
1.0
1.0
1.0


1
0
1
0
0
0
0
0
0
0
0
0
0
0
0
0
0
0
0
0
0
0
0
0
0
0
0
0
0
0


1
0
1
0
0
0
0
0
0
0
0
0
0
0
0
0
0
0
0
0
0
0
0
0
0
0
0
0
0
0

1.0
1.0
1.0
1.0
1.0
1.0
1.0
1.0
1.0
1.0
1.0
1.0
1.0
1.0
1.0
1.0
1.0
1.0
1.0
1.0
1.0
1.0
1.0
1.0
1.0
1.0
1.0
1.0
1.0
1.0


1
0
0
0
0
0
0
0
0
0
0
0
0
0
0
0
1
0
0
0
0
0
0
0
0
0
0
0
0
0


1
0
0
0
0
0
0
0
0
0
0
0
0
0
0
0
1
0
0
0
0
0
0
0
0
0
0
0
0
0

1.0
1.0
1.0
1.0
1.0
1.0
1.0
1.0
1.0
1.0
1.0
1.0
1.0
1.0
1.0
1.0
1.0
1.0
1.0
1.0
1.0
1.0
1.0
1.0
1.0
1.0
1.0
1.0
1.0
1.0


49
3
4
3
3
0
0
2
1
3
2
0
0
0
1
0
0
17
0
3
3
0
0
0
4
0
0
0
0
0

1.0
1.0
1.0
1.0
1.0
1.0
1.0
1.0
1.0
1.0
1.0
1.0
1.0
1.0
1.0
1.0
1.0
1.0
1.0
1.0
1.0
1.0
1.0
1.0
1.0
1.0
1.0
1.0
1.0
1.0


2
0
0
0
0
0
0
0
1
1
0
0
0
0
0
0
0
0
0
0
0
0
0
0
0
0
0
0
0
0

0.77
0.77
0.77
0.77
0.77
0.77
0.77
0.77
0.77
0.77
0.77
0.77
0.77
0.77
0.77
0.77
0.77
0.77
0.77
0.77
0.77
0.77
0.77
0.77
0.77
0.77
0.77
0.77
0.77
0.77


11
1
2
2
1
0
0
2
0
1
2
0
0
0
0
0
0
0
0
0
0
0
0
0
0
0
0
0
0
0


11
1
2
2
1
0
0
2
0
1
2
0
0
0
0
0
0
0
0
0
0
0
0
0
0
0
0
0
0
0

1.0
1.0
1.0
1.0
1.0
1.0
1.0
1.0
1.0
1.0
1.0
1.0
1.0
1.0
1.0
1.0
1.0
1.0
1.0
1.0
1.0
1.0
1.0
1.0
1.0
1.0
1.0
1.0
1.0
1.0


2
1
0
0
0
0
0
0
0
0
0
0
0
0
0
0
0
1
0
0
0
0
0
0
0
0
0
0
0
0

0.75
0.75
0.75
0.75
0.75
0.75
0.75
0.75
0.75
0.75
0.75
0.75
0.75
0.75
0.75
0.75
0.75
0.75
0.75
0.75
0.75
0.75
0.75
0.75
0.75
0.75
0.75
0.75
0.75
0.75


1
1
0
0
0
0
0
0
0
0
0
0
0
0
0
0
0
0
0
0
0
0
0
0
0
0
0
0
0
0

0.99
0.99
0.99
0.99
0.99
0.99
0.99
0.99
0.99
0.99
0.99
0.99
0.99
0.99
0.99
0.99
0.99
0.99
0.99
0.99
0.99
0.99
0.99
0.99
0.99
0.99
0.99
0.99
0.99
0.99


2
0
0
0
2
0
0
0
0
0
0
0
0
0
0
0
0
0
0
0
0
0
0
0
0
0
0
0
0
0

1.0
1.0
1.0
1.0
1.0
1.0
1.0
1.0
1.0
1.0
1.0
1.0
1.0
1.0
1.0
1.0
1.0
1.0
1.0
1.0
1.0
1.0
1.0
1.0
1.0
1.0
1.0
1.0
1.0
1.0


5
0
0
1
0
0
0
0
0
0
0
0
0
0
0
0
0
0
0
0
0
0
0
0
4
0
0
0
0
0

1.0
1.0
1.0
1.0
1.0
1.0
1.0
1.0
1.0
1.0
1.0
1.0
1.0
1.0
1.0
1.0
1.0
1.0
1.0
1.0
1.0
1.0
1.0
1.0
1.0
1.0
1.0
1.0
1.0
1.0


2
0
1
0
0
0
0
0
0
1
0
0
0
0
0
0
0
0
0
0
0
0
0
0
0
0
0
0
0
0


2
0
1
0
0
0
0
0
0
1
0
0
0
0
0
0
0
0
0
0
0
0
0
0
0
0
0
0
0
0

0.98
0.98
0.98
0.98
0.98
0.98
0.98
0.98
0.98
0.98
0.98
0.98
0.98
0.98
0.98
0.98
0.98
0.98
0.98
0.98
0.98
0.98
0.98
0.98
0.98
0.98
0.98
0.98
0.98
0.98


1
1
0
0
0
0
0
0
0
0
0
0
0
0
0
0
0
0
0
0
0
0
0
0
0
0
0
0
0
0

0.71
0.71
0.71
0.71
0.71
0.71
0.71
0.71
0.71
0.71
0.71
0.71
0.71
0.71
0.71
0.71
0.71
0.71
0.71
0.71
0.71
0.71
0.71
0.71
0.71
0.71
0.71
0.71
0.71
0.71


4
0
0
0
0
0
0
0
0
0
0
0
0
0
1
0
0
0
0
0
3
0
0
0
0
0
0
0
0
0


4
0
0
0
0
0
0
0
0
0
0
0
0
0
1
0
0
0
0
0
3
0
0
0
0
0
0
0
0
0

1.0
1.0
1.0
1.0
1.0
1.0
1.0
1.0
1.0
1.0
1.0
1.0
1.0
1.0
1.0
1.0
1.0
1.0
1.0
1.0
1.0
1.0
1.0
1.0
1.0
1.0
1.0
1.0
1.0
1.0


9
0
0
0
0
0
0
0
0
0
0
0
0
0
0
0
0
9
0
0
0
0
0
0
0
0
0
0
0
0


9
0
0
0
0
0
0
0
0
0
0
0
0
0
0
0
0
9
0
0
0
0
0
0
0
0
0
0
0
0

1.0
1.0
1.0
1.0
1.0
1.0
1.0
1.0
1.0
1.0
1.0
1.0
1.0
1.0
1.0
1.0
1.0
1.0
1.0
1.0
1.0
1.0
1.0
1.0
1.0
1.0
1.0
1.0
1.0
1.0


3
0
0
0
0
0
0
0
0
0
0
0
0
0
0
0
0
0
0
3
0
0
0
0
0
0
0
0
0
0


3
0
0
0
0
0
0
0
0
0
0
0
0
0
0
0
0
0
0
3
0
0
0
0
0
0
0
0
0
0

1.0
1.0
1.0
1.0
1.0
1.0
1.0
1.0
1.0
1.0
1.0
1.0
1.0
1.0
1.0
1.0
1.0
1.0
1.0
1.0
1.0
1.0
1.0
1.0
1.0
1.0
1.0
1.0
1.0
1.0


45
3
4
1
0
5
3
0
0
2
1
1
0
0
0
0
17
0
0
0
0
0
0
0
8
0
0
0
0
0

1.0
1.0
1.0
1.0
1.0
1.0
1.0
1.0
1.0
1.0
1.0
1.0
1.0
1.0
1.0
1.0
1.0
1.0
1.0
1.0
1.0
1.0
1.0
1.0
1.0
1.0
1.0
1.0
1.0
1.0


17
1
0
1
0
4
1
0
0
1
0
1
0
0
0
0
0
0
0
0
0
0
0
0
8
0
0
0
0
0


1
0
0
0
0
0
1
0
0
0
0
0
0
0
0
0
0
0
0
0
0
0
0
0
0
0
0
0
0
0

1.0
1.0
1.0
1.0
1.0
1.0
1.0
1.0
1.0
1.0
1.0
1.0
1.0
1.0
1.0
1.0
1.0
1.0
1.0
1.0
1.0
1.0
1.0
1.0
1.0
1.0
1.0
1.0
1.0
1.0


5
0
0
0
0
4
0
0
0
0
0
0
0
0
0
0
0
0
0
0
0
0
0
0
1
0
0
0
0
0

1.0
1.0
1.0
1.0
1.0
1.0
1.0
1.0
1.0
1.0
1.0
1.0
1.0
1.0
1.0
1.0
1.0
1.0
1.0
1.0
1.0
1.0
1.0
1.0
1.0
1.0
1.0
1.0
1.0
1.0


11
1
0
1
0
0
0
0
0
1
0
1
0
0
0
0
0
0
0
0
0
0
0
0
7
0
0
0
0
0

1.0
1.0
1.0
1.0
1.0
1.0
1.0
1.0
1.0
1.0
1.0
1.0
1.0
1.0
1.0
1.0
1.0
1.0
1.0
1.0
1.0
1.0
1.0
1.0
1.0
1.0
1.0
1.0
1.0
1.0


2
0
1
0
0
0
1
0
0
0
0
0
0
0
0
0
0
0
0
0
0
0
0
0
0
0
0
0
0
0


2
0
1
0
0
0
1
0
0
0
0
0
0
0
0
0
0
0
0
0
0
0
0
0
0
0
0
0
0
0

1.0
1.0
1.0
1.0
1.0
1.0
1.0
1.0
1.0
1.0
1.0
1.0
1.0
1.0
1.0
1.0
1.0
1.0
1.0
1.0
1.0
1.0
1.0
1.0
1.0
1.0
1.0
1.0
1.0
1.0


4
2
1
0
0
0
0
0
0
0
1
0
0
0
0
0
0
0
0
0
0
0
0
0
0
0
0
0
0
0


4
2
1
0
0
0
0
0
0
0
1
0
0
0
0
0
0
0
0
0
0
0
0
0
0
0
0
0
0
0

0.98
0.98
0.98
0.98
0.98
0.98
0.98
0.98
0.98
0.98
0.98
0.98
0.98
0.98
0.98
0.98
0.98
0.98
0.98
0.98
0.98
0.98
0.98
0.98
0.98
0.98
0.98
0.98
0.98
0.98


17
0
0
0
0
0
1
0
0
0
0
0
0
0
0
0
16
0
0
0
0
0
0
0
0
0
0
0
0
0

1.0
1.0
1.0
1.0
1.0
1.0
1.0
1.0
1.0
1.0
1.0
1.0
1.0
1.0
1.0
1.0
1.0
1.0
1.0
1.0
1.0
1.0
1.0
1.0
1.0
1.0
1.0
1.0
1.0
1.0


16
0
0
0
0
0
1
0
0
0
0
0
0
0
0
0
15
0
0
0
0
0
0
0
0
0
0
0
0
0

1.0
1.0
1.0
1.0
1.0
1.0
1.0
1.0
1.0
1.0
1.0
1.0
1.0
1.0
1.0
1.0
1.0
1.0
1.0
1.0
1.0
1.0
1.0
1.0
1.0
1.0
1.0
1.0
1.0
1.0


1
0
0
0
0
1
0
0
0
0
0
0
0
0
0
0
0
0
0
0
0
0
0
0
0
0
0
0
0
0


1
0
0
0
0
1
0
0
0
0
0
0
0
0
0
0
0
0
0
0
0
0
0
0
0
0
0
0
0
0

1.0
1.0
1.0
1.0
1.0
1.0
1.0
1.0
1.0
1.0
1.0
1.0
1.0
1.0
1.0
1.0
1.0
1.0
1.0
1.0
1.0
1.0
1.0
1.0
1.0
1.0
1.0
1.0
1.0
1.0


1
0
1
0
0
0
0
0
0
0
0
0
0
0
0
0
0
0
0
0
0
0
0
0
0
0
0
0
0
0


1
0
1
0
0
0
0
0
0
0
0
0
0
0
0
0
0
0
0
0
0
0
0
0
0
0
0
0
0
0

1.0
1.0
1.0
1.0
1.0
1.0
1.0
1.0
1.0
1.0
1.0
1.0
1.0
1.0
1.0
1.0
1.0
1.0
1.0
1.0
1.0
1.0
1.0
1.0
1.0
1.0
1.0
1.0
1.0
1.0


2
0
1
0
0
0
0
0
0
1
0
0
0
0
0
0
0
0
0
0
0
0
0
0
0
0
0
0
0
0

0.99
0.99
0.99
0.99
0.99
0.99
0.99
0.99
0.99
0.99
0.99
0.99
0.99
0.99
0.99
0.99
0.99
0.99
0.99
0.99
0.99
0.99
0.99
0.99
0.99
0.99
0.99
0.99
0.99
0.99


1
0
1
0
0
0
0
0
0
0
0
0
0
0
0
0
0
0
0
0
0
0
0
0
0
0
0
0
0
0

1.0
1.0
1.0
1.0
1.0
1.0
1.0
1.0
1.0
1.0
1.0
1.0
1.0
1.0
1.0
1.0
1.0
1.0
1.0
1.0
1.0
1.0
1.0
1.0
1.0
1.0
1.0
1.0
1.0
1.0


9
1
1
1
0
0
1
1
1
1
0
0
0
1
0
0
0
0
0
0
0
0
0
0
1
0
0
0
0
0

0.81
0.81
0.81
0.81
0.81
0.81
0.81
0.81
0.81
0.81
0.81
0.81
0.81
0.81
0.81
0.81
0.81
0.81
0.81
0.81
0.81
0.81
0.81
0.81
0.81
0.81
0.81
0.81
0.81
0.81


5
1
0
0
0
0
1
1
1
1
0
0
0
0
0
0
0
0
0
0
0
0
0
0
0
0
0
0
0
0


5
1
0
0
0
0
1
1
1
1
0
0
0
0
0
0
0
0
0
0
0
0
0
0
0
0
0
0
0
0

0.97
0.97
0.97
0.97
0.97
0.97
0.97
0.97
0.97
0.97
0.97
0.97
0.97
0.97
0.97
0.97
0.97
0.97
0.97
0.97
0.97
0.97
0.97
0.97
0.97
0.97
0.97
0.97
0.97
0.97


2
0
1
1
0
0
0
0
0
0
0
0
0
0
0
0
0
0
0
0
0
0
0
0
0
0
0
0
0
0


2
0
1
1
0
0
0
0
0
0
0
0
0
0
0
0
0
0
0
0
0
0
0
0
0
0
0
0
0
0

1.0
1.0
1.0
1.0
1.0
1.0
1.0
1.0
1.0
1.0
1.0
1.0
1.0
1.0
1.0
1.0
1.0
1.0
1.0
1.0
1.0
1.0
1.0
1.0
1.0
1.0
1.0
1.0
1.0
1.0


13
1
1
1
1
2
1
0
0
0
0
1
0
0
0
0
0
0
0
0
0
0
0
0
2
1
2
0
0
0


13
1
1
1
1
2
1
0
0
0
0
1
0
0
0
0
0
0
0
0
0
0
0
0
2
1
2
0
0
0


13
1
1
1
1
2
1
0
0
0
0
1
0
0
0
0
0
0
0
0
0
0
0
0
2
1
2
0
0
0

1.0
1.0
1.0
1.0
1.0
1.0
1.0
1.0
1.0
1.0
1.0
1.0
1.0
1.0
1.0
1.0
1.0
1.0
1.0
1.0
1.0
1.0
1.0
1.0
1.0
1.0
1.0
1.0
1.0
1.0


5
0
0
0
0
0
0
1
0
1
0
1
0
0
0
0
0
0
2
0
0
0
0
0
0
0
0
0
0
0


1
0
0
0
0
0
0
1
0
0
0
0
0
0
0
0
0
0
0
0
0
0
0
0
0
0
0
0
0
0

1.0
1.0
1.0
1.0
1.0
1.0
1.0
1.0
1.0
1.0
1.0
1.0
1.0
1.0
1.0
1.0
1.0
1.0
1.0
1.0
1.0
1.0
1.0
1.0
1.0
1.0
1.0
1.0
1.0
1.0


1
0
0
0
0
0
0
0
0
0
0
1
0
0
0
0
0
0
0
0
0
0
0
0
0
0
0
0
0
0


1
0
0
0
0
0
0
0
0
0
0
1
0
0
0
0
0
0
0
0
0
0
0
0
0
0
0
0
0
0

1.0
1.0
1.0
1.0
1.0
1.0
1.0
1.0
1.0
1.0
1.0
1.0
1.0
1.0
1.0
1.0
1.0
1.0
1.0
1.0
1.0
1.0
1.0
1.0
1.0
1.0
1.0
1.0
1.0
1.0


1
0
0
0
0
0
0
0
0
1
0
0
0
0
0
0
0
0
0
0
0
0
0
0
0
0
0
0
0
0

0.83
0.83
0.83
0.83
0.83
0.83
0.83
0.83
0.83
0.83
0.83
0.83
0.83
0.83
0.83
0.83
0.83
0.83
0.83
0.83
0.83
0.83
0.83
0.83
0.83
0.83
0.83
0.83
0.83
0.83


2
0
0
0
0
0
0
0
0
0
0
0
0
0
0
0
0
0
2
0
0
0
0
0
0
0
0
0
0
0


2
0
0
0
0
0
0
0
0
0
0
0
0
0
0
0
0
0
2
0
0
0
0
0
0
0
0
0
0
0

1.0
1.0
1.0
1.0
1.0
1.0
1.0
1.0
1.0
1.0
1.0
1.0
1.0
1.0
1.0
1.0
1.0
1.0
1.0
1.0
1.0
1.0
1.0
1.0
1.0
1.0
1.0
1.0
1.0
1.0


15
3
1
0
0
0
0
2
1
4
3
1
0
0
0
0
0
0
0
0
0
0
0
0
0
0
0
0
0
0


13
3
1
0
0
0
0
2
1
3
2
1
0
0
0
0
0
0
0
0
0
0
0
0
0
0
0
0
0
0


13
3
1
0
0
0
0
2
1
3
2
1
0
0
0
0
0
0
0
0
0
0
0
0
0
0
0
0
0
0

1.0
1.0
1.0
1.0
1.0
1.0
1.0
1.0
1.0
1.0
1.0
1.0
1.0
1.0
1.0
1.0
1.0
1.0
1.0
1.0
1.0
1.0
1.0
1.0
1.0
1.0
1.0
1.0
1.0
1.0


2
0
0
0
0
0
0
0
0
1
1
0
0
0
0
0
0
0
0
0
0
0
0
0
0
0
0
0
0
0

0.86
0.86
0.86
0.86
0.86
0.86
0.86
0.86
0.86
0.86
0.86
0.86
0.86
0.86
0.86
0.86
0.86
0.86
0.86
0.86
0.86
0.86
0.86
0.86
0.86
0.86
0.86
0.86
0.86
0.86


4
0
0
0
0
0
2
0
0
0
0
0
0
0
0
0
0
0
0
0
0
0
0
0
0
0
0
2
0
0


4
0
0
0
0
0
2
0
0
0
0
0
0
0
0
0
0
0
0
0
0
0
0
0
0
0
0
2
0
0


4
0
0
0
0
0
2
0
0
0
0
0
0
0
0
0
0
0
0
0
0
0
0
0
0
0
0
2
0
0

1.0
1.0
1.0
1.0
1.0
1.0
1.0
1.0
1.0
1.0
1.0
1.0
1.0
1.0
1.0
1.0
1.0
1.0
1.0
1.0
1.0
1.0
1.0
1.0
1.0
1.0
1.0
1.0
1.0
1.0


9
1
2
3
0
1
0
0
0
0
0
0
0
0
0
0
0
0
0
0
0
0
0
0
2
0
0
0
0
0


8
1
2
2
0
1
0
0
0
0
0
0
0
0
0
0
0
0
0
0
0
0
0
0
2
0
0
0
0
0

1.0
1.0
1.0
1.0
1.0
1.0
1.0
1.0
1.0
1.0
1.0
1.0
1.0
1.0
1.0
1.0
1.0
1.0
1.0
1.0
1.0
1.0
1.0
1.0
1.0
1.0
1.0
1.0
1.0
1.0


6
1
2
2
0
0
0
0
0
0
0
0
0
0
0
0
0
0
0
0
0
0
0
0
1
0
0
0
0
0

1.0
1.0
1.0
1.0
1.0
1.0
1.0
1.0
1.0
1.0
1.0
1.0
1.0
1.0
1.0
1.0
1.0
1.0
1.0
1.0
1.0
1.0
1.0
1.0
1.0
1.0
1.0
1.0
1.0
1.0


1
0
0
0
0
1
0
0
0
0
0
0
0
0
0
0
0
0
0
0
0
0
0
0
0
0
0
0
0
0

0.82
0.82
0.82
0.82
0.82
0.82
0.82
0.82
0.82
0.82
0.82
0.82
0.82
0.82
0.82
0.82
0.82
0.82
0.82
0.82
0.82
0.82
0.82
0.82
0.82
0.82
0.82
0.82
0.82
0.82


1
0
0
1
0
0
0
0
0
0
0
0
0
0
0
0
0
0
0
0
0
0
0
0
0
0
0
0
0
0

1.0
1.0
1.0
1.0
1.0
1.0
1.0
1.0
1.0
1.0
1.0
1.0
1.0
1.0
1.0
1.0
1.0
1.0
1.0
1.0
1.0
1.0
1.0
1.0
1.0
1.0
1.0
1.0
1.0
1.0


11
1
2
1
1
1
1
0
0
0
0
2
0
0
0
0
0
0
0
0
2
0
0
0
0
0
0
0
0
0


3
1
1
0
0
0
0
0
0
0
0
1
0
0
0
0
0
0
0
0
0
0
0
0
0
0
0
0
0
0


3
1
1
0
0
0
0
0
0
0
0
1
0
0
0
0
0
0
0
0
0
0
0
0
0
0
0
0
0
0

0.92
0.92
0.92
0.92
0.92
0.92
0.92
0.92
0.92
0.92
0.92
0.92
0.92
0.92
0.92
0.92
0.92
0.92
0.92
0.92
0.92
0.92
0.92
0.92
0.92
0.92
0.92
0.92
0.92
0.92


5
0
1
1
1
1
1
0
0
0
0
0
0
0
0
0
0
0
0
0
0
0
0
0
0
0
0
0
0
0


5
0
1
1
1
1
1
0
0
0
0
0
0
0
0
0
0
0
0
0
0
0
0
0
0
0
0
0
0
0

0.89
0.89
0.89
0.89
0.89
0.89
0.89
0.89
0.89
0.89
0.89
0.89
0.89
0.89
0.89
0.89
0.89
0.89
0.89
0.89
0.89
0.89
0.89
0.89
0.89
0.89
0.89
0.89
0.89
0.89


1
0
0
0
0
0
0
0
0
0
0
1
0
0
0
0
0
0
0
0
0
0
0
0
0
0
0
0
0
0


1
0
0
0
0
0
0
0
0
0
0
1
0
0
0
0
0
0
0
0
0
0
0
0
0
0
0
0
0
0

1.0
1.0
1.0
1.0
1.0
1.0
1.0
1.0
1.0
1.0
1.0
1.0
1.0
1.0
1.0
1.0
1.0
1.0
1.0
1.0
1.0
1.0
1.0
1.0
1.0
1.0
1.0
1.0
1.0
1.0


2
0
0
0
0
0
0
0
0
0
0
0
0
0
0
0
0
0
0
0
2
0
0
0
0
0
0
0
0
0


2
0
0
0
0
0
0
0
0
0
0
0
0
0
0
0
0
0
0
0
2
0
0
0
0
0
0
0
0
0

1.0
1.0
1.0
1.0
1.0
1.0
1.0
1.0
1.0
1.0
1.0
1.0
1.0
1.0
1.0
1.0
1.0
1.0
1.0
1.0
1.0
1.0
1.0
1.0
1.0
1.0
1.0
1.0
1.0
1.0


4
1
1
1
0
0
0
0
0
0
0
0
0
0
0
0
0
0
0
0
0
0
0
0
0
0
1
0
0
0


4
1
1
1
0
0
0
0
0
0
0
0
0
0
0
0
0
0
0
0
0
0
0
0
0
0
1
0
0
0


4
1
1
1
0
0
0
0
0
0
0
0
0
0
0
0
0
0
0
0
0
0
0
0
0
0
1
0
0
0

1.0
1.0
1.0
1.0
1.0
1.0
1.0
1.0
1.0
1.0
1.0
1.0
1.0
1.0
1.0
1.0
1.0
1.0
1.0
1.0
1.0
1.0
1.0
1.0
1.0
1.0
1.0
1.0
1.0
1.0


7
0
0
0
0
0
0
0
0
0
0
0
0
0
0
0
0
0
3
0
0
0
0
0
0
0
0
0
0
4


7
0
0
0
0
0
0
0
0
0
0
0
0
0
0
0
0
0
3
0
0
0
0
0
0
0
0
0
0
4

1.0
1.0
1.0
1.0
1.0
1.0
1.0
1.0
1.0
1.0
1.0
1.0
1.0
1.0
1.0
1.0
1.0
1.0
1.0
1.0
1.0
1.0
1.0
1.0
1.0
1.0
1.0
1.0
1.0
1.0


26
0
0
0
0
0
0
0
0
0
0
0
0
0
0
0
0
0
0
0
0
0
0
0
14
0
10
0
0
2

0.97
0.97
0.97
0.97
0.97
0.97
0.97
0.97
0.97
0.97
0.97
0.97
0.97
0.97
0.97
0.97
0.97
0.97
0.97
0.97
0.97
0.97
0.97
0.97
0.97
0.97
0.97
0.97
0.97
0.97


19
0
0
0
0
0
0
0
0
0
0
0
0
0
0
0
0
0
0
0
0
0
0
0
11
0
7
0
0
1

1.0
1.0
1.0
1.0
1.0
1.0
1.0
1.0
1.0
1.0
1.0
1.0
1.0
1.0
1.0
1.0
1.0
1.0
1.0
1.0
1.0
1.0
1.0
1.0
1.0
1.0
1.0
1.0
1.0
1.0


1
0
0
0
0
0
0
0
0
0
0
0
0
0
0
0
0
0
0
0
0
0
0
0
1
0
0
0
0
0


1
0
0
0
0
0
0
0
0
0
0
0
0
0
0
0
0
0
0
0
0
0
0
0
1
0
0
0
0
0


1
0
0
0
0
0
0
0
0
0
0
0
0
0
0
0
0
0
0
0
0
0
0
0
1
0
0
0
0
0

1.0
1.0
1.0
1.0
1.0
1.0
1.0
1.0
1.0
1.0
1.0
1.0
1.0
1.0
1.0
1.0
1.0
1.0
1.0
1.0
1.0
1.0
1.0
1.0
1.0
1.0
1.0
1.0
1.0
1.0


1
0
0
0
0
0
0
0
0
0
0
0
0
0
0
0
0
0
0
0
0
0
0
0
0
0
0
0
1
0

0.88
0.88
0.88
0.88
0.88
0.88
0.88
0.88
0.88
0.88
0.88
0.88
0.88
0.88
0.88
0.88
0.88
0.88
0.88
0.88
0.88
0.88
0.88
0.88
0.88
0.88
0.88
0.88
0.88
0.88


2860
84
75
48
36
51
48
36
63
97
96
105
10
34
52
90
80
117
132
162
102
56
41
10
306
211
195
192
170
161

1.0
1.0
1.0
1.0
1.0
1.0
1.0
1.0
1.0
1.0
1.0
1.0
1.0
1.0
1.0
1.0
1.0
1.0
1.0
1.0
1.0
1.0
1.0
1.0
1.0
1.0
1.0
1.0
1.0
1.0


217
3
0
1
0
2
1
0
0
9
7
2
0
0
0
0
0
3
8
33
30
30
0
0
0
18
11
20
28
11

1.0
1.0
1.0
1.0
1.0
1.0
1.0
1.0
1.0
1.0
1.0
1.0
1.0
1.0
1.0
1.0
1.0
1.0
1.0
1.0
1.0
1.0
1.0
1.0
1.0
1.0
1.0
1.0
1.0
1.0


186
3
0
1
0
1
1
0
0
9
7
2
0
0
0
0
0
1
5
33
30
30
0
0
0
17
11
13
11
11

1.0
1.0
1.0
1.0
1.0
1.0
1.0
1.0
1.0
1.0
1.0
1.0
1.0
1.0
1.0
1.0
1.0
1.0
1.0
1.0
1.0
1.0
1.0
1.0
1.0
1.0
1.0
1.0
1.0
1.0


102
2
0
0
0
0
0
0
0
5
4
1
0
0
0
0
0
0
0
32
29
29
0
0
0
0
0
0
0
0

0.99
0.99
0.99
0.99
0.99
0.99
0.99
0.99
0.99
0.99
0.99
0.99
0.99
0.99
0.99
0.99
0.99
0.99
0.99
0.99
0.99
0.99
0.99
0.99
0.99
0.99
0.99
0.99
0.99
0.99


2
0
0
1
0
0
0
0
0
0
1
0
0
0
0
0
0
0
0
0
0
0
0
0
0
0
0
0
0
0

1.0
1.0
1.0
1.0
1.0
1.0
1.0
1.0
1.0
1.0
1.0
1.0
1.0
1.0
1.0
1.0
1.0
1.0
1.0
1.0
1.0
1.0
1.0
1.0
1.0
1.0
1.0
1.0
1.0
1.0


1
0
0
0
0
0
0
0
0
1
0
0
0
0
0
0
0
0
0
0
0
0
0
0
0
0
0
0
0
0

0.98
0.98
0.98
0.98
0.98
0.98
0.98
0.98
0.98
0.98
0.98
0.98
0.98
0.98
0.98
0.98
0.98
0.98
0.98
0.98
0.98
0.98
0.98
0.98
0.98
0.98
0.98
0.98
0.98
0.98


4
0
0
0
0
0
0
0
0
1
1
0
0
0
0
0
0
1
0
0
0
0
0
0
0
0
0
0
1
0

0.9
0.9
0.9
0.9
0.9
0.9
0.9
0.9
0.9
0.9
0.9
0.9
0.9
0.9
0.9
0.9
0.9
0.9
0.9
0.9
0.9
0.9
0.9
0.9
0.9
0.9
0.9
0.9
0.9
0.9


16
0
0
0
0
1
1
0
0
0
0
0
0
0
0
0
0
0
0
0
0
0
0
0
0
4
3
3
0
4

1.0
1.0
1.0
1.0
1.0
1.0
1.0
1.0
1.0
1.0
1.0
1.0
1.0
1.0
1.0
1.0
1.0
1.0
1.0
1.0
1.0
1.0
1.0
1.0
1.0
1.0
1.0
1.0
1.0
1.0


1
0
0
0
0
0
0
0
0
1
0
0
0
0
0
0
0
0
0
0
0
0
0
0
0
0
0
0
0
0

1.0
1.0
1.0
1.0
1.0
1.0
1.0
1.0
1.0
1.0
1.0
1.0
1.0
1.0
1.0
1.0
1.0
1.0
1.0
1.0
1.0
1.0
1.0
1.0
1.0
1.0
1.0
1.0
1.0
1.0


1
0
0
0
0
0
0
0
0
0
0
0
0
0
0
0
0
0
0
0
0
0
0
0
0
0
0
0
1
0

1.0
1.0
1.0
1.0
1.0
1.0
1.0
1.0
1.0
1.0
1.0
1.0
1.0
1.0
1.0
1.0
1.0
1.0
1.0
1.0
1.0
1.0
1.0
1.0
1.0
1.0
1.0
1.0
1.0
1.0


2
0
0
0
0
0
0
0
0
0
0
0
0
0
0
0
0
0
0
0
0
0
0
0
0
1
1
0
0
0

0.95
0.95
0.95
0.95
0.95
0.95
0.95
0.95
0.95
0.95
0.95
0.95
0.95
0.95
0.95
0.95
0.95
0.95
0.95
0.95
0.95
0.95
0.95
0.95
0.95
0.95
0.95
0.95
0.95
0.95


4
0
0
0
0
0
0
0
0
0
0
0
0
0
0
0
0
0
4
0
0
0
0
0
0
0
0
0
0
0

1.0
1.0
1.0
1.0
1.0
1.0
1.0
1.0
1.0
1.0
1.0
1.0
1.0
1.0
1.0
1.0
1.0
1.0
1.0
1.0
1.0
1.0
1.0
1.0
1.0
1.0
1.0
1.0
1.0
1.0


1
0
0
0
0
0
0
0
0
0
0
0
0
0
0
0
0
0
0
0
0
0
0
0
0
1
0
0
0
0

0.88
0.88
0.88
0.88
0.88
0.88
0.88
0.88
0.88
0.88
0.88
0.88
0.88
0.88
0.88
0.88
0.88
0.88
0.88
0.88
0.88
0.88
0.88
0.88
0.88
0.88
0.88
0.88
0.88
0.88


6
0
0
0
0
0
0
0
0
0
0
0
0
0
0
0
0
0
0
0
0
0
0
0
0
0
0
0
6
0

1.0
1.0
1.0
1.0
1.0
1.0
1.0
1.0
1.0
1.0
1.0
1.0
1.0
1.0
1.0
1.0
1.0
1.0
1.0
1.0
1.0
1.0
1.0
1.0
1.0
1.0
1.0
1.0
1.0
1.0


27
0
0
0
0
1
0
0
0
0
0
0
0
0
0
0
0
2
0
0
0
0
0
0
0
0
0
7
17
0

1.0
1.0
1.0
1.0
1.0
1.0
1.0
1.0
1.0
1.0
1.0
1.0
1.0
1.0
1.0
1.0
1.0
1.0
1.0
1.0
1.0
1.0
1.0
1.0
1.0
1.0
1.0
1.0
1.0
1.0


14
0
0
0
0
1
0
0
0
0
0
0
0
0
0
0
0
0
0
0
0
0
0
0
0
0
0
6
7
0

1.0
1.0
1.0
1.0
1.0
1.0
1.0
1.0
1.0
1.0
1.0
1.0
1.0
1.0
1.0
1.0
1.0
1.0
1.0
1.0
1.0
1.0
1.0
1.0
1.0
1.0
1.0
1.0
1.0
1.0


1
0
0
0
0
0
0
0
0
0
0
0
0
0
0
0
0
1
0
0
0
0
0
0
0
0
0
0
0
0

1.0
1.0
1.0
1.0
1.0
1.0
1.0
1.0
1.0
1.0
1.0
1.0
1.0
1.0
1.0
1.0
1.0
1.0
1.0
1.0
1.0
1.0
1.0
1.0
1.0
1.0
1.0
1.0
1.0
1.0


2
0
0
0
0
0
0
0
0
0
0
0
0
0
0
0
0
0
1
0
0
0
0
0
0
1
0
0
0
0


1
0
0
0
0
0
0
0
0
0
0
0
0
0
0
0
0
0
1
0
0
0
0
0
0
0
0
0
0
0

1.0
1.0
1.0
1.0
1.0
1.0
1.0
1.0
1.0
1.0
1.0
1.0
1.0
1.0
1.0
1.0
1.0
1.0
1.0
1.0
1.0
1.0
1.0
1.0
1.0
1.0
1.0
1.0
1.0
1.0


1
0
0
0
0
0
0
0
0
0
0
0
0
0
0
0
0
0
0
0
0
0
0
0
0
1
0
0
0
0

1.0
1.0
1.0
1.0
1.0
1.0
1.0
1.0
1.0
1.0
1.0
1.0
1.0
1.0
1.0
1.0
1.0
1.0
1.0
1.0
1.0
1.0
1.0
1.0
1.0
1.0
1.0
1.0
1.0
1.0


37
2
1
0
0
1
1
0
1
2
3
1
0
0
0
3
3
1
0
0
1
0
0
0
1
5
3
3
1
4


19
2
0
0
0
1
1
0
0
1
2
1
0
0
0
2
2
1
0
0
1
0
0
0
0
1
1
1
1
1


19
2
0
0
0
1
1
0
0
1
2
1
0
0
0
2
2
1
0
0
1
0
0
0
0
1
1
1
1
1

1.0
1.0
1.0
1.0
1.0
1.0
1.0
1.0
1.0
1.0
1.0
1.0
1.0
1.0
1.0
1.0
1.0
1.0
1.0
1.0
1.0
1.0
1.0
1.0
1.0
1.0
1.0
1.0
1.0
1.0


3
0
0
0
0
0
0
0
1
1
1
0
0
0
0
0
0
0
0
0
0
0
0
0
0
0
0
0
0
0


2
0
0
0
0
0
0
0
0
1
1
0
0
0
0
0
0
0
0
0
0
0
0
0
0
0
0
0
0
0

0.95
0.95
0.95
0.95
0.95
0.95
0.95
0.95
0.95
0.95
0.95
0.95
0.95
0.95
0.95
0.95
0.95
0.95
0.95
0.95
0.95
0.95
0.95
0.95
0.95
0.95
0.95
0.95
0.95
0.95


1
0
0
0
0
0
0
0
1
0
0
0
0
0
0
0
0
0
0
0
0
0
0
0
0
0
0
0
0
0

1.0
1.0
1.0
1.0
1.0
1.0
1.0
1.0
1.0
1.0
1.0
1.0
1.0
1.0
1.0
1.0
1.0
1.0
1.0
1.0
1.0
1.0
1.0
1.0
1.0
1.0
1.0
1.0
1.0
1.0


10
0
1
0
0
0
0
0
0
0
0
0
0
0
0
0
0
0
0
0
0
0
0
0
0
3
1
2
0
3


4
0
1
0
0
0
0
0
0
0
0
0
0
0
0
0
0
0
0
0
0
0
0
0
0
1
0
1
0
1

1.0
1.0
1.0
1.0
1.0
1.0
1.0
1.0
1.0
1.0
1.0
1.0
1.0
1.0
1.0
1.0
1.0
1.0
1.0
1.0
1.0
1.0
1.0
1.0
1.0
1.0
1.0
1.0
1.0
1.0


3
0
0
0
0
0
0
0
0
0
0
0
0
0
0
0
0
0
0
0
0
0
0
0
0
1
1
0
0
1

0.98
0.98
0.98
0.98
0.98
0.98
0.98
0.98
0.98
0.98
0.98
0.98
0.98
0.98
0.98
0.98
0.98
0.98
0.98
0.98
0.98
0.98
0.98
0.98
0.98
0.98
0.98
0.98
0.98
0.98


3
0
0
0
0
0
0
0
0
0
0
0
0
0
0
0
0
0
0
0
0
0
0
0
0
1
0
1
0
1

0.99
0.99
0.99
0.99
0.99
0.99
0.99
0.99
0.99
0.99
0.99
0.99
0.99
0.99
0.99
0.99
0.99
0.99
0.99
0.99
0.99
0.99
0.99
0.99
0.99
0.99
0.99
0.99
0.99
0.99


3
0
0
0
0
0
0
0
0
0
0
0
0
0
0
1
1
0
0
0
0
0
0
0
0
0
1
0
0
0


3
0
0
0
0
0
0
0
0
0
0
0
0
0
0
1
1
0
0
0
0
0
0
0
0
0
1
0
0
0

0.99
0.99
0.99
0.99
0.99
0.99
0.99
0.99
0.99
0.99
0.99
0.99
0.99
0.99
0.99
0.99
0.99
0.99
0.99
0.99
0.99
0.99
0.99
0.99
0.99
0.99
0.99
0.99
0.99
0.99


1
0
0
0
0
0
0
0
0
0
0
0
0
0
0
0
0
0
0
0
0
0
0
0
1
0
0
0
0
0


1
0
0
0
0
0
0
0
0
0
0
0
0
0
0
0
0
0
0
0
0
0
0
0
1
0
0
0
0
0

1.0
1.0
1.0
1.0
1.0
1.0
1.0
1.0
1.0
1.0
1.0
1.0
1.0
1.0
1.0
1.0
1.0
1.0
1.0
1.0
1.0
1.0
1.0
1.0
1.0
1.0
1.0
1.0
1.0
1.0


1
0
0
0
0
0
0
0
0
0
0
0
0
0
0
0
0
0
0
0
0
0
0
0
0
1
0
0
0
0


1
0
0
0
0
0
0
0
0
0
0
0
0
0
0
0
0
0
0
0
0
0
0
0
0
1
0
0
0
0

1.0
1.0
1.0
1.0
1.0
1.0
1.0
1.0
1.0
1.0
1.0
1.0
1.0
1.0
1.0
1.0
1.0
1.0
1.0
1.0
1.0
1.0
1.0
1.0
1.0
1.0
1.0
1.0
1.0
1.0


188
5
1
0
3
1
3
7
9
6
5
7
0
0
0
1
0
1
1
1
12
1
0
0
1
34
29
31
20
9


148
2
0
0
3
1
2
2
2
2
2
2
0
0
0
1
0
1
1
0
12
1
0
0
0
30
29
27
20
8


148
2
0
0
3
1
2
2
2
2
2
2
0
0
0
1
0
1
1
0
12
1
0
0
0
30
29
27
20
8

1.0
1.0
1.0
1.0
1.0
1.0
1.0
1.0
1.0
1.0
1.0
1.0
1.0
1.0
1.0
1.0
1.0
1.0
1.0
1.0
1.0
1.0
1.0
1.0
1.0
1.0
1.0
1.0
1.0
1.0


5
1
0
0
0
0
0
1
0
2
0
1
0
0
0
0
0
0
0
0
0
0
0
0
0
0
0
0
0
0

1.0
1.0
1.0
1.0
1.0
1.0
1.0
1.0
1.0
1.0
1.0
1.0
1.0
1.0
1.0
1.0
1.0
1.0
1.0
1.0
1.0
1.0
1.0
1.0
1.0
1.0
1.0
1.0
1.0
1.0


2
1
0
0
0
0
0
0
0
1
0
0
0
0
0
0
0
0
0
0
0
0
0
0
0
0
0
0
0
0

0.93
0.93
0.93
0.93
0.93
0.93
0.93
0.93
0.93
0.93
0.93
0.93
0.93
0.93
0.93
0.93
0.93
0.93
0.93
0.93
0.93
0.93
0.93
0.93
0.93
0.93
0.93
0.93
0.93
0.93


22
2
0
0
0
0
0
4
7
2
3
4
0
0
0
0
0
0
0
0
0
0
0
0
0
0
0
0
0
0

1.0
1.0
1.0
1.0
1.0
1.0
1.0
1.0
1.0
1.0
1.0
1.0
1.0
1.0
1.0
1.0
1.0
1.0
1.0
1.0
1.0
1.0
1.0
1.0
1.0
1.0
1.0
1.0
1.0
1.0


3
0
0
0
0
0
0
0
1
1
1
0
0
0
0
0
0
0
0
0
0
0
0
0
0
0
0
0
0
0

1.0
1.0
1.0
1.0
1.0
1.0
1.0
1.0
1.0
1.0
1.0
1.0
1.0
1.0
1.0
1.0
1.0
1.0
1.0
1.0
1.0
1.0
1.0
1.0
1.0
1.0
1.0
1.0
1.0
1.0


3
0
0
0
0
0
0
0
0
1
1
1
0
0
0
0
0
0
0
0
0
0
0
0
0
0
0
0
0
0

1.0
1.0
1.0
1.0
1.0
1.0
1.0
1.0
1.0
1.0
1.0
1.0
1.0
1.0
1.0
1.0
1.0
1.0
1.0
1.0
1.0
1.0
1.0
1.0
1.0
1.0
1.0
1.0
1.0
1.0


1
0
0
0
0
0
0
0
1
0
0
0
0
0
0
0
0
0
0
0
0
0
0
0
0
0
0
0
0
0

1.0
1.0
1.0
1.0
1.0
1.0
1.0
1.0
1.0
1.0
1.0
1.0
1.0
1.0
1.0
1.0
1.0
1.0
1.0
1.0
1.0
1.0
1.0
1.0
1.0
1.0
1.0
1.0
1.0
1.0


12
0
1
0
0
0
1
0
0
0
0
0
0
0
0
0
0
0
0
0
0
0
0
0
1
4
0
4
0
1

1.0
1.0
1.0
1.0
1.0
1.0
1.0
1.0
1.0
1.0
1.0
1.0
1.0
1.0
1.0
1.0
1.0
1.0
1.0
1.0
1.0
1.0
1.0
1.0
1.0
1.0
1.0
1.0
1.0
1.0


7
0
1
0
0
0
1
0
0
0
0
0
0
0
0
0
0
0
0
0
0
0
0
0
0
2
0
3
0
0

1.0
1.0
1.0
1.0
1.0
1.0
1.0
1.0
1.0
1.0
1.0
1.0
1.0
1.0
1.0
1.0
1.0
1.0
1.0
1.0
1.0
1.0
1.0
1.0
1.0
1.0
1.0
1.0
1.0
1.0


1
0
0
0
0
0
0
0
0
0
0
0
0
0
0
0
0
0
0
1
0
0
0
0
0
0
0
0
0
0

1.0
1.0
1.0
1.0
1.0
1.0
1.0
1.0
1.0
1.0
1.0
1.0
1.0
1.0
1.0
1.0
1.0
1.0
1.0
1.0
1.0
1.0
1.0
1.0
1.0
1.0
1.0
1.0
1.0
1.0


283
13
12
17
10
14
9
1
5
10
15
16
2
11
12
1
4
5
3
13
5
5
12
2
17
17
13
15
16
8

1.0
1.0
1.0
1.0
1.0
1.0
1.0
1.0
1.0
1.0
1.0
1.0
1.0
1.0
1.0
1.0
1.0
1.0
1.0
1.0
1.0
1.0
1.0
1.0
1.0
1.0
1.0
1.0
1.0
1.0


25
1
5
5
3
1
0
0
1
1
1
1
0
0
0
0
0
0
0
0
0
0
1
0
2
1
0
1
0
1

0.89
0.89
0.89
0.89
0.89
0.89
0.89
0.89
0.89
0.89
0.89
0.89
0.89
0.89
0.89
0.89
0.89
0.89
0.89
0.89
0.89
0.89
0.89
0.89
0.89
0.89
0.89
0.89
0.89
0.89


19
1
5
5
3
1
0
0
1
1
1
1
0
0
0
0
0
0
0
0
0
0
0
0
0
0
0
0
0
0

1.0
1.0
1.0
1.0
1.0
1.0
1.0
1.0
1.0
1.0
1.0
1.0
1.0
1.0
1.0
1.0
1.0
1.0
1.0
1.0
1.0
1.0
1.0
1.0
1.0
1.0
1.0
1.0
1.0
1.0


1
0
0
0
0
0
0
0
0
0
0
0
0
0
0
0
0
0
0
0
0
0
0
0
0
1
0
0
0
0

1.0
1.0
1.0
1.0
1.0
1.0
1.0
1.0
1.0
1.0
1.0
1.0
1.0
1.0
1.0
1.0
1.0
1.0
1.0
1.0
1.0
1.0
1.0
1.0
1.0
1.0
1.0
1.0
1.0
1.0


54
1
1
1
2
1
1
1
2
2
2
2
2
0
3
1
2
3
3
1
3
3
3
2
2
2
2
2
2
2


54
1
1
1
2
1
1
1
2
2
2
2
2
0
3
1
2
3
3
1
3
3
3
2
2
2
2
2
2
2

1.0
1.0
1.0
1.0
1.0
1.0
1.0
1.0
1.0
1.0
1.0
1.0
1.0
1.0
1.0
1.0
1.0
1.0
1.0
1.0
1.0
1.0
1.0
1.0
1.0
1.0
1.0
1.0
1.0
1.0


25
0
1
1
0
2
2
0
0
3
4
2
0
0
1
0
0
0
0
0
1
1
2
0
2
1
0
2
0
0


15
0
1
1
0
2
2
0
0
0
2
0
0
0
0
0
0
0
0
0
1
1
2
0
0
1
0
2
0
0

1.0
1.0
1.0
1.0
1.0
1.0
1.0
1.0
1.0
1.0
1.0
1.0
1.0
1.0
1.0
1.0
1.0
1.0
1.0
1.0
1.0
1.0
1.0
1.0
1.0
1.0
1.0
1.0
1.0
1.0


9
0
0
0
0
0
0
0
0
3
2
1
0
0
1
0
0
0
0
0
0
0
0
0
2
0
0
0
0
0

1.0
1.0
1.0
1.0
1.0
1.0
1.0
1.0
1.0
1.0
1.0
1.0
1.0
1.0
1.0
1.0
1.0
1.0
1.0
1.0
1.0
1.0
1.0
1.0
1.0
1.0
1.0
1.0
1.0
1.0


1
0
0
0
0
0
0
0
0
0
0
1
0
0
0
0
0
0
0
0
0
0
0
0
0
0
0
0
0
0

1.0
1.0
1.0
1.0
1.0
1.0
1.0
1.0
1.0
1.0
1.0
1.0
1.0
1.0
1.0
1.0
1.0
1.0
1.0
1.0
1.0
1.0
1.0
1.0
1.0
1.0
1.0
1.0
1.0
1.0


19
0
0
2
1
2
2
0
0
0
1
1
0
0
0
0
0
0
0
1
0
1
0
0
2
2
1
1
1
1

1.0
1.0
1.0
1.0
1.0
1.0
1.0
1.0
1.0
1.0
1.0
1.0
1.0
1.0
1.0
1.0
1.0
1.0
1.0
1.0
1.0
1.0
1.0
1.0
1.0
1.0
1.0
1.0
1.0
1.0


18
3
2
0
0
1
0
0
2
1
3
2
0
0
0
0
0
0
0
0
0
0
0
0
3
0
0
0
1
0

1.0
1.0
1.0
1.0
1.0
1.0
1.0
1.0
1.0
1.0
1.0
1.0
1.0
1.0
1.0
1.0
1.0
1.0
1.0
1.0
1.0
1.0
1.0
1.0
1.0
1.0
1.0
1.0
1.0
1.0


1
0
0
0
0
0
0
0
0
0
0
0
0
0
0
0
0
0
0
0
0
0
0
0
1
0
0
0
0
0

0.93
0.93
0.93
0.93
0.93
0.93
0.93
0.93
0.93
0.93
0.93
0.93
0.93
0.93
0.93
0.93
0.93
0.93
0.93
0.93
0.93
0.93
0.93
0.93
0.93
0.93
0.93
0.93
0.93
0.93


62
6
0
1
1
1
1
0
0
3
4
6
0
11
7
0
0
0
0
9
0
0
6
0
2
0
2
0
2
0

1.0
1.0
1.0
1.0
1.0
1.0
1.0
1.0
1.0
1.0
1.0
1.0
1.0
1.0
1.0
1.0
1.0
1.0
1.0
1.0
1.0
1.0
1.0
1.0
1.0
1.0
1.0
1.0
1.0
1.0


15
2
0
0
0
0
0
0
0
2
2
2
0
3
1
0
0
0
0
2
0
0
1
0
0
0
0
0
0
0

1.0
1.0
1.0
1.0
1.0
1.0
1.0
1.0
1.0
1.0
1.0
1.0
1.0
1.0
1.0
1.0
1.0
1.0
1.0
1.0
1.0
1.0
1.0
1.0
1.0
1.0
1.0
1.0
1.0
1.0


6
0
0
0
0
0
0
0
0
0
0
0
0
0
0
0
0
0
0
0
0
0
0
0
2
0
2
0
2
0

1.0
1.0
1.0
1.0
1.0
1.0
1.0
1.0
1.0
1.0
1.0
1.0
1.0
1.0
1.0
1.0
1.0
1.0
1.0
1.0
1.0
1.0
1.0
1.0
1.0
1.0
1.0
1.0
1.0
1.0


11
2
0
1
0
2
1
0
0
0
0
0
0
0
1
0
0
1
0
0
0
0
0
0
1
0
0
1
1
0

0.8
0.8
0.8
0.8
0.8
0.8
0.8
0.8
0.8
0.8
0.8
0.8
0.8
0.8
0.8
0.8
0.8
0.8
0.8
0.8
0.8
0.8
0.8
0.8
0.8
0.8
0.8
0.8
0.8
0.8


8
1
0
1
0
0
1
0
0
0
0
0
0
0
1
0
0
1
0
0
0
0
0
0
1
0
0
1
1
0

0.97
0.97
0.97
0.97
0.97
0.97
0.97
0.97
0.97
0.97
0.97
0.97
0.97
0.97
0.97
0.97
0.97
0.97
0.97
0.97
0.97
0.97
0.97
0.97
0.97
0.97
0.97
0.97
0.97
0.97


4
0
1
1
0
0
0
0
0
0
0
2
0
0
0
0
0
0
0
0
0
0
0
0
0
0
0
0
0
0

0.97
0.97
0.97
0.97
0.97
0.97
0.97
0.97
0.97
0.97
0.97
0.97
0.97
0.97
0.97
0.97
0.97
0.97
0.97
0.97
0.97
0.97
0.97
0.97
0.97
0.97
0.97
0.97
0.97
0.97


3
0
0
1
0
0
0
0
0
0
0
0
0
0
0
0
0
0
0
0
0
0
0
0
0
0
0
0
1
1


3
0
0
1
0
0
0
0
0
0
0
0
0
0
0
0
0
0
0
0
0
0
0
0
0
0
0
0
1
1

1.0
1.0
1.0
1.0
1.0
1.0
1.0
1.0
1.0
1.0
1.0
1.0
1.0
1.0
1.0
1.0
1.0
1.0
1.0
1.0
1.0
1.0
1.0
1.0
1.0
1.0
1.0
1.0
1.0
1.0


20
0
1
1
1
1
2
0
0
0
0
0
0
0
0
0
2
0
0
0
0
0
0
0
2
2
2
2
2
2


20
0
1
1
1
1
2
0
0
0
0
0
0
0
0
0
2
0
0
0
0
0
0
0
2
2
2
2
2
2

1.0
1.0
1.0
1.0
1.0
1.0
1.0
1.0
1.0
1.0
1.0
1.0
1.0
1.0
1.0
1.0
1.0
1.0
1.0
1.0
1.0
1.0
1.0
1.0
1.0
1.0
1.0
1.0
1.0
1.0


4
0
0
1
1
1
0
0
0
0
0
0
0
0
0
0
0
1
0
0
0
0
0
0
0
0
0
0
0
0

1.0
1.0
1.0
1.0
1.0
1.0
1.0
1.0
1.0
1.0
1.0
1.0
1.0
1.0
1.0
1.0
1.0
1.0
1.0
1.0
1.0
1.0
1.0
1.0
1.0
1.0
1.0
1.0
1.0
1.0


4
0
0
0
0
0
0
0
0
0
0
0
0
0
0
0
0
0
0
0
0
0
0
0
0
2
1
1
0
0


4
0
0
0
0
0
0
0
0
0
0
0
0
0
0
0
0
0
0
0
0
0
0
0
0
2
1
1
0
0

0.95
0.95
0.95
0.95
0.95
0.95
0.95
0.95
0.95
0.95
0.95
0.95
0.95
0.95
0.95
0.95
0.95
0.95
0.95
0.95
0.95
0.95
0.95
0.95
0.95
0.95
0.95
0.95
0.95
0.95


22
0
0
0
0
0
0
0
0
0
0
0
0
0
0
0
0
0
0
0
0
0
0
0
0
7
5
4
6
0

0.95
0.95
0.95
0.95
0.95
0.95
0.95
0.95
0.95
0.95
0.95
0.95
0.95
0.95
0.95
0.95
0.95
0.95
0.95
0.95
0.95
0.95
0.95
0.95
0.95
0.95
0.95
0.95
0.95
0.95


481
31
20
11
3
4
1
13
23
34
36
39
7
0
5
16
18
49
37
35
25
14
8
7
36
0
1
1
3
4

1.0
1.0
1.0
1.0
1.0
1.0
1.0
1.0
1.0
1.0
1.0
1.0
1.0
1.0
1.0
1.0
1.0
1.0
1.0
1.0
1.0
1.0
1.0
1.0
1.0
1.0
1.0
1.0
1.0
1.0


6
1
1
0
0
0
0
0
1
1
1
1
0
0
0
0
0
0
0
0
0
0
0
0
0
0
0
0
0
0


6
1
1
0
0
0
0
0
1
1
1
1
0
0
0
0
0
0
0
0
0
0
0
0
0
0
0
0
0
0

1.0
1.0
1.0
1.0
1.0
1.0
1.0
1.0
1.0
1.0
1.0
1.0
1.0
1.0
1.0
1.0
1.0
1.0
1.0
1.0
1.0
1.0
1.0
1.0
1.0
1.0
1.0
1.0
1.0
1.0


4
1
0
0
0
0
0
0
0
1
1
1
0
0
0
0
0
0
0
0
0
0
0
0
0
0
0
0
0
0


4
1
0
0
0
0
0
0
0
1
1
1
0
0
0
0
0
0
0
0
0
0
0
0
0
0
0
0
0
0

1.0
1.0
1.0
1.0
1.0
1.0
1.0
1.0
1.0
1.0
1.0
1.0
1.0
1.0
1.0
1.0
1.0
1.0
1.0
1.0
1.0
1.0
1.0
1.0
1.0
1.0
1.0
1.0
1.0
1.0


17
2
1
1
0
0
0
2
2
1
3
3
0
0
0
0
0
0
0
0
0
0
0
0
2
0
0
0
0
0


9
1
0
0
0
0
0
2
2
0
2
2
0
0
0
0
0
0
0
0
0
0
0
0
0
0
0
0
0
0

1.0
1.0
1.0
1.0
1.0
1.0
1.0
1.0
1.0
1.0
1.0
1.0
1.0
1.0
1.0
1.0
1.0
1.0
1.0
1.0
1.0
1.0
1.0
1.0
1.0
1.0
1.0
1.0
1.0
1.0


8
1
1
1
0
0
0
0
0
1
1
1
0
0
0
0
0
0
0
0
0
0
0
0
2
0
0
0
0
0

1.0
1.0
1.0
1.0
1.0
1.0
1.0
1.0
1.0
1.0
1.0
1.0
1.0
1.0
1.0
1.0
1.0
1.0
1.0
1.0
1.0
1.0
1.0
1.0
1.0
1.0
1.0
1.0
1.0
1.0


12
2
2
1
0
0
0
0
1
2
2
2
0
0
0
0
0
0
0
0
0
0
0
0
0
0
0
0
0
0

1.0
1.0
1.0
1.0
1.0
1.0
1.0
1.0
1.0
1.0
1.0
1.0
1.0
1.0
1.0
1.0
1.0
1.0
1.0
1.0
1.0
1.0
1.0
1.0
1.0
1.0
1.0
1.0
1.0
1.0


18
0
0
0
0
0
0
0
1
1
0
1
0
0
0
0
0
4
3
4
4
0
0
0
0
0
0
0
0
0

0.9
0.9
0.9
0.9
0.9
0.9
0.9
0.9
0.9
0.9
0.9
0.9
0.9
0.9
0.9
0.9
0.9
0.9
0.9
0.9
0.9
0.9
0.9
0.9
0.9
0.9
0.9
0.9
0.9
0.9


14
0
0
0
0
0
0
0
1
1
0
1
0
0
0
0
0
3
2
3
3
0
0
0
0
0
0
0
0
0

1.0
1.0
1.0
1.0
1.0
1.0
1.0
1.0
1.0
1.0
1.0
1.0
1.0
1.0
1.0
1.0
1.0
1.0
1.0
1.0
1.0
1.0
1.0
1.0
1.0
1.0
1.0
1.0
1.0
1.0


7
0
1
0
0
0
0
0
0
1
0
1
0
0
0
0
0
2
0
0
2
0
0
0
0
0
0
0
0
0


7
0
1
0
0
0
0
0
0
1
0
1
0
0
0
0
0
2
0
0
2
0
0
0
0
0
0
0
0
0

1.0
1.0
1.0
1.0
1.0
1.0
1.0
1.0
1.0
1.0
1.0
1.0
1.0
1.0
1.0
1.0
1.0
1.0
1.0
1.0
1.0
1.0
1.0
1.0
1.0
1.0
1.0
1.0
1.0
1.0


3
0
0
0
0
0
0
0
1
0
1
1
0
0
0
0
0
0
0
0
0
0
0
0
0
0
0
0
0
0


3
0
0
0
0
0
0
0
1
0
1
1
0
0
0
0
0
0
0
0
0
0
0
0
0
0
0
0
0
0

1.0
1.0
1.0
1.0
1.0
1.0
1.0
1.0
1.0
1.0
1.0
1.0
1.0
1.0
1.0
1.0
1.0
1.0
1.0
1.0
1.0
1.0
1.0
1.0
1.0
1.0
1.0
1.0
1.0
1.0


12
1
1
1
0
1
0
0
0
0
0
1
0
0
0
0
1
1
2
0
1
0
1
0
1
0
0
0
0
0


12
1
1
1
0
1
0
0
0
0
0
1
0
0
0
0
1
1
2
0
1
0
1
0
1
0
0
0
0
0

0.97
0.97
0.97
0.97
0.97
0.97
0.97
0.97
0.97
0.97
0.97
0.97
0.97
0.97
0.97
0.97
0.97
0.97
0.97
0.97
0.97
0.97
0.97
0.97
0.97
0.97
0.97
0.97
0.97
0.97


7
1
1
0
0
0
0
0
0
1
1
1
0
0
0
0
0
0
2
0
0
0
0
0
0
0
0
0
0
0


7
1
1
0
0
0
0
0
0
1
1
1
0
0
0
0
0
0
2
0
0
0
0
0
0
0
0
0
0
0

1.0
1.0
1.0
1.0
1.0
1.0
1.0
1.0
1.0
1.0
1.0
1.0
1.0
1.0
1.0
1.0
1.0
1.0
1.0
1.0
1.0
1.0
1.0
1.0
1.0
1.0
1.0
1.0
1.0
1.0


11
0
1
0
0
0
0
2
2
2
3
1
0
0
0
0
0
0
0
0
0
0
0
0
0
0
0
0
0
0

1.0
1.0
1.0
1.0
1.0
1.0
1.0
1.0
1.0
1.0
1.0
1.0
1.0
1.0
1.0
1.0
1.0
1.0
1.0
1.0
1.0
1.0
1.0
1.0
1.0
1.0
1.0
1.0
1.0
1.0


2
1
0
0
0
0
0
0
0
1
0
0
0
0
0
0
0
0
0
0
0
0
0
0
0
0
0
0
0
0


2
1
0
0
0
0
0
0
0
1
0
0
0
0
0
0
0
0
0
0
0
0
0
0
0
0
0
0
0
0

1.0
1.0
1.0
1.0
1.0
1.0
1.0
1.0
1.0
1.0
1.0
1.0
1.0
1.0
1.0
1.0
1.0
1.0
1.0
1.0
1.0
1.0
1.0
1.0
1.0
1.0
1.0
1.0
1.0
1.0


13
1
0
0
0
0
0
0
0
0
0
1
0
0
0
0
0
3
3
3
2
0
0
0
0
0
0
0
0
0


13
1
0
0
0
0
0
0
0
0
0
1
0
0
0
0
0
3
3
3
2
0
0
0
0
0
0
0
0
0

1.0
1.0
1.0
1.0
1.0
1.0
1.0
1.0
1.0
1.0
1.0
1.0
1.0
1.0
1.0
1.0
1.0
1.0
1.0
1.0
1.0
1.0
1.0
1.0
1.0
1.0
1.0
1.0
1.0
1.0


10
1
1
1
0
0
0
0
1
2
2
2
0
0
0
0
0
0
0
0
0
0
0
0
0
0
0
0
0
0


10
1
1
1
0
0
0
0
1
2
2
2
0
0
0
0
0
0
0
0
0
0
0
0
0
0
0
0
0
0

0.86
0.86
0.86
0.86
0.86
0.86
0.86
0.86
0.86
0.86
0.86
0.86
0.86
0.86
0.86
0.86
0.86
0.86
0.86
0.86
0.86
0.86
0.86
0.86
0.86
0.86
0.86
0.86
0.86
0.86


56
1
1
0
0
0
0
0
0
0
0
3
0
0
0
0
0
7
8
11
7
7
1
0
8
0
1
0
1
0

1.0
1.0
1.0
1.0
1.0
1.0
1.0
1.0
1.0
1.0
1.0
1.0
1.0
1.0
1.0
1.0
1.0
1.0
1.0
1.0
1.0
1.0
1.0
1.0
1.0
1.0
1.0
1.0
1.0
1.0


1
1
0
0
0
0
0
0
0
0
0
0
0
0
0
0
0
0
0
0
0
0
0
0
0
0
0
0
0
0

1.0
1.0
1.0
1.0
1.0
1.0
1.0
1.0
1.0
1.0
1.0
1.0
1.0
1.0
1.0
1.0
1.0
1.0
1.0
1.0
1.0
1.0
1.0
1.0
1.0
1.0
1.0
1.0
1.0
1.0


5
1
0
1
1
0
0
0
0
0
1
0
0
0
0
0
0
1
0
0
0
0
0
0
0
0
0
0
0
0


5
1
0
1
1
0
0
0
0
0
1
0
0
0
0
0
0
1
0
0
0
0
0
0
0
0
0
0
0
0

1.0
1.0
1.0
1.0
1.0
1.0
1.0
1.0
1.0
1.0
1.0
1.0
1.0
1.0
1.0
1.0
1.0
1.0
1.0
1.0
1.0
1.0
1.0
1.0
1.0
1.0
1.0
1.0
1.0
1.0


4
1
0
0
0
0
0
0
1
1
1
0
0
0
0
0
0
0
0
0
0
0
0
0
0
0
0
0
0
0


4
1
0
0
0
0
0
0
1
1
1
0
0
0
0
0
0
0
0
0
0
0
0
0
0
0
0
0
0
0

1.0
1.0
1.0
1.0
1.0
1.0
1.0
1.0
1.0
1.0
1.0
1.0
1.0
1.0
1.0
1.0
1.0
1.0
1.0
1.0
1.0
1.0
1.0
1.0
1.0
1.0
1.0
1.0
1.0
1.0


69
1
0
1
1
0
0
0
0
0
0
1
0
0
0
11
12
14
14
0
0
0
0
0
14
0
0
0
0
0


69
1
0
1
1
0
0
0
0
0
0
1
0
0
0
11
12
14
14
0
0
0
0
0
14
0
0
0
0
0

1.0
1.0
1.0
1.0
1.0
1.0
1.0
1.0
1.0
1.0
1.0
1.0
1.0
1.0
1.0
1.0
1.0
1.0
1.0
1.0
1.0
1.0
1.0
1.0
1.0
1.0
1.0
1.0
1.0
1.0


30
1
1
1
0
0
1
0
0
2
1
1
3
0
0
1
1
2
2
3
2
1
0
3
0
0
0
0
2
2

0.99
0.99
0.99
0.99
0.99
0.99
0.99
0.99
0.99
0.99
0.99
0.99
0.99
0.99
0.99
0.99
0.99
0.99
0.99
0.99
0.99
0.99
0.99
0.99
0.99
0.99
0.99
0.99
0.99
0.99


17
1
1
1
0
0
1
0
0
1
1
1
1
0
0
1
1
1
1
1
1
1
0
1
0
0
0
0
0
1

1.0
1.0
1.0
1.0
1.0
1.0
1.0
1.0
1.0
1.0
1.0
1.0
1.0
1.0
1.0
1.0
1.0
1.0
1.0
1.0
1.0
1.0
1.0
1.0
1.0
1.0
1.0
1.0
1.0
1.0


7
0
0
0
0
0
0
0
0
0
0
0
1
0
0
0
0
1
1
1
1
0
0
1
0
0
0
0
0
1

1.0
1.0
1.0
1.0
1.0
1.0
1.0
1.0
1.0
1.0
1.0
1.0
1.0
1.0
1.0
1.0
1.0
1.0
1.0
1.0
1.0
1.0
1.0
1.0
1.0
1.0
1.0
1.0
1.0
1.0


2
0
0
0
0
0
0
0
0
0
0
0
1
0
0
0
0
0
0
0
0
0
0
1
0
0
0
0
0
0

1.0
1.0
1.0
1.0
1.0
1.0
1.0
1.0
1.0
1.0
1.0
1.0
1.0
1.0
1.0
1.0
1.0
1.0
1.0
1.0
1.0
1.0
1.0
1.0
1.0
1.0
1.0
1.0
1.0
1.0


1
0
0
0
0
0
0
0
0
0
0
0
0
0
0
0
0
0
0
1
0
0
0
0
0
0
0
0
0
0

0.97
0.97
0.97
0.97
0.97
0.97
0.97
0.97
0.97
0.97
0.97
0.97
0.97
0.97
0.97
0.97
0.97
0.97
0.97
0.97
0.97
0.97
0.97
0.97
0.97
0.97
0.97
0.97
0.97
0.97


3
1
0
0
0
0
0
0
0
1
1
0
0
0
0
0
0
0
0
0
0
0
0
0
0
0
0
0
0
0


3
1
0
0
0
0
0
0
0
1
1
0
0
0
0
0
0
0
0
0
0
0
0
0
0
0
0
0
0
0

1.0
1.0
1.0
1.0
1.0
1.0
1.0
1.0
1.0
1.0
1.0
1.0
1.0
1.0
1.0
1.0
1.0
1.0
1.0
1.0
1.0
1.0
1.0
1.0
1.0
1.0
1.0
1.0
1.0
1.0


1
0
0
0
0
0
0
0
0
0
1
0
0
0
0
0
0
0
0
0
0
0
0
0
0
0
0
0
0
0

1.0
1.0
1.0
1.0
1.0
1.0
1.0
1.0
1.0
1.0
1.0
1.0
1.0
1.0
1.0
1.0
1.0
1.0
1.0
1.0
1.0
1.0
1.0
1.0
1.0
1.0
1.0
1.0
1.0
1.0


1
0
0
0
0
1
0
0
0
0
0
0
0
0
0
0
0
0
0
0
0
0
0
0
0
0
0
0
0
0


1
0
0
0
0
1
0
0
0
0
0
0
0
0
0
0
0
0
0
0
0
0
0
0
0
0
0
0
0
0

1.0
1.0
1.0
1.0
1.0
1.0
1.0
1.0
1.0
1.0
1.0
1.0
1.0
1.0
1.0
1.0
1.0
1.0
1.0
1.0
1.0
1.0
1.0
1.0
1.0
1.0
1.0
1.0
1.0
1.0


1
0
0
0
0
1
0
0
0
0
0
0
0
0
0
0
0
0
0
0
0
0
0
0
0
0
0
0
0
0

0.74
0.74
0.74
0.74
0.74
0.74
0.74
0.74
0.74
0.74
0.74
0.74
0.74
0.74
0.74
0.74
0.74
0.74
0.74
0.74
0.74
0.74
0.74
0.74
0.74
0.74
0.74
0.74
0.74
0.74


3
0
0
0
0
0
0
0
0
0
2
1
0
0
0
0
0
0
0
0
0
0
0
0
0
0
0
0
0
0


3
0
0
0
0
0
0
0
0
0
2
1
0
0
0
0
0
0
0
0
0
0
0
0
0
0
0
0
0
0

0.97
0.97
0.97
0.97
0.97
0.97
0.97
0.97
0.97
0.97
0.97
0.97
0.97
0.97
0.97
0.97
0.97
0.97
0.97
0.97
0.97
0.97
0.97
0.97
0.97
0.97
0.97
0.97
0.97
0.97


1
1
0
0
0
0
0
0
0
0
0
0
0
0
0
0
0
0
0
0
0
0
0
0
0
0
0
0
0
0


1
1
0
0
0
0
0
0
0
0
0
0
0
0
0
0
0
0
0
0
0
0
0
0
0
0
0
0
0
0

1.0
1.0
1.0
1.0
1.0
1.0
1.0
1.0
1.0
1.0
1.0
1.0
1.0
1.0
1.0
1.0
1.0
1.0
1.0
1.0
1.0
1.0
1.0
1.0
1.0
1.0
1.0
1.0
1.0
1.0


1
0
1
0
0
0
0
0
0
0
0
0
0
0
0
0
0
0
0
0
0
0
0
0
0
0
0
0
0
0


1
0
1
0
0
0
0
0
0
0
0
0
0
0
0
0
0
0
0
0
0
0
0
0
0
0
0
0
0
0

1.0
1.0
1.0
1.0
1.0
1.0
1.0
1.0
1.0
1.0
1.0
1.0
1.0
1.0
1.0
1.0
1.0
1.0
1.0
1.0
1.0
1.0
1.0
1.0
1.0
1.0
1.0
1.0
1.0
1.0


1
0
0
0
0
0
0
0
0
0
0
1
0
0
0
0
0
0
0
0
0
0
0
0
0
0
0
0
0
0


1
0
0
0
0
0
0
0
0
0
0
1
0
0
0
0
0
0
0
0
0
0
0
0
0
0
0
0
0
0

1.0
1.0
1.0
1.0
1.0
1.0
1.0
1.0
1.0
1.0
1.0
1.0
1.0
1.0
1.0
1.0
1.0
1.0
1.0
1.0
1.0
1.0
1.0
1.0
1.0
1.0
1.0
1.0
1.0
1.0


3
0
1
0
0
1
0
0
0
0
1
0
0
0
0
0
0
0
0
0
0
0
0
0
0
0
0
0
0
0

1.0
1.0
1.0
1.0
1.0
1.0
1.0
1.0
1.0
1.0
1.0
1.0
1.0
1.0
1.0
1.0
1.0
1.0
1.0
1.0
1.0
1.0
1.0
1.0
1.0
1.0
1.0
1.0
1.0
1.0


2
0
0
0
0
0
0
0
0
1
0
1
0
0
0
0
0
0
0
0
0
0
0
0
0
0
0
0
0
0


2
0
0
0
0
0
0
0
0
1
0
1
0
0
0
0
0
0
0
0
0
0
0
0
0
0
0
0
0
0

1.0
1.0
1.0
1.0
1.0
1.0
1.0
1.0
1.0
1.0
1.0
1.0
1.0
1.0
1.0
1.0
1.0
1.0
1.0
1.0
1.0
1.0
1.0
1.0
1.0
1.0
1.0
1.0
1.0
1.0


3
1
1
0
0
0
0
0
0
1
0
0
0
0
0
0
0
0
0
0
0
0
0
0
0
0
0
0
0
0


3
1
1
0
0
0
0
0
0
1
0
0
0
0
0
0
0
0
0
0
0
0
0
0
0
0
0
0
0
0

1.0
1.0
1.0
1.0
1.0
1.0
1.0
1.0
1.0
1.0
1.0
1.0
1.0
1.0
1.0
1.0
1.0
1.0
1.0
1.0
1.0
1.0
1.0
1.0
1.0
1.0
1.0
1.0
1.0
1.0


2
0
0
0
0
0
0
0
0
1
1
0
0
0
0
0
0
0
0
0
0
0
0
0
0
0
0
0
0
0


2
0
0
0
0
0
0
0
0
1
1
0
0
0
0
0
0
0
0
0
0
0
0
0
0
0
0
0
0
0

1.0
1.0
1.0
1.0
1.0
1.0
1.0
1.0
1.0
1.0
1.0
1.0
1.0
1.0
1.0
1.0
1.0
1.0
1.0
1.0
1.0
1.0
1.0
1.0
1.0
1.0
1.0
1.0
1.0
1.0


10
0
0
0
0
0
0
0
0
0
0
1
0
0
0
2
2
0
0
1
0
0
0
0
2
0
0
0
0
2

1.0
1.0
1.0
1.0
1.0
1.0
1.0
1.0
1.0
1.0
1.0
1.0
1.0
1.0
1.0
1.0
1.0
1.0
1.0
1.0
1.0
1.0
1.0
1.0
1.0
1.0
1.0
1.0
1.0
1.0


10
0
0
0
0
0
0
0
0
0
0
0
1
0
0
0
0
1
1
0
0
3
3
1
0
0
0
0
0
0

1.0
1.0
1.0
1.0
1.0
1.0
1.0
1.0
1.0
1.0
1.0
1.0
1.0
1.0
1.0
1.0
1.0
1.0
1.0
1.0
1.0
1.0
1.0
1.0
1.0
1.0
1.0
1.0
1.0
1.0


3
0
0
0
0
0
0
0
0
0
0
0
0
0
0
0
0
0
0
0
1
1
1
0
0
0
0
0
0
0


3
0
0
0
0
0
0
0
0
0
0
0
0
0
0
0
0
0
0
0
1
1
1
0
0
0
0
0
0
0

1.0
1.0
1.0
1.0
1.0
1.0
1.0
1.0
1.0
1.0
1.0
1.0
1.0
1.0
1.0
1.0
1.0
1.0
1.0
1.0
1.0
1.0
1.0
1.0
1.0
1.0
1.0
1.0
1.0
1.0


9
0
0
0
0
0
0
0
0
0
0
0
0
0
0
0
0
7
0
0
1
0
0
0
1
0
0
0
0
0

1.0
1.0
1.0
1.0
1.0
1.0
1.0
1.0
1.0
1.0
1.0
1.0
1.0
1.0
1.0
1.0
1.0
1.0
1.0
1.0
1.0
1.0
1.0
1.0
1.0
1.0
1.0
1.0
1.0
1.0


7
0
0
0
0
0
0
0
0
0
0
0
1
0
1
0
0
1
0
2
1
0
0
1
0
0
0
0
0
0

1.0
1.0
1.0
1.0
1.0
1.0
1.0
1.0
1.0
1.0
1.0
1.0
1.0
1.0
1.0
1.0
1.0
1.0
1.0
1.0
1.0
1.0
1.0
1.0
1.0
1.0
1.0
1.0
1.0
1.0


1
0
0
0
0
0
0
0
0
0
0
0
0
0
1
0
0
0
0
0
0
0
0
0
0
0
0
0
0
0

0.96
0.96
0.96
0.96
0.96
0.96
0.96
0.96
0.96
0.96
0.96
0.96
0.96
0.96
0.96
0.96
0.96
0.96
0.96
0.96
0.96
0.96
0.96
0.96
0.96
0.96
0.96
0.96
0.96
0.96


1
0
0
0
0
0
0
0
0
0
0
0
0
0
0
0
0
0
0
1
0
0
0
0
0
0
0
0
0
0

1.0
1.0
1.0
1.0
1.0
1.0
1.0
1.0
1.0
1.0
1.0
1.0
1.0
1.0
1.0
1.0
1.0
1.0
1.0
1.0
1.0
1.0
1.0
1.0
1.0
1.0
1.0
1.0
1.0
1.0


1
0
0
0
0
0
0
0
0
0
0
0
0
0
0
0
0
0
0
0
1
0
0
0
0
0
0
0
0
0

1.0
1.0
1.0
1.0
1.0
1.0
1.0
1.0
1.0
1.0
1.0
1.0
1.0
1.0
1.0
1.0
1.0
1.0
1.0
1.0
1.0
1.0
1.0
1.0
1.0
1.0
1.0
1.0
1.0
1.0


6
0
0
0
0
0
0
0
0
0
0
0
0
0
0
1
1
1
1
1
0
0
0
0
0
0
0
1
0
0


6
0
0
0
0
0
0
0
0
0
0
0
0
0
0
1
1
1
1
1
0
0
0
0
0
0
0
1
0
0

0.92
0.92
0.92
0.92
0.92
0.92
0.92
0.92
0.92
0.92
0.92
0.92
0.92
0.92
0.92
0.92
0.92
0.92
0.92
0.92
0.92
0.92
0.92
0.92
0.92
0.92
0.92
0.92
0.92
0.92


3
0
0
0
0
0
0
0
0
0
0
0
0
0
0
0
0
0
0
0
0
1
2
0
0
0
0
0
0
0

0.87
0.87
0.87
0.87
0.87
0.87
0.87
0.87
0.87
0.87
0.87
0.87
0.87
0.87
0.87
0.87
0.87
0.87
0.87
0.87
0.87
0.87
0.87
0.87
0.87
0.87
0.87
0.87
0.87
0.87


4
0
0
0
0
0
0
0
0
0
0
0
0
0
0
0
0
0
0
2
1
1
0
0
0
0
0
0
0
0


4
0
0
0
0
0
0
0
0
0
0
0
0
0
0
0
0
0
0
2
1
1
0
0
0
0
0
0
0
0

1.0
1.0
1.0
1.0
1.0
1.0
1.0
1.0
1.0
1.0
1.0
1.0
1.0
1.0
1.0
1.0
1.0
1.0
1.0
1.0
1.0
1.0
1.0
1.0
1.0
1.0
1.0
1.0
1.0
1.0


2
0
0
0
0
0
0
0
0
0
0
0
1
0
0
0
0
0
0
0
0
0
0
1
0
0
0
0
0
0

1.0
1.0
1.0
1.0
1.0
1.0
1.0
1.0
1.0
1.0
1.0
1.0
1.0
1.0
1.0
1.0
1.0
1.0
1.0
1.0
1.0
1.0
1.0
1.0
1.0
1.0
1.0
1.0
1.0
1.0


6
0
0
0
0
0
0
0
0
0
0
0
0
0
0
0
0
2
0
2
0
0
0
0
2
0
0
0
0
0


6
0
0
0
0
0
0
0
0
0
0
0
0
0
0
0
0
2
0
2
0
0
0
0
2
0
0
0
0
0

1.0
1.0
1.0
1.0
1.0
1.0
1.0
1.0
1.0
1.0
1.0
1.0
1.0
1.0
1.0
1.0
1.0
1.0
1.0
1.0
1.0
1.0
1.0
1.0
1.0
1.0
1.0
1.0
1.0
1.0


1
0
0
0
0
0
0
0
0
0
0
0
0
0
0
0
0
1
0
0
0
0
0
0
0
0
0
0
0
0

1.0
1.0
1.0
1.0
1.0
1.0
1.0
1.0
1.0
1.0
1.0
1.0
1.0
1.0
1.0
1.0
1.0
1.0
1.0
1.0
1.0
1.0
1.0
1.0
1.0
1.0
1.0
1.0
1.0
1.0


34
3
3
0
0
0
0
1
9
11
4
3
0
0
0
0
0
0
0
0
0
0
0
0
0
0
0
0
0
0


34
3
3
0
0
0
0
1
9
11
4
3
0
0
0
0
0
0
0
0
0
0
0
0
0
0
0
0
0
0


6
0
0
0
0
0
0
0
3
3
0
0
0
0
0
0
0
0
0
0
0
0
0
0
0
0
0
0
0
0

1.0
1.0
1.0
1.0
1.0
1.0
1.0
1.0
1.0
1.0
1.0
1.0
1.0
1.0
1.0
1.0
1.0
1.0
1.0
1.0
1.0
1.0
1.0
1.0
1.0
1.0
1.0
1.0
1.0
1.0


1
0
0
0
0
0
0
1
0
0
0
0
0
0
0
0
0
0
0
0
0
0
0
0
0
0
0
0
0
0

1.0
1.0
1.0
1.0
1.0
1.0
1.0
1.0
1.0
1.0
1.0
1.0
1.0
1.0
1.0
1.0
1.0
1.0
1.0
1.0
1.0
1.0
1.0
1.0
1.0
1.0
1.0
1.0
1.0
1.0


24
2
2
0
0
0
0
0
6
8
3
3
0
0
0
0
0
0
0
0
0
0
0
0
0
0
0
0
0
0

1.0
1.0
1.0
1.0
1.0
1.0
1.0
1.0
1.0
1.0
1.0
1.0
1.0
1.0
1.0
1.0
1.0
1.0
1.0
1.0
1.0
1.0
1.0
1.0
1.0
1.0
1.0
1.0
1.0
1.0


1
0
1
0
0
0
0
0
0
0
0
0
0
0
0
0
0
0
0
0
0
0
0
0
0
0
0
0
0
0

1.0
1.0
1.0
1.0
1.0
1.0
1.0
1.0
1.0
1.0
1.0
1.0
1.0
1.0
1.0
1.0
1.0
1.0
1.0
1.0
1.0
1.0
1.0
1.0
1.0
1.0
1.0
1.0
1.0
1.0


2
1
0
0
0
0
0
0
0
0
1
0
0
0
0
0
0
0
0
0
0
0
0
0
0
0
0
0
0
0

1.0
1.0
1.0
1.0
1.0
1.0
1.0
1.0
1.0
1.0
1.0
1.0
1.0
1.0
1.0
1.0
1.0
1.0
1.0
1.0
1.0
1.0
1.0
1.0
1.0
1.0
1.0
1.0
1.0
1.0


162
0
4
2
4
6
9
0
0
0
1
9
0
0
0
4
1
11
2
5
1
0
0
0
90
6
3
2
1
1

0.98
0.98
0.98
0.98
0.98
0.98
0.98
0.98
0.98
0.98
0.98
0.98
0.98
0.98
0.98
0.98
0.98
0.98
0.98
0.98
0.98
0.98
0.98
0.98
0.98
0.98
0.98
0.98
0.98
0.98


8
0
1
0
1
1
2
0
0
0
0
1
0
0
0
1
0
0
0
0
0
0
0
0
0
0
0
1
0
0

1.0
1.0
1.0
1.0
1.0
1.0
1.0
1.0
1.0
1.0
1.0
1.0
1.0
1.0
1.0
1.0
1.0
1.0
1.0
1.0
1.0
1.0
1.0
1.0
1.0
1.0
1.0
1.0
1.0
1.0


6
0
1
0
1
1
2
0
0
0
0
1
0
0
0
0
0
0
0
0
0
0
0
0
0
0
0
0
0
0

0.72
0.72
0.72
0.72
0.72
0.72
0.72
0.72
0.72
0.72
0.72
0.72
0.72
0.72
0.72
0.72
0.72
0.72
0.72
0.72
0.72
0.72
0.72
0.72
0.72
0.72
0.72
0.72
0.72
0.72


5
0
1
1
1
0
1
0
0
0
0
1
0
0
0
0
0
0
0
0
0
0
0
0
0
0
0
0
0
0

0.71
0.71
0.71
0.71
0.71
0.71
0.71
0.71
0.71
0.71
0.71
0.71
0.71
0.71
0.71
0.71
0.71
0.71
0.71
0.71
0.71
0.71
0.71
0.71
0.71
0.71
0.71
0.71
0.71
0.71


3
0
0
0
0
0
0
0
0
0
0
3
0
0
0
0
0
0
0
0
0
0
0
0
0
0
0
0
0
0


3
0
0
0
0
0
0
0
0
0
0
3
0
0
0
0
0
0
0
0
0
0
0
0
0
0
0
0
0
0

1.0
1.0
1.0
1.0
1.0
1.0
1.0
1.0
1.0
1.0
1.0
1.0
1.0
1.0
1.0
1.0
1.0
1.0
1.0
1.0
1.0
1.0
1.0
1.0
1.0
1.0
1.0
1.0
1.0
1.0


12
0
1
0
1
1
2
0
0
0
0
1
0
0
0
1
1
1
1
1
0
0
0
0
0
1
0
0
0
0


11
0
1
0
1
1
1
0
0
0
0
1
0
0
0
1
1
1
1
1
0
0
0
0
0
1
0
0
0
0

1.0
1.0
1.0
1.0
1.0
1.0
1.0
1.0
1.0
1.0
1.0
1.0
1.0
1.0
1.0
1.0
1.0
1.0
1.0
1.0
1.0
1.0
1.0
1.0
1.0
1.0
1.0
1.0
1.0
1.0


1
0
0
0
0
0
1
0
0
0
0
0
0
0
0
0
0
0
0
0
0
0
0
0
0
0
0
0
0
0

1.0
1.0
1.0
1.0
1.0
1.0
1.0
1.0
1.0
1.0
1.0
1.0
1.0
1.0
1.0
1.0
1.0
1.0
1.0
1.0
1.0
1.0
1.0
1.0
1.0
1.0
1.0
1.0
1.0
1.0


3
0
0
0
0
0
1
0
0
0
0
0
0
0
0
0
0
0
1
1
0
0
0
0
0
0
0
0
0
0


3
0
0
0
0
0
1
0
0
0
0
0
0
0
0
0
0
0
1
1
0
0
0
0
0
0
0
0
0
0

1.0
1.0
1.0
1.0
1.0
1.0
1.0
1.0
1.0
1.0
1.0
1.0
1.0
1.0
1.0
1.0
1.0
1.0
1.0
1.0
1.0
1.0
1.0
1.0
1.0
1.0
1.0
1.0
1.0
1.0


2
0
0
0
0
1
1
0
0
0
0
0
0
0
0
0
0
0
0
0
0
0
0
0
0
0
0
0
0
0


2
0
0
0
0
1
1
0
0
0
0
0
0
0
0
0
0
0
0
0
0
0
0
0
0
0
0
0
0
0

1.0
1.0
1.0
1.0
1.0
1.0
1.0
1.0
1.0
1.0
1.0
1.0
1.0
1.0
1.0
1.0
1.0
1.0
1.0
1.0
1.0
1.0
1.0
1.0
1.0
1.0
1.0
1.0
1.0
1.0


2
0
0
0
0
2
0
0
0
0
0
0
0
0
0
0
0
0
0
0
0
0
0
0
0
0
0
0
0
0


2
0
0
0
0
2
0
0
0
0
0
0
0
0
0
0
0
0
0
0
0
0
0
0
0
0
0
0
0
0

1.0
1.0
1.0
1.0
1.0
1.0
1.0
1.0
1.0
1.0
1.0
1.0
1.0
1.0
1.0
1.0
1.0
1.0
1.0
1.0
1.0
1.0
1.0
1.0
1.0
1.0
1.0
1.0
1.0
1.0


2
0
0
0
0
0
0
0
0
0
1
1
0
0
0
0
0
0
0
0
0
0
0
0
0
0
0
0
0
0


2
0
0
0
0
0
0
0
0
0
1
1
0
0
0
0
0
0
0
0
0
0
0
0
0
0
0
0
0
0

1.0
1.0
1.0
1.0
1.0
1.0
1.0
1.0
1.0
1.0
1.0
1.0
1.0
1.0
1.0
1.0
1.0
1.0
1.0
1.0
1.0
1.0
1.0
1.0
1.0
1.0
1.0
1.0
1.0
1.0


92
0
0
1
0
0
0
0
0
0
0
1
0
0
0
0
0
0
0
0
0
0
0
0
90
0
0
0
0
0

1.0
1.0
1.0
1.0
1.0
1.0
1.0
1.0
1.0
1.0
1.0
1.0
1.0
1.0
1.0
1.0
1.0
1.0
1.0
1.0
1.0
1.0
1.0
1.0
1.0
1.0
1.0
1.0
1.0
1.0


2
0
0
1
0
0
0
0
0
0
0
1
0
0
0
0
0
0
0
0
0
0
0
0
0
0
0
0
0
0

1.0
1.0
1.0
1.0
1.0
1.0
1.0
1.0
1.0
1.0
1.0
1.0
1.0
1.0
1.0
1.0
1.0
1.0
1.0
1.0
1.0
1.0
1.0
1.0
1.0
1.0
1.0
1.0
1.0
1.0


74
0
0
0
0
0
0
0
0
0
0
0
0
0
0
0
0
0
0
0
0
0
0
0
74
0
0
0
0
0

1.0
1.0
1.0
1.0
1.0
1.0
1.0
1.0
1.0
1.0
1.0
1.0
1.0
1.0
1.0
1.0
1.0
1.0
1.0
1.0
1.0
1.0
1.0
1.0
1.0
1.0
1.0
1.0
1.0
1.0


6
0
1
0
0
1
1
0
0
0
0
0
0
0
0
0
0
0
0
0
0
0
0
0
0
3
0
0
0
0


5
0
1
0
0
0
1
0
0
0
0
0
0
0
0
0
0
0
0
0
0
0
0
0
0
3
0
0
0
0

1.0
1.0
1.0
1.0
1.0
1.0
1.0
1.0
1.0
1.0
1.0
1.0
1.0
1.0
1.0
1.0
1.0
1.0
1.0
1.0
1.0
1.0
1.0
1.0
1.0
1.0
1.0
1.0
1.0
1.0


1
0
0
0
0
1
0
0
0
0
0
0
0
0
0
0
0
0
0
0
0
0
0
0
0
0
0
0
0
0

1.0
1.0
1.0
1.0
1.0
1.0
1.0
1.0
1.0
1.0
1.0
1.0
1.0
1.0
1.0
1.0
1.0
1.0
1.0
1.0
1.0
1.0
1.0
1.0
1.0
1.0
1.0
1.0
1.0
1.0


1
0
0
0
0
0
0
0
0
0
0
1
0
0
0
0
0
0
0
0
0
0
0
0
0
0
0
0
0
0


1
0
0
0
0
0
0
0
0
0
0
1
0
0
0
0
0
0
0
0
0
0
0
0
0
0
0
0
0
0

0.98
0.98
0.98
0.98
0.98
0.98
0.98
0.98
0.98
0.98
0.98
0.98
0.98
0.98
0.98
0.98
0.98
0.98
0.98
0.98
0.98
0.98
0.98
0.98
0.98
0.98
0.98
0.98
0.98
0.98


10
0
0
0
0
0
0
0
0
0
0
0
0
0
0
0
0
10
0
0
0
0
0
0
0
0
0
0
0
0

1.0
1.0
1.0
1.0
1.0
1.0
1.0
1.0
1.0
1.0
1.0
1.0
1.0
1.0
1.0
1.0
1.0
1.0
1.0
1.0
1.0
1.0
1.0
1.0
1.0
1.0
1.0
1.0
1.0
1.0


3
0
0
0
0
0
0
0
0
0
0
0
0
0
0
0
0
0
0
3
0
0
0
0
0
0
0
0
0
0

1.0
1.0
1.0
1.0
1.0
1.0
1.0
1.0
1.0
1.0
1.0
1.0
1.0
1.0
1.0
1.0
1.0
1.0
1.0
1.0
1.0
1.0
1.0
1.0
1.0
1.0
1.0
1.0
1.0
1.0


2
0
0
0
0
0
0
0
0
0
0
0
0
0
0
0
0
0
0
2
0
0
0
0
0
0
0
0
0
0

1.0
1.0
1.0
1.0
1.0
1.0
1.0
1.0
1.0
1.0
1.0
1.0
1.0
1.0
1.0
1.0
1.0
1.0
1.0
1.0
1.0
1.0
1.0
1.0
1.0
1.0
1.0
1.0
1.0
1.0


1
0
0
0
0
0
0
0
0
0
0
0
0
0
0
0
0
0
0
0
1
0
0
0
0
0
0
0
0
0


1
0
0
0
0
0
0
0
0
0
0
0
0
0
0
0
0
0
0
0
1
0
0
0
0
0
0
0
0
0

1.0
1.0
1.0
1.0
1.0
1.0
1.0
1.0
1.0
1.0
1.0
1.0
1.0
1.0
1.0
1.0
1.0
1.0
1.0
1.0
1.0
1.0
1.0
1.0
1.0
1.0
1.0
1.0
1.0
1.0


1
0
0
0
0
0
0
0
0
0
0
0
0
0
0
1
0
0
0
0
0
0
0
0
0
0
0
0
0
0


1
0
0
0
0
0
0
0
0
0
0
0
0
0
0
1
0
0
0
0
0
0
0
0
0
0
0
0
0
0

1.0
1.0
1.0
1.0
1.0
1.0
1.0
1.0
1.0
1.0
1.0
1.0
1.0
1.0
1.0
1.0
1.0
1.0
1.0
1.0
1.0
1.0
1.0
1.0
1.0
1.0
1.0
1.0
1.0
1.0


56
2
2
2
2
0
0
0
2
2
0
0
0
9
19
0
0
0
0
0
0
0
0
0
15
1
0
0
0
0

1.0
1.0
1.0
1.0
1.0
1.0
1.0
1.0
1.0
1.0
1.0
1.0
1.0
1.0
1.0
1.0
1.0
1.0
1.0
1.0
1.0
1.0
1.0
1.0
1.0
1.0
1.0
1.0
1.0
1.0


3
1
0
0
0
0
0
0
1
1
0
0
0
0
0
0
0
0
0
0
0
0
0
0
0
0
0
0
0
0


3
1
0
0
0
0
0
0
1
1
0
0
0
0
0
0
0
0
0
0
0
0
0
0
0
0
0
0
0
0

1.0
1.0
1.0
1.0
1.0
1.0
1.0
1.0
1.0
1.0
1.0
1.0
1.0
1.0
1.0
1.0
1.0
1.0
1.0
1.0
1.0
1.0
1.0
1.0
1.0
1.0
1.0
1.0
1.0
1.0


2
1
0
0
0
0
0
0
0
1
0
0
0
0
0
0
0
0
0
0
0
0
0
0
0
0
0
0
0
0


2
1
0
0
0
0
0
0
0
1
0
0
0
0
0
0
0
0
0
0
0
0
0
0
0
0
0
0
0
0

1.0
1.0
1.0
1.0
1.0
1.0
1.0
1.0
1.0
1.0
1.0
1.0
1.0
1.0
1.0
1.0
1.0
1.0
1.0
1.0
1.0
1.0
1.0
1.0
1.0
1.0
1.0
1.0
1.0
1.0


1
0
0
0
0
0
0
0
1
0
0
0
0
0
0
0
0
0
0
0
0
0
0
0
0
0
0
0
0
0


1
0
0
0
0
0
0
0
1
0
0
0
0
0
0
0
0
0
0
0
0
0
0
0
0
0
0
0
0
0

1.0
1.0
1.0
1.0
1.0
1.0
1.0
1.0
1.0
1.0
1.0
1.0
1.0
1.0
1.0
1.0
1.0
1.0
1.0
1.0
1.0
1.0
1.0
1.0
1.0
1.0
1.0
1.0
1.0
1.0


7
0
0
0
0
0
0
0
0
0
0
0
0
2
2
0
0
0
0
0
0
0
0
0
2
1
0
0
0
0


7
0
0
0
0
0
0
0
0
0
0
0
0
2
2
0
0
0
0
0
0
0
0
0
2
1
0
0
0
0

1.0
1.0
1.0
1.0
1.0
1.0
1.0
1.0
1.0
1.0
1.0
1.0
1.0
1.0
1.0
1.0
1.0
1.0
1.0
1.0
1.0
1.0
1.0
1.0
1.0
1.0
1.0
1.0
1.0
1.0


8
0
1
1
0
1
0
0
0
0
1
0
0
0
0
1
0
1
0
0
0
0
0
0
1
0
0
0
1
0

0.83
0.83
0.83
0.83
0.83
0.83
0.83
0.83
0.83
0.83
0.83
0.83
0.83
0.83
0.83
0.83
0.83
0.83
0.83
0.83
0.83
0.83
0.83
0.83
0.83
0.83
0.83
0.83
0.83
0.83


7
0
1
1
0
1
0
0
0
0
0
0
0
0
0
1
0
1
0
0
0
0
0
0
1
0
0
0
1
0


7
0
1
1
0
1
0
0
0
0
0
0
0
0
0
1
0
1
0
0
0
0
0
0
1
0
0
0
1
0

1.0
1.0
1.0
1.0
1.0
1.0
1.0
1.0
1.0
1.0
1.0
1.0
1.0
1.0
1.0
1.0
1.0
1.0
1.0
1.0
1.0
1.0
1.0
1.0
1.0
1.0
1.0
1.0
1.0
1.0


4
0
1
0
1
1
1
0
0
0
0
0
0
0
0
0
0
0
0
0
0
0
0
0
0
0
0
0
0
0


4
0
1
0
1
1
1
0
0
0
0
0
0
0
0
0
0
0
0
0
0
0
0
0
0
0
0
0
0
0


4
0
1
0
1
1
1
0
0
0
0
0
0
0
0
0
0
0
0
0
0
0
0
0
0
0
0
0
0
0

0.92
0.92
0.92
0.92
0.92
0.92
0.92
0.92
0.92
0.92
0.92
0.92
0.92
0.92
0.92
0.92
0.92
0.92
0.92
0.92
0.92
0.92
0.92
0.92
0.92
0.92
0.92
0.92
0.92
0.92


41
1
5
0
0
0
0
0
0
1
2
2
0
0
0
1
0
0
0
0
0
0
0
0
3
0
8
3
0
15

0.74
0.74
0.74
0.74
0.74
0.74
0.74
0.74
0.74
0.74
0.74
0.74
0.74
0.74
0.74
0.74
0.74
0.74
0.74
0.74
0.74
0.74
0.74
0.74
0.74
0.74
0.74
0.74
0.74
0.74


9
1
2
0
0
0
0
0
0
0
1
2
0
0
0
0
0
0
0
0
0
0
0
0
3
0
0
0
0
0

0.87
0.87
0.87
0.87
0.87
0.87
0.87
0.87
0.87
0.87
0.87
0.87
0.87
0.87
0.87
0.87
0.87
0.87
0.87
0.87
0.87
0.87
0.87
0.87
0.87
0.87
0.87
0.87
0.87
0.87


7
1
2
0
0
0
0
0
0
0
1
2
0
0
0
0
0
0
0
0
0
0
0
0
1
0
0
0
0
0

1.0
1.0
1.0
1.0
1.0
1.0
1.0
1.0
1.0
1.0
1.0
1.0
1.0
1.0
1.0
1.0
1.0
1.0
1.0
1.0
1.0
1.0
1.0
1.0
1.0
1.0
1.0
1.0
1.0
1.0


5
0
3
0
0
0
0
0
0
1
1
0
0
0
0
0
0
0
0
0
0
0
0
0
0
0
0
0
0
0


3
0
3
0
0
0
0
0
0
0
0
0
0
0
0
0
0
0
0
0
0
0
0
0
0
0
0
0
0
0

0.83
0.83
0.83
0.83
0.83
0.83
0.83
0.83
0.83
0.83
0.83
0.83
0.83
0.83
0.83
0.83
0.83
0.83
0.83
0.83
0.83
0.83
0.83
0.83
0.83
0.83
0.83
0.83
0.83
0.83


2
0
0
0
0
0
0
0
0
1
1
0
0
0
0
0
0
0
0
0
0
0
0
0
0
0
0
0
0
0

1.0
1.0
1.0
1.0
1.0
1.0
1.0
1.0
1.0
1.0
1.0
1.0
1.0
1.0
1.0
1.0
1.0
1.0
1.0
1.0
1.0
1.0
1.0
1.0
1.0
1.0
1.0
1.0
1.0
1.0


13
0
0
0
0
0
0
0
0
0
0
0
0
0
0
1
0
0
0
0
0
0
0
0
0
0
7
3
0
2

1.0
1.0
1.0
1.0
1.0
1.0
1.0
1.0
1.0
1.0
1.0
1.0
1.0
1.0
1.0
1.0
1.0
1.0
1.0
1.0
1.0
1.0
1.0
1.0
1.0
1.0
1.0
1.0
1.0
1.0


2
0
0
0
0
0
0
0
0
0
0
0
0
0
0
0
0
0
0
0
0
0
0
0
0
0
0
1
0
1

0.84
0.84
0.84
0.84
0.84
0.84
0.84
0.84
0.84
0.84
0.84
0.84
0.84
0.84
0.84
0.84
0.84
0.84
0.84
0.84
0.84
0.84
0.84
0.84
0.84
0.84
0.84
0.84
0.84
0.84


1
0
0
0
0
0
0
0
0
0
0
0
0
0
0
0
0
0
0
0
0
0
0
0
0
0
1
0
0
0

0.98
0.98
0.98
0.98
0.98
0.98
0.98
0.98
0.98
0.98
0.98
0.98
0.98
0.98
0.98
0.98
0.98
0.98
0.98
0.98
0.98
0.98
0.98
0.98
0.98
0.98
0.98
0.98
0.98
0.98


1
0
0
0
0
0
0
0
0
0
0
0
0
0
0
1
0
0
0
0
0
0
0
0
0
0
0
0
0
0

1.0
1.0
1.0
1.0
1.0
1.0
1.0
1.0
1.0
1.0
1.0
1.0
1.0
1.0
1.0
1.0
1.0
1.0
1.0
1.0
1.0
1.0
1.0
1.0
1.0
1.0
1.0
1.0
1.0
1.0


12
0
0
0
0
0
0
0
0
0
0
0
0
0
0
0
0
0
0
0
0
0
0
0
0
0
0
0
0
12


12
0
0
0
0
0
0
0
0
0
0
0
0
0
0
0
0
0
0
0
0
0
0
0
0
0
0
0
0
12

1.0
1.0
1.0
1.0
1.0
1.0
1.0
1.0
1.0
1.0
1.0
1.0
1.0
1.0
1.0
1.0
1.0
1.0
1.0
1.0
1.0
1.0
1.0
1.0
1.0
1.0
1.0
1.0
1.0
1.0


1
0
0
0
0
0
0
0
0
0
0
0
0
0
0
0
0
0
0
0
0
0
0
0
0
0
0
0
0
1


1
0
0
0
0
0
0
0
0
0
0
0
0
0
0
0
0
0
0
0
0
0
0
0
0
0
0
0
0
1

0.97
0.97
0.97
0.97
0.97
0.97
0.97
0.97
0.97
0.97
0.97
0.97
0.97
0.97
0.97
0.97
0.97
0.97
0.97
0.97
0.97
0.97
0.97
0.97
0.97
0.97
0.97
0.97
0.97
0.97


177
15
13
6
3
3
2
9
9
15
15
16
0
3
2
6
6
10
7
5
4
0
1
0
8
5
3
0
5
6

1.0
1.0
1.0
1.0
1.0
1.0
1.0
1.0
1.0
1.0
1.0
1.0
1.0
1.0
1.0
1.0
1.0
1.0
1.0
1.0
1.0
1.0
1.0
1.0
1.0
1.0
1.0
1.0
1.0
1.0


16
2
2
2
0
1
0
2
2
2
2
1
0
0
0
0
0
0
0
0
0
0
0
0
0
0
0
0
0
0


16
2
2
2
0
1
0
2
2
2
2
1
0
0
0
0
0
0
0
0
0
0
0
0
0
0
0
0
0
0

1.0
1.0
1.0
1.0
1.0
1.0
1.0
1.0
1.0
1.0
1.0
1.0
1.0
1.0
1.0
1.0
1.0
1.0
1.0
1.0
1.0
1.0
1.0
1.0
1.0
1.0
1.0
1.0
1.0
1.0


23
1
2
1
1
1
0
0
0
3
2
2
0
0
0
0
0
1
4
2
3
0
0
0
0
0
0
0
0
0

1.0
1.0
1.0
1.0
1.0
1.0
1.0
1.0
1.0
1.0
1.0
1.0
1.0
1.0
1.0
1.0
1.0
1.0
1.0
1.0
1.0
1.0
1.0
1.0
1.0
1.0
1.0
1.0
1.0
1.0


10
1
1
1
1
1
0
0
0
1
1
1
0
0
0
0
0
1
0
0
1
0
0
0
0
0
0
0
0
0

1.0
1.0
1.0
1.0
1.0
1.0
1.0
1.0
1.0
1.0
1.0
1.0
1.0
1.0
1.0
1.0
1.0
1.0
1.0
1.0
1.0
1.0
1.0
1.0
1.0
1.0
1.0
1.0
1.0
1.0


21
3
0
0
1
0
0
4
3
3
5
2
0
0
0
0
0
0
0
0
0
0
0
0
0
0
0
0
0
0

1.0
1.0
1.0
1.0
1.0
1.0
1.0
1.0
1.0
1.0
1.0
1.0
1.0
1.0
1.0
1.0
1.0
1.0
1.0
1.0
1.0
1.0
1.0
1.0
1.0
1.0
1.0
1.0
1.0
1.0


6
1
0
0
0
0
0
1
1
1
1
1
0
0
0
0
0
0
0
0
0
0
0
0
0
0
0
0
0
0

0.98
0.98
0.98
0.98
0.98
0.98
0.98
0.98
0.98
0.98
0.98
0.98
0.98
0.98
0.98
0.98
0.98
0.98
0.98
0.98
0.98
0.98
0.98
0.98
0.98
0.98
0.98
0.98
0.98
0.98


4
1
0
0
0
0
0
1
0
1
1
0
0
0
0
0
0
0
0
0
0
0
0
0
0
0
0
0
0
0

1.0
1.0
1.0
1.0
1.0
1.0
1.0
1.0
1.0
1.0
1.0
1.0
1.0
1.0
1.0
1.0
1.0
1.0
1.0
1.0
1.0
1.0
1.0
1.0
1.0
1.0
1.0
1.0
1.0
1.0


1
0
0
0
0
0
0
0
0
0
1
0
0
0
0
0
0
0
0
0
0
0
0
0
0
0
0
0
0
0

0.97
0.97
0.97
0.97
0.97
0.97
0.97
0.97
0.97
0.97
0.97
0.97
0.97
0.97
0.97
0.97
0.97
0.97
0.97
0.97
0.97
0.97
0.97
0.97
0.97
0.97
0.97
0.97
0.97
0.97


6
1
0
0
0
0
0
1
1
1
1
1
0
0
0
0
0
0
0
0
0
0
0
0
0
0
0
0
0
0


6
1
0
0
0
0
0
1
1
1
1
1
0
0
0
0
0
0
0
0
0
0
0
0
0
0
0
0
0
0

1.0
1.0
1.0
1.0
1.0
1.0
1.0
1.0
1.0
1.0
1.0
1.0
1.0
1.0
1.0
1.0
1.0
1.0
1.0
1.0
1.0
1.0
1.0
1.0
1.0
1.0
1.0
1.0
1.0
1.0


5
2
1
0
0
0
0
0
0
0
0
2
0
0
0
0
0
0
0
0
0
0
0
0
0
0
0
0
0
0


5
2
1
0
0
0
0
0
0
0
0
2
0
0
0
0
0
0
0
0
0
0
0
0
0
0
0
0
0
0

1.0
1.0
1.0
1.0
1.0
1.0
1.0
1.0
1.0
1.0
1.0
1.0
1.0
1.0
1.0
1.0
1.0
1.0
1.0
1.0
1.0
1.0
1.0
1.0
1.0
1.0
1.0
1.0
1.0
1.0


25
0
2
1
0
0
2
0
0
0
0
1
0
1
1
2
5
4
2
0
0
0
0
0
1
0
0
0
2
1

1.0
1.0
1.0
1.0
1.0
1.0
1.0
1.0
1.0
1.0
1.0
1.0
1.0
1.0
1.0
1.0
1.0
1.0
1.0
1.0
1.0
1.0
1.0
1.0
1.0
1.0
1.0
1.0
1.0
1.0


2
0
1
0
0
0
0
0
0
0
0
0
0
0
0
0
0
1
0
0
0
0
0
0
0
0
0
0
0
0

0.99
0.99
0.99
0.99
0.99
0.99
0.99
0.99
0.99
0.99
0.99
0.99
0.99
0.99
0.99
0.99
0.99
0.99
0.99
0.99
0.99
0.99
0.99
0.99
0.99
0.99
0.99
0.99
0.99
0.99


2
0
0
0
0
0
0
0
0
0
0
0
0
0
0
0
1
1
0
0
0
0
0
0
0
0
0
0
0
0

0.98
0.98
0.98
0.98
0.98
0.98
0.98
0.98
0.98
0.98
0.98
0.98
0.98
0.98
0.98
0.98
0.98
0.98
0.98
0.98
0.98
0.98
0.98
0.98
0.98
0.98
0.98
0.98
0.98
0.98


6
1
1
0
0
0
0
0
1
0
1
1
0
0
0
0
0
0
0
0
0
0
0
0
1
0
0
0
0
0

1.0
1.0
1.0
1.0
1.0
1.0
1.0
1.0
1.0
1.0
1.0
1.0
1.0
1.0
1.0
1.0
1.0
1.0
1.0
1.0
1.0
1.0
1.0
1.0
1.0
1.0
1.0
1.0
1.0
1.0


5
1
1
0
0
0
0
0
1
0
1
1
0
0
0
0
0
0
0
0
0
0
0
0
0
0
0
0
0
0

1.0
1.0
1.0
1.0
1.0
1.0
1.0
1.0
1.0
1.0
1.0
1.0
1.0
1.0
1.0
1.0
1.0
1.0
1.0
1.0
1.0
1.0
1.0
1.0
1.0
1.0
1.0
1.0
1.0
1.0


4
0
0
0
0
0
0
1
1
1
1
0
0
0
0
0
0
0
0
0
0
0
0
0
0
0
0
0
0
0

0.89
0.89
0.89
0.89
0.89
0.89
0.89
0.89
0.89
0.89
0.89
0.89
0.89
0.89
0.89
0.89
0.89
0.89
0.89
0.89
0.89
0.89
0.89
0.89
0.89
0.89
0.89
0.89
0.89
0.89


3
0
0
0
0
0
0
0
0
1
1
1
0
0
0
0
0
0
0
0
0
0
0
0
0
0
0
0
0
0


3
0
0
0
0
0
0
0
0
1
1
1
0
0
0
0
0
0
0
0
0
0
0
0
0
0
0
0
0
0

1.0
1.0
1.0
1.0
1.0
1.0
1.0
1.0
1.0
1.0
1.0
1.0
1.0
1.0
1.0
1.0
1.0
1.0
1.0
1.0
1.0
1.0
1.0
1.0
1.0
1.0
1.0
1.0
1.0
1.0


2
1
1
0
0
0
0
0
0
0
0
0
0
0
0
0
0
0
0
0
0
0
0
0
0
0
0
0
0
0


2
1
1
0
0
0
0
0
0
0
0
0
0
0
0
0
0
0
0
0
0
0
0
0
0
0
0
0
0
0

1.0
1.0
1.0
1.0
1.0
1.0
1.0
1.0
1.0
1.0
1.0
1.0
1.0
1.0
1.0
1.0
1.0
1.0
1.0
1.0
1.0
1.0
1.0
1.0
1.0
1.0
1.0
1.0
1.0
1.0


2
1
1
0
0
0
0
0
0
0
0
0
0
0
0
0
0
0
0
0
0
0
0
0
0
0
0
0
0
0


2
1
1
0
0
0
0
0
0
0
0
0
0
0
0
0
0
0
0
0
0
0
0
0
0
0
0
0
0
0

1.0
1.0
1.0
1.0
1.0
1.0
1.0
1.0
1.0
1.0
1.0
1.0
1.0
1.0
1.0
1.0
1.0
1.0
1.0
1.0
1.0
1.0
1.0
1.0
1.0
1.0
1.0
1.0
1.0
1.0


2
0
0
0
0
0
0
0
0
1
0
1
0
0
0
0
0
0
0
0
0
0
0
0
0
0
0
0
0
0


2
0
0
0
0
0
0
0
0
1
0
1
0
0
0
0
0
0
0
0
0
0
0
0
0
0
0
0
0
0

1.0
1.0
1.0
1.0
1.0
1.0
1.0
1.0
1.0
1.0
1.0
1.0
1.0
1.0
1.0
1.0
1.0
1.0
1.0
1.0
1.0
1.0
1.0
1.0
1.0
1.0
1.0
1.0
1.0
1.0


3
0
0
0
0
0
0
0
0
0
0
0
0
1
0
0
1
0
1
0
0
0
0
0
0
0
0
0
0
0

1.0
1.0
1.0
1.0
1.0
1.0
1.0
1.0
1.0
1.0
1.0
1.0
1.0
1.0
1.0
1.0
1.0
1.0
1.0
1.0
1.0
1.0
1.0
1.0
1.0
1.0
1.0
1.0
1.0
1.0


7
0
0
0
0
0
0
0
0
0
0
0
0
0
0
3
0
3
0
1
0
0
0
0
0
0
0
0
0
0

0.71
0.71
0.71
0.71
0.71
0.71
0.71
0.71
0.71
0.71
0.71
0.71
0.71
0.71
0.71
0.71
0.71
0.71
0.71
0.71
0.71
0.71
0.71
0.71
0.71
0.71
0.71
0.71
0.71
0.71


4
0
0
0
0
0
0
0
0
0
0
0
0
0
0
2
0
2
0
0
0
0
0
0
0
0
0
0
0
0

0.96
0.96
0.96
0.96
0.96
0.96
0.96
0.96
0.96
0.96
0.96
0.96
0.96
0.96
0.96
0.96
0.96
0.96
0.96
0.96
0.96
0.96
0.96
0.96
0.96
0.96
0.96
0.96
0.96
0.96


1
0
0
0
0
0
0
0
0
0
0
0
0
0
0
0
0
1
0
0
0
0
0
0
0
0
0
0
0
0

0.91
0.91
0.91
0.91
0.91
0.91
0.91
0.91
0.91
0.91
0.91
0.91
0.91
0.91
0.91
0.91
0.91
0.91
0.91
0.91
0.91
0.91
0.91
0.91
0.91
0.91
0.91
0.91
0.91
0.91


1
0
0
0
0
0
0
0
0
0
0
0
0
0
0
0
0
0
0
1
0
0
0
0
0
0
0
0
0
0

0.96
0.96
0.96
0.96
0.96
0.96
0.96
0.96
0.96
0.96
0.96
0.96
0.96
0.96
0.96
0.96
0.96
0.96
0.96
0.96
0.96
0.96
0.96
0.96
0.96
0.96
0.96
0.96
0.96
0.96


1
0
0
0
0
0
0
0
0
0
0
0
0
0
0
0
0
1
0
0
0
0
0
0
0
0
0
0
0
0


1
0
0
0
0
0
0
0
0
0
0
0
0
0
0
0
0
1
0
0
0
0
0
0
0
0
0
0
0
0

1.0
1.0
1.0
1.0
1.0
1.0
1.0
1.0
1.0
1.0
1.0
1.0
1.0
1.0
1.0
1.0
1.0
1.0
1.0
1.0
1.0
1.0
1.0
1.0
1.0
1.0
1.0
1.0
1.0
1.0


3
0
0
0
0
0
0
0
0
0
0
0
0
0
0
0
0
1
0
0
0
0
0
0
0
1
0
0
0
1

1.0
1.0
1.0
1.0
1.0
1.0
1.0
1.0
1.0
1.0
1.0
1.0
1.0
1.0
1.0
1.0
1.0
1.0
1.0
1.0
1.0
1.0
1.0
1.0
1.0
1.0
1.0
1.0
1.0
1.0


3
0
0
0
0
0
0
0
0
0
0
0
0
1
0
0
0
0
0
1
1
0
0
0
0
0
0
0
0
0

1.0
1.0
1.0
1.0
1.0
1.0
1.0
1.0
1.0
1.0
1.0
1.0
1.0
1.0
1.0
1.0
1.0
1.0
1.0
1.0
1.0
1.0
1.0
1.0
1.0
1.0
1.0
1.0
1.0
1.0


3
0
0
0
0
0
0
0
0
0
0
0
0
0
0
0
0
0
0
0
0
0
0
0
0
1
0
0
0
2

1.0
1.0
1.0
1.0
1.0
1.0
1.0
1.0
1.0
1.0
1.0
1.0
1.0
1.0
1.0
1.0
1.0
1.0
1.0
1.0
1.0
1.0
1.0
1.0
1.0
1.0
1.0
1.0
1.0
1.0


1
0
0
0
0
0
0
0
0
0
0
0
0
0
0
0
0
0
0
0
0
0
0
0
0
1
0
0
0
0

1.0
1.0
1.0
1.0
1.0
1.0
1.0
1.0
1.0
1.0
1.0
1.0
1.0
1.0
1.0
1.0
1.0
1.0
1.0
1.0
1.0
1.0
1.0
1.0
1.0
1.0
1.0
1.0
1.0
1.0


1
0
0
0
0
0
0
0
0
0
0
0
0
0
0
0
0
0
0
0
0
0
0
0
1
0
0
0
0
0


1
0
0
0
0
0
0
0
0
0
0
0
0
0
0
0
0
0
0
0
0
0
0
0
1
0
0
0
0
0

1.0
1.0
1.0
1.0
1.0
1.0
1.0
1.0
1.0
1.0
1.0
1.0
1.0
1.0
1.0
1.0
1.0
1.0
1.0
1.0
1.0
1.0
1.0
1.0
1.0
1.0
1.0
1.0
1.0
1.0


2
0
0
0
0
0
0
0
0
0
0
0
0
0
0
1
0
0
0
0
0
0
0
0
1
0
0
0
0
0


1
0
0
0
0
0
0
0
0
0
0
0
0
0
0
1
0
0
0
0
0
0
0
0
0
0
0
0
0
0

0.99
0.99
0.99
0.99
0.99
0.99
0.99
0.99
0.99
0.99
0.99
0.99
0.99
0.99
0.99
0.99
0.99
0.99
0.99
0.99
0.99
0.99
0.99
0.99
0.99
0.99
0.99
0.99
0.99
0.99


1
0
0
0
0
0
0
0
0
0
0
0
0
0
0
0
0
0
0
0
0
0
0
0
1
0
0
0
0
0

1.0
1.0
1.0
1.0
1.0
1.0
1.0
1.0
1.0
1.0
1.0
1.0
1.0
1.0
1.0
1.0
1.0
1.0
1.0
1.0
1.0
1.0
1.0
1.0
1.0
1.0
1.0
1.0
1.0
1.0


1
0
0
0
0
0
0
0
0
0
0
0
0
0
1
0
0
0
0
0
0
0
0
0
0
0
0
0
0
0

0.97
0.97
0.97
0.97
0.97
0.97
0.97
0.97
0.97
0.97
0.97
0.97
0.97
0.97
0.97
0.97
0.97
0.97
0.97
0.97
0.97
0.97
0.97
0.97
0.97
0.97
0.97
0.97
0.97
0.97


2
0
0
0
0
0
0
0
0
0
0
0
0
0
0
0
0
0
0
0
0
0
0
0
0
0
1
0
0
1

1.0
1.0
1.0
1.0
1.0
1.0
1.0
1.0
1.0
1.0
1.0
1.0
1.0
1.0
1.0
1.0
1.0
1.0
1.0
1.0
1.0
1.0
1.0
1.0
1.0
1.0
1.0
1.0
1.0
1.0


1
0
0
0
0
0
0
0
0
0
0
0
0
0
0
0
0
0
0
0
0
0
1
0
0
0
0
0
0
0


1
0
0
0
0
0
0
0
0
0
0
0
0
0
0
0
0
0
0
0
0
0
1
0
0
0
0
0
0
0

1.0
1.0
1.0
1.0
1.0
1.0
1.0
1.0
1.0
1.0
1.0
1.0
1.0
1.0
1.0
1.0
1.0
1.0
1.0
1.0
1.0
1.0
1.0
1.0
1.0
1.0
1.0
1.0
1.0
1.0


1
0
0
0
0
0
0
0
0
0
0
0
0
0
0
0
0
0
0
0
0
0
0
0
1
0
0
0
0
0


1
0
0
0
0
0
0
0
0
0
0
0
0
0
0
0
0
0
0
0
0
0
0
0
1
0
0
0
0
0

1.0
1.0
1.0
1.0
1.0
1.0
1.0
1.0
1.0
1.0
1.0
1.0
1.0
1.0
1.0
1.0
1.0
1.0
1.0
1.0
1.0
1.0
1.0
1.0
1.0
1.0
1.0
1.0
1.0
1.0


1
0
0
0
0
0
0
0
0
0
0
0
0
0
0
0
0
0
0
1
0
0
0
0
0
0
0
0
0
0


1
0
0
0
0
0
0
0
0
0
0
0
0
0
0
0
0
0
0
1
0
0
0
0
0
0
0
0
0
0

0.99
0.99
0.99
0.99
0.99
0.99
0.99
0.99
0.99
0.99
0.99
0.99
0.99
0.99
0.99
0.99
0.99
0.99
0.99
0.99
0.99
0.99
0.99
0.99
0.99
0.99
0.99
0.99
0.99
0.99


2
0
0
0
0
0
0
0
0
0
0
0
0
0
0
0
0
0
0
0
0
0
0
0
0
1
1
0
0
0

0.99
0.99
0.99
0.99
0.99
0.99
0.99
0.99
0.99
0.99
0.99
0.99
0.99
0.99
0.99
0.99
0.99
0.99
0.99
0.99
0.99
0.99
0.99
0.99
0.99
0.99
0.99
0.99
0.99
0.99


6
2
1
0
0
0
2
0
0
0
0
0
0
0
0
0
0
1
0
0
0
0
0
0
0
0
0
0
0
0


5
2
1
0
0
0
2
0
0
0
0
0
0
0
0
0
0
0
0
0
0
0
0
0
0
0
0
0
0
0

1.0
1.0
1.0
1.0
1.0
1.0
1.0
1.0
1.0
1.0
1.0
1.0
1.0
1.0
1.0
1.0
1.0
1.0
1.0
1.0
1.0
1.0
1.0
1.0
1.0
1.0
1.0
1.0
1.0
1.0


2
0
0
0
0
0
2
0
0
0
0
0
0
0
0
0
0
0
0
0
0
0
0
0
0
0
0
0
0
0

1.0
1.0
1.0
1.0
1.0
1.0
1.0
1.0
1.0
1.0
1.0
1.0
1.0
1.0
1.0
1.0
1.0
1.0
1.0
1.0
1.0
1.0
1.0
1.0
1.0
1.0
1.0
1.0
1.0
1.0


2
1
1
0
0
0
0
0
0
0
0
0
0
0
0
0
0
0
0
0
0
0
0
0
0
0
0
0
0
0

0.98
0.98
0.98
0.98
0.98
0.98
0.98
0.98
0.98
0.98
0.98
0.98
0.98
0.98
0.98
0.98
0.98
0.98
0.98
0.98
0.98
0.98
0.98
0.98
0.98
0.98
0.98
0.98
0.98
0.98


1
0
0
0
0
0
0
0
0
0
0
0
0
0
0
0
0
1
0
0
0
0
0
0
0
0
0
0
0
0


1
0
0
0
0
0
0
0
0
0
0
0
0
0
0
0
0
1
0
0
0
0
0
0
0
0
0
0
0
0

1.0
1.0
1.0
1.0
1.0
1.0
1.0
1.0
1.0
1.0
1.0
1.0
1.0
1.0
1.0
1.0
1.0
1.0
1.0
1.0
1.0
1.0
1.0
1.0
1.0
1.0
1.0
1.0
1.0
1.0


11
0
0
0
1
3
1
0
0
0
0
0
0
0
0
0
2
0
0
0
0
0
0
0
1
0
1
1
1
0


3
0
0
0
1
1
1
0
0
0
0
0
0
0
0
0
0
0
0
0
0
0
0
0
0
0
0
0
0
0

1.0
1.0
1.0
1.0
1.0
1.0
1.0
1.0
1.0
1.0
1.0
1.0
1.0
1.0
1.0
1.0
1.0
1.0
1.0
1.0
1.0
1.0
1.0
1.0
1.0
1.0
1.0
1.0
1.0
1.0


2
0
0
0
1
0
1
0
0
0
0
0
0
0
0
0
0
0
0
0
0
0
0
0
0
0
0
0
0
0

1.0
1.0
1.0
1.0
1.0
1.0
1.0
1.0
1.0
1.0
1.0
1.0
1.0
1.0
1.0
1.0
1.0
1.0
1.0
1.0
1.0
1.0
1.0
1.0
1.0
1.0
1.0
1.0
1.0
1.0


2
0
0
0
0
2
0
0
0
0
0
0
0
0
0
0
0
0
0
0
0
0
0
0
0
0
0
0
0
0


1
0
0
0
0
1
0
0
0
0
0
0
0
0
0
0
0
0
0
0
0
0
0
0
0
0
0
0
0
0

0.72
0.72
0.72
0.72
0.72
0.72
0.72
0.72
0.72
0.72
0.72
0.72
0.72
0.72
0.72
0.72
0.72
0.72
0.72
0.72
0.72
0.72
0.72
0.72
0.72
0.72
0.72
0.72
0.72
0.72


1
0
0
0
0
1
0
0
0
0
0
0
0
0
0
0
0
0
0
0
0
0
0
0
0
0
0
0
0
0

1.0
1.0
1.0
1.0
1.0
1.0
1.0
1.0
1.0
1.0
1.0
1.0
1.0
1.0
1.0
1.0
1.0
1.0
1.0
1.0
1.0
1.0
1.0
1.0
1.0
1.0
1.0
1.0
1.0
1.0


2
0
0
0
0
0
0
0
0
0
0
0
0
0
0
0
1
0
0
0
0
0
0
0
0
0
0
1
0
0

1.0
1.0
1.0
1.0
1.0
1.0
1.0
1.0
1.0
1.0
1.0
1.0
1.0
1.0
1.0
1.0
1.0
1.0
1.0
1.0
1.0
1.0
1.0
1.0
1.0
1.0
1.0
1.0
1.0
1.0


2
0
0
0
0
0
0
0
0
0
0
0
0
0
0
0
1
0
0
0
0
0
0
0
1
0
0
0
0
0


2
0
0
0
0
0
0
0
0
0
0
0
0
0
0
0
1
0
0
0
0
0
0
0
1
0
0
0
0
0

1.0
1.0
1.0
1.0
1.0
1.0
1.0
1.0
1.0
1.0
1.0
1.0
1.0
1.0
1.0
1.0
1.0
1.0
1.0
1.0
1.0
1.0
1.0
1.0
1.0
1.0
1.0
1.0
1.0
1.0


2
0
0
0
0
0
0
0
0
0
0
0
0
0
0
0
0
0
0
0
0
0
0
0
0
0
1
0
1
0


2
0
0
0
0
0
0
0
0
0
0
0
0
0
0
0
0
0
0
0
0
0
0
0
0
0
1
0
1
0

1.0
1.0
1.0
1.0
1.0
1.0
1.0
1.0
1.0
1.0
1.0
1.0
1.0
1.0
1.0
1.0
1.0
1.0
1.0
1.0
1.0
1.0
1.0
1.0
1.0
1.0
1.0
1.0
1.0
1.0


14
0
2
1
0
0
2
0
0
0
0
1
0
0
0
0
1
0
1
0
0
0
0
0
0
1
0
3
0
2


1
0
0
0
0
0
1
0
0
0
0
0
0
0
0
0
0
0
0
0
0
0
0
0
0
0
0
0
0
0

1.0
1.0
1.0
1.0
1.0
1.0
1.0
1.0
1.0
1.0
1.0
1.0
1.0
1.0
1.0
1.0
1.0
1.0
1.0
1.0
1.0
1.0
1.0
1.0
1.0
1.0
1.0
1.0
1.0
1.0


13
0
2
1
0
0
1
0
0
0
0
1
0
0
0
0
1
0
1
0
0
0
0
0
0
1
0
3
0
2

1.0
1.0
1.0
1.0
1.0
1.0
1.0
1.0
1.0
1.0
1.0
1.0
1.0
1.0
1.0
1.0
1.0
1.0
1.0
1.0
1.0
1.0
1.0
1.0
1.0
1.0
1.0
1.0
1.0
1.0


1
0
1
0
0
0
0
0
0
0
0
0
0
0
0
0
0
0
0
0
0
0
0
0
0
0
0
0
0
0

1.0
1.0
1.0
1.0
1.0
1.0
1.0
1.0
1.0
1.0
1.0
1.0
1.0
1.0
1.0
1.0
1.0
1.0
1.0
1.0
1.0
1.0
1.0
1.0
1.0
1.0
1.0
1.0
1.0
1.0


1
0
0
0
0
0
1
0
0
0
0
0
0
0
0
0
0
0
0
0
0
0
0
0
0
0
0
0
0
0

1.0
1.0
1.0
1.0
1.0
1.0
1.0
1.0
1.0
1.0
1.0
1.0
1.0
1.0
1.0
1.0
1.0
1.0
1.0
1.0
1.0
1.0
1.0
1.0
1.0
1.0
1.0
1.0
1.0
1.0


2
0
0
1
0
0
0
0
0
0
0
0
0
0
0
0
0
0
0
0
0
0
0
0
0
0
0
0
0
1

1.0
1.0
1.0
1.0
1.0
1.0
1.0
1.0
1.0
1.0
1.0
1.0
1.0
1.0
1.0
1.0
1.0
1.0
1.0
1.0
1.0
1.0
1.0
1.0
1.0
1.0
1.0
1.0
1.0
1.0


6
0
1
0
0
0
0
0
0
0
0
0
0
0
0
0
1
0
1
0
0
0
0
0
0
1
0
2
0
0

1.0
1.0
1.0
1.0
1.0
1.0
1.0
1.0
1.0
1.0
1.0
1.0
1.0
1.0
1.0
1.0
1.0
1.0
1.0
1.0
1.0
1.0
1.0
1.0
1.0
1.0
1.0
1.0
1.0
1.0


92
5
4
5
4
7
4
0
0
1
3
2
0
0
2
1
1
0
2
0
0
0
0
0
12
12
6
7
6
8

1.0
1.0
1.0
1.0
1.0
1.0
1.0
1.0
1.0
1.0
1.0
1.0
1.0
1.0
1.0
1.0
1.0
1.0
1.0
1.0
1.0
1.0
1.0
1.0
1.0
1.0
1.0
1.0
1.0
1.0


18
2
1
3
2
2
1
0
0
0
0
0
0
0
2
0
0
0
0
0
0
0
0
0
3
0
1
0
1
0

1.0
1.0
1.0
1.0
1.0
1.0
1.0
1.0
1.0
1.0
1.0
1.0
1.0
1.0
1.0
1.0
1.0
1.0
1.0
1.0
1.0
1.0
1.0
1.0
1.0
1.0
1.0
1.0
1.0
1.0


4
1
0
1
0
0
0
0
0
0
0
0
0
0
2
0
0
0
0
0
0
0
0
0
0
0
0
0
0
0

1.0
1.0
1.0
1.0
1.0
1.0
1.0
1.0
1.0
1.0
1.0
1.0
1.0
1.0
1.0
1.0
1.0
1.0
1.0
1.0
1.0
1.0
1.0
1.0
1.0
1.0
1.0
1.0
1.0
1.0


35
3
3
1
0
3
2
0
0
1
3
2
0
0
0
1
1
0
2
0
0
0
0
0
5
0
1
2
1
4

1.0
1.0
1.0
1.0
1.0
1.0
1.0
1.0
1.0
1.0
1.0
1.0
1.0
1.0
1.0
1.0
1.0
1.0
1.0
1.0
1.0
1.0
1.0
1.0
1.0
1.0
1.0
1.0
1.0
1.0


25
0
0
0
1
1
1
0
0
0
0
0
0
0
0
0
0
0
0
0
0
0
0
0
3
10
3
2
2
2

1.0
1.0
1.0
1.0
1.0
1.0
1.0
1.0
1.0
1.0
1.0
1.0
1.0
1.0
1.0
1.0
1.0
1.0
1.0
1.0
1.0
1.0
1.0
1.0
1.0
1.0
1.0
1.0
1.0
1.0


3
0
0
0
0
0
0
0
0
0
0
0
0
0
0
0
0
0
0
0
0
0
0
0
0
3
0
0
0
0

1.0
1.0
1.0
1.0
1.0
1.0
1.0
1.0
1.0
1.0
1.0
1.0
1.0
1.0
1.0
1.0
1.0
1.0
1.0
1.0
1.0
1.0
1.0
1.0
1.0
1.0
1.0
1.0
1.0
1.0


4
0
0
0
0
0
0
0
0
0
0
0
0
0
0
0
0
0
0
0
0
0
0
0
0
1
0
1
1
1

0.84
0.84
0.84
0.84
0.84
0.84
0.84
0.84
0.84
0.84
0.84
0.84
0.84
0.84
0.84
0.84
0.84
0.84
0.84
0.84
0.84
0.84
0.84
0.84
0.84
0.84
0.84
0.84
0.84
0.84


2
0
0
0
0
0
0
0
0
0
0
0
0
0
0
0
0
0
0
0
0
0
0
0
0
1
0
0
0
1


2
0
0
0
0
0
0
0
0
0
0
0
0
0
0
0
0
0
0
0
0
0
0
0
0
1
0
0
0
1

1.0
1.0
1.0
1.0
1.0
1.0
1.0
1.0
1.0
1.0
1.0
1.0
1.0
1.0
1.0
1.0
1.0
1.0
1.0
1.0
1.0
1.0
1.0
1.0
1.0
1.0
1.0
1.0
1.0
1.0


1
0
0
0
0
0
0
0
0
0
0
1
0
0
0
0
0
0
0
0
0
0
0
0
0
0
0
0
0
0


1
0
0
0
0
0
0
0
0
0
0
1
0
0
0
0
0
0
0
0
0
0
0
0
0
0
0
0
0
0


1
0
0
0
0
0
0
0
0
0
0
1
0
0
0
0
0
0
0
0
0
0
0
0
0
0
0
0
0
0

1.0
1.0
1.0
1.0
1.0
1.0
1.0
1.0
1.0
1.0
1.0
1.0
1.0
1.0
1.0
1.0
1.0
1.0
1.0
1.0
1.0
1.0
1.0
1.0
1.0
1.0
1.0
1.0
1.0
1.0


4
0
1
0
1
0
1
0
0
0
0
1
0
0
0
0
0
0
0
0
0
0
0
0
0
0
0
0
0
0


4
0
1
0
1
0
1
0
0
0
0
1
0
0
0
0
0
0
0
0
0
0
0
0
0
0
0
0
0
0


4
0
1
0
1
0
1
0
0
0
0
1
0
0
0
0
0
0
0
0
0
0
0
0
0
0
0
0
0
0

1.0
1.0
1.0
1.0
1.0
1.0
1.0
1.0
1.0
1.0
1.0
1.0
1.0
1.0
1.0
1.0
1.0
1.0
1.0
1.0
1.0
1.0
1.0
1.0
1.0
1.0
1.0
1.0
1.0
1.0


8
1
1
1
0
0
0
1
1
1
1
1
0
0
0
0
0
0
0
0
0
0
0
0
0
0
0
0
0
0

0.96
0.96
0.96
0.96
0.96
0.96
0.96
0.96
0.96
0.96
0.96
0.96
0.96
0.96
0.96
0.96
0.96
0.96
0.96
0.96
0.96
0.96
0.96
0.96
0.96
0.96
0.96
0.96
0.96
0.96


7
1
1
1
0
0
0
0
1
1
1
1
0
0
0
0
0
0
0
0
0
0
0
0
0
0
0
0
0
0

1.0
1.0
1.0
1.0
1.0
1.0
1.0
1.0
1.0
1.0
1.0
1.0
1.0
1.0
1.0
1.0
1.0
1.0
1.0
1.0
1.0
1.0
1.0
1.0
1.0
1.0
1.0
1.0
1.0
1.0


43
0
1
0
0
2
3
0
0
0
0
0
0
0
0
7
0
3
4
4
0
0
0
0
1
4
5
4
1
4

0.98
0.98
0.98
0.98
0.98
0.98
0.98
0.98
0.98
0.98
0.98
0.98
0.98
0.98
0.98
0.98
0.98
0.98
0.98
0.98
0.98
0.98
0.98
0.98
0.98
0.98
0.98
0.98
0.98
0.98


6
0
0
0
0
1
3
0
0
0
0
0
0
0
0
1
0
0
0
1
0
0
0
0
0
0
0
0
0
0

1.0
1.0
1.0
1.0
1.0
1.0
1.0
1.0
1.0
1.0
1.0
1.0
1.0
1.0
1.0
1.0
1.0
1.0
1.0
1.0
1.0
1.0
1.0
1.0
1.0
1.0
1.0
1.0
1.0
1.0


3
0
1
0
0
1
0
0
0
0
0
0
0
0
0
1
0
0
0
0
0
0
0
0
0
0
0
0
0
0

1.0
1.0
1.0
1.0
1.0
1.0
1.0
1.0
1.0
1.0
1.0
1.0
1.0
1.0
1.0
1.0
1.0
1.0
1.0
1.0
1.0
1.0
1.0
1.0
1.0
1.0
1.0
1.0
1.0
1.0


2
0
1
0
0
1
0
0
0
0
0
0
0
0
0
0
0
0
0
0
0
0
0
0
0
0
0
0
0
0

1.0
1.0
1.0
1.0
1.0
1.0
1.0
1.0
1.0
1.0
1.0
1.0
1.0
1.0
1.0
1.0
1.0
1.0
1.0
1.0
1.0
1.0
1.0
1.0
1.0
1.0
1.0
1.0
1.0
1.0


2
0
0
0
0
0
0
0
0
0
0
0
0
0
0
0
0
0
0
1
0
0
0
0
0
0
0
0
0
1

0.8
0.8
0.8
0.8
0.8
0.8
0.8
0.8
0.8
0.8
0.8
0.8
0.8
0.8
0.8
0.8
0.8
0.8
0.8
0.8
0.8
0.8
0.8
0.8
0.8
0.8
0.8
0.8
0.8
0.8


1
0
0
0
0
0
0
0
0
0
0
0
0
0
0
0
0
0
0
0
0
0
0
0
0
0
0
0
0
1

1.0
1.0
1.0
1.0
1.0
1.0
1.0
1.0
1.0
1.0
1.0
1.0
1.0
1.0
1.0
1.0
1.0
1.0
1.0
1.0
1.0
1.0
1.0
1.0
1.0
1.0
1.0
1.0
1.0
1.0


9
0
0
0
0
0
0
1
1
2
3
2
0
0
0
0
0
0
0
0
0
0
0
0
0
0
0
0
0
0


2
0
0
0
0
0
0
1
0
0
1
0
0
0
0
0
0
0
0
0
0
0
0
0
0
0
0
0
0
0

0.98
0.98
0.98
0.98
0.98
0.98
0.98
0.98
0.98
0.98
0.98
0.98
0.98
0.98
0.98
0.98
0.98
0.98
0.98
0.98
0.98
0.98
0.98
0.98
0.98
0.98
0.98
0.98
0.98
0.98


7
0
0
0
0
0
0
0
1
2
2
2
0
0
0
0
0
0
0
0
0
0
0
0
0
0
0
0
0
0


7
0
0
0
0
0
0
0
1
2
2
2
0
0
0
0
0
0
0
0
0
0
0
0
0
0
0
0
0
0

1.0
1.0
1.0
1.0
1.0
1.0
1.0
1.0
1.0
1.0
1.0
1.0
1.0
1.0
1.0
1.0
1.0
1.0
1.0
1.0
1.0
1.0
1.0
1.0
1.0
1.0
1.0
1.0
1.0
1.0


4
0
0
1
0
0
0
0
0
0
0
0
0
0
0
0
0
0
0
0
0
0
0
0
3
0
0
0
0
0

0.72
0.72
0.72
0.72
0.72
0.72
0.72
0.72
0.72
0.72
0.72
0.72
0.72
0.72
0.72
0.72
0.72
0.72
0.72
0.72
0.72
0.72
0.72
0.72
0.72
0.72
0.72
0.72
0.72
0.72


2
0
0
1
0
0
0
0
0
0
0
0
0
0
0
0
0
0
0
0
0
0
0
0
1
0
0
0
0
0


2
0
0
1
0
0
0
0
0
0
0
0
0
0
0
0
0
0
0
0
0
0
0
0
1
0
0
0
0
0

1.0
1.0
1.0
1.0
1.0
1.0
1.0
1.0
1.0
1.0
1.0
1.0
1.0
1.0
1.0
1.0
1.0
1.0
1.0
1.0
1.0
1.0
1.0
1.0
1.0
1.0
1.0
1.0
1.0
1.0


1
0
0
0
0
0
0
0
0
0
0
0
0
0
0
0
0
0
0
0
0
0
0
0
1
0
0
0
0
0


1
0
0
0
0
0
0
0
0
0
0
0
0
0
0
0
0
0
0
0
0
0
0
0
1
0
0
0
0
0

1.0
1.0
1.0
1.0
1.0
1.0
1.0
1.0
1.0
1.0
1.0
1.0
1.0
1.0
1.0
1.0
1.0
1.0
1.0
1.0
1.0
1.0
1.0
1.0
1.0
1.0
1.0
1.0
1.0
1.0


27
0
0
0
0
0
1
0
0
0
0
0
0
0
0
0
0
0
0
0
0
0
0
0
3
5
4
5
4
5

0.83
0.83
0.83
0.83
0.83
0.83
0.83
0.83
0.83
0.83
0.83
0.83
0.83
0.83
0.83
0.83
0.83
0.83
0.83
0.83
0.83
0.83
0.83
0.83
0.83
0.83
0.83
0.83
0.83
0.83


21
0
0
0
0
0
0
0
0
0
0
0
0
0
0
0
0
0
0
0
0
0
0
0
0
5
4
5
2
5


21
0
0
0
0
0
0
0
0
0
0
0
0
0
0
0
0
0
0
0
0
0
0
0
0
5
4
5
2
5

1.0
1.0
1.0
1.0
1.0
1.0
1.0
1.0
1.0
1.0
1.0
1.0
1.0
1.0
1.0
1.0
1.0
1.0
1.0
1.0
1.0
1.0
1.0
1.0
1.0
1.0
1.0
1.0
1.0
1.0


1
0
0
0
0
0
0
0
0
0
0
0
0
0
0
0
0
0
0
0
0
0
0
0
1
0
0
0
0
0

0.99
0.99
0.99
0.99
0.99
0.99
0.99
0.99
0.99
0.99
0.99
0.99
0.99
0.99
0.99
0.99
0.99
0.99
0.99
0.99
0.99
0.99
0.99
0.99
0.99
0.99
0.99
0.99
0.99
0.99


40
1
0
0
0
1
0
1
1
1
0
1
0
0
0
2
0
0
1
0
0
0
0
0
15
1
5
0
6
4

1.0
1.0
1.0
1.0
1.0
1.0
1.0
1.0
1.0
1.0
1.0
1.0
1.0
1.0
1.0
1.0
1.0
1.0
1.0
1.0
1.0
1.0
1.0
1.0
1.0
1.0
1.0
1.0
1.0
1.0


13
1
0
0
0
1
0
1
1
0
0
1
0
0
0
0
0
0
0
0
0
0
0
0
0
0
3
0
4
1

1.0
1.0
1.0
1.0
1.0
1.0
1.0
1.0
1.0
1.0
1.0
1.0
1.0
1.0
1.0
1.0
1.0
1.0
1.0
1.0
1.0
1.0
1.0
1.0
1.0
1.0
1.0
1.0
1.0
1.0


3
1
0
0
0
0
0
1
1
0
0
0
0
0
0
0
0
0
0
0
0
0
0
0
0
0
0
0
0
0

1.0
1.0
1.0
1.0
1.0
1.0
1.0
1.0
1.0
1.0
1.0
1.0
1.0
1.0
1.0
1.0
1.0
1.0
1.0
1.0
1.0
1.0
1.0
1.0
1.0
1.0
1.0
1.0
1.0
1.0


2
0
0
0
0
0
0
0
0
0
0
0
0
0
0
0
0
0
0
0
0
0
0
0
0
0
1
0
1
0

0.8
0.8
0.8
0.8
0.8
0.8
0.8
0.8
0.8
0.8
0.8
0.8
0.8
0.8
0.8
0.8
0.8
0.8
0.8
0.8
0.8
0.8
0.8
0.8
0.8
0.8
0.8
0.8
0.8
0.8


7
0
0
0
0
0
0
0
0
0
0
0
0
0
0
2
0
0
0
0
0
0
0
0
0
0
2
0
1
2


7
0
0
0
0
0
0
0
0
0
0
0
0
0
0
2
0
0
0
0
0
0
0
0
0
0
2
0
1
2

1.0
1.0
1.0
1.0
1.0
1.0
1.0
1.0
1.0
1.0
1.0
1.0
1.0
1.0
1.0
1.0
1.0
1.0
1.0
1.0
1.0
1.0
1.0
1.0
1.0
1.0
1.0
1.0
1.0
1.0


6
0
0
0
0
0
0
0
0
0
0
0
0
0
0
0
0
0
0
0
0
0
0
0
6
0
0
0
0
0


6
0
0
0
0
0
0
0
0
0
0
0
0
0
0
0
0
0
0
0
0
0
0
0
6
0
0
0
0
0

1.0
1.0
1.0
1.0
1.0
1.0
1.0
1.0
1.0
1.0
1.0
1.0
1.0
1.0
1.0
1.0
1.0
1.0
1.0
1.0
1.0
1.0
1.0
1.0
1.0
1.0
1.0
1.0
1.0
1.0


4
0
0
0
0
0
0
0
0
0
0
0
0
0
0
0
0
0
0
0
0
0
0
0
4
0
0
0
0
0


4
0
0
0
0
0
0
0
0
0
0
0
0
0
0
0
0
0
0
0
0
0
0
0
4
0
0
0
0
0

0.86
0.86
0.86
0.86
0.86
0.86
0.86
0.86
0.86
0.86
0.86
0.86
0.86
0.86
0.86
0.86
0.86
0.86
0.86
0.86
0.86
0.86
0.86
0.86
0.86
0.86
0.86
0.86
0.86
0.86


12
0
0
0
0
0
1
0
0
0
0
0
0
0
0
0
0
0
1
1
0
0
0
0
1
2
1
2
0
3

0.71
0.71
0.71
0.71
0.71
0.71
0.71
0.71
0.71
0.71
0.71
0.71
0.71
0.71
0.71
0.71
0.71
0.71
0.71
0.71
0.71
0.71
0.71
0.71
0.71
0.71
0.71
0.71
0.71
0.71


2
0
0
0
0
0
1
0
0
0
0
0
0
0
0
0
0
0
0
0
0
0
0
0
0
0
0
1
0
0


1
0
0
0
0
0
1
0
0
0
0
0
0
0
0
0
0
0
0
0
0
0
0
0
0
0
0
0
0
0

0.92
0.92
0.92
0.92
0.92
0.92
0.92
0.92
0.92
0.92
0.92
0.92
0.92
0.92
0.92
0.92
0.92
0.92
0.92
0.92
0.92
0.92
0.92
0.92
0.92
0.92
0.92
0.92
0.92
0.92


1
0
0
0
0
0
0
0
0
0
0
0
0
0
0
0
0
0
0
0
0
0
0
0
0
0
0
1
0
0

0.85
0.85
0.85
0.85
0.85
0.85
0.85
0.85
0.85
0.85
0.85
0.85
0.85
0.85
0.85
0.85
0.85
0.85
0.85
0.85
0.85
0.85
0.85
0.85
0.85
0.85
0.85
0.85
0.85
0.85


6
0
0
0
0
0
0
0
0
0
0
0
0
0
0
0
0
0
0
1
0
0
0
0
1
1
0
1
0
2

0.94
0.94
0.94
0.94
0.94
0.94
0.94
0.94
0.94
0.94
0.94
0.94
0.94
0.94
0.94
0.94
0.94
0.94
0.94
0.94
0.94
0.94
0.94
0.94
0.94
0.94
0.94
0.94
0.94
0.94


1
0
0
0
0
0
0
0
0
0
0
0
0
0
0
0
0
0
0
0
0
0
0
0
1
0
0
0
0
0

0.97
0.97
0.97
0.97
0.97
0.97
0.97
0.97
0.97
0.97
0.97
0.97
0.97
0.97
0.97
0.97
0.97
0.97
0.97
0.97
0.97
0.97
0.97
0.97
0.97
0.97
0.97
0.97
0.97
0.97


3
0
0
0
0
0
0
0
0
0
0
0
0
0
0
0
0
0
1
0
0
0
0
0
0
0
1
0
0
1


3
0
0
0
0
0
0
0
0
0
0
0
0
0
0
0
0
0
1
0
0
0
0
0
0
0
1
0
0
1

1.0
1.0
1.0
1.0
1.0
1.0
1.0
1.0
1.0
1.0
1.0
1.0
1.0
1.0
1.0
1.0
1.0
1.0
1.0
1.0
1.0
1.0
1.0
1.0
1.0
1.0
1.0
1.0
1.0
1.0


69
0
0
0
2
2
2
0
0
0
0
0
0
0
1
1
3
1
2
1
0
0
0
0
12
11
8
4
7
12

1.0
1.0
1.0
1.0
1.0
1.0
1.0
1.0
1.0
1.0
1.0
1.0
1.0
1.0
1.0
1.0
1.0
1.0
1.0
1.0
1.0
1.0
1.0
1.0
1.0
1.0
1.0
1.0
1.0
1.0


10
0
0
0
2
2
2
0
0
0
0
0
0
0
0
0
0
0
0
0
0
0
0
0
1
0
0
0
2
1

1.0
1.0
1.0
1.0
1.0
1.0
1.0
1.0
1.0
1.0
1.0
1.0
1.0
1.0
1.0
1.0
1.0
1.0
1.0
1.0
1.0
1.0
1.0
1.0
1.0
1.0
1.0
1.0
1.0
1.0


8
0
0
0
0
0
0
0
0
0
0
0
0
0
0
0
1
0
0
0
0
0
0
0
3
0
0
1
3
0

0.99
0.99
0.99
0.99
0.99
0.99
0.99
0.99
0.99
0.99
0.99
0.99
0.99
0.99
0.99
0.99
0.99
0.99
0.99
0.99
0.99
0.99
0.99
0.99
0.99
0.99
0.99
0.99
0.99
0.99


4
0
0
0
0
0
0
0
0
0
0
0
0
0
0
0
0
0
0
0
0
0
0
0
0
1
3
0
0
0

1.0
1.0
1.0
1.0
1.0
1.0
1.0
1.0
1.0
1.0
1.0
1.0
1.0
1.0
1.0
1.0
1.0
1.0
1.0
1.0
1.0
1.0
1.0
1.0
1.0
1.0
1.0
1.0
1.0
1.0


1
0
0
0
0
0
0
0
0
0
0
0
0
0
0
0
0
1
0
0
0
0
0
0
0
0
0
0
0
0


1
0
0
0
0
0
0
0
0
0
0
0
0
0
0
0
0
1
0
0
0
0
0
0
0
0
0
0
0
0

1.0
1.0
1.0
1.0
1.0
1.0
1.0
1.0
1.0
1.0
1.0
1.0
1.0
1.0
1.0
1.0
1.0
1.0
1.0
1.0
1.0
1.0
1.0
1.0
1.0
1.0
1.0
1.0
1.0
1.0


1
0
0
0
0
0
0
0
0
0
0
0
0
0
1
0
0
0
0
0
0
0
0
0
0
0
0
0
0
0


1
0
0
0
0
0
0
0
0
0
0
0
0
0
1
0
0
0
0
0
0
0
0
0
0
0
0
0
0
0

1.0
1.0
1.0
1.0
1.0
1.0
1.0
1.0
1.0
1.0
1.0
1.0
1.0
1.0
1.0
1.0
1.0
1.0
1.0
1.0
1.0
1.0
1.0
1.0
1.0
1.0
1.0
1.0
1.0
1.0


3
0
0
0
0
0
0
0
0
0
0
0
0
0
0
0
0
0
0
0
0
0
0
0
1
0
1
0
0
1

0.79
0.79
0.79
0.79
0.79
0.79
0.79
0.79
0.79
0.79
0.79
0.79
0.79
0.79
0.79
0.79
0.79
0.79
0.79
0.79
0.79
0.79
0.79
0.79
0.79
0.79
0.79
0.79
0.79
0.79


1
0
0
0
0
0
0
0
0
0
0
0
0
0
0
0
0
0
0
0
0
0
0
0
0
0
0
0
0
1

0.71
0.71
0.71
0.71
0.71
0.71
0.71
0.71
0.71
0.71
0.71
0.71
0.71
0.71
0.71
0.71
0.71
0.71
0.71
0.71
0.71
0.71
0.71
0.71
0.71
0.71
0.71
0.71
0.71
0.71


1
0
0
0
0
0
0
0
0
0
0
0
0
0
0
0
0
0
0
0
0
0
0
0
0
1
0
0
0
0


1
0
0
0
0
0
0
0
0
0
0
0
0
0
0
0
0
0
0
0
0
0
0
0
0
1
0
0
0
0

1.0
1.0
1.0
1.0
1.0
1.0
1.0
1.0
1.0
1.0
1.0
1.0
1.0
1.0
1.0
1.0
1.0
1.0
1.0
1.0
1.0
1.0
1.0
1.0
1.0
1.0
1.0
1.0
1.0
1.0


1
0
0
0
0
0
0
0
0
0
0
0
0
0
0
0
0
0
0
0
0
0
0
0
1
0
0
0
0
0


1
0
0
0
0
0
0
0
0
0
0
0
0
0
0
0
0
0
0
0
0
0
0
0
1
0
0
0
0
0

1.0
1.0
1.0
1.0
1.0
1.0
1.0
1.0
1.0
1.0
1.0
1.0
1.0
1.0
1.0
1.0
1.0
1.0
1.0
1.0
1.0
1.0
1.0
1.0
1.0
1.0
1.0
1.0
1.0
1.0


107
0
0
0
0
0
1
0
0
0
0
0
0
5
0
1
1
0
24
30
0
5
5
0
5
9
11
1
6
3


84
0
0
0
0
0
1
0
0
0
0
0
0
5
0
0
0
0
24
30
0
5
5
0
4
1
1
1
5
2

1.0
1.0
1.0
1.0
1.0
1.0
1.0
1.0
1.0
1.0
1.0
1.0
1.0
1.0
1.0
1.0
1.0
1.0
1.0
1.0
1.0
1.0
1.0
1.0
1.0
1.0
1.0
1.0
1.0
1.0


9
0
0
0
0
0
1
0
0
0
0
0
0
0
0
0
0
0
0
0
0
0
0
0
2
1
1
1
1
2

0.97
0.97
0.97
0.97
0.97
0.97
0.97
0.97
0.97
0.97
0.97
0.97
0.97
0.97
0.97
0.97
0.97
0.97
0.97
0.97
0.97
0.97
0.97
0.97
0.97
0.97
0.97
0.97
0.97
0.97


14
0
0
0
0
0
0
0
0
0
0
0
0
0
0
0
0
0
8
6
0
0
0
0
0
0
0
0
0
0

1.0
1.0
1.0
1.0
1.0
1.0
1.0
1.0
1.0
1.0
1.0
1.0
1.0
1.0
1.0
1.0
1.0
1.0
1.0
1.0
1.0
1.0
1.0
1.0
1.0
1.0
1.0
1.0
1.0
1.0


21
0
0
0
0
0
0
0
0
0
0
0
0
0
0
0
0
0
0
9
0
5
5
0
2
0
0
0
0
0

1.0
1.0
1.0
1.0
1.0
1.0
1.0
1.0
1.0
1.0
1.0
1.0
1.0
1.0
1.0
1.0
1.0
1.0
1.0
1.0
1.0
1.0
1.0
1.0
1.0
1.0
1.0
1.0
1.0
1.0


9
0
0
0
0
0
0
0
0
0
0
0
0
5
0
0
0
0
0
0
0
0
0
0
0
0
0
0
4
0

1.0
1.0
1.0
1.0
1.0
1.0
1.0
1.0
1.0
1.0
1.0
1.0
1.0
1.0
1.0
1.0
1.0
1.0
1.0
1.0
1.0
1.0
1.0
1.0
1.0
1.0
1.0
1.0
1.0
1.0


23
0
0
0
0
0
0
0
0
0
0
0
0
0
0
1
1
0
0
0
0
0
0
0
1
8
10
0
1
1

0.99
0.99
0.99
0.99
0.99
0.99
0.99
0.99
0.99
0.99
0.99
0.99
0.99
0.99
0.99
0.99
0.99
0.99
0.99
0.99
0.99
0.99
0.99
0.99
0.99
0.99
0.99
0.99
0.99
0.99


132
0
0
0
0
0
0
0
0
0
0
0
0
0
0
0
0
0
0
3
0
0
1
0
37
17
25
42
6
1


132
0
0
0
0
0
0
0
0
0
0
0
0
0
0
0
0
0
0
3
0
0
1
0
37
17
25
42
6
1

1.0
1.0
1.0
1.0
1.0
1.0
1.0
1.0
1.0
1.0
1.0
1.0
1.0
1.0
1.0
1.0
1.0
1.0
1.0
1.0
1.0
1.0
1.0
1.0
1.0
1.0
1.0
1.0
1.0
1.0


115
0
0
0
0
0
0
0
0
0
0
0
0
0
0
0
0
0
0
0
0
0
0
0
28
17
24
41
4
1

1.0
1.0
1.0
1.0
1.0
1.0
1.0
1.0
1.0
1.0
1.0
1.0
1.0
1.0
1.0
1.0
1.0
1.0
1.0
1.0
1.0
1.0
1.0
1.0
1.0
1.0
1.0
1.0
1.0
1.0


6
0
0
0
0
0
0
0
0
0
0
0
0
0
0
0
0
0
0
2
0
0
1
0
3
0
0
0
0
0

0.8
0.8
0.8
0.8
0.8
0.8
0.8
0.8
0.8
0.8
0.8
0.8
0.8
0.8
0.8
0.8
0.8
0.8
0.8
0.8
0.8
0.8
0.8
0.8
0.8
0.8
0.8
0.8
0.8
0.8


9
0
0
0
0
0
0
0
0
0
0
0
0
0
0
3
6
0
0
0
0
0
0
0
0
0
0
0
0
0


9
0
0
0
0
0
0
0
0
0
0
0
0
0
0
3
6
0
0
0
0
0
0
0
0
0
0
0
0
0


9
0
0
0
0
0
0
0
0
0
0
0
0
0
0
3
6
0
0
0
0
0
0
0
0
0
0
0
0
0

1.0
1.0
1.0
1.0
1.0
1.0
1.0
1.0
1.0
1.0
1.0
1.0
1.0
1.0
1.0
1.0
1.0
1.0
1.0
1.0
1.0
1.0
1.0
1.0
1.0
1.0
1.0
1.0
1.0
1.0


11
0
0
0
0
0
0
0
0
0
0
0
0
0
0
0
0
0
0
0
0
0
0
0
1
1
5
0
0
4

0.97
0.97
0.97
0.97
0.97
0.97
0.97
0.97
0.97
0.97
0.97
0.97
0.97
0.97
0.97
0.97
0.97
0.97
0.97
0.97
0.97
0.97
0.97
0.97
0.97
0.97
0.97
0.97
0.97
0.97


1
0
0
0
0
0
0
0
0
0
0
0
0
0
0
0
0
0
0
0
0
0
0
0
0
0
0
0
0
1

0.99
0.99
0.99
0.99
0.99
0.99
0.99
0.99
0.99
0.99
0.99
0.99
0.99
0.99
0.99
0.99
0.99
0.99
0.99
0.99
0.99
0.99
0.99
0.99
0.99
0.99
0.99
0.99
0.99
0.99


5
0
0
0
0
0
0
0
0
0
0
0
0
0
0
0
0
0
0
0
0
0
0
0
1
0
4
0
0
0


1
0
0
0
0
0
0
0
0
0
0
0
0
0
0
0
0
0
0
0
0
0
0
0
1
0
0
0
0
0


1
0
0
0
0
0
0
0
0
0
0
0
0
0
0
0
0
0
0
0
0
0
0
0
1
0
0
0
0
0

0.98
0.98
0.98
0.98
0.98
0.98
0.98
0.98
0.98
0.98
0.98
0.98
0.98
0.98
0.98
0.98
0.98
0.98
0.98
0.98
0.98
0.98
0.98
0.98
0.98
0.98
0.98
0.98
0.98
0.98


4
0
0
0
0
0
0
0
0
0
0
0
0
0
0
0
0
0
0
0
0
0
0
0
0
0
4
0
0
0

0.96
0.96
0.96
0.96
0.96
0.96
0.96
0.96
0.96
0.96
0.96
0.96
0.96
0.96
0.96
0.96
0.96
0.96
0.96
0.96
0.96
0.96
0.96
0.96
0.96
0.96
0.96
0.96
0.96
0.96


5
0
0
0
0
0
0
0
0
0
0
0
0
0
0
0
0
0
0
0
0
0
0
0
3
0
0
0
2
0


3
0
0
0
0
0
0
0
0
0
0
0
0
0
0
0
0
0
0
0
0
0
0
0
3
0
0
0
0
0

0.94
0.94
0.94
0.94
0.94
0.94
0.94
0.94
0.94
0.94
0.94
0.94
0.94
0.94
0.94
0.94
0.94
0.94
0.94
0.94
0.94
0.94
0.94
0.94
0.94
0.94
0.94
0.94
0.94
0.94


2
0
0
0
0
0
0
0
0
0
0
0
0
0
0
0
0
0
0
0
0
0
0
0
0
0
0
0
2
0

0.89
0.89
0.89
0.89
0.89
0.89
0.89
0.89
0.89
0.89
0.89
0.89
0.89
0.89
0.89
0.89
0.89
0.89
0.89
0.89
0.89
0.89
0.89
0.89
0.89
0.89
0.89
0.89
0.89
0.89


1
0
0
0
0
0
0
0
0
0
0
0
0
0
0
0
1
0
0
0
0
0
0
0
0
0
0
0
0
0


1
0
0
0
0
0
0
0
0
0
0
0
0
0
0
0
1
0
0
0
0
0
0
0
0
0
0
0
0
0

1.0
1.0
1.0
1.0
1.0
1.0
1.0
1.0
1.0
1.0
1.0
1.0
1.0
1.0
1.0
1.0
1.0
1.0
1.0
1.0
1.0
1.0
1.0
1.0
1.0
1.0
1.0
1.0
1.0
1.0


30
0
0
0
0
0
0
0
0
0
0
0
0
0
0
1
0
2
3
1
2
0
0
0
5
4
4
0
4
4

1.0
1.0
1.0
1.0
1.0
1.0
1.0
1.0
1.0
1.0
1.0
1.0
1.0
1.0
1.0
1.0
1.0
1.0
1.0
1.0
1.0
1.0
1.0
1.0
1.0
1.0
1.0
1.0
1.0
1.0


5
0
0
0
0
0
0
0
0
0
0
0
0
0
0
0
0
0
0
0
1
0
0
0
4
0
0
0
0
0

1.0
1.0
1.0
1.0
1.0
1.0
1.0
1.0
1.0
1.0
1.0
1.0
1.0
1.0
1.0
1.0
1.0
1.0
1.0
1.0
1.0
1.0
1.0
1.0
1.0
1.0
1.0
1.0
1.0
1.0


1
0
0
0
0
0
0
0
0
0
0
0
0
0
0
0
0
0
0
0
0
0
0
0
1
0
0
0
0
0

1.0
1.0
1.0
1.0
1.0
1.0
1.0
1.0
1.0
1.0
1.0
1.0
1.0
1.0
1.0
1.0
1.0
1.0
1.0
1.0
1.0
1.0
1.0
1.0
1.0
1.0
1.0
1.0
1.0
1.0


3
0
0
0
0
0
0
0
0
0
0
0
0
0
0
0
0
1
1
0
1
0
0
0
0
0
0
0
0
0

1.0
1.0
1.0
1.0
1.0
1.0
1.0
1.0
1.0
1.0
1.0
1.0
1.0
1.0
1.0
1.0
1.0
1.0
1.0
1.0
1.0
1.0
1.0
1.0
1.0
1.0
1.0
1.0
1.0
1.0


5
0
0
0
0
0
0
0
0
0
0
0
0
0
0
0
0
0
0
0
0
0
0
0
0
0
2
0
0
3


5
0
0
0
0
0
0
0
0
0
0
0
0
0
0
0
0
0
0
0
0
0
0
0
0
0
2
0
0
3

0.96
0.96
0.96
0.96
0.96
0.96
0.96
0.96
0.96
0.96
0.96
0.96
0.96
0.96
0.96
0.96
0.96
0.96
0.96
0.96
0.96
0.96
0.96
0.96
0.96
0.96
0.96
0.96
0.96
0.96


2
0
0
0
0
0
0
0
0
0
0
0
0
0
0
0
0
0
0
0
0
0
0
0
0
1
0
0
1
0

0.72
0.72
0.72
0.72
0.72
0.72
0.72
0.72
0.72
0.72
0.72
0.72
0.72
0.72
0.72
0.72
0.72
0.72
0.72
0.72
0.72
0.72
0.72
0.72
0.72
0.72
0.72
0.72
0.72
0.72


14
0
0
0
0
0
0
0
0
0
0
0
0
0
1
0
0
2
0
1
0
0
0
0
3
3
2
1
1
0


5
0
0
0
0
0
0
0
0
0
0
0
0
0
1
0
0
0
0
1
0
0
0
0
1
1
0
0
1
0

0.92
0.92
0.92
0.92
0.92
0.92
0.92
0.92
0.92
0.92
0.92
0.92
0.92
0.92
0.92
0.92
0.92
0.92
0.92
0.92
0.92
0.92
0.92
0.92
0.92
0.92
0.92
0.92
0.92
0.92


1
0
0
0
0
0
0
0
0
0
0
0
0
0
0
0
0
0
0
0
0
0
0
0
1
0
0
0
0
0

1.0
1.0
1.0
1.0
1.0
1.0
1.0
1.0
1.0
1.0
1.0
1.0
1.0
1.0
1.0
1.0
1.0
1.0
1.0
1.0
1.0
1.0
1.0
1.0
1.0
1.0
1.0
1.0
1.0
1.0


2
0
0
0
0
0
0
0
0
0
0
0
0
0
0
0
0
0
0
0
0
0
0
0
0
1
1
0
0
0

1.0
1.0
1.0
1.0
1.0
1.0
1.0
1.0
1.0
1.0
1.0
1.0
1.0
1.0
1.0
1.0
1.0
1.0
1.0
1.0
1.0
1.0
1.0
1.0
1.0
1.0
1.0
1.0
1.0
1.0


4
0
0
0
0
0
0
0
0
0
0
0
0
0
0
0
0
1
0
0
0
0
0
0
1
1
0
1
0
0

0.74
0.74
0.74
0.74
0.74
0.74
0.74
0.74
0.74
0.74
0.74
0.74
0.74
0.74
0.74
0.74
0.74
0.74
0.74
0.74
0.74
0.74
0.74
0.74
0.74
0.74
0.74
0.74
0.74
0.74


3
0
0
0
0
0
0
0
0
0
0
0
0
0
0
0
0
1
0
0
0
0
0
0
1
0
0
1
0
0

1.0
1.0
1.0
1.0
1.0
1.0
1.0
1.0
1.0
1.0
1.0
1.0
1.0
1.0
1.0
1.0
1.0
1.0
1.0
1.0
1.0
1.0
1.0
1.0
1.0
1.0
1.0
1.0
1.0
1.0


1
0
0
0
0
0
0
0
0
0
0
0
0
0
0
0
0
0
0
0
0
0
0
0
1
0
0
0
0
0

0.87
0.87
0.87
0.87
0.87
0.87
0.87
0.87
0.87
0.87
0.87
0.87
0.87
0.87
0.87
0.87
0.87
0.87
0.87
0.87
0.87
0.87
0.87
0.87
0.87
0.87
0.87
0.87
0.87
0.87


2
0
0
0
0
0
0
0
0
0
0
0
0
0
0
0
0
1
0
0
0
0
0
0
0
0
1
0
0
0

1.0
1.0
1.0
1.0
1.0
1.0
1.0
1.0
1.0
1.0
1.0
1.0
1.0
1.0
1.0
1.0
1.0
1.0
1.0
1.0
1.0
1.0
1.0
1.0
1.0
1.0
1.0
1.0
1.0
1.0


102
0
0
0
0
0
0
0
0
0
0
0
0
0
0
6
0
7
2
0
15
0
9
0
6
5
5
20
14
13

0.94
0.94
0.94
0.94
0.94
0.94
0.94
0.94
0.94
0.94
0.94
0.94
0.94
0.94
0.94
0.94
0.94
0.94
0.94
0.94
0.94
0.94
0.94
0.94
0.94
0.94
0.94
0.94
0.94
0.94


13
0
0
0
0
0
0
0
0
0
0
0
0
0
0
0
0
0
0
0
0
0
0
0
1
0
3
3
3
3

0.97
0.97
0.97
0.97
0.97
0.97
0.97
0.97
0.97
0.97
0.97
0.97
0.97
0.97
0.97
0.97
0.97
0.97
0.97
0.97
0.97
0.97
0.97
0.97
0.97
0.97
0.97
0.97
0.97
0.97


23
0
0
0
0
0
0
0
0
0
0
0
0
0
0
6
0
1
1
0
0
0
0
0
3
3
0
8
1
0


23
0
0
0
0
0
0
0
0
0
0
0
0
0
0
6
0
1
1
0
0
0
0
0
3
3
0
8
1
0

1.0
1.0
1.0
1.0
1.0
1.0
1.0
1.0
1.0
1.0
1.0
1.0
1.0
1.0
1.0
1.0
1.0
1.0
1.0
1.0
1.0
1.0
1.0
1.0
1.0
1.0
1.0
1.0
1.0
1.0


43
0
0
0
0
0
0
0
0
0
0
0
0
0
0
0
0
0
0
0
15
0
0
0
1
2
2
4
9
10

1.0
1.0
1.0
1.0
1.0
1.0
1.0
1.0
1.0
1.0
1.0
1.0
1.0
1.0
1.0
1.0
1.0
1.0
1.0
1.0
1.0
1.0
1.0
1.0
1.0
1.0
1.0
1.0
1.0
1.0


10
0
0
0
0
0
0
0
0
0
0
0
0
0
0
0
0
0
0
0
8
0
0
0
0
0
0
0
1
1

1.0
1.0
1.0
1.0
1.0
1.0
1.0
1.0
1.0
1.0
1.0
1.0
1.0
1.0
1.0
1.0
1.0
1.0
1.0
1.0
1.0
1.0
1.0
1.0
1.0
1.0
1.0
1.0
1.0
1.0


6
0
0
0
0
0
0
0
0
0
0
0
0
0
0
0
0
6
0
0
0
0
0
0
0
0
0
0
0
0

1.0
1.0
1.0
1.0
1.0
1.0
1.0
1.0
1.0
1.0
1.0
1.0
1.0
1.0
1.0
1.0
1.0
1.0
1.0
1.0
1.0
1.0
1.0
1.0
1.0
1.0
1.0
1.0
1.0
1.0


5
0
0
0
0
0
0
0
0
0
0
0
0
0
0
0
0
5
0
0
0
0
0
0
0
0
0
0
0
0

1.0
1.0
1.0
1.0
1.0
1.0
1.0
1.0
1.0
1.0
1.0
1.0
1.0
1.0
1.0
1.0
1.0
1.0
1.0
1.0
1.0
1.0
1.0
1.0
1.0
1.0
1.0
1.0
1.0
1.0


11
0
0
0
0
0
0
0
0
0
0
0
0
0
0
0
0
0
1
0
0
0
9
0
1
0
0
0
0
0

1.0
1.0
1.0
1.0
1.0
1.0
1.0
1.0
1.0
1.0
1.0
1.0
1.0
1.0
1.0
1.0
1.0
1.0
1.0
1.0
1.0
1.0
1.0
1.0
1.0
1.0
1.0
1.0
1.0
1.0


2
0
0
0
0
0
0
0
0
0
0
0
0
0
0
0
0
0
0
0
0
0
2
0
0
0
0
0
0
0

0.9
0.9
0.9
0.9
0.9
0.9
0.9
0.9
0.9
0.9
0.9
0.9
0.9
0.9
0.9
0.9
0.9
0.9
0.9
0.9
0.9
0.9
0.9
0.9
0.9
0.9
0.9
0.9
0.9
0.9


3
0
0
0
0
0
0
0
0
0
0
0
0
0
0
0
0
0
0
0
0
0
3
0
0
0
0
0
0
0

1.0
1.0
1.0
1.0
1.0
1.0
1.0
1.0
1.0
1.0
1.0
1.0
1.0
1.0
1.0
1.0
1.0
1.0
1.0
1.0
1.0
1.0
1.0
1.0
1.0
1.0
1.0
1.0
1.0
1.0


11
0
0
0
0
0
0
0
0
0
0
0
0
0
0
5
0
0
0
0
0
0
0
0
1
5
0
0
0
0


11
0
0
0
0
0
0
0
0
0
0
0
0
0
0
5
0
0
0
0
0
0
0
0
1
5
0
0
0
0

0.97
0.97
0.97
0.97
0.97
0.97
0.97
0.97
0.97
0.97
0.97
0.97
0.97
0.97
0.97
0.97
0.97
0.97
0.97
0.97
0.97
0.97
0.97
0.97
0.97
0.97
0.97
0.97
0.97
0.97


2
0
0
0
0
0
0
0
0
0
0
0
0
0
0
0
0
0
1
0
0
0
0
0
0
1
0
0
0
0


2
0
0
0
0
0
0
0
0
0
0
0
0
0
0
0
0
0
1
0
0
0
0
0
0
1
0
0
0
0

1.0
1.0
1.0
1.0
1.0
1.0
1.0
1.0
1.0
1.0
1.0
1.0
1.0
1.0
1.0
1.0
1.0
1.0
1.0
1.0
1.0
1.0
1.0
1.0
1.0
1.0
1.0
1.0
1.0
1.0


4
0
0
0
0
0
0
0
0
0
0
0
0
0
0
0
0
0
0
0
0
0
0
0
4
0
0
0
0
0


4
0
0
0
0
0
0
0
0
0
0
0
0
0
0
0
0
0
0
0
0
0
0
0
4
0
0
0
0
0


4
0
0
0
0
0
0
0
0
0
0
0
0
0
0
0
0
0
0
0
0
0
0
0
4
0
0
0
0
0

1.0
1.0
1.0
1.0
1.0
1.0
1.0
1.0
1.0
1.0
1.0
1.0
1.0
1.0
1.0
1.0
1.0
1.0
1.0
1.0
1.0
1.0
1.0
1.0
1.0
1.0
1.0
1.0
1.0
1.0


11
0
0
0
0
0
0
0
0
0
0
0
1
0
0
0
0
0
2
0
0
1
5
1
0
1
0
0
0
0

0.7
0.7
0.7
0.7
0.7
0.7
0.7
0.7
0.7
0.7
0.7
0.7
0.7
0.7
0.7
0.7
0.7
0.7
0.7
0.7
0.7
0.7
0.7
0.7
0.7
0.7
0.7
0.7
0.7
0.7


5
0
0
0
0
0
0
0
0
0
0
0
0
0
0
0
0
0
0
0
0
0
5
0
0
0
0
0
0
0

0.98
0.98
0.98
0.98
0.98
0.98
0.98
0.98
0.98
0.98
0.98
0.98
0.98
0.98
0.98
0.98
0.98
0.98
0.98
0.98
0.98
0.98
0.98
0.98
0.98
0.98
0.98
0.98
0.98
0.98


3
0
0
0
0
0
0
0
0
0
0
0
1
0
0
0
0
0
0
0
0
1
0
1
0
0
0
0
0
0

1.0
1.0
1.0
1.0
1.0
1.0
1.0
1.0
1.0
1.0
1.0
1.0
1.0
1.0
1.0
1.0
1.0
1.0
1.0
1.0
1.0
1.0
1.0
1.0
1.0
1.0
1.0
1.0
1.0
1.0


2
0
0
0
0
0
0
0
0
0
0
0
0
0
0
0
0
0
2
0
0
0
0
0
0
0
0
0
0
0

1.0
1.0
1.0
1.0
1.0
1.0
1.0
1.0
1.0
1.0
1.0
1.0
1.0
1.0
1.0
1.0
1.0
1.0
1.0
1.0
1.0
1.0
1.0
1.0
1.0
1.0
1.0
1.0
1.0
1.0


6
0
0
0
0
0
0
0
0
0
0
0
0
0
0
0
2
0
0
2
0
0
0
0
2
0
0
0
0
0

0.94
0.94
0.94
0.94
0.94
0.94
0.94
0.94
0.94
0.94
0.94
0.94
0.94
0.94
0.94
0.94
0.94
0.94
0.94
0.94
0.94
0.94
0.94
0.94
0.94
0.94
0.94
0.94
0.94
0.94


2
0
0
0
0
0
0
0
0
0
0
0
0
0
0
0
1
0
0
0
0
0
0
0
1
0
0
0
0
0

1.0
1.0
1.0
1.0
1.0
1.0
1.0
1.0
1.0
1.0
1.0
1.0
1.0
1.0
1.0
1.0
1.0
1.0
1.0
1.0
1.0
1.0
1.0
1.0
1.0
1.0
1.0
1.0
1.0
1.0


2
0
0
0
0
0
0
0
0
0
0
0
0
0
0
0
0
0
0
2
0
0
0
0
0
0
0
0
0
0

0.94
0.94
0.94
0.94
0.94
0.94
0.94
0.94
0.94
0.94
0.94
0.94
0.94
0.94
0.94
0.94
0.94
0.94
0.94
0.94
0.94
0.94
0.94
0.94
0.94
0.94
0.94
0.94
0.94
0.94


1
0
0
0
0
0
0
0
0
0
0
0
0
0
0
0
0
0
0
0
0
0
0
0
0
0
0
0
1
0

0.92
0.92
0.92
0.92
0.92
0.92
0.92
0.92
0.92
0.92
0.92
0.92
0.92
0.92
0.92
0.92
0.92
0.92
0.92
0.92
0.92
0.92
0.92
0.92
0.92
0.92
0.92
0.92
0.92
0.92


7
0
0
0
0
0
0
0
0
0
0
0
0
0
0
0
0
0
0
0
0
0
0
0
0
0
7
0
0
0


7
0
0
0
0
0
0
0
0
0
0
0
0
0
0
0
0
0
0
0
0
0
0
0
0
0
7
0
0
0


7
0
0
0
0
0
0
0
0
0
0
0
0
0
0
0
0
0
0
0
0
0
0
0
0
0
7
0
0
0

1.0
1.0
1.0
1.0
1.0
1.0
1.0
1.0
1.0
1.0
1.0
1.0
1.0
1.0
1.0
1.0
1.0
1.0
1.0
1.0
1.0
1.0
1.0
1.0
1.0
1.0
1.0
1.0
1.0
1.0


2175
7
73
80
96
150
119
0
0
27
54
38
0
91
0
2
0
7
317
0
0
0
72
0
1040
0
1
0
1
0

0.73
0.73
0.73
0.73
0.73
0.73
0.73
0.73
0.73
0.73
0.73
0.73
0.73
0.73
0.73
0.73
0.73
0.73
0.73
0.73
0.73
0.73
0.73
0.73
0.73
0.73
0.73
0.73
0.73
0.73


1954
0
67
73
77
111
102
0
0
12
45
33
0
1
0
2
0
0
317
0
0
0
72
0
1040
0
1
0
1
0

1.0
1.0
1.0
1.0
1.0
1.0
1.0
1.0
1.0
1.0
1.0
1.0
1.0
1.0
1.0
1.0
1.0
1.0
1.0
1.0
1.0
1.0
1.0
1.0
1.0
1.0
1.0
1.0
1.0
1.0


357
0
34
28
41
0
32
0
0
11
20
32
0
1
0
1
0
0
1
0
0
0
67
0
89
0
0
0
0
0

1.0
1.0
1.0
1.0
1.0
1.0
1.0
1.0
1.0
1.0
1.0
1.0
1.0
1.0
1.0
1.0
1.0
1.0
1.0
1.0
1.0
1.0
1.0
1.0
1.0
1.0
1.0
1.0
1.0
1.0


272
0
24
20
31
0
28
0
0
8
16
23
0
1
0
1
0
0
1
0
0
0
52
0
67
0
0
0
0
0

1.0
1.0
1.0
1.0
1.0
1.0
1.0
1.0
1.0
1.0
1.0
1.0
1.0
1.0
1.0
1.0
1.0
1.0
1.0
1.0
1.0
1.0
1.0
1.0
1.0
1.0
1.0
1.0
1.0
1.0


4
0
0
0
0
0
0
0
0
0
0
0
0
0
0
0
0
0
0
0
0
0
3
0
1
0
0
0
0
0

0.95
0.95
0.95
0.95
0.95
0.95
0.95
0.95
0.95
0.95
0.95
0.95
0.95
0.95
0.95
0.95
0.95
0.95
0.95
0.95
0.95
0.95
0.95
0.95
0.95
0.95
0.95
0.95
0.95
0.95


741
0
23
11
1
0
0
0
0
1
16
0
0
0
0
0
0
0
205
0
0
0
0
0
484
0
0
0
0
0

1.0
1.0
1.0
1.0
1.0
1.0
1.0
1.0
1.0
1.0
1.0
1.0
1.0
1.0
1.0
1.0
1.0
1.0
1.0
1.0
1.0
1.0
1.0
1.0
1.0
1.0
1.0
1.0
1.0
1.0


208
0
2
2
0
0
0
0
0
0
1
0
0
0
0
0
0
0
63
0
0
0
0
0
140
0
0
0
0
0

1.0
1.0
1.0
1.0
1.0
1.0
1.0
1.0
1.0
1.0
1.0
1.0
1.0
1.0
1.0
1.0
1.0
1.0
1.0
1.0
1.0
1.0
1.0
1.0
1.0
1.0
1.0
1.0
1.0
1.0


7
0
0
0
0
0
0
0
0
0
0
0
0
0
0
0
0
0
2
0
0
0
0
0
5
0
0
0
0
0

0.97
0.97
0.97
0.97
0.97
0.97
0.97
0.97
0.97
0.97
0.97
0.97
0.97
0.97
0.97
0.97
0.97
0.97
0.97
0.97
0.97
0.97
0.97
0.97
0.97
0.97
0.97
0.97
0.97
0.97


4
0
0
0
0
0
0
0
0
0
0
0
0
0
0
0
0
0
4
0
0
0
0
0
0
0
0
0
0
0

0.99
0.99
0.99
0.99
0.99
0.99
0.99
0.99
0.99
0.99
0.99
0.99
0.99
0.99
0.99
0.99
0.99
0.99
0.99
0.99
0.99
0.99
0.99
0.99
0.99
0.99
0.99
0.99
0.99
0.99


158
0
0
13
18
50
36
0
0
0
0
0
0
0
0
1
0
0
0
0
0
0
0
0
38
0
1
0
1
0

1.0
1.0
1.0
1.0
1.0
1.0
1.0
1.0
1.0
1.0
1.0
1.0
1.0
1.0
1.0
1.0
1.0
1.0
1.0
1.0
1.0
1.0
1.0
1.0
1.0
1.0
1.0
1.0
1.0
1.0


127
0
0
9
14
39
29
0
0
0
0
0
0
0
0
1
0
0
0
0
0
0
0
0
33
0
1
0
1
0

1.0
1.0
1.0
1.0
1.0
1.0
1.0
1.0
1.0
1.0
1.0
1.0
1.0
1.0
1.0
1.0
1.0
1.0
1.0
1.0
1.0
1.0
1.0
1.0
1.0
1.0
1.0
1.0
1.0
1.0


1
0
0
0
0
0
1
0
0
0
0
0
0
0
0
0
0
0
0
0
0
0
0
0
0
0
0
0
0
0


1
0
0
0
0
0
1
0
0
0
0
0
0
0
0
0
0
0
0
0
0
0
0
0
0
0
0
0
0
0

0.94
0.94
0.94
0.94
0.94
0.94
0.94
0.94
0.94
0.94
0.94
0.94
0.94
0.94
0.94
0.94
0.94
0.94
0.94
0.94
0.94
0.94
0.94
0.94
0.94
0.94
0.94
0.94
0.94
0.94


3
0
1
1
0
0
0
0
0
0
0
1
0
0
0
0
0
0
0
0
0
0
0
0
0
0
0
0
0
0


3
0
1
1
0
0
0
0
0
0
0
1
0
0
0
0
0
0
0
0
0
0
0
0
0
0
0
0
0
0

1.0
1.0
1.0
1.0
1.0
1.0
1.0
1.0
1.0
1.0
1.0
1.0
1.0
1.0
1.0
1.0
1.0
1.0
1.0
1.0
1.0
1.0
1.0
1.0
1.0
1.0
1.0
1.0
1.0
1.0


27
0
0
2
5
12
8
0
0
0
0
0
0
0
0
0
0
0
0
0
0
0
0
0
0
0
0
0
0
0

0.97
0.97
0.97
0.97
0.97
0.97
0.97
0.97
0.97
0.97
0.97
0.97
0.97
0.97
0.97
0.97
0.97
0.97
0.97
0.97
0.97
0.97
0.97
0.97
0.97
0.97
0.97
0.97
0.97
0.97


2
0
0
0
0
0
0
0
0
0
0
0
0
0
0
0
0
0
1
0
0
0
0
0
1
0
0
0
0
0


2
0
0
0
0
0
0
0
0
0
0
0
0
0
0
0
0
0
1
0
0
0
0
0
1
0
0
0
0
0

1.0
1.0
1.0
1.0
1.0
1.0
1.0
1.0
1.0
1.0
1.0
1.0
1.0
1.0
1.0
1.0
1.0
1.0
1.0
1.0
1.0
1.0
1.0
1.0
1.0
1.0
1.0
1.0
1.0
1.0


7
0
0
0
0
0
0
0
0
0
0
0
0
0
0
0
0
0
2
0
0
0
0
0
5
0
0
0
0
0

0.87
0.87
0.87
0.87
0.87
0.87
0.87
0.87
0.87
0.87
0.87
0.87
0.87
0.87
0.87
0.87
0.87
0.87
0.87
0.87
0.87
0.87
0.87
0.87
0.87
0.87
0.87
0.87
0.87
0.87


30
0
0
0
0
0
0
0
0
0
0
0
0
0
0
0
0
0
9
0
0
0
0
0
21
0
0
0
0
0

0.97
0.97
0.97
0.97
0.97
0.97
0.97
0.97
0.97
0.97
0.97
0.97
0.97
0.97
0.97
0.97
0.97
0.97
0.97
0.97
0.97
0.97
0.97
0.97
0.97
0.97
0.97
0.97
0.97
0.97


2
0
0
0
0
0
0
0
0
0
0
0
0
0
0
0
0
0
0
0
0
0
0
0
2
0
0
0
0
0

0.77
0.77
0.77
0.77
0.77
0.77
0.77
0.77
0.77
0.77
0.77
0.77
0.77
0.77
0.77
0.77
0.77
0.77
0.77
0.77
0.77
0.77
0.77
0.77
0.77
0.77
0.77
0.77
0.77
0.77


22
0
0
0
0
0
0
0
0
0
0
0
0
0
0
0
0
0
3
0
0
0
0
0
19
0
0
0
0
0

0.89
0.89
0.89
0.89
0.89
0.89
0.89
0.89
0.89
0.89
0.89
0.89
0.89
0.89
0.89
0.89
0.89
0.89
0.89
0.89
0.89
0.89
0.89
0.89
0.89
0.89
0.89
0.89
0.89
0.89


15
0
0
0
0
0
0
0
0
0
0
0
0
0
0
0
0
0
0
0
0
0
0
0
15
0
0
0
0
0

0.89
0.89
0.89
0.89
0.89
0.89
0.89
0.89
0.89
0.89
0.89
0.89
0.89
0.89
0.89
0.89
0.89
0.89
0.89
0.89
0.89
0.89
0.89
0.89
0.89
0.89
0.89
0.89
0.89
0.89


194
7
6
7
14
39
5
0
0
15
9
5
0
87
0
0
0
0
0
0
0
0
0
0
0
0
0
0
0
0

1.0
1.0
1.0
1.0
1.0
1.0
1.0
1.0
1.0
1.0
1.0
1.0
1.0
1.0
1.0
1.0
1.0
1.0
1.0
1.0
1.0
1.0
1.0
1.0
1.0
1.0
1.0
1.0
1.0
1.0


11
0
0
0
0
6
5
0
0
0
0
0
0
0
0
0
0
0
0
0
0
0
0
0
0
0
0
0
0
0


11
0
0
0
0
6
5
0
0
0
0
0
0
0
0
0
0
0
0
0
0
0
0
0
0
0
0
0
0
0

1.0
1.0
1.0
1.0
1.0
1.0
1.0
1.0
1.0
1.0
1.0
1.0
1.0
1.0
1.0
1.0
1.0
1.0
1.0
1.0
1.0
1.0
1.0
1.0
1.0
1.0
1.0
1.0
1.0
1.0


5
0
5
0
0
0
0
0
0
0
0
0
0
0
0
0
0
0
0
0
0
0
0
0
0
0
0
0
0
0


5
0
5
0
0
0
0
0
0
0
0
0
0
0
0
0
0
0
0
0
0
0
0
0
0
0
0
0
0
0

1.0
1.0
1.0
1.0
1.0
1.0
1.0
1.0
1.0
1.0
1.0
1.0
1.0
1.0
1.0
1.0
1.0
1.0
1.0
1.0
1.0
1.0
1.0
1.0
1.0
1.0
1.0
1.0
1.0
1.0


14
0
0
0
0
0
0
0
0
0
9
5
0
0
0
0
0
0
0
0
0
0
0
0
0
0
0
0
0
0

1.0
1.0
1.0
1.0
1.0
1.0
1.0
1.0
1.0
1.0
1.0
1.0
1.0
1.0
1.0
1.0
1.0
1.0
1.0
1.0
1.0
1.0
1.0
1.0
1.0
1.0
1.0
1.0
1.0
1.0


12
0
0
0
0
0
0
0
0
0
7
5
0
0
0
0
0
0
0
0
0
0
0
0
0
0
0
0
0
0

1.0
1.0
1.0
1.0
1.0
1.0
1.0
1.0
1.0
1.0
1.0
1.0
1.0
1.0
1.0
1.0
1.0
1.0
1.0
1.0
1.0
1.0
1.0
1.0
1.0
1.0
1.0
1.0
1.0
1.0


14
0
0
0
0
0
0
0
0
14
0
0
0
0
0
0
0
0
0
0
0
0
0
0
0
0
0
0
0
0


14
0
0
0
0
0
0
0
0
14
0
0
0
0
0
0
0
0
0
0
0
0
0
0
0
0
0
0
0
0

1.0
1.0
1.0
1.0
1.0
1.0
1.0
1.0
1.0
1.0
1.0
1.0
1.0
1.0
1.0
1.0
1.0
1.0
1.0
1.0
1.0
1.0
1.0
1.0
1.0
1.0
1.0
1.0
1.0
1.0


1
0
1
0
0
0
0
0
0
0
0
0
0
0
0
0
0
0
0
0
0
0
0
0
0
0
0
0
0
0


1
0
1
0
0
0
0
0
0
0
0
0
0
0
0
0
0
0
0
0
0
0
0
0
0
0
0
0
0
0

0.72
0.72
0.72
0.72
0.72
0.72
0.72
0.72
0.72
0.72
0.72
0.72
0.72
0.72
0.72
0.72
0.72
0.72
0.72
0.72
0.72
0.72
0.72
0.72
0.72
0.72
0.72
0.72
0.72
0.72


6
0
0
0
2
4
0
0
0
0
0
0
0
0
0
0
0
0
0
0
0
0
0
0
0
0
0
0
0
0

0.92
0.92
0.92
0.92
0.92
0.92
0.92
0.92
0.92
0.92
0.92
0.92
0.92
0.92
0.92
0.92
0.92
0.92
0.92
0.92
0.92
0.92
0.92
0.92
0.92
0.92
0.92
0.92
0.92
0.92


2
0
0
0
0
2
0
0
0
0
0
0
0
0
0
0
0
0
0
0
0
0
0
0
0
0
0
0
0
0

1.0
1.0
1.0
1.0
1.0
1.0
1.0
1.0
1.0
1.0
1.0
1.0
1.0
1.0
1.0
1.0
1.0
1.0
1.0
1.0
1.0
1.0
1.0
1.0
1.0
1.0
1.0
1.0
1.0
1.0


3
1
0
1
0
0
0
0
0
0
0
0
0
1
0
0
0
0
0
0
0
0
0
0
0
0
0
0
0
0

0.99
0.99
0.99
0.99
0.99
0.99
0.99
0.99
0.99
0.99
0.99
0.99
0.99
0.99
0.99
0.99
0.99
0.99
0.99
0.99
0.99
0.99
0.99
0.99
0.99
0.99
0.99
0.99
0.99
0.99


34
2
0
2
0
0
0
0
0
0
0
0
0
30
0
0
0
0
0
0
0
0
0
0
0
0
0
0
0
0

1.0
1.0
1.0
1.0
1.0
1.0
1.0
1.0
1.0
1.0
1.0
1.0
1.0
1.0
1.0
1.0
1.0
1.0
1.0
1.0
1.0
1.0
1.0
1.0
1.0
1.0
1.0
1.0
1.0
1.0


17
0
0
0
5
0
12
0
0
0
0
0
0
0
0
0
0
0
0
0
0
0
0
0
0
0
0
0
0
0


17
0
0
0
5
0
12
0
0
0
0
0
0
0
0
0
0
0
0
0
0
0
0
0
0
0
0
0
0
0


17
0
0
0
5
0
12
0
0
0
0
0
0
0
0
0
0
0
0
0
0
0
0
0
0
0
0
0
0
0

1.0
1.0
1.0
1.0
1.0
1.0
1.0
1.0
1.0
1.0
1.0
1.0
1.0
1.0
1.0
1.0
1.0
1.0
1.0
1.0
1.0
1.0
1.0
1.0
1.0
1.0
1.0
1.0
1.0
1.0


7
0
0
0
0
0
0
0
0
0
0
0
0
0
0
0
0
7
0
0
0
0
0
0
0
0
0
0
0
0


7
0
0
0
0
0
0
0
0
0
0
0
0
0
0
0
0
7
0
0
0
0
0
0
0
0
0
0
0
0

1.0
1.0
1.0
1.0
1.0
1.0
1.0
1.0
1.0
1.0
1.0
1.0
1.0
1.0
1.0
1.0
1.0
1.0
1.0
1.0
1.0
1.0
1.0
1.0
1.0
1.0
1.0
1.0
1.0
1.0


6
0
0
0
0
0
0
0
0
0
0
0
0
0
0
0
0
6
0
0
0
0
0
0
0
0
0
0
0
0

1.0
1.0
1.0
1.0
1.0
1.0
1.0
1.0
1.0
1.0
1.0
1.0
1.0
1.0
1.0
1.0
1.0
1.0
1.0
1.0
1.0
1.0
1.0
1.0
1.0
1.0
1.0
1.0
1.0
1.0


123
1
1
2
6
9
9
0
0
3
2
2
0
0
14
7
5
0
0
0
0
1
0
0
7
19
17
15
2
1

0.89
0.89
0.89
0.89
0.89
0.89
0.89
0.89
0.89
0.89
0.89
0.89
0.89
0.89
0.89
0.89
0.89
0.89
0.89
0.89
0.89
0.89
0.89
0.89
0.89
0.89
0.89
0.89
0.89
0.89


44
0
0
2
4
4
5
0
0
0
0
0
0
0
12
1
4
0
0
0
0
1
0
0
4
6
1
0
0
0


4
0
0
0
1
0
3
0
0
0
0
0
0
0
0
0
0
0
0
0
0
0
0
0
0
0
0
0
0
0


4
0
0
0
1
0
3
0
0
0
0
0
0
0
0
0
0
0
0
0
0
0
0
0
0
0
0
0
0
0

1.0
1.0
1.0
1.0
1.0
1.0
1.0
1.0
1.0
1.0
1.0
1.0
1.0
1.0
1.0
1.0
1.0
1.0
1.0
1.0
1.0
1.0
1.0
1.0
1.0
1.0
1.0
1.0
1.0
1.0


28
0
0
2
3
4
2
0
0
0
0
0
0
0
0
1
4
0
0
0
0
1
0
0
4
6
1
0
0
0

0.99
0.99
0.99
0.99
0.99
0.99
0.99
0.99
0.99
0.99
0.99
0.99
0.99
0.99
0.99
0.99
0.99
0.99
0.99
0.99
0.99
0.99
0.99
0.99
0.99
0.99
0.99
0.99
0.99
0.99


4
0
0
0
0
0
0
0
0
0
0
0
0
0
0
0
0
0
0
0
0
0
0
0
2
2
0
0
0
0

0.94
0.94
0.94
0.94
0.94
0.94
0.94
0.94
0.94
0.94
0.94
0.94
0.94
0.94
0.94
0.94
0.94
0.94
0.94
0.94
0.94
0.94
0.94
0.94
0.94
0.94
0.94
0.94
0.94
0.94


12
0
0
0
0
0
0
0
0
0
0
0
0
0
12
0
0
0
0
0
0
0
0
0
0
0
0
0
0
0


12
0
0
0
0
0
0
0
0
0
0
0
0
0
12
0
0
0
0
0
0
0
0
0
0
0
0
0
0
0

1.0
1.0
1.0
1.0
1.0
1.0
1.0
1.0
1.0
1.0
1.0
1.0
1.0
1.0
1.0
1.0
1.0
1.0
1.0
1.0
1.0
1.0
1.0
1.0
1.0
1.0
1.0
1.0
1.0
1.0


7
0
0
0
1
2
4
0
0
0
0
0
0
0
0
0
0
0
0
0
0
0
0
0
0
0
0
0
0
0


7
0
0
0
1
2
4
0
0
0
0
0
0
0
0
0
0
0
0
0
0
0
0
0
0
0
0
0
0
0


7
0
0
0
1
2
4
0
0
0
0
0
0
0
0
0
0
0
0
0
0
0
0
0
0
0
0
0
0
0

1.0
1.0
1.0
1.0
1.0
1.0
1.0
1.0
1.0
1.0
1.0
1.0
1.0
1.0
1.0
1.0
1.0
1.0
1.0
1.0
1.0
1.0
1.0
1.0
1.0
1.0
1.0
1.0
1.0
1.0


48
1
0
0
0
2
0
0
0
3
2
2
0
0
2
0
0
0
0
0
0
0
0
0
0
10
13
10
2
1

0.99
0.99
0.99
0.99
0.99
0.99
0.99
0.99
0.99
0.99
0.99
0.99
0.99
0.99
0.99
0.99
0.99
0.99
0.99
0.99
0.99
0.99
0.99
0.99
0.99
0.99
0.99
0.99
0.99
0.99


28
1
0
0
0
1
0
0
0
3
2
2
0
0
2
0
0
0
0
0
0
0
0
0
0
5
3
7
2
0

1.0
1.0
1.0
1.0
1.0
1.0
1.0
1.0
1.0
1.0
1.0
1.0
1.0
1.0
1.0
1.0
1.0
1.0
1.0
1.0
1.0
1.0
1.0
1.0
1.0
1.0
1.0
1.0
1.0
1.0


13
0
0
0
0
1
0
0
0
0
0
0
0
0
0
0
0
0
0
0
0
0
0
0
0
5
3
4
0
0

1.0
1.0
1.0
1.0
1.0
1.0
1.0
1.0
1.0
1.0
1.0
1.0
1.0
1.0
1.0
1.0
1.0
1.0
1.0
1.0
1.0
1.0
1.0
1.0
1.0
1.0
1.0
1.0
1.0
1.0


19
0
0
0
0
1
0
0
0
0
0
0
0
0
0
0
0
0
0
0
0
0
0
0
0
5
10
2
0
1

1.0
1.0
1.0
1.0
1.0
1.0
1.0
1.0
1.0
1.0
1.0
1.0
1.0
1.0
1.0
1.0
1.0
1.0
1.0
1.0
1.0
1.0
1.0
1.0
1.0
1.0
1.0
1.0
1.0
1.0


1
0
0
0
0
1
0
0
0
0
0
0
0
0
0
0
0
0
0
0
0
0
0
0
0
0
0
0
0
0

0.95
0.95
0.95
0.95
0.95
0.95
0.95
0.95
0.95
0.95
0.95
0.95
0.95
0.95
0.95
0.95
0.95
0.95
0.95
0.95
0.95
0.95
0.95
0.95
0.95
0.95
0.95
0.95
0.95
0.95


7
0
0
0
0
0
0
0
0
0
0
0
0
0
0
0
0
0
0
0
0
0
0
0
0
3
2
2
0
0

1.0
1.0
1.0
1.0
1.0
1.0
1.0
1.0
1.0
1.0
1.0
1.0
1.0
1.0
1.0
1.0
1.0
1.0
1.0
1.0
1.0
1.0
1.0
1.0
1.0
1.0
1.0
1.0
1.0
1.0


2
0
0
0
0
0
0
0
0
0
0
0
0
0
0
0
0
0
0
0
0
0
0
0
0
0
1
0
0
1

0.95
0.95
0.95
0.95
0.95
0.95
0.95
0.95
0.95
0.95
0.95
0.95
0.95
0.95
0.95
0.95
0.95
0.95
0.95
0.95
0.95
0.95
0.95
0.95
0.95
0.95
0.95
0.95
0.95
0.95


1
0
0
0
0
0
0
0
0
0
0
0
0
0
0
0
0
0
0
0
0
0
0
0
0
0
1
0
0
0

0.99
0.99
0.99
0.99
0.99
0.99
0.99
0.99
0.99
0.99
0.99
0.99
0.99
0.99
0.99
0.99
0.99
0.99
0.99
0.99
0.99
0.99
0.99
0.99
0.99
0.99
0.99
0.99
0.99
0.99


15
0
1
0
1
1
0
0
0
0
0
0
0
0
0
4
0
0
0
0
0
0
0
0
0
0
3
5
0
0


15
0
1
0
1
1
0
0
0
0
0
0
0
0
0
4
0
0
0
0
0
0
0
0
0
0
3
5
0
0


15
0
1
0
1
1
0
0
0
0
0
0
0
0
0
4
0
0
0
0
0
0
0
0
0
0
3
5
0
0

1.0
1.0
1.0
1.0
1.0
1.0
1.0
1.0
1.0
1.0
1.0
1.0
1.0
1.0
1.0
1.0
1.0
1.0
1.0
1.0
1.0
1.0
1.0
1.0
1.0
1.0
1.0
1.0
1.0
1.0


21
2
0
0
0
0
0
0
0
0
3
3
0
0
0
0
0
8
0
5
0
0
0
0
0
0
0
0
0
0


9
2
0
0
0
0
0
0
0
0
3
3
0
0
0
0
0
1
0
0
0
0
0
0
0
0
0
0
0
0

1.0
1.0
1.0
1.0
1.0
1.0
1.0
1.0
1.0
1.0
1.0
1.0
1.0
1.0
1.0
1.0
1.0
1.0
1.0
1.0
1.0
1.0
1.0
1.0
1.0
1.0
1.0
1.0
1.0
1.0


8
2
0
0
0
0
0
0
0
0
3
3
0
0
0
0
0
0
0
0
0
0
0
0
0
0
0
0
0
0

1.0
1.0
1.0
1.0
1.0
1.0
1.0
1.0
1.0
1.0
1.0
1.0
1.0
1.0
1.0
1.0
1.0
1.0
1.0
1.0
1.0
1.0
1.0
1.0
1.0
1.0
1.0
1.0
1.0
1.0


5
0
0
0
0
0
0
0
0
0
0
0
0
0
0
0
0
0
0
5
0
0
0
0
0
0
0
0
0
0

0.78
0.78
0.78
0.78
0.78
0.78
0.78
0.78
0.78
0.78
0.78
0.78
0.78
0.78
0.78
0.78
0.78
0.78
0.78
0.78
0.78
0.78
0.78
0.78
0.78
0.78
0.78
0.78
0.78
0.78


1
0
0
0
0
0
0
0
0
0
0
0
0
0
0
0
0
0
0
1
0
0
0
0
0
0
0
0
0
0

0.74
0.74
0.74
0.74
0.74
0.74
0.74
0.74
0.74
0.74
0.74
0.74
0.74
0.74
0.74
0.74
0.74
0.74
0.74
0.74
0.74
0.74
0.74
0.74
0.74
0.74
0.74
0.74
0.74
0.74


7
0
0
0
0
0
0
0
0
0
0
0
0
0
0
0
0
7
0
0
0
0
0
0
0
0
0
0
0
0


7
0
0
0
0
0
0
0
0
0
0
0
0
0
0
0
0
7
0
0
0
0
0
0
0
0
0
0
0
0


7
0
0
0
0
0
0
0
0
0
0
0
0
0
0
0
0
7
0
0
0
0
0
0
0
0
0
0
0
0

0.96
0.96
0.96
0.96
0.96
0.96
0.96
0.96
0.96
0.96
0.96
0.96
0.96
0.96
0.96
0.96
0.96
0.96
0.96
0.96
0.96
0.96
0.96
0.96
0.96
0.96
0.96
0.96
0.96
0.96


48
1
1
1
0
0
1
0
0
0
0
1
0
0
0
0
11
0
0
22
10
0
0
0
0
0
0
0
0
0

0.94
0.94
0.94
0.94
0.94
0.94
0.94
0.94
0.94
0.94
0.94
0.94
0.94
0.94
0.94
0.94
0.94
0.94
0.94
0.94
0.94
0.94
0.94
0.94
0.94
0.94
0.94
0.94
0.94
0.94


37
1
1
1
0
0
1
0
0
0
0
1
0
0
0
0
0
0
0
22
10
0
0
0
0
0
0
0
0
0


37
1
1
1
0
0
1
0
0
0
0
1
0
0
0
0
0
0
0
22
10
0
0
0
0
0
0
0
0
0

1.0
1.0
1.0
1.0
1.0
1.0
1.0
1.0
1.0
1.0
1.0
1.0
1.0
1.0
1.0
1.0
1.0
1.0
1.0
1.0
1.0
1.0
1.0
1.0
1.0
1.0
1.0
1.0
1.0
1.0


20
1
1
0
0
0
1
0
0
0
0
1
0
0
0
0
0
0
0
16
0
0
0
0
0
0
0
0
0
0

1.0
1.0
1.0
1.0
1.0
1.0
1.0
1.0
1.0
1.0
1.0
1.0
1.0
1.0
1.0
1.0
1.0
1.0
1.0
1.0
1.0
1.0
1.0
1.0
1.0
1.0
1.0
1.0
1.0
1.0


10
0
0
1
0
0
0
0
0
0
0
0
0
0
0
0
0
0
0
0
9
0
0
0
0
0
0
0
0
0

1.0
1.0
1.0
1.0
1.0
1.0
1.0
1.0
1.0
1.0
1.0
1.0
1.0
1.0
1.0
1.0
1.0
1.0
1.0
1.0
1.0
1.0
1.0
1.0
1.0
1.0
1.0
1.0
1.0
1.0


10
0
0
0
0
0
0
0
0
0
0
0
0
0
0
0
10
0
0
0
0
0
0
0
0
0
0
0
0
0


10
0
0
0
0
0
0
0
0
0
0
0
0
0
0
0
10
0
0
0
0
0
0
0
0
0
0
0
0
0


10
0
0
0
0
0
0
0
0
0
0
0
0
0
0
0
10
0
0
0
0
0
0
0
0
0
0
0
0
0

1.0
1.0
1.0
1.0
1.0
1.0
1.0
1.0
1.0
1.0
1.0
1.0
1.0
1.0
1.0
1.0
1.0
1.0
1.0
1.0
1.0
1.0
1.0
1.0
1.0
1.0
1.0
1.0
1.0
1.0


47
7
2
3
4
0
1
4
0
6
1
6
0
0
1
0
0
0
1
0
5
1
1
0
3
1
0
0
0
0


7
0
0
0
0
0
0
2
0
1
0
0
0
0
0
0
0
0
1
0
1
1
1
0
0
0
0
0
0
0


6
0
0
0
0
0
0
1
0
1
0
0
0
0
0
0
0
0
1
0
1
1
1
0
0
0
0
0
0
0


6
0
0
0
0
0
0
1
0
1
0
0
0
0
0
0
0
0
1
0
1
1
1
0
0
0
0
0
0
0

1.0
1.0
1.0
1.0
1.0
1.0
1.0
1.0
1.0
1.0
1.0
1.0
1.0
1.0
1.0
1.0
1.0
1.0
1.0
1.0
1.0
1.0
1.0
1.0
1.0
1.0
1.0
1.0
1.0
1.0


1
0
0
0
0
0
0
1
0
0
0
0
0
0
0
0
0
0
0
0
0
0
0
0
0
0
0
0
0
0


1
0
0
0
0
0
0
1
0
0
0
0
0
0
0
0
0
0
0
0
0
0
0
0
0
0
0
0
0
0

1.0
1.0
1.0
1.0
1.0
1.0
1.0
1.0
1.0
1.0
1.0
1.0
1.0
1.0
1.0
1.0
1.0
1.0
1.0
1.0
1.0
1.0
1.0
1.0
1.0
1.0
1.0
1.0
1.0
1.0


8
1
0
0
0
0
0
0
0
1
0
2
0
0
0
0
0
0
0
0
4
0
0
0
0
0
0
0
0
0


4
0
0
0
0
0
0
0
0
1
0
1
0
0
0
0
0
0
0
0
2
0
0
0
0
0
0
0
0
0

1.0
1.0
1.0
1.0
1.0
1.0
1.0
1.0
1.0
1.0
1.0
1.0
1.0
1.0
1.0
1.0
1.0
1.0
1.0
1.0
1.0
1.0
1.0
1.0
1.0
1.0
1.0
1.0
1.0
1.0


3
0
0
0
0
0
0
0
0
1
0
1
0
0
0
0
0
0
0
0
1
0
0
0
0
0
0
0
0
0

1.0
1.0
1.0
1.0
1.0
1.0
1.0
1.0
1.0
1.0
1.0
1.0
1.0
1.0
1.0
1.0
1.0
1.0
1.0
1.0
1.0
1.0
1.0
1.0
1.0
1.0
1.0
1.0
1.0
1.0


4
1
0
0
0
0
0
0
0
0
0
1
0
0
0
0
0
0
0
0
2
0
0
0
0
0
0
0
0
0


4
1
0
0
0
0
0
0
0
0
0
1
0
0
0
0
0
0
0
0
2
0
0
0
0
0
0
0
0
0

1.0
1.0
1.0
1.0
1.0
1.0
1.0
1.0
1.0
1.0
1.0
1.0
1.0
1.0
1.0
1.0
1.0
1.0
1.0
1.0
1.0
1.0
1.0
1.0
1.0
1.0
1.0
1.0
1.0
1.0


29
6
2
3
4
0
1
2
0
4
1
4
0
0
1
0
0
0
0
0
0
0
0
0
1
0
0
0
0
0


29
6
2
3
4
0
1
2
0
4
1
4
0
0
1
0
0
0
0
0
0
0
0
0
1
0
0
0
0
0


27
6
2
3
4
0
1
2
0
4
1
3
0
0
1
0
0
0
0
0
0
0
0
0
0
0
0
0
0
0

1.0
1.0
1.0
1.0
1.0
1.0
1.0
1.0
1.0
1.0
1.0
1.0
1.0
1.0
1.0
1.0
1.0
1.0
1.0
1.0
1.0
1.0
1.0
1.0
1.0
1.0
1.0
1.0
1.0
1.0


2
0
0
0
0
0
0
0
0
0
0
1
0
0
0
0
0
0
0
0
0
0
0
0
1
0
0
0
0
0

1.0
1.0
1.0
1.0
1.0
1.0
1.0
1.0
1.0
1.0
1.0
1.0
1.0
1.0
1.0
1.0
1.0
1.0
1.0
1.0
1.0
1.0
1.0
1.0
1.0
1.0
1.0
1.0
1.0
1.0


1
0
0
0
0
0
0
0
0
0
0
0
0
0
0
0
0
0
0
0
0
0
0
0
1
0
0
0
0
0


1
0
0
0
0
0
0
0
0
0
0
0
0
0
0
0
0
0
0
0
0
0
0
0
1
0
0
0
0
0


1
0
0
0
0
0
0
0
0
0
0
0
0
0
0
0
0
0
0
0
0
0
0
0
1
0
0
0
0
0

1.0
1.0
1.0
1.0
1.0
1.0
1.0
1.0
1.0
1.0
1.0
1.0
1.0
1.0
1.0
1.0
1.0
1.0
1.0
1.0
1.0
1.0
1.0
1.0
1.0
1.0
1.0
1.0
1.0
1.0


1
0
0
0
0
0
0
0
0
0
0
0
0
0
0
0
0
0
0
0
0
0
0
0
1
0
0
0
0
0


1
0
0
0
0
0
0
0
0
0
0
0
0
0
0
0
0
0
0
0
0
0
0
0
1
0
0
0
0
0


1
0
0
0
0
0
0
0
0
0
0
0
0
0
0
0
0
0
0
0
0
0
0
0
1
0
0
0
0
0

1.0
1.0
1.0
1.0
1.0
1.0
1.0
1.0
1.0
1.0
1.0
1.0
1.0
1.0
1.0
1.0
1.0
1.0
1.0
1.0
1.0
1.0
1.0
1.0
1.0
1.0
1.0
1.0
1.0
1.0


1
0
0
0
0
0
0
0
0
0
0
0
0
0
0
0
0
0
0
0
0
0
0
0
0
1
0
0
0
0


1
0
0
0
0
0
0
0
0
0
0
0
0
0
0
0
0
0
0
0
0
0
0
0
0
1
0
0
0
0


1
0
0
0
0
0
0
0
0
0
0
0
0
0
0
0
0
0
0
0
0
0
0
0
0
1
0
0
0
0

1.0
1.0
1.0
1.0
1.0
1.0
1.0
1.0
1.0
1.0
1.0
1.0
1.0
1.0
1.0
1.0
1.0
1.0
1.0
1.0
1.0
1.0
1.0
1.0
1.0
1.0
1.0
1.0
1.0
1.0


32
4
1
0
0
0
0
1
3
5
5
1
1
0
0
0
0
0
0
0
0
1
0
1
0
1
2
2
2
2


13
2
0
0
0
0
0
0
2
3
3
0
1
0
0
0
0
0
0
0
0
1
0
1
0
0
0
0
0
0

0.81
0.81
0.81
0.81
0.81
0.81
0.81
0.81
0.81
0.81
0.81
0.81
0.81
0.81
0.81
0.81
0.81
0.81
0.81
0.81
0.81
0.81
0.81
0.81
0.81
0.81
0.81
0.81
0.81
0.81


4
1
0
0
0
0
0
0
1
1
1
0
0
0
0
0
0
0
0
0
0
0
0
0
0
0
0
0
0
0

0.99
0.99
0.99
0.99
0.99
0.99
0.99
0.99
0.99
0.99
0.99
0.99
0.99
0.99
0.99
0.99
0.99
0.99
0.99
0.99
0.99
0.99
0.99
0.99
0.99
0.99
0.99
0.99
0.99
0.99


6
1
0
0
0
0
0
0
1
2
2
0
0
0
0
0
0
0
0
0
0
0
0
0
0
0
0
0
0
0


2
0
0
0
0
0
0
0
0
1
1
0
0
0
0
0
0
0
0
0
0
0
0
0
0
0
0
0
0
0

1.0
1.0
1.0
1.0
1.0
1.0
1.0
1.0
1.0
1.0
1.0
1.0
1.0
1.0
1.0
1.0
1.0
1.0
1.0
1.0
1.0
1.0
1.0
1.0
1.0
1.0
1.0
1.0
1.0
1.0


4
1
0
0
0
0
0
0
1
1
1
0
0
0
0
0
0
0
0
0
0
0
0
0
0
0
0
0
0
0

1.0
1.0
1.0
1.0
1.0
1.0
1.0
1.0
1.0
1.0
1.0
1.0
1.0
1.0
1.0
1.0
1.0
1.0
1.0
1.0
1.0
1.0
1.0
1.0
1.0
1.0
1.0
1.0
1.0
1.0


8
1
0
0
0
0
0
1
1
2
2
1
0
0
0
0
0
0
0
0
0
0
0
0
0
0
0
0
0
0


8
1
0
0
0
0
0
1
1
2
2
1
0
0
0
0
0
0
0
0
0
0
0
0
0
0
0
0
0
0


8
1
0
0
0
0
0
1
1
2
2
1
0
0
0
0
0
0
0
0
0
0
0
0
0
0
0
0
0
0

1.0
1.0
1.0
1.0
1.0
1.0
1.0
1.0
1.0
1.0
1.0
1.0
1.0
1.0
1.0
1.0
1.0
1.0
1.0
1.0
1.0
1.0
1.0
1.0
1.0
1.0
1.0
1.0
1.0
1.0


2
1
1
0
0
0
0
0
0
0
0
0
0
0
0
0
0
0
0
0
0
0
0
0
0
0
0
0
0
0


2
1
1
0
0
0
0
0
0
0
0
0
0
0
0
0
0
0
0
0
0
0
0
0
0
0
0
0
0
0


2
1
1
0
0
0
0
0
0
0
0
0
0
0
0
0
0
0
0
0
0
0
0
0
0
0
0
0
0
0

1.0
1.0
1.0
1.0
1.0
1.0
1.0
1.0
1.0
1.0
1.0
1.0
1.0
1.0
1.0
1.0
1.0
1.0
1.0
1.0
1.0
1.0
1.0
1.0
1.0
1.0
1.0
1.0
1.0
1.0


9
0
0
0
0
0
0
0
0
0
0
0
0
0
0
0
0
0
0
0
0
0
0
0
0
1
2
2
2
2


5
0
0
0
0
0
0
0
0
0
0
0
0
0
0
0
0
0
0
0
0
0
0
0
0
1
1
1
1
1

0.97
0.97
0.97
0.97
0.97
0.97
0.97
0.97
0.97
0.97
0.97
0.97
0.97
0.97
0.97
0.97
0.97
0.97
0.97
0.97
0.97
0.97
0.97
0.97
0.97
0.97
0.97
0.97
0.97
0.97


4
0
0
0
0
0
0
0
0
0
0
0
0
0
0
0
0
0
0
0
0
0
0
0
0
0
1
1
1
1


4
0
0
0
0
0
0
0
0
0
0
0
0
0
0
0
0
0
0
0
0
0
0
0
0
0
1
1
1
1

1.0
1.0
1.0
1.0
1.0
1.0
1.0
1.0
1.0
1.0
1.0
1.0
1.0
1.0
1.0
1.0
1.0
1.0
1.0
1.0
1.0
1.0
1.0
1.0
1.0
1.0
1.0
1.0
1.0
1.0


1
1
0
0
0
0
0
0
0
0
0
0
0
0
0
0
0
0
0
0
0
0
0
0
0
0
0
0
0
0


1
1
0
0
0
0
0
0
0
0
0
0
0
0
0
0
0
0
0
0
0
0
0
0
0
0
0
0
0
0


1
1
0
0
0
0
0
0
0
0
0
0
0
0
0
0
0
0
0
0
0
0
0
0
0
0
0
0
0
0


1
1
0
0
0
0
0
0
0
0
0
0
0
0
0
0
0
0
0
0
0
0
0
0
0
0
0
0
0
0

0.95
0.95
0.95
0.95
0.95
0.95
0.95
0.95
0.95
0.95
0.95
0.95
0.95
0.95
0.95
0.95
0.95
0.95
0.95
0.95
0.95
0.95
0.95
0.95
0.95
0.95
0.95
0.95
0.95
0.95


1
0
0
0
0
0
0
0
0
0
1
0
0
0
0
0
0
0
0
0
0
0
0
0
0
0
0
0
0
0


1
0
0
0
0
0
0
0
0
0
1
0
0
0
0
0
0
0
0
0
0
0
0
0
0
0
0
0
0
0


1
0
0
0
0
0
0
0
0
0
1
0
0
0
0
0
0
0
0
0
0
0
0
0
0
0
0
0
0
0


1
0
0
0
0
0
0
0
0
0
1
0
0
0
0
0
0
0
0
0
0
0
0
0
0
0
0
0
0
0

1.0
1.0
1.0
1.0
1.0
1.0
1.0
1.0
1.0
1.0
1.0
1.0
1.0
1.0
1.0
1.0
1.0
1.0
1.0
1.0
1.0
1.0
1.0
1.0
1.0
1.0
1.0
1.0
1.0
1.0


1
0
0
0
0
0
0
0
0
1
0
0
0
0
0
0
0
0
0
0
0
0
0
0
0
0
0
0
0
0


1
0
0
0
0
0
0
0
0
1
0
0
0
0
0
0
0
0
0
0
0
0
0
0
0
0
0
0
0
0


1
0
0
0
0
0
0
0
0
1
0
0
0
0
0
0
0
0
0
0
0
0
0
0
0
0
0
0
0
0

0.93
0.93
0.93
0.93
0.93
0.93
0.93
0.93
0.93
0.93
0.93
0.93
0.93
0.93
0.93
0.93
0.93
0.93
0.93
0.93
0.93
0.93
0.93
0.93
0.93
0.93
0.93
0.93
0.93
0.93


89
5
4
3
2
0
0
4
2
4
4
5
5
1
0
1
0
0
0
5
6
3
12
5
6
0
3
5
1
3

0.74
0.74
0.74
0.74
0.74
0.74
0.74
0.74
0.74
0.74
0.74
0.74
0.74
0.74
0.74
0.74
0.74
0.74
0.74
0.74
0.74
0.74
0.74
0.74
0.74
0.74
0.74
0.74
0.74
0.74


84
5
4
3
2
0
0
4
2
4
4
5
4
1
0
1
0
0
0
5
6
3
12
4
6
0
2
4
1
2

0.9
0.9
0.9
0.9
0.9
0.9
0.9
0.9
0.9
0.9
0.9
0.9
0.9
0.9
0.9
0.9
0.9
0.9
0.9
0.9
0.9
0.9
0.9
0.9
0.9
0.9
0.9
0.9
0.9
0.9


75
4
3
3
2
0
0
3
2
3
4
4
3
1
0
1
0
0
0
5
6
3
10
3
6
0
2
4
1
2

0.79
0.79
0.79
0.79
0.79
0.79
0.79
0.79
0.79
0.79
0.79
0.79
0.79
0.79
0.79
0.79
0.79
0.79
0.79
0.79
0.79
0.79
0.79
0.79
0.79
0.79
0.79
0.79
0.79
0.79


74
4
3
3
2
0
0
3
2
3
4
4
3
1
0
1
0
0
0
5
6
3
10
3
5
0
2
4
1
2

0.95
0.95
0.95
0.95
0.95
0.95
0.95
0.95
0.95
0.95
0.95
0.95
0.95
0.95
0.95
0.95
0.95
0.95
0.95
0.95
0.95
0.95
0.95
0.95
0.95
0.95
0.95
0.95
0.95
0.95


54
4
3
3
2
0
0
3
2
2
4
4
2
0
0
0
0
0
0
4
4
3
7
2
5
0
0
0
0
0

1.0
1.0
1.0
1.0
1.0
1.0
1.0
1.0
1.0
1.0
1.0
1.0
1.0
1.0
1.0
1.0
1.0
1.0
1.0
1.0
1.0
1.0
1.0
1.0
1.0
1.0
1.0
1.0
1.0
1.0


9
0
0
0
0
0
0
0
0
0
0
0
0
0
0
1
0
0
0
0
0
0
0
0
0
0
2
3
1
2

0.94
0.94
0.94
0.94
0.94
0.94
0.94
0.94
0.94
0.94
0.94
0.94
0.94
0.94
0.94
0.94
0.94
0.94
0.94
0.94
0.94
0.94
0.94
0.94
0.94
0.94
0.94
0.94
0.94
0.94


1
0
0
0
0
0
0
0
0
0
0
0
0
1
0
0
0
0
0
0
0
0
0
0
0
0
0
0
0
0

0.86
0.86
0.86
0.86
0.86
0.86
0.86
0.86
0.86
0.86
0.86
0.86
0.86
0.86
0.86
0.86
0.86
0.86
0.86
0.86
0.86
0.86
0.86
0.86
0.86
0.86
0.86
0.86
0.86
0.86


1
0
0
0
0
0
0
0
0
0
0
0
0
0
0
0
0
0
0
0
1
0
0
0
0
0
0
0
0
0

0.72
0.72
0.72
0.72
0.72
0.72
0.72
0.72
0.72
0.72
0.72
0.72
0.72
0.72
0.72
0.72
0.72
0.72
0.72
0.72
0.72
0.72
0.72
0.72
0.72
0.72
0.72
0.72
0.72
0.72


1
0
0
0
0
0
0
0
0
0
0
0
0
0
0
0
0
0
0
0
0
0
1
0
0
0
0
0
0
0

0.82
0.82
0.82
0.82
0.82
0.82
0.82
0.82
0.82
0.82
0.82
0.82
0.82
0.82
0.82
0.82
0.82
0.82
0.82
0.82
0.82
0.82
0.82
0.82
0.82
0.82
0.82
0.82
0.82
0.82


5
1
1
0
0
0
0
1
0
1
0
1
0
0
0
0
0
0
0
0
0
0
0
0
0
0
0
0
0
0


5
1
1
0
0
0
0
1
0
1
0
1
0
0
0
0
0
0
0
0
0
0
0
0
0
0
0
0
0
0


5
1
1
0
0
0
0
1
0
1
0
1
0
0
0
0
0
0
0
0
0
0
0
0
0
0
0
0
0
0

0.82
0.82
0.82
0.82
0.82
0.82
0.82
0.82
0.82
0.82
0.82
0.82
0.82
0.82
0.82
0.82
0.82
0.82
0.82
0.82
0.82
0.82
0.82
0.82
0.82
0.82
0.82
0.82
0.82
0.82


196
9
7
11
8
5
7
12
6
7
12
15
4
5
12
10
1
5
4
2
6
2
3
4
14
5
6
5
7
2

0.72
0.72
0.72
0.72
0.72
0.72
0.72
0.72
0.72
0.72
0.72
0.72
0.72
0.72
0.72
0.72
0.72
0.72
0.72
0.72
0.72
0.72
0.72
0.72
0.72
0.72
0.72
0.72
0.72
0.72


180
7
7
11
8
5
7
11
6
7
9
13
4
4
12
10
1
5
3
1
5
1
2
4
13
5
6
5
6
2

1.0
1.0
1.0
1.0
1.0
1.0
1.0
1.0
1.0
1.0
1.0
1.0
1.0
1.0
1.0
1.0
1.0
1.0
1.0
1.0
1.0
1.0
1.0
1.0
1.0
1.0
1.0
1.0
1.0
1.0


22
2
2
2
1
1
0
1
0
1
1
2
1
1
1
0
0
1
0
0
1
0
0
1
3
0
0
0
0
0

0.72
0.72
0.72
0.72
0.72
0.72
0.72
0.72
0.72
0.72
0.72
0.72
0.72
0.72
0.72
0.72
0.72
0.72
0.72
0.72
0.72
0.72
0.72
0.72
0.72
0.72
0.72
0.72
0.72
0.72


16
2
2
2
1
1
0
1
0
1
1
2
0
1
0
0
0
0
0
0
0
0
0
0
2
0
0
0
0
0

0.8
0.8
0.8
0.8
0.8
0.8
0.8
0.8
0.8
0.8
0.8
0.8
0.8
0.8
0.8
0.8
0.8
0.8
0.8
0.8
0.8
0.8
0.8
0.8
0.8
0.8
0.8
0.8
0.8
0.8


7
1
1
1
0
0
0
0
0
0
0
1
0
1
0
0
0
0
0
0
0
0
0
0
2
0
0
0
0
0

1.0
1.0
1.0
1.0
1.0
1.0
1.0
1.0
1.0
1.0
1.0
1.0
1.0
1.0
1.0
1.0
1.0
1.0
1.0
1.0
1.0
1.0
1.0
1.0
1.0
1.0
1.0
1.0
1.0
1.0


31
1
3
2
1
1
2
2
1
2
1
2
0
0
5
0
0
0
0
0
0
0
0
0
5
0
0
0
3
0


27
1
2
1
1
1
2
2
1
2
1
1
0
0
4
0
0
0
0
0
0
0
0
0
5
0
0
0
3
0


23
1
2
1
1
1
2
2
1
2
1
1
0
0
4
0
0
0
0
0
0
0
0
0
2
0
0
0
2
0

1.0
1.0
1.0
1.0
1.0
1.0
1.0
1.0
1.0
1.0
1.0
1.0
1.0
1.0
1.0
1.0
1.0
1.0
1.0
1.0
1.0
1.0
1.0
1.0
1.0
1.0
1.0
1.0
1.0
1.0


4
0
0
0
0
0
0
0
0
0
0
0
0
0
0
0
0
0
0
0
0
0
0
0
3
0
0
0
1
0

0.9
0.9
0.9
0.9
0.9
0.9
0.9
0.9
0.9
0.9
0.9
0.9
0.9
0.9
0.9
0.9
0.9
0.9
0.9
0.9
0.9
0.9
0.9
0.9
0.9
0.9
0.9
0.9
0.9
0.9


4
0
1
1
0
0
0
0
0
0
0
1
0
0
1
0
0
0
0
0
0
0
0
0
0
0
0
0
0
0


4
0
1
1
0
0
0
0
0
0
0
1
0
0
1
0
0
0
0
0
0
0
0
0
0
0
0
0
0
0

1.0
1.0
1.0
1.0
1.0
1.0
1.0
1.0
1.0
1.0
1.0
1.0
1.0
1.0
1.0
1.0
1.0
1.0
1.0
1.0
1.0
1.0
1.0
1.0
1.0
1.0
1.0
1.0
1.0
1.0


25
1
0
1
3
1
3
2
2
1
0
3
0
1
2
0
0
1
0
0
1
0
0
0
0
1
0
1
0
1


6
1
0
0
0
0
0
0
1
1
0
1
0
0
1
0
0
1
0
0
0
0
0
0
0
0
0
0
0
0


6
1
0
0
0
0
0
0
1
1
0
1
0
0
1
0
0
1
0
0
0
0
0
0
0
0
0
0
0
0

0.93
0.93
0.93
0.93
0.93
0.93
0.93
0.93
0.93
0.93
0.93
0.93
0.93
0.93
0.93
0.93
0.93
0.93
0.93
0.93
0.93
0.93
0.93
0.93
0.93
0.93
0.93
0.93
0.93
0.93


7
0
0
0
1
1
2
2
0
0
0
0
0
0
0
0
0
0
0
0
0
0
0
0
0
1
0
0
0
0


7
0
0
0
1
1
2
2
0
0
0
0
0
0
0
0
0
0
0
0
0
0
0
0
0
1
0
0
0
0

1.0
1.0
1.0
1.0
1.0
1.0
1.0
1.0
1.0
1.0
1.0
1.0
1.0
1.0
1.0
1.0
1.0
1.0
1.0
1.0
1.0
1.0
1.0
1.0
1.0
1.0
1.0
1.0
1.0
1.0


5
0
0
0
2
0
0
0
0
0
0
0
0
1
1
0
0
0
0
0
1
0
0
0
0
0
0
0
0
0

0.82
0.82
0.82
0.82
0.82
0.82
0.82
0.82
0.82
0.82
0.82
0.82
0.82
0.82
0.82
0.82
0.82
0.82
0.82
0.82
0.82
0.82
0.82
0.82
0.82
0.82
0.82
0.82
0.82
0.82


2
0
0
0
0
0
0
0
0
0
0
0
0
1
0
0
0
0
0
0
1
0
0
0
0
0
0
0
0
0

0.95
0.95
0.95
0.95
0.95
0.95
0.95
0.95
0.95
0.95
0.95
0.95
0.95
0.95
0.95
0.95
0.95
0.95
0.95
0.95
0.95
0.95
0.95
0.95
0.95
0.95
0.95
0.95
0.95
0.95


5
0
0
1
0
0
1
0
1
0
0
1
0
0
0
0
0
0
0
0
0
0
0
0
0
0
0
0
0
1

0.94
0.94
0.94
0.94
0.94
0.94
0.94
0.94
0.94
0.94
0.94
0.94
0.94
0.94
0.94
0.94
0.94
0.94
0.94
0.94
0.94
0.94
0.94
0.94
0.94
0.94
0.94
0.94
0.94
0.94


1
0
0
0
0
0
0
0
0
0
0
0
0
0
0
0
0
0
0
0
0
0
0
0
0
0
0
0
0
1

0.93
0.93
0.93
0.93
0.93
0.93
0.93
0.93
0.93
0.93
0.93
0.93
0.93
0.93
0.93
0.93
0.93
0.93
0.93
0.93
0.93
0.93
0.93
0.93
0.93
0.93
0.93
0.93
0.93
0.93


1
0
0
0
0
0
0
0
0
0
0
1
0
0
0
0
0
0
0
0
0
0
0
0
0
0
0
0
0
0


1
0
0
0
0
0
0
0
0
0
0
1
0
0
0
0
0
0
0
0
0
0
0
0
0
0
0
0
0
0

1.0
1.0
1.0
1.0
1.0
1.0
1.0
1.0
1.0
1.0
1.0
1.0
1.0
1.0
1.0
1.0
1.0
1.0
1.0
1.0
1.0
1.0
1.0
1.0
1.0
1.0
1.0
1.0
1.0
1.0


1
0
0
0
0
0
0
0
0
0
0
0
0
0
0
0
0
0
0
0
0
0
0
0
0
0
0
1
0
0

1.0
1.0
1.0
1.0
1.0
1.0
1.0
1.0
1.0
1.0
1.0
1.0
1.0
1.0
1.0
1.0
1.0
1.0
1.0
1.0
1.0
1.0
1.0
1.0
1.0
1.0
1.0
1.0
1.0
1.0


16
0
0
0
1
1
0
3
1
2
3
1
1
0
0
0
0
0
0
0
0
0
0
1
0
1
1
0
0
0

0.85
0.85
0.85
0.85
0.85
0.85
0.85
0.85
0.85
0.85
0.85
0.85
0.85
0.85
0.85
0.85
0.85
0.85
0.85
0.85
0.85
0.85
0.85
0.85
0.85
0.85
0.85
0.85
0.85
0.85


6
0
0
0
0
0
0
2
0
0
1
1
1
0
0
0
0
0
0
0
0
0
0
1
0
0
0
0
0
0


6
0
0
0
0
0
0
2
0
0
1
1
1
0
0
0
0
0
0
0
0
0
0
1
0
0
0
0
0
0

1.0
1.0
1.0
1.0
1.0
1.0
1.0
1.0
1.0
1.0
1.0
1.0
1.0
1.0
1.0
1.0
1.0
1.0
1.0
1.0
1.0
1.0
1.0
1.0
1.0
1.0
1.0
1.0
1.0
1.0


4
0
0
0
1
1
0
0
0
0
1
0
0
0
0
0
0
0
0
0
0
0
0
0
0
0
1
0
0
0


4
0
0
0
1
1
0
0
0
0
1
0
0
0
0
0
0
0
0
0
0
0
0
0
0
0
1
0
0
0

1.0
1.0
1.0
1.0
1.0
1.0
1.0
1.0
1.0
1.0
1.0
1.0
1.0
1.0
1.0
1.0
1.0
1.0
1.0
1.0
1.0
1.0
1.0
1.0
1.0
1.0
1.0
1.0
1.0
1.0


1
0
0
0
0
0
0
0
0
0
0
0
0
0
0
0
0
0
0
0
0
0
0
0
0
1
0
0
0
0


1
0
0
0
0
0
0
0
0
0
0
0
0
0
0
0
0
0
0
0
0
0
0
0
0
1
0
0
0
0

1.0
1.0
1.0
1.0
1.0
1.0
1.0
1.0
1.0
1.0
1.0
1.0
1.0
1.0
1.0
1.0
1.0
1.0
1.0
1.0
1.0
1.0
1.0
1.0
1.0
1.0
1.0
1.0
1.0
1.0


10
1
0
2
0
0
0
2
0
0
0
0
0
0
1
0
0
1
1
0
0
0
0
0
2
0
0
0
0
0

0.98
0.98
0.98
0.98
0.98
0.98
0.98
0.98
0.98
0.98
0.98
0.98
0.98
0.98
0.98
0.98
0.98
0.98
0.98
0.98
0.98
0.98
0.98
0.98
0.98
0.98
0.98
0.98
0.98
0.98


3
1
0
0
0
0
0
1
0
0
0
0
0
0
1
0
0
0
0
0
0
0
0
0
0
0
0
0
0
0


3
1
0
0
0
0
0
1
0
0
0
0
0
0
1
0
0
0
0
0
0
0
0
0
0
0
0
0
0
0

1.0
1.0
1.0
1.0
1.0
1.0
1.0
1.0
1.0
1.0
1.0
1.0
1.0
1.0
1.0
1.0
1.0
1.0
1.0
1.0
1.0
1.0
1.0
1.0
1.0
1.0
1.0
1.0
1.0
1.0


1
0
0
1
0
0
0
0
0
0
0
0
0
0
0
0
0
0
0
0
0
0
0
0
0
0
0
0
0
0


1
0
0
1
0
0
0
0
0
0
0
0
0
0
0
0
0
0
0
0
0
0
0
0
0
0
0
0
0
0

1.0
1.0
1.0
1.0
1.0
1.0
1.0
1.0
1.0
1.0
1.0
1.0
1.0
1.0
1.0
1.0
1.0
1.0
1.0
1.0
1.0
1.0
1.0
1.0
1.0
1.0
1.0
1.0
1.0
1.0


4
0
0
1
0
0
0
0
0
0
0
0
0
0
0
0
0
0
1
0
0
0
0
0
2
0
0
0
0
0


3
0
0
1
0
0
0
0
0
0
0
0
0
0
0
0
0
0
0
0
0
0
0
0
2
0
0
0
0
0

1.0
1.0
1.0
1.0
1.0
1.0
1.0
1.0
1.0
1.0
1.0
1.0
1.0
1.0
1.0
1.0
1.0
1.0
1.0
1.0
1.0
1.0
1.0
1.0
1.0
1.0
1.0
1.0
1.0
1.0


1
0
0
0
0
0
0
0
0
0
0
0
0
0
0
0
0
0
1
0
0
0
0
0
0
0
0
0
0
0

1.0
1.0
1.0
1.0
1.0
1.0
1.0
1.0
1.0
1.0
1.0
1.0
1.0
1.0
1.0
1.0
1.0
1.0
1.0
1.0
1.0
1.0
1.0
1.0
1.0
1.0
1.0
1.0
1.0
1.0


1
0
0
0
0
0
0
0
0
0
0
0
0
0
0
0
0
1
0
0
0
0
0
0
0
0
0
0
0
0


1
0
0
0
0
0
0
0
0
0
0
0
0
0
0
0
0
1
0
0
0
0
0
0
0
0
0
0
0
0

1.0
1.0
1.0
1.0
1.0
1.0
1.0
1.0
1.0
1.0
1.0
1.0
1.0
1.0
1.0
1.0
1.0
1.0
1.0
1.0
1.0
1.0
1.0
1.0
1.0
1.0
1.0
1.0
1.0
1.0


9
0
0
0
0
0
0
1
0
0
0
0
1
0
1
0
0
1
1
1
1
0
0
1
1
0
0
0
0
0

1.0
1.0
1.0
1.0
1.0
1.0
1.0
1.0
1.0
1.0
1.0
1.0
1.0
1.0
1.0
1.0
1.0
1.0
1.0
1.0
1.0
1.0
1.0
1.0
1.0
1.0
1.0
1.0
1.0
1.0


3
0
0
0
0
0
0
1
0
0
0
0
0
0
0
0
0
1
0
0
0
0
0
0
1
0
0
0
0
0


1
0
0
0
0
0
0
1
0
0
0
0
0
0
0
0
0
0
0
0
0
0
0
0
0
0
0
0
0
0

1.0
1.0
1.0
1.0
1.0
1.0
1.0
1.0
1.0
1.0
1.0
1.0
1.0
1.0
1.0
1.0
1.0
1.0
1.0
1.0
1.0
1.0
1.0
1.0
1.0
1.0
1.0
1.0
1.0
1.0


2
0
0
0
0
0
0
0
0
0
0
0
0
0
0
0
0
1
0
0
0
0
0
0
1
0
0
0
0
0

0.79
0.79
0.79
0.79
0.79
0.79
0.79
0.79
0.79
0.79
0.79
0.79
0.79
0.79
0.79
0.79
0.79
0.79
0.79
0.79
0.79
0.79
0.79
0.79
0.79
0.79
0.79
0.79
0.79
0.79


1
0
0
0
0
0
0
0
0
0
0
0
0
0
0
0
0
0
1
0
0
0
0
0
0
0
0
0
0
0


1
0
0
0
0
0
0
0
0
0
0
0
0
0
0
0
0
0
1
0
0
0
0
0
0
0
0
0
0
0

1.0
1.0
1.0
1.0
1.0
1.0
1.0
1.0
1.0
1.0
1.0
1.0
1.0
1.0
1.0
1.0
1.0
1.0
1.0
1.0
1.0
1.0
1.0
1.0
1.0
1.0
1.0
1.0
1.0
1.0


2
0
0
1
0
0
0
0
0
0
1
0
0
0
0
0
0
0
0
0
0
0
0
0
0
0
0
0
0
0


2
0
0
1
0
0
0
0
0
0
1
0
0
0
0
0
0
0
0
0
0
0
0
0
0
0
0
0
0
0


2
0
0
1
0
0
0
0
0
0
1
0
0
0
0
0
0
0
0
0
0
0
0
0
0
0
0
0
0
0

1.0
1.0
1.0
1.0
1.0
1.0
1.0
1.0
1.0
1.0
1.0
1.0
1.0
1.0
1.0
1.0
1.0
1.0
1.0
1.0
1.0
1.0
1.0
1.0
1.0
1.0
1.0
1.0
1.0
1.0


7
0
0
0
0
0
0
0
1
0
1
2
0
1
0
0
0
0
0
0
0
0
1
0
0
0
0
0
1
0

0.7
0.7
0.7
0.7
0.7
0.7
0.7
0.7
0.7
0.7
0.7
0.7
0.7
0.7
0.7
0.7
0.7
0.7
0.7
0.7
0.7
0.7
0.7
0.7
0.7
0.7
0.7
0.7
0.7
0.7


2
0
0
0
0
0
0
0
0
0
0
1
0
1
0
0
0
0
0
0
0
0
0
0
0
0
0
0
0
0


2
0
0
0
0
0
0
0
0
0
0
1
0
1
0
0
0
0
0
0
0
0
0
0
0
0
0
0
0
0

0.95
0.95
0.95
0.95
0.95
0.95
0.95
0.95
0.95
0.95
0.95
0.95
0.95
0.95
0.95
0.95
0.95
0.95
0.95
0.95
0.95
0.95
0.95
0.95
0.95
0.95
0.95
0.95
0.95
0.95


1
0
0
0
0
0
0
0
0
0
1
0
0
0
0
0
0
0
0
0
0
0
0
0
0
0
0
0
0
0

0.99
0.99
0.99
0.99
0.99
0.99
0.99
0.99
0.99
0.99
0.99
0.99
0.99
0.99
0.99
0.99
0.99
0.99
0.99
0.99
0.99
0.99
0.99
0.99
0.99
0.99
0.99
0.99
0.99
0.99


1
0
0
0
0
0
0
0
0
0
0
0
0
0
0
0
0
0
0
0
0
0
1
0
0
0
0
0
0
0


1
0
0
0
0
0
0
0
0
0
0
0
0
0
0
0
0
0
0
0
0
0
1
0
0
0
0
0
0
0

1.0
1.0
1.0
1.0
1.0
1.0
1.0
1.0
1.0
1.0
1.0
1.0
1.0
1.0
1.0
1.0
1.0
1.0
1.0
1.0
1.0
1.0
1.0
1.0
1.0
1.0
1.0
1.0
1.0
1.0


1
0
0
0
0
0
0
0
0
0
0
0
0
0
0
0
0
0
0
0
0
0
0
0
0
0
0
0
1
0


1
0
0
0
0
0
0
0
0
0
0
0
0
0
0
0
0
0
0
0
0
0
0
0
0
0
0
0
1
0

1.0
1.0
1.0
1.0
1.0
1.0
1.0
1.0
1.0
1.0
1.0
1.0
1.0
1.0
1.0
1.0
1.0
1.0
1.0
1.0
1.0
1.0
1.0
1.0
1.0
1.0
1.0
1.0
1.0
1.0


6
0
0
0
0
0
0
0
0
1
0
0
0
1
0
0
0
0
0
0
0
0
0
0
0
0
2
1
1
0


1
0
0
0
0
0
0
0
0
1
0
0
0
0
0
0
0
0
0
0
0
0
0
0
0
0
0
0
0
0


1
0
0
0
0
0
0
0
0
1
0
0
0
0
0
0
0
0
0
0
0
0
0
0
0
0
0
0
0
0

1.0
1.0
1.0
1.0
1.0
1.0
1.0
1.0
1.0
1.0
1.0
1.0
1.0
1.0
1.0
1.0
1.0
1.0
1.0
1.0
1.0
1.0
1.0
1.0
1.0
1.0
1.0
1.0
1.0
1.0


2
0
0
0
0
0
0
0
0
0
0
0
0
0
0
0
0
0
0
0
0
0
0
0
0
0
1
1
0
0


2
0
0
0
0
0
0
0
0
0
0
0
0
0
0
0
0
0
0
0
0
0
0
0
0
0
1
1
0
0

0.73
0.73
0.73
0.73
0.73
0.73
0.73
0.73
0.73
0.73
0.73
0.73
0.73
0.73
0.73
0.73
0.73
0.73
0.73
0.73
0.73
0.73
0.73
0.73
0.73
0.73
0.73
0.73
0.73
0.73


2
0
0
0
0
0
0
0
0
0
0
0
0
0
0
0
0
0
0
0
0
0
0
0
0
0
1
0
1
0


2
0
0
0
0
0
0
0
0
0
0
0
0
0
0
0
0
0
0
0
0
0
0
0
0
0
1
0
1
0

1.0
1.0
1.0
1.0
1.0
1.0
1.0
1.0
1.0
1.0
1.0
1.0
1.0
1.0
1.0
1.0
1.0
1.0
1.0
1.0
1.0
1.0
1.0
1.0
1.0
1.0
1.0
1.0
1.0
1.0


1
0
0
0
0
0
0
0
0
0
0
0
0
1
0
0
0
0
0
0
0
0
0
0
0
0
0
0
0
0


1
0
0
0
0
0
0
0
0
0
0
0
0
1
0
0
0
0
0
0
0
0
0
0
0
0
0
0
0
0

0.87
0.87
0.87
0.87
0.87
0.87
0.87
0.87
0.87
0.87
0.87
0.87
0.87
0.87
0.87
0.87
0.87
0.87
0.87
0.87
0.87
0.87
0.87
0.87
0.87
0.87
0.87
0.87
0.87
0.87


1
0
0
0
0
0
0
0
0
0
1
0
0
0
0
0
0
0
0
0
0
0
0
0
0
0
0
0
0
0


1
0
0
0
0
0
0
0
0
0
1
0
0
0
0
0
0
0
0
0
0
0
0
0
0
0
0
0
0
0


1
0
0
0
0
0
0
0
0
0
1
0
0
0
0
0
0
0
0
0
0
0
0
0
0
0
0
0
0
0

0.95
0.95
0.95
0.95
0.95
0.95
0.95
0.95
0.95
0.95
0.95
0.95
0.95
0.95
0.95
0.95
0.95
0.95
0.95
0.95
0.95
0.95
0.95
0.95
0.95
0.95
0.95
0.95
0.95
0.95


2
0
0
0
0
0
0
0
0
0
0
0
0
0
0
2
0
0
0
0
0
0
0
0
0
0
0
0
0
0


2
0
0
0
0
0
0
0
0
0
0
0
0
0
0
2
0
0
0
0
0
0
0
0
0
0
0
0
0
0


2
0
0
0
0
0
0
0
0
0
0
0
0
0
0
2
0
0
0
0
0
0
0
0
0
0
0
0
0
0

1.0
1.0
1.0
1.0
1.0
1.0
1.0
1.0
1.0
1.0
1.0
1.0
1.0
1.0
1.0
1.0
1.0
1.0
1.0
1.0
1.0
1.0
1.0
1.0
1.0
1.0
1.0
1.0
1.0
1.0


2
0
0
0
0
0
0
0
0
0
0
0
0
0
0
0
0
0
0
0
1
0
0
0
0
1
0
0
0
0


2
0
0
0
0
0
0
0
0
0
0
0
0
0
0
0
0
0
0
0
1
0
0
0
0
1
0
0
0
0


2
0
0
0
0
0
0
0
0
0
0
0
0
0
0
0
0
0
0
0
1
0
0
0
0
1
0
0
0
0

1.0
1.0
1.0
1.0
1.0
1.0
1.0
1.0
1.0
1.0
1.0
1.0
1.0
1.0
1.0
1.0
1.0
1.0
1.0
1.0
1.0
1.0
1.0
1.0
1.0
1.0
1.0
1.0
1.0
1.0


2
0
0
0
0
0
0
0
0
0
0
0
0
0
0
0
0
0
1
0
0
0
0
0
0
0
0
1
0
0


2
0
0
0
0
0
0
0
0
0
0
0
0
0
0
0
0
0
1
0
0
0
0
0
0
0
0
1
0
0


2
0
0
0
0
0
0
0
0
0
0
0
0
0
0
0
0
0
1
0
0
0
0
0
0
0
0
1
0
0

0.84
0.84
0.84
0.84
0.84
0.84
0.84
0.84
0.84
0.84
0.84
0.84
0.84
0.84
0.84
0.84
0.84
0.84
0.84
0.84
0.84
0.84
0.84
0.84
0.84
0.84
0.84
0.84
0.84
0.84


10
2
0
0
0
0
0
1
0
0
3
2
0
1
0
0
0
0
0
0
0
0
0
0
1
0
0
0
0
0


8
2
0
0
0
0
0
1
0
0
3
2
0
0
0
0
0
0
0
0
0
0
0
0
0
0
0
0
0
0


8
2
0
0
0
0
0
1
0
0
3
2
0
0
0
0
0
0
0
0
0
0
0
0
0
0
0
0
0
0


8
2
0
0
0
0
0
1
0
0
3
2
0
0
0
0
0
0
0
0
0
0
0
0
0
0
0
0
0
0

1.0
1.0
1.0
1.0
1.0
1.0
1.0
1.0
1.0
1.0
1.0
1.0
1.0
1.0
1.0
1.0
1.0
1.0
1.0
1.0
1.0
1.0
1.0
1.0
1.0
1.0
1.0
1.0
1.0
1.0


2
0
0
0
0
0
0
0
0
0
0
0
0
1
0
0
0
0
0
0
0
0
0
0
1
0
0
0
0
0

0.74
0.74
0.74
0.74
0.74
0.74
0.74
0.74
0.74
0.74
0.74
0.74
0.74
0.74
0.74
0.74
0.74
0.74
0.74
0.74
0.74
0.74
0.74
0.74
0.74
0.74
0.74
0.74
0.74
0.74


1
0
0
0
0
0
0
0
0
0
0
0
0
0
0
0
0
0
0
0
0
0
0
0
1
0
0
0
0
0


1
0
0
0
0
0
0
0
0
0
0
0
0
0
0
0
0
0
0
0
0
0
0
0
1
0
0
0
0
0

0.96
0.96
0.96
0.96
0.96
0.96
0.96
0.96
0.96
0.96
0.96
0.96
0.96
0.96
0.96
0.96
0.96
0.96
0.96
0.96
0.96
0.96
0.96
0.96
0.96
0.96
0.96
0.96
0.96
0.96


4
0
0
0
0
0
0
0
0
0
0
0
0
0
0
0
0
0
1
1
1
0
0
0
0
0
0
0
1
0


2
0
0
0
0
0
0
0
0
0
0
0
0
0
0
0
0
0
1
1
0
0
0
0
0
0
0
0
0
0


2
0
0
0
0
0
0
0
0
0
0
0
0
0
0
0
0
0
1
1
0
0
0
0
0
0
0
0
0
0


2
0
0
0
0
0
0
0
0
0
0
0
0
0
0
0
0
0
1
1
0
0
0
0
0
0
0
0
0
0

1.0
1.0
1.0
1.0
1.0
1.0
1.0
1.0
1.0
1.0
1.0
1.0
1.0
1.0
1.0
1.0
1.0
1.0
1.0
1.0
1.0
1.0
1.0
1.0
1.0
1.0
1.0
1.0
1.0
1.0


1
0
0
0
0
0
0
0
0
0
0
0
0
0
0
0
0
0
0
0
1
0
0
0
0
0
0
0
0
0


1
0
0
0
0
0
0
0
0
0
0
0
0
0
0
0
0
0
0
0
1
0
0
0
0
0
0
0
0
0


1
0
0
0
0
0
0
0
0
0
0
0
0
0
0
0
0
0
0
0
1
0
0
0
0
0
0
0
0
0

0.77
0.77
0.77
0.77
0.77
0.77
0.77
0.77
0.77
0.77
0.77
0.77
0.77
0.77
0.77
0.77
0.77
0.77
0.77
0.77
0.77
0.77
0.77
0.77
0.77
0.77
0.77
0.77
0.77
0.77


1
0
0
0
0
0
0
0
0
0
0
0
0
0
0
0
0
0
0
0
0
0
0
0
0
0
0
0
1
0


1
0
0
0
0
0
0
0
0
0
0
0
0
0
0
0
0
0
0
0
0
0
0
0
0
0
0
0
1
0


1
0
0
0
0
0
0
0
0
0
0
0
0
0
0
0
0
0
0
0
0
0
0
0
0
0
0
0
1
0

1.0
1.0
1.0
1.0
1.0
1.0
1.0
1.0
1.0
1.0
1.0
1.0
1.0
1.0
1.0
1.0
1.0
1.0
1.0
1.0
1.0
1.0
1.0
1.0
1.0
1.0
1.0
1.0
1.0
1.0


7
0
0
0
0
0
0
0
0
0
0
0
0
0
0
0
7
0
0
0
0
0
0
0
0
0
0
0
0
0


7
0
0
0
0
0
0
0
0
0
0
0
0
0
0
0
7
0
0
0
0
0
0
0
0
0
0
0
0
0


7
0
0
0
0
0
0
0
0
0
0
0
0
0
0
0
7
0
0
0
0
0
0
0
0
0
0
0
0
0

0.73
0.73
0.73
0.73
0.73
0.73
0.73
0.73
0.73
0.73
0.73
0.73
0.73
0.73
0.73
0.73
0.73
0.73
0.73
0.73
0.73
0.73
0.73
0.73
0.73
0.73
0.73
0.73
0.73
0.73


6
0
0
0
0
0
0
0
0
0
0
0
0
0
0
0
6
0
0
0
0
0
0
0
0
0
0
0
0
0


6
0
0
0
0
0
0
0
0
0
0
0
0
0
0
0
6
0
0
0
0
0
0
0
0
0
0
0
0
0

1.0
1.0
1.0
1.0
1.0
1.0
1.0
1.0
1.0
1.0
1.0
1.0
1.0
1.0
1.0
1.0
1.0
1.0
1.0
1.0
1.0
1.0
1.0
1.0
1.0
1.0
1.0
1.0
1.0
1.0


35
2
4
3
3
0
0
2
1
0
5
7
0
0
0
0
0
0
0
0
3
0
0
0
4
0
0
0
1
0


35
2
4
3
3
0
0
2
1
0
5
7
0
0
0
0
0
0
0
0
3
0
0
0
4
0
0
0
1
0


35
2
4
3
3
0
0
2
1
0
5
7
0
0
0
0
0
0
0
0
3
0
0
0
4
0
0
0
1
0


29
1
4
3
3
0
0
2
1
0
2
7
0
0
0
0
0
0
0
0
1
0
0
0
4
0
0
0
1
0


10
0
1
0
0
0
0
1
1
0
2
2
0
0
0
0
0
0
0
0
1
0
0
0
1
0
0
0
1
0


10
0
1
0
0
0
0
1
1
0
2
2
0
0
0
0
0
0
0
0
1
0
0
0
1
0
0
0
1
0

1.0
1.0
1.0
1.0
1.0
1.0
1.0
1.0
1.0
1.0
1.0
1.0
1.0
1.0
1.0
1.0
1.0
1.0
1.0
1.0
1.0
1.0
1.0
1.0
1.0
1.0
1.0
1.0
1.0
1.0


4
0
1
0
0
0
0
1
0
0
0
1
0
0
0
0
0
0
0
0
0
0
0
0
1
0
0
0
0
0


4
0
1
0
0
0
0
1
0
0
0
1
0
0
0
0
0
0
0
0
0
0
0
0
1
0
0
0
0
0

1.0
1.0
1.0
1.0
1.0
1.0
1.0
1.0
1.0
1.0
1.0
1.0
1.0
1.0
1.0
1.0
1.0
1.0
1.0
1.0
1.0
1.0
1.0
1.0
1.0
1.0
1.0
1.0
1.0
1.0


12
1
0
3
3
0
0
0
0
0
0
3
0
0
0
0
0
0
0
0
0
0
0
0
2
0
0
0
0
0

1.0
1.0
1.0
1.0
1.0
1.0
1.0
1.0
1.0
1.0
1.0
1.0
1.0
1.0
1.0
1.0
1.0
1.0
1.0
1.0
1.0
1.0
1.0
1.0
1.0
1.0
1.0
1.0
1.0
1.0


4
1
0
1
1
0
0
0
0
0
0
1
0
0
0
0
0
0
0
0
0
0
0
0
0
0
0
0
0
0

0.7
0.7
0.7
0.7
0.7
0.7
0.7
0.7
0.7
0.7
0.7
0.7
0.7
0.7
0.7
0.7
0.7
0.7
0.7
0.7
0.7
0.7
0.7
0.7
0.7
0.7
0.7
0.7
0.7
0.7


5
0
0
1
1
0
0
0
0
0
0
1
0
0
0
0
0
0
0
0
0
0
0
0
2
0
0
0
0
0

1.0
1.0
1.0
1.0
1.0
1.0
1.0
1.0
1.0
1.0
1.0
1.0
1.0
1.0
1.0
1.0
1.0
1.0
1.0
1.0
1.0
1.0
1.0
1.0
1.0
1.0
1.0
1.0
1.0
1.0


3
0
2
0
0
0
0
0
0
0
0
1
0
0
0
0
0
0
0
0
0
0
0
0
0
0
0
0
0
0

0.87
0.87
0.87
0.87
0.87
0.87
0.87
0.87
0.87
0.87
0.87
0.87
0.87
0.87
0.87
0.87
0.87
0.87
0.87
0.87
0.87
0.87
0.87
0.87
0.87
0.87
0.87
0.87
0.87
0.87


6
1
0
0
0
0
0
0
0
0
3
0
0
0
0
0
0
0
0
0
2
0
0
0
0
0
0
0
0
0


6
1
0
0
0
0
0
0
0
0
3
0
0
0
0
0
0
0
0
0
2
0
0
0
0
0
0
0
0
0


5
1
0
0
0
0
0
0
0
0
2
0
0
0
0
0
0
0
0
0
2
0
0
0
0
0
0
0
0
0

1.0
1.0
1.0
1.0
1.0
1.0
1.0
1.0
1.0
1.0
1.0
1.0
1.0
1.0
1.0
1.0
1.0
1.0
1.0
1.0
1.0
1.0
1.0
1.0
1.0
1.0
1.0
1.0
1.0
1.0


1
0
0
0
0
0
0
0
0
0
1
0
0
0
0
0
0
0
0
0
0
0
0
0
0
0
0
0
0
0

0.92
0.92
0.92
0.92
0.92
0.92
0.92
0.92
0.92
0.92
0.92
0.92
0.92
0.92
0.92
0.92
0.92
0.92
0.92
0.92
0.92
0.92
0.92
0.92
0.92
0.92
0.92
0.92
0.92
0.92


20
0
1
0
1
0
1
2
2
3
2
1
0
0
1
0
0
1
0
0
0
1
1
0
0
1
1
1
0
0


20
0
1
0
1
0
1
2
2
3
2
1
0
0
1
0
0
1
0
0
0
1
1
0
0
1
1
1
0
0


20
0
1
0
1
0
1
2
2
3
2
1
0
0
1
0
0
1
0
0
0
1
1
0
0
1
1
1
0
0

1.0
1.0
1.0
1.0
1.0
1.0
1.0
1.0
1.0
1.0
1.0
1.0
1.0
1.0
1.0
1.0
1.0
1.0
1.0
1.0
1.0
1.0
1.0
1.0
1.0
1.0
1.0
1.0
1.0
1.0


2
0
0
0
0
0
0
1
0
0
0
0
0
0
0
0
0
0
0
0
0
1
0
0
0
0
0
0
0
0


2
0
0
0
0
0
0
1
0
0
0
0
0
0
0
0
0
0
0
0
0
1
0
0
0
0
0
0
0
0


2
0
0
0
0
0
0
1
0
0
0
0
0
0
0
0
0
0
0
0
0
1
0
0
0
0
0
0
0
0

0.93
0.93
0.93
0.93
0.93
0.93
0.93
0.93
0.93
0.93
0.93
0.93
0.93
0.93
0.93
0.93
0.93
0.93
0.93
0.93
0.93
0.93
0.93
0.93
0.93
0.93
0.93
0.93
0.93
0.93


1
0
0
0
0
0
0
0
0
0
0
0
0
0
0
0
0
0
0
0
0
0
1
0
0
0
0
0
0
0


1
0
0
0
0
0
0
0
0
0
0
0
0
0
0
0
0
0
0
0
0
0
1
0
0
0
0
0
0
0


1
0
0
0
0
0
0
0
0
0
0
0
0
0
0
0
0
0
0
0
0
0
1
0
0
0
0
0
0
0

0.76
0.76
0.76
0.76
0.76
0.76
0.76
0.76
0.76
0.76
0.76
0.76
0.76
0.76
0.76
0.76
0.76
0.76
0.76
0.76
0.76
0.76
0.76
0.76
0.76
0.76
0.76
0.76
0.76
0.76


1
0
0
0
0
0
0
0
0
0
0
0
0
0
0
0
0
1
0
0
0
0
0
0
0
0
0
0
0
0


1
0
0
0
0
0
0
0
0
0
0
0
0
0
0
0
0
1
0
0
0
0
0
0
0
0
0
0
0
0

0.88
0.88
0.88
0.88
0.88
0.88
0.88
0.88
0.88
0.88
0.88
0.88
0.88
0.88
0.88
0.88
0.88
0.88
0.88
0.88
0.88
0.88
0.88
0.88
0.88
0.88
0.88
0.88
0.88
0.88


1
0
0
0
0
0
0
0
0
0
0
0
0
0
0
0
0
0
0
1
0
0
0
0
0
0
0
0
0
0


1
0
0
0
0
0
0
0
0
0
0
0
0
0
0
0
0
0
0
1
0
0
0
0
0
0
0
0
0
0


1
0
0
0
0
0
0
0
0
0
0
0
0
0
0
0
0
0
0
1
0
0
0
0
0
0
0
0
0
0


1
0
0
0
0
0
0
0
0
0
0
0
0
0
0
0
0
0
0
1
0
0
0
0
0
0
0
0
0
0


1
0
0
0
0
0
0
0
0
0
0
0
0
0
0
0
0
0
0
1
0
0
0
0
0
0
0
0
0
0


1
0
0
0
0
0
0
0
0
0
0
0
0
0
0
0
0
0
0
1
0
0
0
0
0
0
0
0
0
0

1.0
1.0
1.0
1.0
1.0
1.0
1.0
1.0
1.0
1.0
1.0
1.0
1.0
1.0
1.0
1.0
1.0
1.0
1.0
1.0
1.0
1.0
1.0
1.0
1.0
1.0
1.0
1.0
1.0
1.0


3
0
0
0
0
0
0
1
1
1
0
0
0
0
0
0
0
0
0
0
0
0
0
0
0
0
0
0
0
0


3
0
0
0
0
0
0
1
1
1
0
0
0
0
0
0
0
0
0
0
0
0
0
0
0
0
0
0
0
0


3
0
0
0
0
0
0
1
1
1
0
0
0
0
0
0
0
0
0
0
0
0
0
0
0
0
0
0
0
0

0.73
0.73
0.73
0.73
0.73
0.73
0.73
0.73
0.73
0.73
0.73
0.73
0.73
0.73
0.73
0.73
0.73
0.73
0.73
0.73
0.73
0.73
0.73
0.73
0.73
0.73
0.73
0.73
0.73
0.73


208
24
21
12
17
17
22
11
12
16
19
21
1
0
0
0
0
2
3
2
2
1
1
1
1
0
0
1
0
1


208
24
21
12
17
17
22
11
12
16
19
21
1
0
0
0
0
2
3
2
2
1
1
1
1
0
0
1
0
1


208
24
21
12
17
17
22
11
12
16
19
21
1
0
0
0
0
2
3
2
2
1
1
1
1
0
0
1
0
1


208
24
21
12
17
17
22
11
12
16
19
21
1
0
0
0
0
2
3
2
2
1
1
1
1
0
0
1
0
1


208
24
21
12
17
17
22
11
12
16
19
21
1
0
0
0
0
2
3
2
2
1
1
1
1
0
0
1
0
1


208
24
21
12
17
17
22
11
12
16
19
21
1
0
0
0
0
2
3
2
2
1
1
1
1
0
0
1
0
1

1.0
1.0
1.0
1.0
1.0
1.0
1.0
1.0
1.0
1.0
1.0
1.0
1.0
1.0
1.0
1.0
1.0
1.0
1.0
1.0
1.0
1.0
1.0
1.0
1.0
1.0
1.0
1.0
1.0
1.0


170
20
18
10
14
13
20
10
11
14
17
16
1
0
0
0
0
0
1
0
1
1
1
1
0
0
0
0
0
1

1.0
1.0
1.0
1.0
1.0
1.0
1.0
1.0
1.0
1.0
1.0
1.0
1.0
1.0
1.0
1.0
1.0
1.0
1.0
1.0
1.0
1.0
1.0
1.0
1.0
1.0
1.0
1.0
1.0
1.0


48
4
1
0
1
3
3
3
3
3
3
4
0
0
0
0
1
3
1
1
0
0
1
0
3
4
2
0
2
2


48
4
1
0
1
3
3
3
3
3
3
4
0
0
0
0
1
3
1
1
0
0
1
0
3
4
2
0
2
2

0.99
0.99
0.99
0.99
0.99
0.99
0.99
0.99
0.99
0.99
0.99
0.99
0.99
0.99
0.99
0.99
0.99
0.99
0.99
0.99
0.99
0.99
0.99
0.99
0.99
0.99
0.99
0.99
0.99
0.99


41
3
0
0
1
3
3
3
3
3
3
3
0
0
0
0
1
3
1
1
0
0
1
0
2
3
1
0
2
1


40
3
0
0
1
3
3
3
3
3
3
3
0
0
0
0
1
3
1
1
0
0
1
0
2
2
1
0
2
1


40
3
0
0
1
3
3
3
3
3
3
3
0
0
0
0
1
3
1
1
0
0
1
0
2
2
1
0
2
1


40
3
0
0
1
3
3
3
3
3
3
3
0
0
0
0
1
3
1
1
0
0
1
0
2
2
1
0
2
1

0.84
0.84
0.84
0.84
0.84
0.84
0.84
0.84
0.84
0.84
0.84
0.84
0.84
0.84
0.84
0.84
0.84
0.84
0.84
0.84
0.84
0.84
0.84
0.84
0.84
0.84
0.84
0.84
0.84
0.84


17
2
0
0
1
2
2
2
2
2
2
2
0
0
0
0
0
0
0
0
0
0
0
0
0
0
0
0
0
0

1.0
1.0
1.0
1.0
1.0
1.0
1.0
1.0
1.0
1.0
1.0
1.0
1.0
1.0
1.0
1.0
1.0
1.0
1.0
1.0
1.0
1.0
1.0
1.0
1.0
1.0
1.0
1.0
1.0
1.0


8
0
0
0
0
0
0
0
0
0
0
0
0
0
0
0
1
2
0
0
0
0
1
0
1
1
1
0
0
1

0.83
0.83
0.83
0.83
0.83
0.83
0.83
0.83
0.83
0.83
0.83
0.83
0.83
0.83
0.83
0.83
0.83
0.83
0.83
0.83
0.83
0.83
0.83
0.83
0.83
0.83
0.83
0.83
0.83
0.83


6
0
0
0
0
0
0
0
0
0
0
0
0
0
0
0
0
1
1
1
0
0
0
0
1
1
0
0
1
0

0.8
0.8
0.8
0.8
0.8
0.8
0.8
0.8
0.8
0.8
0.8
0.8
0.8
0.8
0.8
0.8
0.8
0.8
0.8
0.8
0.8
0.8
0.8
0.8
0.8
0.8
0.8
0.8
0.8
0.8


1
0
0
0
0
0
0
0
0
0
0
0
0
0
0
0
0
0
0
0
0
0
0
0
0
0
0
0
1
0

0.92
0.92
0.92
0.92
0.92
0.92
0.92
0.92
0.92
0.92
0.92
0.92
0.92
0.92
0.92
0.92
0.92
0.92
0.92
0.92
0.92
0.92
0.92
0.92
0.92
0.92
0.92
0.92
0.92
0.92


1
0
0
0
0
0
0
0
0
0
0
0
0
0
0
0
0
0
0
0
0
0
0
0
0
1
0
0
0
0


1
0
0
0
0
0
0
0
0
0
0
0
0
0
0
0
0
0
0
0
0
0
0
0
0
1
0
0
0
0


1
0
0
0
0
0
0
0
0
0
0
0
0
0
0
0
0
0
0
0
0
0
0
0
0
1
0
0
0
0


1
0
0
0
0
0
0
0
0
0
0
0
0
0
0
0
0
0
0
0
0
0
0
0
0
1
0
0
0
0

0.87
0.87
0.87
0.87
0.87
0.87
0.87
0.87
0.87
0.87
0.87
0.87
0.87
0.87
0.87
0.87
0.87
0.87
0.87
0.87
0.87
0.87
0.87
0.87
0.87
0.87
0.87
0.87
0.87
0.87


4
1
1
0
0
0
0
0
0
0
0
1
0
0
0
0
0
0
0
0
0
0
0
0
1
0
0
0
0
0


4
1
1
0
0
0
0
0
0
0
0
1
0
0
0
0
0
0
0
0
0
0
0
0
1
0
0
0
0
0


4
1
1
0
0
0
0
0
0
0
0
1
0
0
0
0
0
0
0
0
0
0
0
0
1
0
0
0
0
0

0.9
0.9
0.9
0.9
0.9
0.9
0.9
0.9
0.9
0.9
0.9
0.9
0.9
0.9
0.9
0.9
0.9
0.9
0.9
0.9
0.9
0.9
0.9
0.9
0.9
0.9
0.9
0.9
0.9
0.9


1
0
0
0
0
0
0
0
0
0
0
0
0
0
0
0
0
0
0
0
0
0
0
0
1
0
0
0
0
0

0.84
0.84
0.84
0.84
0.84
0.84
0.84
0.84
0.84
0.84
0.84
0.84
0.84
0.84
0.84
0.84
0.84
0.84
0.84
0.84
0.84
0.84
0.84
0.84
0.84
0.84
0.84
0.84
0.84
0.84


2
0
0
0
0
0
0
0
0
0
0
0
0
0
0
0
0
0
0
0
0
0
0
0
0
0
1
0
0
1


1
0
0
0
0
0
0
0
0
0
0
0
0
0
0
0
0
0
0
0
0
0
0
0
0
0
0
0
0
1


1
0
0
0
0
0
0
0
0
0
0
0
0
0
0
0
0
0
0
0
0
0
0
0
0
0
0
0
0
1


1
0
0
0
0
0
0
0
0
0
0
0
0
0
0
0
0
0
0
0
0
0
0
0
0
0
0
0
0
1

0.76
0.76
0.76
0.76
0.76
0.76
0.76
0.76
0.76
0.76
0.76
0.76
0.76
0.76
0.76
0.76
0.76
0.76
0.76
0.76
0.76
0.76
0.76
0.76
0.76
0.76
0.76
0.76
0.76
0.76


1
0
0
0
0
0
0
0
0
0
0
0
0
0
0
0
0
0
0
0
0
0
0
0
0
0
1
0
0
0


1
0
0
0
0
0
0
0
0
0
0
0
0
0
0
0
0
0
0
0
0
0
0
0
0
0
1
0
0
0


1
0
0
0
0
0
0
0
0
0
0
0
0
0
0
0
0
0
0
0
0
0
0
0
0
0
1
0
0
0


1
0
0
0
0
0
0
0
0
0
0
0
0
0
0
0
0
0
0
0
0
0
0
0
0
0
1
0
0
0

1.0
1.0
1.0
1.0
1.0
1.0
1.0
1.0
1.0
1.0
1.0
1.0
1.0
1.0
1.0
1.0
1.0
1.0
1.0
1.0
1.0
1.0
1.0
1.0
1.0
1.0
1.0
1.0
1.0
1.0
