## Supplementary material for "Charting the Insect Biodiversity of Crete: Insights from a Pilot Metabarcoding Survey": README.md

# Supplementary Data

## RaxTax results above 70 and filtered using phylogenetic results and exact matches
- `raxtax_filtered_DatasetA.tsv`
- `raxtax_filtered_DatasetB.tsv`

## ESVs from the analysis in FASTA format
- `ESVs_DatasetA.fasta`
- `ESVs_DatasetB.fasta`

## Latest checklist for the 14 insect orders of Crete
- `Insect_Species_Checklist_of_Crete.xlsx`

## Abundance of each ESV in each library
- `ESVs.lib.stats_DatasetA.tsv`
- `ESVs.lib.stats_DatasetB.tsv`

## Raw RaxTax results
- `raxtax_raw_DatasetA.tsv`
- `raxtax_raw_DatasetB.tsv`

## List of species present in the samples
- `Species_present_in_the_samples.xlsx`

## Krona plots using either ESV counts or sequence abundance
- `Krona_esvs.html`
- `Krona_abundances.html`

## Lifemap visualizations using either ESV counts or sequence abundance
- `Lifemap_esvs.html`
- `Lifemap_abundances.html`
